# Transcriptome profiling of the dorsomedial prefrontal cortex in suicide victims

**DOI:** 10.1101/2022.04.19.488577

**Authors:** Fanni Dóra, Éva Renner, Dávid Keller, Miklós Palkovits, Árpád Dobolyi

## Abstract

The default mode network (DMN) plays an outstanding role in psychiatric disorders. Still, gene expressional changes in its major component, the dorsomedial prefrontal cortex (DMPFC), have not been characterized. We used RNA-sequencing in postmortem DMPFC samples to investigate suicide victims compared to control subjects. Most of the data variance (79%) was associated with expression changes between suicide and control samples. 1400 genes differed using log2FC > ± 1 and adjusted *p*-value < 0.05 criteria between groups. Genes associated with depressive disorder, schizophrenia and impaired cognition were strongly overexpressed in top differentially expressed genes. Gene set enrichment and protein-protein interaction network analysis revealed that pathways related to cytokine receptor signaling were enriched in downregulated while glutamatergic synaptic signaling in upregulated genes in suicide individuals. A validated differentially expressed gene, which is known to be associated with mGluR5, was the N-terminal EF-hand calcium-binding proteins 2 (NECAB2). *In situ* hybridization histochemistry and immunohistochemistry proved that NECAB2 is expressed in 2 different types of inhibitory neurons located in layers II-IV and VI, respectively. Our results imply extensive gene expressional alterations in the DMPFC related to suicidal behavior. Some of these genes may contribute to the altered mental state and behavior of suicide victims.

## INTRODUCTION

Neuroimaging studies revealed abnormal resting-state functional connectivity (RSFC) between distributed brain areas in major depression. The connection between the medial prefrontal cortex (MPFC) and the medial posterior parietal cortex (MPPC) has been named as resting-state network (RSN) since they have temporal correlation in the absence of a variety of tasks in resting condition (Biswal et al. 1995, Greicius et al. 2003, Greicius et al. 2009, Raichle and Snyder 2007). Since the RSN includes distinct networks such as the *default-mode (DMN)*, *salience* and *attention networks* (Isaacs et al. 2018, Vincent et al. 2008, Figley et al. 2015, Cao et al. 2016, Zhang, Geng, and Lee 2017, Hilger et al. 2017, Anticevic et al. 2012). Presently, terms RSN and RSFC are used as alternatives.

In the default-mode network, the *dorsomedial prefrontal cortex (DMPFC)* represents one of the nodes. It includes the superior frontal and the paracingulate gyrus (also called dorsal anterior cingulate cortex) on the medial surface of the frontal cortex (*see Methods).* The DMPFC is involved in a variety of functions, combining sensory signals from the outside world transferred by the lateral prefrontal (cognitive) and the orbitofrontal (motivational) cortices, as well as viscero-sensory signals through the insula and the anterior cingulate cortex (Cauda et al. 2011, Seeley et al. 2007). The DMPFC prepares executive motor programs for the supplementary and pre-supplementary motor cortices (Isaacs et al. 2018, Cacciola et al. 2017, Kotz 2011). The functions of the DMPFC include activation of working memories, decision making and planning (Buckner, Andrews-Hanna, and Schacter 2008, Cavanna and Trimble 2006, Báez-Mendoza et al. 2021, Anticevic et al. 2012, Kolling and O’Reilly 2018). The DMPFC is linked to social processes (Lieberman et al., 2019), emotional processing (Raschle et al. 2015), novelty processing and dynamic contextual updating (Solbakk and Lovstad 2014), conscious threat appraisal (Kalisch and Gerlicher 2014, Northoff et al. 2006).

The other node in the DMN is present in the MPPC including the precuneus and the posterior cingulate cortex (Figley et al. 2015, Chen et al. 2021, Utevsky, Smith, and Huettel 2014). The precuneus plays a significant role in the *human brain* mental imagination, individualization of cognitive processes, as reflexions to signals from the external world, introspection, self-awareness, creativity and intelligence (Bruner et al. 2017, Bruner et al. 2014, Bruton 2021, Cavanna 2007, Margulies et al. 2009). The precuneus directly projects to the supplementary and the pre-supplementary motor cortices (Beaty, Seli, and Schacter 2019, Fuentealba-Villarroel et al. 2021). In addition, the precuneus participates in the control of the motor executive program also through connections with the DMPFC (Wu et al. 2016).

One of the outputs of the RSN is connected with the cortico-striato (including nucleus accumbens/ventral striatum)-thalamo-cortical loop (circuit) that connects the DMPFC and the precuneus with the reward-association system (Chen et al. 2021, Haber and Knutson 2010, Isaacs et al. 2018). Dysfunction in the reward-association system results in major depression (Coenen et al. 2018, Xu, Nan, and Lan 2020, Chen et al. 2021).

Abnormal communication within the DMN has been reported in depression using RSFC analysis (Liu, Jiang, and Yuan 2018). Meta-analysis revealed that clinical response to several different treatments, including repetitive transcranial magnetic stimulation (TMS), pharmacotherapy, cognitive behavioral therapy, electroconvulsive therapy and transcutaneous vagal nerve stimulation could be predicted by baseline DMN connectivity in patients with depression (Long et al. 2020). In the DMPFC, depressed patients demonstrated increased functional connectivity with sensorimotor, visual and salience network regions (Yang et al. 2021). Furthermore, a major symptom of depression, anhedonia was associated with increased RSFC between seed regions of bilateral nucleus accumbens and areas of right DMPFC (Olson et al. 2018). Another recent line of research, which suggest that the DMPFC may be involved in depression, is that repetitive TMS of the DMPFC was found as a novel intervention for treatment-refractory depression (Kreuzer et al. 2019, Schulze et al. 2018).

Although depression is the most common psychiatric disorder in people who die by suicide (Hawton et al. 2013, Franklin et al. 2017, Turecki and Brent 2016), there may also be differences in the underlying mechanisms. A study found evidence of different patterns of connectivity strength within the DMN of depressed-suicidal and depressed-non-suicidal adult participants (Malhi et al. 2020). The involvement of the DMN in suicide is predicated on their key role in self-referential thinking, which may be a consequence of altered integrity within the DMN. On the basis of subregions previously reported to show structural and functional alterations within the DMN, functional activation studies applying cognitive or motor tasks generally showed altered involvement of the DMPFC linked with suicidal thoughts among depressed individuals (Dombrovski and Hallquist 2017, Marchand et al. 2012, Reisch et al. 2010).

Neuronal plasticity but also other pathological alterations associated with depression and suicidal behavior may involve gene expressional alterations (Bai et al. 2019). Initially, individual systems, such as GABAergic function (Ghosal, Hare, and Duman 2017), neurotrophic factors (Zhang, Yao, and Hashimoto 2016), glial cells (O’Leary et al. 2021) were examined. More recently, high throughput methods were applied to establish a wider variety of genes significantly altered in suicide victims. Microarray studies identified a number of altered genes, regulating e.g. glial, endothelial, and mitochondrial activities (Pantazatos et al. 2017) some of them showing sexual dimorphism (Labonte et al. 2017) and comorbidity with substance use disorder (Cabrera et al. 2019, Cabrera-Mendoza et al. 2020). More recently, sequencing approaches identified additional new susceptibility genes and pathways in depression using animal models of chronic stress (Li et al. 2020), and genome-wide association studies related to depression (Yang et al. 2021) and suicide (Romero-Pimentel et al. 2021). Although these gene expressional studies provided valuable data on the neuronal mechanisms of suicidal behavior, they were all performed in the dorsolateral prefrontal cortex. Given the above described roles of the DMPFC, we hypothesized that gene expressional changes associated with suicidal behavior take place in the DMPFC as well. We obtained DMPFC samples of suicide victims with relatively short postmortem delay whose investigation provided useful insight into the underlying reasons of suicidal behavior.

## RESULTS

### Transcriptome sequencing in the DMPFC of the suicide brains

The mean age, the sex ratios, PMI and the RNA quality scores did not show detectable differences between the suicide and control groups (8 individuals/groups) (Supplementary file 1, Supplementary figure 1). Samples with mean RNA integrity number (RIN) values > 4 were used in the study, which provided very good quality for sequencing based on the QC (Supplementary file 2, Supplementary figure 3) as expected (Puchta, Boczkowska, and Groszyk 2020, Purves-Tyson et al. 2021, Sonntag et al. 2016). In average, 62.9 ± 3.6 Mb total raw reads per sample were generated. The reads demonstrating inadequate-quality, adaptor-containing or unknown sequences were not included in downstream analysis. The average clean reads ratio after quality control (QC) was 91.87 ± 0.46%, and the average mapping ratio with the reference genome was 91.28 ± 0.79%. In parallel, Q20 and Q30 of the clean data were calculated (Supplementary file 2).

The average ratio of clean reads Q20 (the percentage of bases number which is higher than 20 in the total number of bases) was 97.46 ± 0.079%, the average ratio of clean reads Q30 (the percentage of bases number which is higher than 30 in the total number of bases) was 89.80 ± 0.19%. These analyses indicated the high-quality of library construction and sequencing data of postmortem human samples. The average mapping ratio with gene was 72.22 ± 0.96%. These high alignment percentages suggest the suitability of the sequenced data for downstream analysis. Totally, 29 791 genes were identified, out of which 28 088 were known genes. About 67% of the fragments were attributed to protein-coding genes. The remaining fragments were attributed to other RNA classes including long intergenic noncoding RNAs, pseudogenes, etc. The amount of total raw reads, clean reads and their percentages are shown in Supplementary file 2.

### Comparison of suicide and control samples based on differentially expressed genes (DEGs)

After counting the reads aligning to each gene, we used DESeq2 to analyze gene expression in suicide victims compared to matched controls. We detected expression of 19 692 protein-coding genes in the DMPFC, out of which 1400 were differentially expressed between the two groups, 1262 genes were down- and 138 genes were upregulated at the adjusted *p*-value < 0.05 and log2 FC > ±1 (Supplementary file 3). All samples were compared to one another using Euclidean distance. The correlation matrix based on all gene expression data indicated that samples mostly cluster by individual and diagnostic group (Figure 1A). Hierarchical clustering of suicide and control samples using the 1400 genes differentially expressed revealed that these genes distinguished the two groups with 3 misclassified control samples (Figure 1B).

**Figure 1.**
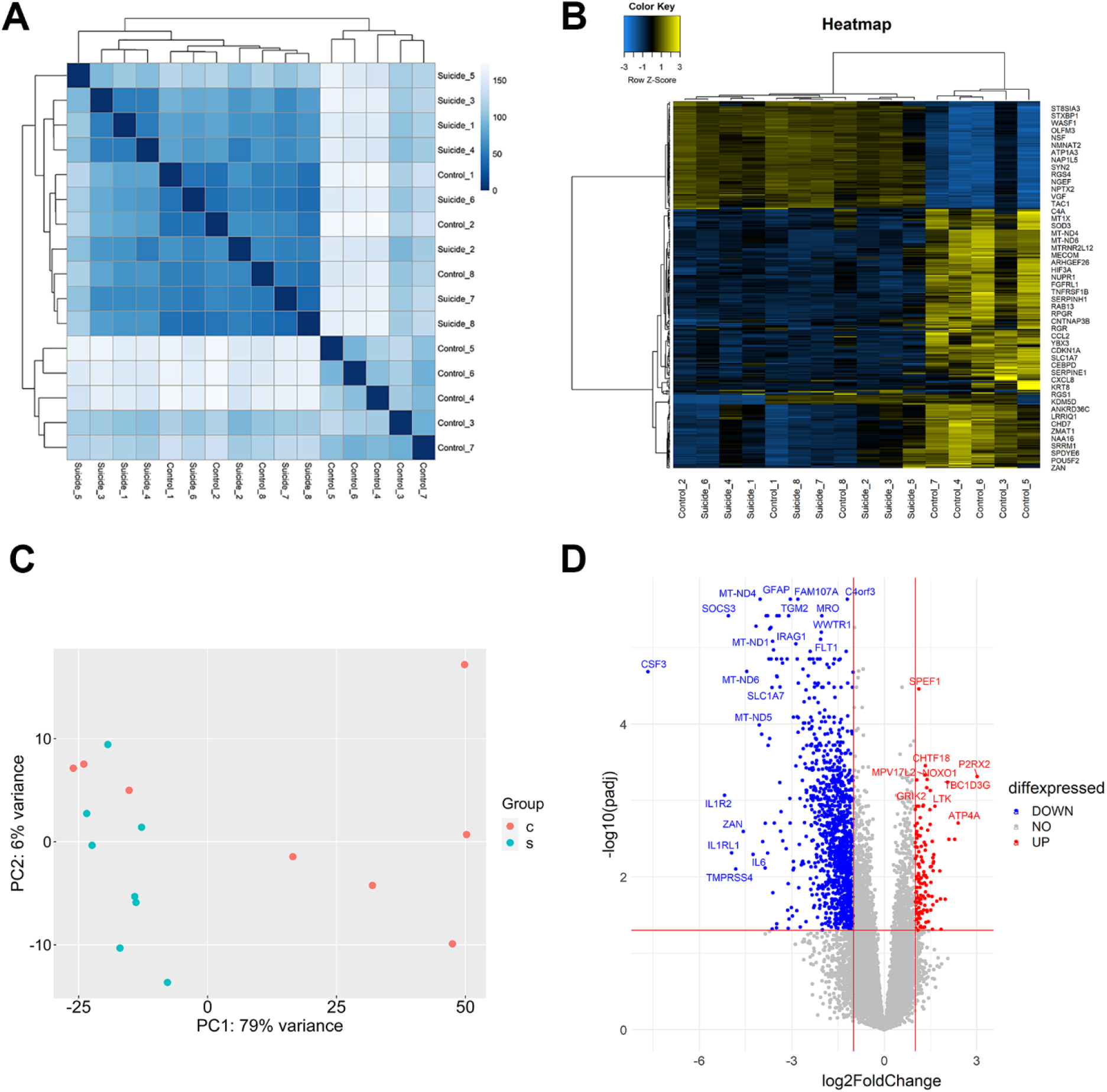
Analysis of gene expression between suicide and control groups. (A) Hierarchical clustering of 16 RNA-seq samples based on the Euclidean distance. A heatmap of the distance matrix shows the similarities and dissimilarities between the samples as calculated from the rlog transformation of the count data. (B) Hierarchical clustering heatmap of Pearson correlation coefficients between suicide and control individuals. The heatmap represents the top 500 protein coding gene expression for the different groups in columns, with a dendogram presented at the top of the heatmap. Proteins were selected based on their variability between all samples. Counts were log transformed, normalized, and used for clustering based on similarity in expression patterns. X-axis represents the sample. Y-axis represents the top 500 differentially expressed genes. The color represents the log10 transformed gene expression level. The scale shows the level of expression: the yellow color means high, the blue color means low, while black color represents medium expression level. (C) Principal Component Analysis plot (PCA) of the samples showing the clustering of VST (variance stabilizing transformation) transcriptomic data for visualizing the overall effect of experimental covariates and batch effects. Red dots represent control (c) while blue dots refer to suicide (s) individuals. (D) Volcano plot showing the log2 fold change (log2FC) of gene expression and the statistical significance of the differential expression (DE) analysis performed between suicide and control individuals. The x-axis represents the log2 fold change of genes, while the y-axis represents the −log10 of the corrected *p*-values (padj) for the different pairs of conditions. Each dot represents a gene and the colored areas represent the DEGs that met the following selection criteria: log2FC of at least ± 1 (log2FC ≥ 1 or ≤ −1) and Benjamini and Hochberg-adjusted *p*-value < 0.05. Upregulated genes are shown in red, while the downregulated ones are blue. The top 40 significant DEGs are labeled.

Principal component analysis (PCA) based on the expression of the DEGs confirmed that most of the variance in the data (79%, PC1) was associated with expression changes between the suicide and control samples (Figure 1C). Among the DEGs, 1262 genes were downregulated and 138 were upregulated in suicide victims, as shown in the volcano plot (Figure 1D).

### Functional annotation and classification of the DEGs

To understand how the DEGs from the DMPFC relate to suicidal behavior, Gene Ontology (GO) classification was performed. Figure 2A shows the top 10 down- and upregulated DEGs. The top 3 downregulated genes were CSF3, IL1R2 and SOCS3. They all play a critical role in inflammation. The top 3 upregulated genes were P2RX2, ATP4A and BX276092.9. P2RX2 and ATP4A are found in the plasma membrane and both of them have role in ATP binding and ion transport. BX276092.9 is a barely characterized protein. Additional GO functional enrichment analysis identified significant differences in more than 500 functional pathways of downregulated genes and 37 functional pathways of upregulated genes at adjusted *p*-value < 0.05 (Supplementary file 4). The top 3 enriched GO terms within biological process, molecular function, and cellular component categories were visualized (Figure 2B and C). A number of pathways related to cell surface receptor signaling (GO:0007166) and growth factor binding (GO:0019838) were enriched in downregulated genes in suicide individuals. In contrast, pathways pertaining to synaptic signaling (GO:0099536) and sodium channel activity (GO:0005272) were enriched in upregulated genes of suicide individuals (Figure 2B and C). To further identify the pathways involved in suicidal behavior, pathway analysis of DEGs was performed using pathway classifications from the Kyoto Encyclopedia of Genes and Genomes (KEGG) and Reactome resources (Statistics of Pathway Enrichment, Figure 2C). The top 3 enriched KEGG and Reactome pathways are shown in Figure 2C. A total of 23 significantly enriched downregulated KEGG pathways and 25 significantly enriched downregulated Reactome pathways were found in downregulated genes, while there was 1 significantly enriched upregulated KEGG pathway and 2 upregulated Reactome pathways at adjusted *p*-value < 0.05. The complete list of significantly enriched pathways can be found in Supplementary file 4.

**Figure 2.**
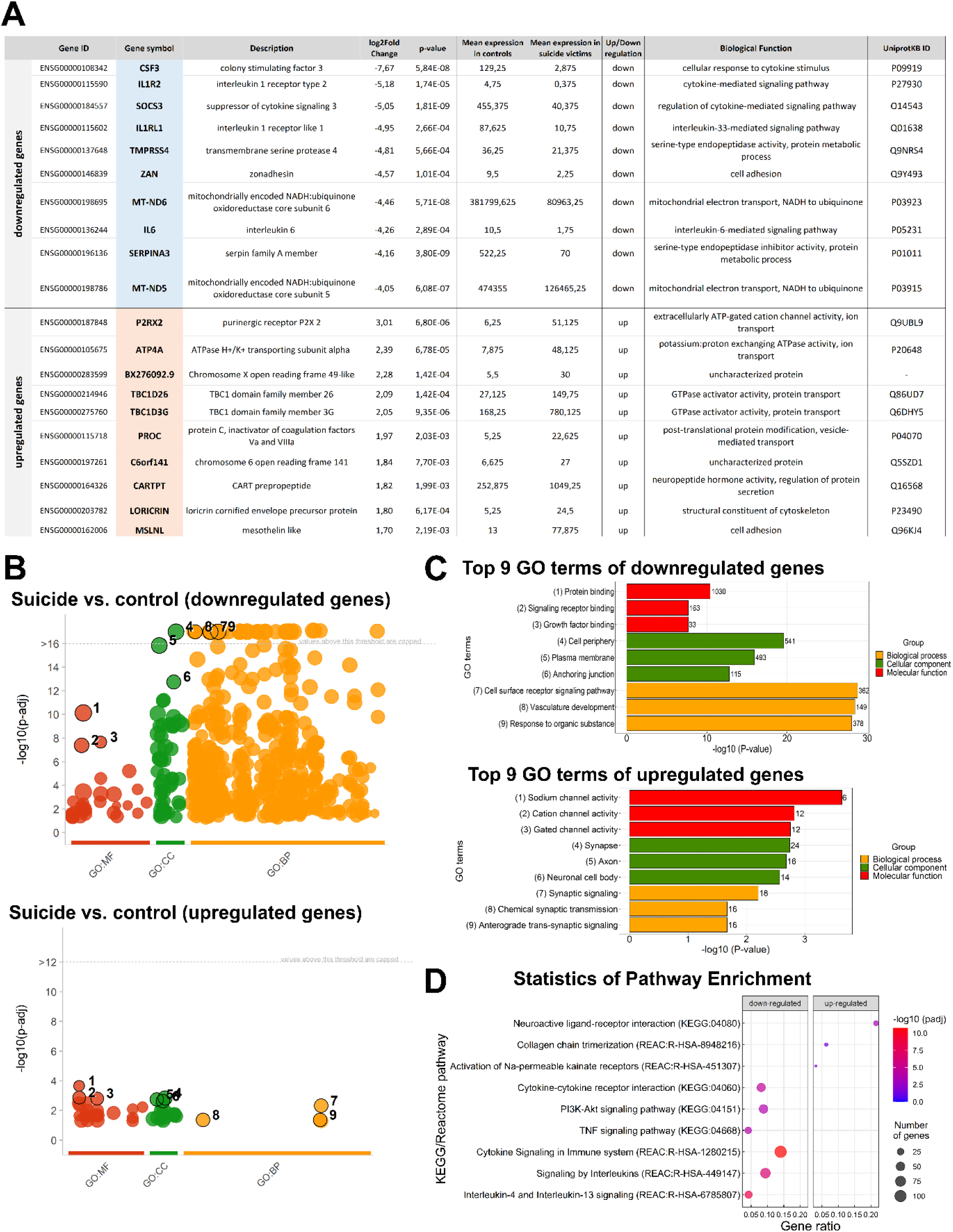
Functional enrichment of differentially expressed genes. (A) Top 10 down- and upregulated differentially expressed genes ranked by the value of the log2 fold change (log2FC) and their biological functions. (B) Manhattan plot shows the enrichment results of down- and upregulated genes. The x-axis shows the terms and y-axis shows the enrichment *p*-values on −log10 scale. Each circle on plot corresponds to a single term. Circles are colored according to the origin of annotation and size-scaled according to the total number of genes annotated to the corresponding term. The locations on the x-axis are fixed. Terms from the same GO subtree are located closer to each other on the x-axis, which helps to highlight different enriched GO sub-branches making plots from different queries comparable. The top 3 terms from each GO category are indicated with numbers on the plot. The corresponding statistics of annotation are listed in the Supplementary file 4. (C) GO classification and scatter plot-enriched Kyoto Encyclopedia of Genes and Genomes (KEGG) pathways of DEGs. Bar plots reporting the top 9 significantly enriched GO terms related to down- and upregulated genes with the adjusted *p*-value < 0.05, respectively. The table describes, for each GO term, the number of mapped annotated genes in the reference data set and its −log10 *p*-value. The color code is proportional to the three ontologies. (D) Dot plots of top 3 pathway annotations from KEGG and Reactome pathways illustrate the distributions of gene sets among up- and downregulated differentially expressed genes. The x-axis represents the gene ratio. The y-axis represents the pathway term. Gene ratio refers to the value of enrichment, which is the ratio of DEGs annotated in the pathway to total gene amount annotated in the pathway. The larger the value, the more significant the enrichment. The circle size indicates the DEG number associated with each significant pathway. The color indicates the adjusted *p*-value, the lower *p*-value indicates the more significant enrichment.

### Protein-protein interaction (PPI) network analysis of differentially expressed genes

Computational methods analyzing PPI networks are useful for understanding biological meaning of gene expressional alterations. Therefore, we used the STRING online database (http://string-db.org) to construct the PPI networks of down- and upregulated DEGs, which was then visualized and optimized using Cytoscape software (Figure 3A and C). The PPI network of downregulated genes is identified in 7826 interactions between the 1234 proteins found differentially expressed in suicide, with a PPI enrichment *p*-value < 1.0 × 10^−16^ (Figure 3A). The PPI network of downregulated genes was grouped into 234 relevant protein clusters using the Markov Clustering algorithm (MCL) clustering (Supplementary file 5), but for clarity, clusters with less than 10 genes were not shown in Figure 3B. Many of the clusters share interactions among them indicating that these proteins play key roles in diverse pathways. To conclude the functional associations, the 9 main clusters were classified according to GO Biological processes (Supplementary file 5). The largest cluster, Cluster 1 (red) is associated with cell surface receptor signaling pathway (GO:0007166) and cytokine-mediated signaling pathway (GO:0019221) including the most ranked three genes, the epidermal growth factor receptor (EGFR), fibronectin (FN1) and interleukin-6 (IL6). Cluster 7 (dark violet) represents proteins specifically involved in neurotransmitter uptake (GO:0001504), sodium ion transport (GO:0006814) and L-glutamate import across plasma membrane (GO:0098712), including the most ranked proteins excitatory amino acid transporter 1 (SLC1A3), glutamine synthase (GLUL) and excitatory amino acid transporter 5 (SLC1A7). Cluster 4 (pink) showed a significant enrichment of oxidative phosphorylation and mitochondrial electron transport (GO:0006119), NADH to ubiquinone (GO:0006120) related to the mitochondrial gene NADH dehydrogenase 1 (MT-ND1), cytochrome C Oxidase I (MT-CO1) and cytochrome C Oxidase II (MT-CO2). The PPI network of upregulated genes was grouped into 12 clusters using the MCL clustering (Supplementary file 5), and Figure 3D represents the three biggest clusters. Functional annotation of upregulated modules shows that the largest module, the Cluster 1 (soft red) represents a significant enrichment of chemical synaptic transmission (GO:0007268), regulation of biological quality (GO:0065008) and regulation of membrane potential (GO:0042391) related to ionotropic and metabotropic glutamate receptors (GRIK1, GRIK2 and GRM2) and voltage-sensitive calcium channel subunits (CACNA1G and CACNG8). Cluster 2 (blue) represents proteins involved in positive regulation of cytosolic calcium ion concentration (GO:0007204) including the polycystin-1 (PKD1), P2X purinoceptor 2 (P2RX2) and transient receptor potential cation channel subfamily M member 2 (TRPM2), which are essential for Ca^2+^ ion increases. Analysis of hub genes was achieved using the degree method in cytoHubba where the top 10 down- and upregulated DEGs were identified as hub genes (Figure 3E and G). Enrichment analysis of top 10 hub genes of downregulated DEGs (Figure 3E) through stringApp revealed that these genes are mainly associated with astrocyte differentiation (GO:0048708), regulation of MAPK cascade (GO:0043408) and cell surface receptor signaling pathway (GO:0007166) (Figure 3F), meanwhile the top 10 hub genes of upregulated DEGs (Figure 3G) were associated with regulation of membrane potential (GO:0042391), nervous system process (GO:0050877) and cell-cell signaling (GO:0007267) (Figure 3H). The complete list of hub genes and their associated pathways can be found in Supplementary file 6.

**Figure 3.**
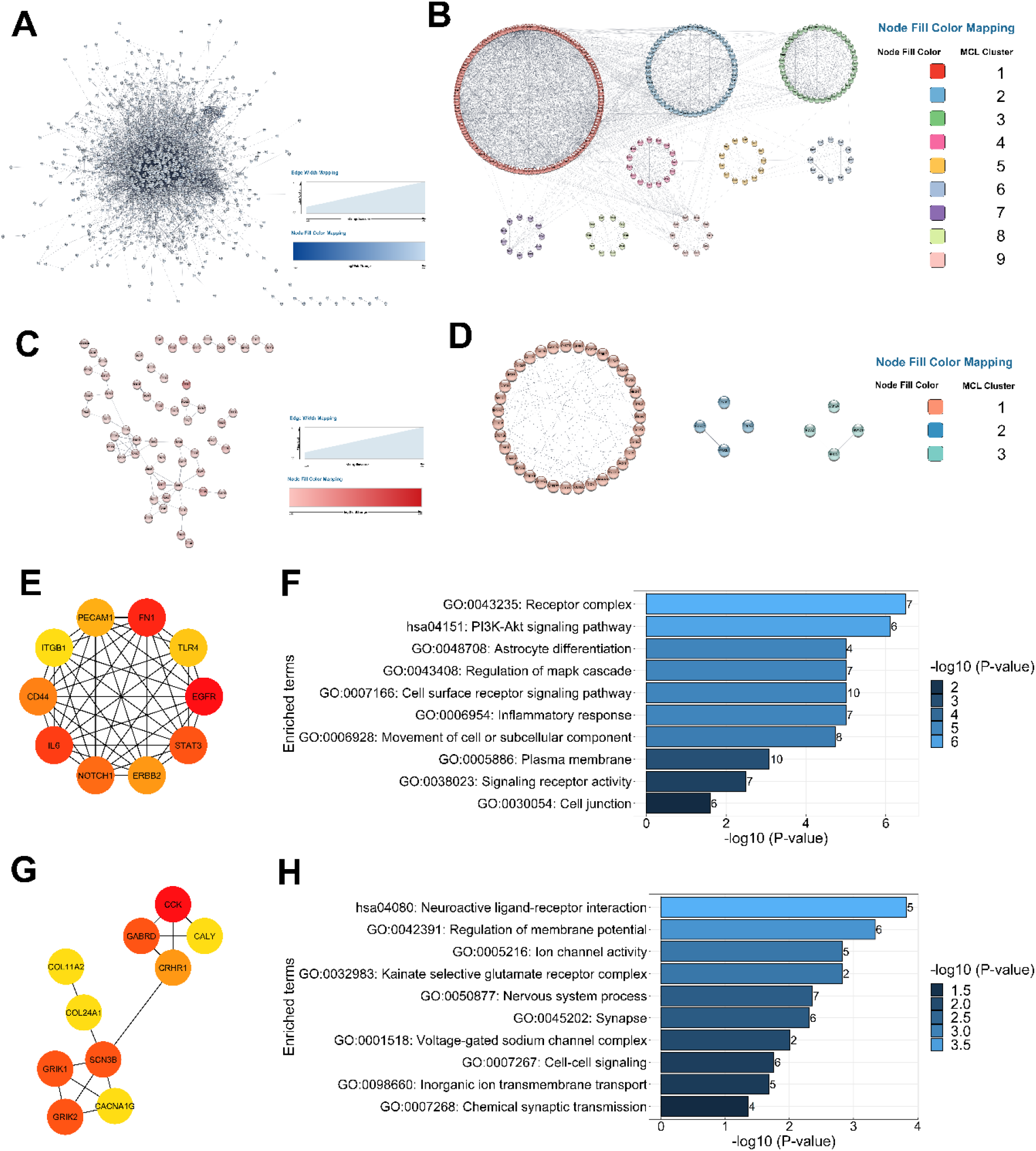
PPI network construction and hub gene screening of down- and upregulated differentially expressed genes. (A, C) PPI networks are constructed using STRING Online Database. (A) PPI network of downregulated genes contains 1234 nodes and 7826 edges, while (C) PPI network of upregulated genes contains 131 nodes and 74 edges. The networks were optimized through Cytoscape software. The color of the nodes represents the fold change of the genes. Upregulated genes (C) are in red color, downregulated genes (A) are in blue color. The deeper the color is, the higher the fold changes are. The width of the edge (grey lines) represents stringdb score between two genes. The thickness of the lines represents the strength of data supporting a protein-protein interaction. The PPI enrichment *p*-value of downregulated genes for the number of identified edges compared to the expected number (4413) was < 1.0 × 10^−16^, significantly more than expected with a medium interaction score (0.4). Likewise, the PPI enrichment *p*-value of upregulated genes for the number of identified edges compared to the expected number (24) was < 1.0 × 10^−15^, significantly more than expected with a medium interaction score. (B, D) Clustering PPI networks with Markov Clustering algorithm (MCL) using Cytoscape software. (B) represents the first 9 functional clusters of PPI network of downregulated genes. For clarity, clusters with less than 10 genes are not shown. Clusters with 10 or more genes were functionally annotated. (D) represents the 3 main clusters of PPI network of upregulated genes. Clusters were functionally annotated using the srtingApp plugin. (E, G) The top 10 ranked hub genes of PPI networks were obtained from CytoHubba analysis based on degree method. (E) represents the top 10 hub genes of downregulated genes and (G) represents the top 10 hub genes of upregulated genes. The change in color from red to yellow represents a change in degree score from high to low. (F, H) Gene Ontology (GO) functional and KEGG pathway classification of top 10 hub genes analyzed through stringApp with the FDR-corrected *p*-value < 0.05. (F) shows the enriched terms of top 10 hub genes of downregulated genes, while (H) shows enriched terms of top 10 hub genes of upregulated genes. On the graph, y-axis represents the significantly enrichment terms. Each bar describes the number of mapped annotated genes in the reference data set while the x-axis and color gradient indicate the significance (-log10 *p*-value) in each category.

### Validation of RNA-seq data

To verify changes in gene expression associated with suicidal behavior we performed qRT-PCR. We have chosen fifteen functionally relevant genes from the DEGs, which have been potentially implicated in depressive behavior based on the literature: GRIK1, GRIK2, GRM2, NRGN, SYT5 and NECAB2 as upregulated genes and AQP1, ITPKB, ITGB4, EPHA2, SLCO2B1, GJA1, PRKCH, GLUL and S100B as downregulated genes. As shown in Supplementary figure 5, results of quantitative PCR analysis of the selected genes confirmed our RNA-seq results. Thus, we verified the increased expression of GRIK1, GRIK2 and GRM2 involved in glutamate signaling, as well as NECAB2, while the reduced expression of AQP1 and astrocyte-related genes such as S100B and GJA1, the growth factor receptor EPHA2 and the integrin ITGB4 genes in suicide victims (Table 1).

**Table 1.**
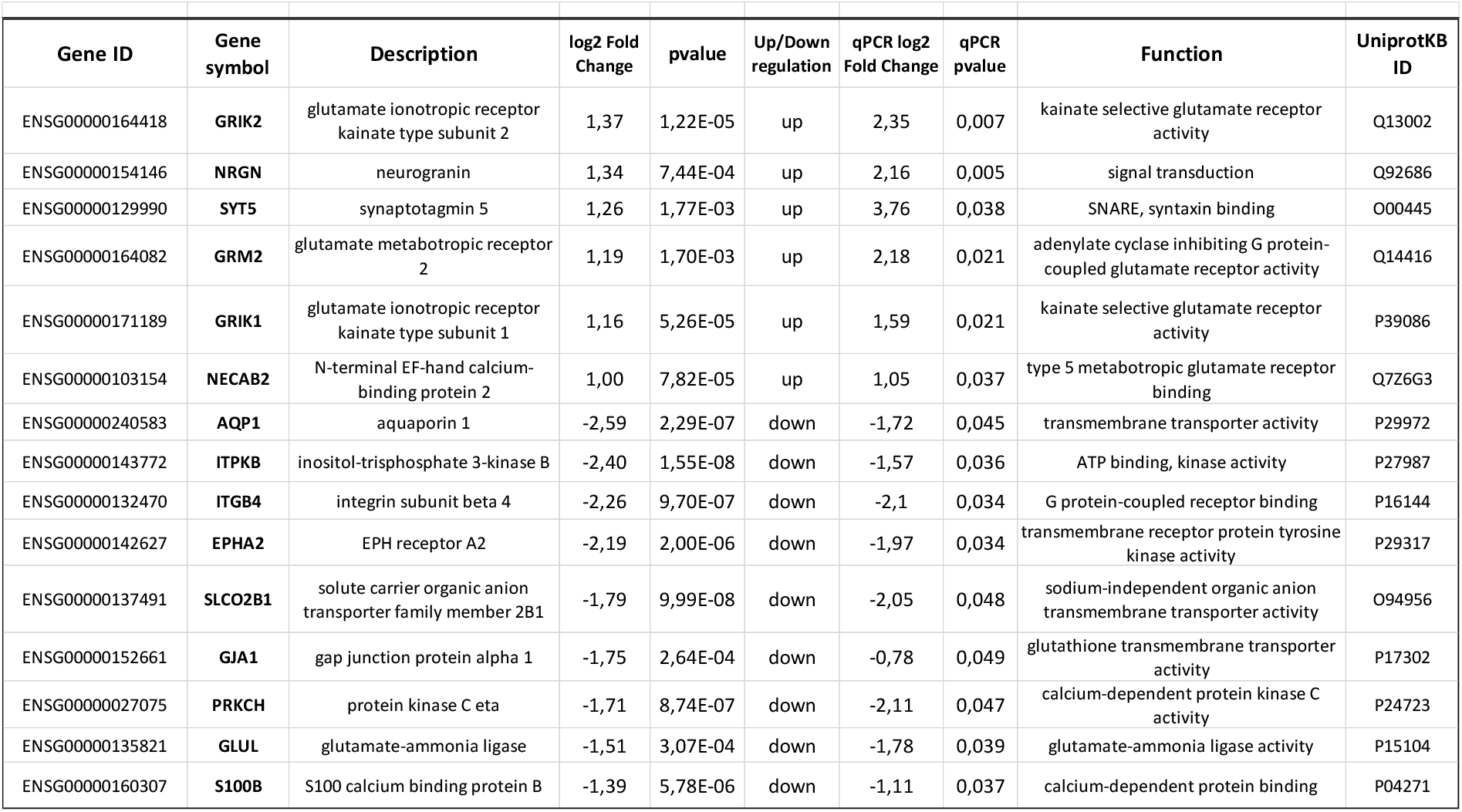
Validation of DEGs with qRT-PCR. Based on the RNA-seq analysis, 15 differentially expressed genes were selected. For validation, quantitative real-time polymerase chain reaction (qRT-PCR) was used, and the results showed significantly altered expression of all genes in the expected direction based on RNA-seq.

### Depression focused gene set enrichment

The top 10 down- and upregulated DEGs were investigated in the DisGeNET database for searching significant enrichments associated with depression and other mental disorder using the disease classification terms ‘Mental Disorders” and “Behavior and Behavior Mechanisms”. We found 30 enriched categories associated with more than 1 gene. We found 10 genes which have been previously associated with depression including CSF3, IL1R2, IL6, SERPINA3, MT-ND5, MT-ND6, P2RX2, ATP4, PROC and CARTPT. Genes associated with depressive disorder, schizophrenia and impaired cognition were strongly expressed in top DEGs in the DMPFC (Supplementary file 7). Although the database does not contain information specific to DMPFC, our data suggest that these genes are altered in this brain region, too.

### Co-expression network analysis in the DMPFC of suicide individuals compared to controls

We built two independent gene co-expression networks for down- and upregulated genes to identify gene co-expression communities (Figure 4 and Supplementary file 8). Analysis was conducted using genes with correlation > 0.9 and Padj < 0.01. In the networks, the DEGs were represented by nodes, and pairwise co-expression relationships between genes by edges. The co-expression network of downregulated DEGs revealed the top 10 highest ranked hub genes as SORBS3, RHOC, S100A16, SZRD1, TRIP6, AHNAK, GRIN2C, CHDH, MAPKAPK2 and FTL with higher degree and betweenness centrality than others, indicating a more critical role played by them in the network (Supplementary file 8). The network was further subdivided into 21 functional clusters composed of a total of 710 downregulated DEGs (nodes) surrounding their hub proteins (Figure 4A). There was a significant difference in the biological processes as different modules were enriched (Figure 4B and C). For example, the biggest module Cluster 1 (yellow) was significantly enriched using biological terms of the GO classification such as “gliogenesis” (GO:0042063) and “response to cholesterol” (GO:0070723). The second biggest module, Cluster 2 (red) was significantly enriched using biological terms such as “regulation of epithelial cell differentiation” (GO:0030856) and “type I interferon signaling pathway” (GO:0060337) (Figure 4B). The co-expression network of upregulated DEGs revealed the top 10 most ranked hub genes as CALY, SOHLH1, CPNE9, NRGN, MAST1, GRIK1, SYT5, TRPM2, BX276092.9 and MYO15A (Supplementary file 8). The network was further subdivided into 6 functional clusters of a total of 97 upregulated DEGs surrounding the hub proteins (Figure 4A), where Cluster 1 (blue) was significantly enriched using biological terms “neurotransmitter receptor internalization” (GO:0099590) and “biotin metabolic process” (GO:0006768). Cluster 2 (red) was significantly enriched using biological terms, such as “sodium channel activity” (GO:0005272) and “neuropeptide hormone activity” (GO:0005184) (Figure 4C). The hub proteins of clusters were specifically identified as most ranked, based on their degree and betweenness centrality. For instance, the hub protein of downregulated co-expression network in Cluster 1 is the SORBS3 encoding vinexin protein, which is implicated in promoting upregulation of actin stress fiber formation. An additional hub protein of downregulated network in Cluster 2 is RHOC encoding the Rho-related GTP-binding protein which is known to regulate signal transduction pathway linking plasma membrane receptors to actin stress fibers. The hub protein of upregulated co-expression network in Cluster 1 is CALY encoding the neuron-specific vesicular protein calcyon, which interacts with clathrin light chain A and stimulates clathrin-mediated endocytosis. Another hub protein from upregulated network in Cluster 2 is SOHLH1 encoding the spermatogenesis- and oogenesis-specific basic helix-loop-helix-containing protein 1, which was implicated in transcriptional regulation of both male and female germline differentiations (Figure 4A). The entire co-expression networks and the annotation information of the top 3 clusters are available in Supplementary file 8.

**Figure 4.**
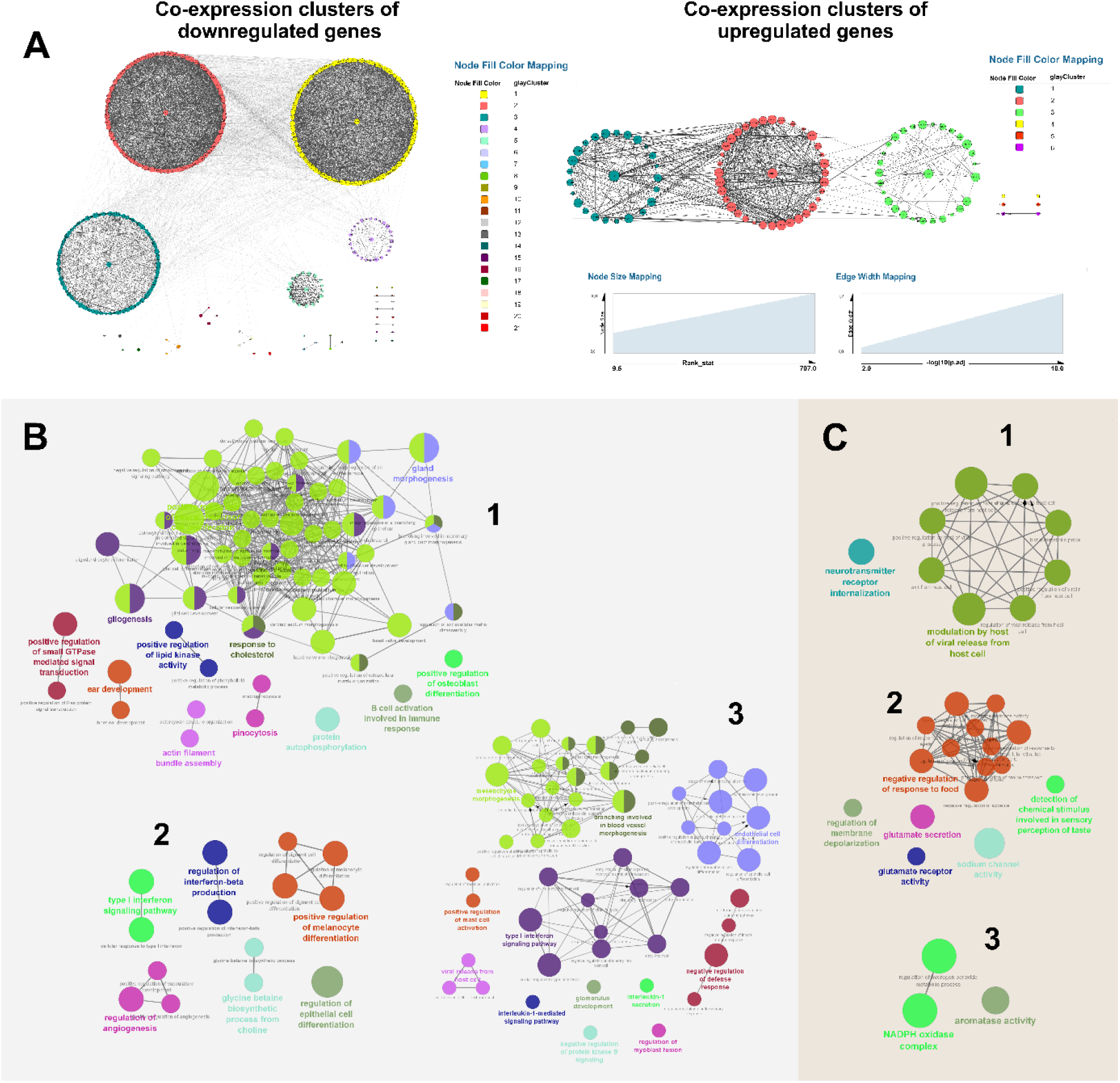
Co-expression network analysis and functional annotation of DEGs. (A) Co-expression networks of down- and upregulated genes constructed using DEGs with absolute correlation ≥ 0.9 and adjusted *p*-value ≤ 0.01 by Cytoscape. For analysing functional modules, a Cytoscape plugin, Community Clusters GLay clustering algorithm with the default options was used for clustering networks. Node size is represented according to their rank (Node Size Mapping). The width of the edge represents significance of the correlation between the two genes (Edge Width Mapping). The larger the width is, the smaller the Padj is. Circle layout is represented a hub gene placed at the center of the main clusters. (B and C) Visualization of statistically overrepresented GO terms in a given set of genes from the three main co-expression network clusters (1-3) using ClueGO plugin. Functional annotation of DEGs using ClueGO enriched by downregulated (B) and upregulated DEGs (C). Each node refers to a specific Gene Ontology (GO) Biological process term and are grouped based on the similarity of their associated genes. The size of the nodes is equivalent with the statistical power of significance of each term: the larger the node size is, the smaller the *p*-value is. Node colors represent different functional groups. The emphasized term of each functional group is given by the most significant term of the group.

### Distribution of NECAB2 in the DMPFC

As a functionally intriguing validated DEG, NECAB2 was selected for analysis of its distribution within the DMPFC. *In situ* hybridization histochemistry was applied in a control and a suicide individual to localize the expression of NECAB2 mRNA. The hybridization signal was abundant in layers II-VI (Figure 5A and B). NECAB2 immunolabeled cells showed the same distribution (Figure 5C). Immunolabeling also revealed that NECAB2 is present in two different interneuron subtypes. Most of the NECAB2-positive interneurons had small size cell body, while some of them demonstrated large soma (Figure 5C3). The larger interneurons were located in deep layers, meanwhile the smaller ones were distributed in upper layers, predominantly in layer II (Figure 5C, C1 and C3). Immunolabeling also revealed that NECAB2 is present not only in the somatodendritic compartments but also in axon terminals in neurons (Figure 5C2-3). To determine whether NECAB2 protein is also present in glial cells we performed combined immunolabeling for Iba1 (a microglia marker) and S100B (an astrocyte marker) with NECAB2. The lack of colocalization indicates that NECAB2 is not expressed in glial cells (Figure 5D-F).

**Figure 5.**
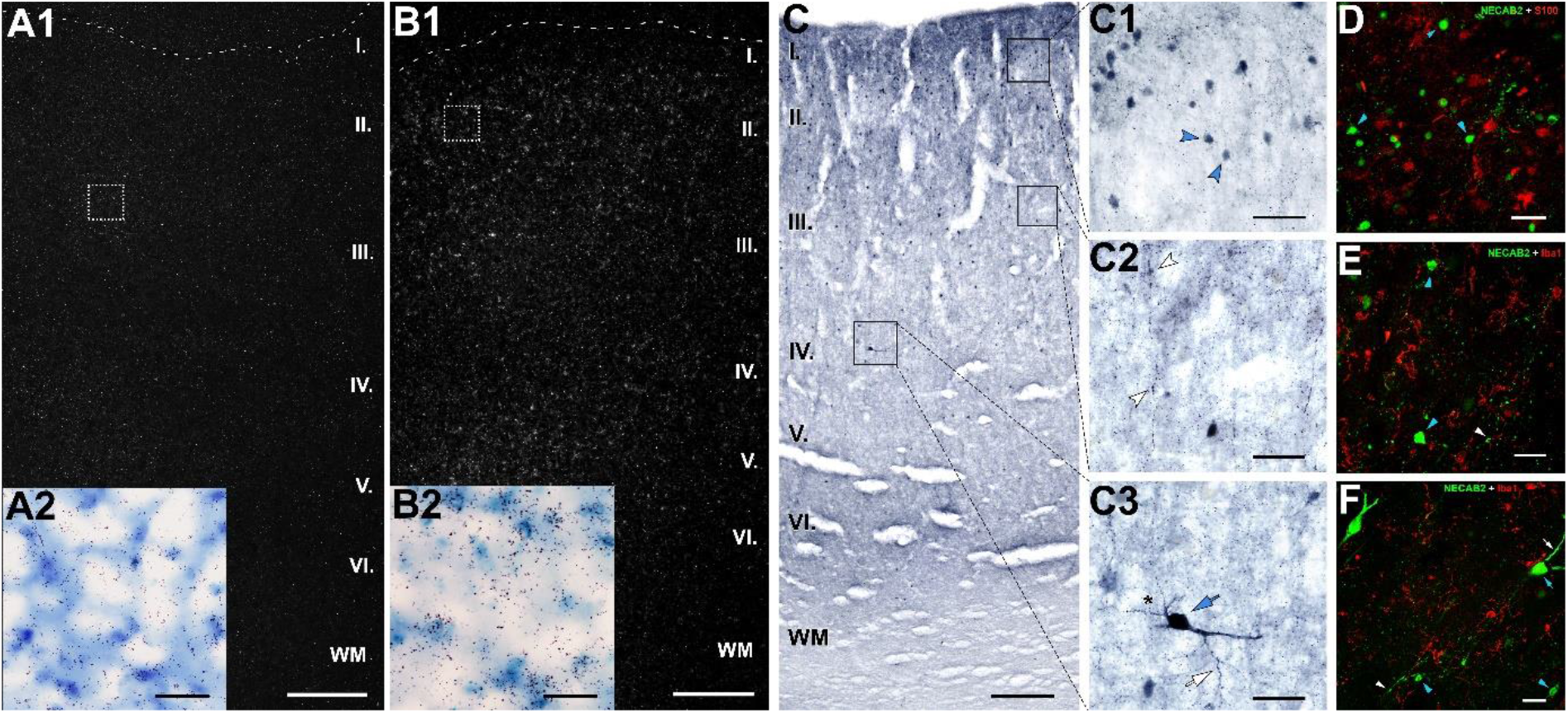
Characterization of NECAB2-immunoreactive (NECAB2-ir) neurons in the DMPFC. (A-B) NECAB2 mRNA expression in the DMPFC of control (A1) and suicide (B1) individuals visualized by *in situ* hybridization histochemistry. A dark-field photomicrograph shows high intensity of NECAB2 hybridization signal in the DMPFC. NECAB2 signal is prominent in cortical layers II and IV in the DMPFC of suicide victims (B1) as compared to the representative control (A1). High magnification bright-field microphotographs of tissue sections (A2 and B2) show the area assigned by the dashed boxes in A1 and B1 and developed for silver grains in the control (A2) and suicide (B2) victims. Cortical layers are indicated on the left. (C) Laminar distribution of NECAB2-ir neurons across cortical layers in the left DMPFC of a female control individual (62 years old) visualized by DAB immunostaining. NECAB2-ir neurons are mainly located in II-V layers and as it is shown on C1 and C3, NECAB2 is located in at least 2 different interneuron subtypes based on the shape and size of labeled neurons. (C1-3) Higher magnification of the boxed area in C. (C1) NECAB2-ir neurons are presented in cortical layer II of the DMPFC (blue arrowhead). (C2) NECAB2-ir axons give rise to varicosities (white arrowheads) along the section. (C3) A higher magnification photomicrograph of a layer IV NECAB2-immunopositive interneuron shows the soma (blue arrow), the varicose dendritic tree (asterisk) and a part of the axon (white arrow). Cortical layers are indicated on the left. (D-F) Representative confocal microscopy images of double immunolabeling of Iba1 and S100B glial markers (red) with NECAB2 (green) in DMPFC control sections. (D) High-magnification confocal image of a section double-labeled with S100B (red) indicates that NECAB2 (green) does not colocalize with S100-positive astrocytes. (E-F) Likewise, NECAB2-positive cells and Iba1-positive microglia do not have overlapping distributions. Scale bar represents 500 µm in A1-B1, 250 µm in C, and 50 µm in A2-B2, C1-3 and D-F.

## DISCUSSION

The underlying pathophysiological basis of suicidal behavior remains basically obscure. Neuroimaging studies revealed that committing suicide, similar to MDD, may be a neural network-level disturbance (Hamilton, Chen, and Gotlib 2013, Drevets, Price, and Furey 2008). The default mode network (DMN) has been highlighted in neuroimaging studies, since its discovery has grown the interest due to its implications in MDD and suicide-related behavior. The DMN is involved through convergent findings of increased resting-state functional connectivity between two core DMN regions, namely the dorsomedial prefrontal cortex (DMPFC) and precuneus (Sheline et al. 2010). Even though the DMPFC is a major component of the DMN, no attention has been assigned to examine transcriptome changes in this region in suicide victims or depressed patients (Cabrera et al. 2019, Pantazatos et al. 2017, Romero-Pimentel et al. 2021). We present here the results of RNA-seq analyses in the DMPFC from sixteen subjects, eight with fatal suicide action and eight controls. The number of samples in both groups are sufficiently high to conclude significant results and draw proper conclusions (Schurch et al. 2016). We identified more than 1000 differentially expressed genes (DEGs). The number of DEGs is in line with previous studies investigating suicide victims (Cabrera et al. 2019) and schizophrenic individuals (Ramaker et al. 2017). In our study, a high number of genes decreased its expression level. Since our RNA-seq data fit for the DESeq2 model (Supplementary figure 4), technical issues are not likely. Therefore, we conclude that suicidal behavior is accompanied with a generally reduced gene expression in the DMPFC.

### Conclusions based on individual DEGs

Among the DEGs the top 10 downregulated genes were involved in immune response, neurodegeneration, mitochondrial electron transport and cell adhesion. The role of immune dysfunction in suicide victims has been reported in the prefrontal cortex (Pantazatos et al. 2017) and insula (Jabbi et al. 2020). It has also been proposed that neurodegeneration and impaired structural neuroplasticity (Jabbi et al. 2020, Underwood and Arango 2011), as well as mitochondrial dysfunction (Punzi et al. 2022, Cabrera et al. 2019) are associated with suicide completion. The top 10 upregulated genes were involved in ATP signaling, protein and vesicle-mediated transport, component of membrane and cytoskeleton and regulating protein secretion. Changes in protein and vesicle-mediated transport in suicide have also been observed previously (Jabbi et al. 2020, Sequeira et al. 2009, Pantazatos et al. 2017).

Among the top DEGs, 10 genes have been previously associated with depression, including CSF3, IL1R2, IL6, SERPINA3, MT-ND5, MT-ND6, P2RX2, ATP4, PROC and CARTPT, based on the DisGeNET database (Pinero et al. 2017). Studies of genetic polymorphisms in psychiatric disorders have shown that CARTPT gene mutation exhibits increased anxiety and depression (Miraglia del Giudice et al. 2006). Furthermore, the peptide encoded by CARTPT is a candidate biomarker for MDD because of its effects on mood regulation (Ahmadian-Moghadam, Sadat-Shirazi, and Zarrindast 2018, Wiehager et al. 2009).

In addition to the DEGs with the highest fold changes, we also focus on the individually validated DEGs. For qRT-PCR validation we selected 15 genes whose alterations were reported in human depression and suicide or rodent depression-like behavior. For ITPKB, EPHA2, GJA1 and GLUL genes, the previously reported alteration in the DLPFC was demonstrated to also occur in the DMPFC. For AQP1 and ITGB4, studied only in rodent models of depression, the present study was the first to implicate them in human suicidal behavior.

Several genes connected to astrocyte function, such as ATP1A2, ALDH1L1, GFAP, S100B, GJA1 and AQP1 showed decreased expression levels in suicide victims. These observations suggest that genes promoting astrocyte functions may be involved in the pathophysiology of depression and suicidal behavior as shown previously for depression (Cotter et al. 2002, Öngür, Drevets, and Price 1998, Rajkowska et al. 1999).

### Conclusion based on the distribution of NECAB2

NECAB2 is one of the validated genes whose level increased in suicide victims. It was shown to bind specifically to the type 5 metabotropic glutamate receptor (mGluR5) to modulate its function (Canela et al. 2009). Our findings, consistent with a previous study (Zhao et al. 2018), provided evidence that the glutamatergic signaling system is involved in the pathogenesis of suicidal behavior. Therefore, it is likely that NECAB2 exerts its role on suicidal behavior by increasing the activity of mGluR5. The Allen Human Brain Atlas suggests that NECAB2 is expressed in interneurons located in layers II-VI in human cerebral cortex. The majority of NECAB2 expressing cells are GABAergic interneurons belonging to the vasoactive intestinal peptide, somatostatin, cholecystokinin and lysosome-associated membrane glycoprotein 5 expressing interneurons (Supplementary file 10). Our *in situ* hybridization and immunohistochemistry study confirmed that NECAB2 is not expressed in glial cells. We found that NECAB2 was presented in the cell body of two morphologically different interneuron subtypes in the DMPFC. The interneurons with larger cell body were located in deep layers, while the smaller, more abundant cell types were distributed in upper layers, mainly in layer II. Based on the distributional and morphological data, the larger cell type may correspond to SST+ interneurons while the smaller one to VIP+ interneurons (Mazuir, Fricker, and Sol-Foulon 2021, Zhu et al. 2018). The presence of NECAB2 in somatodendritic compartments as well as in axon terminals in DMPFC interneurons is consistent with previous data in the hippocampus (Miczán et al. 2021). Based on these data, it is likely that at least one of the cell types expressing NECAB2 is implicated in suicidal behavior possibly by the regulation of mGluR5.

### Functional implications based on pathway analyses

Our integrated analyses using GO functional annotation, as well as KEGG and Reactome pathways revealed a number of pathways altered in suicide victims. The downregulated DEGs were significantly enriched in cell surface receptor signaling pathway of biological process and growth factor binding of molecular function, in PI3K-Akt signaling pathway, in TNF signaling pathway and in cytokine–cytokine receptor interaction. Alterations in inflammatory responses have been related to mental illnesses (Dowlati et al. 2010, Farooq et al. 2017, Kitagishi et al. 2012). However, our study is the first to suggest that suicidal behavior is linked to reduced inflammatory ability in the DMPFC suggesting reduced microglial function in suicidal behavior.

Upregulated DEGs were enriched in axons and synapses as cell components, in synaptic function as biological process and in sodium channel activity as molecular function. Pathways found among upregulated DEGs were neuroactive ligand-receptor interactions and activation of Na-permeable kainate receptor pathways, which all suggest an increased glutamatergic, specifically kainatergic signaling. This change had been implicated in mood disorders (Zarate et al. 2003, Nagy et al. 2015), addictive disorders (Li, Mao, and Wei 2008) and in alcohol dependence (Biernacka et al. 2013). Further evidence from postmortem studies revealed higher expression levels of the majority of glutamatergic genes in the DLPFC associated with MDD and suicide (Gray et al. 2015, Sequeira et al. 2009), and now we provide evidence that it also holds for the DMPFC. Target genes of high interest included GRIK2, which likely plays a role in emerging suicidal thoughts after antidepressant treatment (Laje et al. 2007). In a subsequent study, polymorphisms in GRIK2 gene were further associated with suicidal ideation in MDD patients following antidepressant treatment (Menke et al. 2008). A rodent study showed that GRIK2 deficient mice were more impulsive and aggressive, suggesting that it has a unique role in controlling the behavioral symptoms of mania (Shaltiel et al. 2008). In particular, the outcome of completed suicide has been associated with increased expression of GRIK2 (Punzi et al. 2022). The present data further suggest that elevated kainate signaling in the DMPFC can predispose the development of committing suicide and hence can be a potential biomarker.

### Functional cluster analysis of gene expressions in the DMPFC

Co-expression analysis on the DEGs was performed to identify the key modules of highly co-expressed genes. The largest cluster in downregulated gene network was associated with gliogenesis while another cluster was associated with type-I interferon signaling pathway suggesting reduced glial function and the role of interferon-gamma in reduced inflammatory response ability. In turn, modules in upregulated gene networks were associated with neurotransmitter receptor internalization, sodium channel activity and peptide neuromodulator function. Alteration in neurotransmitter receptor internalization pathway is consistent with the assumption that dysfunction of neurotransmitter receptors may contribute to the pathophysiology of MDD (Cannon et al. 2009, Kang et al. 2012, Nagy et al. 2015).

Intramodular hubs have an important role in the enriched pathways of the modules in co-expression networks. The major hub gene of Cluster 1 in the downregulated gene network was SORBS3 (vinexin). This protein modulates actin cytoskeleton, and negatively regulates autophagy. Its expression increases with age in mouse and human brain tissue contributing to autophagic decline in mammalian brain ageing (Kioka, Ueda, and Amachi 2002, Park et al. 2021). Moreover, a meta-analysis provided evidence that SORB3 has strong relationship with MDD (Howard et al. 2019).

The hub gene from upregulated gene network, calcyon neuron-specific vesicular protein (CALY) regulates dopamine-related signaling (Li et al. 2011) and dopamine D1 receptor internalization of pyramidal cells in the prefrontal cortex and dorsal striatum (Ha et al. 2012). Other studies reported that CALY has trafficking functions primarily involved with neural development and synaptic plasticity (Chander et al. 2019, Davidson et al. 2009). These results suggest that intramodular hub genes could function as potential biomarkers for future therapeutic interventions.

### Functions supported by known PPIs of DEGs

The clustering result of MCL on PPI network revealed major clusters of downregulated DEGs involved in receptor and cytokine-mediated signaling pathways, glutamate metabolism and mitochondrial respiration. The major clusters of upregulated DEGs were associated with short-term glutamate receptor signaling and regulation of cytosolic calcium ion concentration. Analysis of top 10 downregulated hub genes (PECAM1, ERBB2, ITGB1, EGFR, STAT3, NOTCH1, CD44, IL6, TLR4 and FN1) from PPI network revealed that these genes were strongly enriched in cell-surface receptor signaling pathway, astrocyte differentiation and regulation of MAPK cascade, suggesting that these pathways might be implicated in suicidal behavior. As previously defined, two of the top hub genes, the adhesive stress-response protein PECAM1 and the proto-oncogene ERBB2 were reported decreasing in the DLPFC in suicide victims (Cabrera-Mendoza et al. 2020). Analysis of top 10 upregulated hub genes (CCK, GABRD, CRHR1, CALY, COL11A2, COL24A1, SCN3B, GRIK1, GRIK2, CACNA1G) revealed that these genes were associated with regulation of membrane potential, nervous system processes and cell-cell signaling. A pilot study examining the GABAergic system in suicide victims found GABRD variants in the prefrontal cortex of patients associated with depressive disorder (Yin et al. 2016). Recently, a strong association between CRHR1 polymorphism and suicide attempts was shown (De la Cruz-Cano 2017). Löfberg et al. (1998) found that MDD patients associated with the number of previous suicide attempts had higher CCK levels. A more recent study showed that CCK levels were higher among first suicide attempters after the first 12 hours following the attempt (Jahangard et al. 2018). Likewise, a study investigated postmortem brain tissues of completed suicides observed higher number of CCK receptors in the frontal cortex compared to healthy controls (Harro, Marcusson, and Oreland 1992).

A multi-locus GWAS study demonstrated that several allelic variations in glutamatergic synaptic signaling, such as GRIK1, GRIK2, GRIK3, GRIN1, GRIN2A, GRIN2C or GRM7 were associated with MDD (Lee et al. 2012). Moreover, several studies found decreases in mRNA expression of NMDA and AMPA receptor subunits associated with schizophrenia in frontal cortex (Scarr et al. 2005, Sokolov 1998, Weickert et al. 2013). The involvement of glutamate in psychiatric and medical conditions has been intensively examined, however, earlier studies mostly focused on the biology and pathophysiology of ionotropic glutamate receptors. In turn, metabotropic receptors (mGluR) can also modify neuronal activity through G-protein coupled signaling. Indeed, strong interactions have been reported between mGluR5 and NMDA receptors suggesting that mGluR5 might be implicated in mediating neural plasticity as well as learning and memory processes (Cleva and Olive 2011). Two proteins from upregulated DEGs, the CACNG8 and CALY are considered as neurotransmitter receptor regulatory proteins. CALY regulates dopamine D1 receptor internalization in the prefrontal cortex and dorsal striatum (Ha et al., 2012), and CACNG8 (also known as TARP-γ8) negatively modulate AMPA receptor signaling (Maher et al., 2016).

In summary we performed a comprehensive transcriptome study of DMPFC in suicide victims, using RNA-seq and observed significant associations between specific transcriptome changes in DMPFC and suicidal behavior. They include but are not limited to the suppression of genes, that regulate inactivation of immune responses and glial cell differentiation, and upregulation of genes involved in glutamatergic synaptic transmission. However, in this study gene expression alterations may reveal dysregulated pathways and functional cascades causative for the pathophysiology of suicidal behavior, we conclude that the DEGs and pathways identified in this study are causally linked to depression. Our results suggesting that the activation or inhibition of these processes may be a common denominator of suicide behavior in the DMPFC. Taken together, these findings may contribute to the pathophysiology of depressive or suicidal behavior. We conclude that our study provides new insights into the genomic underpinnings of major depressive disorder and may allow for novel precision therapeutic development strategies targeting depression to be implemented in future studies.

## MATERIALS AND METHODS

### Human brain tissue samples

Human brain samples were collected in accordance with the Ethical Rules for Using Human Tissues for Medical Research in Hungary (HM 34/1999) and the Code of Ethics of the World Medical Association (Declaration of Helsinki). Tissue samples were taken during brain autopsy at the Department of Forensic Medicine of Semmelweis University in the framework of the Human Brain Tissue Bank (HBTB), Budapest. The activity of the HBTB has been authorized by the Committee of Science and Research Ethic of the Ministry of Health Hungary (ETT TUKEB: 189/KO/02.6008/2002/ETT) and the Semmelweis University Regional Committee of Science and Research Ethic (No. 32/1992/TUKEB). The study reported in the manuscript was performed according to protocol approved by the Committee of Science and Research Ethics, Semmelweis University (TUKEB 189/2015). The medical history of the subjects was obtained from clinical records, interviews with family members and relatives, as well as from pathological and neuropathological reports. All personal data are stored in a strict ethical control, and samples were coded before the analyses of tissue.

### Tissue preparation for RNA sequencing

Postmortem human brain tissue samples from the dorsomedial prefrontal cortex (DMPFC) (Brodmann area 9) were acquired from the Human Brain Tissue Bank (Semmelweis University, Budapest, Hungary). There is considerable variability in how the topographical extensions of the DMPFC is defined. The medial prefrontal cortex (MPFC), as its name indicates, occupies the medial surface of the frontal lobe over the anterior cingulate gyrus. The MPFC is divided into dorsal and ventral prefrontal cortex with a theoretical horizontal line through the most rostral (genual) point of the corpus callosum. (These two parts are frequently called anterior and posterior prefrontal cortex.) The posterior border of the DMPFC is defined by the precentral sulcus which separates it from the premotor cortex. The border of the dorsomedial and dorsolateral prefrontal cortex in coronal sections can be defined by the deep superior frontal sulcus (Supplementary figure 2). The white matter also serves as an excellent landmark: the superior longitudinal cortical pathway, as a well visible white matter bundles is divided into a medial and a lateral portion in the dorsal prefrontal cortex. The cortical samples were collected from 16 subjects: 8 control subjects with no history of psychiatric or neurological diseases (2 females and 6 males, mean age of 65.4 ± 5.6) and 8 suicide victims (3 females and 5 males, mean age of 53.6 ± 4.8). The age of the subjects ranged from 31 to 89 years (Supplementary file 1). The selected control subjects were not diagnosed with any psychiatric disorder while the suicide victims without evidence of acute or chronic depression. The exact time of death, the time of autopsy and the removal of the brain were known and recorded for all subjects. The medical history of the victims was obtained from medical or hospital records, interviews with family members and relatives, as well as from pathological and neuropathological reports while also from forensic records in the case of suicide victims. Brains were removed from the skull with a postmortem delay of 1 to 10 hours, frozen rapidly on dry ice and stored at −80°C until microdissection. Serial coronal sections were cut from the frontal lobe. Sections between 50.0 and 30.0 mm from the origin of AC-PC coordinates (see Mai and Majtanik 2017) (practically at the first appearance of the genu corporis callosi or the first appearance of the anterior pole of the lateral ventricle) were punched out. Special microdissection needles with 8 and 15 mm inside diameters were used (Palkovits 1973, Palkovits 1985). Tissue pellets (1-3 per brain) include samples from the superior frontal gyrus (Brodmann area 9) (both gray and white portion within the gyrus) and from the paracingulate cortex (also called as the pre- and supragenual dorsal part of the anterior cingulate cortex, Brodmann area 32), the upper bank and the deep portion of the gyrus (Supplementary figure 2). In samples, the superior frontal gyrus/paracingulate gyrus ratio were about 80-85%, respectively. The microdissected tissue pellets were collected in 1.5 ml Eppendorf tubes and kept until use at −80°C. During each step of the microdissection procedure (slicing, micropunch, storage) the tissue samples were kept frozen.

### RNA sequencing analysis and data

We performed RNA sequencing on cortical samples to determine RNA expression changes in the dorsomedial prefrontal cortex related to suicidal behavior. Sample preparation, sequencing library construction and RNA sequencing were conducted by the Beijing Genomics Institute’s.(BGI, Hongkong, Shenzhen, China) standard procedure, To extract total RNA, 20 mg of postmortem brain tissue was subjected to RNA sequencing on the BGISEQ-500 platform (BGI) using the 100-bp paired-end sequencing strategy. RNA size, concentration and integrity were verified using Agilent 2100 Bioanalyzer (Agilent Technologies). All the generated raw sequencing reads were filtered using SOAPnuke (1.5.2), a filter software developed by BGI company, to remove reads containing adapters, reads in which unknown bases are more than 5%, and low-quality reads (> 20% of the bases with a quality score lower than 15). After filtering, the remaining clean reads were obtained and stored in FASTQ format (Cock et al. 2010). HISAT2 (2.0.4) software in combination with the Bowtie index (2.2.5) (Langmead and Salzberg 2012, Kim, Langmead, and Salzberg 2015) were used to map clean reads to the human reference genome (hg19), respectively. The average mapping ratio with genes was 76.12%, and the average mapping ratio with reference genome was 91.28% (Supplementary file 2). The output of HISAT2 was imported into the RStudio environment (R version 4.0.4; RStudio version 1.4.1106) using Rsubread package (2.4.3), which converted data from the transcript level to the gene level, then aligned to the human genome (GRCh38) and counted into genes focusing on all annotated protein coding genes. The Rsubread facilitates the RNA-seq read data analyses, producing the quality assessment of sequence reads, read alignment and read summarization among others (Liao, Smyth, and Shi 2013). To identify differentially expressed genes (DEGs), DESeq2 package (1.34.0) as the standard workflow was used for detection of DEGs (Love, Huber, and Anders 2014). Before using DESeq2, we performed a minimal pre-filtering to keep the rows that have at least one read total. To increase power, stricter filtering is automatically applied via independent filtering on the mean of normalized counts within the *results* function. The complete dataset of DEGs is listed in Supplementary file 3. The DEGs were identified by Benjamini and Hochberg-adjusted *p*-value (*p*adj < 0.05) (Benjamini and Hochberg 1995) and log2 FC ratio ≥ ± 1 was defined as the thresholds to discriminate significant DEGs. DEGs were visualized in a volcano plot created with the R package ggplot2 (3.3.5). Hierarchical clustering of DEGs were analyzed using the *heatmap.2* function in the gplots (3.1.1) package for R. Heatmap was created by two color arrays. Pathway-based and co-expression analysis helps to further understand genes biological functions.

### Gene ontology and pathway enrichment analysis of DEGs

To assess the function of the DEGs, functional enrichment analysis was performed using the gprofiler2 R package (version 0.2.0). The Gene Ontology (GO) analysis was queried using the *gost* function in “ordered” mode (Kolberg et al. 2020). The differentially expressed genes were ranked by the adjusted *p*-value significance (*p*.adjust < 0.05). Functional enrichment was performed for upregulated and downregulated DEGs separately. The Kyoto Encyclopedia of Genes and Genomes (KEGG) and Reactome resources were used for functional annotation and pathway analysis (Kanehisa and Goto 2000, Jassal et al. 2020). The results of the functional enrichment were visualized in Manhattan plots created with the gprofiler2. The significant GO terms were listed in Supplementary file 4. The functional enrichment analysis of the top 10 down- and upregulated genes was carried out using Database for Annotation, Visualization and Integration Discovery (DAVID) v6.8 online server to annotate gene-related biological mechanisms using standardized gene terminology.

### Protein-protein interaction network construction and hub gene screening

As an alternative approach to protein-protein interaction (PPI) network analysis, STRING version 11.5 (Szklarczyk et al. 2019) was used to perform gene clustering with a Markov clustering (MCL) algorithm with a minimum required interaction score of 0.4 and the inflation factor set at 2.5. The results were then uploaded to Cytoscape software version 3.8.2 (Smoot et al. 2011). Downregulated clusters containing at least 10 genes and three out of twelve upregulated clusters were functionally annotated with GO and KEGG pathway using stringApp plugin in Cytoscape. CytoHubba (Chin et al. 2014), a Cytoscape Plugin, was used to evaluate PPI network hub genes. The top 10 hub genes were screened out by node degree from both down- and upregulated gene networks. Red colors denoted the higher degree of a gene. Gene list enrichments of the top 10 hub genes were identified through the stringApp. Functional annotation terms were considered significantly enriched with an FDR-corrected *p*-value < 0.05.

### Disease-associated gene sets

Human disease-associated gene sets were obtained from the DisGeNET database, which is known to integrate disease-gene links from several sources (Pinero et al. 2017). The list of top 10 down- and upregulated genes was imported into the DisGeNET database browser (http://www.disgenet.org/), then the curated gene-disease association file was downloaded. Keywords with special emphasis on mood disorders and associated comorbidities were used for filtering categories to gather those validated genes which involved in the pathophysiology of depression. The result was imported and listed in Supplementary file 7.

### Co-expression network construction and functional annotation

Coexpression correlation was computed using the Pearson correlation coefficient as described previously (Contreras-López et al. 2018). Read counts of each gene and each sample were normalized by median normalization using the EBSeq R package (Leng et al. 2013). The correlation (Pearson’s) and correlation significance of every pair DEG (for downregulated genes adjusted *p*-value < 0.01, for upregulated genes adjusted *p*-value < 0.05) was calculated using logarithmic matrix of read counts with the psych R package (Revelle 2017). A network table containing the statistically significant correlations across the whole data set for every pair of DEGs was generated, and for calculating network statistics, igraph R package (Csardi and Nepusz 2006) was used. The network statistics (degree and betweenness centrality for each node) were uploaded to Cytoscape. The gene symbols were designated as the identifiers of nodes. The correlation, degree and betweenness were mapped to the edge color, edge width and node size respectively. Regarding the topological analysis, Markov Cluster Algorithm (MCL) clustering evaluation (Van Dongen 2008) was performed in order to identify densely connected nodes. The functional annotation analysis of the network was performed using the ClueGO application (Bindea et al. 2009). Functional annotation terms were considered significantly enriched with an Bonferroni-corrected *p*-value < 0.05. Further details on the interplay of the reconstructed networks were examined using the stringApp. The network datasets and functional annotation results of the networks were shown in Supplementary file 8.

### Validation of expression changes by qRT-PCR

To confirm the RNA-seq data, a subset of differentially expressed genes whose selection was based on the known or potentially probable involvement in depression and neuron-specific functions were validated by quantitative real time PCR (qRT-PCR). The procedure was carried out as described previously (Dobolyi 2009). Briefly, total RNA was isolated from approximately 20 mg of frozen postmortem brain tissue samples using TRIzol reagent (Invitrogen, Carlsbad, CA, USA) as lysis buffer combined with RNeasy Mini kit (Qiagen, Germany) following the manufacturer’s instructions. The quality and quantity of extracted RNA was determined using NanoDrop ND-1000 Spectrophotometer (Thermo Fisher Scientific, Waltham, MA, USA), and only those with A260/A280 ratio between 1.8 and 2.1 were used in subsequent experiments. The concentration of RNA was adjusted to 500 ng/µL, and it was treated with Amplification Grade DNase I (Invitrogen, Carlsbad, CA, USA). The isolated RNA concentration was calculated and normalized with RNase-free water and reverse transcribed into cDNA using SuperScript II Reverse Transcriptase Kit (Invitrogen, Carlsbad, CA, USA). After 10-fold dilution, 2.5 μl of the resulting cDNA was used as template in PCR performed in duplicates using SYBR Green dye (Sigma, St Louis, MO, USA). The PCR reactions were performed on CFX-96 C1000 Touch Real-Time System (Bio-Rad Laboratories, Hercules, CA, USA) with iTaq DNA polymerase (Bio-Rad Laboratories, Hercules, CA, USA) in total volumes of 12.5 μl under the following conditions: 95°C for 3 min, followed by 35 cycles of 95°C for 0.5 min, 60°C for 0.5 min and 72°C for 1 min. A melting curve was performed at the end of amplification cycles to verify the specificity of the PCR products. The primers used for qRT-PCR were synthesized by Integrated DNA Technologies, Inc., (IDT, Coralville, IA USA) and used at 300 nM final concentration. Sequences of primers are listed in Supplementary file 9. Housekeeping genes ACTB, GAPDH and LDHA were used as internal controls and the relative gene expression values were calculated from their averages using the 2−△△Ct method.

### Preparation of *in situ* hybridization probes

A mixture of cDNAs prepared with RT-PCR from the human DMPFC was used as template in PCR reactions with primers for NECAB2 (CAGGATCTTGGTGCCAGCT and TGTGGTCAGTGTGGGTCATG) yielding a probe that corresponds to the GenBank accession number NM_019065.2. The purified PCR products were applied as templates in a PCR reaction with the primer pairs specific for the probe and also containing T7 RNA polymerase recognition site added to the reverse primers. Finally, the identities of the cDNA probes were verified by sequencing them with T7 primers.

### *In situ* hybridization histochemistry

Two fresh-frozen DMPFC brain blocks of subjects were used: one from a 75 years old female control subject with negative clinical reports for any major diseases and one from a 72 years old female suicide individual. Using a cryostat, serial coronal sections (12 μm) were cut and mounted on positively-charged slides (SuperfrostPlus®, Fisher Scientific), dried and stored at −80°C until use. Further steps were performed according to the procedure described previously by Dobolyi et al. (2015). Briefly, antisense [35S]UTP-labeled riboprobes were generated from the above-described DNA probes using T7 RNA polymerase of the MAXIscript Transcription Kit (Ambion, Austin, TX, USA) and used for hybridization at 1 million DPM (discharges per minute) activity per slide. Washing procedures included a 30 min incubation in RNase A followed by decreasing concentrations of sodium-citrate buffer (pH 7.4) at room temperature and subsequently at 65 °C. Following hybridization and washes, slides were dipped in NTB nuclear track emulsion (Eastman Kodak) and stored at 4°C for 3 weeks for autoradiography. Then, the slides were developed and fixed with Kodak Dektol developer and Kodak fixer, respectively, counterstained with Giemsa and coverslipped with Cytoseal 60 (Stephens Scientific).

### Tissue collection for immunolabeling

Immunohistochemistry was used to assess the distribution of NECAB2 protein in the DMPFC. For immunolabeling, one DMPFC brain block from a 62 years old female control individual was cut into 10 mm thick coronal slice and immersion fixed in 4% paraformaldehyde in 0.1 M phosphate-buffered saline (PBS) for 5 days. Subsequently, the block was transferred to PBS containing 0.1% sodium azide for 2 days to remove excess paraformaldehyde. Then, the block was placed in PBS containing 20% sucrose for 2 days cryoprotection. The block was frozen and cut into 60 μm thick serial coronal sections on a sliding microtome. Sections were collected in PBS containing 0.1% sodium azide and stored at 4 °C until further processing.

### DAB immunolabeling

Free-floating DMPFC brain sections were immunolabeled for NECAB2 (Thermo Fisher Scientific, Cat. No. PA5-53108). The antibody (1:250 dilution) was applied for 24 h at room temperature, followed by incubation of the sections in biotinylated anti-rabbit secondary antibody (1:1,000 dilution, Vector Laboratories, Burlingame, CA) and then in avidin–biotin-peroxidase complex (1:500, Vector Laboratories) for 2 h. Subsequently, the labeling was visualized by incubation in 0.02% 3,3-diaminobenzidine (DAB; Sigma), 0.08% nickel (II) sulfate and 0.001% hydrogen peroxide in PBS, pH 7.4 for 5 min. Sections were mounted, dehydrated and coverslipped with Cytoseal 60 (Stephens Scientific, Riverdale, NJ, USA).

### Double labeling of NECAB2

Double immunofluorescence staining was used to clarified the colocalization of NECAB2 in the DMPFC. To reduce autofluorescence, tissue sections were treated with 0.15% Sudan Black B (in 70% ethanol) after antigen retrieval (0.05 M Tris buffer, pH = 9.0) procedures. Slides were blocked by incubation in 3% bovine serum albumin (with 0.5% Triton X-100 dissolved in 0.1 M PB, Sigma) for 1 h at room temperature, followed by washing with washing buffer (10 min × 3). NECAB2 was immunolabeled as for single labeling using 1:250 dilution except for the visualization, which was performed with fluorescein isothiocyanate (FITC)-tyramide (1:8,000 dilution) and H_2_O_2_ (0.003%) in 100 mM Trizma buffer (pH 8.0 adjusted with HCl) for 6 min. Subsequently, sections were placed in goat anti-ionized calcium-binding adapter molecule 1 (Iba1) (1:500 dilution, Abcam, Cat. No. ab107159), and mouse anti-S100 (1:250 dilution, Millipore, Cat. No. MAB079-1) for 24 h at room temperature. The sections were then incubated in Alexa 594 donkey anti-goat/mouse secondary antibody (1:500 dilution, Vector Laboratories) followed by 2 h incubation in a solution containing avidin–biotin–peroxidase complex (ABC, 1:300 dilution, Vector Laboratories). Finally, all sections with fluorescent labels were mounted on positively charged slides (Superfrost Plus, Fisher Scientific, Pittsburgh, PA) and coverslipped in antifade medium (Prolong Antifade Kit, Molecular Probes).

### Microscopy and photography

Sections were examined using an Olympus BX60 light microscope also equipped with fluorescent epi-illumination and a dark-field condensor. Images were captured at 2048 × 2048 pixel resolution with a SPOT Xplorer digital CCD camera (Diagnostic Instruments, Sterling Heights, MI, USA) using a 4× objective for dark-field images, and 4–40× objectives for bright- field and fluorescent images. Confocal images were acquired with a Zeiss LSM 70 Confocal Microscope System using a 40-63X objectives at an optical thickness of 1 µm for counting varicosities and 3 µm for counting labeled cell bodies. Contrast and sharpness of the images were adjusted using the ‘levels’ and ‘sharpness’ commands in Adobe Photoshop CS 8.0. Full resolution was maintained until the photomicrographs were cropped at which point the images were adjusted to a resolution of at least 300 dpi.

### Statistical analysis

Demographics were contrasted using the Chi-Square test or Welch’s unequal variances t-test (p<0.05). The plot and heatmap were generated using R version 4.0.4 (https://www.r-project.org/). Quantitative statistics of qPCR results were performed with the GraphPad Prism version 8.0.1 (GraphPad Software). Normality distribution was assessed via Shapiro-Wilk test. For comparisons between values of the two groups, Welch’s unequal variances t-test was applied. The nonparametric Mann-Whitney U test was used for data that were not normally distributed. Differences were considered statistically significant when *p* < 0.05 and plotted as mean ± S.E.M (standard error of the mean).

## Author Contributions

**Fanni Dóra**: Conceptualization, Methodology, Validation, Formal analysis, Investigation, Resources, Data Curation, Writing - Original Draft, Writing - Review & Editing, Visualization **Éva Renner**: Methodology, Investigation, **Dávid Keller**: Methodology, Investigation, **Miklós Palkovits**: Conceptualization, Writing - Original Draft, Writing - Review & Editing, Supervision, Project administration, Funding acquisition, **Arpád Dobolyi**: Conceptualization, Methodology, Resources, Writing - Original Draft, Writing - Review & Editing, Supervision, Project administration, Funding acquisition

## Data availability

The data presented in this study are available in the supplementary material of this article.

## Acknowledgements

We appreciate the technical assistance of Szilvia Deák and Viktória Dellaszéga-Lábas.

## SUPPLEMENTARY MATERIAL

**Supplementary figure 1.**
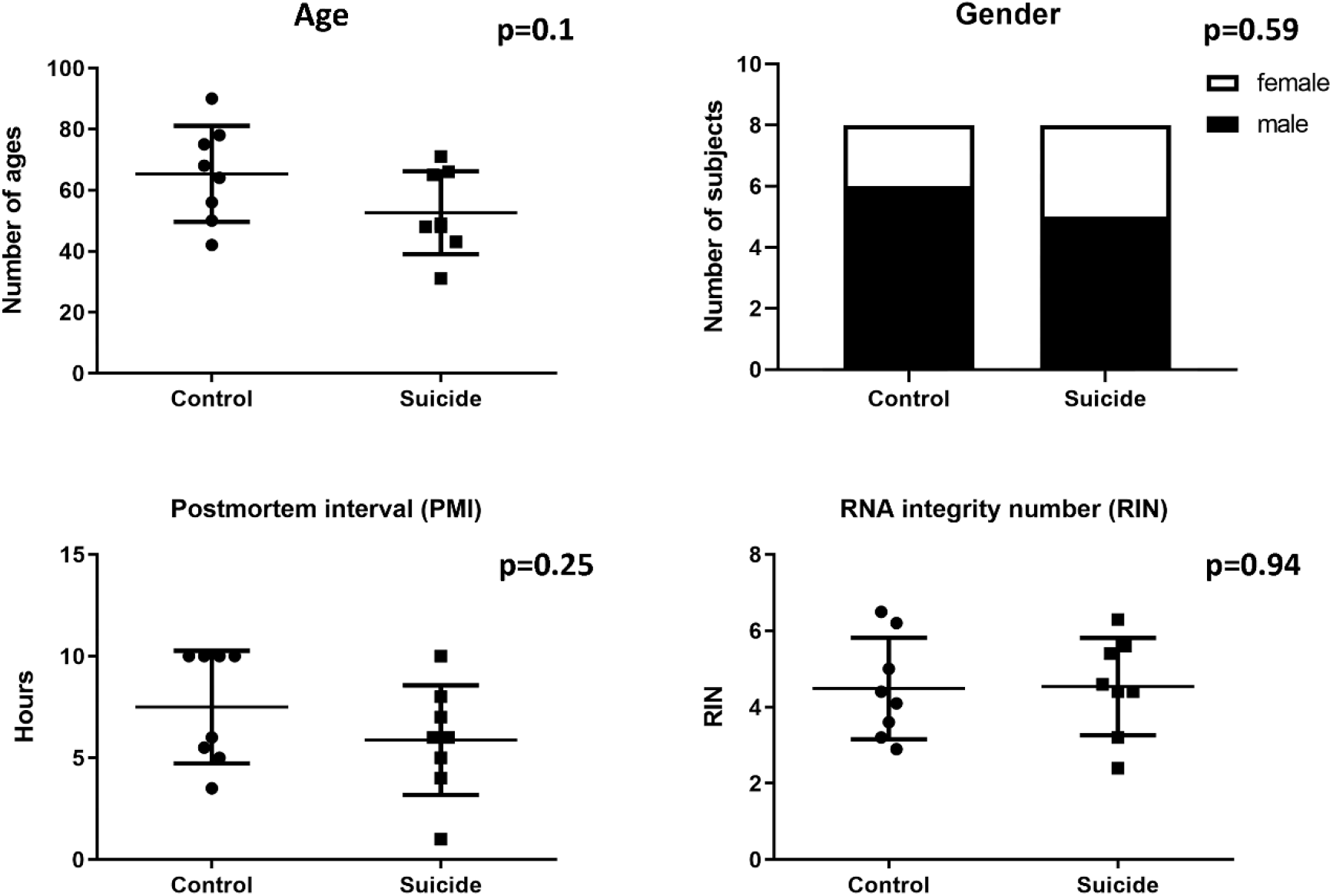
Characteristics of samples. Mean age, gender, PMI and RIN proportions of samples in control (n=8) and suicide (n=8) groups. There were no significant differences between the groups for any covariates tested by Welch’s unequal variances t-test (age, PMI, RIN) and Chi-square test (gender). (Age: p=0.1; Gender: p = 0.59; PMI: p = 0.25; RIN: p = 0.94)

**Supplementary figure 2.**
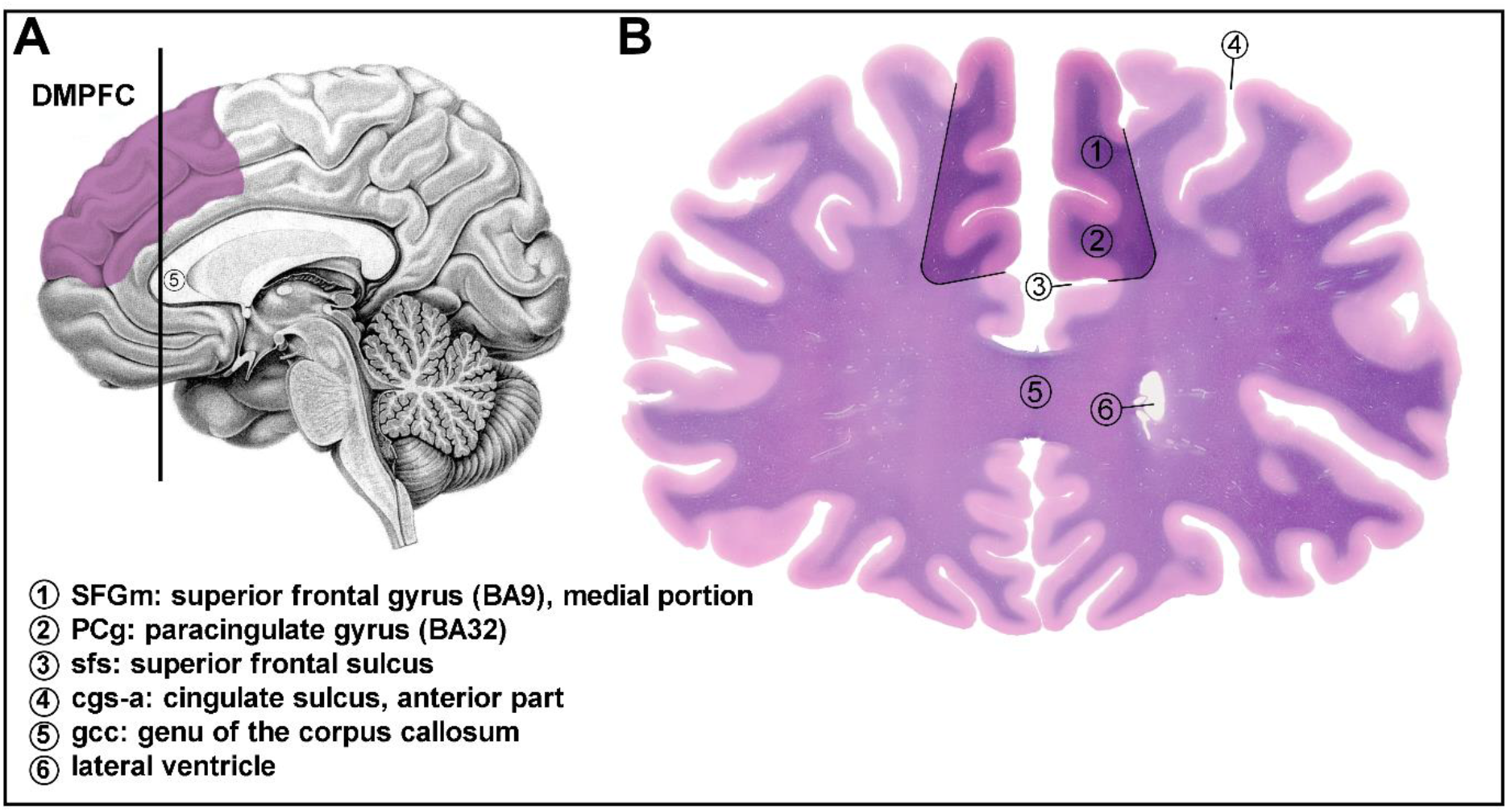
Representative view of human brain region dissected in the study. Details on the anatomical origin and structure of the dissected region. (A) The location and topographical extension of the dissected area, the dorsomedial prefrontal cortex (DMPFC) is shown by lilac color on the medial surface of human brain. The vertical black line represents the cutting line used to obtain the coronal section illustrated on the right panel. (B) Coronal section of human brain represents the DMPFC (1+2). The dissected area analyzed in this study are highlighted and demarcated by black lines. Landmarks and boundaries are notified by numbers. The section was stained using Levanol-Fast Cyanine 5RN method.

**Supplementary figure 3.**
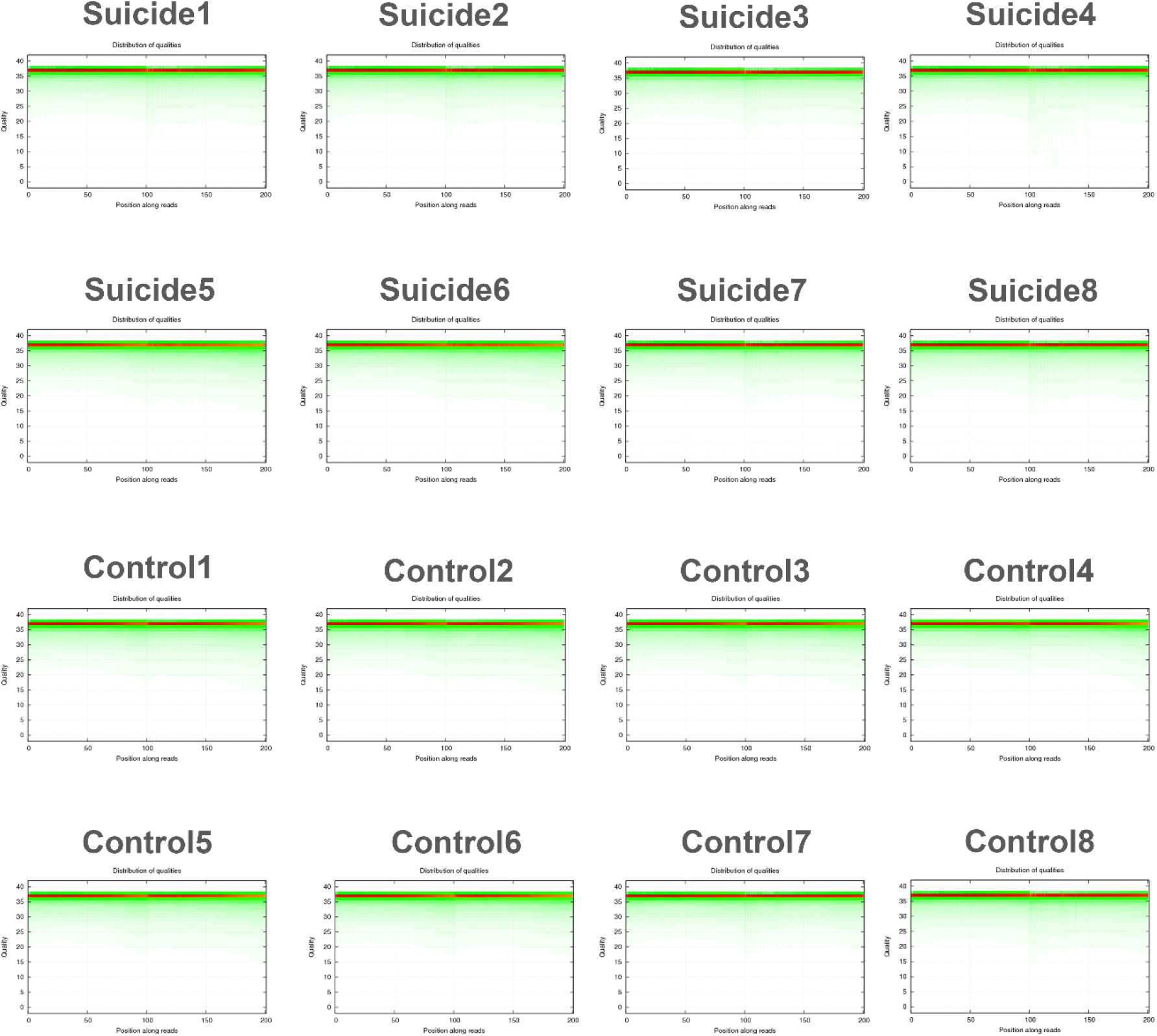
Distribution of base quality on clean reads. Quality control (QC) of RNA-seq dataset showed good sequencing quality reads and congruence of the biological samples. Analysis of the sequenced library showing that each sequenced base had a mean value score of > 30, indicating the good quality sequencing of control and suicide samples. The x-axis represents base positions along reads. The y-axis represents base quality value. Each dot in the image represents the number of total bases with certain quality value of the corresponding base along reads. Darker dot color means greater base number. The proportion of the bases with low quality (< 20) is very low indicating that the sequencing quality of this lane is good.

**Supplementary figure 4.**
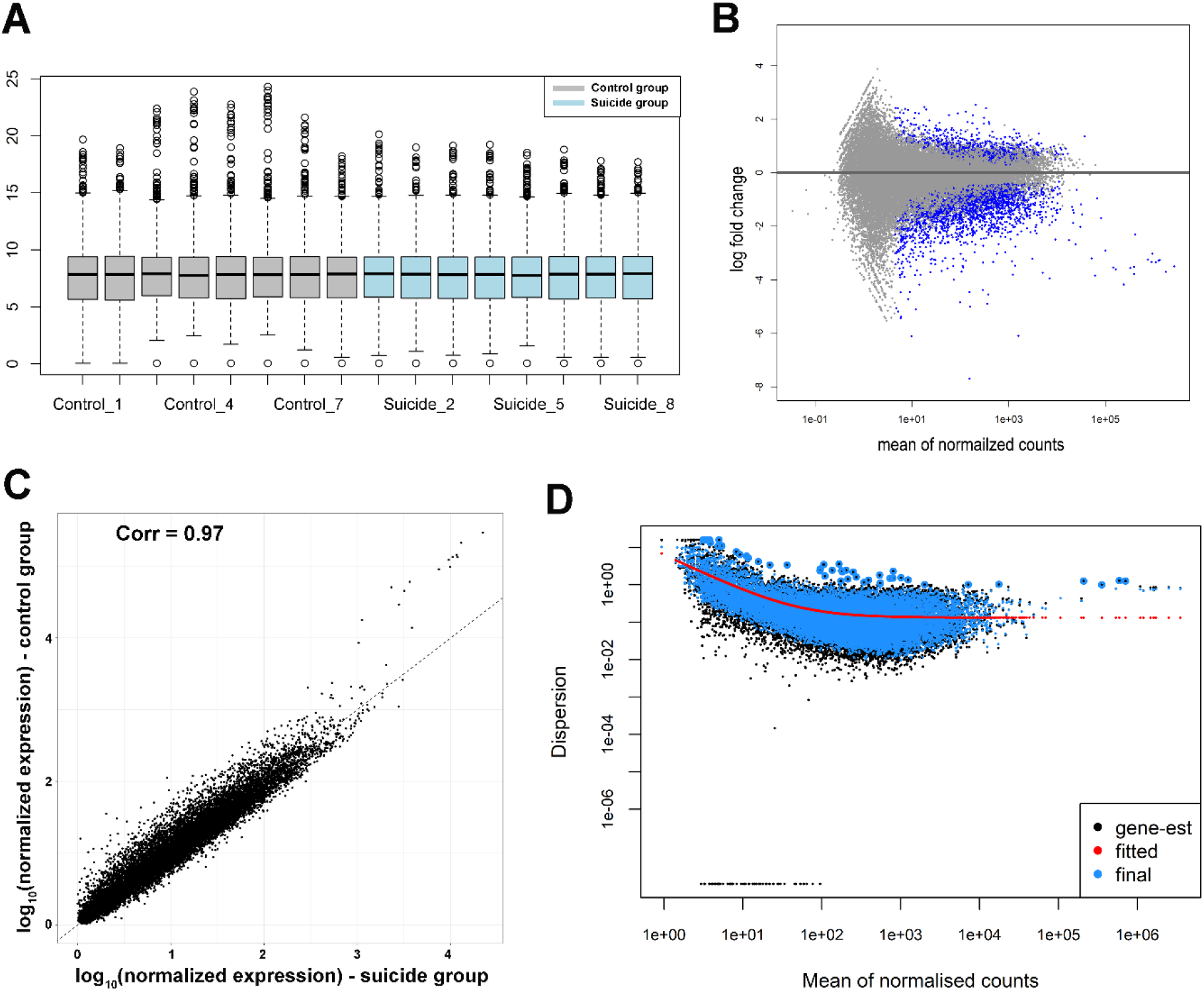
Overview of RNA-seq data. (A) Sample level box plots provided a visual means of comparing the distributions of counts between groups. The line dividing the box represents the median of the data and top and bottom of the box shows the upper and lower quartiles, respectively. The whiskers show the highest and lowest values, excluding the outliers, which are shown as circles. (B) MA plot displays the deferentially expressed genes versus expression strength (log2 fold change) between control and suicide. On the x-axis, the base 2 logarithm of average expression across all samples is plotted versus the base 2 logarithm of fold-change. Points that are significantly different represent the genes either up- or down-regulated with an adjusted *p*-value less than 0.05 are in blue, all others are in gray. (C) Scatter plot is shown comparing the RNA-seq data of the average normalized expression for suicide replicates versus the average normalized expression for control replicates. The correlation coefficient was calculated using R version 4.0.4. (D) Dispersion plot for mean of normalized read counts. The plot provides a visual means of examining dispersion estimates for each gene. The red curve plots the estimate for the expected dispersion value for genes of a given expression strength relative to the mean of the normalized counts. Each black dot is a gene with an associated mean expression level and maximum likelihood estimation (MLE) of the dispersion, while blue dots represent moderated estimates calculated by DESeq2. Black points circled in blue with extremely high dispersion are outliers that are not shrunk towards the curve.

**Supplementary figure 5.**
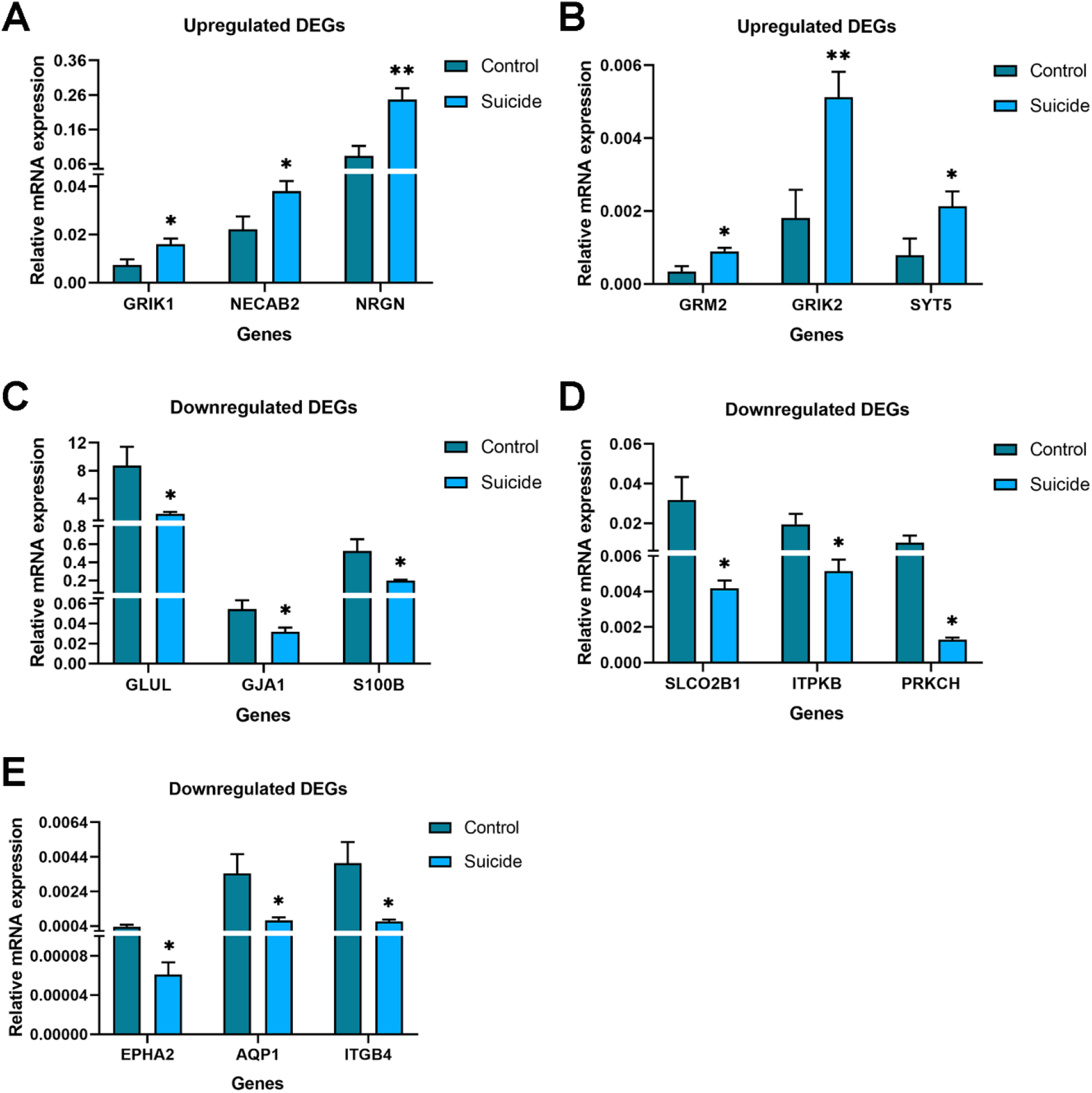
Validation of the RNA-seq results with quantitative PCR. The expression level changes in suicides victims are shown as determined by RT-qPCR. The mRNA expression of upregulated genes is elevated (A and B), while the expression of downregulated genes is reduced (C-E) in suicide victims, which confirm the expressional changes in gene expression profiles deduced from RNA-seq results. Bar graphs represent mean ± SEM of 8 control and 8 suicide individuals (*p < 0.05, **p < 0.01).

**Supplementary figure 6.**
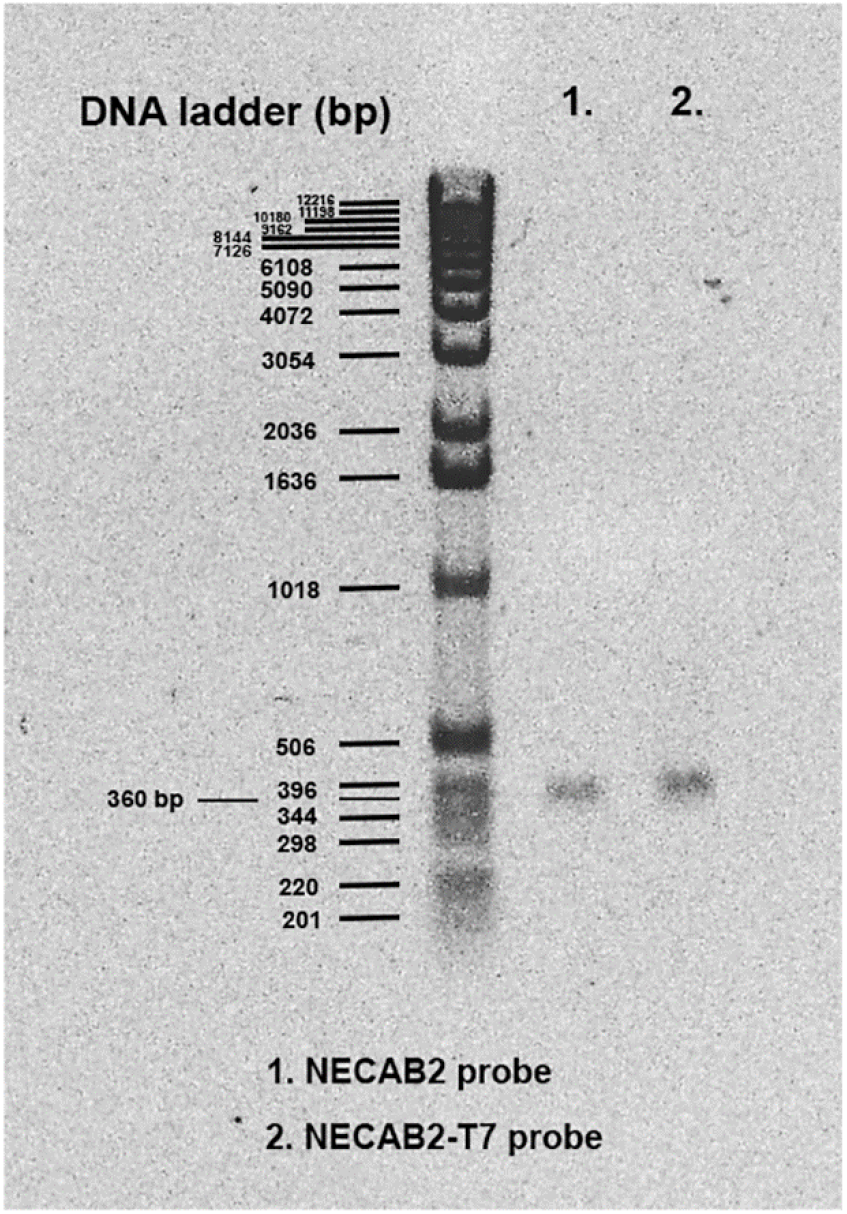
Preparation and visualization of NECAB2 probe in the DMPFC. Two-step polymerase chain reaction (PCR) were performed and PCR products were examined by agarose gel electrophoresis. The products of NECAB2 from the first PCR (1) and products of NECAB2 containing the T7 promoter from the second PCR (2) are indicated. The exact molecular weight of NECAB2 product is 360 bp.

