## Supplementary tables for "Transcriptome profiling of the dorsomedial prefrontal cortex in suicide victims": S1_Demographic_data.pdf

Supplementary file 1.: Demographic data of individuals.

| #Donor | Sex | Age | Post mortem interval (PMI) | RIN | Cause of death | Clinical and pathological diagnosis |
|---|---|---|---|---|---|---|
| #1 | female | 48 | 6-7 h | 6,3 | suicide (drug overdose) | - |
| #2 | male | 71 | 1 h | 3,2 | suicide (jumping from a height) | without any clinical care during the past 6 months |
| #3 | male | 48 | 6 h | 4,4 | suicide (hanging - asphyxia) | without known drug treatment |
| #4 | female | 65 | 5 h | 4,4 | suicide (hanging - asphyxia) | pathological diagnosis: negative status (no pathological sign for any diseases) |
| #5 | male | 31 | 8 h | 2,4 | suicide (hanging - asphyxia) | without known drug treatment |
| #6 | female | 49 | 6 h | 5,4 | suicide (drug overdose) | without known drug treatment |
| #7 | male | 43 | 4 h | 4,6 | suicide (hanging) | without any clinical care |
| #8 | male | 66 | 8-10 h | 5,6 | suicide (hanging - asphyxia) | Laboratory test: alcohol: negative |
| #9 | male | 42 | 3,5 h | 6,5 | acute respiratory insufficiency | - |
| #10 | female | 56 | 6 h | 5 | cardiorespiratory insufficiency, oedema cerebri | oedema cerebri, coarctatio aortae, hepatitis alcoholica |
| #11 | male | 50 | 5,5 h | 4,1 | stroke, brain hemorrhage | large cortical and subcortical hemorrhage in the parietal lobe |
| #12 | male | 68 | 10 h | 2,9 | acute heart failure | acute pulmonary oedema, serious arteriosclerosis (especially in the heart and kidney), periferial arterial shunt, cerebral sclerosis. left coronary occlusion |
| #13 | female | 75 | 10 h | 3,6 | stroke (right side arteria cerebri media) | diabetes, stroke, hypertonia, mamma carcinoma, emolitio arteriae cerebri mediae lateralis dextri, cortical infarction, general atherosclerosis |
| #14 | male | 64 | 10 h | 4,4 | stroke (arteria cerebri media on the left side), bronchopneumonia | cardiomyopathia, coronary sclerosis, hypertonia, infarctus myocardii, bronchopneumonia, cardiorespiratory insufficiency, femoralis amputatio, aphasia, carotis stenosis, pneumonia |
| #15 | male | 90 | 4-5 h | 3,2 | stroke (cerebri media and posterior) | stroke, infarctus lacunaris multiplex cerebri, Parkinson's disease, emolitio, tracheobronchitis, cardiopulmonary insufficiency, carotis stenosis |
| #16 | male | 78 | 10 h | 6,2 | cardiorespiratory insufficiency, stroke | stroke, dementia, diabetes, hypertonia, carotis interna occlusio, polyneuropathia |
