## Supplementary tables for "Transcriptome profiling of the dorsomedial prefrontal cortex in suicide victims": S2_Gene_mapping.pdf

Supplementary file 2: Gene mapping, read data, quality of sequencing.

| Sample | Total Raw Reads (M) | Total Clean Reads (M) | Clean Reads Ratio(%) | Clean Reads Q20 (%) |
|---|---|---|---|---|
| **Suicide_1** | 72,24 | 66,92 | 92,64 | 97,62 |
| **Suicide_2** | 37,18 | 34,31 | 92,29 | 97,73 |
| **Suicide_3** | 75,2 | 70,26 | 93,43 | 97,74 |
| **Suicide_4** | 61,23 | 55,19 | 90,14 | 97,26 |
| **Suicide_5** | 41,98 | 37,87 | 90,21 | 96,89 |
| **Suicide_6** | 72,69 | 68,14 | 93,74 | 97,15 |
| **Suicide_7** | 72,69 | 67,83 | 93,32 | 97,73 |
| **Suicide_8** | 74,79 | 69,53 | 92,96 | 97,68 |
| **Control_1** | 74,2 | 67,04 | 90,36 | 97,19 |
| **Control_2** | 75,78 | 68,31 | 90,14 | 97,06 |
| **Control_3** | 37,53 | 33,71 | 89,83 | 97,27 |
| **Control_4** | 49,23 | 46,85 | 95,15 | 97,85 |
| **Control_5** | 56,51 | 50,2 | 88,82 | 97,27 |
| **Control_6** | 58,45 | 52,75 | 90,25 | 97,41 |
| **Control_7** | 71,61 | 66,66 | 93,1 | 97,96 |
| **Control_8** | 75,2 | 70,3 | 93,49 | 97,59 |

| Clean Reads Q30 (%) | Total Mapping Ratio (%) | Uniquely Mapping Ratio (%) | Total Gene Number | Total Gene Mapping Ratio (%) |
|---|---|---|---|---|
| 90,04 | 92,34 | 74,74 | 22157 | 75,2 |
| 90,35 | 91,75 | 74,9 | 20928 | 72,9 |
| 90,39 | 90,35 | 73,4 | 22294 | 66,1 |
| 89,88 | 89,53 | 72,67 | 21594 | 67,5 |
| 88,62 | 80,85 | 62,02 | 20878 | 41,8 |
| 89,1 | 93,76 | 73,13 | 21694 | 87,6 |
| 90,41 | 93,17 | 76,04 | 22518 | 80,2 |
| 90,2 | 93,58 | 76,4 | 22300 | 83,4 |
| 89,06 | 94,04 | 74,13 | 21492 | 89,8 |
| 88,63 | 93,28 | 73,12 | 21468 | 87,9 |
| 89,21 | 91,63 | 71,81 | 21216 | 75,2 |
| 91,07 | 90,57 | 71,5 | 19122 | 74,9 |
| 89,3 | 90,86 | 67,08 | 19799 | 81,2 |
| 89,63 | 91,26 | 67,97 | 19034 | 83,2 |
| 91,13 | 89,88 | 70,45 | 20935 | 68,2 |
| 89,8 | 93,57 | 76,17 | 22538 | 82,8 |
