## Supplementary tables for "Transcriptome profiling of the dorsomedial prefrontal cortex in suicide victims": S3_List_of_differentially_expressed_genes.pdf

Supplementary file 3: List of differentially expressed genes with a cutoff padj<0,05 and log2FC>±1.

| ID | Gene_symbol | baseMean | log2FoldChange | lfcSE | stat | pvalue | padj |
|---|---|---|---|---|---|---|---|
| ENSG00000108342 | CSF3 | 166,08 | -7,67 | 1,41 | -5,42 | 5,84E-08 | 2,06E-05 |
| ENSG00000115590 | IL1R2 | 5,14 | -5,18 | 1,21 | -4,30 | 1,74E-05 | 8,54E-04 |
| ENSG00000184557 | SOCS3 | 391,12 | -5,06 | 0,84 | -6,01 | 1,81E-09 | 3,85E-06 |
| ENSG00000115602 | IL1RL1 | 106,11 | -4,95 | 1,36 | -3,65 | 2,66E-04 | 4,86E-03 |
| ENSG00000137648 | TMPRSS4 | 4,19 | -4,81 | 1,40 | -3,45 | 5,66E-04 | 7,92E-03 |
| ENSG00000146839 | ZAN | 13,31 | -4,57 | 1,18 | -3,89 | 1,01E-04 | 2,56E-03 |
| ENSG00000198695 | MT-ND6 | 589943,91 | -4,46 | 0,82 | -5,43 | 5,71E-08 | 2,06E-05 |
| ENSG00000136244 | IL6 | 10,79 | -4,26 | 1,17 | -3,62 | 2,89E-04 | 5,08E-03 |
| ENSG00000196136 | SERPINA3 | 343,07 | -4,16 | 0,71 | -5,89 | 3,80E-09 | 5,25E-06 |
| ENSG00000198786 | MT-ND5 | 710966,26 | -4,05 | 0,81 | -4,99 | 6,08E-07 | 1,03E-04 |
| ENSG00000198886 | MT-ND4 | 2498832,28 | -4,03 | 0,65 | -6,20 | 5,62E-10 | 2,33E-06 |
| ENSG00000212907 | MT-ND4L | 209027,22 | -3,98 | 0,81 | -4,90 | 9,63E-07 | 1,36E-04 |
| ENSG00000174898 | CATSPERD | 6,69 | -3,94 | 1,04 | -3,78 | 1,58E-04 | 3,45E-03 |
| ENSG00000153684 | GOLGA8F | 6,92 | -3,87 | 1,12 | -3,46 | 5,38E-04 | 7,64E-03 |
| ENSG00000124253 | PCK1 | 14,05 | -3,86 | 0,97 | -3,98 | 6,88E-05 | 1,99E-03 |
| ENSG00000198840 | MT-ND3 | 537201,00 | -3,83 | 0,64 | -5,96 | 2,55E-09 | 3,85E-06 |
| ENSG00000269720 | CCDC194 | 6,85 | -3,78 | 1,04 | -3,64 | 2,68E-04 | 4,88E-03 |
| ENSG00000165507 | DEPP1 | 1320,75 | -3,78 | 0,63 | -6,01 | 1,81E-09 | 3,85E-06 |
| ENSG00000159167 | STC1 | 44,71 | -3,77 | 0,79 | -4,80 | 1,63E-06 | 1,91E-04 |
| ENSG00000090339 | ICAM1 | 197,94 | -3,74 | 0,68 | -5,54 | 3,06E-08 | 1,41E-05 |
| ENSG00000198804 | MT-CO1 | 3499659,02 | -3,72 | 0,64 | -5,83 | 5,49E-09 | 5,69E-06 |
| ENSG00000173530 | TNFRSF10D | 58,46 | -3,72 | 0,76 | -4,87 | 1,14E-06 | 1,55E-04 |
| ENSG00000198763 | MT-ND2 | 1465435,39 | -3,68 | 0,63 | -5,86 | 4,73E-09 | 5,47E-06 |
| ENSG00000198899 | MT-ATP6 | 1606472,14 | -3,68 | 0,63 | -5,87 | 4,39E-09 | 5,47E-06 |
| ENSG00000269028 | MTRNR2L12 | 125,24 | -3,67 | 0,66 | -5,57 | 2,60E-08 | 1,41E-05 |
| ENSG00000162383 | SLC1A7 | 76,44 | -3,64 | 0,69 | -5,27 | 1,36E-07 | 3,32E-05 |
| ENSG00000105392 | CRX | 6,26 | -3,63 | 1,36 | -2,67 | 7,53E-03 | 4,80E-02 |
| ENSG00000198888 | MT-ND1 | 1720014,21 | -3,62 | 0,63 | -5,74 | 9,45E-09 | 8,25E-06 |
| ENSG00000163734 | CXCL3 | 10,15 | -3,62 | 1,14 | -3,17 | 1,51E-03 | 1,61E-02 |
| ENSG00000198727 | MT-CYB | 1088352,47 | -3,59 | 0,63 | -5,68 | 1,35E-08 | 1,07E-05 |
| ENSG00000205502 | C2CD4B | 33,10 | -3,57 | 0,89 | -3,98 | 6,77E-05 | 1,98E-03 |
| ENSG00000169245 | CXCL10 | 6,71 | -3,49 | 1,30 | -2,69 | 7,20E-03 | 4,66E-02 |
| ENSG00000198712 | MT-CO2 | 1618642,29 | -3,49 | 0,62 | -5,61 | 2,06E-08 | 1,41E-05 |
| ENSG00000125735 | TNFSF14 | 6,74 | -3,49 | 1,27 | -2,75 | 6,04E-03 | 4,11E-02 |
| ENSG00000255823 | MTRNR2L8 | 320,93 | -3,49 | 0,65 | -5,39 | 7,08E-08 | 2,35E-05 |
| ENSG00000214688 | C10orf105 | 169,17 | -3,48 | 0,58 | -5,97 | 2,43E-09 | 3,85E-06 |
| ENSG00000128016 | ZFP36 | 506,47 | -3,47 | 0,65 | -5,38 | 7,45E-08 | 2,42E-05 |
| ENSG00000177464 | GPR4 | 140,29 | -3,41 | 0,57 | -5,97 | 2,36E-09 | 3,85E-06 |
| ENSG00000143546 | S100A8 | 32,51 | -3,39 | 0,77 | -4,43 | 9,64E-06 | 5,84E-04 |
| ENSG00000228253 | MT-ATP8 | 354482,94 | -3,38 | 0,64 | -5,30 | 1,17E-07 | 3,28E-05 |
| ENSG00000205362 | MT1A | 213,99 | -3,36 | 0,86 | -3,89 | 9,97E-05 | 2,54E-03 |
| ENSG00000100336 | APOL4 | 28,40 | -3,35 | 0,81 | -4,11 | 3,99E-05 | 1,40E-03 |
| ENSG00000108691 | CCL2 | 52,52 | -3,33 | 0,60 | -5,54 | 3,00E-08 | 1,41E-05 |
| ENSG00000198938 | MT-CO3 | 1193741,94 | -3,30 | 0,60 | -5,50 | 3,74E-08 | 1,59E-05 |
| ENSG00000093009 | CDC45 | 32,45 | -3,29 | 0,87 | -3,79 | 1,53E-04 | 3,37E-03 |
| ENSG00000170421 | KRT8 | 23,49 | -3,18 | 0,86 | -3,70 | 2,19E-04 | 4,27E-03 |
| ENSG00000124762 | CDKN1A | 611,07 | -3,15 | 0,56 | -5,60 | 2,12E-08 | 1,41E-05 |
| ENSG00000070915 | SLC12A3 | 5,42 | -3,13 | 1,07 | -2,94 | 3,32E-03 | 2,74E-02 |
| ENSG00000109321 | AREG | 5,00 | -3,12 | 1,16 | -2,68 | 7,36E-03 | 4,73E-02 |
| ENSG00000106366 | SERPINE1 | 109,80 | -3,11 | 0,95 | -3,28 | 1,04E-03 | 1,23E-02 |
| ENSG00000198959 | TGM2 | 910,97 | -3,10 | 0,52 | -5,99 | 2,12E-09 | 3,85E-06 |
| ENSG00000117501 | MROH9 | 4,39 | -3,09 | 1,10 | -2,81 | 4,95E-03 | 3,58E-02 |
| ENSG00000064886 | CHI3L2 | 30,78 | -3,06 | 0,72 | -4,25 | 2,15E-05 | 9,44E-04 |
| ENSG00000187922 | LCN10 | 39,11 | -3,05 | 0,77 | -3,98 | 6,83E-05 | 1,98E-03 |

| | | | | | | | |
|---|---|---|---|---|---|---|---|
| ENSG00000131095 | GFAP | 106411,60 | -3,05 | 0,48 | -6,39 | 1,68E-10 | 2,33E-06 |
| ENSG00000128342 | LIF | 17,15 | -3,02 | 1,01 | -2,97 | 2,97E-03 | 2,53E-02 |
| ENSG00000166523 | CLEC4E | 6,96 | -2,98 | 1,04 | -2,85 | 4,31E-03 | 3,27E-02 |
| ENSG00000127533 | F2RL3 | 11,17 | -2,96 | 0,86 | -3,44 | 5,84E-04 | 8,11E-03 |
| ENSG00000162366 | PDZK1IP1 | 6,09 | -2,95 | 0,83 | -3,54 | 3,96E-04 | 6,31E-03 |
| ENSG00000175591 | P2RY2 | 27,44 | -2,95 | 0,58 | -5,06 | 4,13E-07 | 8,14E-05 |
| ENSG00000188282 | RUFY4 | 6,31 | -2,93 | 0,85 | -3,46 | 5,44E-04 | 7,70E-03 |
| ENSG00000072952 | IRAG1 | 1490,29 | -2,87 | 0,50 | -5,72 | 1,08E-08 | 8,96E-06 |
| ENSG00000012223 | LTF | 15,99 | -2,86 | 0,62 | -4,62 | 3,91E-06 | 3,53E-04 |
| ENSG00000126262 | FFAR2 | 4,73 | -2,84 | 0,97 | -2,93 | 3,38E-03 | 2,77E-02 |
| ENSG00000134531 | EMP1 | 108,18 | -2,83 | 0,56 | -5,07 | 3,96E-07 | 8,03E-05 |
| ENSG00000137507 | LRRC32 | 325,33 | -2,82 | 0,51 | -5,55 | 2,85E-08 | 1,41E-05 |
| ENSG00000168309 | FAM107A | 25868,98 | -2,80 | 0,45 | -6,21 | 5,28E-10 | 2,33E-06 |
| ENSG00000077238 | IL4R | 218,78 | -2,79 | 0,57 | -4,92 | 8,58E-07 | 1,28E-04 |
| ENSG00000152049 | KCNE4 | 51,84 | -2,79 | 0,55 | -5,05 | 4,36E-07 | 8,22E-05 |
| ENSG00000163638 | ADAMTS9 | 155,69 | -2,79 | 0,52 | -5,33 | 9,94E-08 | 2,91E-05 |
| ENSG00000104537 | ANXA13 | 3,93 | -2,77 | 0,86 | -3,23 | 1,24E-03 | 1,39E-02 |
| ENSG00000149573 | MPZL2 | 18,41 | -2,75 | 0,74 | -3,75 | 1,80E-04 | 3,78E-03 |
| ENSG00000144837 | PLA1A | 42,82 | -2,73 | 0,64 | -4,25 | 2,11E-05 | 9,40E-04 |
| ENSG00000196954 | CASP4 | 60,25 | -2,72 | 0,58 | -4,69 | 2,70E-06 | 2,69E-04 |
| ENSG00000163739 | CXCL1 | 16,53 | -2,72 | 1,01 | -2,70 | 6,92E-03 | 4,54E-02 |
| ENSG00000163220 | S100A9 | 26,57 | -2,69 | 0,77 | -3,50 | 4,62E-04 | 6,97E-03 |
| ENSG00000188015 | S100A3 | 7,44 | -2,66 | 0,75 | -3,53 | 4,10E-04 | 6,45E-03 |
| ENSG00000129654 | FOXJ1 | 83,61 | -2,65 | 0,60 | -4,45 | 8,73E-06 | 5,48E-04 |
| ENSG00000132530 | XAF1 | 884,48 | -2,65 | 0,60 | -4,45 | 8,48E-06 | 5,45E-04 |
| ENSG00000267206 | LCN6 | 11,12 | -2,64 | 0,75 | -3,55 | 3,91E-04 | 6,27E-03 |
| ENSG00000187513 | GJA4 | 89,64 | -2,64 | 0,48 | -5,45 | 4,96E-08 | 1,91E-05 |
| ENSG00000129467 | ADCY4 | 229,68 | -2,63 | 0,53 | -4,96 | 7,18E-07 | 1,15E-04 |
| ENSG00000148926 | ADM | 95,97 | -2,63 | 0,55 | -4,77 | 1,85E-06 | 2,10E-04 |
| ENSG00000169679 | BUB1 | 6,13 | -2,62 | 0,75 | -3,52 | 4,39E-04 | 6,74E-03 |
| ENSG00000149257 | SERPINH1 | 285,09 | -2,62 | 0,52 | -5,00 | 5,68E-07 | 9,71E-05 |
| ENSG00000240583 | AQP1 | 1217,97 | -2,60 | 0,50 | -5,17 | 2,29E-07 | 5,13E-05 |
| ENSG00000175793 | SFN | 32,18 | -2,59 | 0,65 | -3,99 | 6,64E-05 | 1,95E-03 |
| ENSG00000168874 | ATOH8 | 361,23 | -2,58 | 0,51 | -5,00 | 5,65E-07 | 9,71E-05 |
| ENSG00000027869 | SH2D2A | 12,34 | -2,57 | 0,62 | -4,16 | 3,24E-05 | 1,20E-03 |
| ENSG00000182718 | ANXA2 | 320,94 | -2,57 | 0,55 | -4,63 | 3,59E-06 | 3,31E-04 |
| ENSG00000163909 | HEYL | 86,57 | -2,54 | 0,48 | -5,25 | 1,51E-07 | 3,59E-05 |
| ENSG00000126778 | SIX1 | 10,02 | -2,54 | 0,91 | -2,78 | 5,51E-03 | 3,86E-02 |
| ENSG00000124440 | HIF3A | 940,67 | -2,53 | 0,58 | -4,35 | 1,38E-05 | 7,31E-04 |
| ENSG00000118503 | TNFAIP3 | 36,21 | -2,53 | 0,66 | -3,84 | 1,25E-04 | 2,96E-03 |
| ENSG00000179627 | ZBTB42 | 23,95 | -2,53 | 0,55 | -4,60 | 4,30E-06 | 3,74E-04 |
| ENSG00000141469 | SLC14A1 | 721,60 | -2,52 | 0,58 | -4,32 | 1,57E-05 | 7,86E-04 |
| ENSG00000187193 | MT1X | 1528,98 | -2,52 | 0,52 | -4,80 | 1,61E-06 | 1,91E-04 |
| ENSG00000101276 | SLC52A3 | 96,94 | -2,51 | 0,58 | -4,34 | 1,44E-05 | 7,43E-04 |
| ENSG00000205364 | MT1M | 1035,37 | -2,50 | 0,59 | -4,23 | 2,35E-05 | 9,93E-04 |
| ENSG00000130589 | HELZ2 | 111,50 | -2,47 | 0,46 | -5,39 | 6,90E-08 | 2,33E-05 |
| ENSG00000115604 | IL18R1 | 10,82 | -2,45 | 0,69 | -3,54 | 4,06E-04 | 6,40E-03 |
| ENSG00000128274 | A4GALT | 103,92 | -2,44 | 0,49 | -5,00 | 5,68E-07 | 9,71E-05 |
| ENSG00000136205 | TNS3 | 1607,36 | -2,43 | 0,44 | -5,56 | 2,64E-08 | 1,41E-05 |
| ENSG00000028137 | TNFRSF1B | 194,72 | -2,43 | 0,47 | -5,20 | 2,02E-07 | 4,58E-05 |
| ENSG00000185201 | IFITM2 | 640,79 | -2,43 | 0,49 | -4,95 | 7,35E-07 | 1,16E-04 |
| ENSG00000103241 | FOXF1 | 56,32 | -2,42 | 0,53 | -4,58 | 4,66E-06 | 3,87E-04 |
| ENSG00000145623 | OSMR | 58,46 | -2,41 | 0,43 | -5,56 | 2,71E-08 | 1,41E-05 |
| ENSG00000143772 | ITPKB | 3962,07 | -2,40 | 0,42 | -5,66 | 1,55E-08 | 1,12E-05 |
| ENSG00000075275 | CELSR1 | 174,06 | -2,40 | 0,72 | -3,34 | 8,44E-04 | 1,06E-02 |
| ENSG00000128602 | SMO | 263,11 | -2,40 | 0,50 | -4,80 | 1,55E-06 | 1,91E-04 |
| ENSG00000127418 | FGFRL1 | 775,65 | -2,38 | 0,47 | -5,06 | 4,17E-07 | 8,14E-05 |

| | | | | | | | |
|---|---|---|---|---|---|---|---|
| ENSG00000161638 | ITGA5 | 193,42 | -2,38 | 0,54 | -4,38 | 1,19E-05 | 6,68E-04 |
| ENSG00000115648 | MLPH | 20,49 | -2,38 | 0,74 | -3,21 | 1,32E-03 | 1,46E-02 |
| ENSG00000183615 | FAM167B | 41,86 | -2,38 | 0,57 | -4,17 | 3,03E-05 | 1,17E-03 |
| ENSG00000145888 | GLRA1 | 10,34 | -2,37 | 0,68 | -3,48 | 5,09E-04 | 7,42E-03 |
| ENSG00000102362 | SYTL4 | 139,46 | -2,37 | 0,55 | -4,29 | 1,80E-05 | 8,68E-04 |
| ENSG00000105559 | PLEKHA4 | 214,88 | -2,37 | 0,56 | -4,25 | 2,15E-05 | 9,44E-04 |
| ENSG00000154016 | GRAP | 73,21 | -2,37 | 0,48 | -4,96 | 7,10E-07 | 1,15E-04 |
| ENSG00000176046 | NUPR1 | 975,05 | -2,37 | 0,49 | -4,79 | 1,65E-06 | 1,93E-04 |
| ENSG00000116117 | PARD3B | 420,81 | -2,37 | 0,60 | -3,96 | 7,60E-05 | 2,09E-03 |
| ENSG00000025708 | TYMP | 239,58 | -2,36 | 0,60 | -3,95 | 7,82E-05 | 2,13E-03 |
| ENSG00000125148 | MT2A | 4574,93 | -2,36 | 0,51 | -4,59 | 4,44E-06 | 3,77E-04 |
| ENSG00000184113 | CLDN5 | 3033,76 | -2,35 | 0,49 | -4,82 | 1,46E-06 | 1,84E-04 |
| ENSG00000099998 | GGT5 | 540,57 | -2,35 | 0,47 | -4,98 | 6,47E-07 | 1,08E-04 |
| ENSG00000176771 | NCKAP5 | 382,93 | -2,35 | 0,57 | -4,13 | 3,66E-05 | 1,32E-03 |
| ENSG00000168404 | MLKL | 85,17 | -2,35 | 0,62 | -3,79 | 1,53E-04 | 3,37E-03 |
| ENSG00000060138 | YBX3 | 206,88 | -2,34 | 0,47 | -5,03 | 4,90E-07 | 9,03E-05 |
| ENSG00000105374 | NKG7 | 23,34 | -2,34 | 0,74 | -3,14 | 1,68E-03 | 1,73E-02 |
| ENSG00000197142 | ACSL5 | 125,68 | -2,34 | 0,50 | -4,64 | 3,45E-06 | 3,22E-04 |
| ENSG00000130303 | BST2 | 652,11 | -2,33 | 0,51 | -4,57 | 4,98E-06 | 4,01E-04 |
| ENSG00000091879 | ANGPT2 | 119,54 | -2,32 | 0,51 | -4,59 | 4,45E-06 | 3,77E-04 |
| ENSG00000271425 | NBPF10 | 357,92 | -2,31 | 0,50 | -4,59 | 4,35E-06 | 3,74E-04 |
| ENSG00000124942 | AHNAK | 1903,00 | -2,31 | 0,42 | -5,55 | 2,94E-08 | 1,41E-05 |
| ENSG00000177989 | ODF3B | 56,34 | -2,31 | 0,54 | -4,29 | 1,80E-05 | 8,68E-04 |
| ENSG00000198417 | MT1F | 1016,72 | -2,31 | 0,55 | -4,22 | 2,42E-05 | 9,99E-04 |
| ENSG00000171517 | LPAR3 | 115,38 | -2,31 | 0,54 | -4,28 | 1,90E-05 | 8,95E-04 |
| ENSG00000107738 | VSIR | 1181,07 | -2,31 | 0,50 | -4,57 | 4,85E-06 | 3,95E-04 |
| ENSG00000125144 | MT1G | 1126,29 | -2,30 | 0,61 | -3,79 | 1,48E-04 | 3,31E-03 |
| ENSG00000185507 | IRF7 | 197,86 | -2,30 | 0,54 | -4,29 | 1,80E-05 | 8,68E-04 |
| ENSG00000148498 | PARD3 | 574,02 | -2,30 | 0,54 | -4,26 | 2,02E-05 | 9,23E-04 |
| ENSG00000114790 | ARHGEF26 | 630,14 | -2,30 | 0,49 | -4,73 | 2,25E-06 | 2,39E-04 |
| ENSG00000111057 | KRT18 | 20,18 | -2,30 | 0,70 | -3,28 | 1,04E-03 | 1,22E-02 |
| ENSG00000286106 | NOTCH2NLR | 9,60 | -2,29 | 0,77 | -2,97 | 2,95E-03 | 2,52E-02 |
| ENSG00000154102 | C16orf74 | 96,41 | -2,29 | 0,53 | -4,35 | 1,34E-05 | 7,26E-04 |
| ENSG00000128849 | CGNL1 | 454,50 | -2,27 | 0,41 | -5,51 | 3,64E-08 | 1,59E-05 |
| ENSG00000125810 | CD93 | 32,88 | -2,27 | 0,52 | -4,39 | 1,16E-05 | 6,61E-04 |
| ENSG00000132470 | ITGB4 | 897,70 | -2,27 | 0,46 | -4,90 | 9,70E-07 | 1,36E-04 |
| ENSG00000064205 | CCN5 | 54,06 | -2,27 | 0,43 | -5,30 | 1,19E-07 | 3,28E-05 |
| ENSG00000161940 | BCL6B | 57,92 | -2,27 | 0,44 | -5,20 | 1,97E-07 | 4,54E-05 |
| ENSG00000176692 | FOXC2 | 26,13 | -2,26 | 0,76 | -3,00 | 2,74E-03 | 2,38E-02 |
| ENSG00000018280 | SLC11A1 | 118,55 | -2,26 | 0,55 | -4,10 | 4,09E-05 | 1,43E-03 |
| ENSG00000162692 | VCAM1 | 33,10 | -2,25 | 0,64 | -3,53 | 4,21E-04 | 6,57E-03 |
| ENSG00000185022 | MAFF | 208,11 | -2,25 | 0,42 | -5,34 | 9,29E-08 | 2,91E-05 |
| ENSG00000150048 | CLEC1A | 19,06 | -2,25 | 0,56 | -3,99 | 6,55E-05 | 1,94E-03 |
| ENSG00000085276 | MECOM | 108,94 | -2,24 | 0,55 | -4,04 | 5,28E-05 | 1,68E-03 |
| ENSG00000143545 | RAB13 | 991,00 | -2,23 | 0,47 | -4,78 | 1,71E-06 | 1,97E-04 |
| ENSG00000285526 | ENSG000002855 | 89,04 | -2,23 | 0,51 | -4,37 | 1,26E-05 | 6,96E-04 |
| ENSG00000185860 | CCDC190 | 44,85 | -2,23 | 0,64 | -3,48 | 5,11E-04 | 7,42E-03 |
| ENSG00000154025 | SLC5A10 | 80,80 | -2,23 | 0,46 | -4,87 | 1,12E-06 | 1,54E-04 |
| ENSG00000110852 | CLEC2B | 33,81 | -2,21 | 0,51 | -4,34 | 1,45E-05 | 7,46E-04 |
| ENSG00000134817 | APLNR | 666,88 | -2,21 | 0,52 | -4,26 | 2,05E-05 | 9,27E-04 |
| ENSG00000168913 | ENHO | 6717,43 | -2,21 | 0,48 | -4,63 | 3,59E-06 | 3,31E-04 |
| ENSG00000020633 | RUNX3 | 43,95 | -2,20 | 0,66 | -3,35 | 7,98E-04 | 1,01E-02 |
| ENSG00000142089 | IFITM3 | 2552,69 | -2,20 | 0,45 | -4,86 | 1,16E-06 | 1,57E-04 |
| ENSG00000185499 | MUC1 | 62,11 | -2,20 | 0,59 | -3,70 | 2,17E-04 | 4,26E-03 |
| ENSG00000140022 | STON2 | 640,81 | -2,20 | 0,45 | -4,86 | 1,19E-06 | 1,59E-04 |
| ENSG00000137959 | IFI44L | 418,15 | -2,20 | 0,61 | -3,62 | 2,93E-04 | 5,13E-03 |
| ENSG00000188039 | NWD1 | 441,66 | -2,20 | 0,47 | -4,66 | 3,12E-06 | 3,01E-04 |

| | | | | | | | |
|---|---|---|---|---|---|---|---|
| ENSG00000142627 | EPHA2 | 58,45 | -2,19 | 0,46 | -4,75 | 2,00E-06 | 2,20E-04 |
| ENSG00000275395 | FCGBP | 151,01 | -2,19 | 0,61 | -3,62 | 2,98E-04 | 5,19E-03 |
| ENSG00000244414 | CFHR1 | 12,75 | -2,19 | 0,69 | -3,15 | 1,63E-03 | 1,70E-02 |
| ENSG00000100249 | C22orf31 | 12,66 | -2,19 | 0,61 | -3,59 | 3,29E-04 | 5,57E-03 |
| ENSG00000168899 | VAMP5 | 522,23 | -2,18 | 0,46 | -4,78 | 1,73E-06 | 1,98E-04 |
| ENSG00000102287 | GABRE | 33,33 | -2,18 | 0,55 | -3,98 | 7,02E-05 | 2,01E-03 |
| ENSG00000129116 | PALLD | 417,52 | -2,18 | 0,41 | -5,31 | 1,09E-07 | 3,11E-05 |
| ENSG00000270629 | NBPF14 | 575,16 | -2,17 | 0,51 | -4,28 | 1,90E-05 | 8,95E-04 |
| ENSG00000139832 | RAB20 | 33,60 | -2,17 | 0,45 | -4,79 | 1,68E-06 | 1,95E-04 |
| ENSG00000198851 | CD3E | 9,39 | -2,17 | 0,76 | -2,84 | 4,56E-03 | 3,39E-02 |
| ENSG00000161509 | GRIN2C | 1317,73 | -2,17 | 0,46 | -4,75 | 2,01E-06 | 2,20E-04 |
| ENSG00000110799 | VWF | 908,32 | -2,17 | 0,39 | -5,57 | 2,49E-08 | 1,41E-05 |
| ENSG00000172766 | NAA16 | 491,77 | -2,16 | 0,58 | -3,74 | 1,81E-04 | 3,79E-03 |
| ENSG00000106571 | GLI3 | 141,51 | -2,16 | 0,51 | -4,26 | 2,04E-05 | 9,27E-04 |
| ENSG00000099860 | GADD45B | 551,35 | -2,15 | 0,48 | -4,46 | 8,27E-06 | 5,42E-04 |
| ENSG00000196616 | ADH1B | 11,29 | -2,15 | 0,67 | -3,22 | 1,29E-03 | 1,44E-02 |
| ENSG00000087086 | FTL | 32125,17 | -2,15 | 0,42 | -5,17 | 2,36E-07 | 5,22E-05 |
| ENSG00000160808 | MYL3 | 121,39 | -2,14 | 0,51 | -4,23 | 2,32E-05 | 9,84E-04 |
| ENSG00000127472 | PLA2G5 | 140,43 | -2,14 | 0,48 | -4,45 | 8,76E-06 | 5,48E-04 |
| ENSG00000181856 | SLC2A4 | 52,83 | -2,14 | 0,57 | -3,73 | 1,92E-04 | 3,97E-03 |
| ENSG00000135245 | HILPDA | 317,02 | -2,14 | 0,53 | -4,06 | 4,91E-05 | 1,60E-03 |
| ENSG00000135926 | TMBIM1 | 1421,37 | -2,14 | 0,40 | -5,33 | 1,00E-07 | 2,91E-05 |
| ENSG00000006327 | TNFRSF12A | 25,77 | -2,13 | 0,60 | -3,54 | 4,06E-04 | 6,40E-03 |
| ENSG00000196329 | GIMAP5 | 173,43 | -2,13 | 0,47 | -4,57 | 4,83E-06 | 3,95E-04 |
| ENSG00000167157 | PRRX2 | 13,58 | -2,13 | 0,65 | -3,26 | 1,13E-03 | 1,30E-02 |
| ENSG00000241839 | PLEKHO2 | 647,41 | -2,12 | 0,45 | -4,75 | 2,00E-06 | 2,20E-04 |
| ENSG00000109610 | SOD3 | 455,65 | -2,11 | 0,51 | -4,10 | 4,05E-05 | 1,42E-03 |
| ENSG00000161958 | FGF11 | 240,61 | -2,10 | 0,50 | -4,18 | 2,88E-05 | 1,14E-03 |
| ENSG00000174990 | CA5A | 9,61 | -2,10 | 0,68 | -3,08 | 2,07E-03 | 2,00E-02 |
| ENSG00000243649 | CFB | 131,29 | -2,10 | 0,50 | -4,23 | 2,31E-05 | 9,84E-04 |
| ENSG00000156313 | RPGR | 334,08 | -2,10 | 0,51 | -4,11 | 3,98E-05 | 1,40E-03 |
| ENSG00000130052 | STARD8 | 57,21 | -2,10 | 0,54 | -3,91 | 9,21E-05 | 2,40E-03 |
| ENSG00000137693 | YAP1 | 296,41 | -2,10 | 0,44 | -4,82 | 1,42E-06 | 1,80E-04 |
| ENSG00000144908 | ALDH1L1 | 1530,42 | -2,10 | 0,52 | -4,02 | 5,87E-05 | 1,80E-03 |
| ENSG00000171773 | NXNL1 | 18,14 | -2,10 | 0,71 | -2,96 | 3,06E-03 | 2,58E-02 |
| ENSG00000123374 | CDK2 | 27,02 | -2,09 | 0,48 | -4,32 | 1,53E-05 | 7,72E-04 |
| ENSG00000185339 | TCN2 | 300,00 | -2,09 | 0,41 | -5,10 | 3,41E-07 | 7,15E-05 |
| ENSG00000135111 | TBX3 | 41,66 | -2,09 | 0,42 | -4,97 | 6,85E-07 | 1,13E-04 |
| ENSG00000168077 | SCARA3 | 1278,16 | -2,09 | 0,43 | -4,82 | 1,40E-06 | 1,79E-04 |
| ENSG00000165917 | RAPSN | 8,28 | -2,08 | 0,69 | -3,00 | 2,70E-03 | 2,36E-02 |
| ENSG00000146535 | GNA12 | 1231,81 | -2,08 | 0,37 | -5,57 | 2,61E-08 | 1,41E-05 |
| ENSG00000103260 | METRN | 7437,87 | -2,08 | 0,47 | -4,42 | 1,00E-05 | 5,95E-04 |
| ENSG00000122786 | CALD1 | 4006,11 | -2,08 | 0,54 | -3,86 | 1,12E-04 | 2,73E-03 |
| ENSG00000132031 | MATN3 | 18,00 | -2,07 | 0,64 | -3,23 | 1,24E-03 | 1,39E-02 |
| ENSG00000172361 | CFAP53 | 129,60 | -2,07 | 0,60 | -3,43 | 6,13E-04 | 8,35E-03 |
| ENSG00000178175 | ZNF366 | 126,31 | -2,06 | 0,57 | -3,60 | 3,17E-04 | 5,41E-03 |
| ENSG00000102755 | FLT1 | 534,93 | -2,06 | 0,36 | -5,76 | 8,44E-09 | 7,77E-06 |
| ENSG00000119938 | PPP1R3C | 499,82 | -2,06 | 0,47 | -4,37 | 1,26E-05 | 6,96E-04 |
| ENSG00000151929 | BAG3 | 344,21 | -2,06 | 0,50 | -4,16 | 3,14E-05 | 1,19E-03 |
| ENSG00000121068 | TBX2 | 226,98 | -2,06 | 0,48 | -4,24 | 2,22E-05 | 9,61E-04 |
| ENSG00000197324 | LRP10 | 772,76 | -2,06 | 0,43 | -4,80 | 1,58E-06 | 1,91E-04 |
| ENSG00000096696 | DSP | 60,57 | -2,05 | 0,57 | -3,57 | 3,62E-04 | 5,95E-03 |
| ENSG00000102359 | SRPX2 | 107,83 | -2,04 | 0,51 | -3,96 | 7,58E-05 | 2,09E-03 |
| ENSG00000018408 | WWTR1 | 176,43 | -2,04 | 0,35 | -5,80 | 6,46E-09 | 6,30E-06 |
| ENSG00000100060 | MFNG | 68,61 | -2,04 | 0,46 | -4,42 | 1,00E-05 | 5,94E-04 |
| ENSG00000203811 | H3C14 | 14,75 | -2,03 | 0,63 | -3,25 | 1,17E-03 | 1,34E-02 |
| ENSG00000162618 | ADGRL4 | 71,17 | -2,03 | 0,40 | -5,02 | 5,18E-07 | 9,33E-05 |

| | | | | | | | |
|---|---|---|---|---|---|---|---|
| ENSG00000225781 | OR6V1 | 14,15 | -2,03 | 0,74 | -2,73 | 6,27E-03 | 4,21E-02 |
| ENSG00000181826 | RELL1 | 182,21 | -2,03 | 0,38 | -5,28 | 1,32E-07 | 3,30E-05 |
| ENSG00000134042 | MRO | 519,62 | -2,03 | 0,33 | -6,08 | 1,21E-09 | 3,85E-06 |
| ENSG00000160801 | PTH1R | 721,18 | -2,02 | 0,48 | -4,18 | 2,87E-05 | 1,14E-03 |
| ENSG00000265972 | TXNIP | 1505,32 | -2,02 | 0,50 | -4,07 | 4,76E-05 | 1,57E-03 |
| ENSG00000167676 | PLIN4 | 320,33 | -2,02 | 0,53 | -3,85 | 1,18E-04 | 2,86E-03 |
| ENSG00000129450 | SIGLEC9 | 9,39 | -2,01 | 0,74 | -2,74 | 6,15E-03 | 4,15E-02 |
| ENSG00000148604 | RGR | 139,23 | -2,01 | 0,58 | -3,45 | 5,59E-04 | 7,88E-03 |
| ENSG00000089356 | FXYD3 | 83,08 | -2,01 | 0,53 | -3,75 | 1,76E-04 | 3,72E-03 |
| ENSG00000152207 | CYSLTR2 | 19,98 | -2,01 | 0,49 | -4,13 | 3,56E-05 | 1,29E-03 |
| ENSG00000198604 | BAZ1A | 615,32 | -2,00 | 0,54 | -3,68 | 2,35E-04 | 4,48E-03 |
| ENSG00000108848 | LUC7L3 | 23108,56 | -2,00 | 0,61 | -3,26 | 1,10E-03 | 1,28E-02 |
| ENSG00000161955 | TNFSF13 | 696,47 | -2,00 | 0,44 | -4,58 | 4,71E-06 | 3,89E-04 |
| ENSG00000164761 | TNFRSF11B | 10,05 | -2,00 | 0,75 | -2,66 | 7,87E-03 | 4,94E-02 |
| ENSG00000203852 | H3C15 | 15,01 | -2,00 | 0,63 | -3,16 | 1,59E-03 | 1,67E-02 |
| ENSG00000171316 | CHD7 | 1893,98 | -1,99 | 0,56 | -3,55 | 3,91E-04 | 6,27E-03 |
| ENSG00000185338 | SOCS1 | 13,62 | -1,98 | 0,62 | -3,22 | 1,30E-03 | 1,44E-02 |
| ENSG00000151414 | NEK7 | 443,50 | -1,98 | 0,37 | -5,29 | 1,22E-07 | 3,30E-05 |
| ENSG00000101096 | NFATC2 | 49,63 | -1,98 | 0,46 | -4,30 | 1,68E-05 | 8,32E-04 |
| ENSG00000187720 | THSD4 | 137,27 | -1,98 | 0,50 | -3,95 | 7,88E-05 | 2,14E-03 |
| ENSG00000181019 | NQO1 | 404,18 | -1,98 | 0,47 | -4,18 | 2,93E-05 | 1,15E-03 |
| ENSG00000100427 | MLC1 | 5066,01 | -1,97 | 0,44 | -4,50 | 6,72E-06 | 4,82E-04 |
| ENSG00000155850 | SLC26A2 | 114,81 | -1,97 | 0,54 | -3,65 | 2,60E-04 | 4,81E-03 |
| ENSG00000251493 | FOXD1 | 24,09 | -1,97 | 0,51 | -3,87 | 1,10E-04 | 2,70E-03 |
| ENSG00000064393 | HIPK2 | 8287,86 | -1,97 | 0,42 | -4,70 | 2,63E-06 | 2,64E-04 |
| ENSG00000077150 | NFKB2 | 168,66 | -1,97 | 0,44 | -4,43 | 9,55E-06 | 5,84E-04 |
| ENSG00000135094 | SDS | 182,67 | -1,96 | 0,51 | -3,83 | 1,26E-04 | 2,96E-03 |
| ENSG00000067066 | SP100 | 146,76 | -1,96 | 0,45 | -4,35 | 1,37E-05 | 7,28E-04 |
| ENSG00000165424 | ZCCHC24 | 3530,73 | -1,96 | 0,39 | -5,06 | 4,18E-07 | 8,14E-05 |
| ENSG00000189152 | GRAPL | 8,48 | -1,96 | 0,63 | -3,12 | 1,79E-03 | 1,80E-02 |
| ENSG00000174348 | PODN | 91,91 | -1,96 | 0,42 | -4,67 | 3,06E-06 | 3,00E-04 |
| ENSG00000261701 | HPR | 237,44 | -1,95 | 0,57 | -3,42 | 6,15E-04 | 8,37E-03 |
| ENSG00000185650 | ZFP36L1 | 1474,46 | -1,95 | 0,48 | -4,10 | 4,07E-05 | 1,42E-03 |
| ENSG00000136297 | MMD2 | 260,84 | -1,95 | 0,45 | -4,32 | 1,57E-05 | 7,86E-04 |
| ENSG00000124145 | SDC4 | 1762,02 | -1,94 | 0,48 | -4,05 | 5,05E-05 | 1,64E-03 |
| ENSG00000177409 | SAMD9L | 35,41 | -1,94 | 0,46 | -4,25 | 2,16E-05 | 9,45E-04 |
| ENSG00000086062 | B4GALT1 | 93,38 | -1,93 | 0,42 | -4,64 | 3,44E-06 | 3,22E-04 |
| ENSG00000162645 | GBP2 | 79,12 | -1,93 | 0,41 | -4,74 | 2,11E-06 | 2,26E-04 |
| ENSG00000175592 | FOSL1 | 14,57 | -1,93 | 0,65 | -2,99 | 2,79E-03 | 2,42E-02 |
| ENSG00000115594 | IL1R1 | 41,67 | -1,93 | 0,47 | -4,09 | 4,28E-05 | 1,47E-03 |
| ENSG00000155366 | RHOC | 1539,22 | -1,93 | 0,38 | -5,12 | 3,09E-07 | 6,56E-05 |
| ENSG00000288681 | ENSG000002886 | 516,39 | -1,92 | 0,39 | -4,90 | 9,62E-07 | 1,36E-04 |
| ENSG00000162825 | NBPF20 | 771,80 | -1,92 | 0,63 | -3,04 | 2,39E-03 | 2,19E-02 |
| ENSG00000163874 | ZC3H12A | 13,13 | -1,92 | 0,59 | -3,25 | 1,17E-03 | 1,34E-02 |
| ENSG00000221869 | CEBPD | 607,73 | -1,92 | 0,49 | -3,90 | 9,82E-05 | 2,51E-03 |
| ENSG00000185885 | IFITM1 | 610,23 | -1,91 | 0,38 | -5,02 | 5,14E-07 | 9,33E-05 |
| ENSG00000134201 | GSTM5 | 976,63 | -1,91 | 0,61 | -3,11 | 1,88E-03 | 1,86E-02 |
| ENSG00000059378 | PARP12 | 170,92 | -1,91 | 0,46 | -4,17 | 3,09E-05 | 1,18E-03 |
| ENSG00000205517 | RGL3 | 183,32 | -1,90 | 0,52 | -3,65 | 2,62E-04 | 4,83E-03 |
| ENSG00000181626 | ANKRD62 | 43,78 | -1,90 | 0,69 | -2,75 | 5,92E-03 | 4,06E-02 |
| ENSG00000102265 | TIMP1 | 457,60 | -1,90 | 0,44 | -4,37 | 1,25E-05 | 6,96E-04 |
| ENSG00000185561 | TLCD2 | 69,94 | -1,90 | 0,58 | -3,29 | 9,86E-04 | 1,18E-02 |
| ENSG00000110077 | MS4A6A | 12,97 | -1,90 | 0,67 | -2,83 | 4,67E-03 | 3,44E-02 |
| ENSG00000159423 | ALDH4A1 | 1652,08 | -1,90 | 0,45 | -4,25 | 2,11E-05 | 9,40E-04 |
| ENSG00000262165 | C17orf114 | 57,35 | -1,90 | 0,53 | -3,55 | 3,79E-04 | 6,14E-03 |
| ENSG00000130055 | GDPD2 | 119,00 | -1,89 | 0,46 | -4,07 | 4,77E-05 | 1,57E-03 |
| ENSG00000101187 | SLCO4A1 | 353,60 | -1,89 | 0,50 | -3,80 | 1,45E-04 | 3,25E-03 |

| | | | | | | | |
|---|---|---|---|---|---|---|---|
| ENSG00000281887 | GIMAP1-GIMAP5 | 153,28 | -1,89 | 0,44 | -4,28 | 1,84E-05 | 8,80E-04 |
| ENSG00000183148 | ANKRD20A2P | 2459,46 | -1,89 | 0,65 | -2,90 | 3,68E-03 | 2,95E-02 |
| ENSG00000173548 | SNX33 | 274,07 | -1,89 | 0,42 | -4,49 | 7,06E-06 | 4,94E-04 |
| ENSG00000112343 | TRIM38 | 75,19 | -1,88 | 0,48 | -3,95 | 7,82E-05 | 2,13E-03 |
| ENSG00000187479 | C11orf96 | 384,29 | -1,88 | 0,46 | -4,08 | 4,48E-05 | 1,52E-03 |
| ENSG00000221963 | APOL6 | 113,40 | -1,88 | 0,46 | -4,07 | 4,75E-05 | 1,57E-03 |
| ENSG00000177426 | TGIF1 | 77,38 | -1,88 | 0,39 | -4,80 | 1,60E-06 | 1,91E-04 |
| ENSG00000163239 | TDRD10 | 27,35 | -1,88 | 0,60 | -3,15 | 1,63E-03 | 1,70E-02 |
| ENSG00000129250 | KIF1C | 5558,54 | -1,88 | 0,41 | -4,59 | 4,48E-06 | 3,78E-04 |
| ENSG00000064787 | BCAS1 | 9755,80 | -1,88 | 0,41 | -4,60 | 4,29E-06 | 3,74E-04 |
| ENSG00000198113 | TOR4A | 39,22 | -1,88 | 0,54 | -3,50 | 4,73E-04 | 7,05E-03 |
| ENSG00000276203 | ANKRD20A3P | 2295,25 | -1,88 | 0,65 | -2,89 | 3,82E-03 | 3,02E-02 |
| ENSG00000177791 | MYOZ1 | 22,80 | -1,87 | 0,45 | -4,17 | 3,04E-05 | 1,17E-03 |
| ENSG00000163513 | TGFBR2 | 161,75 | -1,87 | 0,43 | -4,40 | 1,06E-05 | 6,23E-04 |
| ENSG00000100065 | CARD10 | 60,41 | -1,87 | 0,49 | -3,79 | 1,52E-04 | 3,36E-03 |
| ENSG00000169871 | TRIM56 | 1437,05 | -1,87 | 0,50 | -3,77 | 1,63E-04 | 3,52E-03 |
| ENSG00000170425 | ADORA2B | 90,37 | -1,87 | 0,49 | -3,84 | 1,24E-04 | 2,95E-03 |
| ENSG00000214456 | PLIN5 | 320,30 | -1,87 | 0,57 | -3,27 | 1,07E-03 | 1,25E-02 |
| ENSG00000111783 | RFX4 | 303,73 | -1,87 | 0,42 | -4,48 | 7,58E-06 | 5,15E-04 |
| ENSG00000286019 | NOTCH2NLB | 79,06 | -1,87 | 0,48 | -3,87 | 1,10E-04 | 2,70E-03 |
| ENSG00000164330 | EBF1 | 103,19 | -1,86 | 0,47 | -3,94 | 8,10E-05 | 2,18E-03 |
| ENSG00000155659 | VSIG4 | 28,83 | -1,86 | 0,62 | -3,03 | 2,47E-03 | 2,23E-02 |
| ENSG00000074047 | GLI2 | 26,29 | -1,86 | 0,57 | -3,26 | 1,11E-03 | 1,28E-02 |
| ENSG00000113504 | SLC12A7 | 328,94 | -1,86 | 0,40 | -4,65 | 3,33E-06 | 3,17E-04 |
| ENSG00000213390 | ARHGAP19 | 130,10 | -1,86 | 0,34 | -5,49 | 4,09E-08 | 1,65E-05 |
| ENSG00000142102 | PGGHG | 344,23 | -1,85 | 0,46 | -3,99 | 6,63E-05 | 1,95E-03 |
| ENSG00000016391 | CHDH | 969,18 | -1,85 | 0,44 | -4,25 | 2,09E-05 | 9,39E-04 |
| ENSG00000169715 | MT1E | 1894,06 | -1,85 | 0,47 | -3,95 | 7,98E-05 | 2,16E-03 |
| ENSG00000169291 | SHE | 77,90 | -1,85 | 0,42 | -4,39 | 1,12E-05 | 6,45E-04 |
| ENSG00000158195 | WASF2 | 1765,77 | -1,84 | 0,40 | -4,58 | 4,60E-06 | 3,84E-04 |
| ENSG00000124374 | PAIP2B | 2160,01 | -1,84 | 0,39 | -4,71 | 2,44E-06 | 2,49E-04 |
| ENSG00000132321 | IQCA1 | 2043,52 | -1,84 | 0,56 | -3,30 | 9,77E-04 | 1,18E-02 |
| ENSG00000147509 | RGS20 | 244,53 | -1,84 | 0,44 | -4,21 | 2,61E-05 | 1,05E-03 |
| ENSG00000244255 | AL645922.1 | 140,15 | -1,84 | 0,45 | -4,05 | 5,03E-05 | 1,63E-03 |
| ENSG00000197971 | MBP | 196107,50 | -1,84 | 0,44 | -4,18 | 2,87E-05 | 1,14E-03 |
| ENSG00000136830 | NIBAN2 | 648,57 | -1,84 | 0,38 | -4,91 | 9,30E-07 | 1,34E-04 |
| ENSG00000188313 | PLSCR1 | 73,38 | -1,84 | 0,39 | -4,71 | 2,48E-06 | 2,52E-04 |
| ENSG00000139352 | ASCL1 | 771,34 | -1,83 | 0,54 | -3,38 | 7,29E-04 | 9,50E-03 |
| ENSG00000258659 | TRIM34 | 15,53 | -1,83 | 0,56 | -3,28 | 1,04E-03 | 1,23E-02 |
| ENSG00000133858 | ZFC3H1 | 1201,87 | -1,82 | 0,53 | -3,42 | 6,31E-04 | 8,52E-03 |
| ENSG00000001617 | SEMA3F | 188,53 | -1,82 | 0,44 | -4,11 | 3,88E-05 | 1,38E-03 |
| ENSG00000173156 | RHOD | 44,78 | -1,82 | 0,66 | -2,78 | 5,51E-03 | 3,86E-02 |
| ENSG00000124782 | RREB1 | 75,33 | -1,82 | 0,49 | -3,70 | 2,17E-04 | 4,26E-03 |
| ENSG00000123636 | BAZ2B | 3957,26 | -1,82 | 0,57 | -3,21 | 1,31E-03 | 1,45E-02 |
| ENSG00000135917 | SLC19A3 | 53,63 | -1,82 | 0,43 | -4,24 | 2,27E-05 | 9,74E-04 |
| ENSG00000119927 | GPAM | 305,55 | -1,81 | 0,43 | -4,18 | 2,97E-05 | 1,16E-03 |
| ENSG00000205929 | C21orf62 | 38,42 | -1,81 | 0,66 | -2,72 | 6,45E-03 | 4,30E-02 |
| ENSG00000133640 | LRRIQ1 | 205,58 | -1,81 | 0,57 | -3,15 | 1,61E-03 | 1,68E-02 |
| ENSG00000100599 | RIN3 | 221,98 | -1,81 | 0,47 | -3,87 | 1,07E-04 | 2,66E-03 |
| ENSG00000163346 | PBXIP1 | 2889,46 | -1,81 | 0,42 | -4,26 | 2,04E-05 | 9,27E-04 |
| ENSG00000139567 | ACVRL1 | 230,18 | -1,81 | 0,36 | -5,07 | 3,97E-07 | 8,03E-05 |
| ENSG00000106624 | AEBP1 | 1039,79 | -1,80 | 0,44 | -4,07 | 4,65E-05 | 1,56E-03 |
| ENSG00000197747 | S100A10 | 192,25 | -1,80 | 0,42 | -4,27 | 1,96E-05 | 9,11E-04 |
| ENSG00000086544 | ITPKC | 332,91 | -1,80 | 0,43 | -4,24 | 2,24E-05 | 9,64E-04 |
| ENSG00000128917 | DLL4 | 114,29 | -1,80 | 0,47 | -3,87 | 1,07E-04 | 2,66E-03 |
| ENSG00000140931 | CMTM3 | 107,38 | -1,80 | 0,43 | -4,23 | 2,32E-05 | 9,84E-04 |
| ENSG00000134508 | CABLES1 | 1644,09 | -1,80 | 0,42 | -4,34 | 1,43E-05 | 7,41E-04 |

| | | | | | | | |
|---|---|---|---|---|---|---|---|
| ENSG00000251184 | AL672142.1 | 170,20 | -1,80 | 0,54 | -3,37 | 7,64E-04 | 9,83E-03 |
| ENSG00000259207 | ITGB3 | 21,77 | -1,80 | 0,54 | -3,33 | 8,79E-04 | 1,09E-02 |
| ENSG00000137491 | SLCO2B1 | 538,99 | -1,80 | 0,34 | -5,33 | 9,99E-08 | 2,91E-05 |
| ENSG00000101400 | SNTA1 | 4741,04 | -1,80 | 0,42 | -4,26 | 2,06E-05 | 9,29E-04 |
| ENSG00000160233 | LRRC3 | 23,51 | -1,79 | 0,60 | -2,98 | 2,92E-03 | 2,50E-02 |
| ENSG00000069399 | BCL3 | 121,07 | -1,79 | 0,46 | -3,92 | 8,80E-05 | 2,32E-03 |
| ENSG00000235750 | KIAA0040 | 32,90 | -1,79 | 0,37 | -4,83 | 1,35E-06 | 1,74E-04 |
| ENSG00000020577 | SAMD4A | 464,93 | -1,79 | 0,33 | -5,47 | 4,61E-08 | 1,82E-05 |
| ENSG00000168461 | RAB31 | 2079,97 | -1,79 | 0,41 | -4,34 | 1,44E-05 | 7,43E-04 |
| ENSG00000254469 | AP002495.1 | 110,68 | -1,79 | 0,60 | -2,96 | 3,06E-03 | 2,58E-02 |
| ENSG00000164976 | MYORG | 768,80 | -1,79 | 0,41 | -4,38 | 1,17E-05 | 6,65E-04 |
| ENSG00000090376 | IRAK3 | 31,69 | -1,79 | 0,51 | -3,49 | 4,80E-04 | 7,12E-03 |
| ENSG00000008517 | IL32 | 190,62 | -1,79 | 0,46 | -3,87 | 1,07E-04 | 2,66E-03 |
| ENSG00000078804 | TP53INP2 | 5176,56 | -1,79 | 0,41 | -4,40 | 1,06E-05 | 6,23E-04 |
| ENSG00000087266 | SH3BP2 | 519,76 | -1,79 | 0,45 | -3,99 | 6,73E-05 | 1,97E-03 |
| ENSG00000258588 | TRIM6-TRIM34 | 12,11 | -1,78 | 0,49 | -3,61 | 3,05E-04 | 5,27E-03 |
| ENSG00000151012 | SLC7A11 | 734,98 | -1,78 | 0,43 | -4,15 | 3,37E-05 | 1,24E-03 |
| ENSG00000116016 | EPAS1 | 1086,01 | -1,78 | 0,35 | -5,06 | 4,27E-07 | 8,14E-05 |
| ENSG00000164849 | GPR146 | 188,83 | -1,78 | 0,41 | -4,38 | 1,18E-05 | 6,66E-04 |
| ENSG00000275993 | SIK1B | 193,79 | -1,78 | 0,39 | -4,59 | 4,43E-06 | 3,77E-04 |
| ENSG00000101850 | GPR143 | 101,78 | -1,78 | 0,43 | -4,11 | 3,91E-05 | 1,39E-03 |
| ENSG00000175556 | LONRF3 | 6,95 | -1,77 | 0,63 | -2,82 | 4,76E-03 | 3,49E-02 |
| ENSG00000184371 | CSF1 | 286,69 | -1,77 | 0,39 | -4,56 | 5,13E-06 | 4,03E-04 |
| ENSG00000138696 | BMPR1B | 295,08 | -1,77 | 0,45 | -3,95 | 7,95E-05 | 2,15E-03 |
| ENSG00000244731 | C4A | 1653,26 | -1,77 | 0,49 | -3,65 | 2,59E-04 | 4,80E-03 |
| ENSG00000224389 | C4B | 1741,46 | -1,77 | 0,49 | -3,64 | 2,73E-04 | 4,91E-03 |
| ENSG00000106560 | GIMAP2 | 18,24 | -1,77 | 0,45 | -3,94 | 8,02E-05 | 2,16E-03 |
| ENSG00000133048 | CHI3L1 | 516,58 | -1,77 | 0,48 | -3,71 | 2,04E-04 | 4,09E-03 |
| ENSG00000163072 | NOSTRIN | 111,59 | -1,77 | 0,47 | -3,76 | 1,73E-04 | 3,69E-03 |
| ENSG00000144579 | CTDSP1 | 1022,84 | -1,77 | 0,38 | -4,64 | 3,41E-06 | 3,21E-04 |
| ENSG00000169908 | TM4SF1 | 172,61 | -1,77 | 0,39 | -4,56 | 5,06E-06 | 4,02E-04 |
| ENSG00000175287 | PHYHD1 | 493,48 | -1,76 | 0,50 | -3,52 | 4,26E-04 | 6,62E-03 |
| ENSG00000177674 | AGTRAP | 340,67 | -1,76 | 0,46 | -3,85 | 1,19E-04 | 2,86E-03 |
| ENSG00000145555 | MYO10 | 6343,45 | -1,76 | 0,53 | -3,30 | 9,75E-04 | 1,17E-02 |
| ENSG00000188060 | RAB42 | 9,63 | -1,76 | 0,64 | -2,77 | 5,60E-03 | 3,90E-02 |
| ENSG00000288656 | ENSG000002886 | 1102,36 | -1,76 | 0,42 | -4,16 | 3,13E-05 | 1,19E-03 |
| ENSG00000081320 | STK17B | 86,68 | -1,76 | 0,50 | -3,52 | 4,36E-04 | 6,72E-03 |
| ENSG00000026508 | CD44 | 107,05 | -1,76 | 0,49 | -3,58 | 3,49E-04 | 5,78E-03 |
| ENSG00000143382 | ADAMTSL4 | 38,24 | -1,76 | 0,48 | -3,68 | 2,37E-04 | 4,51E-03 |
| ENSG00000130158 | DOCK6 | 331,68 | -1,76 | 0,43 | -4,10 | 4,20E-05 | 1,45E-03 |
| ENSG00000007866 | TEAD3 | 54,51 | -1,75 | 0,47 | -3,72 | 1,98E-04 | 4,02E-03 |
| ENSG00000196209 | SIRPB2 | 9,85 | -1,75 | 0,53 | -3,33 | 8,57E-04 | 1,07E-02 |
| ENSG00000160712 | IL6R | 184,43 | -1,75 | 0,48 | -3,64 | 2,70E-04 | 4,89E-03 |
| ENSG00000256235 | SMIM3 | 107,47 | -1,75 | 0,38 | -4,57 | 4,98E-06 | 4,01E-04 |
| ENSG00000152661 | GJA1 | 2495,03 | -1,75 | 0,48 | -3,65 | 2,64E-04 | 4,84E-03 |
| ENSG00000213719 | CLIC1 | 177,85 | -1,75 | 0,35 | -4,95 | 7,52E-07 | 1,18E-04 |
| ENSG00000068079 | IFI35 | 177,56 | -1,74 | 0,42 | -4,13 | 3,70E-05 | 1,33E-03 |
| ENSG00000074219 | TEAD2 | 70,77 | -1,74 | 0,48 | -3,60 | 3,16E-04 | 5,41E-03 |
| ENSG00000187800 | PEAR1 | 41,43 | -1,74 | 0,47 | -3,67 | 2,41E-04 | 4,57E-03 |
| ENSG00000160781 | PAQR6 | 8575,89 | -1,74 | 0,37 | -4,66 | 3,12E-06 | 3,01E-04 |
| ENSG00000100968 | NFATC4 | 511,34 | -1,74 | 0,48 | -3,59 | 3,34E-04 | 5,63E-03 |
| ENSG00000167261 | DPEP2 | 39,96 | -1,74 | 0,48 | -3,64 | 2,72E-04 | 4,91E-03 |
| ENSG00000056487 | PHF21B | 135,49 | -1,73 | 0,46 | -3,79 | 1,52E-04 | 3,37E-03 |
| ENSG00000182902 | SLC25A18 | 2190,79 | -1,73 | 0,44 | -3,98 | 6,86E-05 | 1,99E-03 |
| ENSG00000182541 | LIMK2 | 273,19 | -1,73 | 0,36 | -4,76 | 1,98E-06 | 2,20E-04 |
| ENSG00000125733 | TRIP10 | 214,35 | -1,73 | 0,43 | -4,07 | 4,80E-05 | 1,58E-03 |
| ENSG00000140459 | CYP11A1 | 58,06 | -1,73 | 0,50 | -3,48 | 4,93E-04 | 7,28E-03 |

| | | | | | | | |
|---|---|---|---|---|---|---|---|
| ENSG00000174501 | ANKRD36C | 2872,61 | -1,72 | 0,59 | -2,90 | 3,75E-03 | 2,98E-02 |
| ENSG00000171903 | CYP4F11 | 75,43 | -1,72 | 0,47 | -3,67 | 2,42E-04 | 4,57E-03 |
| ENSG00000205336 | ADGRG1 | 2479,78 | -1,72 | 0,41 | -4,22 | 2,42E-05 | 9,99E-04 |
| ENSG00000148671 | ADIRF | 947,05 | -1,72 | 0,40 | -4,34 | 1,42E-05 | 7,41E-04 |
| ENSG00000183943 | PRKX | 311,37 | -1,72 | 0,48 | -3,62 | 2,90E-04 | 5,09E-03 |
| ENSG00000160183 | TMPRSS3 | 35,12 | -1,72 | 0,64 | -2,70 | 6,90E-03 | 4,53E-02 |
| ENSG00000187837 | H1-2 | 184,03 | -1,72 | 0,48 | -3,57 | 3,56E-04 | 5,87E-03 |
| ENSG00000114737 | CISH | 15,07 | -1,72 | 0,58 | -2,98 | 2,91E-03 | 2,50E-02 |
| ENSG00000173511 | VEGFB | 2545,24 | -1,71 | 0,39 | -4,45 | 8,69E-06 | 5,48E-04 |
| ENSG00000119630 | PGF | 82,69 | -1,71 | 0,47 | -3,63 | 2,81E-04 | 4,97E-03 |
| ENSG00000177045 | SIX5 | 99,04 | -1,71 | 0,46 | -3,71 | 2,06E-04 | 4,10E-03 |
| ENSG00000027075 | PRKCH | 62,03 | -1,71 | 0,35 | -4,92 | 8,74E-07 | 1,29E-04 |
| ENSG00000213949 | ITGA1 | 104,88 | -1,71 | 0,44 | -3,88 | 1,03E-04 | 2,60E-03 |
| ENSG00000178602 | OTOS | 54,30 | -1,71 | 0,56 | -3,05 | 2,32E-03 | 2,15E-02 |
| ENSG00000114738 | MAPKAPK3 | 277,30 | -1,70 | 0,42 | -4,09 | 4,35E-05 | 1,48E-03 |
| ENSG00000270882 | H4C14 | 86,13 | -1,70 | 0,55 | -3,09 | 2,00E-03 | 1,95E-02 |
| ENSG00000102174 | PHEX | 28,87 | -1,70 | 0,52 | -3,26 | 1,13E-03 | 1,30E-02 |
| ENSG00000104518 | GSDMD | 249,39 | -1,70 | 0,45 | -3,80 | 1,42E-04 | 3,22E-03 |
| ENSG00000153822 | KCNJ16 | 98,65 | -1,70 | 0,43 | -3,91 | 9,25E-05 | 2,40E-03 |
| ENSG00000168062 | BATF2 | 30,37 | -1,70 | 0,48 | -3,57 | 3,55E-04 | 5,86E-03 |
| ENSG00000134470 | IL15RA | 33,91 | -1,70 | 0,47 | -3,65 | 2,65E-04 | 4,86E-03 |
| ENSG00000197249 | SERPINA1 | 22,34 | -1,70 | 0,45 | -3,77 | 1,64E-04 | 3,53E-03 |
| ENSG00000132840 | BHMT2 | 61,94 | -1,69 | 0,41 | -4,18 | 2,98E-05 | 1,16E-03 |
| ENSG00000006534 | ALDH3B1 | 105,03 | -1,69 | 0,44 | -3,87 | 1,10E-04 | 2,70E-03 |
| ENSG00000000938 | FGR | 77,43 | -1,69 | 0,40 | -4,18 | 2,88E-05 | 1,14E-03 |
| ENSG00000050327 | ARHGEF5 | 19,47 | -1,69 | 0,56 | -3,02 | 2,53E-03 | 2,26E-02 |
| ENSG00000131941 | RHPN2 | 108,22 | -1,69 | 0,42 | -4,07 | 4,72E-05 | 1,57E-03 |
| ENSG00000270276 | H4C15 | 87,16 | -1,69 | 0,55 | -3,06 | 2,23E-03 | 2,10E-02 |
| ENSG00000197905 | TEAD4 | 26,93 | -1,68 | 0,45 | -3,77 | 1,62E-04 | 3,50E-03 |
| ENSG00000179604 | CDC42EP4 | 1324,52 | -1,68 | 0,38 | -4,42 | 9,86E-06 | 5,90E-04 |
| ENSG00000204592 | HLA-E | 2890,72 | -1,68 | 0,36 | -4,61 | 4,09E-06 | 3,65E-04 |
| ENSG00000109062 | SLC9A3R1 | 1197,40 | -1,68 | 0,41 | -4,13 | 3,68E-05 | 1,32E-03 |
| ENSG00000271447 | MMP28 | 479,20 | -1,67 | 0,42 | -4,00 | 6,46E-05 | 1,92E-03 |
| ENSG00000075234 | TTC38 | 165,45 | -1,67 | 0,41 | -4,03 | 5,63E-05 | 1,76E-03 |
| ENSG00000125089 | SH3TC1 | 104,07 | -1,67 | 0,43 | -3,91 | 9,11E-05 | 2,38E-03 |
| ENSG00000106211 | HSPB1 | 4549,98 | -1,66 | 0,43 | -3,85 | 1,18E-04 | 2,86E-03 |
| ENSG00000154274 | C4orf19 | 96,28 | -1,66 | 0,50 | -3,30 | 9,61E-04 | 1,16E-02 |
| ENSG00000166432 | ZMAT1 | 1299,57 | -1,66 | 0,54 | -3,09 | 2,03E-03 | 1,96E-02 |
| ENSG00000142961 | MOB3C | 137,44 | -1,66 | 0,37 | -4,48 | 7,61E-06 | 5,15E-04 |
| ENSG00000105852 | PON3 | 15,70 | -1,66 | 0,51 | -3,23 | 1,23E-03 | 1,38E-02 |
| ENSG00000277196 | AC007325.2 | 2490,69 | -1,66 | 0,50 | -3,32 | 9,05E-04 | 1,11E-02 |
| ENSG00000167601 | AXL | 446,66 | -1,66 | 0,43 | -3,86 | 1,12E-04 | 2,73E-03 |
| ENSG00000101849 | TBL1X | 423,54 | -1,66 | 0,37 | -4,51 | 6,50E-06 | 4,71E-04 |
| ENSG00000134250 | NOTCH2 | 372,45 | -1,66 | 0,43 | -3,83 | 1,26E-04 | 2,96E-03 |
| ENSG00000170370 | EMX2 | 797,20 | -1,66 | 0,45 | -3,66 | 2,48E-04 | 4,65E-03 |
| ENSG00000125430 | HS3ST3B1 | 21,57 | -1,66 | 0,51 | -3,24 | 1,20E-03 | 1,36E-02 |
| ENSG00000170759 | KIF5B | 4724,03 | -1,65 | 0,36 | -4,55 | 5,35E-06 | 4,09E-04 |
| ENSG00000183876 | ARSI | 56,08 | -1,65 | 0,50 | -3,29 | 9,96E-04 | 1,19E-02 |
| ENSG00000156535 | CD109 | 40,81 | -1,65 | 0,42 | -3,93 | 8,67E-05 | 2,29E-03 |
| ENSG00000122122 | SASH3 | 35,08 | -1,65 | 0,52 | -3,17 | 1,51E-03 | 1,61E-02 |
| ENSG00000078401 | EDN1 | 84,65 | -1,65 | 0,42 | -3,98 | 6,94E-05 | 2,00E-03 |
| ENSG00000131981 | LGALS3 | 367,28 | -1,65 | 0,34 | -4,81 | 1,53E-06 | 1,89E-04 |
| ENSG00000173473 | SMARCC1 | 1098,53 | -1,65 | 0,43 | -3,80 | 1,43E-04 | 3,23E-03 |
| ENSG00000134569 | LRP4 | 828,34 | -1,65 | 0,39 | -4,28 | 1,84E-05 | 8,80E-04 |
| ENSG00000125878 | TCF15 | 18,99 | -1,65 | 0,50 | -3,31 | 9,46E-04 | 1,15E-02 |
| ENSG00000089159 | PXN | 660,72 | -1,65 | 0,38 | -4,38 | 1,19E-05 | 6,66E-04 |
| ENSG00000068078 | FGFR3 | 3486,38 | -1,65 | 0,44 | -3,71 | 2,04E-04 | 4,09E-03 |

| | | | | | | | |
|---|---|---|---|---|---|---|---|
| ENSG00000174059 | CD34 | 265,01 | -1,65 | 0,30 | -5,49 | 3,92E-08 | 1,62E-05 |
| ENSG00000188643 | S100A16 | 750,38 | -1,65 | 0,38 | -4,38 | 1,18E-05 | 6,65E-04 |
| ENSG00000184232 | OAF | 868,03 | -1,64 | 0,41 | -3,97 | 7,30E-05 | 2,05E-03 |
| ENSG00000132256 | TRIM5 | 21,45 | -1,64 | 0,41 | -4,02 | 5,77E-05 | 1,79E-03 |
| ENSG00000196139 | AKR1C3 | 162,76 | -1,64 | 0,44 | -3,75 | 1,78E-04 | 3,75E-03 |
| ENSG00000183801 | OLFML1 | 29,06 | -1,64 | 0,55 | -2,97 | 3,02E-03 | 2,56E-02 |
| ENSG00000187942 | LDLRAD2 | 194,90 | -1,64 | 0,47 | -3,52 | 4,38E-04 | 6,74E-03 |
| ENSG00000148841 | ITPRIP | 67,57 | -1,64 | 0,41 | -3,96 | 7,60E-05 | 2,09E-03 |
| ENSG00000002586 | CD99 | 945,48 | -1,64 | 0,37 | -4,43 | 9,62E-06 | 5,84E-04 |
| ENSG00000169515 | CCDC8 | 75,55 | -1,63 | 0,48 | -3,43 | 6,12E-04 | 8,34E-03 |
| ENSG00000105137 | SYDE1 | 133,61 | -1,63 | 0,41 | -3,96 | 7,52E-05 | 2,09E-03 |
| ENSG00000213853 | EMP2 | 354,18 | -1,63 | 0,39 | -4,21 | 2,53E-05 | 1,04E-03 |
| ENSG00000248871 | TNFSF12-TNFSF1 | 906,49 | -1,63 | 0,36 | -4,48 | 7,46E-06 | 5,09E-04 |
| ENSG00000138823 | MTTP | 22,61 | -1,63 | 0,55 | -2,98 | 2,92E-03 | 2,50E-02 |
| ENSG00000074181 | NOTCH3 | 428,78 | -1,63 | 0,32 | -5,01 | 5,44E-07 | 9,59E-05 |
| ENSG00000261371 | PECAM1 | 341,74 | -1,63 | 0,31 | -5,28 | 1,30E-07 | 3,30E-05 |
| ENSG00000126016 | AMOT | 587,75 | -1,63 | 0,35 | -4,63 | 3,67E-06 | 3,36E-04 |
| ENSG00000099377 | HSD3B7 | 119,64 | -1,63 | 0,46 | -3,51 | 4,47E-04 | 6,80E-03 |
| ENSG00000158710 | TAGLN2 | 575,02 | -1,63 | 0,32 | -5,06 | 4,22E-07 | 8,14E-05 |
| ENSG00000142173 | COL6A2 | 445,96 | -1,62 | 0,46 | -3,57 | 3,63E-04 | 5,97E-03 |
| ENSG00000149489 | ROM1 | 517,56 | -1,62 | 0,40 | -4,02 | 5,81E-05 | 1,79E-03 |
| ENSG00000164850 | GPER1 | 185,11 | -1,62 | 0,36 | -4,46 | 8,16E-06 | 5,39E-04 |
| ENSG00000150907 | FOXO1 | 234,72 | -1,62 | 0,38 | -4,22 | 2,43E-05 | 9,99E-04 |
| ENSG00000126562 | WNK4 | 34,95 | -1,62 | 0,43 | -3,71 | 2,04E-04 | 4,08E-03 |
| ENSG00000138496 | PARP9 | 85,60 | -1,62 | 0,40 | -4,01 | 6,20E-05 | 1,87E-03 |
| ENSG00000100342 | APOL1 | 141,92 | -1,62 | 0,41 | -3,95 | 7,79E-05 | 2,13E-03 |
| ENSG00000181781 | ODF3L2 | 37,45 | -1,62 | 0,59 | -2,75 | 6,05E-03 | 4,11E-02 |
| ENSG00000047849 | MAP4 | 10581,38 | -1,61 | 0,29 | -5,59 | 2,33E-08 | 1,41E-05 |
| ENSG00000092820 | EZR | 1777,64 | -1,61 | 0,42 | -3,87 | 1,09E-04 | 2,69E-03 |
| ENSG00000162733 | DDR2 | 212,38 | -1,61 | 0,37 | -4,39 | 1,16E-05 | 6,61E-04 |
| ENSG00000189129 | PLAC9 | 128,05 | -1,61 | 0,50 | -3,24 | 1,21E-03 | 1,37E-02 |
| ENSG00000185291 | IL3RA | 411,16 | -1,61 | 0,52 | -3,07 | 2,14E-03 | 2,04E-02 |
| ENSG00000162889 | MAPKAPK2 | 696,63 | -1,61 | 0,35 | -4,55 | 5,26E-06 | 4,05E-04 |
| ENSG00000005243 | COPZ2 | 146,05 | -1,61 | 0,41 | -3,91 | 9,19E-05 | 2,40E-03 |
| ENSG00000140853 | NLRC5 | 105,73 | -1,61 | 0,46 | -3,53 | 4,20E-04 | 6,56E-03 |
| ENSG00000071054 | MAP4K4 | 10340,31 | -1,61 | 0,44 | -3,61 | 3,02E-04 | 5,24E-03 |
| ENSG00000108375 | RNF43 | 70,03 | -1,61 | 0,49 | -3,26 | 1,11E-03 | 1,28E-02 |
| ENSG00000111907 | TPD52L1 | 671,26 | -1,61 | 0,39 | -4,17 | 3,09E-05 | 1,18E-03 |
| ENSG00000181722 | ZBTB20 | 2617,97 | -1,60 | 0,46 | -3,51 | 4,57E-04 | 6,91E-03 |
| ENSG00000117228 | GBP1 | 39,14 | -1,60 | 0,45 | -3,58 | 3,49E-04 | 5,78E-03 |
| ENSG00000174963 | ZIC4 | 134,87 | -1,60 | 0,41 | -3,93 | 8,57E-05 | 2,27E-03 |
| ENSG00000087077 | TRIP6 | 419,45 | -1,60 | 0,36 | -4,43 | 9,37E-06 | 5,77E-04 |
| ENSG00000183230 | CTNNA3 | 291,72 | -1,60 | 0,46 | -3,47 | 5,16E-04 | 7,45E-03 |
| ENSG00000106714 | CNTNAP3 | 172,86 | -1,60 | 0,52 | -3,09 | 2,03E-03 | 1,96E-02 |
| ENSG00000131196 | NFATC1 | 113,44 | -1,60 | 0,42 | -3,79 | 1,51E-04 | 3,36E-03 |
| ENSG00000000971 | CFH | 88,78 | -1,60 | 0,39 | -4,07 | 4,75E-05 | 1,57E-03 |
| ENSG00000149548 | CCDC15 | 50,95 | -1,60 | 0,57 | -2,81 | 4,98E-03 | 3,59E-02 |
| ENSG00000137198 | GMPR | 182,82 | -1,60 | 0,47 | -3,37 | 7,44E-04 | 9,63E-03 |
| ENSG00000164708 | PGAM2 | 315,43 | -1,60 | 0,44 | -3,66 | 2,51E-04 | 4,69E-03 |
| ENSG00000134548 | SPX | 218,71 | -1,60 | 0,31 | -5,21 | 1,91E-07 | 4,47E-05 |
| ENSG00000186998 | EMID1 | 1158,06 | -1,60 | 0,42 | -3,81 | 1,38E-04 | 3,16E-03 |
| ENSG00000165795 | NDRG2 | 23573,48 | -1,60 | 0,33 | -4,81 | 1,51E-06 | 1,88E-04 |
| ENSG00000037280 | FLT4 | 77,11 | -1,60 | 0,44 | -3,65 | 2,64E-04 | 4,84E-03 |
| ENSG00000090924 | PLEKHG2 | 65,91 | -1,60 | 0,44 | -3,61 | 3,05E-04 | 5,27E-03 |
| ENSG00000117318 | ID3 | 283,42 | -1,59 | 0,47 | -3,37 | 7,43E-04 | 9,63E-03 |
| ENSG00000115414 | FN1 | 627,82 | -1,59 | 0,37 | -4,35 | 1,37E-05 | 7,28E-04 |
| ENSG00000129151 | BBOX1 | 135,16 | -1,59 | 0,36 | -4,46 | 8,38E-06 | 5,43E-04 |

| | | | | | | | |
|---|---|---|---|---|---|---|---|
| ENSG00000003989 | SLC7A2 | 174,92 | -1,59 | 0,42 | -3,82 | 1,36E-04 | 3,12E-03 |
| ENSG00000163362 | INAVA | 35,91 | -1,59 | 0,54 | -2,95 | 3,14E-03 | 2,63E-02 |
| ENSG00000196586 | MYO6 | 6320,86 | -1,59 | 0,58 | -2,72 | 6,49E-03 | 4,32E-02 |
| ENSG00000104870 | FCGRT | 579,40 | -1,59 | 0,41 | -3,88 | 1,03E-04 | 2,58E-03 |
| ENSG00000133226 | SRRM1 | 4773,51 | -1,58 | 0,55 | -2,89 | 3,85E-03 | 3,02E-02 |
| ENSG00000171940 | ZNF217 | 40,38 | -1,58 | 0,36 | -4,40 | 1,09E-05 | 6,34E-04 |
| ENSG00000169249 | ZRSR2 | 1078,96 | -1,58 | 0,50 | -3,16 | 1,58E-03 | 1,67E-02 |
| ENSG00000219607 | PPP1R3G | 121,75 | -1,58 | 0,46 | -3,40 | 6,67E-04 | 8,86E-03 |
| ENSG00000158423 | RIBC1 | 17,38 | -1,58 | 0,48 | -3,32 | 8,93E-04 | 1,10E-02 |
| ENSG00000160951 | PTGER1 | 16,06 | -1,58 | 0,55 | -2,86 | 4,26E-03 | 3,24E-02 |
| ENSG00000135063 | FAM189A2 | 236,50 | -1,58 | 0,45 | -3,54 | 4,01E-04 | 6,36E-03 |
| ENSG00000135976 | ANKRD36 | 4411,61 | -1,58 | 0,57 | -2,77 | 5,55E-03 | 3,87E-02 |
| ENSG00000100767 | PAPLN | 281,30 | -1,57 | 0,48 | -3,30 | 9,81E-04 | 1,18E-02 |
| ENSG00000154133 | ROBO4 | 241,59 | -1,57 | 0,37 | -4,26 | 2,01E-05 | 9,23E-04 |
| ENSG00000172638 | EFEMP2 | 766,13 | -1,57 | 0,39 | -4,04 | 5,33E-05 | 1,69E-03 |
| ENSG00000198075 | SULT1C4 | 121,48 | -1,57 | 0,35 | -4,55 | 5,28E-06 | 4,05E-04 |
| ENSG00000008394 | MGST1 | 448,35 | -1,57 | 0,40 | -3,97 | 7,06E-05 | 2,02E-03 |
| ENSG00000125347 | IRF1 | 211,73 | -1,57 | 0,43 | -3,68 | 2,33E-04 | 4,47E-03 |
| ENSG00000101017 | CD40 | 54,12 | -1,57 | 0,40 | -3,96 | 7,40E-05 | 2,07E-03 |
| ENSG00000154945 | ANKRD40 | 2084,53 | -1,57 | 0,33 | -4,69 | 2,71E-06 | 2,69E-04 |
| ENSG00000143819 | EPHX1 | 2579,16 | -1,57 | 0,41 | -3,80 | 1,45E-04 | 3,26E-03 |
| ENSG00000135269 | TES | 13,92 | -1,57 | 0,47 | -3,32 | 9,13E-04 | 1,12E-02 |
| ENSG00000153914 | SREK1 | 5020,77 | -1,56 | 0,56 | -2,82 | 4,86E-03 | 3,53E-02 |
| ENSG00000188176 | SMTNL2 | 11,97 | -1,56 | 0,53 | -2,96 | 3,09E-03 | 2,60E-02 |
| ENSG00000137767 | SQOR | 58,63 | -1,56 | 0,37 | -4,25 | 2,17E-05 | 9,45E-04 |
| ENSG00000138119 | MYOF | 41,61 | -1,56 | 0,51 | -3,08 | 2,04E-03 | 1,97E-02 |
| ENSG00000188338 | SLC38A3 | 595,21 | -1,56 | 0,36 | -4,33 | 1,47E-05 | 7,52E-04 |
| ENSG00000286219 | NOTCH2NLC | 151,93 | -1,56 | 0,40 | -3,90 | 9,57E-05 | 2,47E-03 |
| ENSG00000170412 | GPRC5C | 195,89 | -1,56 | 0,40 | -3,89 | 9,97E-05 | 2,54E-03 |
| ENSG00000007372 | PAX6 | 678,98 | -1,56 | 0,41 | -3,82 | 1,33E-04 | 3,10E-03 |
| ENSG00000114857 | NKTR | 2992,16 | -1,56 | 0,54 | -2,88 | 3,95E-03 | 3,07E-02 |
| ENSG00000130222 | GADD45G | 281,05 | -1,56 | 0,42 | -3,68 | 2,32E-04 | 4,46E-03 |
| ENSG00000125319 | HROB | 73,47 | -1,55 | 0,58 | -2,69 | 7,19E-03 | 4,66E-02 |
| ENSG00000157833 | GAREM2 | 652,69 | -1,55 | 0,44 | -3,50 | 4,68E-04 | 7,02E-03 |
| ENSG00000205669 | ACOT6 | 24,17 | -1,55 | 0,56 | -2,75 | 6,05E-03 | 4,11E-02 |
| ENSG00000135736 | CCDC102A | 32,59 | -1,55 | 0,45 | -3,43 | 6,05E-04 | 8,32E-03 |
| ENSG00000139800 | ZIC5 | 44,66 | -1,55 | 0,45 | -3,42 | 6,23E-04 | 8,45E-03 |
| ENSG00000184154 | LRTOMT | 242,63 | -1,55 | 0,34 | -4,56 | 5,18E-06 | 4,05E-04 |
| ENSG00000153944 | MSI2 | 1126,22 | -1,55 | 0,35 | -4,43 | 9,22E-06 | 5,73E-04 |
| ENSG00000177575 | CD163 | 11,04 | -1,54 | 0,55 | -2,79 | 5,28E-03 | 3,74E-02 |
| ENSG00000271254 | AC240274.1 | 637,42 | -1,54 | 0,55 | -2,79 | 5,23E-03 | 3,71E-02 |
| ENSG00000186951 | PPARA | 321,17 | -1,54 | 0,41 | -3,72 | 1,99E-04 | 4,02E-03 |
| ENSG00000172987 | HPSE2 | 213,31 | -1,54 | 0,48 | -3,24 | 1,19E-03 | 1,36E-02 |
| ENSG00000166927 | MS4A7 | 33,55 | -1,54 | 0,54 | -2,83 | 4,61E-03 | 3,43E-02 |
| ENSG00000067900 | ROCK1 | 1051,33 | -1,54 | 0,43 | -3,61 | 3,07E-04 | 5,29E-03 |
| ENSG00000107736 | CDH23 | 191,46 | -1,54 | 0,42 | -3,64 | 2,77E-04 | 4,95E-03 |
| ENSG00000128284 | APOL3 | 104,90 | -1,54 | 0,32 | -4,75 | 2,05E-06 | 2,22E-04 |
| ENSG00000264343 | NOTCH2NLA | 157,00 | -1,54 | 0,42 | -3,68 | 2,36E-04 | 4,50E-03 |
| ENSG00000105974 | CAV1 | 169,26 | -1,54 | 0,56 | -2,75 | 5,93E-03 | 4,06E-02 |
| ENSG00000173641 | HSPB7 | 48,06 | -1,54 | 0,39 | -3,92 | 8,99E-05 | 2,36E-03 |
| ENSG00000198399 | ITSN2 | 2421,35 | -1,54 | 0,48 | -3,17 | 1,53E-03 | 1,62E-02 |
| ENSG00000140511 | HAPLN3 | 62,84 | -1,54 | 0,46 | -3,36 | 7,73E-04 | 9,91E-03 |
| ENSG00000082074 | FYB1 | 210,71 | -1,53 | 0,51 | -3,02 | 2,54E-03 | 2,27E-02 |
| ENSG00000078177 | N4BP2 | 43,33 | -1,53 | 0,44 | -3,47 | 5,25E-04 | 7,52E-03 |
| ENSG00000164430 | CGAS | 17,19 | -1,53 | 0,53 | -2,89 | 3,91E-03 | 3,05E-02 |
| ENSG00000205730 | ITPRIPL2 | 126,09 | -1,53 | 0,42 | -3,68 | 2,33E-04 | 4,47E-03 |
| ENSG00000187144 | SPATA21 | 245,15 | -1,52 | 0,34 | -4,47 | 7,94E-06 | 5,34E-04 |

| | | | | | | | |
|---|---|---|---|---|---|---|---|
| ENSG00000133321 | PLAAT4 | 280,86 | -1,52 | 0,46 | -3,32 | 9,11E-04 | 1,12E-02 |
| ENSG00000106538 | RARRES2 | 627,44 | -1,52 | 0,42 | -3,59 | 3,32E-04 | 5,61E-03 |
| ENSG00000140682 | TGFB1I1 | 73,44 | -1,52 | 0,41 | -3,71 | 2,09E-04 | 4,14E-03 |
| ENSG00000134184 | GSTM1 | 1943,15 | -1,52 | 0,51 | -2,97 | 3,01E-03 | 2,55E-02 |
| ENSG00000198142 | SOWAHC | 86,26 | -1,52 | 0,35 | -4,29 | 1,75E-05 | 8,55E-04 |
| ENSG00000105939 | ZC3HAV1 | 145,48 | -1,52 | 0,38 | -4,03 | 5,68E-05 | 1,77E-03 |
| ENSG00000140450 | ARRDC4 | 263,26 | -1,52 | 0,32 | -4,76 | 1,90E-06 | 2,14E-04 |
| ENSG00000100439 | ABHD4 | 259,52 | -1,52 | 0,35 | -4,32 | 1,58E-05 | 7,88E-04 |
| ENSG00000130283 | GDF1 | 4657,36 | -1,52 | 0,37 | -4,12 | 3,71E-05 | 1,33E-03 |
| ENSG00000107438 | PDLIM1 | 64,38 | -1,52 | 0,41 | -3,71 | 2,08E-04 | 4,14E-03 |
| ENSG00000184481 | FOXO4 | 1082,25 | -1,52 | 0,33 | -4,58 | 4,56E-06 | 3,82E-04 |
| ENSG00000197712 | FAM114A1 | 210,10 | -1,52 | 0,44 | -3,46 | 5,32E-04 | 7,58E-03 |
| ENSG00000198805 | PNP | 92,31 | -1,51 | 0,44 | -3,47 | 5,13E-04 | 7,43E-03 |
| ENSG00000156218 | ADAMTSL3 | 41,71 | -1,51 | 0,50 | -3,01 | 2,57E-03 | 2,29E-02 |
| ENSG00000169006 | NTSR2 | 632,23 | -1,51 | 0,41 | -3,73 | 1,91E-04 | 3,95E-03 |
| ENSG00000168497 | CAVIN2 | 167,28 | -1,51 | 0,33 | -4,60 | 4,14E-06 | 3,67E-04 |
| ENSG00000113916 | BCL6 | 894,39 | -1,51 | 0,35 | -4,37 | 1,25E-05 | 6,96E-04 |
| ENSG00000103257 | SLC7A5 | 2269,97 | -1,51 | 0,32 | -4,66 | 3,16E-06 | 3,02E-04 |
| ENSG00000135821 | GLUL | 22062,59 | -1,51 | 0,42 | -3,61 | 3,07E-04 | 5,29E-03 |
| ENSG00000148411 | NACC2 | 1695,37 | -1,51 | 0,31 | -4,84 | 1,32E-06 | 1,73E-04 |
| ENSG00000170989 | S1PR1 | 496,79 | -1,51 | 0,38 | -3,95 | 7,85E-05 | 2,14E-03 |
| ENSG00000114698 | PLSCR4 | 148,74 | -1,51 | 0,33 | -4,52 | 6,32E-06 | 4,64E-04 |
| ENSG00000173918 | C1QTNF1 | 161,74 | -1,51 | 0,39 | -3,90 | 9,79E-05 | 2,51E-03 |
| ENSG00000112559 | MDFI | 50,61 | -1,51 | 0,37 | -4,05 | 5,22E-05 | 1,67E-03 |
| ENSG00000144485 | HES6 | 1194,98 | -1,50 | 0,44 | -3,43 | 6,01E-04 | 8,29E-03 |
| ENSG00000120896 | SORBS3 | 1431,28 | -1,50 | 0,34 | -4,43 | 9,39E-06 | 5,77E-04 |
| ENSG00000164023 | SGMS2 | 190,31 | -1,50 | 0,36 | -4,17 | 3,04E-05 | 1,17E-03 |
| ENSG00000170915 | PAQR8 | 2279,94 | -1,50 | 0,37 | -4,02 | 5,80E-05 | 1,79E-03 |
| ENSG00000116754 | SRSF11 | 10838,62 | -1,50 | 0,56 | -2,68 | 7,27E-03 | 4,69E-02 |
| ENSG00000055070 | SZRD1 | 1026,57 | -1,50 | 0,33 | -4,56 | 5,21E-06 | 4,05E-04 |
| ENSG00000181284 | TMEM102 | 30,50 | -1,50 | 0,47 | -3,19 | 1,42E-03 | 1,54E-02 |
| ENSG00000142227 | EMP3 | 163,96 | -1,49 | 0,43 | -3,50 | 4,64E-04 | 6,98E-03 |
| ENSG00000031081 | ARHGAP31 | 182,42 | -1,49 | 0,38 | -3,97 | 7,29E-05 | 2,05E-03 |
| ENSG00000158769 | F11R | 167,34 | -1,49 | 0,40 | -3,69 | 2,24E-04 | 4,35E-03 |
| ENSG00000043355 | ZIC2 | 211,48 | -1,49 | 0,41 | -3,62 | 2,91E-04 | 5,09E-03 |
| ENSG00000223953 | C1QTNF5 | 271,44 | -1,49 | 0,41 | -3,60 | 3,20E-04 | 5,45E-03 |
| ENSG00000119408 | NEK6 | 350,57 | -1,49 | 0,37 | -4,05 | 5,07E-05 | 1,64E-03 |
| ENSG00000142798 | HSPG2 | 248,08 | -1,49 | 0,42 | -3,55 | 3,83E-04 | 6,19E-03 |
| ENSG00000168306 | ACOX2 | 49,48 | -1,49 | 0,47 | -3,19 | 1,45E-03 | 1,56E-02 |
| ENSG00000126458 | RRAS | 366,28 | -1,49 | 0,35 | -4,21 | 2,61E-05 | 1,05E-03 |
| ENSG00000160447 | PKN3 | 99,62 | -1,49 | 0,45 | -3,29 | 1,02E-03 | 1,21E-02 |
| ENSG00000126870 | DYNC2I1 | 3008,09 | -1,49 | 0,53 | -2,81 | 4,98E-03 | 3,59E-02 |
| ENSG00000091986 | CCDC80 | 291,78 | -1,49 | 0,49 | -3,06 | 2,19E-03 | 2,07E-02 |
| ENSG00000146250 | PRSS35 | 45,26 | -1,49 | 0,42 | -3,51 | 4,55E-04 | 6,89E-03 |
| ENSG00000106991 | ENG | 670,22 | -1,49 | 0,34 | -4,32 | 1,55E-05 | 7,79E-04 |
| ENSG00000142632 | ARHGEF19 | 27,91 | -1,48 | 0,46 | -3,20 | 1,36E-03 | 1,49E-02 |
| ENSG00000140067 | FAM181A | 133,73 | -1,48 | 0,40 | -3,69 | 2,26E-04 | 4,36E-03 |
| ENSG00000148488 | ST8SIA6 | 37,37 | -1,48 | 0,37 | -4,01 | 6,09E-05 | 1,86E-03 |
| ENSG00000235718 | MFRP | 272,14 | -1,48 | 0,41 | -3,58 | 3,45E-04 | 5,75E-03 |
| ENSG00000142910 | TINAGL1 | 211,87 | -1,48 | 0,39 | -3,82 | 1,34E-04 | 3,10E-03 |
| ENSG00000161714 | PLCD3 | 1174,04 | -1,48 | 0,34 | -4,34 | 1,40E-05 | 7,39E-04 |
| ENSG00000132424 | PNISR | 8168,12 | -1,48 | 0,54 | -2,75 | 6,00E-03 | 4,09E-02 |
| ENSG00000162614 | NEXN | 186,63 | -1,48 | 0,49 | -3,05 | 2,28E-03 | 2,13E-02 |
| ENSG00000133739 | LRRCC1 | 377,85 | -1,48 | 0,40 | -3,67 | 2,44E-04 | 4,60E-03 |
| ENSG00000233670 | PIRT | 24,80 | -1,48 | 0,45 | -3,32 | 8,87E-04 | 1,09E-02 |
| ENSG00000115325 | DOK1 | 105,69 | -1,48 | 0,41 | -3,61 | 3,11E-04 | 5,35E-03 |
| ENSG00000179776 | CDH5 | 132,88 | -1,48 | 0,30 | -4,91 | 9,26E-07 | 1,34E-04 |

| | | | | | | | |
|---|---|---|---|---|---|---|---|
| ENSG00000168314 | MOBP | 11308,81 | -1,48 | 0,42 | -3,53 | 4,11E-04 | 6,46E-03 |
| ENSG00000119699 | TGFB3 | 490,81 | -1,48 | 0,38 | -3,90 | 9,73E-05 | 2,50E-03 |
| ENSG00000159388 | BTG2 | 322,77 | -1,48 | 0,32 | -4,60 | 4,32E-06 | 3,74E-04 |
| ENSG00000164035 | EMCN | 116,05 | -1,48 | 0,40 | -3,69 | 2,23E-04 | 4,32E-03 |
| ENSG00000165449 | SLC16A9 | 120,64 | -1,47 | 0,45 | -3,29 | 9,85E-04 | 1,18E-02 |
| ENSG00000176105 | YES1 | 180,34 | -1,47 | 0,27 | -5,53 | 3,25E-08 | 1,46E-05 |
| ENSG00000138449 | SLC40A1 | 58,36 | -1,47 | 0,33 | -4,45 | 8,42E-06 | 5,44E-04 |
| ENSG00000113578 | FGF1 | 2700,79 | -1,47 | 0,33 | -4,46 | 8,10E-06 | 5,39E-04 |
| ENSG00000168918 | INPP5D | 271,23 | -1,47 | 0,46 | -3,16 | 1,59E-03 | 1,67E-02 |
| ENSG00000021300 | PLEKHB1 | 12015,07 | -1,47 | 0,30 | -4,94 | 7,78E-07 | 1,21E-04 |
| ENSG00000170075 | GPR37L1 | 4589,21 | -1,47 | 0,40 | -3,70 | 2,18E-04 | 4,27E-03 |
| ENSG00000092969 | TGFB2 | 247,91 | -1,47 | 0,36 | -4,10 | 4,14E-05 | 1,44E-03 |
| ENSG00000143514 | TP53BP2 | 681,86 | -1,46 | 0,37 | -4,01 | 6,18E-05 | 1,87E-03 |
| ENSG00000002933 | TMEM176A | 236,26 | -1,46 | 0,41 | -3,56 | 3,73E-04 | 6,07E-03 |
| ENSG00000131097 | HIGD1B | 165,47 | -1,46 | 0,36 | -4,03 | 5,58E-05 | 1,76E-03 |
| ENSG00000105227 | PRX | 168,39 | -1,46 | 0,42 | -3,45 | 5,54E-04 | 7,82E-03 |
| ENSG00000054598 | FOXC1 | 85,94 | -1,46 | 0,45 | -3,26 | 1,13E-03 | 1,30E-02 |
| ENSG00000143036 | SLC44A3 | 57,74 | -1,46 | 0,32 | -4,49 | 6,97E-06 | 4,92E-04 |
| ENSG00000178764 | ZHX2 | 339,23 | -1,46 | 0,31 | -4,65 | 3,37E-06 | 3,20E-04 |
| ENSG00000111321 | LTBR | 179,32 | -1,46 | 0,36 | -4,09 | 4,32E-05 | 1,47E-03 |
| ENSG00000136938 | ANP32B | 2374,78 | -1,46 | 0,40 | -3,64 | 2,70E-04 | 4,89E-03 |
| ENSG00000178075 | GRAMD1C | 152,63 | -1,45 | 0,39 | -3,70 | 2,19E-04 | 4,27E-03 |
| ENSG00000114315 | HES1 | 496,53 | -1,45 | 0,46 | -3,16 | 1,57E-03 | 1,66E-02 |
| ENSG00000091409 | ITGA6 | 249,05 | -1,45 | 0,35 | -4,16 | 3,15E-05 | 1,19E-03 |
| ENSG00000124508 | BTN2A2 | 46,97 | -1,45 | 0,35 | -4,15 | 3,29E-05 | 1,22E-03 |
| ENSG00000178878 | APOLD1 | 802,27 | -1,45 | 0,32 | -4,57 | 4,85E-06 | 3,95E-04 |
| ENSG00000172728 | FUT10 | 42,68 | -1,45 | 0,37 | -3,96 | 7,48E-05 | 2,08E-03 |
| ENSG00000186198 | SLC51B | 37,36 | -1,45 | 0,48 | -3,05 | 2,32E-03 | 2,15E-02 |
| ENSG00000064655 | EYA2 | 114,57 | -1,44 | 0,39 | -3,73 | 1,94E-04 | 3,97E-03 |
| ENSG00000104218 | CSPP1 | 1818,05 | -1,44 | 0,53 | -2,75 | 6,04E-03 | 4,11E-02 |
| ENSG00000142694 | EVA1B | 210,32 | -1,44 | 0,47 | -3,07 | 2,17E-03 | 2,06E-02 |
| ENSG00000186470 | BTN3A2 | 215,28 | -1,44 | 0,40 | -3,64 | 2,75E-04 | 4,91E-03 |
| ENSG00000100906 | NFKBIA | 748,39 | -1,44 | 0,42 | -3,43 | 6,07E-04 | 8,32E-03 |
| ENSG00000067182 | TNFRSF1A | 295,84 | -1,44 | 0,36 | -3,99 | 6,68E-05 | 1,96E-03 |
| ENSG00000132613 | MTSS2 | 4848,60 | -1,44 | 0,30 | -4,74 | 2,09E-06 | 2,25E-04 |
| ENSG00000126603 | GLIS2 | 227,78 | -1,44 | 0,33 | -4,35 | 1,34E-05 | 7,26E-04 |
| ENSG00000158825 | CDA | 29,24 | -1,44 | 0,43 | -3,36 | 7,85E-04 | 1,00E-02 |
| ENSG00000117523 | PRRC2C | 9292,77 | -1,44 | 0,48 | -3,01 | 2,65E-03 | 2,33E-02 |
| ENSG00000091317 | CMTM6 | 251,75 | -1,44 | 0,39 | -3,65 | 2,59E-04 | 4,80E-03 |
| ENSG00000148175 | STOM | 669,87 | -1,44 | 0,31 | -4,69 | 2,77E-06 | 2,74E-04 |
| ENSG00000134824 | FADS2 | 2407,02 | -1,44 | 0,32 | -4,49 | 7,25E-06 | 5,03E-04 |
| ENSG00000173193 | PARP14 | 140,10 | -1,44 | 0,35 | -4,15 | 3,29E-05 | 1,22E-03 |
| ENSG00000162496 | DHRS3 | 505,32 | -1,44 | 0,38 | -3,81 | 1,39E-04 | 3,16E-03 |
| ENSG00000132950 | ZMYM5 | 146,60 | -1,44 | 0,41 | -3,52 | 4,35E-04 | 6,72E-03 |
| ENSG00000143384 | MCL1 | 863,69 | -1,44 | 0,30 | -4,80 | 1,63E-06 | 1,91E-04 |
| ENSG00000154319 | FAM167A | 363,28 | -1,44 | 0,31 | -4,59 | 4,35E-06 | 3,74E-04 |
| ENSG00000278662 | GOLGA6L10 | 63,78 | -1,44 | 0,49 | -2,90 | 3,72E-03 | 2,97E-02 |
| ENSG00000288678 | ENSG000002886 | 61,29 | -1,43 | 0,38 | -3,78 | 1,57E-04 | 3,44E-03 |
| ENSG00000084093 | REST | 269,90 | -1,43 | 0,42 | -3,43 | 6,09E-04 | 8,34E-03 |
| ENSG00000117394 | SLC2A1 | 1084,09 | -1,43 | 0,29 | -4,88 | 1,06E-06 | 1,46E-04 |
| ENSG00000005981 | ASB4 | 22,28 | -1,43 | 0,49 | -2,91 | 3,61E-03 | 2,91E-02 |
| ENSG00000147065 | MSN | 712,47 | -1,43 | 0,26 | -5,56 | 2,67E-08 | 1,41E-05 |
| ENSG00000168994 | PXDC1 | 259,98 | -1,43 | 0,34 | -4,18 | 2,87E-05 | 1,14E-03 |
| ENSG00000179144 | GIMAP7 | 98,93 | -1,43 | 0,41 | -3,47 | 5,18E-04 | 7,45E-03 |
| ENSG00000214447 | FAM187A | 181,62 | -1,43 | 0,41 | -3,52 | 4,26E-04 | 6,62E-03 |
| ENSG00000100033 | PRODH | 1757,75 | -1,43 | 0,50 | -2,89 | 3,90E-03 | 3,04E-02 |
| ENSG00000131634 | TMEM204 | 120,66 | -1,43 | 0,39 | -3,63 | 2,88E-04 | 5,08E-03 |

| | | | | | | | |
|---|---|---|---|---|---|---|---|
| ENSG00000082781 | ITGB5 | 411,35 | -1,43 | 0,33 | -4,27 | 1,93E-05 | 9,02E-04 |
| ENSG00000150760 | DOCK1 | 605,96 | -1,43 | 0,29 | -5,01 | 5,43E-07 | 9,59E-05 |
| ENSG00000081059 | TCF7 | 49,28 | -1,43 | 0,38 | -3,76 | 1,70E-04 | 3,64E-03 |
| ENSG00000132164 | SLC6A11 | 476,48 | -1,43 | 0,36 | -3,96 | 7,42E-05 | 2,07E-03 |
| ENSG00000166825 | ANPEP | 41,25 | -1,43 | 0,51 | -2,80 | 5,14E-03 | 3,66E-02 |
| ENSG00000147588 | PMP2 | 2521,19 | -1,43 | 0,32 | -4,53 | 5,94E-06 | 4,42E-04 |
| ENSG00000163132 | MSX1 | 180,34 | -1,43 | 0,46 | -3,10 | 1,94E-03 | 1,91E-02 |
| ENSG00000119686 | FLVCR2 | 19,86 | -1,43 | 0,50 | -2,84 | 4,55E-03 | 3,39E-02 |
| ENSG00000186994 | KANK3 | 276,33 | -1,43 | 0,33 | -4,31 | 1,60E-05 | 7,96E-04 |
| ENSG00000167131 | CCDC103 | 183,95 | -1,43 | 0,40 | -3,54 | 4,06E-04 | 6,40E-03 |
| ENSG00000198121 | LPAR1 | 2226,14 | -1,42 | 0,40 | -3,56 | 3,67E-04 | 6,00E-03 |
| ENSG00000132669 | RIN2 | 249,34 | -1,42 | 0,33 | -4,25 | 2,13E-05 | 9,43E-04 |
| ENSG00000234465 | PINLYP | 178,02 | -1,42 | 0,51 | -2,79 | 5,26E-03 | 3,73E-02 |
| ENSG00000127241 | MASP1 | 108,90 | -1,42 | 0,35 | -4,03 | 5,65E-05 | 1,76E-03 |
| ENSG00000132000 | PODNL1 | 51,05 | -1,42 | 0,47 | -3,01 | 2,65E-03 | 2,33E-02 |
| ENSG00000213654 | GPSM3 | 163,16 | -1,42 | 0,40 | -3,52 | 4,27E-04 | 6,63E-03 |
| ENSG00000183770 | FOXL2 | 21,68 | -1,42 | 0,50 | -2,84 | 4,49E-03 | 3,36E-02 |
| ENSG00000096060 | FKBP5 | 279,77 | -1,42 | 0,51 | -2,77 | 5,67E-03 | 3,93E-02 |
| ENSG00000100106 | TRIOBP | 392,10 | -1,42 | 0,39 | -3,67 | 2,43E-04 | 4,57E-03 |
| ENSG00000109113 | RAB34 | 354,70 | -1,42 | 0,41 | -3,43 | 5,93E-04 | 8,21E-03 |
| ENSG00000125637 | PSD4 | 150,44 | -1,42 | 0,43 | -3,26 | 1,13E-03 | 1,30E-02 |
| ENSG00000080493 | SLC4A4 | 776,63 | -1,41 | 0,40 | -3,58 | 3,43E-04 | 5,73E-03 |
| ENSG00000084453 | SLCO1A2 | 1475,03 | -1,41 | 0,40 | -3,54 | 3,95E-04 | 6,31E-03 |
| ENSG00000138795 | LEF1 | 68,88 | -1,41 | 0,29 | -4,84 | 1,32E-06 | 1,73E-04 |
| ENSG00000182492 | BGN | 1049,18 | -1,41 | 0,32 | -4,41 | 1,05E-05 | 6,17E-04 |
| ENSG00000105854 | PON2 | 1197,50 | -1,41 | 0,38 | -3,74 | 1,83E-04 | 3,82E-03 |
| ENSG00000167693 | NXN | 270,57 | -1,41 | 0,43 | -3,25 | 1,16E-03 | 1,33E-02 |
| ENSG00000196411 | EPHB4 | 78,33 | -1,41 | 0,40 | -3,56 | 3,66E-04 | 6,00E-03 |
| ENSG00000136802 | LRRC8A | 2047,68 | -1,41 | 0,35 | -4,00 | 6,33E-05 | 1,90E-03 |
| ENSG00000101605 | MYOM1 | 214,50 | -1,41 | 0,35 | -4,05 | 5,12E-05 | 1,65E-03 |
| ENSG00000167291 | TBC1D16 | 765,92 | -1,41 | 0,33 | -4,22 | 2,41E-05 | 9,99E-04 |
| ENSG00000106565 | TMEM176B | 224,80 | -1,41 | 0,40 | -3,52 | 4,35E-04 | 6,72E-03 |
| ENSG00000167772 | ANGPTL4 | 454,05 | -1,41 | 0,50 | -2,83 | 4,70E-03 | 3,46E-02 |
| ENSG00000183549 | ACSM5 | 57,79 | -1,41 | 0,44 | -3,22 | 1,29E-03 | 1,43E-02 |
| ENSG00000162174 | ASRGL1 | 2057,86 | -1,41 | 0,39 | -3,63 | 2,82E-04 | 4,99E-03 |
| ENSG00000134769 | DTNA | 2508,96 | -1,41 | 0,27 | -5,28 | 1,32E-07 | 3,30E-05 |
| ENSG00000170276 | HSPB2 | 241,41 | -1,40 | 0,43 | -3,23 | 1,23E-03 | 1,38E-02 |
| ENSG00000044459 | CNTLN | 74,23 | -1,40 | 0,44 | -3,22 | 1,30E-03 | 1,44E-02 |
| ENSG00000163191 | S100A11 | 138,50 | -1,40 | 0,41 | -3,40 | 6,80E-04 | 8,98E-03 |
| ENSG00000134186 | PRPF38B | 2684,13 | -1,40 | 0,49 | -2,85 | 4,36E-03 | 3,29E-02 |
| ENSG00000176845 | METRNL | 244,03 | -1,40 | 0,37 | -3,78 | 1,58E-04 | 3,45E-03 |
| ENSG00000104972 | LILRB1 | 58,01 | -1,40 | 0,49 | -2,87 | 4,16E-03 | 3,19E-02 |
| ENSG00000185942 | NKAIN3 | 1188,25 | -1,40 | 0,44 | -3,17 | 1,52E-03 | 1,62E-02 |
| ENSG00000135899 | SP110 | 193,00 | -1,40 | 0,47 | -2,96 | 3,09E-03 | 2,60E-02 |
| ENSG00000198624 | CCDC69 | 209,85 | -1,40 | 0,34 | -4,11 | 4,04E-05 | 1,42E-03 |
| ENSG00000152518 | ZFP36L2 | 420,57 | -1,40 | 0,38 | -3,64 | 2,67E-04 | 4,87E-03 |
| ENSG00000111181 | SLC6A12 | 439,83 | -1,40 | 0,32 | -4,42 | 9,68E-06 | 5,84E-04 |
| ENSG00000078549 | ADCYAP1R1 | 1695,85 | -1,39 | 0,32 | -4,40 | 1,07E-05 | 6,24E-04 |
| ENSG00000163285 | GABRG1 | 504,00 | -1,39 | 0,37 | -3,77 | 1,65E-04 | 3,54E-03 |
| ENSG00000148700 | ADD3 | 1640,20 | -1,39 | 0,28 | -5,04 | 4,66E-07 | 8,68E-05 |
| ENSG00000151322 | NPAS3 | 657,80 | -1,39 | 0,37 | -3,72 | 1,98E-04 | 4,02E-03 |
| ENSG00000074410 | CA12 | 77,74 | -1,39 | 0,50 | -2,80 | 5,06E-03 | 3,62E-02 |
| ENSG00000160307 | S100B | 5487,73 | -1,39 | 0,31 | -4,53 | 5,78E-06 | 4,36E-04 |
| ENSG00000180354 | MTURN | 21899,74 | -1,39 | 0,29 | -4,72 | 2,31E-06 | 2,42E-04 |
| ENSG00000172380 | GNG12 | 193,22 | -1,39 | 0,32 | -4,39 | 1,11E-05 | 6,41E-04 |
| ENSG00000144040 | SFXN5 | 4946,82 | -1,39 | 0,38 | -3,61 | 3,04E-04 | 5,27E-03 |
| ENSG00000157554 | ERG | 50,07 | -1,39 | 0,40 | -3,50 | 4,70E-04 | 7,02E-03 |

| | | | | | | | |
|---|---|---|---|---|---|---|---|
| ENSG00000104881 | PPP1R13L | 110,42 | -1,39 | 0,40 | -3,45 | 5,70E-04 | 7,96E-03 |
| ENSG00000163820 | FYCO1 | 137,58 | -1,39 | 0,31 | -4,53 | 5,93E-06 | 4,42E-04 |
| ENSG00000143178 | TBX19 | 27,07 | -1,39 | 0,50 | -2,78 | 5,42E-03 | 3,81E-02 |
| ENSG00000196091 | MYBPC1 | 544,72 | -1,39 | 0,38 | -3,65 | 2,60E-04 | 4,81E-03 |
| ENSG00000173457 | PPP1R14B | 614,62 | -1,39 | 0,39 | -3,51 | 4,42E-04 | 6,78E-03 |
| ENSG00000174807 | CD248 | 47,74 | -1,38 | 0,31 | -4,44 | 9,05E-06 | 5,64E-04 |
| ENSG00000132688 | NES | 606,19 | -1,38 | 0,31 | -4,51 | 6,53E-06 | 4,71E-04 |
| ENSG00000135272 | MDFIC | 71,89 | -1,38 | 0,33 | -4,23 | 2,32E-05 | 9,84E-04 |
| ENSG00000113721 | PDGFRB | 637,43 | -1,38 | 0,34 | -4,08 | 4,57E-05 | 1,54E-03 |
| ENSG00000168884 | TNIP2 | 199,27 | -1,38 | 0,34 | -4,07 | 4,63E-05 | 1,55E-03 |
| ENSG00000111145 | ELK3 | 55,64 | -1,38 | 0,30 | -4,67 | 3,07E-06 | 3,00E-04 |
| ENSG00000079482 | OPHN1 | 608,22 | -1,38 | 0,46 | -3,02 | 2,54E-03 | 2,27E-02 |
| ENSG00000077585 | GPR137B | 273,59 | -1,38 | 0,32 | -4,29 | 1,77E-05 | 8,63E-04 |
| ENSG00000168394 | TAP1 | 266,14 | -1,38 | 0,37 | -3,75 | 1,75E-04 | 3,71E-03 |
| ENSG00000072274 | TFRC | 452,80 | -1,38 | 0,32 | -4,27 | 1,91E-05 | 8,99E-04 |
| ENSG00000274290 | H2BC6 | 15,64 | -1,38 | 0,49 | -2,79 | 5,29E-03 | 3,75E-02 |
| ENSG00000165895 | ARHGAP42 | 38,47 | -1,38 | 0,41 | -3,40 | 6,84E-04 | 9,01E-03 |
| ENSG00000012211 | PRICKLE3 | 82,15 | -1,38 | 0,39 | -3,49 | 4,87E-04 | 7,21E-03 |
| ENSG00000172183 | ISG20 | 66,55 | -1,38 | 0,34 | -4,03 | 5,56E-05 | 1,76E-03 |
| ENSG00000183486 | MX2 | 56,44 | -1,37 | 0,44 | -3,12 | 1,80E-03 | 1,81E-02 |
| ENSG00000131771 | PPP1R1B | 4886,83 | -1,37 | 0,40 | -3,44 | 5,71E-04 | 7,97E-03 |
| ENSG00000109911 | ELP4 | 565,44 | -1,37 | 0,37 | -3,69 | 2,20E-04 | 4,29E-03 |
| ENSG00000196935 | SRGAP1 | 297,86 | -1,37 | 0,39 | -3,50 | 4,62E-04 | 6,97E-03 |
| ENSG00000235387 | SPAAR | 24,70 | -1,37 | 0,49 | -2,80 | 5,10E-03 | 3,65E-02 |
| ENSG00000018625 | ATP1A2 | 6666,60 | -1,37 | 0,40 | -3,41 | 6,43E-04 | 8,66E-03 |
| ENSG00000181449 | SOX2 | 2834,36 | -1,37 | 0,47 | -2,94 | 3,32E-03 | 2,73E-02 |
| ENSG00000121742 | GJB6 | 573,10 | -1,37 | 0,47 | -2,93 | 3,41E-03 | 2,79E-02 |
| ENSG00000154930 | ACSS1 | 1572,94 | -1,37 | 0,39 | -3,50 | 4,68E-04 | 7,02E-03 |
| ENSG00000139146 | SINHCAF | 10,69 | -1,37 | 0,45 | -3,04 | 2,38E-03 | 2,18E-02 |
| ENSG00000152977 | ZIC1 | 251,57 | -1,37 | 0,35 | -3,89 | 1,02E-04 | 2,57E-03 |
| ENSG00000140522 | RLBP1 | 115,01 | -1,36 | 0,47 | -2,89 | 3,90E-03 | 3,05E-02 |
| ENSG00000183888 | SRARP | 95,73 | -1,36 | 0,43 | -3,18 | 1,47E-03 | 1,57E-02 |
| ENSG00000129219 | PLD2 | 397,79 | -1,36 | 0,36 | -3,78 | 1,59E-04 | 3,46E-03 |
| ENSG00000026025 | VIM | 2116,19 | -1,36 | 0,32 | -4,24 | 2,23E-05 | 9,62E-04 |
| ENSG00000100003 | SEC14L2 | 1257,86 | -1,36 | 0,37 | -3,64 | 2,73E-04 | 4,91E-03 |
| ENSG00000188783 | PRELP | 432,78 | -1,36 | 0,32 | -4,21 | 2,54E-05 | 1,04E-03 |
| ENSG00000170439 | METTL7B | 133,69 | -1,36 | 0,41 | -3,35 | 8,05E-04 | 1,02E-02 |
| ENSG00000173546 | CSPG4 | 468,55 | -1,36 | 0,35 | -3,90 | 9,81E-05 | 2,51E-03 |
| ENSG00000135838 | NPL | 131,60 | -1,36 | 0,36 | -3,79 | 1,49E-04 | 3,32E-03 |
| ENSG00000161798 | AQP5 | 30,62 | -1,36 | 0,44 | -3,11 | 1,87E-03 | 1,86E-02 |
| ENSG00000011422 | PLAUR | 64,67 | -1,36 | 0,45 | -3,04 | 2,40E-03 | 2,19E-02 |
| ENSG00000169604 | ANTXR1 | 237,50 | -1,36 | 0,27 | -4,96 | 6,90E-07 | 1,13E-04 |
| ENSG00000150281 | CTF1 | 54,39 | -1,36 | 0,45 | -2,99 | 2,79E-03 | 2,42E-02 |
| ENSG00000131055 | COX4I2 | 32,33 | -1,36 | 0,44 | -3,10 | 1,90E-03 | 1,88E-02 |
| ENSG00000003402 | CFLAR | 542,43 | -1,36 | 0,36 | -3,82 | 1,35E-04 | 3,12E-03 |
| ENSG00000129680 | MAP7D3 | 119,03 | -1,35 | 0,45 | -3,01 | 2,61E-03 | 2,31E-02 |
| ENSG00000122257 | RBBP6 | 2913,16 | -1,35 | 0,47 | -2,90 | 3,70E-03 | 2,96E-02 |
| ENSG00000134802 | SLC43A3 | 59,35 | -1,35 | 0,35 | -3,90 | 9,68E-05 | 2,49E-03 |
| ENSG00000168268 | NT5DC2 | 552,62 | -1,35 | 0,36 | -3,71 | 2,07E-04 | 4,12E-03 |
| ENSG00000173110 | HSPA6 | 22,45 | -1,35 | 0,47 | -2,90 | 3,78E-03 | 3,00E-02 |
| ENSG00000101144 | BMP7 | 537,63 | -1,35 | 0,33 | -4,06 | 4,82E-05 | 1,58E-03 |
| ENSG00000039560 | RAI14 | 119,18 | -1,35 | 0,31 | -4,33 | 1,52E-05 | 7,72E-04 |
| ENSG00000164292 | RHOBTB3 | 897,87 | -1,35 | 0,30 | -4,42 | 9,72E-06 | 5,84E-04 |
| ENSG00000216490 | IFI30 | 61,01 | -1,35 | 0,44 | -3,05 | 2,32E-03 | 2,15E-02 |
| ENSG00000183963 | SMTN | 398,08 | -1,34 | 0,41 | -3,26 | 1,13E-03 | 1,30E-02 |
| ENSG00000138646 | HERC5 | 43,07 | -1,34 | 0,31 | -4,30 | 1,72E-05 | 8,46E-04 |
| ENSG00000148655 | LRMDA | 50,59 | -1,34 | 0,45 | -2,98 | 2,87E-03 | 2,47E-02 |

| | | | | | | | |
|---|---|---|---|---|---|---|---|
| ENSG00000255690 | TRIL | 776,38 | -1,34 | 0,41 | -3,30 | 9,58E-04 | 1,16E-02 |
| ENSG00000138434 | ITPRID2 | 802,18 | -1,34 | 0,28 | -4,83 | 1,34E-06 | 1,74E-04 |
| ENSG00000102007 | PLP2 | 77,72 | -1,34 | 0,38 | -3,55 | 3,82E-04 | 6,19E-03 |
| ENSG00000166292 | TMEM100 | 27,88 | -1,34 | 0,44 | -3,03 | 2,43E-03 | 2,22E-02 |
| ENSG00000122863 | CHST3 | 225,08 | -1,34 | 0,35 | -3,79 | 1,49E-04 | 3,31E-03 |
| ENSG00000188549 | CCDC9B | 191,70 | -1,34 | 0,37 | -3,67 | 2,46E-04 | 4,62E-03 |
| ENSG00000132205 | EMILIN2 | 37,06 | -1,34 | 0,33 | -4,05 | 5,07E-05 | 1,64E-03 |
| ENSG00000239282 | CASTOR1 | 255,92 | -1,34 | 0,45 | -2,99 | 2,83E-03 | 2,44E-02 |
| ENSG00000132881 | CPLANE2 | 76,39 | -1,34 | 0,44 | -3,04 | 2,34E-03 | 2,16E-02 |
| ENSG00000003436 | TFPI | 25,80 | -1,34 | 0,48 | -2,81 | 4,94E-03 | 3,58E-02 |
| ENSG00000147872 | PLIN2 | 110,26 | -1,34 | 0,38 | -3,53 | 4,19E-04 | 6,55E-03 |
| ENSG00000150457 | LATS2 | 44,05 | -1,34 | 0,33 | -4,00 | 6,30E-05 | 1,90E-03 |
| ENSG00000173281 | PPP1R3B | 29,18 | -1,33 | 0,45 | -2,95 | 3,19E-03 | 2,66E-02 |
| ENSG00000120129 | DUSP1 | 643,59 | -1,33 | 0,39 | -3,43 | 5,98E-04 | 8,27E-03 |
| ENSG00000223802 | CERS1 | 6171,60 | -1,33 | 0,36 | -3,75 | 1,79E-04 | 3,76E-03 |
| ENSG00000221932 | HEPN1 | 3591,15 | -1,33 | 0,38 | -3,51 | 4,45E-04 | 6,80E-03 |
| ENSG00000163083 | INHBB | 74,31 | -1,33 | 0,33 | -4,05 | 5,20E-05 | 1,67E-03 |
| ENSG00000137331 | IER3 | 196,79 | -1,33 | 0,37 | -3,58 | 3,39E-04 | 5,70E-03 |
| ENSG00000119711 | ALDH6A1 | 1806,78 | -1,33 | 0,35 | -3,78 | 1,55E-04 | 3,41E-03 |
| ENSG00000137601 | NEK1 | 521,74 | -1,33 | 0,44 | -3,00 | 2,66E-03 | 2,34E-02 |
| ENSG00000125676 | THOC2 | 2341,91 | -1,33 | 0,49 | -2,70 | 7,03E-03 | 4,59E-02 |
| ENSG00000137834 | SMAD6 | 71,93 | -1,33 | 0,39 | -3,41 | 6,48E-04 | 8,71E-03 |
| ENSG00000137628 | DDX60 | 30,44 | -1,33 | 0,41 | -3,20 | 1,39E-03 | 1,52E-02 |
| ENSG00000072840 | EVC | 28,79 | -1,33 | 0,46 | -2,90 | 3,67E-03 | 2,95E-02 |
| ENSG00000107249 | GLIS3 | 88,44 | -1,32 | 0,41 | -3,27 | 1,08E-03 | 1,26E-02 |
| ENSG00000254979 | AP000781.2 | 26,71 | -1,32 | 0,45 | -2,92 | 3,48E-03 | 2,83E-02 |
| ENSG00000165949 | IFI27 | 588,83 | -1,32 | 0,33 | -4,02 | 5,82E-05 | 1,79E-03 |
| ENSG00000163884 | KLF15 | 298,60 | -1,32 | 0,42 | -3,11 | 1,84E-03 | 1,84E-02 |
| ENSG00000088888 | MAVS | 1212,24 | -1,32 | 0,36 | -3,62 | 2,89E-04 | 5,08E-03 |
| ENSG00000127528 | KLF2 | 283,71 | -1,32 | 0,40 | -3,34 | 8,44E-04 | 1,06E-02 |
| ENSG00000125398 | SOX9 | 730,46 | -1,32 | 0,40 | -3,31 | 9,45E-04 | 1,15E-02 |
| ENSG00000021355 | SERPINB1 | 168,68 | -1,32 | 0,36 | -3,64 | 2,74E-04 | 4,91E-03 |
| ENSG00000140368 | PSTPIP1 | 86,88 | -1,32 | 0,40 | -3,29 | 1,00E-03 | 1,20E-02 |
| ENSG00000131503 | ANKHD1 | 2667,76 | -1,32 | 0,46 | -2,86 | 4,29E-03 | 3,25E-02 |
| ENSG00000033327 | GAB2 | 1303,81 | -1,32 | 0,34 | -3,83 | 1,29E-04 | 3,02E-03 |
| ENSG00000257594 | GALNT4 | 55,97 | -1,32 | 0,43 | -3,07 | 2,15E-03 | 2,05E-02 |
| ENSG00000103196 | CRISPLD2 | 119,34 | -1,31 | 0,43 | -3,09 | 2,01E-03 | 1,96E-02 |
| ENSG00000197245 | FAM110D | 55,03 | -1,31 | 0,42 | -3,13 | 1,74E-03 | 1,77E-02 |
| ENSG00000166535 | A2ML1 | 71,34 | -1,31 | 0,44 | -2,96 | 3,09E-03 | 2,60E-02 |
| ENSG00000126777 | KTN1 | 3629,89 | -1,31 | 0,35 | -3,71 | 2,06E-04 | 4,10E-03 |
| ENSG00000138080 | EMILIN1 | 195,26 | -1,31 | 0,38 | -3,43 | 6,10E-04 | 8,34E-03 |
| ENSG00000118707 | TGIF2 | 60,10 | -1,31 | 0,43 | -3,02 | 2,51E-03 | 2,26E-02 |
| ENSG00000179761 | PIPOX | 154,37 | -1,31 | 0,36 | -3,60 | 3,16E-04 | 5,40E-03 |
| ENSG00000148737 | TCF7L2 | 302,31 | -1,31 | 0,33 | -3,96 | 7,53E-05 | 2,09E-03 |
| ENSG00000224383 | PRR29 | 267,63 | -1,31 | 0,32 | -4,11 | 3,97E-05 | 1,40E-03 |
| ENSG00000174021 | GNG5 | 181,71 | -1,31 | 0,29 | -4,56 | 5,07E-06 | 4,02E-04 |
| ENSG00000129667 | RHBDF2 | 132,56 | -1,31 | 0,45 | -2,89 | 3,89E-03 | 3,04E-02 |
| ENSG00000185483 | ROR1 | 54,63 | -1,31 | 0,48 | -2,74 | 6,14E-03 | 4,14E-02 |
| ENSG00000006756 | ARSD | 265,42 | -1,31 | 0,38 | -3,45 | 5,69E-04 | 7,94E-03 |
| ENSG00000184384 | MAML2 | 60,08 | -1,31 | 0,35 | -3,77 | 1,63E-04 | 3,52E-03 |
| ENSG00000159792 | PSKH1 | 300,00 | -1,31 | 0,30 | -4,40 | 1,09E-05 | 6,34E-04 |
| ENSG00000266964 | FXYD1 | 4683,73 | -1,31 | 0,45 | -2,89 | 3,81E-03 | 3,01E-02 |
| ENSG00000103710 | RASL12 | 151,64 | -1,30 | 0,38 | -3,42 | 6,32E-04 | 8,53E-03 |
| ENSG00000090776 | EFNB1 | 214,23 | -1,30 | 0,38 | -3,46 | 5,35E-04 | 7,61E-03 |
| ENSG00000158966 | CACHD1 | 205,11 | -1,30 | 0,31 | -4,21 | 2,61E-05 | 1,05E-03 |
| ENSG00000144283 | PKP4 | 3785,98 | -1,30 | 0,31 | -4,23 | 2,31E-05 | 9,84E-04 |
| ENSG00000101888 | NXT2 | 52,92 | -1,30 | 0,36 | -3,58 | 3,40E-04 | 5,70E-03 |

| | | | | | | | |
|---|---|---|---|---|---|---|---|
| ENSG00000110900 | TSPAN11 | 149,13 | -1,30 | 0,43 | -3,04 | 2,37E-03 | 2,17E-02 |
| ENSG00000143570 | SLC39A1 | 407,94 | -1,30 | 0,30 | -4,26 | 2,08E-05 | 9,33E-04 |
| ENSG00000104419 | NDRG1 | 4602,50 | -1,30 | 0,33 | -3,93 | 8,41E-05 | 2,23E-03 |
| ENSG00000136869 | TLR4 | 163,59 | -1,30 | 0,36 | -3,58 | 3,46E-04 | 5,76E-03 |
| ENSG00000146112 | PPP1R18 | 168,88 | -1,30 | 0,32 | -4,03 | 5,68E-05 | 1,77E-03 |
| ENSG00000271810 | AL603832.3 | 506,35 | -1,30 | 0,28 | -4,62 | 3,86E-06 | 3,50E-04 |
| ENSG00000213366 | GSTM2 | 2442,58 | -1,30 | 0,44 | -2,92 | 3,50E-03 | 2,83E-02 |
| ENSG00000103335 | PIEZO1 | 546,96 | -1,30 | 0,37 | -3,48 | 5,01E-04 | 7,38E-03 |
| ENSG00000122862 | SRGN | 120,74 | -1,30 | 0,32 | -4,07 | 4,62E-05 | 1,55E-03 |
| ENSG00000168610 | STAT3 | 1481,04 | -1,30 | 0,33 | -3,91 | 9,25E-05 | 2,40E-03 |
| ENSG00000150893 | FREM2 | 25,71 | -1,29 | 0,41 | -3,13 | 1,76E-03 | 1,79E-02 |
| ENSG00000087903 | RFX2 | 194,95 | -1,29 | 0,43 | -3,04 | 2,37E-03 | 2,17E-02 |
| ENSG00000254996 | ANKHD1-EIF4EBI | 2796,23 | -1,29 | 0,45 | -2,88 | 3,92E-03 | 3,06E-02 |
| ENSG00000248235 | AC037459.1 | 191,88 | -1,29 | 0,39 | -3,32 | 9,11E-04 | 1,12E-02 |
| ENSG00000130164 | LDLR | 342,25 | -1,29 | 0,35 | -3,73 | 1,89E-04 | 3,92E-03 |
| ENSG00000137269 | LRRC1 | 189,59 | -1,29 | 0,43 | -2,97 | 2,96E-03 | 2,52E-02 |
| ENSG00000166689 | PLEKHA7 | 74,80 | -1,29 | 0,37 | -3,48 | 5,03E-04 | 7,39E-03 |
| ENSG00000060339 | CCAR1 | 2103,22 | -1,29 | 0,44 | -2,96 | 3,10E-03 | 2,60E-02 |
| ENSG00000143416 | SELENBP1 | 620,03 | -1,29 | 0,40 | -3,21 | 1,35E-03 | 1,49E-02 |
| ENSG00000132481 | TRIM47 | 548,98 | -1,29 | 0,35 | -3,72 | 2,01E-04 | 4,05E-03 |
| ENSG00000188613 | NANOS1 | 59,46 | -1,29 | 0,37 | -3,46 | 5,37E-04 | 7,63E-03 |
| ENSG00000101439 | CST3 | 32620,37 | -1,29 | 0,44 | -2,96 | 3,10E-03 | 2,60E-02 |
| ENSG00000135540 | NHSL1 | 148,01 | -1,29 | 0,34 | -3,77 | 1,62E-04 | 3,50E-03 |
| ENSG00000125817 | CENPB | 3352,94 | -1,29 | 0,33 | -3,95 | 7,79E-05 | 2,13E-03 |
| ENSG00000143171 | RXRG | 174,32 | -1,29 | 0,42 | -3,04 | 2,33E-03 | 2,15E-02 |
| ENSG00000187164 | SHTN1 | 3000,73 | -1,29 | 0,34 | -3,77 | 1,63E-04 | 3,52E-03 |
| ENSG00000166508 | MCM7 | 1051,37 | -1,29 | 0,33 | -3,87 | 1,10E-04 | 2,70E-03 |
| ENSG00000069702 | TGFBR3 | 154,17 | -1,29 | 0,33 | -3,93 | 8,64E-05 | 2,29E-03 |
| ENSG00000164736 | SOX17 | 31,43 | -1,29 | 0,40 | -3,20 | 1,39E-03 | 1,51E-02 |
| ENSG00000024422 | EHD2 | 569,55 | -1,28 | 0,36 | -3,55 | 3,86E-04 | 6,23E-03 |
| ENSG00000110324 | IL10RA | 50,95 | -1,28 | 0,43 | -2,96 | 3,06E-03 | 2,58E-02 |
| ENSG00000120913 | PDLIM2 | 432,98 | -1,28 | 0,39 | -3,30 | 9,64E-04 | 1,16E-02 |
| ENSG00000172340 | SUCLG2 | 135,64 | -1,28 | 0,30 | -4,27 | 1,98E-05 | 9,16E-04 |
| ENSG00000144730 | IL17RD | 124,45 | -1,28 | 0,28 | -4,55 | 5,25E-06 | 4,05E-04 |
| ENSG00000132793 | LPIN3 | 224,98 | -1,28 | 0,41 | -3,14 | 1,71E-03 | 1,75E-02 |
| ENSG00000110719 | TCIRG1 | 236,68 | -1,28 | 0,42 | -3,04 | 2,34E-03 | 2,16E-02 |
| ENSG00000117298 | ECE1 | 1113,49 | -1,28 | 0,31 | -4,06 | 4,88E-05 | 1,60E-03 |
| ENSG00000165478 | HEPACAM | 3956,45 | -1,28 | 0,36 | -3,59 | 3,25E-04 | 5,52E-03 |
| ENSG00000187244 | BCAM | 653,62 | -1,28 | 0,35 | -3,67 | 2,40E-04 | 4,56E-03 |
| ENSG00000164938 | TP53INP1 | 99,47 | -1,27 | 0,35 | -3,64 | 2,73E-04 | 4,91E-03 |
| ENSG00000170345 | FOS | 306,27 | -1,27 | 0,37 | -3,47 | 5,18E-04 | 7,45E-03 |
| ENSG00000131435 | PDLIM4 | 190,71 | -1,27 | 0,39 | -3,24 | 1,20E-03 | 1,36E-02 |
| ENSG00000115380 | EFEMP1 | 350,65 | -1,27 | 0,42 | -3,05 | 2,25E-03 | 2,11E-02 |
| ENSG00000153048 | CARHSP1 | 936,00 | -1,27 | 0,35 | -3,64 | 2,70E-04 | 4,89E-03 |
| ENSG00000149564 | ESAM | 286,43 | -1,27 | 0,31 | -4,16 | 3,19E-05 | 1,19E-03 |
| ENSG00000213398 | LCAT | 320,49 | -1,27 | 0,44 | -2,89 | 3,81E-03 | 3,01E-02 |
| ENSG00000060237 | WNK1 | 2975,06 | -1,27 | 0,27 | -4,72 | 2,33E-06 | 2,43E-04 |
| ENSG00000177685 | CRACR2B | 98,34 | -1,27 | 0,44 | -2,89 | 3,84E-03 | 3,02E-02 |
| ENSG00000259075 | POC1B-GALNT4 | 61,90 | -1,27 | 0,40 | -3,21 | 1,31E-03 | 1,45E-02 |
| ENSG00000083857 | FAT1 | 270,29 | -1,27 | 0,37 | -3,43 | 6,07E-04 | 8,32E-03 |
| ENSG00000162599 | NFIA | 3065,00 | -1,27 | 0,44 | -2,90 | 3,77E-03 | 3,00E-02 |
| ENSG00000120729 | MYOT | 58,28 | -1,27 | 0,44 | -2,89 | 3,85E-03 | 3,02E-02 |
| ENSG00000197360 | ZNF98 | 32,81 | -1,27 | 0,46 | -2,73 | 6,33E-03 | 4,24E-02 |
| ENSG00000103876 | FAH | 205,98 | -1,27 | 0,39 | -3,26 | 1,11E-03 | 1,28E-02 |
| ENSG00000167107 | ACSF2 | 242,12 | -1,26 | 0,38 | -3,34 | 8,32E-04 | 1,05E-02 |
| ENSG00000214063 | TSPAN4 | 222,37 | -1,26 | 0,35 | -3,58 | 3,46E-04 | 5,76E-03 |
| ENSG00000171791 | BCL2 | 267,45 | -1,26 | 0,32 | -3,94 | 8,31E-05 | 2,22E-03 |

| | | | | | | | |
|---|---|---|---|---|---|---|---|
| ENSG00000144476 | ACKR3 | 63,58 | -1,26 | 0,41 | -3,11 | 1,86E-03 | 1,86E-02 |
| ENSG00000172037 | LAMB2 | 817,21 | -1,26 | 0,37 | -3,41 | 6,60E-04 | 8,81E-03 |
| ENSG00000231852 | CYP21A2 | 493,95 | -1,26 | 0,47 | -2,67 | 7,49E-03 | 4,79E-02 |
| ENSG00000066056 | TIE1 | 281,94 | -1,26 | 0,35 | -3,61 | 3,03E-04 | 5,26E-03 |
| ENSG00000187079 | TEAD1 | 307,98 | -1,26 | 0,32 | -3,88 | 1,05E-04 | 2,63E-03 |
| ENSG00000204301 | NOTCH4 | 401,88 | -1,26 | 0,33 | -3,82 | 1,33E-04 | 3,10E-03 |
| ENSG00000103742 | IGDCC4 | 63,29 | -1,26 | 0,35 | -3,56 | 3,71E-04 | 6,06E-03 |
| ENSG00000010379 | SLC6A13 | 184,52 | -1,25 | 0,36 | -3,47 | 5,27E-04 | 7,55E-03 |
| ENSG00000087245 | MMP2 | 47,81 | -1,25 | 0,44 | -2,82 | 4,77E-03 | 3,50E-02 |
| ENSG00000116962 | NID1 | 49,86 | -1,25 | 0,38 | -3,28 | 1,05E-03 | 1,23E-02 |
| ENSG00000168710 | AHCYL1 | 5309,05 | -1,25 | 0,35 | -3,61 | 3,12E-04 | 5,35E-03 |
| ENSG00000126653 | NSRP1 | 2095,15 | -1,25 | 0,40 | -3,13 | 1,73E-03 | 1,77E-02 |
| ENSG00000115109 | EPB41L5 | 189,23 | -1,25 | 0,34 | -3,65 | 2,66E-04 | 4,86E-03 |
| ENSG00000213977 | TAX1BP3 | 621,67 | -1,25 | 0,33 | -3,85 | 1,18E-04 | 2,86E-03 |
| ENSG00000171223 | JUNB | 1130,99 | -1,25 | 0,35 | -3,58 | 3,49E-04 | 5,78E-03 |
| ENSG00000167034 | NKX3-1 | 17,83 | -1,25 | 0,28 | -4,49 | 7,06E-06 | 4,94E-04 |
| ENSG00000183255 | PTTG1IP | 1285,48 | -1,25 | 0,29 | -4,35 | 1,37E-05 | 7,28E-04 |
| ENSG00000130413 | STK33 | 49,79 | -1,25 | 0,41 | -3,07 | 2,14E-03 | 2,04E-02 |
| ENSG00000197256 | KANK2 | 284,69 | -1,25 | 0,34 | -3,72 | 1,99E-04 | 4,02E-03 |
| ENSG00000111961 | SASH1 | 1002,84 | -1,25 | 0,29 | -4,36 | 1,29E-05 | 7,07E-04 |
| ENSG00000104313 | EYA1 | 54,05 | -1,25 | 0,44 | -2,86 | 4,25E-03 | 3,24E-02 |
| ENSG00000137404 | NRM | 52,97 | -1,25 | 0,41 | -3,07 | 2,17E-03 | 2,06E-02 |
| ENSG00000183779 | ZNF703 | 683,54 | -1,24 | 0,37 | -3,35 | 8,22E-04 | 1,04E-02 |
| ENSG00000159176 | CSRP1 | 10443,15 | -1,24 | 0,34 | -3,61 | 3,06E-04 | 5,27E-03 |
| ENSG00000111331 | OAS3 | 80,30 | -1,24 | 0,32 | -3,89 | 1,00E-04 | 2,54E-03 |
| ENSG00000115461 | IGFBP5 | 1043,53 | -1,24 | 0,41 | -3,06 | 2,24E-03 | 2,10E-02 |
| ENSG00000143845 | ETNK2 | 191,63 | -1,24 | 0,31 | -4,00 | 6,36E-05 | 1,91E-03 |
| ENSG00000148143 | ZNF462 | 451,92 | -1,24 | 0,43 | -2,87 | 4,12E-03 | 3,17E-02 |
| ENSG00000079308 | TNS1 | 827,12 | -1,24 | 0,29 | -4,28 | 1,86E-05 | 8,86E-04 |
| ENSG00000146122 | DAAM2 | 2991,40 | -1,24 | 0,33 | -3,80 | 1,44E-04 | 3,24E-03 |
| ENSG00000076555 | ACACB | 450,35 | -1,24 | 0,36 | -3,41 | 6,61E-04 | 8,81E-03 |
| ENSG00000132274 | TRIM22 | 62,97 | -1,24 | 0,35 | -3,54 | 4,03E-04 | 6,37E-03 |
| ENSG00000141485 | SLC13A5 | 212,38 | -1,24 | 0,40 | -3,06 | 2,24E-03 | 2,10E-02 |
| ENSG00000204264 | PSMB8 | 158,28 | -1,24 | 0,32 | -3,82 | 1,32E-04 | 3,08E-03 |
| ENSG00000107829 | FBXW4 | 1944,98 | -1,24 | 0,28 | -4,34 | 1,43E-05 | 7,41E-04 |
| ENSG00000151702 | FLI1 | 76,16 | -1,24 | 0,35 | -3,52 | 4,31E-04 | 6,68E-03 |
| ENSG00000147113 | DIPK2B | 158,59 | -1,24 | 0,30 | -4,17 | 3,01E-05 | 1,17E-03 |
| ENSG00000179399 | GPC5 | 84,73 | -1,24 | 0,40 | -3,09 | 2,00E-03 | 1,95E-02 |
| ENSG00000047457 | CP | 138,64 | -1,24 | 0,37 | -3,36 | 7,72E-04 | 9,91E-03 |
| ENSG00000108622 | ICAM2 | 245,91 | -1,23 | 0,30 | -4,07 | 4,66E-05 | 1,56E-03 |
| ENSG00000112977 | DAP | 406,22 | -1,23 | 0,26 | -4,73 | 2,26E-06 | 2,39E-04 |
| ENSG00000152284 | TCF7L1 | 139,26 | -1,23 | 0,43 | -2,90 | 3,77E-03 | 3,00E-02 |
| ENSG00000148730 | EIF4EBP2 | 903,41 | -1,23 | 0,30 | -4,06 | 4,91E-05 | 1,60E-03 |
| ENSG00000131724 | IL13RA1 | 134,56 | -1,23 | 0,31 | -3,99 | 6,59E-05 | 1,95E-03 |
| ENSG00000108349 | CASC3 | 1601,35 | -1,23 | 0,22 | -5,66 | 1,49E-08 | 1,12E-05 |
| ENSG00000123080 | CDKN2C | 119,14 | -1,23 | 0,30 | -4,17 | 3,05E-05 | 1,17E-03 |
| ENSG00000176076 | KCNE5 | 27,57 | -1,23 | 0,41 | -2,97 | 2,95E-03 | 2,52E-02 |
| ENSG00000173805 | HAP1 | 258,06 | -1,23 | 0,44 | -2,79 | 5,28E-03 | 3,74E-02 |
| ENSG00000213445 | SIPA1 | 388,10 | -1,23 | 0,33 | -3,73 | 1,92E-04 | 3,96E-03 |
| ENSG00000101265 | RASSF2 | 1564,13 | -1,23 | 0,29 | -4,30 | 1,70E-05 | 8,40E-04 |
| ENSG00000123243 | ITIH5 | 349,42 | -1,23 | 0,33 | -3,68 | 2,31E-04 | 4,46E-03 |
| ENSG00000117519 | CNN3 | 821,34 | -1,22 | 0,30 | -4,05 | 5,08E-05 | 1,64E-03 |
| ENSG00000112851 | ERBIN | 1027,95 | -1,22 | 0,34 | -3,65 | 2,62E-04 | 4,83E-03 |
| ENSG00000153902 | LGI4 | 3393,13 | -1,22 | 0,42 | -2,92 | 3,53E-03 | 2,86E-02 |
| ENSG00000117280 | RAB29 | 391,42 | -1,22 | 0,43 | -2,86 | 4,22E-03 | 3,22E-02 |
| ENSG00000168209 | DDIT4 | 811,39 | -1,22 | 0,45 | -2,72 | 6,62E-03 | 4,39E-02 |
| ENSG00000138336 | TET1 | 22,79 | -1,22 | 0,41 | -2,96 | 3,03E-03 | 2,56E-02 |

| | | | | | | | |
|---|---|---|---|---|---|---|---|
| ENSG00000240065 | PSMB9 | 117,64 | -1,22 | 0,36 | -3,38 | 7,14E-04 | 9,33E-03 |
| ENSG00000117525 | F3 | 413,83 | -1,22 | 0,36 | -3,39 | 7,08E-04 | 9,28E-03 |
| ENSG00000137819 | PAQR5 | 41,63 | -1,22 | 0,44 | -2,80 | 5,13E-03 | 3,66E-02 |
| ENSG00000172936 | MYD88 | 41,14 | -1,22 | 0,45 | -2,74 | 6,19E-03 | 4,17E-02 |
| ENSG00000107562 | CXCL12 | 110,22 | -1,22 | 0,31 | -3,93 | 8,35E-05 | 2,23E-03 |
| ENSG00000069869 | NEDD4 | 19,74 | -1,22 | 0,40 | -3,04 | 2,37E-03 | 2,17E-02 |
| ENSG00000162551 | ALPL | 394,24 | -1,22 | 0,33 | -3,69 | 2,24E-04 | 4,35E-03 |
| ENSG00000137561 | TTPA | 28,84 | -1,22 | 0,44 | -2,75 | 5,98E-03 | 4,08E-02 |
| ENSG00000076067 | RBMS2 | 365,96 | -1,22 | 0,37 | -3,34 | 8,53E-04 | 1,06E-02 |
| ENSG00000168792 | ABHD15 | 66,39 | -1,22 | 0,35 | -3,48 | 5,08E-04 | 7,42E-03 |
| ENSG00000198838 | RYR3 | 417,98 | -1,22 | 0,39 | -3,14 | 1,66E-03 | 1,71E-02 |
| ENSG00000204580 | DDR1 | 2061,59 | -1,22 | 0,35 | -3,46 | 5,44E-04 | 7,70E-03 |
| ENSG00000167315 | ACAA2 | 416,98 | -1,21 | 0,28 | -4,26 | 2,02E-05 | 9,23E-04 |
| ENSG00000088826 | SMOX | 986,53 | -1,21 | 0,33 | -3,70 | 2,13E-04 | 4,19E-03 |
| ENSG00000198000 | NOL8 | 403,66 | -1,21 | 0,36 | -3,40 | 6,79E-04 | 8,98E-03 |
| ENSG00000164188 | RANBP3L | 451,82 | -1,21 | 0,37 | -3,28 | 1,03E-03 | 1,22E-02 |
| ENSG00000186918 | ZNF395 | 895,69 | -1,21 | 0,40 | -3,02 | 2,54E-03 | 2,27E-02 |
| ENSG00000147127 | RAB41 | 61,82 | -1,21 | 0,37 | -3,24 | 1,20E-03 | 1,37E-02 |
| ENSG00000088280 | ASAP3 | 347,92 | -1,21 | 0,29 | -4,12 | 3,71E-05 | 1,33E-03 |
| ENSG00000183864 | TOB2 | 1070,40 | -1,21 | 0,40 | -3,03 | 2,47E-03 | 2,24E-02 |
| ENSG00000198363 | ASPH | 1893,95 | -1,21 | 0,31 | -3,85 | 1,19E-04 | 2,86E-03 |
| ENSG00000106003 | LFNG | 237,33 | -1,21 | 0,32 | -3,80 | 1,45E-04 | 3,25E-03 |
| ENSG00000177469 | CAVIN1 | 929,65 | -1,21 | 0,32 | -3,80 | 1,47E-04 | 3,29E-03 |
| ENSG00000162367 | TAL1 | 51,01 | -1,21 | 0,43 | -2,83 | 4,64E-03 | 3,43E-02 |
| ENSG00000107104 | KANK1 | 532,38 | -1,20 | 0,29 | -4,08 | 4,44E-05 | 1,51E-03 |
| ENSG00000163702 | IL17RC | 470,09 | -1,20 | 0,40 | -3,05 | 2,33E-03 | 2,15E-02 |
| ENSG00000169105 | CHST14 | 72,91 | -1,20 | 0,34 | -3,49 | 4,79E-04 | 7,12E-03 |
| ENSG00000155324 | GRAMD2B | 665,32 | -1,20 | 0,32 | -3,76 | 1,71E-04 | 3,65E-03 |
| ENSG00000091656 | ZFHX4 | 542,33 | -1,20 | 0,43 | -2,82 | 4,87E-03 | 3,54E-02 |
| ENSG00000214290 | COLCA2 | 67,19 | -1,20 | 0,39 | -3,12 | 1,82E-03 | 1,83E-02 |
| ENSG00000071967 | CYBRD1 | 328,93 | -1,20 | 0,34 | -3,51 | 4,45E-04 | 6,80E-03 |
| ENSG00000153029 | MR1 | 61,75 | -1,20 | 0,40 | -3,00 | 2,69E-03 | 2,36E-02 |
| ENSG00000150551 | LYPD1 | 436,16 | -1,20 | 0,30 | -4,07 | 4,75E-05 | 1,57E-03 |
| ENSG00000164096 | C4orf3 | 1331,48 | -1,20 | 0,19 | -6,28 | 3,37E-10 | 2,33E-06 |
| ENSG00000164089 | ETNPPL | 894,27 | -1,20 | 0,38 | -3,15 | 1,63E-03 | 1,70E-02 |
| ENSG00000140836 | ZFHX3 | 497,83 | -1,20 | 0,42 | -2,83 | 4,62E-03 | 3,43E-02 |
| ENSG00000186350 | RXRA | 1336,39 | -1,20 | 0,32 | -3,70 | 2,13E-04 | 4,19E-03 |
| ENSG00000164199 | ADGRV1 | 471,30 | -1,19 | 0,37 | -3,19 | 1,43E-03 | 1,55E-02 |
| ENSG00000172201 | ID4 | 700,23 | -1,19 | 0,34 | -3,47 | 5,22E-04 | 7,50E-03 |
| ENSG00000133106 | EPSTI1 | 19,46 | -1,19 | 0,43 | -2,79 | 5,25E-03 | 3,73E-02 |
| ENSG00000127920 | GNG11 | 143,27 | -1,19 | 0,24 | -4,93 | 8,25E-07 | 1,24E-04 |
| ENSG00000099875 | MKNK2 | 763,71 | -1,19 | 0,36 | -3,35 | 8,16E-04 | 1,03E-02 |
| ENSG00000144749 | LRIG1 | 1011,51 | -1,19 | 0,36 | -3,33 | 8,65E-04 | 1,07E-02 |
| ENSG00000042493 | CAPG | 290,95 | -1,19 | 0,34 | -3,51 | 4,49E-04 | 6,83E-03 |
| ENSG00000142611 | PRDM16 | 208,72 | -1,19 | 0,35 | -3,36 | 7,89E-04 | 1,01E-02 |
| ENSG00000147883 | CDKN2B | 25,75 | -1,19 | 0,36 | -3,34 | 8,39E-04 | 1,05E-02 |
| ENSG00000012048 | BRCA1 | 110,18 | -1,19 | 0,36 | -3,27 | 1,09E-03 | 1,27E-02 |
| ENSG00000111077 | TNS2 | 1845,72 | -1,19 | 0,31 | -3,86 | 1,12E-04 | 2,74E-03 |
| ENSG00000079215 | SLC1A3 | 4676,18 | -1,19 | 0,39 | -3,02 | 2,49E-03 | 2,25E-02 |
| ENSG00000186654 | PRR5 | 711,66 | -1,19 | 0,44 | -2,70 | 6,91E-03 | 4,53E-02 |
| ENSG00000254087 | LYN | 63,83 | -1,19 | 0,34 | -3,53 | 4,11E-04 | 6,46E-03 |
| ENSG00000123104 | ITPR2 | 273,31 | -1,19 | 0,36 | -3,31 | 9,44E-04 | 1,15E-02 |
| ENSG00000197879 | MYO1C | 412,22 | -1,19 | 0,35 | -3,43 | 6,05E-04 | 8,32E-03 |
| ENSG00000171634 | BPTF | 2089,60 | -1,19 | 0,42 | -2,82 | 4,74E-03 | 3,49E-02 |
| ENSG00000264522 | OTUD7B | 725,99 | -1,18 | 0,34 | -3,44 | 5,91E-04 | 8,18E-03 |
| ENSG00000176623 | RMDN1 | 360,41 | -1,18 | 0,23 | -5,08 | 3,78E-07 | 7,83E-05 |
| ENSG00000183840 | GPR39 | 347,21 | -1,18 | 0,29 | -4,12 | 3,86E-05 | 1,37E-03 |

| | | | | | | | |
|---|---|---|---|---|---|---|---|
| ENSG00000100292 | HMOX1 | 85,89 | -1,18 | 0,40 | -2,93 | 3,44E-03 | 2,81E-02 |
| ENSG00000149499 | EML3 | 600,84 | -1,18 | 0,37 | -3,20 | 1,36E-03 | 1,49E-02 |
| ENSG00000123094 | RASSF8 | 167,18 | -1,18 | 0,22 | -5,34 | 9,54E-08 | 2,91E-05 |
| ENSG00000040531 | CTNS | 411,62 | -1,18 | 0,34 | -3,52 | 4,35E-04 | 6,72E-03 |
| ENSG00000169554 | ZEB2 | 2625,38 | -1,18 | 0,31 | -3,77 | 1,65E-04 | 3,54E-03 |
| ENSG00000086159 | AQP6 | 41,33 | -1,18 | 0,38 | -3,09 | 1,99E-03 | 1,95E-02 |
| ENSG00000163430 | FSTL1 | 230,28 | -1,18 | 0,26 | -4,51 | 6,36E-06 | 4,65E-04 |
| ENSG00000185633 | NDUFA4L2 | 428,90 | -1,18 | 0,31 | -3,83 | 1,27E-04 | 2,98E-03 |
| ENSG00000167614 | TTYH1 | 9456,47 | -1,18 | 0,35 | -3,36 | 7,85E-04 | 1,00E-02 |
| ENSG00000203814 | H2BC18 | 10,93 | -1,18 | 0,42 | -2,81 | 4,95E-03 | 3,58E-02 |
| ENSG00000176244 | ACBD7 | 953,16 | -1,18 | 0,34 | -3,50 | 4,70E-04 | 7,02E-03 |
| ENSG00000017483 | SLC38A5 | 333,26 | -1,18 | 0,37 | -3,17 | 1,53E-03 | 1,62E-02 |
| ENSG00000154856 | APCDD1 | 650,90 | -1,17 | 0,34 | -3,49 | 4,81E-04 | 7,12E-03 |
| ENSG00000101871 | MID1 | 129,46 | -1,17 | 0,32 | -3,66 | 2,56E-04 | 4,77E-03 |
| ENSG00000182511 | FES | 207,32 | -1,17 | 0,39 | -2,98 | 2,87E-03 | 2,47E-02 |
| ENSG00000138592 | USP8 | 2187,56 | -1,17 | 0,43 | -2,70 | 7,01E-03 | 4,58E-02 |
| ENSG00000100100 | PIK3IP1 | 569,94 | -1,17 | 0,35 | -3,34 | 8,40E-04 | 1,05E-02 |
| ENSG00000149932 | TMEM219 | 2466,10 | -1,17 | 0,34 | -3,48 | 5,09E-04 | 7,42E-03 |
| ENSG00000143409 | MINDY1 | 234,71 | -1,17 | 0,33 | -3,59 | 3,34E-04 | 5,63E-03 |
| ENSG00000173947 | PIFO | 64,58 | -1,17 | 0,35 | -3,32 | 8,94E-04 | 1,10E-02 |
| ENSG00000145012 | LPP | 446,22 | -1,16 | 0,35 | -3,38 | 7,37E-04 | 9,58E-03 |
| ENSG00000183762 | KREMEN1 | 329,49 | -1,16 | 0,27 | -4,24 | 2,21E-05 | 9,61E-04 |
| ENSG00000122432 | SPATA1 | 50,77 | -1,16 | 0,26 | -4,45 | 8,54E-06 | 5,46E-04 |
| ENSG00000143842 | SOX13 | 230,65 | -1,16 | 0,36 | -3,22 | 1,28E-03 | 1,42E-02 |
| ENSG00000198826 | ARHGAP11A | 29,23 | -1,16 | 0,29 | -4,01 | 6,16E-05 | 1,87E-03 |
| ENSG00000135720 | DYNC1LI2 | 3500,69 | -1,16 | 0,24 | -4,76 | 1,98E-06 | 2,20E-04 |
| ENSG00000182118 | FAM89A | 95,59 | -1,16 | 0,34 | -3,38 | 7,37E-04 | 9,58E-03 |
| ENSG00000126878 | AIF1L | 2333,11 | -1,16 | 0,33 | -3,51 | 4,47E-04 | 6,80E-03 |
| ENSG00000144655 | CSRNP1 | 297,64 | -1,16 | 0,30 | -3,87 | 1,08E-04 | 2,68E-03 |
| ENSG00000187840 | EIF4EBP1 | 113,85 | -1,16 | 0,42 | -2,80 | 5,16E-03 | 3,68E-02 |
| ENSG00000174099 | MSRB3 | 255,12 | -1,16 | 0,33 | -3,52 | 4,28E-04 | 6,64E-03 |
| ENSG00000163840 | DTX3L | 58,09 | -1,16 | 0,38 | -3,04 | 2,37E-03 | 2,17E-02 |
| ENSG00000183773 | AIFM3 | 2316,34 | -1,16 | 0,36 | -3,21 | 1,34E-03 | 1,48E-02 |
| ENSG00000125753 | VASP | 192,49 | -1,16 | 0,33 | -3,48 | 5,08E-04 | 7,42E-03 |
| ENSG00000115355 | CCDC88A | 7009,81 | -1,16 | 0,43 | -2,68 | 7,31E-03 | 4,71E-02 |
| ENSG00000198844 | ARHGEF15 | 85,63 | -1,15 | 0,41 | -2,84 | 4,48E-03 | 3,36E-02 |
| ENSG00000157600 | TMEM164 | 242,76 | -1,15 | 0,29 | -3,99 | 6,63E-05 | 1,95E-03 |
| ENSG00000126785 | RHOJ | 206,35 | -1,15 | 0,35 | -3,26 | 1,11E-03 | 1,29E-02 |
| ENSG00000164379 | FOXQ1 | 61,63 | -1,15 | 0,35 | -3,27 | 1,07E-03 | 1,25E-02 |
| ENSG00000099385 | BCL7C | 817,72 | -1,15 | 0,41 | -2,83 | 4,62E-03 | 3,43E-02 |
| ENSG00000128271 | ADORA2A | 139,63 | -1,15 | 0,38 | -3,01 | 2,64E-03 | 2,33E-02 |
| ENSG00000064961 | HMG20B | 501,15 | -1,15 | 0,36 | -3,22 | 1,30E-03 | 1,45E-02 |
| ENSG00000173926 | MARCHF3 | 46,88 | -1,15 | 0,36 | -3,15 | 1,61E-03 | 1,68E-02 |
| ENSG00000137831 | UACA | 209,74 | -1,15 | 0,39 | -2,92 | 3,48E-03 | 2,83E-02 |
| ENSG00000173269 | MMRN2 | 101,52 | -1,15 | 0,26 | -4,42 | 9,93E-06 | 5,92E-04 |
| ENSG00000204219 | TCEA3 | 93,83 | -1,15 | 0,34 | -3,37 | 7,44E-04 | 9,63E-03 |
| ENSG00000183196 | CHST6 | 254,24 | -1,14 | 0,38 | -3,04 | 2,35E-03 | 2,16E-02 |
| ENSG00000173402 | DAG1 | 787,07 | -1,14 | 0,31 | -3,72 | 1,97E-04 | 4,00E-03 |
| ENSG00000166801 | FAM111A | 70,04 | -1,14 | 0,38 | -2,98 | 2,92E-03 | 2,51E-02 |
| ENSG00000128567 | PODXL | 470,60 | -1,14 | 0,25 | -4,53 | 5,83E-06 | 4,38E-04 |
| ENSG00000163406 | SLC15A2 | 236,19 | -1,14 | 0,38 | -3,02 | 2,49E-03 | 2,25E-02 |
| ENSG00000105329 | TGFB1 | 323,70 | -1,14 | 0,32 | -3,56 | 3,71E-04 | 6,06E-03 |
| ENSG00000004799 | PDK4 | 462,24 | -1,14 | 0,40 | -2,85 | 4,33E-03 | 3,28E-02 |
| ENSG00000090554 | FLT3LG | 94,92 | -1,14 | 0,43 | -2,65 | 7,96E-03 | 4,99E-02 |
| ENSG00000168389 | MFSD2A | 345,28 | -1,14 | 0,35 | -3,21 | 1,34E-03 | 1,47E-02 |
| ENSG00000234745 | HLA-B | 1330,41 | -1,14 | 0,35 | -3,23 | 1,23E-03 | 1,38E-02 |
| ENSG00000175938 | ORAI3 | 186,70 | -1,13 | 0,33 | -3,44 | 5,75E-04 | 8,01E-03 |

| | | | | | | | |
|---|---|---|---|---|---|---|---|
| ENSG00000188931 | CFAP126 | 75,70 | -1,13 | 0,26 | -4,35 | 1,39E-05 | 7,33E-04 |
| ENSG00000089472 | HEPH | 270,24 | -1,13 | 0,40 | -2,82 | 4,85E-03 | 3,53E-02 |
| ENSG00000196196 | HRCT1 | 45,99 | -1,13 | 0,42 | -2,68 | 7,28E-03 | 4,70E-02 |
| ENSG00000068001 | HYAL2 | 223,58 | -1,13 | 0,32 | -3,58 | 3,42E-04 | 5,72E-03 |
| ENSG00000180481 | GLIPR1L2 | 124,64 | -1,13 | 0,40 | -2,86 | 4,26E-03 | 3,24E-02 |
| ENSG00000079337 | RAPGEF3 | 1818,08 | -1,13 | 0,38 | -2,97 | 2,94E-03 | 2,52E-02 |
| ENSG00000181192 | DHTKD1 | 557,82 | -1,13 | 0,38 | -2,99 | 2,83E-03 | 2,44E-02 |
| ENSG00000101162 | TUBB1 | 24,36 | -1,13 | 0,33 | -3,40 | 6,83E-04 | 9,01E-03 |
| ENSG00000248751 | AC004997.1 | 317,59 | -1,13 | 0,38 | -2,99 | 2,77E-03 | 2,40E-02 |
| ENSG00000146648 | EGFR | 180,09 | -1,12 | 0,36 | -3,09 | 2,01E-03 | 1,96E-02 |
| ENSG00000114853 | ZBTB47 | 2283,65 | -1,12 | 0,27 | -4,17 | 3,04E-05 | 1,17E-03 |
| ENSG00000164181 | ELOVL7 | 124,37 | -1,12 | 0,26 | -4,28 | 1,84E-05 | 8,80E-04 |
| ENSG00000127914 | AKAP9 | 2705,32 | -1,12 | 0,39 | -2,84 | 4,55E-03 | 3,39E-02 |
| ENSG00000267534 | S1PR2 | 61,81 | -1,12 | 0,41 | -2,72 | 6,44E-03 | 4,30E-02 |
| ENSG00000204128 | C2orf72 | 1045,08 | -1,12 | 0,29 | -3,84 | 1,24E-04 | 2,95E-03 |
| ENSG00000067177 | PHKA1 | 83,74 | -1,12 | 0,36 | -3,08 | 2,05E-03 | 1,97E-02 |
| ENSG00000140299 | BNIP2 | 180,43 | -1,12 | 0,24 | -4,56 | 5,09E-06 | 4,02E-04 |
| ENSG00000259417 | CTXND1 | 233,50 | -1,12 | 0,26 | -4,24 | 2,28E-05 | 9,79E-04 |
| ENSG00000114166 | KAT2B | 443,63 | -1,12 | 0,27 | -4,21 | 2,57E-05 | 1,05E-03 |
| ENSG00000119471 | HSDL2 | 408,66 | -1,12 | 0,27 | -4,14 | 3,41E-05 | 1,25E-03 |
| ENSG00000182704 | TSKU | 52,79 | -1,12 | 0,31 | -3,65 | 2,67E-04 | 4,87E-03 |
| ENSG00000130517 | PGPEP1 | 1038,72 | -1,12 | 0,41 | -2,74 | 6,07E-03 | 4,11E-02 |
| ENSG00000140545 | MFGE8 | 1250,65 | -1,11 | 0,32 | -3,48 | 5,01E-04 | 7,38E-03 |
| ENSG00000136044 | APPL2 | 353,90 | -1,11 | 0,29 | -3,90 | 9,66E-05 | 2,49E-03 |
| ENSG00000135686 | KLHL36 | 621,25 | -1,11 | 0,35 | -3,16 | 1,60E-03 | 1,67E-02 |
| ENSG00000167470 | MIDN | 865,42 | -1,11 | 0,32 | -3,47 | 5,17E-04 | 7,45E-03 |
| ENSG00000138835 | RGS3 | 400,05 | -1,11 | 0,27 | -4,09 | 4,23E-05 | 1,46E-03 |
| ENSG00000054179 | ENTPD2 | 224,65 | -1,11 | 0,39 | -2,86 | 4,22E-03 | 3,23E-02 |
| ENSG00000137393 | RNF144B | 71,60 | -1,11 | 0,30 | -3,73 | 1,90E-04 | 3,94E-03 |
| ENSG00000122986 | HVCN1 | 156,51 | -1,11 | 0,30 | -3,73 | 1,88E-04 | 3,92E-03 |
| ENSG00000249751 | ECSCR | 23,26 | -1,11 | 0,38 | -2,91 | 3,58E-03 | 2,89E-02 |
| ENSG00000148482 | SLC39A12 | 231,28 | -1,11 | 0,40 | -2,73 | 6,30E-03 | 4,22E-02 |
| ENSG00000130429 | ARPC1B | 336,79 | -1,10 | 0,37 | -2,97 | 3,03E-03 | 2,56E-02 |
| ENSG00000206190 | ATP10A | 247,02 | -1,10 | 0,35 | -3,19 | 1,44E-03 | 1,55E-02 |
| ENSG00000141401 | IMPA2 | 91,29 | -1,10 | 0,41 | -2,67 | 7,65E-03 | 4,84E-02 |
| ENSG00000176971 | FIBIN | 142,10 | -1,10 | 0,34 | -3,27 | 1,09E-03 | 1,27E-02 |
| ENSG00000184985 | SORCS2 | 443,31 | -1,10 | 0,30 | -3,63 | 2,79E-04 | 4,96E-03 |
| ENSG00000123453 | SARDH | 105,57 | -1,10 | 0,40 | -2,78 | 5,39E-03 | 3,81E-02 |
| ENSG00000198157 | HMGN5 | 474,46 | -1,10 | 0,38 | -2,92 | 3,45E-03 | 2,82E-02 |
| ENSG00000269713 | NBPF9 | 254,87 | -1,10 | 0,40 | -2,74 | 6,12E-03 | 4,14E-02 |
| ENSG00000157191 | NECAP2 | 318,97 | -1,10 | 0,28 | -3,89 | 1,00E-04 | 2,54E-03 |
| ENSG00000156049 | GNA14 | 89,77 | -1,10 | 0,33 | -3,30 | 9,57E-04 | 1,16E-02 |
| ENSG00000133131 | MORC4 | 54,44 | -1,10 | 0,31 | -3,50 | 4,57E-04 | 6,91E-03 |
| ENSG00000046889 | PREX2 | 213,83 | -1,10 | 0,26 | -4,27 | 1,97E-05 | 9,16E-04 |
| ENSG00000141526 | SLC16A3 | 123,49 | -1,10 | 0,35 | -3,12 | 1,80E-03 | 1,81E-02 |
| ENSG00000149090 | PAMR1 | 338,42 | -1,10 | 0,33 | -3,29 | 1,00E-03 | 1,20E-02 |
| ENSG00000154217 | PITPNC1 | 602,84 | -1,09 | 0,29 | -3,78 | 1,55E-04 | 3,41E-03 |
| ENSG00000102996 | MMP15 | 376,36 | -1,09 | 0,35 | -3,16 | 1,59E-03 | 1,67E-02 |
| ENSG00000172458 | IL17D | 1283,02 | -1,09 | 0,23 | -4,72 | 2,35E-06 | 2,44E-04 |
| ENSG00000124615 | MOCS1 | 1756,38 | -1,09 | 0,31 | -3,57 | 3,60E-04 | 5,92E-03 |
| ENSG00000183111 | ARHGEF37 | 487,85 | -1,09 | 0,30 | -3,58 | 3,39E-04 | 5,69E-03 |
| ENSG00000050555 | LAMC3 | 143,55 | -1,09 | 0,39 | -2,81 | 4,94E-03 | 3,57E-02 |
| ENSG00000111254 | AKAP3 | 23,49 | -1,09 | 0,29 | -3,72 | 2,01E-04 | 4,05E-03 |
| ENSG00000160325 | CACFD1 | 392,41 | -1,09 | 0,30 | -3,61 | 3,03E-04 | 5,26E-03 |
| ENSG00000158813 | EDA | 28,82 | -1,09 | 0,30 | -3,67 | 2,40E-04 | 4,56E-03 |
| ENSG00000166925 | TSC22D4 | 5570,43 | -1,09 | 0,33 | -3,27 | 1,07E-03 | 1,25E-02 |
| ENSG00000162337 | LRP5 | 137,19 | -1,09 | 0,32 | -3,41 | 6,54E-04 | 8,75E-03 |

| | | | | | | | |
|---|---|---|---|---|---|---|---|
| ENSG00000187068 | C3orf70 | 244,54 | -1,09 | 0,24 | -4,47 | 7,99E-06 | 5,35E-04 |
| ENSG00000154553 | PDLIM3 | 131,03 | -1,09 | 0,26 | -4,23 | 2,37E-05 | 9,93E-04 |
| ENSG00000125520 | SLC2A4RG | 839,91 | -1,09 | 0,35 | -3,11 | 1,87E-03 | 1,86E-02 |
| ENSG00000171388 | APLN | 273,66 | -1,09 | 0,32 | -3,42 | 6,29E-04 | 8,51E-03 |
| ENSG00000173905 | GOLIM4 | 433,59 | -1,09 | 0,28 | -3,82 | 1,35E-04 | 3,12E-03 |
| ENSG00000123609 | NMI | 23,45 | -1,08 | 0,39 | -2,82 | 4,88E-03 | 3,54E-02 |
| ENSG00000164904 | ALDH7A1 | 1444,64 | -1,08 | 0,32 | -3,41 | 6,47E-04 | 8,71E-03 |
| ENSG00000204520 | MICA | 72,25 | -1,08 | 0,39 | -2,75 | 5,93E-03 | 4,06E-02 |
| ENSG00000239779 | WBP1 | 1608,25 | -1,08 | 0,28 | -3,86 | 1,14E-04 | 2,76E-03 |
| ENSG00000286185 | ENSG000002861 | 339,35 | -1,08 | 0,39 | -2,79 | 5,30E-03 | 3,75E-02 |
| ENSG00000132002 | DNAJB1 | 1729,50 | -1,08 | 0,33 | -3,29 | 9,88E-04 | 1,18E-02 |
| ENSG00000159403 | C1R | 243,44 | -1,08 | 0,34 | -3,15 | 1,63E-03 | 1,70E-02 |
| ENSG00000103966 | EHD4 | 123,68 | -1,08 | 0,29 | -3,71 | 2,09E-04 | 4,14E-03 |
| ENSG00000151233 | GXYLT1 | 221,50 | -1,07 | 0,39 | -2,75 | 5,92E-03 | 4,06E-02 |
| ENSG00000135046 | ANXA1 | 96,91 | -1,07 | 0,28 | -3,78 | 1,60E-04 | 3,48E-03 |
| ENSG00000163659 | TIPARP | 141,10 | -1,07 | 0,31 | -3,45 | 5,65E-04 | 7,92E-03 |
| ENSG00000133789 | SWAP70 | 233,07 | -1,07 | 0,27 | -4,03 | 5,59E-05 | 1,76E-03 |
| ENSG00000016602 | CLCA4 | 47,50 | -1,07 | 0,33 | -3,20 | 1,35E-03 | 1,49E-02 |
| ENSG00000189221 | MAOA | 355,72 | -1,07 | 0,29 | -3,67 | 2,42E-04 | 4,57E-03 |
| ENSG00000113645 | WWC1 | 759,93 | -1,07 | 0,25 | -4,36 | 1,30E-05 | 7,11E-04 |
| ENSG00000164574 | GALNT10 | 209,67 | -1,07 | 0,25 | -4,36 | 1,32E-05 | 7,19E-04 |
| ENSG00000173041 | ZNF680 | 181,64 | -1,07 | 0,32 | -3,39 | 6,97E-04 | 9,15E-03 |
| ENSG00000137502 | RAB30 | 642,44 | -1,07 | 0,30 | -3,58 | 3,50E-04 | 5,78E-03 |
| ENSG00000197696 | NMB | 112,87 | -1,07 | 0,39 | -2,71 | 6,80E-03 | 4,49E-02 |
| ENSG00000144642 | RBMS3 | 170,96 | -1,07 | 0,35 | -3,04 | 2,34E-03 | 2,16E-02 |
| ENSG00000072163 | LIMS2 | 772,75 | -1,07 | 0,36 | -2,94 | 3,28E-03 | 2,71E-02 |
| ENSG00000148400 | NOTCH1 | 423,50 | -1,07 | 0,28 | -3,75 | 1,75E-04 | 3,71E-03 |
| ENSG00000110057 | UNC93B1 | 196,43 | -1,07 | 0,38 | -2,77 | 5,61E-03 | 3,90E-02 |
| ENSG00000165886 | UBTD1 | 283,19 | -1,06 | 0,27 | -3,98 | 6,95E-05 | 2,00E-03 |
| ENSG00000140807 | NKD1 | 198,76 | -1,06 | 0,24 | -4,45 | 8,57E-06 | 5,46E-04 |
| ENSG00000165801 | ARHGEF40 | 682,45 | -1,06 | 0,32 | -3,32 | 9,05E-04 | 1,11E-02 |
| ENSG00000236609 | ZNF853 | 671,65 | -1,06 | 0,24 | -4,48 | 7,40E-06 | 5,08E-04 |
| ENSG00000160678 | S100A1 | 4016,16 | -1,06 | 0,34 | -3,08 | 2,08E-03 | 2,00E-02 |
| ENSG00000186815 | TPCN1 | 1176,28 | -1,06 | 0,33 | -3,25 | 1,17E-03 | 1,34E-02 |
| ENSG00000157657 | ZNF618 | 160,40 | -1,06 | 0,39 | -2,75 | 6,01E-03 | 4,09E-02 |
| ENSG00000146966 | DENND2A | 625,25 | -1,06 | 0,24 | -4,35 | 1,36E-05 | 7,28E-04 |
| ENSG00000101384 | JAG1 | 141,70 | -1,06 | 0,33 | -3,19 | 1,43E-03 | 1,54E-02 |
| ENSG00000010704 | HFE | 27,34 | -1,06 | 0,40 | -2,66 | 7,85E-03 | 4,94E-02 |
| ENSG00000198585 | NUDT16 | 1238,87 | -1,06 | 0,21 | -4,93 | 8,21E-07 | 1,24E-04 |
| ENSG00000136425 | CIB2 | 258,87 | -1,06 | 0,31 | -3,39 | 6,88E-04 | 9,05E-03 |
| ENSG00000139350 | NEDD1 | 34,84 | -1,06 | 0,28 | -3,81 | 1,38E-04 | 3,16E-03 |
| ENSG00000173821 | RNF213 | 877,66 | -1,06 | 0,37 | -2,88 | 4,04E-03 | 3,12E-02 |
| ENSG00000130307 | USHBP1 | 103,76 | -1,06 | 0,40 | -2,67 | 7,52E-03 | 4,80E-02 |
| ENSG00000165175 | MID1IP1 | 1602,07 | -1,06 | 0,39 | -2,69 | 7,23E-03 | 4,68E-02 |
| ENSG00000150093 | ITGB1 | 682,38 | -1,05 | 0,24 | -4,46 | 8,28E-06 | 5,42E-04 |
| ENSG00000213722 | DDAH2 | 687,62 | -1,05 | 0,30 | -3,56 | 3,74E-04 | 6,09E-03 |
| ENSG00000101198 | NKAIN4 | 558,19 | -1,05 | 0,38 | -2,77 | 5,54E-03 | 3,87E-02 |
| ENSG00000166341 | DCHS1 | 370,09 | -1,05 | 0,30 | -3,53 | 4,12E-04 | 6,46E-03 |
| ENSG00000010319 | SEMA3G | 134,57 | -1,05 | 0,37 | -2,85 | 4,40E-03 | 3,31E-02 |
| ENSG00000149131 | SERPING1 | 587,12 | -1,05 | 0,29 | -3,64 | 2,73E-04 | 4,91E-03 |
| ENSG00000197496 | SLC2A10 | 23,34 | -1,05 | 0,39 | -2,73 | 6,41E-03 | 4,28E-02 |
| ENSG00000256514 | AP003419.1 | 536,67 | -1,05 | 0,33 | -3,15 | 1,64E-03 | 1,70E-02 |
| ENSG00000129353 | SLC44A2 | 2033,45 | -1,05 | 0,29 | -3,63 | 2,78E-04 | 4,96E-03 |
| ENSG00000177954 | RPS27 | 3787,43 | -1,05 | 0,30 | -3,46 | 5,41E-04 | 7,67E-03 |
| ENSG00000187091 | PLCD1 | 232,79 | -1,04 | 0,35 | -2,97 | 2,93E-03 | 2,51E-02 |
| ENSG00000085063 | CD59 | 2734,11 | -1,04 | 0,28 | -3,68 | 2,34E-04 | 4,48E-03 |
| ENSG00000120885 | CLU | 69171,86 | -1,04 | 0,35 | -3,01 | 2,59E-03 | 2,30E-02 |

| | | | | | | | |
|---|---|---|---|---|---|---|---|
| ENSG00000162433 | AK4 | 586,81 | -1,04 | 0,26 | -3,97 | 7,13E-05 | 2,03E-03 |
| ENSG00000184221 | OLIG1 | 2261,79 | -1,04 | 0,23 | -4,46 | 8,34E-06 | 5,42E-04 |
| ENSG00000158270 | COLEC12 | 148,74 | -1,04 | 0,24 | -4,35 | 1,36E-05 | 7,28E-04 |
| ENSG00000130202 | NECTIN2 | 387,31 | -1,04 | 0,30 | -3,47 | 5,12E-04 | 7,42E-03 |
| ENSG00000157110 | RBPMS | 110,00 | -1,04 | 0,38 | -2,77 | 5,54E-03 | 3,87E-02 |
| ENSG00000150403 | TMCO3 | 324,16 | -1,04 | 0,21 | -4,91 | 9,25E-07 | 1,34E-04 |
| ENSG00000173575 | CHD2 | 3173,24 | -1,04 | 0,39 | -2,69 | 7,10E-03 | 4,62E-02 |
| ENSG00000213694 | S1PR3 | 62,26 | -1,04 | 0,27 | -3,84 | 1,25E-04 | 2,95E-03 |
| ENSG00000100300 | TSPO | 400,49 | -1,04 | 0,36 | -2,85 | 4,34E-03 | 3,28E-02 |
| ENSG00000145390 | USP53 | 246,21 | -1,04 | 0,37 | -2,84 | 4,45E-03 | 3,34E-02 |
| ENSG00000091436 | MAP3K20 | 306,10 | -1,04 | 0,35 | -2,93 | 3,34E-03 | 2,74E-02 |
| ENSG00000110651 | CD81 | 7017,13 | -1,04 | 0,31 | -3,39 | 6,98E-04 | 9,15E-03 |
| ENSG00000141736 | ERBB2 | 686,06 | -1,04 | 0,27 | -3,78 | 1,58E-04 | 3,44E-03 |
| ENSG00000153446 | C16orf89 | 386,31 | -1,04 | 0,39 | -2,67 | 7,58E-03 | 4,81E-02 |
| ENSG00000085563 | ABCB1 | 133,31 | -1,04 | 0,20 | -5,29 | 1,25E-07 | 3,30E-05 |
| ENSG00000105835 | NAMPT | 382,72 | -1,04 | 0,25 | -4,23 | 2,39E-05 | 9,99E-04 |
| ENSG00000105699 | LSR | 201,98 | -1,03 | 0,26 | -4,01 | 6,20E-05 | 1,87E-03 |
| ENSG00000168961 | LGALS9 | 181,55 | -1,03 | 0,37 | -2,82 | 4,81E-03 | 3,51E-02 |
| ENSG00000102935 | ZNF423 | 166,76 | -1,03 | 0,26 | -3,94 | 7,99E-05 | 2,16E-03 |
| ENSG00000164050 | PLXNB1 | 2861,79 | -1,03 | 0,34 | -3,06 | 2,22E-03 | 2,10E-02 |
| ENSG00000067082 | KLF6 | 1145,05 | -1,03 | 0,36 | -2,88 | 4,01E-03 | 3,11E-02 |
| ENSG00000176894 | PXMP2 | 181,70 | -1,03 | 0,32 | -3,22 | 1,27E-03 | 1,42E-02 |
| ENSG00000112378 | PERP | 68,28 | -1,03 | 0,26 | -3,93 | 8,41E-05 | 2,23E-03 |
| ENSG00000147475 | ERLIN2 | 236,74 | -1,03 | 0,29 | -3,51 | 4,44E-04 | 6,79E-03 |
| ENSG00000104728 | ARHGEF10 | 531,37 | -1,03 | 0,32 | -3,24 | 1,18E-03 | 1,35E-02 |
| ENSG00000070526 | ST6GALNAC1 | 33,34 | -1,03 | 0,37 | -2,74 | 6,05E-03 | 4,11E-02 |
| ENSG00000093072 | ADA2 | 154,38 | -1,03 | 0,28 | -3,72 | 1,97E-04 | 4,00E-03 |
| ENSG00000164867 | NOS3 | 155,85 | -1,03 | 0,32 | -3,26 | 1,11E-03 | 1,28E-02 |
| ENSG00000103852 | TTC23 | 100,03 | -1,03 | 0,28 | -3,72 | 1,97E-04 | 4,00E-03 |
| ENSG00000148053 | NTRK2 | 8660,33 | -1,03 | 0,30 | -3,47 | 5,14E-04 | 7,43E-03 |
| ENSG00000182240 | BACE2 | 63,61 | -1,03 | 0,31 | -3,33 | 8,76E-04 | 1,08E-02 |
| ENSG00000038219 | BOD1L1 | 3072,43 | -1,03 | 0,38 | -2,71 | 6,72E-03 | 4,44E-02 |
| ENSG00000175215 | CTDSP2 | 1002,89 | -1,03 | 0,26 | -3,96 | 7,37E-05 | 2,07E-03 |
| ENSG00000070778 | PTPN21 | 119,07 | -1,02 | 0,23 | -4,50 | 6,81E-06 | 4,85E-04 |
| ENSG00000106689 | LHX2 | 1068,10 | -1,02 | 0,31 | -3,30 | 9,82E-04 | 1,18E-02 |
| ENSG00000183688 | RFLNB | 152,43 | -1,02 | 0,31 | -3,34 | 8,37E-04 | 1,05E-02 |
| ENSG00000069122 | ADGRF5 | 388,54 | -1,02 | 0,32 | -3,19 | 1,41E-03 | 1,53E-02 |
| ENSG00000151491 | EPS8 | 110,37 | -1,02 | 0,29 | -3,48 | 5,09E-04 | 7,42E-03 |
| ENSG00000137776 | SLTM | 2795,74 | -1,02 | 0,36 | -2,83 | 4,64E-03 | 3,43E-02 |
| ENSG00000100987 | VSX1 | 21,58 | -1,02 | 0,38 | -2,69 | 7,20E-03 | 4,66E-02 |
| ENSG00000213563 | C8orf82 | 562,98 | -1,02 | 0,32 | -3,23 | 1,25E-03 | 1,40E-02 |
| ENSG00000135414 | GDF11 | 881,27 | -1,02 | 0,30 | -3,40 | 6,77E-04 | 8,96E-03 |
| ENSG00000049239 | H6PD | 534,10 | -1,02 | 0,33 | -3,10 | 1,95E-03 | 1,92E-02 |
| ENSG00000132819 | RBM38 | 260,13 | -1,02 | 0,29 | -3,50 | 4,69E-04 | 7,02E-03 |
| ENSG00000075426 | FOSL2 | 485,77 | -1,02 | 0,23 | -4,45 | 8,66E-06 | 5,48E-04 |
| ENSG00000170275 | CRTAP | 1380,46 | -1,02 | 0,27 | -3,79 | 1,51E-04 | 3,36E-03 |
| ENSG00000146063 | TRIM41 | 1216,77 | -1,02 | 0,24 | -4,25 | 2,14E-05 | 9,44E-04 |
| ENSG00000185187 | SIGIRR | 706,85 | -1,01 | 0,35 | -2,90 | 3,74E-03 | 2,98E-02 |
| ENSG00000144677 | CTDSPL | 762,50 | -1,01 | 0,38 | -2,70 | 7,03E-03 | 4,59E-02 |
| ENSG00000144674 | GOLGA4 | 1240,53 | -1,01 | 0,38 | -2,70 | 6,87E-03 | 4,52E-02 |
| ENSG00000105643 | ARRDC2 | 952,08 | -1,01 | 0,32 | -3,14 | 1,70E-03 | 1,74E-02 |
| ENSG00000114439 | BBX | 1417,10 | -1,01 | 0,37 | -2,75 | 5,98E-03 | 4,08E-02 |
| ENSG00000138166 | DUSP5 | 164,54 | -1,01 | 0,31 | -3,23 | 1,22E-03 | 1,38E-02 |
| ENSG00000187498 | COL4A1 | 81,52 | -1,01 | 0,38 | -2,68 | 7,30E-03 | 4,71E-02 |
| ENSG00000122756 | CNTFR | 916,70 | -1,01 | 0,32 | -3,17 | 1,51E-03 | 1,61E-02 |
| ENSG00000140905 | GCSH | 563,41 | -1,01 | 0,19 | -5,42 | 6,06E-08 | 2,09E-05 |
| ENSG00000069431 | ABCC9 | 96,52 | -1,01 | 0,33 | -3,10 | 1,94E-03 | 1,91E-02 |

| | | | | | | | |
|---|---|---|---|---|---|---|---|
| ENSG00000155760 | FZD7 | 41,95 | -1,01 | 0,38 | -2,67 | 7,65E-03 | 4,84E-02 |
| ENSG00000177303 | CASKIN2 | 750,23 | -1,01 | 0,28 | -3,66 | 2,49E-04 | 4,65E-03 |
| ENSG00000165092 | ALDH1A1 | 548,65 | -1,01 | 0,26 | -3,83 | 1,31E-04 | 3,06E-03 |
| ENSG00000273136 | NBPF26 | 492,76 | -1,01 | 0,33 | -3,10 | 1,97E-03 | 1,92E-02 |
| ENSG00000144821 | MYH15 | 56,06 | -1,01 | 0,33 | -3,04 | 2,36E-03 | 2,16E-02 |
| ENSG00000161921 | CXCL16 | 128,84 | -1,01 | 0,32 | -3,18 | 1,50E-03 | 1,60E-02 |
| ENSG00000142733 | MAP3K6 | 265,45 | -1,00 | 0,36 | -2,76 | 5,74E-03 | 3,96E-02 |
| ENSG00000175482 | POLD4 | 581,18 | -1,00 | 0,32 | -3,12 | 1,84E-03 | 1,84E-02 |
| ENSG00000173801 | JUP | 508,53 | -1,00 | 0,35 | -2,86 | 4,29E-03 | 3,26E-02 |
| ENSG00000109790 | KLHL5 | 1224,83 | -1,00 | 0,25 | -4,02 | 5,83E-05 | 1,79E-03 |
| ENSG00000165806 | CASP7 | 57,18 | -1,00 | 0,35 | -2,88 | 3,93E-03 | 3,06E-02 |
| ENSG00000253598 | SLC10A5 | 10,91 | 1,00 | 0,36 | 2,80 | 5,13E-03 | 3,66E-02 |
| ENSG00000103154 | NECAB2 | 1410,59 | 1,00 | 0,25 | 3,95 | 7,82E-05 | 2,13E-03 |
| ENSG00000120875 | DUSP4 | 152,71 | 1,00 | 0,34 | 2,92 | 3,56E-03 | 2,87E-02 |
| ENSG00000157778 | PSMG3 | 553,04 | 1,00 | 0,27 | 3,77 | 1,66E-04 | 3,56E-03 |
| ENSG00000204052 | LRRC73 | 287,90 | 1,00 | 0,31 | 3,24 | 1,21E-03 | 1,37E-02 |
| ENSG00000089847 | ANKRD24 | 1681,00 | 1,01 | 0,24 | 4,13 | 3,62E-05 | 1,31E-03 |
| ENSG00000146221 | TCTE1 | 1137,24 | 1,01 | 0,33 | 3,05 | 2,29E-03 | 2,13E-02 |
| ENSG00000100276 | RASL10A | 570,53 | 1,01 | 0,28 | 3,58 | 3,48E-04 | 5,78E-03 |
| ENSG00000111218 | PRMT8 | 437,78 | 1,02 | 0,32 | 3,17 | 1,54E-03 | 1,63E-02 |
| ENSG00000120088 | CRHR1 | 219,80 | 1,02 | 0,35 | 2,89 | 3,86E-03 | 3,03E-02 |
| ENSG00000172824 | CES4A | 731,26 | 1,02 | 0,33 | 3,14 | 1,68E-03 | 1,73E-02 |
| ENSG00000237763 | AMY1A | 86,17 | 1,02 | 0,38 | 2,69 | 7,20E-03 | 4,66E-02 |
| ENSG00000204248 | COL11A2 | 227,68 | 1,02 | 0,28 | 3,72 | 2,01E-04 | 4,05E-03 |
| ENSG00000164076 | CAMKV | 2800,25 | 1,02 | 0,34 | 3,00 | 2,66E-03 | 2,34E-02 |
| ENSG00000105613 | MAST1 | 1768,12 | 1,03 | 0,32 | 3,17 | 1,54E-03 | 1,63E-02 |
| ENSG00000125931 | CITED1 | 98,84 | 1,03 | 0,38 | 2,68 | 7,27E-03 | 4,69E-02 |
| ENSG00000102879 | CORO1A | 1189,44 | 1,04 | 0,27 | 3,86 | 1,13E-04 | 2,75E-03 |
| ENSG00000048991 | R3HDM1 | 1165,64 | 1,04 | 0,37 | 2,80 | 5,07E-03 | 3,63E-02 |
| ENSG00000173267 | SNCG | 4174,63 | 1,04 | 0,31 | 3,36 | 7,73E-04 | 9,91E-03 |
| ENSG00000187559 | FOXD4L3 | 20,22 | 1,04 | 0,35 | 2,95 | 3,20E-03 | 2,67E-02 |
| ENSG00000204149 | AGAP6 | 247,52 | 1,04 | 0,33 | 3,18 | 1,45E-03 | 1,56E-02 |
| ENSG00000176155 | CCDC57 | 581,93 | 1,04 | 0,33 | 3,14 | 1,67E-03 | 1,72E-02 |
| ENSG00000162931 | TRIM17 | 155,88 | 1,05 | 0,31 | 3,34 | 8,44E-04 | 1,06E-02 |
| ENSG00000142185 | TRPM2 | 841,75 | 1,05 | 0,24 | 4,46 | 8,13E-06 | 5,39E-04 |
| ENSG00000260001 | TGFBR3L | 799,51 | 1,05 | 0,27 | 3,89 | 1,01E-04 | 2,56E-03 |
| ENSG00000103723 | AP3B2 | 1522,23 | 1,05 | 0,38 | 2,81 | 5,02E-03 | 3,61E-02 |
| ENSG00000117425 | PTCH2 | 205,63 | 1,05 | 0,29 | 3,61 | 3,12E-04 | 5,35E-03 |
| ENSG00000008710 | PKD1 | 4151,80 | 1,05 | 0,25 | 4,16 | 3,19E-05 | 1,19E-03 |
| ENSG00000166257 | SCN3B | 3898,30 | 1,06 | 0,34 | 3,10 | 1,94E-03 | 1,91E-02 |
| ENSG00000178440 | TIMM23B-AGAP6 | 241,67 | 1,06 | 0,33 | 3,23 | 1,22E-03 | 1,38E-02 |
| ENSG00000130822 | PNCK | 1756,84 | 1,06 | 0,28 | 3,82 | 1,35E-04 | 3,12E-03 |
| ENSG00000100156 | SLC16A8 | 54,92 | 1,06 | 0,39 | 2,74 | 6,06E-03 | 4,11E-02 |
| ENSG00000176454 | LPCAT4 | 1505,62 | 1,06 | 0,37 | 2,90 | 3,71E-03 | 2,96E-02 |
| ENSG00000137968 | SLC44A5 | 54,88 | 1,06 | 0,37 | 2,89 | 3,82E-03 | 3,02E-02 |
| ENSG00000155428 | TRIM74 | 50,00 | 1,07 | 0,37 | 2,89 | 3,79E-03 | 3,00E-02 |
| ENSG00000142408 | CACNG8 | 3699,34 | 1,07 | 0,29 | 3,71 | 2,06E-04 | 4,10E-03 |
| ENSG00000102385 | DRP2 | 445,12 | 1,07 | 0,34 | 3,12 | 1,78E-03 | 1,80E-02 |
| ENSG00000176956 | LY6H | 4734,55 | 1,08 | 0,27 | 3,98 | 6,90E-05 | 1,99E-03 |
| ENSG00000205702 | CYP2D7 | 138,63 | 1,08 | 0,38 | 2,82 | 4,86E-03 | 3,53E-02 |
| ENSG00000149527 | PLCH2 | 907,67 | 1,09 | 0,36 | 3,04 | 2,36E-03 | 2,16E-02 |
| ENSG00000188011 | RTP5 | 337,69 | 1,09 | 0,31 | 3,51 | 4,52E-04 | 6,86E-03 |
| ENSG00000132386 | SERPINF1 | 678,89 | 1,10 | 0,41 | 2,65 | 7,97E-03 | 4,99E-02 |
| ENSG00000105278 | ZFR2 | 439,17 | 1,10 | 0,29 | 3,85 | 1,19E-04 | 2,86E-03 |
| ENSG00000006283 | CACNA1G | 430,93 | 1,10 | 0,40 | 2,76 | 5,79E-03 | 3,99E-02 |
| ENSG00000022556 | NLRP2 | 51,46 | 1,11 | 0,33 | 3,39 | 6,90E-04 | 9,07E-03 |
| ENSG00000184515 | BEX5 | 2131,56 | 1,11 | 0,36 | 3,10 | 1,93E-03 | 1,90E-02 |

| | | | | | | | |
|---|---|---|---|---|---|---|---|
| ENSG00000258366 | RTEL1 | 439,29 | 1,12 | 0,27 | 4,16 | 3,18E-05 | 1,19E-03 |
| ENSG00000184709 | LRRC26 | 20,29 | 1,12 | 0,33 | 3,37 | 7,44E-04 | 9,63E-03 |
| ENSG00000171462 | DLK2 | 157,05 | 1,12 | 0,32 | 3,51 | 4,47E-04 | 6,80E-03 |
| ENSG00000077080 | ACTL6B | 938,83 | 1,12 | 0,39 | 2,87 | 4,12E-03 | 3,17E-02 |
| ENSG00000220201 | ZGLP1 | 264,99 | 1,12 | 0,38 | 2,97 | 2,98E-03 | 2,54E-02 |
| ENSG00000165568 | AKR1E2 | 18,82 | 1,12 | 0,39 | 2,91 | 3,65E-03 | 2,93E-02 |
| ENSG00000172260 | NEGR1 | 3989,34 | 1,12 | 0,34 | 3,30 | 9,59E-04 | 1,16E-02 |
| ENSG00000086506 | HBQ1 | 69,24 | 1,12 | 0,37 | 3,05 | 2,33E-03 | 2,15E-02 |
| ENSG00000101222 | SPEF1 | 191,65 | 1,12 | 0,21 | 5,26 | 1,44E-07 | 3,47E-05 |
| ENSG00000172137 | CALB2 | 326,98 | 1,12 | 0,31 | 3,66 | 2,51E-04 | 4,69E-03 |
| ENSG00000261793 | AL929554.1 | 17,48 | 1,13 | 0,37 | 3,05 | 2,26E-03 | 2,11E-02 |
| ENSG00000124839 | RAB17 | 22,05 | 1,14 | 0,39 | 2,89 | 3,83E-03 | 3,02E-02 |
| ENSG00000129951 | PLPPR3 | 821,31 | 1,14 | 0,34 | 3,33 | 8,59E-04 | 1,07E-02 |
| ENSG00000165434 | PGM2L1 | 2358,52 | 1,15 | 0,33 | 3,47 | 5,18E-04 | 7,45E-03 |
| ENSG00000100197 | CYP2D6 | 99,21 | 1,16 | 0,43 | 2,70 | 6,89E-03 | 4,52E-02 |
| ENSG00000144214 | LYG1 | 16,02 | 1,16 | 0,36 | 3,23 | 1,22E-03 | 1,38E-02 |
| ENSG00000129749 | CHRNA10 | 12,39 | 1,16 | 0,41 | 2,82 | 4,79E-03 | 3,50E-02 |
| ENSG00000171189 | GRIK1 | 205,27 | 1,16 | 0,29 | 4,04 | 5,26E-05 | 1,68E-03 |
| ENSG00000128482 | RNF112 | 802,64 | 1,16 | 0,30 | 3,84 | 1,21E-04 | 2,90E-03 |
| ENSG00000139044 | B4GALNT3 | 200,16 | 1,17 | 0,35 | 3,28 | 1,02E-03 | 1,21E-02 |
| ENSG00000162711 | NLRP3 | 40,17 | 1,17 | 0,40 | 2,91 | 3,67E-03 | 2,95E-02 |
| ENSG00000203499 | IQANK1 | 34,86 | 1,18 | 0,43 | 2,71 | 6,69E-03 | 4,43E-02 |
| ENSG00000135127 | BICDL1 | 400,29 | 1,18 | 0,39 | 3,03 | 2,46E-03 | 2,23E-02 |
| ENSG00000120903 | CHRNA2 | 89,90 | 1,19 | 0,41 | 2,92 | 3,52E-03 | 2,85E-02 |
| ENSG00000164082 | GRM2 | 327,01 | 1,19 | 0,38 | 3,14 | 1,70E-03 | 1,74E-02 |
| ENSG00000119973 | PRLHR | 14,03 | 1,19 | 0,44 | 2,69 | 7,07E-03 | 4,61E-02 |
| ENSG00000187730 | GABRD | 1841,00 | 1,19 | 0,39 | 3,09 | 2,00E-03 | 1,95E-02 |
| ENSG00000215018 | COL28A1 | 28,89 | 1,20 | 0,36 | 3,35 | 8,16E-04 | 1,03E-02 |
| ENSG00000172247 | C1QTNF4 | 1290,79 | 1,20 | 0,32 | 3,72 | 1,96E-04 | 4,00E-03 |
| ENSG00000167968 | DNASE1L2 | 110,01 | 1,20 | 0,33 | 3,60 | 3,23E-04 | 5,50E-03 |
| ENSG00000091536 | MYO15A | 134,76 | 1,20 | 0,31 | 3,92 | 8,77E-05 | 2,31E-03 |
| ENSG00000082684 | SEMA5B | 168,86 | 1,22 | 0,34 | 3,56 | 3,76E-04 | 6,10E-03 |
| ENSG00000149742 | SLC22A9 | 32,65 | 1,23 | 0,38 | 3,28 | 1,04E-03 | 1,22E-02 |
| ENSG00000171502 | COL24A1 | 69,66 | 1,24 | 0,43 | 2,89 | 3,79E-03 | 3,00E-02 |
| ENSG00000188747 | NOXA1 | 965,13 | 1,25 | 0,35 | 3,54 | 4,01E-04 | 6,36E-03 |
| ENSG00000234965 | SHISA8 | 80,92 | 1,25 | 0,30 | 4,16 | 3,18E-05 | 1,19E-03 |
| ENSG00000129990 | SYT5 | 2788,59 | 1,26 | 0,40 | 3,13 | 1,77E-03 | 1,79E-02 |
| ENSG00000136883 | KIF12 | 70,66 | 1,27 | 0,30 | 4,18 | 2,93E-05 | 1,15E-03 |
| ENSG00000132744 | ACY3 | 50,47 | 1,27 | 0,43 | 2,94 | 3,28E-03 | 2,71E-02 |
| ENSG00000243480 | AMY2A | 87,19 | 1,28 | 0,47 | 2,69 | 7,12E-03 | 4,63E-02 |
| ENSG00000138100 | TRIM54 | 186,30 | 1,28 | 0,37 | 3,41 | 6,39E-04 | 8,62E-03 |
| ENSG00000161944 | ASGR2 | 20,95 | 1,29 | 0,38 | 3,38 | 7,31E-04 | 9,52E-03 |
| ENSG00000135596 | MICAL1 | 1070,53 | 1,30 | 0,34 | 3,77 | 1,64E-04 | 3,52E-03 |
| ENSG00000187094 | CCK | 3101,92 | 1,30 | 0,48 | 2,70 | 6,88E-03 | 4,52E-02 |
| ENSG00000059588 | TARBP1 | 498,41 | 1,31 | 0,40 | 3,29 | 1,01E-03 | 1,20E-02 |
| ENSG00000063438 | AHRR | 53,62 | 1,31 | 0,44 | 3,00 | 2,69E-03 | 2,36E-02 |
| ENSG00000128422 | KRT17 | 569,79 | 1,31 | 0,42 | 3,10 | 1,95E-03 | 1,92E-02 |
| ENSG00000107593 | PKD2L1 | 53,18 | 1,32 | 0,39 | 3,37 | 7,57E-04 | 9,76E-03 |
| ENSG00000254858 | MPV17L2 | 1492,20 | 1,32 | 0,29 | 4,52 | 6,32E-06 | 4,64E-04 |
| ENSG00000127586 | CHTF18 | 316,10 | 1,33 | 0,29 | 4,62 | 3,85E-06 | 3,50E-04 |
| ENSG00000268041 | ERFL | 17,42 | 1,33 | 0,36 | 3,68 | 2,34E-04 | 4,48E-03 |
| ENSG00000118194 | TNNT2 | 608,31 | 1,34 | 0,36 | 3,74 | 1,84E-04 | 3,85E-03 |
| ENSG00000165643 | SOHLH1 | 835,47 | 1,34 | 0,42 | 3,18 | 1,47E-03 | 1,57E-02 |
| ENSG00000154146 | NRGN | 43007,74 | 1,34 | 0,40 | 3,37 | 7,44E-04 | 9,63E-03 |
| ENSG00000091651 | ORC6 | 51,18 | 1,35 | 0,49 | 2,77 | 5,66E-03 | 3,93E-02 |
| ENSG00000167549 | CORO6 | 1501,92 | 1,36 | 0,42 | 3,26 | 1,11E-03 | 1,29E-02 |
| ENSG00000188886 | ASTL | 10,86 | 1,37 | 0,41 | 3,35 | 8,05E-04 | 1,02E-02 |

| | | | | | | | |
|---|---|---|---|---|---|---|---|
| ENSG00000164418 | GRIK2 | 409,58 | 1,37 | 0,31 | 4,37 | 1,22E-05 | 6,82E-04 |
| ENSG00000169436 | COL22A1 | 40,46 | 1,38 | 0,47 | 2,94 | 3,32E-03 | 2,74E-02 |
| ENSG00000107831 | FGF8 | 11,94 | 1,39 | 0,47 | 2,92 | 3,49E-03 | 2,83E-02 |
| ENSG00000196408 | NOXO1 | 70,17 | 1,39 | 0,31 | 4,47 | 7,95E-06 | 5,34E-04 |
| ENSG00000179698 | WDR97 | 105,50 | 1,41 | 0,36 | 3,90 | 9,51E-05 | 2,46E-03 |
| ENSG00000198883 | PNMA5 | 216,62 | 1,42 | 0,45 | 3,14 | 1,70E-03 | 1,74E-02 |
| ENSG00000140798 | ABCC12 | 87,28 | 1,43 | 0,51 | 2,78 | 5,48E-03 | 3,85E-02 |
| ENSG00000179673 | RPRML | 345,52 | 1,45 | 0,46 | 3,13 | 1,75E-03 | 1,78E-02 |
| ENSG00000241563 | CORT | 279,49 | 1,48 | 0,36 | 4,13 | 3,66E-05 | 1,32E-03 |
| ENSG00000130643 | CALY | 4185,83 | 1,49 | 0,37 | 3,97 | 7,24E-05 | 2,05E-03 |
| ENSG00000183837 | PNMA3 | 1614,82 | 1,49 | 0,43 | 3,48 | 5,08E-04 | 7,42E-03 |
| ENSG00000062524 | LTK | 82,92 | 1,49 | 0,34 | 4,34 | 1,42E-05 | 7,41E-04 |
| ENSG00000254656 | RTL1 | 9,24 | 1,51 | 0,51 | 2,94 | 3,29E-03 | 2,72E-02 |
| ENSG00000130649 | CYP2E1 | 432,32 | 1,51 | 0,46 | 3,29 | 9,84E-04 | 1,18E-02 |
| ENSG00000171772 | SYCE1 | 337,77 | 1,53 | 0,44 | 3,48 | 4,95E-04 | 7,30E-03 |
| ENSG00000124713 | GNMT | 15,26 | 1,55 | 0,50 | 3,12 | 1,79E-03 | 1,80E-02 |
| ENSG00000144550 | CPNE9 | 455,04 | 1,56 | 0,41 | 3,78 | 1,57E-04 | 3,44E-03 |
| ENSG00000181085 | MAPK15 | 108,07 | 1,56 | 0,59 | 2,66 | 7,75E-03 | 4,88E-02 |
| ENSG00000166863 | TAC3 | 126,13 | 1,58 | 0,52 | 3,01 | 2,63E-03 | 2,32E-02 |
| ENSG00000185615 | PDIA2 | 1315,76 | 1,58 | 0,44 | 3,59 | 3,35E-04 | 5,64E-03 |
| ENSG00000092295 | TGM1 | 23,22 | 1,58 | 0,46 | 3,43 | 6,10E-04 | 8,34E-03 |
| ENSG00000215788 | TNFRSF25 | 570,96 | 1,58 | 0,45 | 3,52 | 4,27E-04 | 6,64E-03 |
| ENSG00000177984 | LCN15 | 67,95 | 1,60 | 0,45 | 3,54 | 3,98E-04 | 6,34E-03 |
| ENSG00000226490 | AC138647.1 | 52,06 | 1,61 | 0,44 | 3,62 | 2,93E-04 | 5,12E-03 |
| ENSG00000261587 | TMEM249 | 52,19 | 1,64 | 0,39 | 4,16 | 3,18E-05 | 1,19E-03 |
| ENSG00000162006 | MSLNL | 35,54 | 1,70 | 0,56 | 3,06 | 2,19E-03 | 2,07E-02 |
| ENSG00000203782 | LORICRIN | 11,30 | 1,80 | 0,53 | 3,42 | 6,17E-04 | 8,38E-03 |
| ENSG00000164326 | CARTPT | 391,77 | 1,82 | 0,59 | 3,09 | 1,99E-03 | 1,95E-02 |
| ENSG00000197261 | C6orf141 | 12,30 | 1,84 | 0,69 | 2,66 | 7,70E-03 | 4,86E-02 |
| ENSG00000115718 | PROC | 7,91 | 1,97 | 0,64 | 3,09 | 2,03E-03 | 1,96E-02 |
| ENSG00000260287 | TBC1D3G | 311,47 | 2,05 | 0,46 | 4,43 | 9,35E-06 | 5,77E-04 |
| ENSG00000214946 | TBC1D26 | 57,72 | 2,09 | 0,55 | 3,80 | 1,42E-04 | 3,22E-03 |
| ENSG00000283599 | BX276092.9 | 11,96 | 2,28 | 0,60 | 3,80 | 1,42E-04 | 3,22E-03 |
| ENSG00000105675 | ATP4A | 20,11 | 2,39 | 0,60 | 3,98 | 6,78E-05 | 1,98E-03 |
| ENSG00000187848 | P2RX2 | 24,31 | 3,01 | 0,67 | 4,50 | 6,80E-06 | 4,85E-04 |
