## Supplementary tables for "Transcriptome profiling of the dorsomedial prefrontal cortex in suicide victims": S4_Functional_enrichment_of_DEGs.pdf

Supplementary file 4: Functional enrichment of down- and upregulated genes.

| Cluster | Category | ID | Description | p.adjust | query_size | Count | term_size | effective_d | geneID | GeneRatio | BgRatio |
|---|---|---|---|---|---|---|---|---|---|---|---|
| down-regulated | CORUM | CORUM:2383 | ITGA5-ITGB1-FN1-TGM2 complex | 0.00373459429787653 | 255 | 4 | 4 | 3627 | FN1/ITGA5 | 4/255 | 4/3627 |
| down-regulated | GO:BP | GO:0007166 | cell surface receptor signaling pathway | 1.65140377035333e-29 | 1134 | 362 | 3226 | 18123 | ACKR3/AC | 362/1134 | 3226/18123 |
| down-regulated | GO:BP | GO:0001944 | vasculature development | 3.48324300679902e-29 | 1134 | 149 | 834 | 18123 | ACKR3/AC | 149/1134 | 834/18123 |
| down-regulated | GO:BP | GO:0010033 | response to organic substance | 8.49577212833826e-29 | 1134 | 378 | 3457 | 18123 | ABCC9/AC | 378/1134 | 3457/18123 |
| down-regulated | GO:BP | GO:0072359 | circulatory system development | 1.08416571141918e-28 | 1134 | 187 | 1215 | 18123 | ACACB/AC | 187/1134 | 1215/18123 |
| down-regulated | GO:BP | GO:0048513 | animal organ development | 1.59969687881114e-27 | 1134 | 393 | 3701 | 18123 | ACACB/AC | 393/1134 | 3701/18123 |
| down-regulated | GO:BP | GO:0009653 | anatomical structure morphogenesis | 1.65329247100238e-27 | 1134 | 324 | 2816 | 18123 | ACKR3/AC | 324/1134 | 2816/18123 |
| down-regulated | GO:BP | GO:0001568 | blood vessel development | 1.66258001820629e-27 | 1134 | 142 | 797 | 18123 | ACKR3/AC | 142/1134 | 797/18123 |
| down-regulated | GO:BP | GO:0048731 | system development | 2.57654926971967e-26 | 1134 | 491 | 5095 | 18123 | ACACB/AC | 491/1134 | 5095/18123 |
| down-regulated | GO:BP | GO:0070887 | cellular response to chemical stimulus | 4.88661024236411e-26 | 1134 | 369 | 3443 | 18123 | ACAA2/AC | 369/1134 | 3443/18123 |
| down-regulated | GO:BP | GO:0007165 | signal transduction | 7.98855752624899e-26 | 1134 | 582 | 6447 | 18123 | ACAA2/AC | 582/1134 | 6447/18123 |
| down-regulated | GO:BP | GO:0050896 | response to stimulus | 1.29287075357542e-25 | 1134 | 773 | 9487 | 18123 | ABCB1/AB | 773/1134 | 9487/18123 |
| down-regulated | GO:BP | GO:0050793 | regulation of developmental process | 1.41965522951415e-25 | 1134 | 303 | 2621 | 18123 | ACACB/AC | 303/1134 | 2621/18123 |
| down-regulated | GO:BP | GO:0071310 | cellular response to organic substance | 1.50836924314677e-25 | 1134 | 319 | 2821 | 18123 | ACKR3/AC | 319/1134 | 2821/18123 |
| down-regulated | GO:BP | GO:0023052 | signaling | 2.04254734578716e-25 | 1134 | 613 | 6937 | 18123 | ABCC9/AC | 613/1134 | 6937/18123 |
| down-regulated | GO:BP | GO:0007275 | multicellular organism development | 2.17116113752663e-25 | 1134 | 528 | 5670 | 18123 | ACACB/AC | 528/1134 | 5670/18123 |
| down-regulated | GO:BP | GO:0007154 | cell communication | 5.99301351644535e-25 | 1134 | 613 | 6961 | 18123 | ACAA2/AC | 613/1134 | 6961/18123 |
| down-regulated | GO:BP | GO:0051239 | regulation of multicellular organismal process | 9.99894776725061e-25 | 1134 | 322 | 2888 | 18123 | ABCC9/AC | 322/1134 | 2888/18123 |
| down-regulated | GO:BP | GO:0023051 | regulation of signaling | 1.20729135327945e-24 | 1134 | 382 | 3669 | 18123 | ABCC9/AC | 382/1134 | 3669/18123 |
| down-regulated | GO:BP | GO:0048856 | anatomical structure development | 1.30215939527507e-24 | 1134 | 559 | 6164 | 18123 | ACACB/AC | 559/1134 | 6164/18123 |
| down-regulated | GO:BP | GO:0032502 | developmental process | 3.07536029249744e-24 | 1134 | 592 | 6678 | 18123 | ACACB/AC | 592/1134 | 6678/18123 |
| down-regulated | GO:BP | GO:0009966 | regulation of signal transduction | 1.35236423494065e-23 | 1134 | 347 | 3251 | 18123 | ACAA2/AC | 347/1134 | 3251/18123 |
| down-regulated | GO:BP | GO:0048514 | blood vessel morphogenesis | 1.47024240996376e-23 | 1134 | 126 | 715 | 18123 | ACKR3/AC | 126/1134 | 715/18123 |
| down-regulated | GO:BP | GO:0048583 | regulation of response to stimulus | 2.3586764292265e-23 | 1134 | 433 | 4416 | 18123 | ABCB1/AC | 433/1134 | 4416/18123 |
| down-regulated | GO:BP | GO:0048646 | anatomical structure formation involved in morphogenesis | 2.57987517128582e-23 | 1134 | 175 | 1209 | 18123 | ACKR3/AC | 175/1134 | 1209/18123 |
| down-regulated | GO:BP | GO:0010646 | regulation of cell communication | 3.13829485210255e-23 | 1134 | 375 | 3634 | 18123 | ACAA2/AC | 375/1134 | 3634/18123 |
| down-regulated | GO:BP | GO:0035239 | tube morphogenesis | 4.24788477650198e-23 | 1134 | 150 | 956 | 18123 | ACKR3/AC | 150/1134 | 956/18123 |
| down-regulated | GO:BP | GO:0048518 | positive regulation of biological process | 6.13543300249754e-23 | 1134 | 574 | 6474 | 18123 | ABCB1/AC | 574/1134 | 6474/18123 |
| down-regulated | GO:BP | GO:0009967 | positive regulation of signal transduction | 7.29863259819095e-23 | 1134 | 212 | 1626 | 18123 | ACKR3/AC | 212/1134 | 1626/18123 |
| down-regulated | GO:BP | GO:0035295 | tube development | 8.68260219962858e-23 | 1134 | 169 | 1158 | 18123 | ACKR3/AC | 169/1134 | 1158/18123 |
| down-regulated | GO:BP | GO:0048522 | positive regulation of cellular process | 1.138640843175e-22 | 1134 | 532 | 5863 | 18123 | ABCB1/AC | 532/1134 | 5863/18123 |
| down-regulated | GO:BP | GO:0051716 | cellular response to stimulus | 6.95283177508684e-22 | 1134 | 661 | 7873 | 18123 | ACAA2/AC | 661/1134 | 7873/18123 |
| down-regulated | GO:BP | GO:0048584 | positive regulation of response to stimulus | 8.06239858397139e-22 | 1134 | 272 | 2368 | 18123 | ACKR3/AC | 272/1134 | 2368/18123 |
| down-regulated | GO:BP | GO:0022610 | biological adhesion | 8.43868712881391e-22 | 1134 | 199 | 1509 | 18123 | ACKR3/AC | 199/1134 | 1509/18123 |
| down-regulated | GO:BP | GO:0007155 | cell adhesion | 3.06408224045942e-21 | 1134 | 197 | 1502 | 18123 | ACKR3/AC | 197/1134 | 1502/18123 |
| down-regulated | GO:BP | GO:0023056 | positive regulation of signaling | 3.70949962450978e-21 | 1134 | 224 | 1814 | 18123 | ACKR3/AC | 224/1134 | 1814/18123 |
| down-regulated | GO:BP | GO:0042221 | response to chemical | 3.99215825938387e-21 | 1134 | 455 | 4829 | 18123 | ABCB1/AB | 455/1134 | 4829/18123 |
| down-regulated | GO:BP | GO:2000026 | regulation of multicellular organismal development | 9.10562160738664e-21 | 1134 | 192 | 1459 | 18123 | ACACB/AC | 192/1134 | 1459/18123 |
| down-regulated | GO:BP | GO:0048869 | cellular developmental process | 9.60264978663947e-21 | 1134 | 428 | 4468 | 18123 | ACKR3/AC | 428/1134 | 4468/18123 |
| down-regulated | GO:BP | GO:0030154 | cell differentiation | 1.38297077777399e-20 | 1134 | 422 | 4392 | 18123 | ACKR3/AC | 422/1134 | 4392/18123 |
| down-regulated | GO:BP | GO:0010647 | positive regulation of cell communication | 1.45751128176454e-20 | 1134 | 222 | 1809 | 18123 | ACKR3/AC | 222/1134 | 1809/18123 |
| down-regulated | GO:BP | GO:0032501 | multicellular organismal process | 3.12937128809702e-20 | 1134 | 663 | 8000 | 18123 | ABCB1/AB | 663/1134 | 8000/18123 |
| down-regulated | GO:BP | GO:0006950 | response to stress | 2.43553059000398e-18 | 1134 | 409 | 4321 | 18123 | ABCB1/AB | 409/1134 | 4321/18123 |
| down-regulated | GO:BP | GO:0001525 | angiogenesis | 3.06028238583952e-18 | 1134 | 106 | 621 | 18123 | ACKR3/AC | 106/1134 | 621/18123 |
| down-regulated | GO:BP | GO:0016477 | cell migration | 3.15096462615355e-18 | 1134 | 203 | 1660 | 18123 | ACKR3/AC | 203/1134 | 1660/18123 |
| down-regulated | GO:BP | GO:0035556 | intracellular signal transduction | 3.44425196304271e-18 | 1134 | 302 | 2886 | 18123 | ACKR3/AD | 302/1134 | 2886/18123 |
| down-regulated | GO:BP | GO:0042127 | regulation of cell population proliferation | 5.59100537201081e-18 | 1134 | 212 | 1774 | 18123 | ACKR3/AC | 212/1134 | 1774/18123 |
| down-regulated | GO:BP | GO:0008283 | cell population proliferation | 8.46825709796981e-18 | 1134 | 235 | 2057 | 18123 | ABCB1/AC | 235/1134 | 2057/18123 |
| down-regulated | GO:BP | GO:0045595 | regulation of cell differentiation | 1.01544169909335e-17 | 1134 | 204 | 1688 | 18123 | ACVRL1/A | 204/1134 | 1688/18123 |
| down-regulated | GO:BP | GO:0009611 | response to wounding | 1.19928282833241e-17 | 1134 | 111 | 680 | 18123 | ACVRL1/A | 111/1134 | 680/18123 |
| down-regulated | GO:BP | GO:0040011 | locomotion | 1.30239486058213e-16 | 1134 | 230 | 2039 | 18123 | ACKR3/AC | 230/1134 | 2039/18123 |
| down-regulated | GO:BP | GO:0034097 | response to cytokine | 1.5651775310473e-16 | 1134 | 162 | 1242 | 18123 | ACKR3/AN | 162/1134 | 1242/18123 |
| down-regulated | GO:BP | GO:0048519 | negative regulation of biological process | 1.62057931971357e-16 | 1134 | 509 | 5861 | 18123 | A2ML1/AC | 509/1134 | 5861/18123 |
| down-regulated | GO:BP | GO:0009888 | tissue development | 1.74979394068517e-16 | 1134 | 234 | 2093 | 18123 | ACVRL1/A | 234/1134 | 2093/18123 |
| down-regulated | GO:BP | GO:0051094 | positive regulation of developmental process | 3.5538326244206e-16 | 1134 | 171 | 1354 | 18123 | ACACB/AC | 171/1134 | 1354/18123 |
| down-regulated | GO:BP | GO:0071345 | cellular response to cytokine stimulus | 6.39184218913364e-16 | 1134 | 152 | 1148 | 18123 | ACKR3/AN | 152/1134 | 1148/18123 |
| down-regulated | GO:BP | GO:0002376 | immune system process | 1.16601272679902e-15 | 1134 | 335 | 3438 | 18123 | ABCC9/AC | 335/1134 | 3438/18123 |
| down-regulated | GO:BP | GO:0048523 | negative regulation of cellular process | 1.37630378017462e-15 | 1134 | 457 | 5153 | 18123 | A2ML1/AC | 457/1134 | 5153/18123 |
| down-regulated | GO:BP | GO:0030155 | regulation of cell adhesion | 5.70673183489685e-15 | 1134 | 113 | 756 | 18123 | ACVRL1/A | 113/1134 | 756/18123 |
| down-regulated | GO:BP | GO:0051240 | positive regulation of multicellular organismal process | 6.07162029119063e-15 | 1134 | 180 | 1497 | 18123 | ACACB/AC | 180/1134 | 1497/18123 |
| down-regulated | GO:BP | GO:1901700 | response to oxygen-containing compound | 6.51730275665344e-15 | 1134 | 201 | 1748 | 18123 | ABCC9/AD | 201/1134 | 1748/18123 |
| down-regulated | GO:BP | GO:0051674 | localization of cell | 1.01782956788413e-14 | 1134 | 208 | 1840 | 18123 | ACKR3/AC | 208/1134 | 1840/18123 |
| down-regulated | GO:BP | GO:0048870 | cell motility | 1.01782956788413e-14 | 1134 | 208 | 1840 | 18123 | ACKR3/AC | 208/1134 | 1840/18123 |
| down-regulated | GO:BP | GO:0019221 | cytokine-mediated signaling pathway | 1.03173705827548e-14 | 1134 | 119 | 824 | 18123 | ACKR3/AN | 119/1134 | 824/18123 |
| down-regulated | GO:BP | GO:0042060 | wound healing | 1.26908693430165e-14 | 1134 | 92 | 555 | 18123 | ACVRL1/A | 92/1134 | 555/18123 |
| down-regulated | GO:BP | GO:0045597 | positive regulation of cell differentiation | 1.6079483383035e-14 | 1134 | 125 | 892 | 18123 | ACVRL1/A | 125/1134 | 892/18123 |
| down-regulated | GO:BP | GO:0033993 | response to lipid | 4.01974482179658e-14 | 1134 | 130 | 956 | 18123 | ADCYAP1R | 130/1134 | 956/18123 |
| down-regulated | GO:BP | GO:0009887 | animal organ morphogenesis | 4.86331607539209e-14 | 1134 | 141 | 1079 | 18123 | ACVRL1/A | 141/1134 | 1079/18123 |
| down-regulated | GO:BP | GO:0051093 | negative regulation of developmental process | 6.85532061898829e-14 | 1134 | 132 | 984 | 18123 | ACVRL1/A | 132/1134 | 984/18123 |
| down-regulated | GO:BP | GO:0030334 | regulation of cell migration | 8.1682737526648e-14 | 1134 | 134 | 1008 | 18123 | ACKR3/AC | 134/1134 | 1008/18123 |
| down-regulated | GO:BP | GO:0009605 | response to external stimulus | 9.01605999954915e-14 | 1134 | 304 | 3114 | 18123 | ABCC9/AC | 304/1134 | 3114/18123 |
| down-regulated | GO:BP | GO:0006928 | movement of cell or subcellular component | 1.10825306172462e-13 | 1134 | 243 | 2319 | 18123 | ACKR3/AC | 243/1134 | 2319/18123 |
| down-regulated | GO:BP | GO:0008284 | positive regulation of cell population proliferation | 4.62639234505563e-13 | 1134 | 130 | 985 | 18123 | ACVRL1/A | 130/1134 | 985/18123 |
| down-regulated | GO:BP | GO:0065009 | regulation of molecular function | 4.77180900444864e-13 | 1134 | 307 | 3191 | 18123 | A2ML1/AB | 307/1134 | 3191/18123 |
| down-regulated | GO:BP | GO:0032879 | regulation of localization | 7.53312876635381e-13 | 1134 | 285 | 2906 | 18123 | ABCB1/AC | 285/1134 | 2906/18123 |
| down-regulated | GO:BP | GO:0051270 | regulation of cellular component movement | 8.10388005513777e-13 | 1134 | 144 | 1149 | 18123 | ACKR3/AC | 144/1134 | 1149/18123 |
| down-regulated | GO:BP | GO:2000145 | regulation of cell motility | 8.16867947359421e-13 | 1134 | 137 | 1070 | 18123 | ACKR3/AC | 137/1134 | 1070/18123 |
| down-regulated | GO:BP | GO:0009719 | response to endogenous stimulus | 8.28577190192808e-13 | 1134 | 193 | 1729 | 18123 | ACVRL1/A | 193/1134 | 1729/18123 |
| down-regulated | GO:BP | GO:0051241 | negative regulation of multicellular organismal process | 2.19755881149986e-12 | 1134 | 145 | 1174 | 18123 | ACVRL1/A | 145/1134 | 1174/18123 |
| down-regulated | GO:BP | GO:1902533 | positive regulation of intracellular signal transduction | 2.687843884203e-12 | 1134 | 135 | 1063 | 18123 | ACKR3/AD | 135/1134 | 1063/18123 |
| down-regulated | GO:BP | GO:0098609 | cell-cell adhesion | 3.02540686931741e-12 | 1134 | 120 | 898 | 18123 | ADGRV1/A | 120/1134 | 898/18123 |
| down-regulated | GO:BP | GO:0010941 | regulation of cell death | 3.2777174523168e-12 | 1134 | 192 | 1740 | 18123 | ACAA2/AC | 192/1134 | 1740/18123 |
| down-regulated | GO:BP | GO:0007167 | enzyme linked receptor protein signaling pathway | 5.68544626113705e-12 | 1134 | 138 | 1107 | 18123 | ACVRL1/A | 138/1134 | 1107/18123 |
| down-regulated | GO:BP | GO:0040012 | regulation of locomotion | 8.82029379738001e-12 | 1134 | 138 | 1113 | 18123 | ACKR3/AC | 138/1134 | 1113/18123 |
| down-regulated | GO:BP | GO:1902531 | regulation of intracellular signal transduction | 9.55956891694928e-12 | 1134 | 202 | 1883 | 18123 | ACKR3/AD | 202/1134 | 1883/18123 |
| down-regulated | GO:BP | GO:0060429 | epithelium development | 1.16426582580041e-11 | 1134 | 157 | 1338 | 18123 | ACVRL1/A | 157/1134 | 1338/18123 |
| down-regulated | GO:BP | GO:0002520 | immune system development | 2.66895458580147e-11 | 1134 | 130 | 1037 | 18123 | ADGRF5/A | 130/1134 | 1037/18123 |
| down-regulated | GO:BP | GO:0008219 | cell death | 3.97879539986576e-11 | 1134 | 233 | 2306 | 18123 | ACAA2/AC | 233/1134 | 2306/18123 |
| down-regulated | GO:BP | GO:0030335 | positive regulation of cell migration | 4.05468043462635e-11 | 1134 | 86 | 568 | 18123 | ACKR3/AM | 86/1134 | 568/18123 |
| down-regulated | GO:BP | GO:0009790 | embryo development | 4.34484716460526e-11 | 1134 | 131 | 1055 | 18123 | ACVRL1/A | 131/1134 | 1055/18123 |
| down-regulated | GO:BP | GO:0031325 | positive regulation of cellular metabolic process | 5.17642518329718e-11 | 1134 | 310 | 3342 | 18123 | ACACB/AC | 310/1134 | 3342/18123 |
| down-regulated | GO:BP | GO:0009893 | positive regulation of metabolic process | 5.82523577971033e-11 | 1134 | 348 | 3873 | 18123 | ACACB/AC | 348/1134 | 3873/18123 |
| down-regulated | GO:BP | GO:0014070 | response to organic cyclic compound | 6.66022320789162e-11 | 1134 | 126 | 1004 | 18123 | ABCC9/AC | 126/1134 | 1004/18123 |
| down-regulated | GO:BP | GO:0006915 | apoptotic process | 7.72800265149181e-11 | 1134 | 208 | 1998 | 18123 | ACAA2/AC | 208/1134 | 1998/18123 |
| down-regulated | GO:BP | GO:0002682 | regulation of immune system process | 7.82340293779386e-11 | 1134 | 187 | 1734 | 18123 | ADCY4/AD | 187/1134 | 1734/18123 |
| down-regulated | GO:BP | GO:0048534 | hematopoietic or lymphoid organ development | 8.09261892631614e-11 | 1134 | 124 | 984 | 18123 | ADGRF5/A | 124/1134 | 984/18123 |
| down-regulated | GO:BP | GO:0022603 | regulation of anatomical structure morphogenesis | 1.36399870636808e-10 | 1134 | 129 | 1048 | 18123 | ACVRL1/A | 129/1134 | 1048/18123 |
| down-regulated | GO:BP | GO:0030097 | hemopoiesis | 1.37699217199851e-10 | 1134 | 120 | 946 | 18123 | ADGRF5/A | 120/1134 | 946/18123 |
| down-regulated | GO:BP | GO:0003013 | circulatory system process | 1.43771746359699e-10 | 1134 | 91 | 632 | 18123 | ABCB1/AB | 91/1134 | 632/18123 |
| down-regulated | GO:BP | GO:0045785 | positive regulation of cell adhesion | 1.54767433618848e-10 | 1134 | 72 | 442 | 18123 | ADGRG1/A | 72/1134 | 442/18123 |
| down-regulated | GO:BP | GO:0048729 | tissue morphogenesis | 1.59377493301611e-10 | 1134 | 95 | 675 | 18123 | ACVRL1/A | 95/1134 | 675/18123 |
| down-regulated | GO:BP | GO:2000147 | positive regulation of cell motility | 1.63631565426951e-10 | 1134 | 87 | 592 | 18123 | ACKR3/AM | 87/1134 | 592/18123 |
| down-regulated | GO:BP | GO:0001570 | vasculogenesis | 2.04978415259737e-10 | 1134 | 28 | 84 | 18123 | ACKR3/AD | 28/1134 | 84/18123 |
| down-regulated | GO:BP | GO:0051272 | positive regulation of cellular component movement | 2.12889832650113e-10 | 1134 | 88 | 605 | 18123 | ACKR3/AM | 88/1134 | 605/18123 |
| down-regulated | GO:BP | GO:0042981 | regulation of apoptotic process | 2.98212669173984e-10 | 1134 | 171 | 1560 | 18123 | ACAA2/AC | 171/1134 | 1560/18123 |
| down-regulated | GO:BP | GO:0042325 | regulation of phosphorylation | 4.1058682706479e-10 | 1134 | 159 | 1419 | 18123 | ACVRL1/A | 159/1134 | 1419/18123 |
| down-regulated | GO:BP | GO:0043067 | regulation of programmed cell death | 4.18503550652538e-10 | 1134 | 174 | 1603 | 18123 | ACAA2/AC | 174/1134 | 1603/18123 |
| down-regulated | GO:BP | GO:0010001 | glial cell differentiation | 4.48436656169862e-10 | 1134 | 48 | 233 | 18123 | ADORA2A/ | 48/1134 | 233/18123 |
| down-regulated | GO:BP | GO:0001775 | cell activation | 4.80760584301333e-10 | 1134 | 166 | 1507 | 18123 | ADA2/ADG | 166/1134 | 1507/18123 |

| | | | | | | | | | | | |
|---|---|---|---|---|---|---|---|---|---|---|---|
| down-regulated | GO:BP | GO:0061061 | muscle structure development | 5.29473298895892e-10 | 1134 | 91 | 646 | 18123 | ADM/APLN | 91/1134 | 646/18123 |
| down-regulated | GO:BP | GO:0007423 | sensory organ development | 6.51822229535043e-10 | 1134 | 83 | 565 | 18123 | ACVRL1/AI | 83/1134 | 565/18123 |
| down-regulated | GO:BP | GO:0044093 | positive regulation of molecular function | 6.73826236005831e-10 | 1134 | 187 | 1774 | 18123 | ABCB1/AD | 187/1134 | 1774/18123 |
| down-regulated | GO:BP | GO:0040017 | positive regulation of locomotion | 6.9283135911714e-10 | 1134 | 87 | 607 | 18123 | ACKR3/AM | 87/1134 | 607/18123 |
| down-regulated | GO:BP | GO:0007507 | heart development | 7.91222773455031e-10 | 1134 | 86 | 598 | 18123 | ACACB/AC | 86/1134 | 598/18123 |
| down-regulated | GO:BP | GO:0010604 | positive regulation of macromolecule metabolic process | 9.08151252767244e-10 | 1134 | 321 | 3568 | 18123 | ACVRL1/AI | 321/1134 | 3568/18123 |
| down-regulated | GO:BP | GO:0030198 | extracellular matrix organization | 9.49942040280961e-10 | 1134 | 68 | 419 | 18123 | ADAMTS9/ | 68/1134 | 419/18123 |
| down-regulated | GO:BP | GO:0043062 | extracellular structure organization | 1.06667122930409e-09 | 1134 | 68 | 420 | 18123 | ADAMTS9/ | 68/1134 | 420/18123 |
| down-regulated | GO:BP | GO:0042063 | gliogenesis | 1.23255679892103e-09 | 1134 | 57 | 318 | 18123 | ADGRG1/A | 57/1134 | 318/18123 |
| down-regulated | GO:BP | GO:0045229 | external encapsulating structure organization | 1.34307003678262e-09 | 1134 | 68 | 422 | 18123 | ADAMTS9/ | 68/1134 | 422/18123 |
| down-regulated | GO:BP | GO:0051179 | localization | 1.364698828061e-09 | 1134 | 549 | 6938 | 18123 | ABCB1/AB | 549/1134 | 6938/18123 |
| down-regulated | GO:BP | GO:0051174 | regulation of phosphorus metabolic process | 1.60493639134372e-09 | 1134 | 175 | 1640 | 18123 | ACVRL1/AI | 175/1134 | 1640/18123 |
| down-regulated | GO:BP | GO:0019220 | regulation of phosphate metabolic process | 1.60493639134372e-09 | 1134 | 175 | 1640 | 18123 | ACVRL1/AI | 175/1134 | 1640/18123 |
| down-regulated | GO:BP | GO:0045937 | positive regulation of phosphate metabolic process | 1.65464649130664e-09 | 1134 | 122 | 1003 | 18123 | ACVRL1/AI | 122/1134 | 1003/18123 |
| down-regulated | GO:BP | GO:0010562 | positive regulation of phosphorus metabolic process | 1.65464649130664e-09 | 1134 | 122 | 1003 | 18123 | ACVRL1/AI | 122/1134 | 1003/18123 |
| down-regulated | GO:BP | GO:0044419 | biological process involved in interspecies interaction between organ | 1.82784359441747e-09 | 1134 | 185 | 1768 | 18123 | ABCC9/ADI | 185/1134 | 1768/18123 |
| down-regulated | GO:BP | GO:0042327 | positive regulation of phosphorylation | 2.37519804329357e-09 | 1134 | 115 | 928 | 18123 | ACVRL1/AI | 115/1134 | 928/18123 |
| down-regulated | GO:BP | GO:0048771 | tissue remodeling | 3.15687029666688e-09 | 1134 | 41 | 187 | 18123 | ACVRL1/A | 41/1134 | 187/18123 |
| down-regulated | GO:BP | GO:0065008 | regulation of biological quality | 3.73080636308152e-09 | 1134 | 357 | 4116 | 18123 | ABCB1/ABI | 357/1134 | 4116/18123 |
| down-regulated | GO:BP | GO:0009628 | response to abiotic stimulus | 4.02762250000802e-09 | 1134 | 143 | 1264 | 18123 | ABCB1/AC | 143/1134 | 1264/18123 |
| down-regulated | GO:BP | GO:0022612 | gland morphogenesis | 4.63738290514245e-09 | 1134 | 32 | 121 | 18123 | AREG/BCL | 32/1134 | 121/18123 |
| down-regulated | GO:BP | GO:0003158 | endothelium development | 6.1612217053855e-09 | 1134 | 35 | 144 | 18123 | ACVRL1/AI | 35/1134 | 144/18123 |
| down-regulated | GO:BP | GO:0071495 | cellular response to endogenous stimulus | 6.42403532087858e-09 | 1134 | 159 | 1467 | 18123 | ACVRL1/AI | 159/1134 | 1467/18123 |
| down-regulated | GO:BP | GO:0012501 | programmed cell death | 7.73470328374326e-09 | 1134 | 213 | 2158 | 18123 | ACAA2/AC | 213/1134 | 2158/18123 |
| down-regulated | GO:BP | GO:0030029 | actin filament-based process | 8.57507880474248e-09 | 1134 | 103 | 810 | 18123 | ADD3/AIF1 | 103/1134 | 810/18123 |
| down-regulated | GO:BP | GO:0051173 | positive regulation of nitrogen compound metabolic process | 8.7942421443291e-09 | 1134 | 286 | 3143 | 18123 | ACVRL1/AI | 286/1134 | 3143/18123 |
| down-regulated | GO:BP | GO:0071396 | cellular response to lipid | 1.16322274317936e-08 | 1134 | 85 | 617 | 18123 | AKR1C3/AI | 85/1134 | 617/18123 |
| down-regulated | GO:BP | GO:0050794 | regulation of cellular process | 1.2753430365929e-08 | 1134 | 826 | 11517 | 18123 | A2ML1/AB | 826/1134 | 11517/18123 |
| down-regulated | GO:BP | GO:0048598 | embryonic morphogenesis | 1.84964185085132e-08 | 1134 | 83 | 601 | 18123 | ADM/AMC | 83/1134 | 601/18123 |
| down-regulated | GO:BP | GO:0072001 | renal system development | 2.19356960998244e-08 | 1134 | 53 | 303 | 18123 | ANGPT2/B | 53/1134 | 303/18123 |
| down-regulated | GO:BP | GO:0032835 | glomerulus development | 2.32083222826242e-08 | 1134 | 22 | 62 | 18123 | ANGPT2/B | 22/1134 | 62/18123 |
| down-regulated | GO:BP | GO:0070848 | response to growth factor | 2.64335404118451e-08 | 1134 | 98 | 769 | 18123 | ACVRL1/AI | 98/1134 | 769/18123 |
| down-regulated | GO:BP | GO:0048585 | negative regulation of response to stimulus | 2.67696206157203e-08 | 1134 | 194 | 1938 | 18123 | ACAA2/AC | 194/1134 | 1938/18123 |
| down-regulated | GO:BP | GO:0002064 | epithelial cell development | 2.96101240853405e-08 | 1134 | 44 | 225 | 18123 | ADAMTSL4 | 44/1134 | 225/18123 |
| down-regulated | GO:BP | GO:0072006 | nephron development | 3.03663343762032e-08 | 1134 | 34 | 144 | 18123 | ANGPT2/B | 34/1134 | 144/18123 |
| down-regulated | GO:BP | GO:0050789 | regulation of biological process | 3.34250988597036e-08 | 1134 | 859 | 12122 | 18123 | A2ML1/AB | 859/1134 | 12122/18123 |
| down-regulated | GO:BP | GO:0048545 | response to steroid hormone | 4.53804554067851e-08 | 1134 | 61 | 385 | 18123 | ADM/AKR1 | 61/1134 | 385/18123 |
| down-regulated | GO:BP | GO:0001894 | tissue homeostasis | 4.5629450783774e-08 | 1134 | 49 | 272 | 18123 | ADGRF5/A | 49/1134 | 272/18123 |
| down-regulated | GO:BP | GO:0001667 | ameboidal-type cell migration | 4.73658232232852e-08 | 1134 | 72 | 496 | 18123 | ACVRL1/AI | 72/1134 | 496/18123 |
| down-regulated | GO:BP | GO:0045596 | negative regulation of cell differentiation | 5.57557231387763e-08 | 1134 | 91 | 701 | 18123 | ACVRL1/AI | 91/1134 | 701/18123 |
| down-regulated | GO:BP | GO:0006952 | defense response | 6.10844757732437e-08 | 1134 | 194 | 1956 | 18123 | ABCC9/AD | 194/1134 | 1956/18123 |
| down-regulated | GO:BP | GO:0006955 | immune response | 7.54832941874496e-08 | 1134 | 238 | 2544 | 18123 | ABCC9/ACI | 238/1134 | 2544/18123 |
| down-regulated | GO:BP | GO:0065007 | biological regulation | 7.82233102059987e-08 | 1134 | 896 | 12802 | 18123 | A2ML1/AB | 896/1134 | 12802/18123 |
| down-regulated | GO:BP | GO:0034341 | response to interferon-gamma | 9.0187263325427e-08 | 1134 | 40 | 198 | 18123 | BST2/CCL2 | 40/1134 | 198/18123 |
| down-regulated | GO:BP | GO:0009725 | response to hormone | 9.0499175358929e-08 | 1134 | 113 | 958 | 18123 | ADCY4/AD | 113/1134 | 958/18123 |
| down-regulated | GO:BP | GO:0001932 | regulation of protein phosphorylation | 9.78186501474923e-08 | 1134 | 137 | 1246 | 18123 | ACVRL1/AI | 137/1134 | 1246/18123 |
| down-regulated | GO:BP | GO:0008285 | negative regulation of cell population proliferation | 1.50498372850162e-07 | 1134 | 99 | 804 | 18123 | ACKR3/AC | 99/1134 | 804/18123 |
| down-regulated | GO:BP | GO:0007219 | Notch signaling pathway | 1.57568824866187e-07 | 1134 | 39 | 193 | 18123 | ASCL1/BCL | 39/1134 | 193/18123 |
| down-regulated | GO:BP | GO:0031589 | cell-substrate adhesion | 1.78782314599775e-07 | 1134 | 58 | 368 | 18123 | ACVRL1/AI | 58/1134 | 368/18123 |
| down-regulated | GO:BP | GO:0001655 | urogenital system development | 1.8176906175326e-07 | 1134 | 55 | 339 | 18123 | ANGPT2/A | 55/1134 | 339/18123 |
| down-regulated | GO:BP | GO:0048754 | branching morphogenesis of an epithelial tube | 2.1847499883781e-07 | 1134 | 34 | 154 | 18123 | AREG/BCL | 34/1134 | 154/18123 |
| down-regulated | GO:BP | GO:0002684 | positive regulation of immune system process | 2.23867659592576e-07 | 1134 | 122 | 1079 | 18123 | ANXA1/AP | 122/1134 | 1079/18123 |
| down-regulated | GO:BP | GO:0045944 | positive regulation of transcription by RNA polymerase II | 2.26998603427756e-07 | 1134 | 132 | 1200 | 18123 | ACVRL1/AI | 132/1134 | 1200/18123 |
| down-regulated | GO:BP | GO:0070482 | response to oxygen levels | 2.28210412069053e-07 | 1134 | 62 | 410 | 18123 | ACAA2/AC | 62/1134 | 410/18123 |
| down-regulated | GO:BP | GO:0002521 | leukocyte differentiation | 2.28964464601825e-07 | 1134 | 77 | 566 | 18123 | ANXA1/AN | 77/1134 | 566/18123 |
| down-regulated | GO:BP | GO:0002009 | morphogenesis of an epithelium | 2.28964464601825e-07 | 1134 | 77 | 566 | 18123 | ACVRL1/AI | 77/1134 | 566/18123 |
| down-regulated | GO:BP | GO:0050790 | regulation of catalytic activity | 2.39940900466141e-07 | 1134 | 229 | 2452 | 18123 | A2ML1/AC | 229/1134 | 2452/18123 |
| down-regulated | GO:BP | GO:0045765 | regulation of angiogenesis | 2.43491920197797e-07 | 1134 | 57 | 361 | 18123 | ACVRL1/AI | 57/1134 | 361/18123 |
| down-regulated | GO:BP | GO:0006935 | chemotaxis | 2.49999470633525e-07 | 1134 | 85 | 654 | 18123 | ACKR3/AM | 85/1134 | 654/18123 |
| down-regulated | GO:BP | GO:0001934 | positive regulation of protein phosphorylation | 2.5592346688939e-07 | 1134 | 102 | 846 | 18123 | ACVRL1/AI | 102/1134 | 846/18123 |
| down-regulated | GO:BP | GO:0048468 | cell development | 2.7943905887131e-07 | 1134 | 204 | 2122 | 18123 | ADAMTSL4 | 204/1134 | 2122/18123 |
| down-regulated | GO:BP | GO:0060548 | negative regulation of cell death | 2.82306776669301e-07 | 1134 | 119 | 1047 | 18123 | ACAA2/AC | 119/1134 | 1047/18123 |
| down-regulated | GO:BP | GO:0030036 | actin cytoskeleton organization | 3.0212783053883e-07 | 1134 | 90 | 712 | 18123 | ADD3/AIF1 | 90/1134 | 712/18123 |
| down-regulated | GO:BP | GO:0060485 | mesenchyme development | 3.13682189766127e-07 | 1134 | 50 | 296 | 18123 | ACVRL1/AI | 50/1134 | 296/18123 |
| down-regulated | GO:BP | GO:0042330 | taxis | 3.16035155883683e-07 | 1134 | 85 | 657 | 18123 | ACKR3/AM | 85/1134 | 657/18123 |
| down-regulated | GO:BP | GO:0030856 | regulation of epithelial cell differentiation | 4.4676441648131e-07 | 1134 | 35 | 166 | 18123 | ACVRL1/AI | 35/1134 | 166/18123 |
| down-regulated | GO:BP | GO:1901701 | cellular response to oxygen-containing compound | 4.55354836685337e-07 | 1134 | 134 | 1237 | 18123 | ADCY4/AG | 134/1134 | 1237/18123 |
| down-regulated | GO:BP | GO:0010648 | negative regulation of cell communication | 4.6357413395495e-07 | 1134 | 155 | 1499 | 18123 | ACAA2/AC | 155/1134 | 1499/18123 |
| down-regulated | GO:BP | GO:0009607 | response to biotic stimulus | 4.67333895490179e-07 | 1134 | 166 | 1639 | 18123 | ABCC9/ADI | 166/1134 | 1639/18123 |
| down-regulated | GO:BP | GO:1901342 | regulation of vasculature development | 4.68758055831149e-07 | 1134 | 57 | 367 | 18123 | ACVRL1/AI | 57/1134 | 367/18123 |
| down-regulated | GO:BP | GO:0071363 | cellular response to growth factor stimulus | 4.85818614427315e-07 | 1134 | 92 | 741 | 18123 | ACVRL1/AI | 92/1134 | 741/18123 |
| down-regulated | GO:BP | GO:0072132 | mesenchyme morphogenesis | 5.01592653732772e-07 | 1134 | 19 | 53 | 18123 | ACVRL1/AI | 19/1134 | 53/18123 |
| down-regulated | GO:BP | GO:0003170 | heart valve development | 5.26863093826227e-07 | 1134 | 21 | 65 | 18123 | ADAMTS9/ | 21/1134 | 65/18123 |
| down-regulated | GO:BP | GO:0048871 | multicellular organismal homeostasis | 5.29176524234887e-07 | 1134 | 73 | 533 | 18123 | ACACB/AD | 73/1134 | 533/18123 |
| down-regulated | GO:BP | GO:0023057 | negative regulation of signaling | 5.37314731071208e-07 | 1134 | 155 | 1502 | 18123 | ACAA2/AC | 155/1134 | 1502/18123 |
| down-regulated | GO:BP | GO:0061138 | morphogenesis of a branching epithelium | 5.66222754373921e-07 | 1134 | 37 | 184 | 18123 | ADM/AREG | 37/1134 | 184/18123 |
| down-regulated | GO:BP | GO:0006468 | protein phosphorylation | 6.89990544866582e-07 | 1134 | 168 | 1673 | 18123 | ACVRL1/AI | 168/1134 | 1673/18123 |
| down-regulated | GO:BP | GO:0071407 | cellular response to organic cyclic compound | 7.51223537799424e-07 | 1134 | 81 | 624 | 18123 | AKAP9/AK | 81/1134 | 624/18123 |
| down-regulated | GO:BP | GO:0001822 | kidney development | 7.7875537609016e-07 | 1134 | 49 | 294 | 18123 | ANGPT2/B | 49/1134 | 294/18123 |
| down-regulated | GO:BP | GO:0051707 | response to other organism | 8.27761725800919e-07 | 1134 | 162 | 1600 | 18123 | ABCC9/ADI | 162/1134 | 1600/18123 |
| down-regulated | GO:BP | GO:0043207 | response to external biotic stimulus | 8.67551884535855e-07 | 1134 | 162 | 1601 | 18123 | ABCC9/ADI | 162/1134 | 1601/18123 |
| down-regulated | GO:BP | GO:0003018 | vascular process in circulatory system | 9.22339828318818e-07 | 1134 | 45 | 258 | 18123 | ABCB1/AB | 45/1134 | 258/18123 |
| down-regulated | GO:BP | GO:0001816 | cytokine production | 9.43165518304766e-07 | 1134 | 101 | 854 | 18123 | ADGRG1/A | 101/1134 | 854/18123 |
| down-regulated | GO:BP | GO:0021782 | glial cell development | 1.02878127546781e-06 | 1134 | 29 | 123 | 18123 | ADORA2A/ | 29/1134 | 123/18123 |
| down-regulated | GO:BP | GO:0048568 | embryonic organ development | 1.08971604845378e-06 | 1134 | 63 | 436 | 18123 | ADM/APLN | 63/1134 | 436/18123 |
| down-regulated | GO:BP | GO:0072073 | kidney epithelium development | 1.13831303730482e-06 | 1134 | 31 | 139 | 18123 | BCL2/BMP | 31/1134 | 139/18123 |
| down-regulated | GO:BP | GO:0001817 | regulation of cytokine production | 1.27645080144144e-06 | 1134 | 100 | 847 | 18123 | ANXA1/AP | 100/1134 | 847/18123 |
| down-regulated | GO:BP | GO:0009968 | negative regulation of signal transduction | 1.31236520608469e-06 | 1134 | 145 | 1394 | 18123 | ACAA2/AC | 145/1134 | 1394/18123 |
| down-regulated | GO:BP | GO:0060333 | interferon-gamma-mediated signaling pathway | 1.39899989690855e-06 | 1134 | 24 | 88 | 18123 | CD44/GBP | 24/1134 | 88/18123 |
| down-regulated | GO:BP | GO:0071384 | cellular response to corticosteroid stimulus | 1.47073673143117e-06 | 1134 | 20 | 62 | 18123 | AKR1C3/AI | 20/1134 | 62/18123 |
| down-regulated | GO:BP | GO:0032496 | response to lipopolysaccharide | 1.6182347390093e-06 | 1134 | 53 | 339 | 18123 | ADM/ALPL | 53/1134 | 339/18123 |
| down-regulated | GO:BP | GO:0003197 | endocardial cushion development | 1.76709556774136e-06 | 1134 | 17 | 45 | 18123 | ACVRL1/AI | 17/1134 | 45/18123 |
| down-regulated | GO:BP | GO:0008015 | blood circulation | 1.87043512201685e-06 | 1134 | 73 | 548 | 18123 | ABCC9/AC | 73/1134 | 548/18123 |
| down-regulated | GO:BP | GO:0030855 | epithelial cell differentiation | 1.95126022753281e-06 | 1134 | 96 | 807 | 18123 | ACVRL1/AI | 96/1134 | 807/18123 |
| down-regulated | GO:BP | GO:0045321 | leukocyte activation | 2.13679413940041e-06 | 1134 | 140 | 1341 | 18123 | ADA2/ADG | 140/1134 | 1341/18123 |
| down-regulated | GO:BP | GO:0036293 | response to decreased oxygen levels | 2.23084641362317e-06 | 1134 | 57 | 382 | 18123 | ACAA2/AC | 57/1134 | 382/18123 |
| down-regulated | GO:BP | GO:0002237 | response to molecule of bacterial origin | 2.26080754339288e-06 | 1134 | 55 | 362 | 18123 | ADM/ALPL | 55/1134 | 362/18123 |
| down-regulated | GO:BP | GO:0048708 | astrocyte differentiation | 2.26958496813971e-06 | 1134 | 23 | 83 | 18123 | ADORA2A/ | 23/1134 | 83/18123 |
| down-regulated | GO:BP | GO:0007399 | nervous system development | 2.2709518807447e-06 | 1134 | 227 | 2484 | 18123 | ADGRG1/A | 227/1134 | 2484/18123 |
| down-regulated | GO:BP | GO:0072009 | nephron epithelium development | 2.43692546870446e-06 | 1134 | 27 | 112 | 18123 | BCL2/CD34 | 27/1134 | 112/18123 |
| down-regulated | GO:BP | GO:1905314 | semi-lunar valve development | 3.21417720416995e-06 | 1134 | 16 | 41 | 18123 | DLL4/EMIL | 16/1134 | 41/18123 |
| down-regulated | GO:BP | GO:0016310 | phosphorylation | 3.22844474795503e-06 | 1134 | 190 | 1996 | 18123 | ACVRL1/AI | 190/1134 | 1996/18123 |
| down-regulated | GO:BP | GO:0003203 | endocardial cushion morphogenesis | 3.58177273581094e-06 | 1134 | 15 | 36 | 18123 | ACVRL1/AI | 15/1134 | 36/18123 |
| down-regulated | GO:BP | GO:0007159 | leukocyte cell-cell adhesion | 3.72829303647163e-06 | 1134 | 56 | 377 | 18123 | ADORA2A/ | 56/1134 | 377/18123 |
| down-regulated | GO:BP | GO:0045893 | positive regulation of transcription, DNA-templated | 4.27279643718005e-06 | 1134 | 161 | 1623 | 18123 | ACVRL1/AI | 161/1134 | 1623/18123 |
| down-regulated | GO:BP | GO:1903508 | positive regulation of nucleic acid-templated transcription | 4.27279643718005e-06 | 1134 | 161 | 1623 | 18123 | ACVRL1/AI | 161/1134 | 1623/18123 |
| down-regulated | GO:BP | GO:1902680 | positive regulation of RNA biosynthetic process | 4.46748474412174e-06 | 1134 | 161 | 1624 | 18123 | ACVRL1/AI | 161/1134 | 1624/18123 |
| down-regulated | GO:BP | GO:0001763 | morphogenesis of a branching structure | 4.88842142360781e-06 | 1134 | 37 | 198 | 18123 | ADM/AREG | 37/1134 | 198/18123 |
| down-regulated | GO:BP | GO:0043066 | negative regulation of apoptotic process | 5.02842794701098e-06 | 1134 | 105 | 928 | 18123 | ACAA2/AC | 105/1134 | 928/18123 |
| down-regulated | GO:BP | GO:0043069 | negative regulation of programmed cell death | 5.05515234791136e-06 | 1134 | 107 | 952 | 18123 | ACAA2/AC | 107/1134 | 952/18123 |
| down-regulated | GO:BP | GO:1903053 | regulation of extracellular matrix organization | 5.68367391465632e-06 | 1134 | 17 | 48 | 18123 | AEBP1/AN | 17/1134 | 48/18123 |
| down-regulated | GO:BP | GO:0001501 | skeletal system development | 6.47413951036829e-06 | 1134 | 68 | 509 | 18123 | ACVRL1/AI | 68/1134 | 509/18123 |
| down-regulated | GO:BP | GO:0060562 | epithelial tube morphogenesis | 7.85620787905634e-06 | 1134 | 51 | 334 | 18123 | ACVRL1/AI | 51/1134 | 334/18123 |

| | | | | | | | | | | | |
|---|---|---|---|---|---|---|---|---|---|---|---|
| down-regulated | GO:BP | GO:0003179 | heart valve morphogenesis | 8.26918322448338e-06 | 1134 | 18 | 55 | 18123 | ADAMTS9/ | 18/1134 | 55/18123 |
| down-regulated | GO:BP | GO:0045446 | endothelial cell differentiation | 8.66074222381237e-06 | 1134 | 28 | 126 | 18123 | ACVRL1/AF | 28/1134 | 126/18123 |
| down-regulated | GO:BP | GO:0010942 | positive regulation of cell death | 8.66184112144859e-06 | 1134 | 84 | 691 | 18123 | ACOX2/AD | 84/1134 | 691/18123 |
| down-regulated | GO:BP | GO:0043085 | positive regulation of catalytic activity | 8.95239898127886e-06 | 1134 | 141 | 1383 | 18123 | ADCY4/AD( | 141/1134 | 1383/18123 |
| down-regulated | GO:BP | GO:0006954 | inflammatory response | 9.02914834255205e-06 | 1134 | 107 | 962 | 18123 | ADCY4/AD( | 107/1134 | 962/18123 |
| down-regulated | GO:BP | GO:0001503 | ossification | 9.7484273028934e-06 | 1134 | 59 | 418 | 18123 | ADGRV1/A | 59/1134 | 418/18123 |
| down-regulated | GO:BP | GO:0031960 | response to corticosteroid | 9.9908302768445e-06 | 1134 | 33 | 168 | 18123 | ADM/AKR1 | 33/1134 | 168/18123 |
| down-regulated | GO:BP | GO:0022407 | regulation of cell-cell adhesion | 1.01045984766013e-05 | 1134 | 62 | 450 | 18123 | ADORA2A/ | 62/1134 | 450/18123 |
| down-regulated | GO:BP | GO:0001666 | response to hypoxia | 1.03625449367126e-05 | 1134 | 54 | 367 | 18123 | ACAA2/AC' | 54/1134 | 367/18123 |
| down-regulated | GO:BP | GO:0007369 | gastrulation | 1.28204134859545e-05 | 1134 | 35 | 187 | 18123 | AMOT/APL | 35/1134 | 187/18123 |
| down-regulated | GO:BP | GO:0071385 | cellular response to glucocorticoid stimulus | 1.58014415713322e-05 | 1134 | 18 | 57 | 18123 | ANXA1/AQ | 18/1134 | 57/18123 |
| down-regulated | GO:BP | GO:0042592 | homeostatic process | 1.71860229249924e-05 | 1134 | 186 | 1985 | 18123 | ABHD4/AC | 186/1134 | 1985/18123 |
| down-regulated | GO:BP | GO:0007389 | pattern specification process | 1.79380580246975e-05 | 1134 | 61 | 446 | 18123 | ACVRL1/AI | 61/1134 | 446/18123 |
| down-regulated | GO:BP | GO:0042476 | odontogenesis | 1.81681374521249e-05 | 1134 | 28 | 130 | 18123 | ADM/ALPL | 28/1134 | 130/18123 |
| down-regulated | GO:BP | GO:0050900 | leukocyte migration | 1.83416126109681e-05 | 1134 | 67 | 511 | 18123 | ANGPT2/A | 67/1134 | 511/18123 |
| down-regulated | GO:BP | GO:0051254 | positive regulation of RNA metabolic process | 1.87037006577889e-05 | 1134 | 170 | 1775 | 18123 | ACVRL1/AI | 170/1134 | 1775/18123 |
| down-regulated | GO:BP | GO:0071496 | cellular response to external stimulus | 1.93883719842675e-05 | 1134 | 52 | 353 | 18123 | AKR1C3/A( | 52/1134 | 353/18123 |
| down-regulated | GO:BP | GO:0071383 | cellular response to steroid hormone stimulus | 1.96730500998358e-05 | 1134 | 41 | 245 | 18123 | AKR1C3/AI | 41/1134 | 245/18123 |
| down-regulated | GO:BP | GO:0045766 | positive regulation of angiogenesis | 1.97489318684676e-05 | 1134 | 35 | 190 | 18123 | ACVRL1/AI | 35/1134 | 190/18123 |
| down-regulated | GO:BP | GO:1904018 | positive regulation of vasculature development | 1.97489318684676e-05 | 1134 | 35 | 190 | 18123 | ACVRL1/AI | 35/1134 | 190/18123 |
| down-regulated | GO:BP | GO:0007517 | muscle organ development | 1.99000045843841e-05 | 1134 | 50 | 333 | 18123 | BCL2/BTG2 | 50/1134 | 333/18123 |
| down-regulated | GO:BP | GO:0031399 | regulation of protein modification process | 2.0215126396542e-05 | 1134 | 163 | 1685 | 18123 | ACVRL1/AI | 163/1134 | 1685/18123 |
| down-regulated | GO:BP | GO:0003007 | heart morphogenesis | 2.0947221151873e-05 | 1134 | 42 | 255 | 18123 | ACVRL1/AF | 42/1134 | 255/18123 |
| down-regulated | GO:BP | GO:0061005 | cell differentiation involved in kidney development | 2.15814705037801e-05 | 1134 | 18 | 58 | 18123 | CD34/FOX( | 18/1134 | 58/18123 |
| down-regulated | GO:BP | GO:0072012 | glomerulus vasculature development | 2.29476799096382e-05 | 1134 | 12 | 25 | 18123 | ANGPT2/BI | 12/1134 | 25/18123 |
| down-regulated | GO:BP | GO:0031323 | regulation of cellular metabolic process | 2.40273923153867e-05 | 1134 | 484 | 6309 | 18123 | A2ML1/AC | 484/1134 | 6309/18123 |
| down-regulated | GO:BP | GO:0035909 | aorta morphogenesis | 2.42535984991446e-05 | 1134 | 14 | 35 | 18123 | ACVRL1/AI | 14/1134 | 35/18123 |
| down-regulated | GO:BP | GO:0001654 | eye development | 2.4571343117779e-05 | 1134 | 54 | 376 | 18123 | ACVRL1/A( | 54/1134 | 376/18123 |
| down-regulated | GO:BP | GO:0051246 | regulation of protein metabolic process | 2.46259548195e-05 | 1134 | 242 | 2759 | 18123 | A2ML1/AC | 242/1134 | 2759/18123 |
| down-regulated | GO:BP | GO:0010243 | response to organonitrogen compound | 2.70133882937587e-05 | 1134 | 116 | 1092 | 18123 | ABCC9/AD( | 116/1134 | 1092/18123 |
| down-regulated | GO:BP | GO:0043010 | camera-type eye development | 3.32730823212073e-05 | 1134 | 49 | 328 | 18123 | ACVRL1/A( | 49/1134 | 328/18123 |
| down-regulated | GO:BP | GO:0043009 | chordate embryonic development | 3.4095998350107e-05 | 1134 | 77 | 631 | 18123 | ACVRL1/AI | 77/1134 | 631/18123 |
| down-regulated | GO:BP | GO:0150063 | visual system development | 3.56454743093489e-05 | 1134 | 54 | 380 | 18123 | ACVRL1/A( | 54/1134 | 380/18123 |
| down-regulated | GO:BP | GO:0007264 | small GTPase mediated signal transduction | 3.70779232456136e-05 | 1134 | 67 | 520 | 18123 | ADCYAP1R | 67/1134 | 520/18123 |
| down-regulated | GO:BP | GO:0071346 | cellular response to interferon-gamma | 3.88794760383815e-05 | 1134 | 33 | 177 | 18123 | CCL2/CD44 | 33/1134 | 177/18123 |
| down-regulated | GO:BP | GO:0043065 | positive regulation of apoptotic process | 4.1255439318902e-05 | 1134 | 75 | 611 | 18123 | ADAMTSL4 | 75/1134 | 611/18123 |
| down-regulated | GO:BP | GO:0060840 | artery development | 4.28236380238923e-05 | 1134 | 24 | 103 | 18123 | ACVRL1/AI | 24/1134 | 103/18123 |
| down-regulated | GO:BP | GO:1901698 | response to nitrogen compound | 4.76007994546702e-05 | 1134 | 122 | 1178 | 18123 | ABCC9/AD( | 122/1134 | 1178/18123 |
| down-regulated | GO:BP | GO:0051247 | positive regulation of protein metabolic process | 5.21405431799595e-05 | 1134 | 153 | 1577 | 18123 | ACVRL1/AI | 153/1134 | 1577/18123 |
| down-regulated | GO:BP | GO:0031328 | positive regulation of cellular biosynthetic process | 5.62117756633466e-05 | 1134 | 183 | 1976 | 18123 | ACVRL1/AI | 183/1134 | 1976/18123 |
| down-regulated | GO:BP | GO:0003176 | aortic valve development | 5.6723515200106e-05 | 1134 | 14 | 37 | 18123 | DLL4/EMIL | 14/1134 | 37/18123 |
| down-regulated | GO:BP | GO:0043068 | positive regulation of programmed cell death | 6.07422091725696e-05 | 1134 | 76 | 628 | 18123 | ADAMTSL4 | 76/1134 | 628/18123 |
| down-regulated | GO:BP | GO:0009792 | embryo development ending in birth or egg hatching | 6.09529962431997e-05 | 1134 | 78 | 651 | 18123 | ACVRL1/AI | 78/1134 | 651/18123 |
| down-regulated | GO:BP | GO:0097435 | supramolecular fiber organization | 6.10193617972976e-05 | 1134 | 96 | 863 | 18123 | ADD3/AEB | 96/1134 | 863/18123 |
| down-regulated | GO:BP | GO:0048880 | sensory system development | 6.14783834353676e-05 | 1134 | 54 | 386 | 18123 | ACVRL1/A( | 54/1134 | 386/18123 |
| down-regulated | GO:BP | GO:0031401 | positive regulation of protein modification process | 6.3784470928862e-05 | 1134 | 114 | 1084 | 18123 | ACVRL1/AI | 114/1134 | 1084/18123 |
| down-regulated | GO:BP | GO:0061440 | kidney vasculature development | 6.82247634377841e-05 | 1134 | 12 | 27 | 18123 | ANGPT2/BI | 12/1134 | 27/18123 |
| down-regulated | GO:BP | GO:0061437 | renal system vasculature development | 6.82247634377841e-05 | 1134 | 12 | 27 | 18123 | ANGPT2/BI | 12/1134 | 27/18123 |
| down-regulated | GO:BP | GO:0003272 | endocardial cushion formation | 6.82247634377841e-05 | 1134 | 12 | 27 | 18123 | APLNR/BM | 12/1134 | 27/18123 |
| down-regulated | GO:BP | GO:0042110 | T cell activation | 6.89819544510677e-05 | 1134 | 64 | 495 | 18123 | ADORA2A/ | 64/1134 | 495/18123 |
| down-regulated | GO:BP | GO:1901652 | response to peptide | 7.13928827439383e-05 | 1134 | 69 | 551 | 18123 | ADCY4/AD | 69/1134 | 551/18123 |
| down-regulated | GO:BP | GO:0060537 | muscle tissue development | 7.24035127712505e-05 | 1134 | 56 | 409 | 18123 | ADAMTS9/ | 56/1134 | 409/18123 |
| down-regulated | GO:BP | GO:0009891 | positive regulation of biosynthetic process | 8.17092611786862e-05 | 1134 | 185 | 2013 | 18123 | ACVRL1/AI | 185/1134 | 2013/18123 |
| down-regulated | GO:BP | GO:0048732 | gland development | 8.44188758989186e-05 | 1134 | 59 | 443 | 18123 | AK4/ANXA | 59/1134 | 443/18123 |
| down-regulated | GO:BP | GO:0003206 | cardiac chamber morphogenesis | 9.41294932833553e-05 | 1134 | 26 | 123 | 18123 | APLNR/BM | 26/1134 | 123/18123 |
| down-regulated | GO:BP | GO:0022008 | neurogenesis | 9.988234012714e-05 | 1134 | 161 | 1698 | 18123 | ADGRG1/A | 161/1134 | 1698/18123 |
| down-regulated | GO:BP | GO:1903706 | regulation of hemopoiesis | 0.000104275217821192 | 1134 | 57 | 424 | 18123 | ANXA1/AXI | 57/1134 | 424/18123 |
| down-regulated | GO:BP | GO:0043583 | ear development | 0.00010473808855655 | 1134 | 37 | 221 | 18123 | ADGRV1/A | 37/1134 | 221/18123 |
| down-regulated | GO:BP | GO:0051384 | response to glucocorticoid | 0.000108444465123052 | 1134 | 29 | 149 | 18123 | ADM/ALPL | 29/1134 | 149/18123 |
| down-regulated | GO:BP | GO:0051090 | regulation of DNA-binding transcription factor activity | 0.000108820643146808 | 1134 | 60 | 457 | 18123 | ARHGEF5/I | 60/1134 | 457/18123 |
| down-regulated | GO:BP | GO:0045747 | positive regulation of Notch signaling pathway | 0.000109285777017708 | 1134 | 17 | 57 | 18123 | ASCL1/DLL | 17/1134 | 57/18123 |
| down-regulated | GO:BP | GO:0001704 | formation of primary germ layer | 0.000112118183088461 | 1134 | 26 | 124 | 18123 | ATOH8/BM | 26/1134 | 124/18123 |
| down-regulated | GO:BP | GO:1902105 | regulation of leukocyte differentiation | 0.000116136175214336 | 1134 | 44 | 290 | 18123 | ANXA1/AX | 44/1134 | 290/18123 |
| down-regulated | GO:BP | GO:0045935 | positive regulation of nucleobase-containing compound metabolic pr | 0.000121144647691157 | 1134 | 179 | 1943 | 18123 | ACVRL1/AI | 179/1134 | 1943/18123 |
| down-regulated | GO:BP | GO:0050673 | epithelial cell proliferation | 0.000137992838682466 | 1134 | 60 | 460 | 18123 | ACVRL1/AF | 60/1134 | 460/18123 |
| down-regulated | GO:BP | GO:0043408 | regulation of MAPK cascade | 0.00013943798704377 | 1134 | 84 | 734 | 18123 | ACKR3/AD( | 84/1134 | 734/18123 |
| down-regulated | GO:BP | GO:0008593 | regulation of Notch signaling pathway | 0.00016626303131484 | 1134 | 24 | 110 | 18123 | ASCL1/BCL | 24/1134 | 110/18123 |
| down-regulated | GO:BP | GO:0001974 | blood vessel remodeling | 0.000181846175401421 | 1134 | 15 | 46 | 18123 | ACVRL1/A) | 15/1134 | 46/18123 |
| down-regulated | GO:BP | GO:0035904 | aorta development | 0.000192764456401559 | 1134 | 17 | 59 | 18123 | ACVRL1/AI | 17/1134 | 59/18123 |
| down-regulated | GO:BP | GO:0060284 | regulation of cell development | 0.000198048991570907 | 1134 | 64 | 509 | 18123 | APPL2/ASC | 64/1134 | 509/18123 |
| down-regulated | GO:BP | GO:0001701 | in utero embryonic development | 0.000212949723908524 | 1134 | 52 | 379 | 18123 | ACVRL1/AI | 52/1134 | 379/18123 |
| down-regulated | GO:BP | GO:0050678 | regulation of epithelial cell proliferation | 0.000225024844877166 | 1134 | 54 | 401 | 18123 | ACVRL1/AF | 54/1134 | 401/18123 |
| down-regulated | GO:BP | GO:0002252 | immune effector process | 0.000226643170175065 | 1134 | 116 | 1135 | 18123 | ADA2/ALDI | 116/1134 | 1135/18123 |
| down-regulated | GO:BP | GO:0090130 | tissue migration | 0.000232129114912263 | 1134 | 52 | 380 | 18123 | ACVRL1/AI | 52/1134 | 380/18123 |
| down-regulated | GO:BP | GO:0006793 | phosphorus metabolic process | 0.000232698175412608 | 1134 | 255 | 3014 | 18123 | ABHD4/AC | 255/1134 | 3014/18123 |
| down-regulated | GO:BP | GO:0080090 | regulation of primary metabolic process | 0.000233777961557149 | 1134 | 462 | 6071 | 18123 | A2ML1/AC | 462/1134 | 6071/18123 |
| down-regulated | GO:BP | GO:0006796 | phosphate-containing compound metabolic process | 0.000241842544277278 | 1134 | 253 | 2987 | 18123 | ABHD4/AC | 253/1134 | 2987/18123 |
| down-regulated | GO:BP | GO:0032101 | regulation of response to external stimulus | 0.000242190555138568 | 1134 | 124 | 1238 | 18123 | ADCY4/AD( | 124/1134 | 1238/18123 |
| down-regulated | GO:BP | GO:0007162 | negative regulation of cell adhesion | 0.000255530789315451 | 1134 | 45 | 308 | 18123 | ACVRL1/AI | 45/1134 | 308/18123 |
| down-regulated | GO:BP | GO:0010631 | epithelial cell migration | 0.000266944425168969 | 1134 | 51 | 371 | 18123 | ACVRL1/AI | 51/1134 | 371/18123 |
| down-regulated | GO:BP | GO:0061572 | actin filament bundle organization | 0.000272505679849201 | 1134 | 30 | 164 | 18123 | AIF1L/AM( | 30/1134 | 164/18123 |
| down-regulated | GO:BP | GO:0032270 | positive regulation of cellular protein metabolic process | 0.000291113256620948 | 1134 | 143 | 1488 | 18123 | ACVRL1/AI | 143/1134 | 1488/18123 |
| down-regulated | GO:BP | GO:0071559 | response to transforming growth factor beta | 0.000294334139779422 | 1134 | 41 | 269 | 18123 | ACVRL1/AF | 41/1134 | 269/18123 |
| down-regulated | GO:BP | GO:0007169 | transmembrane receptor protein tyrosine kinase signaling pathway | 0.000303250801504102 | 1134 | 85 | 759 | 18123 | ANGPT2/A | 85/1134 | 759/18123 |
| down-regulated | GO:BP | GO:0060326 | cell chemotaxis | 0.000309766466928104 | 1134 | 45 | 310 | 18123 | ACKR3/AN | 45/1134 | 310/18123 |
| down-regulated | GO:BP | GO:0003151 | outflow tract morphogenesis | 0.000323289873909208 | 1134 | 19 | 75 | 18123 | BMP7/CLD | 19/1134 | 75/18123 |
| down-regulated | GO:BP | GO:0009892 | negative regulation of metabolic process | 0.000330498062133157 | 1134 | 288 | 3498 | 18123 | A2ML1/AC | 288/1134 | 3498/18123 |
| down-regulated | GO:BP | GO:0090100 | positive regulation of transmembrane receptor protein serine/threor | 0.000341159045337467 | 1134 | 24 | 114 | 18123 | ACVRL1/BI | 24/1134 | 114/18123 |
| down-regulated | GO:BP | GO:0035850 | epithelial cell differentiation involved in kidney development | 0.000345068748474057 | 1134 | 15 | 48 | 18123 | CD34/FOX( | 15/1134 | 48/18123 |
| down-regulated | GO:BP | GO:0090132 | epithelium migration | 0.000345632617810934 | 1134 | 51 | 374 | 18123 | ACVRL1/AI | 51/1134 | 374/18123 |
| down-regulated | GO:BP | GO:0007178 | transmembrane receptor protein serine/threonine kinase signaling p | 0.000345632617810934 | 1134 | 51 | 374 | 18123 | ACVRL1/AI | 51/1134 | 374/18123 |
| down-regulated | GO:BP | GO:0001818 | negative regulation of cytokine production | 0.000355061626932075 | 1134 | 52 | 385 | 18123 | ANXA1/AP | 52/1134 | 385/18123 |
| down-regulated | GO:BP | GO:0097190 | apoptotic signaling pathway | 0.000366117213139605 | 1134 | 72 | 609 | 18123 | ACAA2/AC | 72/1134 | 609/18123 |
| down-regulated | GO:BP | GO:0098542 | defense response to other organism | 0.000371628149516316 | 1134 | 122 | 1222 | 18123 | ABCC9/AN | 122/1134 | 1222/18123 |
| down-regulated | GO:BP | GO:0000902 | cell morphogenesis | 0.000375976647566493 | 1134 | 108 | 1045 | 18123 | ADORA2A/ | 108/1134 | 1045/18123 |
| down-regulated | GO:BP | GO:1903131 | mononuclear cell differentiation | 0.000377541519554579 | 1134 | 57 | 440 | 18123 | ANXA1/AX | 57/1134 | 440/18123 |
| down-regulated | GO:BP | GO:0048010 | vascular endothelial growth factor receptor signaling pathway | 0.00041623577538497 | 1134 | 22 | 99 | 18123 | AXL/DOCK | 22/1134 | 99/18123 |
| down-regulated | GO:BP | GO:0040007 | growth | 0.000432640455195916 | 1134 | 102 | 973 | 18123 | ACACB/AC' | 102/1134 | 973/18123 |
| down-regulated | GO:BP | GO:0003177 | pulmonary valve development | 0.000459203818044308 | 1134 | 10 | 21 | 18123 | HEYL/JAG1 | 10/1134 | 21/18123 |
| down-regulated | GO:BP | GO:0010557 | positive regulation of macromolecule biosynthetic process | 0.000465404015001803 | 1134 | 170 | 1859 | 18123 | ACVRL1/AI | 170/1134 | 1859/18123 |
| down-regulated | GO:BP | GO:0002683 | negative regulation of immune system process | 0.000504858292125113 | 1134 | 72 | 614 | 18123 | ADCY4/AD( | 72/1134 | 614/18123 |
| down-regulated | GO:BP | GO:0048844 | artery morphogenesis | 0.000507721896882481 | 1134 | 19 | 77 | 18123 | ACVRL1/AI | 19/1134 | 77/18123 |
| down-regulated | GO:BP | GO:0007417 | central nervous system development | 0.00052609380617923 | 1134 | 108 | 1052 | 18123 | ADGRG1/A | 108/1134 | 1052/18123 |
| down-regulated | GO:BP | GO:0032268 | regulation of cellular protein metabolic process | 0.000526564242017464 | 1134 | 223 | 2591 | 18123 | A2ML1/AC | 223/1134 | 2591/18123 |
| down-regulated | GO:BP | GO:0060411 | cardiac septum morphogenesis | 0.00054308192072449 | 1134 | 18 | 70 | 18123 | APLNR/BM | 18/1134 | 70/18123 |
| down-regulated | GO:BP | GO:0001885 | endothelial cell development | 0.00054308192072449 | 1134 | 18 | 70 | 18123 | ARHGEF26 | 18/1134 | 70/18123 |
| down-regulated | GO:BP | GO:0050865 | regulation of cell activation | 0.000543167185763496 | 1134 | 76 | 662 | 18123 | ADGRF5/A | 76/1134 | 662/18123 |
| down-regulated | GO:BP | GO:0007160 | cell-matrix adhesion | 0.000592530449471456 | 1134 | 37 | 236 | 18123 | ACVRL1/AI | 37/1134 | 236/18123 |
| down-regulated | GO:BP | GO:0046718 | viral entry into host cell | 0.000605364829366893 | 1134 | 27 | 143 | 18123 | ANPEP/AXI | 27/1134 | 143/18123 |
| down-regulated | GO:BP | GO:0010605 | negative regulation of macromolecule metabolic process | 0.00061255508184771 | 1134 | 270 | 3262 | 18123 | A2ML1/AC | 270/1134 | 3262/18123 |
| down-regulated | GO:BP | GO:0022408 | negative regulation of cell-cell adhesion | 0.000629236300681117 | 1134 | 33 | 198 | 18123 | ADORA2A/ | 33/1134 | 198/18123 |
| down-regulated | GO:BP | GO:0003180 | aortic valve morphogenesis | 0.000661276173651965 | 1134 | 12 | 32 | 18123 | DLL4/EMIL | 12/1134 | 32/18123 |
| down-regulated | GO:BP | GO:0032970 | regulation of actin filament-based process | 0.00069375091961775 | 1134 | 53 | 404 | 18123 | ADD3/AKA | 53/1134 | 404/18123 |

| | | | | | | | | | | | |
|---|---|---|---|---|---|---|---|---|---|---|---|
| down-regulated | GO:BP | GO:0014002 | astrocyte development | 0.000706161331825323 | 1134 | 14 | 44 | 18123 | ADORA2A/ | 14/1134 | 44/18123 |
| down-regulated | GO:BP | GO:0031324 | negative regulation of cellular metabolic process | 0.000738712410937126 | 1134 | 236 | 2785 | 18123 | A2ML1/AC | 236/1134 | 2785/18123 |
| down-regulated | GO:BP | GO:0019222 | regulation of metabolic process | 0.000742729020512687 | 1134 | 530 | 7175 | 18123 | A2ML1/AC | 530/1134 | 7175/18123 |
| down-regulated | GO:BP | GO:1903037 | regulation of leukocyte cell-cell adhesion | 0.000782605143186241 | 1134 | 47 | 341 | 18123 | ADORA2A/ | 47/1134 | 341/18123 |
| down-regulated | GO:BP | GO:0048762 | mesenchymal cell differentiation | 0.000819791234995014 | 1134 | 37 | 239 | 18123 | BCL2/BMP | 37/1134 | 239/18123 |
| down-regulated | GO:BP | GO:0032870 | cellular response to hormone stimulus | 0.000835114778215454 | 1134 | 77 | 681 | 18123 | ADCY4/AG | 77/1134 | 681/18123 |
| down-regulated | GO:BP | GO:0061028 | establishment of endothelial barrier | 0.000842619744906265 | 1134 | 15 | 51 | 18123 | CDH5/CLDI | 15/1134 | 51/18123 |
| down-regulated | GO:BP | GO:0001656 | metanephros development | 0.000846398728917233 | 1134 | 20 | 87 | 18123 | BCL2/BMP | 20/1134 | 87/18123 |
| down-regulated | GO:BP | GO:0043410 | positive regulation of MAPK cascade | 0.000901646515790158 | 1134 | 65 | 542 | 18123 | ACKR3/AD | 65/1134 | 542/18123 |
| down-regulated | GO:BP | GO:0002274 | myeloid leukocyte activation | 0.000932801509925701 | 1134 | 76 | 671 | 18123 | ADA2/ADG | 76/1134 | 671/18123 |
| down-regulated | GO:BP | GO:0071260 | cellular response to mechanical stimulus | 0.000968693731175167 | 1134 | 19 | 80 | 18123 | AQP1/ATP | 19/1134 | 80/18123 |
| down-regulated | GO:BP | GO:0033627 | cell adhesion mediated by integrin | 0.00107644847857239 | 1134 | 18 | 73 | 18123 | CD3E/EPH | 18/1134 | 73/18123 |
| down-regulated | GO:BP | GO:0051091 | positive regulation of DNA-binding transcription factor activity | 0.0011033576385173 | 1134 | 40 | 272 | 18123 | ARHGEF5/ | 40/1134 | 272/18123 |
| down-regulated | GO:BP | GO:0010718 | positive regulation of epithelial to mesenchymal transition | 0.00111616008373596 | 1134 | 15 | 52 | 18123 | BMP7/ENG | 15/1134 | 52/18123 |
| down-regulated | GO:BP | GO:0051171 | regulation of nitrogen compound metabolic process | 0.00119528341795279 | 1134 | 445 | 5886 | 18123 | A2ML1/AC | 445/1134 | 5886/18123 |
| down-regulated | GO:BP | GO:0010810 | regulation of cell-substrate adhesion | 0.00122038523925122 | 1134 | 35 | 223 | 18123 | ACVRL1/AI | 35/1134 | 223/18123 |
| down-regulated | GO:BP | GO:0002694 | regulation of leukocyte activation | 0.00126413556552414 | 1134 | 71 | 617 | 18123 | ADGRF5/A | 71/1134 | 617/18123 |
| down-regulated | GO:BP | GO:0014706 | striated muscle tissue development | 0.0012879194107052 | 1134 | 51 | 390 | 18123 | ADAMTS9/ | 51/1134 | 390/18123 |
| down-regulated | GO:BP | GO:0071294 | cellular response to zinc ion | 0.00132701551486436 | 1134 | 10 | 23 | 18123 | GLRA1/HV | 10/1134 | 23/18123 |
| down-regulated | GO:BP | GO:0030099 | myeloid cell differentiation | 0.00148780513633646 | 1134 | 55 | 436 | 18123 | ADGRF5/A | 55/1134 | 436/18123 |
| down-regulated | GO:BP | GO:0097191 | extrinsic apoptotic signaling pathway | 0.00168934117068009 | 1134 | 35 | 226 | 18123 | ACSL5/BA | 35/1134 | 226/18123 |
| down-regulated | GO:BP | GO:0045087 | innate immune response | 0.00183616297787197 | 1134 | 102 | 1003 | 18123 | ANKHD1/A | 102/1134 | 1003/18123 |
| down-regulated | GO:BP | GO:0009612 | response to mechanical stimulus | 0.00184998595914234 | 1134 | 34 | 217 | 18123 | ADGRV1/A | 34/1134 | 217/18123 |
| down-regulated | GO:BP | GO:0051172 | negative regulation of nitrogen compound metabolic process | 0.00189207789540358 | 1134 | 220 | 2592 | 18123 | A2ML1/AC | 220/1134 | 2592/18123 |
| down-regulated | GO:BP | GO:0050863 | regulation of T cell activation | 0.00194453944924369 | 1134 | 46 | 341 | 18123 | ADORA2A/ | 46/1134 | 341/18123 |
| down-regulated | GO:BP | GO:0035633 | maintenance of blood-brain barrier | 0.00205076288014294 | 1134 | 12 | 35 | 18123 | CDH5/CLDI | 12/1134 | 35/18123 |
| down-regulated | GO:BP | GO:0044403 | biological process involved in symbiotic interaction | 0.00206064310079672 | 1134 | 45 | 331 | 18123 | ANPEP/AN | 45/1134 | 331/18123 |
| down-regulated | GO:BP | GO:0051017 | actin filament bundle assembly | 0.00211295065705049 | 1134 | 28 | 161 | 18123 | AIF1L/AM | 28/1134 | 161/18123 |
| down-regulated | GO:BP | GO:0007010 | cytoskeleton organization | 0.00214125011922531 | 1134 | 138 | 1473 | 18123 | ADD3/AIF1 | 138/1134 | 1473/18123 |
| down-regulated | GO:BP | GO:0072010 | glomerular epithelium development | 0.00214734077274942 | 1134 | 10 | 24 | 18123 | CD34/FOX | 10/1134 | 24/18123 |
| down-regulated | GO:BP | GO:0043549 | regulation of kinase activity | 0.00232131198792033 | 1134 | 95 | 920 | 18123 | ADCY4/AD | 95/1134 | 920/18123 |
| down-regulated | GO:BP | GO:0051336 | regulation of hydrolase activity | 0.00232740109329445 | 1134 | 121 | 1252 | 18123 | A2ML1/AD | 121/1134 | 1252/18123 |
| down-regulated | GO:BP | GO:0022614 | membrane to membrane docking | 0.00264251306799086 | 1134 | 5 | 5 | 18123 | EZR/ICAM1 | 5/1134 | 5/18123 |
| down-regulated | GO:BP | GO:0010035 | response to inorganic substance | 0.00280098702638862 | 1134 | 67 | 583 | 18123 | ADGRV1/A | 67/1134 | 583/18123 |
| down-regulated | GO:BP | GO:0034340 | response to type I interferon | 0.00304833429969373 | 1134 | 21 | 102 | 18123 | BST2/GBP | 21/1134 | 102/18123 |
| down-regulated | GO:BP | GO:0051050 | positive regulation of transport | 0.00307031855807222 | 1134 | 97 | 951 | 18123 | ABCB1/AC | 97/1134 | 951/18123 |
| down-regulated | GO:BP | GO:0000165 | MAPK cascade | 0.00327539753446802 | 1134 | 98 | 965 | 18123 | ACKR3/AD | 98/1134 | 965/18123 |
| down-regulated | GO:BP | GO:0071560 | cellular response to transforming growth factor beta stimulus | 0.0033358897923967 | 1134 | 38 | 263 | 18123 | ACVRL1/AI | 38/1134 | 263/18123 |
| down-regulated | GO:BP | GO:0070663 | regulation of leukocyte proliferation | 0.00341684391317529 | 1134 | 37 | 253 | 18123 | ANXA1/BC | 37/1134 | 253/18123 |
| down-regulated | GO:BP | GO:0044409 | entry into host | 0.00365039121756314 | 1134 | 27 | 156 | 18123 | ANPEP/AX | 27/1134 | 156/18123 |
| down-regulated | GO:BP | GO:0030217 | T cell differentiation | 0.00366716910592888 | 1134 | 38 | 264 | 18123 | ANXA1/BC | 38/1134 | 264/18123 |
| down-regulated | GO:BP | GO:0006820 | anion transport | 0.00369306732620621 | 1134 | 62 | 529 | 18123 | ABCB1/AD | 62/1134 | 529/18123 |
| down-regulated | GO:BP | GO:0002718 | regulation of cytokine production involved in immune response | 0.00377744889902858 | 1134 | 20 | 95 | 18123 | BCL6/BST2 | 20/1134 | 95/18123 |
| down-regulated | GO:BP | GO:0002367 | cytokine production involved in immune response | 0.00377744889902858 | 1134 | 20 | 95 | 18123 | BCL6/BST2 | 20/1134 | 95/18123 |
| down-regulated | GO:BP | GO:0032944 | regulation of mononuclear cell proliferation | 0.00388314765860022 | 1134 | 35 | 234 | 18123 | ANXA1/BC | 35/1134 | 234/18123 |
| down-regulated | GO:BP | GO:0003002 | regionalization | 0.00400857623406696 | 1134 | 45 | 339 | 18123 | ACVRL1/A | 45/1134 | 339/18123 |
| down-regulated | GO:BP | GO:0045601 | regulation of endothelial cell differentiation | 0.00404476570294564 | 1134 | 14 | 50 | 18123 | ACVRL1/AI | 14/1134 | 50/18123 |
| down-regulated | GO:BP | GO:0050878 | regulation of body fluid levels | 0.00409226634144892 | 1134 | 61 | 519 | 18123 | ADCY4/AD | 61/1134 | 519/18123 |
| down-regulated | GO:BP | GO:0022409 | positive regulation of cell-cell adhesion | 0.0040967708926243 | 1134 | 40 | 286 | 18123 | ANXA1/BC | 40/1134 | 286/18123 |
| down-regulated | GO:BP | GO:0046631 | alpha-beta T cell activation | 0.00415035832031754 | 1134 | 27 | 157 | 18123 | ADORA2A/ | 27/1134 | 157/18123 |
| down-regulated | GO:BP | GO:1904019 | epithelial cell apoptotic process | 0.00428948394664532 | 1134 | 24 | 130 | 18123 | AKR1C3/AI | 24/1134 | 130/18123 |
| down-regulated | GO:BP | GO:0060337 | type I interferon signaling pathway | 0.00449445168502301 | 1134 | 20 | 96 | 18123 | BST2/GBP | 20/1134 | 96/18123 |
| down-regulated | GO:BP | GO:0010232 | vascular transport | 0.00461723691592491 | 1134 | 19 | 88 | 18123 | ABCB1/AB | 19/1134 | 88/18123 |
| down-regulated | GO:BP | GO:0048839 | inner ear development | 0.00463480146212696 | 1134 | 31 | 196 | 18123 | ADGRV1/A | 31/1134 | 196/18123 |
| down-regulated | GO:BP | GO:0060541 | respiratory system development | 0.0047363812303489 | 1134 | 32 | 206 | 18123 | ASCL1/CEL | 32/1134 | 206/18123 |
| down-regulated | GO:BP | GO:0050866 | negative regulation of cell activation | 0.00478531241652712 | 1134 | 33 | 216 | 18123 | ADGRF5/A | 33/1134 | 216/18123 |
| down-regulated | GO:BP | GO:0019058 | viral life cycle | 0.00513768291758959 | 1134 | 49 | 386 | 18123 | ANPEP/AX | 49/1134 | 386/18123 |
| down-regulated | GO:BP | GO:0072109 | glomerular mesangium development | 0.00521538813196226 | 1134 | 8 | 16 | 18123 | BMP7/CD3 | 8/1134 | 16/18123 |
| down-regulated | GO:BP | GO:0030595 | leukocyte chemotaxis | 0.00529851183567158 | 1134 | 34 | 227 | 18123 | ANXA1/CC | 34/1134 | 227/18123 |
| down-regulated | GO:BP | GO:0071357 | cellular response to type I interferon | 0.00533313603906498 | 1134 | 20 | 97 | 18123 | BST2/GBP | 20/1134 | 97/18123 |
| down-regulated | GO:BP | GO:0061326 | renal tubule development | 0.00533313603906498 | 1134 | 20 | 97 | 18123 | BCL2/COL4 | 20/1134 | 97/18123 |
| down-regulated | GO:BP | GO:0048589 | developmental growth | 0.00547080932009302 | 1134 | 72 | 654 | 18123 | ACACB/AD | 72/1134 | 654/18123 |
| down-regulated | GO:BP | GO:0018108 | peptidyl-tyrosine phosphorylation | 0.00553505595940075 | 1134 | 49 | 387 | 18123 | AREG/AXL/ | 49/1134 | 387/18123 |
| down-regulated | GO:BP | GO:0051145 | smooth muscle cell differentiation | 0.00554897960956277 | 1134 | 18 | 81 | 18123 | ADM/APLN | 18/1134 | 81/18123 |
| down-regulated | GO:BP | GO:0002697 | regulation of immune effector process | 0.00571059744083896 | 1134 | 52 | 421 | 18123 | ANXA1/AP | 52/1134 | 421/18123 |
| down-regulated | GO:BP | GO:1901699 | cellular response to nitrogen compound | 0.00582678898046652 | 1134 | 78 | 728 | 18123 | ADCY4/AG | 78/1134 | 728/18123 |
| down-regulated | GO:BP | GO:0009615 | response to virus | 0.00602542977501969 | 1134 | 47 | 366 | 18123 | ABCC9/BCI | 47/1134 | 366/18123 |
| down-regulated | GO:BP | GO:0001837 | epithelial to mesenchymal transition | 0.00605268356410553 | 1134 | 27 | 160 | 18123 | BMP7/DAG | 27/1134 | 160/18123 |
| down-regulated | GO:BP | GO:1902337 | regulation of apoptotic process involved in morphogenesis | 0.00614082626819921 | 1134 | 7 | 12 | 18123 | BMP7/FOX | 7/1134 | 12/18123 |
| down-regulated | GO:BP | GO:0050776 | regulation of immune response | 0.00619207313046625 | 1134 | 112 | 1159 | 18123 | ADCY4/AD | 112/1134 | 1159/18123 |
| down-regulated | GO:BP | GO:0060443 | mammary gland morphogenesis | 0.00650133867052337 | 1134 | 13 | 45 | 18123 | AREG/CAV | 13/1134 | 45/18123 |
| down-regulated | GO:BP | GO:0007498 | mesoderm development | 0.00653878923334052 | 1134 | 24 | 133 | 18123 | BMP7/EPB | 24/1134 | 133/18123 |
| down-regulated | GO:BP | GO:0046649 | lymphocyte activation | 0.0067270130233992 | 1134 | 82 | 780 | 18123 | ADORA2A/ | 82/1134 | 780/18123 |
| down-regulated | GO:BP | GO:0033674 | positive regulation of kinase activity | 0.00685196179335455 | 1134 | 68 | 610 | 18123 | ADCY4/AD | 68/1134 | 610/18123 |
| down-regulated | GO:BP | GO:1904035 | regulation of epithelial cell apoptotic process | 0.00689641327299589 | 1134 | 21 | 107 | 18123 | AKR1C3/AI | 21/1134 | 107/18123 |
| down-regulated | GO:BP | GO:0018212 | peptidyl-tyrosine modification | 0.00690588159833753 | 1134 | 49 | 390 | 18123 | AREG/AXL/ | 49/1134 | 390/18123 |
| down-regulated | GO:BP | GO:0071417 | cellular response to organonitrogen compound | 0.00716563420226549 | 1134 | 73 | 671 | 18123 | ADCY4/AG | 73/1134 | 671/18123 |
| down-regulated | GO:BP | GO:0060255 | regulation of macromolecule metabolic process | 0.00738056610982439 | 1134 | 487 | 6622 | 18123 | A2ML1/AC | 487/1134 | 6622/18123 |
| down-regulated | GO:BP | GO:0010811 | positive regulation of cell-substrate adhesion | 0.00747400560516465 | 1134 | 23 | 125 | 18123 | CCDC80/CI | 23/1134 | 125/18123 |
| down-regulated | GO:BP | GO:0048699 | generation of neurons | 0.00793862900282251 | 1134 | 143 | 1576 | 18123 | ADGRG1/A | 143/1134 | 1576/18123 |
| down-regulated | GO:BP | GO:0048017 | inositol lipid-mediated signaling | 0.00798677599436629 | 1134 | 31 | 201 | 18123 | CSF3/EDN1 | 31/1134 | 201/18123 |
| down-regulated | GO:BP | GO:0002573 | myeloid leukocyte differentiation | 0.00799890516303909 | 1134 | 33 | 221 | 18123 | ANXA2/BA | 33/1134 | 221/18123 |
| down-regulated | GO:BP | GO:0045667 | regulation of osteoblast differentiation | 0.00858992982175129 | 1134 | 24 | 135 | 18123 | AREG/BMP | 24/1134 | 135/18123 |
| down-regulated | GO:BP | GO:0050670 | regulation of lymphocyte proliferation | 0.00870792144490479 | 1134 | 34 | 232 | 18123 | ANXA1/BC | 34/1134 | 232/18123 |
| down-regulated | GO:BP | GO:0070661 | leukocyte proliferation | 0.00874675386714005 | 1134 | 43 | 327 | 18123 | ANXA1/BC | 43/1134 | 327/18123 |
| down-regulated | GO:BP | GO:0002685 | regulation of leukocyte migration | 0.00890481322392358 | 1134 | 32 | 212 | 18123 | ANXA1/CC | 32/1134 | 212/18123 |
| down-regulated | GO:BP | GO:0003184 | pulmonary valve morphogenesis | 0.00931006235368772 | 1134 | 8 | 17 | 18123 | HEYL/JAG1 | 8/1134 | 17/18123 |
| down-regulated | GO:BP | GO:0030038 | contractile actin filament bundle assembly | 0.00940720458551563 | 1134 | 21 | 109 | 18123 | AMOT/ARH | 21/1134 | 109/18123 |
| down-regulated | GO:BP | GO:0043149 | stress fiber assembly | 0.00940720458551563 | 1134 | 21 | 109 | 18123 | AMOT/ARH | 21/1134 | 109/18123 |
| down-regulated | GO:BP | GO:0002695 | negative regulation of leukocyte activation | 0.0097641660198439 | 1134 | 30 | 193 | 18123 | ADGRF5/A | 30/1134 | 193/18123 |
| down-regulated | GO:BP | GO:0051128 | regulation of cellular component organization | 0.00986208501234754 | 1134 | 205 | 2439 | 18123 | ACAA2/AC | 205/1134 | 2439/18123 |
| down-regulated | GO:BP | GO:0015849 | organic acid transport | 0.0101482591506174 | 1134 | 42 | 318 | 18123 | ABCB1/AC | 42/1134 | 318/18123 |
| down-regulated | GO:BP | GO:0060341 | regulation of cellular localization | 0.0102470886418206 | 1134 | 87 | 851 | 18123 | ADCYAP1R | 87/1134 | 851/18123 |
| down-regulated | GO:BP | GO:0051056 | regulation of small GTPase mediated signal transduction | 0.0102541050026081 | 1134 | 43 | 329 | 18123 | ADCYAP1R | 43/1134 | 329/18123 |
| down-regulated | GO:BP | GO:0051249 | regulation of lymphocyte activation | 0.0108799285244214 | 1134 | 60 | 523 | 18123 | ADORA2A/ | 60/1134 | 523/18123 |
| down-regulated | GO:BP | GO:0061448 | connective tissue development | 0.0108926072424031 | 1134 | 36 | 255 | 18123 | ACVRL1/B | 36/1134 | 255/18123 |
| down-regulated | GO:BP | GO:0051493 | regulation of cytoskeleton organization | 0.0110291883937638 | 1134 | 61 | 535 | 18123 | ADD3/AKA | 61/1134 | 535/18123 |
| down-regulated | GO:BP | GO:0043123 | positive regulation of I-kappaB kinase/NF-kappaB signaling | 0.0119095673462299 | 1134 | 29 | 185 | 18123 | APOL3/BS | 29/1134 | 185/18123 |
| down-regulated | GO:BP | GO:0044092 | negative regulation of molecular function | 0.0120219754927464 | 1134 | 116 | 1228 | 18123 | A2ML1/AC | 116/1134 | 1228/18123 |
| down-regulated | GO:BP | GO:0009617 | response to bacterium | 0.0125412022159386 | 1134 | 78 | 743 | 18123 | ADM/ALPL | 78/1134 | 743/18123 |
| down-regulated | GO:BP | GO:2001233 | regulation of apoptotic signaling pathway | 0.0128316335795969 | 1134 | 46 | 365 | 18123 | ACAA2/AC | 46/1134 | 365/18123 |
| down-regulated | GO:BP | GO:0032956 | regulation of actin cytoskeleton organization | 0.0128316335795969 | 1134 | 46 | 365 | 18123 | ADD3/AM | 46/1134 | 365/18123 |
| down-regulated | GO:BP | GO:0072080 | nephron tubule development | 0.0130517995543879 | 1134 | 19 | 94 | 18123 | BCL2/DCH | 19/1134 | 94/18123 |
| down-regulated | GO:BP | GO:0051492 | regulation of stress fiber assembly | 0.0130517995543879 | 1134 | 19 | 94 | 18123 | AMOT/ARH | 19/1134 | 94/18123 |
| down-regulated | GO:BP | GO:0010038 | response to metal ion | 0.0136332341157165 | 1134 | 47 | 377 | 18123 | ADGRV1/A | 47/1134 | 377/18123 |
| down-regulated | GO:BP | GO:0051338 | regulation of transferase activity | 0.0137688540329874 | 1134 | 101 | 1036 | 18123 | ADCY4/AD | 101/1134 | 1036/18123 |
| down-regulated | GO:BP | GO:0003208 | cardiac ventricle morphogenesis | 0.0138097657845823 | 1134 | 16 | 70 | 18123 | CHD7/DLL | 16/1134 | 70/18123 |
| down-regulated | GO:BP | GO:0003229 | ventricular cardiac muscle tissue development | 0.013989899475463 | 1134 | 14 | 55 | 18123 | ADAMTS9/ | 14/1134 | 55/18123 |
| down-regulated | GO:BP | GO:0140546 | defense response to symbiont | 0.0148726178359205 | 1134 | 37 | 269 | 18123 | ABCC9/BCI | 37/1134 | 269/18123 |
| down-regulated | GO:BP | GO:0051607 | defense response to virus | 0.0148726178359205 | 1134 | 37 | 269 | 18123 | ABCC9/BCI | 37/1134 | 269/18123 |
| down-regulated | GO:BP | GO:0060348 | bone development | 0.0148989430203412 | 1134 | 31 | 207 | 18123 | ALPL/ANX | 31/1134 | 207/18123 |
| down-regulated | GO:BP | GO:0048015 | phosphatidylinositol-mediated signaling | 0.0149294798111841 | 1134 | 30 | 197 | 18123 | CSF3/EDN1 | 30/1134 | 197/18123 |
| down-regulated | GO:BP | GO:2000811 | negative regulation of anoikis | 0.0158381119551638 | 1134 | 8 | 18 | 18123 | BCL2/CAV1 | 8/1134 | 18/18123 |

| | | | | | | | | | | | |
|---|---|---|---|---|---|---|---|---|---|---|---|
| down-regulated | GO:BP | GO:0150104 | transport across blood-brain barrier | 0.0163544947227207 | 1134 | 18 | 87 | 18123 | ABCB1/AB | 18/1134 | 87/18123 |
| down-regulated | GO:BP | GO:0002507 | tolerance induction | 0.0164420635527771 | 1134 | 10 | 29 | 18123 | CD3E/FOXJ | 10/1134 | 29/18123 |
| down-regulated | GO:BP | GO:0110020 | regulation of actomyosin structure organization | 0.0164615445142842 | 1134 | 20 | 104 | 18123 | AMOT/ARH | 20/1134 | 104/18123 |
| down-regulated | GO:BP | GO:0008630 | intrinsic apoptotic signaling pathway in response to DNA damage | 0.0164615445142842 | 1134 | 20 | 104 | 18123 | ACKR3/BCL | 20/1134 | 104/18123 |
| down-regulated | GO:BP | GO:0032880 | regulation of protein localization | 0.0164645395040091 | 1134 | 91 | 912 | 18123 | ADORA2A/ | 91/1134 | 912/18123 |
| down-regulated | GO:BP | GO:0080134 | regulation of response to stress | 0.0171940275558639 | 1134 | 145 | 1626 | 18123 | ABCB1/ACI | 145/1134 | 1626/18123 |
| down-regulated | GO:BP | GO:0033002 | muscle cell proliferation | 0.0178416094350791 | 1134 | 35 | 250 | 18123 | APLN/CDKI | 35/1134 | 250/18123 |
| down-regulated | GO:BP | GO:0043434 | response to peptide hormone | 0.0179956947476981 | 1134 | 54 | 461 | 18123 | ADCY4/AD | 54/1134 | 461/18123 |
| down-regulated | GO:BP | GO:0046688 | response to copper ion | 0.0180652505471189 | 1134 | 12 | 42 | 18123 | AQP1/ICAN | 12/1134 | 42/18123 |
| down-regulated | GO:BP | GO:0003008 | system process | 0.0181039169278876 | 1134 | 194 | 2306 | 18123 | ABCB1/AB | 194/1134 | 2306/18123 |
| down-regulated | GO:BP | GO:0031099 | regeneration | 0.0182158889217783 | 1134 | 31 | 209 | 18123 | ADM/ANGI | 31/1134 | 209/18123 |
| down-regulated | GO:BP | GO:0031032 | actomyosin structure organization | 0.0183603964465186 | 1134 | 30 | 199 | 18123 | AMOT/ARH | 30/1134 | 199/18123 |
| down-regulated | GO:BP | GO:0030879 | mammary gland development | 0.0187644706746381 | 1134 | 24 | 141 | 18123 | APLN/AREG | 24/1134 | 141/18123 |
| down-regulated | GO:BP | GO:1902107 | positive regulation of leukocyte differentiation | 0.0189222778287117 | 1134 | 26 | 160 | 18123 | ANXA1/AX | 26/1134 | 160/18123 |
| down-regulated | GO:BP | GO:1903708 | positive regulation of hemopoiesis | 0.0189222778287117 | 1134 | 26 | 160 | 18123 | ANXA1/AX | 26/1134 | 160/18123 |
| down-regulated | GO:BP | GO:0051347 | positive regulation of transferase activity | 0.019413024814906 | 1134 | 73 | 690 | 18123 | ADCY4/AD | 73/1134 | 690/18123 |
| down-regulated | GO:BP | GO:0060249 | anatomical structure homeostasis | 0.019973893494454 | 1134 | 56 | 486 | 18123 | ADGRF5/A | 56/1134 | 486/18123 |
| down-regulated | GO:BP | GO:0048562 | embryonic organ morphogenesis | 0.0218466156549289 | 1134 | 39 | 295 | 18123 | APLNR/BM | 39/1134 | 295/18123 |
| down-regulated | GO:BP | GO:0090596 | sensory organ morphogenesis | 0.0221506655017489 | 1134 | 36 | 263 | 18123 | AQP5/BCL | 36/1134 | 263/18123 |
| down-regulated | GO:BP | GO:0045621 | positive regulation of lymphocyte differentiation | 0.0222421628929668 | 1134 | 20 | 106 | 18123 | ANXA1/AX | 20/1134 | 106/18123 |
| down-regulated | GO:BP | GO:0032231 | regulation of actin filament bundle assembly | 0.0222421628929668 | 1134 | 20 | 106 | 18123 | AMOT/ARH | 20/1134 | 106/18123 |
| down-regulated | GO:BP | GO:0002526 | acute inflammatory response | 0.0227024633947411 | 1134 | 21 | 115 | 18123 | B4GALT1/C | 21/1134 | 115/18123 |
| down-regulated | GO:BP | GO:0030278 | regulation of ossification | 0.0227024633947411 | 1134 | 21 | 115 | 18123 | ADGRV1/B | 21/1134 | 115/18123 |
| down-regulated | GO:BP | GO:0060325 | face morphogenesis | 0.0232846704410076 | 1134 | 10 | 30 | 18123 | CLDN5/CRI | 10/1134 | 30/18123 |
| down-regulated | GO:BP | GO:1904748 | regulation of apoptotic process involved in development | 0.0238054477723536 | 1134 | 7 | 14 | 18123 | BMP7/FOX | 7/1134 | 14/18123 |
| down-regulated | GO:BP | GO:1990169 | stress response to copper ion | 0.0238054477723536 | 1134 | 7 | 14 | 18123 | MT1A/MT1 | 7/1134 | 14/18123 |
| down-regulated | GO:BP | GO:0010273 | detoxification of copper ion | 0.0238054477723536 | 1134 | 7 | 14 | 18123 | MT1A/MT1 | 7/1134 | 14/18123 |
| down-regulated | GO:BP | GO:1903829 | positive regulation of cellular protein localization | 0.0238990673696802 | 1134 | 40 | 307 | 18123 | ANXA13/B | 40/1134 | 307/18123 |
| down-regulated | GO:BP | GO:0007265 | Ras protein signal transduction | 0.0253566239336702 | 1134 | 44 | 352 | 18123 | ADGRG1/A | 44/1134 | 352/18123 |
| down-regulated | GO:BP | GO:0051345 | positive regulation of hydrolase activity | 0.0259767275532447 | 1134 | 76 | 733 | 18123 | ADCYAP1R | 76/1134 | 733/18123 |
| down-regulated | GO:BP | GO:0042475 | odontogenesis of dentin-containing tooth | 0.0269514503812591 | 1134 | 18 | 90 | 18123 | ADM/BMP | 18/1134 | 90/18123 |
| down-regulated | GO:BP | GO:0050679 | positive regulation of epithelial cell proliferation | 0.0269634271281779 | 1134 | 31 | 213 | 18123 | ACVRL1/AI | 31/1134 | 213/18123 |
| down-regulated | GO:BP | GO:0030098 | lymphocyte differentiation | 0.0275310945750429 | 1134 | 47 | 387 | 18123 | ANXA1/AX | 47/1134 | 387/18123 |
| down-regulated | GO:BP | GO:0045619 | regulation of lymphocyte differentiation | 0.0277423000203817 | 1134 | 28 | 183 | 18123 | ANXA1/AX | 28/1134 | 183/18123 |
| down-regulated | GO:BP | GO:0007015 | actin filament organization | 0.0284625100839779 | 1134 | 52 | 445 | 18123 | ADD3/AIF1 | 52/1134 | 445/18123 |
| down-regulated | GO:BP | GO:0048878 | chemical homeostasis | 0.0287234210593059 | 1134 | 115 | 1238 | 18123 | ABHD4/AC | 115/1134 | 1238/18123 |
| down-regulated | GO:BP | GO:0060389 | pathway-restricted SMAD protein phosphorylation | 0.0292344730222236 | 1134 | 15 | 66 | 18123 | ACVRL1/BN | 15/1134 | 66/18123 |
| down-regulated | GO:BP | GO:0032943 | mononuclear cell proliferation | 0.0300650790976616 | 1134 | 39 | 299 | 18123 | ANXA1/BCI | 39/1134 | 299/18123 |
| down-regulated | GO:BP | GO:0001649 | osteoblast differentiation | 0.0304522343353553 | 1134 | 33 | 235 | 18123 | ALPL/AREG | 33/1134 | 235/18123 |
| down-regulated | GO:BP | GO:0045165 | cell fate commitment | 0.0311330096361631 | 1134 | 36 | 267 | 18123 | ASCL1/BCL | 36/1134 | 267/18123 |
| down-regulated | GO:BP | GO:0007599 | hemostasis | 0.0314647360246838 | 1134 | 44 | 355 | 18123 | ADORA2A/ | 44/1134 | 355/18123 |
| down-regulated | GO:BP | GO:1900182 | positive regulation of protein localization to nucleus | 0.0316581172171805 | 1134 | 18 | 91 | 18123 | BAG3/CAR | 18/1134 | 91/18123 |
| down-regulated | GO:BP | GO:0007266 | Rho protein signal transduction | 0.0323477509642675 | 1134 | 23 | 136 | 18123 | ADGRG1/A | 23/1134 | 136/18123 |
| down-regulated | GO:BP | GO:0060561 | apoptotic process involved in morphogenesis | 0.0323993898170164 | 1134 | 9 | 25 | 18123 | BMP7/FOX | 9/1134 | 25/18123 |
| down-regulated | GO:BP | GO:0032103 | positive regulation of response to external stimulus | 0.0328712024222191 | 1134 | 58 | 518 | 18123 | C2CD4B/CI | 58/1134 | 518/18123 |
| down-regulated | GO:BP | GO:0009991 | response to extracellular stimulus | 0.0349801252536463 | 1134 | 60 | 543 | 18123 | ACACB/AD | 60/1134 | 543/18123 |
| down-regulated | GO:BP | GO:1902903 | regulation of supramolecular fiber organization | 0.036103824464155 | 1134 | 47 | 391 | 18123 | ADD3/AEB | 47/1134 | 391/18123 |
| down-regulated | GO:BP | GO:0003205 | cardiac chamber development | 0.0372805148775908 | 1134 | 26 | 166 | 18123 | APLNR/BM | 26/1134 | 166/18123 |
| down-regulated | GO:BP | GO:0043542 | endothelial cell migration | 0.0378198387295403 | 1134 | 38 | 291 | 18123 | ACVRL1/AI | 38/1134 | 291/18123 |
| down-regulated | GO:BP | GO:0040008 | regulation of growth | 0.0404834452312379 | 1134 | 71 | 680 | 18123 | ACACB/AC | 71/1134 | 680/18123 |
| down-regulated | GO:BP | GO:0032495 | response to muramyl dipeptide | 0.040731159792642 | 1134 | 8 | 20 | 18123 | ERBIN/INA | 8/1134 | 20/18123 |
| down-regulated | GO:BP | GO:0045216 | cell-cell junction organization | 0.0432494815864884 | 1134 | 31 | 218 | 18123 | AMOT/APL | 31/1134 | 218/18123 |
| down-regulated | GO:BP | GO:0002221 | pattern recognition receptor signaling pathway | 0.0432494815864884 | 1134 | 31 | 218 | 18123 | APPL2/CAV | 31/1134 | 218/18123 |
| down-regulated | GO:BP | GO:0060317 | cardiac epithelial to mesenchymal transition | 0.0445683625743564 | 1134 | 10 | 32 | 18123 | ENG/HEYL/ | 10/1134 | 32/18123 |
| down-regulated | GO:BP | GO:0006357 | regulation of transcription by RNA polymerase II | 0.0445781056982732 | 1134 | 210 | 2567 | 18123 | ACVRL1/AI | 210/1134 | 2567/18123 |
| down-regulated | GO:BP | GO:0045859 | regulation of protein kinase activity | 0.047847064986836 | 1134 | 81 | 809 | 18123 | ADCY4/AD | 81/1134 | 809/18123 |
| down-regulated | GO:CC | GO:0071944 | cell periphery | 2.65056427553784e-20 | 1180 | 541 | 6178 | 18964 | ABCB1/AB | 541/1180 | 6178/18964 |
| down-regulated | GO:CC | GO:0005886 | plasma membrane | 1.15625466884372e-16 | 1180 | 493 | 5680 | 18964 | ABCB1/AB | 493/1180 | 5680/18964 |
| down-regulated | GO:CC | GO:0070161 | anchoring junction | 1.62071880865443e-13 | 1180 | 115 | 838 | 18964 | ADCYAP1R | 115/1180 | 838/18964 |
| down-regulated | GO:CC | GO:0031012 | extracellular matrix | 5.76463165739226e-12 | 1180 | 85 | 562 | 18964 | ADAMTS9/ | 85/1180 | 562/18964 |
| down-regulated | GO:CC | GO:0030312 | external encapsulating structure | 6.39298780587138e-12 | 1180 | 85 | 563 | 18964 | ADAMTS9/ | 85/1180 | 563/18964 |
| down-regulated | GO:CC | GO:0005576 | extracellular region | 2.70705483659837e-11 | 1180 | 393 | 4564 | 18964 | A2ML1/AB | 393/1180 | 4564/18964 |
| down-regulated | GO:CC | GO:0070062 | extracellular exosome | 7.85894288562558e-11 | 1180 | 218 | 2178 | 18964 | A2ML1/AB | 218/1180 | 2178/18964 |
| down-regulated | GO:CC | GO:0098590 | plasma membrane region | 3.52110897549613e-10 | 1180 | 141 | 1240 | 18964 | ABCB1/ACI | 141/1180 | 1240/18964 |
| down-regulated | GO:CC | GO:1903561 | extracellular vesicle | 3.52499465998251e-10 | 1180 | 222 | 2263 | 18964 | A2ML1/AB | 222/1180 | 2263/18964 |
| down-regulated | GO:CC | GO:0043230 | extracellular organelle | 3.85520738347383e-10 | 1180 | 222 | 2265 | 18964 | A2ML1/AB | 222/1180 | 2265/18964 |
| down-regulated | GO:CC | GO:0065010 | extracellular membrane-bounded organelle | 3.85520738347383e-10 | 1180 | 222 | 2265 | 18964 | A2ML1/AB | 222/1180 | 2265/18964 |
| down-regulated | GO:CC | GO:0030055 | cell-substrate junction | 6.8425023664858e-10 | 1180 | 67 | 427 | 18964 | AHNAK/AII | 67/1180 | 427/18964 |
| down-regulated | GO:CC | GO:0005925 | focal adhesion | 9.36889228700992e-10 | 1180 | 66 | 420 | 18964 | AHNAK/AII | 66/1180 | 420/18964 |
| down-regulated | GO:CC | GO:0062023 | collagen-containing extracellular matrix | 1.04582373848408e-09 | 1180 | 66 | 421 | 18964 | ADAMTS9/ | 66/1180 | 421/18964 |
| down-regulated | GO:CC | GO:0005615 | extracellular space | 1.43033510597274e-09 | 1180 | 317 | 3595 | 18964 | A2ML1/AB | 317/1180 | 3595/18964 |
| down-regulated | GO:CC | GO:0009986 | cell surface | 6.38240917676087e-09 | 1180 | 108 | 896 | 18964 | ABCB1/ACI | 108/1180 | 896/18964 |
| down-regulated | GO:CC | GO:0031982 | vesicle | 3.05255283953937e-08 | 1180 | 343 | 4058 | 18964 | A2ML1/AB | 343/1180 | 4058/18964 |
| down-regulated | GO:CC | GO:0045177 | apical part of cell | 8.10648429229834e-08 | 1180 | 63 | 433 | 18964 | ABCB1/AD | 63/1180 | 433/18964 |
| down-regulated | GO:CC | GO:0005737 | cytoplasm | 9.53590066316906e-08 | 1180 | 842 | 11951 | 18964 | A4GALT/AI | 842/1180 | 11951/18964 |
| down-regulated | GO:CC | GO:0015629 | actin cytoskeleton | 5.06535059792348e-07 | 1180 | 69 | 517 | 18964 | AHNAK/AII | 69/1180 | 517/18964 |
| down-regulated | GO:CC | GO:0005887 | integral component of plasma membrane | 5.14548579924999e-07 | 1180 | 162 | 1643 | 18964 | ABCC9/AC | 162/1180 | 1643/18964 |
| down-regulated | GO:CC | GO:0005911 | cell-cell junction | 5.32242118855538e-07 | 1180 | 67 | 496 | 18964 | ADCYAP1R | 67/1180 | 496/18964 |
| down-regulated | GO:CC | GO:0031226 | intrinsic component of plasma membrane | 6.0816542484253e-07 | 1180 | 168 | 1725 | 18964 | ABCC9/AC | 168/1180 | 1725/18964 |
| down-regulated | GO:CC | GO:0016324 | apical plasma membrane | 7.81559088980763e-07 | 1180 | 54 | 364 | 18964 | ABCB1/AH | 54/1180 | 364/18964 |
| down-regulated | GO:CC | GO:0001726 | ruffle | 6.36876187024353e-06 | 1180 | 33 | 180 | 18964 | AIF1L/AMC | 33/1180 | 180/18964 |
| down-regulated | GO:CC | GO:0031252 | cell leading edge | 1.25207006677379e-05 | 1180 | 57 | 426 | 18964 | ADGRV1/A | 57/1180 | 426/18964 |
| down-regulated | GO:CC | GO:0048471 | perinuclear region of cytoplasm | 2.63150634514158e-05 | 1180 | 84 | 745 | 18964 | ACKR3/AN | 84/1180 | 745/18964 |
| down-regulated | GO:CC | GO:0098857 | membrane microdomain | 2.65544667344928e-05 | 1180 | 48 | 339 | 18964 | ADCYAP1R | 48/1180 | 339/18964 |
| down-regulated | GO:CC | GO:0045121 | membrane raft | 2.65544667344928e-05 | 1180 | 48 | 339 | 18964 | ADCYAP1R | 48/1180 | 339/18964 |
| down-regulated | GO:CC | GO:0030054 | cell junction | 4.67067482230533e-05 | 1180 | 188 | 2107 | 18964 | ADCYAP1R | 188/1180 | 2107/18964 |
| down-regulated | GO:CC | GO:0005912 | adherens junction | 7.63506094101491e-05 | 1180 | 30 | 171 | 18964 | ANXA1/AN | 30/1180 | 171/18964 |
| down-regulated | GO:CC | GO:0045178 | basal part of cell | 7.83855140084485e-05 | 1180 | 40 | 268 | 18964 | ANXA1/AN | 40/1180 | 268/18964 |
| down-regulated | GO:CC | GO:0043235 | receptor complex | 0.000111326257870465 | 1180 | 63 | 522 | 18964 | ACVRL1/AI | 63/1180 | 522/18964 |
| down-regulated | GO:CC | GO:0009925 | basal plasma membrane | 0.000271644619097775 | 1180 | 37 | 250 | 18964 | ANXA1/AN | 37/1180 | 250/18964 |
| down-regulated | GO:CC | GO:0031253 | cell projection membrane | 0.000308441491609726 | 1180 | 46 | 346 | 18964 | ADGRV1/A | 46/1180 | 346/18964 |
| down-regulated | GO:CC | GO:0009897 | external side of plasma membrane | 0.000563708854482487 | 1180 | 51 | 409 | 18964 | ABCB1/ACI | 51/1180 | 409/18964 |
| down-regulated | GO:CC | GO:0031256 | leading edge membrane | 0.00197630574350228 | 1180 | 28 | 179 | 18964 | ADGRV1/A | 28/1180 | 179/18964 |
| down-regulated | GO:CC | GO:0098552 | side of membrane | 0.00263921234874443 | 1180 | 67 | 622 | 18964 | ABCB1/ACI | 67/1180 | 622/18964 |
| down-regulated | GO:CC | GO:0016323 | basolateral plasma membrane | 0.00343727277584832 | 1180 | 32 | 225 | 18964 | ANXA1/AN | 32/1180 | 225/18964 |
| down-regulated | GO:CC | GO:0098858 | actin-based cell projection | 0.00429391538125545 | 1180 | 31 | 217 | 18964 | ADGRV1/A | 31/1180 | 217/18964 |
| down-regulated | GO:CC | GO:0030027 | lamellipodium | 0.00807859604710833 | 1180 | 29 | 203 | 18964 | AMOT/AN | 29/1180 | 203/18964 |
| down-regulated | GO:CC | GO:0031091 | platelet alpha granule | 0.0116172382314474 | 1180 | 17 | 90 | 18964 | CD109/CLU | 17/1180 | 90/18964 |
| down-regulated | GO:CC | GO:0032587 | ruffle membrane | 0.011795194706438 | 1180 | 18 | 99 | 18964 | AIF1L/APPI | 18/1180 | 99/18964 |
| down-regulated | GO:CC | GO:0016021 | integral component of membrane | 0.0144874337932473 | 1180 | 417 | 5718 | 18964 | A4GALT/AI | 417/1180 | 5718/18964 |
| down-regulated | GO:CC | GO:0042383 | sarcolemma | 0.0198535563962118 | 1180 | 22 | 141 | 18964 | AHNAK/AN | 22/1180 | 141/18964 |
| down-regulated | GO:CC | GO:0031224 | intrinsic component of membrane | 0.0247413494493135 | 1180 | 425 | 5873 | 18964 | A4GALT/AI | 425/1180 | 5873/18964 |
| down-regulated | GO:CC | GO:0008305 | integrin complex | 0.0346087831069241 | 1180 | 9 | 32 | 18964 | EMILIN1/IT | 9/1180 | 32/18964 |
| down-regulated | GO:CC | GO:0005902 | microvillus | 0.036310980994107 | 1180 | 16 | 89 | 18964 | AQP5/CD4 | 16/1180 | 89/18964 |
| down-regulated | GO:CC | GO:0072562 | blood microparticle | 0.0463830910536551 | 1180 | 21 | 139 | 18964 | ANGPTL4/ | 21/1180 | 139/18964 |
| down-regulated | GO:MF | GO:0005515 | protein binding | 4.00453166057001e-11 | 1182 | 1030 | 14767 | 18679 | A2ML1/AB | 1030/1182 | 14767/18679 |
| down-regulated | GO:MF | GO:0005102 | signaling receptor binding | 2.15119432770463e-08 | 1182 | 163 | 1551 | 18679 | ADA2/ADM | 163/1182 | 1551/18679 |
| down-regulated | GO:MF | GO:0019838 | growth factor binding | 2.18441195482495e-08 | 1182 | 33 | 141 | 18679 | ACVRL1/CC | 33/1182 | 141/18679 |
| down-regulated | GO:MF | GO:0050839 | cell adhesion molecule binding | 6.06922378645369e-06 | 1182 | 70 | 544 | 18679 | AHNAK/AN | 70/1182 | 544/18679 |
| down-regulated | GO:MF | GO:0019955 | cytokine binding | 2.62087404657716e-05 | 1182 | 28 | 140 | 18679 | ACKR3/AC | 28/1182 | 140/18679 |
| down-regulated | GO:MF | GO:0005488 | binding | 0.000188419330826853 | 1182 | 1124 | 17070 | 18679 | A2ML1/A4 | 1124/1182 | 17070/18679 |
| down-regulated | GO:MF | GO:0098631 | cell adhesion mediator activity | 0.000216302284620495 | 1182 | 16 | 58 | 18679 | ANXA1/AN | 16/1182 | 58/18679 |
| down-regulated | GO:MF | GO:0019899 | enzyme binding | 0.000524007004267612 | 1182 | 182 | 2048 | 18679 | A2ML1/AB | 182/1182 | 2048/18679 |

| | | | | | | | | | | | |
|---|---|---|---|---|---|---|---|---|---|---|---|
| down-regulated | GO:MF | GO:0042802 | identical protein binding | 0.000548078244404409 | 1182 | 183 | 2063 | 18679 | ACACB/AC | 183/1182 | 2063/18679 |
| down-regulated | GO:MF | GO:0001618 | virus receptor activity | 0.00230170941935707 | 1182 | 17 | 76 | 18679 | ANPEP/AX | 17/1182 | 76/18679 |
| down-regulated | GO:MF | GO:0019199 | transmembrane receptor protein kinase activity | 0.0023960703219422 | 1182 | 25 | 145 | 18679 | ACVRL1/A | 25/1182 | 145/18679 |
| down-regulated | GO:MF | GO:0140272 | exogenous protein binding | 0.00278159616339343 | 1182 | 17 | 77 | 18679 | ANPEP/AX | 17/1182 | 77/18679 |
| down-regulated | GO:MF | GO:0005518 | collagen binding | 0.00326571178302028 | 1182 | 16 | 70 | 18679 | ADGRG1/A | 16/1182 | 70/18679 |
| down-regulated | GO:MF | GO:0050431 | transforming growth factor beta binding | 0.00492366434264561 | 1182 | 9 | 24 | 18679 | ACVRL1/C | 9/1182 | 24/18679 |
| down-regulated | GO:MF | GO:0044548 | S100 protein binding | 0.00524730816618525 | 1182 | 7 | 14 | 18679 | AHNAK/AN | 7/1182 | 14/18679 |
| down-regulated | GO:MF | GO:0005126 | cytokine receptor binding | 0.00602792884445662 | 1182 | 37 | 274 | 18679 | CCDC88A/ | 37/1182 | 274/18679 |
| down-regulated | GO:MF | GO:0005178 | integrin binding | 0.00854208480476489 | 1182 | 24 | 146 | 18679 | CCN5/CD8 | 24/1182 | 146/18679 |
| down-regulated | GO:MF | GO:0008289 | lipid binding | 0.00949019446232925 | 1182 | 79 | 774 | 18679 | ACBD7/AC | 79/1182 | 774/18679 |
| down-regulated | GO:MF | GO:0008514 | organic anion transmembrane transporter activity | 0.0129384045407601 | 1182 | 27 | 179 | 18679 | CTNS/GJA1 | 27/1182 | 179/18679 |
| down-regulated | GO:MF | GO:0008013 | beta-catenin binding | 0.0153945521592522 | 1182 | 17 | 87 | 18679 | CDH5/CTN | 17/1182 | 87/18679 |
| down-regulated | GO:MF | GO:0003779 | actin binding | 0.0206524506757334 | 1182 | 51 | 449 | 18679 | ADD3/AIF1 | 51/1182 | 449/18679 |
| down-regulated | GO:MF | GO:0042803 | protein homodimerization activity | 0.0208638089284288 | 1182 | 70 | 679 | 18679 | ACOX2/AD | 70/1182 | 679/18679 |
| down-regulated | GO:MF | GO:0098632 | cell-cell adhesion mediator activity | 0.0229338192611977 | 1182 | 12 | 49 | 18679 | ANXA1/AN | 12/1182 | 49/18679 |
| down-regulated | GO:MF | GO:0008509 | anion transmembrane transporter activity | 0.0265127200681325 | 1182 | 38 | 304 | 18679 | APOL1/AQ | 38/1182 | 304/18679 |
| down-regulated | GO:MF | GO:0000987 | cis-regulatory region sequence-specific DNA binding | 0.0295677100972041 | 1182 | 110 | 1203 | 18679 | ASCL1/ATC | 110/1182 | 1203/18679 |
| down-regulated | GO:MF | GO:0000978 | RNA polymerase II cis-regulatory region sequence-specific DNA bindir | 0.0382290141047731 | 1182 | 108 | 1184 | 18679 | ASCL1/ATC | 108/1182 | 1184/18679 |
| down-regulated | GO:MF | GO:0045296 | cadherin binding | 0.0415642014970182 | 1182 | 40 | 333 | 18679 | AHNAK/AN | 40/1182 | 333/18679 |
| down-regulated | GO:MF | GO:0004857 | enzyme inhibitor activity | 0.0427134973344219 | 1182 | 45 | 391 | 18679 | A2ML1/AD | 45/1182 | 391/18679 |
| down-regulated | GO:MF | GO:0001216 | DNA-binding transcription activator activity | 0.0482003885492212 | 1182 | 50 | 452 | 18679 | CEBPD/CR | 50/1182 | 452/18679 |
| down-regulated | HP | HP:0001112 | Leber optic atrophy | 2.1550553380292e-08 | 373 | 10 | 10 | 4511 | MT-ATP6/ | 10/373 | 10/4511 |
| down-regulated | HP | HP:0005116 | Arterial tortuosity | 2.18738759374658e-08 | 373 | 14 | 20 | 4511 | EFEMP2/M | 14/373 | 20/4511 |
| down-regulated | HP | HP:0004948 | Vascular tortuosity | 1.5459582828805e-07 | 373 | 14 | 22 | 4511 | EFEMP2/M | 14/373 | 22/4511 |
| down-regulated | HP | HP:0000631 | Retinal arterial tortuosity | 2.19675836401061e-07 | 373 | 10 | 11 | 4511 | MT-ATP6/ | 10/373 | 11/4511 |
| down-regulated | HP | HP:0007768 | Central retinal vessel vascular tortuosity | 2.19675836401061e-07 | 373 | 10 | 11 | 4511 | MT-ATP6/ | 10/373 | 11/4511 |
| down-regulated | HP | HP:0000576 | Centrocecal scotoma | 1.22146330911508e-06 | 373 | 10 | 12 | 4511 | MT-ATP6/ | 10/373 | 12/4511 |
| down-regulated | HP | HP:0007763 | Retinal telangiectasia | 4.11280174025763e-06 | 373 | 12 | 19 | 4511 | ACVRL1/E | 12/373 | 19/4511 |
| down-regulated | HP | HP:0001269 | Hemiparesis | 4.57932938892779e-06 | 373 | 26 | 87 | 4511 | ADA2/ARH | 26/373 | 87/4511 |
| down-regulated | HP | HP:0001427 | Mitochondrial inheritance | 1.35899916330095e-05 | 373 | 11 | 17 | 4511 | MT-ATP6/ | 11/373 | 17/4511 |
| down-regulated | HP | HP:0011025 | Abnormal cardiovascular system physiology | 8.72716661770336e-05 | 373 | 140 | 1150 | 4511 | A2ML1/AB | 140/373 | 1150/4511 |
| down-regulated | HP | HP:0000622 | Blurred vision | 0.000169244784663088 | 373 | 14 | 33 | 4511 | ATP1A2/C | 14/373 | 33/4511 |
| down-regulated | HP | HP:0012766 | Widened cerebral subarachnoid space | 0.000300708873035055 | 373 | 7 | 8 | 4511 | MT-CO1/M | 7/373 | 8/4511 |
| down-regulated | HP | HP:0000822 | Hypertension | 0.000349166380854744 | 373 | 52 | 307 | 4511 | ACVRL1/A | 52/373 | 307/4511 |
| down-regulated | HP | HP:0032263 | Increased blood pressure | 0.00037787754132211 | 373 | 57 | 351 | 4511 | ACVRL1/A | 57/373 | 351/4511 |
| down-regulated | HP | HP:0003572 | Low plasma citrulline | 0.000433617547659317 | 373 | 8 | 11 | 4511 | MT-ATP6/ | 8/373 | 11/4511 |
| down-regulated | HP | HP:0200125 | Mitochondrial respiratory chain defects | 0.000513358225131566 | 373 | 10 | 18 | 4511 | MT-ATP6/ | 10/373 | 18/4511 |
| down-regulated | HP | HP:0002076 | Migraine | 0.000881684659516216 | 373 | 26 | 110 | 4511 | ACVRL1/A | 26/373 | 110/4511 |
| down-regulated | HP | HP:0002572 | Episodic vomiting | 0.000901327237364239 | 373 | 13 | 32 | 4511 | MT-ATP6/ | 13/373 | 32/4511 |
| down-regulated | HP | HP:0004309 | Ventricular preexcitation | 0.000939081683204098 | 373 | 11 | 23 | 4511 | MT-ATP6/ | 11/373 | 23/4511 |
| down-regulated | HP | HP:0012841 | Retinal vascular tortuosity | 0.00124243163791739 | 373 | 15 | 43 | 4511 | COL4A1/L | 15/373 | 43/4511 |
| down-regulated | HP | HP:0002315 | Headache | 0.00145883853550227 | 373 | 43 | 244 | 4511 | ACVRL1/A | 43/373 | 244/4511 |
| down-regulated | HP | HP:0030972 | Abnormal systemic blood pressure | 0.00216784249722344 | 373 | 63 | 424 | 4511 | ACVRL1/A | 63/373 | 424/4511 |
| down-regulated | HP | HP:0011276 | Vascular skin abnormality | 0.00287193014396035 | 373 | 70 | 493 | 4511 | ACVRL1/A | 70/373 | 493/4511 |
| down-regulated | HP | HP:0012429 | Aplasia/Hypoplasia of the cerebral white matter | 0.00354520022384447 | 373 | 9 | 17 | 4511 | MT-CO1/M | 9/373 | 17/4511 |
| down-regulated | HP | HP:0003481 | Segmental peripheral demyelination/remyelination | 0.00354520022384447 | 373 | 9 | 17 | 4511 | MT-ATP6/ | 9/373 | 17/4511 |
| down-regulated | HP | HP:0005157 | Concentric hypertrophic cardiomyopathy | 0.00389291078118545 | 373 | 7 | 10 | 4511 | MT-CO1/M | 7/373 | 10/4511 |
| down-regulated | HP | HP:0100611 | Multiple glomerular cysts | 0.00389291078118545 | 373 | 7 | 10 | 4511 | MT-ATP6/ | 7/373 | 10/4511 |
| down-regulated | HP | HP:0007327 | Mixed demyelinating and axonal polyneuropathy | 0.00389291078118545 | 373 | 7 | 10 | 4511 | MT-CO1/M | 7/373 | 10/4511 |
| down-regulated | HP | HP:0011965 | Abnormal circulating citrulline concentration | 0.00631061763896002 | 373 | 8 | 14 | 4511 | MT-ATP6/ | 8/373 | 14/4511 |
| down-regulated | HP | HP:0000575 | Scotoma | 0.0084366151830419 | 373 | 19 | 74 | 4511 | ADGRV1/A | 19/373 | 74/4511 |
| down-regulated | HP | HP:0033109 | Abnormal circulating non-proteinogenic amino acid concentration | 0.00919009266402545 | 373 | 10 | 23 | 4511 | HFE/MT-A | 10/373 | 23/4511 |
| down-regulated | HP | HP:0002401 | Stroke-like episode | 0.0125531361766901 | 373 | 8 | 15 | 4511 | MT-CO1/M | 8/373 | 15/4511 |
| down-regulated | HP | HP:0002597 | Abnormality of the vasculature | 0.0162638509929202 | 373 | 173 | 1622 | 4511 | A2ML1/AB | 173/373 | 1622/4511 |
| down-regulated | HP | HP:0002922 | Increased CSF protein | 0.0182150055878966 | 373 | 14 | 46 | 4511 | ATP1A2/FT | 14/373 | 46/4511 |
| down-regulated | HP | HP:0100651 | Type I diabetes mellitus | 0.0182150055878966 | 373 | 14 | 46 | 4511 | ADA2/EDA | 14/373 | 46/4511 |
| down-regulated | HP | HP:0033353 | Abnormal blood vessel morphology | 0.0198430936046331 | 373 | 98 | 802 | 4511 | A2ML1/AB | 98/373 | 802/4511 |
| down-regulated | HP | HP:0004374 | Hemiplegia/hemiparesis | 0.0213814257895353 | 373 | 35 | 201 | 4511 | ADA2/ARH | 35/373 | 201/4511 |
| down-regulated | HP | HP:0025268 | Stuttering | 0.0221817328677403 | 373 | 7 | 12 | 4511 | MT-CO1/M | 7/373 | 12/4511 |
| down-regulated | HP | HP:0003200 | Ragged-red muscle fibers | 0.0229139826117643 | 373 | 13 | 41 | 4511 | MT-ATP6/ | 13/373 | 41/4511 |
| down-regulated | HP | HP:0000006 | Autosomal dominant inheritance | 0.0245661502398265 | 373 | 186 | 1783 | 4511 | A2ML1/A4 | 186/373 | 1783/4511 |
| down-regulated | HP | HP:0025015 | Abnormal vascular morphology | 0.0247190471885395 | 373 | 98 | 806 | 4511 | A2ML1/AB | 98/373 | 806/4511 |
| down-regulated | HP | HP:0001892 | Abnormal bleeding | 0.0304494234248582 | 373 | 61 | 439 | 4511 | A2ML1/AC | 61/373 | 439/4511 |
| down-regulated | HP | HP:0025456 | Abnormal CSF protein level | 0.0314056689676015 | 373 | 14 | 48 | 4511 | ATP1A2/FT | 14/373 | 48/4511 |
| down-regulated | HP | HP:0007067 | Distal peripheral sensory neuropathy | 0.0446611720982115 | 373 | 7 | 13 | 4511 | MT-CO1/M | 7/373 | 13/4511 |
| down-regulated | HPA | HPA:0100202 | cerebral cortex; endothelial cells[<U+2265>Medium] | 2.26701504221891e-10 | 770 | 256 | 2450 | 10877 | ABCB1/AB | 256/770 | 2450/10877 |
| down-regulated | HPA | HPA:0100203 | cerebral cortex; endothelial cells[High] | 1.03912510261211e-08 | 770 | 67 | 407 | 10877 | ABCB1/AD | 67/770 | 407/10877 |
| down-regulated | HPA | HPA:0100201 | cerebral cortex; endothelial cells[<U+2265>Low] | 8.74322967156397e-07 | 770 | 402 | 4585 | 10877 | ABCB1/AB | 402/770 | 4585/10877 |
| down-regulated | HPA | HPA:0130203 | colon; endothelial cells[High] | 4.70467003699806e-06 | 770 | 98 | 792 | 10877 | ADIRF/AD | 98/770 | 792/10877 |
| down-regulated | HPA | HPA:0270000 | kidney | 0.00535685313739708 | 770 | 614 | 8002 | 10877 | ABCB1/AB | 614/770 | 8002/10877 |
| down-regulated | HPA | HPA:0090973 | cerebellum; granular cells - cytoplasm/membrane[High] | 0.0114561404624836 | 770 | 23 | 132 | 10877 | ALDH1L1/ | 23/770 | 132/10877 |
| down-regulated | HPA | HPA:0270352 | kidney; cells in glomeruli[<U+2265>Medium] | 0.0126750139537624 | 770 | 257 | 2959 | 10877 | ADCY4/AD | 257/770 | 2959/10877 |
| down-regulated | HPA | HPA:0091033 | cerebellum; processes in molecular layer[High] | 0.0145726102161893 | 770 | 20 | 108 | 10877 | AKAP9/AL | 20/770 | 108/10877 |
| down-regulated | HPA | HPA:0091032 | cerebellum; processes in molecular layer[<U+2265>Medium] | 0.0195529215190623 | 770 | 29 | 192 | 10877 | AKAP9/AL | 29/770 | 192/10877 |
| down-regulated | HPA | HPA:0570772 | testis; peritubular cells[<U+2265>Medium] | 0.0401836156855368 | 770 | 54 | 462 | 10877 | ADD3/AKA | 54/770 | 462/10877 |
| down-regulated | KEGG | KEGG:04060 | Cytokine-cytokine receptor interaction | 1.8604622183566e-05 | 581 | 47 | 293 | 8000 | ACKR3/AC | 47/581 | 293/8000 |
| down-regulated | KEGG | KEGG:04668 | TNF signaling pathway | 0.000135843239910989 | 581 | 24 | 112 | 8000 | BCL3/CASP | 24/581 | 112/8000 |
| down-regulated | KEGG | KEGG:04151 | PI3K-Akt signaling pathway | 0.000152870261871099 | 581 | 51 | 353 | 8000 | ANGPT2/A | 51/581 | 353/8000 |
| down-regulated | KEGG | KEGG:04933 | AGE-RAGE signaling pathway in diabetic complications | 0.000238442793756705 | 581 | 22 | 100 | 8000 | BCL2/CCL2 | 22/581 | 100/8000 |
| down-regulated | KEGG | KEGG:04390 | Hippo signaling pathway | 0.000276179742004787 | 581 | 29 | 157 | 8000 | AMOT/AR | 29/581 | 157/8000 |
| down-regulated | KEGG | KEGG:04064 | NF-kappa B signaling pathway | 0.000338536286300529 | 581 | 22 | 102 | 8000 | BCL2/CAR | 22/581 | 102/8000 |
| down-regulated | KEGG | KEGG:05418 | Fluid shear stress and atherosclerosis | 0.00195311235397992 | 581 | 25 | 138 | 8000 | BCL2/BMP | 25/581 | 138/8000 |
| down-regulated | KEGG | KEGG:04392 | Hippo signaling pathway - multiple species | 0.00254804286196863 | 581 | 10 | 29 | 8000 | DCHS1/LA | 10/581 | 29/8000 |
| down-regulated | KEGG | KEGG:05144 | Malaria | 0.00374184125868721 | 581 | 13 | 49 | 8000 | CCL2/CD4 | 13/581 | 49/8000 |
| down-regulated | KEGG | KEGG:04010 | MAPK signaling pathway | 0.00407902021985123 | 581 | 41 | 294 | 8000 | ANGPT2/A | 41/581 | 294/8000 |
| down-regulated | KEGG | KEGG:04512 | ECM-receptor interaction | 0.00574105801541251 | 581 | 18 | 88 | 8000 | CD44/COL | 18/581 | 88/8000 |
| down-regulated | KEGG | KEGG:05166 | Human T-cell leukemia virus 1 infection | 0.00935661402505226 | 581 | 32 | 216 | 8000 | ADCY4/CD | 32/581 | 216/8000 |
| down-regulated | KEGG | KEGG:04066 | HIF-1 signaling pathway | 0.0112766024114627 | 581 | 20 | 109 | 8000 | ANGPT2/B | 20/581 | 109/8000 |
| down-regulated | KEGG | KEGG:05412 | Arrhythmogenic right ventricular cardiomyopathy | 0.0122961898553359 | 581 | 16 | 77 | 8000 | CTNNA3/D | 16/581 | 77/8000 |
| down-regulated | KEGG | KEGG:04610 | Complement and coagulation cascades | 0.0124493007850196 | 581 | 17 | 85 | 8000 | C1R/C4A/ | 17/581 | 85/8000 |
| down-regulated | KEGG | KEGG:05165 | Human papillomavirus infection | 0.0142719748782244 | 581 | 43 | 331 | 8000 | CDK2/CDK | 43/581 | 331/8000 |
| down-regulated | KEGG | KEGG:05167 | Kaposi sarcoma-associated herpesvirus infection | 0.0156129046265092 | 581 | 29 | 193 | 8000 | ANGPT2/C | 29/581 | 193/8000 |
| down-regulated | KEGG | KEGG:05224 | Breast cancer | 0.0161662496352166 | 581 | 24 | 147 | 8000 | BRCA1/CD | 24/581 | 147/8000 |
| down-regulated | KEGG | KEGG:04640 | Hematopoietic cell lineage | 0.0163422004491412 | 581 | 18 | 95 | 8000 | ANPEP/CD | 18/581 | 95/8000 |
| down-regulated | KEGG | KEGG:05226 | Gastric cancer | 0.0180096622115202 | 581 | 24 | 148 | 8000 | ABCB1/BC | 24/581 | 148/8000 |
| down-regulated | KEGG | KEGG:04330 | Notch signaling pathway | 0.0293924366363049 | 581 | 13 | 59 | 8000 | DLL4/DTX3 | 13/581 | 59/8000 |
| down-regulated | KEGG | KEGG:04978 | Mineral absorption | 0.0293924366363049 | 581 | 13 | 59 | 8000 | ATP1A2/C | 13/581 | 59/8000 |
| down-regulated | KEGG | KEGG:04218 | Cellular senescence | 0.0408507334904592 | 581 | 24 | 156 | 8000 | CDK2/CDK | 24/581 | 156/8000 |
| down-regulated | MIRNA | MIRNA:hsa-miR-1 | hsa-miR-124-3p | 6.21114117622648e-14 | 1058 | 187 | 1441 | 14823 | ABHD4/AC | 187/1058 | 1441/14823 |
| down-regulated | MIRNA | MIRNA:hsa-miR-1 | hsa-miR-146a-5p | 0.000193712096077719 | 1058 | 36 | 200 | 14823 | BRCA1/BT | 36/1058 | 200/14823 |
| down-regulated | MIRNA | MIRNA:hsa-miR-2 | hsa-miR-200c-3p | 0.00756572188060313 | 1058 | 34 | 214 | 14823 | BBX/BCL2/ | 34/1058 | 214/14823 |
| down-regulated | MIRNA | MIRNA:hsa-miR-2 | hsa-miR-21-5p | 0.011371274545669 | 1058 | 72 | 607 | 14823 | ABCB1/AK | 72/1058 | 607/14823 |
| down-regulated | MIRNA | MIRNA:hsa-miR-1 | hsa-miR-145-5p | 0.0256560614687277 | 1058 | 35 | 236 | 14823 | ADD3/AGT | 35/1058 | 236/14823 |
| down-regulated | MIRNA | MIRNA:hsa-miR-2 | hsa-miR-223-3p | 0.0486677052313676 | 1058 | 19 | 98 | 14823 | ABCB1/CD | 19/1058 | 98/14823 |
| down-regulated | REAC | REAC:R-HSA-1280 | Cytokine Signaling in Immune system | 1.77183823328045e-11 | 734 | 103 | 696 | 10622 | ANXA1/AN | 103/734 | 696/10622 |
| down-regulated | REAC | REAC:R-HSA-6785 | Interleukin-4 and Interleukin-13 signaling | 3.21393925260189e-09 | 734 | 31 | 110 | 10622 | ANXA1/BC | 31/734 | 110/10622 |
| down-regulated | REAC | REAC:R-HSA-4491 | Signaling by Interleukins | 2.19953459206504e-07 | 734 | 69 | 459 | 10622 | ANXA1/AN | 69/734 | 459/10622 |
| down-regulated | REAC | REAC:R-HSA-1682 | Immune System | 1.78154518537386e-05 | 734 | 199 | 2038 | 10622 | ADA2/AHC | 199/734 | 2038/10622 |
| down-regulated | REAC | REAC:R-HSA-8951 | RUNX3 regulates YAP1-mediated transcription | 2.70051362129065e-05 | 734 | 7 | 8 | 10622 | RUNX3/TE | 7/734 | 8/10622 |
| down-regulated | REAC | REAC:R-HSA-6783 | Interleukin-10 signaling | 8.1135412953472e-05 | 734 | 15 | 45 | 10622 | CCL2/CXCL | 15/734 | 45/10622 |
| down-regulated | REAC | REAC:R-HSA-9097 | Interferon alpha/beta signaling | 0.000245600818907636 | 734 | 18 | 68 | 10622 | BST2/GBP | 18/734 | 68/10622 |
| down-regulated | REAC | REAC:R-HSA-1625 | Signal Transduction | 0.000298338306429336 | 734 | 246 | 2730 | 10622 | ACKR3/AC | 246/734 | 2730/10622 |
| down-regulated | REAC | REAC:R-HSA-2032 | YAP1- and WWTR1 (TAZ)-stimulated gene expression | 0.000507251264109985 | 734 | 8 | 14 | 10622 | HIPK2/KAT | 8/734 | 14/10622 |

| | | | | | | | | | | | |
|---|---|---|---|---|---|---|---|---|---|---|---|
| down-regulated | REAC | REAC:R-HSA-5660 | Response to metal ions | 0.000507251264109985 | 734 | 8 | 14 | 10622 | CSRP1/MT | 8/734 | 14/10622 |
| down-regulated | REAC | REAC:R-HSA-8773 | Interferon gamma signaling | 0.000671912312998067 | 734 | 20 | 87 | 10622 | CD44/GBP | 20/734 | 87/10622 |
| down-regulated | REAC | REAC:R-HSA-5083 | Defective LFNG causes SCDO3 | 0.000762603195586092 | 734 | 5 | 5 | 10622 | LFNG/NOT | 5/734 | 5/10622 |
| down-regulated | REAC | REAC:R-HSA-9006 | Signaling by Receptor Tyrosine Kinases | 0.000763284049947671 | 734 | 62 | 486 | 10622 | ADCYAP1R | 62/734 | 486/10622 |
| down-regulated | REAC | REAC:R-HSA-5661 | Metallothioneins bind metals | 0.000925786037659445 | 734 | 7 | 11 | 10622 | MT1A/MT | 7/734 | 11/10622 |
| down-regulated | REAC | REAC:R-HSA-9135 | Interferon Signaling | 0.00145328934911648 | 734 | 32 | 193 | 10622 | BST2/CD44 | 32/734 | 193/10622 |
| down-regulated | REAC | REAC:R-HSA-8878 | Transcriptional regulation by RUNX3 | 0.00241150624823296 | 734 | 20 | 94 | 10622 | CDKN1A/H | 20/734 | 94/10622 |
| down-regulated | REAC | REAC:R-HSA-1941 | Signaling by VEGF | 0.00336749515814057 | 734 | 21 | 104 | 10622 | AHCYL1/A | 21/734 | 104/10622 |
| down-regulated | REAC | REAC:R-HSA-1474 | Extracellular matrix organization | 0.00344417465480104 | 734 | 42 | 298 | 10622 | ADAMTS9/ | 42/734 | 298/10622 |
| down-regulated | REAC | REAC:R-HSA-2129 | Molecules associated with elastic fibres | 0.0126451721556407 | 734 | 11 | 37 | 10622 | BMP7/EFE | 11/734 | 37/10622 |
| down-regulated | REAC | REAC:R-HSA-1566 | Elastic fibre formation | 0.0142967436935335 | 734 | 12 | 44 | 10622 | BMP7/EFE | 12/734 | 44/10622 |
| down-regulated | REAC | REAC:R-HSA-3000 | Non-integrin membrane-ECM interactions | 0.0142999363012042 | 734 | 14 | 58 | 10622 | COL4A1/D | 14/734 | 58/10622 |
| down-regulated | REAC | REAC:R-HSA-2160 | Integrin cell surface interactions | 0.0242083999713144 | 734 | 17 | 84 | 10622 | CD44/COL | 17/734 | 84/10622 |
| down-regulated | REAC | REAC:R-HSA-8951 | RUNX3 regulates WNT signaling | 0.0357972541884252 | 734 | 5 | 8 | 10622 | LEF1/RUN | 5/734 | 8/10622 |
| down-regulated | REAC | REAC:R-HSA-4411 | Binding of TCF/LEF:CTNNB1 to target gene promoters | 0.0357972541884252 | 734 | 5 | 8 | 10622 | LEF1/RUN | 5/734 | 8/10622 |
| down-regulated | REAC | REAC:R-HSA-3560 | Diseases associated with glycosaminoglycan metabolism | 0.0362193045889268 | 734 | 11 | 41 | 10622 | B4GALT1/ | 11/734 | 41/10622 |
| down-regulated | TF | TF:M09984 | Factor: MAZ; motif: GGGGGAGGGGGNGRGRRRGNRG | 5.88740984078488e-20 | 1241 | 769 | 9627 | 19917 | ABHD15/A | 769/1241 | 9627/19917 |
| down-regulated | TF | TF:M01100_1 | Factor: LRF; motif: GGGGKYNNB; match class: 1 | 2.79776174787221e-14 | 1241 | 281 | 2777 | 19917 | A4GALT/A | 281/1241 | 2777/19917 |
| down-regulated | TF | TF:M01857 | Factor: AP-2alpha; motif: NGCCYSNNGSN | 3.23882692402222e-14 | 1241 | 720 | 9223 | 19917 | A4GALT/A | 720/1241 | 9223/19917 |
| down-regulated | TF | TF:M07289_1 | Factor: GKLF; motif: NNNRGGNGNGGSN; match class: 1 | 5.88521588527209e-14 | 1241 | 806 | 10645 | 19917 | ABHD15/A | 806/1241 | 10645/19917 |
| down-regulated | TF | TF:M07289 | Factor: GKLF; motif: NNNRGGNGNGGSN | 8.48712553851538e-14 | 1241 | 1041 | 14779 | 19917 | A4GALT/A | 1041/1241 | 14779/19917 |
| down-regulated | TF | TF:M10432 | Factor: MAZ; motif: GGGMGGGGS | 3.13501496496215e-13 | 1241 | 722 | 9328 | 19917 | A4GALT/A | 722/1241 | 9328/19917 |
| down-regulated | TF | TF:M01857_1 | Factor: AP-2alpha; motif: NGCCYSNNGSN; match class: 1 | 8.93673694947662e-13 | 1241 | 410 | 4625 | 19917 | ABHD15/A | 410/1241 | 4625/19917 |
| down-regulated | TF | TF:M07040 | Factor: GKLF; motif: NNRRGRRNGNSNNN | 1.43513477441703e-12 | 1241 | 927 | 12809 | 19917 | A4GALT/A | 927/1241 | 12809/19917 |
| down-regulated | TF | TF:M00470_1 | Factor: AP-2gamma; motif: GCCYNNGGS; match class: 1 | 1.04796076667856e-11 | 1241 | 475 | 5637 | 19917 | A4GALT/A | 475/1241 | 5637/19917 |
| down-regulated | TF | TF:M10112 | Factor: Miz-1; motif: NNRGGWGGGGGAGGGGMRR | 1.38266159401843e-11 | 1241 | 679 | 8768 | 19917 | A4GALT/A | 679/1241 | 8768/19917 |
| down-regulated | TF | TF:M03876_1 | Factor: Kaiso; motif: GCMGGGRGCRGS; match class: 1 | 4.58668640350492e-11 | 1241 | 701 | 9160 | 19917 | A4GALT/A | 701/1241 | 9160/19917 |
| down-regulated | TF | TF:M01100 | Factor: LRF; motif: GGGGKYNNB | 6.11029784019944e-11 | 1241 | 601 | 7597 | 19917 | A4GALT/A | 601/1241 | 7597/19917 |
| down-regulated | TF | TF:M07261 | Factor: LKLF; motif: GGGGTGGKSN | 6.8304554751405e-11 | 1241 | 744 | 9867 | 19917 | A4GALT/A | 744/1241 | 9867/19917 |
| down-regulated | TF | TF:M10432_1 | Factor: MAZ; motif: GGGMGGGGS; match class: 1 | 7.36463156535762e-11 | 1241 | 385 | 4390 | 19917 | ABHD15/A | 385/1241 | 4390/19917 |
| down-regulated | TF | TF:M01973 | Factor: PLAG1; motif: CCCCCKWNNNGGSCCC | 8.44410820591936e-11 | 1241 | 509 | 6207 | 19917 | ABHD15/A | 509/1241 | 6207/19917 |
| down-regulated | TF | TF:M12351_1 | Factor: TIEG1; motif: NCCCNSNCCCCGCCCCC; match class: 1 | 4.36260308652698e-10 | 1241 | 638 | 8242 | 19917 | A4GALT/A | 638/1241 | 8242/19917 |
| down-regulated | TF | TF:M07039_1 | Factor: ETF; motif: CCCCGCCCCYN; match class: 1 | 2.49326119774735e-09 | 1241 | 965 | 13735 | 19917 | A4GALT/A | 965/1241 | 13735/19917 |
| down-regulated | TF | TF:M09826_1 | Factor: BTEB3; motif: CCNNSCCNSCCCCKCCCCC; match class: 1 | 2.50614183102805e-09 | 1241 | 590 | 7557 | 19917 | A4GALT/A | 590/1241 | 7557/19917 |
| down-regulated | TF | TF:M09636_1 | Factor: MAZ; motif: GGGMGGGGSSGGGGGGGGGGGG; match class: 1 | 2.83639926604788e-09 | 1241 | 991 | 14202 | 19917 | A4GALT/A | 991/1241 | 14202/19917 |
| down-regulated | TF | TF:M09984_1 | Factor: MAZ; motif: GGGGGAGGGGGNGRGRRRGNRG; match class: 1 | 3.53350397732283e-09 | 1241 | 460 | 5599 | 19917 | ABHD15/A | 460/1241 | 5599/19917 |
| down-regulated | TF | TF:M08867_1 | Factor: AP2; motif: GCCYGSGGSN; match class: 1 | 3.65792293760414e-09 | 1241 | 433 | 5202 | 19917 | A4GALT/A | 433/1241 | 5202/19917 |
| down-regulated | TF | TF:M00915 | Factor: AP-2; motif: SNNNCCNCAGGCN | 4.64287855187496e-09 | 1241 | 723 | 9683 | 19917 | A4GALT/A | 723/1241 | 9683/19917 |
| down-regulated | TF | TF:M04595_1 | Factor: SALL2; motif: GGGTGGG; match class: 1 | 5.06385158507417e-09 | 1241 | 580 | 7428 | 19917 | A4GALT/A | 580/1241 | 7428/19917 |
| down-regulated | TF | TF:M01104_1 | Factor: MOVO-B; motif: GNGGGGG; match class: 1 | 6.6808359072459e-09 | 1241 | 462 | 5650 | 19917 | ABHD15/A | 462/1241 | 5650/19917 |
| down-regulated | TF | TF:M07436 | Factor: WT1; motif: NNGGGNGGGSGN | 8.61031323014229e-09 | 1241 | 500 | 6227 | 19917 | A4GALT/A | 500/1241 | 6227/19917 |
| down-regulated | TF | TF:M08819 | Factor: LKLF; motif: CNCCACCCS | 8.93795279773581e-09 | 1241 | 461 | 5645 | 19917 | ABHD15/A | 461/1241 | 5645/19917 |
| down-regulated | TF | TF:M00470 | Factor: AP-2gamma; motif: GCCYNNGGS | 1.05368472338808e-08 | 1241 | 708 | 9473 | 19917 | A4GALT/A | 708/1241 | 9473/19917 |
| down-regulated | TF | TF:M12351 | Factor: TIEG1; motif: NCCCNSNCCCCGCCCCC | 1.10257862476485e-08 | 1241 | 885 | 12403 | 19917 | A4GALT/A | 885/1241 | 12403/19917 |
| down-regulated | TF | TF:M12354_1 | Factor: ZNF37A; motif: CCYYGGCTCCNTSCCMN; match class: 1 | 1.50834873011053e-08 | 1241 | 460 | 5648 | 19917 | ABHD15/A | 460/1241 | 5648/19917 |
| down-regulated | TF | TF:M00800_1 | Factor: AP-2; motif: GSCCSCRGGCNRNRNN; match class: 1 | 1.65312631377062e-08 | 1241 | 366 | 4278 | 19917 | A4GALT/A | 366/1241 | 4278/19917 |
| down-regulated | TF | TF:M07461 | Factor: KLF; motif: GGGNGGGG | 2.21837121950002e-08 | 1241 | 666 | 8831 | 19917 | A4GALT/A | 666/1241 | 8831/19917 |
| down-regulated | TF | TF:M07261_1 | Factor: LKLF; motif: GGGGTGGKSN; match class: 1 | 2.96006273004687e-08 | 1241 | 353 | 4110 | 19917 | A4GALT/A | 353/1241 | 4110/19917 |
| down-regulated | TF | TF:M00800 | Factor: AP-2; motif: GSCCSCRGGCNRNRNN | 3.91286311894947e-08 | 1241 | 686 | 9172 | 19917 | A4GALT/A | 686/1241 | 9172/19917 |
| down-regulated | TF | TF:M01835_1 | Factor: GKLF; motif: CCTCCYN; match class: 1 | 4.93411854008705e-08 | 1241 | 750 | 10214 | 19917 | A2ML1/A4 | 750/1241 | 10214/19917 |
| down-regulated | TF | TF:M09727 | Factor: GEMIN3; motif: NCWGGRARRGRGNGNG | 5.78028734501607e-08 | 1241 | 651 | 8631 | 19917 | ABHD15/A | 651/1241 | 8631/19917 |
| down-regulated | TF | TF:M12354 | Factor: ZNF37A; motif: CCYYGGCTCCNTSCCMN | 7.06812694055355e-08 | 1241 | 799 | 11033 | 19917 | A4GALT/A | 799/1241 | 11033/19917 |
| down-regulated | TF | TF:M01721 | Factor: PUR1; motif: GGGNCAGNN | 1.16636555204521e-07 | 1241 | 685 | 9200 | 19917 | A2ML1/A4 | 685/1241 | 9200/19917 |
| down-regulated | TF | TF:M01858 | Factor: AP-2beta; motif: GCNNNGGSCNGVGGGN | 1.36791985971696e-07 | 1241 | 574 | 7460 | 19917 | A4GALT/A | 574/1241 | 7460/19917 |
| down-regulated | TF | TF:M01175 | Factor: CKROX; motif: SCCCTCCCC | 2.14702854065076e-07 | 1241 | 566 | 7354 | 19917 | A4GALT/A | 566/1241 | 7354/19917 |
| down-regulated | TF | TF:M00469 | Factor: AP-2alpha; motif: GCCNNNRGS | 2.43523328772694e-07 | 1241 | 422 | 5182 | 19917 | A4GALT/A | 422/1241 | 5182/19917 |
| down-regulated | TF | TF:M01865 | Factor: BTEB3; motif: BNRNGGGAGGNGT | 2.75517554170085e-07 | 1241 | 731 | 9976 | 19917 | A2ML1/AB | 731/1241 | 9976/19917 |
| down-regulated | TF | TF:M01973_1 | Factor: PLAG1; motif: CCCCCKWNNNGGSCCC; match class: 1 | 3.01624462705112e-07 | 1241 | 226 | 2413 | 19917 | ABHD15/A | 226/1241 | 2413/19917 |
| down-regulated | TF | TF:M09729 | Factor: LUMAN; motif: CYCAGCYYCY | 4.70056591776782e-07 | 1241 | 779 | 10782 | 19917 | A2ML1/A4 | 779/1241 | 10782/19917 |
| down-regulated | TF | TF:M00986 | Factor: Churchill; motif: CGGGNN | 4.76312847367892e-07 | 1241 | 986 | 14305 | 19917 | A4GALT/A | 986/1241 | 14305/19917 |
| down-regulated | TF | TF:M00469_1 | Factor: AP-2alpha; motif: GCCNNNRGS; match class: 1 | 5.48202316743068e-07 | 1241 | 293 | 3350 | 19917 | A4GALT/A | 293/1241 | 3350/19917 |
| down-regulated | TF | TF:M12160 | Factor: KLF15; motif: RCCMCRCCCMCN | 5.93700092588438e-07 | 1241 | 901 | 12836 | 19917 | A4GALT/A | 901/1241 | 12836/19917 |
| down-regulated | TF | TF:M07461_1 | Factor: KLF; motif: GGGNGGGG; match class: 1 | 6.01455070786696e-07 | 1241 | 298 | 3423 | 19917 | A4GALT/A | 298/1241 | 3423/19917 |
| down-regulated | TF | TF:M03876 | Factor: Kaiso; motif: GCMGGGRGCRGS | 6.41133810223489e-07 | 1241 | 953 | 13737 | 19917 | A4GALT/A | 953/1241 | 13737/19917 |
| down-regulated | TF | TF:M00189_1 | Factor: AP-2; motif: MKCCCSCNGGCG; match class: 1 | 6.48251355173237e-07 | 1241 | 482 | 6114 | 19917 | A4GALT/A | 482/1241 | 6114/19917 |
| down-regulated | TF | TF:M09826 | Factor: BTEB3; motif: CCNNSCCNSCCCCKCCCCC | 6.94364672749162e-07 | 1241 | 825 | 11560 | 19917 | A4GALT/A | 825/1241 | 11560/19917 |
| down-regulated | TF | TF:M11478 | Factor: AP-2beta; motif: NSCCNNNGGSN | 7.77782397996285e-07 | 1241 | 731 | 10020 | 19917 | A4GALT/A | 731/1241 | 10020/19917 |
| down-regulated | TF | TF:M10026 | Factor: PATZ; motif: GGGGNGGGGGMKGGRRNGGNRN | 8.35049048005471e-07 | 1241 | 632 | 8440 | 19917 | A4GALT/A | 632/1241 | 8440/19917 |
| down-regulated | TF | TF:M04953 | Factor: Sp1; motif: GGNDGGRGGCGGGG | 8.68812138323591e-07 | 1241 | 652 | 8758 | 19917 | ABHD15/A | 652/1241 | 8758/19917 |
| down-regulated | TF | TF:M01721_1 | Factor: PUR1; motif: GGGNCAGNN; match class: 1 | 9.80858217024217e-07 | 1241 | 298 | 3438 | 19917 | A4GALT/A | 298/1241 | 3438/19917 |
| down-regulated | TF | TF:M09636 | Factor: MAZ; motif: GGGMGGGGSSGGGGGGGGGGGG | 1.06047012718167e-06 | 1241 | 1098 | 16380 | 19917 | A4GALT/A | 1098/1241 | 16380/19917 |
| down-regulated | TF | TF:M08867 | Factor: AP2; motif: GCCYGSGGSN | 1.74814004517705e-06 | 1241 | 735 | 10120 | 19917 | A4GALT/A | 735/1241 | 10120/19917 |
| down-regulated | TF | TF:M02023 | Factor: MAZ; motif: NKGGGAGGGGRGGR | 2.06398301529394e-06 | 1241 | 549 | 7183 | 19917 | A4GALT/A | 549/1241 | 7183/19917 |
| down-regulated | TF | TF:M00915_1 | Factor: AP-2; motif: SNNNCCNCAGGCN; match class: 1 | 2.32119487180211e-06 | 1241 | 375 | 4572 | 19917 | A4GALT/A | 375/1241 | 4572/19917 |
| down-regulated | TF | TF:M01303 | Factor: SP1; motif: GGGGYGGGGNS | 2.39880034618656e-06 | 1241 | 597 | 7935 | 19917 | A4GALT/A | 597/1241 | 7935/19917 |
| down-regulated | TF | TF:M00706 | Factor: TFII-I; motif: RGAGGKAGG | 2.72052190134069e-06 | 1241 | 454 | 5748 | 19917 | A2ML1/A4 | 454/1241 | 5748/19917 |
| down-regulated | TF | TF:M07039 | Factor: ETF; motif: CCCCGCCCCYN | 2.73062201739603e-06 | 1241 | 1104 | 16527 | 19917 | A4GALT/A | 1104/1241 | 16527/19917 |
| down-regulated | TF | TF:M10108 | Factor: WT1; motif: RGGNGGGGGAGGRGGNGGRG | 3.03687061848301e-06 | 1241 | 509 | 6585 | 19917 | ACSF2/ACS | 509/1241 | 6585/19917 |
| down-regulated | TF | TF:M01858_1 | Factor: AP-2beta; motif: GCNNNGGSCNGVGGGN; match class: 1 | 3.25479009447265e-06 | 1241 | 275 | 3152 | 19917 | ABHD15/A | 275/1241 | 3152/19917 |
| down-regulated | TF | TF:M00649 | Factor: MAZ; motif: GGGGAGGG | 3.38486557816056e-06 | 1241 | 689 | 9407 | 19917 | A4GALT/A | 689/1241 | 9407/19917 |
| down-regulated | TF | TF:M00444_1 | Factor: VDR; motif: GGGKNARNRRGGWSA; match class: 1 | 3.55032841899933e-06 | 1241 | 373 | 4558 | 19917 | A4GALT/A | 373/1241 | 4558/19917 |
| down-regulated | TF | TF:M09973 | Factor: CPBP; motif: GNNRGGGHGGGGNNGGGRN | 3.76937430913931e-06 | 1241 | 782 | 10922 | 19917 | A4GALT/A | 782/1241 | 10922/19917 |
| down-regulated | TF | TF:M09729_1 | Factor: LUMAN; motif: CYCAGCYYCY; match class: 1 | 4.02798915598417e-06 | 1241 | 380 | 4665 | 19917 | A4GALT/A | 380/1241 | 4665/19917 |
| down-regulated | TF | TF:M05327 | Factor: WT1; motif: NGCGGGGGGGTSMMCYN | 4.51528486119276e-06 | 1241 | 439 | 5543 | 19917 | ACAA2/AC | 439/1241 | 5543/19917 |
| down-regulated | TF | TF:M01118 | Factor: WT1; motif: SMCNCCNSC | 5.38832837175969e-06 | 1241 | 526 | 6869 | 19917 | ABHD15/A | 526/1241 | 6869/19917 |
| down-regulated | TF | TF:M11478_1 | Factor: AP-2beta; motif: NSCCNNNGGSN; match class: 1 | 6.28697122981078e-06 | 1241 | 605 | 8102 | 19917 | A4GALT/A | 605/1241 | 8102/19917 |
| down-regulated | TF | TF:M07348 | Factor: AP-2alpha; motif: NSCCNCRGGSN | 6.57978891104923e-06 | 1241 | 605 | 8104 | 19917 | A4GALT/A | 605/1241 | 8104/19917 |
| down-regulated | TF | TF:M09862_1 | Factor: ZNF614; motif: NCYCWGCCYYNNN; match class: 1 | 8.80779321644583e-06 | 1241 | 347 | 4212 | 19917 | A4GALT/A | 347/1241 | 4212/19917 |
| down-regulated | TF | TF:M04595 | Factor: SALL2; motif: GGGTGGG | 8.88176039612927e-06 | 1241 | 911 | 13123 | 19917 | A4GALT/A | 911/1241 | 13123/19917 |
| down-regulated | TF | TF:M09973_1 | Factor: CPBP; motif: GNNRGGGHGGGGNNGGGRN; match class: 1 | 1.37497877965238e-05 | 1241 | 509 | 6648 | 19917 | A4GALT/A | 509/1241 | 6648/19917 |
| down-regulated | TF | TF:M00749_1 | Factor: SREBP-1; motif: CACSCCA; match class: 1 | 1.4751435283687e-05 | 1241 | 246 | 2794 | 19917 | A4GALT/A | 246/1241 | 2794/19917 |
| down-regulated | TF | TF:M04148 | Factor: TFAP2A; motif: NGCCCYNNGGGCN | 1.66253071025803e-05 | 1241 | 728 | 10108 | 19917 | A4GALT/A | 728/1241 | 10108/19917 |
| down-regulated | TF | TF:M07040_1 | Factor: GKLF; motif: NNRRGRRNGNSNNN; match class: 1 | 1.70426879037806e-05 | 1241 | 610 | 8225 | 19917 | A4GALT/A | 610/1241 | 8225/19917 |
| down-regulated | TF | TF:M04863_1 | Factor: TF3C-beta; motif: CCNGGAGGGCTTCCTGGAGGAG; match clas | 1.72429405488252e-05 | 1241 | 635 | 8620 | 19917 | A4GALT/A | 635/1241 | 8620/19917 |
| down-regulated | TF | TF:M10529 | Factor: Sp1; motif: RGGGMGGRGSNGGGG | 2.39620162193852e-05 | 1241 | 526 | 6933 | 19917 | ABHD15/A | 526/1241 | 6933/19917 |
| down-regulated | TF | TF:M10026_1 | Factor: PATZ; motif: GGGGNGGGGGMKGGRRNGGNRN; match class: | 2.40985159554527e-05 | 1241 | 393 | 4924 | 19917 | A4GALT/A | 393/1241 | 4924/19917 |
| down-regulated | TF | TF:M04148_1 | Factor: TFAP2A; motif: NGCCCYNNGGGCN; match class: 1 | 2.92238528424679e-05 | 1241 | 699 | 9666 | 19917 | A4GALT/A | 699/1241 | 9666/19917 |
| down-regulated | TF | TF:M00444 | Factor: VDR; motif: GGGKNARNRRGGWSA | 3.29405918569493e-05 | 1241 | 744 | 10400 | 19917 | A2ML1/A4 | 744/1241 | 10400/19917 |
| down-regulated | TF | TF:M01104 | Factor: MOVO-B; motif: GNGGGGG | 3.37007536526046e-05 | 1241 | 740 | 10336 | 19917 | A4GALT/A | 740/1241 | 10336/19917 |
| down-regulated | TF | TF:M01303_1 | Factor: SP1; motif: GGGGYGGGGNS; match class: 1 | 3.38073721853868e-05 | 1241 | 296 | 3525 | 19917 | A4GALT/A | 296/1241 | 3525/19917 |
| down-regulated | TF | TF:M09734 | Factor: ZNF692; motif: SYNGGSCCCASCCNC | 3.68488677414954e-05 | 1241 | 828 | 11788 | 19917 | ABHD15/A | 828/1241 | 11788/19917 |
| down-regulated | TF | TF:M04896 | Factor: AP-2alpha; motif: NNNGSCCTGRGGSN | 3.84208035294512e-05 | 1241 | 451 | 5810 | 19917 | A4GALT/A | 451/1241 | 5810/19917 |
| down-regulated | TF | TF:M07397 | Factor: ZBP89; motif: CCCCKCCCCCNN | 4.25320573998366e-05 | 1241 | 541 | 7190 | 19917 | ABHD15/A | 541/1241 | 7190/19917 |
| down-regulated | TF | TF:M00189 | Factor: AP-2; motif: MKCCCSCNGGCG | 5.67619322395499e-05 | 1241 | 760 | 10687 | 19917 | A4GALT/A | 760/1241 | 10687/19917 |
| down-regulated | TF | TF:M08952 | Factor: NF-KAPPAB1; motif: NGGKRNTTYCCCN | 5.83426191449954e-05 | 1241 | 580 | 7811 | 19917 | A4GALT/A | 580/1241 | 7811/19917 |
| down-regulated | TF | TF:M09723 | Factor: BTEB1; motif: GGGGGCGGGGCNGSGGGNGS | 5.95185156746119e-05 | 1241 | 731 | 10217 | 19917 | A4GALT/A | 731/1241 | 10217/19917 |
| down-regulated | TF | TF:M09972 | Factor: BTEB2; motif: WGGGTGKGGCNGGN | 5.99840889522133e-05 | 1241 | 673 | 9283 | 19917 | A4GALT/A | 673/1241 | 9283/19917 |
| down-regulated | TF | TF:M09890_1 | Factor: ctcf; motif: YGGCCACCAGRKGGCRSYN; match class: 1 | 6.09995501230977e-05 | 1241 | 250 | 2894 | 19917 | A4GALT/A | 250/1241 | 2894/19917 |
| down-regulated | TF | TF:M10107 | Factor: DB1; motif: GGRRRRGRRGGAGGGGGNGRRR | 6.10707584874767e-05 | 1241 | 454 | 5875 | 19917 | ACAA2/AC | 454/1241 | 5875/19917 |
| down-regulated | TF | TF:M09762_1 | Factor: NF1B; motif: CYTGGCNYNCWGCCAN; match class: 1 | 7.02594307389246e-05 | 1241 | 149 | 1529 | 19917 | ACAA2/AC | 149/1241 | 1529/19917 |
| down-regulated | TF | TF:M10002 | Factor: NF-1C; motif: NYTGGCNNYNNGCCARN | 7.41777986283252e-05 | 1241 | 515 | 6814 | 19917 | ACAA2/AC | 515/1241 | 6814/19917 |
| down-regulated | TF | TF:M10072 | Factor: sp4; motif: NNGNARGRGGCGGRGCNNRR | 7.68075627090313e-05 | 1241 | 752 | 10571 | 19917 | A4GALT/A | 752/1241 | 10571/19917 |
| down-regulated | TF | TF:M04453_1 | Factor: AR; motif: RGGWACAYNGTGTWCYN; match class: 1 | 8.09865771207057e-05 | 1241 | 315 | 3829 | 19917 | ACACB/AD | 315/1241 | 3829/19917 |

| | | | | | | | | | | | |
|---|---|---|---|---|---|---|---|---|---|---|---|
| down-regulated | TF | TF:M09910_1 | Factor: ER-beta; motif: RGGTCASCNTGMCCY; match class: 1 | 8.87537623855226e-05 | 1241 | 334 | 4108 | 19917 | A4GALT/AI | 334/1241 | 4108/19917 |
| down-regulated | TF | TF:M09658 | Factor: Sp2; motif: GGSNNGGGGGCGGGGCCNGNGS | 0.000106852381332629 | 1241 | 414 | 5297 | 19917 | ACSF2/ACS | 414/1241 | 5297/19917 |
| down-regulated | TF | TF:M09734_1 | Factor: ZNF692; motif: SYNGGSCCCASCCNC; match class: 1 | 0.000108524660436163 | 1241 | 480 | 6295 | 19917 | ABHD15/A | 480/1241 | 6295/19917 |
| down-regulated | TF | TF:M02089_1 | Factor: E2F-3; motif: GGCGGGN; match class: 1 | 0.000111354899360942 | 1241 | 680 | 9425 | 19917 | A4GALT/AI | 680/1241 | 9425/19917 |
| down-regulated | TF | TF:M07329 | Factor: Osx; motif: CCNCCCCCNNN | 0.000119259811251944 | 1241 | 516 | 6851 | 19917 | ACVRL1/A( | 516/1241 | 6851/19917 |
| down-regulated | TF | TF:M02036_1 | Factor: WT1; motif: CGCCCCCNCN; match class: 1 | 0.000138624290099743 | 1241 | 414 | 5308 | 19917 | ABHD15/A | 414/1241 | 5308/19917 |
| down-regulated | TF | TF:M10112_1 | Factor: Miz-1; motif: NNRGGWGGGGGAGGGGMRR; match class: 1 | 0.000139089706651235 | 1241 | 332 | 4096 | 19917 | A4GALT/A( | 332/1241 | 4096/19917 |
| down-regulated | TF | TF:M10134 | Factor: FPM315; motif: GGGAGGAGGRRGRGGRGGRR | 0.000142289543458212 | 1241 | 221 | 2517 | 19917 | ABHD15/A | 221/1241 | 2517/19917 |
| down-regulated | TF | TF:M09723_1 | Factor: BTEB1; motif: GGGGGCGGGGCNGSGGGNGS; match class: 1 | 0.000150395092834405 | 1241 | 469 | 6142 | 19917 | A4GALT/AI | 469/1241 | 6142/19917 |
| down-regulated | TF | TF:M01587 | Factor: FPM315; motif: SRGGGAGGAGGN | 0.000157595098039112 | 1241 | 275 | 3279 | 19917 | ABHD15/A | 275/1241 | 3279/19917 |
| down-regulated | TF | TF:M01778 | Factor: PLAG1; motif: GRGGCNNHNNNRRGGG | 0.000179835496990828 | 1241 | 238 | 2761 | 19917 | ABHD15/A | 238/1241 | 2761/19917 |
| down-regulated | TF | TF:M01201 | Factor: AR; motif: GGNACRNNRTGTWCT | 0.000193288142896651 | 1241 | 487 | 6428 | 19917 | ACAA2/AC. | 487/1241 | 6428/19917 |
| down-regulated | TF | TF:M01733 | Factor: MZF-1; motif: TGGGGAR | 0.000200267518343445 | 1241 | 804 | 11472 | 19917 | A2ML1/A4 | 804/1241 | 11472/19917 |
| down-regulated | TF | TF:M10426_1 | Factor: ctcf; motif: CCRSCAGGGGGCGCN; match class: 1 | 0.000208721573166799 | 1241 | 351 | 4390 | 19917 | A4GALT/AI | 351/1241 | 4390/19917 |
| down-regulated | TF | TF:M11482 | Factor: AP-2gamma; motif: NSCCYNNRGSN | 0.000211690435434022 | 1241 | 507 | 6739 | 19917 | A4GALT/AI | 507/1241 | 6739/19917 |
| down-regulated | TF | TF:M12158 | Factor: KLF15; motif: NCCMCGCCCMCN | 0.000226821489432962 | 1241 | 785 | 11165 | 19917 | A4GALT/AI | 785/1241 | 11165/19917 |
| down-regulated | TF | TF:M12227 | Factor: ZIC4; motif: NNCCNCCCRYNGYGN | 0.000238131842758113 | 1241 | 784 | 11151 | 19917 | A2ML1/A4 | 784/1241 | 11151/19917 |
| down-regulated | TF | TF:M07277 | Factor: BTEB2; motif: RGGGNGKGGN | 0.000238718513781272 | 1241 | 622 | 8539 | 19917 | A4GALT/AI | 622/1241 | 8539/19917 |
| down-regulated | TF | TF:M00378_1 | Factor: Pax-4; motif: NNNNNYCACCCB; match class: 1 | 0.000242647681505219 | 1241 | 1002 | 14850 | 19917 | A4GALT/AI | 1002/1241 | 14850/19917 |
| down-regulated | TF | TF:M09862 | Factor: ZNF614; motif: NCYCWGCCYYNNN | 0.000245190294784014 | 1241 | 678 | 9432 | 19917 | A4GALT/AI | 678/1241 | 9432/19917 |
| down-regulated | TF | TF:M09834 | Factor: ZNF148; motif: NNNNNNCCNNCCCCTCCCCCACCCN | 0.000275292350655174 | 1241 | 523 | 6998 | 19917 | ABHD15/A | 523/1241 | 6998/19917 |
| down-regulated | TF | TF:M09744 | Factor: CTCF; motif: NNYGCCCYCTRSTGGN | 0.000340108243916734 | 1241 | 313 | 3854 | 19917 | A2ML1/A4 | 313/1241 | 3854/19917 |
| down-regulated | TF | TF:M12160_1 | Factor: KLF15; motif: RCCMCRCCCMCN; match class: 1 | 0.000350482161172904 | 1241 | 592 | 8085 | 19917 | A4GALT/AI | 592/1241 | 8085/19917 |
| down-regulated | TF | TF:M01047 | Factor: AP-2alphaA; motif: ANMGCCTNAGGCKNT | 0.000351255192275741 | 1241 | 669 | 9306 | 19917 | A4GALT/AI | 669/1241 | 9306/19917 |
| down-regulated | TF | TF:M01047_1 | Factor: AP-2alphaA; motif: ANMGCCTNAGGCKNT; match class: 1 | 0.000351255192275741 | 1241 | 669 | 9306 | 19917 | A4GALT/AI | 669/1241 | 9306/19917 |
| down-regulated | TF | TF:M02036 | Factor: WT1; motif: CGCCCCCNCN | 0.000352604810618356 | 1241 | 701 | 9820 | 19917 | ABHD15/A | 701/1241 | 9820/19917 |
| down-regulated | TF | TF:M00932_1 | Factor: Sp1; motif: NNGGGGCGGGGNN; match class: 1 | 0.000367847769048045 | 1241 | 462 | 6076 | 19917 | A4GALT/AI | 462/1241 | 6076/19917 |
| down-regulated | TF | TF:M00051 | Factor: NF-kappaB; motif: GGGGATYCCC | 0.000369640274052658 | 1241 | 487 | 6458 | 19917 | A4GALT/AI | 487/1241 | 6458/19917 |
| down-regulated | TF | TF:M09760_1 | Factor: DPF2; motif: NYCACYTCCYCNYYCY; match class: 1 | 0.000375156573380305 | 1241 | 254 | 3011 | 19917 | ACAA2/AC' | 254/1241 | 3011/19917 |
| down-regulated | TF | TF:M01733_1 | Factor: MZF-1; motif: TGGGGAR; match class: 1 | 0.000402505758590237 | 1241 | 388 | 4965 | 19917 | ABHD15/A | 388/1241 | 4965/19917 |
| down-regulated | TF | TF:M10071_1 | Factor: Sp1; motif: NGGGGGCGGGGCCNGGGGGGGG; match class: 1 | 0.000430909560752517 | 1241 | 366 | 4641 | 19917 | A4GALT/AI | 366/1241 | 4641/19917 |
| down-regulated | TF | TF:M00933 | Factor: Sp1; motif: CCCCGCCCCN | 0.000486221718179899 | 1241 | 697 | 9772 | 19917 | A4GALT/AI | 697/1241 | 9772/19917 |
| down-regulated | TF | TF:M01072 | Factor: HIC1; motif: NSNNNNTGCCCSSNN | 0.000496421350799838 | 1241 | 335 | 4190 | 19917 | A4GALT/AI | 335/1241 | 4190/19917 |
| down-regulated | TF | TF:M09907_1 | Factor: Erg; motif: NRRSAGGAAGNGG; match class: 1 | 0.000550756370385112 | 1241 | 398 | 5128 | 19917 | ACOX2/AC' | 398/1241 | 5128/19917 |
| down-regulated | TF | TF:M01837 | Factor: FKLF; motif: BGGGNGGVMD | 0.000557067895097191 | 1241 | 422 | 5489 | 19917 | ABHD15/A | 422/1241 | 5489/19917 |
| down-regulated | TF | TF:M00933_1 | Factor: Sp1; motif: CCCCGCCCCN; match class: 1 | 0.000590699805614477 | 1241 | 402 | 5191 | 19917 | A4GALT/A( | 402/1241 | 5191/19917 |
| down-regulated | TF | TF:M02089 | Factor: E2F-3; motif: GGCGGGN | 0.000622696649893048 | 1241 | 908 | 13271 | 19917 | A4GALT/AI | 908/1241 | 13271/19917 |
| down-regulated | TF | TF:M00982_1 | Factor: KROX; motif: CCCGCCCCCRCCCC; match class: 1 | 0.000642640353409619 | 1241 | 289 | 3531 | 19917 | ABHD15/A | 289/1241 | 3531/19917 |
| down-regulated | TF | TF:M00450_1 | Factor: Zic3; motif: NGGGKGGTC; match class: 1 | 0.000660213733580268 | 1241 | 584 | 7991 | 19917 | A4GALT/A( | 584/1241 | 7991/19917 |
| down-regulated | TF | TF:M00749 | Factor: SREBP-1; motif: CACSCCA | 0.000666772132070848 | 1241 | 626 | 8654 | 19917 | A4GALT/A( | 626/1241 | 8654/19917 |
| down-regulated | TF | TF:M00986_1 | Factor: Churchill; motif: CGGGNN; match class: 1 | 0.00080376087606272 | 1241 | 736 | 10429 | 19917 | A4GALT/AI | 736/1241 | 10429/19917 |
| down-regulated | TF | TF:M07249_1 | Factor: ctcf; motif: CCNCNAGRKGGCRSTN; match class: 1 | 0.000897104665407214 | 1241 | 185 | 2078 | 19917 | A4GALT/AI | 185/1241 | 2078/19917 |
| down-regulated | TF | TF:M05444_1 | Factor: CPBP; motif: NGGGCGG; match class: 1 | 0.000919191418840031 | 1241 | 275 | 3343 | 19917 | ACVRL1/A( | 275/1241 | 3343/19917 |
| down-regulated | TF | TF:M05332_1 | Factor: Sp2; motif: WGGGCGG; match class: 1 | 0.000919191418840031 | 1241 | 275 | 3343 | 19917 | ACVRL1/A( | 275/1241 | 3343/19917 |
| down-regulated | TF | TF:M05361_1 | Factor: Sp6; motif: WGGGCGG; match class: 1 | 0.000919191418840031 | 1241 | 275 | 3343 | 19917 | ACVRL1/A( | 275/1241 | 3343/19917 |
| down-regulated | TF | TF:M11482_1 | Factor: AP-2gamma; motif: NSCCYNNRGSN; match class: 1 | 0.00094832591471259 | 1241 | 184 | 2066 | 19917 | ACAA2/AD | 184/1241 | 2066/19917 |
| down-regulated | TF | TF:M00481_1 | Factor: AR; motif: GGTACANNRTGTTCT; match class: 1 | 0.000960607053893955 | 1241 | 202 | 2314 | 19917 | ABHD15/A | 202/1241 | 2314/19917 |
| down-regulated | TF | TF:M04617 | Factor: LRF; motif: NGNAGNGGGTYN | 0.00125340001527837 | 1241 | 467 | 6210 | 19917 | ABCB1/ABI | 467/1241 | 6210/19917 |
| down-regulated | TF | TF:M07354_1 | Factor: Egr-1; motif: GCGGGGGCGG; match class: 1 | 0.0013114936650109 | 1241 | 290 | 3573 | 19917 | ACBD7/AD. | 290/1241 | 3573/19917 |
| down-regulated | TF | TF:M08911 | Factor: CTCF; motif: NCCRSTAGGGGGCGC | 0.00135048916152598 | 1241 | 656 | 9168 | 19917 | A4GALT/AI | 656/1241 | 9168/19917 |
| down-regulated | TF | TF:M12227_1 | Factor: ZIC4; motif: NNCCNCCCRYNGYGN; match class: 1 | 0.0013843478812859 | 1241 | 448 | 5925 | 19917 | A4GALT/AI | 448/1241 | 5925/19917 |
| down-regulated | TF | TF:M09727_1 | Factor: GEMIN3; motif: NCWGGRARRGRGNGNG; match class: 1 | 0.00141280802748408 | 1241 | 266 | 3230 | 19917 | ABHD15/A | 266/1241 | 3230/19917 |
| down-regulated | TF | TF:M04476_1 | Factor: NR3C1; motif: NRGWACAYNRTGTWCYN; match class: 1 | 0.0015767258351721 | 1241 | 414 | 5416 | 19917 | ABCC9/ABI | 414/1241 | 5416/19917 |
| down-regulated | TF | TF:M01224 | Factor: P50:RELA-P65; motif: GGANTTYCCCWN | 0.00163922603290865 | 1241 | 204 | 2359 | 19917 | ACKR3/AC! | 204/1241 | 2359/19917 |
| down-regulated | TF | TF:M00033 | Factor: p300; motif: NNNGGGAGTNNNNS | 0.00169748190226492 | 1241 | 624 | 8671 | 19917 | ABHD15/A | 624/1241 | 8671/19917 |
| down-regulated | TF | TF:M00774 | Factor: NF-kappaB; motif: NGGGANTTYCCMNNNN | 0.00183408741899489 | 1241 | 165 | 1827 | 19917 | ABCB1/AC\ | 165/1241 | 1827/19917 |
| down-regulated | TF | TF:M07348_1 | Factor: AP-2alpha; motif: NSCCNCRGGSN; match class: 1 | 0.00199274681191867 | 1241 | 246 | 2957 | 19917 | A4GALT/AI | 246/1241 | 2957/19917 |
| down-regulated | TF | TF:M12313 | Factor: ZNF460; motif: NNACNCCCCCCNN | 0.00247847856965135 | 1241 | 443 | 5877 | 19917 | ABHD15/A | 443/1241 | 5877/19917 |
| down-regulated | TF | TF:M07226 | Factor: SP1; motif: NCCCCKCCCCC | 0.00275754690152366 | 1241 | 601 | 8333 | 19917 | A4GALT/A( | 601/1241 | 8333/19917 |
| down-regulated | TF | TF:M04855 | Factor: IRF-4; motif: AAGTTTC | 0.00276229339379139 | 1241 | 565 | 7767 | 19917 | A4GALT/AI | 565/1241 | 7767/19917 |
| down-regulated | TF | TF:M08952_1 | Factor: NF-KAPPAB1; motif: NGGKRNTTYCCCN; match class: 1 | 0.00282116185503228 | 1241 | 281 | 3473 | 19917 | ACSS1/AD/ | 281/1241 | 3473/19917 |
| down-regulated | TF | TF:M01200 | Factor: CTCF; motif: NNNGCCASCAGRKGGCRSNN | 0.00320787147610729 | 1241 | 315 | 3974 | 19917 | A2ML1/AB | 315/1241 | 3974/19917 |
| down-regulated | TF | TF:M09972_1 | Factor: BTEB2; motif: WGGGTGKGGCNGGN; match class: 1 | 0.00338959037420747 | 1241 | 313 | 3947 | 19917 | A4GALT/A( | 313/1241 | 3947/19917 |
| down-regulated | TF | TF:M00051_1 | Factor: NF-kappaB; motif: GGGGATYCCC; match class: 1 | 0.00349748184876144 | 1241 | 234 | 2807 | 19917 | ADGRG1/A | 234/1241 | 2807/19917 |
| down-regulated | TF | TF:M03811 | Factor: AP-2gamma; motif: GCCYNCRGSN | 0.00366473509179797 | 1241 | 637 | 8919 | 19917 | ABCB1/ABI | 637/1241 | 8919/19917 |
| down-regulated | TF | TF:M03893_1 | Factor: WT1; motif: GNGGGGGCGGGG; match class: 1 | 0.00389735865822118 | 1241 | 316 | 3997 | 19917 | ACSF2/ACS | 316/1241 | 3997/19917 |
| down-regulated | TF | TF:M09739 | Factor: ZGPAT; motif: GRGGCWGNGGNG | 0.00398654733535439 | 1241 | 596 | 8274 | 19917 | A4GALT/AI | 596/1241 | 8274/19917 |
| down-regulated | TF | TF:M11664 | Factor: IRF-2; motif: NGAAASYGAAAS | 0.00407884892075226 | 1241 | 151 | 1661 | 19917 | ABCB1/AC! | 151/1241 | 1661/19917 |
| down-regulated | TF | TF:M09725 | Factor: DREF; motif: CTYYCWCTTCCY | 0.0042089734804855 | 1241 | 552 | 7586 | 19917 | ABCB1/AB( | 552/1241 | 7586/19917 |
| down-regulated | TF | TF:M12313_1 | Factor: ZNF460; motif: NNACNCCCCCCNN; match class: 1 | 0.00442959963963267 | 1241 | 192 | 2225 | 19917 | ADAMTS9/ | 192/1241 | 2225/19917 |
| down-regulated | TF | TF:M01199 | Factor: RNF96; motif: BCCCGCRGCC | 0.00462592283924981 | 1241 | 612 | 8535 | 19917 | A4GALT/AI | 612/1241 | 8535/19917 |
| down-regulated | TF | TF:M07277_1 | Factor: BTEB2; motif: RGGGNGKGGN; match class: 1 | 0.00490527469986318 | 1241 | 327 | 4169 | 19917 | ACVRL1/A( | 327/1241 | 4169/19917 |
| down-regulated | TF | TF:M09828 | Factor: CTCF; motif: RSYGCCMYCTRSTGGN | 0.00493042472109885 | 1241 | 310 | 3919 | 19917 | A2ML1/AB | 310/1241 | 3919/19917 |
| down-regulated | TF | TF:M10426 | Factor: ctcf; motif: CCRSCAGGGGGCGCN | 0.00555615829070397 | 1241 | 665 | 9389 | 19917 | A2ML1/A4 | 665/1241 | 9389/19917 |
| down-regulated | TF | TF:M00931_1 | Factor: Sp1; motif: GGGGCGGGGC; match class: 1 | 0.00575044468349205 | 1241 | 444 | 5934 | 19917 | A4GALT/AI | 444/1241 | 5934/19917 |
| down-regulated | TF | TF:M07602 | Factor: CP2; motif: NNNNCCAGNCNN | 0.00580532454549778 | 1241 | 775 | 11173 | 19917 | A4GALT/AI | 775/1241 | 11173/19917 |
| down-regulated | TF | TF:M04756_1 | Factor: sin3A; motif: TGTCCNNGGTGCTG; match class: 1 | 0.00607353044577778 | 1241 | 563 | 7778 | 19917 | ABHD4/AC | 563/1241 | 7778/19917 |
| down-regulated | TF | TF:M10107_1 | Factor: DB1; motif: GGRRRRGRRGGAGGGGGNGRRR; match class: 1 | 0.00617980313048358 | 1241 | 224 | 2686 | 19917 | ACSF2/AD/ | 224/1241 | 2686/19917 |
| down-regulated | TF | TF:M03563_1 | Factor: RelA-p65; motif: GGGANTTTCCNN; match class: 1 | 0.00660780893251664 | 1241 | 91 | 889 | 19917 | ADCYAP1R | 91/1241 | 889/19917 |
| down-regulated | TF | TF:M06948_1 | Factor: Sp2; motif: TGGGCGCGCCCA; match class: 1 | 0.00676943845491264 | 1241 | 463 | 6233 | 19917 | ABHD15/A | 463/1241 | 6233/19917 |
| down-regulated | TF | TF:M11529_1 | Factor: E2F-2; motif: GCGCGCGCNCS; match class: 1 | 0.00677397928654239 | 1241 | 978 | 14592 | 19917 | A4GALT/AI | 978/1241 | 14592/19917 |
| down-regulated | TF | TF:M07216 | Factor: IRF1; motif: NNNYASTTTCACTTTCNNTTT | 0.0068446491119539 | 1241 | 54 | 444 | 19917 | ACSM5/AN | 54/1241 | 444/19917 |
| down-regulated | TF | TF:M05775_1 | Factor: ZXDL; motif: NGGGGWS; match class: 1 | 0.0074983716482749 | 1241 | 155 | 1733 | 19917 | ACAA2/AC' | 155/1241 | 1733/19917 |
| down-regulated | TF | TF:M01219 | Factor: SP1:SP3; motif: CCSCCCCCYCC | 0.0076817346895675 | 1241 | 503 | 6856 | 19917 | ABHD15/A | 503/1241 | 6856/19917 |
| down-regulated | TF | TF:M10108_1 | Factor: WT1; motif: RGGNGGGGGAGGRGGNGGRG; match class: 1 | 0.00804580110129235 | 1241 | 237 | 2881 | 19917 | ACSF2/AD/ | 237/1241 | 2881/19917 |
| down-regulated | TF | TF:M09910 | Factor: ER-beta; motif: RGGTCASCNTGMCCY | 0.00814016918127658 | 1241 | 677 | 9603 | 19917 | A4GALT/AI | 677/1241 | 9603/19917 |
| down-regulated | TF | TF:M12092 | Factor: ZNF787; motif: TGCCTCAGTTTMCCN | 0.00887839950975165 | 1241 | 70 | 636 | 19917 | ANPEP/AP( | 70/1241 | 636/19917 |
| down-regulated | TF | TF:M08905 | Factor: TIEG1; motif: GGSGGKGNNN | 0.00888361344475827 | 1241 | 394 | 5197 | 19917 | A4GALT/A( | 394/1241 | 5197/19917 |
| down-regulated | TF | TF:M01073 | Factor: HIC1; motif: NNNGGKTGCCCSNNNNNN | 0.00915864436139045 | 1241 | 322 | 4123 | 19917 | A4GALT/AI | 322/1241 | 4123/19917 |
| down-regulated | TF | TF:M07297 | Factor: MAZ; motif: CCCTCCCYCYN | 0.00932807935851701 | 1241 | 244 | 2987 | 19917 | A4GALT/AI | 244/1241 | 2987/19917 |
| down-regulated | TF | TF:M01072_1 | Factor: HIC1; motif: NSNNNNTGCCCSSNN; match class: 1 | 0.00959493613953286 | 1241 | 91 | 897 | 19917 | ABHD15/A | 91/1241 | 897/19917 |
| down-regulated | TF | TF:M04855_1 | Factor: IRF-4; motif: AAGTTTC; match class: 1 | 0.00967736336341146 | 1241 | 176 | 2029 | 19917 | ACKR3/AD( | 176/1241 | 2029/19917 |
| down-regulated | TF | TF:M10014 | Factor: FXR; motif: NRGGKCANTGRCCNNNNGG | 0.0098397959205103 | 1241 | 426 | 5687 | 19917 | ACVRL1/A( | 426/1241 | 5687/19917 |
| down-regulated | TF | TF:M07395_1 | Factor: Sp1; motif: NGGGGCGGGGN; match class: 1 | 0.00994871883636562 | 1241 | 471 | 6376 | 19917 | A4GALT/AI | 471/1241 | 6376/19917 |
| down-regulated | TF | TF:M00706_1 | Factor: TFII-I; motif: RGAGGKAGG; match class: 1 | 0.0099738270766805 | 1241 | 145 | 1606 | 19917 | ADAMTS9/ | 145/1241 | 1606/19917 |
| down-regulated | TF | TF:M01175_1 | Factor: CKROX; motif: SCCCTCCCC; match class: 1 | 0.0100796976466131 | 1241 | 223 | 2690 | 19917 | ACACB/AC! | 223/1241 | 2690/19917 |
| down-regulated | TF | TF:M09892_1 | Factor: E2F-1; motif: NNNNGGCGGGAARN; match class: 1 | 0.0102791442516365 | 1241 | 654 | 9248 | 19917 | A4GALT/AI | 654/1241 | 9248/19917 |
| down-regulated | TF | TF:M12158_1 | Factor: KLF15; motif: NCCMCGCCCMCN; match class: 1 | 0.0104344836566232 | 1241 | 449 | 6041 | 19917 | ABHD15/A | 449/1241 | 6041/19917 |
| down-regulated | TF | TF:M09762 | Factor: NF1B; motif: CYTGGCNYNCWGCCAN | 0.0105487721128795 | 1241 | 428 | 5721 | 19917 | ABCB1/ABI | 428/1241 | 5721/19917 |
| down-regulated | TF | TF:M10072_1 | Factor: sp4; motif: NNGNARGRGGCGGRGCNNRR; match class: 1 | 0.0106450307582747 | 1241 | 464 | 6272 | 19917 | A4GALT/AI | 464/1241 | 6272/19917 |
| down-regulated | TF | TF:M00772 | Factor: IRF; motif: RRAANTGAAASYGNV | 0.0108259885279978 | 1241 | 102 | 1040 | 19917 | ABCB1/AC! | 102/1241 | 1040/19917 |
| down-regulated | TF | TF:M07141 | Factor: FPM315; motif: GGAGGAGGRRGRGGRGGRRGR | 0.0109927181928596 | 1241 | 188 | 2200 | 19917 | ADAMTS9/ | 188/1241 | 2200/19917 |
| down-regulated | TF | TF:M07615 | Factor: Sp3; motif: GGGGCGGGGSNN | 0.0111289369422152 | 1241 | 501 | 6845 | 19917 | A4GALT/AI | 501/1241 | 6845/19917 |
| down-regulated | TF | TF:M10002_1 | Factor: NF-1C; motif: NYTGGCNNYNNGCCARN; match class: 1 | 0.0114448105262693 | 1241 | 152 | 1705 | 19917 | ACAA2/AC! | 152/1241 | 1705/19917 |
| down-regulated | TF | TF:M09724 | Factor: DATF1; motif: SNGGRRGCWGNGGG | 0.0122524093018376 | 1241 | 604 | 8463 | 19917 | A2ML1/A4 | 604/1241 | 8463/19917 |
| down-regulated | TF | TF:M10008 | Factor: CSX; motif: NKRAGWGS | 0.0131571741540397 | 1241 | 714 | 10227 | 19917 | A4GALT/AI | 714/1241 | 10227/19917 |
| down-regulated | TF | TF:M05327_1 | Factor: WT1; motif: NGCGGGGGGGTSMMCYN; match class: 1 | 0.0132295455832935 | 1241 | 156 | 1764 | 19917 | ACVRL1/AI | 156/1241 | 1764/19917 |
| down-regulated | TF | TF:M08819_1 | Factor: LKLF; motif: CNCCACCCS; match class: 1 | 0.0133226633832909 | 1241 | 140 | 1547 | 19917 | ALDH3B1/ | 140/1241 | 1547/19917 |
| down-regulated | TF | TF:M09589 | Factor: AR; motif: NGNACANNNTGTTCYNN | 0.0133421639470388 | 1241 | 392 | 5187 | 19917 | ABCC9/ABI | 392/1241 | 5187/19917 |
| down-regulated | TF | TF:M00649_1 | Factor: MAZ; motif: GGGGAGGG; match class: 1 | 0.013473651923478 | 1241 | 325 | 4185 | 19917 | ACACB/AC' | 325/1241 | 4185/19917 |
| down-regulated | TF | TF:M09591 | Factor: AP-2gamma; motif: NTGSCCTGRGGSNN | 0.013669504570724 | 1241 | 600 | 8406 | 19917 | A4GALT/AI | 600/1241 | 8406/19917 |
| down-regulated | TF | TF:M08487_1 | Factor: GCMa:Erg; motif: ATGCGGGCGGAARKG; match class: 1 | 0.0143972201584136 | 1241 | 628 | 8853 | 19917 | A4GALT/AI | 628/1241 | 8853/19917 |
| down-regulated | TF | TF:M04788 | Factor: IRF-1; motif: STTTCACTTTCNNT | 0.0153658965357292 | 1241 | 61 | 538 | 19917 | ACSM5/AP | 61/1241 | 538/19917 |

| | | | | | | | | | | | |
|---|---|---|---|---|---|---|---|---|---|---|---|
| down-regulated | TF | TF:M04934_1 | Factor: TR4; motif: ACCCCGS; match class: 1 | 0.015651039834231 | 1241 | 1033 | 15594 | 19917 | A2ML1/A4 | 1033/1241 | 15594/19917 |
| down-regulated | TF | TF:M11390 | Factor: Erg; motif: NACAGGAARTN | 0.0167236350819301 | 1241 | 229 | 2795 | 19917 | ABCB1/AC | 229/1241 | 2795/19917 |
| down-regulated | TF | TF:M01865_1 | Factor: BTEB3; motif: BNRNGGGAGGNGT; match class: 1 | 0.0180034884176671 | 1241 | 328 | 4243 | 19917 | ACSF2/AD | 328/1241 | 4243/19917 |
| down-regulated | TF | TF:M00054 | Factor: NF-kappaB; motif: GGGAMTTYCC | 0.0182804925160137 | 1241 | 205 | 2457 | 19917 | ABCB1/ACI | 205/1241 | 2457/19917 |
| down-regulated | TF | TF:M07249 | Factor: ctcf; motif: CCNCNAGRKGGCRSTN | 0.0185017695506461 | 1241 | 493 | 6749 | 19917 | A2ML1/A4 | 493/1241 | 6749/19917 |
| down-regulated | TF | TF:M02281 | Factor: SP1; motif: CCCCKCCCCC | 0.0187669666361043 | 1241 | 478 | 6518 | 19917 | ACOX2/AC | 478/1241 | 6518/19917 |
| down-regulated | TF | TF:M01199_1 | Factor: RNF96; motif: BCCCGCRGCC; match class: 1 | 0.0191071392280348 | 1241 | 334 | 4335 | 19917 | A4GALT/AI | 334/1241 | 4335/19917 |
| down-regulated | TF | TF:M10056 | Factor: DRRS; motif: GNNGGGWGGG | 0.0193378069358994 | 1241 | 368 | 4844 | 19917 | A4GALT/AI | 368/1241 | 4844/19917 |
| down-regulated | TF | TF:M09658_1 | Factor: Sp2; motif: GGSNNGGGGGCGGGGCCNGNGS; match class: 1 | 0.0203404695673353 | 1241 | 175 | 2040 | 19917 | ACSF2/ACS | 175/1241 | 2040/19917 |
| down-regulated | TF | TF:M08911_1 | Factor: CTCF; motif: NCCRSTAGGGGGCGC; match class: 1 | 0.0219280000448291 | 1241 | 308 | 3956 | 19917 | A4GALT/A | 308/1241 | 3956/19917 |
| down-regulated | TF | TF:M09738 | Factor: ZNF511; motif: GGRRGRGGCWGNG | 0.0219993041263576 | 1241 | 757 | 10956 | 19917 | A2ML1/A4 | 757/1241 | 10956/19917 |
| down-regulated | TF | TF:M01835 | Factor: GKLF; motif: CCTCCYN | 0.023135211188021 | 1241 | 1040 | 15737 | 19917 | A2ML1/A4 | 1040/1241 | 15737/19917 |
| down-regulated | TF | TF:M00054_1 | Factor: NF-kappaB; motif: GGGAMTTYCC; match class: 1 | 0.0252089740872958 | 1241 | 75 | 717 | 19917 | ACVRL1/AI | 75/1241 | 717/19917 |
| down-regulated | TF | TF:M09970_1 | Factor: KLF3; motif: NNNNNNGGGCGGGGCNNGN; match class: 1 | 0.0268131356456394 | 1241 | 297 | 3803 | 19917 | A4GALT/AI | 297/1241 | 3803/19917 |
| down-regulated | TF | TF:M09610 | Factor: ER-beta; motif: NRGGTCANKSTGACCTNN | 0.0273740516028131 | 1241 | 435 | 5878 | 19917 | A4GALT/AI | 435/1241 | 5878/19917 |
| down-regulated | TF | TF:M03557_1 | Factor: P50; motif: GGRRANTCCCNN; match class: 1 | 0.0278566457308077 | 1241 | 120 | 1300 | 19917 | A4GALT/AI | 120/1241 | 1300/19917 |
| down-regulated | TF | TF:M05318 | Factor: KLF8; motif: NGGGCGGGGG | 0.0280987492123494 | 1241 | 84 | 832 | 19917 | ABHD4/AD | 84/1241 | 832/19917 |
| down-regulated | TF | TF:M05319 | Factor: KLF7; motif: NGGGCGGGGG | 0.0280987492123494 | 1241 | 84 | 832 | 19917 | ABHD4/AD | 84/1241 | 832/19917 |
| down-regulated | TF | TF:M08878_1 | Factor: EGR; motif: CGCCCCCGCNN; match class: 1 | 0.0284709113706372 | 1241 | 258 | 3235 | 19917 | ACBD7/AEI | 258/1241 | 3235/19917 |
| down-regulated | TF | TF:M08905_1 | Factor: TIEG1; motif: GGSGGKGNNN; match class: 1 | 0.0302779622920265 | 1241 | 99 | 1026 | 19917 | ADAMTSL4 | 99/1241 | 1026/19917 |
| down-regulated | TF | TF:M00807 | Factor: Egr; motif: GTGGGSGCRRS | 0.0308615507525155 | 1241 | 335 | 4373 | 19917 | ACBD7/AD | 335/1241 | 4373/19917 |
| down-regulated | TF | TF:M00084 | Factor: MZF-1; motif: KNGNKAGGGGNAA | 0.0343565988360722 | 1241 | 481 | 6598 | 19917 | A2ML1/AB | 481/1241 | 6598/19917 |
| down-regulated | TF | TF:M09668 | Factor: RelA-p65; motif: GGGRNTTTCCM | 0.0345394362844699 | 1241 | 282 | 3594 | 19917 | ABCB1/ABI | 282/1241 | 3594/19917 |
| down-regulated | TF | TF:M04953_1 | Factor: Sp1; motif: GGNDGGRGGCGGGG; match class: 1 | 0.0369438163814177 | 1241 | 316 | 4099 | 19917 | ACSF2/AD | 316/1241 | 4099/19917 |
| down-regulated | TF | TF:M01224_1 | Factor: P50:RELA-P65; motif: GGANTTYCCCWN; match class: 1 | 0.0398554172312734 | 1241 | 33 | 236 | 19917 | ARHGEF40 | 33/1241 | 236/19917 |
| down-regulated | TF | TF:M10071 | Factor: Sp1; motif: NGGGGGCGGGGCCNGGGGGGGG | 0.0410577787962974 | 1241 | 605 | 8550 | 19917 | A4GALT/AI | 605/1241 | 8550/19917 |
| down-regulated | TF | TF:M09915_1 | Factor: PEA3; motif: NNCAGGAARNN; match class: 1 | 0.0424612635095399 | 1241 | 425 | 5749 | 19917 | ABCB1/AC | 425/1241 | 5749/19917 |
| down-regulated | TF | TF:M09968 | Factor: KLF15; motif: RGGGMGGRGNNGGGGNGG | 0.0428657291820047 | 1241 | 253 | 3180 | 19917 | ABHD15/A | 253/1241 | 3180/19917 |
| down-regulated | TF | TF:M02023_1 | Factor: MAZ; motif: NKGGGAGGGGRGGR; match class: 1 | 0.0444639653966423 | 1241 | 250 | 3138 | 19917 | ACVRL1/A | 250/1241 | 3138/19917 |
| down-regulated | TF | TF:M01837_1 | Factor: FKLF; motif: BGGGNGGVMD; match class: 1 | 0.0450209021438259 | 1241 | 128 | 1421 | 19917 | ADCY4/AEI | 128/1241 | 1421/19917 |
| down-regulated | TF | TF:M07226_1 | Factor: SP1; motif: NCCCCKCCCCC; match class: 1 | 0.0454147045980054 | 1241 | 304 | 3931 | 19917 | ACSF2/ACS | 304/1241 | 3931/19917 |
| down-regulated | TF | TF:M07063 | Factor: Sp1; motif: GGGGCGGGGC | 0.0454583800651347 | 1241 | 553 | 7736 | 19917 | A4GALT/AI | 553/1241 | 7736/19917 |
| down-regulated | TF | TF:M10435 | Factor: Sp2; motif: GGGGCGGGG | 0.0454583800651347 | 1241 | 553 | 7736 | 19917 | A4GALT/AI | 553/1241 | 7736/19917 |
| down-regulated | TF | TF:M09737 | Factor: ZNF644; motif: TCCWGCCTCTSN | 0.0462616302557778 | 1241 | 527 | 7330 | 19917 | A4GALT/AI | 527/1241 | 7330/19917 |
| down-regulated | TF | TF:M00339_1 | Factor: c-Ets-1; motif: RCAGGAAGTGNNTNS; match class: 1 | 0.0479760462462777 | 1241 | 132 | 1477 | 19917 | ABCC9/ACS | 132/1241 | 1477/19917 |
| down-regulated | WP | WP:WP2882 | Nuclear Receptors Meta-Pathway | 1.08563453141997e-05 | 665 | 58 | 321 | 7562 | ABCB1/ACI | 58/665 | 321/7562 |
| down-regulated | WP | WP:WP4331 | Neovascularisation processes | 0.0014786325791834 | 665 | 13 | 37 | 7562 | ACVRL1/CX | 13/665 | 37/7562 |
| down-regulated | WP | WP:WP2431 | Spinal Cord Injury | 0.00151810376133972 | 665 | 26 | 118 | 7562 | ANXA1/AQ | 26/665 | 118/7562 |
| down-regulated | WP | WP:WP236 | Adipogenesis | 0.00374481893599094 | 665 | 27 | 131 | 7562 | CDKN1A/C | 27/665 | 131/7562 |
| down-regulated | WP | WP:WP2880 | Glucocorticoid Receptor Pathway | 0.00514986170248671 | 665 | 18 | 71 | 7562 | ACKR3/AN | 18/665 | 71/7562 |
| down-regulated | WP | WP:WP3678 | Amplification and Expansion of Oncogenic Pathways as Metastatic Tr | 0.00736789950153853 | 665 | 8 | 17 | 7562 | EPAS1/JAG | 8/665 | 17/7562 |
| down-regulated | WP | WP:WP4462 | Platelet-mediated interactions with vascular and circulating cells | 0.00736789950153853 | 665 | 8 | 17 | 7562 | CCL2/CD4 | 8/665 | 17/7562 |
| down-regulated | WP | WP:WP3845 | Canonical and Non-canonical Notch signaling | 0.00990804529211763 | 665 | 10 | 27 | 7562 | DLL4/HES1 | 10/665 | 27/7562 |
| down-regulated | WP | WP:WP3888 | VEGFA-VEGFR2 Signaling Pathway | 0.0153696149786186 | 665 | 62 | 437 | 7562 | ACACB/ACI | 62/665 | 437/7562 |
| down-regulated | WP | WP:WP4542 | Overview of leukocyte-intrinsic Hippo pathway functions | 0.0314632291145704 | 665 | 11 | 36 | 7562 | FOXO1/FO | 11/665 | 36/7562 |
| down-regulated | WP | WP:WP2840 | Hair Follicle Development: Cytodifferentiation - Part 3 of 3 | 0.0382508563697309 | 665 | 19 | 89 | 7562 | CD34/EGFI | 19/665 | 89/7562 |
| down-regulated | WP | WP:WP4823 | Genes controlling nephrogenesis | 0.0419082854158895 | 665 | 12 | 43 | 7562 | CXCL12/EN | 12/665 | 43/7562 |
| up-regulated | GO:BP | GO:0099536 | synaptic signaling | 0.00628908240609847 | 124 | 18 | 747 | 18123 | CACNA1G/ | 18/124 | 747/18123 |
| up-regulated | GO:BP | GO:0007268 | chemical synaptic transmission | 0.0213217490904402 | 116 | 16 | 713 | 18123 | CACNA1G/ | 16/116 | 713/18123 |
| up-regulated | GO:BP | GO:0098916 | anterograde trans-synaptic signaling | 0.0213217490904402 | 116 | 16 | 713 | 18123 | CACNA1G/ | 16/116 | 713/18123 |
| up-regulated | GO:BP | GO:0099537 | trans-synaptic signaling | 0.0244734807654426 | 116 | 16 | 721 | 18123 | CACNA1G/ | 16/116 | 721/18123 |
| up-regulated | GO:CC | GO:0045202 | synapse | 0.00180851368110797 | 123 | 24 | 1348 | 18964 | CACNA1G/ | 24/123 | 1348/18964 |
| up-regulated | GO:CC | GO:0030424 | axon | 0.00207955900629833 | 123 | 16 | 661 | 18964 | AP3B2/CAI | 16/123 | 661/18964 |
| up-regulated | GO:CC | GO:0043025 | neuronal cell body | 0.00270063396411897 | 123 | 14 | 523 | 18964 | CCK/CHRN | 14/123 | 523/18964 |
| up-regulated | GO:CC | GO:0043005 | neuron projection | 0.00282715232168121 | 123 | 24 | 1384 | 18964 | AP3B2/CA | 24/123 | 1384/18964 |
| up-regulated | GO:CC | GO:0036477 | somatodendritic compartment | 0.00486658848901195 | 123 | 18 | 875 | 18964 | CALB2/CCK | 18/123 | 875/18964 |
| up-regulated | GO:CC | GO:0034702 | ion channel complex | 0.00984610631670279 | 123 | 10 | 302 | 18964 | CACNA1G/ | 10/123 | 302/18964 |
| up-regulated | GO:CC | GO:0044297 | cell body | 0.0113455885480604 | 123 | 14 | 595 | 18964 | CCK/CHRN | 14/123 | 595/18964 |
| up-regulated | GO:CC | GO:1902495 | transmembrane transporter complex | 0.0183015607388494 | 123 | 10 | 325 | 18964 | CACNA1G/ | 10/123 | 325/18964 |
| up-regulated | GO:CC | GO:0120025 | plasma membrane bounded cell projection | 0.025611370080386 | 123 | 30 | 2234 | 18964 | AP3B2/CA | 30/123 | 2234/18964 |
| up-regulated | GO:CC | GO:1990351 | transporter complex | 0.02665626488872 | 123 | 10 | 340 | 18964 | CACNA1G/ | 10/123 | 340/18964 |
| up-regulated | GO:CC | GO:0045211 | postsynaptic membrane | 0.0332862400600858 | 123 | 9 | 282 | 18964 | CACNG8/C | 9/123 | 282/18964 |
| up-regulated | GO:CC | GO:0031226 | intrinsic component of plasma membrane | 0.0358773495127966 | 123 | 25 | 1725 | 18964 | ATP4A/CA | 25/123 | 1725/18964 |
| up-regulated | GO:CC | GO:0034703 | cation channel complex | 0.0417778501081608 | 123 | 8 | 227 | 18964 | CACNA1G/ | 8/123 | 227/18964 |
| up-regulated | GO:CC | GO:0005887 | integral component of plasma membrane | 0.0449544973661173 | 123 | 24 | 1643 | 18964 | ATP4A/CA | 24/123 | 1643/18964 |
| up-regulated | GO:MF | GO:0005272 | sodium channel activity | 0.000232178108774484 | 126 | 6 | 44 | 18679 | CACNA1G/ | 6/126 | 44/18679 |
| up-regulated | GO:MF | GO:0005261 | cation channel activity | 0.00151348907585196 | 126 | 12 | 338 | 18679 | CACNA1G/ | 12/126 | 338/18679 |
| up-regulated | GO:MF | GO:0022836 | gated channel activity | 0.00175976175180365 | 126 | 12 | 343 | 18679 | CACNA1G/ | 12/126 | 343/18679 |
| up-regulated | GO:MF | GO:0015081 | sodium ion transmembrane transporter activity | 0.0032207344863595 | 126 | 8 | 146 | 18679 | ATP4A/CA | 8/126 | 146/18679 |
| up-regulated | GO:MF | GO:0005216 | ion channel activity | 0.0036875623223811 | 126 | 13 | 434 | 18679 | CACNA1G/ | 13/126 | 434/18679 |
| up-regulated | GO:MF | GO:0005230 | extracellular ligand-gated ion channel activity | 0.00477318221180534 | 126 | 6 | 73 | 18679 | CHRNA10/ | 6/126 | 73/18679 |
| up-regulated | GO:MF | GO:1904315 | transmitter-gated ion channel activity involved in regulation of posts | 0.006161687357991 | 126 | 5 | 45 | 18679 | CHRNA10/ | 5/126 | 45/18679 |
| up-regulated | GO:MF | GO:0099529 | neurotransmitter receptor activity involved in regulation of postsyna | 0.0084971573222746 | 126 | 5 | 48 | 18679 | CHRNA10/ | 5/126 | 48/18679 |
| up-regulated | GO:MF | GO:0022803 | passive transmembrane transporter activity | 0.0114283143437341 | 126 | 13 | 483 | 18679 | CACNA1G/ | 13/126 | 483/18679 |
| up-regulated | GO:MF | GO:0015267 | channel activity | 0.0114283143437341 | 126 | 13 | 483 | 18679 | CACNA1G/ | 13/126 | 483/18679 |
| up-regulated | GO:MF | GO:0046873 | metal ion transmembrane transporter activity | 0.0164658760159257 | 126 | 12 | 429 | 18679 | ATP4A/CA | 12/126 | 429/18679 |
| up-regulated | GO:MF | GO:0005231 | excitatory extracellular ligand-gated ion channel activity | 0.0229223082377133 | 126 | 4 | 30 | 18679 | CHRNA10/ | 4/126 | 30/18679 |
| up-regulated | GO:MF | GO:0015276 | ligand-gated ion channel activity | 0.0233372140625653 | 126 | 7 | 141 | 18679 | CHRNA10/ | 7/126 | 141/18679 |
| up-regulated | GO:MF | GO:0022834 | ligand-gated channel activity | 0.0233372140625653 | 126 | 7 | 141 | 18679 | CHRNA10/ | 7/126 | 141/18679 |
| up-regulated | GO:MF | GO:0022890 | inorganic cation transmembrane transporter activity | 0.0243769236559167 | 126 | 14 | 596 | 18679 | ATP4A/CA | 14/126 | 596/18679 |
| up-regulated | GO:MF | GO:0022824 | transmitter-gated ion channel activity | 0.0254065410218709 | 126 | 5 | 60 | 18679 | CHRNA10/ | 5/126 | 60/18679 |
| up-regulated | GO:MF | GO:0022835 | transmitter-gated channel activity | 0.0254065410218709 | 126 | 5 | 60 | 18679 | CHRNA10/ | 5/126 | 60/18679 |
| up-regulated | GO:MF | GO:0015075 | ion transmembrane transporter activity | 0.0258760958756596 | 126 | 17 | 845 | 18679 | ATP4A/CA | 17/126 | 845/18679 |
| up-regulated | GO:MF | GO:0098960 | postsynaptic neurotransmitter receptor activity | 0.0321742225798795 | 126 | 5 | 63 | 18679 | CHRNA10/ | 5/126 | 63/18679 |
| up-regulated | HPA | HPA:0260000 | hypothalamus | 0.0206765188086552 | 71 | 4 | 39 | 10877 | CALY/CART | 4/71 | 39/10877 |
| up-regulated | HPA | HPA:0090941 | cerebellum; Purkinje cells - cytoplasm/membrane[<U+2265>Low] | 0.0251250901181051 | 71 | 11 | 448 | 10877 | AP3B2/C1 | 11/71 | 448/10877 |
| up-regulated | KEGG | KEGG:04080 | Neuroactive ligand-receptor interaction | 0.000576124345706354 | 51 | 11 | 340 | 8000 | CCK/CHRN | 11/51 | 340/8000 |
| up-regulated | REAC | REAC:R-HSA-4513 | Activation of Na-permeable kainate receptors | 0.0135244797421138 | 63 | 2 | 2 | 10622 | GRIK1/GRI | 2/63 | 2/10622 |
| up-regulated | REAC | REAC:R-HSA-8948 | Collagen chain trimerization | 0.0498780448387853 | 63 | 4 | 44 | 10622 | COL11A2/ | 4/63 | 44/10622 |
| up-regulated | TF | TF:M07461_1 | Factor: KLF; motif: GGGNGGGG; match class: 1 | 2.1948321330361e-06 | 131 | 52 | 3423 | 19917 | ACTL6B/A | 52/131 | 3423/19917 |
| up-regulated | TF | TF:M07261_1 | Factor: LKLF; motif: GGGGTGGKSN; match class: 1 | 2.23534512758732e-06 | 131 | 58 | 4110 | 19917 | ACTL6B/AI | 58/131 | 4110/19917 |
| up-regulated | TF | TF:M09734_1 | Factor: ZNF692; motif: SYNGGSCCCASCCNC; match class: 1 | 2.40926824668873e-05 | 131 | 73 | 6295 | 19917 | ACTL6B/A | 73/131 | 6295/19917 |
| up-regulated | TF | TF:M08819 | Factor: LKLF; motif: CNCCACCCS | 2.77537029674663e-05 | 131 | 68 | 5645 | 19917 | ACTL6B/A | 68/131 | 5645/19917 |
| up-regulated | TF | TF:M09729_1 | Factor: LUMAN; motif: CYCAGCYYCY; match class: 1 | 0.000106760130727501 | 131 | 59 | 4665 | 19917 | ACTL6B/AI | 59/131 | 4665/19917 |
| up-regulated | TF | TF:M09591 | Factor: AP-2gamma; motif: NTGSCCTGRGGSNN | 0.000129169215974285 | 131 | 86 | 8406 | 19917 | ABCC12/A | 86/131 | 8406/19917 |
| up-regulated | TF | TF:M00450_1 | Factor: Zic3; motif: NGGGKGGTC; match class: 1 | 0.000162104996666665 | 131 | 83 | 7991 | 19917 | ABCC12/A | 83/131 | 7991/19917 |
| up-regulated | TF | TF:M07461 | Factor: KLF; motif: GGGNGGG | 0.000275702211568732 | 131 | 88 | 8831 | 19917 | ACTL6B/A | 88/131 | 8831/19917 |
| up-regulated | TF | TF:M07436 | Factor: WT1; motif: NNGGGNGGGSGN | 0.000723248005533735 | 131 | 69 | 6227 | 19917 | ACTL6B/A | 69/131 | 6227/19917 |
| up-regulated | TF | TF:M07261 | Factor: LKLF; motif: GGGGTGGKSN | 0.00122501298765914 | 131 | 93 | 9867 | 19917 | ABCC12/A | 93/131 | 9867/19917 |
| up-regulated | TF | TF:M00378_1 | Factor: Pax-4; motif: NNNNNYCACCCB; match class: 1 | 0.00152500034016686 | 131 | 120 | 14850 | 19917 | ABCC12/A | 120/131 | 14850/19917 |
| up-regulated | TF | TF:M08819_1 | Factor: LKLF; motif: CNCCACCCS; match class: 1 | 0.00191761259553761 | 131 | 28 | 1547 | 19917 | ACTL6B/AT | 28/131 | 1547/19917 |
| up-regulated | TF | TF:M07289_1 | Factor: GKLF; motif: NNNRGGNGNGGSN; match class: 1 | 0.00248392064179001 | 131 | 97 | 10645 | 19917 | ACTL6B/AI | 97/131 | 10645/19917 |
| up-regulated | TF | TF:M04595 | Factor: SALL2; motif: GGGTGGG | 0.00256507757915453 | 131 | 111 | 13123 | 19917 | ACTL6B/A | 111/131 | 13123/19917 |
| up-regulated | TF | TF:M04595_1 | Factor: SALL2; motif: GGGTGGG; match class: 1 | 0.00291706004824285 | 131 | 76 | 7428 | 19917 | ACTL6B/A | 76/131 | 7428/19917 |
| up-regulated | TF | TF:M00257_1 | Factor: RREB-1; motif: CCCCAAACMMCCCC; match class: 1 | 0.00398173107690659 | 131 | 54 | 4513 | 19917 | ABCC12/A | 54/131 | 4513/19917 |
| up-regulated | TF | TF:M04896 | Factor: AP-2alpha; motif: NNNGSCCTGRGGSN | 0.00400070048825331 | 131 | 64 | 5810 | 19917 | AGAP6/AH | 64/131 | 5810/19917 |
| up-regulated | TF | TF:M11595_1 | Factor: GCMa; motif: RTGCGGGTN; match class: 1 | 0.00520905636391492 | 131 | 64 | 5851 | 19917 | AHRR/AKR | 64/131 | 5851/19917 |
| up-regulated | TF | TF:M00721 | Factor: CACCC-binding; motif: CANCCNNWGGGTGDGG | 0.00845341412785439 | 131 | 40 | 2940 | 19917 | AGAP6/AK | 40/131 | 2940/19917 |
| up-regulated | TF | TF:M00257 | Factor: RREB-1; motif: CCCCAAACMMCCCC | 0.0116396162919053 | 131 | 95 | 10599 | 19917 | ABCC12/A | 95/131 | 10599/19917 |
| up-regulated | TF | TF:M11477 | Factor: AP-2beta; motif: NSCCYNRGGSN | 0.0157062260300855 | 131 | 80 | 8310 | 19917 | ACTL6B/AK | 80/131 | 8310/19917 |
| up-regulated | TF | TF:M11477_1 | Factor: AP-2beta; motif: NSCCYNRGGSN; match class: 1 | 0.0159187520934954 | 131 | 71 | 7003 | 19917 | ACTL6B/AK | 71/131 | 7003/19917 |
| up-regulated | TF | TF:M10112 | Factor: Miz-1; motif: NNRGGWGGGGGAGGGGMRR | 0.0162141313116773 | 131 | 83 | 8768 | 19917 | ABCC12/A | 83/131 | 8768/19917 |
| up-regulated | TF | TF:M11479 | Factor: AP-2alpha; motif: NGCCTNAGGCN | 0.0172542274900993 | 131 | 53 | 4598 | 19917 | ACTL6B/AK | 53/131 | 4598/19917 |

| | | | | | | | | | | | |
|---|---|---|---|---|---|---|---|---|---|---|---|
| up-regulated | TF | TF:M09972 | Factor: BTEB2; motif: WGGGTGKGGCNGGN | 0.0216119813613964 | 131 | 86 | 9283 | 19917 | ACTL6B/A | 86/131 | 9283/19917 |
| up-regulated | TF | TF:M12160_1 | Factor: KLF15; motif: RCCMCRCCCMCN; match class: 1 | 0.0231997512563556 | 131 | 78 | 8085 | 19917 | ACTL6B/A | 78/131 | 8085/19917 |
| up-regulated | TF | TF:M09591_1 | Factor: AP-2gamma; motif: NTGSCCTGRGGSNN; match class: 1 | 0.0250884599170063 | 131 | 38 | 2843 | 19917 | AKR1E2/A | 38/131 | 2843/19917 |
| up-regulated | TF | TF:M07436_1 | Factor: WT1; motif: NNGGGNGGGSGN; match class: 1 | 0.0316395655883718 | 131 | 34 | 2424 | 19917 | ACTL6B/A | 34/131 | 2424/19917 |
| up-regulated | TF | TF:M09972_1 | Factor: BTEB2; motif: WGGGTGKGGCNGGN; match class: 1 | 0.0336760094686207 | 131 | 47 | 3947 | 19917 | AGAP6/AH | 47/131 | 3947/19917 |
| up-regulated | TF | TF:M09834 | Factor: ZNF148; motif: NNNNNNCCNNCCCCTCCCCCACCCN | 0.0342282182296972 | 131 | 70 | 6998 | 19917 | ANKRD24/ | 70/131 | 6998/19917 |
