## Supplementary tables for "Transcriptome profiling of the dorsomedial prefrontal cortex in suicide victims": S5_PPI_network_data_of_down-_and_upregulated_genes1.pdf

Supplementary file 5. : Downregulated DEGs assigned to clusters by STRING using MCL algorithm. Functional annotation of clusters with 10 or more genes was performed based on GO-Biological process ontology.

| MCL cluster number | Number of genes | Functional annotation of cluster | Protein name | Protein identifier | Protein description |
|---|---|---|---|---|---|
| 1 | 152 | Cell surface receptor signaling pathway, Signal transduction, Cytokine-mediated signaling pathway | A2ML1 | ENSP00000299698 | C3 and PZP-like alpha-2-macroglobulin domain-containing protein 9; Is able to inhibit all four classes of proteinases by a unique 'trapping' mechanism. This protein has a pep |
| | | | ACACB | ENSP00000341044 | acetyl-CoA carboxylase beta; Catalyzes the ATP-dependent carboxylation of acetyl-CoA to malonyl-CoA. Carries out three functions: biotin carboxyl carrier protein, biotin car |
| | | | ACKR3 | ENSP00000272928 | G-protein coupled receptor RDC1 homolog; Atypical chemokine receptor that controls chemokine levels and localization via high-affinity chemokine binding that is uncouple |
| | | | ACSL5 | ENSP00000348429 | acyl-CoA synthetase long-chain family member 5; Acyl-CoA synthetases (ACSL) activate long-chain fatty acids for both synthesis of cellular lipids, and degradation via beta-oxi |
| | | | ADM | ENSP00000436607 | Adrenomedullin; AM and PAMP are potent hypotensive and vasodilatator agents. Numerous actions have been reported most related to the physiologic control of fluid and |
| | | | ANGPT2 | ENSP00000314897 | Angiopoietin 2; Binds to TEK/TIE2, competing for the ANGPT1 binding site, and modulating ANGPT1 signaling. Can induce tyrosine phosphorylation of TEK/TIE2 in the absen |
| | | | APLN | ENSP00000391800 | Apelin; Endogenous ligands |
| | | | AREG | ENSP00000379097 | Colorectum cell-derived growth factor; Ligand of the EGF receptor/EGFR. Autocrine growth factor as well as a mitogen for a broad range of target cells including astrocytes, |
| | | | AXL | ENSP00000301178 | Tyrosine-protein kinase receptor UFO; Receptor tyrosine kinase that transduces signals from the extracellular matrix into the cytoplasm by binding growth factor GAS6 and w |
| | | | BCL6 | ENSP00000384371 | Zinc finger and BTB domain-containing protein 27; Transcriptional repressor mainly required for germinal center (GC) formation and antibody affinity maturation which has |
| | | | CASP4 | ENSP00000388566 | Caspase 4, apoptosis-related cysteine peptidase; Inflammatory caspase. Essential effector of NLRP3 inflammasome- dependent CASP1 activation and IL1B and IL18 secretion |
| | | | CCL2 | ENSP00000225831 | Monocyte chemotactic and activating factor; Chemotactic factor that attracts monocytes and basophils but not neutrophils or eosinophils. Augments monocyte anti-tumor a |
| | | | CD163 | ENSP00000352071 | Scavenger receptor cysteine-rich type 1 protein M130; Acute phase-regulated receptor involved in clearance and endocytosis of hemoglobin/haptoglobin complexes by mac |
| | | | CD248 | ENSP00000308117 | CD248 molecule, endosialin; May play a role in tumor angiogenesis; C-type lectin domain containing |
| | | | CD34 | ENSP00000310036 | Hematopoietic progenitor cell antigen CD34; Possible adhesion molecule with a role in early hematopoiesis by mediating the attachment of stem cells to the bone marrow e |
| | | | CD40 | ENSP00000361359 | Tumor necrosis factor receptor superfamily member 5; Receptor for TNFSF5/CD40LG. Transduces TRAF6- and MAP3K8-mediated signals that activate ERK in macrophages ar |
| | | | CD44 | ENSP00000398632 | GP90 lymphocyte homing/adhesion receptor; Receptor for hyaluronic acid (HA). Mediates cell-cell and cell-matrix interactions through its affinity for HA, and possibly also th |
| | | | CDKN2B | ENSP00000276925 | Cyclin-dependent kinase inhibitor 2B (p15, inhibits CDK4); Interacts strongly with CDK4 and CDK6. Potent inhibitor. Potential effector of TGF-beta induced cell cycle arrest; B |
| | | | CISH | ENSP00000409346 | Cytokine inducible SH2-containing protein; SOCS family proteins form part of a classical negative feedback system that regulates cytokine signal transduction. CIS is involved |
| | | | CLIC1 | ENSP00000364935 | Regulatory nuclear chloride ion channel protein; Can insert into membranes and form chloride ion channels. Channel activity depends on the pH. Membrane insertion seem |
| | | | CNTFR | ENSP00000368265 | Ciliary neurotrophic factor receptor subunit alpha; Binds to CNTF. The alpha subunit provides the receptor specificity; Belongs to the type I cytokine receptor family. Type 3 |
| | | | COL4A1 | ENSP00000364979 | Collagen alpha-1(IV) chain; Type IV collagen is the major structural component of glomerular basement membranes (GBM), forming a 'chicken-wire' meshwork together with |
| | | | CP | ENSP00000264613 | Ceruloplasmin (ferroxidase); Ceruloplasmin is a blue, copper-binding (6-7 atoms per molecule) glycoprotein. It has ferroxidase activity oxidizing Fe(2+) to Fe(3+) without rele |
| | | | CTF1 | ENSP00000279804 | Cardiotrophin 1; Induces cardiac myocyte hypertrophy in vitro. Binds to and activates the ILST/gp130 receptor; Interleukin 6 type cytokine family |
| | | | CXCL1 | ENSP00000379110 | Chemokine (C-X-C motif) ligand 1 (melanoma growth stimulating activity, alpha); Has chemotactic activity for neutrophils. May play a role in inflammation and exerts its effec |
| | | | CXCL10 | ENSP00000305651 | 10 kDa interferon gamma-induced protein; Chemotactic for monocytes and T-lymphocytes. Binds to CXCR3; Belongs to the intercrine alpha (chemokine CxC) family. |
| | | | CXCL12 | ENSP00000379140 | Pre-B cell growth-stimulating factor; Chemoattractant active on T-lymphocytes, monocytes, but not neutrophils. Activates the C-X-C chemokine receptor CXCR4 to induce a r |
| | | | CXCL16 | ENSP00000293778 | Scavenger receptor for phosphatidylserine and oxidized low density lipoprotein; Acts as a scavenger receptor on macrophages, which specifically binds to OxLDL (oxidized lo |
| | | | CXCL3 | ENSP00000296026 | Macrophage inflammatory protein 2-beta; Ligand for CXCR2 (By similarity). Has chemotactic activity for neutrophils. May play a role in inflammation and exert its effects on |
| | | | CSF1 | ENSP00000327513 | Colony stimulating factor 1 (macrophage); Cytokine that plays an essential role in the regulation of survival, proliferation and differentiation of hematopoietic precursor cells |
| | | | CSF3 | ENSP00000225474 | Colony stimulating factor 3 (granulocyte); Granulocyte/macrophage colony-stimulating factors are cytokines that act in hematopoiesis by controlling the production, differen |
| | | | CST3 | ENSP00000381448 | Neuroendocrine basic polypeptide; As an inhibitor of cysteine proteinases, this protein is thought to serve an important physiological role as a local regulator of this enzyme |
| | | | DDR1 | ENSP00000427552 | Epithelial discoidin domain-containing receptor 1; Tyrosine kinase that functions as cell surface receptor for fibrillar collagen and regulates cell attachment to the extracellul |
| | | | DDR2 | ENSP00000356899 | Discoidin domain-containing receptor tyrosine kinase 2; Tyrosine kinase that functions as cell surface receptor for fibrillar collagen and regulates cell differentiation, remode |
| | | | DOK1 | ENSP00000233668 | Docking protein 1, 62kDa (downstream of tyrosine kinase 1); DOK proteins are enzymatically inert adaptor or scaffolding proteins. They provide a docking platform for the as |
| | | | EFNB1 | ENSP00000204961 | EPH-related receptor tyrosine kinase ligand 2; Binds to the receptor tyrosine kinases EPHB1 and EPHA1. Binds to, and induce the collapse of, commissural axons/growth con |
| | | | EGFR | ENSP00000275493 | Receptor tyrosine-protein kinase erbB-1; Receptor tyrosine kinase binding ligands of the EGF family and activating several signaling cascades to convert extracellular cues int |
| | | | ENG | ENSP00000362299 | Endoglin; Vascular endothelium glycoprotein that plays an important role in the regulation of angiogenesis. Required for normal structure and integrity of adult vasculature. |
| | | | EPHA2 | ENSP00000351209 | Tyrosine-protein kinase receptor ECK; Receptor tyrosine kinase which binds promiscuously membrane-bound ephrin-A family ligands residing on adjacent cells, leading to co |
| | | | EPHB4 | ENSP00000350896 | Tyrosine-protein kinase TYRO11; Receptor tyrosine kinase which binds promiscuously transmembrane ephrin-B family ligands residing on adjacent cells, leading to contact-d |
| | | | ERBB2 | ENSP00000269571 | V-erb-b2 avian erythroblastic leukemia viral oncogene homolog 2; Protein tyrosine kinase that is part of several cell surface receptor complexes, but that apparently needs a |
| | | | EVA1B | ENSP00000270824 | Eva-1 homolog B; Belongs to the EVA1 family. |
| | | | F11R | ENSP00000357005 | Junctional adhesion molecule 1; Seems to play a role in epithelial tight junction formation. Appears early in primordial forms of cell junctions and recruits PARD3. The associ |
| | | | F3 | ENSP00000334145 | Coagulation factor III (thromboplastin, tissue factor); Initiates blood coagulation by forming a complex with circulating factor VII or VIIa. The [TF:VIIa] complex activates facto |
| | | | FFAR2 | ENSP00000473159 | G-protein coupled receptor 43; G protein-coupled receptor that is activated by a major product of dietary fiber digestion, the short chain fatty acids (SCFAs), and that plays a |
| | | | FGF11 | ENSP00000293829 | Fibroblast growth factor homologous factor 3; Probably involved in nervous system development and function. |
| | | | FGFR3 | ENSP00000339824 | Fibroblast growth factor receptor 3; Tyrosine-protein kinase that acts as cell-surface receptor for fibroblast growth factors and plays an essential role in the regulation of cell |
| | | | FLT1 | ENSP00000282397 | Vascular endothelial growth factor receptor 1; Tyrosine-protein kinase that acts as a cell-surface receptor for VEGFA, VEGFB and PGF, and plays an essential role in the devel |
| | | | FLT3LG | ENSP00000469613 | Fms-related tyrosine kinase 3 ligand; Stimulates the proliferation of early hematopoietic cells by activating FLT3. Synergizes well with a number of other colony stimulating f |
| | | | FLT4 | ENSP00000261937 | Vascular endothelial growth factor receptor 3; Tyrosine-protein kinase that acts as a cell-surface receptor for VEGFC and VEGFD, and plays an essential role in adult lymphan |
| | | | FN1 | ENSP00000346839 | Fibronectin type III domain containing; Endogenous ligands |
| | | | FOXO1 | ENSP00000368880 | Forkhead in rhabdomyosarcoma; Transcription factor that is the main target of insulin signaling and regulates metabolic homeostasis in response to oxidative stress. Binds to |
| | | | FOXO4 | ENSP00000363377 | Fork head domain transcription factor AFX1; Transcription factor involved in the regulation of the insulin signaling pathway. Binds to insulin-response elements (IREs) and ca |
| | | | FSTL1 | ENSP00000295633 | Follistatin-related protein 1; May modulate the action of some growth factors on cell proliferation and differentiation. Binds heparin (By similarity); SPARC family |
| | | | FTL | ENSP00000366525 | Ferritin, light polypeptide; Stores iron in a soluble, non-toxic, readily available form. Important for iron homeostasis. Iron is taken up in the ferrous form and deposited as fe |
| | | | GABRE | ENSP00000359353 | Gamma-aminobutyric acid (GABA) A receptor, epsilon; GABA, the major inhibitory neurotransmitter in the vertebrate brain, mediates neuronal inhibition by binding to the |
| | | | GABRG1 | ENSP00000295452 | Gamma-aminobutyric acid (GABA) A receptor, gamma 1; GABA, the major inhibitory neurotransmitter in the vertebrate brain, mediates neuronal inhibition by binding to the |
| | | | GLRA1 | ENSP00000411593 | Glycine receptor strychnine-binding subunit; Glycine receptors are ligand-gated chloride channels. Channel opening is triggered by extracellular glycine. Channel opening is |
| | | | GPAM | ENSP00000265276 | Glycerol-3-phosphate acyltransferase 1, mitochondrial; Esterifies acyl-group from acyl-ACP to the sn-1 position of glycerol-3-phosphate, an essential step in glycerolipid biosy |
| | | | GPER1 | ENSP00000380281 | Flow-induced endothelial G-protein coupled receptor 1; G-protein coupled estrogen receptor that binds to 17- beta-estradiol (E2) with high affinity, leading to rapid and tran |
| | | | HAP1 | ENSP00000334002 | Huntingtin-associated protein 1; Originally identified as neuronal protein that specifically associates with HTT/huntingtin and the binding is enhanced by an expanded polygl |
| | | | HFE | ENSP00000417404 | Hereditary hemochromatosis protein; Binds to transferrin receptor (TFR) and reduces its affinity for iron-loaded transferrin; Belongs to the MHC class I family. |
| | | | ICAM1 | ENSP00000264832 | Intercellular adhesion molecule 1; ICAM proteins are ligands for the leukocyte adhesion protein LFA-1 (integrin alpha-L/beta-2). During leukocyte trans- endothelial migratio |
| | | | ICAM2 | ENSP00000415283 | Intercellular adhesion molecule 2; ICAM proteins are ligands for the leukocyte adhesion protein LFA-1 (integrin alpha-L/beta-2). ICAM2 may play a role in lymphocyte recircu |
| | | | IL10RA | ENSP00000227752 | Interleukin-10 receptor subunit alpha; Receptor for IL10; binds IL10 with a high affinity; CD molecules |
| | | | IL13RA1 | ENSP00000360730 | Interleukin-13 receptor subunit alpha-1; Binds with low affinity to interleukin-13 (IL13). Together with IL4RA can form a functional receptor for IL13. Also serves as an alterna |
| | | | IL15RA | ENSP00000380421 | Interleukin 15 receptor subunit alpha; CD molecules |
| | | | IL17D | ENSP00000302924 | Interleukin 17D; Induces expression of IL6, CXCL8/IL8, and CSF2/GM-CSF from endothelial cells; Interleukins |
| | | | IL17RC | ENSP00000295981 | Interleukin-17 receptor-like protein; Isoform 8: Receptor for both IL17A and IL17F; Interleukin receptors |
| | | | IL17RD | ENSP00000296318 | Interleukin-17 receptor-like protein; Feedback inhibitor of fibroblast growth factor mediated Ras-MAPK signaling and ERK activation. May inhibit FGF-induced FGFR1 tyrosine |
| | | | IL18R1 | ENSP00000387211 | CD218 antigen-like family member A; Within the IL18 receptor complex, responsible for the binding of the proinflammatory cytokine IL18, but not IL1A nor IL1B (Probable). |
| | | | IL1R1 | ENSP00000386380 | CD121 antigen-like family member A; Receptor for IL1A, IL1B and IL1RN. After binding to interleukin-1 associates with the coreceptor IL1RAP to form the high affinity interle |
| | | | IL1R2 | ENSP00000330959 | CD121 antigen-like family member B; Non-signaling receptor for IL1A, IL1B and IL1RN. Reduces IL1B activities. Serves as a decoy receptor by competetive binding to IL1B and |
| | | | IL1RL1 | ENSP00000233954 | Interleukin 1 receptor-like 1; Receptor for interleukin-33 (IL-33); signaling requires association of the coreceptor IL1RAP. Its stimulation recruits MYD88, IRAK1, IRAK4, and T |
| | | | IL32 | ENSP00000432218 | Interleukin 32; Interleukins |
| | | | IL3RA | ENSP00000327890 | Interleukin 3 receptor, alpha (low affinity); This is a receptor for interleukin-3; CD molecules |
| | | | IL4R | ENSP00000379111 | Interleukin-4 receptor subunit alpha; Receptor for both interleukin 4 and interleukin 13. Couples to the JAK1/2/3-STAT6 pathway. The IL4 response is involved in promoting |
| | | | IL6 | ENSP00000385675 | B-cell stimulatory factor 2; Cytokine with a wide variety of biological functions. It is a potent inducer of the acute phase response. Plays an essential role in the final different |
| | | | IL6R | ENSP00000357470 | Interleukin-6 receptor subunit alpha; Part of the receptor for interleukin 6. Binds to IL6 with low affinity, but does not transduce a signal. Signal activation necessitate an ass |
| | | | IRAK3 | ENSP00000261233 | Interleukin-1 receptor-associated kinase 3; Inhibits dissociation of IRAK1 and IRAK4 from the Toll- like receptor signaling complex by either inhibiting the phosphorylation of |
| | | | ITGA1 | ENSP00000282588 | CD49 antigen-like family member A; Integrin alpha-1/beta-1 is a receptor for laminin and collagen. It recognizes the proline-hydroxylated sequence G-F-P-G- E-R in collagen. |
| | | | ITGA5 | ENSP00000293379 | Integrin, alpha 5 (fibronectin receptor, alpha polypeptide); Integrin alpha-5/beta-1 is a receptor for fibronectin and fibrinogen. It recognizes the sequence R-G-D in its ligands |
| | | | ITGA6 | ENSP00000386896 | CD49 antigen-like family member F; Integrin alpha-6/beta-1 is a receptor for laminin on platelets. Integrin alpha-6/beta-4 is a receptor for laminin in epithelial cells and it pla |
| | | | ITGB1 | ENSP00000379350 | Integrin, beta 1 (fibronectin receptor, beta polypeptide, antigen CD29 includes MDF2, MSK12); Integrins alpha-1/beta-1, alpha-2/beta-1, alpha-10/beta- 1 and alpha-11/beta- |
| | | | ITGB4 | ENSP00000200181 | Integrin, beta 4; Integrin alpha-6/beta-4 is a receptor for laminin. Plays a critical structural role in the hemidesmosome of epithelial cells. Is required for the regulation of ker |
| | | | ITGB5 | ENSP00000296181 | Integrin, beta 5; Integrin alpha-V/beta-5 (ITGAV:ITGB5) is a receptor for fibronectin. It recognizes the sequence R-G-D in its ligand. |
| | | | KLF15 | ENSP00000296233 | Kidney-enriched krueppel-like factor; Transcriptional regulator that binds to the GA element of the CLCNKA promoter. Binds to the KCNIP2 promoter and regulates KCNIP2 |
| | | | KRT8 | ENSP00000449404 | Keratin, type II cytoskeletal 8; Together with KRT19, helps to link the contractile apparatus to dystrophin at the costameres of striated muscle; Keratins, type II |
| | | | LAMB2 | ENSP00000388325 | Laminin, beta 2 (laminin S); Binding to cells via a high affinity receptor, laminin is thought to mediate the attachment, migration and organization of cells into tissues during |
| | | | LAMC3 | ENSP00000354360 | Laminin-12 subunit gamma; Binding to cells via a high affinity receptor, laminin is thought to mediate the attachment, migration and organization of cells into tissues during |
| | | | LIF | ENSP00000249075 | Differentiation-stimulating factor; LIF has the capacity to induce terminal differentiation in leukemic cells. Its activities include the induction of hematopoietic differentiation |
| | | | LRIG1 | ENSP00000273261 | Leucine-rich repeats and immunoglobulin-like domains protein 1; Acts as a feedback negative regulator of signaling by receptor tyrosine kinases, through a mechanism that i |
| | | | LTBR | ENSP00000228918 | Lymphotoxin beta receptor (TNFR superfamily, member 3); Receptor for the heterotrimeric lymphotoxin containing LTA and LTB, and for TNFS14/LIGHT. Promotes apoptosis |
| | | | MCL1 | ENSP00000358022 | Induced myeloid leukemia cell differentiation protein Mcl-1; Involved in the regulation of apoptosis versus cell survival, and in the maintenance of viability but not of prolifer |
| | | | MLKL | ENSP00000308351 | Mixed lineage kinase domain-like protein; Pseudokinase that plays a key role in TNF-induced necroptosis, a programmed cell death process. Activated following phosphoryla |
| | | | MMP15 | ENSP00000219271 | Matrix metallopeptidase 15 (membrane-inserted); Endopeptidase that degrades various components of the extracellular matrix. May activate progelatinase A; Belongs to th |
| | | | MMP2 | ENSP00000219070 | Matrix metallopeptidase 2 (gelatinase A, 72kDa gelatinase, 72kDa type IV collagenase); Ubiquitinous metalloproteinase that is involved in diverse functions such as remodeli |
| | | | MMP28 | ENSP00000473853 | Matrix metalloproteinase-28; Can degrade casein. Could play a role in tissues homeostasis and repair; Belongs to the peptidase M10A family. |
| | | | MTTP | ENSP00000427679 | Microsomal triglyceride transfer protein large subunit; Catalyzes the transport of triglyceride, cholesteryl ester, and phospholipid between phospholipid surfaces. Required f |
| | | | MYD88 | ENSP00000401399 | Myeloid differentiation primary response protein MyD88; Adapter protein involved in the Toll-like receptor and IL-1 receptor signaling pathway in the innate immune respo |
| | | | NAMPT | ENSP00000222553 | Nicotinamide phosphoribosyltransferase; Catalyzes the condensation of nicotinamide with 5- phosphoribosyl-1-pyrophosphate to yield nicotinamide mononucleotide, an int |
| | | | NFATC1 | ENSP00000389377 | Nuclear factor of activated T-cells, cytoplasmic, calcineurin-dependent 1; Plays a role in the inducible expression of cytokine genes in T-cells, especially in the induction of the |
| | | | NFKB2 | ENSP00000358983 | Nuclear factor of kappa light polypeptide gene enhancer in B-cells 2 (p49/p100); NF-kappa-B is a pleiotropic transcription factor present in almost all cell types and is the end |
| | | | NKG7 | ENSP00000221978 | Natural killer cell granule protein 7; Belongs to the PMP-22/EMP/MP20 family. |
| | | | OSMR | ENSP00000274276 | Oncostatin-M-specific receptor subunit beta; Associates with IL31RA to form the IL31 receptor. Binds IL31 to activate STAT3 and possibly STAT1 and STAT5. Capable of trans |
| | | | PDGFRB | ENSP00000261799 | Platelet-derived growth factor receptor, beta polypeptide; Tyrosine-protein kinase that acts as cell-surface receptor for homodimeric PDGFB and PDGFD and for heterodime |
| | | | PDK4 | ENSP00000005178 | [Pyruvate dehydrogenase (acetyl-transferring)] kinase isozyme 4, mitochondrial; Kinase that plays a key role in regulation of glucose and fatty acid metabolism and homeosta |
| | | | PECAM1 | ENSP00000457421 | Platelet/endothelial cell adhesion molecule 1; Cell adhesion molecule which is required for leukocyte transendothelial migration (TEM) under most inflammatory conditions |
| | | | PGF | ENSP00000451040 | Placental growth factor; Growth factor active in angiogenesis and endothelial cell growth, stimulating their proliferation and migration. It binds to the receptor FLT1/VEGFR- |
| | | | PLIN5 | ENSP00000371272 | Lipid storage droplet protein 5; Lipid droplet-associated protein that maintains the balance between lipogenesis and lipolysis and also regulates fatty acid oxidation in oxidat |
| | | | PLXNB1 | ENSP00000351338 | Semaphorin receptor SEP; Receptor for SEMA4D. Plays a role in RHOA activation and subsequent changes of the actin cytoskeleton. Plays a role in axon guidance, invasive gr |
| | | | PPARA | ENSP00000385523 | Peroxisome proliferator-activated receptor alpha; Ligand-activated transcription factor. Key regulator of lipid metabolism. Activated by the endogenous ligand 1-palmitoyl- 2 |
| | | | RGL3 | ENSP00000377075 | Ral guanine nucleotide dissociation stimulator-like 3; Guanine nucleotide exchange factor (GEF) for Ral-A. Potential effector of GTPase HRas and Ras-related protein M-Ras. |
| | | | RIN2 | ENSP00000255006 | Ras interaction/interference protein 2; Ras effector protein. May function as an upstream activator and/or downstream effector for RAB5B in endocytic pathway. May functi |
| | | | ROR1 | ENSP00000360120 | Inactive tyrosine-protein kinase transmembrane receptor ROR1; Has very low kinase activity in vitro and is unlikely to function as a tyrosine kinase in vivo. Receptor for ligan |
| | | | RRAS | ENSP00000246792 | Related RAS viral (r-ras) oncogene homolog; Regulates the organization of the actin cytoskeleton. With OSPBL3, modulates integrin beta-1 (ITGB1) activity; Belongs to the sm |
| | | | S100A8 | ENSP00000357722 | Migration inhibitory factor-related protein 8; S100A8 is a calcium- and zinc-binding protein which plays a prominent role in the regulation of inflammatory processes and imr |
| | | | S100A9 | ENSP00000357727 | Migration inhibitory factor-related protein 14; S100A9 is a calcium- and zinc-binding protein which plays a prominent role in the regulation of inflammatory processes and in |
| | | | S1PR1 | ENSP00000305416 | Endothelial differentiation G-protein coupled receptor 1; G-protein coupled receptor for the bioactive lysosphingolipid sphingosine 1-phosphate (S1P) that seems to be coup |
| | | | SEMA3G | ENSP00000231721 | Sema domain, immunoglobulin domain (Ig), short basic domain, secreted, (semaphorin) 3G; Has chemorepulsive activities for sympathetic axons. Ligand of NRP2 (By similari |
| | | | SERPINA1 | ENSP00000416066 | Serpin peptidase inhibitor, clade A (alpha-1 antiproteinase, antitrypsin), member 1; Inhibitor of serine proteases. Its primary target is elastase, but it also has a moderate affi |
| | | | SERPINE1 | ENSP00000223095 | Serpin peptidase inhibitor, clade E (nexin, plasminogen activator inhibitor type 1), member 1; Serine protease inhibitor. This inhibitor acts as 'bait' for tissue plasminogen act |
| | | | SIGIRR | ENSP00000403104 | Single immunoglobulin and toll-interleukin 1 receptor (TIR) domain; Acts as a negative regulator of the Toll-like and IL-1R receptor signaling pathways. Attenuates the recruit |
| | | | SOCS1 | ENSP00000329418 | Suppressor of cytokine signaling 1; SOCS family proteins form part of a classical negative feedback system that regulates cytokine signal transduction. SOCS1 is involved in ne |
| | | | SOCS3 | ENSP00000330341 | Suppressor of cytokine signaling 3; SOCS family proteins form part of a classical negative feedback system that regulates cytokine signal transduction. SOCS3 is involved in ne |
| | | | STAT3 | ENSP00000264657 | Signal transducer and activator of transcription 3 (acute-phase response factor); Signal transducer and transcription activator that mediates cellular responses to interleukins |
| | | | TFRC | ENSP00000353224 | Transferrin receptor protein 1; Cellular uptake of iron occurs via receptor-mediated endocytosis of ligand-occupied transferrin receptor into specialized endosomes. Endoso |
| | | | TGFB1 | ENSP00000221930 | Transforming growth factor, beta 1; Multifunctional protein that controls proliferation, differentiation and other functions in many cell types. Many cells synthesize TGFB1 a |
| | | | TGFB2 | ENSP00000355896 | Glioblastoma-derived T-cell suppressor factor; TGF-beta 2 has suppressive effects on interleukin-2 dependent T-cell growth; Endogenous ligands |
| | | | TGFB3 | ENSP00000238682 | Transforming growth factor, beta 3; Involved in embryogenesis and cell differentiation; Belongs to the TGF-beta family. |
| | | | TGFBR3 | ENSP00000212355 | Transforming growth factor beta receptor type 3; Binds to TGF-beta. Could be involved in capturing and retaining TGF-beta for presentation to the signaling receptors; Prote |

| | | | | | |
|---|---|---|---|---|---|
| | | | TGIF1 | ENSP00000327959 | TGFB-induced factor homeobox 1; Binds to a retinoid X receptor (RXR) responsive element from the cellular retinol-binding protein II promoter (CRBPII- RXRE). Inhibits the 9 |
| | | | TGIF2 | ENSP00000362981 | TGF-beta-induced transcription factor 2; Transcriptional repressor, which probably repress transcription by binding directly the 5'-CTGTCAA-3' DNA sequence or by interactir |
| | | | TGM2 | ENSP00000355330 | Protein-glutamine gamma-glutamyltransferase 2; Catalyzes the cross-linking of proteins and the conjugation of polyamines to proteins; Transglutaminases |
| | | | TIE1 | ENSP00000361554 | Tyrosine kinase with immunoglobulin-like and EGF-like domains 1; Transmembrane tyrosine-protein kinase that may modulate TEK/TIE2 activity and contribute to the regula |
| | | | TIMP1 | ENSP00000218388 | Tissue inhibitor of metalloproteinases 1; Metalloproteinase inhibitor that functions by forming one to one complexes with target metalloproteinases, such as collagenases, a |
| | | | TLR4 | ENSP00000363089 | Toll-like receptor 4; Cooperates with LY96 and CD14 to mediate the innate immune response to bacterial lipopolysaccharide (LPS). Acts via MYD88, TIRAP and TRAF6, leading |
| | | | TNFAIP3 | ENSP00000481570 | Tumor necrosis factor, alpha-induced protein 3; Ubiquitin-editing enzyme that contains both ubiquitin ligase and deubiquitinase activities. Involved in immune and inflamma |
| | | | TNFRSF10D | ENSP00000310263 | Tumor necrosis factor receptor superfamily, member 10d, decoy with truncated death domain; Receptor for the cytotoxic ligand TRAIL. Contains a truncated death domain a |
| | | | TNFRSF1A | ENSP00000162749 | Tumor necrosis factor receptor superfamily, member 1A; Receptor for TNFSF2/TNF-alpha and homotrimeric TNFSF1/lymphotoxin-alpha. The adapter molecule FADD recruit |
| | | | TNFRSF1B | ENSP00000365435 | Tumor necrosis factor receptor superfamily, member 1B; Receptor with high affinity for TNFSF2/TNF-alpha and approximately 5-fold lower affinity for homotrimeric TNFSF1, |
| | | | TNFSF13 | ENSP00000343505 | Tumor necrosis factor (ligand) superfamily, member 13; Cytokine that binds to TNFRSF13B/TACI and to TNFRSF17/BCMA. Plays a role in the regulation of tumor cell growth. |
| | | | TNFSF14 | ENSP00000469049 | Tumor necrosis factor (ligand) superfamily, member 14; Cytokine that binds to TNFRSF3/LTBR. Binding to the decoy receptor TNFRSF6B modulates its effects. Activates NFKE |
| | | | TXNIP | ENSP00000462521 | Vitamin D3 up-regulated protein 1; May act as an oxidative stress mediator by inhibiting thioredoxin activity or by limiting its bioavailability. Interacts with COPS5 and restor |
| | | | TTYH1 | ENSP00000365714 | Protein tweety homolog 1; Probable chloride channel. May be involved in cell adhesion (By similarity). |
| | | | VCAM1 | ENSP00000294728 | Vascular cell adhesion molecule 1; Important in cell-cell recognition. Appears to function in leukocyte-endothelial cell adhesion. Interacts with integrin alpha-4/beta-1 (ITGA |
| | | | VEGFB | ENSP00000311127 | Vascular endothelial growth factor B; Growth factor for endothelial cells. VEGF-B167 binds heparin and neuropilin-1 whereas the binding to neuropilin-1 of VEGF-B186 is reg |
| | | | VIM | ENSP00000446007 | Vimentin; Vimentins are class-III intermediate filaments found in various non-epithelial cells, especially mesenchymal cells. Vimentin is attached to the nucleus, endoplasmic |
| | | | YES1 | ENSP00000462468 | YES proto-oncogene 1, Src family tyrosine kinase; Non-receptor protein tyrosine kinase that is involved in the regulation of cell growth and survival, apoptosis, cell-cell adhes |
| | | | ZC3H12A | ENSP00000362179 | Monocyte chemotactic protein-induced protein 1; Endoribonuclease involved in various biological functions such as cellular inflammatory response and immune homeostasi |
| | | | ZFHX3 | ENSP00000268489 | Alpha-fetoprotein enhancer-binding protein; Transcriptional regulator which can act as an activator or a repressor. Inhibits the enhancer element of the AFP gene by binding |
| | | | ZNF703 | ENSP00000332325 | Zinc finger elbow-related proline domain protein 1; Transcriptional corepressor which does not bind directly to DNA and may regulate transcription through recruitment of |
| 2 | 55 | Regulation of transcription by RNA polymerase II, Anatomical structure morphogenesis, Nervous system development | ASCL1 | ENSP00000266744 | Achaete-scute family bHLH transcription factor 1; Transcription factor that plays a key role in neuronal differentiation: acts as a pioneer transcription factor, accessing closed |
| | | | BCL6B | ENSP00000293805 | B-cell CLL/lymphoma 6 member B protein; Acts as a sequence-specific transcriptional repressor in association with BCL6. May function in a narrow stage or be related to som |
| | | | BMP7 | ENSP00000379204 | Bone morphogenetic protein 7; Induces cartilage and bone formation. May be the osteoinductive factor responsible for the phenomenon of epithelial osteogenesis. Plays a |
| | | | CERS1 | ENSP00000485582 | Embryonic growth/differentiation factor 1; May mediate cell differentiation events during embryonic development; CERS class homeoboxes |
| | | | DLL4 | ENSP00000249749 | Drosophila Delta homolog 4; Involved in the Notch signaling pathway as Notch ligand. Activates NOTCH1 and NOTCH4. Involved in angiogenesis; negatively regulates endoth |
| | | | EBF1 | ENSP00000322898 | Transcription factor COE1; Transcriptional activator which recognizes variations of the palindromic sequence 5'-ATTCCCNNGGGAATT-3'. |
| | | | EMX2 | ENSP00000450962 | Empty spiracles-like protein 2; Transcription factor, which in cooperation with EMX2, acts to generate the boundary between the roof and archipallium in the developing bra |
| | | | ENHO | ENSP00000382675 | Energy homeostasis-associated protein; Involved in the regulation of glucose homeostasis and lipid metabolism. |
| | | | EVC | ENSP00000264956 | Ellis-van Creveld syndrome protein; Component of the EvC complex that positively regulates ciliary Hedgehog (Hh) signaling. Involved in endochondral growth and skeletal d |
| | | | EYA1 | ENSP00000342626 | EYA transcriptional coactivator and phosphatase 1; Functions both as protein phosphatase and as transcriptional coactivator for SIX1, and probably also for SIX2, SIX4 and SI |
| | | | FLI1 | ENSP00000433488 | Friend leukemia integration 1 transcription factor; Sequence-specific transcriptional activator. Recognizes the DNA sequence 5'-C[CA]GGAAGT-3'; Belongs to the ETS family. |
| | | | FOXC1 | ENSP00000370256 | Forkhead-related transcription factor 3; DNA-binding transcriptional factor that plays a role in a broad range of cellular and developmental processes such as eye, bones, car |
| | | | FOXC2 | ENSP00000326371 | Forkhead box C2 (MFH-1, mesenchyme forkhead 1); Transcriptional activator. Might be involved in the formation of special mesenchymal tissues; Forkhead boxes |
| | | | FOXD1 | ENSP00000481581 | Forkhead-related transcription factor 4; Transcription factor involved in regulation of gene expression in a variety of processes, including formation of positional identity in t |
| | | | GDF1 | ENSP00000247005 | Growth differentiation factor 1; May mediate cell differentiation events during embryonic development; Endogenous ligands |
| | | | GFAP | ENSP00000468500 | Glial fibrillary acidic protein; GFAP, a class-III intermediate filament, is a cell- specific marker that, during the development of the central nervous system, distinguishes astroc |
| | | | GLI2 | ENSP00000390436 | GLI family zinc finger protein 2; Functions as transcription regulator in the hedgehog (Hh) pathway. Functions as transcriptional activator. May also function as transcriptiona |
| | | | GLI3 | ENSP00000379258 | Transcriptional activator GLI3; Has a dual function as a transcriptional activator and a repressor of the sonic hedgehog (Shh) pathway, and plays a role in limb development. T |
| | | | GRIN2C | ENSP00000293190 | Glutamate receptor, ionotropic, N-methyl D-aspartate 2C; Component of NMDA receptor complexes that function as heterotetrameric, ligand-gated ion channels with high c |
| | | | HEPN1 | ENSP00000386143 | Putative cancer susceptibility gene HEPN1 protein; Hepatocellular carcinoma, down-regulated 1 |
| | | | HES1 | ENSP00000232424 | Class B basic helix-loop-helix protein 39; Transcriptional repressor of genes that require a bHLH protein for their transcription. May act as a negative regulator of myogenesis |
| | | | HES6 | ENSP00000272937 | Class B basic helix-loop-helix protein 41; Does not bind DNA itself but suppresses both HES1- mediated N box-dependent transcriptional repression and binding of HES1 to E |
| | | | HEYL | ENSP00000361943 | Hes-related family bHLH transcription factor with YRPW motif-like; Downstream effector of Notch signaling which may be required for cardiovascular development (By simila |
| | | | ID4 | ENSP00000367972 | Inhibitor of DNA binding 4, dominant negative helix-loop-helix protein; Transcriptional regulator (lacking a basic DNA binding domain) which negatively regulates the basic h |
| | | | JAG1 | ENSP00000254958 | Protein jagged-1; Ligand for multiple Notch receptors and involved in the mediation of Notch signaling. May be involved in cell-fate decisions during hematopoiesis. Seems t |
| | | | KLF2 | ENSP00000248071 | Lung krueppel-like factor; Transcription factor that binds to the CACCC box in the promoter of target genes such as HBB/beta globin or NOV and activates their transcription; |
| | | | LFNG | ENSP00000222725 | LFNG O-fucosylpeptide 3-beta-N-acetylglucosaminyltransferase; Glycosyltransferase that initiates the elongation of O- linked fucose residues attached to EGF-like repeats in |
| | | | MAML2 | ENSP00000434552 | Mastermind-like 2 (Drosophila); Acts as a transcriptional coactivator for NOTCH proteins. Has been shown to amplify NOTCH-induced transcription of HES1. Potentiates activ |
| | | | MBP | ENSP00000380958 | Myelin membrane encephalitogenic protein; The classic group of MBP isoforms (isoform 4-isoform 14) are with PLP the most abundant protein components of the myelin m |
| | | | MFNG | ENSP00000349490 | MFNG O-fucosylpeptide 3-beta-N-acetylglucosaminyltransferase; Glycosyltransferase that initiates the elongation of O- linked fucose residues attached to EGF-like repeats in |
| | | | MOBP | ENSP00000312293 | Myelin-associated oligodendrocyte basic protein |
| | | | MSX1 | ENSP00000372170 | Msh homeobox 1-like protein; Acts as a transcriptional repressor. May play a role in limb-pattern formation. Acts in cranofacial development and specifically in odontogenes |
| | | | NES | ENSP00000357206 | Nestin; Required for brain and eye development. Promotes the disassembly of phosphorylated vimentin intermediate filaments (IF) during mitosis and may play a role in the |
| | | | NKX3-1 | ENSP00000370253 | Homeobox protein NK-3 homolog A; Transcription factor, which binds preferentially the consensus sequence 5'-TAAGT[AG]-3' and can behave as a transcriptional repressor. |
| | | | NOTCH1 | ENSP00000277541 | Translocation-associated notch protein TAN-1; Functions as a receptor for membrane-bound ligands Jagged1, Jagged2 and Delta1 to regulate cell-fate determination. Upon l |
| | | | NOTCH2 | ENSP00000256646 | Neurogenic locus notch homolog protein 2; Functions as a receptor for membrane-bound ligands Jagged1, Jagged2 and Delta1 to regulate cell-fate determination. Upon liga |
| | | | NOTCH3 | ENSP00000263388 | Neurogenic locus notch homolog protein 3; Functions as a receptor for membrane-bound ligands Jagged1, Jagged2 and Delta1 to regulate cell-fate determination. Upon liga |
| | | | NOTCH4 | ENSP00000364163 | Neurogenic locus notch homolog protein 4; Functions as a receptor for membrane-bound ligands Jagged1, Jagged2 and Delta1 to regulate cell-fate determination. Upon liga |
| | | | NTRK2 | ENSP00000277120 | Neurotrophic tyrosine kinase, receptor, type 2; Receptor tyrosine kinase involved in the development and the maturation of the central and the peripheral nervous systems |
| | | | NTSR2 | ENSP00000303686 | Levocabastine-sensitive neurotensin receptor; Receptor for the tridecapeptide neurotensin. It is associated with G proteins that activate a phosphatidylinositol- calcium seco |
| | | | OLIG1 | ENSP00000371785 | Class E basic helix-loop-helix protein 21; Promotes formation and maturation of oligodendrocytes, especially within the brain. Cooperates with OLIG2 to establish the pMN d |
| | | | PPP1R1B | ENSP00000254079 | Protein phosphatase 1, regulatory (inhibitor) subunit 1B; Inhibitor of protein-phosphatase 1. |
| | | | RHOBTB3 | ENSP00000369318 | Rho-related BTB domain-containing protein 3; Rab9-regulated ATPase required for endosome to Golgi transport. Involved in transport vesicle docking at the Golgi complex, |
| | | | S100B | ENSP00000291700 | S100 calcium binding protein B; Weakly binds calcium but binds zinc very tightly- distinct binding sites with different affinities exist for both ions on each monomer. Physiolo |
| | | | SIX1 | ENSP00000247182 | Sine oculis homeobox homolog 1; Transcription factor that is involved in the regulation of cell proliferation, apoptosis and embryonic development. Plays an important role |
| | | | SLC39A12 | ENSP00000366586 | Solute carrier family 39 (zinc transporter), member 12; Acts as a zinc-influx transporter (Potential). May be partly involved in the outbreak of schizophrenia; Belongs to the ZI |
| | | | SMO | ENSP00000249373 | Smoothened, frizzled class receptor; G protein-coupled receptor that probably associates with the patched protein (PTCH) to transduce the hedgehog's proteins signal. Bindi |
| | | | SOX17 | ENSP00000297316 | SRY (sex determining region Y)-box 17; Acts as transcription regulator that binds target promoter DNA and bends the DNA. Binds to the sequences 5'- AACAAT-'3 or 5'-AACA |
| | | | SOX2 | ENSP00000323588 | SRY (sex determining region Y)-box 2; Transcription factor that forms a trimeric complex with OCT4 on DNA and controls the expression of a number of genes involved in em |
| | | | SOX9 | ENSP00000245479 | SRY (sex determining region Y)-box 9; Transcriptional regulator. Binds to the COL2A1 promoter and activates COL2A1 expression, as part of a complex with ZNF219 (By simila |
| | | | TBX2 | ENSP00000240328 | T-box transcription factor TBX2; Involved in the transcriptional regulation of genes required for mesoderm differentiation. Probably plays a role in limb pattern formation. A |
| | | | TBX3 | ENSP00000257566 | T-box transcription factor TBX3; Transcriptional repressor involved in developmental processes. Probably plays a role in limb pattern formation. Acts as a negative regulator |
| | | | TP53INP2 | ENSP00000363943 | Tumor protein p53 inducible nuclear protein 2; Dual regulator of transcription and autophagy. Positively regulates autophagy and is required for autophagosome formation |
| | | | ZEB2 | ENSP00000454157 | Zinc finger E-box binding homeobox 2; Transcriptional inhibitor that binds to DNA sequence 5'- CACCT-3' in different promoters. Represses transcription of E- cadherin; ZF cl |
| | | | ZIC5 | ENSP00000267294 | Zinc finger protein of the cerebellum 5; Essential for neural crest development, converting cells from an epidermal fate to a neural crest cell fate. Binds to DNA (By similarity |
| 3 | 39 | Innate immune response, Type I interferon signaling pathway, Response to interferon-gamma | BST2 | ENSP00000252593 | Bone marrow stromal cell antigen 2; IFN-induced antiviral host restriction factor which efficiently blocks the release of diverse mammalian enveloped viruses by directly teth |
| | | | CLEC2B | ENSP00000228438 | C-type lectin domain family 2 member B |
| | | | DDX60 | ENSP00000377344 | DEAD (Asp-Glu-Ala-Asp) box polypeptide 60; Positively regulates DDX58/RIG-I- and IFIH1/MDA5- dependent type I interferon and interferon inducible gene expression in res |
| | | | EPSTI1 | ENSP00000318982 | Epithelial stromal interaction 1 |
| | | | FAM89A | ENSP00000355614 | Family with sequence similarity 89 member A; Belongs to the FAM89 family. |
| | | | GBP1 | ENSP00000359504 | Guanylate binding protein 1, interferon-inducible; Hydrolyzes GTP to GMP in 2 consecutive cleavage reactions. Exhibits antiviral activity against influenza virus. Promote oxid |
| | | | GBP2 | ENSP00000359497 | Guanylate binding protein 2, interferon-inducible; Hydrolyzes GTP to GMP in 2 consecutive cleavage reactions, but the major reaction product is GDP. Exhibits antiviral activi |
| | | | GLIS2 | ENSP00000262366 | Neuronal Krueppel-like protein; Can act either as a transcriptional repressor or as a transcriptional activator, depending on the cell context. Acts as a repressor of the Hedge |
| | | | HERC5 | ENSP00000264350 | HECT and RLD domain containing E3 ubiquitin protein ligase 5; Major E3 ligase for ISG15 conjugation. Acts as a positive regulator of innate antiviral response in cells induced |
| | | | HLA-B | ENSP00000399168 | HLA class I histocompatibility antigen, B-7 alpha chain; Involved in the presentation of foreign antigens to the immune system; C1-set domain containing |
| | | | HLA-E | ENSP00000365817 | HLA class I histocompatibility antigen, alpha chain E; Preferably binds to a peptide derived from the signal sequence of most HLA-A, -B, -C and -G molecules; Belongs to the |
| | | | IFI27 | ENSP00000483430 | Interferon alpha-inducible protein 27, mitochondrial; Promotes cell death. Mediates IFN-induced apoptosis characterized by a rapid and robust release of cytochrome C fron |
| | | | IFI30 | ENSP00000384886 | Gamma-interferon-inducible lysosomal thiol reductase; Lysosomal thiol reductase that can reduce protein disulfide bonds. May facilitate the complete unfolding of proteins |
| | | | IFI35 | ENSP00000395590 | Interferon-induced 35 kDa protein; Not yet known. |
| | | | IFI44L | ENSP00000359787 | Interferon-induced protein 44-like; Exhibits a low antiviral activity against hepatitis C virus; Belongs to the IFI44 family. |
| | | | IFITM1 | ENSP00000386187 | Interferon induced transmembrane protein 1; IFN-induced antiviral protein which inhibits the entry of viruses to the host cell cytoplasm, permitting endocytosis, but preven |
| | | | IFITM2 | ENSP00000484689 | Interferon induced transmembrane protein 2; IFN-induced antiviral protein which inhibits the entry of viruses to the host cell cytoplasm, permitting endocytosis, but preven |
| | | | IFITM3 | ENSP00000382707 | Interferon induced transmembrane protein 3; IFN-induced antiviral protein which disrupts intracellular cholesterol homeostasis. Inhibits the entry of viruses to the host cell |
| | | | IRF1 | ENSP00000245414 | Interferon regulatory factor 1; Transcriptional regulator which displays a remarkable functional diversity in the regulation of cellular responses. These include the regulation |
| | | | IRF7 | ENSP00000380697 | Interferon regulatory factor 7; Key transcriptional regulator of type I interferon (IFN)-dependent immune responses and plays a critical role in the innate immune response a |
| | | | ISG20 | ENSP00000306565 | Promyelocytic leukemia nuclear body-associated protein ISG20; Interferon-induced antiviral exoribonuclease that acts on single-stranded RNA and also has minor activity to |
| | | | LILRB1 | ENSP00000315997 | Leukocyte immunoglobulin-like receptor, subfamily B (with TM and ITIM domains), member 1; Receptor for class I MHC antigens. Recognizes a broad spectrum of HLA-A, HL |
| | | | MAVS | ENSP00000401980 | Interferon beta promoter stimulator protein 1; Required for innate immune defense against viruses. Acts downstream of DDX58/RIG-I and IFIH1/MDA5, which detect intrace |
| | | | MICA | ENSP00000413079 | MHC class I polypeptide-related sequence A; Seems to have no role in antigen presentation. Acts as a stress-induced self-antigen that is recognized by gamma delta T- cells. |
| | | | MR1 | ENSP00000477563 | Major histocompatibility complex class I-related gene protein; Antigen-presenting molecule specialized in presenting microbial vitamin B metabolites. Involved in the develo |
| | | | MX2 | ENSP00000333657 | Interferon-regulated resistance GTP-binding protein MxB; Interferon-induced dynamin-like GTPase with potent antiviral activity against human immunodeficiency virus type |
| | | | NLRC5 | ENSP00000262510 | Nucleotide-binding oligomerization domain protein 27; Probable regulator of the NF-kappa-B and type I interferon signaling pathways. May also regulate the type II interfer |
| | | | NWD1 | ENSP00000428579 | NACHT domain- and WD repeat-containing protein 1; May play a role in the control of androgen receptor (AR) protein steady-state levels; WD repeat domain containing |
| | | | OAS3 | ENSP00000228928 | 2'-5'-oligoadenylate synthetase 3, 100kDa; Interferon-induced, dsRNA-activated antiviral enzyme which plays a critical role in cellular innate antiviral response. In addition, it |
| | | | PARP12 | ENSP00000263549 | ADP-ribosyltransferase diphtheria toxin-like 12; poly(ADP-ribose) polymerase family member 12 |
| | | | PARP14 | ENSP00000418194 | ADP-ribosyltransferase diphtheria toxin-like 8; ADP-ribosyltransferase. By mono-ADP-ribosylating STAT1 at 'Glu-657' and 'Glu-705' and thus decreasing STAT1 phosphorylatio |
| | | | PARP9 | ENSP00000353512 | ADP-ribosyltransferase diphtheria toxin-like 9; ADP-ribosyltransferase which, in association with E3 ligase DTX3L, plays a role in DNA damage repair and in immune response |
| | | | PSMB8 | ENSP00000364016 | Proteasome (prosome, macropain) subunit, beta type, 8; The proteasome is a multicatalytic proteinase complex which is characterized by its ability to cleave peptides with A |
| | | | RNF213 | ENSP00000464087 | ALK lymphoma oligomerization partner on chromosome 17; E3 ubiquitin-protein ligase involved in angiogenesis. Involved in the non-canonical Wnt signaling pathway in vas |
| | | | SAMD9L | ENSP00000326247 | Sterile alpha motif domain-containing protein 9-like; May be involved in endosome fusion. Mediates down- regulation of growth factor signaling via internalization of growt |
| | | | SP110 | ENSP00000258381 | Transcriptional coactivator Sp110; Transcription factor. May be a nuclear hormone receptor coactivator. Enhances transcription of genes with retinoic acid response elemen |
| | | | TAP1 | ENSP00000346206 | Transporter 1, ATP-binding cassette, sub-family B (MDR/TAP); Involved in the transport of antigens from the cytoplasm to the endoplasmic reticulum for association with M |
| | | | TRIM22 | ENSP00000369299 | RING-type E3 ubiquitin transferase TRIM22; Interferon-induced antiviral protein involved in cell innate immunity. The antiviral activity could in part be mediated by TRIM22- |
| | | | XAF1 | ENSP00000354822 | XIAP associated factor 1; Seems to function as a negative regulator of members of the IAP (inhibitor of apoptosis protein) family. Inhibits anti- caspase activity of BIRC4. Indu |
| 4 | 15 | Oxidative phosphorylation, Mitochondrial ATP synthesis coupled electron transport, Mitochondrial electron transport, NADH to ubiquinone | COX4I2 | ENSP00000365243 | Cytochrome c oxidase subunit 4 isoform 2, mitochondrial; This protein is one of the nuclear-coded polypeptide chains of cytochrome c oxidase, the terminal oxidase in mito |
| | | | MT-ATP6 | ENSP00000354632 | Mitochondrially encoded ATP synthase 6; Mitochondrial membrane ATP synthase (F(1)F(0) ATP synthase or Complex V) produces ATP from ADP in the presence of a proton |
| | | | MT-ATP8 | ENSP00000355265 | Mitochondrially encoded ATP synthase 8; Mitochondrial membrane ATP synthase (F(1)F(0) ATP synthase or Complex V) produces ATP from ADP in the presence of a proton |
| | | | MT-CO1 | ENSP00000354499 | Mitochondrially encoded cytochrome c oxidase I; Cytochrome c oxidase is the component of the respiratory chain that catalyzes the reduction of oxygen to water. Subunits |
| | | | MT-CO2 | ENSP00000354876 | Mitochondrially encoded cytochrome c oxidase II; Cytochrome c oxidase is the component of the respiratory chain that catalyzes the reduction of oxygen to water. Subunits |
| | | | MT-CO3 | ENSP00000354982 | Mitochondrially encoded cytochrome c oxidase III; Subunits I, II and III form the functional core of the enzyme complex; Mitochondrial complex IV: cytochrome c oxidase su |
| | | | MT-CYB | ENSP00000354554 | Ubiquinol-cytochrome-c reductase complex cytochrome b subunit; Component of the ubiquinol-cytochrome c reductase complex (complex III or cytochrome b-c1 complex) |
| | | | MT-ND1 | ENSP00000354687 | Mitochondrially encoded NADH dehydrogenase 1; Core subunit of the mitochondrial membrane respiratory chain NADH dehydrogenase (Complex I) that is believed to belo |
| | | | MT-ND2 | ENSP00000355046 | Mitochondrially encoded NADH dehydrogenase 2; Core subunit of the mitochondrial membrane respiratory chain NADH dehydrogenase (Complex I) that is believed to belo |
| | | | MT-ND3 | ENSP00000355206 | Mitochondrially encoded NADH dehydrogenase 3; Core subunit of the mitochondrial membrane respiratory chain NADH dehydrogenase (Complex I) that is believed to belo |
| | | | MT-ND4 | ENSP00000354961 | Mitochondrially encoded NADH dehydrogenase 4; Core subunit of the mitochondrial membrane respiratory chain NADH dehydrogenase (Complex I) that is believed to belo |
| | | | MT-ND4L | ENSP00000354728 | Mitochondrially encoded NADH dehydrogenase 4L; Core subunit of the mitochondrial membrane respiratory chain NADH dehydrogenase (Complex I) that is believed to belo |
| | | | MT-ND5 | ENSP00000354813 | Mitochondrially encoded NADH dehydrogenase 5; Core subunit of the mitochondrial membrane respiratory chain NADH dehydrogenase (Complex I) that is believed to belo |
| | | | MT-ND6 | ENSP00000354665 | Mitochondrially encoded NADH dehydrogenase 6; Core subunit of the mitochondrial membrane respiratory chain NADH dehydrogenase (Complex I) that is believed to belo |
| | | | NDUFA4L2 | ENSP00000377411 | NADH dehydrogenase [ubiquinone] 1 alpha subcomplex subunit 4-like 2; NDUFA4, mitochondrial complex associated like 2 |
| 5 | 13 | - | AKAP9 | ENSP00000348573 | Centrosome- and Golgi-localized PKN-associated protein; Scaffolding protein that assembles several protein kinases and phosphatases on the centrosome and Golgi apparat |
| | | | BAZ1A | ENSP00000353458 | Williams syndrome transcription factor-related chromatin-remodeling factor 180; Component of the ACF complex, an ATP-dependent chromatin remodeling complex, that r |
| | | | BAZ2B | ENSP00000376534 | Bromodomain adjacent to zinc finger domain protein 2B; May play a role in transcriptional regulation interacting with ISWI; Belongs to the WAL family. |
| | | | BPTF | ENSP00000307208 | Bromodomain and PHD finger-containing transcription factor; Histone-binding component of NURF (nucleosome-remodeling factor), a complex which catalyzes ATP-depend |

| | | | | | |
|---|---|---|---|---|---|
| | | | CNTLN | ENSP00000370021 | Centlein, centrosomal protein; Required for centrosome cohesion and recruitment of CEP68 to centrosomes. |
| | | | KCNE4 | ENSP00000281830 | Potassium voltage-gated channel, Isk-related family, member 4; Ancillary protein that assembles as a beta subunit with a voltage-gated potassium channel complex of pore-f |
| | | | LRRCC1 | ENSP00000353538 | Centrosomal leucine-rich repeat and coiled-coil domain-containing protein; Required for the organization of the mitotic spindle. Maintains the structural integrity of centros |
| | | | NEDD1 | ENSP00000451211 | Neural precursor cell expressed developmentally down-regulated protein 1; Required for mitosis progression. Promotes the nucleation of microtubules from the spindle; WI |
| | | | PHEX | ENSP00000368682 | Phosphate regulating endopeptidase homolog, X-linked; Probably involved in bone and dentin mineralization and renal phosphate reabsorption; M13 metallopeptidases |
| | | | PRRC2C | ENSP00000343629 | Proline-rich and coiled-coil-containing protein 2C; Proline rich coiled-coil 2C |
| | | | PTH1R | ENSP00000321999 | Parathyroid hormone/parathyroid hormone-related peptide receptor; Receptor for parathyroid hormone and for parathyroid hormone-related peptide. The activity of this r |
| | | | RAB34 | ENSP00000413156 | RAB34, member RAS oncogene family; Protein transport. Involved in the redistribution of lysosomes to the peri-Golgi region (By similarity). Plays a role in the maturation of |
| | | | RBBP6 | ENSP00000317872 | P53-associated cellular protein of testis; E3 ubiquitin-protein ligase which promotes ubiquitination of YBX1, leading to its degradation by the proteasome. May play a role as |
| 6 | 12 | RNA splicing, mRNA processing, mRNA export from nucleus | CASC3 | ENSP00000264645 | Cancer susceptibility candidate gene 3 protein; Core component of the splicing-dependent multiprotein exon junction complex (EJC) deposited at splice junctions on mRNAs |
| | | | GRAP | ENSP00000284154 | GRB2-related adapter protein; Couples signals from receptor and cytoplasmic tyrosine kinases to the Ras signaling pathway; SH2 domain containing |
| | | | LUC7L3 | ENSP00000425092 | Cisplatin resistance-associated-overexpressed protein; Binds cAMP regulatory element DNA sequence. May play a role in RNA splicing; Belongs to the Luc7 family. |
| | | | NKTR | ENSP00000232978 | Natural-killer cells cyclophilin-related protein; Component of a putative tumor-recognition complex. Involved in the function of NK cells; Cyclophilin peptidylprolyl isomerase |
| | | | PNISR | ENSP00000358242 | Serine/arginine-rich-splicing regulatory protein 130; PNN interacting serine and arginine rich protein; Belongs to the splicing factor SR family. |
| | | | PRPF38B | ENSP00000359042 | pre-mRNA processing factor 38B; May be required for pre-mRNA splicing. |
| | | | SLTM | ENSP00000369887 | Modulator of estrogen-induced transcription; When overexpressed, acts as a general inhibitor of transcription that eventually leads to apoptosis; RNA binding motif containi |
| | | | SREK1 | ENSP00000334538 | Serine/arginine-rich-splicing regulatory protein 86; Participates in the regulation of alternative splicing by modulating the activity of other splice facors. Inhibits the splicing a |
| | | | SRRM1 | ENSP00000326261 | SR-related nuclear matrix protein of 160 kDa; Part of pre- and post-splicing multiprotein mRNP complexes. Involved in numerous pre-mRNA processing events. Promotes co |
| | | | SRSF11 | ENSP00000359988 | Splicing factor, arginine/serine-rich 11; May function in pre-mRNA splicing; RNA binding motif containing |
| | | | THOC2 | ENSP00000245838 | THO complex subunit 2; Required for efficient export of polyadenylated RNA and spliced mRNA. Acts as component of the THO subcomplex of the TREX complex which is th |
| | | | ZFC3H1 | ENSP00000368017 | Zinc finger C3H1 domain-containing protein; Subunit of the trimeric poly(A) tail exosome targeting (PAXT) complex, a complex that directs a subset of long and polyadenylat |
| 7 | 11 | Neurotransmitter uptake, Sodium ion transport, L-glutamate import across plasma membrane | ALDH4A1 | ENSP00000364490 | Delta-1-pyrroline-5-carboxylate dehydrogenase, mitochondrial; Irreversible conversion of delta-1-pyrroline-5- carboxylate (P5C), derived either from proline or ornithine, to |
| | | | GLUL | ENSP00000307900 | Glutamate--ammonia ligase; This enzyme has 2 functions: it catalyzes the production of glutamine and 4-aminobutanoate (gamma-aminobutyric acid, GABA), the latter in a p |
| | | | SLC1A3 | ENSP00000265113 | Solute carrier family 1 (glial high affinity glutamate transporter), member 3; Sodium-dependent, high-affinity amino acid transporter that mediates the uptake of L-glutamate |
| | | | SLC1A7 | ENSP00000478639 | Solute carrier family 1 member 7 |
| | | | SLC38A3 | ENSP00000481301 | Sodium-coupled neutral amino acid transporter 3; Sodium-dependent amino acid/proton antiporter. Mediates electrogenic cotransport of glutamine and sodium ions in exc |
| | | | SLC38A5 | ENSP00000471683 | Sodium-coupled neutral amino acid transporter 5; Functions as a sodium-dependent amino acid transporter which countertransport protons. Mediates the saturable, pH- se |
| | | | SLC6A11 | ENSP00000254488 | Solute carrier family 6 (neurotransmitter transporter), member 11; Terminates the action of GABA by its high affinity sodium-dependent reuptake into presynaptic terminals |
| | | | SLC6A12 | ENSP00000399136 | Solute carrier family 6 (neurotransmitter transporter), member 12; Transports betaine and GABA. May have a role in regulation of GABAergic transmission in the brain throu |
| | | | SLC6A13 | ENSP00000339260 | Solute carrier family 6 (neurotransmitter transporter), member 13; Sodium-dependent GABA and taurine transporter. In presynaptic terminals, regulates GABA signaling ter |
| | | | SLC7A11 | ENSP00000280612 | Solute carrier family 7 (anionic amino acid transporter light chain, xc- system), member 11; Sodium-independent, high-affinity exchange of anionic amino acids with high spe |
| | | | SLC7A5 | ENSP00000261622 | Solute carrier family 7 (amino acid transporter light chain, L system), member 5; Sodium-independent, high-affinity transport of large neutral amino acids such as phenylalan |
| 8 | 10 | G protein-coupled receptor signaling pathway | CYSLTR2 | ENSP00000282018 | G-protein coupled receptor GPCR21; Receptor for cysteinyl leukotrienes. The response is mediated via a G-protein that activates a phosphatidylinositol- calcium second mess |
| | | | F2RL3 | ENSP00000248076 | Coagulation factor II (thrombin) receptor-like 3; Receptor for activated thrombin or trypsin coupled to G proteins that stimulate phosphoinositide hydrolysis. May play a role |
| | | | GNG11 | ENSP00000248564 | Guanine nucleotide-binding protein G(I)/G(S)/G(O) subunit gamma-11; Guanine nucleotide-binding proteins (G proteins) are involved as a modulator or transducer in variou |
| | | | GNG12 | ENSP00000360021 | Guanine nucleotide-binding protein G(I)/G(S)/G(O) subunit gamma-12; Guanine nucleotide-binding proteins (G proteins) are involved as a modulator or transducer in variou |
| | | | GNG5 | ENSP00000359675 | Guanine nucleotide-binding protein G(I)/G(S)/G(O) subunit gamma-5; Guanine nucleotide-binding proteins (G proteins) are involved as a modulator or transducer in various |
| | | | KCNJ16 | ENSP00000465295 | Potassium inwardly-rectifying channel, subfamily J, member 16; Inward rectifier potassium channels are characterized by a greater tendency to allow potassium to flow into |
| | | | PRKCH | ENSP00000329127 | Protein kinase C eta type; Calcium-independent, phospholipid- and diacylglycerol (DAG)-dependent serine/threonine-protein kinase that is involved in the regulation of cell |
| | | | PTGER1 | ENSP00000292513 | Prostaglandin E receptor 1 (subtype EP1), 42kDa; Receptor for prostaglandin E2 (PGE2). The activity of this receptor is mediated by G(q) proteins which activate a phosphatid |
| | | | RGR | ENSP00000352427 | RPE-retinal G protein-coupled receptor; Receptor for all-trans- and 11-cis-retinal. Binds preferentially to the former and may catalyze the isomerization of the chromophore l |
| | | | RGS20 | ENSP00000297313 | Regulator of Gz-selective protein signaling 1; Inhibits signal transduction by increasing the GTPase activity of G protein alpha subunits thereby driving them into their inactive |
| 9 | 10 | Regulation of cellular metabolic process, Regulation of molecular function | BTG2 | ENSP00000290551 | NGF-inducible anti-proliferative protein PC3; Anti-proliferative protein; the function is mediated by association with deadenylase subunits of the CCR4-NOT complex. Activat |
| | | | CEBPD | ENSP00000386165 | CCAAT/enhancer binding protein (C/EBP), delta; Transcription activator that recognizes two different DNA motifs: the CCAAT homology common to many promoters and the |
| | | | DUSP1 | ENSP00000239223 | Mitogen-activated protein kinase phosphatase 1; Dual specificity phosphatase that dephosphorylates MAP kinase MAPK1/ERK2 on both 'Thr-183' and 'Tyr-185', regulating it |
| | | | DUSP5 | ENSP00000358596 | Dual specificity protein phosphatase hVH3; Dual specificity protein phosphatase; active with phosphotyrosine, phosphoserine and phosphothreonine residues. The highest r |
| | | | FOS | ENSP00000306245 | FBJ murine osteosarcoma viral oncogene homolog; Nuclear phosphoprotein which forms a tight but non- covalently linked complex with the JUN/AP-1 transcription factor. I |
| | | | FOSL2 | ENSP00000264716 | Fos-related antigen 2; Controls osteoclast survival and size. As a dimer with JUN, activates LIF transcription. Activates CEBPB transcription in PGE2-activated osteoblasts; Basi |
| | | | GADD45B | ENSP00000215631 | Growth arrest and DNA damage-inducible protein GADD45 beta; Involved in the regulation of growth and apoptosis. Mediates activation of stress-responsive MTK1/MEKK4 |
| | | | JUNB | ENSP00000303315 | Transcription factor jun-B; Transcription factor involved in regulating gene activity following the primary growth factor response. Binds to the DNA sequence 5'-TGA[CG]TCA- |
| | | | MAFF | ENSP00000345393 | V-maf avian musculoaponeurotic fibrosarcoma oncogene homolog F; Interacts with the upstream promoter region of the oxytocin receptor gene. May be a transcriptional e |
| | | | ZFP36 | ENSP00000469647 | Growth factor-inducible nuclear protein NUP475; Zinc-finger RNA-binding protein that destabilizes several cytoplasmic AU-rich element (ARE)-containing mRNA transcripts b |
| 10 | 9 | | ATP10A | ENSP00000349325 | P4-ATPase flippase complex alpha subunit ATP10A; Catalytic component of a P4-ATPase flippase complex which catalyzes the hydrolysis of ATP coupled to the transport of a |
| | | | CDH23 | ENSP00000381768 | Cadherin-related 23; Cadherins are calcium-dependent cell adhesion proteins. They preferentially interact with themselves in a homophilic manner in connecting cells. CDH2 |
| | | | CIB2 | ENSP00000258930 | Calcium and integrin binding family member 2; Calcium-binding protein critical for proper photoreceptor cell maintenance and function. Plays a role in intracellular calcium l |
| | | | EHD4 | ENSP00000220325 | Hepatocellular carcinoma-associated protein 10/11; ATP- and membrane-binding protein that probably controls membrane reorganization/tubulation upon ATP hydrolysis. I |
| | | | GJB6 | ENSP00000348521 | Gap junction protein, beta 6, 30kDa; One gap junction consists of a cluster of closely packed pairs of transmembrane channels, the connexons, through which materials of lo |
| | | | LRTOMT | ENSP00000305742 | Leucine rich transmembrane and O-methyltransferase domain containing; Catalyzes the O-methylation, and thereby the inactivation, of catecholamine neurotransmitters an |
| | | | TMPRSS3 | ENSP00000291532 | Tumor-associated differentially-expressed gene 12 protein; Probable serine protease that plays a role in hearing. Acts as a permissive factor for cochlear hair cell survival an |
| | | | TMPRSS4 | ENSP00000477949 | Transmembrane protease, serine 4; Probable protease. Seems to be capable of activating ENaC (By similarity); Scavenger receptor cysteine rich domain containing |
| | | | TRIOBP | ENSP00000384312 | TRIO and F-actin binding protein; May regulate actin cytoskeletal organization, cell spreading and cell contraction by directly binding and stabilizing filamentous F-actin. The |
| 11 | 8 | | ADH1B | ENSP00000306606 | Alcohol dehydrogenase 1B, beta polypeptide; Belongs to the zinc-containing alcohol dehydrogenase family. |
| | | | ALDH3B1 | ENSP00000473990 | Aldehyde dehydrogenase 3 family, member B1; Oxidizes medium and long chain saturated and unsaturated aldehydes. Metabolizes also benzaldehyde. Low activity towards |
| | | | EPHX1 | ENSP00000480004 | Epoxide hydrolase 1, microsomal (xenobiotic); Biotransformation enzyme that catalyzes the hydrolysis of arene and aliphatic epoxides to less reactive and more water solubl |
| | | | FAH | ENSP00000385080 | Fumarylacetoacetate hydrolase |
| | | | GSTM1 | ENSP00000311469 | Glutathione S-transferase Mu 1; Conjugation of reduced glutathione to a wide number of exogenous and endogenous hydrophobic electrophiles; Soluble glutathione S-trans |
| | | | GSTM2 | ENSP00000241337 | Glutathione S-transferase mu 2 (muscle); Conjugation of reduced glutathione to a wide number of exogenous and endogenous hydrophobic electrophiles; Belongs to the GS |
| | | | GSTM5 | ENSP00000256593 | Glutathione S-transferase Mu 5; Conjugation of reduced glutathione to a wide number of exogenous and endogenous hydrophobic electrophiles; Soluble glutathione S-trans |
| | | | MGST1 | ENSP00000379512 | Microsomal glutathione S-transferase 1; Conjugation of reduced glutathione to a wide number of exogenous and endogenous hydrophobic electrophiles. Has a wide substra |
| 12 | 8 | | C4A | ENSP00000396688 | C3 and PZP-like alpha-2-macroglobulin domain-containing protein 2; Non-enzymatic component of C3 and C5 convertases and thus essential for the propagation of the classi |
| | | | C4B | ENSP00000415941 | C3 and PZP-like alpha-2-macroglobulin domain-containing protein 3; Non-enzymatic component of the C3 and C5 convertases and thus essential for the propagation of the c |
| | | | CFB | ENSP00000416561 | Glycine-rich beta glycoprotein; Factor B which is part of the alternate pathway of the complement system is cleaved by factor D into 2 fragments: Ba and Bb. Bb, a serine pro |
| | | | CFH | ENSP00000356399 | Complement factor H; Factor H functions as a cofactor in the inactivation of C3b by factor I and also increases the rate of dissociation of the C3bBb complex (C3 convertase) a |
| | | | CFHR1 | ENSP00000314299 | Complement factor H-related protein 1; Involved in complement regulation. The dimerized forms have avidity for tissue-bound complement fragments and efficiently compe |
| | | | MASP1 | ENSP00000296280 | Mannan-binding lectin serine peptidase 1 (C4/C2 activating component of Ra-reactive factor); Mannan binding lectin serine peptidase 1 |
| | | | RBMS2 | ENSP00000262031 | RNA binding motif single stranded interacting protein 2 |
| | | | SERPING1 | ENSP00000278407 | Serpin peptidase inhibitor, clade G (C1 inhibitor), member 1; Activation of the C1 complex is under control of the C1- inhibitor. It forms a proteolytically inactive stoichiomet |
| 13 | 6 | | BBOX1 | ENSP00000263182 | Butyrobetaine (gamma), 2-oxoglutarate dioxygenase (gamma-butyrobetaine hydroxylase) 1; Catalyzes the formation of L-carnitine from gamma- butyrobetaine; Belongs to th |
| | | | TRIM34 | ENSP00000422947 | Tripartite motif containing 34; May function as antiviral protein and may contribute to the defense against retroviral infections; Belongs to the TRIM/RBCC family. |
| | | | TRIM38 | ENSP00000349596 | RING-type E3 ubiquitin transferase TRIM38; E3 ubiquitin-protein ligase. Mediates 'Lys-48'-linked polyubiquitination and proteasomal degradation of the critical TLR adapter T |
| | | | TRIM41 | ENSP00000320869 | RING finger-interacting protein with C kinase; Functions as an E3 ligase that catalyzes the ubiquitin- mediated degradation of protein kinase C; Ring finger proteins |
| | | | TRIM5 | ENSP00000369373 | RING-type E3 ubiquitin transferase TRIM5; Capsid-specific restriction factor that prevents infection from non-host-adapted retroviruses. Blocks viral replication early in the li |
| | | | TRIM56 | ENSP00000305161 | RING-type E3 ubiquitin transferase TRIM56; E3 ubiquitin-protein ligase that plays a key role in innate antiviral immunity. In response to pathogen- and host-derived double-s |
| 14 | 8 | | MT1A | ENSP00000478425 | Metallothionein 1A; Metallothioneins have a high content of cysteine residues that bind various heavy metals; these proteins are transcriptionally regulated by both heavy m |
| | | | MT1E | ENSP00000307706 | Metallothionein 1E; Metallothioneins have a high content of cysteine residues that bind various heavy metals; these proteins are transcriptionally regulated by both heavy m |
| | | | MT1F | ENSP00000334872 | Metallothionein 1F; Metallothioneins have a high content of cysteine residues that bind various heavy metals; these proteins are transcriptionally regulated by both heavy m |
| | | | MT1G | ENSP00000391397 | Metallothionein 1G; Metallothioneins have a high content of cysteine residues that bind various heavy metals; these proteins are transcriptionally regulated by both heavy m |
| | | | MT1M | ENSP00000369146 | Metallothionein 1M; Metallothioneins have a high content of cysteine residues that bind various heavy metals; these proteins are transcriptionally regulated by both heavy r |
| | | | MT1X | ENSP00000377995 | Metallothionein 1X; Metallothioneins have a high content of cysteine residues that bind various heavy metals; these proteins are transcriptionally regulated by both heavy m |
| | | | MT2A | ENSP00000245185 | Metallothionein 2A; Metallothioneins have a high content of cysteine residues that bind various heavy metals; these proteins are transcriptionally regulated by both heavy m |
| | | | NQO1 | ENSP00000319788 | NAD(P)H dehydrogenase [quinone] 1; The enzyme apparently serves as a quinone reductase in connection with conjugation reactions of hydroquinons involved in detoxifica |
| 15 | 6 | | GPR146 | ENSP00000380283 | Probable G-protein coupled receptor 146; Orphan receptor; G protein-coupled receptors, Class A orphans |
| | | | MS4A6A | ENSP00000392770 | Membrane-spanning 4-domains, subfamily A, member 6A; May be involved in signal transduction as a component of a multimeric receptor complex; Belongs to the MS4A fa |
| | | | MS4A7 | ENSP00000300184 | Membrane-spanning 4-domains, subfamily A, member 7; May be involved in signal transduction as a component of a multimeric receptor complex; Belongs to the MS4A fam |
| | | | PIK3IP1 | ENSP00000215912 | Phosphoinositide-3-kinase interacting protein 1; Negative regulator of hepatic phosphatidylinositol 3- kinase (PI3K) activity. |
| | | | TMEM176A | ENSP00000417626 | Hepatocellular carcinoma-associated antigen 112; Membrane spanning 4-domains |
| | | | TMEM176B | ENSP00000410269 | Transmembrane protein 176B; May play a role in the process of maturation of dendritic cells. Required for the development of cerebellar granule cells (By similarity); Memb |
| 16 | 7 | | AMOT | ENSP00000361027 | Angiomotin; Plays a central role in tight junction maintenance via the complex formed with ARHGAP17, which acts by regulating the uptake of polarity proteins at tight junct |
| | | | TEAD1 | ENSP00000435233 | TEA domain family member 1 (SV40 transcriptional enhancer factor); Transcription factor which plays a key role in the Hippo signaling pathway, a pathway involved in organ |
| | | | TEAD2 | ENSP00000472109 | Transcriptional enhancer factor TEF-4; Transcription factor which plays a key role in the Hippo signaling pathway, a pathway involved in organ size control and tumor suppre |
| | | | TEAD3 | ENSP00000345772 | Transcriptional enhancer factor TEF-5; Transcription factor which plays a key role in the Hippo signaling pathway, a pathway involved in organ size control and tumor suppre |
| | | | WWC1 | ENSP00000427772 | WW domain-containing protein 1; Probable regulator of the Hippo/SWH (Sav/Wts/Hpo) signaling pathway, a signaling pathway that plays a pivotal role in tumor suppression |
| | | | WWTR1 | ENSP00000419465 | WW domain-containing transcription regulator protein 1; Transcriptional coactivator which acts as a downstream regulatory target in the Hippo signaling pathway that plays |
| | | | YAP1 | ENSP00000478927 | Yes-associated protein YAP65 homolog; Transcriptional regulator which can act both as a coactivator and a corepressor and is the critical downstream regulatory target in th |
| 17 | 6 | | APOL1 | ENSP00000317674 | Apolipoprotein L, 1; May play a role in lipid exchange and transport throughout the body. May participate in reverse cholesterol transport from peripheral cells to the liver. |
| | | | CLU | ENSP00000315130 | Testosterone-repressed prostate message 2; Isoform 1 functions as extracellular chaperone that prevents aggregation of nonnative proteins. Prevents stress- induced aggreg |
| | | | HPR | ENSP00000441828 | Haptoglobin-related protein; Primate-specific plasma protein associated with apolipoprotein L-I (apoL-I)-containing high-density lipoprotein (HDL). This HDL particle, termed |
| | | | LCAT | ENSP00000264005 | Phosphatidylcholine-sterol acyltransferase; Central enzyme in the extracellular metabolism of plasma lipoproteins. Synthesized mainly in the liver and secreted into plasma v |
| | | | PLA1A | ENSP00000273371 | Phosphatidylserine-specific phospholipase A1; Hydrolyzes the ester bond at the sn-1 position of glycerophospholipids and produces 2-acyl lysophospholipids. Hydrolyzes pho |
| | | | PON3 | ENSP00000265627 | Serum paraoxonase/lactonase 3; Has low activity towards the organophosphate paraxon and aromatic carboxylic acid esters. Rapidly hydrolyzes lactones such as statin prodr |
| 18 | 6 | | CLEC1A | ENSP00000326407 | C-type lectin domain family 1 member A |
| | | | CLEC4E | ENSP00000299663 | C-type lectin domain family 4, member E; C-type lectin that functions as cell-surface receptor for a wide variety of ligands such as damaged cells, fungi and mycobacteria. Pla |
| | | | CNTNAP3 | ENSP00000297668 | Contactin associated protein like 3 |
| | | | GLIPR1L2 | ENSP00000448248 | GLI pathogenesis-related 1 like 2; GLI pathogenesis related family 1; Belongs to the CRISP family. |
| | | | SEC14L2 | ENSP00000478755 | Alpha-tocopherol-associated protein; Carrier protein. Binds to some hydrophobic molecules and promotes their transfer between the different cellular sites. Binds with high |
| | | | ZNF98 | ENSP00000350418 | Zinc finger protein F7175; May be involved in transcriptional regulation; Zinc fingers C2H2-type |
| 19 | 6 | | EYA2 | ENSP00000333640 | EYA transcriptional coactivator and phosphatase 2; Functions both as protein phosphatase and as transcriptional coactivator for SIX1, and probably also for SIX2, SIX4 and SI |
| | | | FCGRT | ENSP00000221466 | Fc fragment of IgG, receptor, transporter, alpha; Binds to the Fc region of monomeric immunoglobulins gamma. Mediates the selective uptake of IgG from milk and helps ne |
| | | | FREM2 | ENSP00000280481 | FRAS1 related extracellular matrix protein 2; Extracellular matrix protein required for maintenance of the integrity of the skin epithelium and for maintenance of renal epith |
| | | | IGFBP5 | ENSP00000233813 | Insulin-like growth factor binding protein 5; IGF-binding proteins prolong the half-life of the IGFs and have been shown to either inhibit or stimulate the growth promoting e |
| | | | INHBB | ENSP00000295228 | Activin beta-B chain; Inhibins and activins inhibit and activate, respectively, the secretion of follitropin by the pituitary gland. Inhibins/activins are involved in regulating a nu |
| | | | SIX5 | ENSP00000316842 | DM locus-associated homeodomain protein; Transcription factor that is thought to be involved in regulation of organogenesis. May be involved in determination and mainte |
| 20 | 6 | | MYBPC1 | ENSP00000400908 | C-protein, skeletal muscle slow isoform; Thick filament-associated protein located in the crossbridge region of vertebrate striated muscle a bands. In vitro it binds MHC, F-act |
| | | | MYH15 | ENSP00000273353 | Myosin, heavy chain 15; Muscle contraction; Belongs to the TRAFAC class myosin-kinesin ATPase superfamily. Myosin family. |
| | | | MYL3 | ENSP00000379210 | Myosin light chain 1, slow-twitch muscle B/ventricular isoform; Regulatory light chain of myosin. Does not bind calcium; EF-hand domain containing |
| | | | MYOT | ENSP00000239926 | Myofibrillar titin-like Ig domains protein; Component of a complex of multiple actin cross-linking proteins. Involved in the control of myofibril assembly and stability at the Z |
| | | | MYOZ1 | ENSP00000352272 | Filamin-, actinin- and telethonin-binding protein; Myozenins may serve as intracellular binding proteins involved in linking Z-disk proteins such as alpha-actinin, gamma- filan |
| | | | PGAM2 | ENSP00000297283 | Muscle-specific phosphoglycerate mutase; Interconversion of 3- and 2-phosphoglycerate with 2,3- bisphosphoglycerate as the primer of the reaction. Can also catalyze the re |
| 21 | 6 | | EPB41L5 | ENSP00000263713 | Erythrocyte membrane protein band 4.1 like 5; May contribute to the correct positioning of tight junctions during the establishment of polarity in epithelial cells; FERM dom |
| | | | NXN | ENSP00000337443 | Nucleoredoxin; Functions as a redox-dependent negative regulator of the Wnt signaling pathway, possibly by preventing ubiquitination of DVL3 by the BCR(KLHL12) complex |
| | | | PAIP2B | ENSP00000244221 | Polyadenylate-binding protein-interacting protein 2B; Inhibits translation of capped and polyadenylated mRNAs by displacing PABPC1 from the poly(A) tail; Belongs to the P |
| | | | PARD3B | ENSP00000351618 | Amyotrophic lateral sclerosis 2 chromosomal region candidate gene 19 protein; Putative adapter protein involved in asymmetrical cell division and cell polarization processe |
| | | | RBMS3 | ENSP00000373277 | RNA binding motif, single stranded interacting protein 3; Binds poly(A) and poly(U) oligoribonucleotides; RNA binding motif containing |

| | | | |
|---|---|---|---|
| | | ZMYM5 | ENSP00000372361 | Zinc finger MYM-type protein 5; Functions as a transcriptional regulator; Zinc fingers MYM-type |
| 22 | 6 | ADD3 | ENSP00000348381 | Adducin-like protein 70; Membrane-cytoskeleton-associated protein that promotes the assembly of the spectrin-actin network. Plays a role in actin filament capping. Binds t |
| | | ARHGAP19 | ENSP00000351333 | Rho-type GTPase-activating protein 19; GTPase activator for the Rho-type GTPases by converting them to an inactive GDP-bound state. |
| | | FES | ENSP00000331504 | Feline sarcoma/Fujinami avian sarcoma oncogene homolog; Tyrosine-protein kinase that acts downstream of cell surface receptors and plays a role in the regulation of the a |
| | | NOSTRIN | ENSP00000394051 | Nitric oxide synthase trafficking; F-BAR domain containing |
| | | SCARA3 | ENSP00000301904 | Cellular stress response gene protein; Seems to protect cells by scavenging oxidative molecules or harmful products of oxidation; Scavenger receptors |
| | | SRGAP1 | ENSP00000347198 | SLIT-ROBO Rho GTPase activating protein 1; GTPase-activating protein for RhoA and Cdc42 small GTPases. Together with CDC42 seems to be involved in the pathway mediat |
| 23 | 4 | ADORA2A | ENSP00000336630 | Adenosine A2a receptor; Receptor for adenosine. The activity of this receptor is mediated by G proteins which activate adenylyl cyclase; Belongs to the G-protein coupled re |
| | | ADORA2B | ENSP00000304501 | Adenosine A2b receptor; Receptor for adenosine. The activity of this receptor is mediated by G proteins which activate adenylyl cyclase. |
| | | P2RY2 | ENSP00000310305 | Purinergic receptor P2Y, G-protein coupled, 2; Receptor for ATP and UTP coupled to G-proteins that activate a phosphatidylinositol-calcium second messenger system. The a |
| | | RAPGEF3 | ENSP00000395708 | Rap1 guanine-nucleotide-exchange factor directly activated by cAMP; Guanine nucleotide exchange factor (GEF) for RAP1A and RAP2A small GTPases that is activated by bin |
| 24 | 5 | CCDC103 | ENSP00000391692 | Coiled-coil domain containing 103; Dynein-attachment factor required for cilia motility; Belongs to the CCDC103/PR46b family. |
| | | FAM187A | ENSP00000391869 | Family with sequence similarity 187, member A; Dynein-attachment factor required for cilia motility; Belongs to the CCDC103/PR46b family. |
| | | FOXJ1 | ENSP00000323880 | Hepatocyte nuclear factor 3 forkhead homolog 4; Transcription factor specifically required for the formation of motile cilia. Acts by activating transcription of genes that med |
| | | HIGD1B | ENSP00000253410 | HIG1 hypoxia inducible domain family member 1B |
| | | RPGR | ENSP00000367766 | X-linked retinitis pigmentosa GTPase regulator; Could be a guanine-nucleotide releasing factor. Plays a role in ciliogenesis. Probably regulates cilia formation by regulating ac |
| 25 | 5 | CCDC8 | ENSP00000303158 | Coiled-coil domain-containing protein 8; Core component of the 3M complex, a complex required to regulate microtubule dynamics and genome integrity. It is unclear how |
| | | MDFI | ENSP00000230321 | Myogenic repressor I-mf; Inhibits the transactivation activity of the Myod family of myogenic factors and represses myogenesis. Acts by associating with Myod family membe |
| | | PBXIP1 | ENSP00000357448 | Pre-B-cell leukemia transcription factor-interacting protein 1; Regulator of pre-B-cell leukemia transcription factors (BPXs) function. Inhibits the binding of PBX1-HOX comple |
| | | RAB13 | ENSP00000357564 | Cell growth-inhibiting gene 4 protein; The small GTPases Rab are key regulators of intracellular membrane trafficking, from the formation of transport vesicles to their fusion |
| | | RPS27 | ENSP00000357555 | Small ribosomal subunit protein eS27; Component of the small ribosomal subunit. Required for proper rRNA processing and maturation of 18S rRNAs; Belongs to the eukary |
| 26 | 5 | MTURN | ENSP00000324204 | Maturin, neural progenitor differentiation regulator homolog (Xenopus); May be involved in early neuronal development. |
| | | NKD1 | ENSP00000268459 | Naked cuticle homolog 1 (Drosophila); Cell autonomous antagonist of the canonical Wnt signaling pathway. May activate a second Wnt signaling pathway that controls plana |
| | | PPP1R3B | ENSP00000308318 | Hepatic glycogen-targeting protein phosphatase 1 regulatory subunit GL; Acts as a glycogen-targeting subunit for phosphatase PP1. Facilitates interaction of the PP1 with enz |
| | | PPP1R3C | ENSP00000238994 | Protein phosphatase 1, regulatory subunit 3C; Acts as a glycogen-targeting subunit for PP1 and regulates its activity. Activates glycogen synthase, reduces glycogen phosphor |
| | | PPP1R3G | ENSP00000393832 | Protein phosphatase 1, regulatory subunit 3G; Glycogen-targeting subunit for protein phosphatase 1 (PP1). Involved in the regulation of hepatic glycogenesis in a manner co |
| 27 | 5 | ARHGEF15 | ENSP00000355026 | Rho guanine nucleotide exchange factor (GEF) 15; Specific GEF for RhoA activation. Does not activate RAC1 or CDC42. Regulates vascular smooth muscle contractility. Negativ |
| | | ARHGEF40 | ENSP00000298694 | Rho guanine nucleotide exchange factor (GEF) 40; May act as a guanine nucleotide exchange factor (GEF). |
| | | PLEKHG2 | ENSP00000392906 | Pleckstrin homology domain containing, family G (with RhoGef domain) member 2; May be a transforming oncogene with exchange activity for CDC42 (By similarity). May be |
| | | SWAP70 | ENSP00000315630 | SWAP switching B-cell complex 70kDa subunit; Phosphatidylinositol 3,4,5-trisphosphate-dependent guanine nucleotide exchange factor (GEF) which, independently of RAS, t |
| | | TIPARP | ENSP00000420612 | ADP-ribosyltransferase diphtheria toxin-like 14; Poly [ADP-ribose] polymerase using NAD(+) as a substrate to transfer ADP-ribose onto glutamic acid residues of a protein acc |
| 28 | 3 | NBPF10 | ENSP00000463957 | Neuroblastoma breakpoint family, member 10; NBPF member 10 |
| | | NBPF26 | ENSP00000479693 | Neuroblastoma breakpoint family, member 14; NBPF member 14 |
| | | NBPF9 | ENSP00000477979 | Neuroblastoma breakpoint family member 20; NBPF member 9; Belongs to the NBPF family. |
| 29 | 5 | MYO6 | ENSP00000358994 | Unconventional myosin-VI; Myosins are actin-based motor molecules with ATPase activity. Unconventional myosins serve in intracellular movements. Myosin 6 is a reverse-c |
| | | PDLIM1 | ENSP00000360305 | C-terminal LIM domain protein 1; Cytoskeletal protein that may act as an adapter that brings other proteins (like kinases) to the cytoskeleton. Involved in assembly, disassem |
| | | PDLIM2 | ENSP00000312634 | PDZ and LIM domain 2 (mystique); Probable adapter protein located at the actin cytoskeleton that promotes cell attachment. Necessary for the migratory capacity of epithel |
| | | PDLIM3 | ENSP00000284770 | Alpha-actinin-2-associated LIM protein; May play a role in the organization of actin filament arrays within muscle cells; LIM domain containing |
| | | PDLIM4 | ENSP00000253754 | Reversion-induced LIM protein; Isoform 1: Suppresses SRC activation by recognizing and binding to active SRC and facilitating PTPN13-mediated dephosphorylation of SRC 'T |
| 31 | 5 | CTDSP1 | ENSP00000273062 | CTD (carboxy-terminal domain, RNA polymerase II, polypeptide A) small phosphatase 1; Preferentially catalyzes the dephosphorylation of 'Ser- 5' within the tandem 7 residu |
| | | CTDSP2 | ENSP00000381148 | CTD (carboxy-terminal domain, RNA polymerase II, polypeptide A) small phosphatase 2; Preferentially catalyzes the dephosphorylation of 'Ser- 5' within the tandem 7 residu |
| | | CTDSPL | ENSP00000273179 | CTD (carboxy-terminal domain, RNA polymerase II, polypeptide A) small phosphatase-like; Recruited by REST to neuronal genes that contain RE-1 elements, leading to neuro |
| | | LPIN3 | ENSP00000362354 | Phosphatidate phosphatase LPIN3; Regulates fatty acid metabolism. Magnesium-dependent phosphatidate phosphatase enzyme which catalyzes the conversion of phosphat |
| | | REST | ENSP00000311816 | Neural-restrictive silencer factor; Transcriptional repressor which binds neuron-restrictive silencer element (NRSE) and represses neuronal gene transcription in non-neurona |
| 32 | 5 | ABCB1 | ENSP00000478255 | ATP-binding cassette, sub-family B (MDR/TAP), member 1; Energy-dependent efflux pump responsible for decreased drug accumulation in multidrug-resistant cells; ATP binc |
| | | SLC15A2 | ENSP00000417085 | Solute carrier family 15 (oligopeptide transporter), member 2; Proton-coupled intake of oligopeptides of 2 to 4 amino acids with a preference for dipeptides; Belongs to the |
| | | SLC2A1 | ENSP00000416293 | Solute carrier family 2 (facilitated glucose transporter), member 1; Facilitative glucose transporter. This isoform may be responsible for constitutive or basal glucose uptake. |
| | | SLC7A2 | ENSP00000004531 | Solute carrier family 7 (cationic amino acid transporter, y+ system), member 2; Functions as permease involved in the transport of the cationic amino acids (arginine, lysine a |
| | | SLCO4A1 | ENSP00000217159 | Solute carrier organic anion transporter family, member 4A1; Mediates the Na(+)-independent transport of organic anions such as the thyroid hormones T3 (triiodo-L-thyron |
| 33 | 5 | CHST14 | ENSP00000307297 | Carbohydrate (N-acetylgalactosamine 4-0) sulfotransferase 14; Catalyzes the transfer of sulfate to position 4 of the N-acetylgalactosamine (GalNAc) residue of dermatan sulfa |
| | | CHST3 | ENSP00000362207 | Galactose/N-acetylglucosamine/N-acetylglucosamine 6-O-sulfotransferase 0; Sulfotransferase that utilizes 3'-phospho-5'-adenylyl sulfate (PAPS) as sulfonate donor to catalyz |
| | | CHST6 | ENSP00000328983 | Galactose/N-acetylglucosamine/N-acetylglucosamine 6-O-sulfotransferase 4-beta; Sulfotransferase that utilizes 3'-phospho-5'-adenylyl sulfate (PAPS) as sulfonate donor to ca |
| | | PRELP | ENSP00000343924 | Proline-arginine-rich end leucine-rich repeat protein; May anchor basement membranes to the underlying connective tissue; Small leucine rich repeat proteoglycans |
| | | TSKU | ENSP00000434847 | Leucine-rich repeat-containing protein 54; Tsukushi, small leucine rich proteoglycan |
| 34 | 4 | GPC5 | ENSP00000366267 | Glypican 5; Cell surface proteoglycan that bears heparan sulfate; Belongs to the glypican family. |
| | | HPSE2 | ENSP00000359583 | Heparanase 2 (inactive); Binds heparin and heparan sulfate with high affinity, but lacks heparanase activity. Inhibits HPSE, possibly by competing for its substrates (in vitro). |
| | | HSPG2 | ENSP00000363827 | Basement membrane-specific heparan sulfate proteoglycan core protein; Integral component of basement membranes. Component of the glomerular basement membrane |
| | | LRP10 | ENSP00000352601 | Low density lipoprotein receptor-related protein 10; Probable receptor, which is involved in the internalization of lipophilic molecules and/or signal transduction. May be inv |
| 35 | 4 | DAAM2 | ENSP00000381876 | Dishevelled associated activator of morphogenesis 2; Armadillo-like helical domain containing |
| | | PKN3 | ENSP00000291906 | Serine/threonine-protein kinase N3; Contributes to invasiveness in malignant prostate cancer; Belongs to the protein kinase superfamily. AGC Ser/Thr protein kinase family. |
| | | PREX2 | ENSP00000288368 | Phosphatidylinositol 3,4,5-trisphosphate-dependent Rac exchanger 2 protein; Functions as a RAC1 guanine nucleotide exchange factor (GEF), activating Rac proteins by excha |
| | | RHOC | ENSP00000285735 | Rho-related GTP-binding protein RhoC; Regulates a signal transduction pathway linking plasma membrane receptors to the assembly of focal adhesions and actin stress fiber |
| 36 | 4 | HSD3B7 | ENSP00000297679 | Hydroxy-delta-5-steroid dehydrogenase, 3 beta- and steroid delta-isomerase 7; The 3-beta-HSD enzymatic system plays a crucial role in the biosynthesis of all classes of horm |
| | | SLC51B | ENSP00000335292 | Organic solute transporter subunit beta; Essential component of the Ost-alpha/Ost-beta complex, a heterodimer that acts as the intestinal basolateral transporter responsibl |
| | | SLCO1A2 | ENSP00000305974 | Solute carrier organic anion transporter family, member 1A2; Mediates the Na(+)-independent transport of organic anions such as sulfobromophthalein (BSP) and conjugate |
| | | SLCO2B1 | ENSP00000289575 | Solute carrier organic anion transporter family, member 2B1; Mediates the Na(+)-independent transport of organic anions such as taurocholate, the prostaglandins PGD2, PC |
| 37 | 3 | AHCYL1 | ENSP00000358814 | S-adenosylhomocysteine hydrolase-like protein 1; Multifaceted cellular regulator which coordinates several essential cellular functions including regulation of epithelial HCO |
| | | BHMT2 | ENSP00000255192 | S-methylmethionine--homocysteine S-methyltransferase BHMT2; Involved in the regulation of homocysteine metabolism. Converts homocysteine to methionine using S-met |
| | | SDS | ENSP00000257549 | L-serine dehydratase/L-threonine deaminase; Serine dehydratase |
| 38 | 4 | EDA | ENSP00000363680 | Ectodermal dysplasia protein; Cytokine which is involved in epithelial-mesenchymal signaling during morphogenesis of ectodermal organs. Functions as a ligand activating th |
| | | FOXQ1 | ENSP00000296839 | Hepatocyte nuclear factor 3 forkhead homolog 1; Plays a role in hair follicle differentiation; Forkhead boxes |
| | | LRRC1 | ENSP00000359925 | Leucine-rich repeat-containing protein 1; Leucine rich repeat containing 1 |
| | | TCF15 | ENSP00000246080 | Transcription factor 15 (basic helix-loop-helix); May function as an early transcriptional regulator, involved in the patterning of the mesoderm and in lineage determination c |
| 39 | 4 | ADAMTS9 | ENSP00000418735 | A disintegrin and metalloproteinase with thrombospondin motifs 9; Cleaves the large aggregating proteoglycans, aggrecan (at the '1838-Glu-\|-Ala-1839' site) and versican (at |
| | | ADAMTSL3 | ENSP00000286744 | ADAMTS-like protein 3; Immunoglobulin like domain containing |
| | | ADAMTSL4 | ENSP00000358035 | Thrombospondin repeat-containing protein 1; Positive regulation of apoptosis. May facilitate FBN1 microfibril biogenesis; ADAMTS like |
| | | THSD4 | ENSP00000347484 | A disintegrin and metalloproteinase with thrombospondin motifs-like protein 6; Promotes FBN1 matrix assembly. Attenuates TGFB signaling, possibly by accelerating the sec |
| 40 | 4 | ADCYAP1R1 | ENSP00000483721 | Adenylate cyclase activating polypeptide 1 (pituitary) receptor type I; This is a receptor for PACAP-27 and PACAP-38. The activity of this receptor is mediated by G proteins w |
| | | AQP1 | ENSP00000311165 | Water channel protein for red blood cells and kidney proximal tubule; Forms a water-specific channel that provides the plasma membranes of red cells and kidney proximal |
| | | AQP5 | ENSP00000293599 | Aquaporin 5; Forms a water-specific channel. Implicated in the generation of saliva, tears, and pulmonary secretions. Required for TRPV4 activation by hypotonicity. Togethe |
| | | SLC14A1 | ENSP00000390637 | Solute carrier family 14 (urea transporter), member 1 (Kidd blood group); Urea channel that facilitates transmembrane urea transport down a concentration gradient. A cons |
| 41 | 4 | CDH5 | ENSP00000344115 | Cadherin 5, type 2 (vascular endothelium); Cadherins are calcium-dependent cell adhesion proteins. They preferentially interact with themselves in a homophilic manner in c |
| | | ERG | ENSP00000414150 | V-ets avian erythroblastosis virus E26 oncogene homolog; Transcriptional regulator. May participate in transcriptional regulation through the recruitment of SETDB1 histone |
| | | GJA4 | ENSP00000343676 | Gap junction protein, alpha 4, 37kDa; One gap junction consists of a cluster of closely packed pairs of transmembrane channels, the connexons, through which materials of lo |
| | | ROBO4 | ENSP00000304945 | Roundabout, axon guidance receptor, homolog 4 (Drosophila); Receptor for Slit proteins, at least for SLIT2, and seems to be involved in angiogenesis and vascular patterning |
| 42 | 4 | GXYLT1 | ENSP00000381666 | Glycosyltransferase 8 domain-containing protein 3; Glycosyltransferase which elongates the O-linked glucose attached to EGF-like repeats in the extracellular domain of Notc |
| | | NHSL1 | ENSP00000394546 | NHS-like protein 1; NHS like 1; Belongs to the NHS family. |
| | | STON2 | ENSP00000450857 | Stoned B; Adapter protein involved in endocytic machinery. Involved in the synaptic vesicle recycling. May facilitate clathrin-coated vesicle uncoating; Belongs to the Stoned |
| | | ZNF462 | ENSP00000277225 | Zinc finger protein 462; May be involved in transcriptional regulation; Zinc fingers C2H2-type |
| 43 | 4 | CTNNA3 | ENSP00000389714 | Catenin (cadherin-associated protein), alpha 3; May be involved in formation of stretch-resistant cell- cell adhesion complexes; Alpha catenins |
| | | DSP | ENSP00000369129 | 250/210 kDa paraneoplastic pemphigus antigen; Major high molecular weight protein of desmosomes. Involved in the organization of the desmosomal cadherin- plakoglobir |
| | | PKP4 | ENSP00000374409 | Plakophilin 4; Plays a role as a regulator of Rho activity during cytokinesis. May play a role in junctional plaques; Armadillo repeat containing |
| | | PPP1R13L | ENSP00000403902 | Protein phosphatase 1, regulatory subunit 13 like; Regulator that plays a central role in regulation of apoptosis and transcription via its interaction with NF-kappa-B and p53/ |
| 44 | 4 | GALNT4 | ENSP00000436604 | UDP-GalNAc:polypeptide N-acetylgalactosaminyltransferase 4; Catalyzes the initial reaction in O-linked oligosaccharide biosynthesis, the transfer of an N-acetyl-D- galactosan |
| | | MUC1 | ENSP00000484824 | Tumor-associated epithelial membrane antigen; The alpha subunit has cell adhesive properties. Can act both as an adhesion and an anti-adhesion protein. May provide a pro |
| | | SLC25A18 | ENSP00000329033 | Solute carrier family 25 (glutamate carrier), member 18; Involved in the transport of glutamate across the inner mitochondrial membrane. Glutamate is cotransported with H |
| | | ST6GALNAC1 | ENSP00000156626 | ST6 N-acetylgalactosaminide alpha-2,6-sialyltransferase 1; Sialyltransferases |
| 45 | 4 | CCDC88A | ENSP00000338728 | G alpha-interacting vesicle-associated protein; Plays a role as a key modulator of the AKT-mTOR signaling pathway controlling the tempo of the process of newborn neurons |
| | | CENPB | ENSP00000369075 | Major centromere autoantigen B; Interacts with centromeric heterochromatin in chromosomes and binds to a specific 17 bp subset of alphoid satellite DNA, called the CENP |
| | | EML3 | ENSP00000378254 | Echinoderm microtubule associated protein like 3; May modify the assembly dynamics of microtubules, such that microtubules are slightly longer, but more dynamic; WD re |
| | | MAP4 | ENSP00000353375 | Microtubule-associated protein 4; Non-neuronal microtubule-associated protein. Promotes microtubule assembly. |
| 46 | 4 | CELSR1 | ENSP00000262738 | Cadherin, EGF LAG seven-pass G-type receptor 1; Receptor that may have an important role in cell/cell signaling during nervous system formation; Adhesion G protein-coupl |
| | | DCHS1 | ENSP00000299441 | Dachsous cadherin-related 1; Calcium-dependent cell-adhesion protein. Mediates functions in neuroprogenitor cell proliferation and differentiation. In the heart, has a critic |
| | | FZD7 | ENSP00000286201 | Frizzled class receptor 7; Receptor for Wnt proteins. Most of frizzled receptors are coupled to the beta-catenin canonical signaling pathway, which leads to the activation of c |
| | | PRICKLE3 | ENSP00000470248 | Prickle planar cell polarity protein 3; Involved in the planar cell polarity (PCP) pathway that is essential for the polarization of epithelial cells during morphogenetic processes |
| 47 | 3 | GNA12 | ENSP00000275364 | Guanine nucleotide binding protein (G protein) alpha 12; Guanine nucleotide-binding proteins (G proteins) are involved as modulators or transducers in various transmembr |
| | | GNA14 | ENSP00000365807 | Guanine nucleotide binding protein (G protein), alpha 14; Guanine nucleotide-binding proteins (G proteins) are involved as modulators or transducers in various transmemb |
| | | S1PR3 | ENSP00000365006 | Endothelial differentiation G-protein coupled receptor 3; Receptor for the lysosphingolipid sphingosine 1- phosphate (S1P). S1P is a bioactive lysophospholipid that elicits div |
| 48 | 4 | DDAH2 | ENSP00000364945 | N(G),N(G)-dimethylarginine dimethylaminohydrolase 2; Hydrolyzes N(G),N(G)-dimethyl-L-arginine (ADMA) and N(G)-monomethyl-L-arginine (MMA) which act as inhibitors of |
| | | DYNC1LI2 | ENSP00000258198 | Dynein, cytoplasmic 1, light intermediate chain 2; Acts as one of several non-catalytic accessory components of the cytoplasmic dynein 1 complex that are thought to be invo |
| | | KIF5B | ENSP00000307078 | Conventional kinesin heavy chain; Microtubule-dependent motor required for normal distribution of mitochondria and lysosomes. Can induce formation of neurite-like mem |
| | | NOS3 | ENSP00000297494 | Nitric oxide synthase 3 (endothelial cell); Produces nitric oxide (NO) which is implicated in vascular smooth muscle relaxation through a cGMP-mediated signal transduction p |
| 49 | 3 | AKR1C3 | ENSP00000369927 | Trans-1,2-dihydrobenzene-1,2-diol dehydrogenase; Catalyzes the conversion of aldehydes and ketones to alcohols. Catalyzes the reduction of prostaglandin (PG) D2, PGH2 ar |
| | | ARHGEF37 | ENSP00000328083 | Rho guanine nucleotide exchange factor (GEF) 37; May act as a guanine nucleotide exchange factor (GEF); Classical BAR domain containing |
| | | NOL8 | ENSP00000441140 | Nucleolar protein Nop132; Plays an essential role in the survival of diffuse-type gastric cancer cells. Acts as a nucleolar anchoring protein for DDX47. May be involved in regu |
| 50 | 4 | ATP1A2 | ENSP00000354490 | Sodium/potassium-transporting ATPase subunit alpha-2; This is the catalytic component of the active enzyme, which catalyzes the hydrolysis of ATP coupled with the exchan |
| | | FXYD1 | ENSP00000481244 | Sodium/potassium-transporting ATPase subunit FXYD1; Associates with and regulates the activity of the sodium/potassium-transporting ATPase (NKA) which transports Na(+ |
| | | FXYD3 | ENSP00000473929 | Sodium/potassium-transporting ATPase subunit FXYD3; Associates with and regulates the activity of the sodium/potassium-transporting ATPase (NKA) which transports Na(+ |
| | | LGI4 | ENSP00000312273 | Leucine-rich glioma-inactivated protein 4; Leucine rich repeat LGI family member 4 |
| 51 | 4 | HIPK2 | ENSP00000385571 | Homeodomain interacting protein kinase 2; Serine/threonine-protein kinase involved in transcription regulation, p53/TP53-mediated cellular apoptosis and regulation of the |
| | | NUPR1 | ENSP00000379003 | Nuclear protein 1, transcriptional regulator |
| | | SASH3 | ENSP00000349359 | SH3 protein expressed in lymphocytes homolog; May function as a signaling adapter protein in lymphocytes; SAM and SH3 domain containing |
| | | TP53INP1 | ENSP00000344215 | P53-dependent damage-inducible nuclear protein 1; Antiproliferative and proapoptotic protein involved in cell stress response which acts as a dual regulator of transcription |
| 52 | 4 | ADIRF | ENSP00000361083 | Adipose most abundant gene transcript 2 protein; Plays a role in fat cell development; promotes adipogenic differentiation and stimulates transcription initiation of master a |
| | | EMID1 | ENSP00000335481 | Emilin and multimerin domain-containing protein 1; EMI domain containing 1 |
| | | MMRN2 | ENSP00000361097 | Elastin microfibril interface located protein 3; Inhibits endothelial cells motility and acts as a negative regulator of angiogenesis; it downregulates KDR activation by binding V |
| | | USHBP1 | ENSP00000252597 | Usher syndrome type-1C protein-binding protein 1; USH1 protein network component harmonin binding protein 1; Belongs to the MCC family. |
| 53 | 3 | CAV1 | ENSP00000339191 | Caveolin 1, caveolae protein, 22kDa; May act as a scaffolding protein within caveolar membranes. Interacts directly with G-protein alpha subunits and can functionally regula |
| | | EHD2 | ENSP00000263277 | EH domain-containing protein 2; ATP- and membrane-binding protein that controls membrane reorganization/tubulation upon ATP hydrolysis (By similarity). Plays a role in r |
| | | LRRC8A | ENSP00000259324 | Leucine rich repeat containing 8 family, member A; Essential component of the volume-regulated anion channel (VRAC, also named VSOAC channel), an anion channel requi |
| 54 | 4 | LONRF3 | ENSP00000360690 | LON peptidase N-terminal domain and ring finger 3 |
| | | PLEKHA4 | ENSP00000263265 | Pleckstrin homology domain containing, family A (phosphoinositide binding specific) member 4; Binds specifically to phosphatidylinositol 3-phosphate (PtdIns3P), but not to |
| | | SH3TC1 | ENSP00000245105 | SH3 domain and tetratricopeptide repeats 1 |
| | | SORCS2 | ENSP00000422185 | Sortilin related VPS10 domain containing receptor 2; Belongs to the VPS10-related sortilin family. SORCS subfamily. |
| 55 | 3 | CSPP1 | ENSP00000262210 | Centrosome and spindle pole associated protein 1; May play a role in cell-cycle-dependent microtubule organization. |

| | | | | |
|---|---|---|---|---|
| | | NEK1 | ENSP00000424757 | Serine/threonine-protein kinase Nek1; Phosphorylates serines and threonines, but also appears to possess tyrosine kinase activity (By similarity). Involved in DNA damage ch |
| | | NT5DC2 | ENSP00000406933 | 5'-nucleotidase domain containing 2 |
| 56 | 4 | ANXA13 | ENSP00000262219 | Annexin A13; Belongs to the annexin family. |
| | | ANXA2 | ENSP00000346032 | Placental anticoagulant protein IV; Calcium-regulated membrane-binding protein whose affinity for calcium is greatly enhanced by anionic phospholipids. It binds two calciu |
| | | S100A10 | ENSP00000357801 | S100 calcium binding protein A10; Because S100A10 induces the dimerization of ANXA2/p36, it may function as a regulator of protein phosphorylation in that the ANXA2 mo |
| | | S100A11 | ENSP00000271638 | Metastatic lymph node gene 70 protein; Facilitates the differentiation and the cornification of keratinocytes; Belongs to the S-100 family. |
| 57 | 4 | KREMEN1 | ENSP00000331242 | Kringle-containing protein marking the eye and the nose; Receptor for Dickkopf proteins. Cooperates with DKK1/2 to inhibit Wnt/beta-catenin signaling by promoting the en |
| | | LRP4 | ENSP00000367888 | Low density lipoprotein receptor-related protein 4; Mediates SOST-dependent inhibition of bone formation. Functions as a specific facilitator of SOST-mediated inhibition of |
| | | LRP5 | ENSP00000294304 | Low density lipoprotein receptor-related protein 5; Component of the Wnt-Fzd-LRP5-LRP6 complex that triggers beta-catenin signaling through inducing aggregation of recep |
| | | RNF43 | ENSP00000463069 | RING-type E3 ubiquitin transferase RNF43; E3 ubiquitin-protein ligase that acts as a negative regulator of the Wnt signaling pathway by mediating the ubiquitination, endocy |
| 58 | 3 | ASPH | ENSP00000368767 | Aspartyl/asparaginyl beta-hydroxylase; Isoform 1: specifically hydroxylates an Asp or Asn residue in certain epidermal growth factor-like (EGF) domains of a number of prote |
| | | RYR3 | ENSP00000373884 | Brain ryanodine receptor-calcium release channel; Calcium channel that mediates the release of Ca(2+) from the sarcoplasmic reticulum into the cytoplasm in muscle and the |
| | | TET1 | ENSP00000362748 | Leukemia-associated protein with a CXXC domain; Dioxygenase that catalyzes the conversion of the modified genomic base 5-methylcytosine (5mC) into 5- hydroxymethylcyt |
| 59 | 4 | ITPKB | ENSP00000411152 | Inositol-trisphosphate 3-kinase B |
| | | ITPKC | ENSP00000263370 | Inositol 1,4,5-trisphosphate 3-kinase C; Can phosphorylate inositol 2,4,5-triphosphate to inositol 2,4,5,6-tetraphosphate; Belongs to the inositol phosphokinase (IPK) family. |
| | | PLCD1 | ENSP00000430344 | 1-phosphatidylinositol 4,5-bisphosphate phosphodiesterase delta-1; The production of the second messenger molecules diacylglycerol (DAG) and inositol 1,4,5-trisphosphate |
| | | PLCD3 | ENSP00000479636 | 1-phosphatidylinositol 4,5-bisphosphate phosphodiesterase delta-3; Hydrolyzes the phosphatidylinositol 4,5-bisphosphate (PIP2) to generate 2 second messenger molecules |
| 60 | 4 | MAPKAPK2 | ENSP00000356070 | Mitogen-activated protein kinase-activated protein kinase 2; Stress-activated serine/threonine-protein kinase involved in cytokine production, endocytosis, reorganization of |
| | | MAPKAPK3 | ENSP00000396467 | Mitogen-activated protein kinase-activated protein kinase 3; Stress-activated serine/threonine-protein kinase involved in cytokines production, endocytosis, cell migration, c |
| | | RAB29 | ENSP00000356107 | RAB29, member RAS oncogene family; Rab GTPase key regulator in vesicle trafficking. Essential for maintaining the integrity of the endosome-trans- Golgi network structure. |
| | | ZFP36L2 | ENSP00000282388 | Zinc finger protein 36, C3H1 type-like 2; Zinc-finger RNA-binding protein that destabilizes several cytoplasmic AU-rich element (ARE)-containing mRNA transcripts by promoti |
| 61 | 3 | CNN3 | ENSP00000359225 | Calponin, acidic isoform; Thin filament-associated protein that is implicated in the regulation and modulation of smooth muscle contraction. It is capable of binding to actin, |
| | | FLVCR2 | ENSP00000238667 | Feline leukemia virus subgroup C cellular receptor family, member 2; Acts as an importer of heme. Also acts as a transporter for a calcium-chelator complex, important for gr |
| | | RFX4 | ENSP00000350552 | Regulatory factor X, 4 (influences HLA class II expression); May activate transcription by interacting directly with the X-box; Belongs to the RFX family. |
| 62 | 2 | ETNK2 | ENSP00000356169 | Ethanolamine kinase-like protein; Highly specific for ethanolamine phosphorylation. Does not have choline kinase activity (By similarity); Belongs to the choline/ethanolamin |
| | | SOX13 | ENSP00000356172 | SRY (Sex determining region Y)-box 13; Binds to the sequence 5'-AACAAT-3'; SRY-boxes |
| 63 | 3 | CYBRD1 | ENSP00000319141 | Ferric-chelate reductase 3; Ferric-chelate reductase that reduces Fe(3+) to Fe(2+). Present at the brush border of duodenal enterocytes where it probably reduces dietary Fe |
| | | HEPH | ENSP00000430620 | Hephaestin; May function as a ferroxidase for ferrous (II) to ferric ion (III) conversion and may be involved in copper transport and homeostasis. Implicated in iron homeosta |
| | | SLC40A1 | ENSP00000261024 | Solute carrier family 40 (iron-regulated transporter), member 1; May be involved in iron export from duodenal epithelial cell and also in transfer of iron between maternal a |
| 64 | 3 | ABHD15 | ENSP00000302657 | Alpha/beta hydrolase domain-containing protein 15; Abhydrolase domain containing 15 |
| | | ABHD4 | ENSP00000414558 | Alpha/beta hydrolase domain-containing protein 4; Lysophospholipase selective for N-acyl phosphatidylethanolamine (NAPE). Contributes to the biosynthesis of N-acyl ethar |
| | | KANK3 | ENSP00000328923 | KN motif and ankyrin repeat domain-containing protein 3; May be involved in the control of cytoskeleton formation by regulating actin polymerization; Ankyrin repeat doma |
| 65 | 2 | HMG20B | ENSP00000328269 | Structural DNA-binding protein BRAF35; Required for correct progression through G2 phase of the cell cycle and entry into mitosis. Required for RCOR1/CoREST mediated re |
| | | PHF21B | ENSP00000324403 | PHD finger protein 21B |
| 66 | 2 | ITPR2 | ENSP00000370744 | Inositol 1,4,5-trisphosphate receptor, type 2; Receptor for inositol 1,4,5-trisphosphate, a second messenger that mediates the release of intracellular calcium. This release is |
| | | ORAI3 | ENSP00000322249 | ORAI calcium release-activated calcium modulator 3; Key regulator or component of store-operated Ca(2+) channel and transcription factor NFAT nuclear import; Belongs to |
| 67 | 3 | CRX | ENSP00000221996 | Cone-rod homeobox protein; Transcription factor that binds and transactivates the sequence 5'-TAATC[CA]-3' which is found upstream of several photoreceptor-specific gen |
| | | ROM1 | ENSP00000278833 | Retinal outer segment membrane protein 1; May function as an adhesion molecule involved in stabilization and compaction of outer segment disks or in the maintenance of |
| | | TSPAN4 | ENSP00000380553 | Transmembrane 4 superfamily member 7; Tetraspanin 4; Belongs to the tetraspanin (TM4SF) family. |
| 68 | 3 | FAM189A2 | ENSP00000257515 | Family with sequence similarity 189 member A2 |
| | | SNX33 | ENSP00000311427 | SH3 and PX domain-containing protein 3; Plays a role in the reorganization of the cytoskeleton, endocytosis and cellular vesicle trafficking via its interactions with membrane |
| | | TSPO | ENSP00000379563 | Peripheral-type benzodiazepine receptor; Can bind protoporphyrin IX and may play a role in the transport of porphyrins and heme (By similarity). Promotes the transport of |
| 69 | 2 | ANKRD40 | ENSP00000285243 | Ankyrin repeat domain-containing protein 40; Ankyrin repeat domain containing |
| | | PAQR6 | ENSP00000338330 | Progesterone and adipoQ receptor family member 6; Plasma membrane progesterone (P4) receptor coupled to G proteins. Seems to act through a G(s) mediated pathway. I |
| 70 | 3 | ESAM | ENSP00000278927 | Endothelial cell-selective adhesion molecule; Can mediate aggregation most likely through a homophilic molecular interaction; IgCAM CXADR-related subfamily |
| | | SLC44A2 | ENSP00000336888 | Solute carrier family 44 (choline transporter), member 2; Isoform 1, but not isoform 3, exhibits some choline transporter activity; Solute carriers |
| | | VWF | ENSP00000261405 | Von Willebrand factor; Important in the maintenance of hemostasis, it promotes adhesion of platelets to the sites of vascular injury by forming a molecular bridge between s |
| 71 | 2 | ARSI | ENSP00000333395 | Arylsulfatase family, member I; Displays arylsulfatase activity at neutral pH, when co- expressed with SUMF1; arylsulfatase activity is measured in the secretion medium of re |
| | | ST8SIA6 | ENSP00000366827 | ST8 alpha-N-acetyl-neuraminide alpha-2,8-sialyltransferase 6; Prefers O-glycans to N-glycans or glycolipids as acceptor substrates. The minimal acceptor substrate is the Neu/ |
| 72 | 3 | CRACR2B | ENSP00000435299 | Calcium release-activated calcium channel regulator 2B; Plays a role in store-operated Ca(2+) entry (SOCE); EF-hand domain containing |
| | | S100A16 | ENSP00000357693 | Aging-associated gene 13 protein; Calcium-binding protein. Binds one calcium ion per monomer. Can promote differentiation of adipocytes (in vitro) (By similarity). Overexp |
| | | S100A3 | ENSP00000357702 | S100 calcium binding protein A3; Binds both calcium and zinc. May be involved in calcium- dependent cuticle cell differentiation, hair shaft and hair cuticular barrier formatio |
| 73 | 3 | APPL2 | ENSP00000446917 | Adaptor protein, phosphotyrosine interaction, PH domain and leucine zipper containing 2; Required for the regulation of cell proliferation in response to extracellular signals |
| | | RAB31 | ENSP00000461945 | RAB31, member RAS oncogene family; The small GTPases Rab are key regulators of intracellular membrane trafficking, from the formation of transport vesicles to their fusio |
| | | RIN3 | ENSP00000216487 | Ras interaction/interference protein 3; Ras effector protein that functions as a guanine nucleotide exchange (GEF) for RAB5B and RAB31, by exchanging bound GDP for free ( |
| 74 | 3 | CGNL1 | ENSP00000281282 | Junction-associated coiled-coil protein; May be involved in anchoring the apical junctional complex, especially tight junctions, to actin-based cytoskeletons. |
| | | CSRP1 | ENSP00000356275 | Cysteine and glycine-rich protein 1; Could play a role in neuronal development; LIM domain containing |
| | | MRO | ENSP00000397900 | Male-specific transcription in the developing reproductive organs; Maestro heat like repeat containing |
| 75 | 3 | GPR39 | ENSP00000327417 | G protein-coupled receptor 39; Zn(2+) acts as an agonist. This receptor mediates its action by association with G proteins that activate a phosphatidylinositol-calcium second |
| | | GPR4 | ENSP00000319744 | G-protein coupled receptor 19; Proton-sensing receptor coupled to several G-proteins, including G(s), G(13) and G(q)/G(11) proteins, leading to cAMP production; G protein- |
| | | NMB | ENSP00000378089 | Neuromedin B; Stimulates smooth muscle contraction in a manner similar to that of bombesin; Belongs to the bombesin/neuromedin-B/ranatensin family. |
| 76 | 3 | ARHGEF5 | ENSP00000056217 | Rho guanine nucleotide exchange factor (GEF) 5; Guanine nucleotide exchange factor which activates Rho GTPases. Strongly activates RHOA. Also strongly activates RHOB, w |
| | | OPHN1 | ENSP00000347710 | Oligophrenin 1; Stimulates GTP hydrolysis of members of the Rho family. Its action on RHOA activity and signaling is implicated in growth and stabilization of dendritic spines |
| | | PAMR1 | ENSP00000482899 | Peptidase domain-containing protein associated with muscle regeneration 1; May play a role in regeneration of skeletal muscle; Belongs to the peptidase S1 family. |
| 77 | 3 | ARHGEF10 | ENSP00000340297 | Rho guanine nucleotide exchange factor (GEF) 10; May play a role in developmental myelination of peripheral nerves; Rho guanine nucleotide exchange factors |
| | | ARHGEF19 | ENSP00000270747 | Rho guanine nucleotide exchange factor (GEF) 19; Acts as guanine nucleotide exchange factor (GEF) for RhoA GTPase. |
| | | SLC2A4RG | ENSP00000266077 | Huntington disease gene regulatory region-binding protein 1; Transcription factor involved in SLC2A4 and HD gene transactivation. Binds to the consensus sequence 5'-GCCG |
| 78 | 3 | GPR137B | ENSP00000355551 | G protein-coupled receptor 137B; Belongs to the GPR137 family. |
| | | GPRC5C | ENSP00000376403 | G protein-coupled receptor, class C, group 5, member C; This retinoic acid-inducible G-protein coupled receptor provide evidence for a possible interaction between retinoid |
| | | PAQR5 | ENSP00000378803 | Progesterone and adipoQ receptor family member 5; Plasma membrane progesterone (P4) receptor coupled to G proteins. Seems to act through a G(i) mediated pathway. M |
| 79 | 3 | NXNL1 | ENSP00000305631 | Nucleoredoxin-like protein 1; May play a role in cone cell viability, slowing down cone degeneration, does not seem to play a role in degenerating rods; Nucleoredoxin family |
| | | VSX1 | ENSP00000365899 | Retinal inner nuclear layer homeobox protein; Binds to the 37-bp core of the locus control region (LCR) of the red/green visual pigment gene cluster. May regulate the activit |
| | | ZMAT1 | ENSP00000361868 | Zinc finger matrin-type 1 |
| 80 | 3 | BMPR1B | ENSP00000401907 | Bone morphogenetic protein receptor, type IB; On ligand binding, forms a receptor complex consisting of two type II and two type I transmembrane serine/threonine kinase |
| | | GDF11 | ENSP00000257868 | Growth differentiation factor 11; Secreted signal that acts globally to specify positional identity along the anterior/posterior axis during development. May play critical roles |
| | | SMAD6 | ENSP00000288840 | Mothers against decapentaplegic homolog 6; Acts as a mediator of TGF-beta and BMP antiflammatory activity. Suppresses IL1R-TLR signaling through its direct interaction wi |
| 81 | 3 | CCDC102A | ENSP00000258214 | Coiled-coil domain containing 102A |
| | | CDC42EP4 | ENSP00000338258 | CDC42 effector protein (Rho GTPase binding) 4; Probably involved in the organization of the actin cytoskeleton. May act downstream of CDC42 to induce actin filament asser |
| | | METRN | ENSP00000455068 | Meteorin, glial cell differentiation regulator; Involved in both glial cell differentiation and axonal network formation during neurogenesis. Promotes astrocyte differentiation |
| 82 | 2 | EMCN | ENSP00000296420 | Gastric cancer antigen Ga34; Endothelial sialomucin, also called endomucin or mucin- like sialoglycoprotein, which interferes with the assembly of focal adhesion complexes |
| | | ZNF366 | ENSP00000313158 | Dendritic cell-specific transcript protein; Has transcriptional repression activity. Acts as corepressor of ESR1; the function seems to involve CTBP1 and histone deacetylases; Z |
| 83 | 3 | BAG3 | ENSP00000358081 | BAG family molecular chaperone regulator 3; Co-chaperone for HSP70 and HSC70 chaperone proteins. Acts as a nucleotide-exchange factor (NEF) promoting the release of A |
| | | HSPA6 | ENSP00000310219 | Heat shock 70kDa protein 6 (HSP70B'); Molecular chaperone implicated in a wide variety of cellular processes, including protection of the proteome from stress, folding and |
| | | HSPB7 | ENSP00000310111 | Heat shock protein family B member 7 |
| 84 | 3 | ARHGAP42 | ENSP00000298815 | Rho GTPase-activating protein 10-like; May influence blood pressure by functioning as a GTPase- activating protein for RHOA in vascular smooth muscle. |
| | | PSD4 | ENSP00000245796 | Exchange factor for ADP-ribosylation factor guanine nucleotide factor 6 B; Guanine nucleotide exchange factor for ARF6 and ARL14/ARF7. Through ARL14 activation, controls |
| | | SASH1 | ENSP00000356437 | Proline-glutamate repeat-containing protein; May have a role in a signaling pathway. Could act as a tumor suppressor; SAM and SH3 domain containing |
| 85 | 3 | APOLD1 | ENSP00000324277 | Apolipoprotein L domain-containing protein 1; May be involved in angiogenesis. May play a role in activity-dependent changes of brain vasculature. May affect blood- brain |
| | | MKNK2 | ENSP00000250896 | MAP kinase-interacting serine/threonine-protein kinase 2; Serine/threonine-protein kinase that phosphorylates SFPQ/PSF, HNRNPA1 and EIF4E. May play a role in the respo |
| | | SZRD1 | ENSP00000383866 | Putative MAPK-activating protein PM18/PM20/PM22; SUZ RNA binding domain containing 1 |
| 86 | 2 | NXT2 | ENSP00000218004 | Nuclear transport factor 2-like export factor 2; Regulator of protein export for NES-containing proteins. Also plays a role in mRNA nuclear export. |
| | | TMEM164 | ENSP00000361143 | Transmembrane protein 164; Belongs to the TMEM164 family. |
| 87 | 3 | DHTKD1 | ENSP00000263035 | Probable 2-oxoglutarate dehydrogenase E1 component DHKTD1, mitochondrial; The 2-oxoglutarate dehydrogenase complex catalyzes the overall conversion of 2-oxoglutara |
| | | ELOVL7 | ENSP00000424123 | Elongation of very long chain fatty acids protein 7; Catalyzes the first and rate-limiting reaction of the four that constitute the long-chain fatty acids elongation cycle. This enc |
| | | FADS2 | ENSP00000278840 | Delta(6) fatty acid desaturase; Component of a lipid metabolic pathway that catalyzes biosynthesis of highly unsaturated fatty acids (HUFA) from precursor essential polyunsa |
| 88 | 3 | GOLGA4 | ENSP00000349305 | Golgin subfamily A member 4; May play a role in delivery of transport vesicles containing GPI-linked proteins from the trans-Golgi network through its interaction with MACF |
| | | GOLIM4 | ENSP00000417354 | Golgi-localized phosphoprotein of 130 kDa; Plays a role in endosome to Golgi protein trafficking; mediates protein transport along the late endosome-bypass pathway from t |
| | | MDFIC | ENSP00000484656 | MyoD family inhibitor domain-containing protein; Acts as a transcriptional activator or repressor. Inhibits the transcriptional activation of Zic family proteins ZIC1, ZIC2 and Z |
| 89 | 3 | ARRDC4 | ENSP00000268042 | Arrestin domain-containing protein 4; Functions as an adapter recruiting ubiquitin-protein ligases to their specific substrates (By similarity). Plays a role in endocytosis of acti |
| | | NEDD4 | ENSP00000424827 | Neural precursor cell expressed, developmentally down-regulated 4, E3 ubiquitin protein ligase; E3 ubiquitin-protein ligase which accepts ubiquitin from an E2 ubiquitin-con |
| | | WBP1 | ENSP00000233615 | WW domain binding protein 1; WBP1/VOPP1 family |
| 90 | 2 | GADD45G | ENSP00000252506 | Growth arrest and DNA damage-inducible protein GADD45 gamma; Involved in the regulation of growth and apoptosis. Mediates activation of stress-responsive MTK1/MEKI |
| | | NPAS3 | ENSP00000348460 | Class E basic helix-loop-helix protein 12; May play a broad role in neurogenesis. May control regulatory pathways relevant to schizophrenia and to psychotic illness (By simila |
| 91 | 3 | DPEP2 | ENSP00000458977 | Dipeptidase 2; Probable metalloprotease which hydrolyzes leukotriene D4 (LTD4) into leukotriene E4 (LTE4). |
| | | GGT5 | ENSP00000381340 | Gamma-glutamyl transpeptidase-related enzyme; Cleaves the gamma-glutamyl peptide bond of glutathione conjugates, but maybe not glutathione itself. Converts leukotrien |
| | | HYAL2 | ENSP00000401853 | Hyaluronoglucosaminidase 2; Hydrolyzes high molecular weight hyaluronic acid to produce an intermediate-sized product which is further hydrolyzed by sperm hyaluronidas |
| 92 | 3 | ITPRIP | ENSP00000278071 | Inositol 1,4,5-trisphosphate receptor interacting protein; Enhances Ca(2+)-mediated inhibition of inositol 1,4,5- triphosphate receptor (ITPR) Ca(2+) release; Belongs to the IT |
| | | ITPRIPL2 | ENSP00000370849 | Inositol 1,4,5-trisphosphate receptor interacting protein-like 2; ITPRIP like 2 |
| | | TMEM102 | ENSP00000315387 | Common beta-chain associated protein; Selectively involved in CSF2 deprivation-induced apoptosis via a mitochondria-dependent pathway. |
| 93 | 2 | EIF4EBP1 | ENSP00000340691 | Phosphorylated heat- and acid-stable protein regulated by insulin 1; Repressor of translation initiation that regulates EIF4E activity by preventing its assembly into the eIF4F |
| | | EIF4EBP2 | ENSP00000362314 | Eukaryotic translation initiation factor 4E binding protein 2; Repressor of translation initiation involved in synaptic plasticity, learning and memory formation (By similarity). F |
| 94 | 3 | LEF1 | ENSP00000265165 | T cell-specific transcription factor 1-alpha; Participates in the Wnt signaling pathway. Activates transcription of target genes in the presence of CTNNB1 and EP300. May play |
| | | TCF7 | ENSP00000340347 | Transcription factor 7 (T-cell specific, HMG-box); Transcriptional activator involved in T-cell lymphocyte differentiation. Necessary for the survival of CD4(+) CD8(+) immature |
| | | TCF7L1 | ENSP00000282111 | Transcription factor 7-like 1 (T-cell specific, HMG-box); Participates in the Wnt signaling pathway. Binds to DNA and acts as a repressor in the absence of CTNNB1, and as an a |
| 95 | 2 | HMGN5 | ENSP00000350848 | High mobility group nucleosome-binding domain-containing protein 5; Preferentially binds to euchromatin and modulates cellular transcription by counteracting linker histo |
| | | STC1 | ENSP00000290271 | Stanniocalcin 1; Stimulates renal phosphate reabsorption, and could therefore prevent hypercalcemia. |
| 96 | 3 | RASL12 | ENSP00000220062 | Ras-like protein family member 12; RAS type GTPase family |
| | | SOD3 | ENSP00000371554 | Extracellular superoxide dismutase [Cu-Zn]; Protect the extracellular space from toxic effect of reactive oxygen intermediates by converting superoxide radicals into hydroge |
| | | ZNF853 | ENSP00000455585 | Zinc finger protein 853; Zinc fingers C2H2-type |
| 97 | 2 | SRGN | ENSP00000242465 | Secretory granule proteoglycan core protein; Plays a role in formation of mast cell secretory granules and mediates storage of various compounds in secretory vesicles. Requ |
| | | VSIG4 | ENSP00000363869 | V-set and immunoglobulin domain-containing protein 4; Phagocytic receptor, strong negative regulator of T-cell proliferation and IL2 production. Potent inhibitor of the alte |
| 98 | 3 | COLCA2 | ENSP00000484135 | Cancer susceptibility candidate protein 13; Colorectal cancer associated 2 |
| | | GPR143 | ENSP00000417161 | G protein-coupled receptor 143; Receptor for tyrosine, L-DOPA and dopamine. After binding to L-DOPA, stimulates Ca(2+) influx into the cytoplasm, increases secretion of th |
| | | RHPN2 | ENSP00000254260 | Rhophilin, Rho GTPase binding protein 2; Binds specifically to GTP-Rho. May function in a Rho pathway to limit stress fiber formation and/or increase the turnover of F-actin |
| 99 | 3 | GIMAP2 | ENSP00000223293 | Immunity-associated protein 2; The heterodimer formed by GIMAP2 and GIMAP7 has GTPase activity. In contrast, GIMAP2 has no GTPase activity by itself. |
| | | GIMAP7 | ENSP00000315474 | Immunity-associated nucleotide 7 protein; The dimer has GTPase activity; the active site contains residues from both subunits; GTPases, IMAP |
| | | NFATC4 | ENSP00000388910 | Nuclear factor of activated T-cells, cytoplasmic, calcineurin-dependent 4; Plays a role in the inducible expression of cytokine genes in T-cells, especially in the induction of the |
| 100 | 3 | GDPD2 | ENSP00000414019 | Glycerophosphodiester phosphodiesterase domain-containing protein 2; Has glycerophosphoinositol inositolphosphodiesterase activity and specifically hydrolyzes glyceroph |
| | | RAB30 | ENSP00000435189 | RAB30, member RAS oncogene family; The small GTPases Rab are key regulators of intracellular membrane trafficking, from the formation of transport vesicles to their fusio |
| | | RAB41 | ENSP00000276066 | RAB41, member RAS oncogene family; Required for normal Golgi ribbon organization and ER-to- Golgi trafficking; RAB, member RAS oncogene GTPases |
| 101 | 3 | MFSD2A | ENSP00000361895 | Major facilitator superfamily domain-containing protein 2A; Sodium-dependent lysophosphatidylcholine (LPC) symporter, which plays an essential role for blood-brain barrie |
| | | SLC19A3 | ENSP00000258403 | Solute carrier family 19 (thiamine transporter), member 3; Mediates high affinity thiamine uptake, probably via a proton anti-port mechanism. Has no folate transport activit |
| | | SLC52A3 | ENSP00000217254 | Solute carrier family 52 (riboflavin transporter), member 3; Transporter for riboflavin, which must be obtained as a nutrient via intestinal absorption. Riboflavin transport is N |
| 102 | 3 | SLC16A9 | ENSP00000378757 | Solute carrier family 16, member 9; Proton-linked monocarboxylate transporter. May catalyze the transport of monocarboxylates across the plasma membrane; Solute carrie |
| | | SLC26A2 | ENSP00000286298 | Solute carrier family 26 (anion exchanger), member 2; Sulfate transporter. May play a role in endochondral bone formation; Solute carriers |

| | | | | |
|---|---|---|---|---|
| | | UNC93B1 | ENSP00000227471 | Unc-93 homolog B1 (C. elegans); Plays an important role in innate and adaptive immunity by regulating nucleotide-sensing Toll-like receptor (TLR) signaling. Required for the |
| 103 | 3 | DAP | ENSP00000230895 | Death-associated protein 1; Negative regulator of autophagy. Involved in mediating interferon-gamma-induced cell death. |
| | | HSPB1 | ENSP00000248553 | Estrogen-regulated 24 kDa protein; Small heat shock protein which functions as a molecular chaperone probably maintaining denatured proteins in a folding- competent sta |
| | | HSPB2 | ENSP00000302476 | Heat shock 27kDa protein 2; May regulate the kinase DMPK; Small heat shock proteins |
| 104 | 3 | NPL | ENSP00000258317 | N-acetylneuraminate pyruvate lyase (dihydrodipicolinate synthase); Catalyzes the cleavage of N-acetylneuraminic acid (sialic acid) to form pyruvate and N-acetylmannosamin |
| | | PITPNC1 | ENSP00000464006 | Phosphatidylinositol transfer protein, cytoplasmic 1; Phosphatidylinositol transfer proteins mediate the monomeric transport of lipids by shielding a lipid from the aqueous |
| | | TTPA | ENSP00000260116 | Tocopherol (alpha) transfer protein; Binds alpha-tocopherol, enhances its transfer between separate membranes, and stimulates its release from liver cells. Binds both phos |
| 105 | 3 | FAM107A | ENSP00000419124 | Family with sequence similarity 107, member A; When transfected into cell lines in which it is not expressed, suppresses cell growth. May play a role in tumor development. |
| | | MLC1 | ENSP00000310375 | Megalencephalic leukoencephalopathy with subcortical cysts 1; Regulates the response of astrocytes to hypo-osmosis by promoting calcium influx. |
| | | NEXN | ENSP00000333938 | Nexilin (F actin binding protein); Involved in regulating cell migration through association with the actin cytoskeleton. Has an essential role in the maintenance of Z line and |
| 106 | 3 | ACOX2 | ENSP00000307697 | 3-alpha,7-alpha,12-alpha-trihydroxy-5-beta-cholestanoyl-CoA 24-hydroxylase; Oxidizes the CoA esters of the bile acid intermediates di- and tri-hydroxycholestanoic acids; Be |
| | | ACSF2 | ENSP00000401831 | Acyl-CoA synthetase family member 2, mitochondrial; Acyl-CoA synthases catalyze the initial reaction in fatty acid metabolism, by forming a thioester with CoA. Has some pr |
| | | HSDL2 | ENSP00000381785 | Short chain dehydrogenase/reductase family 13C member 1; Has apparently no steroid dehydrogenase activity; Belongs to the short-chain dehydrogenases/reductases (SDR |
| 107 | 3 | PRKX | ENSP00000262848 | cAMP-dependent protein kinase catalytic subunit PRKX; Serine/threonine protein kinase regulated by and mediating cAMP signaling in cells. Acts through phosphorylation o |
| | | TBL1X | ENSP00000217964 | F-box-like/WD repeat-containing protein TBL1X; F-box-like protein involved in the recruitment of the ubiquitin/19S proteasome complex to nuclear receptor-regulated trans |
| | | TRIM47 | ENSP00000254816 | Gene overexpressed in astrocytoma protein; Tripartite motif containing 47; Ring finger proteins |
| 108 | 3 | SLC2A10 | ENSP00000352216 | Solute carrier family 2 (facilitated glucose transporter), member 10; Facilitative glucose transporter; Solute carriers |
| | | SLC5A10 | ENSP00000379008 | Solute carrier family 5 (sodium/sugar cotransporter), member 10; High capacity transporter for mannose and fructose and, to a lesser extent, glucose, AMG, and galactose; B |
| | | SMOX | ENSP00000478305 | Polyamine oxidase 1; Flavoenzyme which catalyzes the oxidation of spermine to spermidine. Can also use N(1)-acetylspermine and spermidine as substrates, with different a |
| 109 | 3 | PDZK1IP1 | ENSP00000294338 | 17 kDa membrane-associated protein; May play an important role in tumor biology. |
| | | PIEZO1 | ENSP00000301015 | Piezo-type mechanosensitive ion channel component 1; Pore-forming subunit of a mechanosensitive non-specific cation channel. Generates currents characterized by a line |
| | | RBPMS | ENSP00000340176 | RNA binding protein with multiple splicing; Acts as a coactivator of transcriptional activity. Required to increase TGFB1/Smad-mediated transactivation. Acts through SMAD2 |
| 110 | 3 | GPSM3 | ENSP00000364180 | Activator of G-protein signaling 4; Interacts with subunit of G(i) alpha proteins and regulates the activation of G(i) alpha proteins. |
| | | SH2D2A | ENSP00000376123 | SH2 domain-containing adapter protein; Could be a T-cell-specific adapter protein involved in the control of T-cell activation. May play a role in the CD4-p56- LCK-dependent |
| | | SUCLG2 | ENSP00000419325 | Succinate--CoA ligase [GDP-forming] subunit beta, mitochondrial; GTP-specific succinyl-CoA synthetase functions in the citric acid cycle (TCA), coupling the hydrolysis of succi |
| 111 | 3 | COL6A2 | ENSP00000300527 | Collagen alpha-2(VI) chain; Collagen VI acts as a cell-binding protein; Collagens |
| | | CRTAP | ENSP00000323696 | Cartilage associated protein; Necessary for efficient 3-hydroxylation of fibrillar collagen prolyl residues; Belongs to the leprecan family. |
| | | SERPINH1 | ENSP00000434412 | Serpin peptidase inhibitor, clade H (heat shock protein 47), member 1, (collagen binding protein 1); Binds specifically to collagen. Could be involved as a chaperone in the bi |
| 112 | 3 | CMTM3 | ENSP00000400482 | CKLF like MARVEL transmembrane domain containing 3; Belongs to the chemokine-like factor family. |
| | | CMTM6 | ENSP00000205636 | CKLF like MARVEL transmembrane domain containing 6; Belongs to the chemokine-like factor family. |
| | | TSPAN11 | ENSP00000261177 | Tetraspanin 11; Tetraspanins |
| 113 | 3 | APCDD1 | ENSP00000347433 | Adenomatosis polyposis coli down-regulated 1 protein; Negative regulator of the Wnt signaling pathway. Inhibits Wnt signaling in a cell-autonomous manner and functions |
| | | MMD2 | ENSP00000384690 | Monocyte to macrophage differentiation associated 2; Progestin and adipoQ receptor family |
| | | PAQR8 | ENSP00000406197 | Progesterone and adipoQ receptor family member 8; Plasma membrane progesterone (P4) receptor coupled to G proteins. Seems to act through a G(i) mediated pathway. |
| 114 | 3 | BOD1L1 | ENSP00000040738 | Biorientation of chromosomes in cell division protein 1-like 1; Component of the fork protection machinery required to protect stalled/damaged replication forks from unco |
| | | C21orf62 | ENSP00000444950 | Chromosome 21 open reading frame 62 |
| | | TNS3 | ENSP00000312143 | Tensin-like SH2 domain-containing protein 1; May play a role in actin remodeling. Involved in the dissociation of the integrin-tensin-actin complex. EGF activates TNS4 and d |
| 115 | 3 | BATF2 | ENSP00000301887 | Basic leucine zipper transcriptional factor ATF-like 2; AP-1 family transcription factor that controls the differentiation of lineage-specific cells in the immune system. Followin |
| | | LYPD1 | ENSP00000380605 | Ly6/PLAUR domain-containing protein 1; Believed to act as a modulator of nicotinic acetylcholine receptors (nAChRs) activity. In vitro increases receptor desensitization and |
| | | PLAC9 | ENSP00000361337 | Placenta-specific protein 9; Placenta specific 9; Belongs to the PLAC9 family. |
| 116 | 3 | LATS2 | ENSP00000372035 | Kinase phosphorylated during mitosis protein; Negative regulator of YAP1 in the Hippo signaling pathway that plays a pivotal role in organ size control and tumor suppressio |
| | | MOB3C | ENSP00000271139 | Mps one binder kinase activator-like 2C; May regulate the activity of kinases; MOB kinase activators |
| | | TNIP2 | ENSP00000321203 | A20-binding inhibitor of NF-kappa-B activation 2; Inhibits NF-kappa-B activation by blocking the interaction of RIPK1 with its downstream effector NEMO/IKBKG. Forms a ter |
| 117 | 2 | ZBTB47 | ENSP00000232974 | Zinc finger and BTB domain-containing protein 47; May be involved in transcriptional regulation; Belongs to the krueppel C2H2-type zinc-finger protein family. |
| | | ZNF395 | ENSP00000340494 | Huntington disease gene regulatory region-binding protein 2; Plays a role in papillomavirus genes transcription; Zinc fingers C2H2-type |
| 118 | 2 | MPZL2 | ENSP00000278937 | Myelin protein zero-like protein 2; Mediates homophilic cell-cell adhesion; Belongs to the myelin P0 protein family. |
| | | ZBTB20 | ENSP00000419153 | Zinc finger and BTB domain-containing protein 20; May be a transcription factor that may be involved in hematopoiesis, oncogenesis, and immune responses. Plays a role in |
| 119 | 2 | MID1 | ENSP00000312678 | RING-type E3 ubiquitin transferase Midline-1; Has E3 ubiquitin ligase activity towards IGBP1, promoting its monoubiquitination, which results in deprotection of the catalytic |
| | | UBTD1 | ENSP00000359698 | Ubiquitin domain-containing protein 1; May be involved in the regulation of cellular senescence through a positive feedback loop with TP53. Is a TP53 downstream target ge |
| 120 | 1 | TINAGL1 | ENSP00000271064 | Tubulointerstitial nephritis antigen-related protein; May be implicated in the adrenocortical zonation and in mechanisms for repressing the CYP11B1 gene expression in adr |
| 121 | 2 | CABLES1 | ENSP00000256925 | CDK5 and ABL1 enzyme substrate 1; Cyclin-dependent kinase binding protein. Enhances cyclin-dependent kinase tyrosine phosphorylation by nonreceptor tyrosine kinases, |
| | | EFEMP1 | ENSP00000378058 | EGF containing fibulin-like extracellular matrix protein 1; Binds EGFR, the EGF receptor, inducing EGFR autophosphorylation and the activation of downstream signaling path |
| 122 | 1 | GRAMD1C | ENSP00000350881 | GRAM domain containing 1C |
| 123 | 2 | YBX3 | ENSP00000228251 | Single-strand DNA-binding protein NF-GMB; Binds to the GM-CSF promoter. Seems to act as a repressor. Binds also to full-length mRNA and to short RNA sequences contain |
| | | ZC3HAV1 | ENSP00000242351 | ADP-ribosyltransferase diphtheria toxin-like 13; Antiviral protein which inhibits the replication of viruses by recruiting the cellular RNA degradation machineries to degrade |
| 124 | 2 | SERPINB1 | ENSP00000370115 | Serpin peptidase inhibitor, clade B (ovalbumin), member 1; Regulates the activity of the neutrophil proteases elastase, cathepsin G, proteinase-3, chymase, chymotrypsin, an |
| | | TES | ENSP00000350937 | Testis derived transcript (3 LIM domains); Scaffold protein that may play a role in cell adhesion, cell spreading and in the reorganization of the actin cytoskeleton. Plays a rol |
| 125 | 2 | ACSM5 | ENSP00000327916 | Acyl-CoA synthetase medium-chain family member 5; Has medium-chain fatty acid:CoA ligase activity with broad substrate specificity (in vitro). Acts on acids from C(4) to C(1 |
| | | CACFD1 | ENSP00000440832 | Calcium channel flower domain containing 1 |
| 126 | 2 | C2CD4B | ENSP00000369755 | C2 calcium-dependent domain-containing protein 4B; May be involved in inflammatory process. May regulate cell architecture and adhesion; Belongs to the C2CD4 family. |
| | | HILPDA | ENSP00000257696 | Hypoxia-inducible lipid droplet-associated protein; Increases intracellular lipid accumulation. Stimulates expression of cytokines including IL6, MIF and VEGFA. Enhances cell |
| 127 | 2 | DAG1 | ENSP00000442600 | Dystroglycan 1 (dystrophin-associated glycoprotein 1); The dystroglycan complex is involved in a number of processes including laminin and basement membrane assembly, |
| | | RAPSN | ENSP00000298854 | 43 kDa receptor-associated protein of the synapse; Postsynaptic protein required for clustering of nicotinic acetylcholine receptors (nAChRs) at the neuromuscular junction. |
| 128 | 2 | GCSH | ENSP00000319531 | Glycine cleavage system protein H (aminomethyl carrier); The glycine cleavage system catalyzes the degradation of glycine. The H protein (GCSH) shuttles the methylamine g |
| | | SARDH | ENSP00000360938 | Sarcosine dehydrogenase, mitochondrial; Sarcosine dehydrogenase; Belongs to the GcvT family. |
| 129 | 2 | METRNL | ENSP00000315731 | Meteorin, glial cell differentiation regulator-like; Hormone induced following exercise or cold exposure that promotes energy expenditure. Induced either in the skeletal mu |
| | | OTOS | ENSP00000375849 | Otospiralin; May be essential for the survival of the neurosensory epithelium of the inner ear. |
| 130 | 2 | COLEC12 | ENSP00000383115 | Scavenger receptor with C-type lectin; Scavenger receptor that displays several functions associated with host defense. Promotes binding and phagocytosis of Gram-positive, |
| | | LIMK2 | ENSP00000339916 | LIM domain kinase 2; Displays serine/threonine-specific phosphorylation of myelin basic protein and histone (MBP) in vitro; LIM domain containing |
| 131 | 2 | MAP3K6 | ENSP00000419591 | Mitogen-activated protein kinase kinase kinase 6; Component of a protein kinase signal transduction cascade. Activates the JNK, but not ERK or p38 kinase pathways. |
| | | POLD4 | ENSP00000311368 | Polymerase (DNA-directed), delta 4, accessory subunit; As a component of the tetrameric DNA polymerase delta complex (Pol-delta4), plays a role in high fidelity genome re |
| 132 | 2 | MTRNR2L12 | ENSP00000468991 | MT-RNR2-like protein 8; Plays a role as a neuroprotective and antiapoptotic factor; Belongs to the humanin family. |
| | | MTRNR2L8 | ENSP00000439666 | MT-RNR2-like 8; Plays a role as a neuroprotective and antiapoptotic factor; Belongs to the humanin family. |
| 133 | 2 | EMILIN2 | ENSP00000254528 | Elastin microfibril interface-located protein 2; May be responsible for anchoring smooth muscle cells to elastic fibers, and may be involved not only in the formation of the el |
| | | MYOM1 | ENSP00000348821 | 190 kDa connectin-associated protein; Major component of the vertebrate myofibrillar M band. Binds myosin, titin, and light meromyosin. This binding is dose dependent; F |
| 134 | 2 | ANKHD1 | ENSP00000354085 | Ankyrin repeat and KH domain-containing protein 1; May play a role as a scaffolding protein that may be associated with the abnormal phenotype of leukemia cells. Isoform |
| | | SIPA1 | ENSP00000377771 | Signal-induced proliferation-associated protein 1; GTPase activator for the nuclear Ras-related regulatory proteins Rap1 and Rap2 in vitro, converting them to the putatively |
| 135 | 2 | RXRA | ENSP00000419692 | Nuclear receptor subfamily 2 group B member 1; Receptor for retinoic acid. Retinoic acid receptors bind as heterodimers to their target response elements in response to th |
| | | RXRG | ENSP00000352900 | Nuclear receptor subfamily 2 group B member 3; Receptor for retinoic acid. Retinoic acid receptors bind as heterodimers to their target response elements in response to th |
| 136 | 2 | MSRB3 | ENSP00000347324 | Methionine-R-sulfoxide reductase B3; Catalyzes the reduction of free and protein-bound methionine sulfoxide to methionine. Isoform 2 is essential for hearing; Deafness as |
| | | RFX2 | ENSP00000306335 | Regulatory factor X, 2 (influences HLA class II expression); Transcription factor that acts as a key regulator of spermatogenesis. Acts by regulating expression of genes require |
| 137 | 2 | BRCA1 | ENSP00000418960 | Breast cancer type 1 susceptibility protein; E3 ubiquitin-protein ligase that specifically mediates the formation of 'Lys-6'-linked polyubiquitin chains and plays a central role i |
| | | MCM7 | ENSP00000307288 | Minichromosome maintenance complex component 7; Acts as component of the MCM2-7 complex (MCM complex) which is the putative replicative helicase essential for 'o |
| 138 | 2 | CCDC80 | ENSP00000206423 | Up-regulated in BRS-3 deficient mouse homolog; Promotes cell adhesion and matrix assembly. |
| | | SAMD4A | ENSP00000375919 | Sterile alpha motif domain-containing protein 4A; Acts as a translational repressor of SRE-containing messengers; Belongs to the SMAUG family. |
| 139 | 2 | ASRGL1 | ENSP00000400057 | Isoaspartyl peptidase/L-asparaginase; Has both L-asparaginase and beta-aspartyl peptidase activity. May be involved in the production of L-aspartate, which can act as an ex |
| | | ECE1 | ENSP00000364028 | Endothelin converting enzyme 1; Converts big endothelin-1 to endothelin-1; Belongs to the peptidase M13 family. |
| 140 | 2 | DTNA | ENSP00000470152 | Dystrophin-related protein 3; May be involved in the formation and stability of synapses as well as being involved in the clustering of nicotinic acetylcholine receptors; Belon |
| | | FYCO1 | ENSP00000296137 | FYVE and coiled-coil domain-containing protein 1; May mediate microtubule plus end-directed vesicle transport; GOLD domain containing |
| 141 | 2 | EZR | ENSP00000356042 | Cytovillin; Probably involved in connections of major cytoskeletal structures to the plasma membrane. In epithelial cells, required for the formation of microvilli and membra |
| | | MSN | ENSP00000353408 | Membrane-organizing extension spike protein; Probably involved in connections of major cytoskeletal structures to the plasma membrane. May inhibit herpes simplex virus |
| 142 | 2 | ALDH1L1 | ENSP00000273450 | Cytosolic 10-formyltetrahydrofolate dehydrogenase; Aldehyde dehydrogenase 1 family member L1; In the N-terminal section; belongs to the GART family. |
| | | SULT1C4 | ENSP00000272452 | Sulfotransferase family, cytosolic, 1C, member 4; Sulfotransferase that utilizes 3'-phospho-5'-adenylyl sulfate (PAPS) as sulfonate donor to catalyze the sulfate conjugation of |
| 143 | 2 | PERP | ENSP00000397157 | P53 apoptosis effector related to PMP-22; Component of intercellular desmosome junctions. Plays a role in stratified epithelial integrity and cell-cell adhesion by promoting |
| | | RNF144B | ENSP00000259939 | E3 ubiquitin-protein ligase RNF144B; E3 ubiquitin-protein ligase which accepts ubiquitin from E2 ubiquitin-conjugating enzymes UBE2L3 and UBE2L6 in the form of a thioest |
| 144 | 2 | MROH9 | ENSP00000356733 | Maestro heat like repeat family member 9 |
| | | PTPN21 | ENSP00000452414 | Protein tyrosine phosphatase, non-receptor type 21; FERM domain containing |
| 145 | 2 | ITGB3 | ENSP00000452786 | Integrin, beta 3 (platelet glycoprotein IIIa, antigen CD61); Integrin alpha-V/beta-3 (ITGAV:ITGB3) is a receptor for cytotactin, fibronectin, laminin, matrix metalloproteinase-2, |
| | | SYTL4 | ENSP00000362080 | Synaptotagmin-like protein 4; Modulates exocytosis of dense-core granules and secretion of hormones in the pancreas and the pituitary. Interacts with vesicles containing n |
| 146 | 2 | FAM181A | ENSP00000267594 | Family with sequence similarity 181 member A |
| | | TEAD4 | ENSP00000352926 | Transcriptional enhancer factor TEF-3; Transcription factor which plays a key role in the Hippo signaling pathway, a pathway involved in organ size control and tumor suppre |
| 147 | 2 | CCDC69 | ENSP00000347586 | Coiled-coil domain-containing protein 69; May act as a scaffold to regulate the recruitment and assembly of spindle midzone components. Required for the localization of A |
| | | CRISPLD2 | ENSP00000262424 | Cysteine-rich secretory protein LCCL domain containing 2; Promotes matrix assembly; Belongs to the CRISP family. |
| 148 | 2 | METTL7B | ENSP00000377796 | Methyltransferase-like protein 7B; Probable methyltransferase. |
| | | RARRES2 | ENSP00000418009 | Retinoic acid receptor responder (tazarotene induced) 2; Adipocyte-secreted protein (adipokine) that regulates adipogenesis, metabolism and inflammation through activati |
| 149 | 2 | CASKIN2 | ENSP00000325355 | CASK interacting protein 2; Sterile alpha motif domain containing |
| | | LIMS2 | ENSP00000326888 | LIM and senescent cell antigen-like-containing domain protein 2; Adapter protein in a cytoplasmic complex linking beta- integrins to the actin cytoskeleton, bridges the comp |
| 150 | 2 | NANOS1 | ENSP00000393275 | Nanos homolog 1 (Drosophila); May act as a translational repressor which regulates translation of specific mRNAs by forming a complex with PUM2 that associates with the |
| | | PLA2G5 | ENSP00000364249 | Phosphatidylcholine 2-acylhydrolase 5; PA2 catalyzes the calcium-dependent hydrolysis of the 2- acyl groups in 3-sn-phosphoglycerides. This isozyme hydrolyzes more efficie |
| 151 | 2 | CATSPERD | ENSP00000371037 | Cation channel sperm-associated protein subunit delta; Auxiliary component of the CatSper complex, a complex involved in sperm cell hyperactivation. Sperm cell hyperacti |
| | | OR6V1 | ENSP00000396085 | Olfactory receptor, family 6, subfamily V, member 1; Odorant receptor; Olfactory receptors, family 6 |
| 152 | 2 | CHI3L1 | ENSP00000255409 | Chitinase 3-like 1 (cartilage glycoprotein-39); Carbohydrate-binding lectin with a preference for chitin. Has no chitinase activity. May play a role in tissue remodeling and in t |
| | | TMEM219 | ENSP00000457492 | Insulin-like growth factor-binding protein 3 receptor; Cell death receptor specific for IGFBP3, may mediate caspase-8-dependent apoptosis upon ligand binding. |
| 153 | 2 | NDRG1 | ENSP00000404854 | Reducing agents and tunicamycin-responsive protein; Stress-responsive protein involved in hormone responses, cell growth, and differentiation. Acts as a tumor suppressor |
| | | NDRG2 | ENSP00000451712 | N-myc downstream-regulated gene 2 protein; Contributes to the regulation of the Wnt signaling pathway. Down-regulates CTNNB1-mediated transcriptional activation of ta |
| 154 | 2 | CD99 | ENSP00000370588 | T-cell surface glycoprotein E2; Involved in T-cell adhesion processes and in spontaneous rosette formation with erythrocytes. Plays a role in a late step of leukocyte extravas |
| | | N4BP2 | ENSP00000261435 | NEDD4 binding protein 2; Has 5'-polynucleotide kinase and nicking endonuclease activity. May play a role in DNA repair or recombination. |
| 155 | 2 | MORC4 | ENSP00000347821 | Zinc finger CW-type coiled-coil domain protein 2; MORC family CW-type zinc finger 4 |
| | | TPD52L1 | ENSP00000434142 | Tumor protein D52 like 1 |
| 156 | 1 | TRIL | ENSP00000479256 | Leucine-rich repeat-containing protein KIAA0644; Component of the TLR4 signaling complex. Mediate the innate immune response to bacterial lipopolysaccharide (LPS) lead |
| 157 | 2 | FAM167B | ENSP00000362684 | Family with sequence similarity 167 member B; Belongs to the FAM167 (SEC) family. |
| | | STK33 | ENSP00000416750 | Serine/threonine-protein kinase 33; Serine/threonine protein kinase which phosphorylates VIME. May play a specific role in the dynamic behavior of the intermediate filam |
| 158 | 2 | LHX2 | ENSP00000362717 | LIM/homeobox protein Lhx2; Acts as a transcriptional activator. Stimulates the promoter of the alpha-glycoprotein gene. Transcriptional regulatory protein involved in the c |
| | | NEK7 | ENSP00000356355 | Serine/threonine-protein kinase Nek7; Protein kinase which plays an important role in mitotic cell cycle progression. Required for microtubule nucleation activity of the cent |
| 159 | 1 | APOL3 | ENSP00000344577 | TNF-inducible protein CG12-1; May affect the movement of lipids in the cytoplasm or allow the binding of lipids to organelles; Apolipoproteins |
| 160 | 2 | BNIP2 | ENSP00000267859 | BCL2/adenovirus E1B 19 kDa protein-interacting protein 2; Implicated in the suppression of cell death. Interacts with the BCL-2 and adenovirus E1B 19 kDa proteins; BCH do |
| | | PRR5 | ENSP00000384848 | Protein observed with Rictor-1; Subunit of mTORC2, which regulates cell growth and survival in response to hormonal signals. mTORC2 is activated by growth factors, but, in |
| 161 | 2 | LGALS3 | ENSP00000254301 | Lectin, galactoside-binding, soluble, 3; Galactose-specific lectin which binds IgE. May mediate with the alpha-3, beta-1 integrin the stimulation by CSPG4 of endothelial cells |
| | | UACA | ENSP00000314556 | Uveal autoantigen with coiled-coil domains and ankyrin repeats; Regulates APAF1 expression and plays an important role in the regulation of stress-induced apoptosis. Prom |
| 162 | 2 | MID1IP1 | ENSP00000483547 | Gastrulation-specific G12-like protein; Plays a role in the regulation of lipogenesis in liver. Up-regulates ACACA enzyme activity. Required for efficient lipid biosynthesis, inclu |
| | | PLIN4 | ENSP00000301286 | Adipocyte protein S3-12; May play a role in triacylglycerol packaging into adipocytes. May function as a coat protein involved in the biogenesis of lipid droplets (By similarity |
| 163 | 1 | APOL4 | ENSP00000338260 | Apolipoprotein L, 4; May play a role in lipid exchange and transport throughout the body. May participate in reverse cholesterol transport from peripheral cells to the liver ( |
| 164 | 2 | ARSD | ENSP00000370546 | Arylsulfatase D; Sulfatases |
| | | C8orf82 | ENSP00000436621 | Chromosome 8 open reading frame 82 |
| 165 | 2 | CD109 | ENSP00000287097 | C3 and PZP-like alpha-2-macroglobulin domain-containing protein 7; Modulates negatively TGFB1 signaling in keratinocytes; Belongs to the protease inhibitor I39 (alpha-2- |

| | | | | |
|---|---|---|---|---|
| | | PEAR1 | ENSP00000344465 | Multiple epidermal growth factor-like domains protein 12; When overexpressed, reduces the number of both early and late non-adherent myeloid progenitor cells; Belongs |
| 166 | 2 | EMP1 | ENSP00000256951 | Epithelial membrane protein 1 |
| | | SMIM3 | ENSP00000436897 | NGF-induced differentiation clone 67 protein; Small integral membrane protein 3 |
| 167 | 2 | SLC4A4 | ENSP00000393557 | Solute carrier family 4 (sodium bicarbonate cotransporter), member 4; Electrogenic sodium/bicarbonate cotransporter with a Na(+):HCO3(-) stoichiometry varying from 1:2 t |
| | | WNK4 | ENSP00000246914 | WNK lysine deficient protein kinase 4; Serine/threonine kinase which plays an important role in the regulation of electrolyte homeostasis, cell signaling, survival and prolifera |
| 168 | 2 | ARHGEF26 | ENSP00000348828 | Rho guanine nucleotide exchange factor (GEF) 26; Activates RhoG GTPase by promoting the exchange of GDP by GTP. Required for the formation of membrane ruffles during |
| | | DOCK1 | ENSP00000280333 | Dedicator of cytokinesis protein 1; Involved in cytoskeletal rearrangements required for phagocytosis of apoptotic cells and cell motility. Along with DOCK1, mediates CRK/C |
| 169 | 2 | MSI2 | ENSP00000284073 | RNA-binding protein Musashi homolog 2; RNA binding protein that regulates the expression of target mRNAs at the translation level. May play a role in the proliferation and |
| | | SNTA1 | ENSP00000217381 | 59 kDa dystrophin-associated protein A1 acidic component 1; Adapter protein that binds to and probably organizes the subcellular localization of a variety of membrane pro |
| 170 | 2 | LPP | ENSP00000482472 | LIM domain containing preferred translocation partner in lipoma; May play a structural role at sites of cell adhesion in maintaining cell shape and motility. In addition to the |
| | | VASP | ENSP00000245932 | Vasodilator-stimulated phosphoprotein; Ena/VASP proteins are actin-associated proteins involved in a range of processes dependent on cytoskeleton remodeling and cell po |
| 171 | 1 | ARHGAP11A | ENSP00000355090 | Rho GTPase activating protein 11A |
| 172 | 2 | C11orf96 | ENSP00000479976 | Chromosome 11 open reading frame 96 |
| | | TMCO3 | ENSP00000389399 | Transmembrane and coiled-coil domain-containing protein 3; Probable Na(+)/H(+) antiporter; Belongs to the monovalent cation:proton antiporter 2 (CPA2) transporter (TC |
| 173 | 2 | AEBP1 | ENSP00000223357 | Adipocyte enhancer-binding protein 1; May positively regulate MAP-kinase activity in adipocytes, leading to enhanced adipocyte proliferation and reduced adipocyte differe |
| | | MOCS1 | ENSP00000362282 | Molybdenum cofactor synthesis-step 1 protein A-B; Isoform MOCS1A and isoform MOCS1B probably form a complex that catalyzes the conversion of 5'-GTP to cyclic pyrano |
| 174 | 1 | ITSN2 | ENSP00000347244 | SH3P18-like WASP-associated protein; Adapter protein that may provide indirect link between the endocytic membrane traffic and the actin assembly machinery. May regul |
| 175 | 2 | CYP21A2 | ENSP00000408860 | Cytochrome P450, family 21, subfamily A, polypeptide 2; Specifically catalyzes the 21-hydroxylation of steroids. Required for the adrenal synthesis of mineralocorticoids and |
| | | CYP4F11 | ENSP00000384588 | Cytochrome P450, family 4, subfamily F, polypeptide 11; Omega-hydroxylase that oxidizes a variety of structurally unrelated compounds, including fatty acids and xenobiotic |
| 176 | 1 | PLEKHB1 | ENSP00000346127 | Pleckstrin homology domain containing, family B (evectins) member 1; Required for proper localization of retinogeniculate projections but not for eye-specific segregation; |
| 177 | 2 | FAM114A1 | ENSP00000351740 | Family with sequence similarity 114, member A1; May play a role in neuronal cell development; Belongs to the FAM114 family. |
| | | KLHL5 | ENSP00000423897 | Kelch like family member 5; BTB domain containing |
| 178 | 2 | CTNS | ENSP00000371294 | Cystinosin, lysosomal cystine transporter; Cystine/H(+) symporter thought to transport cystine out of lysosomes. Plays an important role in melanin synthesis, possibly by pre |
| | | PIRT | ENSP00000462046 | Phosphoinositide-interacting regulator of transient receptor potential channels; Regulatory subunit of TRPV1, a molecular sensor of noxious heat and capsaicin. Positively re |
| 179 | 2 | BTN2A2 | ENSP00000349143 | Butyrophilin, subfamily 2, member A2; Inhibits the proliferation of CD4 and CD8 T-cells activated by anti-CD3 antibodies, T-cell metabolism and IL2 and IFNG secretion; Belor |
| | | TTC23 | ENSP00000377690 | Cervical cancer proto-oncogene 8 protein; Tetratricopeptide repeat domain containing |
| 180 | 2 | GSDMD | ENSP00000433209 | Gasdermin domain-containing protein 1; Gasdermin-D, N-terminal: Promotes pyroptosis in response to microbial infection and danger signals. Produced by the cleavage of |
| | | LRRC3 | ENSP00000291592 | Leucine-rich repeat-containing protein 3; Leucine rich repeat containing 3 |
| 181 | 2 | SOWAHC | ENSP00000365830 | Sosondowah ankyrin repeat domain family member C; Belongs to the SOWAH family. |
| | | SYDE1 | ENSP00000341489 | Synapse defective 1, Rho GTPase, homolog 1 (C. elegans); GTPase activator for the Rho-type GTPases. As a GCM1 downstream effector, it is involved in placental developmen |
| 182 | 2 | CARD10 | ENSP00000384570 | Caspase recruitment domain-containing protein 10; Activates NF-kappa-B via BCL10 and IKK; Caspase recruitment domain containing |
| | | LPAR1 | ENSP00000363553 | Lysophosphatidic acid receptor Edg-2; Receptor for lysophosphatidic acid (LPA). Plays a role in the reorganization of the actin cytoskeleton, cell migration, differentiation and |
| 183 | 2 | CCDC15 | ENSP00000341684 | Coiled-coil domain containing 15 |
| | | NSRP1 | ENSP00000247026 | Nuclear speckle splicing regulatory protein 1; RNA-binding protein that mediates pre-mRNA alternative splicing regulation. |
| 184 | 2 | BACE2 | ENSP00000332979 | Beta-site amyloid precursor protein cleaving enzyme 2; Responsible for the proteolytic processing of the amyloid precursor protein (APP). Cleaves APP, between residues 69 |
| | | CACHD1 | ENSP00000290039 | VWFA and cache domain-containing protein 1; May regulate voltage-dependent calcium channels. |
| 185 | 2 | GMPR | ENSP00000259727 | Guanosine 5'-monophosphate oxidoreductase 1; Catalyzes the irreversible NADPH-dependent deamination of GMP to IMP. It functions in the conversion of nucleobase, nuc |
| | | NUDT16 | ENSP00000422375 | Nudix (nucleoside diphosphate linked moiety X)-type motif 16; RNA-binding and decapping enzyme that catalyzes the cleavage of the cap structure of snoRNAs and mRNAs i |
| 186 | 2 | AIFM3 | ENSP00000382120 | Apoptosis-inducing factor, mitochondrion-associated, 3; Induces apoptosis through a caspase dependent pathway. Reduces mitochondrial membrane potential. |
| | | SMTN | ENSP00000484398 | Smoothelin; Structural protein of the cytoskeleton; Belongs to the smoothelin family. |
| 187 | 2 | FIBIN | ENSP00000321962 | Fin bud initiation factor homolog; Belongs to the FIBIN family. |
| | | HVCN1 | ENSP00000349181 | Voltage sensor domain-only protein; Mediates the voltage-dependent proton permeability of excitable membranes. Forms a proton-selective channel through which proton |
| 188 | 2 | MIDN | ENSP00000300952 | Midbrain nucleolar protein; Facilitates ubiquitin-independent proteasomal degradation of polycomb protein CBX4. Plays a role in inhibiting the activity of glucokinase GCK a |
| | | ODF3L2 | ENSP00000318029 | Outer dense fiber of sperm tails 3 like 2 |
| 189 | 2 | DHRS3 | ENSP00000480439 | Short chain dehydrogenase/reductase family 16C member 1; Catalyzes the reduction of all-trans-retinal to all- trans-retinol in the presence of NADPH; Short chain dehydroge |
| | | ELP4 | ENSP00000379267 | Elongator acetyltransferase complex subunit 4; Acts as subunit of the RNA polymerase II elongator complex, which is a histone acetyltransferase component of the RNA poly |
| 190 | 2 | ATOH8 | ENSP00000304676 | Class A basic helix-loop-helix protein 21; Transcription factor that binds a palindromic (canonical) core consensus DNA sequence 5'-CANNTG- 3' known as an E-box element, |
| | | OLFML1 | ENSP00000332511 | Olfactomedin-like protein 1; Olfactomedin like 1 |
| 191 | 2 | LRRIQ1 | ENSP00000376910 | Leucine rich repeats and IQ motif containing 1 |
| | | SLC44A3 | ENSP00000271227 | Choline transporter-like protein 3; Solute carrier family 44 member 3 |
| 192 | 2 | CLCA4 | ENSP00000359594 | Calcium-activated chloride channel family member 4; May be involved in mediating calcium-activated chloride conductance; Chloride channel accessory |
| | | FCGBP | ENSP00000481056 | Fc fragment of IgG binding protein |
| 193 | 2 | FGF1 | ENSP00000480791 | Fibroblast growth factor 1 (acidic); Plays an important role in the regulation of cell survival, cell division, angiogenesis, cell differentiation and cell migration. Functions as pot |
| | | FGFRL1 | ENSP00000381498 | Fibroblast growth factor receptor-like 1; Has a negative effect on cell proliferation; I-set domain containing |
| 194 | 2 | FAM111A | ENSP00000434435 | Family with sequence similarity 111, member A; Chromatin-associated protein required for PCNA loading on replication sites. Promotes S-phase entry and DNA synthesis. M |
| | | NKAIN4 | ENSP00000359340 | Sodium/potassium-transporting ATPase subunit beta-1-interacting protein 4; Sodium/potassium transporting ATPase interacting 4; Belongs to the NKAIN family. |
| 195 | 2 | PSKH1 | ENSP00000291041 | Serine/threonine-protein kinase H1; May be a SFC-associated serine kinase (splicing factor compartment-associated serine kinase) with a role in intranuclear SR protein (non |
| | | ZCCHC24 | ENSP00000361411 | Zinc finger CCHC-type containing 24; Zinc fingers 3CxxC-type |
| 196 | 2 | ETNPPL | ENSP00000296486 | Alanine--glyoxylate aminotransferase 2-like 1; Catalyzes the pyridoxal-phosphate-dependent breakdown of phosphoethanolamine, converting it to ammonia, inorganic phos |
| | | PHYHD1 | ENSP00000309515 | Phytanoyl-CoA dioxygenase domain-containing protein 1; Isoform 1 has alpha-ketoglutarate-dependent dioxygenase activity. Does not show detectable activity towards fatty |
| 197 | 2 | FGR | ENSP00000363117 | Gardner-Rasheed feline sarcoma viral (v-fgr) oncogene homolog; Non-receptor tyrosine-protein kinase that transmits signals from cell surface receptors devoid of kinase acti |
| | | SIGLEC9 | ENSP00000413861 | Sialic acid binding Ig-like lectin 9; Putative adhesion molecule that mediates sialic-acid dependent binding to cells. Preferentially binds to alpha-2,3- or alpha-2,6-linked sialic |
| 198 | 2 | ANKRD36 | ENSP00000391950 | Ankyrin repeat domain-containing protein 36A; Ankyrin repeat domain containing |
| | | ANKRD36C | ENSP00000403302 | Protein immuno-reactive with anti-PTH polyclonal antibodies; Ankyrin repeat domain containing |
| 199 | 2 | RUFY4 | ENSP00000363270 | RUN and FYVE domain containing 4; Zinc fingers FYVE-type |
| | | TMBIM1 | ENSP00000409738 | Transmembrane BAX inhibitor motif-containing protein 1; Negatively regulates aortic matrix metalloproteinase-9 (MMP9) production and may play a protective role in vascu |
| 200 | 2 | CSRNP1 | ENSP00000273153 | Cysteine-serine-rich nuclear protein 1; Binds to the consensus sequence 5'-AGAGTG-3' and has transcriptional activator activity (By similarity). May have a tumor-suppressor |
| | | RASSF2 | ENSP00000368710 | Ras association (RalGDS/AF-6) domain family member 2; Potential tumor suppressor. Acts as a KRAS-specific effector protein. May promote apoptosis and cell cycle arrest. S |
| 201 | 1 | SELENBP1 | ENSP00000397261 | 56 kDa selenium-binding protein; Selenium-binding protein which may be involved in the sensing of reactive xenobiotics in the cytoplasm. May be involved in intra-Golgi pro |
| 202 | 2 | LGALS9 | ENSP00000378856 | Lectin, galactoside-binding, soluble, 9; Binds galactosides. Has high affinity for the Forssman pentasaccharide. Ligand for HAVCR2/TIM3. Binding to HAVCR2 induces T-helper |
| | | SLC12A7 | ENSP00000264930 | Solute carrier family 12 (potassium/chloride transporter), member 7; Mediates electroneutral potassium-chloride cotransport when activated by cell swelling. May mediate |
| 203 | 2 | LCN10 | ENSP00000418491 | Epididymal-specific lipocalin-10; Lipocalin 10; Belongs to the calycin superfamily. Lipocalin family. |
| | | LCN6 | ENSP00000339621 | Epididymal-specific lipocalin-6; May play a role in male fertility; Lipocalins |
| 204 | 2 | IMPA2 | ENSP00000269159 | Inositol(myo)-1(or 4)-monophosphatase 2; Can use myo-inositol monophosphates, scylloinositol 1,4- diphosphate, glucose-1-phosphate, beta-glycerophosphate, and 2'- AMP |
| | | PXDC1 | ENSP00000369636 | PX domain containing 1 |
| 205 | 2 | PTTG1IP | ENSP00000328325 | Pituitary tumor-transforming gene 1 protein-interacting protein; May facilitate PTTG1 nuclear translocation. |
| | | TPCN1 | ENSP00000448083 | Voltage-dependent calcium channel protein TPC1; Nicotinic acid adenine dinucleotide phosphate (NAADP) receptor that may function as one of the major voltage-gated Ca( |
| 206 | 2 | PRSS35 | ENSP00000358714 | Serine protease 35 |
| | | RANBP3L | ENSP00000421853 | RAN binding protein 3-like; Nuclear export factor for BMP-specific SMAD1/5/8 that plays a critical role in terminating BMP signaling and regulating mesenchymal stem cell di |
| 207 | 2 | ARPC1B | ENSP00000389631 | Actin related protein 2/3 complex, subunit 1B, 41kDa; Functions as component of the Arp2/3 complex which is involved in regulation of actin polymerization and together w |
| | | CAPG | ENSP00000263867 | Capping protein (actin filament), gelsolin-like; Calcium-sensitive protein which reversibly blocks the barbed ends of actin filaments but does not sever preformed actin filame |
| 208 | 2 | KTN1 | ENSP00000378725 | Kinectin 1 (kinesin receptor); Receptor for kinesin thus involved in kinesin-driven vesicle motility. Accumulates in integrin-based adhesion complexes (IAC) upon integrin agg |
| | | PPP1R14B | ENSP00000310117 | Protein phosphatase 1, regulatory (inhibitor) subunit 14B; Inhibitor of PPP1CA. Has over 50-fold higher inhibitory activity when phosphorylated (By similarity). |
| 209 | 2 | ASAP3 | ENSP00000338769 | Arf-GAP with SH3 domain, ANK repeat and PH domain-containing protein 3; Promotes cell proliferation; Ankyrin repeat domain containing |
| | | TCEA3 | ENSP00000406293 | Transcription elongation factor S-II protein 3; Necessary for efficient RNA polymerase II transcription elongation past template-encoded arresting sites. The arresting sites in |
| 210 | 2 | ANPEP | ENSP00000300060 | Myeloid plasma membrane glycoprotein CD13; Broad specificity aminopeptidase which plays a role in the final digestion of peptides generated from hydrolysis of proteins b |
| | | PGPEP1 | ENSP00000269919 | Pyrrolidone-carboxylate peptidase; Removes 5-oxoproline from various penultimate amino acid residues except L-proline. |
| 211 | 1 | ZNF423 | ENSP00000455426 | Smad- and Olf-interacting zinc finger protein; Transcription factor that can both act as an activator or a repressor depending on the context. Plays a central role in BMP signa |
| 212 | 2 | BBX | ENSP00000319974 | HMG box transcription factor BBX; Transcription factor that is necessary for cell cycle progression from G1 to S phase. |
| | | NACC2 | ENSP00000277554 | NACC family member 2, BEN and BTB (POZ) domain containing; Functions as a transcriptional repressor through its association with the NuRD complex. Recruits the NuRD c |
| 213 | 2 | PLSCR4 | ENSP00000347038 | Ca(2+)-dependent phospholipid scramblase 4; May mediate accelerated ATP-independent bidirectional transbilayer migration of phospholipids upon binding calcium ions th |
| | | RELL1 | ENSP00000398778 | RELT-like protein 1; RELT family; Belongs to the RELT family. |
| 214 | 2 | BUB1 | ENSP00000302530 | Mitotic checkpoint serine/threonine-protein kinase BUB1; Serine/threonine-protein kinase that performs 2 crucial functions during mitosis: it is essential for spindle-assemb |
| | | CDC45 | ENSP00000405726 | Cell division control protein 45 homolog; Required for initiation of chromosomal DNA replication. |
| 215 | 1 | TNFRSF12A | ENSP00000326737 | Fibroblast growth factor-inducible immediate-early response protein 14; Receptor for TNFSF12/TWEAK. Weak inducer of apoptosis in some cell types. Promotes angiogenes |
| 216 | 2 | PCK1 | ENSP00000319814 | Phosphoenolpyruvate carboxykinase, cytosolic [GTP]; Catalyzes the conversion of oxaloacetate (OAA) to phosphoenolpyruvate (PEP), the rate-limiting step in the metabolic |
| | | RBM38 | ENSP00000348538 | RNA-binding region-containing protein 1; RNA-binding protein that specifically bind the 3'-UTR of CDKN1A transcripts, leading to maintain the stability of CDKN1A transcripts |
| 217 | 2 | SGMS2 | ENSP00000378176 | Phosphatidylcholine:ceramide cholinephosphotransferase 2; Sphingomyelin synthases synthesize the sphingolipid, sphingomyelin, through transfer of the phosphatidyl head |
| | | USP53 | ENSP00000409906 | Inactive ubiquitin carboxyl-terminal hydrolase 53; Tight junction-associated protein that is involved in the survival of auditory hair cells and hearing. Maybe by modulating th |
| 218 | 2 | ACVRL1 | ENSP00000373574 | Serine/threonine-protein kinase receptor R3; Type I receptor for TGF-beta family ligands BMP9/GDF2 and BMP10 and important regulator of normal blood vessel developm |
| | | TMEM100 | ENSP00000465638 | Transmembrane protein 100; Plays a role during embryonic arterial endothelium differentiation and vascular morphogenesis through the ACVRL1 receptor-dependent signa |
| 219 | 2 | C16orf74 | ENSP00000284245 | Chromosome 16 open reading frame 74 |
| | | FOXF1 | ENSP00000262426 | Forkhead-related transcription factor 1; Probable transcription activator for a number of lung- specific genes; Forkhead boxes |
| 220 | 2 | C1R | ENSP00000438615 | Complement component 1, r subcomponent; C1r B chain is a serine protease that combines with C1q and C1s to form C1, the first component of the classical pathway of the |
| | | VAMP5 | ENSP00000305647 | Vesicle-associated membrane protein 5; May participate in trafficking events that are associated with myogenesis, such as myoblast fusion and/or GLUT4 trafficking; Belongs |
| 221 | 1 | ZNF618 | ENSP00000288466 | Zinc finger protein 618; Zinc fingers C2H2-type |
| 222 | 2 | BCL7C | ENSP00000369674 | B-cell CLL/lymphoma 7 protein family member C; May play an anti-apoptotic role; BAF complex |
| | | SMARCC1 | ENSP00000254480 | SWI/SNF related, matrix associated, actin dependent regulator of chromatin, subfamily c, member 1; Involved in transcriptional activation and repression of select genes by |
| 223 | 1 | BCAM | ENSP00000270233 | Basal cell adhesion molecule (Lutheran blood group); Laminin alpha-5 receptor. May mediate intracellular signaling; Blood group antigens |
| 224 | 2 | C1QTNF1 | ENSP00000340864 | Complement C1q tumor necrosis factor-related protein 1; C1q and TNF related 1 |
| | | CSPG4 | ENSP00000312506 | Melanoma-associated chondroitin sulfate proteoglycan; Proteoglycan playing a role in cell proliferation and migration which stimulates endothelial cells motility during micr |
| 225 | 2 | CDA | ENSP00000364212 | Cytidine aminohydrolase; This enzyme scavenges exogenous and endogenous cytidine and 2'-deoxycytidine for UMP synthesis. |
| | | TTC38 | ENSP00000370419 | Tetratricopeptide repeat domain containing; Belongs to the TTC38 family. |
| 226 | 2 | ARHGAP31 | ENSP00000264245 | Rho GTPase activating protein 31; Functions as a GTPase-activating protein (GAP) for RAC1 and CDC42. Required for cell spreading, polarized lamellipodia formation and cell |
| | | DOCK6 | ENSP00000294618 | Dedicator of cytokinesis protein 6; Acts as guanine nucleotide exchange factor (GEF) for CDC42 and RAC1 small GTPases. Through its activation of CDC42 and RAC1, may regu |
| 227 | 2 | CHI3L2 | ENSP00000437082 | Chitinase-3-like protein 2; Lectin that binds chitooligosaccharides and other glycans with high affinity, but not heparin. Has no chitinase activity; Belongs to the glycosyl hydro |
| | | PIFO | ENSP00000358753 | Primary cilia formation; During primary cilia disassembly, involved in cilia disassembly. Required specifically to control cilia retraction as well as the liberation and duplicatior |
| 228 | 2 | NID1 | ENSP00000264187 | Nidogen 1; Sulfated glycoprotein widely distributed in basement membranes and tightly associated with laminin. Also binds to collagen IV and perlecan. It probably has a rol |
| | | ZAN | ENSP00000480750 | Zonadhesin (gene/pseudogene); Binds in a species-specific manner to the zona pellucida of the egg. May be involved in gamete recognition and/or signaling. |
| 229 | 2 | IER3 | ENSP00000259874 | Radiation-inducible immediate-early gene IEX-1; May play a role in the ERK signaling pathway by inhibiting the dephosphorylation of ERK by phosphatase PP2A- PPP2R5C ho |
| | | PPP1R18 | ENSP00000274853 | Protein phosphatase 1 F-actin cytoskeleton-targeting subunit; Isoform 1: May target protein phosphatase 1 to F-actin cytoskeleton. |
| 230 | 2 | RIBC1 | ENSP00000364476 | RIB43A domain with coiled-coils 1; Belongs to the RIB43A family. |
| | | TOB2 | ENSP00000331305 | Transducer of ERBB2, 2; Anti-proliferative protein inhibits cell cycle progression from the G0/G1 to S phases; Belongs to the BTG family. |
| 231 | 2 | EPAS1 | ENSP00000263734 | Endothelial PAS domain-containing protein 1; Transcription factor involved in the induction of oxygen regulated genes. Binds to core DNA sequence 5'-[AG]CGTG-3' within th |
| | | HIF3A | ENSP00000366898 | Class E basic helix-loop-helix protein 17; Isoform 5: Attenuates the ability of transcription factor HIF1A to bind to hypoxia-responsive elements (HRE) located within the enha |
| 232 | 2 | HAPLN3 | ENSP00000352606 | Hyaluronan and proteoglycan link protein 3; May function in hyaluronic acid binding; V-set domain containing |
| | | MFGE8 | ENSP00000268150 | Milk fat globule-EGF factor 8 protein; Plays an important role in the maintenance of intestinal epithelial homeostasis and the promotion of mucosal healing. Promotes VEGF- |
| 233 | 2 | EMP3 | ENSP00000270221 | Hematopoietic neural membrane protein 1; Probably involved in cell proliferation and cell-cell interactions. |
| | | PLP2 | ENSP00000365505 | Proteolipid protein 2 (colonic epithelium-enriched); May play a role in cell differentiation in the intestinal epithelium. |
| 234 | 2 | IQCA1 | ENSP00000407213 | IQ motif containing with AAA domain 1 |
| | | STK17B | ENSP00000263955 | DAP kinase-related apoptosis-inducing protein kinase 2; Phosphorylates myosin light chains (By similarity). Acts as a positive regulator of apoptosis; Belongs to the protein ki |
