## Supplementary tables for "Transcriptome profiling of the dorsomedial prefrontal cortex in suicide victims": S5_PPI_network_data_of_down-_and_upregulated_genes2.pdf

Supplementary file 5. : Downregulated DEGs assigned to clusters by STRING using MCL algorithm.

| Gene nam | String ID | Cluster | Cluster col | Protein ID | AverageSh | Clustering | Closeness | Eccentricit | Stress | Degree | Betweenn | Neighborh | Numberof | Numberof | Radiality | TopologicalCoefficient | Description |
|---|---|---|---|---|---|---|---|---|---|---|---|---|---|---|---|---|---|
| A2ML1 | Q96LQ8 | 1 | Red | 9606.ENSP | 3.8134263 | 0.0 | 0.2622313 | 8 | 160 | 2 | 1.7527705 | 17.0 | 0 | 2 | 0.9846260 | 0.5 | C3 and PZP-like alpha-2-macroglobulin domain-containing protein 9; |
| ACACB | O00763 | 1 | Red | 9606.ENSP | 3.3836094 | 0.3333333 | 0.2955423 | 8 | 7418 | 12 | 5.6475638 | 16.666666 | 0 | 12 | 0.9869748 | 0.150150150150150 | acetyl-CoA carboxylase beta; Catalyzes the ATP-dependent carboxyla |
| ACKR3 | P25106 | 1 | Red | 9606.ENSP | 3.0078465 | 0.7 | 0.3324637 | 7 | 340 | 5 | 3.1397567 | 101.4 | 0 | 5 | 0.9890281 | 0.3608540925266904 | G-protein coupled receptor RDC1 homolog; Atypical chemokine rece |
| ACSL5 | Q9ULC5 | 1 | Red | 9606.ENSP | 3.1778552 | 0.2285714 | 0.3146776 | 7 | 20554 | 15 | 0.0019626 | 19.0 | 0 | 15 | 0.9880991 | 0.107575757575757 | acyl-CoA synthetase long-chain family member 5; Acyl-CoA synthetas |
| ADM | P35318 | 1 | Red | 9606.ENSP | 3.0470793 | 0.4181818 | 0.3281831 | 7 | 4038 | 11 | 1.6025662 | 58.090909 | 0 | 11 | 0.9888137 | 0.214814814814814 | Adrenomedullin; AM and PAMP are potent hypotensive and vasodila |
| ANGPT2 | O15123 | 1 | Red | 9606.ENSP | 2.6713164 | 0.4746300 | 0.3743472 | 7 | 53710 | 44 | 0.0019258 | 62.295454 | 0 | 44 | 0.9908671 | 0.1258494031221304 | Angiopoietin 2; Binds to TEK/TIE2, competing for the ANGPT1 bindin |
| APLN | | 1 | Red | 9606.ENSP | 3.1002615 | 0.4642857 | 0.3225534 | 7 | 13848 | 8 | 0.0010084 | 52.0 | 0 | 8 | 0.9885231 | 0.221688034188034 | Apelin; Endogenous ligands |
| AREG | P15514 | 1 | Red | 9606.ENSP | 2.7550130 | 0.3913043 | 0.3629746 | 7 | 18184 | 24 | 6.4591484 | 68.666666 | 0 | 24 | 0.9904097 | 0.153961136023916 | Colorectum cell-derived growth factor; Ligand of the EGF receptor/E |
| AXL | P30530 | 1 | Red | 9606.ENSP | 2.7271142 | 0.2953846 | 0.3666879 | 7 | 30808 | 26 | 0.0012918 | 62.192307 | 0 | 26 | 0.9905622 | 0.133459887751733 | Tyrosine-protein kinase receptor UFO; Receptor tyrosine kinase that |
| BCL6 | P41182 | 1 | Red | 9606.ENSP | 2.6843940 | 0.2807807 | 0.3725235 | 7 | 60116 | 37 | 0.0026155 | 53.378378 | 0 | 37 | 0.9907956 | 0.108880308880308 | Zinc finger and BTB domain-containing protein 27; Transcriptional re |
| CASP4 | Q9UG96 | 1 | Red | 9606.ENSP | 3.1194420 | 0.3611111 | 0.3205701 | 7 | 2786 | 9 | 1.0649268 | 51.777777 | 0 | 9 | 0.9884183 | 0.2203309692671395 | Caspase 4, apoptosis-related cysteine peptidase; Inflammatory caspa |
| CCL2 | P13500 | 1 | Red | 9606.ENSP | 2.4472537 | 0.2909698 | 0.4086213 | 7 | 263238 | 92 | 0.0090796 | 52.304347 | 0 | 92 | 0.9920915 | 0.080592215448516 | Monocyte chemotactic and activating factor; Chemotactic factor that |
| CD163 | Q86VB7 | 1 | Red | 9606.ENSP | 2.7349607 | 0.3492063 | 0.3656359 | 6 | 134924 | 36 | 0.0066330 | 53.888888 | 0 | 36 | 0.9905193 | 0.117404986686032 | Scavenger receptor cysteine-rich type 1 protein M130; Acute phase-r |
| CD248 | Q9HCU0 | 1 | Red | 9606.ENSP | 3.0714908 | 0.3406593 | 0.3255747 | 7 | 4766 | 14 | 1.8701219 | 45.785714 | 0 | 14 | 0.9886803 | 0.1840437788018433 | CD248 molecule, endosialin; May play a role in tumor angiogenesis; ( |
| CD34 | P28906 | 1 | Red | 9606.ENSP | 2.4768962 | 0.3049382 | 0.4037310 | 6 | 211616 | 81 | 0.0081576 | 54.123456 | 0 | 81 | 0.9919295 | 0.086736308958531 | Hematopoietic progenitor cell antigen CD34; Possible adhesion mole |
| CD40 | P25942 | 1 | Red | 9606.ENSP | 2.6303400 | 0.3282485 | 0.3801789 | 7 | 150334 | 60 | 0.0063606 | 53.116666 | 0 | 60 | 0.9910910 | 0.100727387729285 | Tumor necrosis factor receptor superfamily member 5; Receptor for |
| CD44 | Q96J24 | 1 | Red | 9606.ENSP | 2.3356582 | 0.2007864 | 0.4281448 | 6 | 588668 | 132 | 0.0250233 | 46.659090 | 0 | 132 | 0.9927013 | 0.065532431052093 | GP90 lymphocyte homing/adhesion receptor; Receptor for hyaluroni |
| CDKN2B | P42772 | 1 | Red | 9606.ENSP | 2.9372275 | 0.5555555 | 0.3404571 | 7 | 2430 | 10 | 6.9100968 | 72.8 | 0 | 10 | 0.9894140 | 0.2233128834355828 | Cyclin-dependent kinase inhibitor 2B (p15, inhibits CDK4); Interacts s |
| CISH | Q9NSE2 | 1 | Red | 9606.ENSP | 2.8544027 | 0.4782608 | 0.3503359 | 7 | 27752 | 24 | 0.0012622 | 61.0 | 0 | 24 | 0.9898666 | 0.1586371527777778 | Cytokine inducible SH2-containing protein; SOCS family proteins form |
| CLIC1 | O00299 | 1 | Red | 9606.ENSP | 2.9895379 | 0.1969696 | 0.3344998 | 7 | 46220 | 12 | 0.0038830 | 37.5 | 0 | 12 | 0.9891282 | 0.134963768115942 | Regulatory nuclear chloride ion channel protein; Can insert into mem |
| CNTFR | P26992 | 1 | Red | 9606.ENSP | 3.0793374 | 0.4727272 | 0.3247451 | 7 | 5120 | 11 | 2.3813025 | 50.181818 | 0 | 11 | 0.9886375 | 0.197208303507516 | Ciliary neurotrophic factor receptor subunit alpha; Binds to CNTF. Th |
| COL4A1 | P02462 | 1 | Red | 9606.ENSP | 2.7663469 | 0.3357041 | 0.3614875 | 7 | 84690 | 38 | 0.0042398 | 47.263157 | 0 | 38 | 0.9903478 | 0.112674287149855 | Collagen alpha-1(IV) chain; Type IV collagen is the major structural co |
| CP | P00450 | 1 | Red | 9606.ENSP | 2.9145597 | 0.3333333 | 0.3431049 | 7 | 22708 | 22 | 0.0012311 | 34.045454 | 0 | 22 | 0.9895379 | 0.101325757575757 | Ceruloplasmin (ferroxidase); Ceruloplasmin is a blue, copper-binding |
| CTF1 | Q16619 | 1 | Red | 9606.ENSP | 2.9625108 | 0.6373626 | 0.3375515 | 7 | 2456 | 14 | 6.2626298 | 59.357142 | 0 | 14 | 0.9892758 | 0.188435374149659 | Cardiotrophin 1; Induces cardiac myocyte hypertrophy in vitro. Binds |
| CXCL1 | P09341 | 1 | Red | 9606.ENSP | 2.6704446 | 0.4546938 | 0.3744694 | 6 | 62120 | 50 | 0.0018230 | 62.1 | 0 | 50 | 0.9908718 | 0.1242 | Chemokine (C-X-C motif) ligand 1 (melanoma growth stimulating acti |
| CXCL10 | P02778 | 1 | Red | 9606.ENSP | 2.6896251 | 0.2923395 | 0.3717990 | 7 | 286334 | 70 | 0.0120195 | 46.785714 | 0 | 70 | 0.9907670 | 0.097643900984193 | 10 kDa interferon gamma-induced protein; Chemotactic for monocyt |
| CXCL12 | Q9H554 | 1 | Red | 9606.ENSP | 2.4934612 | 0.3646797 | 0.4010489 | 7 | 160148 | 74 | 0.0055862 | 59.405405 | 0 | 74 | 0.9918390 | 0.095660878269573 | Pre-B cell growth-stimulating factor; Chemoattractant active on T-lyn |
| CXCL16 | Q9H2A7 | 1 | Red | 9606.ENSP | 3.1578029 | 1.0 | 0.3166758 | 7 | 0 | 8 | 0.0 | 79.25 | 0 | 8 | 0.9882087 | 0.356981981981982 | Scavenger receptor for phosphatidylserine and oxidized low density l |
| CXCL3 | P19876 | 1 | Red | 9606.ENSP | 3.0087183 | 0.8823529 | 0.3323674 | 7 | 560 | 17 | 1.5087437 | 72.235294 | 0 | 17 | 0.9890233 | 0.244038155802861 | Macrophage inflammatory protein 2-beta; Ligand for CXCR2 (By simil |
| CSF1 | P09603 | 1 | Red | 9606.ENSP | 2.6486486 | 0.4906122 | 0.3775510 | 7 | 52152 | 50 | 0.0014781 | 66.54 | 0 | 50 | 0.9909909 | 0.128208092485549 | Colony stimulating factor 1 (macrophage); Cytokine that plays an ess |
| CSF3 | P09919 | 1 | Red | 9606.ENSP | 2.6765475 | 0.4532428 | 0.3736156 | 6 | 77216 | 52 | 0.0033401 | 61.903846 | 0 | 52 | 0.9908385 | 0.125272500778573 | Colony stimulating factor 3 (granulocyte); Granulocyte/macrophage ( |
| CST3 | P01034 | 1 | Red | 9606.ENSP | 3.0383609 | 0.3083333 | 0.3291248 | 7 | 23512 | 16 | 0.0013536 | 41.625 | 0 | 16 | 0.9888614 | 0.151136363636363 | Neuroendocrine basic polypeptide; As an inhibitor of cysteine protei |
| DDR1 | Q5ST12 | 1 | Red | 9606.ENSP | 3.0217959 | 0.1648351 | 0.3309290 | 7 | 10590 | 14 | 5.9709276 | 37.0 | 0 | 14 | 0.9889519 | 0.135241596638655 | Epithelial discoidin domain-containing receptor 1; Tyrosine kinase th |
| DDR2 | Q16832 | 1 | Red | 9606.ENSP | 2.9075850 | 0.6181818 | 0.3439280 | 7 | 1502 | 11 | 3.3542135 | 82.363636 | 0 | 11 | 0.9895760 | 0.240829346092503 | Discoidin domain-containing receptor tyrosine kinase 2; Tyrosine kin |
| DOK1 | Q99704 | 1 | Red | 9606.ENSP | 2.9511769 | 0.3787878 | 0.3388478 | 6 | 19468 | 12 | 0.0012367 | 53.5 | 0 | 12 | 0.9893378 | 0.177464008859357 | Docking protein 1, 62kDa (downstream of tyrosine kinase 1); DOK pr |
| EFNB1 | P98172 | 1 | Red | 9606.ENSP | 2.7637314 | 0.3007246 | 0.3618296 | 7 | 93422 | 24 | 0.0058894 | 53.583333 | 0 | 24 | 0.9903621 | 0.1256846635367762 | EPH-related receptor tyrosine kinase ligand 2; Binds to the receptor t |
| EGFR | Q9H2C9 | 1 | Red | 9606.ENSP | 2.1778552 | 0.1197381 | 0.4591673 | 6 | 1513486 | 183 | 0.0984402 | 37.229508 | 0 | 183 | 0.9935636 | 0.045985396529480 | Receptor tyrosine-protein kinase erbB-1; Receptor tyrosine kinase bi |
| ENG | P17813 | 1 | Red | 9606.ENSP | 2.6303400 | 0.4285714 | 0.3801789 | 6 | 63034 | 57 | 0.0018804 | 60.280701 | 0 | 57 | 0.9910910 | 0.115006026516673 | Endoglin; Vascular endothelium glycoprotein that plays an important |
| EPHA2 | P29317 | 1 | Red | 9606.ENSP | 2.6765475 | 0.3143669 | 0.3736156 | 7 | 100694 | 38 | 0.0050674 | 58.5 | 0 | 38 | 0.9908385 | 0.118261240144896 | Tyrosine-protein kinase receptor ECK; Receptor tyrosine kinase whic |
| EPHB4 | P54760 | 1 | Red | 9606.ENSP | 2.8081952 | 0.4095238 | 0.3561005 | 7 | 23362 | 21 | 0.0014606 | 61.476190 | 0 | 21 | 0.9901191 | 0.157387057387057 | Tyrosine-protein kinase TYRO11; Receptor tyrosine kinase which bin |
| ERBB2 | P04626 | 1 | Red | 9606.ENSP | 2.3975588 | 0.1886465 | 0.4170909 | 7 | 460210 | 108 | 0.0232899 | 44.638888 | 0 | 108 | 0.9923630 | 0.066983650316983 | V-erb-b2 avian erythroblastic leukemia viral oncogene homolog 2; Pr |
| EVA1B | Q9NVM1 | 1 | Red | 9606.ENSP | 3.6207497 | 0.0 | 0.2761858 | 8 | 0 | 1 | 0.0 | 53.0 | 0 | 1 | 0.9856789 | 0.0 | Eva-1 homolog B; Belongs to the EVA1 family. |
| F11R | Q9Y624 | 1 | Red | 9606.ENSP | 2.8570183 | 0.3666666 | 0.3500152 | 7 | 29404 | 25 | 0.0015862 | 51.88 | 0 | 25 | 0.9898523 | 0.139838274932614 | Junctional adhesion molecule 1; Seems to play a role in epithelial tigh |
| F3 | P13726 | 1 | Red | 9606.ENSP | 2.7707061 | 0.5770114 | 0.3609188 | 7 | 15806 | 30 | 4.4275345 | 70.1 | 0 | 30 | 0.9903240 | 0.163403263403263 | Coagulation factor III (thromboplastin, tissue factor); Initiates blood c |
| FFAR2 | O15552 | 1 | Red | 9606.ENSP | 3.1490845 | 0.6666666 | 0.3175526 | 7 | 150 | 4 | 4.1432240 | 89.0 | 0 | 4 | 0.9882563 | 0.4158878504672897 | G-protein coupled receptor 43; G protein-coupled receptor that is ac |
| FGF11 | Q92914 | 1 | Red | 9606.ENSP | 2.8081952 | 0.6666666 | 0.3561005 | 7 | 3388 | 15 | 1.0908782 | 89.266666 | 0 | 15 | 0.9901191 | 0.219328419328419 | Fibroblast growth factor homologous factor 3; Probably involved in n |
| FGFR3 | P22607 | 1 | Red | 9606.ENSP | 2.7497820 | 0.2906403 | 0.3636651 | 7 | 34482 | 29 | 0.0016235 | 54.931034 | 0 | 29 | 0.9904383 | 0.124560168895144 | Fibroblast growth factor receptor 3; Tyrosine-protein kinase that act |
| FLT1 | P17948 | 1 | Red | 9606.ENSP | 2.6355710 | 0.4230769 | 0.3794244 | 7 | 91722 | 53 | 0.0031406 | 61.641509 | 0 | 53 | 0.9910624 | 0.119460289600702 | Vascular endothelial growth factor receptor 1; Tyrosine-protein kina |
| FLT3LG | P49771 | 1 | Red | 9606.ENSP | 2.8378378 | 0.5019762 | 0.3523809 | 7 | 10882 | 23 | 3.2645315 | 67.956521 | 0 | 23 | 0.9899571 | 0.171607378129117 | Fms-related tyrosine kinase 3 ligand; Stimulates the proliferation of e |
| FLT4 | P35916 | 1 | Red | 9606.ENSP | 2.7244986 | 0.3704994 | 0.3670400 | 7 | 61876 | 42 | 0.0021694 | 55.142857 | 0 | 42 | 0.9905765 | 0.1217281614632608 | Vascular endothelial growth factor receptor 3; Tyrosine-protein kinas |
| FN1 | H0Y4K8 | 1 | Red | 9606.ENSP | 2.2083696 | 0.1465018 | 0.4528227 | 6 | 1220576 | 174 | 0.0696078 | 40.120689 | 0 | 174 | 0.9933968 | 0.0507214787043899 | Fibronectin type III domain containing; Endogenous ligands |
| FOXO1 | Q12778 | 1 | Red | 9606.ENSP | 2.6094158 | 0.2728954 | 0.3832275 | 7 | 112154 | 47 | 0.0049274 | 50.893617 | 0 | 47 | 0.9912053 | 0.094734339712349 | Forkhead in rhabdomyosarcoma; Transcription factor that is the mai |
| FOXO4 | P98177 | 1 | Red | 9606.ENSP | 3.0775937 | 0.3454545 | 0.3249291 | 7 | 12890 | 11 | 8.0733355 | 41.0 | 0 | 11 | 0.9886470 | 0.171886936592818 | Fork head domain transcription factor AFX1; Transcription factor inv |
| FSTL1 | Q12841 | 1 | Red | 9606.ENSP | 2.9285091 | 0.4833333 | 0.3414706 | 7 | 6144 | 16 | 1.4833555 | 61.9375 | 0 | 16 | 0.9894616 | 0.185441616766470 | Follistatin-related protein 1; May modulate the action of some growt |
| FTL | P02792 | 1 | Red | 9606.ENSP | 3.2571926 | 0.4181818 | 0.3070128 | 7 | 6040 | 11 | 4.8514093 | 22.454545 | 0 | 11 | 0.9876656 | 0.146531100478468 | Ferritin, light polypeptide; Stores iron in a soluble, non-toxic, readily |
| GABRE | P78334 | 1 | Red | 9606.ENSP | 3.7558849 | 0.4666666 | 0.2662488 | 7 | 1630 | 6 | 1.8482954 | 8.6666666 | 0 | 6 | 0.9849405 | 0.229729729729729 | Gamma-aminobutyric acid (GABA) A receptor, epsilon; GABA, the ma |
| GABRG1 | Q8N1C3 | 1 | Red | 9606.ENSP | 3.3478639 | 0.1904761 | 0.2986979 | 6 | 15838 | 15 | 0.0014877 | 11.266666 | 0 | 15 | 0.9871701 | 0.1080906148867314 | Gamma-aminobutyric acid (GABA) A receptor, gamma 1; GABA, the r |
| GLRA1 | P23415 | 1 | Red | 9606.ENSP | 3.8884045 | 1.0 | 0.2571748 | 7 | 0 | 4 | 0.0 | 10.0 | 0 | 4 | 0.9842163 | 0.370370370370370 | Glycine receptor strychnine-binding subunit; Glycine receptors are lig |
| GPAM | Q9HCL2 | 1 | Red | 9606.ENSP | 3.4463818 | 0.5 | 0.2901593 | 8 | 2386 | 9 | 1.6743477 | 16.666666 | 0 | 9 | 0.9866317 | 0.185185185185185 | Glycerol-3-phosphate acyltransferase 1, mitochondrial; Esterifies acy |
| GPER1 | Q99527 | 1 | Red | 9606.ENSP | 2.9808195 | 0.5238095 | 0.3354782 | 7 | 1008 | 7 | 3.3568105 | 81.571428 | 0 | 7 | 0.9891758 | 0.2842210054753609 | Flow-induced endothelial G-protein coupled receptor 1; G-protein co |
| HAP1 | P54257 | 1 | Red | 9606.ENSP | 3.6669572 | 0.1666666 | 0.2727056 | 7 | 2036 | 4 | 2.0278322 | 15.75 | 0 | 4 | 0.9854264 | 0.299019607843137 | Huntingtin-associated protein 1; Originally identified as neuronal pro |
| HFE | Q96KU8 | 1 | Red | 9606.ENSP | 3.0148212 | 0.3047619 | 0.3316946 | 7 | 35162 | 15 | 0.0027084 | 30.4 | 0 | 15 | 0.9889900 | 0.107328605200945 | Hereditary hemochromatosis protein; Binds to transferrin receptor ( |
| ICAM1 | P05362 | 1 | Red | 9606.ENSP | 2.4847428 | 0.3194879 | 0.4024561 | 6 | 238208 | 88 | 0.0093992 | 54.579545 | 0 | 88 | 0.9918866 | 0.088013196480938 | Intercellular adhesion molecule 1; ICAM proteins are ligands for the l |
| ICAM2 | P13598 | 1 | Red | 9606.ENSP | 2.9311246 | 0.6 | 0.3411659 | 7 | 7910 | 20 | 2.3975814 | 65.45 | 0 | 20 | 0.9894474 | 0.197734138972809 | Intercellular adhesion molecule 2; ICAM proteins are ligands for the l |
| IL10RA | Q13651 | 1 | Red | 9606.ENSP | 2.9616390 | 0.3709677 | 0.3376508 | 7 | 64402 | 32 | 0.0028582 | 40.46875 | 0 | 32 | 0.9892806 | 0.128373015873015 | Interleukin-10 receptor subunit alpha; Receptor for IL10; binds IL10 v |
| IL13RA1 | P78552 | 1 | Red | 9606.ENSP | 3.1142109 | 0.5384615 | 0.3211086 | 7 | 4960 | 13 | 2.4840359 | 46.0 | 0 | 13 | 0.9884469 | 0.195417348608837 | Interleukin-13 receptor subunit alpha-1; Binds with low affinity to int |
| IL15RA | A0A0A0M | 1 | Red | 9606.ENSP | 3.2197035 | 0.3928571 | 0.3105875 | 8 | 1132 | 8 | 7.0058057 | 42.125 | 0 | 8 | 0.9878704 | 0.214285714285714 | Interleukin 15 receptor subunit alpha; CD molecules |
| IL17D | Q8TAD2 | 1 | Red | 9606.ENSP | 3.0 | 0.3717948 | 0.3333333 | 7 | 3712 | 13 | 1.2463546 | 49.076923 | 0 | 13 | 0.9890710 | 0.168649220195611 | Interleukin 17D; Induces expression of IL6, CXCL8/IL8, and CSF2/GM- |
| IL17RC | Q9BR97 | 1 | Red | 9606.ENSP | 3.1386224 | 0.5277777 | 0.3186111 | 7 | 784 | 9 | 3.1265837 | 57.333333 | 0 | 9 | 0.9883135 | 0.2571001494768311 | Interleukin-17 receptor-like protein; Isoform 8: Receptor for both IL1 |
| IL17RD | Q8NFM7 | 1 | Red | 9606.ENSP | 3.1238012 | 0.3333333 | 0.3201228 | 7 | 1316 | 6 | 4.7053313 | 62.833333 | 0 | 6 | 0.9883945 | 0.2805059523809524 | Interleukin-17 receptor-like protein; Feedback inhibitor of fibroblast |
| IL18R1 | Q13478 | 1 | Red | 9606.ENSP | 3.1142109 | 0.4909090 | 0.3211086 | 7 | 11428 | 11 | 9.0990432 | 47.454545 | 0 | 11 | 0.9884469 | 0.203277409285929 | CD218 antigen-like family member A; Within the IL18 receptor compl |
| IL1R1 | P14778 | 1 | Red | 9606.ENSP | 2.7986050 | 0.5405405 | 0.3573208 | 7 | 27256 | 37 | 7.3214315 | 58.729729 | 0 | 37 | 0.9901715 | 0.139500545676317 | CD121 antigen-like family member A; Receptor for IL1A, IL1B and IL1 |
| IL1R2 | P27930 | 1 | Red | 9606.ENSP | 2.9616390 | 0.5133333 | 0.3376508 | 7 | 26138 | 25 | 0.0011247 | 51.56 | 0 | 25 | 0.9892806 | 0.162649842271293 | CD121 antigen-like family member B; Non-signaling receptor for IL1A |
| IL1RL1 | Q01638 | 1 | Red | 9606.ENSP | 3.1979075 | 0.4 | 0.3127044 | 7 | 1114 | 6 | 4.7692549 | 62.666666 | 0 | 6 | 0.9879895 | 0.322164948453608 | Interleukin 1 receptor-like 1; Receptor for interleukin-33 (IL-33); sign |
| IL32 | F8W1V1 | 1 | Red | 9606.ENSP | 2.9642545 | 0.6428571 | 0.3373529 | 7 | 1726 | 8 | 3.2145437 | 99.375 | 0 | 8 | 0.9892663 | 0.308618012422360 | Interleukin 32; Interleukins |
| IL3RA | P26951 | 1 | Red | 9606.ENSP | 2.9189189 | 0.4047619 | 0.3425925 | 6 | 14168 | 21 | 5.9203193 | 51.285714 | 0 | 21 | 0.9895141 | 0.148654244306418 | Interleukin 3 receptor, alpha (low affinity); This is a receptor for inter |
| IL4R | Q9H185 | 1 | Red | 9606.ENSP | 2.9973844 | 0.4576719 | 0.3336242 | 7 | 6724 | 28 | 3.4037250 | 45.428571 | 0 | 28 | 0.9890853 | 0.155046318868844 | Interleukin-4 receptor subunit alpha; Receptor for both interleukin 4 |
| IL6 | P05231 | 1 | Red | 9606.ENSP | 2.2519616 | 0.1452140 | 0.4440572 | 6 | 1084460 | 170 | 0.0567781 | 39.235294 | 0 | 170 | 0.9931586 | 0.0518222084078017 | B-cell stimulatory factor 2; Cytokine with a wide variety of biological f |
| IL6R | P08887 | 1 | Red | 9606.ENSP | 2.8404533 | 0.4677419 | 0.3520564 | 7 | 11960 | 32 | 4.1620828 | 55.59375 | 0 | 32 | 0.9899428 | 0.141459923664122 | Interleukin-6 receptor subunit alpha; Part of the receptor for interleu |
| IRAK3 | Q9Y616 | 1 | Red | 9606.ENSP | 2.9721011 | 0.5490196 | 0.3364623 | 7 | 6222 | 18 | 2.5712680 | 52.388888 | 0 | 18 | 0.9892234 | 0.165264633718892 | Interleukin-1 receptor-associated kinase 3; Inhibits dissociation of IR |
| ITGA1 | P56199 | 1 | Red | 9606.ENSP | 2.7271142 | 0.3968705 | 0.3666879 | 6 | 90692 | 38 | 0.0045941 | 52.315789 | 0 | 38 | 0.9905622 | 0.117240972386122 | CD49 antigen-like family member A; Integrin alpha-1/beta-1 is a rece |
| ITGA5 | P08648 | 1 | Red | 9606.ENSP | 2.7131647 | 0.4357487 | 0.3685732 | 6 | 63640 | 46 | 0.0020905 | 57.195652 | 0 | 46 | 0.9906384 | 0.124609263995453 | Integrin, alpha 5 (fibronectin receptor, alpha polypeptide); Integrin a |
| ITGA6 | Q16508 | 1 | Red | 9606.ENSP | 2.6355710 | 0.3828282 | 0.3794244 | 6 | 63996 | 45 | 0.0021165 | 56.666666 | 0 | 45 | 0.9910624 | 0.111329404060248 | CD49 antigen-like family member F; Integrin alpha-6/beta-1 is a rece |
| ITGB1 | Q13090 | 1 | Red | 9606.ENSP | 2.4498692 | 0.2409059 | 0.4081850 | 6 | 379230 | 94 | 0.0184966 | 47.414893 | 0 | 94 | 0.9920772 | 0.074618864131345 | Integrin, beta 1 (fibronectin receptor, beta polypeptide, antigen CD2 |
| ITGB4 | P16144 | 1 | Red | 9606.ENSP | 2.7349607 | 0.3863636 | 0.3656359 | 7 | 50578 | 33 | 0.0025167 | 52.575757 | 0 | 33 | 0.9905193 | 0.118612764210958 | Integrin, beta 4; Integrin alpha-6/beta-4 is a receptor for laminin. Pla |
| ITGB5 | P18084 | 1 | Red | 9606.ENSP | 2.8404533 | 0.3770161 | 0.3520564 | 6 | 28742 | 32 | 0.0010787 | 46.21875 | 0 | 32 | 0.9899428 | 0.122596153846153 | Integrin, beta 5; Integrin alpha-V/beta-5 (ITGAV:ITGB5) is a receptor |
| KLF15 | Q9UIH9 | 1 | Red | 9606.ENSP | 3.1089799 | 0.4285714 | 0.3216489 | 8 | 3904 | 8 | 2.0631847 | 45.0 | 0 | 8 | 0.9884755 | 0.200334821428571 | Kidney-enriched krueppel-like factor; Transcriptional regulator that b |
| KRT8 | Q6GMY0 | 1 | Red | 9606.ENSP | 2.7384481 | 0.47 | 0.3651703 | 7 | 42384 | 25 | 0.0017043 | 67.2 | 0 | 25 | 0.9905002 | 0.150246085011185 | Keratin, type II cytoskeletal 8; Together with KRT19, helps to link the |
| LAMB2 | P55268 | 1 | Red | 9606.ENSP | 3.0941586 | 0.725 | 0.3231896 | 7 | 1286 | 16 | 4.6920272 | 45.5 | 0 | 16 | 0.9885565 | 0.190376569037656 | Laminin, beta 2 (laminin S); Binding to cells via a high affinity recepto |
| LAMC3 | Q9Y6N6 | 1 | Red | 9606.ENSP | 3.0880557 | 0.7 | 0.3238283 | 6 | 2368 | 16 | 1.0345543 | 44.9375 | 0 | 16 | 0.9885898 | 0.188550420168067 | Laminin-12 subunit gamma; Binding to cells via a high affinity recept |
| LIF | P15018 | 1 | Red | 9606.ENSP | 2.6931124 | 0.4102564 | 0.3713175 | 7 | 35402 | 39 | 0.0011349 | 62.512820 | 0 | 39 | 0.9907480 | 0.130506932177078 | Differentiation-stimulating factor; LIF has the capacity to induce term |
| LRIG1 | Q96JA1 | 1 | Red | 9606.ENSP | 2.9938971 | 0.3333333 | 0.3340128 | 7 | 9440 | 10 | 4.9610823 | 58.9 | 0 | 10 | 0.9891043 | 0.203114186851211 | Leucine-rich repeats and immunoglobulin-like domains protein 1; Ac |
| LTBR | P36941 | 1 | Red | 9606.ENSP | 3.0645161 | 0.4952380 | 0.3263157 | 7 | 4894 | 15 | 1.4621310 | 50.533333 | 0 | 15 | 0.9887184 | 0.196627756160830 | Lymphotoxin beta receptor (TNFR superfamily, member 3); Receptor |
| MCL1 | Q9UHR9 | 1 | Red | 9606.ENSP | 2.7149084 | 0.4229885 | 0.3683365 | 7 | 38376 | 30 | 0.0014893 | 66.6 | 0 | 30 | 0.9906289 | 0.138970076548364 | Induced myeloid leukemia cell differentiation protein Mcl-1; Involved |
| MLKL | Q8NB16 | 1 | Red | 9606.ENSP | 3.2711421 | 0.5 | 0.3057036 | 7 | 3456 | 8 | 2.0202210 | 34.875 | 0 | 8 | 0.9875893 | 0.228618421052631 | Mixed lineage kinase domain-like protein; Pseudokinase that plays a |
| MMP15 | P51511 | 1 | Red | 9606.ENSP | 2.8561464 | 0.7948717 | 0.3501221 | 7 | 3138 | 13 | 5.7157855 | 102.23076 | 0 | 13 | 0.9898571 | 0.266225961538461 | Matrix metallopeptidase 15 (membrane-inserted); Endopeptidase th |
| MMP2 | P08253 | 1 | Red | 9606.ENSP | 2.4402789 | 0.3143544 | 0.4097892 | 7 | 232636 | 87 | 0.0091146 | 56.195402 | 0 | 87 | 0.9921296 | 0.084998348027196 | Matrix metallopeptidase 2 (gelatinase A, 72kDa gelatinase, 72kDa typ |
| MMP28 | Q9H239 | 1 | Red | 9606.ENSP | 3.5623365 | 0.0 | 0.2807146 | 8 | 0 | 1 | 0.0 | 67.0 | 0 | 1 | 0.9859981 | 0.0 | Matrix metalloproteinase-28; Can degrade casein. Could play a role in |
| MTTP | P55157 | 1 | Red | 9606.ENSP | 3.0488230 | 0.3809523 | 0.3279954 | 7 | 14496 | 7 | 0.0010232 | 49.714285 | 0 | 7 | 0.9888042 | 0.203161592505854 | Microsomal triglyceride transfer protein large subunit; Catalyzes the |
| MYD88 | Q99836 | 1 | Red | 9606.ENSP | 2.6442894 | 0.3095238 | 0.3781734 | 6 | 134142 | 57 | 0.0057472 | 50.824561 | 0 | 57 | 0.9910148 | 0.098846337633968 | Myeloid differentiation primary response protein MyD88; Adapter pr |
| NAMPT | P43490 | 1 | Red | 9606.ENSP | 2.8666085 | 0.4152046 | 0.3488442 | 7 | 69426 | 19 | 0.0042217 | 57.631578 | 0 | 19 | 0.9897999 | 0.154640633842671 | Nicotinamide phosphoribosyltransferase; Catalyzes the condensation |
| NFATC1 | Q15793 | 1 | Red | 9606.ENSP | 2.7802964 | 0.3514492 | 0.3596738 | 7 | 22774 | 24 | 9.0158570 | 59.5 | 0 | 24 | 0.9902716 | 0.1382751937984496 | Nuclear factor of activated T-cells, cytoplasmic, calcineurin-depende |
| NFKB2 | Q00653 | 1 | Red | 9606.ENSP | 2.9023539 | 0.51 | 0.3445479 | 7 | 43888 | 25 | 0.0019774 | 54.72 | 0 | 25 | 0.9896046 | 0.154463276836158 | Nuclear factor of kappa light polypeptide gene enhancer in B-cells 2 ( |
| NKG7 | Q16617 | 1 | Red | 9606.ENSP | 3.4716652 | 0.3 | 0.2880462 | 7 | 326 | 5 | 1.5460018 | 30.0 | 0 | 5 | 0.9864936 | 0.2777777777777778 | Natural killer cell granule protein 7; Belongs to the PMP-22/EMP/MP |
| OSMR | Q99650 | 1 | Red | 9606.ENSP | 2.9764603 | 0.7636363 | 0.3359695 | 7 | 652 | 11 | 2.0180355 | 74.272727 | 0 | 11 | 0.9891996 | 0.243517138599105 | Oncostatin-M-specific receptor subunit beta; Associates with IL31RA |
| PDGFRB | P09619 | 1 | Red | 9606.ENSP | 2.5353095 | 0.3032081 | 0.3944291 | 6 | 181988 | 72 | 0.0084228 | 54.069444 | 0 | 72 | 0.9916103 | 0.094289315493503 | Platelet-derived growth factor receptor, beta polypeptide; Tyrosine- |
| PDK4 | Q16654 | 1 | Red | 9606.ENSP | 3.2414995 | 0.4181818 | 0.3084991 | 8 | 5146 | 11 | 3.1976895 | 25.454545 | 0 | 11 | 0.9877513 | 0.168573148705599 | [Pyruvate dehydrogenase (acetyl-transferring)] kinase isozyme 4, mit |
| PECAM1 | Q9NPB7 | 1 | Red | 9606.ENSP | 2.4158674 | 0.2618908 | 0.4139299 | 6 | 315752 | 102 | 0.0123553 | 49.941176 | 0 | 102 | 0.9922630 | 0.075553973480466 | Platelet/endothelial cell adhesion molecule 1; Cell adhesion molecule |
| PGF | P49763 | 1 | Red | 9606.ENSP | 2.7192676 | 0.5133565 | 0.3677460 | 7 | 43000 | 42 | 0.0017177 | 62.952380 | 0 | 42 | 0.9906050 | 0.135621921182266 | Placental growth factor; Growth factor active in angiogenesis and en |
| PLIN5 | Q00G26 | 1 | Red | 9606.ENSP | 3.5727986 | 0.0 | 0.2798926 | 8 | 152 | 2 | 4.7928627 | 36.0 | 0 | 2 | 0.9859409 | 0.538461538461538 | Lipid storage droplet protein 5; Lipid droplet-associated protein that |
| PLXNB1 | O43157 | 1 | Red | 9606.ENSP | 3.0601569 | 0.3111111 | 0.3267806 | 7 | 9574 | 10 | 3.7404751 | 46.0 | 0 | 10 | 0.9887423 | 0.185483870967741 | Semaphorin receptor SEP; Receptor for SEMA4D. Plays a role in RHO |
| PPARA | Q07869 | 1 | Red | 9606.ENSP | 2.6216216 | 0.1981132 | 0.3814432 | 7 | 331896 | 53 | 0.0196915 | 40.641509 | 0 | 53 | 0.9911386 | 0.076431084106172 | Peroxisome proliferator-activated receptor alpha; Ligand-activated tr |
| RGL3 | Q3MIN7 | 1 | Red | 9606.ENSP | 4.3722755 | 0.0 | 0.2287138 | 8 | 0 | 1 | 0.0 | 16.0 | 0 | 1 | 0.9815722 | 0.0 | Ral guanine nucleotide dissociation stimulator-like 3; Guanine nucleo |
| RIN2 | Q8WYP3 | 1 | Red | 9606.ENSP | 4.0340017 | 0.1666666 | 0.2478928 | 8 | 1184 | 4 | 1.6062417 | 9.0 | 0 | 4 | 0.9834207 | 0.265625 | Ras interaction/interference protein 2; Ras effector protein. May fun |
| ROR1 | Q01973 | 1 | Red | 9606.ENSP | 2.9816913 | 0.4722222 | 0.3353801 | 6 | 2352 | 9 | 7.5582056 | 61.333333 | 0 | 9 | 0.9891710 | 0.210045662100456 | Inactive tyrosine-protein kinase transmembrane receptor ROR1; Has |
| RRAS | P10301 | 1 | Red | 9606.ENSP | 3.7731473 | 0.075 | 0.2964590 | 7 | 64242 | 16 | 0.0034904 | 12.5 | 0 | 16 | 0.9870319 | 0.114485981308411 | Related RAS viral (r-ras) oncogene homolog; Regulates the organizati |
| S100A8 | P05109 | 1 | Red | 9606.ENSP | 2.8674803 | 0.2875816 | 0.3487382 | 7 | 16080 | 18 | 6.2773124 | 60.944444 | 0 | 18 | 0.9897951 | 0.164120994309673 | Migration inhibitory factor-related protein 8; S100A8 is a calcium- an |
| S100A9 | P06702 | 1 | Red | 9606.ENSP | 2.9424585 | 0.2867647 | 0.3398518 | 7 | 47280 | 17 | 0.0028130 | 55.529411 | 0 | 17 | 0.9893854 | 0.167080085046066 | Migration inhibitory factor-related protein 14; S100A9 is a calcium- a |
| S1PR1 | P21453 | 1 | Red | 9606.ENSP | 2.8177855 | 0.4466666 | 0.3548886 | 7 | 20490 | 25 | 8.8740169 | 64.28 | 0 | 25 | 0.9900667 | 0.158226600985221 | Endothelial differentiation G-protein coupled receptor 1; G-protein c |
| SEMA3G | Q9NS98 | 1 | Red | 9606.ENSP | 3.1037489 | 0.3611111 | 0.3221910 | 7 | 7422 | 9 | 2.3936589 | 49.666666 | 0 | 9 | 0.9885040 | 0.218306412139011 | Sema domain, immunoglobulin domain (Ig), short basic domain, secr |
| SERPINA1 | Q86U19 | 1 | Red | 9606.ENSP | 2.8788142 | 0.2820512 | 0.3473652 | 7 | 35616 | 27 | 0.0018401 | 41.111111 | 0 | 27 | 0.9897332 | 0.116252295966701 | Serpin peptidase inhibitor, clade A (alpha-1 antiproteinase, antitrypsi |
| SERPINE1 | P05121 | 1 | Red | 9606.ENSP | 2.5396687 | 0.2776659 | 0.3937521 | 6 | 191306 | 71 | 0.0095010 | 51.366197 | 0 | 71 | 0.9915865 | 0.087655626592318 | Serpin peptidase inhibitor, clade E (nexin, plasminogen activator inhi |
| SIGIRR | Q6IA17 | 1 | Red | 9606.ENSP | 3.0749782 | 0.6727272 | 0.3252055 | 7 | 1988 | 11 | 7.2360096 | 58.090909 | 0 | 11 | 0.9886613 | 0.229608336327703 | Single immunoglobulin and toll-interleukin 1 receptor (TIR) domain; |
| SOCS1 | O15524 | 1 | Red | 9606.ENSP | 2.7846556 | 0.3939393 | 0.3591108 | 7 | 46796 | 33 | 0.0021529 | 54.727272 | 0 | 33 | 0.9902477 | 0.127725856697819 | Suppressor of cytokine signaling 1; SOCS family proteins form part of |
| SOCS3 | O14543 | 1 | Red | 9606.ENSP | 2.7157802 | 0.3626271 | 0.3682182 | 7 | 57168 | 47 | 0.0022011 | 48.765957 | 0 | 47 | 0.9906241 | 0.104873026767330 | Suppressor of cytokine signaling 3; SOCS family proteins form part of |
| STAT3 | P40763 | 1 | Red | 9606.ENSP | 2.3060156 | 0.1633572 | 0.4336483 | 7 | 813768 | 148 | 0.0415164 | 41.756756 | 0 | 148 | 0.9928633 | 0.057437079445332 | Signal transducer and activator of transcription 3 (acute-phase respo |

| | | | | | | | | | | | | | | | | | |
|---|---|---|---|---|---|---|---|---|---|---|---|---|---|---|---|---|---|
| TFRC | P02786 | 1 | Red | 9606.ENSP | 2.5754141 | 0.2293178 | 0.3882870 | 6 | 269922 | 53 | 0.0167685 | 49.415094 | 0 | 53 | 0.9913911 | 0.0861075438769798 | Transferrin receptor protein 1; Cellular uptake of iron occurs via rece |
| TGFB1 | P01137 | 1 | Red | 9606.ENSP | 2.4864864 | 0.3306182 | 0.4021739 | 7 | 205998 | 74 | 0.0091777 | 58.175675 | 0 | 74 | 0.9918771 | 0.0917384261232841 | Transforming growth factor, beta 1; Multifunctional protein that con |
| TGFB2 | P61812 | 1 | Red | 9606.ENSP | 2.6408020 | 0.4202020 | 0.3786728 | 7 | 92890 | 45 | 0.0037952 | 63.888888 | 0 | 45 | 0.9910338 | 0.1228205128205128 | Glioblastoma-derived T-cell suppressor factor; TGF-beta 2 has suppre |
| TGFB3 | P10600 | 1 | Red | 9606.ENSP | 2.7061900 | 0.4534412 | 0.3695231 | 7 | 43616 | 39 | 0.0013948 | 65.897435 | 0 | 39 | 0.9906765 | 0.1378607445553052 | Transforming growth factor, beta 3; Involved in embryogenesis and c |
| TGFBR3 | Q03167 | 1 | Red | 9606.ENSP | 3.1002615 | 0.5054945 | 0.3225534 | 7 | 2736 | 14 | 1.7105906 | 44.785714 | 0 | 14 | 0.9885231 | 0.1905775075987842 | Transforming growth factor beta receptor type 3; Binds to TGF-beta. |
| TGIF1 | Q15583 | 1 | Red | 9606.ENSP | 3.3391455 | 0.3 | 0.2994778 | 7 | 1500 | 5 | 8.7416718 | 29.8 | 0 | 5 | 0.9872177 | 0.2422764227642276 | TGFB-induced factor homeobox 1; Binds to a retinoid X receptor (RX |
| TGIF2 | Q9GZN2 | 1 | Red | 9606.ENSP | 3.4882301 | 0.3333333 | 0.2866783 | 7 | 94 | 3 | 5.9799380 | 28.666666 | 0 | 3 | 0.9864031 | 0.3497942386831276 | TGF-beta-induced transcription factor 2; Transcriptional repressor, w |
| TGM2 | P21980 | 1 | Red | 9606.ENSP | 2.8003487 | 0.3384615 | 0.3570983 | 6 | 62096 | 26 | 0.0032898 | 56.538461 | 0 | 26 | 0.9901620 | 0.1377110694183864 | Protein-glutamine gamma-glutamyltransferase 2; Catalyzes the cross |
| TIE1 | P35590 | 1 | Red | 9606.ENSP | 3.0095902 | 0.6043956 | 0.3322711 | 7 | 13500 | 14 | 6.4318553 | 59.785714 | 0 | 14 | 0.9890186 | 0.2179353493222106 | Tyrosine kinase with immunoglobulin-like and EGF-like domains 1; Tr |
| TIMP1 | P01033 | 1 | Red | 9606.ENSP | 2.5632083 | 0.3871551 | 0.3901360 | 7 | 150072 | 67 | 0.0066844 | 59.313432 | 0 | 67 | 0.9914578 | 0.1029228855721393 | Tissue inhibitor of metalloproteinases 1; Metalloproteinase inhibitor |
| TLR4 | Q5VZI9 | 1 | Red | 9606.ENSP | 2.4228421 | 0.2178932 | 0.4127383 | 6 | 464360 | 99 | 0.0238347 | 45.858585 | 0 | 99 | 0.9922249 | 0.0696938994811335 | Toll-like receptor 4; Cooperates with LY96 and CD14 to mediate the i |
| TNFAIP3 | P21580 | 1 | Red | 9606.ENSP | 2.8901482 | 0.3798319 | 0.3460030 | 7 | 82512 | 35 | 0.0038897 | 44.257142 | 0 | 35 | 0.9896713 | 0.1251315256980979 | Tumor necrosis factor, alpha-induced protein 3; Ubiquitin-editing en |
| TNFRSF10I | Q9UBN6 | 1 | Red | 9606.ENSP | 3.5300784 | 0.3333333 | 0.2832798 | 7 | 858 | 4 | 6.5739329 | 27.25 | 0 | 4 | 0.9861744 | 0.3333333333333333 | Tumor necrosis factor receptor superfamily, member 10d, decoy wit |
| TNFRSF1A | P19438 | 1 | Red | 9606.ENSP | 2.6399302 | 0.3640810 | 0.3787978 | 7 | 153484 | 54 | 0.0066553 | 54.722222 | 0 | 54 | 0.9910386 | 0.1051994301994302 | Tumor necrosis factor receptor superfamily, member 1A; Receptor f |
| TNFRSF1B | P20333 | 1 | Red | 9606.ENSP | 2.9093286 | 0.4729064 | 0.3437219 | 7 | 13646 | 29 | 5.1909478 | 52.103448 | 0 | 29 | 0.9895665 | 0.1488669950738916 | Tumor necrosis factor receptor superfamily, member 1B; Receptor w |
| TNFSF13 | O75888 | 1 | Red | 9606.ENSP | 3.6024411 | 0.3333333 | 0.2775895 | 8 | 410 | 3 | 1.9750751 | 24.333333 | 0 | 3 | 0.9857790 | 0.3636363636363636 | Tumor necrosis factor (ligand) superfamily, member 13; Cytokine tha |
| TNFSF14 | O43557 | 1 | Red | 9606.ENSP | 3.3766346 | 0.4642857 | 0.2961528 | 8 | 1456 | 8 | 8.9679523 | 29.0 | 0 | 8 | 0.9870129 | 0.2377049180327868 | Tumor necrosis factor (ligand) superfamily, member 14; Cytokine tha |
| TXNIP | Q9H3M7 | 1 | Red | 9606.ENSP | 3.0453356 | 0.2727272 | 0.3283710 | 7 | 9898 | 11 | 4.5802254 | 44.454545 | 0 | 11 | 0.9888233 | 0.1664388243335611 | Vitamin D3 up-regulated protein 1; May act as an oxidative stress me |
| TTYH1 | Q9H313 | 1 | Red | 9606.ENSP | 3.7837837 | 0.2857142 | 0.2642857 | 7 | 1360 | 7 | 1.8546698 | 8.7142857 | 0 | 7 | 0.9847880 | 0.2124542124542124 | Protein tweety homolog 1; Probable chloride channel. May be involv |
| VCAM1 | P19320 | 1 | Red | 9606.ENSP | 2.5152571 | 0.3710045 | 0.3975736 | 6 | 143918 | 73 | 0.0041053 | 59.054794 | 0 | 73 | 0.9917199 | 0.0980976653165248 | Vascular cell adhesion molecule 1; Important in cell-cell recognition. |
| VEGFB | P49765 | 1 | Red | 9606.ENSP | 2.9171752 | 0.5684210 | 0.3427973 | 7 | 7670 | 20 | 3.2527236 | 58.5 | 0 | 20 | 0.9895236 | 0.1756756756756756 | Vascular endothelial growth factor B; Growth factor for endothelial c |
| VIM | Q96ML2 | 1 | Red | 9606.ENSP | 2.5771578 | 0.3171262 | 0.3880243 | 7 | 166386 | 53 | 0.0082987 | 56.301886 | 0 | 53 | 0.9913816 | 0.1003262368412201 | Vimentin; Vimentins are class-III intermediate filaments found in vari |
| YES1 | P07947 | 1 | Red | 9606.ENSP | 2.8177855 | 0.3260869 | 0.3548886 | 6 | 73682 | 24 | 0.0037022 | 53.25 | 0 | 24 | 0.9900667 | 0.1364316239316239 | YES proto-oncogene 1, Src family tyrosine kinase; Non-receptor prot |
| ZC3H12A | Q5D1E8 | 1 | Red | 9606.ENSP | 3.1002615 | 0.3333333 | 0.3225534 | 7 | 8224 | 13 | 3.7957880 | 41.769230 | 0 | 13 | 0.9885231 | 0.1759169100941252 | Monocyte chemotactic protein-induced protein 1; Endoribonuclease |
| ZFHX3 | Q15911 | 1 | Red | 9606.ENSP | 3.1455972 | 0.6 | 0.3179046 | 8 | 238 | 6 | 1.0481640 | 57.5 | 0 | 6 | 0.9882754 | 0.2625570776255708 | Alpha-fetoprotein enhancer-binding protein; Transcriptional regulat |
| ZNF703 | Q9H7S9 | 1 | Red | 9606.ENSP | 3.3234524 | 0.3333333 | 0.3008919 | 8 | 1366 | 4 | 8.2001997 | 40.75 | 0 | 4 | 0.9873035 | 0.3022388059701492 | Zinc finger elbow-related proline domain protein 1; Transcriptional c |
| ASCL1 | P50553 | 2 | Blue | 9606.ENSP | 2.8369659 | 0.3840579 | 0.3524892 | 7 | 24912 | 24 | 0.0010598 | 43.083333 | 0 | 24 | 0.9899619 | 0.1183608058608058 | Achaete-scute family bHLH transcription factor 1; Transcription facto |
| BCL6B | Q8N143 | 2 | Blue | 9606.ENSP | 3.2319093 | 0.2 | 0.3094146 | 7 | 6960 | 5 | 4.4936631 | 39.8 | 0 | 5 | 0.9878037 | 0.2387878787878788 | B-cell CLL/lymphoma 6 member B protein; Acts as a sequence-specifi |
| BMP7 | P18075 | 2 | Blue | 9606.ENSP | 2.7863993 | 0.3399014 | 0.3588861 | 7 | 35560 | 29 | 0.0013932 | 51.586206 | 0 | 29 | 0.9902382 | 0.1246043644844244 | Bone morphogenetic protein 7; Induces cartilage and bone formatio |
| CERS1 | P27539 | 2 | Blue | 9606.ENSP | 3.9834350 | 1.0 | 0.2510396 | 8 | 0 | 2 | 0.0 | 6.5 | 0 | 2 | 0.9836970 | 0.5909090909090909 | Embryonic growth/differentiation factor 1; May mediate cell differen |
| DLL4 | Q9NR61 | 2 | Blue | 9606.ENSP | 2.7741935 | 0.3217189 | 0.3604651 | 7 | 119470 | 42 | 0.0048357 | 47.738095 | 0 | 42 | 0.9903049 | 0.1136054421768707 | Drosophila Delta homolog 4; Involved in the Notch signaling pathway |
| EBF1 | Q9UH73 | 2 | Blue | 9606.ENSP | 3.0549258 | 0.2417582 | 0.3273401 | 6 | 8596 | 14 | 5.1676790 | 34.857142 | 0 | 14 | 0.9887708 | 0.1397016637980493 | Transcription factor COE1; Transcriptional activator which recognizes |
| EMX2 | Q04743 | 2 | Blue | 9606.ENSP | 3.2990409 | 0.3888888 | 0.3031183 | 7 | 7950 | 9 | 4.6256764 | 31.888888 | 0 | 9 | 0.9874369 | 0.2184170471841704 | Empty spiracles-like protein 2; Transcription factor, which in coopera |
| ENHO | Q6UWT2 | 2 | Blue | 9606.ENSP | 3.3487358 | 0.0 | 0.2986201 | 7 | 0 | 1 | 0.0 | 135.0 | 0 | 1 | 0.9871653 | 0.0 | Energy homeostasis-associated protein; Involved in the regulation of |
| EVC | P57679 | 2 | Blue | 9606.ENSP | 3.5675675 | 0.5 | 0.2803030 | 7 | 154 | 4 | 9.9836763 | 25.0 | 0 | 4 | 0.9859695 | 0.375 | Ellis-van Creveld syndrome protein; Component of the EvC complex t |
| EYA1 | Q99502 | 2 | Blue | 9606.ENSP | 2.9459459 | 0.4705882 | 0.3394495 | 7 | 14310 | 17 | 6.4110110 | 47.647058 | 0 | 17 | 0.9893664 | 0.1532059769245318 | EYA transcriptional coactivator and phosphatase 1; Functions both as |
| FLI1 | Q01543 | 2 | Blue | 9606.ENSP | 3.1150828 | 0.3055555 | 0.3210187 | 7 | 3192 | 9 | 1.5760115 | 45.888888 | 0 | 9 | 0.9884421 | 0.1999029597282872 | Friend leukemia integration 1 transcription factor; Sequence-specific |
| FOXC1 | Q12948 | 2 | Blue | 9606.ENSP | 2.8631211 | 0.4526315 | 0.3492691 | 7 | 19160 | 20 | 7.0142102 | 55.45 | 0 | 20 | 0.9898190 | 0.1531767955801105 | Forkhead-related transcription factor 3; DNA-binding transcriptional |
| FOXC2 | Q99958 | 2 | Blue | 9606.ENSP | 2.8953792 | 0.3478260 | 0.3453778 | 7 | 22874 | 24 | 9.0860134 | 49.25 | 0 | 24 | 0.9896427 | 0.1444281524926686 | Forkhead box C2 (MFH-1, mesenchyme forkhead 1); Transcriptional |
| FOXD1 | Q16676 | 2 | Blue | 9606.ENSP | 3.3775065 | 0.4285714 | 0.2960764 | 7 | 1266 | 7 | 5.2898537 | 29.714285 | 0 | 7 | 0.9870081 | 0.2506053268765133 | Forkhead-related transcription factor 4; Transcription factor involved |
| GDF1 | P27539 | 2 | Blue | 9606.ENSP | 3.9834350 | 1.0 | 0.2510396 | 8 | 0 | 2 | 0.0 | 6.5 | 0 | 2 | 0.9836970 | 0.5909090909090909 | Growth differentiation factor 1; May mediate cell differentiation eve |
| GFAP | Q7Z5J7 | 2 | Blue | 9606.ENSP | 2.4856146 | 0.2053571 | 0.4023149 | 6 | 500702 | 64 | 0.0312171 | 46.796875 | 0 | 64 | 0.9918818 | 0.0741135103011093 | Glial fibrillary acidic protein; GFAP, a class-III intermediate filament, i |
| GLI2 | P10070 | 2 | Blue | 9606.ENSP | 2.7637314 | 0.2761904 | 0.3618296 | 6 | 82800 | 36 | 0.0038908 | 41.611111 | 0 | 36 | 0.9903621 | 0.0988756613756613 | GLI family zinc finger protein 2; Functions as transcription regulator i |
| GLI3 | P10071 | 2 | Blue | 9606.ENSP | 2.8090671 | 0.2807881 | 0.3559900 | 6 | 68654 | 29 | 0.0034319 | 39.793103 | 0 | 29 | 0.9901143 | 0.1014250527797325 | Transcriptional activator GLI3; Has a dual function as a transcriptiona |
| GRIN2C | Q14957 | 2 | Blue | 9606.ENSP | 3.3469921 | 0.4 | 0.2987757 | 6 | 1556 | 5 | 1.0784133 | 31.4 | 0 | 5 | 0.9871749 | 0.2915887850467289 | Glutamate receptor, ionotropic, N-methyl D-aspartate 2C; Compone |
| HEPN1 | Q6WQI6 | 2 | Blue | 9606.ENSP | 3.4420226 | 0.8 | 0.2905268 | 6 | 402 | 5 | 5.1124950 | 22.4 | 0 | 5 | 0.9866556 | 0.2731707317073171 | Putative cancer susceptibility gene HEPN1 protein; Hepatocellular ca |
| HES1 | Q14469 | 2 | Blue | 9606.ENSP | 2.9938971 | 0.4444444 | 0.3340128 | 7 | 12982 | 18 | 6.5733535 | 41.333333 | 0 | 18 | 0.9891043 | 0.1443278943278943 | Class B basic helix-loop-helix protein 39; Transcriptional repressor of |
| HES6 | Q96HZ4 | 2 | Blue | 9606.ENSP | 3.2423714 | 0.8888888 | 0.3084162 | 7 | 144 | 9 | 3.8280640 | 43.888888 | 0 | 9 | 0.9877466 | 0.2612433862433862 | Class B basic helix-loop-helix protein 41; Does not bind DNA itself but |
| HEYL | Q9NQ87 | 2 | Blue | 9606.ENSP | 3.2266782 | 0.7333333 | 0.3099162 | 7 | 1494 | 10 | 8.5289400 | 40.6 | 0 | 10 | 0.9878323 | 0.2314285714285714 | Hes-related family bHLH transcription factor with YRPW motif-like; D |
| ID4 | P47928 | 2 | Blue | 9606.ENSP | 3.1360069 | 0.6111111 | 0.3188768 | 7 | 1662 | 9 | 9.8462386 | 48.333333 | 0 | 9 | 0.9883278 | 0.2290679304897314 | Inhibitor of DNA binding 4, dominant negative helix-loop-helix protei |
| JAG1 | P78504 | 2 | Blue | 9606.ENSP | 2.5876198 | 0.3079249 | 0.3864555 | 6 | 230642 | 58 | 0.0105025 | 53.810344 | 0 | 58 | 0.9913244 | 0.0978056426332288 | Protein jagged-1; Ligand for multiple Notch receptors and involved in |
| KLF2 | Q9Y5W3 | 2 | Blue | 9606.ENSP | 2.7776809 | 0.3760683 | 0.3600125 | 7 | 19150 | 27 | 5.9753992 | 60.296296 | 0 | 27 | 0.9902858 | 0.1415406016344983 | Lung krueppel-like factor; Transcription factor that binds to the CACC |
| LFNG | Q8NES3 | 2 | Blue | 9606.ENSP | 3.1909328 | 0.3676470 | 0.3133879 | 7 | 41994 | 17 | 0.0017152 | 28.117647 | 0 | 17 | 0.9880276 | 0.1487706193588546 | LFNG O-fucosylpeptide 3-beta-N-acetylglucosaminyltransferase; Glyc |
| MAML2 | Q8IZL2 | 2 | Blue | 9606.ENSP | 3.1272885 | 0.5128205 | 0.3197658 | 7 | 3590 | 13 | 1.8249067 | 37.846153 | 0 | 13 | 0.9883754 | 0.1802197802197802 | Mastermind-like 2 (Drosophila); Acts as a transcriptional coactivator |
| MBP | Q15338 | 2 | Blue | 9606.ENSP | 2.8125544 | 0.3162055 | 0.3555486 | 7 | 52964 | 23 | 0.0026211 | 50.521739 | 0 | 23 | 0.9900953 | 0.1259893743901116 | Myelin membrane encephalitogenic protein; The classic group of MB |
| MFNG | O00587 | 2 | Blue | 9606.ENSP | 3.2188317 | 0.5454545 | 0.3106717 | 7 | 7484 | 12 | 3.4459250 | 34.25 | 0 | 12 | 0.9878752 | 0.1913407821229050 | MFNG O-fucosylpeptide 3-beta-N-acetylglucosaminyltransferase; Gly |
| MOBP | C9JAR7 | 2 | Blue | 9606.ENSP | 3.4158674 | 0.4761904 | 0.2927514 | 7 | 612 | 7 | 6.5943406 | 21.428571 | 0 | 7 | 0.9867985 | 0.2491694352159468 | Myelin-associated oligodendrocyte basic protein |
| MSX1 | P28360 | 2 | Blue | 9606.ENSP | 2.9006102 | 0.2919540 | 0.3447550 | 7 | 50368 | 30 | 0.0028537 | 37.0 | 0 | 30 | 0.9896141 | 0.1103482587064676 | Msh homeobox 1-like protein; Acts as a transcriptional repressor. Ma |
| NES | P48681 | 2 | Blue | 9606.ENSP | 2.6111595 | 0.3968547 | 0.3829716 | 7 | 88680 | 47 | 0.0027201 | 61.872340 | 0 | 47 | 0.9911958 | 0.1156492344402465 | Nestin; Required for brain and eye development. Promotes the disas |
| NKX3-1 | Q99801 | 2 | Blue | 9606.ENSP | 3.1822144 | 0.2777777 | 0.3142465 | 7 | 4184 | 9 | 2.6394128 | 34.111111 | 0 | 9 | 0.9880753 | 0.1855921855921856 | Homeobox protein NK-3 homolog A; Transcription factor, which bind |
| NOTCH1 | P46531 | 2 | Blue | 9606.ENSP | 2.3496076 | 0.1614151 | 0.4256029 | 6 | 812800 | 135 | 0.0424525 | 41.103703 | 0 | 135 | 0.9926250 | 0.0597222222222222 | Translocation-associated notch protein TAN-1; Functions as a recept |
| NOTCH2 | Q04721 | 2 | Blue | 9606.ENSP | 2.7401918 | 0.3351158 | 0.3649379 | 7 | 57754 | 34 | 0.0023751 | 48.264705 | 0 | 34 | 0.9904907 | 0.1106988667026443 | Neurogenic locus notch homolog protein 2; Functions as a receptor f |
| NOTCH3 | Q9UM47 | 2 | Blue | 9606.ENSP | 2.6817785 | 0.3573573 | 0.3728868 | 7 | 66738 | 37 | 0.0022594 | 57.243243 | 0 | 37 | 0.9908099 | 0.1185160315595098 | Neurogenic locus notch homolog protein 3; Functions as a receptor f |
| NOTCH4 | Q9UIJ0 | 2 | Blue | 9606.ENSP | 2.7925021 | 0.3655172 | 0.3581017 | 7 | 60596 | 30 | 0.0027967 | 50.9 | 0 | 30 | 0.9902049 | 0.1222756410256410 | Neurogenic locus notch homolog protein 4; Functions as a receptor f |
| NTRK2 | Q16620 | 2 | Blue | 9606.ENSP | 2.7933740 | 0.2093596 | 0.3579900 | 7 | 127392 | 29 | 0.0079890 | 43.344827 | 0 | 29 | 0.9902001 | 0.1071275776503807 | Neurotrophic tyrosine kinase, receptor, type 2; Receptor tyrosine kin |
| NTSR2 | O95665 | 2 | Blue | 9606.ENSP | 3.3653007 | 0.3454545 | 0.2971502 | 6 | 3884 | 11 | 4.5657093 | 16.818181 | 0 | 11 | 0.9870748 | 0.1501623376623376 | Levocabastine-sensitive neurotensin receptor; Receptor for the tride |
| OLIG1 | Q8TAK6 | 2 | Blue | 9606.ENSP | 3.0296425 | 0.4416666 | 0.3300719 | 7 | 34730 | 16 | 0.0018934 | 38.5625 | 0 | 16 | 0.9889090 | 0.1469465648854961 | Class E basic helix-loop-helix protein 21; Promotes formation and ma |
| PPP1R1B | Q9UD71 | 2 | Blue | 9606.ENSP | 2.9529206 | 0.4 | 0.3386477 | 7 | 6426 | 10 | 3.1657396 | 57.1 | 0 | 10 | 0.9893283 | 0.1812698412698412 | Protein phosphatase 1, regulatory (inhibitor) subunit 1B; Inhibitor of |
| RHOBTB3 | O94955 | 2 | Blue | 9606.ENSP | 3.7602441 | 0.0 | 0.2659401 | 8 | 12806 | 5 | 8.2874416 | 8.4 | 0 | 5 | 0.9849166 | 0.2466666666666666 | Rho-related BTB domain-containing protein 3; Rab9-regulated ATPas |
| S100B | P04271 | 2 | Blue | 9606.ENSP | 2.7279860 | 0.2873563 | 0.3665707 | 7 | 97380 | 30 | 0.0052097 | 47.833333 | 0 | 30 | 0.9905574 | 0.1057522123893805 | S100 calcium binding protein B; Weakly binds calcium but binds zinc |
| SIX1 | Q15475 | 2 | Blue | 9606.ENSP | 2.9625108 | 0.3473684 | 0.3375515 | 7 | 19994 | 20 | 0.0010113 | 38.95 | 0 | 20 | 0.9892758 | 0.1347750865051903 | Sine oculis homeobox homolog 1; Transcription factor that is involve |
| SLC39A12 | Q6NZY5 | 2 | Blue | 9606.ENSP | 3.3897122 | 0.4 | 0.2950102 | 6 | 10318 | 6 | 9.0418244 | 19.833333 | 0 | 6 | 0.9869414 | 0.2166666666666666 | Solute carrier family 39 (zinc transporter), member 12; Acts as a zinc- |
| SMO | Q99835 | 2 | Blue | 9606.ENSP | 2.9851787 | 0.2909090 | 0.3349883 | 7 | 41260 | 11 | 0.0038847 | 44.545454 | 0 | 11 | 0.9891520 | 0.1527647610121837 | Smoothened, frizzled class receptor; G protein-coupled receptor tha |
| SOX17 | Q9H6I2 | 2 | Blue | 9606.ENSP | 2.8666085 | 0.2984615 | 0.3488442 | 7 | 26902 | 26 | 0.0010372 | 44.576923 | 0 | 26 | 0.9897999 | 0.1228014409832591 | SRY (sex determining region Y)-box 17; Acts as transcription regulato |
| SOX2 | P48431 | 2 | Blue | 9606.ENSP | 2.4507410 | 0.1698518 | 0.4080398 | 6 | 566574 | 92 | 0.0315273 | 40.673913 | 0 | 92 | 0.9920724 | 0.0645100069013112 | SRY (sex determining region Y)-box 2; Transcription factor that forms |
| SOX9 | P48436 | 2 | Blue | 9606.ENSP | 2.5597210 | 0.2265734 | 0.3906675 | 6 | 237292 | 66 | 0.0109813 | 46.242424 | 0 | 66 | 0.9914769 | 0.0824285637119861 | SRY (sex determining region Y)-box 9; Transcriptional regulator. Bind |
| TBX2 | Q13207 | 2 | Blue | 9606.ENSP | 2.9468177 | 0.3516483 | 0.3393491 | 7 | 10730 | 14 | 4.3044559 | 49.857142 | 0 | 14 | 0.9893616 | 0.1624011167985109 | T-box transcription factor TBX2; Involved in the transcriptional regula |
| TBX3 | O15119 | 2 | Blue | 9606.ENSP | 2.9032258 | 0.3976608 | 0.3444444 | 7 | 18646 | 19 | 8.7664227 | 48.473684 | 0 | 19 | 0.9895998 | 0.1449732114717932 | T-box transcription factor TBX3; Transcriptional repressor involved in |
| TP53INP2 | Q8IXH6 | 2 | Blue | 9606.ENSP | 3.7925021 | 0.0 | 0.2636781 | 8 | 0 | 1 | 0.0 | 29.0 | 0 | 1 | 0.9847404 | 0.0 | Tumor protein p53 inducible nuclear protein 2; Dual regulator of tra |
| ZEB2 | O60315 | 2 | Blue | 9606.ENSP | 2.7410636 | 0.3916256 | 0.3648218 | 7 | 43234 | 29 | 0.0019770 | 61.655172 | 0 | 29 | 0.9904859 | 0.1378538918460232 | Zinc finger E-box binding homeobox 2; Transcriptional inhibitor that |
| ZIC5 | Q96T25 | 2 | Blue | 9606.ENSP | 3.3156059 | 0.3333333 | 0.3016039 | 7 | 7834 | 6 | 7.0393159 | 32.5 | 0 | 6 | 0.9873464 | 0.2330917874396135 | Zinc finger protein of the cerebellum 5; Essential for neural crest dev |
| BST2 | Q10589 | 3 | Lime greer | 9606.ENSP | 2.9755884 | 0.3841269 | 0.3360679 | 7 | 69244 | 36 | 0.0039542 | 30.555555 | 0 | 36 | 0.9892044 | 0.1071150097465886 | Bone marrow stromal cell antigen 2; IFN-induced antiviral host restri |
| CLEC2B | Q92478 | 3 | Lime greer | 9606.ENSP | 3.6713164 | 0.0 | 0.2723818 | 7 | 598 | 3 | 2.4361006 | 20.666666 | 0 | 3 | 0.9854026 | 0.3575757575757576 | C-type lectin domain family 2 member B |
| DDX60 | Q8IY21 | 3 | Lime greer | 9606.ENSP | 3.2711421 | 0.5886699 | 0.3057036 | 8 | 10724 | 29 | 4.2546420 | 27.241379 | 0 | 29 | 0.9875893 | 0.1746242263483642 | DEAD (Asp-Glu-Ala-Asp) box polypeptide 60; Positively regulates DD |
| EPSTI1 | Q96J88 | 3 | Lime greer | 9606.ENSP | 3.4978204 | 0.8545454 | 0.2858923 | 8 | 1102 | 11 | 3.0495056 | 30.545454 | 0 | 11 | 0.9863507 | 0.2909090909090909 | Epithelial stromal interaction 1 |
| FAM89A | Q96GI7 | 3 | Lime greer | 9606.ENSP | 4.2981691 | 0.0 | 0.2326572 | 9 | 0 | 1 | 0.0 | 28.0 | 0 | 1 | 0.9819772 | 0.0 | Family with sequence similarity 89 member A; Belongs to the FAM89 |
| GBP1 | P32455 | 3 | Lime greer | 9606.ENSP | 3.0470793 | 0.5100806 | 0.3281831 | 7 | 48186 | 32 | 0.0027318 | 30.4375 | 0 | 32 | 0.9888137 | 0.1183122568093385 | Guanylate binding protein 1, interferon-inducible; Hydrolyzes GTP to |
| GBP2 | P32456 | 3 | Lime greer | 9606.ENSP | 3.2048823 | 0.5470085 | 0.3120239 | 7 | 21924 | 27 | 0.0011306 | 30.185185 | 0 | 27 | 0.9879514 | 0.1558242180003838 | Guanylate binding protein 2, interferon-inducible; Hydrolyzes GTP to |
| GLIS2 | Q9BZE0 | 3 | Lime greer | 9606.ENSP | 3.6329555 | 0.0952380 | 0.2752579 | 8 | 26680 | 7 | 0.0023681 | 9.8571428 | 0 | 7 | 0.9856122 | 0.1629072681704260 | Neuronal Krueppel-like protein; Can act either as a transcriptional re |
| HERC5 | Q9UII4 | 3 | Lime greer | 9606.ENSP | 3.3818657 | 0.7359307 | 0.2956947 | 8 | 22864 | 22 | 0.0013526 | 28.863636 | 0 | 22 | 0.9869843 | 0.2233967582804792 | HECT and RLD domain containing E3 ubiquitin protein ligase 5; Major |
| HLA-B | Q29681 | 3 | Lime greer | 9606.ENSP | 3.2162162 | 0.3732193 | 0.3109243 | 8 | 64034 | 27 | 0.0032383 | 24.518518 | 0 | 27 | 0.9878895 | 0.1375364128173117 | HLA class I histocompatibility antigen, B-7 alpha chain; Involved in the |
| HLA-E | P13747 | 3 | Lime greer | 9606.ENSP | 3.3234524 | 0.4033333 | 0.3008919 | 7 | 40874 | 25 | 0.0021843 | 23.28 | 0 | 25 | 0.9873035 | 0.1589041095890411 | HLA class I histocompatibility antigen, alpha chain E; Preferably binds |
| IFI27 | P40305 | 3 | Lime greer | 9606.ENSP | 3.3156059 | 0.7107692 | 0.3016039 | 8 | 7620 | 26 | 2.1948996 | 29.384615 | 0 | 26 | 0.9873464 | 0.1946001018848701 | Interferon alpha-inducible protein 27, mitochondrial; Promotes cell c |
| IFI30 | P13284 | 3 | Lime greer | 9606.ENSP | 3.6024411 | 0.6222222 | 0.2775895 | 8 | 6630 | 10 | 5.2470668 | 28.0 | 0 | 10 | 0.9857790 | 0.3088888888888889 | Gamma-interferon-inducible lysosomal thiol reductase; Lysosomal th |
| IFI35 | P80217 | 3 | Lime greer | 9606.ENSP | 3.1839581 | 0.5454545 | 0.3140744 | 8 | 23252 | 33 | 0.0013294 | 27.727272 | 0 | 33 | 0.9880658 | 0.1420357420357420 | Interferon-induced 35 kDa protein; Not yet known. |
| IFI44L | Q53G44 | 3 | Lime greer | 9606.ENSP | 3.2990409 | 0.6164021 | 0.3031183 | 8 | 38372 | 28 | 0.0021436 | 27.25 | 0 | 28 | 0.9874369 | 0.1802270577105014 | Interferon-induced protein 44-like; Exhibits a low antiviral activity ag |
| IFITM1 | P13164 | 3 | Lime greer | 9606.ENSP | 3.0993897 | 0.5763546 | 0.3226441 | 7 | 45986 | 29 | 0.0024094 | 30.758620 | 0 | 29 | 0.9885279 | 0.1347549909255898 | Interferon induced transmembrane protein 1; IFN-induced antiviral |
| IFITM2 | Q01629 | 3 | Lime greer | 9606.ENSP | 3.4106364 | 0.7777777 | 0.2932004 | 8 | 9752 | 19 | 5.8472416 | 30.263157 | 0 | 19 | 0.9868271 | 0.2360197368421052 | Interferon induced transmembrane protein 2; IFN-induced antiviral |
| IFITM3 | Q01628 | 3 | Lime greer | 9606.ENSP | 3.0845684 | 0.5584045 | 0.3241944 | 8 | 42520 | 27 | 0.0020913 | 33.148148 | 0 | 27 | 0.9886089 | 0.1286928952298602 | Interferon induced transmembrane protein 3; IFN-induced antiviral |
| IRF1 | P10914 | 3 | Lime greer | 9606.ENSP | 2.8343504 | 0.3502040 | 0.3528145 | 7 | 204944 | 50 | 0.0071471 | 42.72 | 0 | 50 | 0.9899762 | 0.1114882506527415 | Interferon regulatory factor 1; Transcriptional regulator which displa |
| IRF7 | Q92985 | 3 | Lime greer | 9606.ENSP | 2.7986050 | 0.2659968 | 0.3573208 | 7 | 256208 | 62 | 0.0116615 | 33.693548 | 0 | 62 | 0.9901715 | 0.0856931790199458 | Interferon regulatory factor 7; Key transcriptional regulator of type I |
| ISG20 | Q96AZ6 | 3 | Lime greer | 9606.ENSP | 3.2911944 | 0.6707692 | 0.3038410 | 8 | 9286 | 26 | 3.4141267 | 29.615384 | 0 | 26 | 0.9874798 | 0.1850961538461538 | Promyelocytic leukemia nuclear body-associated protein ISG20; Inte |
| LILRB1 | Q8NHL6 | 3 | Lime greer | 9606.ENSP | 3.3251961 | 0.2142857 | 0.3007341 | 7 | 8182 | 8 | 5.4144717 | 23.125 | 0 | 8 | 0.9872940 | 0.1811023622047244 | Leukocyte immunoglobulin-like receptor, subfamily B (with TM and I |
| MAVS | Q86VY7 | 3 | Lime greer | 9606.ENSP | 3.0226678 | 0.4069264 | 0.3308335 | 7 | 48072 | 22 | 0.0018241 | 41.863636 | 0 | 22 | 0.9889471 | 0.1504251144538914 | Interferon beta promoter stimulator protein 1; Required for innate i |
| MICA | Q9UE06 | 3 | Lime greer | 9606.ENSP | 3.3400174 | 0.2380952 | 0.2993996 | 7 | 8034 | 7 | 5.6224844 | 25.428571 | 0 | 7 | 0.9872130 | 0.1901181525241675 | MHC class I polypeptide-related sequence A; Seems to have no role i |
| MR1 | Q9TQB3 | 3 | Lime greer | 9606.ENSP | 4.1700087 | 1.0 | 0.2398076 | 8 | 0 | 2 | 0.0 | 26.0 | 0 | 2 | 0.9826775 | 0.8387096774193549 | Major histocompatibility complex class I-related gene protein; Antige |
| MX2 | P20592 | 3 | Lime greer | 9606.ENSP | 3.1665213 | 0.5042016 | 0.3158039 | 7 | 26990 | 35 | 0.0017506 | 27.485714 | 0 | 35 | 0.9881610 | 0.1379755922469490 | Interferon-regulated resistance GTP-binding protein MxB; Interferon |
| NLRC5 | Q86WI3 | 3 | Lime greer | 9606.ENSP | 3.6172624 | 0.5833333 | 0.2764521 | 8 | 58248 | 9 | 0.0016452 | 27.444444 | 0 | 9 | 0.9856980 | 0.2939068100358423 | Nucleotide-binding oligomerization domain protein 27; Probable reg |
| NWD1 | Q149M9 | 3 | Lime greer | 9606.ENSP | 4.5414123 | 0.0 | 0.2201958 | 9 | 934 | 2 | 6.2294746 | 6.5 | 0 | 2 | 0.9806480 | 0.5 | NACHT domain- and WD repeat-containing protein 1; May play a role |
| OAS3 | Q9Y6K5 | 3 | Lime greer | 9606.ENSP | 3.2458587 | 0.6279569 | 0.3080848 | 8 | 11964 | 31 | 3.8649306 | 27.903225 | 0 | 31 | 0.9877275 | 0.1691104594330400 | 2'-5'-oligoadenylate synthetase 3, 100kDa; Interferon-induced, dsRN |
| PARP12 | Q9H0J9 | 3 | Lime greer | 9606.ENSP | 3.4184829 | 0.7426470 | 0.2925274 | 8 | 3712 | 17 | 1.2282108 | 27.058823 | 0 | 17 | 0.9867842 | 0.2528862012094557 | ADP-ribosyltransferase diphtheria toxin-like 12; poly(ADP-ribose) pol |
| PARP14 | Q460N5 | 3 | Lime greer | 9606.ENSP | 3.2589363 | 0.5281385 | 0.3068485 | 8 | 23144 | 22 | 0.0014967 | 28.181818 | 0 | 22 | 0.9876560 | 0.1702479338842975 | ADP-ribosyltransferase diphtheria toxin-like 8; ADP-ribosyltransfera |
| PARP9 | Q8IXQ6 | 3 | Lime greer | 9606.ENSP | 3.1743679 | 0.5138461 | 0.3150233 | 7 | 28410 | 26 | 0.0016181 | 27.5 | 0 | 26 | 0.9881182 | 0.1415543219666931 | ADP-ribosyltransferase diphtheria toxin-like 9; ADP-ribosyltransfera |
| PSMB8 | P28062 | 3 | Lime greer | 9606.ENSP | 3.0932868 | 0.4928774 | 0.3232807 | 7 | 42192 | 27 | 0.0015993 | 32.074074 | 0 | 27 | 0.9885612 | 0.1298545509071824 | Proteasome (prosome, macropain) subunit, beta type, 8; The protea |
| RNF213 | Q8IZX2 | 3 | Lime greer | 9606.ENSP | 3.4812554 | 0.6794871 | 0.2872526 | 8 | 8538 | 13 | 5.6862953 | 27.307692 | 0 | 13 | 0.9864412 | 0.2778649921507064 | ALK lymphoma oligomerization partner on chromosome 17; E3 ubiqu |
| SAMD9L | Q8IWI4 | 3 | Lime greer | 9606.ENSP | 3.1918047 | 0.6536796 | 0.3133023 | 7 | 15672 | 22 | 0.0010144 | 29.590909 | 0 | 22 | 0.9880229 | 0.1605731225296442 | Sterile alpha motif domain-containing protein 9-like; May be involve |
| SP110 | Q9HB58 | 3 | Lime greer | 9606.ENSP | 3.4551002 | 0.6373626 | 0.2894272 | 8 | 6598 | 14 | 4.3554108 | 27.5 | 0 | 14 | 0.9865841 | 0.2493506493506493 | Transcriptional coactivator Sp110; Transcription factor. May be a nu |
| TAP1 | Q03518 | 3 | Lime greer | 9606.ENSP | 3.6469049 | 0.6666666 | 0.2742051 | 8 | 3388 | 7 | 2.4003303 | 24.142857 | 0 | 7 | 0.9855360 | 0.3037974683544304 | Transporter 1, ATP-binding cassette, sub-family B (MDR/TAP); Involv |
| TRIM22 | Q8IYM9 | 3 | Lime greer | 9606.ENSP | 3.4428945 | 0.7 | 0.2904532 | 8 | 8456 | 20 | 6.6280400 | 28.25 | 0 | 20 | 0.9866508 | 0.2522321428571428 | RING-type E3 ubiquitin transferase TRIM22; Interferon-induced antiv |
| XAF1 | Q6GPH4 | 3 | Lime greer | 9606.ENSP | 3.2650392 | 0.6043010 | 0.3062750 | 8 | 12266 | 31 | 5.1555332 | 27.645161 | 0 | 31 | 0.9876227 | 0.1663427905169063 | XIAP associated factor 1; Seems to function as a negative regulator o |
| COX4I2 | Q96KJ9 | 4 | Pink | 9606.ENSP | 3.6460331 | 0.8974358 | 0.2742706 | 8 | 704 | 13 | 2.5262583 | 16.153846 | 0 | 13 | 0.9855408 | 0.3296703296703296 | Cytochrome c oxidase subunit 4 isoform 2, mitochondrial; This protei |
| MT-ATP6 | Q8WCX6 | 4 | Pink | 9606.ENSP | 3.6190061 | 0.8571428 | 0.2763189 | 7 | 804 | 14 | 6.9022131 | 16.0 | 0 | 14 | 0.9856884 | 0.3018867924528302 | Mitochondrially encoded ATP synthase 6; Mitochondrial membrane |
| MT-ATP8 | P03928 | 4 | Pink | 9606.ENSP | 3.6198779 | 0.8589743 | 0.2762524 | 7 | 716 | 13 | 5.9693595 | 16.307692 | 0 | 13 | 0.9856837 | 0.3076923076923077 | Mitochondrially encoded ATP synthase 8; Mitochondrial membrane |
| MT-CO1 | P00395 | 4 | Pink | 9606.ENSP | 3.2432432 | 0.4 | 0.3083333 | 8 | 35946 | 21 | 0.0031340 | 17.333333 | 0 | 21 | 0.9877418 | 0.1190476190476190 | Mitochondrially encoded cytochrome c oxidase I; Cytochrome c oxid |
| MT-CO2 | P00403 | 4 | Pink | 9606.ENSP | 2.9555361 | 0.4523809 | 0.3383480 | 7 | 73748 | 21 | 0.0066078 | 28.0 | 0 | 21 | 0.9893140 | 0.0933333333333333 | Mitochondrially encoded cytochrome c oxidase II; Cytochrome c oxid |
| MT-CO3 | P00414 | 4 | Pink | 9606.ENSP | 3.5823888 | 0.7166666 | 0.2791433 | 8 | 4328 | 16 | 2.8667599 | 15.1875 | 0 | 16 | 0.9858885 | 0.2410714285714285 | Mitochondrially encoded cytochrome c oxidase III; Subunits I, II and I |
| MT-CYB | Q9B2Z0 | 4 | Pink | 9606.ENSP | 3.5405405 | 0.4853801 | 0.2824427 | 8 | 16742 | 19 | 0.0015530 | 13.157894 | 0 | 19 | 0.9861172 | 0.1904024767801857 | Ubiquinol-cytochrome-c reductase complex cytochrome b subunit; C |
| MT-ND1 | P03886 | 4 | Pink | 9606.ENSP | 2.8709677 | 0.3514492 | 0.3483146 | 7 | 120408 | 24 | 0.0086644 | 38.666666 | 0 | 24 | 0.9897761 | 0.1081932773109243 | Mitochondrially encoded NADH dehydrogenase 1; Core subunit of th |
| MT-ND2 | P03891 | 4 | Pink | 9606.ENSP | 3.3478639 | 0.6176470 | 0.2986979 | 8 | 15018 | 17 | 9.8950448 | 17.882352 | 0 | 17 | 0.9871701 | 0.1510468594217348 | Mitochondrially encoded NADH dehydrogenase 2; Core subunit of th |

| | | | | | | | | | | | | | | | | | |
|---|---|---|---|---|---|---|---|---|---|---|---|---|---|---|---|---|---|
| MT-ND3 | P03897 | 4 | Pink | 9606.ENSP | 3.5945945 | 0.8901098 | 0.2781954 | 8 | 1060 | 14 | 5.7960227 | 16.714285 | 0 | 14 | 0.9858218 | 0.2832929782082324 | Mitochondrially encoded NADH dehydrogenase 3; Core subunit of th |
| MT-ND4 | P03905 | 4 | Pink | 9606.ENSP | 3.5309503 | 0.7619047 | 0.2832098 | 8 | 3768 | 15 | 2.4419512 | 16.2 | 0 | 15 | 0.9861696 | 0.2347826086956521 | Mitochondrially encoded NADH dehydrogenase 4; Core subunit of th |
| MT-ND4L | P03901 | 4 | Pink | 9606.ENSP | 3.5945945 | 0.8901098 | 0.2781954 | 8 | 1060 | 14 | 5.7960227 | 16.714285 | 0 | 14 | 0.9858218 | 0.2832929782082324 | Mitochondrially encoded NADH dehydrogenase 4L; Core subunit of t |
| MT-ND5 | P03915 | 4 | Pink | 9606.ENSP | 3.5309503 | 0.7619047 | 0.2832098 | 8 | 3768 | 15 | 2.4419512 | 16.2 | 0 | 15 | 0.9861696 | 0.2347826086956521 | Mitochondrially encoded NADH dehydrogenase 5; Core subunit of th |
| MT-ND6 | P03923 | 4 | Pink | 9606.ENSP | 3.6530078 | 0.9871794 | 0.2737470 | 8 | 2 | 13 | 1.2679458 | 16.615384 | 0 | 13 | 0.9855026 | 0.353518821603928 | Mitochondrially encoded NADH dehydrogenase 6; Core subunit of th |
| NDUFA4L2 | Q9NRX3 | 4 | Pink | 9606.ENSP | 3.6965998 | 0.4761904 | 0.2705188 | 8 | 6924 | 7 | 5.0952225 | 15.0 | 0 | 7 | 0.9852644 | 0.326984126984127 | NADH dehydrogenase [ubiquinone] 1 alpha subcomplex subunit 4-lik |
| AKAP9 | Q9UQQ4 | 5 | Soft orang | 9606.ENSP | 2.9433304 | 0.1045751 | 0.3397511 | 7 | 106350 | 18 | 0.0101886 | 26.055555 | 0 | 18 | 0.9893807 | 0.0873822975517890 | Centrosome- and Golgi-localized PKN-associated protein; Scaffolding |
| BAZ1A | Q9NRL2 | 5 | Soft orang | 9606.ENSP | 3.6033129 | 0.5333333 | 0.2775223 | 8 | 14862 | 6 | 0.0012148 | 16.166666 | 0 | 6 | 0.9857742 | 0.2424242424242424 | Williams syndrome transcription factor-related chromatin-remodelin |
| BAZ2B | Q9UIF8 | 5 | Soft orang | 9606.ENSP | 3.7506538 | 0.3333333 | 0.2666201 | 8 | 400 | 3 | 4.1737416 | 20.333333 | 0 | 3 | 0.9849691 | 0.4 | Bromodomain adjacent to zinc finger domain protein 2B; May play a |
| BPTF | Q12830 | 5 | Soft orang | 9606.ENSP | 3.2127288 | 0.0676923 | 0.3112618 | 8 | 112340 | 26 | 0.0103468 | 9.9615384 | 0 | 26 | 0.9879085 | 0.0608568646543330 | Bromodomain and PHD finger-containing transcription factor; Histon |
| CNTLN | Q9NXG0 | 5 | Soft orang | 9606.ENSP | 3.3931996 | 0.0714285 | 0.2947070 | 8 | 55502 | 8 | 0.0050788 | 13.875 | 0 | 8 | 0.9869224 | 0.1394736842105263 | Centlein, centrosomal protein; Required for centrosome cohesion an |
| KCNE4 | Q8WWG9 | 5 | Soft orang | 9606.ENSP | 3.8735832 | 0.3333333 | 0.2581589 | 8 | 552 | 3 | 8.9180827 | 10.0 | 0 | 3 | 0.9842973 | 0.3717948717948718 | Potassium voltage-gated channel, Isk-related family, member 4; Anci |
| LRRCC1 | Q9C099 | 5 | Soft orang | 9606.ENSP | 4.3923278 | 0.0 | 0.2276697 | 9 | 0 | 1 | 0.0 | 8.0 | 0 | 1 | 0.9814626 | 0.0 | Centrosomal leucine-rich repeat and coiled-coil domain-containing p |
| NEDD1 | Q8NHV4 | 5 | Soft orang | 9606.ENSP | 3.8561464 | 0.3333333 | 0.2593262 | 8 | 732 | 3 | 9.0383665 | 10.0 | 0 | 3 | 0.9843926 | 0.3717948717948718 | Neural precursor cell expressed developmentally down-regulated pro |
| PHEX | P78562 | 5 | Soft orang | 9606.ENSP | 3.6643417 | 0.5 | 0.2729003 | 7 | 138 | 4 | 1.5371578 | 13.0 | 0 | 4 | 0.9854407 | 0.3109756097560975 | Phosphate regulating endopeptidase homolog, X-linked; Probably inv |
| PRRC2C | Q9Y520 | 5 | Soft orang | 9606.ENSP | 3.7602441 | 0.0 | 0.2659401 | 8 | 946 | 2 | 9.1456067 | 22.0 | 0 | 2 | 0.9849166 | 0.5121951219512195 | Proline-rich and coiled-coil-containing protein 2C; Proline rich coiled- |
| PTH1R | Q03431 | 5 | Soft orang | 9606.ENSP | 3.0941586 | 0.1208791 | 0.3231896 | 6 | 19966 | 14 | 0.0016229 | 22.428571 | 0 | 14 | 0.9885565 | 0.1044474393530997 | Parathyroid hormone/parathyroid hormone-related peptide recepto |
| RAB34 | Q9BZG1 | 5 | Soft orang | 9606.ENSP | 3.5196163 | 0.0 | 0.2841218 | 7 | 2636 | 4 | 3.0406413 | 20.75 | 0 | 4 | 0.9862316 | 0.2564935064935065 | RAB34, member RAS oncogene family; Protein transport. Involved in |
| RBBP6 | Q9H3I8 | 5 | Soft orang | 9606.ENSP | 3.6904969 | 0.0 | 0.2709662 | 7 | 1652 | 3 | 1.7863732 | 19.333333 | 0 | 3 | 0.9852978 | 0.3333333333333333 | P53-associated cellular protein of testis; E3 ubiquitin-protein ligase w |
| CASC3 | O15234 | 6 | Soft blue | 9606.ENSP | 4.2606800 | 1.0 | 0.2347043 | 8 | 0 | 3 | 0.0 | 10.333333 | 0 | 3 | 0.9821820 | 0.5740740740740741 | Cancer susceptibility candidate gene 3 protein; Core component of th |
| GRAP | Q13588 | 6 | Soft blue | 9606.ENSP | 3.7776809 | 0.1666666 | 0.2647126 | 8 | 4316 | 4 | 2.4419324 | 13.0 | 0 | 4 | 0.9848214 | 0.3048780487804878 | GRB2-related adapter protein; Couples signals from receptor and cyt |
| LUC7L3 | O95232 | 6 | Soft blue | 9606.ENSP | 3.6965998 | 0.2909090 | 0.2705188 | 8 | 25148 | 11 | 0.0023397 | 9.6363636 | 0 | 11 | 0.9852644 | 0.1486291486291486 | Cisplatin resistance-associated-overexpressed protein; Binds cAMP r |
| NKTR | P30414 | 6 | Soft blue | 9606.ENSP | 3.3993025 | 0.1666666 | 0.2941779 | 8 | 20122 | 9 | 0.0020569 | 13.333333 | 0 | 9 | 0.9868890 | 0.1400966183574879 | Natural-killer cells cyclophilin-related protein; Component of a putati |
| PNISR | Q8TF01 | 6 | Soft blue | 9606.ENSP | 4.1246730 | 0.6666666 | 0.2424434 | 8 | 958 | 7 | 8.0510316 | 9.7142857 | 0 | 7 | 0.9829252 | 0.3349753694581280 | Serine/arginine-rich-splicing regulatory protein 130; PNN interacting |
| PRPF38B | Q5VTL8 | 6 | Soft blue | 9606.ENSP | 4.3836094 | 1.0 | 0.2281225 | 8 | 0 | 3 | 0.0 | 12.0 | 0 | 3 | 0.9815103 | 0.5714285714285714 | pre-mRNA processing factor 38B; May be required for pre-mRNA spl |
| SLTM | Q9NWH9 | 6 | Soft blue | 9606.ENSP | 3.8221447 | 0.5714285 | 0.2616332 | 8 | 5482 | 7 | 3.5268012 | 10.857142 | 0 | 7 | 0.9845784 | 0.2613240418118467 | Modulator of estrogen-induced transcription; When overexpressed, |
| SREK1 | Q8WXA9 | 6 | Soft blue | 9606.ENSP | 3.9843068 | 0.2888888 | 0.2509846 | 7 | 6026 | 10 | 6.9983756 | 6.9 | 0 | 10 | 0.9836923 | 0.20625 | Serine/arginine-rich-splicing regulatory protein 86; Participates in the |
| SRRM1 | Q8IYB3 | 6 | Soft blue | 9606.ENSP | 3.9616390 | 0.4166666 | 0.2524207 | 8 | 6150 | 9 | 6.6713442 | 10.0 | 0 | 9 | 0.9838161 | 0.2040816326530612 | SR-related nuclear matrix protein of 160 kDa; Part of pre- and post-sp |
| SRSF11 | Q05519 | 6 | Soft blue | 9606.ENSP | 3.5884917 | 0.2285714 | 0.2786686 | 8 | 46594 | 15 | 0.0034528 | 7.8666666 | 0 | 15 | 0.9858552 | 0.1208333333333333 | Splicing factor, arginine/serine-rich 11; May function in pre-mRNA sp |
| THOC2 | Q8NI27 | 6 | Soft blue | 9606.ENSP | 3.3661726 | 0.2857142 | 0.2970732 | 7 | 32574 | 7 | 0.0020688 | 21.571428 | 0 | 7 | 0.9870700 | 0.1834975369458128 | THO complex subunit 2; Required for efficient export of polyadenylat |
| ZFC3H1 | O60293 | 6 | Soft blue | 9606.ENSP | 3.8770706 | 0.2857142 | 0.2579266 | 9 | 6098 | 7 | 0.0010037 | 11.428571 | 0 | 7 | 0.9842783 | 0.2025974025974026 | Zinc finger C3H1 domain-containing protein; Subunit of the trimeric p |
| ALDH4A1 | Q9UDI6 | 7 | Dark violet | 9606.ENSP | 3.6477768 | 0.3333333 | 0.2741395 | 7 | 1570 | 6 | 1.2406388 | 14.166666 | 0 | 6 | 0.9855312 | 0.2146464646464646 | Delta-1-pyrroline-5-carboxylate dehydrogenase, mitochondrial; Irrev |
| GLUL | P15104 | 7 | Dark violet | 9606.ENSP | 2.8849171 | 0.2391304 | 0.3466304 | 7 | 89498 | 24 | 0.0058837 | 29.916666 | 0 | 24 | 0.9896999 | 0.0895708582834331 | Glutamate--ammonia ligase; This enzyme has 2 functions: it catalyzes |
| SLC1A3 | P43003 | 7 | Dark violet | 9606.ENSP | 2.9049694 | 0.1747899 | 0.3442376 | 7 | 99008 | 35 | 0.0073564 | 23.371428 | 0 | 35 | 0.9895903 | 0.0718694885361552 | Solute carrier family 1 (glial high affinity glutamate transporter), men |
| SLC1A7 | A0A087WU | 7 | Dark violet | 9606.ENSP | 2.9694856 | 0.1857142 | 0.3367586 | 7 | 61732 | 21 | 0.0038434 | 27.190476 | 0 | 21 | 0.9892377 | 0.0937602627257799 | Solute carrier family 1 member 7 |
| SLC38A3 | Q99624 | 7 | Dark violet | 9606.ENSP | 3.4899738 | 0.4666666 | 0.2865350 | 8 | 2442 | 10 | 2.3668474 | 15.0 | 0 | 10 | 0.9863935 | 0.18625 | Sodium-coupled neutral amino acid transporter 3; Sodium-depender |
| SLC38A5 | Q8WUX1 | 7 | Dark violet | 9606.ENSP | 3.4411508 | 0.5277777 | 0.2906004 | 8 | 13266 | 9 | 7.3581719 | 18.555555 | 0 | 9 | 0.9866603 | 0.2049382716049382 | Sodium-coupled neutral amino acid transporter 5; Functions as a sod |
| SLC6A11 | P48066 | 7 | Dark violet | 9606.ENSP | 3.3618134 | 0.3571428 | 0.2974585 | 7 | 14856 | 8 | 8.2219499 | 21.5 | 0 | 8 | 0.9870939 | 0.2063106796116504 | Solute carrier family 6 (neurotransmitter transporter), member 11; T |
| SLC6A12 | P48065 | 7 | Dark violet | 9606.ENSP | 3.6033129 | 0.5 | 0.2775223 | 8 | 696 | 5 | 4.3206155 | 18.4 | 0 | 5 | 0.9857742 | 0.3033333333333333 | Solute carrier family 6 (neurotransmitter transporter), member 12; T |
| SLC6A13 | Q9NSD5 | 7 | Dark violet | 9606.ENSP | 3.6913687 | 0.5 | 0.2709022 | 8 | 544 | 5 | 7.0179163 | 16.0 | 0 | 5 | 0.9852930 | 0.2925925925925925 | Solute carrier family 6 (neurotransmitter transporter), member 13; S |
| SLC7A11 | Q9UPY5 | 7 | Dark violet | 9606.ENSP | 2.9520488 | 0.2190476 | 0.3387477 | 7 | 31304 | 15 | 0.0017901 | 39.266666 | 0 | 15 | 0.9893330 | 0.1283060109289617 | Solute carrier family 7 (anionic amino acid transporter light chain, xc- |
| SLC7A5 | Q01650 | 7 | Dark violet | 9606.ENSP | 3.3714036 | 0.4363636 | 0.2966123 | 8 | 6370 | 11 | 4.0744971 | 17.909090 | 0 | 11 | 0.9870415 | 0.1722027972027972 | Solute carrier family 7 (amino acid transporter light chain, L system), |
| CYSLTR2 | Q9NS75 | 8 | Soft green | 9606.ENSP | 3.8849171 | 0.8666666 | 0.2574057 | 8 | 106 | 6 | 6.2305745 | 15.833333 | 0 | 6 | 0.9842354 | 0.4279279279279279 | G-protein coupled receptor GPCR21; Receptor for cysteinyl leukotrie |
| F2RL3 | Q96RI0 | 8 | Soft green | 9606.ENSP | 3.6556233 | 0.4761904 | 0.2735511 | 8 | 1152 | 7 | 8.5566924 | 17.285714 | 0 | 7 | 0.9854883 | 0.2741935483870967 | Coagulation factor II (thrombin) receptor-like 3; Receptor for activate |
| GNG11 | P61952 | 8 | Soft green | 9606.ENSP | 3.4333042 | 0.2807017 | 0.2912646 | 8 | 18868 | 19 | 0.0010378 | 12.052631 | 0 | 19 | 0.9867032 | 0.1137040714995034 | Guanine nucleotide-binding protein G(I)/G(S)/G(O) subunit gamma-1 |
| GNG12 | Q9UBI6 | 8 | Soft green | 9606.ENSP | 3.0313862 | 0.2380952 | 0.3298820 | 7 | 74988 | 21 | 0.0050793 | 19.714285 | 0 | 21 | 0.9888995 | 0.0768229166666666 | Guanine nucleotide-binding protein G(I)/G(S)/G(O) subunit gamma-1 |
| GNG5 | P63218 | 8 | Soft green | 9606.ENSP | 3.4088927 | 0.2526315 | 0.2933503 | 8 | 21082 | 20 | 0.0013690 | 11.6 | 0 | 20 | 0.9868366 | 0.1069444444444444 | Guanine nucleotide-binding protein G(I)/G(S)/G(O) subunit gamma-5 |
| KCNJ16 | Q9NPI9 | 8 | Soft green | 9606.ENSP | 3.9468177 | 0.5 | 0.2533686 | 8 | 928 | 4 | 9.9653395 | 17.25 | 0 | 4 | 0.9838971 | 0.5666666666666667 | Potassium inwardly-rectifying channel, subfamily J, member 16; Inwa |
| PRKCH | P24723 | 8 | Soft green | 9606.ENSP | 3.2938099 | 0.1428571 | 0.3035997 | 7 | 8062 | 7 | 4.7757043 | 25.285714 | 0 | 7 | 0.9874655 | 0.2009291521486643 | Protein kinase C eta type; Calcium-independent, phospholipid- and d |
| PTGER1 | P34995 | 8 | Soft green | 9606.ENSP | 3.3548387 | 0.5 | 0.2980769 | 8 | 9592 | 8 | 3.7042523 | 28.0 | 0 | 8 | 0.9871320 | 0.2276422764227642 | Prostaglandin E receptor 1 (subtype EP1), 42kDa; Receptor for prosta |
| RGR | P47804 | 8 | Soft green | 9606.ENSP | 3.7680906 | 0.5 | 0.2653863 | 8 | 1110 | 4 | 6.8683875 | 18.5 | 0 | 4 | 0.9848738 | 0.5214285714285715 | RPE-retinal G protein-coupled receptor; Receptor for all-trans- and 1 |
| RGS20 | O76081 | 8 | Soft green | 9606.ENSP | 3.6198779 | 0.4 | 0.2762524 | 8 | 1698 | 6 | 1.1622772 | 17.166666 | 0 | 6 | 0.9856837 | 0.2953216374269006 | Regulator of Gz-selective protein signaling 1; Inhibits signal transduct |
| BTG2 | P78543 | 9 | Soft red | 9606.ENSP | 3.2720139 | 0.4393939 | 0.3056221 | 8 | 9556 | 12 | 6.3756142 | 24.083333 | 0 | 12 | 0.9875846 | 0.1621621621621621 | NGF-inducible anti-proliferative protein PC3; Anti-proliferative protei |
| CEBPD | P49716 | 9 | Soft red | 9606.ENSP | 2.8910200 | 0.4912280 | 0.3458986 | 7 | 8430 | 19 | 3.1293461 | 52.315789 | 0 | 19 | 0.9896665 | 0.1486244019138756 | CCAAT/enhancer binding protein (C/EBP), delta; Transcription activat |
| DUSP1 | P28562 | 9 | Soft red | 9606.ENSP | 2.7663469 | 0.2850574 | 0.3614875 | 7 | 73990 | 30 | 0.0038209 | 48.266666 | 0 | 30 | 0.9903478 | 0.1120930232558139 | Mitogen-activated protein kinase phosphatase 1; Dual specificity pho |
| DUSP5 | Q16690 | 9 | Soft red | 9606.ENSP | 3.0967741 | 0.2777777 | 0.3229166 | 7 | 3782 | 9 | 1.6044021 | 44.222222 | 0 | 9 | 0.9885422 | 0.1946902654867256 | Dual specificity protein phosphatase hVH3; Dual specificity protein ph |
| FOS | P01100 | 9 | Soft red | 9606.ENSP | 2.4350479 | 0.1880877 | 0.4106695 | 7 | 460482 | 88 | 0.0252069 | 44.068181 | 0 | 88 | 0.9921582 | 0.0658377496942519 | FBJ murine osteosarcoma viral oncogene homolog; Nuclear phospho |
| FOSL2 | P15408 | 9 | Soft red | 9606.ENSP | 3.0976460 | 0.5111111 | 0.3228257 | 8 | 1204 | 10 | 5.3085147 | 45.0 | 0 | 10 | 0.9885374 | 0.1931330472103004 | Fos-related antigen 2; Controls osteoclast survival and size. As a dime |
| GADD45B | O75293 | 9 | Soft red | 9606.ENSP | 3.0244115 | 0.3666666 | 0.3306428 | 7 | 9874 | 16 | 5.5521693 | 39.9375 | 0 | 16 | 0.9889376 | 0.1396929824561403 | Growth arrest and DNA damage-inducible protein GADD45 beta; Invo |
| JUNB | P17275 | 9 | Soft red | 9606.ENSP | 2.8979947 | 0.4047619 | 0.3450661 | 7 | 17392 | 21 | 7.3964542 | 45.857142 | 0 | 21 | 0.9896284 | 0.1339459760512392 | Transcription factor jun-B; Transcription factor involved in regulating |
| MAFF | Q9ULX9 | 9 | Soft red | 9606.ENSP | 3.2519616 | 0.4666666 | 0.3075067 | 8 | 1042 | 6 | 5.0014635 | 41.5 | 0 | 6 | 0.9876941 | 0.266025641025641 | V-maf avian musculoaponeurotic fibrosarcoma oncogene homolog F; |
| ZFP36 | P26651 | 9 | Soft red | 9606.ENSP | 3.0183086 | 0.4190476 | 0.3313113 | 7 | 11298 | 15 | 6.1030964 | 40.333333 | 0 | 15 | 0.9889709 | 0.1446043165467626 | Growth factor-inducible nuclear protein NUP475; Zinc-finger RNA-bin |
| ATP10A | O60312 | 10 | | 9606.ENSP | 4.1647776 | 0.0 | 0.2401088 | 8 | 0 | 1 | 0.0 | 20.0 | 0 | 1 | 0.9827061 | 0.0 | P4-ATPase flippase complex alpha subunit ATP10A; Catalytic compon |
| CDH23 | Q9H251 | 10 | | 9606.ENSP | 3.1656495 | 0.1052631 | 0.3158909 | 7 | 80836 | 20 | 0.0068620 | 13.4 | 0 | 20 | 0.9881658 | 0.0705405405405405 | Cadherin-related 23; Cadherins are calcium-dependent cell adhesion |
| CIB2 | O75838 | 10 | | 9606.ENSP | 3.6913687 | 0.3333333 | 0.2709022 | 7 | 2772 | 6 | 2.1536233 | 11.166666 | 0 | 6 | 0.9852930 | 0.2340425531914893 | Calcium and integrin binding family member 2; Calcium-binding prote |
| EHD4 | Q9H223 | 10 | | 9606.ENSP | 3.6050566 | 0.0 | 0.2773881 | 8 | 698 | 3 | 4.0790799 | 21.666666 | 0 | 3 | 0.9857647 | 0.3563218390804598 | Hepatocellular carcinoma-associated protein 10/11; ATP- and membr |
| GJB6 | O95452 | 10 | | 9606.ENSP | 3.1508282 | 0.3076923 | 0.3173768 | 7 | 14900 | 14 | 0.0012086 | 24.714285 | 0 | 14 | 0.9882468 | 0.1328725038402457 | Gap junction protein, beta 6, 30kDa; One gap junction consists of a cl |
| LRTOMT | Q8WZ04 | 10 | | 9606.ENSP | 3.8334786 | 0.2857142 | 0.2608596 | 8 | 2470 | 7 | 3.0713482 | 9.8571428 | 0 | 7 | 0.9845165 | 0.2292358803986711 | Leucine rich transmembrane and O-methyltransferase domain conta |
| TMPRSS3 | P57727 | 10 | | 9606.ENSP | 3.4751525 | 0.4 | 0.2877571 | 7 | 4070 | 6 | 2.9518324 | 16.833333 | 0 | 6 | 0.9864745 | 0.2136752136752136 | Tumor-associated differentially-expressed gene 12 protein; Probable |
| TMPRSS4 | Q9NRS4 | 10 | | 9606.ENSP | 3.4777680 | 0.4 | 0.2875407 | 7 | 4420 | 6 | 2.6816980 | 17.0 | 0 | 6 | 0.9864602 | 0.2130801687763713 | Transmembrane protease, serine 4; Probable protease. Seems to be |
| TRIOBP | Q9H2D6 | 10 | | 9606.ENSP | 3.6163905 | 0.3214285 | 0.2765188 | 6 | 3710 | 8 | 4.3502532 | 11.125 | 0 | 8 | 0.9857027 | 0.1843220338983051 | TRIO and F-actin binding protein; May regulate actin cytoskeletal org |
| ADH1B | P00325 | 11 | | 9606.ENSP | 3.3469921 | 0.1923076 | 0.2987757 | 7 | 19090 | 13 | 0.0016351 | 12.307692 | 0 | 13 | 0.9871749 | 0.1100917431192660 | Alcohol dehydrogenase 1B, beta polypeptide; Belongs to the zinc-cor |
| ALDH3B1 | P43353 | 11 | | 9606.ENSP | 3.6931124 | 0.3111111 | 0.2707743 | 8 | 5896 | 10 | 5.7221867 | 10.3 | 0 | 10 | 0.9852835 | 0.1758620689655172 | Aldehyde dehydrogenase 3 family, member B1; Oxidizes medium and |
| EPHX1 | Q9NQV0 | 11 | | 9606.ENSP | 3.3504795 | 0.3611111 | 0.2984647 | 7 | 6776 | 9 | 5.1094635 | 18.111111 | 0 | 9 | 0.9871558 | 0.1626262626262626 | Epoxide hydrolase 1, microsomal (xenobiotic); Biotransformation enz |
| FAH | P16930 | 11 | | 9606.ENSP | 3.5501307 | 0.0 | 0.2816797 | 7 | 630 | 3 | 4.3876082 | 24.333333 | 0 | 3 | 0.9860648 | 0.3645833333333333 | Fumarylacetoacetate hydrolase |
| GSTM1 | P09488 | 11 | | 9606.ENSP | 3.1761115 | 0.2747252 | 0.3148503 | 7 | 26718 | 14 | 0.0017450 | 22.5 | 0 | 14 | 0.9881086 | 0.1212355212355212 | Glutathione S-transferase Mu 1; Conjugation of reduced glutathione |
| GSTM2 | P28161 | 11 | | 9606.ENSP | 3.4272013 | 0.5111111 | 0.2917832 | 7 | 6540 | 10 | 3.4715046 | 15.7 | 0 | 10 | 0.9867366 | 0.1652631578947368 | Glutathione S-transferase mu 2 (muscle); Conjugation of reduced glu |
| GSTM5 | P46439 | 11 | | 9606.ENSP | 3.7933740 | 0.7619047 | 0.2636175 | 7 | 326 | 7 | 2.4663272 | 11.285714 | 0 | 7 | 0.9847356 | 0.2893772893772894 | Glutathione S-transferase Mu 5; Conjugation of reduced glutathione |
| MGST1 | P10620 | 11 | | 9606.ENSP | 3.0610287 | 0.3205128 | 0.3266875 | 7 | 32152 | 13 | 0.0020075 | 31.153846 | 0 | 13 | 0.9887375 | 0.1248069199876428 | Microsomal glutathione S-transferase 1; Conjugation of reduced glut |
| C4A | Q9UIP5 | 12 | | 9606.ENSP | 2.8857890 | 0.3623188 | 0.3465256 | 7 | 63580 | 24 | 0.0033015 | 35.125 | 0 | 24 | 0.9896951 | 0.0962328767123287 | C3 and PZP-like alpha-2-macroglobulin domain-containing protein 2; |
| C4B | Q9UIP5 | 12 | | 9606.ENSP | 3.0087183 | 0.4 | 0.3323674 | 7 | 39500 | 20 | 0.0018505 | 32.3 | 0 | 20 | 0.9890233 | 0.1076666666666666 | C3 and PZP-like alpha-2-macroglobulin domain-containing protein 3; |
| CFB | Q9BX92 | 12 | | 9606.ENSP | 2.8945074 | 0.4509803 | 0.3454819 | 7 | 30776 | 18 | 0.0016689 | 48.111111 | 0 | 18 | 0.9896475 | 0.1334876543209876 | Glycine-rich beta glycoprotein; Factor B which is part of the alternate |
| CFH | P08603 | 12 | | 9606.ENSP | 2.9276373 | 0.3809523 | 0.3415723 | 7 | 54676 | 22 | 0.0021615 | 50.409090 | 0 | 22 | 0.9894664 | 0.1481283422459893 | Complement factor H; Factor H functions as a cofactor in the inactiva |
| CFHR1 | Q03591 | 12 | | 9606.ENSP | 3.4768962 | 0.6727272 | 0.2876128 | 7 | 2558 | 11 | 1.3845153 | 18.727272 | 0 | 11 | 0.9864650 | 0.2167019027484144 | Complement factor H-related protein 1; Involved in complement reg |
| MASP1 | F8W876 | 12 | | 9606.ENSP | 3.5780296 | 0.9285714 | 0.2794834 | 7 | 202 | 8 | 1.0210079 | 20.875 | 0 | 8 | 0.9859124 | 0.2899305555555556 | Mannan-binding lectin serine peptidase 1 (C4/C2 activating compone |
| RBMS2 | Q15434 | 12 | | 9606.ENSP | 3.8116826 | 1.0 | 0.2623513 | 8 | 0 | 4 | 0.0 | 19.25 | 0 | 4 | 0.9846356 | 0.5661764705882353 | RNA binding motif single stranded interacting protein 2 |
| SERPING1 | Q7Z455 | 12 | | 9606.ENSP | 3.1046207 | 0.3274853 | 0.3221005 | 7 | 16178 | 19 | 8.2995297 | 27.526315 | 0 | 19 | 0.9884993 | 0.1166853303471444 | Serpin peptidase inhibitor, clade G (C1 inhibitor), member 1; Activati |
| BBOX1 | O75936 | 13 | | 9606.ENSP | 3.0601569 | 0.0583333 | 0.3267806 | 8 | 92182 | 16 | 0.0082152 | 19.6875 | 0 | 16 | 0.9887423 | 0.0779352226720647 | Butyrobetaine (gamma), 2-oxoglutarate dioxygenase (gamma-butyro |
| TRIM34 | Q9BYJ4 | 13 | | 9606.ENSP | 3.9372275 | 0.6666666 | 0.2539858 | 8 | 446 | 3 | 2.9709717 | 10.666666 | 0 | 3 | 0.9839495 | 0.3950617283950617 | Tripartite motif containing 34; May function as antiviral protein and r |
| TRIM38 | O00635 | 13 | | 9606.ENSP | 3.7332170 | 0.3333333 | 0.2678654 | 8 | 700 | 4 | 7.1355283 | 13.25 | 0 | 4 | 0.9850643 | 0.3095238095238095 | RING-type E3 ubiquitin transferase TRIM38; E3 ubiquitin-protein liga |
| TRIM41 | Q8WV44 | 13 | | 9606.ENSP | 3.6503923 | 0.2 | 0.2739431 | 8 | 1980 | 5 | 1.8698326 | 11.6 | 0 | 5 | 0.9855169 | 0.2545454545454545 | RING finger-interacting protein with C kinase; Functions as an E3 liga |
| TRIM5 | Q9C035 | 13 | | 9606.ENSP | 3.2807323 | 0.3186813 | 0.3048099 | 7 | 20350 | 14 | 0.0014354 | 24.714285 | 0 | 14 | 0.9875369 | 0.1621240601503759 | RING-type E3 ubiquitin transferase TRIM5; Capsid-specific restriction |
| TRIM56 | Q9BRZ2 | 13 | | 9606.ENSP | 3.6931124 | 0.3214285 | 0.2707743 | 8 | 7542 | 8 | 6.0031200 | 13.125 | 0 | 8 | 0.9852835 | 0.203125 | RING-type E3 ubiquitin transferase TRIM56; E3 ubiquitin-protein liga |
| MT1A | P04731 | 14 | | 9606.ENSP | 3.6408020 | 0.4 | 0.2746647 | 8 | 1636 | 5 | 1.1247410 | 14.2 | 0 | 5 | 0.9855693 | 0.2592592592592592 | Metallothionein 1A; Metallothioneins have a high content of cysteine |
| MT1E | P04732 | 14 | | 9606.ENSP | 3.8456843 | 0.6190476 | 0.2600317 | 8 | 21378 | 7 | 6.3022701 | 8.2857142 | 0 | 7 | 0.9844498 | 0.2959183673469387 | Metallothionein 1E; Metallothioneins have a high content of cysteine |
| MT1F | P04733 | 14 | | 9606.ENSP | 3.8605056 | 0.8 | 0.2590334 | 8 | 16862 | 5 | 5.1120780 | 9.4 | 0 | 5 | 0.9843688 | 0.3615384615384615 | Metallothionein 1F; Metallothioneins have a high content of cysteine |
| MT1G | P13640 | 14 | | 9606.ENSP | 3.8465562 | 0.7333333 | 0.2599728 | 8 | 20756 | 6 | 6.0627715 | 9.0 | 0 | 6 | 0.9844450 | 0.3214285714285714 | Metallothionein 1G; Metallothioneins have a high content of cysteine |
| MT1M | Q8N339 | 14 | | 9606.ENSP | 4.8430688 | 1.0 | 0.2064806 | 9 | 0 | 3 | 0.0 | 6.0 | 0 | 3 | 0.9789996 | 0.8571428571428571 | Metallothionein 1M; Metallothioneins have a high content of cystein |
| MT1X | P80297 | 14 | | 9606.ENSP | 3.1307759 | 0.5238095 | 0.3194096 | 7 | 34008 | 7 | 0.0024240 | 34.0 | 0 | 7 | 0.9883564 | 0.1659663865546218 | Metallothionein 1X; Metallothioneins have a high content of cysteine |
| MT2A | P02795 | 14 | | 9606.ENSP | 3.3068875 | 0.3928571 | 0.3023991 | 8 | 16874 | 8 | 7.6225168 | 26.125 | 0 | 8 | 0.9873940 | 0.1906934306569343 | Metallothionein 2A; Metallothioneins have a high content of cysteine |
| NQO1 | P15559 | 14 | | 9606.ENSP | 2.9808195 | 0.1304347 | 0.3354782 | 7 | 165930 | 24 | 0.0113634 | 24.708333 | 0 | 24 | 0.9891758 | 0.0819397993311036 | NAD(P)H dehydrogenase [quinone] 1; The enzyme apparently serves |
| GPR146 | Q96CH1 | 15 | | 9606.ENSP | 4.6198779 | 0.0 | 0.2164559 | 8 | 0 | 1 | 0.0 | 7.0 | 0 | 1 | 0.9802192 | 0.0 | Probable G-protein coupled receptor 146; Orphan receptor; G protei |
| MS4A6A | Q9H2W1 | 15 | | 9606.ENSP | 3.4376634 | 0.2888888 | 0.2908952 | 7 | 26710 | 10 | 0.0013706 | 18.8 | 0 | 10 | 0.9866794 | 0.1528455284552845 | Membrane-spanning 4-domains, subfamily A, member 6A; May be in |
| MS4A7 | Q9GZW8 | 15 | | 9606.ENSP | 3.6207497 | 0.3333333 | 0.2761858 | 7 | 35948 | 7 | 0.0021790 | 12.0 | 0 | 7 | 0.9856789 | 0.1859410430839002 | Membrane-spanning 4-domains, subfamily A, member 7; May be inv |
| PIK3IP1 | Q96FE7 | 15 | | 9606.ENSP | 4.4167393 | 0.3333333 | 0.2264113 | 7 | 480 | 3 | 8.2536517 | 5.0 | 0 | 3 | 0.9813292 | 0.3888888888888889 | Phosphoinositide-3-kinase interacting protein 1; Negative regulator c |
| TMEM176 | Q96HP8 | 15 | | 9606.ENSP | 4.2824760 | 1.0 | 0.2335097 | 7 | 0 | 3 | 0.0 | 8.6666666 | 0 | 3 | 0.9820629 | 0.5098039215686274 | Hepatocellular carcinoma-associated antigen 112; Membrane spanni |
| TMEM176 | Q3YBM2 | 15 | | 9606.ENSP | 3.5457715 | 0.2222222 | 0.2820260 | 6 | 41740 | 9 | 0.0030268 | 11.222222 | 0 | 9 | 0.9860886 | 0.1406469760900140 | Transmembrane protein 176B; May play a role in the process of matu |
| AMOT | Q4VCS5 | 16 | | 9606.ENSP | 3.0959023 | 0.3161764 | 0.3230076 | 7 | 12478 | 17 | 9.1857764 | 23.882352 | 0 | 17 | 0.9885469 | 0.1224984839296543 | Angiomotin; Plays a central role in tight junction maintenance via the |
| TEAD1 | P28347 | 16 | | 9606.ENSP | 2.8326068 | 0.3369565 | 0.3530317 | 7 | 56294 | 24 | 0.0035339 | 38.916666 | 0 | 24 | 0.9899857 | 0.1045026881720430 | TEA domain family member 1 (SV40 transcriptional enhancer factor); |
| TEAD2 | Q15562 | 16 | | 9606.ENSP | 3.0549258 | 0.5 | 0.3273401 | 7 | 33342 | 17 | 0.0019510 | 32.705882 | 0 | 17 | 0.9887708 | 0.1329507412721186 | Transcriptional enhancer factor TEF-4; Transcription factor which pla |
| TEAD3 | Q99594 | 16 | | 9606.ENSP | 3.2092414 | 0.5428571 | 0.3116001 | 7 | 6654 | 15 | 3.8728049 | 26.666666 | 0 | 15 | 0.9879276 | 0.1568627450980392 | Transcriptional enhancer factor TEF-5; Transcription factor which pla |
| WWC1 | Q8IX03 | 16 | | 9606.ENSP | 3.0662598 | 0.3071895 | 0.3261302 | 7 | 51358 | 18 | 0.0029732 | 29.833333 | 0 | 18 | 0.9887089 | 0.1243607624360762 | WW domain-containing protein 1; Probable regulator of the Hippo/S |
| WWTR1 | Q9GZV5 | 16 | | 9606.ENSP | 2.8500435 | 0.2982456 | 0.3508718 | 7 | 52172 | 19 | 0.0034266 | 46.0 | 0 | 19 | 0.9898904 | 0.1243759806019112 | WW domain-containing transcription regulator protein 1; Transcripti |
| YAP1 | P46937 | 16 | | 9606.ENSP | 2.6460331 | 0.2043740 | 0.3779242 | 7 | 154360 | 52 | 0.0084117 | 40.288461 | 0 | 52 | 0.9910052 | 0.0822715117193644 | Yes-associated protein YAP65 homolog; Transcriptional regulator wh |
| APOL1 | Q9BQ03 | 17 | | 9606.ENSP | 3.2772449 | 0.3454545 | 0.3051343 | 7 | 14442 | 11 | 9.1684664 | 21.636363 | 0 | 11 | 0.9875560 | 0.1572660915726609 | Apolipoprotein L, 1; May play a role in lipid exchange and transport t |
| CLU | P10909 | 17 | | 9606.ENSP | 2.7776809 | 0.2158730 | 0.3600125 | 7 | 100386 | 36 | 0.0059055 | 36.25 | 0 | 36 | 0.9902858 | 0.084696261682243 | Testosterone-repressed prostate message 2; Isoform 1 functions as e |
| HPR | P00739 | 17 | | 9606.ENSP | 3.3190932 | 0.3555555 | 0.3012871 | 7 | 2100 | 10 | 1.6447315 | 21.4 | 0 | 10 | 0.9873273 | 0.1752066115702479 | Haptoglobin-related protein; Primate-specific plasma protein associa |
| LCAT | P04180 | 17 | | 9606.ENSP | 3.0217959 | 0.1323529 | 0.3309290 | 7 | 54522 | 17 | 0.0041789 | 23.588235 | 0 | 17 | 0.9889519 | 0.0927058823529411 | Phosphatidylcholine-sterol acyltransferase; Central enzyme in the ex |
| PLA1A | Q53H76 | 17 | | 9606.ENSP | 3.5448997 | 0.0 | 0.2820954 | 8 | 280 | 3 | 1.5051646 | 20.666666 | 0 | 3 | 0.9860934 | 0.3575757575757576 | Phosphatidylserine-specific phospholipase A1; Hydrolyzes the ester b |
| PON3 | Q15166 | 17 | | 9606.ENSP | 3.5152571 | 0.4 | 0.2844742 | 8 | 1570 | 6 | 1.4430771 | 15.833333 | 0 | 6 | 0.9862554 | 0.2206572769953051 | Serum paraoxonase/lactonase 3; Has low activity towards the organo |
| CLEC1A | Q8NC01 | 18 | | 9606.ENSP | 4.0619006 | 0.0 | 0.2461901 | 8 | 29968 | 2 | 0.0024233 | 6.5 | 0 | 2 | 0.9832683 | 0.5 | C-type lectin domain family 1 member A |
| CLEC4E | Q9ULY5 | 18 | | 9606.ENSP | 3.0723626 | 0.4722222 | 0.3254824 | 7 | 66130 | 9 | 0.0044758 | 65.555555 | 0 | 9 | 0.9886756 | 0.2623828647925033 | C-type lectin domain family 4, member E; C-type lectin that functions |
| CNTNAP3 | Q9BZ76 | 18 | | 9606.ENSP | 5.7279860 | 0.0 | 0.1745814 | 10 | 0 | 1 | 0.0 | 4.0 | 0 | 1 | 0.9741640 | 0.0 | Contactin associated protein like 3 |
| GLIPR1L2 | Q4G1C9 | 18 | | 9606.ENSP | 4.7288578 | 0.0 | 0.2114675 | 9 | 35512 | 4 | 0.0035001 | 1.5 | 0 | 4 | 0.9796237 | 0.25 | GLI pathogenesis-related 1 like 2; GLI pathogenesis related family 1; |
| SEC14L2 | O76054 | 18 | | 9606.ENSP | 3.8927637 | 0.0 | 0.2568868 | 8 | 23232 | 2 | 0.0028068 | 9.5 | 0 | 2 | 0.9841925 | 0.5 | Alpha-tocopherol-associated protein; Carrier protein. Binds to some |
| ZNF98 | A6NK75 | 18 | | 9606.ENSP | 5.7279860 | 0.0 | 0.1745814 | 10 | 0 | 1 | 0.0 | 4.0 | 0 | 1 | 0.9741640 | 0.0 | Zinc finger protein F7175; May be involved in transcriptional regulati |
| EYA2 | Q9H4S9 | 19 | | 9606.ENSP | 3.4646904 | 0.5 | 0.2886260 | 7 | 656 | 5 | 3.6081105 | 24.0 | 0 | 5 | 0.9865317 | 0.2696629213483146 | EYA transcriptional coactivator and phosphatase 2; Functions both as |
| FCGRT | P55899 | 19 | | 9606.ENSP | 3.0697471 | 0.4285714 | 0.3257597 | 7 | 6988 | 7 | 3.0740104 | 63.857142 | 0 | 7 | 0.9886899 | 0.2435686918445539 | Fc fragment of IgG, receptor, transporter, alpha; Binds to the Fc regio |
| FREM2 | Q5SZK8 | 19 | | 9606.ENSP | 3.0828247 | 0.3 | 0.3243778 | 7 | 5998 | 5 | 3.8102323 | 80.6 | 0 | 5 | 0.9886184 | 0.3355648535564853 | FRAS1 related extracellular matrix protein 2; Extracellular matrix pro |

| | | | | | | | | | | | | | | | |
|---|---|---|---|---|---|---|---|---|---|---|---|---|---|---|---|
| IGFBP5 | P24593 | 19 | 9606.ENSP | 3.1107236 | 0.2666666 | 0.3214686 | 7 | 3064 | 6 | 1.4253919 | 56.0 | 0 | 6 | 0.9884659 | 0.249255952380952 | Insulin-like growth factor binding protein 5; IGF-binding proteins pro |
| INHBB | P09529 | 19 | 9606.ENSP | 3.3417611 | 0.1388888 | 0.2992434 | 8 | 4454 | 9 | 3.4167836 | 19.555555 | 0 | 9 | 0.9872034 | 0.161616161616161 | Activin beta-B chain; Inhibins and activins inhibit and activate, respec |
| SIX5 | Q8N196 | 19 | 9606.ENSP | 3.4149956 | 0.0952380 | 0.2928261 | 8 | 1630 | 7 | 1.2436242 | 15.714285 | 0 | 7 | 0.9868033 | 0.169841269841269 | DM locus-associated homeodomain protein; Transcription factor tha |
| MYBPC1 | Q15497 | 20 | 9606.ENSP | 3.5196163 | 0.3333333 | 0.2841218 | 8 | 7682 | 4 | 4.5078839 | 18.75 | 0 | 4 | 0.9862316 | 0.2803030303030303 | C-protein, skeletal muscle slow isoform; Thick filament-associated pr |
| MYH15 | Q9Y2K3 | 20 | 9606.ENSP | 3.6364428 | 0.0 | 0.2749940 | 7 | 502 | 3 | 5.2571136 | 18.333333 | 0 | 3 | 0.9855932 | 0.353741496598639 | Myosin, heavy chain 15; Muscle contraction; Belongs to the TRAFAC |
| MYL3 | P08590 | 20 | 9606.ENSP | 3.3574542 | 0.1777777 | 0.2978447 | 7 | 19810 | 10 | 0.0012529 | 14.1 | 0 | 10 | 0.9871177 | 0.130188679245283 | Myosin light chain 1, slow-twitch muscle B/ventricular isoform; Regu |
| MYOT | Q9UBF9 | 20 | 9606.ENSP | 3.5257192 | 0.2666666 | 0.2836300 | 8 | 4476 | 6 | 3.0402870 | 18.0 | 0 | 6 | 0.9861982 | 0.212851405622489 | Myofibrillar titin-like Ig domains protein; Component of a complex of |
| MYOZ1 | Q9NP98 | 20 | 9606.ENSP | 3.9302528 | 0.4666666 | 0.2544365 | 8 | 1158 | 6 | 1.1714591 | 8.0 | 0 | 6 | 0.9839876 | 0.25 | Filamin-, actinin- and telethonin-binding protein; Myozenins may ser |
| PGAM2 | P15259 | 20 | 9606.ENSP | 3.8230165 | 0.1 | 0.2615735 | 8 | 6848 | 5 | 9.2103107 | 10.0 | 0 | 5 | 0.9845736 | 0.229268292682926 | Muscle-specific phosphoglycerate mutase; Interconversion of 3- and |
| EPB41L5 | Q9HCM4 | 21 | 9606.ENSP | 3.4795117 | 0.0666666 | 0.2873966 | 8 | 54772 | 6 | 0.0044701 | 17.0 | 0 | 6 | 0.9864507 | 0.1814814814814815 | Erythrocyte membrane protein band 4.1 like 5; May contribute to the |
| NXN | Q6DKJ4 | 21 | 9606.ENSP | 3.4568439 | 0.0 | 0.2892812 | 7 | 2406 | 3 | 1.6650436 | 30.666666 | 0 | 3 | 0.9865746 | 0.3449612403100775 | Nucleoredoxin; Functions as a redox-dependent negative regulator o |
| PAIP2B | Q9ULR5 | 21 | 9606.ENSP | 4.2327811 | 0.3333333 | 0.2362512 | 9 | 2000 | 3 | 3.0046082 | 5.6666666 | 0 | 3 | 0.9823345 | 0.380952380952380 | Polyadenylate-binding protein-interacting protein 2B; Inhibits transla |
| PARD3B | Q8TEW8 | 21 | 9606.ENSP | 4.4193548 | 0.3333333 | 0.2262773 | 9 | 22396 | 4 | 0.0019382 | 3.5 | 0 | 4 | 0.9813150 | 0.35 | Amyotrophic lateral sclerosis 2 chromosomal region candidate gene |
| RBMS3 | Q6XE24 | 21 | 9606.ENSP | 5.0 | 0.3333333 | 0.2 | 8 | 1842 | 3 | 2.0335318 | 3.6666666 | 0 | 3 | 0.9781420 | 0.4166666666666667 | RNA binding motif, single stranded interacting protein 3; Binds poly( |
| ZMYM5 | Q9UJ78 | 21 | 9606.ENSP | 5.3783783 | 1.0 | 0.1859296 | 9 | 0 | 2 | 0.0 | 3.5 | 0 | 2 | 0.9760744 | 0.7 | Zinc finger MYM-type protein 5; Functions as a transcriptional regula |
| ADD3 | Q9UEY8 | 22 | 9606.ENSP | 4.5387968 | 0.0 | 0.2203227 | 9 | 0 | 1 | 0.0 | 10.0 | 0 | 1 | 0.9806623 | 0.0 | Adducin-like protein 70; Membrane-cytoskeleton-associated protein |
| ARHGAP19 | Q8NF34 | 22 | 9606.ENSP | 4.5387968 | 0.0 | 0.2203227 | 9 | 0 | 1 | 0.0 | 10.0 | 0 | 1 | 0.9806623 | 0.0 | Rho-type GTPase-activating protein 19; GTPase activator for the Rho |
| FES | P07332 | 22 | 9606.ENSP | 2.9049694 | 0.1758241 | 0.3442376 | 7 | 47954 | 14 | 0.0032883 | 44.142857 | 0 | 14 | 0.9895903 | 0.132561132561132 | Feline sarcoma/Fujinami avian sarcoma oncogene homolog; Tyrosine |
| NOSTRIN | A0A0G2JP | 22 | 9606.ENSP | 3.2528334 | 0.4 | 0.3074242 | 8 | 7310 | 5 | 5.3977428 | 38.8 | 0 | 5 | 0.9876894 | 0.269444444444444 | Nitric oxide synthase trafficking; F-BAR domain containing |
| SCARA3 | Q6AZY7 | 22 | 9606.ENSP | 3.3653007 | 0.0 | 0.2971502 | 8 | 4750 | 2 | 3.3239130 | 59.0 | 0 | 2 | 0.9870748 | 0.5 | Cellular stress response gene protein; Seems to protect cells by scave |
| SRGAP1 | Q7Z6B7 | 22 | 9606.ENSP | 3.5396687 | 0.1111111 | 0.2825123 | 8 | 56102 | 10 | 0.0054793 | 7.9 | 0 | 10 | 0.9861220 | 0.123728813559322 | SLIT-ROBO Rho GTPase activating protein 1; GTPase-activating protei |
| ADORA2A | P29274 | 23 | 9606.ENSP | 3.0087183 | 0.3076923 | 0.3323674 | 7 | 18624 | 13 | 0.0010503 | 37.692307 | 0 | 13 | 0.9890233 | 0.1350978770333609 | Adenosine A2a receptor; Receptor for adenosine. The activity of this |
| ADORA2B | P29275 | 23 | 9606.ENSP | 2.9442022 | 0.1985294 | 0.3396505 | 7 | 45070 | 17 | 0.0026306 | 32.705882 | 0 | 17 | 0.9893759 | 0.107039537126325 | Adenosine A2b receptor; Receptor for adenosine. The activity of this |
| P2RY2 | P41231 | 23 | 9606.ENSP | 3.0462074 | 0.3928571 | 0.3282770 | 7 | 5232 | 8 | 2.6223638 | 55.875 | 0 | 8 | 0.9888185 | 0.215733590733590 | Purinergic receptor P2Y, G-protein coupled, 2; Receptor for ATP and |
| RAPGEF3 | O95398 | 23 | 9606.ENSP | 3.1900610 | 0.1944444 | 0.3134736 | 7 | 9152 | 9 | 5.9380084 | 29.777777 | 0 | 9 | 0.9880324 | 0.160898603521554 | Rap1 guanine-nucleotide-exchange factor directly activated by cAMP |
| CCDC103 | Q8IW40 | 24 | 9606.ENSP | 3.8761987 | 0.5 | 0.2579847 | 8 | 1432 | 4 | 1.2304516 | 9.5 | 0 | 4 | 0.9842830 | 0.296875 | Coiled-coil domain containing 103; Dynein-attachment factor require |
| FAM187A | Q8IW40 | 24 | 9606.ENSP | 3.8761987 | 0.5 | 0.2579847 | 8 | 1432 | 4 | 1.2304516 | 9.5 | 0 | 4 | 0.9842830 | 0.296875 | Family with sequence similarity 187, member A; Dynein-attachment |
| FOXJ1 | Q92949 | 24 | 9606.ENSP | 3.0662598 | 0.1691176 | 0.3261302 | 7 | 102622 | 17 | 0.0066022 | 31.352941 | 0 | 17 | 0.9887089 | 0.128540305010893 | Hepatocyte nuclear factor 3 forkhead homolog 4; Transcription facto |
| HIGD1B | Q9P298 | 24 | 9606.ENSP | 3.9110723 | 0.3333333 | 0.2556843 | 8 | 1916 | 4 | 1.6362723 | 11.25 | 0 | 4 | 0.9840925 | 0.45 | HIG1 hypoxia inducible domain family member 1B |
| RPGR | Q9UMR1 | 24 | 9606.ENSP | 3.4062772 | 0.0769230 | 0.2935756 | 8 | 20782 | 13 | 0.0027842 | 9.6153846 | 0 | 13 | 0.9868509 | 0.094017094017094 | X-linked retinitis pigmentosa GTPase regulator; Could be a guanine-n |
| CCDC8 | Q9H0W5 | 25 | 9606.ENSP | 4.3156059 | 0.0 | 0.2317171 | 8 | 0 | 1 | 0.0 | 9.0 | 0 | 1 | 0.9818819 | 0.0 | Coiled-coil domain-containing protein 8; Core component of the 3M |
| MDFI | Q99750 | 25 | 9606.ENSP | 3.6015693 | 0.0357142 | 0.2776567 | 8 | 26578 | 8 | 0.0024253 | 6.875 | 0 | 8 | 0.9857837 | 0.139204545454545 | Myogenic repressor I-mf; Inhibits the transactivation activity of the M |
| PBXIP1 | Q96AQ6 | 25 | 9606.ENSP | 4.2101133 | 0.0 | 0.2375232 | 8 | 280 | 2 | 4.6813822 | 8.5 | 0 | 2 | 0.9824583 | 0.5 | Pre-B-cell leukemia transcription factor-interacting protein 1; Regula |
| RAB13 | P51153 | 25 | 9606.ENSP | 3.3164777 | 0.0833333 | 0.3015247 | 7 | 44234 | 9 | 0.0034713 | 15.666666 | 0 | 9 | 0.9873416 | 0.126984126984126 | Cell growth-inhibiting gene 4 protein; The small GTPases Rab are key |
| RPS27 | P42677 | 25 | 9606.ENSP | 3.3931996 | 0.0 | 0.2947070 | 8 | 1626 | 3 | 9.2753543 | 31.333333 | 0 | 3 | 0.9869224 | 0.337037037037037 | Small ribosomal subunit protein eS27; Component of the small ribos |
| MTURN | Q8N3F0 | 26 | 9606.ENSP | 4.1813426 | 0.0 | 0.2391576 | 7 | 0 | 1 | 0.0 | 13.0 | 0 | 1 | 0.9826156 | 0.0 | Maturin, neural progenitor differentiation regulator homolog (Xenop |
| NKD1 | Q969G9 | 26 | 9606.ENSP | 3.1822144 | 0.1410256 | 0.3142465 | 6 | 136884 | 13 | 0.0090984 | 22.923076 | 0 | 13 | 0.9880753 | 0.120065120065120 | Naked cuticle homolog 1 (Drosophila); Cell autonomous antagonist o |
| PPP1R3B | Q86XI6 | 26 | 9606.ENSP | 3.8352223 | 0.2666666 | 0.2607410 | 7 | 9812 | 6 | 8.9699265 | 7.5 | 0 | 6 | 0.9845069 | 0.2107843137254902 | Hepatic glycogen-targeting protein phosphatase 1 regulatory subunit |
| PPP1R3C | Q9UQK1 | 26 | 9606.ENSP | 4.1211857 | 0.6666666 | 0.2426486 | 7 | 2180 | 4 | 1.5117689 | 6.75 | 0 | 4 | 0.9829443 | 0.355263157894736 | Protein phosphatase 1, regulatory subunit 3C; Acts as a glycogen-targ |
| PPP1R3G | B7ZBB8 | 26 | 9606.ENSP | 4.1438535 | 1.0 | 0.2413212 | 7 | 0 | 3 | 0.0 | 7.6666666 | 0 | 3 | 0.9828204 | 0.4791666666666667 | Protein phosphatase 1, regulatory subunit 3G; Glycogen-targeting su |
| ARHGEF15 | O94989 | 27 | 9606.ENSP | 4.0322580 | 0.3333333 | 0.248 | 8 | 2434 | 3 | 1.1564966 | 8.0 | 0 | 3 | 0.9834302 | 0.425925925925925 | Rho guanine nucleotide exchange factor (GEF) 15; Specific GEF for Rh |
| ARHGEF40 | Q8TER5 | 27 | 9606.ENSP | 4.2850915 | 0.1071428 | 0.2333672 | 9 | 5142 | 8 | 7.7852921 | 4.375 | 0 | 8 | 0.9820486 | 0.166666666666666 | Rho guanine nucleotide exchange factor (GEF) 40; May act as a guani |
| PLEKHG2 | Q9H7P9 | 27 | 9606.ENSP | 4.7462946 | 1.0 | 0.2106906 | 9 | 0 | 2 | 0.0 | 7.0 | 0 | 2 | 0.9795284 | 0.6363636363636364 | Pleckstrin homology domain containing, family G (with RhoGef doma |
| SWAP70 | Q9UH65 | 27 | 9606.ENSP | 3.8413251 | 0.2 | 0.2603268 | 8 | 23982 | 6 | 0.0019924 | 7.8333333 | 0 | 6 | 0.9844736 | 0.219047619047619 | SWAP switching B-cell complex 70kDa subunit; Phosphatidylinositol 3 |
| TIPARP | Q7Z3E1 | 27 | 9606.ENSP | 3.9529206 | 0.0 | 0.2529775 | 8 | 4452 | 2 | 3.1039080 | 12.0 | 0 | 2 | 0.9838638 | 0.5 | ADP-ribosyltransferase diphtheria toxin-like 14; Poly [ADP-ribose] po |
| NBPF10 | A0A075B7 | 28 | 9606.ENSP | 4.4594594 | 0.0 | 0.2242424 | 8 | 0 | 1 | 0.0 | 9.0 | 0 | 1 | 0.9810958 | 0.0 | Neuroblastoma breakpoint family, member 10; NBPF member 10 |
| NBPF26 | A0A087W | 28 | 9606.ENSP | 4.4594594 | 0.0 | 0.2242424 | 8 | 0 | 1 | 0.0 | 9.0 | 0 | 1 | 0.9810958 | 0.0 | Neuroblastoma breakpoint family, member 14; NBPF member 14 |
| NBPF9 | Q3BBV1 | 28 | 9606.ENSP | 4.4594594 | 0.0 | 0.2242424 | 8 | 0 | 1 | 0.0 | 9.0 | 0 | 1 | 0.9810958 | 0.0 | Neuroblastoma breakpoint family member 20; NBPF member 9; Belo |
| MYO6 | Q9UM54 | 29 | 9606.ENSP | 3.2292938 | 0.2426470 | 0.3096652 | 7 | 25066 | 17 | 0.0020942 | 15.588235 | 0 | 17 | 0.9878180 | 0.098164106406893 | Unconventional myosin-VI; Myosins are actin-based motor molecules |
| PDLIM1 | O00151 | 29 | 9606.ENSP | 3.1046207 | 0.3076923 | 0.3221005 | 6 | 14356 | 14 | 0.0011038 | 25.714285 | 0 | 14 | 0.9884993 | 0.117840684660961 | C-terminal LIM domain protein 1; Cytoskeletal protein that may act a |
| PDLIM2 | Q96JY6 | 29 | 9606.ENSP | 3.2040104 | 0.3818181 | 0.3121088 | 7 | 10538 | 11 | 8.0291276 | 22.181818 | 0 | 11 | 0.9879562 | 0.127591706539074 | PDZ and LIM domain 2 (mystique); Probable adapter protein located |
| PDLIM3 | Q53GG5 | 29 | 9606.ENSP | 3.2571926 | 0.2647058 | 0.3070128 | 7 | 21972 | 17 | 0.0022390 | 14.941176 | 0 | 17 | 0.9876656 | 0.097020626432391 | Alpha-actinin-2-associated LIM protein; May play a role in the organi |
| PDLIM4 | P50479 | 29 | 9606.ENSP | 3.3025283 | 0.3939393 | 0.3027983 | 7 | 5406 | 12 | 5.3337570 | 16.166666 | 0 | 12 | 0.9874178 | 0.126322751322751 | Reversion-induced LIM protein; Isoform 1: Suppresses SRC activation |
| CTDSP1 | Q9GZU7 | 31 | 9606.ENSP | 3.2571926 | 0.2857142 | 0.3070128 | 7 | 6772 | 7 | 4.4551993 | 24.0 | 0 | 7 | 0.9876656 | 0.166833166833166 | CTD (carboxy-terminal domain, RNA polymerase II, polypeptide A) sm |
| CTDSP2 | O14595 | 31 | 9606.ENSP | 3.6782911 | 0.2 | 0.2718653 | 8 | 980 | 5 | 1.0539849 | 10.2 | 0 | 5 | 0.9853645 | 0.239024390243902 | CTD (carboxy-terminal domain, RNA polymerase II, polypeptide A) sm |
| CTDSPL | O15194 | 31 | 9606.ENSP | 3.4829991 | 0.2666666 | 0.2871088 | 8 | 2774 | 6 | 2.4473617 | 15.0 | 0 | 6 | 0.9864316 | 0.200450450450450 | CTD (carboxy-terminal domain, RNA polymerase II, polypeptide A) sm |
| LPIN3 | Q9BQK8 | 31 | 9606.ENSP | 3.4655623 | 0.2380952 | 0.2885534 | 8 | 4184 | 7 | 3.8258187 | 15.285714 | 0 | 7 | 0.9865269 | 0.199248120300751 | Phosphatidate phosphatase LPIN3; Regulates fatty acid metabolism. |
| REST | Q13127 | 31 | 9606.ENSP | 3.1037489 | 0.1978021 | 0.3221910 | 7 | 40758 | 14 | 0.0029229 | 26.285714 | 0 | 14 | 0.9885041 | 0.120803159973668 | Neural-restrictive silencer factor; Transcriptional repressor which bin |
| ABCB1 | P08183 | 32 | 9606.ENSP | 2.6887532 | 0.4031620 | 0.3719195 | 7 | 95334 | 23 | 0.0048324 | 72.956521 | 0 | 23 | 0.9907718 | 0.147685266678401 | ATP-binding cassette, sub-family B (MDR/TAP), member 1; Energy-de |
| SLC15A2 | Q16348 | 32 | 9606.ENSP | 3.4341761 | 0.25 | 0.2911906 | 8 | 33210 | 8 | 0.0020752 | 14.875 | 0 | 8 | 0.9866984 | 0.1828125 | Solute carrier family 15 (oligopeptide transporter), member 2; Proto |
| SLC2A1 | P11166 | 32 | 9606.ENSP | 2.6660854 | 0.1817073 | 0.3750817 | 7 | 255442 | 41 | 0.0147235 | 44.926829 | 0 | 41 | 0.9908957 | 0.088995354239256 | Solute carrier family 2 (facilitated glucose transporter), member 1; Fa |
| SLC7A2 | P52569 | 32 | 9606.ENSP | 3.4925893 | 0.3333333 | 0.2863205 | 8 | 258 | 3 | 7.0091701 | 32.333333 | 0 | 3 | 0.9863792 | 0.432432432432432 | Solute carrier family 7 (cationic amino acid transporter, y+ system), m |
| SLCO4A1 | Q96BD0 | 32 | 9606.ENSP | 4.4333042 | 0.0 | 0.2255653 | 9 | 0 | 1 | 0.0 | 8.0 | 0 | 1 | 0.9812387 | 0.0 | Solute carrier organic anion transporter family, member 4A1; Mediat |
| CHST14 | Q8NCH0 | 33 | 9606.ENSP | 3.5902353 | 0.2666666 | 0.2785332 | 7 | 9820 | 6 | 6.1011535 | 14.333333 | 0 | 6 | 0.9858457 | 0.2153846153846154 | Carbohydrate (N-acetylgalactosamine 4-0) sulfotransferase 14; Catal |
| CHST3 | Q7LGC8 | 33 | 9606.ENSP | 3.5736704 | 0.4 | 0.2798243 | 7 | 2232 | 5 | 1.6278889 | 17.6 | 0 | 5 | 0.9859362 | 0.259701492537313 | Galactose/N-acetylglucosamine/N-acetylglucosamine 6-O-sulfotransf |
| CHST6 | Q9GZX3 | 33 | 9606.ENSP | 4.1185701 | 0.0 | 0.2428027 | 8 | 32 | 2 | 4.9387894 | 6.5 | 0 | 2 | 0.9829586 | 0.5 | Galactose/N-acetylglucosamine/N-acetylglucosamine 6-O-sulfotransf |
| PRELP | P51888 | 33 | 9606.ENSP | 3.1473408 | 0.1904761 | 0.3177285 | 7 | 19114 | 7 | 0.0017470 | 39.142857 | 0 | 7 | 0.9882658 | 0.188850174216027 | Proline-arginine-rich end leucine-rich repeat protein; May anchor ba |
| TSKU | Q8WUA8 | 33 | 9606.ENSP | 3.7698343 | 0.0 | 0.2652636 | 7 | 12 | 2 | 9.1814083 | 23.0 | 0 | 2 | 0.9848642 | 0.5365853658536586 | Leucine-rich repeat-containing protein 54; Tsukushi, small leucine ric |
| GPC5 | P78333 | 34 | 9606.ENSP | 3.1857018 | 0.3636363 | 0.3139025 | 7 | 10606 | 12 | 5.1163836 | 29.5 | 0 | 12 | 0.9880562 | 0.150085034013605 | Glypican 5; Cell surface proteoglycan that bears heparan sulfate; Bel |
| HPSE2 | Q8WWQ2 | 34 | 9606.ENSP | 3.2781168 | 0.1904761 | 0.3050531 | 7 | 2656 | 7 | 1.3956014 | 31.0 | 0 | 7 | 0.9875512 | 0.201127819548872 | Heparanase 2 (inactive); Binds heparin and heparan sulfate with high |
| HSPG2 | P98160 | 34 | 9606.ENSP | 2.8299912 | 0.2890756 | 0.3533579 | 7 | 84912 | 35 | 0.0041962 | 41.971428 | 0 | 35 | 0.9900000 | 0.1124473381846036 | Basement membrane-specific heparan sulfate proteoglycan core pro |
| LRP10 | Q7Z4F1 | 34 | 9606.ENSP | 3.6687009 | 1.0 | 0.2725760 | 8 | 0 | 3 | 0.0 | 27.333333 | 0 | 3 | 0.9854169 | 0.5061728395061729 | Low density lipoprotein receptor-related protein 10; Probable recept |
| DAAM2 | Q86T65 | 35 | 9606.ENSP | 3.5318221 | 0.1 | 0.2831399 | 7 | 4432 | 5 | 4.8652568 | 13.4 | 0 | 5 | 0.9861649 | 0.213333333333333 | Dishevelled associated activator of morphogenesis 2; Armadillo-like |
| PKN3 | Q6P5Z2 | 35 | 9606.ENSP | 3.6399302 | 0.1666666 | 0.2747305 | 8 | 1684 | 4 | 1.5733083 | 14.5 | 0 | 4 | 0.9855741 | 0.3181818181818182 | Serine/threonine-protein kinase N3; Contributes to invasiveness in m |
| PREX2 | Q70Z35 | 35 | 9606.ENSP | 3.7088055 | 0.0 | 0.2696285 | 8 | 0 | 1 | 0.0 | 31.0 | 0 | 1 | 0.9851977 | 0.0 | Phosphatidylinositol 3,4,5-trisphosphate-dependent Rac exchanger 2 |
| RHOC | P08134 | 35 | 9606.ENSP | 2.7096774 | 0.2043010 | 0.3690476 | 7 | 191924 | 31 | 0.0119435 | 47.322580 | 0 | 31 | 0.9906575 | 0.104037598803674 | Rho-related GTP-binding protein RhoC; Regulates a signal transducti |
| HSD3B7 | Q9H2F3 | 36 | 9606.ENSP | 4.0784655 | 0.0 | 0.2451902 | 8 | 602 | 3 | 1.1444087 | 8.3333333 | 0 | 3 | 0.9831777 | 0.3333333333333333 | Hydroxy-delta-5-steroid dehydrogenase, 3 beta- and steroid delta-iso |
| SLC51B | Q86UW2 | 36 | 9606.ENSP | 4.3496076 | 0.0 | 0.2299057 | 9 | 110 | 2 | 9.0422553 | 6.5 | 0 | 2 | 0.9816961 | 0.6111111111111112 | Organic solute transporter subunit beta; Essential component of the |
| SLCO1A2 | P46721 | 36 | 9606.ENSP | 3.5065387 | 0.0357142 | 0.2851815 | 8 | 37092 | 8 | 0.0022212 | 9.5 | 0 | 8 | 0.9863030 | 0.145833333333333 | Solute carrier organic anion transporter family, member 1A2; Mediat |
| SLCO2B1 | O94956 | 36 | 9606.ENSP | 3.4725370 | 0.3 | 0.2879738 | 8 | 21344 | 5 | 0.0011137 | 18.6 | 0 | 5 | 0.9864888 | 0.2493150684931507 | Solute carrier organic anion transporter family, member 2B1; Mediat |
| AHCYL1 | O43865 | 37 | 9606.ENSP | 3.1621621 | 0.0439560 | 0.3162393 | 7 | 65982 | 14 | 0.0063118 | 15.714285 | 0 | 14 | 0.9881849 | 0.082747853239656 | S-adenosylhomocysteine hydrolase-like protein 1; Multifaceted cellul |
| BHMT2 | Q9H2M3 | 37 | 9606.ENSP | 3.8282476 | 0.3333333 | 0.2612161 | 8 | 1914 | 7 | 2.0000702 | 8.1428571 | 0 | 7 | 0.9845450 | 0.214285714285714 | S-methylmethionine--homocysteine S-methyltransferase BHMT2; Inv |
| SDS | P20132 | 37 | 9606.ENSP | 4.0078465 | 0.3333333 | 0.2495105 | 8 | 748 | 4 | 9.6512086 | 9.75 | 0 | 4 | 0.9835636 | 0.325 | L-serine dehydratase/L-threonine deaminase; Serine dehydratase |
| EDA | Q9Y6L0 | 38 | 9606.ENSP | 2.9973844 | 0.1111111 | 0.3336242 | 6 | 13098 | 9 | 8.8915318 | 45.222222 | 0 | 9 | 0.9890853 | 0.160493827160493 | Ectodermal dysplasia protein; Cytokine which is involved in epithelial |
| FOXQ1 | Q9C009 | 38 | 9606.ENSP | 4.1098517 | 1.0 | 0.2433177 | 8 | 0 | 2 | 0.0 | 13.0 | 0 | 2 | 0.9830062 | 0.6190476190476191 | Hepatocyte nuclear factor 3 forkhead homolog 1; Plays a role in hair |
| LRRC1 | Q9BTT6 | 38 | 9606.ENSP | 3.7018308 | 0.0 | 0.2701365 | 7 | 284 | 3 | 2.7521066 | 15.666666 | 0 | 3 | 0.9852358 | 0.3492063492063492 | Leucine-rich repeat-containing protein 1; Leucine rich repeat contain |
| TCF15 | Q12870 | 38 | 9606.ENSP | 3.1883173 | 0.2222222 | 0.3136450 | 7 | 25614 | 9 | 0.0012698 | 29.0 | 0 | 9 | 0.9880419 | 0.154954954954954 | Transcription factor 15 (basic helix-loop-helix); May function as an ea |
| ADAMTS9 | Q9P2N4 | 39 | 9606.ENSP | 3.2807323 | 0.0909090 | 0.3048099 | 7 | 37848 | 11 | 0.0028751 | 17.818181 | 0 | 11 | 0.9875369 | 0.111175448755066 | A disintegrin and metalloproteinase with thrombospondin motifs 9; |
| ADAMTSL3 | P82987 | 39 | 9606.ENSP | 3.7942458 | 0.4 | 0.2635569 | 8 | 978 | 5 | 7.3476504 | 7.8 | 0 | 5 | 0.9847308 | 0.2689655172413793 | ADAMTS-like protein 3; Immunoglobulin like domain containing |
| ADAMTSL4 | Q6UY14 | 39 | 9606.ENSP | 3.8108108 | 0.5 | 0.2624113 | 8 | 1288 | 4 | 1.6458769 | 10.0 | 0 | 4 | 0.9846403 | 0.295454545454545 | Thrombospondin repeat-containing protein 1; Positive regulation of |
| THSD4 | Q6ZMP0 | 39 | 9606.ENSP | 3.6399302 | 0.2 | 0.2747305 | 7 | 14426 | 6 | 0.0010358 | 9.1666666 | 0 | 6 | 0.9855741 | 0.201550387596899 | A disintegrin and metalloproteinase with thrombospondin motifs-like |
| ADCYAP1R | P41586 | 40 | 9606.ENSP | 3.4533565 | 0.1785714 | 0.2895733 | 8 | 6748 | 8 | 5.6265906 | 17.625 | 0 | 8 | 0.9865936 | 0.190217391304347 | Adenylate cyclase activating polypeptide 1 (pituitary) receptor type I; |
| AQP1 | P29972 | 40 | 9606.ENSP | 2.9485614 | 0.1111111 | 0.3391484 | 7 | 75944 | 18 | 0.0062958 | 26.888888 | 0 | 18 | 0.9893521 | 0.0927216415021293 | Water channel protein for red blood cells and kidney proximal tubule |
| AQP5 | P55064 | 40 | 9606.ENSP | 3.0871839 | 0.1333333 | 0.3239197 | 7 | 14318 | 6 | 0.0010155 | 40.5 | 0 | 6 | 0.9885946 | 0.180180180180180 | Aquaporin 5; Forms a water-specific channel. Implicated in the gener |
| SLC14A1 | Q13336 | 40 | 9606.ENSP | 3.6878814 | 0.0 | 0.2711583 | 8 | 136 | 2 | 1.1663295 | 18.0 | 0 | 2 | 0.9853121 | 0.5 | Solute carrier family 14 (urea transporter), member 1 (Kidd blood gr |
| CDH5 | P33151 | 41 | 9606.ENSP | 2.5422842 | 0.2462025 | 0.3933470 | 6 | 314116 | 80 | 0.0155889 | 46.325 | 0 | 80 | 0.9915722 | 0.081514084507042 | Cadherin 5, type 2 (vascular endothelium); Cadherins are calcium-de |
| ERG | P11308 | 41 | 9606.ENSP | 3.5414123 | 0.0 | 0.2823732 | 7 | 0 | 1 | 0.0 | 80.0 | 0 | 1 | 0.9861125 | 0.0 | V-ets avian erythroblastosis virus E26 oncogene homolog; Transcripti |
| GJA4 | P35212 | 41 | 9606.ENSP | 3.0531822 | 0.3717948 | 0.3275271 | 7 | 6946 | 13 | 3.8723934 | 41.384615 | 0 | 13 | 0.9887804 | 0.165894346617238 | Gap junction protein, alpha 4, 37kDa; One gap junction consists of a |
| ROBO4 | Q8WZ75 | 41 | 9606.ENSP | 2.9459459 | 0.3859649 | 0.3394495 | 7 | 38336 | 19 | 0.0017834 | 47.789473 | 0 | 19 | 0.9893664 | 0.1517126148705096 | Roundabout, axon guidance receptor, homolog 4 (Drosophila); Rece |
| GXYLT1 | Q4G148 | 42 | 9606.ENSP | 3.3060156 | 0.4 | 0.3024789 | 7 | 2586 | 5 | 2.2402100 | 38.4 | 0 | 5 | 0.9873988 | 0.263013698630136 | Glycosyltransferase 8 domain-containing protein 3; Glycosyltransfera |
| NHSL1 | Q5SYE7 | 42 | 9606.ENSP | 4.3391455 | 0.0 | 0.2304601 | 8 | 0 | 1 | 0.0 | 8.0 | 0 | 1 | 0.9817533 | 0.0 | NHS-like protein 1; NHS like 1; Belongs to the NHS family. |
| STON2 | Q8WXE9 | 42 | 9606.ENSP | 3.3400174 | 0.0714285 | 0.2993996 | 7 | 37618 | 8 | 0.0030918 | 15.0 | 0 | 8 | 0.9872130 | 0.143564356435643 | Stoned B; Adapter protein involved in endocytic machinery. Involved |
| ZNF462 | Q96JM2 | 42 | 9606.ENSP | 3.4149956 | 0.3333333 | 0.2928261 | 7 | 1362 | 3 | 1.5723609 | 35.0 | 0 | 3 | 0.9868033 | 0.3466666666666667 | Zinc finger protein 462; May be involved in transcriptional regulation |
| CTNNA3 | Q9UI47 | 43 | 9606.ENSP | 3.3321708 | 0.9 | 0.3001046 | 7 | 242 | 5 | 9.2034452 | 39.4 | 0 | 5 | 0.9872558 | 0.291851851851851 | Catenin (cadherin-associated protein), alpha 3; May be involved in fo |
| DSP | P15924 | 43 | 9606.ENSP | 2.8997384 | 0.2810457 | 0.3448586 | 7 | 36260 | 18 | 0.0022546 | 44.0 | 0 | 18 | 0.9896189 | 0.131798465131798 | 250/210 kDa paraneoplastic pemphigus antigen; Major high molecul |
| PKP4 | Q99569 | 43 | 9606.ENSP | 3.2659110 | 0.1818181 | 0.3061932 | 7 | 9592 | 11 | 8.9332256 | 20.0 | 0 | 11 | 0.9876179 | 0.126686217008797 | Plakophilin 4; Plays a role as a regulator of Rho activity during cytokin |
| PPP1R13L | Q8WUF5 | 43 | 9606.ENSP | 3.2545771 | 0.3333333 | 0.3072595 | 7 | 124 | 3 | 3.5419543 | 79.333333 | 0 | 3 | 0.9876799 | 0.478787878787878 | Protein phosphatase 1, regulatory subunit 13 like; Regulator that pla |
| GALNT4 | Q8N4A0 | 44 | 9606.ENSP | 3.7061900 | 1.0 | 0.2698188 | 8 | 0 | 2 | 0.0 | 18.5 | 0 | 2 | 0.9852120 | 0.5441176470588235 | UDP-GalNAc:polypeptide N-acetylgalactosaminyltransferase 4; Catal |
| MUC1 | Q9UQL1 | 44 | 9606.ENSP | 2.7297297 | 0.3011494 | 0.3663366 | 7 | 94930 | 30 | 0.0058770 | 58.433333 | 0 | 30 | 0.9905479 | 0.128560939794419 | Tumor-associated epithelial membrane antigen; The alpha subunit h |
| SLC25A18 | Q9H1K4 | 44 | 9606.ENSP | 4.0610287 | 0.3333333 | 0.2462430 | 8 | 482 | 3 | 6.7751768 | 8.6666666 | 0 | 3 | 0.9832730 | 0.362318840579710 | Solute carrier family 25 (glutamate carrier), member 18; Involved in t |
| ST6GALNA | Q9NSC7 | 44 | 9606.ENSP | 3.5614646 | 0.1428571 | 0.2807833 | 8 | 27192 | 7 | 0.0020183 | 10.142857 | 0 | 7 | 0.9860029 | 0.169950738916256 | ST6 N-acetylgalactosaminide alpha-2,6-sialyltransferase 1; Sialyltrans |
| CCDC88A | Q3V6T2 | 45 | 9606.ENSP | 3.0217959 | 0.1428571 | 0.3309290 | 7 | 19036 | 7 | 0.0014141 | 50.285714 | 0 | 7 | 0.9889519 | 0.191023535851122 | G alpha-interacting vesicle-associated protein; Plays a role as a key m |
| CENPB | P07199 | 45 | 9606.ENSP | 3.9093286 | 0.0 | 0.2557983 | 8 | 652 | 3 | 9.5699645 | 10.0 | 0 | 3 | 0.9841020 | 0.3333333333333333 | Major centromere autoantigen B; Interacts with centromeric hetero |
| EML3 | Q32P44 | 45 | 9606.ENSP | 3.8020924 | 0.0 | 0.2630130 | 8 | 2156 | 4 | 1.8257829 | 9.5 | 0 | 4 | 0.9846880 | 0.25 | Echinoderm microtubule associated protein like 3; May modify the a |
| MAP4 | P27816 | 45 | 9606.ENSP | 3.6948561 | 0.0 | 0.2706465 | 8 | 1222 | 2 | 1.1636907 | 17.0 | 0 | 2 | 0.9852740 | 0.5 | Microtubule-associated protein 4; Non-neuronal microtubule-associa |
| CELSR1 | Q9NYQ6 | 46 | 9606.ENSP | 3.7018308 | 0.3333333 | 0.2701365 | 7 | 2078 | 4 | 1.9669690 | 14.0 | 0 | 4 | 0.9852358 | 0.280612244897959 | Cadherin, EGF LAG seven-pass G-type receptor 1; Receptor that may |
| DCHS1 | Q96JQ0 | 46 | 9606.ENSP | 3.2859633 | 0.2857142 | 0.3043247 | 7 | 6256 | 7 | 4.6340539 | 30.714285 | 0 | 7 | 0.9875083 | 0.192273135669362 | Dachsous cadherin-related 1; Calcium-dependent cell-adhesion prote |
| FZD7 | O75084 | 46 | 9606.ENSP | 2.9006102 | 0.25 | 0.3447550 | 6 | 43726 | 17 | 0.0026122 | 45.352941 | 0 | 17 | 0.9896141 | 0.132610939112487 | Frizzled class receptor 7; Receptor for Wnt proteins. Most of frizzled |
| PRICKLE3 | O43900 | 46 | 9606.ENSP | 3.8256320 | 0.3333333 | 0.2613947 | 7 | 174 | 3 | 1.3520114 | 9.3333333 | 0 | 3 | 0.9845593 | 0.391304347826087 | Prickle planar cell polarity protein 3; Involved in the planar cell polari |
| GNA12 | Q03113 | 47 | 9606.ENSP | 3.1124673 | 0.1238095 | 0.3212885 | 7 | 53542 | 21 | 0.0035261 | 21.190476 | 0 | 21 | 0.9884564 | 0.091495113329174 | Guanine nucleotide binding protein (G protein) alpha 12; Guanine nu |
| GNA14 | O95837 | 47 | 9606.ENSP | 3.3251961 | 0.2857142 | 0.3007341 | 8 | 13090 | 14 | 8.2400203 | 16.357142 | 0 | 14 | 0.9872940 | 0.1256921373200443 | Guanine nucleotide binding protein (G protein), alpha 14; Guanine nu |
| S1PR3 | Q99500 | 47 | 9606.ENSP | 3.4803836 | 0.3333333 | 0.2873246 | 8 | 266 | 3 | 8.8362258 | 32.666666 | 0 | 3 | 0.9864459 | 0.384920634920634 | Endothelial differentiation G-protein coupled receptor 3; Receptor fo |
| DDAH2 | O95865 | 48 | 9606.ENSP | 3.2850915 | 0.1333333 | 0.3044055 | 8 | 2328 | 6 | 2.0650749 | 28.0 | 0 | 6 | 0.9875131 | 0.205223880597014 | N(G),N(G)-dimethylarginine dimethylaminohydrolase 2; Hydrolyzes N |
| DYNC1LI2 | O43237 | 48 | 9606.ENSP | 3.9145597 | 0.1666666 | 0.2554565 | 8 | 30522 | 4 | 0.0027446 | 7.0 | 0 | 4 | 0.9840734 | 0.2826086956521739 | Dynein, cytoplasmic 1, light intermediate chain 2; Acts as one of seve |
| KIF5B | P33176 | 48 | 9606.ENSP | 3.0078465 | 0.0641025 | 0.3324637 | 7 | 110180 | 13 | 0.0093176 | 30.153846 | 0 | 13 | 0.9890281 | 0.114201183431952 | Conventional kinesin heavy chain; Microtubule-dependent motor re |
| NOS3 | Q548C1 | 48 | 9606.ENSP | 2.5544899 | 0.3051715 | 0.3914675 | 7 | 195684 | 63 | 0.0095345 | 53.904761 | 0 | 63 | 0.9915055 | 0.093883081687959 | Nitric oxide synthase 3 (endothelial cell); Produces nitric oxide (NO) v |
| AKR1C3 | Q9UKL9 | 49 | 9606.ENSP | 3.7863993 | 0.1 | 0.2641031 | 8 | 13470 | 5 | 0.0014458 | 9.0 | 0 | 5 | 0.9847737 | 0.21 | Trans-1,2-dihydrobenzene-1,2-diol dehydrogenase; Catalyzes the co |
| ARHGEF37 | A1IGU5 | 49 | 9606.ENSP | 4.0313862 | 0.0666666 | 0.2480536 | 8 | 10558 | 6 | 8.9950458 | 5.8333333 | 0 | 6 | 0.9834350 | 0.191358024691358 | Rho guanine nucleotide exchange factor (GEF) 37; May act as a guani |
| NOL8 | Q9H586 | 49 | 9606.ENSP | 3.7951176 | 0.0 | 0.2634964 | 7 | 2052 | 3 | 2.8049058 | 13.666666 | 0 | 3 | 0.9847261 | 0.3333333333333333 | Nucleolar protein Nop132; Plays an essential role in the survival of di |

| | | | | | | | | | | | | | | | | |
|---|---|---|---|---|---|---|---|---|---|---|---|---|---|---|---|---|
| ATP1A2 | P50993 | 50 | 9606.ENSP | 3.3374019 | 0.1111111 | 0.2996342 | 7 | 27134 | 9 | 0.0022766 | 16.888888 | 0 | 9 | 0.9872273 | 0.146509341199606 | Sodium/potassium-transporting ATPase subunit alpha-2; This is the c |
| FXYD1 | O00168 | 50 | 9606.ENSP | 3.8221447 | 0.1666666 | 0.2616332 | 8 | 1128 | 4 | 1.4516077 | 10.0 | 0 | 4 | 0.9845784 | 0.2714285714285714 | Sodium/potassium-transporting ATPase subunit FXYD1; Associates w |
| FXYD3 | Q14802 | 50 | 9606.ENSP | 4.1194420 | 0.0 | 0.2427513 | 8 | 54 | 2 | 6.4416200 | 6.5 | 0 | 2 | 0.9829538 | 0.55 | Sodium/potassium-transporting ATPase subunit FXYD3; Associates w |
| LGI4 | Q8N135 | 50 | 9606.ENSP | 3.3461203 | 0.0 | 0.2988535 | 8 | 15246 | 4 | 0.0012850 | 31.25 | 0 | 4 | 0.9871796 | 0.2520833333333333 | Leucine-rich glioma-inactivated protein 4; Leucine rich repeat LGI fan |
| HIPK2 | Q9H2X6 | 51 | 9606.ENSP | 3.1089799 | 0.1333333 | 0.3216489 | 7 | 7346 | 6 | 4.9207483 | 51.0 | 0 | 6 | 0.9884755 | 0.215602836879432 | Homeodomain interacting protein kinase 2; Serine/threonine-protein |
| NUPR1 | H3BS92 | 51 | 9606.ENSP | 4.5501307 | 0.0 | 0.2197739 | 9 | 0 | 1 | 0.0 | 5.0 | 0 | 1 | 0.9806003 | 0.0 | Nuclear protein 1, transcriptional regulator |
| SASH3 | O75995 | 51 | 9606.ENSP | 3.8936355 | 0.0 | 0.2568293 | 8 | 1314 | 3 | 1.1219901 | 14.666666 | 0 | 3 | 0.9841877 | 0.3416666666666667 | SH3 protein expressed in lymphocytes homolog; May function as a si |
| TP53INP1 | Q96A56 | 51 | 9606.ENSP | 3.5510026 | 0.0 | 0.2816106 | 8 | 24996 | 5 | 0.0019094 | 13.6 | 0 | 5 | 0.9860600 | 0.2135593220338983 | P53-dependent damage-inducible nuclear protein 1; Antiproliferative |
| ADIRF | Q15847 | 52 | 9606.ENSP | 3.9503051 | 0.0 | 0.2531450 | 8 | 626 | 2 | 3.9637097 | 13.0 | 0 | 2 | 0.9838781 | 0.5 | Adipose most abundant gene transcript 2 protein; Plays a role in fat c |
| EMID1 | Q96A84 | 52 | 9606.ENSP | 3.8212728 | 0.0 | 0.2616929 | 8 | 826 | 3 | 1.0599548 | 12.666666 | 0 | 3 | 0.9845832 | 0.3333333333333333 | Emilin and multimerin domain-containing protein 1; EMI domain con |
| MMRN2 | Q9H8L6 | 52 | 9606.ENSP | 3.4795117 | 0.0555555 | 0.2873966 | 8 | 55890 | 9 | 0.0023977 | 17.666666 | 0 | 9 | 0.9864507 | 0.1728395061728395 | Elastin microfibril interface located protein 3; Inhibits endothelial cell |
| USHBP1 | Q8N6Y0 | 52 | 9606.ENSP | 4.4786399 | 0.0 | 0.2232820 | 9 | 0 | 1 | 0.0 | 9.0 | 0 | 1 | 0.9809910 | 0.0 | Usher syndrome type-1C protein-binding protein 1; USH1 protein net |
| CAV1 | Q03135 | 53 | 9606.ENSP | 2.4202266 | 0.2147634 | 0.4131844 | 7 | 470750 | 92 | 0.0276347 | 47.097826 | 0 | 92 | 0.9922391 | 0.0705951372140017 | Caveolin 1, caveolae protein, 22kDa; May act as a scaffolding protein |
| EHD2 | Q9NZN4 | 53 | 9606.ENSP | 3.1979075 | 0.3333333 | 0.3127044 | 7 | 4036 | 7 | 3.2063062 | 35.142857 | 0 | 7 | 0.9879895 | 0.2046783625730994 | EH domain-containing protein 2; ATP- and membrane-binding protei |
| LRRC8A | Q8IWT6 | 53 | 9606.ENSP | 3.3417611 | 0.3333333 | 0.2992434 | 8 | 276 | 3 | 1.8935565 | 36.666666 | 0 | 3 | 0.9872034 | 0.3395638629283489 | Leucine rich repeat containing 8 family, member A; Essential compor |
| LONRF3 | Q496Y0 | 54 | 9606.ENSP | 3.8142981 | 0.3333333 | 0.2621714 | 8 | 670 | 3 | 4.0967756 | 14.666666 | 0 | 3 | 0.9846213 | 0.398148148148148 | LON peptidase N-terminal domain and ring finger 3 |
| PLEKHA4 | Q9H4M7 | 54 | 9606.ENSP | 3.8177855 | 0.0 | 0.2619319 | 8 | 656 | 2 | 6.3946563 | 14.0 | 0 | 2 | 0.9846022 | 0.5 | Pleckstrin homology domain containing, family A (phosphoinositide b |
| SH3TC1 | Q8TE82 | 54 | 9606.ENSP | 4.4167393 | 0.0 | 0.2264113 | 9 | 208 | 2 | 4.8373777 | 5.5 | 0 | 2 | 0.9813292 | 0.5 | SH3 domain and tetratricopeptide repeats 1 |
| SORCS2 | Q96PQ0 | 54 | 9606.ENSP | 3.7585004 | 0.0 | 0.2660635 | 8 | 10212 | 4 | 6.8656700 | 9.0 | 0 | 4 | 0.9849262 | 0.25 | Sortilin related VPS10 domain containing receptor 2; Belongs to the V |
| CSPP1 | Q1MSJ5 | 55 | 9606.ENSP | 4.3661726 | 0.0 | 0.2290335 | 9 | 5706 | 2 | 5.8058696 | 6.0 | 0 | 2 | 0.9816056 | 0.5 | Centrosome and spindle pole associated protein 1; May play a role in |
| NEK1 | Q96PY6 | 55 | 9606.ENSP | 4.6015693 | 0.0 | 0.2173171 | 9 | 3070 | 2 | 3.0354925 | 5.5 | 0 | 2 | 0.9803192 | 0.5 | Serine/threonine-protein kinase Nek1; Phosphorylates serines and th |
| NT5DC2 | Q9H857 | 55 | 9606.ENSP | 5.7750653 | 0.0 | 0.1731582 | 10 | 0 | 1 | 0.0 | 4.0 | 0 | 1 | 0.9739067 | 0.0 | 5'-nucleotidase domain containing 2 |
| ANXA13 | P27216 | 56 | 9606.ENSP | 3.2885789 | 0.3055555 | 0.3040827 | 7 | 1698 | 9 | 9.9310526 | 23.222222 | 0 | 9 | 0.9874941 | 0.176420695504664 | Annexin A13; Belongs to the annexin family. |
| ANXA2 | P07355 | 56 | 9606.ENSP | 2.5945945 | 0.2693877 | 0.3854166 | 7 | 174894 | 50 | 0.0096636 | 51.18 | 0 | 50 | 0.9912863 | 0.095270018621973 | Placental anticoagulant protein IV; Calcium-regulated membrane-bin |
| S100A10 | P60903 | 56 | 9606.ENSP | 3.0435919 | 0.3818181 | 0.3285591 | 7 | 20802 | 11 | 0.0017203 | 35.090909 | 0 | 11 | 0.9888328 | 0.149572649572649 | S100 calcium binding protein A10; Because S100A10 induces the dim |
| S100A11 | P31949 | 56 | 9606.ENSP | 3.0313862 | 0.4888888 | 0.3298820 | 7 | 2350 | 10 | 8.6213315 | 47.6 | 0 | 10 | 0.9888995 | 0.1904 | Metastatic lymph node gene 70 protein; Facilitates the differentiatior |
| KREMEN1 | Q96MU8 | 57 | 9606.ENSP | 3.5588491 | 0.3333333 | 0.2809897 | 7 | 226 | 3 | 1.2022945 | 23.0 | 0 | 3 | 0.9860172 | 0.384180790960452 | Kringle-containing protein marking the eye and the nose; Receptor fc |
| LRP4 | O75096 | 57 | 9606.ENSP | 3.1691368 | 0.2666666 | 0.3155433 | 6 | 2710 | 6 | 1.4303207 | 44.0 | 0 | 6 | 0.9881467 | 0.223639455782312 | Low density lipoprotein receptor-related protein 4; Mediates SOST-d |
| LRP5 | O75197 | 57 | 9606.ENSP | 2.7680906 | 0.1825396 | 0.3612598 | 6 | 110230 | 28 | 0.0069381 | 43.321428 | 0 | 28 | 0.9903383 | 0.101680672268907 | Low density lipoprotein receptor-related protein 5; Component of th |
| RNF43 | Q68DV7 | 57 | 9606.ENSP | 3.3984306 | 0.4 | 0.2942534 | 6 | 1660 | 5 | 7.4620383 | 28.4 | 0 | 5 | 0.9868938 | 0.2635514018691589 | RING-type E3 ubiquitin transferase RNF43; E3 ubiquitin-protein ligase |
| ASPH | Q6NXR7 | 58 | 9606.ENSP | 3.3993025 | 0.0 | 0.2941779 | 8 | 10832 | 5 | 8.9541500 | 20.6 | 0 | 5 | 0.9868890 | 0.2063157894736842 | Aspartyl/asparaginyl beta-hydroxylase; Isoform 1: specifically hydrox |
| RYR3 | Q15413 | 58 | 9606.ENSP | 3.7628596 | 0.0666666 | 0.2657553 | 8 | 45474 | 6 | 0.0038833 | 6.8333333 | 0 | 6 | 0.9849024 | 0.1761904761904762 | Brain ryanodine receptor-calcium release channel; Calcium channel t |
| TET1 | Q8NFU7 | 58 | 9606.ENSP | 3.3060156 | 0.0666666 | 0.3024789 | 7 | 6614 | 6 | 5.4541320 | 23.833333 | 0 | 6 | 0.9873988 | 0.183862433862433 | Leukemia-associated protein with a CXXC domain; Dioxygenase that |
| ITPKB | P27987 | 59 | 9606.ENSP | 3.8735832 | 0.5 | 0.2581589 | 8 | 1540 | 5 | 1.2650506 | 10.2 | 0 | 5 | 0.9842973 | 0.268421052631578 | Inositol-trisphosphate 3-kinase B |
| ITPKC | Q96DU7 | 59 | 9606.ENSP | 3.5074106 | 0.2666666 | 0.2851106 | 8 | 11336 | 6 | 7.5271795 | 18.166666 | 0 | 6 | 0.9862983 | 0.202651515151515 | Inositol 1,4,5-trisphosphate 3-kinase C; Can phosphorylate inositol 2, |
| PLCD1 | P51178 | 59 | 9606.ENSP | 3.5021795 | 0.1333333 | 0.2855364 | 7 | 27552 | 10 | 0.0021195 | 9.0 | 0 | 10 | 0.9863268 | 0.126086956521739 | 1-phosphatidylinositol 4,5-bisphosphate phosphodiesterase delta-1; |
| PLCD3 | Q8N3E9 | 59 | 9606.ENSP | 3.9485614 | 0.2380952 | 0.2532567 | 8 | 8752 | 7 | 8.7320699 | 6.7142857 | 0 | 7 | 0.9838876 | 0.189075630252100 | 1-phosphatidylinositol 4,5-bisphosphate phosphodiesterase delta-3; |
| MAPKAPK | P49137 | 60 | 9606.ENSP | 3.3461203 | 0.0833333 | 0.2988535 | 8 | 22628 | 9 | 0.0013420 | 17.222222 | 0 | 9 | 0.9871796 | 0.152525252525252 | Mitogen-activated protein kinase-activated protein kinase 2; Stress-a |
| MAPKAPK | Q16644 | 60 | 9606.ENSP | 3.5100261 | 0.3333333 | 0.2848981 | 8 | 1762 | 4 | 7.5265843 | 20.0 | 0 | 4 | 0.9862840 | 0.286231884057971 | Mitogen-activated protein kinase-activated protein kinase 3; Stress-a |
| RAB29 | O14966 | 60 | 9606.ENSP | 3.8448125 | 0.0 | 0.2600907 | 7 | 184 | 2 | 1.3446950 | 12.5 | 0 | 2 | 0.9844545 | 0.5 | RAB29, member RAS oncogene family; Rab GTPase key regulator in v |
| ZFP36L2 | P47974 | 60 | 9606.ENSP | 4.0680034 | 0.3333333 | 0.2458208 | 8 | 218 | 3 | 3.7611063 | 8.0 | 0 | 3 | 0.9832349 | 0.365079365079365 | Zinc finger protein 36, C3H1 type-like 2; Zinc-finger RNA-binding prot |
| CNN3 | Q15417 | 61 | 9606.ENSP | 3.0749782 | 0.5714285 | 0.3252055 | 7 | 11726 | 7 | 9.4439143 | 62.0 | 0 | 7 | 0.9886613 | 0.259903961584633 | Calponin, acidic isoform; Thin filament-associated protein that is imp |
| FLVCR2 | Q9UPI3 | 61 | 9606.ENSP | 3.8273757 | 0.0 | 0.2612756 | 8 | 254 | 3 | 2.6824376 | 8.6666666 | 0 | 3 | 0.9845498 | 0.3484848484848485 | Feline leukemia virus subgroup C cellular receptor family, member 2; |
| RFX4 | Q8NDF9 | 61 | 9606.ENSP | 3.8605056 | 0.1666666 | 0.2590334 | 8 | 1558 | 4 | 2.4814779 | 10.75 | 0 | 4 | 0.9843688 | 0.292857142857142 | Regulatory factor X, 4 (influences HLA class II expression); May activa |
| ETNK2 | Q9NVF9 | 62 | 9606.ENSP | 3.8221447 | 0.0666666 | 0.2616332 | 8 | 36074 | 6 | 0.0023342 | 8.1666666 | 0 | 6 | 0.9845784 | 0.234375 | Ethanolamine kinase-like protein; Highly specific for ethanolamine ph |
| SOX13 | Q9UN79 | 62 | 9606.ENSP | 3.4673060 | 0.2666666 | 0.2884083 | 7 | 6548 | 6 | 3.8143112 | 25.666666 | 0 | 6 | 0.9865174 | 0.2783882783882784 | SRY (Sex determining region Y)-box 13; Binds to the sequence 5'-AAC |
| CYBRD1 | Q53TN4 | 63 | 9606.ENSP | 3.2275501 | 0.3846153 | 0.3098325 | 7 | 13696 | 13 | 0.0011874 | 20.0 | 0 | 13 | 0.9878275 | 0.128039702233250 | Ferric-chelate reductase 3; Ferric-chelate reductase that reduces Fe( |
| HEPH | Q9BQS7 | 63 | 9606.ENSP | 3.2911944 | 0.5272727 | 0.3038410 | 7 | 14514 | 11 | 0.0010298 | 19.909090 | 0 | 11 | 0.9874798 | 0.150589868147120 | Hephaestin; May function as a ferroxidase for ferrous (II) to ferric ion |
| SLC40A1 | Q9NP59 | 63 | 9606.ENSP | 3.0549258 | 0.3 | 0.3273401 | 7 | 41038 | 16 | 0.0034236 | 26.1875 | 0 | 16 | 0.9887708 | 0.1015625 | Solute carrier family 40 (iron-regulated transporter), member 1; May |
| ABHD15 | Q6UXT9 | 64 | 9606.ENSP | 5.6739319 | 0.0 | 0.1762446 | 10 | 0 | 1 | 0.0 | 2.0 | 0 | 1 | 0.9744593 | 0.0 | Alpha/beta hydrolase domain-containing protein 15; Abhydrolase do |
| ABHD4 | Q8TB40 | 64 | 9606.ENSP | 4.6748038 | 0.0 | 0.2139127 | 9 | 26686 | 2 | 0.0017436 | 2.0 | 0 | 2 | 0.9799191 | 0.5 | Alpha/beta hydrolase domain-containing protein 4; Lysophospholipa |
| KANK3 | Q6NY19 | 64 | 9606.ENSP | 3.6774193 | 0.0 | 0.2719298 | 8 | 53482 | 3 | 0.0034910 | 16.0 | 0 | 3 | 0.9853692 | 0.3333333333333333 | KN motif and ankyrin repeat domain-containing protein 3; May be in |
| HMG20B | Q9P0W2 | 65 | 9606.ENSP | 3.6006974 | 0.0833333 | 0.2777239 | 8 | 28182 | 9 | 0.0025075 | 8.5555555 | 0 | 9 | 0.9857885 | 0.135185185185185 | Structural DNA-binding protein BRAF35; Required for correct progres |
| PHF21B | Q96EK2 | 65 | 9606.ENSP | 3.8761987 | 0.1666666 | 0.2579847 | 9 | 1890 | 4 | 2.6763447 | 12.0 | 0 | 4 | 0.9842830 | 0.273809523809523 | PHD finger protein 21B |
| ITPR2 | Q14571 | 66 | 9606.ENSP | 3.3591979 | 0.0909090 | 0.2976901 | 8 | 50486 | 11 | 0.0043470 | 14.727272 | 0 | 11 | 0.9871082 | 0.1305785123966942 | Inositol 1,4,5-trisphosphate receptor, type 2; Receptor for inositol 1, |
| ORAI3 | Q9BRQ5 | 66 | 9606.ENSP | 4.3583260 | 0.0 | 0.2294458 | 9 | 0 | 1 | 0.0 | 11.0 | 0 | 1 | 0.9816484 | 0.0 | ORAI calcium release-activated calcium modulator 3; Key regulator o |
| CRX | O43186 | 67 | 9606.ENSP | 3.1517000 | 0.1142857 | 0.3172890 | 7 | 99970 | 15 | 0.0090678 | 17.066666 | 0 | 15 | 0.9882420 | 0.086355785837651 | Cone-rod homeobox protein; Transcription factor that binds and trar |
| ROM1 | Q03395 | 67 | 9606.ENSP | 3.3984306 | 0.2857142 | 0.2942534 | 8 | 3386 | 7 | 2.6430878 | 16.285714 | 0 | 7 | 0.9868938 | 0.1717325227963526 | Retinal outer segment membrane protein 1; May function as an adhe |
| TSPAN4 | O14817 | 67 | 9606.ENSP | 3.2432432 | 0.1333333 | 0.3083333 | 7 | 2456 | 6 | 1.4454293 | 41.333333 | 0 | 6 | 0.9877418 | 0.2536231884057971 | Transmembrane 4 superfamily member 7; Tetraspanin 4; Belongs to |
| FAM189A | Q15884 | 68 | 9606.ENSP | 3.9904097 | 0.0 | 0.2506008 | 8 | 194 | 2 | 1.0072276 | 6.0 | 0 | 2 | 0.9836589 | 0.5 | Family with sequence similarity 189 member A2 |
| SNX33 | Q8WV41 | 68 | 9606.ENSP | 3.6285963 | 0.1904761 | 0.2755886 | 8 | 5718 | 7 | 6.2675656 | 14.285714 | 0 | 7 | 0.9856360 | 0.216517857142857 | SH3 and PX domain-containing protein 3; Plays a role in the reorganiz |
| TSPO | P30536 | 68 | 9606.ENSP | 3.0932868 | 0.3 | 0.3232807 | 7 | 16950 | 5 | 0.0016220 | 61.0 | 0 | 5 | 0.9885612 | 0.2623376623376623 | Peripheral-type benzodiazepine receptor; Can bind protoporphyrin I |
| ANKRD40 | Q6AI12 | 69 | 9606.ENSP | 4.4760244 | 0.0 | 0.2234125 | 7 | 796 | 2 | 1.5062107 | 6.5 | 0 | 2 | 0.9810053 | 0.5 | Ankyrin repeat domain-containing protein 40; Ankyrin repeat domair |
| PAQR6 | Q8N3M2 | 69 | 9606.ENSP | 4.1935483 | 0.0 | 0.2384615 | 7 | 21926 | 3 | 0.0019695 | 4.6666666 | 0 | 3 | 0.9825489 | 0.3333333333333333 | Progesterone and adipoQ receptor family member 6; Plasma membr |
| ESAM | Q96AP7 | 70 | 9606.ENSP | 3.1211857 | 0.5274725 | 0.3203910 | 7 | 7546 | 14 | 2.2890447 | 51.285714 | 0 | 14 | 0.9884088 | 0.223955084217092 | Endothelial cell-selective adhesion molecule; Can mediate aggregatio |
| SLC44A2 | Q9NY68 | 70 | 9606.ENSP | 3.5135135 | 0.0 | 0.2846153 | 7 | 0 | 1 | 0.0 | 81.0 | 0 | 1 | 0.9862649 | 0.0 | Solute carrier family 44 (choline transporter), member 2; Isoform 1, |
| VWF | P04275 | 70 | 9606.ENSP | 2.5143853 | 0.2404320 | 0.3977115 | 6 | 299732 | 81 | 0.0154248 | 46.827160 | 0 | 81 | 0.9917246 | 0.078792035582159 | Von Willebrand factor; Important in the maintenance of hemostasis, |
| ARSI | Q5FYB1 | 71 | 9606.ENSP | 4.2746294 | 0.0 | 0.2339384 | 8 | 23968 | 4 | 0.0018994 | 3.75 | 0 | 4 | 0.9821058 | 0.25 | Arylsulfatase family, member I; Displays arylsulfatase activity at neutr |
| ST8SIA6 | P61647 | 71 | 9606.ENSP | 5.2737576 | 0.0 | 0.1896181 | 9 | 0 | 1 | 0.0 | 4.0 | 0 | 1 | 0.9766461 | 0.0 | ST8 alpha-N-acetyl-neuraminide alpha-2,8-sialyltransferase 6; Prefers |
| CRACR2B | Q8N4Y2 | 72 | 9606.ENSP | 4.1246730 | 0.0 | 0.2424434 | 8 | 540 | 2 | 9.1999973 | 12.0 | 0 | 2 | 0.9829252 | 0.5 | Calcium release-activated calcium channel regulator 2B; Plays a role i |
| S100A16 | Q96FQ6 | 72 | 9606.ENSP | 4.8012205 | 0.0 | 0.2082803 | 9 | 0 | 1 | 0.0 | 4.0 | 0 | 1 | 0.9792283 | 0.0 | Aging-associated gene 13 protein; Calcium-binding protein. Binds one |
| S100A3 | P33764 | 72 | 9606.ENSP | 3.8020924 | 0.0 | 0.2630130 | 8 | 21584 | 4 | 0.0019315 | 6.5 | 0 | 4 | 0.9846880 | 0.275 | S100 calcium binding protein A3; Binds both calcium and zinc. May be |
| APPL2 | Q8NEU8 | 73 | 9606.ENSP | 3.3182214 | 0.0 | 0.3013662 | 7 | 1774 | 2 | 1.4904765 | 71.5 | 0 | 2 | 0.9873321 | 0.5 | Adaptor protein, phosphotyrosine interaction, PH domain and leucin |
| RAB31 | Q13636 | 73 | 9606.ENSP | 3.1176983 | 0.3333333 | 0.3207494 | 7 | 3034 | 4 | 2.2606475 | 65.75 | 0 | 4 | 0.9884278 | 0.304651162790697 | RAB31, member RAS oncogene family; The small GTPases Rab are ke |
| RIN3 | Q8TB24 | 73 | 9606.ENSP | 3.4943330 | 0.1785714 | 0.2861776 | 8 | 8190 | 8 | 6.3547058 | 12.875 | 0 | 8 | 0.9863697 | 0.145114942528735 | Ras interaction/interference protein 3; Ras effector protein that func |
| CGNL1 | Q0VF96 | 74 | 9606.ENSP | 3.5292066 | 0.1666666 | 0.2833498 | 8 | 12854 | 4 | 9.5281531 | 18.75 | 0 | 4 | 0.9861791 | 0.2644927536231884 | Junction-associated coiled-coil protein; May be involved in anchoring |
| CSRP1 | P21291 | 74 | 9606.ENSP | 3.5701830 | 0.0 | 0.2800976 | 7 | 13098 | 4 | 8.6506310 | 21.0 | 0 | 4 | 0.9859552 | 0.2898550724637681 | Cysteine and glycine-rich protein 1; Could play a role in neuronal dev |
| MRO | Q9BYG7 | 74 | 9606.ENSP | 4.3461203 | 0.0 | 0.2300902 | 8 | 56 | 2 | 6.0220790 | 4.0 | 0 | 2 | 0.9817151 | 0.5 | Male-specific transcription in the developing reproductive organs; M |
| GPR39 | O43194 | 75 | 9606.ENSP | 3.1081081 | 0.0476190 | 0.3217391 | 7 | 20292 | 7 | 0.0019346 | 31.571428 | 0 | 7 | 0.9884802 | 0.149068322981366 | G protein-coupled receptor 39; Zn(2+) acts as an agonist. This recept |
| GPR4 | P46093 | 75 | 9606.ENSP | 3.5544899 | 0.0 | 0.2813343 | 8 | 654 | 3 | 4.9471031 | 30.333333 | 0 | 3 | 0.9860410 | 0.341085271317829 | G-protein coupled receptor 19; Proton-sensing receptor coupled to s |
| NMB | P08949 | 75 | 9606.ENSP | 3.3513513 | 0.3 | 0.2983870 | 8 | 1690 | 5 | 1.2173561 | 29.0 | 0 | 5 | 0.9871510 | 0.232258064516129 | Neuromedin B; Stimulates smooth muscle contraction in a manner si |
| ARHGEF5 | Q12774 | 76 | 9606.ENSP | 3.8108108 | 0.0 | 0.2624113 | 8 | 9802 | 4 | 7.0886153 | 13.0 | 0 | 4 | 0.9846403 | 0.2727272727272727 | Rho guanine nucleotide exchange factor (GEF) 5; Guanine nucleotide |
| OPHN1 | O60890 | 76 | 9606.ENSP | 3.6399302 | 0.0 | 0.2747305 | 8 | 31140 | 7 | 0.0027590 | 7.5714285 | 0 | 7 | 0.9855741 | 0.142857142857142 | Oligophrenin 1; Stimulates GTP hydrolysis of members of the Rho far |
| PAMR1 | Q6UXH9 | 76 | 9606.ENSP | 4.6390584 | 0.0 | 0.2155609 | 9 | 0 | 1 | 0.0 | 7.0 | 0 | 1 | 0.9801144 | 0.0 | Peptidase domain-containing protein associated with muscle regener |
| ARHGEF10 | O15013 | 77 | 9606.ENSP | 3.8857890 | 0.0 | 0.2573479 | 8 | 6830 | 6 | 8.6298699 | 6.1666666 | 0 | 6 | 0.9842306 | 0.172222222222222 | Rho guanine nucleotide exchange factor (GEF) 10; May play a role in |
| ARHGEF19 | Q8IW93 | 77 | 9606.ENSP | 3.7384481 | 0.0 | 0.2674906 | 8 | 2218 | 2 | 1.1278647 | 19.0 | 0 | 2 | 0.9850358 | 0.5142857142857142 | Rho guanine nucleotide exchange factor (GEF) 19; Acts as guanine nu |
| SLC2A4RG | Q9NR83 | 77 | 9606.ENSP | 3.5710549 | 0.0 | 0.2800292 | 7 | 3520 | 3 | 2.5713120 | 18.333333 | 0 | 3 | 0.9859505 | 0.3466666666666667 | Huntington disease gene regulatory region-binding protein 1; Transcr |
| GPR137B | O60478 | 78 | 9606.ENSP | 4.4952048 | 0.0 | 0.2224592 | 7 | 564 | 3 | 1.3207381 | 3.3333333 | 0 | 3 | 0.9809005 | 0.3333333333333333 | G protein-coupled receptor 137B; Belongs to the GPR137 family. |
| GPRC5C | Q9NQ84 | 78 | 9606.ENSP | 3.9581517 | 0.0 | 0.2526431 | 8 | 3896 | 3 | 4.3071159 | 6.0 | 0 | 3 | 0.9838352 | 0.3333333333333333 | G protein-coupled receptor, class C, group 5, member C; This retinoic |
| PAQR5 | Q9NXK6 | 78 | 9606.ENSP | 4.1081081 | 0.0 | 0.2434210 | 7 | 4022 | 3 | 4.5350358 | 7.0 | 0 | 3 | 0.9830158 | 0.3333333333333333 | Progesterone and adipoQ receptor family member 5; Plasma membr |
| NXNL1 | Q96CM4 | 79 | 9606.ENSP | 4.0296425 | 0.0 | 0.2481609 | 8 | 40892 | 3 | 0.0035687 | 8.0 | 0 | 3 | 0.9834445 | 0.3333333333333333 | Nucleoredoxin-like protein 1; May play a role in cone cell viability, slo |
| VSX1 | Q9NZR4 | 79 | 9606.ENSP | 5.0270270 | 0.0 | 0.1989247 | 9 | 20116 | 2 | 0.0017436 | 2.0 | 0 | 2 | 0.9779943 | 0.5 | Retinal inner nuclear layer homeobox protein; Binds to the 37-bp cor |
| ZMAT1 | Q5H9K5 | 79 | 9606.ENSP | 6.0261551 | 0.0 | 0.1659432 | 10 | 0 | 1 | 0.0 | 2.0 | 0 | 1 | 0.9725346 | 0.0 | Zinc finger matrin-type 1 |
| BMPR1B | O00238 | 80 | 9606.ENSP | 3.0453356 | 0.275 | 0.3283710 | 7 | 41036 | 16 | 0.0020731 | 38.0625 | 0 | 16 | 0.9888233 | 0.146394230769230 | Bone morphogenetic protein receptor, type IB; On ligand binding, fo |
| GDF11 | O95390 | 80 | 9606.ENSP | 3.7707061 | 0.3333333 | 0.2652023 | 8 | 138 | 3 | 9.1034999 | 17.333333 | 0 | 3 | 0.9848595 | 0.4146341463414634 | Growth differentiation factor 11; Secreted signal that acts globally to |
| SMAD6 | O43541 | 80 | 9606.ENSP | 2.9119442 | 0.2964426 | 0.3434131 | 7 | 119672 | 23 | 0.0069681 | 42.391304 | 0 | 23 | 0.9895522 | 0.120993788819875 | Mothers against decapentaplegic homolog 6; Acts as a mediator of T |
| CCDC102A | Q96A19 | 81 | 9606.ENSP | 4.3522231 | 0.0 | 0.2297676 | 9 | 386 | 3 | 4.2422352 | 4.0 | 0 | 3 | 0.9816818 | 0.375 | Coiled-coil domain containing 102A |
| CDC42EP4 | Q9H3Q1 | 81 | 9606.ENSP | 3.4751525 | 0.0 | 0.2877571 | 8 | 15108 | 5 | 0.0012696 | 17.8 | 0 | 5 | 0.9864745 | 0.212658227848101 | CDC42 effector protein (Rho GTPase binding) 4; Probably involved in |
| METRN | Q9UJH8 | 81 | 9606.ENSP | 3.9851787 | 0.0 | 0.2509297 | 8 | 2380 | 3 | 1.8413327 | 6.3333333 | 0 | 3 | 0.9836875 | 0.355555555555555 | Meteorin, glial cell differentiation regulator; Involved in both glial cel |
| EMCN | Q9ULC0 | 82 | 9606.ENSP | 3.1734960 | 0.5 | 0.3151098 | 7 | 14670 | 12 | 4.8807541 | 46.833333 | 0 | 12 | 0.9881229 | 0.2247596153846154 | Gastric cancer antigen Ga34; Endothelial sialomucin, also called endo |
| ZNF366 | Q8N895 | 82 | 9606.ENSP | 4.3888404 | 0.0 | 0.2278506 | 9 | 0 | 1 | 0.0 | 10.0 | 0 | 1 | 0.9814817 | 0.0 | Dendritic cell-specific transcript protein; Has transcriptional repressio |
| BAG3 | O95817 | 83 | 9606.ENSP | 3.0366172 | 0.1105263 | 0.3293138 | 7 | 54022 | 20 | 0.0038700 | 21.35 | 0 | 20 | 0.9888709 | 0.0854251012145749 | BAG family molecular chaperone regulator 3; Co-chaperone for HSP7 |
| HSPA6 | P17066 | 83 | 9606.ENSP | 3.3731473 | 0.2857142 | 0.2964590 | 8 | 2226 | 7 | 1.7740868 | 21.714285 | 0 | 7 | 0.9870319 | 0.215714285714285 | Heat shock 70kDa protein 6 (HSP70B'); Molecular chaperone implicat |
| HSPB7 | Q9UBY9 | 83 | 9606.ENSP | 3.5815170 | 0.6666666 | 0.2792112 | 8 | 254 | 4 | 1.8543381 | 18.25 | 0 | 4 | 0.9858933 | 0.337962962962962 | Heat shock protein family B member 7 |
| ARHGAP4 | A6NI28 | 84 | 9606.ENSP | 3.8369659 | 0.0666666 | 0.2606225 | 8 | 16464 | 6 | 0.0013778 | 7.8333333 | 0 | 6 | 0.9844974 | 0.183760683760683 | Rho GTPase-activating protein 10-like; May influence blood pressure |
| PSD4 | Q8NDX1 | 84 | 9606.ENSP | 4.4210985 | 0.0 | 0.2261881 | 9 | 332 | 2 | 5.3542738 | 6.0 | 0 | 2 | 0.9813054 | 0.5 | Exchange factor for ADP-ribosylation factor guanine nucleotide facto |
| SASH1 | O94885 | 84 | 9606.ENSP | 4.2606800 | 0.0 | 0.2347043 | 8 | 474 | 2 | 6.3626295 | 6.5 | 0 | 2 | 0.9821820 | 0.5 | Proline-glutamate repeat-containing protein; May have a role in a sig |
| APOLD1 | Q96LR9 | 85 | 9606.ENSP | 4.4812554 | 0.0 | 0.2231517 | 9 | 0 | 1 | 0.0 | 6.0 | 0 | 1 | 0.9809767 | 0.0 | Apolipoprotein L domain-containing protein 1; May be involved in an |
| MKNK2 | Q9HBH9 | 85 | 9606.ENSP | 3.4821272 | 0.0 | 0.2871807 | 8 | 52964 | 6 | 0.0035669 | 15.166666 | 0 | 6 | 0.9864364 | 0.188888888888888 | MAP kinase-interacting serine/threonine-protein kinase 2; Serine/thr |
| SZRD1 | Q7Z422 | 85 | 9606.ENSP | 4.4812554 | 0.0 | 0.2231517 | 9 | 0 | 1 | 0.0 | 6.0 | 0 | 1 | 0.9809767 | 0.0 | Putative MAPK-activating protein PM18/PM20/PM22; SUZ RNA bindi |
| NXT2 | Q9Y3M4 | 86 | 9606.ENSP | 4.3626852 | 0.3333333 | 0.2292166 | 9 | 1798 | 3 | 2.4710225 | 6.3333333 | 0 | 3 | 0.9816246 | 0.375 | Nuclear transport factor 2-like export factor 2; Regulator of protein e |
| TMEM164 | Q5U3C3 | 86 | 9606.ENSP | 4.7419354 | 1.0 | 0.2108843 | 9 | 0 | 2 | 0.0 | 4.5 | 0 | 2 | 0.9795522 | 0.6428571428571429 | Transmembrane protein 164; Belongs to the TMEM164 family. |
| DHTKD1 | Q96HY7 | 87 | 9606.ENSP | 3.6006974 | 0.0833333 | 0.2777239 | 8 | 11242 | 9 | 0.0014956 | 11.0 | 0 | 9 | 0.9857885 | 0.135327635327635 | Probable 2-oxoglutarate dehydrogenase E1 component DHKTD1, mit |
| ELOVL7 | A1L3X0 | 87 | 9606.ENSP | 4.1325196 | 0.0 | 0.2419831 | 9 | 228 | 2 | 2.5017049 | 10.5 | 0 | 2 | 0.9828824 | 0.5277777777777778 | Elongation of very long chain fatty acids protein 7; Catalyzes the first |
| FADS2 | O95864 | 87 | 9606.ENSP | 3.2615518 | 0.1666666 | 0.3066025 | 8 | 78908 | 12 | 0.0057645 | 17.166666 | 0 | 12 | 0.9876417 | 0.119642857142857 | Delta(6) fatty acid desaturase; Component of a lipid metabolic pathw |
| GOLGA4 | Q13439 | 88 | 9606.ENSP | 3.8090671 | 0.0 | 0.2625314 | 7 | 23152 | 6 | 0.0015389 | 9.3333333 | 0 | 6 | 0.9846499 | 0.1773049645390071 | Golgin subfamily A member 4; May play a role in delivery of transpor |
| GOLIM4 | O00461 | 88 | 9606.ENSP | 3.8823016 | 0.0 | 0.2575791 | 8 | 1128 | 2 | 7.7983974 | 9.5 | 0 | 2 | 0.9842497 | 0.5 | Golgi-localized phosphoprotein of 130 kDa; Plays a role in endosome |
| MDFIC | Q9P1T7 | 88 | 9606.ENSP | 3.7201394 | 0.0 | 0.2688071 | 7 | 2694 | 4 | 2.1054311 | 11.0 | 0 | 4 | 0.9851358 | 0.2564102564102564 | MyoD family inhibitor domain-containing protein; Acts as a transcript |
| ARRDC4 | Q8NCT1 | 89 | 9606.ENSP | 3.8544027 | 0.0 | 0.2594435 | 8 | 0 | 1 | 0.0 | 22.0 | 0 | 1 | 0.9844021 | 0.0 | Arrestin domain-containing protein 4; Functions as an adapter recrui |
| NEDD4 | P46934 | 89 | 9606.ENSP | 2.8552746 | 0.0995670 | 0.3502290 | 7 | 108846 | 22 | 0.0083545 | 33.818181 | 0 | 22 | 0.9898618 | 0.091530054644808 | Neural precursor cell expressed, developmentally down-regulated 4, |
| WBP1 | Q96G27 | 89 | 9606.ENSP | 3.5736704 | 0.0 | 0.2798243 | 8 | 104 | 2 | 3.4104063 | 37.0 | 0 | 2 | 0.9859362 | 0.5625 | WW domain binding protein 1; WBP1/VOPP1 family |
| GADD45G | O95257 | 90 | 9606.ENSP | 3.1944202 | 0.4 | 0.3130458 | 7 | 26214 | 6 | 0.0018836 | 49.0 | 0 | 6 | 0.9880086 | 0.244166666666666 | Growth arrest and DNA damage-inducible protein GADD45 gamma; I |
| NPAS3 | Q8IXF0 | 90 | 9606.ENSP | 4.1830863 | 0.0 | 0.2390579 | 8 | 2300 | 2 | 1.4654519 | 4.0 | 0 | 2 | 0.9826060 | 0.5 | Class E basic helix-loop-helix protein 12; May play a broad role in neu |
| DPEP2 | Q9H4A9 | 91 | 9606.ENSP | 3.9921534 | 0.0 | 0.2504913 | 7 | 38 | 2 | 6.0044243 | 8.0 | 0 | 2 | 0.9836494 | 0.5 | Dipeptidase 2; Probable metalloprotease which hydrolyzes leukotrier |
| GGT5 | P36269 | 91 | 9606.ENSP | 3.4210985 | 0.1555555 | 0.2923037 | 7 | 10624 | 10 | 8.5725813 | 13.0 | 0 | 10 | 0.9867699 | 0.14 | Gamma-glutamyl transpeptidase-related enzyme; Cleaves the gamma |
| HYAL2 | Q12891 | 91 | 9606.ENSP | 3.0985178 | 0.4 | 0.3227349 | 7 | 21404 | 6 | 0.0014087 | 84.0 | 0 | 6 | 0.9885326 | 0.3545197740112994 | Hyaluronoglucosaminidase 2; Hydrolyzes high molecular weight hyalu |
| ITPRIP | Q8IWB1 | 92 | 9606.ENSP | 5.3897122 | 0.0 | 0.1855386 | 8 | 54756 | 2 | 0.0034843 | 2.5 | 0 | 2 | 0.9760125 | 0.5 | Inositol 1,4,5-trisphosphate receptor interacting protein; Enhances C |
| ITPRIPL2 | Q3MIP1 | 92 | 9606.ENSP | 7.3862249 | 0.0 | 0.1353871 | 10 | 0 | 1 | 0.0 | 2.0 | 0 | 1 | 0.9651025 | 0.0 | Inositol 1,4,5-trisphosphate receptor interacting protein-like 2; ITPRII |
| TMEM102 | Q8N9M5 | 92 | 9606.ENSP | 6.3870967 | 0.0 | 0.1565656 | 9 | 27380 | 2 | 0.0017436 | 1.5 | 0 | 2 | 0.9705623 | 0.5 | Common beta-chain associated protein; Selectively involved in CSF2 |

| | | | | | | | | | | | | | | | |
|---|---|---|---|---|---|---|---|---|---|---|---|---|---|---|---|
| EIF4EBP1 | Q13541 | 93 | 9606.ENSP | 2.8256320 | 0.2571428 | 0.3539031 | 7 | 96568 | 21 | 0.0060349 | 48.285714 | 0 | 21 | 0.9900238 | 0.1233710875654609 | Phosphorylated heat- and acid-stable protein regulated by insulin 1; |
| EIF4EBP2 | Q13542 | 93 | 9606.ENSP | 3.5579773 | 0.1428571 | 0.2810585 | 8 | 14848 | 7 | 0.0014012 | 16.285714 | 0 | 7 | 0.9860219 | 0.1686930091185410 | Eukaryotic translation initiation factor 4E binding protein 2; Represso |
| LEF1 | Q9UJU2 | 94 | 9606.ENSP | 2.6146469 | 0.2075471 | 0.3824608 | 6 | 199476 | 53 | 0.0117480 | 38.962264 | 0 | 53 | 0.9911767 | 0.0741778975741239 | T cell-specific transcription factor 1-alpha; Participates in the Wnt sig |
| TCF7 | P36402 | 94 | 9606.ENSP | 2.8927637 | 0.2307692 | 0.3456901 | 6 | 45862 | 27 | 0.0027612 | 31.111111 | 0 | 27 | 0.9896570 | 0.0913943355119825 | Transcription factor 7 (T-cell specific, HMG-box); Transcriptional activ |
| TCF7L1 | Q9HCS4 | 94 | 9606.ENSP | 3.0270270 | 0.3006535 | 0.3303571 | 6 | 20668 | 18 | 0.0013060 | 28.5 | 0 | 18 | 0.9889233 | 0.1126543209876543 | Transcription factor 7-like 1 (T-cell specific, HMG-box); Participates in |
| HMGN5 | P82970 | 95 | 9606.ENSP | 4.0183086 | 0.0 | 0.2488609 | 8 | 468 | 3 | 7.3773866 | 8.0 | 0 | 3 | 0.9835065 | 0.35 | High mobility group nucleosome-binding domain-containing protein |
| STC1 | P52823 | 95 | 9606.ENSP | 3.2502179 | 0.0952380 | 0.3076716 | 7 | 23204 | 7 | 0.0018769 | 27.714285 | 0 | 7 | 0.9877037 | 0.1635111876075731 | Stanniocalcin 1; Stimulates renal phosphate reabsorption, and could |
| RASL12 | Q9NYN1 | 96 | 9606.ENSP | 3.9590235 | 0.0 | 0.2525875 | 8 | 20630 | 3 | 0.0018070 | 5.6666666 | 0 | 3 | 0.9838304 | 0.3333333333333333 | Ras-like protein family member 12; RAS type GTPase family |
| SOD3 | P08294 | 96 | 9606.ENSP | 3.0409764 | 0.3333333 | 0.3288417 | 7 | 42004 | 9 | 0.0032665 | 55.888888 | 0 | 9 | 0.9888471 | 0.2028282828282828 | Extracellular superoxide dismutase [Cu-Zn]; Protect the extracellular |
| ZNF853 | P0CG23 | 96 | 9606.ENSP | 4.9581517 | 0.0 | 0.2016880 | 9 | 0 | 1 | 0.0 | 3.0 | 0 | 1 | 0.9783707 | 0.0 | Zinc finger protein 853; Zinc fingers C2H2-type |
| SRGN | P10124 | 97 | 9606.ENSP | 2.9163034 | 0.25 | 0.3428998 | 7 | 35584 | 17 | 0.0020696 | 42.0 | 0 | 17 | 0.9895283 | 0.1246498599439775 | Secretory granule proteoglycan core protein; Plays a role in formatio |
| VSIG4 | Q9Y279 | 97 | 9606.ENSP | 3.2441150 | 0.2163742 | 0.3082504 | 7 | 52144 | 19 | 0.0039484 | 15.631578 | 0 | 19 | 0.9877370 | 0.1021468144044321 | V-set and immunoglobulin domain-containing protein 4; Phagocytic |
| COLCA2 | A8K830 | 98 | 9606.ENSP | 4.4551002 | 0.0 | 0.2244618 | 9 | 138 | 2 | 1.7133011 | 3.5 | 0 | 2 | 0.9811196 | 0.5 | Cancer susceptibility candidate protein 13; Colorectal cancer associat |
| GPR143 | P51810 | 98 | 9606.ENSP | 3.7262423 | 0.0 | 0.2683668 | 8 | 25384 | 4 | 0.0022090 | 9.75 | 0 | 4 | 0.9851025 | 0.25 | G protein-coupled receptor 143; Receptor for tyrosine, L-DOPA and |
| RHPN2 | Q8IUC4 | 98 | 9606.ENSP | 3.6460331 | 0.0 | 0.2742706 | 8 | 16256 | 3 | 0.0010953 | 14.666666 | 0 | 3 | 0.9855408 | 0.3504273504273504 | Rhophilin, Rho GTPase binding protein 2; Binds specifically to GTP-Rh |
| GIMAP2 | Q9UG22 | 99 | 9606.ENSP | 4.0828247 | 0.0 | 0.2449284 | 8 | 336 | 2 | 4.9783423 | 9.5 | 0 | 2 | 0.9831539 | 0.5 | Immunity-associated protein 2; The heterodimer formed by GIMAP2 |
| GIMAP7 | Q8NHV1 | 99 | 9606.ENSP | 3.7401918 | 0.1428571 | 0.2673659 | 7 | 5142 | 7 | 5.6616507 | 7.1428571 | 0 | 7 | 0.9850262 | 0.1758241758241758 | Immunity-associated nucleotide 7 protein; The dimer has GTPase act |
| NFATC4 | Q96H68 | 99 | 9606.ENSP | 3.0043591 | 0.2380952 | 0.3328496 | 7 | 14224 | 7 | 0.0011614 | 52.0 | 0 | 7 | 0.9890472 | 0.1956873315363881 | Nuclear factor of activated T-cells, cytoplasmic, calcineurin-depender |
| GDPD2 | Q9HCC8 | 100 | 9606.ENSP | 4.3574542 | 0.0 | 0.2294917 | 8 | 5334 | 2 | 3.1935467 | 7.0 | 0 | 2 | 0.9816532 | 0.5 | Glycerophosphodiester phosphodiesterase domain-containing protei |
| RAB30 | Q15771 | 100 | 9606.ENSP | 4.7297297 | 0.0 | 0.2114285 | 8 | 870 | 3 | 9.5520150 | 3.3333333 | 0 | 3 | 0.9796189 | 0.3888888888888889 | RAB30, member RAS oncogene family; The small GTPases Rab are ke |
| RAB41 | Q5JT25 | 100 | 9606.ENSP | 4.1996512 | 0.0 | 0.2381150 | 9 | 5760 | 2 | 3.2438765 | 14.5 | 0 | 2 | 0.9825155 | 0.5192307692307693 | RAB41, member RAS oncogene family; Required for normal Golgi rib |
| MFSD2A | Q8NA29 | 101 | 9606.ENSP | 3.5518744 | 0.0 | 0.2815414 | 8 | 8262 | 4 | 5.4177412 | 16.75 | 0 | 4 | 0.9860553 | 0.2540322580645161 | Major facilitator superfamily domain-containing protein 2A; Sodium- |
| SLC19A3 | Q9BZV2 | 101 | 9606.ENSP | 4.1011333 | 0.0 | 0.2438350 | 9 | 1890 | 5 | 3.1099247 | 6.0 | 0 | 5 | 0.9830539 | 0.2 | Solute carrier family 19 (thiamine transporter), member 3; Mediates |
| SLC52A3 | Q9NQ40 | 101 | 9606.ENSP | 4.2074978 | 0.0 | 0.2376709 | 8 | 576 | 2 | 7.5638393 | 8.0 | 0 | 2 | 0.9824726 | 0.5 | Solute carrier family 52 (riboflavin transporter), member 3; Transpor |
| SLC16A9 | Q7RTY1 | 102 | 9606.ENSP | 3.8788142 | 0.1666666 | 0.2578107 | 8 | 1058 | 4 | 1.1101999 | 8.75 | 0 | 4 | 0.9842687 | 0.275 | Solute carrier family 16, member 9; Proton-linked monocarboxylate |
| SLC26A2 | P50443 | 102 | 9606.ENSP | 3.6695727 | 0.0476190 | 0.2725112 | 8 | 8312 | 7 | 8.4393866 | 8.2857142 | 0 | 7 | 0.9854121 | 0.1545189504373177 | Solute carrier family 26 (anion exchanger), member 2; Sulfate transp |
| UNC93B1 | Q9H1C4 | 102 | 9606.ENSP | 3.3016564 | 0.4 | 0.3028782 | 7 | 8834 | 6 | 6.0489270 | 42.166666 | 0 | 6 | 0.9874226 | 0.2716450216450216 | Unc-93 homolog B1 (C. elegans); Plays an important role in innate an |
| DAP | P51397 | 103 | 9606.ENSP | 3.6094158 | 0.0 | 0.2770531 | 8 | 0 | 1 | 0.0 | 40.0 | 0 | 1 | 0.9857408 | 0.0 | Death-associated protein 1; Negative regulator of autophagy. Involve |
| HSPB1 | P04792 | 103 | 9606.ENSP | 2.6102877 | 0.2166666 | 0.3830995 | 7 | 152246 | 40 | 0.0083518 | 51.4 | 0 | 40 | 0.9912006 | 0.0954460966542750 | Estrogen-regulated 24 kDa protein; Small heat shock protein which fu |
| HSPB2 | Q16082 | 103 | 9606.ENSP | 2.7733217 | 0.2490118 | 0.3605784 | 7 | 57054 | 23 | 0.0027236 | 54.391304 | 0 | 23 | 0.9903097 | 0.1248375812093953 | Heat shock 27kDa protein 2; May regulate the kinase DMPK; Small he |
| NPL | Q9BXD5 | 104 | 9606.ENSP | 3.9790758 | 0.0 | 0.2513146 | 8 | 1042 | 3 | 1.4223669 | 10.666666 | 0 | 3 | 0.9837208 | 0.3333333333333333 | N-acetylneuraminate pyruvate lyase (dihydrodipicolinate synthase); |
| PITPNC1 | Q9UKF7 | 104 | 9606.ENSP | 3.6774193 | 0.0 | 0.2719298 | 8 | 3264 | 4 | 3.1466167 | 9.75 | 0 | 4 | 0.9853692 | 0.25 | Phosphatidylinositol transfer protein, cytoplasmic 1; Phosphatidylino |
| TTPA | P49638 | 104 | 9606.ENSP | 4.0017436 | 0.0 | 0.2498910 | 8 | 2546 | 6 | 3.1903334 | 5.5 | 0 | 6 | 0.9835970 | 0.1875 | Tocopherol (alpha) transfer protein; Binds alpha-tocopherol, enhanc |
| FAM107A | O95990 | 105 | 9606.ENSP | 3.4158674 | 0.0 | 0.2927514 | 7 | 15556 | 4 | 0.0011216 | 23.75 | 0 | 4 | 0.9867985 | 0.2645348837209302 | Family with sequence similarity 107, member A; When transfected in |
| MLC1 | Q15049 | 105 | 9606.ENSP | 3.3243243 | 0.4 | 0.3008130 | 7 | 14264 | 5 | 9.3033629 | 43.2 | 0 | 5 | 0.9872987 | 0.3359375 | Megalencephalic leukoencephalopathy with subcortical cysts 1; Regu |
| NEXN | Q9Y2V1 | 105 | 9606.ENSP | 4.0479511 | 0.0 | 0.2470385 | 8 | 400 | 3 | 6.0207246 | 5.3333333 | 0 | 3 | 0.9833445 | 0.3333333333333333 | Nexilin (F actin binding protein); Involved in regulating cell migration |
| ACOX2 | Q99424 | 106 | 9606.ENSP | 3.4324324 | 0.1388888 | 0.2913385 | 8 | 18298 | 9 | 0.0012825 | 13.444444 | 0 | 9 | 0.9867080 | 0.1428571428571428 | 3-alpha,7-alpha,12-alpha-trihydroxy-5-beta-cholestanoyl-CoA 24-hyd |
| ACSF2 | Q96CM8 | 106 | 9606.ENSP | 3.5945945 | 0.2 | 0.2781954 | 8 | 3230 | 5 | 2.3179924 | 15.0 | 0 | 5 | 0.9858218 | 0.2317460317460317 | Acyl-CoA synthetase family member 2, mitochondrial; Acyl-CoA synth |
| HSDL2 | Q6YN16 | 106 | 9606.ENSP | 3.9590235 | 0.4 | 0.2525875 | 9 | 610 | 5 | 7.2180483 | 9.2 | 0 | 5 | 0.9838304 | 0.2727272727272727 | Short chain dehydrogenase/reductase family 13C member 1; Has app |
| PRKX | P51817 | 107 | 9606.ENSP | 3.2484742 | 0.1785714 | 0.3078368 | 7 | 9742 | 8 | 5.7644252 | 23.625 | 0 | 8 | 0.9877132 | 0.1570945945945946 | cAMP-dependent protein kinase catalytic subunit PRKX; Serine/threo |
| TBL1X | O60907 | 107 | 9606.ENSP | 2.7672188 | 0.0942528 | 0.3613736 | 7 | 229624 | 30 | 0.0162149 | 33.433333 | 0 | 30 | 0.9903430 | 0.0775700934579439 | F-box-like/WD repeat-containing protein TBL1X; F-box-like protein in |
| TRIM47 | Q96LD4 | 107 | 9606.ENSP | 3.7663469 | 0.0 | 0.2655092 | 8 | 0 | 1 | 0.0 | 30.0 | 0 | 1 | 0.9848833 | 0.0 | Gene overexpressed in astrocytoma protein; Tripartite motif contain |
| SLC2A10 | O95528 | 108 | 9606.ENSP | 3.5396687 | 0.3 | 0.2825123 | 8 | 16422 | 5 | 8.4932912 | 21.0 | 0 | 5 | 0.9861220 | 0.3029411764705882 | Solute carrier family 2 (facilitated glucose transporter), member 10; |
| SLC5A10 | A0PJK1 | 108 | 9606.ENSP | 3.6495204 | 0.0 | 0.2740086 | 8 | 508 | 2 | 6.0144545 | 23.0 | 0 | 2 | 0.9855217 | 0.5 | Solute carrier family 5 (sodium/sugar cotransporter), member 10; Hi |
| SMOX | Q9NWM0 | 108 | 9606.ENSP | 4.2179598 | 0.0 | 0.2370814 | 9 | 148 | 2 | 2.0118247 | 6.5 | 0 | 2 | 0.9824155 | 0.5 | Polyamine oxidase 1; Flavoenzyme which catalyzes the oxidation of s |
| PDZK1IP1 | Q13113 | 109 | 9606.ENSP | 3.9816913 | 0.0 | 0.2511495 | 8 | 186 | 2 | 1.4265280 | 9.0 | 0 | 2 | 0.9837066 | 0.5 | 17 kDa membrane-associated protein; May play an important role in |
| PIEZO1 | Q92508 | 109 | 9606.ENSP | 3.8735832 | 0.0 | 0.2581589 | 8 | 456 | 3 | 6.2486720 | 8.0 | 0 | 3 | 0.9842973 | 0.3333333333333333 | Piezo-type mechanosensitive ion channel component 1; Pore-forming |
| RBPMS | Q93062 | 109 | 9606.ENSP | 3.6416739 | 0.0 | 0.2745989 | 8 | 4338 | 5 | 4.2226176 | 12.0 | 0 | 5 | 0.9855646 | 0.2037037037037037 | RNA binding protein with multiple splicing; Acts as a coactivator of tr |
| GPSM3 | Q9Y4H4 | 110 | 9606.ENSP | 3.6904969 | 0.0 | 0.2709662 | 8 | 228 | 2 | 1.3760295 | 21.0 | 0 | 2 | 0.9852978 | 0.5 | Activator of G-protein signaling 4; Interacts with subunit of G(i) alpha |
| SH2D2A | Q9NP31 | 110 | 9606.ENSP | 3.2371403 | 0.1666666 | 0.3089146 | 7 | 1232 | 4 | 6.5195780 | 50.0 | 0 | 4 | 0.9877751 | 0.3152866242038216 | SH2 domain-containing adapter protein; Could be a T-cell-specific ad |
| SUCLG2 | Q96I99 | 110 | 9606.ENSP | 3.2842197 | 0.0606060 | 0.3044863 | 7 | 16086 | 12 | 0.0015041 | 14.25 | 0 | 12 | 0.9875179 | 0.1076115485564304 | Succinate--CoA ligase [GDP-forming] subunit beta, mitochondrial; GT |
| COL6A2 | Q9UML3 | 111 | 9606.ENSP | 2.9947689 | 0.3723076 | 0.3339155 | 7 | 44668 | 26 | 0.0017396 | 42.576923 | 0 | 26 | 0.9890996 | 0.1440677966101695 | Collagen alpha-2(VI) chain; Collagen VI acts as a cell-binding protein; |
| CRTAP | O75718 | 111 | 9606.ENSP | 3.6643417 | 1.0 | 0.2729003 | 8 | 0 | 3 | 0.0 | 27.666666 | 0 | 3 | 0.9854407 | 0.5123456790123457 | Cartilage associated protein; Necessary for efficient 3-hydroxylation |
| SERPINH1 | P50454 | 111 | 9606.ENSP | 2.9232781 | 0.2923976 | 0.3420817 | 7 | 47068 | 19 | 0.0023886 | 46.157894 | 0 | 19 | 0.9894902 | 0.1428571428571428 | Serpin peptidase inhibitor, clade H (heat shock protein 47), member |
| CMTM3 | Q96MX0 | 112 | 9606.ENSP | 44566 | 0.0 | 0.6666666 | 2 | 0 | 1 | 0.0 | 2.0 | 0 | 1 | 0.75 | 0.0 | CKLF like MARVEL transmembrane domain containing 3; Belongs to t |
| CMTM6 | Q9NX76 | 112 | 9606.ENSP | 1.0 | 0.0 | 1.0 | 1 | 2 | 2 | 1.0 | 1.0 | 0 | 2 | 1.0 | 0.0 | CKLF like MARVEL transmembrane domain containing 6; Belongs to t |
| TSPAN11 | A1L157 | 112 | 9606.ENSP | 44566 | 0.0 | 0.6666666 | 2 | 0 | 1 | 0.0 | 2.0 | 0 | 1 | 0.75 | 0.0 | Tetraspanin 11; Tetraspanins |
| APCDD1 | Q8J025 | 113 | 9606.ENSP | 3.6721883 | 0.1666666 | 0.2723171 | 7 | 36276 | 4 | 0.0025318 | 11.75 | 0 | 4 | 0.9853978 | 0.2616279069767442 | Adenomatosis polyposis coli down-regulated 1 protein; Negative regu |
| MMD2 | Q8IY49 | 113 | 9606.ENSP | 3.2275501 | 0.2 | 0.3098325 | 6 | 105310 | 11 | 0.0086958 | 19.636363 | 0 | 11 | 0.9878275 | 0.1217838765008576 | Monocyte to macrophage differentiation associated 2; Progestin and |
| PAQR8 | Q8TEZ7 | 113 | 9606.ENSP | 4.2266782 | 0.0 | 0.2365924 | 7 | 0 | 1 | 0.0 | 11.0 | 0 | 1 | 0.9823678 | 0.0 | Progesterone and adipoQ receptor family member 8; Plasma membr |
| BOD1L1 | Q8NFC6 | 114 | 9606.ENSP | 3.5353095 | 0.0476190 | 0.2828606 | 8 | 6906 | 7 | 8.3001826 | 11.0 | 0 | 7 | 0.9861458 | 0.1607142857142857 | Biorientation of chromosomes in cell division protein 1-like 1; Compo |
| C21orf62 | Q9NYP8 | 114 | 9606.ENSP | 3.9415867 | 0.0 | 0.2537049 | 8 | 196 | 2 | 1.9416285 | 11.5 | 0 | 2 | 0.9839257 | 0.5 | Chromosome 21 open reading frame 62 |
| TNS3 | Q68CZ2 | 114 | 9606.ENSP | 3.0706190 | 0.3 | 0.3256672 | 7 | 5254 | 5 | 2.6878135 | 81.2 | 0 | 5 | 0.9886851 | 0.3438297872340425 | Tensin-like SH2 domain-containing protein 1; May play a role in actin |
| BATF2 | Q8N1L9 | 115 | 9606.ENSP | 3.5013077 | 0.2666666 | 0.2856075 | 8 | 4538 | 6 | 3.1444221 | 31.0 | 0 | 6 | 0.9863316 | 0.2752976190476190 | Basic leucine zipper transcriptional factor ATF-like 2; AP-1 family tran |
| LYPD1 | Q8N2G4 | 115 | 9606.ENSP | 3.5030514 | 0.0952380 | 0.2854654 | 7 | 14114 | 7 | 8.7204155 | 13.0 | 0 | 7 | 0.9863221 | 0.1676190476190476 | Ly6/PLAUR domain-containing protein 1; Believed to act as a modula |
| PLAC9 | Q5JTB6 | 115 | 9606.ENSP | 4.1726242 | 0.0 | 0.2396573 | 8 | 66 | 2 | 1.0841943 | 9.0 | 0 | 2 | 0.9826632 | 0.5 | Placenta-specific protein 9; Placenta specific 9; Belongs to the PLAC9 |
| LATS2 | Q9NRM7 | 116 | 9606.ENSP | 2.9511769 | 0.1956521 | 0.3388478 | 7 | 47436 | 24 | 0.0031285 | 29.5 | 0 | 24 | 0.9893378 | 0.0951995685005393 | Kinase phosphorylated during mitosis protein; Negative regulator of |
| MOB3C | Q70IA8 | 116 | 9606.ENSP | 3.8587619 | 0.0 | 0.2591504 | 8 | 194 | 2 | 1.5607957 | 15.5 | 0 | 2 | 0.9843783 | 0.5 | Mps one binder kinase activator-like 2C; May regulate the activity of |
| TNIP2 | Q8NFZ5 | 116 | 9606.ENSP | 3.3783783 | 0.5714285 | 0.296 | 8 | 8236 | 7 | 4.4824817 | 34.857142 | 0 | 7 | 0.9870033 | 0.2892857142857143 | A20-binding inhibitor of NF-kappa-B activation 2; Inhibits NF-kappa-B |
| ZBTB47 | Q9UFB7 | 117 | 9606.ENSP | 4.4986922 | 0.0 | 0.2222868 | 9 | 322 | 2 | 4.2237558 | 5.0 | 0 | 2 | 0.9808814 | 0.5 | Zinc finger and BTB domain-containing protein 47; May be involved i |
| ZNF395 | Q9H8N7 | 117 | 9606.ENSP | 3.6617262 | 0.0 | 0.2730952 | 8 | 14302 | 3 | 0.0012745 | 15.333333 | 0 | 3 | 0.9854550 | 0.3495934959349593 | Huntington disease gene regulatory region-binding protein 2; Plays a |
| MPZL2 | O60487 | 118 | 9606.ENSP | 3.5065387 | 0.0666666 | 0.2851815 | 8 | 3430 | 6 | 2.9069778 | 12.666666 | 0 | 6 | 0.9863030 | 0.1791044776119403 | Myelin protein zero-like protein 2; Mediates homophilic cell-cell adhe |
| ZBTB20 | Q9HC78 | 118 | 9606.ENSP | 3.8151700 | 0.0 | 0.2621115 | 8 | 318 | 3 | 4.4542365 | 9.0 | 0 | 3 | 0.9846165 | 0.3333333333333333 | Zinc finger and BTB domain-containing protein 20; May be a transcri |
| MID1 | O15344 | 119 | 9606.ENSP | 3.4734088 | 0.0 | 0.2879016 | 8 | 37286 | 5 | 0.0031080 | 15.6 | 0 | 5 | 0.9864841 | 0.2027777777777777 | RING-type E3 ubiquitin transferase Midline-1; Has E3 ubiquitin ligase |
| UBTD1 | Q9HAC8 | 119 | 9606.ENSP | 4.4725370 | 0.0 | 0.2235867 | 9 | 0 | 1 | 0.0 | 5.0 | 0 | 1 | 0.9810243 | 0.0 | Ubiquitin domain-containing protein 1; May be involved in the regula |
| TINAGL1 | Q9GZM7 | 120 | 9606.ENSP | 3.5013077 | 0.1666666 | 0.2856075 | 7 | 1130 | 4 | 9.6386057 | 29.75 | 0 | 4 | 0.9863316 | 0.3214285714285714 | Tubulointerstitial nephritis antigen-related protein; May be implicate |
| CABLES1 | Q8TDN4 | 121 | 9606.ENSP | 3.4062772 | 0.2 | 0.2935756 | 7 | 3958 | 6 | 2.8011993 | 22.666666 | 0 | 6 | 0.9868509 | 0.2393617021276596 | CDK5 and ABL1 enzyme substrate 1; Cyclin-dependent kinase binding |
| EFEMP1 | Q12805 | 121 | 9606.ENSP | 2.9668700 | 0.2878787 | 0.3370555 | 7 | 26092 | 12 | 0.0017698 | 51.25 | 0 | 12 | 0.9892520 | 0.1694260485651214 | EGF containing fibulin-like extracellular matrix protein 1; Binds EGFR, |
| GRAMD1C | Q8IYS0 | 122 | 9606.ENSP | 1.0 | 0.0 | 1.0 | 1 | 0 | 1 | 0.0 | 1.0 | 0 | 1 | 1.0 | 0.0 | GRAM domain containing 1C |
| YBX3 | P16989 | 123 | 9606.ENSP | 3.3504795 | 0.3333333 | 0.2984647 | 8 | 526 | 3 | 3.2635928 | 38.666666 | 0 | 3 | 0.9871558 | 0.3650793650793650 | Single-strand DNA-binding protein NF-GMB; Binds to the GM-CSF pro |
| ZC3HAV1 | Q7Z2W4 | 123 | 9606.ENSP | 2.9930252 | 0.2333333 | 0.3341101 | 7 | 62830 | 16 | 0.0045887 | 35.5625 | 0 | 16 | 0.9891091 | 0.1272482014388489 | ADP-ribosyltransferase diphtheria toxin-like 13; Antiviral protein whi |
| SERPINB1 | P30740 | 124 | 9606.ENSP | 3.2266782 | 0.0 | 0.3099162 | 7 | 620 | 2 | 2.2828277 | 89.5 | 0 | 2 | 0.9878323 | 0.5057142857142857 | Serpin peptidase inhibitor, clade B (ovalbumin), member 1; Regulate |
| TES | Q9UGI8 | 124 | 9606.ENSP | 3.1691368 | 0.0555555 | 0.3155433 | 7 | 10914 | 9 | 8.4704635 | 25.0 | 0 | 9 | 0.9881467 | 0.1423001949317738 | Testis derived transcript (3 LIM domains); Scaffold protein that may |
| ACSM5 | Q6NUN0 | 125 | 9606.ENSP | 4.2589363 | 0.0 | 0.2348004 | 9 | 24190 | 2 | 0.0017436 | 6.5 | 0 | 2 | 0.9821916 | 0.5 | Acyl-CoA synthetase medium-chain family member 5; Has medium-c |
| CACFD1 | Q9UGQ2 | 125 | 9606.ENSP | 5.2580645 | 0.0 | 0.1901840 | 10 | 0 | 1 | 0.0 | 2.0 | 0 | 1 | 0.9767318 | 0.0 | Calcium channel flower domain containing 1 |
| C2CD4B | A6NLJ0 | 126 | 9606.ENSP | 3.6713164 | 0.3333333 | 0.2723818 | 7 | 3094 | 3 | 2.7454976 | 17.0 | 0 | 3 | 0.9854026 | 0.3546099290780142 | C2 calcium-dependent domain-containing protein 4B; May be involve |
| HILPDA | Q9Y5L2 | 126 | 9606.ENSP | 4.0174367 | 0.0 | 0.2489149 | 7 | 3516 | 3 | 4.2247565 | 7.3333333 | 0 | 3 | 0.9835112 | 0.3333333333333333 | Hypoxia-inducible lipid droplet-associated protein; Increases intracell |
| DAG1 | Q14118 | 127 | 9606.ENSP | 2.8177855 | 0.1970443 | 0.3548886 | 7 | 89664 | 29 | 0.0059754 | 36.827586 | 0 | 29 | 0.9900667 | 0.0948053105230330 | Dystroglycan 1 (dystrophin-associated glycoprotein 1); The dystrogly |
| RAPSN | Q13702 | 127 | 9606.ENSP | 3.0401046 | 0.5333333 | 0.3289360 | 7 | 10106 | 10 | 6.1159569 | 47.9 | 0 | 10 | 0.9888518 | 0.1881889763779527 | 43 kDa receptor-associated protein of the synapse; Postsynaptic prot |
| GCSH | P23434 | 128 | 9606.ENSP | 3.2066259 | 0.1272727 | 0.3118542 | 7 | 34586 | 11 | 0.0027260 | 17.909090 | 0 | 11 | 0.9879419 | 0.1087562744004461 | Glycine cleavage system protein H (aminomethyl carrier); The glycine |
| SARDH | Q9UL12 | 128 | 9606.ENSP | 3.8360941 | 0.3 | 0.2606818 | 8 | 1396 | 5 | 1.3775240 | 10.6 | 0 | 5 | 0.9845022 | 0.2736842105263158 | Sarcosine dehydrogenase, mitochondrial; Sarcosine dehydrogenase; |
| METRNL | Q641Q3 | 129 | 9606.ENSP | 3.7724498 | 0.0 | 0.2650797 | 8 | 31022 | 4 | 0.0022380 | 8.25 | 0 | 4 | 0.9848500 | 0.25 | Meteorin, glial cell differentiation regulator-like; Hormone induced f |
| OTOS | Q8NHW6 | 129 | 9606.ENSP | 4.7715780 | 0.0 | 0.2095742 | 9 | 0 | 1 | 0.0 | 4.0 | 0 | 1 | 0.9793902 | 0.0 | Otospiralin; May be essential for the survival of the neurosensory epi |
| COLEC12 | Q5KU26 | 130 | 9606.ENSP | 4.4838709 | 0.0 | 0.2230215 | 9 | 0 | 1 | 0.0 | 7.0 | 0 | 1 | 0.9809624 | 0.0 | Scavenger receptor with C-type lectin; Scavenger receptor that displa |
| LIMK2 | P53671 | 130 | 9606.ENSP | 3.4847428 | 0.0952380 | 0.2869652 | 8 | 32812 | 7 | 0.0020081 | 14.857142 | 0 | 7 | 0.9864221 | 0.1831501831501831 | LIM domain kinase 2; Displays serine/threonine-specific phosphoryla |
| MAP3K6 | O95382 | 131 | 9606.ENSP | 4.0156931 | 0.0 | 0.2490230 | 8 | 746 | 3 | 1.1287192 | 8.6666666 | 0 | 3 | 0.9835208 | 0.3333333333333333 | Mitogen-activated protein kinase kinase kinase 6; Component of a pr |
| POLD4 | Q9HCU8 | 131 | 9606.ENSP | 3.6076721 | 0.0 | 0.2771870 | 8 | 3910 | 2 | 2.4568301 | 22.5 | 0 | 2 | 0.9857504 | 0.5 | Polymerase (DNA-directed), delta 4, accessory subunit; As a compon |
| MTRNR2L1 | P0CJ75 | 132 | 9606.ENSP | 1.0 | 0.0 | 1.0 | 1 | 0 | 1 | 0.0 | 1.0 | 0 | 1 | 1.0 | 0.0 | MT-RNR2-like protein 8; Plays a role as a neuroprotective and antiap |
| MTRNR2L | P0CJ75 | 132 | 9606.ENSP | 1.0 | 0.0 | 1.0 | 1 | 0 | 1 | 0.0 | 1.0 | 0 | 1 | 1.0 | 0.0 | MT-RNR2-like 8; Plays a role as a neuroprotective and antiapoptotic |
| EMILIN2 | Q9BXX0 | 133 | 9606.ENSP | 4.4446381 | 0.0 | 0.2249901 | 8 | 0 | 1 | 0.0 | 6.0 | 0 | 1 | 0.9811768 | 0.0 | Elastin microfibril interface-located protein 2; May be responsible fo |
| MYOM1 | P52179 | 133 | 9606.ENSP | 3.4455100 | 0.2 | 0.2902327 | 7 | 33772 | 6 | 0.0025036 | 18.166666 | 0 | 6 | 0.9866365 | 0.1879432624113475 | 190 kDa connectin-associated protein; Major component of the verte |
| ANKHD1 | Q9NWG4 | 134 | 9606.ENSP | 3.3801220 | 0.0666666 | 0.2958473 | 8 | 4320 | 6 | 2.9118768 | 23.5 | 0 | 6 | 0.9869938 | 0.2038690476190476 | Ankyrin repeat and KH domain-containing protein 1; May play a role |
| SIPA1 | Q96FS4 | 134 | 9606.ENSP | 3.7576285 | 0.0 | 0.2661252 | 8 | 86 | 2 | 5.1290844 | 17.5 | 0 | 2 | 0.9849309 | 0.515625 | Signal-induced proliferation-associated protein 1; GTPase activator f |
| RXRA | P19793 | 135 | 9606.ENSP | 2.9537925 | 0.1476190 | 0.3385478 | 6 | 40154 | 21 | 0.0033917 | 23.761904 | 0 | 21 | 0.9893235 | 0.0781078107810781 | Nuclear receptor subfamily 2 group B member 1; Receptor for retino |
| RXRG | P48443 | 135 | 9606.ENSP | 3.3251961 | 0.2666666 | 0.3007341 | 6 | 4376 | 6 | 4.1275662 | 23.5 | 0 | 6 | 0.9872940 | 0.2028985507246377 | Nuclear receptor subfamily 2 group B member 3; Receptor for retino |
| MSRB3 | Q8IXL7 | 136 | 9606.ENSP | 4.5213600 | 0.0 | 0.2211723 | 9 | 0 | 1 | 0.0 | 8.0 | 0 | 1 | 0.9807575 | 0.0 | Methionine-R-sulfoxide reductase B3; Catalyzes the reduction of free |
| RFX2 | P48378 | 136 | 9606.ENSP | 3.5222319 | 0.1071428 | 0.2839108 | 8 | 75470 | 8 | 0.0054015 | 10.875 | 0 | 8 | 0.9862173 | 0.1440972222222222 | Regulatory factor X, 2 (influences HLA class II expression); Transcripti |
| BRCA1 | P38398 | 137 | 9606.ENSP | 2.6242371 | 0.1730545 | 0.3810631 | 7 | 230202 | 42 | 0.0153766 | 43.238095 | 0 | 42 | 0.9911243 | 0.0815104869925285 | Breast cancer type 1 susceptibility protein; E3 ubiquitin-protein ligase |
| MCM7 | P33993 | 137 | 9606.ENSP | 3.2999128 | 0.3111111 | 0.3030383 | 7 | 13980 | 10 | 0.0011140 | 22.1 | 0 | 10 | 0.9874321 | 0.1639097744360902 | Minichromosome maintenance complex component 7; Acts as comp |
| CCDC80 | Q76M96 | 138 | 9606.ENSP | 3.1900610 | 0.3333333 | 0.3134736 | 7 | 8364 | 3 | 8.9219394 | 63.666666 | 0 | 3 | 0.9880324 | 0.3558052434456928 | Up-regulated in BRS-3 deficient mouse homolog; Promotes cell adhe |
| SAMD4A | Q9UPU9 | 138 | 9606.ENSP | 3.9877942 | 0.0 | 0.2507651 | 8 | 1576 | 3 | 1.4908659 | 5.6666666 | 0 | 3 | 0.9836732 | 0.3333333333333333 | Sterile alpha motif domain-containing protein 4A; Acts as a translatio |
| ASRGL1 | Q7L266 | 139 | 9606.ENSP | 4.0401046 | 0.0 | 0.2475183 | 8 | 140 | 2 | 1.4463018 | 9.0 | 0 | 2 | 0.9833874 | 0.5 | Isoaspartyl peptidase/L-asparaginase; Has both L-asparaginase and b |
| ECE1 | Q5THM8 | 139 | 9606.ENSP | 3.3173496 | 0.1 | 0.3014454 | 7 | 21576 | 5 | 0.0012673 | 37.8 | 0 | 5 | 0.9873368 | 0.2929133858267717 | Endothelin converting enzyme 1; Converts big endothelin-1 to endot |
| DTNA | Q9Y4J8 | 140 | 9606.ENSP | 3.6172624 | 0.1333333 | 0.2764521 | 8 | 14536 | 6 | 0.0012435 | 11.166666 | 0 | 6 | 0.9856980 | 0.2012578616352201 | Dystrophin-related protein 3; May be involved in the formation and |
| FYCO1 | Q9BQS8 | 140 | 9606.ENSP | 3.5972101 | 0.0 | 0.2779932 | 8 | 1180 | 3 | 1.0553238 | 19.0 | 0 | 3 | 0.9858075 | 0.3529411764705882 | FYVE and coiled-coil domain-containing protein 1; May mediate micr |
| EZR | P15311 | 141 | 9606.ENSP | 2.5379250 | 0.1754098 | 0.3940226 | 6 | 278782 | 61 | 0.0164203 | 40.950819 | 0 | 61 | 0.9915960 | 0.0711617961511047 | Cytovillin; Probably involved in connections of major cytoskeletal stru |
| MSN | P26038 | 141 | 9606.ENSP | 2.6678291 | 0.3076923 | 0.3748366 | 6 | 75280 | 40 | 0.0030840 | 54.3 | 0 | 40 | 0.9908861 | 0.1114989733059548 | Membrane-organizing extension spike protein; Probably involved in |
| ALDH1L1 | O75891 | 142 | 9606.ENSP | 3.1194420 | 0.3166666 | 0.3205701 | 7 | 117956 | 16 | 0.0065277 | 26.8125 | 0 | 16 | 0.9884183 | 0.1302083333333333 | Cytosolic 10-formyltetrahydrofolate dehydrogenase; Aldehyde dehy |
| SULT1C4 | O75897 | 142 | 9606.ENSP | 4.1185701 | 0.0 | 0.2428027 | 8 | 0 | 1 | 0.0 | 16.0 | 0 | 1 | 0.9829586 | 0.0 | Sulfotransferase family, cytosolic, 1C, member 4; Sulfotransferase th |
| PERP | Q96FX8 | 143 | 9606.ENSP | 3.4298169 | 0.2857142 | 0.2915607 | 8 | 32448 | 7 | 0.0013931 | 21.0 | 0 | 7 | 0.9867223 | 0.2327447833065810 | P53 apoptosis effector related to PMP-22; Component of intercellula |
| RNF144B | Q7Z419 | 143 | 9606.ENSP | 4.3025283 | 0.0 | 0.2324214 | 9 | 186 | 2 | 1.9933212 | 7.0 | 0 | 2 | 0.9819533 | 0.5 | E3 ubiquitin-protein ligase RNF144B; E3 ubiquitin-protein ligase whic |
| MROH9 | Q5TGP6 | 144 | 9606.ENSP | 4.6957279 | 0.0 | 0.2129595 | 8 | 0 | 1 | 0.0 | 4.0 | 0 | 1 | 0.9798047 | 0.0 | Maestro heat like repeat family member 9 |
| PTPN21 | Q16825 | 144 | 9606.ENSP | 3.6965998 | 0.0 | 0.2705188 | 7 | 28652 | 4 | 0.0017913 | 12.25 | 0 | 4 | 0.9852644 | 0.2678571428571428 | Protein tyrosine phosphatase, non-receptor type 21; FERM domain c |
| ITGB3 | P05106 | 145 | 9606.ENSP | 2.5422842 | 0.2788732 | 0.3933470 | 6 | 274974 | 71 | 0.0144329 | 49.563380 | 0 | 71 | 0.9915722 | 0.0861481935088793 | Integrin, beta 3 (platelet glycoprotein IIIa, antigen CD61); Integrin alp |
| SYTL4 | Q96C24 | 145 | 9606.ENSP | 3.5414123 | 0.0 | 0.2823732 | 7 | 0 | 1 | 0.0 | 71.0 | 0 | 1 | 0.9861125 | 0.0 | Synaptotagmin-like protein 4; Modulates exocytosis of dense-core gr |
| FAM181A | Q8N9Y4 | 146 | 9606.ENSP | 3.9354838 | 0.0 | 0.2540983 | 8 | 0 | 1 | 0.0 | 23.0 | 0 | 1 | 0.9839591 | 0.0 | Family with sequence similarity 181 member A |
| TEAD4 | Q15561 | 146 | 9606.ENSP | 2.9363557 | 0.3715415 | 0.3405581 | 7 | 46734 | 23 | 0.0027623 | 36.217391 | 0 | 23 | 0.9894188 | 0.1163148329372291 | Transcriptional enhancer factor TEF-3; Transcription factor which pla |
| CCDC69 | A6NI79 | 147 | 9606.ENSP | 5.1700087 | 0.0 | 0.1934232 | 9 | 0 | 1 | 0.0 | 2.0 | 0 | 1 | 0.9772130 | 0.0 | Coiled-coil domain-containing protein 69; May act as a scaffold to reg |
| CRISPLD2 | Q9H0B8 | 147 | 9606.ENSP | 4.1708805 | 0.0 | 0.2397575 | 8 | 23998 | 2 | 0.0017436 | 6.0 | 0 | 2 | 0.9826727 | 0.5 | Cysteine-rich secretory protein LCCL domain containing 2; Promotes |

| | | | | | | | | | | | | | | | | |
|---|---|---|---|---|---|---|---|---|---|---|---|---|---|---|---|---|
| METTL7B | Q6UX53 | 148 | 9606.ENSP | 4.1639058 | 0.0 | 0.2401591 | 8 | 0 | 1 | 0.0 | 7.0 | 0 | 1 | 0.9827108 | 0.0 | Methyltransferase-like protein 7B; Probable methyltransferase. |
| RARRES2 | Q99969 | 148 | 9606.ENSP | 3.1647776 | 0.6190476 | 0.3159779 | 7 | 27706 | 7 | 0.0017454 | 62.142857 | 0 | 7 | 0.9881706 | 0.2980769230769231 | Retinoic acid receptor responder (tazarotene induced) 2; Adipocyte-s |
| CASKIN2 | Q8WXE0 | 149 | 9606.ENSP | 4.3740191 | 0.0 | 0.2286226 | 8 | 0 | 1 | 0.0 | 4.0 | 0 | 1 | 0.9815627 | 0.0 | CASK interacting protein 2; Sterile alpha motif domain containing |
| LIMS2 | Q7Z4I7 | 149 | 9606.ENSP | 3.3748910 | 0.1666666 | 0.2963058 | 7 | 29656 | 4 | 0.0018241 | 43.25 | 0 | 4 | 0.9870224 | 0.3717391304347826 | LIM and senescent cell antigen-like-containing domain protein 2; Ada |
| NANOS1 | Q8WY41 | 150 | 9606.ENSP | 4.1011333 | 0.0666666 | 0.2438350 | 7 | 2716 | 6 | 5.2785456 | 5.8333333 | 0 | 6 | 0.9830539 | 0.1845238095238095 | Nanos homolog 1 (Drosophila); May act as a translational repressor w |
| PLA2G5 | P39877 | 150 | 9606.ENSP | 3.7096774 | 0.2 | 0.2695652 | 8 | 4834 | 5 | 6.5416166 | 9.8 | 0 | 5 | 0.9851930 | 0.2181818181818181 | Phosphatidylcholine 2-acylhydrolase 5; PA2 catalyzes the calcium-dep |
| CATSPERD | Q86XM0 | 151 | 9606.ENSP | 4.4829991 | 0.0 | 0.2230649 | 9 | 24098 | 3 | 0.0018140 | 4.0 | 0 | 3 | 0.9809672 | 0.3333333333333333 | Cation channel sperm-associated protein subunit delta; Auxiliary com |
| OR6V1 | Q8N148 | 151 | 9606.ENSP | 5.4821272 | 0.0 | 0.1824109 | 10 | 0 | 1 | 0.0 | 3.0 | 0 | 1 | 0.9755075 | 0.0 | Olfactory receptor, family 6, subfamily V, member 1; Odorant recept |
| CHI3L1 | P36222 | 152 | 9606.ENSP | 2.8020924 | 0.625 | 0.3568761 | 7 | 26622 | 16 | 0.0018403 | 86.5625 | 0 | 16 | 0.9901525 | 0.2084337349397590 | Chitinase 3-like 1 (cartilage glycoprotein-39); Carbohydrate-binding le |
| TMEM219 | Q86XT9 | 152 | 9606.ENSP | 3.8012205 | 0.0 | 0.2630733 | 8 | 0 | 1 | 0.0 | 16.0 | 0 | 1 | 0.9846927 | 0.0 | Insulin-like growth factor-binding protein 3 receptor; Cell death recep |
| NDRG1 | Q92597 | 153 | 9606.ENSP | 3.4176111 | 0.1428571 | 0.2926020 | 7 | 33818 | 8 | 0.0030166 | 14.25 | 0 | 8 | 0.9867890 | 0.1491935483870967 | Reducing agents and tunicamycin-responsive protein; Stress-respons |
| NDRG2 | Q9UN36 | 153 | 9606.ENSP | 4.4167393 | 0.0 | 0.2264113 | 8 | 0 | 1 | 0.0 | 8.0 | 0 | 1 | 0.9813292 | 0.0 | N-myc downstream-regulated gene 2 protein; Contributes to the reg |
| CD99 | P14209 | 154 | 9606.ENSP | 3.2589363 | 0.1785714 | 0.3068485 | 7 | 18596 | 8 | 0.0010102 | 30.875 | 0 | 8 | 0.9876560 | 0.2006578947368421 | T-cell surface glycoprotein E2; Involved in T-cell adhesion processes a |
| N4BP2 | Q86UW6 | 154 | 9606.ENSP | 3.7384481 | 0.0 | 0.2674906 | 8 | 490 | 3 | 8.6137718 | 13.666666 | 0 | 3 | 0.9850358 | 0.3333333333333333 | NEDD4 binding protein 2; Has 5'-polynucleotide kinase and nicking en |
| MORC4 | Q8TE76 | 155 | 9606.ENSP | 4.5719267 | 0.0 | 0.2187261 | 8 | 0 | 1 | 0.0 | 2.0 | 0 | 1 | 0.9804812 | 0.0 | Zinc finger CW-type coiled-coil domain protein 2; MORC family CW-ty |
| TPD52L1 | Q5TDQ0 | 155 | 9606.ENSP | 3.5727986 | 0.0 | 0.2798926 | 7 | 21938 | 2 | 0.0017436 | 27.0 | 0 | 2 | 0.9859409 | 0.5 | Tumor protein D52 like 1 |
| TRIL | Q7L0X0 | 156 | 9606.ENSP | 3.0557977 | 0.5 | 0.3272467 | 7 | 854 | 4 | 3.8108792 | 89.75 | 0 | 4 | 0.9887661 | 0.358 | Leucine-rich repeat-containing protein KIAA0644; Component of the |
| FAM167B | Q9BTA0 | 157 | 9606.ENSP | 5.1203138 | 0.0 | 0.1953005 | 8 | 0 | 1 | 0.0 | 3.0 | 0 | 1 | 0.9774846 | 0.0 | Family with sequence similarity 167 member B; Belongs to the FAM1 |
| STK33 | Q9BYT3 | 157 | 9606.ENSP | 4.1211857 | 0.0 | 0.2426486 | 7 | 26522 | 3 | 0.0017658 | 6.6666666 | 0 | 3 | 0.9829443 | 0.3541666666666667 | Serine/threonine-protein kinase 33; Serine/threonine protein kinase |
| LHX2 | P50458 | 158 | 9606.ENSP | 2.9564080 | 0.2687747 | 0.3382483 | 7 | 57286 | 23 | 0.0036121 | 35.434782 | 0 | 23 | 0.9893092 | 0.1220389805097451 | LIM/homeobox protein Lhx2; Acts as a transcriptional activator. Stim |
| NEK7 | Q8TDX7 | 158 | 9606.ENSP | 3.6495204 | 0.1666666 | 0.2740086 | 7 | 1204 | 4 | 9.3297708 | 11.75 | 0 | 4 | 0.9855217 | 0.2616279069767442 | Serine/threonine-protein kinase Nek7; Protein kinase which plays an |
| APOL3 | O95236 | 159 | 9606.ENSP | 4.0078465 | 0.0 | 0.2495105 | 8 | 86 | 2 | 5.1458794 | 18.0 | 0 | 2 | 0.9835636 | 0.5151515151515151 | TNF-inducible protein CG12-1; May affect the movement of lipids in t |
| BNIP2 | Q12982 | 160 | 9606.ENSP | 3.5649520 | 0.0 | 0.2805086 | 8 | 6246 | 5 | 5.6507628 | 11.2 | 0 | 5 | 0.9859838 | 0.204 | BCL2/adenovirus E1B 19 kDa protein-interacting protein 2; Implicated |
| PRR5 | Q9NSG0 | 160 | 9606.ENSP | 3.7846556 | 0.0 | 0.2642248 | 8 | 288 | 2 | 3.3898671 | 13.0 | 0 | 2 | 0.9847833 | 0.5 | Protein observed with Rictor-1; Subunit of mTORC2, which regulates |
| LGALS3 | P17931 | 161 | 9606.ENSP | 2.5815170 | 0.2543675 | 0.3873691 | 6 | 200862 | 54 | 0.0111460 | 51.462962 | 0 | 54 | 0.9913578 | 0.0939467804184440 | Lectin, galactoside-binding, soluble, 3; Galactose-specific lectin which |
| UACA | Q9BZF9 | 161 | 9606.ENSP | 3.5309503 | 1.0 | 0.2832098 | 7 | 0 | 2 | 0.0 | 32.5 | 0 | 2 | 0.9861696 | 0.5158730158730159 | Uveal autoantigen with coiled-coil domains and ankyrin repeats; Regu |
| MID1IP1 | Q9NPA3 | 162 | 9606.ENSP | 4.3129904 | 0.0 | 0.2318576 | 9 | 124 | 2 | 1.5573037 | 3.5 | 0 | 2 | 0.9818962 | 0.5 | Gastrulation-specific G12-like protein; Plays a role in the regulation o |
| PLIN4 | Q96Q06 | 162 | 9606.ENSP | 3.6120313 | 0.0 | 0.2768525 | 8 | 8336 | 2 | 6.8410838 | 27.5 | 0 | 2 | 0.9857266 | 0.5 | Adipocyte protein S3-12; May play a role in triacylglycerol packaging i |
| APOL4 | Q9BPW4 | 163 | 9606.ENSP | 4.7035745 | 0.0 | 0.2126042 | 9 | 0 | 1 | 0.0 | 8.0 | 0 | 1 | 0.9797618 | 0.0 | Apolipoprotein L, 4; May play a role in lipid exchange and transport t |
| ARSD | P51689 | 164 | 9606.ENSP | 3.6843940 | 0.1 | 0.2714150 | 8 | 40862 | 5 | 0.0028982 | 10.2 | 0 | 5 | 0.9853311 | 0.2232558139534883 | Arylsulfatase D; Sulfatases |
| C8orf82 | Q6P1X6 | 164 | 9606.ENSP | 4.6835222 | 0.0 | 0.2135145 | 9 | 0 | 1 | 0.0 | 5.0 | 0 | 1 | 0.9798714 | 0.0 | Chromosome 8 open reading frame 82 |
| CD109 | Q8N3A7 | 165 | 9606.ENSP | 3.0026155 | 0.4285714 | 0.3330429 | 7 | 17524 | 7 | 0.0014231 | 74.0 | 0 | 7 | 0.9890567 | 0.2670807453416149 | C3 and PZP-like alpha-2-macroglobulin domain-containing protein 7; |
| PEAR1 | Q5VY43 | 165 | 9606.ENSP | 3.9119442 | 0.0 | 0.2556273 | 8 | 326 | 3 | 3.0641597 | 5.3333333 | 0 | 3 | 0.9840877 | 0.3333333333333333 | Multiple epidermal growth factor-like domains protein 12; When ove |
| EMP1 | P54849 | 166 | 9606.ENSP | 3.1098517 | 0.1333333 | 0.3215587 | 7 | 19814 | 6 | 0.0020474 | 44.0 | 0 | 6 | 0.9884707 | 0.2042253521126760 | Epithelial membrane protein 1 |
| SMIM3 | Q9BZL3 | 166 | 9606.ENSP | 4.1089799 | 0.0 | 0.2433694 | 8 | 0 | 1 | 0.0 | 6.0 | 0 | 1 | 0.9830110 | 0.0 | NGF-induced differentiation clone 67 protein; Small integral membra |
| SLC4A4 | Q9Y6R1 | 167 | 9606.ENSP | 3.1002615 | 0.0808823 | 0.3225534 | 8 | 58488 | 17 | 0.0056892 | 15.941176 | 0 | 17 | 0.9885231 | 0.0762421473443746 | Solute carrier family 4 (sodium bicarbonate cotransporter), member |
| WNK4 | Q96J92 | 167 | 9606.ENSP | 3.7035745 | 0.3333333 | 0.2700094 | 7 | 1444 | 4 | 1.5064236 | 12.5 | 0 | 4 | 0.9852263 | 0.30625 | WNK lysine deficient protein kinase 4; Serine/threonine kinase which |
| ARHGEF26 | Q9H9R2 | 168 | 9606.ENSP | 3.3539668 | 0.1428571 | 0.2981544 | 7 | 24096 | 7 | 0.0014198 | 23.428571 | 0 | 7 | 0.9871367 | 0.1825396825396825 | Rho guanine nucleotide exchange factor (GEF) 26; Activates RhoG GT |
| DOCK1 | Q14185 | 168 | 9606.ENSP | 2.9677419 | 0.1052631 | 0.3369565 | 7 | 114662 | 20 | 0.0088683 | 25.25 | 0 | 20 | 0.9892473 | 0.0863321799307958 | Dedicator of cytokinesis protein 1; Involved in cytoskeletal rearrange |
| MSI2 | Q96DH6 | 169 | 9606.ENSP | 3.4080209 | 0.0 | 0.2934254 | 7 | 878 | 2 | 6.2512592 | 50.5 | 0 | 2 | 0.9868414 | 0.5 | RNA-binding protein Musashi homolog 2; RNA binding protein that re |
| SNTA1 | Q13424 | 169 | 9606.ENSP | 3.2528334 | 0.1111111 | 0.3074242 | 8 | 29694 | 9 | 0.0020416 | 20.888888 | 0 | 9 | 0.9876894 | 0.1407914764079147 | 59 kDa dystrophin-associated protein A1 acidic component 1; Adapte |
| LPP | Q93052 | 170 | 9606.ENSP | 3.6102877 | 0.3333333 | 0.2769862 | 6 | 198 | 3 | 1.3617401 | 23.0 | 0 | 3 | 0.9857361 | 0.3908045977011494 | LIM domain containing preferred translocation partner in lipoma; Ma |
| VASP | P50552 | 170 | 9606.ENSP | 2.8474280 | 0.2229885 | 0.3511941 | 6 | 69426 | 30 | 0.0042148 | 35.033333 | 0 | 30 | 0.9899047 | 0.0991477272727272 | Vasodilator-stimulated phosphoprotein; Ena/VASP proteins are actin |
| ARHGAP11 | Q6P4F7 | 171 | 9606.ENSP | 3.6564952 | 0.0476190 | 0.2734859 | 7 | 26744 | 7 | 0.0028454 | 7.2857142 | 0 | 7 | 0.9854836 | 0.1602787456445993 | Rho GTPase activating protein 11A |
| C11orf96 | Q7Z7L8 | 172 | 9606.ENSP | 4.9773321 | 0.0 | 0.2009108 | 9 | 0 | 1 | 0.0 | 2.0 | 0 | 1 | 0.9782659 | 0.0 | Chromosome 11 open reading frame 96 |
| TMCO3 | Q6UWJ1 | 172 | 9606.ENSP | 3.9782040 | 0.0 | 0.2513697 | 8 | 21288 | 2 | 0.0017436 | 12.5 | 0 | 2 | 0.9837256 | 0.5 | Transmembrane and coiled-coil domain-containing protein 3; Probab |
| AEBP1 | Q8IUX7 | 173 | 9606.ENSP | 3.3400174 | 0.1 | 0.2993996 | 7 | 5932 | 5 | 4.1689286 | 32.4 | 0 | 5 | 0.9872130 | 0.2427480916030534 | Adipocyte enhancer-binding protein 1; May positively regulate MAP- |
| MOCS1 | Q9NZB8 | 173 | 9606.ENSP | 3.7445510 | 0.0476190 | 0.2670547 | 8 | 4150 | 7 | 6.0371679 | 9.2857142 | 0 | 7 | 0.9850024 | 0.1503759398496240 | Molybdenum cofactor synthesis-step 1 protein A-B; Isoform MOCS1A |
| ITSN2 | Q9NZM3 | 174 | 9606.ENSP | 2.9319965 | 0.0441176 | 0.3410645 | 6 | 79980 | 17 | 0.0082595 | 24.647058 | 0 | 17 | 0.9894426 | 0.0802675585284281 | SH3P18-like WASP-associated protein; Adapter protein that may prov |
| CYP21A2 | P08686 | 175 | 9606.ENSP | 3.5248474 | 0.1785714 | 0.2837002 | 8 | 27930 | 8 | 0.0023893 | 14.875 | 0 | 8 | 0.9862030 | 0.1657303370786516 | Cytochrome P450, family 21, subfamily A, polypeptide 2; Specifically |
| CYP4F11 | Q9HBI6 | 175 | 9606.ENSP | 4.5239755 | 0.0 | 0.2210445 | 9 | 0 | 1 | 0.0 | 8.0 | 0 | 1 | 0.9807433 | 0.0 | Cytochrome P450, family 4, subfamily F, polypeptide 11; Omega-hydr |
| PLEKHB1 | Q9UF11 | 176 | 9606.ENSP | 4.0627724 | 0.0 | 0.2461373 | 8 | 19382 | 4 | 0.0019509 | 6.5 | 0 | 4 | 0.9832635 | 0.25 | Pleckstrin homology domain containing, family B (evectins) member |
| FAM114A1 | Q8IWE2 | 177 | 9606.ENSP | 3.9093286 | 0.0 | 0.2557983 | 8 | 26256 | 2 | 0.0017436 | 12.0 | 0 | 2 | 0.9841020 | 0.5 | Family with sequence similarity 114, member A1; May play a role in r |
| KLHL5 | Q9NV27 | 177 | 9606.ENSP | 4.9084568 | 0.0 | 0.2037300 | 9 | 0 | 1 | 0.0 | 2.0 | 0 | 1 | 0.9786423 | 0.0 | Kelch like family member 5; BTB domain containing |
| CTNS | O60931 | 178 | 9606.ENSP | 3.4873583 | 0.0 | 0.28675 | 7 | 11188 | 4 | 6.7634173 | 21.25 | 0 | 4 | 0.9864078 | 0.262987012987013 | Cystinosin, lysosomal cystine transporter; Cystine/H(+) symporter tho |
| PIRT | P0C851 | 178 | 9606.ENSP | 4.1769834 | 0.0 | 0.2394072 | 8 | 896 | 3 | 1.0994965 | 6.3333333 | 0 | 3 | 0.9826394 | 0.3333333333333333 | Phosphoinositide-interacting regulator of transient receptor potentia |
| BTN2A2 | Q8WVV5 | 179 | 9606.ENSP | 4.0854402 | 0.0 | 0.2447716 | 8 | 15720 | 2 | 0.0012023 | 9.0 | 0 | 2 | 0.9831396 | 0.5 | Butyrophilin, subfamily 2, member A2; Inhibits the proliferation of CD |
| TTC23 | Q5W5X9 | 179 | 9606.ENSP | 4.9093286 | 0.0 | 0.2036938 | 9 | 166 | 2 | 3.0599318 | 3.5 | 0 | 2 | 0.9786375 | 0.5 | Cervical cancer proto-oncogene 8 protein; Tetratricopeptide repeat d |
| GSDMD | P57764 | 180 | 9606.ENSP | 3.2676547 | 0.25 | 0.3060298 | 7 | 5716 | 8 | 5.1562038 | 28.25 | 0 | 8 | 0.9876084 | 0.1707317073170731 | Gasdermin domain-containing protein 1; Gasdermin-D, N-terminal: P |
| LRRC3 | Q9BY71 | 180 | 9606.ENSP | 3.6425457 | 0.0 | 0.2745332 | 7 | 68 | 2 | 3.1294388 | 30.0 | 0 | 2 | 0.9855598 | 0.5087719298245614 | Leucine-rich repeat-containing protein 3; Leucine rich repeat contain |
| SOWAHC | Q53LP3 | 181 | 9606.ENSP | 1.0 | 0.0 | 1.0 | 1 | 0 | 1 | 0.0 | 1.0 | 0 | 1 | 1.0 | 0.0 | Sosondowah ankyrin repeat domain family member C; Belongs to the |
| SYDE1 | Q6ZW31 | 181 | 9606.ENSP | 1.0 | 0.0 | 1.0 | 1 | 0 | 1 | 0.0 | 1.0 | 0 | 1 | 1.0 | 0.0 | Synapse defective 1, Rho GTPase, homolog 1 (C. elegans); GTPase act |
| CARD10 | Q9BWT7 | 182 | 9606.ENSP | 3.8108108 | 0.0 | 0.2624113 | 8 | 272 | 2 | 2.6706983 | 15.5 | 0 | 2 | 0.9846403 | 0.5 | Caspase recruitment domain-containing protein 10; Activates NF-kap |
| LPAR1 | Q92633 | 182 | 9606.ENSP | 2.9433304 | 0.2747252 | 0.3397511 | 7 | 47048 | 14 | 0.0031632 | 39.071428 | 0 | 14 | 0.9893807 | 0.1286660359508041 | Lysophosphatidic acid receptor Edg-2; Receptor for lysophosphatidic |
| CCDC15 | Q0P6D6 | 183 | 9606.ENSP | 3.7000871 | 0.0 | 0.2702639 | 8 | 7700 | 4 | 5.8735849 | 11.5 | 0 | 4 | 0.9852454 | 0.2625 | Coiled-coil domain containing 15 |
| NSRP1 | Q9H0G5 | 183 | 9606.ENSP | 4.2659110 | 0.1666666 | 0.2344165 | 9 | 886 | 4 | 1.4026293 | 8.25 | 0 | 4 | 0.9821534 | 0.3369565217391304 | Nuclear speckle splicing regulatory protein 1; RNA-binding protein th |
| BACE2 | Q9Y5Z0 | 184 | 9606.ENSP | 4.0043591 | 0.0 | 0.2497278 | 8 | 4652 | 2 | 4.0608392 | 10.5 | 0 | 2 | 0.9835827 | 0.5 | Beta-site amyloid precursor protein cleaving enzyme 2; Responsible f |
| CACHD1 | Q5VU97 | 184 | 9606.ENSP | 4.5196163 | 0.0 | 0.2212577 | 9 | 1508 | 3 | 2.2714681 | 4.3333333 | 0 | 3 | 0.9807671 | 0.3333333333333333 | VWFA and cache domain-containing protein 1; May regulate voltage- |
| GMPR | P36959 | 185 | 9606.ENSP | 3.6198779 | 0.0 | 0.2762524 | 8 | 3530 | 6 | 3.9570644 | 10.666666 | 0 | 6 | 0.9856837 | 0.1726190476190476 | Guanosine 5'-monophosphate oxidoreductase 1; Catalyzes the irreve |
| NUDT16 | Q96DE0 | 185 | 9606.ENSP | 3.6085440 | 0.3333333 | 0.2771200 | 7 | 1414 | 3 | 1.0591250 | 19.333333 | 0 | 3 | 0.9857456 | 0.3653846153846153 | Nudix (nucleoside diphosphate linked moiety X)-type motif 16; RNA-b |
| AIFM3 | Q96NN9 | 186 | 9606.ENSP | 4.1551874 | 0.0 | 0.2406630 | 8 | 0 | 1 | 0.0 | 13.0 | 0 | 1 | 0.9827585 | 0.0 | Apoptosis-inducing factor, mitochondrion-associated, 3; Induces apo |
| SMTN | Q9UIT2 | 186 | 9606.ENSP | 3.1560592 | 0.3076923 | 0.3168508 | 7 | 26480 | 13 | 0.0023197 | 21.923076 | 0 | 13 | 0.9882182 | 0.1141700404858299 | Smoothelin; Structural protein of the cytoskeleton; Belongs to the sm |
| FIBIN | Q8TAL6 | 187 | 9606.ENSP | 3.7872711 | 0.0 | 0.2640423 | 8 | 6072 | 6 | 9.6317287 | 6.6666666 | 0 | 6 | 0.9847690 | 0.1666666666666666 | Fin bud initiation factor homolog; Belongs to the FIBIN family. |
| HVCN1 | Q96D96 | 187 | 9606.ENSP | 3.9354838 | 0.0 | 0.2540983 | 9 | 1466 | 3 | 2.4276125 | 14.0 | 0 | 3 | 0.9839591 | 0.3333333333333333 | Voltage sensor domain-only protein; Mediates the voltage-depender |
| MIDN | Q504T8 | 188 | 9606.ENSP | 3.9450741 | 0.0 | 0.2534806 | 8 | 12 | 2 | 2.1708770 | 7.5 | 0 | 2 | 0.9839066 | 0.5 | Midbrain nucleolar protein; Facilitates ubiquitin-independent protea |
| ODF3L2 | Q3SX64 | 188 | 9606.ENSP | 3.1743679 | 0.0 | 0.3150233 | 7 | 5996 | 2 | 5.3966972 | 92.5 | 0 | 2 | 0.9881182 | 0.5 | Outer dense fiber of sperm tails 3 like 2 |
| DHRS3 | O75911 | 189 | 9606.ENSP | 3.6408020 | 0.0 | 0.2746647 | 8 | 870 | 3 | 5.1783268 | 15.0 | 0 | 3 | 0.9855693 | 0.3414634146341463 | Short chain dehydrogenase/reductase family 16C member 1; Catalyze |
| ELP4 | Q96EB1 | 189 | 9606.ENSP | 3.5518744 | 0.0476190 | 0.2815414 | 7 | 6262 | 7 | 6.6523214 | 12.285714 | 0 | 7 | 0.9860553 | 0.150278293135436 | Elongator acetyltransferase complex subunit 4; Acts as subunit of the |
| ATOH8 | Q96SQ7 | 190 | 9606.ENSP | 3.6809067 | 0.0 | 0.2716721 | 7 | 2186 | 4 | 2.1595928 | 12.75 | 0 | 4 | 0.9853502 | 0.25 | Class A basic helix-loop-helix protein 21; Transcription factor that bin |
| OLFML1 | Q6UWY5 | 190 | 9606.ENSP | 3.6904969 | 0.0 | 0.2709662 | 8 | 3904 | 6 | 4.7997590 | 8.6666666 | 0 | 6 | 0.9852978 | 0.1666666666666666 | Olfactomedin-like protein 1; Olfactomedin like 1 |
| LRRIQ1 | Q96JM4 | 191 | 9606.ENSP | 1.0 | 0.0 | 1.0 | 1 | 0 | 1 | 0.0 | 1.0 | 0 | 1 | 1.0 | 0.0 | Leucine rich repeats and IQ motif containing 1 |
| SLC44A3 | Q8N4M1 | 191 | 9606.ENSP | 1.0 | 0.0 | 1.0 | 1 | 0 | 1 | 0.0 | 1.0 | 0 | 1 | 1.0 | 0.0 | Choline transporter-like protein 3; Solute carrier family 44 member 3 |
| CLCA4 | Q14CN2 | 192 | 9606.ENSP | 3.9311246 | 0.0 | 0.2543801 | 7 | 3834 | 5 | 5.0409670 | 7.4 | 0 | 5 | 0.9839829 | 0.2133333333333333 | Calcium-activated chloride channel family member 4; May be involve |
| FCGBP | A0A087W | 192 | 9606.ENSP | 3.6782911 | 0.0 | 0.2718653 | 7 | 3898 | 3 | 2.8334674 | 15.333333 | 0 | 3 | 0.9853645 | 0.3495934959349593 | Fc fragment of IgG binding protein |
| FGF1 | P05230 | 193 | 9606.ENSP | 2.6634699 | 0.3554006 | 0.3754500 | 7 | 170090 | 42 | 0.0100098 | 58.047619 | 0 | 42 | 0.9909100 | 0.1157684630738523 | Fibroblast growth factor 1 (acidic); Plays an important role in the regu |
| FGFRL1 | Q8N441 | 193 | 9606.ENSP | 3.5666957 | 1.0 | 0.2803715 | 8 | 0 | 2 | 0.0 | 35.5 | 0 | 2 | 0.9859743 | 0.6120689655172413 | Fibroblast growth factor receptor-like 1; Has a negative effect on cell |
| FAM111A | Q96PZ2 | 194 | 9606.ENSP | 3.6216216 | 0.0 | 0.2761194 | 8 | 20618 | 2 | 0.0017436 | 21.5 | 0 | 2 | 0.9856741 | 0.5 | Family with sequence similarity 111, member A; Chromatin-associate |
| NKAIN4 | Q8IVV8 | 194 | 9606.ENSP | 4.6207497 | 0.0 | 0.2164150 | 9 | 0 | 1 | 0.0 | 2.0 | 0 | 1 | 0.9802144 | 0.0 | Sodium/potassium-transporting ATPase subunit beta-1-interacting pr |
| PSKH1 | P11801 | 195 | 9606.ENSP | 3.8936355 | 0.0 | 0.2568293 | 7 | 1660 | 3 | 2.2004562 | 10.333333 | 0 | 3 | 0.9841877 | 0.3333333333333333 | Serine/threonine-protein kinase H1; May be a SFC-associated serine |
| ZCCHC24 | Q8N2G6 | 195 | 9606.ENSP | 4.0148212 | 0.1 | 0.2490770 | 7 | 2196 | 5 | 3.4414480 | 5.4 | 0 | 5 | 0.9835255 | 0.2181818181818181 | Zinc finger CCHC-type containing 24; Zinc fingers 3CxxC-type |
| ETNPPL | Q8TBG4 | 196 | 9606.ENSP | 4.0802092 | 0.0 | 0.2450854 | 8 | 33740 | 3 | 0.0018413 | 7.6666666 | 0 | 3 | 0.9831682 | 0.3333333333333333 | Alanine--glyoxylate aminotransferase 2-like 1; Catalyzes the pyridoxa |
| PHYHD1 | Q96GM4 | 196 | 9606.ENSP | 5.0793374 | 0.0 | 0.1968760 | 9 | 0 | 1 | 0.0 | 3.0 | 0 | 1 | 0.9777085 | 0.0 | Phytanoyl-CoA dioxygenase domain-containing protein 1; Isoform 1 h |
| FGR | P09769 | 197 | 9606.ENSP | 2.7916303 | 0.1862068 | 0.3582136 | 6 | 75934 | 30 | 0.0047366 | 39.9 | 0 | 30 | 0.9902096 | 0.0992518703241895 | Gardner-Rasheed feline sarcoma viral (v-fgr) oncogene homolog; Nor |
| SIGLEC9 | Q9Y336 | 197 | 9606.ENSP | 3.3147340 | 0.3333333 | 0.3016833 | 7 | 120 | 3 | 4.2842575 | 53.0 | 0 | 3 | 0.9873511 | 0.4082687338501292 | Sialic acid binding Ig-like lectin 9; Putative adhesion molecule that me |
| ANKRD36 | A6QL64 | 198 | 9606.ENSP | 1.0 | 0.0 | 1.0 | 1 | 0 | 1 | 0.0 | 1.0 | 0 | 1 | 1.0 | 0.0 | Ankyrin repeat domain-containing protein 36A; Ankyrin repeat doma |
| ANKRD36C | Q5JPF3 | 198 | 9606.ENSP | 1.0 | 0.0 | 1.0 | 1 | 0 | 1 | 0.0 | 1.0 | 0 | 1 | 1.0 | 0.0 | Protein immuno-reactive with anti-PTH polyclonal antibodies; Ankyri |
| RUFY4 | Q6ZNE9 | 199 | 9606.ENSP | 4.3827375 | 0.0 | 0.2281678 | 9 | 0 | 1 | 0.0 | 3.0 | 0 | 1 | 0.9815150 | 0.0 | RUN and FYVE domain containing 4; Zinc fingers FYVE-type |
| TMBIM1 | Q969X1 | 199 | 9606.ENSP | 3.3836094 | 0.0 | 0.2955423 | 8 | 17764 | 3 | 0.0017488 | 35.666666 | 0 | 3 | 0.9869748 | 0.3365695792880259 | Transmembrane BAX inhibitor motif-containing protein 1; Negatively |
| CSRNP1 | Q96S65 | 200 | 9606.ENSP | 3.7890148 | 0.0 | 0.2639208 | 8 | 9116 | 3 | 7.2212966 | 15.666666 | 0 | 3 | 0.9847594 | 0.3333333333333333 | Cysteine-serine-rich nuclear protein 1; Binds to the consensus sequen |
| RASSF2 | P50749 | 200 | 9606.ENSP | 4.0575414 | 0.0 | 0.2464546 | 7 | 62620 | 5 | 0.0050950 | 4.4 | 0 | 5 | 0.9832921 | 0.2 | Ras association (RalGDS/AF-6) domain family member 2; Potential tu |
| SELENBP1 | Q13228 | 201 | 9606.ENSP | 3.4847428 | 0.0666666 | 0.2869652 | 8 | 17342 | 6 | 0.0011831 | 11.833333 | 0 | 6 | 0.9864221 | 0.1801075268817204 | 56 kDa selenium-binding protein; Selenium-binding protein which ma |
| LGALS9 | O00182 | 202 | 9606.ENSP | 3.0026155 | 0.4166666 | 0.3330429 | 7 | 15354 | 9 | 9.0894844 | 72.0 | 0 | 9 | 0.9890567 | 0.2396296296296296 | Lectin, galactoside-binding, soluble, 9; Binds galactosides. Has high af |
| SLC12A7 | Q9Y666 | 202 | 9606.ENSP | 3.5754141 | 0.1428571 | 0.2796878 | 8 | 8872 | 7 | 7.4974720 | 9.1428571 | 0 | 7 | 0.9859266 | 0.1736694677871148 | Solute carrier family 12 (potassium/chloride transporter), member 7; |
| LCN10 | E9PRR3 | 203 | 9606.ENSP | 1.0 | 0.0 | 1.0 | 1 | 0 | 1 | 0.0 | 1.0 | 0 | 1 | 1.0 | 0.0 | Epididymal-specific lipocalin-10; Lipocalin 10; Belongs to the calycin s |
| LCN6 | P62502 | 203 | 9606.ENSP | 1.0 | 0.0 | 1.0 | 1 | 0 | 1 | 0.0 | 1.0 | 0 | 1 | 1.0 | 0.0 | Epididymal-specific lipocalin-6; May play a role in male fertility; Lipoca |
| IMPA2 | O14732 | 204 | 9606.ENSP | 4.2807323 | 0.1666666 | 0.2336048 | 8 | 760 | 4 | 1.2386727 | 5.75 | 0 | 4 | 0.9820725 | 0.328125 | Inositol(myo)-1(or 4)-monophosphatase 2; Can use myo-inositol mon |
| PXDC1 | Q5TGL8 | 204 | 9606.ENSP | 3.9180470 | 0.0 | 0.2552291 | 8 | 3110 | 3 | 3.3676761 | 9.6666666 | 0 | 3 | 0.9840543 | 0.3333333333333333 | PX domain containing 1 |
| PTTG1IP | P53801 | 205 | 9606.ENSP | 5.7593722 | 0.0 | 0.1736300 | 10 | 0 | 1 | 0.0 | 2.0 | 0 | 1 | 0.9739925 | 0.0 | Pituitary tumor-transforming gene 1 protein-interacting protein; May |
| TPCN1 | Q9ULQ1 | 205 | 9606.ENSP | 4.7602441 | 0.0 | 0.2100732 | 9 | 21218 | 2 | 0.0017436 | 3.5 | 0 | 2 | 0.9794522 | 0.5 | Voltage-dependent calcium channel protein TPC1; Nicotinic acid aden |
| PRSS35 | Q8N3Z0 | 206 | 9606.ENSP | 4.3504795 | 0.0 | 0.2298597 | 7 | 1138 | 2 | 1.1835964 | 6.5 | 0 | 2 | 0.9816913 | 0.5 | Serine protease 35 |
| RANBP3L | Q86VV4 | 206 | 9606.ENSP | 4.4995640 | 0.0 | 0.2222437 | 7 | 386 | 2 | 4.9359991 | 4.5 | 0 | 2 | 0.9808766 | 0.5 | RAN binding protein 3-like; Nuclear export factor for BMP-specific SM |
| ARPC1B | O15143 | 207 | 9606.ENSP | 3.2894507 | 0.0357142 | 0.3040021 | 7 | 7328 | 8 | 6.6392246 | 20.875 | 0 | 8 | 0.9874893 | 0.1490740740740740 | Actin related protein 2/3 complex, subunit 1B, 41kDa; Functions as co |
| CAPG | P40121 | 207 | 9606.ENSP | 3.3748910 | 0.5 | 0.2963058 | 7 | 614 | 4 | 4.6763487 | 38.0 | 0 | 4 | 0.9870224 | 0.3595238095238095 | Capping protein (actin filament), gelsolin-like; Calcium-sensitive prote |
| KTN1 | Q86UP2 | 208 | 9606.ENSP | 4.0052310 | 0.0 | 0.2496734 | 8 | 17730 | 2 | 0.0017436 | 7.0 | 0 | 2 | 0.9835779 | 0.5 | Kinectin 1 (kinesin receptor); Receptor for kinesin thus involved in kin |
| PPP1R14B | Q96C90 | 208 | 9606.ENSP | 5.0043591 | 0.0 | 0.1998257 | 9 | 0 | 1 | 0.0 | 2.0 | 0 | 1 | 0.9781182 | 0.0 | Protein phosphatase 1, regulatory (inhibitor) subunit 14B; Inhibitor o |
| ASAP3 | Q8TDY4 | 209 | 9606.ENSP | 3.5283347 | 0.0 | 0.2834198 | 7 | 1090 | 2 | 6.5506125 | 32.0 | 0 | 2 | 0.9861839 | 0.5081967213114754 | Arf-GAP with SH3 domain, ANK repeat and PH domain-containing pro |
| TCEA3 | O75764 | 209 | 9606.ENSP | 3.6669572 | 0.0 | 0.2727056 | 8 | 8628 | 3 | 5.7589508 | 15.333333 | 0 | 3 | 0.9854264 | 0.3412698412698413 | Transcription elongation factor S-II protein 3; Necessary for efficient |
| ANPEP | P15144 | 210 | 9606.ENSP | 2.7489102 | 0.4344086 | 0.3637805 | 6 | 145324 | 31 | 0.0080142 | 60.709677 | 0 | 31 | 0.9904431 | 0.1361362812613265 | Myeloid plasma membrane glycoprotein CD13; Broad specificity amin |
| PGPEP1 | Q9NXJ5 | 210 | 9606.ENSP | 3.5675675 | 0.0 | 0.2803030 | 7 | 26 | 2 | 9.0976993 | 36.5 | 0 | 2 | 0.9859695 | 0.6016949152542372 | Pyrrolidone-carboxylate peptidase; Removes 5-oxoproline from vario |
| ZNF423 | Q2M1K9 | 211 | 9606.ENSP | 3.1987794 | 0.1785714 | 0.3126192 | 6 | 14706 | 8 | 0.0010885 | 31.375 | 0 | 8 | 0.9879848 | 0.1778571428571428 | Smad- and Olf-interacting zinc finger protein; Transcription factor tha |
| BBX | Q8WY36 | 212 | 9606.ENSP | 3.7253705 | 0.0 | 0.2684296 | 8 | 27378 | 5 | 0.0021123 | 10.4 | 0 | 5 | 0.9851072 | 0.2088888888888889 | HMG box transcription factor BBX; Transcription factor that is necess |
| NACC2 | Q96BF6 | 212 | 9606.ENSP | 4.7244986 | 0.0 | 0.2116626 | 9 | 0 | 1 | 0.0 | 5.0 | 0 | 1 | 0.9796475 | 0.0 | NACC family member 2, BEN and BTB (POZ) domain containing; Func |
| PLSCR4 | Q9NRQ2 | 213 | 9606.ENSP | 4.0924149 | 0.0 | 0.2443544 | 7 | 434 | 2 | 4.2981091 | 8.5 | 0 | 2 | 0.9831015 | 0.5 | Ca(2+)-dependent phospholipid scramblase 4; May mediate accelera |
| RELL1 | Q8IUW5 | 213 | 9606.ENSP | 3.8526591 | 0.0 | 0.2595609 | 8 | 5766 | 4 | 5.5391506 | 12.75 | 0 | 4 | 0.9844116 | 0.2554347826086957 | RELT-like protein 1; RELT family; Belongs to the RELT family. |
| BUB1 | O43683 | 214 | 9606.ENSP | 3.1194420 | 0.1818181 | 0.3205701 | 7 | 40968 | 12 | 0.0031631 | 25.75 | 0 | 12 | 0.9884183 | 0.1272802653399668 | Mitotic checkpoint serine/threonine-protein kinase BUB1; Serine/thr |
| CDC45 | O75419 | 214 | 9606.ENSP | 3.3269398 | 0.4761904 | 0.3005765 | 6 | 8808 | 7 | 6.1826094 | 26.142857 | 0 | 7 | 0.9872844 | 0.2203389830508474 | Cell division control protein 45 homolog; Required for initiation of ch |
| TNFRSF12A | Q9NP84 | 215 | 9606.ENSP | 2.9598953 | 0.3267973 | 0.3378497 | 7 | 37126 | 18 | 0.0022204 | 50.222222 | 0 | 18 | 0.9892901 | 0.1600993965211217 | Fibroblast growth factor-inducible immediate-early response protein |
| PCK1 | P35558 | 216 | 9606.ENSP | 3.1830863 | 0.2205882 | 0.3141605 | 7 | 32200 | 17 | 0.0023686 | 20.235294 | 0 | 17 | 0.9880705 | 0.1117647058823529 | Phosphoenolpyruvate carboxykinase, cytosolic [GTP]; Catalyzes the c |
| RBM38 | Q9H0Z9 | 216 | 9606.ENSP | 3.5361813 | 0.0 | 0.2827909 | 8 | 288 | 2 | 1.0708510 | 31.0 | 0 | 2 | 0.9861410 | 0.5172413793103449 | RNA-binding region-containing protein 1; RNA-binding protein that sp |

| | | | | | | | | | | | | | | | | | |
|---|---|---|---|---|---|---|---|---|---|---|---|---|---|---|---|---|---|
| SGMS2 | Q8NHU3 | 217 | | 9606.ENSP | 3.4176111 | 0.0 | 0.2926020 | 8 | 20282 | 2 | 0.0017436 | 46.5 | 0 | 2 | 0.9867890 | 0.5 | Phosphatidylcholine:ceramide cholinephosphotransferase 2; Sphingo |
| USP53 | Q70EK8 | 217 | | 9606.ENSP | 4.4167393 | 0.0 | 0.2264113 | 9 | 0 | 1 | 0.0 | 2.0 | 0 | 1 | 0.9813292 | 0.0 | Inactive ubiquitin carboxyl-terminal hydrolase 53; Tight junction-asso |
| ACVRL1 | P37023 | 218 | | 9606.ENSP | 2.9755884 | 0.4327485 | 0.3360679 | 7 | 54342 | 19 | 0.0027690 | 49.736842 | 0 | 19 | 0.9892044 | 0.1659919028340081 | Serine/threonine-protein kinase receptor R3; Type I receptor for TGF |
| TMEM100 | Q9NV29 | 218 | | 9606.ENSP | 3.9747166 | 0.0 | 0.2515902 | 8 | 0 | 1 | 0.0 | 19.0 | 0 | 1 | 0.9837447 | 0.0 | Transmembrane protein 100; Plays a role during embryonic arterial e |
| C16orf74 | Q96GX8 | 219 | | 9606.ENSP | 4.2929380 | 0.0 | 0.2329406 | 8 | 0 | 1 | 0.0 | 9.0 | 0 | 1 | 0.9820058 | 0.0 | Chromosome 16 open reading frame 74 |
| FOXF1 | Q12946 | 219 | | 9606.ENSP | 3.2938099 | 0.3611111 | 0.3035997 | 7 | 38426 | 9 | 0.0018497 | 32.555555 | 0 | 9 | 0.9874655 | 0.2309523809523809 | Forkhead-related transcription factor 1; Probable transcription activa |
| C1R | P00736 | 220 | | 9606.ENSP | 2.9755884 | 0.3593073 | 0.3360679 | 7 | 29932 | 22 | 0.0017989 | 31.227272 | 0 | 22 | 0.9892044 | 0.1108055645422193 | Complement component 1, r subcomponent; C1r B chain is a serine p |
| VAMP5 | O95183 | 220 | | 9606.ENSP | 3.6521360 | 0.0 | 0.2738123 | 8 | 188 | 2 | 9.0051092 | 36.0 | 0 | 2 | 0.9855074 | 0.5 | Vesicle-associated membrane protein 5; May participate in trafficking |
| ZNF618 | B5MDS3 | 221 | | 9606.ENSP | 4.5719267 | 0.0 | 0.2187261 | 9 | 0 | 1 | 0.0 | 6.0 | 0 | 1 | 0.9804812 | 0.0 | Zinc finger protein 618; Zinc fingers C2H2-type |
| BCL7C | Q8WUZ0 | 222 | | 9606.ENSP | 3.4551002 | 0.1666666 | 0.2894272 | 7 | 522 | 4 | 4.1413896 | 21.5 | 0 | 4 | 0.9865841 | 0.2625 | B-cell CLL/lymphoma 7 protein family member C; May play an anti-ap |
| SMARCC1 | Q92922 | 222 | | 9606.ENSP | 2.9755884 | 0.1754385 | 0.3360679 | 7 | 56198 | 19 | 0.0039067 | 27.894736 | 0 | 19 | 0.9892044 | 0.0953156086091517 | SWI/SNF related, matrix associated, actin dependent regulator of chr |
| BCAM | P50895 | 223 | | 9606.ENSP | 3.4324324 | 0.0 | 0.2913385 | 7 | 1576 | 4 | 1.3654601 | 19.0 | 0 | 4 | 0.9867080 | 0.2535211267605634 | Basal cell adhesion molecule (Lutheran blood group); Laminin alpha-5 |
| C1QTNF1 | Q9BXJ1 | 224 | | 9606.ENSP | 3.7968613 | 0.0 | 0.2633754 | 8 | 0 | 1 | 0.0 | 29.0 | 0 | 1 | 0.9847166 | 0.0 | Complement C1q tumor necrosis factor-related protein 1; C1q and TI |
| CSPG4 | Q6UVK1 | 224 | | 9606.ENSP | 2.7977332 | 0.2758620 | 0.3574322 | 7 | 81082 | 29 | 0.0044808 | 51.551724 | 0 | 29 | 0.9901763 | 0.1262677484787018 | Melanoma-associated chondroitin sulfate proteoglycan; Proteoglycan |
| CDA | P32320 | 225 | | 9606.ENSP | 3.5396687 | 0.0952380 | 0.2825123 | 8 | 27364 | 7 | 0.0019646 | 13.428571 | 0 | 7 | 0.9861220 | 0.1688311688311688 | Cytidine aminohydrolase; This enzyme scavenges exogenous and enc |
| TTC38 | Q5R3I4 | 225 | | 9606.ENSP | 4.5387968 | 0.0 | 0.2203227 | 9 | 0 | 1 | 0.0 | 7.0 | 0 | 1 | 0.9806623 | 0.0 | Tetratricopeptide repeat domain containing; Belongs to the TTC38 fa |
| ARHGAP31 | Q2M1Z3 | 226 | | 9606.ENSP | 3.5030514 | 0.0 | 0.2854654 | 7 | 2476 | 4 | 2.2465340 | 23.25 | 0 | 4 | 0.9863221 | 0.2816455696202531 | Rho GTPase activating protein 31; Functions as a GTPase-activating p |
| DOCK6 | Q96HP0 | 226 | | 9606.ENSP | 3.7061900 | 0.1 | 0.2698188 | 8 | 6234 | 5 | 5.5877590 | 9.4 | 0 | 5 | 0.9852120 | 0.2256410256410256 | Dedicator of cytokinesis protein 6; Acts as guanine nucleotide exchan |
| CHI3L2 | Q15782 | 227 | | 9606.ENSP | 1.0 | 0.0 | 1.0 | 1 | 0 | 1 | 0.0 | 1.0 | 0 | 1 | 1.0 | 0.0 | Chitinase-3-like protein 2; Lectin that binds chitooligosaccharides and |
| PIFO | Q8TCI5 | 227 | | 9606.ENSP | 1.0 | 0.0 | 1.0 | 1 | 0 | 1 | 0.0 | 1.0 | 0 | 1 | 1.0 | 0.0 | Primary cilia formation; During primary cilia disassembly, involved in |
| NID1 | P14543 | 228 | | 9606.ENSP | 2.9546643 | 0.4492753 | 0.3384479 | 7 | 78366 | 24 | 0.0038290 | 47.833333 | 0 | 24 | 0.9893187 | 0.1554017372421281 | Nidogen 1; Sulfated glycoprotein widely distributed in basement mer |
| ZAN | Q9Y493 | 228 | | 9606.ENSP | 3.9537925 | 0.0 | 0.2529217 | 8 | 0 | 1 | 0.0 | 24.0 | 0 | 1 | 0.9838590 | 0.0 | Zonadhesin (gene/pseudogene); Binds in a species-specific manner to |
| IER3 | P46695 | 229 | | 9606.ENSP | 3.2231909 | 0.4545454 | 0.3102515 | 7 | 38066 | 12 | 0.0017833 | 33.083333 | 0 | 12 | 0.9878514 | 0.1941176470588235 | Radiation-inducible immediate-early gene IEX-1; May play a role in th |
| PPP1R18 | Q6ZUJ6 | 229 | | 9606.ENSP | 4.2223190 | 0.0 | 0.2368366 | 8 | 0 | 1 | 0.0 | 12.0 | 0 | 1 | 0.9823916 | 0.0 | Protein phosphatase 1 F-actin cytoskeleton-targeting subunit; Isoforr |
| RIBC1 | Q8N443 | 230 | | 9606.ENSP | 4.8334786 | 0.0 | 0.2068903 | 9 | 0 | 1 | 0.0 | 3.0 | 0 | 1 | 0.9790520 | 0.0 | RIB43A domain with coiled-coils 1; Belongs to the RIB43A family. |
| TOB2 | Q14106 | 230 | | 9606.ENSP | 3.8343504 | 0.3333333 | 0.2608003 | 8 | 32534 | 3 | 0.0017436 | 13.333333 | 0 | 3 | 0.9845117 | 0.4193548387096774 | Transducer of ERBB2, 2; Anti-proliferative protein inhibits cell cycle p |
| EPAS1 | Q99814 | 231 | | 9606.ENSP | 2.7646033 | 0.2315789 | 0.3617155 | 7 | 50664 | 20 | 0.0034866 | 53.65 | 0 | 20 | 0.9903573 | 0.1228735632183908 | Endothelial PAS domain-containing protein 1; Transcription factor inv |
| HIF3A | Q9Y2N7 | 231 | | 9606.ENSP | 3.7637314 | 0.0 | 0.2656937 | 8 | 0 | 1 | 0.0 | 20.0 | 0 | 1 | 0.9848976 | 0.0 | Class E basic helix-loop-helix protein 17; Isoform 5: Attenuates the ab |
| HAPLN3 | Q96S86 | 232 | | 9606.ENSP | 3.8134263 | 0.0 | 0.2622313 | 8 | 86 | 2 | 4.5687668 | 17.5 | 0 | 2 | 0.9846260 | 0.5 | Hyaluronan and proteoglycan link protein 3; May function in hyaluron |
| MFGE8 | Q08431 | 232 | | 9606.ENSP | 3.0845684 | 0.1241830 | 0.3241944 | 7 | 25112 | 18 | 0.0016941 | 27.722222 | 0 | 18 | 0.9886089 | 0.1169962335216572 | Milk fat globule-EGF factor 8 protein; Plays an important role in the n |
| EMP3 | P54852 | 233 | | 9606.ENSP | 3.1543156 | 0.1555555 | 0.3170259 | 7 | 34614 | 10 | 0.0023993 | 34.0 | 0 | 10 | 0.9882277 | 0.1658415841584158 | Hematopoietic neural membrane protein 1; Probably involved in cell |
| PLP2 | Q04941 | 233 | | 9606.ENSP | 4.1534437 | 0.0 | 0.2407640 | 8 | 0 | 1 | 0.0 | 10.0 | 0 | 1 | 0.9827680 | 0.0 | Proteolipid protein 2 (colonic epithelium-enriched); May play a role in |
| IQCA1 | Q86XH1 | 234 | | 9606.ENSP | 6.0540540 | 0.0 | 0.1651785 | 9 | 0 | 1 | 0.0 | 2.0 | 0 | 1 | 0.9723822 | 0.0 | IQ motif containing with AAA domain 1 |
| STK17B | O94768 | 234 | | 9606.ENSP | 5.0549258 | 0.0 | 0.1978268 | 8 | 21000 | 2 | 0.0017436 | 3.0 | 0 | 2 | 0.9778419 | 0.5 | DAP kinase-related apoptosis-inducing protein kinase 2; Phosphoryla |
