## Supplementary tables for "Transcriptome profiling of the dorsomedial prefrontal cortex in suicide victims": S5_PPI_network_data_of_down-_and_upregulated_genes3.pdf

Supplementary file 5. : Upregulated DEGs assigned to clusters by STRING using MCL algorithm. Functional annotation of clusters was performed based on GO-Biological proces

| MCL cluster number | Number of genes | Functional annotation of cluster | Protein name | Protein identifier | Protein description |
|---|---|---|---|---|---|
| 1 | 34 | Synaptic signaling, Regulation of biological quality, Regulation of membrane potential | ACTL6B | ENSP00000160382 | 53 kDa BRG1-associated factor B; Involved in transcriptio |
| | | | ATP4A | ENSP00000262623 | ATPase, H+/K+ exchanging, alpha polypeptide; Catalyzes |
| | | | CACNA1G | ENSP00000352011 | Calcium channel, voltage-dependent, T type, alpha 1G su |
| | | | CACNG8 | ENSP00000270458 | Neuronal voltage-gated calcium channel gamma-8 subur |
| | | | CALB2 | ENSP00000307508 | 29 kDa calbindin; Calretinin is a calcium-binding protein |
| | | | CALY | ENSP00000252939 | Calcyon neuron-specific vesicular protein; Interacts with |
| | | | CAMKV | ENSP00000419195 | CaM kinase-like vesicle-associated protein; Does not app |
| | | | CARTPT | ENSP00000296777 | Cocaine- and amphetamine-regulated transcript protein; |
| | | | CCK | ENSP00000379472 | Cholecystokinin; This peptide hormone induces gall blad |
| | | | CHRNA10 | ENSP00000250699 | Cholinergic receptor, nicotinic, alpha 10 (neuronal); Iono |
| | | | COL11A2 | ENSP00000363840 | Collagen alpha-2(XI) chain; May play an important role ir |
| | | | COL22A1 | ENSP00000303153 | Collagen alpha-1(XXII) chain; Acts as a cell adhesion ligar |
| | | | COL24A1 | ENSP00000359603 | Collagen alpha-1(XXIV) chain; May participate in regulati |
| | | | COL28A1 | ENSP00000382356 | Collagen alpha-1(XXVIII) chain; May act as a cell-binding |
| | | | CRHR1 | ENSP00000381333 | Corticotropin releasing hormone receptor 1; G-protein c |
| | | | CYP2D6 | ENSP00000353820 | Cytochrome P450, family 2, subfamily D, polypeptide 6; |
| | | | CYP2E1 | ENSP00000440689 | Cytochrome P450, family 2, subfamily E, polypeptide 1; |
| | | | FGF8 | ENSP00000321797 | Fibroblast growth factor 8 (androgen-induced); Plays an |
| | | | GABRD | ENSP00000367848 | Gamma-aminobutyric acid (GABA) A receptor, delta; GA |
| | | | GRIK1 | ENSP00000382791 | Glutamate receptor, ionotropic, kainate 1; Ionotropic glu |
| | | | GRIK2 | ENSP00000397026 | Glutamate receptor, ionotropic, kainate 2; Ionotropic glu |
| | | | GRM2 | ENSP00000378492 | Glutamate receptor, metabotropic 2; G-protein coupled |
| | | | LPCAT4 | ENSP00000317300 | 1-alkenylglycerophosphoethanolamine O-acyltransferas |
| | | | LY6H | ENSP00000399485 | Lymphocyte antigen 6 complex, locus H; Believed to act |
| | | | MYO15A | ENSP00000205890 | Unconventional myosin-15; Myosins are actin-based mo |
| | | | NEGR1 | ENSP00000350364 | Neuronal growth regulator 1; May be involved in cell-ad |
| | | | NRGN | ENSP00000284292 | Neurogranin (protein kinase C substrate, RC3); Acts as a |
| | | | SCN3B | ENSP00000376523 | Sodium channel, voltage-gated, type III, beta subunit; M |
| | | | SHISA8 | ENSP00000481203 | Putative protein shisa-8; Shisa family member 8 |
| | | | SLC44A5 | ENSP00000359892 | Choline transporter-like protein 5; Solute carrier family 4 |
| | | | SNCG | ENSP00000361087 | Synuclein, gamma (breast cancer-specific protein 1); Pla |
| | | | SOHLH1 | ENSP00000404438 | Spermatogenesis- and oogenesis-specific basic helix-loo |
| | | | SYCE1 | ENSP00000341282 | Synaptonemal complex central element protein 1; Major |
| | | | TAC3 | ENSP00000483110 | Tachykinin 3; Tachykinins are active peptides which excit |
| 2 | 4 | Protein tetramerization, Positive regulation of cytosolic calcium ion concentration | P2RX2 | ENSP00000343339 | Purinergic receptor P2X, ligand-gated ion channel, 2; Ion |
| | | | PKD1 | ENSP00000262304 | Autosomal dominant polycystic kidney disease 1 protein |
| | | | PKD2L1 | ENSP00000325296 | Polycystic kidney disease 2-like 1 protein; Pore-forming |
| | | | TRPM2 | ENSP00000381023 | Transient receptor potential cation channel, subfamily M |
| 3 | 4 | NADPH oxidase complex | AMY2A | ENSP00000481450 | 1,4-alpha-D-glucan glucanohydrolase; Amylase, alpha 2A |
| | | | NOXA1 | ENSP00000342848 | NADPH oxidase activator 1; Functions as an activator of |
| | | | NOXO1 | ENSP00000380450 | SH3 and PX domain-containing protein 5; Constitutively |
| | | | PDIA2 | ENSP00000219406 | Protein disulfide isomerase family A, member 2; Acts as |
| 4 | 3 | | AP3B2 | ENSP00000440984 | Clathrin assembly protein complex 3 beta-2 large chain; |
| | | | MAST1 | ENSP00000251472 | Microtubule-associated serine/threonine-protein kinase |
| | | | TCTE1 | ENSP00000360560 | T-complex-associated-testis-expressed 1; Dynein regulat |
| 5 | 2 | | KRT17 | ENSP00000308452 | Keratin, type I cytoskeletal 17; Type I keratin involved in |
| | | | TGM1 | ENSP00000206765 | Protein-glutamine gamma-glutamyltransferase K; Cataly |
| 6 | 2 | | CORO1A | ENSP00000219150 | Tryptophan aspartate-containing coat protein; May be a |
| | | | CORO6 | ENSP00000344562 | Coronin-like protein E; WD repeat domain containing; C |
| 7 | 2 | | PNMA5 | ENSP00000388850 | Paraneoplastic Ma antigens; Belongs to the PNMA famil |
| | | | RTL1 | ENSP00000435342 | Mammalian retrotransposon derived protein 1; Plays an |
| 8 | 2 | | PROC | ENSP00000234071 | Protein C (inactivator of coagulation factors Va and VIIIa) |
| | | | TNNT2 | ENSP00000236918 | Troponin T type 2 (cardiac); Troponin T is the tropomyos |
| 9 | 2 | | MICAL1 | ENSP00000351664 | Microtubule associated monooxygenase, calponin and L |
| | | | SEMA5B | ENSP00000389588 | Semaphorin-5B; May act as positive axonal guidance cue |
| 10 | 2 | | CHTF18 | ENSP00000262315 | CTF18, chromosome transmission fidelity factor 18 hom |
| | | | RTEL1 | ENSP00000353332 | Regulator of telomere elongation helicase 1; ATP-depen |
| 11 | 1 | | TMEM249 | ENSP00000454468 | Transmembrane protein 249 |
| 12 | 2 | | B4GALNT3 | ENSP00000266383 | N-acetyl-beta-glucosaminyl-glycoprotein 4-beta-N-acety |
| | | | MSLNL | ENSP00000441381 | Pre-pro-megakaryocyte-potentiating-factor-like; May pla |
