## Supplementary tables for "Transcriptome profiling of the dorsomedial prefrontal cortex in suicide victims": S5_PPI_network_data_of_down-_and_upregulated_genes4.pdf

Supplementary file 5. : Upregulated DEGs assigned to clusters by STRING using MCL algorithm.

| Gene nam | String ID | Cluster | Cluster col | Protein ID | AverageSh | Clustering | Closeness | Eccentricit | Stress | Degree | Betweenn | Neighborh | Numberofl | Numberofl | Radiality | TopologicalCoefficient | Description |
|---|---|---|---|---|---|---|---|---|---|---|---|---|---|---|---|---|---|
| ACTL6B | O94805 | 1 | Soft red | 9606.ENSP | 4.0 | 1.0 | 0.25 | 8 | 0 | 2 | 0.0 | 5.0 | 0 | 2 | 0.6666666 | 0.8333333333333334 | 53 kDa BRG1-associated factor B; Involved in |
| ATP4A | P20648 | 1 | Soft red | 9606.ENSP | 3.7647058 | 0.0 | 0.265625 | 8 | 76 | 2 | 0.0588235 | 5.0 | 0 | 2 | 0.6928104 | 0.5 | ATPase, H+/K+ exchanging, alpha polypeptide |
| CACNA1G | Q9UHP0 | 1 | Soft red | 9606.ENSP | 3.3235294 | 0.8333333 | 0.3008849 | 6 | 66 | 4 | 0.0152109 | 5.5 | 0 | 4 | 0.7418300 | 0.6111111111111112 | Calcium channel, voltage-dependent, T type, |
| CACNG8 | Q8WXS5 | 1 | Soft red | 9606.ENSP | 4.0588235 | 0.8333333 | 0.2463768 | 7 | 2 | 4 | 5.9417706 | 4.75 | 0 | 4 | 0.6601307 | 0.7916666666666666 | Neuronal voltage-gated calcium channel gam |
| CALB2 | P22676 | 1 | Soft red | 9606.ENSP | 3.5588235 | 0.0 | 0.2809917 | 8 | 52 | 2 | 0.0395127 | 6.0 | 0 | 2 | 0.7156862 | 0.5 | 29 kDa calbindin; Calretinin is a calcium-bindi |
| CALY | Q9NYX4 | 1 | Soft red | 9606.ENSP | 3.7058823 | 0.6666666 | 0.2698412 | 8 | 8 | 4 | 0.0021984 | 5.5 | 0 | 4 | 0.6993464 | 0.55 | Calcyon neuron-specific vesicular protein; Int |
| CAMKV | Q8NCB2 | 1 | Soft red | 9606.ENSP | 3.6176470 | 0.0 | 0.2764227 | 7 | 0 | 1 | 0.0 | 5.0 | 0 | 1 | 0.7091503 | 0.0 | CaM kinase-like vesicle-associated protein; D |
| CARTPT | Q16568 | 1 | Soft red | 9606.ENSP | 3.2352941 | 0.6666666 | 0.3090909 | 7 | 46 | 3 | 0.0112893 | 6.0 | 0 | 3 | 0.7516339 | 0.5454545454545454 | Cocaine- and amphetamine-regulated transc |
| CCK | P06307 | 1 | Soft red | 9606.ENSP | 2.8529411 | 0.1944444 | 0.3505154 | 7 | 608 | 9 | 0.3461081 | 3.4444444 | 0 | 9 | 0.7941176 | 0.2393162393162393 | Cholecystokinin; This peptide hormone induc |
| CHRNA10 | Q9GZZ6 | 1 | Soft red | 9606.ENSP | 5.4705882 | 0.0 | 0.1827956 | 9 | 0 | 1 | 0.0 | 3.0 | 0 | 1 | 0.5032679 | 0.0 | Cholinergic receptor, nicotinic, alpha 10 (neur |
| COL11A2 | Q7Z6C3 | 1 | Soft red | 9606.ENSP | 3.7058823 | 0.3333333 | 0.2698412 | 7 | 426 | 4 | 0.2388591 | 3.0 | 0 | 4 | 0.6993464 | 0.3928571428571428 | Collagen alpha-2(XI) chain; May play an impo |
| COL22A1 | Q8NFW1 | 1 | Soft red | 9606.ENSP | 3.8235294 | 0.6666666 | 0.2615384 | 7 | 82 | 3 | 0.0249554 | 3.3333333 | 0 | 3 | 0.6862745 | 0.5555555555555556 | Collagen alpha-1(XXII) chain; Acts as a cell ad |
| COL24A1 | Q17RW2 | 1 | Soft red | 9606.ENSP | 3.0588235 | 0.1666666 | 0.3269230 | 6 | 714 | 4 | 0.3796791 | 3.75 | 0 | 4 | 0.7712418 | 0.325 | Collagen alpha-1(XXIV) chain; May participate |
| COL28A1 | Q2UY09 | 1 | Soft red | 9606.ENSP | 4.6470588 | 1.0 | 0.2151898 | 8 | 0 | 2 | 0.0 | 3.5 | 0 | 2 | 0.5947712 | 0.875 | Collagen alpha-1(XXVIII) chain; May act as a c |
| CRHR1 | P34998 | 1 | Soft red | 9606.ENSP | 2.6470588 | 0.2 | 0.3777777 | 6 | 914 | 5 | 0.4524064 | 5.0 | 0 | 5 | 0.8169934 | 0.3066666666666666 | Corticotropin releasing hormone receptor 1; |
| CYP2D6 | P10635 | 1 | Soft red | 9606.ENSP | 4.7352941 | 0.0 | 0.2111801 | 9 | 0 | 1 | 0.0 | 2.0 | 0 | 1 | 0.5849673 | 0.0 | Cytochrome P450, family 2, subfamily D, poly |
| CYP2E1 | P05181 | 1 | Soft red | 9606.ENSP | 5.5294117 | 0.0 | 0.1808510 | 10 | 0 | 1 | 0.0 | 2.0 | 0 | 1 | 0.4967320 | 0.0 | Cytochrome P450, family 2, subfamily E, poly |
| FGF8 | P55075 | 1 | Soft red | 9606.ENSP | 4.2647058 | 0.0 | 0.2344827 | 9 | 16 | 2 | 0.0069518 | 2.5 | 0 | 2 | 0.6372549 | 0.5 | Fibroblast growth factor 8 (androgen-induced |
| GABRD | O14764 | 1 | Soft red | 9606.ENSP | 3.0588235 | 0.4 | 0.3269230 | 7 | 322 | 6 | 0.1331253 | 4.5 | 0 | 6 | 0.7712418 | 0.3461538461538461 | Gamma-aminobutyric acid (GABA) A receptor |
| GRIK1 | P39086 | 1 | Soft red | 9606.ENSP | 3.1764705 | 0.5333333 | 0.3148148 | 6 | 250 | 6 | 0.0816696 | 4.3333333 | 0 | 6 | 0.7581699 | 0.4333333333333333 | Glutamate receptor, ionotropic, kainate 1; Io |
| GRIK2 | Q96KS6 | 1 | Soft red | 9606.ENSP | 3.1764705 | 0.5333333 | 0.3148148 | 6 | 250 | 6 | 0.0816696 | 4.3333333 | 0 | 6 | 0.7581699 | 0.4333333333333333 | Glutamate receptor, ionotropic, kainate 2; Io |
| GRM2 | Q14416 | 1 | Soft red | 9606.ENSP | 3.6470588 | 0.3333333 | 0.2741935 | 7 | 164 | 3 | 0.0651812 | 5.0 | 0 | 3 | 0.7058823 | 0.5833333333333334 | Glutamate receptor, metabotropic 2; G-prote |
| LPCAT4 | Q643R3 | 1 | Soft red | 9606.ENSP | 6.3823529 | 0.0 | 0.1566820 | 10 | 0 | 1 | 0.0 | 2.0 | 0 | 1 | 0.4019607 | 0.0 | 1-alkenylglycerophosphoethanolamine O-acy |
| LY6H | O94772 | 1 | Soft red | 9606.ENSP | 3.7058823 | 0.6666666 | 0.2698412 | 8 | 22 | 4 | 0.0082887 | 5.25 | 0 | 4 | 0.6993464 | 0.525 | Lymphocyte antigen 6 complex, locus H; Beli |
| MYO15A | Q9UKN7 | 1 | Soft red | 9606.ENSP | 4.5 | 0.0 | 0.2222222 | 8 | 268 | 3 | 0.1693404 | 2.3333333 | 0 | 3 | 0.6111111 | 0.3333333333333333 | Unconventional myosin-15; Myosins are actin |
| NEGR1 | Q7Z3B1 | 1 | Soft red | 9606.ENSP | 3.9411764 | 0.0 | 0.2537313 | 7 | 18 | 2 | 0.0142602 | 3.0 | 0 | 2 | 0.6732026 | 0.5 | Neuronal growth regulator 1; May be involve |
| NRGN | Q92686 | 1 | Soft red | 9606.ENSP | 3.7352941 | 0.0 | 0.2677165 | 8 | 128 | 3 | 0.0583481 | 2.3333333 | 0 | 3 | 0.6960784 | 0.3333333333333333 | Neurogranin (protein kinase C substrate, RC3 |
| SCN3B | Q9NY72 | 1 | Soft red | 9606.ENSP | 2.5882352 | 0.2 | 0.3863636 | 5 | 1224 | 6 | 0.5940582 | 4.5 | 0 | 6 | 0.8235294 | 0.2857142857142857 | Sodium channel, voltage-gated, type III, beta |
| SHISA8 | B8ZZ34 | 1 | Soft red | 9606.ENSP | 4.0882352 | 1.0 | 0.2446043 | 7 | 0 | 3 | 0.0 | 5.3333333 | 0 | 3 | 0.6568627 | 0.8888888888888888 | Putative protein shisa-8; Shisa family membe |
| SLC44A5 | Q8NCS7 | 1 | Soft red | 9606.ENSP | 4.1470588 | 0.0 | 0.2411347 | 7 | 2 | 2 | 0.0017825 | 2.0 | 0 | 2 | 0.6503267 | 0.5 | Choline transporter-like protein 5; Solute car |
| SNCG | O76070 | 1 | Soft red | 9606.ENSP | 3.4705882 | 0.0 | 0.2881355 | 6 | 74 | 2 | 0.0427807 | 4.0 | 0 | 2 | 0.7254901 | 0.5 | Synuclein, gamma (breast cancer-specific pro |
| SOHLH1 | Q5JUK2 | 1 | Soft red | 9606.ENSP | 3.6470588 | 0.3333333 | 0.2741935 | 8 | 244 | 3 | 0.1140819 | 5.6666666 | 0 | 3 | 0.7058823 | 0.4848484848484848 | Spermatogenesis- and oogenesis-specific bas |
| SYCE1 | Q8N0S2 | 1 | Soft red | 9606.ENSP | 4.5588235 | 0.0 | 0.2193548 | 9 | 124 | 2 | 0.0588235 | 2.0 | 0 | 2 | 0.6045751 | 0.5 | Synaptonemal complex central element prote |
| TAC3 | Q9UHF0 | 1 | Soft red | 9606.ENSP | 3.7058823 | 0.0 | 0.2698412 | 8 | 54 | 2 | 0.0303030 | 5.5 | 0 | 2 | 0.6993464 | 0.5 | Tachykinin 3; Tachykinins are active peptides |
| P2RX2 | Q9UHD5 | 2 | Blue | 9606.ENSP | 1.6666666 | 0.0 | 0.6 | 2 | 0 | 1 | 0.0 | 3.0 | 0 | 1 | 0.7777777 | 0.0 | Purinergic receptor P2X, ligand-gated ion cha |
| PKD1 | P98161 | 2 | Blue | 9606.ENSP | 1.3333333 | 1.0 | 0.75 | 2 | 0 | 2 | 0.0 | 2.5 | 0 | 2 | 0.8888888 | 0.8333333333333334 | Autosomal dominant polycystic kidney diseas |
| PKD2L1 | Q9P0L9 | 2 | Blue | 9606.ENSP | 1.0 | 0.3333333 | 1.0 | 1 | 4 | 3 | 0.6666666 | 1.6666666 | 0 | 3 | 1.0 | 0.6666666666666666 | Polycystic kidney disease 2-like 1 protein; Po |
| TRPM2 | O94759 | 2 | Blue | 9606.ENSP | 1.3333333 | 1.0 | 0.75 | 2 | 0 | 2 | 0.0 | 2.5 | 0 | 2 | 0.8888888 | 0.8333333333333334 | Transient receptor potential cation channel, s |
| AMY2A | P04746 | 3 | Cyan | 9606.ENSP | 2.0 | 0.0 | 0.5 | 3 | 0 | 1 | 0.0 | 2.0 | 0 | 1 | 0.5 | 0.0 | 1,4-alpha-D-glucan glucanohydrolase; Amylas |
| NOXA1 | Q86UR1 | 3 | Cyan | 9606.ENSP | 1.3333333 | 0.0 | 0.75 | 2 | 4 | 2 | 0.6666666 | 1.5 | 0 | 2 | 0.8333333 | 0.5 | NADPH oxidase activator 1; Functions as an a |
| NOXO1 | Q8NFA2 | 3 | Cyan | 9606.ENSP | 2.0 | 0.0 | 0.5 | 3 | 0 | 1 | 0.0 | 2.0 | 0 | 1 | 0.5 | 0.0 | SH3 and PX domain-containing protein 5; Con |
| PDIA2 | Q13087 | 3 | Cyan | 9606.ENSP | 1.3333333 | 0.0 | 0.75 | 2 | 4 | 2 | 0.6666666 | 1.5 | 0 | 2 | 0.8333333 | 0.5 | Protein disulfide isomerase family A, member |
| AP3B2 | Q13367 | 4 | | 9606.ENSP | 1.5 | 0.0 | 0.6666666 | 2 | 0 | 1 | 0.0 | 2.0 | 0 | 1 | 0.75 | 0.0 | Clathrin assembly protein complex 3 beta-2 la |
| MAST1 | Q9Y2H9 | 4 | | 9606.ENSP | 1.5 | 0.0 | 0.6666666 | 2 | 0 | 1 | 0.0 | 2.0 | 0 | 1 | 0.75 | 0.0 | Microtubule-associated serine/threonine-pro |
| TCTE1 | Q5JU00 | 4 | | 9606.ENSP | 1.0 | 0.0 | 1.0 | 1 | 2 | 2 | 1.0 | 1.0 | 0 | 2 | 1.0 | 0.0 | T-complex-associated-testis-expressed 1; Dyn |
| KRT17 | Q04695 | 5 | | 9606.ENSP | 1.0 | 1.0 | 1.0 | 1 | 0 | 2 | 0.0 | 2.0 | 0 | 2 | 1.0 | 1.0 | Keratin, type I cytoskeletal 17; Type I keratin i |
| TGM1 | P22735 | 5 | | 9606.ENSP | 1.0 | 1.0 | 1.0 | 1 | 0 | 2 | 0.0 | 2.0 | 0 | 2 | 1.0 | 1.0 | Protein-glutamine gamma-glutamyltransferas |
| CORO1A | P31146 | 6 | | 9606.ENSP | 1.0 | 0.0 | 1.0 | 1 | 0 | 1 | 0.0 | 1.0 | 0 | 1 | 1.0 | 0.0 | Tryptophan aspartate-containing coat protein |
| CORO6 | Q6QEF8 | 6 | | 9606.ENSP | 1.0 | 0.0 | 1.0 | 1 | 0 | 1 | 0.0 | 1.0 | 0 | 1 | 1.0 | 0.0 | Coronin-like protein E; WD repeat domain co |
| PNMA5 | Q96PV4 | 7 | | 9606.ENSP | 1.0 | 0.0 | 1.0 | 1 | 0 | 1 | 0.0 | 1.0 | 0 | 1 | 1.0 | 0.0 | Paraneoplastic Ma antigens; Belongs to the P |
| RTL1 | A6NKG5 | 7 | | 9606.ENSP | 1.0 | 0.0 | 1.0 | 1 | 0 | 1 | 0.0 | 1.0 | 0 | 1 | 1.0 | 0.0 | Mammalian retrotransposon derived protein |
| PROC | P04070 | 8 | | 9606.ENSP | 1.0 | 0.0 | 1.0 | 1 | 0 | 1 | 0.0 | 1.0 | 0 | 1 | 1.0 | 0.0 | Protein C (inactivator of coagulation factors V |
| TNNT2 | P45379 | 8 | | 9606.ENSP | 1.0 | 0.0 | 1.0 | 1 | 0 | 1 | 0.0 | 1.0 | 0 | 1 | 1.0 | 0.0 | Troponin T type 2 (cardiac); Troponin T is the |
| MICAL1 | Q9UFF7 | 9 | | 9606.ENSP | 1.0 | 0.0 | 1.0 | 1 | 0 | 1 | 0.0 | 1.0 | 0 | 1 | 1.0 | 0.0 | Microtubule associated monooxygenase, calp |
| SEMA5B | Q9P283 | 9 | | 9606.ENSP | 1.0 | 0.0 | 1.0 | 1 | 0 | 1 | 0.0 | 1.0 | 0 | 1 | 1.0 | 0.0 | Semaphorin-5B; May act as positive axonal gu |
| CHTF18 | Q8WVB6 | 10 | | 9606.ENSP | 1.0 | 0.0 | 1.0 | 1 | 0 | 1 | 0.0 | 1.0 | 0 | 1 | 1.0 | 0.0 | CTF18, chromosome transmission fidelity fact |
| RTEL1 | Q9NZ71 | 10 | | 9606.ENSP | 1.0 | 0.0 | 1.0 | 1 | 0 | 1 | 0.0 | 1.0 | 0 | 1 | 1.0 | 0.0 | Regulator of telomere elongation helicase 1; |
| TMEM249 | Q2WGJ8 | 11 | | 9606.ENSP | 1.0 | 0.0 | 1.0 | 1 | 0 | 1 | 0.0 | 1.0 | 0 | 1 | 1.0 | 0.0 | Transmembrane protein 249 |
| B4GALNT3 | Q6L9W6 | 12 | | 9606.ENSP | 1.0 | 0.0 | 1.0 | 1 | 0 | 1 | 0.0 | 1.0 | 0 | 1 | 1.0 | 0.0 | N-acetyl-beta-glucosaminyl-glycoprotein 4-be |
| MSLNL | Q96KJ4 | 12 | | 9606.ENSP | 1.0 | 0.0 | 1.0 | 1 | 0 | 1 | 0.0 | 1.0 | 0 | 1 | 1.0 | 0.0 | Pre-pro-megakaryocyte-potentiating-factor-li |
