## Supplementary tables for "Transcriptome profiling of the dorsomedial prefrontal cortex in suicide victims": S6_Top_10_hub_genes_of_PPI_networks.pdf

Supplementary file 6: Protein-protein interaction network of downregulated genes; the top 10 most ranked hub genes are highlighted.

| node_name | MCC | DMNC | MNC | Degree | EPC | BottleNeck | EcCentricit | Closeness | Radiality | Betweenne | Stress | ClusteringCoefficient |
|---|---|---|---|---|---|---|---|---|---|---|---|---|
| EGFR | 9,22E+13 | 0,29786 | 178 | 183 | 73,399 | 54 | 0,15505 | 605,25 | 8,21623 | 129396 | 1513486 | 0,11974 |
| FN1 | 9,22E+13 | 0,34235 | 174 | 174 | 74,977 | 28 | 0,15505 | 596,7333 | 8,18785 | 91496,87 | 1220576 | 0,1465 |
| IL6 | 9,22E+13 | 0,34034 | 169 | 170 | 74,154 | 26 | 0,15505 | 588,15 | 8,14729 | 74632,71 | 1084460 | 0,14521 |
| STAT3 | 9,22E+13 | 0,36328 | 148 | 148 | 70,81 | 32 | 0,1329 | 571,4762 | 8,097 | 54571,83 | 813768 | 0,16336 |
| NOTCH1 | 6,85E+13 | 0,35794 | 133 | 135 | 66,77 | 33 | 0,15505 | 558,6333 | 8,05645 | 55802,28 | 812800 | 0,16142 |
| CD44 | 9,22E+13 | 0,4311 | 132 | 132 | 70,192 | 25 | 0,15505 | 560,5167 | 8,06943 | 32892,27 | 588668 | 0,20079 |
| ERBB2 | 2,34E+13 | 0,3994 | 105 | 108 | 62,178 | 20 | 0,1329 | 541,1929 | 8,01184 | 30613,78 | 460210 | 0,18865 |
| PECAM1 | 9,22E+13 | 0,51927 | 102 | 102 | 66,26 | 8 | 0,15505 | 536,1333 | 7,99481 | 16240,63 | 315752 | 0,26189 |
| TLR4 | 9,22E+13 | 0,42805 | 99 | 99 | 60,059 | 7 | 0,15505 | 533,9 | 7,98832 | 31329,91 | 464360 | 0,21789 |
| ITGB1 | 5,19E+13 | 0,49211 | 91 | 94 | 59,802 | 12 | 0,15505 | 527,6333 | 7,96318 | 24313,2 | 379230 | 0,24091 |
| CCL2 | 9,22E+13 | 0,55874 | 92 | 92 | 63,035 | 24 | 0,1329 | 528,1595 | 7,96561 | 11934,82 | 263238 | 0,29097 |
| SOX2 | 2,20E+11 | 0,34507 | 89 | 92 | 53,333 | 16 | 0,15505 | 526,3333 | 7,96237 | 41441,48 | 566574 | 0,16985 |
| CAV1 | 9,22E+13 | 0,43583 | 89 | 92 | 60,037 | 18 | 0,1329 | 532,2095 | 7,99075 | 36324,77 | 470750 | 0,21476 |
| ICAM1 | 9,22E+13 | 0,61694 | 87 | 88 | 63,605 | 29 | 0,15505 | 520,5667 | 7,93073 | 12354,92 | 238208 | 0,31949 |
| FOS | 2,66E+11 | 0,37041 | 86 | 88 | 52,38 | 2 | 0,1329 | 529,5095 | 7,97696 | 33133,53 | 460482 | 0,18809 |
| MMP2 | 9,22E+13 | 0,60501 | 86 | 87 | 61,149 | 4 | 0,1329 | 527,7762 | 7,9721 | 11980,89 | 232636 | 0,31435 |
| VWF | 9,22E+13 | 0,46299 | 79 | 81 | 52,472 | 8 | 0,15505 | 512,8 | 7,90316 | 20275,36 | 299732 | 0,24043 |
| CD34 | 9,22E+13 | 0,56277 | 81 | 81 | 58,424 | 4 | 0,15505 | 518,6167 | 7,93803 | 10722,89 | 211616 | 0,30494 |
| CDH5 | 4,62E+13 | 0,47252 | 78 | 80 | 52,884 | 12 | 0,15505 | 507,3333 | 7,8772 | 20491,05 | 314116 | 0,2462 |
| NFKBIA | 4,09E+12 | 0,42216 | 77 | 78 | 50,879 | 33 | 0,1329 | 509,2262 | 7,87801 | 18964,74 | 299056 | 0,22644 |
| TGFB1 | 9,22E+13 | 0,60702 | 73 | 74 | 57,7 | 7 | 0,1329 | 515,8095 | 7,92911 | 12063,82 | 205998 | 0,33062 |
| CXCL12 | 9,22E+13 | 0,65425 | 74 | 74 | 59,696 | 3 | 0,1329 | 513,8262 | 7,92262 | 7342,909 | 160148 | 0,36468 |
| VCAM1 | 9,22E+13 | 0,66276 | 73 | 73 | 57,596 | 2 | 0,15505 | 509,75 | 7,90235 | 5396,366 | 143918 | 0,371 |
| PDGFRB | 5,76E+13 | 0,57977 | 69 | 72 | 58,577 | 13 | 0,15505 | 505,1667 | 7,88369 | 11071,58 | 181988 | 0,30321 |
| SERPINE1 | 9,22E+13 | 0,51544 | 69 | 71 | 54,257 | 10 | 0,15505 | 505,35 | 7,87964 | 12488,75 | 191306 | 0,27767 |
| ITGB3 | 2,09E+12 | 0,51843 | 69 | 71 | 51,517 | 10 | 0,15505 | 504,2667 | 7,8772 | 18971,54 | 274974 | 0,27887 |
| CXCL10 | 1,91E+13 | 0,52815 | 69 | 70 | 48,113 | 26 | 0,1329 | 482,1429 | 7,74013 | 15799,21 | 286334 | 0,29234 |
| TIMP1 | 9,22E+13 | 0,7088 | 65 | 67 | 56,69 | 5 | 0,1329 | 500,9095 | 7,85774 | 8786,397 | 150072 | 0,38716 |
| SOX9 | 9,41E+08 | 0,39211 | 66 | 66 | 44,987 | 27 | 0,15505 | 499,1167 | 7,86098 | 14434,56 | 237292 | 0,22657 |
| GFAP | 7,66E+09 | 0,37148 | 62 | 64 | 43,152 | 24 | 0,15505 | 511,3 | 7,92992 | 41033,79 | 500702 | 0,20536 |
| NOS3 | 9,22E+13 | 0,53479 | 62 | 63 | 50,13 | 5 | 0,1329 | 499,7595 | 7,86585 | 12532,86 | 195684 | 0,30517 |
| EDN1 | 9,22E+13 | 0,5218 | 60 | 62 | 50,298 | 2 | 0,1329 | 499,3929 | 7,86504 | 14845,43 | 246346 | 0,29085 |
| IRF7 | 1,94E+12 | 0,46399 | 61 | 62 | 29,691 | 9 | 0,1329 | 461,1762 | 7,63874 | 15328,65 | 256208 | 0,266 |
| GJA1 | 3,35E+12 | 0,40611 | 59 | 61 | 43,262 | 5 | 0,1329 | 502,8262 | 7,89099 | 18515,44 | 259284 | 0,22732 |
| EZR | 6264952 | 0,31337 | 59 | 61 | 40,091 | 9 | 0,15505 | 500,4167 | 7,88126 | 21583,97 | 278782 | 0,17541 |
| CD40 | 1,12E+13 | 0,58391 | 58 | 60 | 50,179 | 6 | 0,1329 | 487,5095 | 7,79528 | 8360,843 | 150334 | 0,32825 |
| JAG1 | 5,96E+10 | 0,5269 | 57 | 58 | 44,887 | 1 | 0,15505 | 492,4 | 7,83503 | 13805,14 | 230642 | 0,30792 |
| ENG | 9,22E+13 | 0,72969 | 56 | 57 | 53,303 | 3 | 0,15505 | 485,7833 | 7,79528 | 2471,778 | 63034 | 0,42857 |
| MYD88 | 6,55E+12 | 0,527 | 56 | 57 | 42,989 | 3 | 0,15505 | 483,6833 | 7,78231 | 7554,529 | 134142 | 0,30952 |
| PXN | 1,73E+08 | 0,44984 | 53 | 56 | 42,191 | 14 | 0,1329 | 482,2762 | 7,78474 | 14737,88 | 195196 | 0,24935 |
| TNFRSF1A | 1,40E+13 | 0,61034 | 53 | 54 | 46,873 | 5 | 0,1329 | 483,3595 | 7,78636 | 8748,21 | 153484 | 0,36408 |
| LGALS3 | 8,91E+11 | 0,50326 | 48 | 54 | 42,912 | 4 | 0,15505 | 491,2833 | 7,8407 | 14651,1 | 200862 | 0,25437 |
| LYN | 1,25E+08 | 0,4102 | 52 | 54 | 39,538 | 10 | 0,1329 | 490,3095 | 7,83097 | 14648,13 | 215144 | 0,2369 |
| LEF1 | 210951 | 0,34607 | 52 | 53 | 31,812 | 8 | 0,15505 | 485,5333 | 7,80988 | 15442,34 | 199476 | 0,20755 |
| FLT1 | 4,00E+13 | 0,68297 | 53 | 53 | 47,845 | 1 | 0,1329 | 482,9262 | 7,79042 | 4128,29 | 91722 | 0,42308 |
| TFRC | 1,40E+09 | 0,42301 | 49 | 53 | 40,346 | 8 | 0,15505 | 493,8333 | 7,84638 | 22041,67 | 269922 | 0,22932 |
| PPARA | 1,53E+10 | 0,35311 | 50 | 53 | 35,458 | 14 | 0,1329 | 485,7929 | 7,80339 | 25883,74 | 331896 | 0,19811 |
| VIM | 9,73E+09 | 0,56395 | 50 | 53 | 44,39 | 7 | 0,1329 | 492,2929 | 7,84476 | 10908,35 | 166386 | 0,31713 |
| CSF3 | 4,49E+13 | 0,75163 | 51 | 52 | 51,232 | 9 | 0,15505 | 476,9833 | 7,7523 | 4390,482 | 77216 | 0,45324 |
| YAP1 | 1160263 | 0,36277 | 49 | 52 | 32,933 | 6 | 0,1329 | 479,4262 | 7,78068 | 11056,88 | 154360 | 0,20437 |
| IRF1 | 2,13E+12 | 0,57428 | 49 | 50 | 29,845 | 3 | 0,1329 | 452,0262 | 7,60549 | 9394,606 | 204944 | 0,3502 |
| CSF1 | 5,02E+13 | 0,77737 | 50 | 50 | 50,666 | 1 | 0,1329 | 480,9262 | 7,77825 | 1942,994 | 52152 | 0,49061 |
| ANXA2 | 3,14E+10 | 0,44175 | 49 | 50 | 39,338 | 5 | 0,1329 | 487,2762 | 7,82854 | 12702,46 | 174894 | 0,26939 |
| CXCL1 | 2,45E+13 | 0,72045 | 50 | 50 | 48,987 | 1 | 0,15505 | 477,0833 | 7,75797 | 2396,394 | 62120 | 0,45469 |
| HMOX1 | 4,22E+13 | 0,51298 | 47 | 48 | 42,392 | 14 | 0,1329 | 486,2429 | 7,81394 | 11214,87 | 171984 | 0,31649 |
| SOCS3 | 3,95E+11 | 0,56327 | 47 | 47 | 39,867 | 1 | 0,1329 | 468,6429 | 7,7158 | 2893,323 | 57168 | 0,36263 |
| NES | 1,72E+12 | 0,61644 | 47 | 47 | 41,659 | 1 | 0,1329 | 484,2929 | 7,81313 | 3575,532 | 88680 | 0,39685 |
| FOXO1 | 3556355 | 0,43968 | 46 | 47 | 38,555 | 1 | 0,1329 | 484,6762 | 7,81475 | 6476,885 | 112154 | 0,2729 |
| ITGA5 | 4,40E+10 | 0,67218 | 46 | 46 | 41,172 | 10 | 0,15505 | 467,8667 | 7,71823 | 2747,969 | 63640 | 0,43575 |
| PAX6 | 45353 | 0,34193 | 45 | 46 | 25,973 | 12 | 0,1329 | 464,2595 | 7,69714 | 9633,548 | 122138 | 0,21353 |
| TGFB2 | 3,65E+11 | 0,66869 | 44 | 45 | 45,071 | 4 | 0,1329 | 479,6095 | 7,78555 | 4988,69 | 92890 | 0,4202 |
| CDKN1A | 3,64E+07 | 0,42623 | 43 | 45 | 35,919 | 7 | 0,1329 | 480,0262 | 7,79123 | 12273,46 | 174268 | 0,25758 |
| ITGA6 | 1,33E+10 | 0,58638 | 45 | 45 | 40,762 | 2 | 0,15505 | 478,8833 | 7,79042 | 2782,175 | 63996 | 0,38283 |
| ANGPT2 | 1,67E+13 | 0,72173 | 44 | 44 | 44,439 | 2 | 0,1329 | 474,2595 | 7,75716 | 2531,435 | 53710 | 0,47463 |
| TGFBR2 | 6,47E+09 | 0,56193 | 42 | 43 | 40,098 | 3 | 0,1329 | 477,1429 | 7,77338 | 6024,688 | 103014 | 0,3577 |
| DLL4 | 1,85E+08 | 0,50205 | 41 | 42 | 34,659 | 2 | 0,1329 | 457,6095 | 7,66145 | 6356,396 | 119470 | 0,32172 |
| FLT4 | 1,46E+10 | 0,55497 | 42 | 42 | 40,682 | 6 | 0,1329 | 464,8095 | 7,70769 | 2851,715 | 61876 | 0,3705 |
| BRCA1 | 135751 | 0,35103 | 35 | 42 | 28,201 | 11 | 0,1329 | 480,9262 | 7,80096 | 20211,99 | 230202 | 0,17305 |
| PGF | 2,20E+13 | 0,80111 | 41 | 42 | 44,814 | 2 | 0,1329 | 466,4095 | 7,71255 | 2257,856 | 43000 | 0,51336 |
| FGF1 | 8,16E+09 | 0,57839 | 40 | 42 | 39,218 | 9 | 0,1329 | 474,7429 | 7,76446 | 13157,57 | 170090 | 0,3554 |
| TCF7L2 | 3768 | 0,22495 | 39 | 41 | 22,832 | 7 | 0,15505 | 463,8 | 7,71174 | 14125,54 | 153520 | 0,14024 |
| SLC2A1 | 1139611 | 0,30729 | 38 | 41 | 29,684 | 9 | 0,1329 | 474,5595 | 7,76203 | 19353,61 | 255442 | 0,18171 |
| HSPB1 | 1074794 | 0,37984 | 36 | 40 | 33,418 | 21 | 0,1329 | 481,8762 | 7,81394 | 10978,16 | 152246 | 0,21667 |
| MSN | 1,70E+07 | 0,45364 | 40 | 40 | 33,368 | 2 | 0,15505 | 471,95 | 7,76041 | 4053,906 | 75280 | 0,30769 |
| ANXA1 | 2,15E+07 | 0,41242 | 39 | 40 | 35,105 | 6 | 0,1329 | 469,5929 | 7,74013 | 7476,442 | 104678 | 0,26795 |
| TGFB3 | 7,64E+11 | 0,66302 | 39 | 39 | 42,343 | 1 | 0,1329 | 467,3762 | 7,72472 | 1833,413 | 43616 | 0,45344 |

| | | | | | | | | | | | | |
|---|---|---|---|---|---|---|---|---|---|---|---|---|
| LIF | 3,99E+11 | 0,59988 | 39 | 39 | 37,985 | 2 | 0,1329 | 468,4929 | 7,73689 | 1491,854 | 35402 | 0,41026 |
| CDK2 | 88609 | 0,2743 | 38 | 39 | 24,372 | 7 | 0,1329 | 474,8762 | 7,77744 | 12122,88 | 158982 | 0,17949 |
| BGN | 1,15E+07 | 0,42279 | 38 | 39 | 29,501 | 3 | 0,15505 | 454,35 | 7,64686 | 6682,861 | 101206 | 0,27665 |
| ITGA1 | 6,30E+09 | 0,60209 | 37 | 38 | 32,988 | 3 | 0,15505 | 462,1167 | 7,70525 | 6038,811 | 90692 | 0,39687 |
| EPHA2 | 1,81E+07 | 0,52418 | 35 | 38 | 34,195 | 5 | 0,1329 | 471,2429 | 7,7523 | 6661,03 | 100694 | 0,31437 |
| COL4A1 | 2,54E+07 | 0,58554 | 34 | 38 | 32,088 | 1 | 0,1329 | 456,4929 | 7,66875 | 5573,068 | 84690 | 0,3357 |
| NOTCH3 | 9,59E+07 | 0,51361 | 37 | 37 | 33,3 | 3 | 0,1329 | 469,1262 | 7,74743 | 2969,924 | 66738 | 0,35736 |
| BCL6 | 331211 | 0,42279 | 36 | 37 | 31,28 | 2 | 0,1329 | 469,5762 | 7,745 | 3438,053 | 60116 | 0,28078 |
| IL1R1 | 3,29E+12 | 0,77689 | 37 | 37 | 39,062 | 2 | 0,1329 | 453,0762 | 7,63874 | 962,3744 | 27256 | 0,54054 |
| BST2 | 1,84E+12 | 0,57398 | 35 | 36 | 16,707 | 1 | 0,1329 | 424,7595 | 7,4741 | 5197,673 | 69244 | 0,38413 |
| CLU | 57766 | 0,30749 | 36 | 36 | 22,35 | 4 | 0,1329 | 455,3595 | 7,65821 | 7762,66 | 100386 | 0,21587 |
| PLAUR | 3,82E+11 | 0,68417 | 33 | 36 | 36,955 | 5 | 0,1329 | 469,4429 | 7,75067 | 9544,622 | 134044 | 0,41429 |
| CD163 | 8,78E+10 | 0,4974 | 36 | 36 | 31,656 | 5 | 0,15505 | 461,7667 | 7,69795 | 8718,953 | 134924 | 0,34921 |
| GLI2 | 448413 | 0,45612 | 33 | 36 | 23,098 | 1 | 0,15505 | 456,0167 | 7,67119 | 5114,419 | 82800 | 0,27619 |
| SLC1A3 | 17241 | 0,28726 | 32 | 35 | 14,582 | 4 | 0,1329 | 434,5762 | 7,53979 | 9669,733 | 99008 | 0,17479 |
| MX2 | 1,85E+12 | 0,74749 | 34 | 35 | 14,743 | 1 | 0,1329 | 399,8262 | 7,29647 | 2301,203 | 26990 | 0,5042 |
| SDC4 | 1872136 | 0,45874 | 33 | 35 | 30,392 | 3 | 0,1329 | 461,2262 | 7,69958 | 6376,868 | 102264 | 0,29412 |
| HSPG2 | 2373907 | 0,42856 | 34 | 35 | 27,017 | 3 | 0,1329 | 445,1595 | 7,60955 | 5515,769 | 84912 | 0,28908 |
| CLDN5 | 3,60E+08 | 0,53214 | 33 | 35 | 34,398 | 6 | 0,1329 | 463,0595 | 7,71174 | 8097,825 | 110796 | 0,34118 |
| LDLR | 9,99E+08 | 0,41417 | 33 | 35 | 31,193 | 9 | 0,1329 | 469,3429 | 7,74581 | 10610,43 | 139022 | 0,26555 |
| TNFAIP3 | 1,88E+11 | 0,62424 | 32 | 35 | 26,793 | 8 | 0,1329 | 438,1429 | 7,55358 | 5112,916 | 82512 | 0,37983 |
| NOTCH2 | 634712 | 0,46843 | 34 | 34 | 24,731 | 3 | 0,1329 | 458,3929 | 7,69309 | 3122,052 | 57754 | 0,33512 |
| KAT2B | 6244 | 0,24307 | 32 | 34 | 17,834 | 20 | 0,1329 | 451,6262 | 7,64848 | 8350,395 | 105898 | 0,15686 |
| ITGB4 | 2,12E+07 | 0,56348 | 32 | 33 | 31,776 | 2 | 0,1329 | 459,1429 | 7,69795 | 3308,237 | 50578 | 0,38636 |
| CD81 | 1,55E+07 | 0,50552 | 30 | 33 | 29,861 | 1 | 0,15505 | 466,9833 | 7,74013 | 6744,914 | 101020 | 0,31061 |
| SOCS1 | 1,88E+10 | 0,60639 | 31 | 33 | 30,276 | 2 | 0,1329 | 453,4929 | 7,65172 | 2830,011 | 46796 | 0,39394 |
| IFI35 | 1,85E+12 | 0,7955 | 32 | 33 | 14,122 | 1 | 0,11629 | 397,5512 | 7,28025 | 1747,531 | 23252 | 0,54545 |
| IL10RA | 1,50E+08 | 0,53642 | 31 | 32 | 25,067 | 2 | 0,1329 | 426,9762 | 7,48707 | 3757,092 | 64402 | 0,37097 |
| ITGB5 | 1,57E+07 | 0,51652 | 32 | 32 | 25,909 | 1 | 0,15505 | 443,3167 | 7,59981 | 1417,936 | 28742 | 0,37702 |
| SLC2A4 | 22348 | 0,30154 | 28 | 32 | 20,188 | 4 | 0,1329 | 452,9429 | 7,66308 | 11274,25 | 133618 | 0,1754 |
| IL6R | 1,19E+09 | 0,64082 | 32 | 32 | 32,477 | 2 | 0,1329 | 444,7595 | 7,59981 | 547,09 | 11960 | 0,46774 |
| GBP1 | 5,38E+11 | 0,73757 | 31 | 32 | 12,066 | 2 | 0,1329 | 414,4595 | 7,40759 | 3590,931 | 48186 | 0,51008 |
| OAS3 | 1,85E+12 | 0,85127 | 31 | 31 | 11,435 | 1 | 0,11629 | 388,2845 | 7,22266 | 508,0305 | 11964 | 0,62796 |
| RHOC | 535192 | 0,45519 | 23 | 31 | 22,524 | 20 | 0,1329 | 461,9762 | 7,72147 | 15699,39 | 191924 | 0,2043 |
| ANPEP | 1,89E+10 | 0,74479 | 27 | 31 | 32,264 | 4 | 0,15505 | 457,5833 | 7,68498 | 10534,45 | 145324 | 0,43441 |
| XAF1 | 1,85E+12 | 0,86616 | 30 | 31 | 12,071 | 2 | 0,11629 | 387,0845 | 7,20482 | 677,6753 | 12266 | 0,6043 |
| TBL1X | 229 | 0,19854 | 23 | 30 | 17,355 | 13 | 0,1329 | 454,3262 | 7,66794 | 21313,94 | 229624 | 0,09425 |
| DUSP1 | 395296 | 0,42979 | 28 | 30 | 23,191 | 2 | 0,1329 | 454,0095 | 7,66875 | 5022,497 | 73990 | 0,28506 |
| VASP | 370274 | 0,38134 | 26 | 30 | 19,589 | 2 | 0,15505 | 439,9 | 7,59332 | 5540,291 | 69426 | 0,22299 |
| S100B | 38701 | 0,40816 | 29 | 30 | 23,244 | 5 | 0,1329 | 459,3929 | 7,70444 | 6848,027 | 97380 | 0,28736 |
| F3 | 3,26E+12 | 0,77369 | 30 | 30 | 36,983 | 2 | 0,1329 | 453,5095 | 7,6647 | 581,9826 | 15806 | 0,57701 |
| MCL1 | 2627665 | 0,60082 | 29 | 30 | 29,85 | 2 | 0,1329 | 463,0595 | 7,71661 | 1957,682 | 38376 | 0,42299 |
| FGR | 11611 | 0,3362 | 25 | 30 | 23,721 | 8 | 0,15505 | 449,3167 | 7,64523 | 6226,141 | 75934 | 0,18621 |
| NOTCH4 | 2398611 | 0,58256 | 27 | 30 | 23,729 | 2 | 0,1329 | 450,5595 | 7,64442 | 3676,203 | 60596 | 0,36552 |
| MSX1 | 5439 | 0,41469 | 29 | 30 | 20,988 | 5 | 0,1329 | 433,3929 | 7,54385 | 3751,107 | 50368 | 0,29195 |
| CD59 | 464566 | 0,42632 | 28 | 30 | 23,584 | 2 | 0,15505 | 443,8 | 7,60468 | 4781,066 | 72782 | 0,28276 |
| MUC1 | 1249240 | 0,54212 | 25 | 30 | 30,973 | 4 | 0,1329 | 459,4762 | 7,70282 | 7725,131 | 94930 | 0,30115 |
| NTRK2 | 5205 | 0,38288 | 24 | 29 | 22,511 | 5 | 0,1329 | 449,0929 | 7,64361 | 10501,25 | 127392 | 0,20936 |
| ALDH1A1 | 6,81E+07 | 0,77963 | 23 | 29 | 28,126 | 4 | 0,1329 | 463,3095 | 7,72959 | 7187,543 | 86330 | 0,40148 |
| FOSL1 | 4359 | 0,31194 | 28 | 29 | 23,424 | 4 | 0,1329 | 451,4262 | 7,65334 | 4982,774 | 63504 | 0,22167 |
| CSPG4 | 110355 | 0,38819 | 28 | 29 | 26,938 | 6 | 0,1329 | 448,6762 | 7,63956 | 5889,856 | 81082 | 0,27586 |
| FGFR3 | 22648 | 0,38531 | 29 | 29 | 29,336 | 2 | 0,1329 | 455,9262 | 7,68417 | 2134,134 | 34482 | 0,29064 |
| PARD3 | 598 | 0,23113 | 25 | 29 | 17,578 | 12 | 0,1329 | 434,4429 | 7,55358 | 10326,16 | 121524 | 0,13547 |
| TNFRSF1B | 6,69E+08 | 0,62694 | 29 | 29 | 28,609 | 1 | 0,1329 | 433,4429 | 7,53574 | 682,3304 | 13646 | 0,47291 |
| DDX60 | 1,89E+11 | 0,78041 | 29 | 29 | 11,77 | 1 | 0,11629 | 384,5607 | 7,19914 | 559,2565 | 10724 | 0,58867 |
| BMP7 | 33968 | 0,45061 | 29 | 29 | 24,789 | 1 | 0,1329 | 450,1762 | 7,6501 | 1831,369 | 35560 | 0,3399 |
| GLI3 | 40569 | 0,39513 | 28 | 29 | 19,197 | 2 | 0,15505 | 446,2167 | 7,62901 | 4511,21 | 68654 | 0,28079 |
| RUNX3 | 42691 | 0,36982 | 25 | 29 | 18,942 | 5 | 0,1329 | 443,5929 | 7,60711 | 4971,331 | 65192 | 0,22167 |
| IFITM1 | 1,85E+12 | 0,81105 | 28 | 29 | 13,59 | 2 | 0,1329 | 405,8095 | 7,35892 | 3167,183 | 45986 | 0,57635 |
| DAG1 | 23480 | 0,3362 | 25 | 29 | 20,069 | 3 | 0,1329 | 445,3262 | 7,6209 | 7854,441 | 89664 | 0,19704 |
| ZEB2 | 1,07E+07 | 0,5511 | 28 | 29 | 27,678 | 4 | 0,1329 | 457,3095 | 7,69228 | 2598,734 | 43234 | 0,39163 |
| LRP5 | 1973 | 0,28997 | 25 | 28 | 19,151 | 7 | 0,15505 | 452,6 | 7,66713 | 9119,917 | 110230 | 0,18254 |
| IFI44L | 3,63E+11 | 0,85909 | 27 | 28 | 10,451 | 2 | 0,11629 | 381,4345 | 7,17319 | 2817,719 | 38372 | 0,6164 |
| IL4R | 6,34E+07 | 0,59962 | 28 | 28 | 28,16 | 1 | 0,1329 | 420,0762 | 7,45382 | 447,4067 | 6724 | 0,45767 |
| HIST1H4F | 325 | 0,25675 | 24 | 27 | 13,412 | 4 | 0,1329 | 430,4595 | 7,53493 | 10238,07 | 129248 | 0,16239 |
| KLF2 | 1480010 | 0,48669 | 27 | 27 | 30,47 | 1 | 0,1329 | 451,4095 | 7,65821 | 785,4435 | 19150 | 0,37607 |
| TCF7 | 41069 | 0,31844 | 26 | 27 | 15,058 | 1 | 0,15505 | 433,3 | 7,55115 | 3629,559 | 45862 | 0,23077 |
| GBP2 | 1,66E+12 | 0,86485 | 24 | 27 | 12,854 | 2 | 0,1329 | 392,6095 | 7,26078 | 1486,144 | 21924 | 0,54701 |
| PSMB8 | 1,48E+12 | 0,63786 | 27 | 27 | 12,933 | 2 | 0,1329 | 407,3429 | 7,3646 | 2102,267 | 42192 | 0,49288 |
| JUP | 1451 | 0,27913 | 26 | 27 | 19,148 | 9 | 0,1329 | 454,2762 | 7,68335 | 7014,895 | 88730 | 0,20228 |
| IFITM3 | 1,83E+12 | 0,82368 | 25 | 27 | 13,021 | 1 | 0,11629 | 409,2345 | 7,37271 | 2749,062 | 42520 | 0,5584 |
| HLA-B | 1,31E+12 | 0,51501 | 26 | 27 | 11,861 | 1 | 0,11629 | 390,8512 | 7,25024 | 4256,627 | 64034 | 0,37322 |
| SERPINA1 | 20218 | 0,41604 | 25 | 27 | 21,036 | 5 | 0,1329 | 436,1262 | 7,56413 | 2418,863 | 35616 | 0,28205 |
| SOX17 | 3836 | 0,38134 | 26 | 26 | 20,198 | 1 | 0,1329 | 437,4429 | 7,57548 | 1363,413 | 26902 | 0,29846 |
| COL6A2 | 4781200 | 0,54504 | 24 | 26 | 21,182 | 4 | 0,1329 | 420,0929 | 7,45625 | 2286,747 | 44668 | 0,37231 |
| AXL | 28858 | 0,37741 | 26 | 26 | 26,262 | 2 | 0,1329 | 458,9595 | 7,70525 | 1698,09 | 30808 | 0,29538 |
| ISG20 | 1,84E+12 | 0,85704 | 26 | 26 | 10,566 | 1 | 0,11629 | 381,9012 | 7,18048 | 448,774 | 9286 | 0,67077 |
| BPTF | 93 | 0,40573 | 9 | 26 | 4,993 | 12 | 0,11629 | 389,1512 | 7,25348 | 13600,55 | 112340 | 0,06769 |
| PARP9 | 7,34E+08 | 0,70181 | 25 | 26 | 11,135 | 2 | 0,1329 | 394,9429 | 7,28917 | 2126,948 | 28410 | 0,51385 |

| | | | | | | | | | | | | |
|---|---|---|---|---|---|---|---|---|---|---|---|---|
| TGM2 | 183170 | 0,49549 | 24 | 26 | 26,393 | 1 | 0,15505 | 447,4833 | 7,63712 | 4324,323 | 62096 | 0,33846 |
| KRT18 | 1,54E+07 | 0,53603 | 24 | 26 | 25,828 | 1 | 0,1329 | 457,9262 | 7,7012 | 4877,993 | 74292 | 0,36615 |
| IFI27 | 1,84E+12 | 0,90815 | 26 | 26 | 10,303 | 2 | 0,11629 | 379,0012 | 7,15777 | 288,5112 | 7620 | 0,71077 |
| ANGPTL4 | 51916 | 0,40192 | 23 | 25 | 20,956 | 5 | 0,1329 | 447,4595 | 7,63388 | 7623,544 | 103994 | 0,27667 |
| S1PR1 | 5479203 | 0,6036 | 24 | 25 | 29,723 | 1 | 0,1329 | 445,1095 | 7,6209 | 1166,456 | 20490 | 0,44667 |
| IL1R2 | 1,02E+09 | 0,64718 | 25 | 25 | 26,197 | 2 | 0,1329 | 424,0429 | 7,48707 | 1478,385 | 26138 | 0,51333 |
| CALD1 | 3512 | 0,36318 | 23 | 25 | 20,201 | 8 | 0,15505 | 444,3833 | 7,62577 | 3560,728 | 43912 | 0,25 |
| F11R | 811818 | 0,46227 | 25 | 25 | 23,037 | 1 | 0,1329 | 438,3429 | 7,5844 | 2085,017 | 29404 | 0,36667 |
| NFKB2 | 1,56E+09 | 0,68918 | 24 | 25 | 26,817 | 2 | 0,1329 | 432,7762 | 7,54223 | 2599,261 | 43888 | 0,51 |
| HLA-E | 1,31E+12 | 0,58593 | 23 | 25 | 8,546 | 2 | 0,1329 | 377,3595 | 7,15047 | 2871,239 | 40874 | 0,40333 |
| ROCK1 | 97753 | 0,45044 | 24 | 25 | 26,472 | 3 | 0,1329 | 458,1429 | 7,70606 | 4043,305 | 61560 | 0,33333 |
| KRT8 | 4,37E+07 | 0,63513 | 24 | 25 | 27,806 | 1 | 0,1329 | 455,8762 | 7,69471 | 2240,36 | 42384 | 0,47 |
| EFNB1 | 72893 | 0,40192 | 23 | 24 | 18,442 | 6 | 0,1329 | 451,3429 | 7,67119 | 7741,511 | 93422 | 0,30072 |
| NID1 | 2689473 | 0,70089 | 21 | 24 | 22,707 | 3 | 0,1329 | 423,9595 | 7,49356 | 5033,138 | 78366 | 0,44928 |
| ASCL1 | 14470 | 0,47747 | 24 | 24 | 19,789 | 1 | 0,1329 | 439,1595 | 7,60306 | 1393,18 | 24912 | 0,38406 |
| GLUL | 4732 | 0,41544 | 19 | 24 | 12,741 | 3 | 0,1329 | 432,3095 | 7,55845 | 7734,01 | 89498 | 0,23913 |
| NQO1 | 155 | 0,24122 | 19 | 24 | 10,477 | 7 | 0,1329 | 421,1595 | 7,46923 | 14936,79 | 165930 | 0,13043 |
| FOXC2 | 18824 | 0,43243 | 24 | 24 | 22,806 | 2 | 0,1329 | 432,0929 | 7,54871 | 1194,322 | 22874 | 0,34783 |
| MT-ND1 | 9,58E+08 | 0,46971 | 23 | 24 | 13,149 | 1 | 0,1329 | 435,6095 | 7,57143 | 11389,13 | 120408 | 0,35145 |
| CASP7 | 4498 | 0,46235 | 19 | 24 | 19,745 | 2 | 0,1329 | 440,9762 | 7,59819 | 2999,822 | 44084 | 0,26087 |
| LATS2 | 40518 | 0,34844 | 19 | 24 | 12,585 | 3 | 0,1329 | 424,4595 | 7,49681 | 4112,309 | 47436 | 0,19565 |
| AREG | 897250 | 0,48648 | 24 | 24 | 28,025 | 1 | 0,1329 | 453,7762 | 7,6793 | 849,0305 | 18184 | 0,3913 |
| NFATC1 | 10161 | 0,46971 | 23 | 24 | 26,72 | 1 | 0,1329 | 450,1262 | 7,65578 | 1185,1 | 22774 | 0,35145 |
| C4A | 62758 | 0,45044 | 24 | 24 | 13,701 | 6 | 0,1329 | 434,9595 | 7,55764 | 4339,735 | 63580 | 0,36232 |
| CISH | 4,23E+07 | 0,68937 | 22 | 24 | 26,393 | 1 | 0,1329 | 439,4762 | 7,58684 | 1659,197 | 27752 | 0,47826 |
| TEAD1 | 83283 | 0,45035 | 23 | 24 | 16,298 | 3 | 0,1329 | 440,2595 | 7,60711 | 4645,288 | 56294 | 0,33696 |
| YES1 | 23657 | 0,43582 | 23 | 24 | 27,704 | 3 | 0,15505 | 443,2 | 7,6209 | 4866,468 | 73682 | 0,32609 |
| SLC9A3R1 | 711 | 0,25068 | 22 | 23 | 18,029 | 10 | 0,1329 | 442,3095 | 7,61766 | 4742,986 | 61082 | 0,18972 |
| SMAD6 | 26541 | 0,39169 | 22 | 23 | 19,957 | 4 | 0,1329 | 430,9262 | 7,5333 | 9159,382 | 119672 | 0,29644 |
| HSPB2 | 3966 | 0,3561 | 21 | 23 | 21,184 | 2 | 0,1329 | 451,0262 | 7,66227 | 3580,136 | 57054 | 0,24901 |
| TEAD4 | 83369 | 0,49092 | 22 | 23 | 15,222 | 2 | 0,1329 | 425,1762 | 7,51059 | 3631,026 | 46734 | 0,37154 |
| LHX2 | 2223 | 0,41146 | 20 | 23 | 14,12 | 6 | 0,1329 | 421,7095 | 7,49194 | 4748,024 | 57286 | 0,26877 |
| PSMB9 | 12161 | 0,3708 | 22 | 23 | 14,283 | 3 | 0,1329 | 426,0262 | 7,50086 | 3175,468 | 66286 | 0,28063 |
| MBP | 19372 | 0,44653 | 21 | 23 | 18,762 | 1 | 0,1329 | 444,1595 | 7,62577 | 3445,349 | 52964 | 0,31621 |
| FLT3LG | 1596222 | 0,61499 | 23 | 23 | 26,707 | 1 | 0,1329 | 441,4762 | 7,60225 | 429,1103 | 10882 | 0,50198 |
| ABCB1 | 868432 | 0,49393 | 23 | 23 | 24,995 | 3 | 0,1329 | 463,5262 | 7,74094 | 6352,12 | 95334 | 0,40316 |
| HERC5 | 9,55E+10 | 0,9609 | 21 | 22 | 8,245 | 2 | 0,11629 | 369,7941 | 7,09613 | 1778,036 | 22864 | 0,73593 |
| CP | 6626 | 0,40214 | 22 | 22 | 12,07 | 2 | 0,1329 | 429,0762 | 7,53087 | 1618,305 | 22708 | 0,33333 |
| SAMD9L | 1,18E+09 | 0,8535 | 21 | 22 | 7,849 | 1 | 0,1329 | 390,7595 | 7,27295 | 1333,497 | 15672 | 0,65368 |
| SFN | 160 | 0,21412 | 22 | 22 | 16,194 | 1 | 0,15505 | 438,9167 | 7,60144 | 3116,446 | 39454 | 0,17749 |
| CFH | 62595 | 0,4974 | 21 | 22 | 20,882 | 2 | 0,1329 | 428,2429 | 7,5187 | 2841,299 | 54676 | 0,38095 |
| MAVS | 460123 | 0,53132 | 21 | 22 | 15,725 | 2 | 0,1329 | 414,0929 | 7,4303 | 2397,728 | 48072 | 0,40693 |
| PARP14 | 1,42E+08 | 0,74922 | 20 | 22 | 9,506 | 3 | 0,11629 | 383,4012 | 7,21049 | 1967,411 | 23144 | 0,52814 |
| NEDD4 | 77 | 0,23034 | 15 | 22 | 14,604 | 9 | 0,1329 | 437,4595 | 7,58602 | 10981,75 | 108846 | 0,09957 |
| C1R | 54506 | 0,50971 | 20 | 22 | 12,006 | 2 | 0,1329 | 418,7429 | 7,4741 | 2364,623 | 29932 | 0,35931 |
| GNA12 | 381 | 0,28153 | 14 | 21 | 8,183 | 4 | 0,1329 | 401,7262 | 7,34676 | 4635,037 | 53542 | 0,12381 |
| ZIC1 | 678 | 0,42905 | 17 | 21 | 10,844 | 4 | 0,1329 | 405,9762 | 7,38893 | 5072,671 | 59862 | 0,25714 |
| TNFRSF11B | 2178156 | 0,49175 | 21 | 21 | 24,627 | 2 | 0,15505 | 435,6167 | 7,57305 | 2217,167 | 33136 | 0,41429 |
| JUNB | 28651 | 0,522 | 20 | 21 | 20,065 | 1 | 0,1329 | 430,3929 | 7,54628 | 972,2358 | 17392 | 0,40476 |
| IL3RA | 16654 | 0,48045 | 21 | 21 | 19,781 | 1 | 0,15505 | 428,7833 | 7,52682 | 778,2035 | 14168 | 0,40476 |
| EIF4EBP1 | 46225 | 0,33162 | 20 | 21 | 15,633 | 8 | 0,1329 | 441,3262 | 7,6136 | 7932,679 | 96568 | 0,25714 |
| EPHB4 | 54130 | 0,57626 | 19 | 21 | 20,847 | 3 | 0,1329 | 443,1095 | 7,62982 | 1919,926 | 23362 | 0,40952 |
| MT-CO1 | 9,58E+08 | 0,83123 | 15 | 21 | 5,367 | 2 | 0,11629 | 382,7512 | 7,22509 | 4119,631 | 35946 | 0,4 |
| MT-CO2 | 9,58E+08 | 0,77178 | 16 | 21 | 8,016 | 12 | 0,1329 | 421,5762 | 7,49275 | 8685,794 | 73748 | 0,45238 |
| GNG12 | 601 | 0,30706 | 20 | 21 | 6,051 | 3 | 0,1329 | 411,1429 | 7,42219 | 6676,582 | 74988 | 0,2381 |
| INPP5D | 85 | 0,19652 | 20 | 21 | 13,104 | 5 | 0,1329 | 414,9595 | 7,44409 | 5404,19 | 64806 | 0,15238 |
| RXRA | 196 | 0,20772 | 19 | 21 | 9,373 | 3 | 0,15505 | 422,2 | 7,49437 | 4458,379 | 40154 | 0,14762 |
| SLC1A7 | 208 | 0,25463 | 19 | 21 | 11,177 | 6 | 0,1329 | 419,4929 | 7,47977 | 5052,122 | 61732 | 0,18571 |
| SIX1 | 2770 | 0,40532 | 20 | 20 | 15,514 | 1 | 0,1329 | 419,3762 | 7,48626 | 1329,369 | 19994 | 0,34737 |
| EPAS1 | 1480 | 0,39487 | 16 | 20 | 18,909 | 8 | 0,1329 | 450,6262 | 7,67038 | 4583,122 | 50664 | 0,23158 |
| DOCK1 | 62 | 0,27806 | 12 | 20 | 11,58 | 5 | 0,1329 | 419,5595 | 7,4814 | 11657,11 | 114662 | 0,10526 |
| EPS8 | 115 | 0,31933 | 13 | 20 | 11,532 | 7 | 0,15505 | 432,5 | 7,56818 | 7394,451 | 79480 | 0,13158 |
| VEGFB | 2080922 | 0,66324 | 20 | 20 | 25,323 | 1 | 0,1329 | 427,6929 | 7,52844 | 427,5582 | 7670 | 0,56842 |
| BAG3 | 53 | 0,21031 | 15 | 20 | 9,933 | 3 | 0,1329 | 409,7429 | 7,41732 | 5087,029 | 54022 | 0,11053 |
| GNG5 | 595 | 0,32163 | 19 | 20 | 4,282 | 1 | 0,11629 | 364,1845 | 7,07099 | 1799,523 | 21082 | 0,25263 |
| TRIM22 | 1,34E+10 | 0,96964 | 18 | 20 | 7,357 | 1 | 0,11629 | 361,8274 | 7,03936 | 871,2307 | 8456 | 0,7 |
| FOXC1 | 4166 | 0,52814 | 20 | 20 | 18,603 | 1 | 0,1329 | 434,9762 | 7,57872 | 921,9913 | 19160 | 0,45263 |
| CDH23 | 59 | 0,38186 | 9 | 20 | 5,633 | 4 | 0,1329 | 392,9595 | 7,29728 | 9019,84 | 80836 | 0,10526 |
| ALDH7A1 | 47 | 0,31026 | 9 | 20 | 7,849 | 1 | 0,1329 | 405,0429 | 7,38082 | 5439,329 | 57812 | 0,10526 |
| ICAM2 | 4,08E+07 | 0,70009 | 20 | 20 | 23,998 | 1 | 0,1329 | 426,2595 | 7,51546 | 315,153 | 7910 | 0,6 |
| C4B | 58744 | 0,46673 | 20 | 20 | 10,809 | 1 | 0,1329 | 416,5762 | 7,44327 | 2432,532 | 39500 | 0,4 |
| NAMPT | 738904 | 0,57477 | 17 | 19 | 19,832 | 6 | 0,1329 | 435,1929 | 7,57548 | 5549,283 | 69426 | 0,4152 |
| SLC11A1 | 969 | 0,5012 | 9 | 19 | 7,43 | 9 | 0,1329 | 398,5095 | 7,33621 | 3623,672 | 33786 | 0,19298 |
| GNG11 | 594 | 0,32163 | 19 | 19 | 3,827 | 5 | 0,11629 | 361,4179 | 7,04828 | 1364,213 | 18868 | 0,2807 |
| SMARCC1 | 115 | 0,26923 | 16 | 19 | 9,593 | 5 | 0,1329 | 418,3095 | 7,4741 | 5135,331 | 56198 | 0,17544 |
| TBX3 | 3311 | 0,49951 | 18 | 19 | 16,467 | 1 | 0,1329 | 428,4095 | 7,54142 | 1152,313 | 18646 | 0,39766 |
| SERPING1 | 46942 | 0,59684 | 14 | 19 | 11,138 | 3 | 0,1329 | 401,9595 | 7,35406 | 1090,942 | 16178 | 0,32749 |
| ROBO4 | 17668 | 0,44224 | 19 | 19 | 16,631 | 2 | 0,1329 | 423,1262 | 7,50167 | 2344,227 | 38336 | 0,38596 |
| CFLAR | 733854 | 0,58287 | 17 | 19 | 18,022 | 1 | 0,1329 | 432,3262 | 7,55439 | 1580,553 | 26254 | 0,42105 |

| | | | | | | | | | | | | |
|---|---|---|---|---|---|---|---|---|---|---|---|---|
| MT-CYB | 9,58E+08 | 0,83123 | 15 | 19 | 2,984 | 1 | 0,11629 | 349,3036 | 6,94852 | 2041,477 | 16742 | 0,48538 |
| VSIG4 | 820 | 0,29953 | 17 | 19 | 5,12 | 4 | 0,1329 | 382,1095 | 7,22428 | 5190,063 | 52144 | 0,21637 |
| ACVRL1 | 52204 | 0,59906 | 17 | 19 | 20,29 | 3 | 0,1329 | 419,1762 | 7,4741 | 3639,765 | 54342 | 0,43275 |
| CEBPD | 41670 | 0,56286 | 19 | 19 | 20,198 | 1 | 0,1329 | 431,1095 | 7,55277 | 411,3407 | 8430 | 0,49123 |
| HELZ2 | 10349 | 0,73986 | 9 | 19 | 8,993 | 3 | 0,1329 | 399,2262 | 7,34676 | 4621,979 | 45364 | 0,26316 |
| WWTR1 | 40594 | 0,41286 | 17 | 19 | 14,806 | 3 | 0,1329 | 436,5262 | 7,59089 | 4504,153 | 52172 | 0,29825 |
| SERPINH1 | 2689 | 0,44871 | 16 | 19 | 16,709 | 2 | 0,1329 | 425,9762 | 7,52276 | 3139,78 | 47068 | 0,2924 |
| IFITM2 | 1,48E+12 | 0,97699 | 18 | 19 | 8,355 | 1 | 0,11629 | 365,6512 | 7,06937 | 768,5977 | 9752 | 0,77778 |
| TNS1 | 12628 | 0,59684 | 14 | 18 | 21,724 | 6 | 0,1329 | 445,0762 | 7,64848 | 7448,464 | 92520 | 0,35294 |
| HES1 | 23861 | 0,55049 | 17 | 18 | 13,597 | 4 | 0,1329 | 415,8929 | 7,45706 | 864,0423 | 12982 | 0,44444 |
| IRAK3 | 4012104 | 0,61704 | 18 | 18 | 20,671 | 1 | 0,1329 | 420,2762 | 7,47734 | 337,9834 | 6222 | 0,54902 |
| MFGE8 | 46 | 0,24879 | 12 | 18 | 12,966 | 3 | 0,1329 | 403,0762 | 7,37271 | 2226,93 | 25112 | 0,12418 |
| TCF7L1 | 41104 | 0,50675 | 14 | 18 | 7,064 | 1 | 0,15505 | 409,5167 | 7,42624 | 1716,72 | 20668 | 0,30065 |
| AQP1 | 139 | 0,47564 | 7 | 18 | 8,86 | 2 | 0,1329 | 420,2595 | 7,49924 | 8275,601 | 75944 | 0,11111 |
| TNFRSF12A | 2668 | 0,44871 | 16 | 18 | 17,724 | 2 | 0,1329 | 421,3762 | 7,4887 | 2918,758 | 37126 | 0,3268 |
| PLSCR1 | 9,58E+08 | 0,98353 | 13 | 18 | 7,727 | 3 | 0,1329 | 410,6929 | 7,42868 | 6577,128 | 64302 | 0,50327 |
| AKAP9 | 137 | 0,43905 | 7 | 18 | 8,427 | 11 | 0,1329 | 421,6262 | 7,50411 | 13392,62 | 106350 | 0,10458 |
| S100A8 | 525 | 0,3562 | 17 | 18 | 20,113 | 1 | 0,1329 | 434,4762 | 7,57467 | 825,1289 | 16080 | 0,28758 |
| MB21D1 | 15860 | 0,49073 | 15 | 18 | 12,073 | 2 | 0,1329 | 414,1095 | 7,4449 | 3438,914 | 44726 | 0,33333 |
| AHNAK | 223 | 0,30045 | 15 | 18 | 12,007 | 1 | 0,1329 | 429,7762 | 7,55683 | 2278,657 | 27464 | 0,19608 |
| DSP | 364 | 0,38589 | 16 | 18 | 14,96 | 3 | 0,1329 | 428,5595 | 7,54466 | 2963,627 | 36260 | 0,28105 |
| CFB | 56219 | 0,55858 | 17 | 18 | 15,479 | 2 | 0,1329 | 431,3095 | 7,54953 | 2193,751 | 30776 | 0,45098 |
| WWC1 | 40510 | 0,42179 | 16 | 18 | 10,278 | 6 | 0,1329 | 405,0429 | 7,38974 | 3908,209 | 51358 | 0,30719 |
| LFNG | 10906 | 0,40477 | 17 | 17 | 10,043 | 2 | 0,1329 | 389,0762 | 7,27376 | 2254,665 | 41994 | 0,36765 |
| SRGN | 344 | 0,34051 | 15 | 17 | 11,492 | 2 | 0,1329 | 426,7762 | 7,52925 | 2720,51 | 35584 | 0,25 |
| CD93 | 5801 | 0,51801 | 14 | 17 | 17,861 | 3 | 0,1329 | 423,6595 | 7,50816 | 3441,396 | 59010 | 0,34559 |
| PARP12 | 5,12E+07 | 0,81763 | 17 | 17 | 6,48 | 1 | 0,11629 | 361,8179 | 7,06207 | 161,4437 | 3712 | 0,74265 |
| LCAT | 149 | 0,35915 | 10 | 17 | 11,384 | 6 | 0,1329 | 410,2762 | 7,43111 | 5493,082 | 54522 | 0,13235 |
| PDLIM3 | 5088 | 0,49207 | 11 | 17 | 4,342 | 4 | 0,1329 | 380,2429 | 7,21212 | 2943,142 | 21972 | 0,26471 |
| FZD7 | 166 | 0,27524 | 17 | 17 | 14,947 | 2 | 0,15505 | 428,5833 | 7,54385 | 3433,76 | 43726 | 0,25 |
| CXCL3 | 1,92E+10 | 0,97144 | 17 | 17 | 26,198 | 1 | 0,1329 | 414,5595 | 7,44327 | 19,83186 | 560 | 0,88235 |
| DTX3L | 179 | 0,38039 | 6 | 17 | 8,871 | 2 | 0,1329 | 411,2929 | 7,42543 | 4374,97 | 52338 | 0,19118 |
| ADORA2B | 101 | 0,2423 | 16 | 17 | 9,485 | 3 | 0,1329 | 421,4762 | 7,50329 | 3457,871 | 45070 | 0,19853 |
| PCK1 | 222 | 0,37042 | 13 | 17 | 7,473 | 4 | 0,1329 | 389,2929 | 7,28106 | 3113,443 | 32200 | 0,22059 |
| FOXJ1 | 98 | 0,28101 | 13 | 17 | 9,467 | 5 | 0,1329 | 405,1095 | 7,38974 | 8678,374 | 102622 | 0,16912 |
| EYA1 | 9328 | 0,5181 | 17 | 17 | 14,529 | 1 | 0,1329 | 421,7262 | 7,50167 | 842,703 | 14310 | 0,47059 |
| ITSN2 | 23 | 0,28529 | 6 | 17 | 6,74 | 13 | 0,15505 | 422,6833 | 7,51465 | 10856,92 | 79980 | 0,04412 |
| MYO1C | 5052 | 0,64146 | 8 | 17 | 4,085 | 2 | 0,1329 | 390,6929 | 7,29322 | 4065,597 | 27200 | 0,17647 |
| GAB2 | 2179 | 0,41281 | 16 | 17 | 18,591 | 2 | 0,1329 | 431,1429 | 7,56088 | 2302,028 | 32226 | 0,33824 |
| MT-ND2 | 9,58E+08 | 0,75383 | 16 | 17 | 4,059 | 1 | 0,11629 | 369,2179 | 7,12776 | 1300,666 | 15018 | 0,61765 |
| S100A9 | 933 | 0,34999 | 16 | 17 | 17,034 | 2 | 0,1329 | 423,9762 | 7,50492 | 3697,668 | 47280 | 0,28676 |
| MYO6 | 5077 | 0,37162 | 14 | 17 | 4,55 | 4 | 0,1329 | 383,1595 | 7,23807 | 2752,876 | 25066 | 0,24265 |
| AMOT | 40402 | 0,43064 | 15 | 17 | 7,824 | 1 | 0,1329 | 397,9595 | 7,36217 | 1207,435 | 12478 | 0,31618 |
| TGFB1I1 | 115 | 0,30733 | 12 | 17 | 13,427 | 4 | 0,15505 | 420,1667 | 7,49194 | 3481,361 | 44482 | 0,15441 |
| CHD7 | 330 | 0,35052 | 15 | 17 | 12,545 | 1 | 0,1329 | 422,0429 | 7,50654 | 2797,07 | 38704 | 0,25735 |
| SLC4A4 | 26 | 0,28529 | 6 | 17 | 4,404 | 7 | 0,11629 | 399,7179 | 7,35811 | 7478,323 | 58488 | 0,08088 |
| FKBP5 | 24 | 0,25611 | 7 | 17 | 6,22 | 3 | 0,1329 | 399,3429 | 7,34513 | 5240,754 | 49026 | 0,05882 |
| TEAD2 | 82804 | 0,55049 | 17 | 17 | 8,599 | 1 | 0,1329 | 406,0929 | 7,40029 | 2564,563 | 33342 | 0,5 |
| GADD45B | 1298 | 0,49549 | 14 | 16 | 11,49 | 4 | 0,1329 | 412,2929 | 7,42868 | 729,8116 | 9874 | 0,36667 |
| ZC3HAV1 | 205 | 0,35765 | 13 | 16 | 9,139 | 4 | 0,1329 | 414,7262 | 7,45787 | 6031,794 | 62830 | 0,23333 |
| RRAS | 22 | 0,25931 | 5 | 16 | 5,181 | 4 | 0,1329 | 365,4429 | 7,10424 | 4588,1 | 64242 | 0,075 |
| CHI3L1 | 4057327 | 0,75111 | 15 | 16 | 23,51 | 2 | 0,1329 | 443,5429 | 7,6355 | 2419,121 | 26622 | 0,625 |
| SLC40A1 | 2337 | 0,59388 | 11 | 16 | 8,203 | 2 | 0,1329 | 407,0929 | 7,40029 | 4500,22 | 41038 | 0,3 |
| BBOX1 | 21 | 0,21953 | 7 | 16 | 5,048 | 9 | 0,11629 | 406,4845 | 7,39542 | 10798,64 | 92182 | 0,05833 |
| ALDH1L1 | 2926 | 0,55613 | 12 | 16 | 8,598 | 4 | 0,1329 | 396,6095 | 7,34027 | 8580,43 | 117956 | 0,31667 |
| PLIN2 | 162 | 0,39905 | 10 | 16 | 6,287 | 4 | 0,1329 | 398,5429 | 7,36136 | 8236,283 | 81710 | 0,16667 |
| FSTL1 | 2904 | 0,5205 | 16 | 16 | 18,207 | 1 | 0,1329 | 424,8929 | 7,51789 | 194,9815 | 6144 | 0,48333 |
| RHOD | 988 | 0,45368 | 12 | 16 | 11,819 | 2 | 0,1329 | 425,8595 | 7,53168 | 4180,457 | 51740 | 0,25833 |
| MYOF | 90 | 0,40246 | 7 | 16 | 8,679 | 5 | 0,11629 | 415,8179 | 7,46274 | 3226,416 | 35664 | 0,16667 |
| LAMC3 | 7999321 | 0,84125 | 15 | 16 | 17,743 | 1 | 0,15505 | 402,1333 | 7,36947 | 135,9882 | 2368 | 0,7 |
| MT-CO3 | 9,58E+08 | 0,77178 | 16 | 16 | 3,224 | 1 | 0,11629 | 343,7702 | 6,90958 | 376,8247 | 4328 | 0,71667 |
| PODXL | 820 | 0,38288 | 14 | 16 | 14,385 | 2 | 0,15505 | 425,0333 | 7,52925 | 2607,002 | 32616 | 0,28333 |
| OLIG1 | 11803 | 0,53079 | 15 | 16 | 12,263 | 3 | 0,1329 | 409,6595 | 7,42381 | 2488,854 | 34730 | 0,44167 |
| CST3 | 353 | 0,37055 | 15 | 16 | 11,659 | 3 | 0,1329 | 409,9595 | 7,4157 | 1779,259 | 23512 | 0,30833 |
| LAMB2 | 7999344 | 0,78076 | 16 | 16 | 16,53 | 1 | 0,1329 | 401,8095 | 7,36379 | 61,67492 | 1286 | 0,725 |
| LPAR3 | 404 | 0,37162 | 14 | 16 | 14,218 | 4 | 0,1329 | 426,1595 | 7,53655 | 3034,694 | 43014 | 0,275 |
| BMPR1B | 232 | 0,29615 | 16 | 16 | 14,27 | 3 | 0,1329 | 408,2262 | 7,40921 | 2725,12 | 41036 | 0,275 |
| APLNR | 1719 | 0,46832 | 12 | 16 | 13,506 | 2 | 0,1329 | 409,3095 | 7,42543 | 3177,747 | 45034 | 0,29167 |
| WASF2 | 342 | 0,43905 | 12 | 16 | 7,798 | 2 | 0,15505 | 391,2333 | 7,30701 | 2947,805 | 32858 | 0,25 |
| CRX | 42 | 0,38896 | 5 | 15 | 4,681 | 8 | 0,1329 | 392,7262 | 7,31026 | 11919,37 | 99970 | 0,11429 |
| LTBR | 41926 | 0,52077 | 15 | 15 | 16,269 | 1 | 0,1329 | 405,3762 | 7,39137 | 192,1916 | 4894 | 0,49524 |
| PLD2 | 21 | 0,28529 | 6 | 15 | 7,219 | 8 | 0,1329 | 426,3929 | 7,54304 | 13194,21 | 103472 | 0,06667 |
| SLC7A11 | 262 | 0,39905 | 10 | 15 | 10,677 | 1 | 0,1329 | 420,1929 | 7,49599 | 2353,13 | 31304 | 0,21905 |
| FGF11 | 3675030 | 0,70104 | 15 | 15 | 20,774 | 1 | 0,1329 | 441,7095 | 7,62982 | 143,3918 | 3388 | 0,66667 |
| GABRG1 | 86 | 0,37904 | 8 | 15 | 4,325 | 3 | 0,15505 | 366,9667 | 7,12776 | 1955,638 | 15838 | 0,19048 |
| TRIP10 | 25 | 0,33284 | 6 | 15 | 4,619 | 2 | 0,1329 | 400,6929 | 7,38244 | 4156,411 | 31260 | 0,08571 |
| MAP4K4 | 1466 | 0,38319 | 13 | 15 | 16,69 | 1 | 0,1329 | 418,0262 | 7,47653 | 1265,015 | 21398 | 0,29524 |
| TEAD3 | 81364 | 0,57085 | 15 | 15 | 7,673 | 1 | 0,1329 | 384,4929 | 7,25673 | 509,0655 | 6654 | 0,54286 |
| ACSL5 | 95 | 0,43736 | 8 | 15 | 6,023 | 1 | 0,1329 | 388,6262 | 7,28593 | 2579,854 | 20554 | 0,22857 |

| | | | | | | | | | | | | |
|---|---|---|---|---|---|---|---|---|---|---|---|---|
| MT-ND5 | 9,58E+08 | 0,80119 | 15 | 15 | 3,856 | 1 | 0,11629 | 348,2869 | 6,95744 | 320,9852 | 3768 | 0,7619 |
| MT-ND4 | 9,58E+08 | 0,80119 | 15 | 15 | 3,822 | 1 | 0,11629 | 348,2869 | 6,95744 | 320,9852 | 3768 | 0,7619 |
| SRSF11 | 158 | 0,30655 | 13 | 15 | 2,087 | 3 | 0,11629 | 342,8774 | 6,90391 | 4538,648 | 46594 | 0,22857 |
| FAT1 | 87 | 0,2927 | 12 | 15 | 9,482 | 2 | 0,1329 | 429,8262 | 7,56169 | 4344,696 | 42856 | 0,19048 |
| HFE | 1614 | 0,52601 | 11 | 15 | 7,518 | 2 | 0,1329 | 412,3595 | 7,4376 | 3560,178 | 35162 | 0,30476 |
| KLF6 | 210 | 0,52506 | 9 | 15 | 10,22 | 3 | 0,1329 | 402,3929 | 7,37596 | 1446,697 | 17404 | 0,21905 |
| PALLD | 76 | 0,28101 | 13 | 15 | 11,106 | 1 | 0,15505 | 425,1333 | 7,53006 | 3495,818 | 46032 | 0,20952 |
| ZFP36 | 6280 | 0,56202 | 13 | 15 | 12,035 | 1 | 0,1329 | 411,8262 | 7,43435 | 802,2288 | 11298 | 0,41905 |
| H6PD | 22 | 0,37893 | 4 | 15 | 4,704 | 4 | 0,11629 | 384,4845 | 7,24618 | 4628,264 | 47870 | 0,07619 |
| TGFBR3 | 10514 | 0,51801 | 14 | 14 | 12,481 | 1 | 0,1329 | 400,0929 | 7,35811 | 224,8506 | 2736 | 0,50549 |
| TBX2 | 324 | 0,36036 | 14 | 14 | 16,957 | 1 | 0,1329 | 419,8262 | 7,50086 | 565,8044 | 10730 | 0,35165 |
| SP110 | 7257627 | 0,74084 | 13 | 14 | 4,698 | 1 | 0,11629 | 358,1107 | 7,028 | 572,5022 | 6598 | 0,63736 |
| RLBP1 | 283 | 0,31933 | 13 | 14 | 9,424 | 3 | 0,1329 | 409,7929 | 7,43922 | 2893,41 | 30914 | 0,27473 |
| ESAM | 46228 | 0,54053 | 14 | 14 | 15,657 | 1 | 0,1329 | 397,5595 | 7,33865 | 300,8862 | 7546 | 0,52747 |
| CTF1 | 21840 | 0,65315 | 14 | 14 | 15,937 | 1 | 0,1329 | 419,3429 | 7,48626 | 82,31989 | 2456 | 0,63736 |
| CD248 | 394 | 0,45368 | 12 | 14 | 13,852 | 1 | 0,1329 | 403,9929 | 7,38488 | 245,8204 | 4766 | 0,34066 |
| GSTM1 | 223 | 0,52506 | 9 | 14 | 8,794 | 2 | 0,1329 | 389,1929 | 7,28755 | 2293,834 | 26718 | 0,27473 |
| REST | 47 | 0,31924 | 10 | 14 | 7,963 | 1 | 0,1329 | 398,1262 | 7,35487 | 3842,118 | 40758 | 0,1978 |
| PTH1R | 26 | 0,2927 | 7 | 14 | 6,784 | 5 | 0,15505 | 398,6667 | 7,36379 | 2133,354 | 19966 | 0,12088 |
| EBF1 | 85 | 0,28101 | 13 | 14 | 11,149 | 1 | 0,15505 | 405,1333 | 7,40029 | 679,2718 | 8596 | 0,24176 |
| FES | 36 | 0,21948 | 10 | 14 | 11,1 | 7 | 0,1329 | 426,0595 | 7,53979 | 4322,425 | 47954 | 0,17582 |
| SP100 | 82 | 0,47564 | 7 | 14 | 5,885 | 2 | 0,11629 | 379,0345 | 7,21049 | 2458,973 | 27018 | 0,16484 |
| GJB6 | 768 | 0,31531 | 14 | 14 | 9,185 | 1 | 0,1329 | 391,3095 | 7,31107 | 1588,67 | 14900 | 0,30769 |
| CD3E | 281 | 0,39027 | 11 | 14 | 14,322 | 2 | 0,1329 | 415,3262 | 7,45301 | 1086,555 | 23914 | 0,26374 |
| MT-ATP6 | 9,58E+08 | 0,87837 | 14 | 14 | 2,607 | 1 | 0,1329 | 338,8452 | 6,87552 | 90,72697 | 804 | 0,85714 |
| MT-ND4L | 9,58E+08 | 0,91215 | 14 | 14 | 3,687 | 1 | 0,11629 | 341,5036 | 6,89823 | 76,18652 | 1060 | 0,89011 |
| MT-ND3 | 9,58E+08 | 0,91215 | 14 | 14 | 2,954 | 1 | 0,11629 | 341,5036 | 6,89823 | 76,18652 | 1060 | 0,89011 |
| AHCYL1 | 16 | 0,2842 | 4 | 14 | 4,497 | 5 | 0,1329 | 390,8095 | 7,30052 | 8296,695 | 65982 | 0,04396 |
| NFIA | 1000 | 0,57863 | 10 | 14 | 9,502 | 5 | 0,1329 | 406,2929 | 7,41245 | 5565,341 | 54034 | 0,31868 |
| PDLIM1 | 5070 | 0,53872 | 10 | 14 | 6,813 | 1 | 0,15505 | 398,3167 | 7,35406 | 1450,995 | 14356 | 0,30769 |
| TIE1 | 404767 | 0,70252 | 13 | 14 | 16,421 | 2 | 0,1329 | 411,6762 | 7,44246 | 845,4429 | 13500 | 0,6044 |
| LPAR1 | 261 | 0,4242 | 11 | 14 | 9,37 | 4 | 0,1329 | 420,1595 | 7,50411 | 4157,993 | 47048 | 0,27473 |
| GNA14 | 372 | 0,38051 | 12 | 14 | 4,734 | 1 | 0,11629 | 371,1512 | 7,14885 | 1083,119 | 13090 | 0,28571 |
| TRIM5 | 1479 | 0,37042 | 13 | 14 | 6,012 | 1 | 0,1329 | 376,8691 | 7,19022 | 1886,843 | 20350 | 0,31868 |
| TYMP | 741 | 0,3733 | 11 | 14 | 12,799 | 3 | 0,1329 | 418,2929 | 7,48302 | 3878,037 | 49230 | 0,25275 |
| DDR1 | 51 | 0,33413 | 9 | 14 | 10,507 | 1 | 0,1329 | 410,5429 | 7,43111 | 784,8558 | 10590 | 0,16484 |
| SEMA3F | 263 | 0,3366 | 12 | 13 | 11,653 | 1 | 0,1329 | 406,3429 | 7,41245 | 2008,525 | 26094 | 0,29487 |
| MMP15 | 7257840 | 0,79193 | 13 | 13 | 21,298 | 1 | 0,1329 | 434,6762 | 7,58521 | 75,13183 | 3138 | 0,79487 |
| NMI | 253 | 0,3791 | 10 | 13 | 8,389 | 2 | 0,11629 | 378,9345 | 7,19914 | 2246,611 | 24304 | 0,24359 |
| DNAJB1 | 34 | 0,31026 | 9 | 13 | 5,416 | 1 | 0,11629 | 386,5845 | 7,28025 | 1807,045 | 15590 | 0,16667 |
| NKD1 | 27 | 0,2927 | 7 | 13 | 7,362 | 7 | 0,15505 | 388,65 | 7,28187 | 11959,58 | 136884 | 0,14103 |
| TAL1 | 309 | 0,42441 | 12 | 13 | 12,018 | 2 | 0,1329 | 407,8595 | 7,41813 | 2228,729 | 30562 | 0,37179 |
| IL17D | 230 | 0,37042 | 13 | 13 | 12,08 | 1 | 0,1329 | 413,4762 | 7,45139 | 163,8286 | 3712 | 0,37179 |
| TM4SF1 | 79 | 0,57059 | 6 | 13 | 8,731 | 2 | 0,15505 | 388,3 | 7,28187 | 3084,997 | 39832 | 0,15385 |
| ADH1B | 134 | 0,358 | 9 | 13 | 3,762 | 1 | 0,1329 | 367,0929 | 7,12858 | 2149,297 | 19090 | 0,19231 |
| KIF5B | 16 | 0,46346 | 3 | 13 | 6,548 | 7 | 0,1329 | 410,6595 | 7,44409 | 12247,66 | 110180 | 0,0641 |
| DDIT4 | 29 | 0,23943 | 10 | 13 | 6,128 | 3 | 0,1329 | 376,1262 | 7,19995 | 1301,844 | 11984 | 0,15385 |
| EFEMP2 | 302 | 0,3321 | 13 | 13 | 11,955 | 1 | 0,1329 | 397,5429 | 7,34351 | 686,5457 | 11678 | 0,33333 |
| ADCY4 | 58 | 0,40246 | 7 | 13 | 5,015 | 2 | 0,11629 | 368,9774 | 7,13588 | 2258,518 | 22544 | 0,15385 |
| CYBRD1 | 2284 | 0,69213 | 9 | 13 | 4,818 | 3 | 0,1329 | 381,7095 | 7,23969 | 1560,873 | 13696 | 0,38462 |
| ADORA2A | 98 | 0,30655 | 13 | 13 | 10,259 | 1 | 0,1329 | 411,8429 | 7,44327 | 1380,679 | 18624 | 0,30769 |
| GJA4 | 423 | 0,42441 | 12 | 13 | 14,054 | 1 | 0,1329 | 404,9595 | 7,40191 | 509,0114 | 6946 | 0,37179 |
| MT-ND6 | 9,58E+08 | 0,98353 | 13 | 13 | 2,909 | 1 | 0,11629 | 335,0869 | 6,84389 | 0,16667 | 2 | 0,98718 |
| MT-ATP8 | 4,79E+08 | 0,8558 | 13 | 13 | 2,903 | 1 | 0,1329 | 338,3452 | 6,87471 | 78,46496 | 716 | 0,85897 |
| SLC39A1 | 129 | 0,52304 | 6 | 13 | 6,532 | 2 | 0,1329 | 395,5595 | 7,33946 | 2587,932 | 23960 | 0,15385 |
| IL13RA1 | 11047 | 0,61467 | 12 | 13 | 11,454 | 2 | 0,1329 | 398,3429 | 7,34513 | 326,5171 | 4960 | 0,53846 |
| ZC3H12A | 189 | 0,38051 | 12 | 13 | 12,075 | 1 | 0,1329 | 399,7929 | 7,35811 | 498,9419 | 8224 | 0,33333 |
| COX4I2 | 4,79E+08 | 0,89412 | 13 | 13 | 3,098 | 1 | 0,11629 | 335,8536 | 6,85038 | 33,20671 | 704 | 0,89744 |
| ZIC2 | 102 | 0,45346 | 9 | 13 | 6,45 | 2 | 0,1329 | 378,6262 | 7,21131 | 1298,946 | 14338 | 0,24359 |
| RPGR | 16 | 0,46346 | 3 | 13 | 2,686 | 2 | 0,11629 | 360,4845 | 7,07342 | 3659,851 | 20782 | 0,07692 |
| B4GALT1 | 856 | 0,4242 | 11 | 13 | 10,475 | 6 | 0,1329 | 419,6929 | 7,49924 | 3221,773 | 31218 | 0,32051 |
| NFATC2 | 851 | 0,45891 | 10 | 13 | 11,048 | 1 | 0,1329 | 417,2595 | 7,47815 | 1090,065 | 16236 | 0,29487 |
| MGST1 | 205 | 0,36587 | 12 | 13 | 7,131 | 1 | 0,1329 | 404,2429 | 7,39461 | 2638,807 | 32152 | 0,32051 |
| MAML2 | 10262 | 0,51092 | 13 | 13 | 8,646 | 1 | 0,1329 | 394,9262 | 7,33297 | 239,8771 | 3590 | 0,51282 |
| OTUD7B | 65 | 0,38896 | 5 | 13 | 5,6 | 3 | 0,1329 | 378,6095 | 7,20806 | 2152,689 | 20950 | 0,20513 |
| SLC16A3 | 41 | 0,29929 | 10 | 13 | 9,218 | 4 | 0,1329 | 410,5929 | 7,43435 | 3116,505 | 34952 | 0,19231 |
| RNF213 | 408361 | 0,77565 | 12 | 13 | 5,027 | 1 | 0,11629 | 354,0012 | 7,00367 | 747,4419 | 8538 | 0,67949 |
| SMTN | 5049 | 0,76834 | 7 | 13 | 4,638 | 2 | 0,1329 | 390,8095 | 7,3062 | 3049,276 | 26480 | 0,30769 |
| TFPI | 23 | 0,45378 | 5 | 12 | 8,557 | 3 | 0,1329 | 403,1929 | 7,39542 | 2123,309 | 23084 | 0,10606 |
| DOK1 | 279 | 0,4242 | 11 | 12 | 11,517 | 1 | 0,15505 | 418,2333 | 7,49681 | 1625,598 | 19468 | 0,37879 |
| SORBS3 | 66 | 0,4082 | 8 | 12 | 7,089 | 2 | 0,1329 | 392,4762 | 7,32323 | 982,0661 | 10960 | 0,21212 |
| PDLIM4 | 5067 | 0,62053 | 9 | 12 | 3,689 | 1 | 0,1329 | 371,7595 | 7,16994 | 701,1021 | 5406 | 0,39394 |
| IER3 | 337 | 0,50904 | 11 | 12 | 9,297 | 2 | 0,1329 | 382,7762 | 7,24375 | 2344,2 | 38066 | 0,45455 |
| FADS2 | 57 | 0,40246 | 7 | 12 | 4,909 | 5 | 0,11629 | 377,3179 | 7,20806 | 7577,314 | 78908 | 0,16667 |
| STOM | 18 | 0,28529 | 6 | 12 | 6,048 | 2 | 0,11629 | 389,8679 | 7,30377 | 4249,102 | 38356 | 0,09091 |
| BTG2 | 343 | 0,49207 | 11 | 12 | 6,928 | 2 | 0,11629 | 376,7512 | 7,19833 | 838,0503 | 9556 | 0,43939 |
| S100A1 | 37 | 0,43905 | 7 | 12 | 10,896 | 4 | 0,1329 | 411,4429 | 7,46031 | 7859,833 | 76320 | 0,18182 |
| EMCN | 5797 | 0,55995 | 11 | 12 | 14,921 | 3 | 0,1329 | 390,3929 | 7,28998 | 641,5566 | 14670 | 0,5 |
| ANTXR1 | 23 | 0,23866 | 9 | 12 | 10,695 | 1 | 0,15505 | 408,9833 | 7,42786 | 1522,411 | 16488 | 0,15152 |

| | | | | | | | | | | | | |
|---|---|---|---|---|---|---|---|---|---|---|---|---|
| BUB1 | 40 | 0,32073 | 8 | 12 | 6,851 | 2 | 0,1329 | 394,3762 | 7,34027 | 4157,887 | 40968 | 0,18182 |
| ACACB | 80 | 0,32197 | 12 | 12 | 3,666 | 1 | 0,11629 | 363,2345 | 7,09451 | 742,3508 | 7418 | 0,33333 |
| MFNG | 5820 | 0,52686 | 12 | 12 | 9,067 | 6 | 0,1329 | 383,5595 | 7,2478 | 452,9538 | 7484 | 0,54545 |
| CLIC1 | 53 | 0,45378 | 5 | 12 | 8,507 | 5 | 0,1329 | 412,6595 | 7,46112 | 5104,118 | 46220 | 0,19697 |
| GPC5 | 285 | 0,40723 | 11 | 12 | 7,085 | 3 | 0,1329 | 388,1262 | 7,27863 | 672,5292 | 10606 | 0,36364 |
| EFEMP1 | 175 | 0,4296 | 9 | 12 | 10,086 | 4 | 0,1329 | 417,1595 | 7,48221 | 2326,423 | 26092 | 0,28788 |
| SUCLG2 | 15 | 0,25931 | 5 | 12 | 3,82 | 1 | 0,1329 | 374,1429 | 7,18697 | 1977,1 | 16086 | 0,06061 |
| ALDH6A1 | 20 | 0,17957 | 10 | 12 | 2,551 | 1 | 0,11629 | 361,1012 | 7,08397 | 1709,294 | 17940 | 0,13636 |
| PSTPIP1 | 16 | 0,37893 | 4 | 12 | 5,654 | 1 | 0,1329 | 384,9762 | 7,26403 | 2340,833 | 18764 | 0,06061 |
| PDK4 | 188 | 0,39027 | 11 | 11 | 6,014 | 1 | 0,11629 | 379,0845 | 7,22672 | 420,3241 | 5146 | 0,41818 |
| BCL3 | 159 | 0,49567 | 8 | 11 | 8,275 | 3 | 0,1329 | 388,9095 | 7,28998 | 5175,947 | 55258 | 0,30909 |
| LTF | 27 | 0,29157 | 8 | 11 | 5,272 | 2 | 0,1329 | 372,2857 | 7,16913 | 1923,974 | 23186 | 0,2 |
| SMO | 69 | 0,43736 | 8 | 11 | 6,34 | 3 | 0,1329 | 414,1262 | 7,46517 | 5106,337 | 41260 | 0,29091 |
| PMP2 | 19 | 0,33284 | 6 | 11 | 3,379 | 4 | 0,1329 | 361,7262 | 7,09451 | 4357,299 | 54474 | 0,12727 |
| SLC7A5 | 272 | 0,40723 | 11 | 11 | 5,277 | 2 | 0,11629 | 363,5679 | 7,10587 | 535,5772 | 6370 | 0,43636 |
| MECOM | 47 | 0,29929 | 10 | 11 | 8,993 | 2 | 0,1329 | 410,7429 | 7,44652 | 1655,264 | 19686 | 0,27273 |
| PRDM16 | 33 | 0,38039 | 6 | 11 | 9,035 | 4 | 0,1329 | 403,3262 | 7,39623 | 3152,544 | 38346 | 0,16364 |
| OSMR | 12288 | 0,71266 | 11 | 11 | 16,361 | 1 | 0,1329 | 415,9262 | 7,47328 | 26,52631 | 652 | 0,76364 |
| ACAA2 | 21 | 0,26242 | 8 | 11 | 3,003 | 1 | 0,11629 | 365,6107 | 7,10992 | 1662,774 | 14014 | 0,16364 |
| NTSR2 | 76 | 0,32239 | 11 | 11 | 4,491 | 1 | 0,15505 | 364,5833 | 7,11154 | 600,1451 | 3884 | 0,34545 |
| PDLIM2 | 5044 | 0,76834 | 7 | 11 | 4,997 | 1 | 0,1329 | 384,4595 | 7,26159 | 1055,398 | 10538 | 0,38182 |
| CFHR1 | 45385 | 0,73825 | 10 | 11 | 5,426 | 1 | 0,1329 | 352,8857 | 7,00773 | 181,9893 | 2558 | 0,67273 |
| ACSS1 | 30 | 0,31026 | 9 | 11 | 2,088 | 2 | 0,11629 | 349,5179 | 6,97447 | 2367,585 | 21550 | 0,23636 |
| APOL1 | 147 | 0,3791 | 10 | 11 | 6,597 | 3 | 0,1329 | 374,3429 | 7,19346 | 1205,16 | 14442 | 0,34545 |
| EPSTI1 | 161400 | 0,7975 | 11 | 11 | 5,107 | 1 | 0,11629 | 352,6369 | 6,98826 | 40,08459 | 1102 | 0,85455 |
| GCSH | 16 | 0,21953 | 7 | 11 | 5,513 | 6 | 0,1329 | 383,1929 | 7,25916 | 3583,279 | 34586 | 0,12727 |
| TP53BP2 | 19 | 0,33284 | 6 | 11 | 5,88 | 2 | 0,1329 | 387,4429 | 7,29322 | 857,0504 | 7650 | 0,14545 |
| PTRF | 45 | 0,29929 | 10 | 11 | 12,827 | 4 | 0,1329 | 418,9095 | 7,50167 | 1657,792 | 21738 | 0,27273 |
| DDR2 | 864 | 0,57691 | 11 | 11 | 14,842 | 1 | 0,1329 | 425,4429 | 7,53736 | 44,08986 | 1502 | 0,61818 |
| S100A10 | 183 | 0,41901 | 10 | 11 | 8,477 | 4 | 0,1329 | 403,5429 | 7,41083 | 2261,333 | 20802 | 0,38182 |
| FOXO4 | 101 | 0,3791 | 10 | 11 | 10,1 | 2 | 0,1329 | 400,9262 | 7,3792 | 1061,209 | 12890 | 0,34545 |
| ALPL | 15 | 0,37893 | 4 | 11 | 6,144 | 2 | 0,1329 | 391,1262 | 7,3135 | 1147,599 | 11164 | 0,07273 |
| FTL | 1448 | 0,54893 | 9 | 11 | 5,225 | 1 | 0,1329 | 378,1429 | 7,21212 | 637,6993 | 6040 | 0,41818 |
| CNTFR | 1473 | 0,51877 | 10 | 11 | 12,027 | 1 | 0,1329 | 402,4929 | 7,37758 | 313,0132 | 5120 | 0,47273 |
| EMILIN1 | 5046 | 0,54893 | 9 | 11 | 9,595 | 2 | 0,1329 | 375,7429 | 7,19346 | 1927,708 | 35738 | 0,41818 |
| ITPR2 | 14 | 0,46346 | 3 | 11 | 3,489 | 3 | 0,11629 | 364,7179 | 7,11722 | 5714,041 | 50486 | 0,09091 |
| SMTNL2 | 5044 | 0,76834 | 7 | 11 | 2,962 | 3 | 0,1329 | 363,2762 | 7,10505 | 2622,298 | 23118 | 0,4 |
| PKP4 | 38 | 0,36588 | 7 | 11 | 4,959 | 2 | 0,1329 | 376,9762 | 7,20401 | 1174,239 | 9592 | 0,18182 |
| MMD2 | 55 | 0,47549 | 6 | 11 | 4,896 | 12 | 0,15505 | 381,0167 | 7,23969 | 11430,31 | 105310 | 0,2 |
| IL18R1 | 973 | 0,53872 | 10 | 11 | 11,234 | 2 | 0,1329 | 397,2429 | 7,34513 | 1196,035 | 11428 | 0,49091 |
| SIGIRR | 10326 | 0,62782 | 11 | 11 | 13,191 | 1 | 0,1329 | 402,4262 | 7,38163 | 95,1146 | 1988 | 0,67273 |
| ADAMTS9 | 14 | 0,46346 | 3 | 11 | 4,379 | 4 | 0,1329 | 375,8929 | 7,19022 | 3779,225 | 37848 | 0,09091 |
| LUC7L3 | 141 | 0,46652 | 8 | 11 | 1,918 | 2 | 0,11629 | 332,2274 | 6,80333 | 3075,507 | 25148 | 0,29091 |
| HEPH | 2282 | 0,69213 | 9 | 11 | 6,307 | 1 | 0,1329 | 373,2857 | 7,18048 | 1353,68 | 14514 | 0,52727 |
| ADM | 205 | 0,45891 | 10 | 11 | 13,806 | 1 | 0,1329 | 406,6929 | 7,40759 | 210,6512 | 4038 | 0,41818 |
| TXNIP | 66 | 0,358 | 9 | 11 | 10,364 | 2 | 0,1329 | 406,5595 | 7,40921 | 602,0532 | 9898 | 0,27273 |
| GIMAP5 | 22 | 0,2842 | 4 | 11 | 4,426 | 2 | 0,1329 | 375,1929 | 7,18778 | 2137,718 | 16782 | 0,18182 |
| LSR | 127 | 0,64826 | 5 | 11 | 6,191 | 2 | 0,11629 | 392,5012 | 7,32729 | 1152,727 | 10794 | 0,21818 |
| CA12 | 12 | 0,2842 | 4 | 10 | 3,45 | 1 | 0,11629 | 369,0512 | 7,14723 | 1867,57 | 17686 | 0,06667 |
| TRIP6 | 11 | 0,30898 | 3 | 10 | 8,644 | 1 | 0,1329 | 399,6929 | 7,37514 | 342,2392 | 5158 | 0,06667 |
| GSTM2 | 218 | 0,45891 | 10 | 10 | 3,957 | 1 | 0,1329 | 357,6191 | 7,05396 | 456,3161 | 6540 | 0,51111 |
| PVRL2 | 14 | 0,23775 | 6 | 10 | 5,589 | 1 | 0,15505 | 379,8 | 7,22185 | 1111,958 | 10146 | 0,11111 |
| PPP1R1B | 88 | 0,35915 | 10 | 10 | 10,728 | 1 | 0,1329 | 418,3429 | 7,49518 | 416,1245 | 6426 | 0,4 |
| FOSL2 | 294 | 0,45891 | 10 | 10 | 10,818 | 1 | 0,11629 | 398,5512 | 7,36054 | 69,77841 | 1204 | 0,51111 |
| EMP3 | 31 | 0,45378 | 5 | 10 | 6,887 | 4 | 0,1329 | 391,1762 | 7,30782 | 3153,791 | 34614 | 0,15556 |
| S100A11 | 268 | 0,43896 | 10 | 10 | 9,683 | 1 | 0,1329 | 405,7429 | 7,42219 | 113,3241 | 2350 | 0,48889 |
| LRIG1 | 82 | 0,43736 | 8 | 10 | 10,901 | 1 | 0,1329 | 412,7429 | 7,45706 | 652,1154 | 9440 | 0,33333 |
| CDKN2B | 860 | 0,49882 | 10 | 10 | 12,628 | 1 | 0,1329 | 420,9929 | 7,50978 | 90,8306 | 2430 | 0,55556 |
| RAPSN | 873 | 0,57279 | 9 | 10 | 11,103 | 2 | 0,1329 | 405,3595 | 7,41408 | 803,9193 | 10106 | 0,53333 |
| MCM7 | 81 | 0,51223 | 7 | 10 | 5,411 | 2 | 0,1329 | 371,6762 | 7,17237 | 1464,428 | 13980 | 0,31111 |
| RHOJ | 66 | 0,4082 | 8 | 10 | 7,864 | 2 | 0,1329 | 398,2762 | 7,36298 | 3049,985 | 31612 | 0,31111 |
| PIPOX | 11 | 0,30898 | 3 | 10 | 2,242 | 1 | 0,11629 | 338,4941 | 6,86579 | 1474,558 | 9782 | 0,06667 |
| RHBDF2 | 16 | 0,32413 | 5 | 10 | 8,473 | 3 | 0,1329 | 423,3595 | 7,53087 | 4283,492 | 48810 | 0,15556 |
| SREK1 | 131 | 0,47564 | 7 | 10 | 1,717 | 2 | 0,1329 | 306,5452 | 6,53568 | 919,9099 | 6026 | 0,28889 |
| SRGAP1 | 18 | 0,47366 | 4 | 10 | 2,086 | 4 | 0,11629 | 344,3345 | 6,94933 | 7202,344 | 56102 | 0,11111 |
| ENTPD2 | 15 | 0,37893 | 4 | 10 | 4,049 | 1 | 0,11629 | 365,6607 | 7,11235 | 1513,914 | 12396 | 0,13333 |
| PLXNB1 | 44 | 0,27934 | 10 | 10 | 8,641 | 1 | 0,1329 | 403,0595 | 7,39542 | 491,6713 | 9574 | 0,31111 |
| ELTD1 | 20 | 0,33284 | 6 | 10 | 6,601 | 2 | 0,11629 | 363,2179 | 7,08883 | 2603,261 | 44528 | 0,17778 |
| PRRX2 | 14 | 0,25931 | 5 | 10 | 4,336 | 2 | 0,1329 | 357,5429 | 7,04585 | 1235,376 | 10344 | 0,13333 |
| HEYL | 15121 | 0,78759 | 9 | 10 | 9,197 | 1 | 0,1329 | 381,9429 | 7,2405 | 112,1097 | 1494 | 0,73333 |
| ID3 | 55 | 0,40246 | 7 | 10 | 7,517 | 1 | 0,1329 | 398,0595 | 7,35081 | 445,0225 | 5918 | 0,24444 |
| GIG25 | 748 | 0,55399 | 8 | 10 | 9,022 | 1 | 0,1329 | 411,8429 | 7,44976 | 650,6268 | 8908 | 0,42222 |
| MYL3 | 21 | 0,2927 | 7 | 10 | 3,421 | 3 | 0,1329 | 364,5762 | 7,11884 | 1646,949 | 19810 | 0,17778 |
| USP8 | 21 | 0,2927 | 7 | 10 | 6,307 | 3 | 0,15505 | 404,5167 | 7,41489 | 1942,209 | 15786 | 0,17778 |
| BCL2 | 20 | 0,33284 | 6 | 10 | 7,907 | 1 | 0,1329 | 403,7429 | 7,40191 | 1130,096 | 12094 | 0,17778 |
| GGT5 | 30 | 0,56839 | 4 | 10 | 3,37 | 2 | 0,1329 | 357,6595 | 7,05963 | 1126,833 | 10624 | 0,15556 |
| IFI30 | 40322 | 0,8164 | 8 | 10 | 4,12 | 1 | 0,11629 | 342,3607 | 6,89093 | 689,707 | 6630 | 0,62222 |
| MS4A6A | 32 | 0,36588 | 7 | 10 | 4,366 | 1 | 0,1329 | 359,2952 | 7,04422 | 1801,646 | 26710 | 0,28889 |
| MYO10 | 176 | 0,52483 | 8 | 10 | 9,465 | 3 | 0,1329 | 388,8262 | 7,2916 | 909,6469 | 11238 | 0,4 |

| | | | | | | | | | | | | |
|---|---|---|---|---|---|---|---|---|---|---|---|---|
| ERBB2IP | 36 | 0,42794 | 6 | 10 | 11,824 | 1 | 0,1329 | 414,9762 | 7,47734 | 1727,451 | 22048 | 0,2 |
| FYB | 18 | 0,23326 | 8 | 10 | 7,298 | 2 | 0,1329 | 380,1262 | 7,23077 | 737,2337 | 10198 | 0,17778 |
| PLCD1 | 15 | 0,32413 | 5 | 10 | 2,672 | 1 | 0,1329 | 348,2595 | 6,9842 | 2786,029 | 27552 | 0,13333 |
| HPR | 140 | 0,46652 | 8 | 10 | 6,856 | 1 | 0,1329 | 369,0762 | 7,15453 | 216,1937 | 2100 | 0,35556 |
| ALDH3B1 | 129 | 0,47564 | 7 | 10 | 1,924 | 1 | 0,11629 | 331,5107 | 6,80658 | 752,1597 | 5896 | 0,31111 |
| SLC38A3 | 267 | 0,5012 | 9 | 10 | 2,377 | 1 | 0,11629 | 350,7012 | 6,99556 | 311,1131 | 2442 | 0,46667 |
| SNTA1 | 11 | 0,2842 | 4 | 9 | 4,904 | 1 | 0,11629 | 376,9179 | 7,21617 | 2683,65 | 29694 | 0,11111 |
| PON2 | 14 | 0,32413 | 5 | 9 | 3,805 | 2 | 0,11629 | 367,2012 | 7,14155 | 1044,631 | 7342 | 0,16667 |
| SEMA3G | 129 | 0,57059 | 6 | 9 | 9,488 | 1 | 0,1329 | 396,9595 | 7,35487 | 314,6374 | 7422 | 0,36111 |
| NKTR | 16 | 0,38896 | 5 | 9 | 2,472 | 1 | 0,11629 | 359,2012 | 7,07991 | 2703,822 | 20122 | 0,16667 |
| TCF15 | 31 | 0,38039 | 6 | 9 | 6,987 | 2 | 0,1329 | 385,8429 | 7,27619 | 1669,217 | 25614 | 0,22222 |
| ANXA13 | 35 | 0,32073 | 8 | 9 | 7,306 | 1 | 0,1329 | 371,6429 | 7,18292 | 130,5399 | 1698 | 0,30556 |
| FOXF1 | 76 | 0,47564 | 7 | 9 | 7,319 | 2 | 0,1329 | 372,3262 | 7,17805 | 2431,485 | 38426 | 0,36111 |
| NLRC5 | 847 | 0,6123 | 8 | 9 | 4,077 | 2 | 0,11629 | 340,7941 | 6,87714 | 2162,612 | 58248 | 0,58333 |
| DHTKD1 | 10 | 0,30898 | 3 | 9 | 1,776 | 1 | 0,11629 | 340,8702 | 6,89255 | 1966,028 | 11242 | 0,08333 |
| GPAM | 78 | 0,4296 | 9 | 9 | 3,958 | 1 | 0,11629 | 354,6179 | 7,03611 | 220,0867 | 2386 | 0,5 |
| TCIRG1 | 11 | 0,2842 | 4 | 9 | 7,259 | 2 | 0,15505 | 407,4333 | 7,42868 | 2067,458 | 20572 | 0,11111 |
| HES6 | 10800 | 0,76373 | 9 | 9 | 7,609 | 1 | 0,1329 | 379,3595 | 7,22591 | 5,03184 | 144 | 0,88889 |
| INHBB | 10 | 0,30898 | 3 | 9 | 4,978 | 1 | 0,11629 | 366,6012 | 7,13344 | 449,1232 | 4454 | 0,13889 |
| IL17RC | 160 | 0,45346 | 9 | 9 | 10,284 | 1 | 0,1329 | 393,3262 | 7,32242 | 41,09776 | 784 | 0,52778 |
| CLEC4E | 266 | 0,62199 | 7 | 9 | 12,392 | 2 | 0,1329 | 401,6762 | 7,38407 | 5883,336 | 66130 | 0,47222 |
| ACOX2 | 14 | 0,32413 | 5 | 9 | 2,986 | 5 | 0,11629 | 356,4012 | 7,04909 | 1685,868 | 18298 | 0,13889 |
| CHDH | 10 | 0,30898 | 3 | 9 | 2,064 | 5 | 0,11629 | 340,7274 | 6,90228 | 1907,782 | 12478 | 0,08333 |
| SRRM1 | 130 | 0,358 | 9 | 9 | 2,004 | 1 | 0,11629 | 309,3988 | 6,55676 | 876,9229 | 6150 | 0,41667 |
| HMG20B | 10 | 0,30898 | 3 | 9 | 1,866 | 2 | 0,11629 | 339,2179 | 6,89255 | 3296,044 | 28182 | 0,08333 |
| FOXL2 | 13 | 0,23775 | 6 | 9 | 5,463 | 1 | 0,1329 | 366,8595 | 7,13263 | 589,565 | 5150 | 0,13889 |
| HIST2H3D | 19 | 0,26242 | 8 | 9 | 4,368 | 4 | 0,1329 | 378,3595 | 7,22347 | 977,0488 | 14246 | 0,25 |
| PLEKHA7 | 9 | 0,30779 | 2 | 9 | 4,745 | 4 | 0,1329 | 372,1262 | 7,17724 | 2374,607 | 24948 | 0,05556 |
| TES | 10 | 0,30898 | 3 | 9 | 5,865 | 1 | 0,1329 | 386,4429 | 7,29404 | 1113,41 | 10914 | 0,05556 |
| ATP1A2 | 11 | 0,2842 | 4 | 9 | 3,5 | 1 | 0,1329 | 366,7595 | 7,1375 | 2992,606 | 27134 | 0,11111 |
| PNP | 10 | 0,30898 | 3 | 9 | 5,909 | 1 | 0,1329 | 401,2429 | 7,39055 | 2135,878 | 21308 | 0,05556 |
| MAPKAPK2 | 10 | 0,30898 | 3 | 9 | 4,973 | 1 | 0,11629 | 365,3512 | 7,12939 | 1764,038 | 22628 | 0,08333 |
| TAGLN2 | 18 | 0,38896 | 5 | 9 | 5,851 | 1 | 0,1329 | 375,8762 | 7,21049 | 745,2132 | 8702 | 0,16667 |
| RAB13 | 11 | 0,2842 | 4 | 9 | 3,745 | 3 | 0,1329 | 368,7262 | 7,15696 | 4562,922 | 44234 | 0,08333 |
| DUSP5 | 28 | 0,36588 | 7 | 9 | 8,302 | 1 | 0,1329 | 397,5762 | 7,36136 | 210,8926 | 3782 | 0,27778 |
| ROR1 | 84 | 0,40573 | 9 | 9 | 9,743 | 1 | 0,15505 | 413,4 | 7,46842 | 99,34974 | 2352 | 0,47222 |
| MMRN2 | 9 | 0,30779 | 2 | 9 | 5,029 | 4 | 0,11629 | 353,1845 | 7,00529 | 3151,812 | 55890 | 0,05556 |
| EDA | 12 | 0,25931 | 5 | 9 | 8,044 | 2 | 0,15505 | 411,0667 | 7,45382 | 1168,758 | 13098 | 0,11111 |
| ID4 | 488 | 0,52506 | 9 | 9 | 7,686 | 1 | 0,1329 | 392,4595 | 7,32486 | 129,4251 | 1662 | 0,61111 |
| NKX3-1 | 53 | 0,47549 | 6 | 9 | 7,308 | 1 | 0,1329 | 386,0762 | 7,28187 | 346,9408 | 4184 | 0,27778 |
| SOD3 | 41 | 0,34989 | 8 | 9 | 9,143 | 3 | 0,1329 | 406,8762 | 7,41326 | 4293,791 | 42004 | 0,33333 |
| LGALS9 | 81 | 0,43736 | 8 | 9 | 12,382 | 2 | 0,1329 | 412,5262 | 7,44895 | 1194,778 | 15354 | 0,41667 |
| CASP4 | 42 | 0,31026 | 9 | 9 | 9,103 | 1 | 0,1329 | 396,0762 | 7,34027 | 139,9806 | 2786 | 0,36111 |
| RAPGEF3 | 17 | 0,33284 | 6 | 9 | 6,569 | 1 | 0,1329 | 385,5762 | 7,27457 | 780,5287 | 9152 | 0,19444 |
| SLC12A3 | 12 | 0,25931 | 5 | 9 | 2,359 | 1 | 0,1329 | 339,4595 | 6,90715 | 1305,638 | 9514 | 0,11111 |
| TMEM176B | 19 | 0,33284 | 6 | 9 | 2,501 | 3 | 0,15505 | 344,9167 | 6,94365 | 3978,659 | 41740 | 0,22222 |
| FLI1 | 37 | 0,32073 | 8 | 9 | 9,364 | 1 | 0,1329 | 396,2095 | 7,34432 | 207,1607 | 3192 | 0,30556 |
| EMX2 | 36 | 0,33413 | 9 | 9 | 7,103 | 3 | 0,1329 | 372,3262 | 7,17319 | 608,0276 | 7950 | 0,38889 |
| NOTCH2NL | 30 | 0,45378 | 5 | 9 | 4,966 | 5 | 0,1329 | 354,4762 | 7,02314 | 9176,657 | 128948 | 0,19444 |
| KANK2 | 10 | 0,30898 | 3 | 9 | 2,643 | 1 | 0,1329 | 352,3762 | 7,01584 | 724,6614 | 5020 | 0,08333 |
| SLC38A5 | 259 | 0,55399 | 8 | 9 | 2,477 | 1 | 0,11629 | 355,4679 | 7,04098 | 967,2037 | 13266 | 0,52778 |
| EPHX1 | 76 | 0,47564 | 7 | 9 | 4,165 | 1 | 0,1329 | 364,6762 | 7,12533 | 671,6196 | 6776 | 0,36111 |
| TCN2 | 10 | 0,2842 | 4 | 8 | 2,446 | 2 | 0,11629 | 347,3345 | 6,98015 | 1067,559 | 9424 | 0,14286 |
| RIN3 | 12 | 0,37893 | 4 | 8 | 3,969 | 1 | 0,11629 | 349,9774 | 6,9915 | 835,3019 | 8190 | 0,17857 |
| MDFI | 8 | 0,30779 | 2 | 8 | 1,706 | 2 | 0,11629 | 337,1179 | 6,89174 | 3188,012 | 26578 | 0,03571 |
| MT2A | 34 | 0,32073 | 8 | 8 | 4,94 | 3 | 0,11629 | 371,0679 | 7,16589 | 1001,951 | 16874 | 0,39286 |
| SLC6A11 | 32 | 0,47549 | 6 | 8 | 3,805 | 3 | 0,1329 | 363,0595 | 7,11479 | 1080,744 | 14856 | 0,35714 |
| PRKX | 13 | 0,32413 | 5 | 8 | 4,723 | 3 | 0,1329 | 376,6095 | 7,22023 | 757,7118 | 9742 | 0,17857 |
| CCAR1 | 11 | 0,46346 | 3 | 8 | 2,954 | 1 | 0,1329 | 358,0762 | 7,07423 | 1659,321 | 16288 | 0,10714 |
| CYP11A1 | 9 | 0,30898 | 3 | 8 | 3,572 | 2 | 0,1329 | 359,3595 | 7,07991 | 2043,669 | 18898 | 0,10714 |
| PTGER1 | 128 | 0,4082 | 8 | 8 | 4,504 | 1 | 0,11629 | 365,1679 | 7,12128 | 486,9099 | 9592 | 0,5 |
| CXCL16 | 40320 | 0,8164 | 8 | 8 | 12,909 | 1 | 0,1329 | 391,0429 | 7,30458 | 0 | 0 | 1 |
| KLF15 | 57 | 0,43905 | 7 | 8 | 7,54 | 2 | 0,11629 | 396,0845 | 7,35 | 271,1978 | 3904 | 0,42857 |
| MASP1 | 5760 | 0,75809 | 8 | 8 | 4,915 | 1 | 0,1329 | 341,5857 | 6,91364 | 13,42076 | 202 | 0,92857 |
| GALNT10 | 8 | 0,30779 | 2 | 8 | 3,902 | 2 | 0,1329 | 354,6262 | 7,03692 | 656,1101 | 4822 | 0,10714 |
| ARHGEF40 | 9 | 0,30898 | 3 | 8 | 1,298 | 2 | 0,10337 | 283,1694 | 6,25585 | 1023,347 | 5142 | 0,10714 |
| TRIM56 | 19 | 0,32929 | 7 | 8 | 2,073 | 1 | 0,11629 | 331,3441 | 6,80658 | 789,0873 | 7542 | 0,32143 |
| SDPR | 11 | 0,46346 | 3 | 8 | 5,26 | 6 | 0,11629 | 377,7679 | 7,23158 | 6653,365 | 61276 | 0,10714 |
| SLCO1A2 | 8 | 0,30779 | 2 | 8 | 3,138 | 1 | 0,11629 | 347,0679 | 6,98015 | 2919,751 | 37092 | 0,03571 |
| RFX2 | 10 | 0,2842 | 4 | 8 | 2,709 | 6 | 0,11629 | 346,5774 | 6,96555 | 7100,109 | 75470 | 0,10714 |
| MLKL | 63 | 0,51223 | 7 | 8 | 7,435 | 1 | 0,1329 | 375,0762 | 7,19914 | 265,5504 | 3456 | 0,5 |
| P2RY2 | 36 | 0,51861 | 5 | 8 | 9,576 | 2 | 0,1329 | 404,4429 | 7,4084 | 344,6998 | 5232 | 0,39286 |
| LILRB1 | 13 | 0,32413 | 5 | 8 | 4,541 | 1 | 0,1329 | 367,7691 | 7,14885 | 711,7117 | 8182 | 0,21429 |
| AGTRAP | 124 | 0,52304 | 6 | 8 | 7,416 | 1 | 0,1329 | 405,1929 | 7,41408 | 943,8972 | 10158 | 0,39286 |
| TENC1 | 19 | 0,2927 | 7 | 8 | 6,988 | 1 | 0,1329 | 387,4929 | 7,29728 | 352,5309 | 4260 | 0,28571 |
| MAOA | 12 | 0,37893 | 4 | 8 | 3,814 | 3 | 0,11629 | 368,0512 | 7,14399 | 2372,005 | 24944 | 0,14286 |
| CNTLN | 9 | 0,30898 | 3 | 8 | 3,134 | 3 | 0,11629 | 359,4512 | 7,08559 | 6675,968 | 55502 | 0,07143 |
| CD99 | 13 | 0,32413 | 5 | 8 | 6,643 | 2 | 0,1329 | 376,1262 | 7,21049 | 1327,897 | 18596 | 0,17857 |
| IL15RA | 39 | 0,40246 | 7 | 8 | 6,348 | 1 | 0,11629 | 383,6345 | 7,24699 | 92,08865 | 1132 | 0,39286 |

| | | | | | | | | | | | | |
|---|---|---|---|---|---|---|---|---|---|---|---|---|
| TRIOBP | 26 | 0,42794 | 6 | 8 | 2,035 | 1 | 0,15505 | 336,5167 | 6,87795 | 571,8243 | 3710 | 0,32143 |
| A4GALT | 12 | 0,37893 | 4 | 8 | 4,162 | 1 | 0,1329 | 370,0929 | 7,16507 | 1034,749 | 10766 | 0,17857 |
| ARPC1B | 8 | 0,30779 | 2 | 8 | 4,444 | 1 | 0,1329 | 372,1595 | 7,18211 | 872,7009 | 7328 | 0,03571 |
| APLN | 75 | 0,47564 | 7 | 8 | 9,803 | 2 | 0,1329 | 397,4095 | 7,35811 | 1325,62 | 13848 | 0,46429 |
| NDRG1 | 11 | 0,46346 | 3 | 8 | 2,768 | 2 | 0,1329 | 357,0595 | 7,06288 | 3965,241 | 33818 | 0,14286 |
| CYP21A2 | 11 | 0,46346 | 3 | 8 | 2,316 | 3 | 0,11629 | 347,9274 | 6,96312 | 3140,756 | 27930 | 0,17857 |
| SLC15A2 | 16 | 0,33284 | 6 | 8 | 3,474 | 2 | 0,11629 | 354,9679 | 7,04747 | 2727,846 | 33210 | 0,25 |
| IL32 | 728 | 0,52483 | 8 | 8 | 11,787 | 1 | 0,1329 | 417,2095 | 7,48464 | 42,25396 | 1726 | 0,64286 |
| GSDMD | 16 | 0,33284 | 6 | 8 | 5,248 | 1 | 0,1329 | 376,5929 | 7,20238 | 677,7634 | 5716 | 0,25 |
| STON2 | 8 | 0,30779 | 2 | 8 | 3,362 | 3 | 0,1329 | 364,4429 | 7,13506 | 4064,09 | 37618 | 0,07143 |
| ZNF423 | 12 | 0,37893 | 4 | 8 | 5,985 | 2 | 0,15505 | 383,1333 | 7,26646 | 1430,919 | 14706 | 0,17857 |
| TNFSF14 | 126 | 0,37904 | 8 | 8 | 6,827 | 1 | 0,11629 | 363,3345 | 7,101 | 117,8803 | 1456 | 0,46429 |
| PIG6 | 8 | 0,30779 | 2 | 8 | 1,903 | 2 | 0,11629 | 329,4441 | 6,79603 | 2845,044 | 27792 | 0,07143 |
| ADCYAP1R1 | 11 | 0,46346 | 3 | 8 | 3,019 | 1 | 0,11629 | 354,1274 | 7,02962 | 739,594 | 6748 | 0,17857 |
| BOD1L1 | 7 | 0,30779 | 2 | 7 | 2,003 | 2 | 0,11629 | 343,8345 | 6,95338 | 1091,027 | 6906 | 0,04762 |
| ST6GALNAC1 | 8 | 0,30898 | 3 | 7 | 2,303 | 2 | 0,11629 | 341,3607 | 6,92905 | 2653,07 | 27192 | 0,14286 |
| FCGRT | 50 | 0,58344 | 5 | 7 | 9,619 | 3 | 0,1329 | 402,0762 | 7,3865 | 404,067 | 6988 | 0,42857 |
| TAX1BP3 | 7 | 0 | 1 | 7 | 3,778 | 1 | 0,11629 | 364,8512 | 7,12858 | 1402,925 | 13746 | 0 |
| THOC2 | 14 | 0,38896 | 5 | 7 | 3,371 | 2 | 0,1329 | 362,9762 | 7,11073 | 2719,459 | 32574 | 0,28571 |
| F2RL3 | 122 | 0,64826 | 5 | 7 | 2,4 | 1 | 0,11629 | 333,5274 | 6,84145 | 112,4745 | 1152 | 0,47619 |
| BHMT2 | 14 | 0,25611 | 7 | 7 | 1,663 | 1 | 0,11629 | 316,6607 | 6,68086 | 262,9016 | 1914 | 0,33333 |
| GSTM5 | 168 | 0,5854 | 7 | 7 | 1,899 | 1 | 0,1329 | 320,4381 | 6,7133 | 32,41893 | 326 | 0,7619 |
| ABCC9 | 7 | 0,30779 | 2 | 7 | 1,942 | 1 | 0,11629 | 339,0845 | 6,90634 | 1876,072 | 14128 | 0,04762 |
| GLIS2 | 8 | 0,30898 | 3 | 7 | 2,221 | 4 | 0,11629 | 335,4441 | 6,86254 | 3112,842 | 26680 | 0,09524 |
| EHD2 | 15 | 0,33284 | 6 | 7 | 6,139 | 1 | 0,1329 | 382,4762 | 7,26727 | 421,4568 | 4036 | 0,33333 |
| SLC12A7 | 8 | 0,30898 | 3 | 7 | 1,957 | 2 | 0,11629 | 339,2774 | 6,91607 | 985,5142 | 8872 | 0,14286 |
| CTDSP1 | 13 | 0,28529 | 6 | 7 | 4,083 | 1 | 0,1329 | 375,1429 | 7,21212 | 585,619 | 6772 | 0,28571 |
| ROM1 | 13 | 0,46346 | 3 | 7 | 2,616 | 1 | 0,11629 | 358,2679 | 7,08072 | 347,4239 | 3386 | 0,28571 |
| SLC26A2 | 7 | 0,30779 | 2 | 7 | 2,057 | 2 | 0,11629 | 331,4869 | 6,82848 | 1109,325 | 8312 | 0,04762 |
| CD109 | 50 | 0,58344 | 5 | 7 | 9,538 | 2 | 0,1329 | 409,4762 | 7,44895 | 1870,73 | 17524 | 0,42857 |
| STC1 | 8 | 0,30898 | 3 | 7 | 4,023 | 4 | 0,1329 | 377,8429 | 7,21861 | 2467,237 | 23204 | 0,09524 |
| HEPACAM | 27 | 0,56839 | 4 | 7 | 3,371 | 1 | 0,11629 | 375,3179 | 7,21049 | 1180,089 | 11400 | 0,33333 |
| DCHS1 | 13 | 0,28529 | 6 | 7 | 3,74 | 1 | 0,1329 | 373,7429 | 7,18535 | 609,1288 | 6256 | 0,28571 |
| MS4A7 | 20 | 0,45378 | 5 | 7 | 2,772 | 3 | 0,1329 | 337,0786 | 6,8739 | 2864,273 | 35948 | 0,33333 |
| LRTOMT | 12 | 0,2842 | 4 | 7 | 2,171 | 1 | 0,11629 | 316,4607 | 6,67599 | 403,7171 | 2470 | 0,28571 |
| MT1E | 60 | 0,47564 | 7 | 7 | 1,961 | 2 | 0,11629 | 315,2036 | 6,66464 | 828,4095 | 21378 | 0,61905 |
| HSPA6 | 15 | 0,47366 | 4 | 7 | 3,599 | 1 | 0,11629 | 361,2179 | 7,10424 | 233,197 | 2226 | 0,28571 |
| SNX33 | 11 | 0,37893 | 4 | 7 | 1,858 | 2 | 0,11629 | 335,3845 | 6,8666 | 823,8477 | 5718 | 0,19048 |
| MOBP | 26 | 0,36588 | 7 | 7 | 4,216 | 1 | 0,1329 | 356,1262 | 7,0645 | 86,6801 | 612 | 0,47619 |
| ZHX2 | 8 | 0,30898 | 3 | 7 | 4,695 | 1 | 0,1329 | 397,0262 | 7,36703 | 1286,469 | 12258 | 0,09524 |
| GIMAP7 | 8 | 0,30898 | 3 | 7 | 1,668 | 2 | 0,1329 | 324,2357 | 6,76278 | 744,2025 | 5142 | 0,14286 |
| SIX5 | 7 | 0,30779 | 2 | 7 | 4,464 | 1 | 0,11629 | 356,6345 | 7,06531 | 163,4697 | 1630 | 0,09524 |
| TNIP2 | 127 | 0,57059 | 6 | 7 | 7,495 | 2 | 0,11629 | 362,7512 | 7,09938 | 589,2052 | 8236 | 0,57143 |
| PRX | 8 | 0,30898 | 3 | 7 | 5,546 | 3 | 0,1329 | 375,4095 | 7,22104 | 1473,776 | 16062 | 0,14286 |
| GPR39 | 7 | 0,30779 | 2 | 7 | 4,4 | 5 | 0,1329 | 394,2929 | 7,35081 | 2542,985 | 20292 | 0,04762 |
| PRKCH | 10 | 0,46346 | 3 | 7 | 3,826 | 1 | 0,1329 | 370,3429 | 7,17805 | 627,7482 | 8062 | 0,14286 |
| CCDC88A | 9 | 0,2842 | 4 | 7 | 6,096 | 4 | 0,1329 | 406,5929 | 7,43111 | 1858,829 | 19036 | 0,14286 |
| LIMK2 | 8 | 0,30898 | 3 | 7 | 4,237 | 2 | 0,11629 | 349,7179 | 7,00043 | 2639,583 | 32812 | 0,09524 |
| PRELP | 11 | 0,37893 | 4 | 7 | 5,611 | 3 | 0,1329 | 390,4262 | 7,31431 | 2296,369 | 19114 | 0,19048 |
| TAP1 | 241 | 0,66569 | 6 | 7 | 3,494 | 1 | 0,11629 | 336,2036 | 6,84956 | 315,5143 | 3388 | 0,66667 |
| OPHN1 | 7 | 0 | 1 | 7 | 2,058 | 2 | 0,11629 | 333,7179 | 6,85605 | 3626,671 | 31140 | 0 |
| ARHGEF26 | 9 | 0,2842 | 4 | 7 | 5,047 | 2 | 0,1329 | 365,1929 | 7,12209 | 1866,402 | 24096 | 0,14286 |
| HS3ST3B1 | 10 | 0,46346 | 3 | 7 | 2,616 | 1 | 0,1329 | 339,4262 | 6,90147 | 911,5944 | 10782 | 0,19048 |
| ARHGAP11A | 7 | 0,30779 | 2 | 7 | 1,596 | 7 | 0,1329 | 331,3929 | 6,84064 | 3740,247 | 26744 | 0,04762 |
| GPR37L1 | 8 | 0,30898 | 3 | 7 | 3,062 | 1 | 0,1329 | 369,3595 | 7,16832 | 891,595 | 10652 | 0,14286 |
| PNISR | 132 | 0,51223 | 7 | 7 | 1,278 | 1 | 0,11629 | 293,406 | 6,40509 | 105,8278 | 958 | 0,66667 |
| CNN3 | 127 | 0,57059 | 6 | 7 | 7,69 | 1 | 0,1329 | 399,4595 | 7,38163 | 1241,367 | 11726 | 0,57143 |
| HPSE2 | 11 | 0,37893 | 4 | 7 | 5,093 | 1 | 0,1329 | 374,2762 | 7,19265 | 183,4465 | 2656 | 0,19048 |
| MOCS1 | 7 | 0,30779 | 2 | 7 | 1,643 | 1 | 0,11629 | 326,0441 | 6,75872 | 793,5628 | 4150 | 0,04762 |
| EIF4EBP2 | 9 | 0,2842 | 4 | 7 | 2,476 | 2 | 0,11629 | 345,2512 | 6,93229 | 1841,865 | 14848 | 0,14286 |
| LPIN3 | 11 | 0,46346 | 3 | 7 | 3,612 | 1 | 0,11629 | 350,7012 | 7,01827 | 502,8893 | 4184 | 0,2381 |
| RGS3 | 7 | 0,30779 | 2 | 7 | 1,944 | 1 | 0,11629 | 346,4607 | 6,96474 | 573,7182 | 4540 | 0,09524 |
| STARD8 | 7 | 0,30779 | 2 | 7 | 2,432 | 6 | 0,11629 | 343,8179 | 6,955 | 4623,276 | 45718 | 0,04762 |
| CDA | 7 | 0,30779 | 2 | 7 | 2,235 | 2 | 0,11629 | 345,2702 | 6,94933 | 2582,494 | 27364 | 0,09524 |
| TTYH1 | 27 | 0,56839 | 4 | 7 | 2,192 | 1 | 0,1329 | 320,3357 | 6,72223 | 243,7893 | 1360 | 0,28571 |
| ZFC3H1 | 16 | 0,38896 | 5 | 7 | 1,487 | 1 | 0,10337 | 314,5742 | 6,63544 | 1319,36 | 6098 | 0,28571 |
| SLTM | 127 | 0,57059 | 6 | 7 | 1,761 | 2 | 0,11629 | 318,0702 | 6,68654 | 463,5846 | 5482 | 0,57143 |
| GLIS3 | 8 | 0,30898 | 3 | 7 | 3,278 | 3 | 0,1329 | 361,9762 | 7,10424 | 2649,231 | 27986 | 0,09524 |
| NDUFA4L2 | 122 | 0,64826 | 5 | 7 | 1,691 | 2 | 0,11629 | 329,1036 | 6,80333 | 669,7476 | 6924 | 0,47619 |
| MT1X | 55 | 0,52304 | 6 | 7 | 3,288 | 3 | 0,1329 | 391,6595 | 7,32972 | 3186,372 | 34008 | 0,52381 |
| ELP4 | 7 | 0,30779 | 2 | 7 | 1,958 | 1 | 0,1329 | 343,9595 | 6,93797 | 874,4224 | 6262 | 0,04762 |
| GPER1 | 32 | 0,40246 | 7 | 7 | 10,667 | 1 | 0,1329 | 412,3095 | 7,46923 | 44,124 | 1008 | 0,52381 |
| LYPD1 | 7 | 0,30779 | 2 | 7 | 2,997 | 1 | 0,1329 | 347,4762 | 6,98339 | 1146,265 | 14114 | 0,09524 |
| GPR98 | 16 | 0,38896 | 5 | 7 | 1,439 | 1 | 0,1329 | 325,2262 | 6,74818 | 742,9395 | 5536 | 0,28571 |
| ZFP36L1 | 8 | 0,30898 | 3 | 7 | 3,153 | 1 | 0,11629 | 356,4345 | 7,0426 | 1003,244 | 10528 | 0,14286 |
| NFATC4 | 11 | 0,37893 | 4 | 7 | 7,679 | 3 | 0,1329 | 408,2095 | 7,44733 | 1526,725 | 14224 | 0,2381 |
| PERP | 14 | 0,38896 | 5 | 7 | 4,321 | 2 | 0,11629 | 355,2679 | 7,05152 | 1831,187 | 32448 | 0,28571 |
| CDC45 | 51 | 0,47549 | 6 | 7 | 4,641 | 1 | 0,15505 | 366,45 | 7,14723 | 812,6805 | 8808 | 0,47619 |
| MICA | 11 | 0,23775 | 6 | 7 | 3,973 | 1 | 0,1329 | 366,6857 | 7,13506 | 739,0542 | 8034 | 0,2381 |

| | | | | | | | | | | | | |
|---|---|---|---|---|---|---|---|---|---|---|---|---|
| RARRES2 | 145 | 0,61814 | 6 | 7 | 9,252 | 2 | 0,1329 | 389,0095 | 7,29809 | 2294,32 | 27706 | 0,61905 |
| SH3BP2 | 8 | 0,30898 | 3 | 7 | 5,818 | 2 | 0,11629 | 363,5845 | 7,10668 | 745,2288 | 10888 | 0,14286 |
| RAI14 | 7 | 0,30779 | 2 | 7 | 3,21 | 3 | 0,15505 | 344,9667 | 6,95825 | 9752,419 | 117294 | 0,04762 |
| MTTP | 19 | 0,38039 | 6 | 7 | 6,553 | 1 | 0,1329 | 402,4429 | 7,40596 | 1344,983 | 14496 | 0,38095 |
| GPR56 | 8 | 0,30898 | 3 | 7 | 4,009 | 1 | 0,1329 | 358,8429 | 7,07018 | 1132,264 | 11720 | 0,09524 |
| S1PR2 | 8 | 0,30898 | 3 | 7 | 8,126 | 2 | 0,1329 | 398,1595 | 7,36866 | 2128,095 | 19908 | 0,14286 |
| PLCD3 | 12 | 0,32413 | 5 | 7 | 1,421 | 2 | 0,11629 | 307,1869 | 6,56893 | 1147,797 | 8752 | 0,2381 |
| FOXD1 | 33 | 0,42794 | 6 | 7 | 4,661 | 1 | 0,1329 | 362,2429 | 7,10019 | 69,53312 | 1266 | 0,42857 |
| UNC93B1 | 26 | 0,56839 | 4 | 6 | 5,924 | 2 | 0,1329 | 371,8429 | 7,17075 | 795,1085 | 8834 | 0,4 |
| IGFBP5 | 9 | 0,46346 | 3 | 6 | 7,729 | 1 | 0,1329 | 394,8595 | 7,34838 | 187,3624 | 3064 | 0,26667 |
| IL1RL1 | 15 | 0,38896 | 5 | 6 | 8,139 | 1 | 0,1329 | 384,3429 | 7,26727 | 62,69004 | 1114 | 0,4 |
| MYOT | 10 | 0,37893 | 4 | 6 | 2,157 | 1 | 0,11629 | 346,0607 | 6,9623 | 399,6342 | 4476 | 0,26667 |
| MKNK2 | 6 | 0 | 1 | 6 | 2,213 | 3 | 0,11629 | 349,6774 | 7,00286 | 4688,56 | 52964 | 0 |
| GADD45G | 13 | 0,38896 | 5 | 6 | 7,579 | 3 | 0,1329 | 385,1762 | 7,27051 | 2475,928 | 26214 | 0,4 |
| CABLES1 | 7 | 0,30898 | 3 | 6 | 3,649 | 1 | 0,1329 | 357,0929 | 7,07342 | 368,207 | 3958 | 0,2 |
| EMP1 | 7 | 0,30898 | 3 | 6 | 4,513 | 2 | 0,1329 | 394,1595 | 7,34919 | 2691,261 | 19814 | 0,13333 |
| CIB2 | 11 | 0,32413 | 5 | 6 | 1,877 | 1 | 0,1329 | 328,4095 | 6,8082 | 283,0856 | 2772 | 0,33333 |
| GMPR | 6 | 0 | 1 | 6 | 1,746 | 1 | 0,11629 | 336,0536 | 6,87471 | 520,1411 | 3530 | 0 |
| TTPA | 6 | 0 | 1 | 6 | 1,252 | 1 | 0,11629 | 301,6369 | 6,51946 | 419,3572 | 2546 | 0 |
| CDKN2C | 6 | 0,30779 | 2 | 6 | 3,105 | 1 | 0,11629 | 355,6845 | 7,05801 | 604,6398 | 5684 | 0,06667 |
| ITPKC | 10 | 0,37893 | 4 | 6 | 3,236 | 3 | 0,11629 | 348,1512 | 6,97934 | 989,4191 | 11336 | 0,26667 |
| EPB41L5 | 6 | 0,30779 | 2 | 6 | 3,356 | 4 | 0,11629 | 350,6845 | 7,00529 | 5875,862 | 54772 | 0,06667 |
| PON3 | 15 | 0,38896 | 5 | 6 | 2,412 | 1 | 0,11629 | 345,7012 | 6,97204 | 189,687 | 1570 | 0,4 |
| ZIC5 | 14 | 0,47366 | 4 | 6 | 4,581 | 1 | 0,1329 | 368,6762 | 7,15777 | 925,2913 | 7834 | 0,33333 |
| ZFHX3 | 32 | 0,42794 | 6 | 6 | 7,157 | 1 | 0,11629 | 391,4012 | 7,31594 | 13,77772 | 238 | 0,6 |
| CTDSPL | 9 | 0,25931 | 5 | 6 | 2,359 | 1 | 0,11629 | 348,8845 | 7,00205 | 321,6964 | 2774 | 0,26667 |
| ERLIN2 | 6 | 0 | 1 | 6 | 2,663 | 1 | 0,11629 | 349,4179 | 7,00286 | 722,8462 | 4360 | 0 |
| MPZL2 | 6 | 0,30779 | 2 | 6 | 3,077 | 1 | 0,11629 | 346,1345 | 6,98015 | 382,1112 | 3430 | 0,06667 |
| CYSLTR2 | 144 | 0,61814 | 6 | 6 | 1,526 | 1 | 0,11629 | 311,9798 | 6,62814 | 8,18985 | 106 | 0,86667 |
| GPR116 | 9 | 0,46346 | 3 | 6 | 2,578 | 1 | 0,11629 | 325,7298 | 6,77089 | 381,5513 | 3360 | 0,2 |
| FAM167A | 6 | 0,30779 | 2 | 6 | 1,745 | 3 | 0,11629 | 323,0702 | 6,72709 | 1193,542 | 10418 | 0,06667 |
| TMPRSS3 | 15 | 0,38896 | 5 | 6 | 3,01 | 1 | 0,1329 | 349,7429 | 7,00935 | 388,0072 | 4070 | 0,4 |
| AQP5 | 7 | 0,30898 | 3 | 6 | 4,057 | 2 | 0,1329 | 396,9429 | 7,37028 | 1334,844 | 14318 | 0,13333 |
| IL17RD | 10 | 0,30898 | 3 | 6 | 6,713 | 1 | 0,1329 | 393,6262 | 7,33621 | 61,84979 | 1316 | 0,33333 |
| RGS20 | 26 | 0,56839 | 4 | 6 | 2,303 | 1 | 0,11629 | 335,6441 | 6,87471 | 152,7769 | 1698 | 0,4 |
| ARHGAP42 | 6 | 0,30779 | 2 | 6 | 1,464 | 4 | 0,11629 | 316,5107 | 6,67275 | 1811,174 | 16464 | 0,06667 |
| BATF2 | 9 | 0,46346 | 3 | 6 | 4,177 | 1 | 0,11629 | 351,0345 | 6,98501 | 413,3223 | 4538 | 0,26667 |
| CHST14 | 10 | 0,37893 | 4 | 6 | 2,781 | 1 | 0,1329 | 338,9595 | 6,90228 | 801,9735 | 9820 | 0,26667 |
| PPP1R3B | 10 | 0,37893 | 4 | 6 | 1,705 | 2 | 0,1329 | 315,6976 | 6,67437 | 1179,063 | 9812 | 0,26667 |
| SWAP70 | 7 | 0,30898 | 3 | 6 | 1,776 | 2 | 0,11629 | 315,4274 | 6,66869 | 2618,94 | 23982 | 0,2 |
| KIF1C | 6 | 0,30779 | 2 | 6 | 2,047 | 6 | 0,1329 | 352,2095 | 7,03449 | 4083,291 | 39688 | 0,06667 |
| FIBIN | 6 | 0 | 1 | 6 | 1,264 | 2 | 0,11629 | 320,2798 | 6,71898 | 1266,054 | 6072 | 0 |
| ARHGEF37 | 6 | 0,30779 | 2 | 6 | 1,595 | 2 | 0,11629 | 300,5298 | 6,49188 | 1182,365 | 10558 | 0,06667 |
| OLFML1 | 6 | 0 | 1 | 6 | 1,908 | 2 | 0,11629 | 329,3607 | 6,80901 | 630,9101 | 3904 | 0 |
| ARHGEF10 | 6 | 0 | 1 | 6 | 1,357 | 2 | 0,11629 | 311,5631 | 6,62733 | 1134,364 | 6830 | 0 |
| MAFF | 14 | 0,33284 | 6 | 6 | 6,395 | 1 | 0,11629 | 376,8012 | 7,21698 | 65,74234 | 1042 | 0,46667 |
| THSD4 | 9 | 0,46346 | 3 | 6 | 2,042 | 3 | 0,1329 | 332,5857 | 6,85605 | 1361,597 | 14426 | 0,2 |
| MYOM1 | 9 | 0,46346 | 3 | 6 | 3,304 | 3 | 0,1329 | 354,0262 | 7,03692 | 3290,908 | 33772 | 0,2 |
| GOLGA4 | 6 | 0 | 1 | 6 | 1,522 | 2 | 0,1329 | 319,4262 | 6,6987 | 2022,85 | 23152 | 0 |
| MYOZ1 | 16 | 0,33284 | 6 | 6 | 1,79 | 1 | 0,11629 | 308,0488 | 6,58596 | 153,9839 | 1158 | 0,46667 |
| EMP2 | 10 | 0,37893 | 4 | 6 | 4,771 | 1 | 0,1329 | 377,6095 | 7,23321 | 393,0326 | 5316 | 0,26667 |
| RXRG | 9 | 0,25931 | 5 | 6 | 3,995 | 1 | 0,15505 | 365,9333 | 7,14885 | 542,5529 | 4376 | 0,26667 |
| BAZ1A | 31 | 0,51861 | 5 | 6 | 2,91 | 2 | 0,11629 | 337,5345 | 6,89012 | 1596,836 | 14862 | 0,53333 |
| ANKHD1 | 6 | 0,30779 | 2 | 6 | 3,851 | 1 | 0,11629 | 361,2179 | 7,09775 | 382,7552 | 4320 | 0,06667 |
| ETNK2 | 6 | 0,30779 | 2 | 6 | 1,562 | 4 | 0,11629 | 316,7798 | 6,68654 | 3068,223 | 36074 | 0,06667 |
| SOX13 | 10 | 0,37893 | 4 | 6 | 3,335 | 1 | 0,1329 | 351,3595 | 7,01665 | 501,3767 | 6548 | 0,26667 |
| GABRE | 27 | 0,45378 | 5 | 6 | 1,982 | 1 | 0,1329 | 321,8191 | 6,74818 | 242,9514 | 1630 | 0,46667 |
| ZNF217 | 7 | 0,30898 | 3 | 6 | 5,297 | 1 | 0,11629 | 376,6679 | 7,21861 | 1362,995 | 14032 | 0,13333 |
| KCNE1L | 7 | 0,30898 | 3 | 6 | 1,599 | 2 | 0,11629 | 319,0631 | 6,7133 | 2905,558 | 23404 | 0,2 |
| SRPX2 | 6 | 0 | 1 | 6 | 2,282 | 1 | 0,11629 | 345,2845 | 6,96393 | 505,4478 | 4166 | 0 |
| TET1 | 6 | 0,30779 | 2 | 6 | 4,041 | 2 | 0,1329 | 368,8762 | 7,1667 | 716,9249 | 6614 | 0,06667 |
| ALDH4A1 | 10 | 0,23775 | 6 | 6 | 2,206 | 1 | 0,1329 | 333,7524 | 6,84875 | 163,0773 | 1570 | 0,33333 |
| DDAH2 | 7 | 0,30898 | 3 | 6 | 4,317 | 1 | 0,11629 | 371,4179 | 7,18616 | 271,4463 | 2328 | 0,13333 |
| SLC39A12 | 26 | 0,56839 | 4 | 6 | 2,459 | 1 | 0,15505 | 358,25 | 7,08883 | 1188,513 | 10318 | 0,4 |
| LRP4 | 9 | 0,25931 | 5 | 6 | 5,747 | 2 | 0,15505 | 386,8667 | 7,29404 | 188,0102 | 2710 | 0,26667 |
| RREB1 | 6 | 0,30779 | 2 | 6 | 1,398 | 2 | 0,11629 | 321,9702 | 6,74169 | 631,9182 | 3736 | 0,13333 |
| RYR3 | 6 | 0,30779 | 2 | 6 | 1,516 | 3 | 0,11629 | 321,6774 | 6,74169 | 5104,579 | 45474 | 0,06667 |
| ENSP0000037774 | 14 | 0,47366 | 4 | 6 | 1,97 | 2 | 0,11629 | 340,7012 | 6,91851 | 2460,906 | 25594 | 0,33333 |
| TSPAN4 | 7 | 0,30898 | 3 | 6 | 6,321 | 1 | 0,1329 | 377,5429 | 7,22509 | 189,9962 | 2456 | 0,13333 |
| ODF3B | 6 | 0,30779 | 2 | 6 | 1,679 | 1 | 0,11629 | 329,1607 | 6,80414 | 591,2488 | 3530 | 0,06667 |
| MATN3 | 9 | 0,46346 | 3 | 6 | 5,129 | 2 | 0,1329 | 390,1762 | 7,31837 | 1189,796 | 11502 | 0,2 |
| RASSF8 | 9 | 0,46346 | 3 | 6 | 3,251 | 2 | 0,15505 | 354,1 | 7,04341 | 2719,382 | 25898 | 0,2 |
| HIPK2 | 6 | 0,30779 | 2 | 6 | 5,348 | 1 | 0,1329 | 395,7762 | 7,35 | 646,8137 | 7346 | 0,13333 |
| MT1G | 54 | 0,52304 | 6 | 6 | 1,512 | 1 | 0,11629 | 314,7036 | 6,66383 | 796,9283 | 20756 | 0,73333 |
| NANOS1 | 6 | 0,30779 | 2 | 6 | 1,242 | 1 | 0,1329 | 294,7881 | 6,42699 | 693,8448 | 2716 | 0,06667 |
| SELENBP1 | 6 | 0,30779 | 2 | 6 | 2,744 | 1 | 0,11629 | 347,6845 | 7,00043 | 1555,189 | 17342 | 0,06667 |
| HYAL2 | 26 | 0,56839 | 4 | 6 | 8,956 | 2 | 0,1329 | 397,1429 | 7,35973 | 1851,728 | 21404 | 0,4 |
| ZFHX4 | 15 | 0,38896 | 5 | 6 | 3,219 | 2 | 0,1329 | 350,7762 | 7,01178 | 828,5853 | 11138 | 0,4 |
| ZIC4 | 7 | 0,30898 | 3 | 6 | 1,933 | 2 | 0,1329 | 348,6262 | 6,9988 | 809,4719 | 5526 | 0,2 |

| | | | | | | | | | | | | |
|---|---|---|---|---|---|---|---|---|---|---|---|---|
| DTNA | 7 | 0,30898 | 3 | 6 | 2,298 | 3 | 0,11629 | 335,2179 | 6,87714 | 1634,602 | 14536 | 0,13333 |
| TMPRSS4 | 15 | 0,38896 | 5 | 6 | 2,844 | 1 | 0,1329 | 349,4762 | 7,00691 | 352,499 | 4420 | 0,4 |
| COPZ2 | 6 | 0,30779 | 2 | 6 | 3,112 | 1 | 0,15505 | 344,3667 | 6,94608 | 403,125 | 3322 | 0,06667 |
| TUBB1 | 5 | 0 | 1 | 5 | 2,416 | 1 | 0,11629 | 350,7512 | 7,02557 | 1283,665 | 12062 | 0 |
| NKG7 | 6 | 0,30898 | 3 | 5 | 4,733 | 1 | 0,1329 | 352,6024 | 7,01259 | 20,32161 | 326 | 0,3 |
| AEBP1 | 5 | 0,30779 | 2 | 5 | 3,452 | 1 | 0,1329 | 366,6262 | 7,13506 | 547,9898 | 5932 | 0,1 |
| SLC19A3 | 5 | 0 | 1 | 5 | 1,559 | 1 | 0,10337 | 295,0028 | 6,42699 | 408,7878 | 1890 | 0 |
| BNIP2 | 5 | 0 | 1 | 5 | 1,704 | 1 | 0,11629 | 339,4845 | 6,92581 | 742,7713 | 6246 | 0 |
| ACKR3 | 18 | 0,45378 | 5 | 5 | 8,473 | 1 | 0,1329 | 408,4929 | 7,44409 | 4,12709 | 340 | 0,7 |
| FREM2 | 8 | 0,46346 | 3 | 5 | 7,37 | 1 | 0,1329 | 398,3595 | 7,37433 | 500,8406 | 5998 | 0,3 |
| ADAMTSL3 | 8 | 0,46346 | 3 | 5 | 1,794 | 1 | 0,11629 | 318,0536 | 6,71249 | 96,58207 | 978 | 0,4 |
| SLCO2B1 | 8 | 0,46346 | 3 | 5 | 2,292 | 2 | 0,11629 | 349,5512 | 7,01178 | 1463,932 | 21344 | 0,3 |
| GRIN2C | 9 | 0,37893 | 4 | 5 | 3,555 | 1 | 0,15505 | 363,05 | 7,12858 | 141,7533 | 1556 | 0,4 |
| BCL6B | 6 | 0,30898 | 3 | 5 | 4,303 | 2 | 0,1329 | 378,4262 | 7,23564 | 590,675 | 6960 | 0,2 |
| DOCK6 | 5 | 0,30779 | 2 | 5 | 1,921 | 1 | 0,11629 | 326,2845 | 6,79441 | 734,4897 | 6234 | 0,1 |
| PGAM2 | 5 | 0,30779 | 2 | 5 | 1,469 | 1 | 0,11629 | 317,2441 | 6,68573 | 1210,66 | 6848 | 0,1 |
| MLC1 | 9 | 0,37893 | 4 | 5 | 5,734 | 1 | 0,1329 | 366,6595 | 7,14966 | 1222,892 | 14264 | 0,4 |
| TNS3 | 8 | 0,46346 | 3 | 5 | 6,83 | 1 | 0,1329 | 398,9929 | 7,38569 | 353,3029 | 5254 | 0,3 |
| MID1 | 5 | 0 | 1 | 5 | 2,187 | 3 | 0,11629 | 349,2179 | 7,01097 | 4085,353 | 37286 | 0 |
| BBX | 5 | 0 | 1 | 5 | 1,398 | 2 | 0,11629 | 325,7202 | 6,77657 | 2776,559 | 27378 | 0 |
| TRIM41 | 6 | 0,30898 | 3 | 5 | 2,156 | 1 | 0,11629 | 331,6274 | 6,84632 | 245,7824 | 1980 | 0,2 |
| ASB4 | 5 | 0,30779 | 2 | 5 | 3,475 | 1 | 0,11629 | 357,6774 | 7,06207 | 272,9982 | 3302 | 0,1 |
| TGIF1 | 6 | 0,30898 | 3 | 5 | 3,69 | 1 | 0,1329 | 365,0595 | 7,13588 | 114,906 | 1500 | 0,3 |
| EYA2 | 10 | 0,32413 | 5 | 5 | 3,586 | 1 | 0,1329 | 350,7762 | 7,01908 | 47,42724 | 656 | 0,5 |
| MT1F | 30 | 0,51861 | 5 | 5 | 1,669 | 1 | 0,11629 | 312,9536 | 6,65085 | 671,9632 | 16862 | 0,8 |
| CDC42EP4 | 5 | 0 | 1 | 5 | 2,357 | 2 | 0,11629 | 349,9179 | 7,00935 | 1668,86 | 15108 | 0 |
| SLC6A13 | 13 | 0,47366 | 4 | 5 | 1,863 | 1 | 0,11629 | 328,9179 | 6,8082 | 92,24784 | 544 | 0,5 |
| RBPMS | 5 | 0 | 1 | 5 | 1,669 | 3 | 0,11629 | 333,2845 | 6,85443 | 555,047 | 4338 | 0 |
| WNK1 | 5 | 0 | 1 | 5 | 1,326 | 1 | 0,1329 | 329,0024 | 6,82442 | 491,9166 | 3194 | 0 |
| TP53INP1 | 5 | 0 | 1 | 5 | 2,806 | 2 | 0,11629 | 341,6845 | 6,93878 | 2509,836 | 24996 | 0 |
| ANP32B | 5 | 0 | 1 | 5 | 1,418 | 1 | 0,1329 | 326,3429 | 6,80414 | 565,7835 | 3508 | 0 |
| SLC2A10 | 8 | 0,46346 | 3 | 5 | 4,373 | 2 | 0,11629 | 343,5774 | 6,94933 | 1116,411 | 16422 | 0,3 |
| CLCA4 | 5 | 0 | 1 | 5 | 1,493 | 1 | 0,1329 | 307,1333 | 6,58515 | 662,616 | 3834 | 0 |
| SARDH | 7 | 0,2842 | 4 | 5 | 1,798 | 1 | 0,11629 | 315,1607 | 6,67356 | 181,0703 | 1396 | 0,3 |
| ZCCHC24 | 5 | 0,30779 | 2 | 5 | 1,436 | 1 | 0,1329 | 300,0548 | 6,50729 | 452,3653 | 2196 | 0,1 |
| CHST3 | 9 | 0,37893 | 4 | 5 | 2,846 | 2 | 0,1329 | 339,9429 | 6,9177 | 213,9798 | 2232 | 0,4 |
| PHKA1 | 5 | 0,30779 | 2 | 5 | 1,179 | 2 | 0,11629 | 280,2417 | 6,25504 | 974,9371 | 6648 | 0,1 |
| ECE1 | 5 | 0,30779 | 2 | 5 | 5,083 | 1 | 0,1329 | 367,5095 | 7,15615 | 1665,921 | 21576 | 0,1 |
| PLA2G5 | 5 | 0,30779 | 2 | 5 | 1,582 | 1 | 0,11629 | 326,2702 | 6,79117 | 859,8707 | 4834 | 0,2 |
| RASSF2 | 5 | 0 | 1 | 5 | 1,569 | 4 | 0,1329 | 297,1643 | 6,46755 | 6697,284 | 62620 | 0 |
| ASPH | 5 | 0 | 1 | 5 | 4,119 | 4 | 0,11629 | 357,8179 | 7,07991 | 1176,989 | 10832 | 0 |
| RHOBTB3 | 5 | 0 | 1 | 5 | 1,808 | 1 | 0,11629 | 321,2369 | 6,74412 | 1089,353 | 12806 | 0 |
| AKR1C3 | 5 | 0,30779 | 2 | 5 | 1,287 | 2 | 0,11629 | 320,8131 | 6,71979 | 1900,56 | 13470 | 0,1 |
| ARSD | 5 | 0,30779 | 2 | 5 | 2,041 | 2 | 0,11629 | 329,1869 | 6,81469 | 3809,692 | 40862 | 0,1 |
| NMB | 7 | 0,2842 | 4 | 5 | 4,871 | 1 | 0,11629 | 364,4512 | 7,12452 | 160,0168 | 1690 | 0,3 |
| AK4 | 5 | 0 | 1 | 5 | 1,498 | 1 | 0,11629 | 310,2155 | 6,61111 | 950,707 | 7210 | 0 |
| BCAS1 | 5 | 0,30779 | 2 | 5 | 2,572 | 1 | 0,1329 | 357,2262 | 7,08316 | 424,3022 | 3238 | 0,1 |
| TSPO | 8 | 0,46346 | 3 | 5 | 5,689 | 2 | 0,1329 | 396,5429 | 7,3646 | 2132,086 | 16950 | 0,3 |
| CTDSP2 | 6 | 0,30898 | 3 | 5 | 1,852 | 1 | 0,11629 | 328,8202 | 6,82037 | 138,5423 | 980 | 0,2 |
| GXYLT1 | 8 | 0,46346 | 3 | 5 | 4,63 | 1 | 0,1329 | 369,9429 | 7,1667 | 294,4671 | 2586 | 0,4 |
| HSDL2 | 9 | 0,37893 | 4 | 5 | 1,957 | 1 | 0,10337 | 305,6409 | 6,5592 | 94,8785 | 610 | 0,4 |
| DAAM2 | 5 | 0,30779 | 2 | 5 | 1,418 | 2 | 0,1329 | 342,8595 | 6,95663 | 639,5195 | 4432 | 0,1 |
| LRRC32 | 6 | 0,30898 | 3 | 5 | 6,738 | 1 | 0,1329 | 366,9691 | 7,14399 | 37,45312 | 726 | 0,2 |
| HEPN1 | 30 | 0,51861 | 5 | 5 | 2,286 | 1 | 0,15505 | 352,4833 | 7,04017 | 67,2018 | 402 | 0,8 |
| CTNNA3 | 48 | 0,58344 | 5 | 5 | 4,227 | 1 | 0,1329 | 366,5262 | 7,14236 | 12,09758 | 242 | 0,9 |
| NECAP2 | 5 | 0 | 1 | 5 | 2,37 | 1 | 0,1329 | 343,0762 | 6,95663 | 458,2312 | 3912 | 0 |
| NOSTRIN | 8 | 0,46346 | 3 | 5 | 5,208 | 1 | 0,11629 | 374,5012 | 7,21617 | 709,5128 | 7310 | 0,4 |
| SLC6A12 | 13 | 0,47366 | 4 | 5 | 1,781 | 1 | 0,11629 | 336,9679 | 6,89012 | 56,79285 | 696 | 0,5 |
| ACSF2 | 6 | 0,30898 | 3 | 5 | 2,517 | 1 | 0,11629 | 337,8607 | 6,89823 | 304,6913 | 3230 | 0,2 |
| SLC13A5 | 5 | 0 | 1 | 5 | 1,353 | 1 | 0,1329 | 298,5571 | 6,48863 | 693,7488 | 2858 | 0 |
| ITPKB | 10 | 0,32413 | 5 | 5 | 1,792 | 1 | 0,11629 | 312,6202 | 6,63868 | 166,2861 | 1540 | 0,5 |
| HIST2H2BF | 8 | 0,25931 | 5 | 5 | 1,972 | 1 | 0,11629 | 341,2179 | 6,93229 | 224,2902 | 1720 | 0,4 |
| TMEM204 | 5 | 0,30779 | 2 | 5 | 1,828 | 1 | 0,11629 | 339,3845 | 6,92499 | 920,168 | 8242 | 0,1 |
| RNF43 | 9 | 0,37893 | 4 | 5 | 4,04 | 1 | 0,15505 | 358,2333 | 7,08072 | 98,08566 | 1660 | 0,4 |
| KANK1 | 5 | 0 | 1 | 5 | 2,304 | 2 | 0,15505 | 355,9667 | 7,0572 | 1491,535 | 19016 | 0 |
| MT1A | 9 | 0,37893 | 4 | 5 | 1,571 | 1 | 0,11629 | 333,4869 | 6,85524 | 147,8429 | 1636 | 0,4 |
| ECSCR | 5 | 0,30779 | 2 | 5 | 3,773 | 3 | 0,1329 | 350,3762 | 7,00367 | 2529,12 | 29674 | 0,1 |
| ARHGEF5 | 4 | 0 | 1 | 4 | 1,545 | 1 | 0,11629 | 318,0869 | 6,69708 | 931,7716 | 9802 | 0 |
| ARRDC2 | 4 | 0 | 1 | 4 | 1,718 | 2 | 0,1329 | 320,7262 | 6,7425 | 1401,658 | 12708 | 0 |
| PPP1R3C | 8 | 0,37893 | 4 | 4 | 1,471 | 2 | 0,1329 | 292,0691 | 6,40834 | 198,7163 | 2180 | 0,66667 |
| WNK4 | 5 | 0,30898 | 3 | 4 | 1,776 | 1 | 0,1329 | 325,4857 | 6,79684 | 198,0137 | 1444 | 0,33333 |
| NSRP1 | 4 | 0,30779 | 2 | 4 | 1,165 | 1 | 0,10337 | 282,7028 | 6,2737 | 184,3703 | 886 | 0,16667 |
| HIGD1B | 4 | 0,30779 | 2 | 4 | 1,473 | 1 | 0,11629 | 307,7869 | 6,60381 | 215,0818 | 1916 | 0,33333 |
| RIN2 | 4 | 0,30779 | 2 | 4 | 1,341 | 1 | 0,11629 | 300,1131 | 6,48945 | 211,1344 | 1184 | 0,16667 |
| RARRES3 | 4 | 0 | 1 | 4 | 2,019 | 1 | 0,1329 | 333,1691 | 6,85524 | 271,2552 | 1808 | 0 |
| SPX | 4 | 0 | 1 | 4 | 1,166 | 1 | 0,10337 | 294,3623 | 6,40266 | 649,8983 | 4290 | 0 |
| SDS | 4 | 0,30779 | 2 | 4 | 1,445 | 1 | 0,11629 | 300,9202 | 6,51378 | 126,8615 | 748 | 0,33333 |
| DYNC1LI2 | 4 | 0,30779 | 2 | 4 | 1,15 | 3 | 0,11629 | 307,8298 | 6,60056 | 3607,722 | 30522 | 0,16667 |
| RBMS2 | 24 | 0,56839 | 4 | 4 | 2,616 | 1 | 0,11629 | 317,6702 | 6,69627 | 0 | 0 | 1 |

| | | | | | | | | | | | | |
|---|---|---|---|---|---|---|---|---|---|---|---|---|
| CELSR1 | 5 | 0,30898 | 3 | 4 | 1,927 | 1 | 0,1329 | 326,9286 | 6,79847 | 258,5506 | 2078 | 0,33333 |
| CAPG | 7 | 0,46346 | 3 | 4 | 4,5 | 1 | 0,1329 | 359,6429 | 7,10262 | 61,46883 | 614 | 0,5 |
| ARHGAP31 | 4 | 0 | 1 | 4 | 3,438 | 1 | 0,1329 | 346,8262 | 6,98339 | 295,2984 | 2476 | 0 |
| EVC | 7 | 0,46346 | 3 | 4 | 2,399 | 1 | 0,1329 | 340,0357 | 6,92337 | 13,12316 | 154 | 0,5 |
| IMPA2 | 4 | 0,30779 | 2 | 4 | 1,364 | 1 | 0,11629 | 280,2155 | 6,25991 | 162,8188 | 760 | 0,16667 |
| BCAM | 4 | 0 | 1 | 4 | 1,945 | 1 | 0,1329 | 352,0095 | 7,04909 | 179,4846 | 1576 | 0 |
| TINAGL1 | 4 | 0,30779 | 2 | 4 | 3,242 | 1 | 0,1329 | 347,7095 | 6,98501 | 126,6958 | 1130 | 0,16667 |
| CGNL1 | 4 | 0,30779 | 2 | 4 | 2,513 | 2 | 0,11629 | 343,3679 | 6,95906 | 1252,44 | 12854 | 0,16667 |
| GRAP | 4 | 0,30779 | 2 | 4 | 1,925 | 1 | 0,11629 | 320,7702 | 6,7279 | 320,9827 | 4316 | 0,16667 |
| PKN3 | 4 | 0,30779 | 2 | 4 | 2,002 | 1 | 0,11629 | 331,8179 | 6,85605 | 206,8054 | 1684 | 0,16667 |
| ATOH8 | 4 | 0 | 1 | 4 | 1,766 | 1 | 0,1329 | 328,8524 | 6,81793 | 283,8703 | 2186 | 0 |
| HSPB7 | 8 | 0,37893 | 4 | 4 | 2,466 | 1 | 0,11629 | 337,9869 | 6,9104 | 24,37457 | 254 | 0,66667 |
| TNFRSF10D | 5 | 0,30898 | 3 | 4 | 3,624 | 1 | 0,1329 | 344,8095 | 6,95825 | 86,41185 | 858 | 0,33333 |
| LGI4 | 4 | 0 | 1 | 4 | 3,052 | 2 | 0,11629 | 363,9679 | 7,12939 | 1689,089 | 15246 | 0 |
| METRNL | 4 | 0 | 1 | 4 | 1,625 | 3 | 0,11629 | 320,2202 | 6,73277 | 2941,886 | 31022 | 0 |
| PHF21B | 4 | 0,30779 | 2 | 4 | 1,266 | 1 | 0,10337 | 312,1218 | 6,63625 | 351,7954 | 1890 | 0,16667 |
| LIMS2 | 4 | 0,30779 | 2 | 4 | 4,556 | 2 | 0,1329 | 360,9429 | 7,10262 | 2397,752 | 29656 | 0,16667 |
| ZNF703 | 5 | 0,30898 | 3 | 4 | 4,549 | 1 | 0,11629 | 367,1845 | 7,15047 | 107,7885 | 1366 | 0,33333 |
| ARSI | 4 | 0 | 1 | 4 | 1,125 | 2 | 0,11629 | 280,0226 | 6,26559 | 2496,693 | 23968 | 0 |
| HAP1 | 4 | 0,30779 | 2 | 4 | 2,363 | 1 | 0,1329 | 330,2286 | 6,83091 | 266,5509 | 2036 | 0,16667 |
| HIST1H1C | 8 | 0,37893 | 4 | 4 | 2,625 | 2 | 0,11629 | 338,7512 | 6,91851 | 81,48217 | 1714 | 0,66667 |
| CCDC15 | 4 | 0 | 1 | 4 | 2,028 | 2 | 0,11629 | 327,0941 | 6,80009 | 772,0604 | 7700 | 0 |
| PAPLN | 4 | 0 | 1 | 4 | 1,905 | 1 | 0,11629 | 320,3631 | 6,71817 | 377,1504 | 3464 | 0 |
| PLEKHB1 | 4 | 0 | 1 | 4 | 1,337 | 2 | 0,11629 | 296,9417 | 6,46268 | 2564,445 | 19382 | 0 |
| APCDD1 | 4 | 0,30779 | 2 | 4 | 1,839 | 6 | 0,1329 | 329,0024 | 6,82604 | 3328,037 | 36276 | 0,16667 |
| TRIM38 | 5 | 0,30898 | 3 | 4 | 1,929 | 1 | 0,11629 | 324,2107 | 6,76927 | 93,79381 | 700 | 0,33333 |
| RFX4 | 4 | 0,30779 | 2 | 4 | 1,734 | 1 | 0,11629 | 313,206 | 6,65085 | 326,1808 | 1558 | 0,16667 |
| PARD3B | 4 | 0,30779 | 2 | 4 | 1,143 | 2 | 0,10337 | 270,9171 | 6,13095 | 2547,75 | 22396 | 0,33333 |
| RGR | 7 | 0,46346 | 3 | 4 | 1,593 | 1 | 0,11629 | 320,1702 | 6,73682 | 90,28234 | 1110 | 0,5 |
| CSRP1 | 4 | 0 | 1 | 4 | 2,066 | 1 | 0,1329 | 340,1595 | 6,92094 | 1137,093 | 13098 | 0 |
| NEK7 | 4 | 0,30779 | 2 | 4 | 1,683 | 1 | 0,1329 | 330,6929 | 6,84713 | 122,6363 | 1204 | 0,16667 |
| S100A3 | 4 | 0 | 1 | 4 | 1,525 | 3 | 0,11629 | 315,7845 | 6,70519 | 2538,999 | 21584 | 0 |
| ADAMTSL4 | 7 | 0,46346 | 3 | 4 | 1,571 | 1 | 0,11629 | 316,4845 | 6,69708 | 216,3443 | 1288 | 0,5 |
| C10orf11 | 4 | 0 | 1 | 4 | 1,128 | 2 | 0,11629 | 281,4929 | 6,26234 | 466,3876 | 3136 | 0 |
| MFSD2A | 4 | 0 | 1 | 4 | 2,774 | 1 | 0,11629 | 341,3441 | 6,93797 | 712,1415 | 8262 | 0 |
| NEK6 | 4 | 0,30779 | 2 | 4 | 1,73 | 1 | 0,11629 | 317,2631 | 6,70438 | 207,2501 | 1628 | 0,16667 |
| PHEX | 7 | 0,46346 | 3 | 4 | 2,186 | 1 | 0,1329 | 329,7619 | 6,83334 | 20,20536 | 138 | 0,5 |
| BCL7C | 4 | 0,30779 | 2 | 4 | 2,087 | 1 | 0,1329 | 350,7929 | 7,028 | 54,43699 | 522 | 0,16667 |
| CTNS | 4 | 0 | 1 | 4 | 3,066 | 1 | 0,1329 | 347,6929 | 6,99799 | 889,0255 | 11188 | 0 |
| SH2D2A | 4 | 0,30779 | 2 | 4 | 4,698 | 1 | 0,1329 | 376,4429 | 7,23077 | 85,69738 | 1232 | 0,16667 |
| EML3 | 4 | 0 | 1 | 4 | 1,529 | 2 | 0,11629 | 317,8107 | 6,70519 | 239,9922 | 2156 | 0 |
| SLC16A9 | 4 | 0,30779 | 2 | 4 | 1,62 | 1 | 0,11629 | 311,8036 | 6,63382 | 145,9316 | 1058 | 0,16667 |
| WDR60 | 4 | 0 | 1 | 4 | 1,153 | 2 | 0,10337 | 250,0433 | 5,79922 | 2441,67 | 19798 | 0 |
| NCKAP5 | 4 | 0 | 1 | 4 | 1,853 | 1 | 0,11629 | 323,6512 | 6,77251 | 215,0682 | 1276 | 0 |
| CCDC103 | 6 | 0,2842 | 4 | 4 | 1,35 | 1 | 0,11629 | 311,1798 | 6,63625 | 161,7382 | 1432 | 0,5 |
| FAM187A | 6 | 0,2842 | 4 | 4 | 1,517 | 1 | 0,11629 | 311,1798 | 6,63625 | 161,7382 | 1432 | 0,5 |
| MAPKAPK3 | 5 | 0,30898 | 3 | 4 | 3,466 | 1 | 0,11629 | 345,3012 | 6,9769 | 98,93409 | 1762 | 0,33333 |
| RELL1 | 4 | 0 | 1 | 4 | 1,642 | 2 | 0,11629 | 315,4131 | 6,65815 | 728,1003 | 5766 | 0 |
| MYBPC1 | 5 | 0,30898 | 3 | 4 | 2,467 | 2 | 0,11629 | 344,1679 | 6,96798 | 592,5442 | 7682 | 0,33333 |
| GLRA1 | 24 | 0,56839 | 4 | 4 | 1,166 | 1 | 0,1329 | 309,3191 | 6,6249 | 0 | 0 | 1 |
| MRVI1 | 4 | 0,30779 | 2 | 4 | 1,956 | 1 | 0,1329 | 322,8095 | 6,75791 | 449,8453 | 3732 | 0,16667 |
| RAB34 | 4 | 0 | 1 | 4 | 2,814 | 1 | 0,1329 | 344,9762 | 6,96798 | 399,6807 | 2636 | 0 |
| GPR143 | 4 | 0 | 1 | 4 | 1,564 | 2 | 0,11629 | 324,4536 | 6,77576 | 2903,734 | 25384 | 0 |
| FAM107A | 4 | 0 | 1 | 4 | 2,774 | 2 | 0,1329 | 354,9762 | 7,0645 | 1474,407 | 15556 | 0 |
| SORCS2 | 4 | 0 | 1 | 4 | 1,659 | 1 | 0,11629 | 320,9941 | 6,74575 | 902,4662 | 10212 | 0 |
| GLIPR1L2 | 4 | 0 | 1 | 4 | 1,125 | 3 | 0,10337 | 251,5111 | 5,84302 | 4600,857 | 35512 | 0 |
| PTPN21 | 4 | 0 | 1 | 4 | 1,949 | 2 | 0,1329 | 326,8691 | 6,80333 | 2354,665 | 28652 | 0 |
| RAB31 | 5 | 0,30898 | 3 | 4 | 5,708 | 1 | 0,1329 | 392,4929 | 7,34189 | 297,1535 | 3034 | 0,33333 |
| PITPNC1 | 4 | 0 | 1 | 4 | 1,762 | 1 | 0,11629 | 328,4702 | 6,82118 | 413,6108 | 3264 | 0 |
| KCNJ16 | 7 | 0,46346 | 3 | 4 | 1,401 | 1 | 0,11629 | 305,8131 | 6,57055 | 130,9906 | 928 | 0,5 |
| FFAR2 | 8 | 0,37893 | 4 | 4 | 7,565 | 1 | 0,1329 | 389,4429 | 7,31269 | 5,44611 | 150 | 0,66667 |
| TRIL | 7 | 0,46346 | 3 | 4 | 6,402 | 1 | 0,1329 | 401,0262 | 7,39948 | 50,09256 | 854 | 0,5 |
| FXYD1 | 4 | 0,30779 | 2 | 4 | 1,422 | 1 | 0,11629 | 316,2655 | 6,68654 | 190,8083 | 1128 | 0,16667 |
| MDFIC | 4 | 0 | 1 | 4 | 1,326 | 1 | 0,1329 | 324,8024 | 6,78143 | 276,7509 | 2694 | 0 |
| SLC7A2 | 3 | 0,30779 | 2 | 3 | 3,142 | 1 | 0,11629 | 346,8512 | 6,99313 | 9,21329 | 258 | 0,33333 |
| CCDC80 | 3 | 0,30779 | 2 | 3 | 4,391 | 1 | 0,1329 | 382,2429 | 7,27457 | 1172,755 | 8364 | 0,33333 |
| PIK3IP1 | 3 | 0,30779 | 2 | 3 | 1,297 | 1 | 0,1329 | 270,7095 | 6,13338 | 108,4911 | 480 | 0,33333 |
| NXT2 | 3 | 0,30779 | 2 | 3 | 1,116 | 1 | 0,10337 | 274,8956 | 6,18367 | 324,8065 | 1798 | 0,33333 |
| RASL12 | 3 | 0 | 1 | 3 | 1,294 | 2 | 0,11629 | 303,4631 | 6,5592 | 2375,261 | 20630 | 0 |
| EHD4 | 3 | 0 | 1 | 3 | 2,428 | 1 | 0,11629 | 335,7679 | 6,8885 | 53,61796 | 698 | 0 |
| YBX3 | 3 | 0,30779 | 2 | 3 | 2,81 | 1 | 0,11629 | 361,7179 | 7,12533 | 42,89869 | 526 | 0,33333 |
| CLEC2B | 3 | 0 | 1 | 3 | 2,751 | 1 | 0,1329 | 330,1691 | 6,82685 | 32,02162 | 598 | 0 |
| FLVCR2 | 3 | 0 | 1 | 3 | 1,399 | 1 | 0,11629 | 314,1774 | 6,68167 | 35,25962 | 254 | 0 |
| PAIP2B | 3 | 0,30779 | 2 | 3 | 1,296 | 1 | 0,10337 | 283,9147 | 6,30452 | 394,9443 | 2000 | 0,33333 |
| RHPN2 | 3 | 0 | 1 | 3 | 2,413 | 2 | 0,11629 | 330,6774 | 6,85038 | 1439,793 | 16256 | 0 |
| HILPDA | 3 | 0 | 1 | 3 | 1,246 | 1 | 0,1329 | 298,5429 | 6,50486 | 555,3282 | 3516 | 0 |
| GDF11 | 3 | 0,30779 | 2 | 3 | 2,045 | 1 | 0,11629 | 321,0036 | 6,73439 | 11,9662 | 138 | 0,33333 |
| CCDC102A | 3 | 0 | 1 | 3 | 1,202 | 1 | 0,10337 | 274,6718 | 6,1934 | 55,76257 | 386 | 0 |
| NPL | 3 | 0 | 1 | 3 | 1,104 | 1 | 0,11629 | 302,8726 | 6,54054 | 186,9647 | 1042 | 0 |

| | | | | | | | | | | | | |
|---|---|---|---|---|---|---|---|---|---|---|---|---|
| LRRC8A | 3 | 0,30779 | 2 | 3 | 3,217 | 1 | 0,11629 | 362,6679 | 7,13344 | 24,89008 | 276 | 0,33333 |
| SQRDL | 3 | 0 | 1 | 3 | 1,297 | 1 | 0,10337 | 289,1837 | 6,36292 | 130,2964 | 644 | 0 |
| N4BP2 | 3 | 0 | 1 | 3 | 1,9 | 1 | 0,11629 | 323,3774 | 6,7644 | 113,2248 | 490 | 0 |
| CASC3 | 6 | 0,46346 | 3 | 3 | 1,221 | 1 | 0,11629 | 281,625 | 6,27856 | 0 | 0 | 1 |
| SLC2A4RG | 3 | 0 | 1 | 3 | 2,128 | 1 | 0,1329 | 337,5857 | 6,92013 | 337,9892 | 3520 | 0 |
| CSRNP1 | 3 | 0 | 1 | 3 | 1,782 | 1 | 0,11629 | 319,5536 | 6,71736 | 949,212 | 9116 | 0 |
| MYH15 | 3 | 0 | 1 | 3 | 1,91 | 1 | 0,1329 | 331,9095 | 6,8593 | 69,10276 | 502 | 0 |
| PLA1A | 3 | 0 | 1 | 3 | 3,152 | 1 | 0,11629 | 340,5845 | 6,94446 | 19,78482 | 280 | 0 |
| DENND2A | 3 | 0 | 1 | 3 | 1,117 | 1 | 0,1329 | 268,8024 | 6,10499 | 115,3304 | 534 | 0 |
| ZNF462 | 3 | 0,30779 | 2 | 3 | 2,817 | 1 | 0,1329 | 355,5595 | 7,06531 | 206,6809 | 1362 | 0,33333 |
| KCNE4 | 3 | 0,30779 | 2 | 3 | 1,683 | 1 | 0,11629 | 310,2941 | 6,63868 | 117,2248 | 552 | 0,33333 |
| ZFP36L2 | 3 | 0,30779 | 2 | 3 | 1,287 | 1 | 0,11629 | 295,5131 | 6,45781 | 49,43831 | 218 | 0,33333 |
| CACHD1 | 3 | 0 | 1 | 3 | 1,211 | 1 | 0,10337 | 264,5671 | 6,03767 | 298,5759 | 1508 | 0 |
| PSKH1 | 3 | 0 | 1 | 3 | 1,435 | 1 | 0,1329 | 308,8452 | 6,62003 | 289,2416 | 1660 | 0 |
| FYCO1 | 3 | 0 | 1 | 3 | 2,01 | 1 | 0,11629 | 336,0107 | 6,8958 | 138,7183 | 1180 | 0 |
| ETNPPL | 3 | 0 | 1 | 3 | 1,47 | 2 | 0,11629 | 294,5869 | 6,44646 | 2420,429 | 33740 | 0 |
| HSD3B7 | 3 | 0 | 1 | 3 | 1,11 | 1 | 0,11629 | 295,1155 | 6,44808 | 150,4282 | 602 | 0 |
| PIEZO1 | 3 | 0 | 1 | 3 | 1,404 | 1 | 0,11629 | 310,5345 | 6,63868 | 82,13642 | 456 | 0 |
| NXNL1 | 3 | 0 | 1 | 3 | 1,149 | 3 | 0,11629 | 298,2726 | 6,4935 | 4690,991 | 40892 | 0 |
| MARCH3 | 3 | 0 | 1 | 3 | 2,701 | 1 | 0,1329 | 341,5357 | 6,95095 | 392,3286 | 4586 | 0 |
| RBBP6 | 3 | 0 | 1 | 3 | 1,746 | 1 | 0,1329 | 328,3714 | 6,80901 | 234,812 | 1652 | 0 |
| GPR4 | 3 | 0 | 1 | 3 | 3,584 | 1 | 0,11629 | 343,2369 | 6,93554 | 65,02779 | 654 | 0 |
| HIST1H2BC | 6 | 0,46346 | 3 | 3 | 1,428 | 1 | 0,11629 | 308,9845 | 6,6176 | 0 | 0 | 1 |
| CRTAP | 6 | 0,46346 | 3 | 3 | 3,991 | 1 | 0,11629 | 330,4441 | 6,83334 | 0 | 0 | 1 |
| KANK3 | 3 | 0 | 1 | 3 | 2,089 | 3 | 0,11629 | 328,8869 | 6,82118 | 4588,816 | 53482 | 0 |
| SLC25A18 | 3 | 0,30779 | 2 | 3 | 1,224 | 1 | 0,11629 | 296,025 | 6,4643 | 89,05713 | 482 | 0,33333 |
| KREMEN1 | 3 | 0,30779 | 2 | 3 | 2,332 | 1 | 0,1329 | 339,9262 | 6,93148 | 15,80371 | 226 | 0,33333 |
| TOB2 | 3 | 0,30779 | 2 | 3 | 1,97 | 2 | 0,11629 | 315,1464 | 6,67518 | 2292 | 32534 | 0,33333 |
| NEXN | 3 | 0 | 1 | 3 | 1,214 | 1 | 0,11629 | 295,806 | 6,47647 | 79,14014 | 400 | 0 |
| EMID1 | 3 | 0 | 1 | 3 | 1,405 | 1 | 0,11629 | 315,8869 | 6,68735 | 139,327 | 826 | 0 |
| NXN | 3 | 0 | 1 | 3 | 3,167 | 1 | 0,1329 | 350,9524 | 7,02638 | 218,8637 | 2406 | 0 |
| PAQR6 | 3 | 0 | 1 | 3 | 1,095 | 2 | 0,1329 | 285,4048 | 6,34102 | 2588,957 | 21926 | 0 |
| ZNF395 | 3 | 0 | 1 | 3 | 1,481 | 2 | 0,11629 | 329,5941 | 6,83578 | 1675,285 | 14302 | 0 |
| TNFSF13 | 3 | 0,30779 | 2 | 3 | 2,832 | 1 | 0,11629 | 337,0512 | 6,89093 | 25,96161 | 410 | 0,33333 |
| PEAR1 | 3 | 0 | 1 | 3 | 1,393 | 1 | 0,11629 | 306,3274 | 6,603 | 40,27722 | 326 | 0 |
| HVCN1 | 3 | 0 | 1 | 3 | 1,236 | 1 | 0,10337 | 307,1194 | 6,5811 | 319,1004 | 1466 | 0 |
| SASH3 | 3 | 0 | 1 | 3 | 2,138 | 1 | 0,11629 | 311,5798 | 6,62003 | 147,4814 | 1314 | 0 |
| HMGN5 | 3 | 0 | 1 | 3 | 1,441 | 1 | 0,11629 | 299,3726 | 6,50404 | 96,97294 | 468 | 0 |
| LRP10 | 6 | 0,46346 | 3 | 3 | 2,914 | 1 | 0,11629 | 330,0512 | 6,82929 | 0 | 0 | 1 |
| ARHGEF15 | 3 | 0,30779 | 2 | 3 | 1,242 | 1 | 0,11629 | 297,7464 | 6,49107 | 152,0171 | 2434 | 0,33333 |
| GPR137B | 3 | 0 | 1 | 3 | 1,101 | 1 | 0,1329 | 265,0786 | 6,06038 | 173,606 | 564 | 0 |
| RPS27 | 3 | 0 | 1 | 3 | 2,995 | 1 | 0,11629 | 357,0179 | 7,08559 | 121,921 | 1626 | 0 |
| PRPF38B | 6 | 0,46346 | 3 | 3 | 1,159 | 1 | 0,11629 | 273,9107 | 6,1642 | 0 | 0 | 1 |
| LRRC1 | 3 | 0 | 1 | 3 | 2,199 | 1 | 0,1329 | 326,2762 | 6,79847 | 36,1754 | 284 | 0 |
| LONRF3 | 3 | 0,30779 | 2 | 3 | 1,591 | 1 | 0,11629 | 316,8941 | 6,69384 | 53,85056 | 670 | 0,33333 |
| WISP2 | 3 | 0 | 1 | 3 | 2,366 | 1 | 0,15505 | 358,0833 | 7,08316 | 31,86861 | 484 | 0 |
| FAM129B | 3 | 0,30779 | 2 | 3 | 1,245 | 1 | 0,1329 | 276,5643 | 6,20962 | 173,7966 | 778 | 0,33333 |
| TGIF2 | 3 | 0,30779 | 2 | 3 | 3,412 | 1 | 0,1329 | 348,0357 | 6,99718 | 7,8604 | 94 | 0,33333 |
| S1PR3 | 3 | 0,30779 | 2 | 3 | 2,971 | 1 | 0,11629 | 348,7512 | 7,00448 | 11,61488 | 266 | 0,33333 |
| CENPB | 3 | 0 | 1 | 3 | 1,429 | 1 | 0,11629 | 307,7941 | 6,60543 | 125,7936 | 652 | 0 |
| MT1M | 6 | 0,46346 | 3 | 3 | 1,103 | 1 | 0,10337 | 245,7718 | 5,73676 | 0 | 0 | 1 |
| PXDC1 | 3 | 0 | 1 | 3 | 1,229 | 2 | 0,11629 | 307,4488 | 6,59732 | 442,6682 | 3110 | 0 |
| C2CD4B | 3 | 0,30779 | 2 | 3 | 1,624 | 2 | 0,1329 | 328,3691 | 6,82685 | 360,8852 | 3094 | 0,33333 |
| CATSPERD | 3 | 0 | 1 | 3 | 1,156 | 2 | 0,10337 | 266,823 | 6,07174 | 2384,52 | 24098 | 0 |
| RBMS3 | 3 | 0,30779 | 2 | 3 | 1,104 | 1 | 0,11629 | 238,3417 | 5,59077 | 267,3 | 1842 | 0,33333 |
| SAMD4A | 3 | 0 | 1 | 3 | 1,312 | 2 | 0,11629 | 300,4036 | 6,53243 | 195,9687 | 1576 | 0 |
| GPRC5C | 3 | 0 | 1 | 3 | 1,259 | 2 | 0,11629 | 302,6274 | 6,56001 | 566,154 | 3896 | 0 |
| BAZ2B | 3 | 0,30779 | 2 | 3 | 1,61 | 1 | 0,11629 | 322,4345 | 6,75305 | 54,86225 | 400 | 0,33333 |
| PAQR5 | 3 | 0 | 1 | 3 | 1,317 | 2 | 0,1329 | 292,4857 | 6,4205 | 596,1132 | 4022 | 0 |
| ITIH5 | 3 | 0 | 1 | 3 | 1,093 | 4 | 0,1329 | 272,4262 | 6,15447 | 6910,819 | 82354 | 0 |
| CXorf36 | 3 | 0 | 1 | 3 | 1,345 | 1 | 0,11629 | 277,9655 | 6,22666 | 271,1199 | 1386 | 0 |
| CECR1 | 3 | 0,30779 | 2 | 3 | 2,082 | 1 | 0,11629 | 317,4869 | 6,71493 | 8,80871 | 46 | 0,33333 |
| GATSL3 | 3 | 0,30779 | 2 | 3 | 1,838 | 1 | 0,1329 | 317,2786 | 6,71168 | 23,18967 | 204 | 0,33333 |
| FAH | 3 | 0 | 1 | 3 | 2,985 | 1 | 0,1329 | 340,9762 | 6,93959 | 57,67344 | 630 | 0 |
| ATHL1 | 6 | 0,46346 | 3 | 3 | 2,019 | 1 | 0,11629 | 307,0464 | 6,57542 | 0 | 0 | 1 |
| PPP1R3G | 6 | 0,46346 | 3 | 3 | 1,152 | 1 | 0,1329 | 289,6524 | 6,38725 | 0 | 0 | 1 |
| PPP1R13L | 3 | 0,30779 | 2 | 3 | 4,899 | 1 | 0,1329 | 375,5595 | 7,21455 | 4,65576 | 124 | 0,33333 |
| TCEA3 | 3 | 0 | 1 | 3 | 2,166 | 2 | 0,11629 | 329,2679 | 6,83091 | 756,9922 | 8628 | 0 |
| TMBIM1 | 3 | 0 | 1 | 3 | 3,221 | 2 | 0,11629 | 359,3512 | 7,09451 | 2298,744 | 17764 | 0 |
| SIGLEC9 | 3 | 0,30779 | 2 | 3 | 4,381 | 1 | 0,1329 | 366,8429 | 7,15859 | 5,63149 | 120 | 0,33333 |
| STK33 | 3 | 0 | 1 | 3 | 1,276 | 2 | 0,1329 | 291,8262 | 6,40834 | 2321,113 | 26522 | 0 |
| TMEM176A | 6 | 0,46346 | 3 | 3 | 1,303 | 1 | 0,1329 | 280,7667 | 6,25829 | 0 | 0 | 1 |
| ZBTB20 | 3 | 0 | 1 | 3 | 1,631 | 1 | 0,11629 | 314,8107 | 6,69303 | 58,54925 | 318 | 0 |
| MAP3K6 | 3 | 0 | 1 | 3 | 1,227 | 1 | 0,11629 | 300,1226 | 6,50648 | 148,3659 | 746 | 0 |
| NUDT16 | 3 | 0,30779 | 2 | 3 | 2,41 | 1 | 0,1329 | 334,7691 | 6,88525 | 139,218 | 1414 | 0,33333 |
| TRIM34 | 4 | 0,30898 | 3 | 3 | 1,246 | 1 | 0,11629 | 306,2298 | 6,57948 | 39,0523 | 446 | 0,66667 |
| ENSP0000043201 | 3 | 0,30779 | 2 | 3 | 1,831 | 1 | 0,11629 | 329,7607 | 6,82442 | 20,74225 | 278 | 0,33333 |
| RAB30 | 3 | 0 | 1 | 3 | 1,104 | 1 | 0,11629 | 252,6107 | 5,8422 | 125,5576 | 870 | 0 |
| NOL8 | 3 | 0 | 1 | 3 | 1,437 | 1 | 0,1329 | 317,8786 | 6,71168 | 368,6942 | 2052 | 0 |

| | | | | | | | | | | | | |
|---|---|---|---|---|---|---|---|---|---|---|---|---|
| NEDD1 | 3 | 0,30779 | 2 | 3 | 1,577 | 1 | 0,11629 | 311,7702 | 6,65491 | 118,8059 | 732 | 0,33333 |
| METRN | 3 | 0 | 1 | 3 | 1,19 | 2 | 0,11629 | 300,9226 | 6,53487 | 242,0362 | 2380 | 0 |
| PIRT | 3 | 0 | 1 | 3 | 1,285 | 1 | 0,11629 | 287,4155 | 6,35643 | 144,5246 | 896 | 0 |
| PRICKLE3 | 3 | 0,30779 | 2 | 3 | 1,778 | 1 | 0,1329 | 313,8714 | 6,68329 | 17,77168 | 174 | 0,33333 |
| DHRS3 | 3 | 0 | 1 | 3 | 1,906 | 1 | 0,11629 | 331,3607 | 6,85524 | 68,06714 | 870 | 0 |
| FCGBP | 3 | 0 | 1 | 3 | 2,966 | 1 | 0,1329 | 328,3452 | 6,82037 | 372,4485 | 3898 | 0 |
| LPP | 3 | 0,30779 | 2 | 3 | 2,628 | 1 | 0,15505 | 334,9167 | 6,88363 | 17,89956 | 198 | 0,33333 |
| CMTM6 | 2 | 0 | 1 | 2 | 1,064 | 3 | 0,00243 | 2 | 0,00851 | 2 | 2 | 0 |
| SLC52A3 | 2 | 0 | 1 | 2 | 1,08 | 1 | 0,11629 | 284,9917 | 6,32804 | 99,42379 | 576 | 0 |
| GIMAP2 | 2 | 0 | 1 | 2 | 1,474 | 1 | 0,11629 | 293,3107 | 6,44402 | 65,43842 | 336 | 0 |
| ZBTB47 | 2 | 0 | 1 | 2 | 1,08 | 1 | 0,10337 | 264,8564 | 6,05714 | 55,51967 | 322 | 0 |
| WBP1 | 2 | 0 | 1 | 2 | 2,619 | 1 | 0,11629 | 338,2679 | 6,9177 | 4,48285 | 104 | 0 |
| SH3TC1 | 2 | 0 | 1 | 2 | 1,059 | 1 | 0,10337 | 269,9194 | 6,13338 | 63,58549 | 208 | 0 |
| PSD4 | 2 | 0 | 1 | 2 | 1,215 | 1 | 0,10337 | 270,2123 | 6,12933 | 70,3799 | 332 | 0 |
| GDF1 | 2 | 0,30779 | 2 | 2 | 1,283 | 1 | 0,11629 | 300,356 | 6,53649 | 0 | 0 | 1 |
| FAM189A2 | 2 | 0 | 1 | 2 | 1,269 | 1 | 0,11629 | 299,6536 | 6,53 | 13,23962 | 194 | 0 |
| RNF144B | 2 | 0 | 1 | 2 | 1,073 | 1 | 0,10337 | 277,8456 | 6,23963 | 26,20145 | 186 | 0 |
| CSPP1 | 2 | 0 | 1 | 2 | 1,078 | 1 | 0,10337 | 273,0861 | 6,18042 | 763,1595 | 5706 | 0 |
| CRISPLD2 | 2 | 0 | 1 | 2 | 1,221 | 2 | 0,11629 | 286,9012 | 6,36211 | 2292 | 23998 | 0 |
| PLEKHA4 | 2 | 0 | 1 | 2 | 1,435 | 1 | 0,11629 | 314,5274 | 6,69059 | 84,05533 | 656 | 0 |
| STK17B | 2 | 0 | 1 | 2 | 1,073 | 2 | 0,11629 | 234,6369 | 5,53967 | 2292 | 21000 | 0 |
| PGPEP1 | 2 | 0 | 1 | 2 | 4,35 | 1 | 0,1329 | 338,6929 | 6,92337 | 1,19586 | 26 | 0 |
| ARHGEF19 | 2 | 0 | 1 | 2 | 1,571 | 1 | 0,11629 | 321,8607 | 6,7644 | 148,2535 | 2218 | 0 |
| MOB3C | 2 | 0 | 1 | 2 | 1,231 | 1 | 0,11629 | 311,9393 | 6,65247 | 20,51607 | 194 | 0 |
| RAB41 | 2 | 0 | 1 | 2 | 1,193 | 1 | 0,10337 | 286,2861 | 6,33534 | 426,3953 | 5760 | 0 |
| ITPRIP | 2 | 0 | 1 | 2 | 1,067 | 3 | 0,11629 | 219,6286 | 5,22822 | 4580 | 54756 | 0 |
| MSI2 | 2 | 0 | 1 | 2 | 2,622 | 1 | 0,1329 | 355,6762 | 7,0718 | 82,17043 | 878 | 0 |
| ANKRD40 | 2 | 0 | 1 | 2 | 1,118 | 1 | 0,1329 | 266,1095 | 6,07823 | 197,9857 | 796 | 0 |
| LRRC3 | 2 | 0 | 1 | 2 | 3,276 | 1 | 0,1329 | 332,7452 | 6,85362 | 4,11353 | 68 | 0 |
| PDZK1IP1 | 2 | 0 | 1 | 2 | 1,96 | 1 | 0,11629 | 301,4655 | 6,53811 | 18,75117 | 186 | 0 |
| FOXQ1 | 2 | 0,30779 | 2 | 2 | 1,596 | 1 | 0,11629 | 292,3083 | 6,41888 | 0 | 0 | 1 |
| A2ML1 | 2 | 0 | 1 | 2 | 1,513 | 1 | 0,11629 | 315,6369 | 6,69465 | 23,0395 | 160 | 0 |
| MIDN | 2 | 0 | 1 | 2 | 1,375 | 1 | 0,11629 | 303,5702 | 6,57218 | 2,85354 | 12 | 0 |
| PLIN4 | 2 | 0 | 1 | 2 | 2,119 | 2 | 0,11629 | 334,5774 | 6,88201 | 899,2345 | 8336 | 0 |
| SCARA3 | 2 | 0 | 1 | 2 | 3,125 | 5 | 0,11629 | 360,9012 | 7,11154 | 436,9157 | 4750 | 0 |
| VAMP5 | 2 | 0 | 1 | 2 | 2,106 | 1 | 0,11629 | 332,8274 | 6,8447 | 11,83687 | 188 | 0 |
| POLD4 | 2 | 0 | 1 | 2 | 2,157 | 1 | 0,11629 | 333,9107 | 6,88606 | 322,941 | 3910 | 0 |
| UACA | 2 | 0,30779 | 2 | 2 | 1,776 | 1 | 0,1329 | 341,5857 | 6,95744 | 0 | 0 | 1 |
| TMEM102 | 2 | 0 | 1 | 2 | 1,082 | 2 | 0,10337 | 184,3103 | 4,30034 | 2292 | 27380 | 0 |
| ODF3L2 | 2 | 0 | 1 | 2 | 3,408 | 2 | 0,1329 | 383,8429 | 7,28917 | 709,3754 | 5996 | 0 |
| CLEC1A | 2 | 0 | 1 | 2 | 1,167 | 1 | 0,11629 | 294,9226 | 6,46349 | 3185,38 | 29968 | 0 |
| ACSM5 | 2 | 0 | 1 | 2 | 1,155 | 2 | 0,10337 | 280,6194 | 6,28019 | 2292 | 24190 | 0 |
| CHST6 | 2 | 0 | 1 | 2 | 1,134 | 1 | 0,11629 | 290,7083 | 6,41077 | 6,49185 | 32 | 0 |
| BACE2 | 2 | 0 | 1 | 2 | 1,459 | 2 | 0,11629 | 299,6226 | 6,51702 | 533,7819 | 4652 | 0 |
| SLC51B | 2 | 0 | 1 | 2 | 1,064 | 1 | 0,10337 | 274,398 | 6,19583 | 11,8857 | 110 | 0 |
| FAM60A | 2 | 0 | 1 | 2 | 1,874 | 1 | 0,11629 | 314,7702 | 6,68248 | 8,73148 | 108 | 0 |
| ASAP3 | 2 | 0 | 1 | 2 | 2,016 | 2 | 0,1329 | 341,6262 | 6,95987 | 86,10531 | 1090 | 0 |
| MTSS1L | 2 | 0 | 1 | 2 | 1,074 | 1 | 0,10337 | 259,8314 | 5,96387 | 106,8052 | 406 | 0 |
| PRRC2C | 2 | 0 | 1 | 2 | 1,362 | 1 | 0,11629 | 320,6441 | 6,74412 | 120,2155 | 946 | 0 |
| APOL3 | 2 | 0 | 1 | 2 | 1,349 | 1 | 0,11629 | 301,1226 | 6,51378 | 6,76406 | 86 | 0 |
| ENSP0000034691 | 2 | 0,30779 | 2 | 2 | 1,143 | 1 | 0,10337 | 295,6861 | 6,46674 | 0 | 0 | 1 |
| PLSCR4 | 2 | 0 | 1 | 2 | 1,078 | 1 | 0,1329 | 293,1405 | 6,4351 | 56,49701 | 434 | 0 |
| KIAA1598 | 2 | 0 | 1 | 2 | 1,344 | 1 | 0,11629 | 295,2988 | 6,4643 | 53,74599 | 258 | 0 |
| NPAS3 | 2 | 0 | 1 | 2 | 1,234 | 2 | 0,11629 | 286,0845 | 6,35075 | 192,6281 | 2300 | 0 |
| RBM38 | 2 | 0 | 1 | 2 | 2,489 | 1 | 0,11629 | 341,1512 | 6,95257 | 14,07593 | 288 | 0 |
| BTN2A2 | 2 | 0 | 1 | 2 | 1,066 | 2 | 0,11629 | 292,6845 | 6,44159 | 1580,43 | 15720 | 0 |
| FAM114A1 | 2 | 0 | 1 | 2 | 1,474 | 2 | 0,11629 | 308,0298 | 6,60543 | 2292 | 26256 | 0 |
| HAPLN3 | 2 | 0 | 1 | 2 | 2,164 | 1 | 0,11629 | 316,0607 | 6,69465 | 6,00547 | 86 | 0 |
| MAP4 | 2 | 0 | 1 | 2 | 1,903 | 1 | 0,11629 | 325,4607 | 6,80496 | 152,9627 | 1222 | 0 |
| RAB29 | 2 | 0 | 1 | 2 | 1,704 | 1 | 0,1329 | 312,1191 | 6,66545 | 17,67551 | 184 | 0 |
| SASH1 | 2 | 0 | 1 | 2 | 1,205 | 1 | 0,11629 | 280,3726 | 6,27856 | 83,63435 | 474 | 0 |
| PBXIP1 | 2 | 0 | 1 | 2 | 1,062 | 1 | 0,11629 | 284,0917 | 6,32561 | 61,53499 | 280 | 0 |
| PRSS35 | 2 | 0 | 1 | 2 | 1,096 | 2 | 0,1329 | 274,1762 | 6,19502 | 155,5793 | 1138 | 0 |
| ADIRF | 2 | 0 | 1 | 2 | 1,282 | 1 | 0,11629 | 304,9369 | 6,56731 | 52,10146 | 626 | 0 |
| TMEM164 | 2 | 0,30779 | 2 | 2 | 1,091 | 1 | 0,10337 | 250,6825 | 5,83085 | 0 | 0 | 1 |
| PLAC9 | 2 | 0 | 1 | 2 | 1,253 | 1 | 0,11629 | 287,3083 | 6,36048 | 14,25132 | 66 | 0 |
| GPSM3 | 2 | 0 | 1 | 2 | 1,704 | 1 | 0,11629 | 326,7964 | 6,80901 | 18,08739 | 228 | 0 |
| VSX1 | 2 | 0 | 1 | 2 | 1,069 | 2 | 0,10337 | 235,6266 | 5,56563 | 2292 | 20116 | 0 |
| SERPINB1 | 2 | 0 | 1 | 2 | 3,743 | 1 | 0,1329 | 378,6429 | 7,2405 | 30,0069 | 620 | 0 |
| PLIN5 | 2 | 0 | 1 | 2 | 2,206 | 1 | 0,11629 | 339,1941 | 6,91851 | 6,30004 | 152 | 0 |
| ZMYM5 | 2 | 0,30779 | 2 | 2 | 1,058 | 1 | 0,10337 | 220,0302 | 5,23876 | 0 | 0 | 1 |
| TTC23 | 2 | 0 | 1 | 2 | 1,063 | 1 | 0,10337 | 241,6433 | 5,67512 | 40,22164 | 166 | 0 |
| SIPA1 | 2 | 0 | 1 | 2 | 2,049 | 1 | 0,11629 | 320,6179 | 6,74656 | 6,74199 | 86 | 0 |
| SGMS2 | 2 | 0 | 1 | 2 | 2,292 | 2 | 0,11629 | 354,4012 | 7,06288 | 2292 | 20282 | 0 |
| KTN1 | 2 | 0 | 1 | 2 | 1,196 | 2 | 0,11629 | 298,606 | 6,51621 | 2292 | 17730 | 0 |
| SLC5A10 | 2 | 0 | 1 | 2 | 2,054 | 1 | 0,11629 | 330,8131 | 6,84713 | 79,05772 | 508 | 0 |
| FGFRL1 | 2 | 0,30779 | 2 | 2 | 3,514 | 1 | 0,11629 | 338,3607 | 6,92418 | 0 | 0 | 1 |
| CARD10 | 2 | 0 | 1 | 2 | 1,764 | 1 | 0,11629 | 315,4274 | 6,69708 | 35,10532 | 272 | 0 |
| PRR5 | 2 | 0 | 1 | 2 | 2,047 | 1 | 0,11629 | 317,4274 | 6,72141 | 44,55852 | 288 | 0 |

| | | | | | | | | | | | | |
|---|---|---|---|---|---|---|---|---|---|---|---|---|
| TMCO3 | 2 | 0 | 1 | 2 | 1,256 | 2 | 0,11629 | 302,7893 | 6,54135 | 2292 | 21288 | 0 |
| SLC14A1 | 2 | 0 | 1 | 2 | 2,024 | 1 | 0,11629 | 326,2107 | 6,81144 | 15,33096 | 136 | 0 |
| C1orf106 | 2 | 0 | 1 | 2 | 1,504 | 1 | 0,11629 | 313,1369 | 6,66139 | 19,34835 | 140 | 0 |
| PLEKHG2 | 2 | 0,30779 | 2 | 2 | 1,089 | 1 | 0,10337 | 251,2575 | 5,82679 | 0 | 0 | 1 |
| MRO | 2 | 0 | 1 | 2 | 1,069 | 1 | 0,11629 | 274,3345 | 6,19908 | 7,91579 | 56 | 0 |
| ASRGL1 | 2 | 0 | 1 | 2 | 1,721 | 1 | 0,11629 | 297,0202 | 6,48377 | 19,01109 | 140 | 0 |
| GDPD2 | 2 | 0 | 1 | 2 | 1,101 | 2 | 0,11629 | 273,9012 | 6,18853 | 419,7796 | 5334 | 0 |
| ABHD4 | 2 | 0 | 1 | 2 | 1,244 | 2 | 0,10337 | 254,079 | 5,8933 | 2292 | 26686 | 0 |
| GOLIM4 | 2 | 0 | 1 | 2 | 1,293 | 1 | 0,11629 | 308,7298 | 6,63057 | 102,507 | 1128 | 0 |
| TIPARP | 2 | 0 | 1 | 2 | 1,535 | 1 | 0,11629 | 304,0321 | 6,56488 | 407,9969 | 4452 | 0 |
| RANBP3L | 2 | 0 | 1 | 2 | 1,067 | 1 | 0,1329 | 264,8691 | 6,05633 | 64,88183 | 386 | 0 |
| ELOVL7 | 2 | 0 | 1 | 2 | 1,086 | 1 | 0,10337 | 290,4433 | 6,39779 | 32,88396 | 228 | 0 |
| NEK1 | 2 | 0 | 1 | 2 | 1,075 | 3 | 0,10337 | 259,4183 | 5,96143 | 399,004 | 3070 | 0 |
| NWD1 | 2 | 0 | 1 | 2 | 1,07 | 1 | 0,10337 | 264,5849 | 6,0174 | 81,88408 | 934 | 0 |
| TPD52L1 | 2 | 0 | 1 | 2 | 2,365 | 2 | 0,1329 | 338,0619 | 6,91851 | 2292 | 21938 | 0 |
| FAM111A | 2 | 0 | 1 | 2 | 1,522 | 2 | 0,11629 | 332,5107 | 6,87309 | 2292 | 20618 | 0 |
| TSKU | 2 | 0 | 1 | 2 | 2,556 | 1 | 0,1329 | 320,6286 | 6,7352 | 1,20686 | 12 | 0 |
| CRACR2B | 2 | 0 | 1 | 2 | 1,184 | 1 | 0,11629 | 291,0631 | 6,40509 | 120,9305 | 540 | 0 |
| GALNT4 | 2 | 0,30779 | 2 | 2 | 1,681 | 1 | 0,11629 | 324,4441 | 6,79441 | 0 | 0 | 1 |
| C21orf62 | 2 | 0 | 1 | 2 | 1,824 | 1 | 0,11629 | 304,4131 | 6,57542 | 25,52197 | 196 | 0 |
| APPL2 | 2 | 0 | 1 | 2 | 3,262 | 1 | 0,1329 | 367,2762 | 7,15534 | 195,9175 | 1774 | 0 |
| ENSP000004478 | 2 | 0 | 1 | 2 | 1,086 | 1 | 0,10337 | 262,1706 | 6,01659 | 32,82811 | 178 | 0 |
| TPCN1 | 2 | 0 | 1 | 2 | 1,097 | 2 | 0,10337 | 249,4564 | 5,81382 | 2292 | 21218 | 0 |
| DPEP2 | 2 | 0 | 1 | 2 | 1,21 | 1 | 0,1329 | 300,2 | 6,52838 | 7,89259 | 38 | 0 |
| FXYD3 | 2 | 0 | 1 | 2 | 1,068 | 1 | 0,11629 | 289,8488 | 6,40996 | 8,46726 | 54 | 0 |
| MR1 | 2 | 0,30779 | 2 | 2 | 1,737 | 1 | 0,11629 | 289,1345 | 6,36292 | 0 | 0 | 1 |
| SMOX | 2 | 0 | 1 | 2 | 1,414 | 1 | 0,10337 | 283,4266 | 6,31831 | 26,44467 | 148 | 0 |
| SEC14L2 | 2 | 0 | 1 | 2 | 1,447 | 4 | 0,11629 | 307,6202 | 6,62084 | 3689,476 | 23232 | 0 |
| MID1IP1 | 2 | 0 | 1 | 2 | 1,073 | 1 | 0,10337 | 275,8766 | 6,2299 | 20,47017 | 124 | 0 |
| COLCA2 | 2 | 0 | 1 | 2 | 1,05 | 1 | 0,10337 | 266,923 | 6,09769 | 22,52069 | 138 | 0 |
| CERS1 | 2 | 0,30779 | 2 | 2 | 1,225 | 1 | 0,11629 | 300,356 | 6,53649 | 0 | 0 | 1 |
| SLCO4A1 | 1 | 0 | 1 | 1 | 1,149 | 1 | 0,10337 | 268,0242 | 6,11797 | 0 | 0 | 0 |
| DAP | 1 | 0 | 1 | 1 | 2,001 | 1 | 0,11629 | 332,5845 | 6,88444 | 0 | 0 | 0 |
| USHBP1 | 1 | 0 | 1 | 1 | 1,316 | 1 | 0,10337 | 266,4218 | 6,07579 | 0 | 0 | 0 |
| EMILIN2 | 1 | 0 | 1 | 1 | 1,163 | 1 | 0,11629 | 267,4964 | 6,10743 | 0 | 0 | 0 |
| TRIM47 | 1 | 0 | 1 | 1 | 1,369 | 1 | 0,11629 | 319,1036 | 6,73845 | 0 | 0 | 0 |
| TSPAN11 | 1 | 0 | 1 | 1 | 1,031 | 1 | 0,00122 | 1,5 | 0,00729 | 0 | 0 | 0 |
| FAM181A | 1 | 0 | 1 | 1 | 1,581 | 1 | 0,11629 | 304,5798 | 6,5811 | 0 | 0 | 0 |
| ARRDC4 | 1 | 0 | 1 | 1 | 1,133 | 1 | 0,11629 | 311,0774 | 6,65653 | 0 | 0 | 0 |
| EVA1B | 1 | 0 | 1 | 1 | 2,054 | 1 | 0,11629 | 333,2941 | 6,8739 | 0 | 0 | 0 |
| SLC44A3 | 1 | 0 | 1 | 1 | 1,029 | 1 | 0,00162 | 1 | 0,00486 | 0 | 0 | 0 |
| SULT1C4 | 1 | 0 | 1 | 1 | 1,644 | 1 | 0,11629 | 290,0536 | 6,41077 | 0 | 0 | 0 |
| PPP1R18 | 1 | 0 | 1 | 1 | 1,163 | 1 | 0,11629 | 282,825 | 6,31425 | 0 | 0 | 0 |
| NACC2 | 1 | 0 | 1 | 1 | 1,031 | 1 | 0,10337 | 251,1694 | 5,84707 | 0 | 0 | 0 |
| C16orf74 | 1 | 0 | 1 | 1 | 1,197 | 1 | 0,11629 | 277,4441 | 6,24855 | 0 | 0 | 0 |
| PREX2 | 1 | 0 | 1 | 1 | 1,89 | 1 | 0,11629 | 323,1845 | 6,79198 | 0 | 0 | 0 |
| ZNF618 | 1 | 0 | 1 | 1 | 1,049 | 1 | 0,10337 | 259,8123 | 5,98901 | 0 | 0 | 0 |
| ENSP000002938 | 1 | 0 | 1 | 1 | 1,402 | 1 | 0,11629 | 302,8893 | 6,5592 | 0 | 0 | 0 |
| KIAA1161 | 1 | 0 | 1 | 1 | 2,843 | 1 | 0,1329 | 341,7095 | 6,95419 | 0 | 0 | 0 |
| CNTNAP3 | 1 | 0 | 1 | 1 | 1,032 | 1 | 0,09303 | 205,1341 | 4,91352 | 0 | 0 | 0 |
| ABHD15 | 1 | 0 | 1 | 1 | 1,029 | 1 | 0,09303 | 207,1321 | 4,96381 | 0 | 0 | 0 |
| CCDC8 | 1 | 0 | 1 | 1 | 1,041 | 1 | 0,11629 | 275,575 | 6,22747 | 0 | 0 | 0 |
| PHYHD1 | 1 | 0 | 1 | 1 | 1,037 | 1 | 0,10337 | 232,6456 | 5,51696 | 0 | 0 | 0 |
| PPP1R14B | 1 | 0 | 1 | 1 | 1,037 | 1 | 0,10337 | 235,5361 | 5,58671 | 0 | 0 | 0 |
| ZNF366 | 1 | 0 | 1 | 1 | 1,256 | 1 | 0,10337 | 272,029 | 6,15934 | 0 | 0 | 0 |
| C17orf53 | 1 | 0 | 1 | 1 | 1,037 | 1 | 0,11629 | 254,5036 | 5,91115 | 0 | 0 | 0 |
| ORAI3 | 1 | 0 | 1 | 1 | 1,036 | 1 | 0,10337 | 273,0647 | 6,18772 | 0 | 0 | 0 |
| MTURN | 1 | 0 | 1 | 1 | 1,236 | 1 | 0,1329 | 285,8857 | 6,35237 | 0 | 0 | 0 |
| APOLD1 | 1 | 0 | 1 | 1 | 1,044 | 1 | 0,10337 | 265,1135 | 6,07336 | 0 | 0 | 0 |
| CASKIN2 | 1 | 0 | 1 | 1 | 1,031 | 1 | 0,11629 | 271,6417 | 6,17312 | 0 | 0 | 0 |
| PTTG1IP | 1 | 0 | 1 | 1 | 1,065 | 1 | 0,09303 | 203,9948 | 4,88432 | 0 | 0 | 0 |
| FAM101B | 1 | 0 | 1 | 1 | 1,176 | 1 | 0,11629 | 281,7917 | 6,29479 | 0 | 0 | 0 |
| SLC44A2 | 1 | 0 | 1 | 1 | 1,976 | 1 | 0,1329 | 344,7357 | 6,97366 | 0 | 0 | 0 |
| APOL4 | 1 | 0 | 1 | 1 | 1,038 | 1 | 0,10337 | 252,8635 | 5,86654 | 0 | 0 | 0 |
| LCN6 | 1 | 0 | 1 | 1 | 1,024 | 1 | 0,00162 | 1 | 0,00486 | 0 | 0 | 0 |
| C1QTNF1 | 1 | 0 | 1 | 1 | 1,614 | 1 | 0,11629 | 316,2941 | 6,71006 | 0 | 0 | 0 |
| SYDE1 | 1 | 0 | 1 | 1 | 1,032 | 1 | 0,00162 | 1 | 0,00486 | 0 | 0 | 0 |
| MSRB3 | 1 | 0 | 1 | 1 | 1,109 | 1 | 0,10337 | 263,0397 | 6,03605 | 0 | 0 | 0 |
| CCDC69 | 1 | 0 | 1 | 1 | 1,023 | 1 | 0,10337 | 228,1397 | 5,43261 | 0 | 0 | 0 |
| MORC4 | 1 | 0 | 1 | 1 | 1,112 | 1 | 0,11629 | 259,0345 | 5,98901 | 0 | 0 | 0 |
| ADD3 | 1 | 0 | 1 | 1 | 1,03 | 1 | 0,10337 | 261,6552 | 6,01983 | 0 | 0 | 0 |
| ATP10A | 1 | 0 | 1 | 1 | 1,31 | 1 | 0,11629 | 287,556 | 6,36778 | 0 | 0 | 0 |
| ZNF98 | 1 | 0 | 1 | 1 | 1,031 | 1 | 0,09303 | 205,1341 | 4,91352 | 0 | 0 | 0 |
| GRAMD1C | 1 | 0 | 1 | 1 | 1,024 | 1 | 0,00162 | 1 | 0,00486 | 0 | 0 | 0 |
| ARHGAP19 | 1 | 0 | 1 | 1 | 1,121 | 1 | 0,10337 | 261,6552 | 6,01983 | 0 | 0 | 0 |
| LRRCC1 | 1 | 0 | 1 | 1 | 1,044 | 1 | 0,10337 | 270,5028 | 6,15609 | 0 | 0 | 0 |
| FAM89A | 1 | 0 | 1 | 1 | 1,097 | 1 | 0,10337 | 280,129 | 6,24369 | 0 | 0 | 0 |
| MROH9 | 1 | 0 | 1 | 1 | 1,032 | 1 | 0,11629 | 252,1357 | 5,87384 | 0 | 0 | 0 |
| C1orf110 | 1 | 0 | 1 | 1 | 1,03 | 1 | 0,11629 | 227,0619 | 5,41152 | 0 | 0 | 0 |

| | | | | | | | | | | | | |
|---|---|---|---|---|---|---|---|---|---|---|---|---|
| S100A16 | 1 | 0 | 1 | 1 | 1,032 | 1 | 0,10337 | 245,7694 | 5,7757 | 0 | 0 | 0 |
| PIFO | 1 | 0 | 1 | 1 | 1,027 | 1 | 0,00162 | 1 | 0,00486 | 0 | 0 | 0 |
| NKAIN4 | 1 | 0 | 1 | 1 | 1,021 | 1 | 0,10337 | 255,8825 | 5,94359 | 0 | 0 | 0 |
| UBTD1 | 1 | 0 | 1 | 1 | 1,109 | 1 | 0,10337 | 265,0885 | 6,08147 | 0 | 0 | 0 |
| ZMAT1 | 1 | 0 | 1 | 1 | 1,029 | 1 | 0,09303 | 194,7306 | 4,63613 | 0 | 0 | 0 |
| SYTL4 | 1 | 0 | 1 | 1 | 2,272 | 1 | 0,1329 | 341,2762 | 6,94771 | 0 | 0 | 0 |
| FAM167B | 1 | 0 | 1 | 1 | 1,039 | 1 | 0,11629 | 230,9286 | 5,47884 | 0 | 0 | 0 |
| RUFY4 | 1 | 0 | 1 | 1 | 1,024 | 1 | 0,10337 | 270,9552 | 6,16501 | 0 | 0 | 0 |
| TP53INP2 | 1 | 0 | 1 | 1 | 1,719 | 1 | 0,11629 | 316,5536 | 6,71411 | 0 | 0 | 0 |
| RIBC1 | 1 | 0 | 1 | 1 | 1,041 | 1 | 0,10337 | 245,1897 | 5,74569 | 0 | 0 | 0 |
| PLP2 | 1 | 0 | 1 | 1 | 1,039 | 1 | 0,11629 | 287,5655 | 6,37833 | 0 | 0 | 0 |
| SOWAHC | 1 | 0 | 1 | 1 | 1,032 | 1 | 0,00162 | 1 | 0,00486 | 0 | 0 | 0 |
| ST8SIA6 | 1 | 0 | 1 | 1 | 1,037 | 1 | 0,10337 | 223,5004 | 5,33609 | 0 | 0 | 0 |
| HIF3A | 1 | 0 | 1 | 1 | 1,709 | 1 | 0,11629 | 318,1607 | 6,74088 | 0 | 0 | 0 |
| TTC38 | 1 | 0 | 1 | 1 | 1,044 | 1 | 0,10337 | 262,2837 | 6,01983 | 0 | 0 | 0 |
| ITPRIPL2 | 1 | 0 | 1 | 1 | 1,043 | 1 | 0,09303 | 158,3698 | 3,37085 | 0 | 0 | 0 |
| OTOS | 1 | 0 | 1 | 1 | 1,144 | 1 | 0,10337 | 248,2052 | 5,80327 | 0 | 0 | 0 |
| LRRIQ1 | 1 | 0 | 1 | 1 | 1,029 | 1 | 0,00162 | 1 | 0,00486 | 0 | 0 | 0 |
| RGL3 | 1 | 0 | 1 | 1 | 1,149 | 1 | 0,11629 | 272,8583 | 6,17475 | 0 | 0 | 0 |
| METTL7B | 1 | 0 | 1 | 1 | 1,402 | 1 | 0,11629 | 286,6393 | 6,36859 | 0 | 0 | 0 |
| NUPR1 | 1 | 0 | 1 | 1 | 1,159 | 1 | 0,10337 | 260,6933 | 6,00929 | 0 | 0 | 0 |
| GPR146 | 1 | 0 | 1 | 1 | 1,035 | 1 | 0,11629 | 257,5345 | 5,9444 | 0 | 0 | 0 |
| CCDC11 | 1 | 0 | 1 | 1 | 1,062 | 1 | 0,10337 | 258,6194 | 5,96305 | 0 | 0 | 0 |
| AIFM3 | 1 | 0 | 1 | 1 | 1,04 | 1 | 0,11629 | 287,206 | 6,37671 | 0 | 0 | 0 |
| ENHO | 1 | 0 | 1 | 1 | 2,915 | 1 | 0,1329 | 363,4929 | 7,12695 | 0 | 0 | 0 |
| COLEC12 | 1 | 0 | 1 | 1 | 1,034 | 1 | 0,10337 | 265,0075 | 6,07093 | 0 | 0 | 0 |
| SZRD1 | 1 | 0 | 1 | 1 | 1,032 | 1 | 0,10337 | 265,1135 | 6,07336 | 0 | 0 | 0 |
| CYP4F11 | 1 | 0 | 1 | 1 | 1,044 | 1 | 0,10337 | 263,5778 | 6,03362 | 0 | 0 | 0 |
| GAREML | 1 | 0 | 1 | 1 | 1,293 | 1 | 0,1329 | 303,9357 | 6,58515 | 0 | 0 | 0 |
| ANKRD36 | 1 | 0 | 1 | 1 | 1,031 | 1 | 0,00162 | 1 | 0,00486 | 0 | 0 | 0 |
| NHSL1 | 1 | 0 | 1 | 1 | 1,033 | 1 | 0,11629 | 273,4583 | 6,20557 | 0 | 0 | 0 |
| OR6V1 | 1 | 0 | 1 | 1 | 1,022 | 1 | 0,09303 | 215,1127 | 5,14224 | 0 | 0 | 0 |
| CMTM3 | 1 | 0 | 1 | 1 | 1,035 | 1 | 0,00122 | 1,5 | 0,00729 | 0 | 0 | 0 |
| ANKRD36C | 1 | 0 | 1 | 1 | 1,031 | 1 | 0,00162 | 1 | 0,00486 | 0 | 0 | 0 |
| PAQR8 | 1 | 0 | 1 | 1 | 1,028 | 1 | 0,1329 | 282,0643 | 6,3102 | 0 | 0 | 0 |
| NT5DC2 | 1 | 0 | 1 | 1 | 1,045 | 1 | 0,09303 | 203,9611 | 4,86972 | 0 | 0 | 0 |
| IQCA1 | 1 | 0 | 1 | 1 | 1,042 | 1 | 0,10337 | 193,9829 | 4,61017 | 0 | 0 | 0 |
| USP53 | 1 | 0 | 1 | 1 | 1,025 | 1 | 0,10337 | 268,3528 | 6,13338 | 0 | 0 | 0 |
| ERG | 1 | 0 | 1 | 1 | 2,477 | 1 | 0,1329 | 342,0429 | 6,94771 | 0 | 0 | 0 |
| LCN10 | 1 | 0 | 1 | 1 | 1,024 | 1 | 0,00162 | 1 | 0,00486 | 0 | 0 | 0 |
| KLHL5 | 1 | 0 | 1 | 1 | 1,039 | 1 | 0,10337 | 241,0754 | 5,67593 | 0 | 0 | 0 |
| GRAMD3 | 1 | 0 | 1 | 1 | 1,024 | 1 | 0,00162 | 1 | 0,00486 | 0 | 0 | 0 |
| C8orf82 | 1 | 0 | 1 | 1 | 1,037 | 1 | 0,10337 | 253,329 | 5,88519 | 0 | 0 | 0 |
| SMIM3 | 1 | 0 | 1 | 1 | 1,224 | 1 | 0,11629 | 289,7726 | 6,41969 | 0 | 0 | 0 |
| CHI3L2 | 1 | 0 | 1 | 1 | 1,027 | 1 | 0,00162 | 1 | 0,00486 | 0 | 0 | 0 |
| MTRNR2L8 | 1 | 0 | 1 | 1 | 1,026 | 1 | 0,00162 | 1 | 0,00486 | 0 | 0 | 0 |
| CACFD1 | 1 | 0 | 1 | 1 | 1,022 | 1 | 0,09303 | 224,1814 | 5,35069 | 0 | 0 | 0 |
| NDRG2 | 1 | 0 | 1 | 1 | 1,058 | 1 | 0,11629 | 269,1036 | 6,13338 | 0 | 0 | 0 |
| C15orf52 | 1 | 0 | 1 | 1 | 1,046 | 1 | 0,10337 | 245,804 | 5,75704 | 0 | 0 | 0 |
| ZNF853 | 1 | 0 | 1 | 1 | 1,034 | 1 | 0,10337 | 238,2873 | 5,6297 | 0 | 0 | 0 |
| TMEM219 | 1 | 0 | 1 | 1 | 1,758 | 1 | 0,11629 | 314,8202 | 6,706 | 0 | 0 | 0 |
| NBPF10 | 1 | 0 | 1 | 1 | 1,247 | 1 | 0,11629 | 267,2536 | 6,09364 | 0 | 0 | 0 |
| TMEM100 | 1 | 0 | 1 | 1 | 1,428 | 1 | 0,11629 | 301,6631 | 6,5446 | 0 | 0 | 0 |
| MTRNR2L12 | 1 | 0 | 1 | 1 | 1,026 | 1 | 0,00162 | 1 | 0,00486 | 0 | 0 | 0 |
| MMP28 | 1 | 0 | 1 | 1 | 2,2 | 1 | 0,11629 | 339,7607 | 6,92824 | 0 | 0 | 0 |
| NBPF9 | 1 | 0 | 1 | 1 | 1,151 | 1 | 0,11629 | 267,2536 | 6,09364 | 0 | 0 | 0 |
| NBPF19 | 1 | 0 | 1 | 1 | 1,209 | 1 | 0,11629 | 267,2536 | 6,09364 | 0 | 0 | 0 |
| NBPF26 | 1 | 0 | 1 | 1 | 1,139 | 1 | 0,11629 | 267,2536 | 6,09364 | 0 | 0 | 0 |
| C11orf96 | 1 | 0 | 1 | 1 | 1,03 | 1 | 0,10337 | 237,8004 | 5,61186 | 0 | 0 | 0 |
| ZAN | 1 | 0 | 1 | 1 | 1,693 | 1 | 0,11629 | 303,7226 | 6,56406 | 0 | 0 | 0 |
| hCG_401294 | 1 | 0 | 1 | 1 | 1,029 | 1 | 0,10337 | 233,8885 | 5,53318 | 0 | 0 | 0 |
| PAMR1 | 1 | 0 | 1 | 1 | 1,048 | 1 | 0,10337 | 255,7671 | 5,92656 | 0 | 0 | 0 |
| C22orf31 | 0 | 0 | 0 | 0 | 1 | 0 | 0 | 0 | 0 | 0 | 0 | 0 |
| ELK3 | 0 | 0 | 0 | 0 | 1 | 0 | 0 | 0 | 0 | 0 | 0 | 0 |
| AKAP3 | 0 | 0 | 0 | 0 | 1 | 0 | 0 | 0 | 0 | 0 | 0 | 0 |
| NRM | 0 | 0 | 0 | 0 | 1 | 0 | 0 | 0 | 0 | 0 | 0 | 0 |
| MLPH | 0 | 0 | 0 | 0 | 1 | 0 | 0 | 0 | 0 | 0 | 0 | 0 |
| RAB20 | 0 | 0 | 0 | 0 | 1 | 0 | 0 | 0 | 0 | 0 | 0 | 0 |
| SFXN5 | 0 | 0 | 0 | 0 | 1 | 0 | 0 | 0 | 0 | 0 | 0 | 0 |
| C4orf19 | 0 | 0 | 0 | 0 | 1 | 0 | 0 | 0 | 0 | 0 | 0 | 0 |
| C10orf10 | 0 | 0 | 0 | 0 | 1 | 0 | 0 | 0 | 0 | 0 | 0 | 0 |
| TSC22D4 | 0 | 0 | 0 | 0 | 1 | 0 | 0 | 0 | 0 | 0 | 0 | 0 |
| ZRSR2 | 0 | 0 | 0 | 0 | 1 | 0 | 0 | 0 | 0 | 0 | 0 | 0 |
| SHE | 0 | 0 | 0 | 0 | 1 | 0 | 0 | 0 | 0 | 0 | 0 | 0 |
| PODN | 0 | 0 | 0 | 0 | 1 | 0 | 0 | 0 | 0 | 0 | 0 | 0 |
| ZNF680 | 0 | 0 | 0 | 0 | 1 | 0 | 0 | 0 | 0 | 0 | 0 | 0 |
| CA5A | 0 | 0 | 0 | 0 | 1 | 0 | 0 | 0 | 0 | 0 | 0 | 0 |
| TBC1D16 | 0 | 0 | 0 | 0 | 1 | 0 | 0 | 0 | 0 | 0 | 0 | 0 |
| MAP7D3 | 0 | 0 | 0 | 0 | 1 | 0 | 0 | 0 | 0 | 0 | 0 | 0 |

| | | | | | | | | | | | | |
|---|---|---|---|---|---|---|---|---|---|---|---|---|
| IGDCC4 | 0 | 0 | 0 | 0 | 1 | 0 | 0 | 0 | 0 | 0 | 0 | 0 |
| PXMP2 | 0 | 0 | 0 | 0 | 1 | 0 | 0 | 0 | 0 | 0 | 0 | 0 |
| C16orf89 | 0 | 0 | 0 | 0 | 1 | 0 | 0 | 0 | 0 | 0 | 0 | 0 |
| PLEKHO2 | 0 | 0 | 0 | 0 | 1 | 0 | 0 | 0 | 0 | 0 | 0 | 0 |
| TLCD2 | 0 | 0 | 0 | 0 | 1 | 0 | 0 | 0 | 0 | 0 | 0 | 0 |
| C1orf64 | 0 | 0 | 0 | 0 | 1 | 0 | 0 | 0 | 0 | 0 | 0 | 0 |
| OAF | 0 | 0 | 0 | 0 | 1 | 0 | 0 | 0 | 0 | 0 | 0 | 0 |
| FUT10 | 0 | 0 | 0 | 0 | 1 | 0 | 0 | 0 | 0 | 0 | 0 | 0 |
| C3orf70 | 0 | 0 | 0 | 0 | 1 | 0 | 0 | 0 | 0 | 0 | 0 | 0 |
| SPATA21 | 0 | 0 | 0 | 0 | 1 | 0 | 0 | 0 | 0 | 0 | 0 | 0 |
| LDLRAD2 | 0 | 0 | 0 | 0 | 1 | 0 | 0 | 0 | 0 | 0 | 0 | 0 |
| PODNL1 | 0 | 0 | 0 | 0 | 1 | 0 | 0 | 0 | 0 | 0 | 0 | 0 |
| HRCT1 | 0 | 0 | 0 | 0 | 1 | 0 | 0 | 0 | 0 | 0 | 0 | 0 |
| BTN3A2 | 0 | 0 | 0 | 0 | 1 | 0 | 0 | 0 | 0 | 0 | 0 | 0 |
| TOR4A | 0 | 0 | 0 | 0 | 1 | 0 | 0 | 0 | 0 | 0 | 0 | 0 |
| SIRPB2 | 0 | 0 | 0 | 0 | 1 | 0 | 0 | 0 | 0 | 0 | 0 | 0 |
| FAM63A | 0 | 0 | 0 | 0 | 1 | 0 | 0 | 0 | 0 | 0 | 0 | 0 |
| TBX19 | 0 | 0 | 0 | 0 | 1 | 0 | 0 | 0 | 0 | 0 | 0 | 0 |
| C1orf192 | 0 | 0 | 0 | 0 | 1 | 0 | 0 | 0 | 0 | 0 | 0 | 0 |
| TDRD10 | 0 | 0 | 0 | 0 | 1 | 0 | 0 | 0 | 0 | 0 | 0 | 0 |
| FBXW4 | 0 | 0 | 0 | 0 | 1 | 0 | 0 | 0 | 0 | 0 | 0 | 0 |
| AIF1L | 0 | 0 | 0 | 0 | 1 | 0 | 0 | 0 | 0 | 0 | 0 | 0 |
| C2orf72 | 0 | 0 | 0 | 0 | 1 | 0 | 0 | 0 | 0 | 0 | 0 | 0 |
| RAB42 | 0 | 0 | 0 | 0 | 1 | 0 | 0 | 0 | 0 | 0 | 0 | 0 |
| FAM110D | 0 | 0 | 0 | 0 | 1 | 0 | 0 | 0 | 0 | 0 | 0 | 0 |
| ZAK | 0 | 0 | 0 | 0 | 1 | 0 | 0 | 0 | 0 | 0 | 0 | 0 |
| RSG1 | 0 | 0 | 0 | 0 | 1 | 0 | 0 | 0 | 0 | 0 | 0 | 0 |
| ACBD7 | 0 | 0 | 0 | 0 | 1 | 0 | 0 | 0 | 0 | 0 | 0 | 0 |
| NAA16 | 0 | 0 | 0 | 0 | 1 | 0 | 0 | 0 | 0 | 0 | 0 | 0 |
| ACOT6 | 0 | 0 | 0 | 0 | 1 | 0 | 0 | 0 | 0 | 0 | 0 | 0 |
| C10orf54 | 0 | 0 | 0 | 0 | 1 | 0 | 0 | 0 | 0 | 0 | 0 | 0 |
| CARHSP1 | 0 | 0 | 0 | 0 | 1 | 0 | 0 | 0 | 0 | 0 | 0 | 0 |
| ENSP0000038202 | 0 | 0 | 0 | 0 | 1 | 0 | 0 | 0 | 0 | 0 | 0 | 0 |
| RMDN1 | 0 | 0 | 0 | 0 | 1 | 0 | 0 | 0 | 0 | 0 | 0 | 0 |
| APOL6 | 0 | 0 | 0 | 0 | 1 | 0 | 0 | 0 | 0 | 0 | 0 | 0 |
| SSFA2 | 0 | 0 | 0 | 0 | 1 | 0 | 0 | 0 | 0 | 0 | 0 | 0 |
| ENSP000003916 | 0 | 0 | 0 | 0 | 1 | 0 | 0 | 0 | 0 | 0 | 0 | 0 |
| PRR29 | 0 | 0 | 0 | 0 | 1 | 0 | 0 | 0 | 0 | 0 | 0 | 0 |
| C10orf105 | 0 | 0 | 0 | 0 | 1 | 0 | 0 | 0 | 0 | 0 | 0 | 0 |
| ZBTB42 | 0 | 0 | 0 | 0 | 1 | 0 | 0 | 0 | 0 | 0 | 0 | 0 |
| NKAIN3 | 0 | 0 | 0 | 0 | 1 | 0 | 0 | 0 | 0 | 0 | 0 | 0 |
| SLC43A3 | 0 | 0 | 0 | 0 | 1 | 0 | 0 | 0 | 0 | 0 | 0 | 0 |
| GRAPL | 0 | 0 | 0 | 0 | 1 | 0 | 0 | 0 | 0 | 0 | 0 | 0 |
| GOLGA8F | 0 | 0 | 0 | 0 | 1 | 0 | 0 | 0 | 0 | 0 | 0 | 0 |
| KLHL36 | 0 | 0 | 0 | 0 | 1 | 0 | 0 | 0 | 0 | 0 | 0 | 0 |
| KIAA0040 | 0 | 0 | 0 | 0 | 1 | 0 | 0 | 0 | 0 | 0 | 0 | 0 |
| ANKRD62 | 0 | 0 | 0 | 0 | 1 | 0 | 0 | 0 | 0 | 0 | 0 | 0 |
| AQP6 | 0 | 0 | 0 | 0 | 1 | 0 | 0 | 0 | 0 | 0 | 0 | 0 |
| PINLYP | 0 | 0 | 0 | 0 | 1 | 0 | 0 | 0 | 0 | 0 | 0 | 0 |
| GOLGA6L10 | 0 | 0 | 0 | 0 | 1 | 0 | 0 | 0 | 0 | 0 | 0 | 0 |
| ENSP000004808 | 0 | 0 | 0 | 0 | 1 | 0 | 0 | 0 | 0 | 0 | 0 | 0 |
| ENSP000004855 | 0 | 0 | 0 | 0 | 1 | 0 | 0 | 0 | 0 | 0 | 0 | 0 |

Supplementary file 6: Functional enrichment of the top 10 downregulated hub genes.

| # backgrou | # genes | category | description | FDR corrected p-value | genes | network.SUID | nodes.SUID | p-value | term name | transferred FDR value |
|---|---|---|---|---|---|---|---|---|---|---|
| 420 | 9 | COMPARTMENTS | Receptor complex | 2,94E-11 | STAT3\|ERE | 50406 | 4127\|4988\|5 | 1,05E-14 | GOCC:0043235 | 1,053165 |
| 675 | 8 | COMPARTMENTS | Plasma membrane protein complex | 1,25E-07 | STAT3\|ERE | 50406 | 4127\|4988\|5 | 8,92E-11 | GOCC:0098797 | 0,690309 |
| 782 | 8 | COMPARTMENTS | Integral component of plasma membrane | 2,65E-07 | STAT3\|EGI | 50406 | 4127\|5306\|3 | 2,85E-10 | GOCC:0005887 | 0,657675 |
| 841 | 8 | COMPARTMENTS | Intrinsic component of plasma membrane | 3,53E-07 | STAT3\|EGI | 50406 | 4127\|5306\|3 | 5,05E-10 | GOCC:0031226 | 0,645223 |
| 1456 | 9 | COMPARTMENTS | Integral component of membrane | 3,75E-07 | STAT3\|ERE | 50406 | 4127\|4988\|5 | 6,71E-10 | GOCC:0016021 | 0,642597 |
| 247 | 6 | COMPARTMENTS | Plasma membrane signaling receptor complex | 4,12E-07 | STAT3\|ERE | 50406 | 4127\|4988\|5 | 8,83E-10 | GOCC:0098802 | 0,63851 |
| 1552 | 9 | COMPARTMENTS | Intrinsic component of membrane | 4,73E-07 | STAT3\|ERE | 50406 | 4127\|4988\|5 | 1,18E-09 | GOCC:0031224 | 0,632514 |
| 1321 | 8 | COMPARTMENTS | Membrane protein complex | 6,15E-06 | STAT3\|ERE | 50406 | 4127\|4988\|5 | 1,76E-08 | GOCC:0098796 | 0,521112 |
| 3531 | 10 | COMPARTMENTS | Plasma membrane | 1,15E-05 | STAT3\|ERE | 50406 | 4127\|4988\|5 | 3,71E-08 | GOCC:0005886 | 0,49393 |
| 3688 | 10 | COMPARTMENTS | Cell periphery | 1,60E-05 | STAT3\|ERE | 50406 | 4127\|4988\|5 | 5,73E-08 | GOCC:0071944 | 0,479588 |
| 23 | 3 | COMPARTMENTS | NF-kappaB complex | 6,30E-05 | STAT3\|TLR | 50406 | 4127\|4079\|5 | 2,48E-07 | GOCC:0071159 | 0,420066 |
| 3080 | 9 | COMPARTMENTS | Endomembrane system | 1,20E-04 | ERBB2\|EGI | 50406 | 4988\|5306\|3 | 5,16E-07 | GOCC:0012505 | 0,392082 |
| 1416 | 7 | COMPARTMENTS | Bounding membrane of organelle | 2,20E-04 | ERBB2\|EGI | 50406 | 4988\|5306\|3 | 1,04E-06 | GOCC:0098588 | 0,365758 |
| 5142 | 10 | COMPARTMENTS | Protein-containing complex | 3,20E-04 | STAT3\|ERE | 50406 | 4127\|4988\|5 | 1,58E-06 | GOCC:0032991 | 0,349485 |
| 985 | 6 | COMPARTMENTS | Extracellular space | 5,40E-04 | EGFR\|FN1 | 50406 | 5306\|5576\|5 | 2,92E-06 | GOCC:0005615 | 0,326761 |
| 4 | 2 | COMPARTMENTS | interleukin-6 receptor complex | 6,10E-04 | STAT3\|IL6 | 50406 | 4127\|5153 | 3,52E-06 | GOCC:0005896 | 0,321467 |
| 5657 | 10 | COMPARTMENTS | Membrane | 6,10E-04 | STAT3\|ERE | 50406 | 4127\|4988\|5 | 4,11E-06 | GOCC:0016020 | 0,321467 |
| 1709 | 7 | COMPARTMENTS | Cytoplasmic vesicle | 6,10E-04 | ERBB2\|EGI | 50406 | 4988\|5306\|5 | 3,72E-06 | GOCC:0031410 | 0,321467 |
| 4 | 2 | COMPARTMENTS | Integrin alpha5-beta1 complex | 6,10E-04 | FN1\|ITGB1 | 50406 | 5576\|5270 | 3,52E-06 | GOCC:0034674 | 0,321467 |
| 540 | 5 | COMPARTMENTS | Anchoring junction | 6,10E-04 | EGFR\|NOT | 50406 | 5306\|3485\|5 | 3,69E-06 | GOCC:0070161 | 0,321467 |
| 1711 | 7 | COMPARTMENTS | Intracellular vesicle | 6,10E-04 | ERBB2\|EGI | 50406 | 4988\|5306\|5 | 3,75E-06 | GOCC:0097708 | 0,321467 |
| 7 | 2 | COMPARTMENTS | Integrin alpha4-beta1 complex | 0,0011 | FN1\|ITGB1 | 50406 | 5576\|5270 | 8,44E-06 | GOCC:0034668 | 0,295861 |
| 1219 | 6 | COMPARTMENTS | Whole membrane | 0,0012 | ERBB2\|EGI | 50406 | 4988\|5306\|4 | 1,00E-05 | GOCC:0098805 | 0,292082 |
| 2035 | 7 | COMPARTMENTS | Extracellular region | 0,0013 | EGFR\|NOT | 50406 | 5306\|3485\|5 | 1,20E-05 | GOCC:0005576 | 0,288606 |
| 676 | 5 | COMPARTMENTS | Plasma membrane region | 0,0013 | ERBB2\|EGI | 50406 | 4988\|5306\|3 | 1,09E-05 | GOCC:0098590 | 0,288606 |
| 2051 | 7 | COMPARTMENTS | Vesicle | 0,0014 | ERBB2\|EGI | 50406 | 4988\|5306\|5 | 1,27E-05 | GOCC:0031982 | 0,285387 |
| 322 | 4 | COMPARTMENTS | Endosome membrane | 0,0015 | ERBB2\|EGI | 50406 | 4988\|5306\|4 | 1,46E-05 | GOCC:0010008 | 0,282391 |
| 2219 | 7 | COMPARTMENTS | Organelle membrane | 0,0021 | ERBB2\|EGI | 50406 | 4988\|5306\|3 | 2,14E-05 | GOCC:0031090 | 0,267778 |
| 12 | 2 | COMPARTMENTS | Spanning component of plasma membrane | 0,0021 | EGFR\|NOT | 50406 | 5306\|3485 | 2,13E-05 | GOCC:0044214 | 0,267778 |
| 368 | 4 | COMPARTMENTS | Extracellular exosome | 0,0023 | FN1\|ITGB1 | 50406 | 5576\|5270\|3 | 2,46E-05 | GOCC:0070062 | 0,263827 |
| 16 | 2 | COMPARTMENTS | Spanning component of membrane | 0,0032 | EGFR\|NOT | 50406 | 5306\|3485 | 3,58E-05 | GOCC:0089717 | 0,249485 |
| 433 | 4 | COMPARTMENTS | Cell surface | 0,004 | EGFR\|TLR4 | 50406 | 5306\|4079\|5 | 4,62E-05 | GOCC:0009986 | 0,239794 |
| 444 | 4 | COMPARTMENTS | Extracellular vesicle | 0,0043 | FN1\|ITGB1 | 50406 | 5576\|5270\|3 | 5,09E-05 | GOCC:1903561 | 0,236653 |
| 451 | 4 | COMPARTMENTS | Extracellular organelle | 0,0045 | FN1\|ITGB1 | 50406 | 5576\|5270\|3 | 5,41E-05 | GOCC:0043230 | 0,234679 |
| 26 | 2 | COMPARTMENTS | Cell wall | 0,007 | TLR4\|IL6 | 50406 | 4079\|5153 | 8,82E-05 | GOCC:0005618 | 0,21549 |
| 1045 | 5 | COMPARTMENTS | Cell junction | 0,007 | EGFR\|NOT | 50406 | 5306\|3485\|5 | 8,84E-05 | GOCC:0030054 | 0,21549 |
| 7871 | 10 | COMPARTMENTS | Cytoplasm | 0,0084 | STAT3\|ERE | 50406 | 4127\|4988\|5 | 1,10E-04 | GOCC:0005737 | 0,207572 |
| 656 | 4 | COMPARTMENTS | Endosome | 0,0168 | ERBB2\|EGI | 50406 | 4988\|5306\|4 | 2,30E-04 | GOCC:0005768 | 0,177469 |
| 255 | 3 | COMPARTMENTS | Focal adhesion | 0,0182 | EGFR\|ITGE | 50406 | 5306\|5270\|3 | 2,50E-04 | GOCC:0005925 | 0,173993 |
| 262 | 3 | COMPARTMENTS | Cell-substrate junction | 0,0192 | EGFR\|ITGE | 50406 | 5306\|5270\|3 | 2,70E-04 | GOCC:0030055 | 0,17167 |
| 8685 | 10 | COMPARTMENTS | Membrane-bounded organelle | 0,0203 | STAT3\|ERE | 50406 | 4127\|4988\|5 | 3,00E-04 | GOCC:0043227 | 0,16925 |
| 50 | 2 | COMPARTMENTS | Integrin complex | 0,0205 | FN1\|ITGB1 | 50406 | 5576\|5270 | 3,10E-04 | GOCC:0008305 | 0,168825 |
| 61 | 2 | COMPARTMENTS | External encapsulating structure | 0,0293 | TLR4\|IL6 | 50406 | 4079\|5153 | 4,50E-04 | GOCC:0030312 | 0,153313 |
| 9242 | 10 | COMPARTMENTS | Intracellular organelle | 0,0352 | STAT3\|ERE | 50406 | 4127\|4988\|5 | 5,50E-04 | GOCC:0043229 | 0,145346 |
| 73 | 2 | COMPARTMENTS | Protein complex involved in cell adhesion | 0,0397 | FN1\|ITGB1 | 50406 | 5576\|5270 | 6,40E-04 | GOCC:0098636 | 0,140121 |
| 510 | 6 | DISEASES | Gastrointestinal system disease | 2,70E-04 | STAT3\|ERE | 50406 | 4127\|4988\|5 | 6,25E-08 | DOID:77 | 0,356864 |
| 677 | 6 | DISEASES | Organ system cancer | 7,20E-04 | STAT3\|ERE | 50406 | 4127\|4988\|5 | 3,29E-07 | DOID:0050686 | 0,314267 |
| 2 | 2 | DISEASES | Bacterial sepsis | 0,0021 | TLR4\|IL6 | 50406 | 4079\|5153 | 1,41E-06 | DOID:0040085 | 0,267778 |
| 3 | 2 | DISEASES | Perinatal necrotizing enterocolitis | 0,0021 | TLR4\|IL6 | 50406 | 4079\|5153 | 2,35E-06 | DOID:8677 | 0,267778 |
| 58 | 3 | DISEASES | Breast cancer | 0,0025 | ERBB2\|EGI | 50406 | 4988\|5306\|5 | 3,40E-06 | DOID:1612 | 0,260206 |
| 611 | 5 | DISEASES | Immune system disease | 0,0033 | STAT3\|NO | 50406 | 4127\|3485\|5 | 6,71E-06 | DOID:2914 | 0,248149 |
| 10 | 2 | DISEASES | Head and neck squamous cell carcinoma | 0,0056 | EGFR\|NOT | 50406 | 5306\|3485 | 1,55E-05 | DOID:5520 | 0,225181 |
| 2132 | 7 | DISEASES | Nervous system disease | 0,0056 | STAT3\|ERE | 50406 | 4127\|4988\|5 | 1,64E-05 | DOID:863 | 0,225181 |
| 14 | 2 | DISEASES | In situ carcinoma | 0,0077 | ERBB2\|EGI | 50406 | 4988\|5306 | 2,81E-05 | DOID:8719 | 0,211351 |
| 406 | 4 | DISEASES | Cell type cancer | 0,0093 | ERBB2\|EGI | 50406 | 4988\|5306\|3 | 3,60E-05 | DOID:0050687 | 0,203152 |
| 19 | 2 | DISEASES | Lung non-small cell carcinoma | 0,0113 | ERBB2\|EGI | 50406 | 4988\|5306 | 4,91E-05 | DOID:3908 | 0,194692 |
| 22 | 2 | DISEASES | Colitis | 0,0141 | STAT3\|IL6 | 50406 | 4127\|5153 | 6,44E-05 | DOID:0060180 | 0,185078 |
| 23 | 2 | DISEASES | Ischemia | 0,0146 | IL6\|PECAN | 50406 | 5153\|5090 | 7,00E-05 | DOID:326 | 0,183565 |
| 172 | 3 | DISEASES | Lung disease | 0,016 | ERBB2\|EGI | 50406 | 4988\|5306\|5 | 8,05E-05 | DOID:850 | 0,179588 |
| 27 | 2 | DISEASES | Stomach cancer | 0,0177 | ERBB2\|EGI | 50406 | 4988\|5306 | 9,47E-05 | DOID:10534 | 0,175203 |
| 181 | 3 | DISEASES | Intestinal disease | 0,0177 | STAT3\|EGI | 50406 | 4127\|5306\|5 | 9,34E-05 | DOID:5295 | 0,175203 |
| 223 | 3 | DISEASES | Vascular disease | 0,0278 | NOTCH1\|II | 50406 | 3485\|5153\|5 | 1,70E-04 | DOID:178 | 0,155596 |
| 4452 | 8 | DISEASES | Disease of anatomical entity | 0,0323 | STAT3\|ERE | 50406 | 4127\|4988\|5 | 2,10E-04 | DOID:7 | 0,14908 |
| 275 | 3 | DISEASES | Carcinoma | 0,0431 | ERBB2\|EGI | 50406 | 4988\|5306\|3 | 3,20E-04 | DOID:305 | 0,136552 |
| 1459 | 9 | GO Biological Proces | Regulation of protein phosphorylation | 1,00E-05 | STAT3\|ERE | 50406 | 4127\|4988\|5 | 6,84E-10 | GO:0001932 | 0,5 |
| 1019 | 8 | GO Biological Proces | Positive regulation of protein phosphorylation | 1,00E-05 | STAT3\|ERE | 50406 | 4127\|4988\|5 | 2,29E-09 | GO:0001934 | 0,5 |
| 1296 | 8 | GO Biological Proces | Defense response | 1,00E-05 | STAT3\|EGI | 50406 | 4127\|5306\|3 | 1,51E-08 | GO:0006952 | 0,5 |
| 515 | 7 | GO Biological Proces | Inflammatory response | 1,00E-05 | STAT3\|EGI | 50406 | 4127\|5306\|3 | 1,03E-09 | GO:0006954 | 0,5 |
| 2325 | 10 | GO Biological Proces | Cell surface receptor signaling pathway | 1,00E-05 | STAT3\|ERE | 50406 | 4127\|4988\|5 | 5,73E-10 | GO:0007166 | 0,5 |
| 919 | 8 | GO Biological Proces | Positive regulation of cell population proliferation | 1,00E-05 | STAT3\|ERE | 50406 | 4127\|4988\|5 | 1,02E-09 | GO:0008284 | 0,5 |
| 3107 | 10 | GO Biological Proces | Regulation of signal transduction | 1,00E-05 | STAT3\|ERE | 50406 | 4127\|4988\|5 | 1,03E-08 | GO:0009966 | 0,5 |
| 1654 | 9 | GO Biological Proces | Positive regulation of signal transduction | 1,00E-05 | STAT3\|ERE | 50406 | 4127\|4988\|5 | 2,09E-09 | GO:0009967 | 0,5 |
| 178 | 5 | GO Biological Proces | Glial cell differentiation | 1,00E-05 | STAT3\|ERE | 50406 | 4127\|4988\|5 | 1,64E-08 | GO:0010001 | 0,5 |
| 522 | 7 | GO Biological Proces | Positive regulation of cell migration | 1,00E-05 | STAT3\|EGI | 50406 | 4127\|5306\|3 | 1,13E-09 | GO:0030335 | 0,5 |
| 1635 | 9 | GO Biological Proces | Positive regulation of cellular protein metabolic pro | 1,00E-05 | STAT3\|ERE | 50406 | 4127\|4988\|5 | 1,88E-09 | GO:0032270 | 0,5 |
| 1251 | 8 | GO Biological Proces | Locomotion | 1,00E-05 | ERBB2\|EGI | 50406 | 4988\|5306\|3 | 1,15E-08 | GO:0040011 | 0,5 |
| 725 | 7 | GO Biological Proces | Regulation of mapk cascade | 1,00E-05 | ERBB2\|EGI | 50406 | 4988\|5306\|3 | 1,08E-08 | GO:0043408 | 0,5 |
| 728 | 7 | GO Biological Proces | Negative regulation of cell differentiation | 1,00E-05 | STAT3\|ERE | 50406 | 4127\|4988\|5 | 1,11E-08 | GO:0045596 | 0,5 |
| 55 | 4 | GO Biological Proces | Astrocyte differentiation | 1,00E-05 | STAT3\|EGI | 50406 | 4127\|5306\|3 | 1,54E-08 | GO:0048708 | 0,5 |
| 352 | 6 | GO Biological Proces | Multicellular organismal homeostasis | 1,00E-05 | STAT3\|NO | 50406 | 4127\|3485\|4 | 7,08E-09 | GO:0048871 | 0,5 |
| 292 | 6 | GO Biological Proces | Regulation of erk1 and erk2 cascade | 1,00E-05 | ERBB2\|EGI | 50406 | 4988\|5306\|3 | 2,36E-09 | GO:0070372 | 0,5 |
| 929 | 8 | GO Biological Proces | Regulation of cell motility | 1,00E-05 | STAT3\|ERE | 50406 | 4127\|4988\|5 | 1,11E-09 | GO:2000145 | 0,5 |
| 192 | 5 | GO Biological Proces | Positive regulation of epithelial cell proliferation | 1,19E-05 | STAT3\|ERE | 50406 | 4127\|4988\|5 | 2,37E-08 | GO:0050679 | 0,492445 |
| 439 | 6 | GO Biological Proces | Wound healing | 1,26E-05 | ERBB2\|EGI | 50406 | 4988\|5306\|5 | 2,59E-08 | GO:0042060 | 0,489963 |
| 3485 | 10 | GO Biological Proces | Response to stress | 1,54E-05 | STAT3\|ERE | 50406 | 4127\|4988\|5 | 3,26E-08 | GO:0006950 | 0,481248 |
| 1447 | 8 | GO Biological Proces | Response to endogenous stimulus | 1,59E-05 | STAT3\|ERE | 50406 | 4127\|4988\|5 | 3,59E-08 | GO:0009719 | 0,47986 |
| 219 | 5 | GO Biological Proces | Tissue homeostasis | 1,84E-05 | NOTCH1\|T | 50406 | 3485\|4079\|5 | 4,51E-08 | GO:0001894 | 0,473518 |
| 896 | 7 | GO Biological Proces | Cell migration | 1,84E-05 | EGFR\|NOT | 50406 | 5306\|3485\|5 | 4,61E-08 | GO:0016477 | 0,473518 |
| 1501 | 8 | GO Biological Proces | Movement of cell or subcellular component | 1,86E-05 | ERBB2\|EGI | 50406 | 4988\|5306\|3 | 4,79E-08 | GO:0006928 | 0,473049 |
| 2369 | 9 | GO Biological Proces | Cellular response to organic substance | 1,93E-05 | STAT3\|ERE | 50406 | 4127\|4988\|5 | 5,06E-08 | GO:0071310 | 0,471444 |
| 85 | 4 | GO Biological Proces | Positive regulation of receptor signaling pathway via | 2,93E-05 | STAT3\|NO | 50406 | 4127\|3485\|5 | 8,21E-08 | GO:0046427 | 0,453313 |
| 543 | 6 | GO Biological Proces | Positive regulation of mapk cascade | 3,10E-05 | ERBB2\|EGI | 50406 | 4988\|5306\|3 | 9,04E-08 | GO:0043410 | 0,450864 |
| 1629 | 8 | GO Biological Proces | Cell development | 3,10E-05 | ERBB2\|EGI | 50406 | 4988\|5306\|3 | 9,08E-08 | GO:0048468 | 0,450864 |
| 999 | 7 | GO Biological Proces | Negative regulation of cell death | 3,13E-05 | STAT3\|EGI | 50406 | 4127\|5306\|3 | 9,71E-08 | GO:0060548 | 0,450446 |
| 258 | 5 | GO Biological Proces | Regulation of peptidyl-tyrosine phosphorylation | 3,14E-05 | STAT3\|EGI | 50406 | 4127\|5306\|5 | 1,01E-07 | GO:0050730 | 0,450307 |

| | | | | | | | | | | |
|---|---|---|---|---|---|---|---|---|---|---|
| 1657 | 8 | GO Biological Proces | Neurogenesis | 3,19E-05 | STAT3\|ERE | 50406 | 4127\|4988\|5 | 1,04E-07 | GO:0022008 | 0,449621 |
| 1696 | 8 | GO Biological Proces | Regulation of cell death | 3,63E-05 | STAT3\|EGI | 50406 | 4127\|5306\|3 | 1,24E-07 | GO:0010941 | 0,444009 |
| 1041 | 7 | GO Biological Proces | Positive regulation of intracellular signal transductic | 3,69E-05 | ERBB2\|EGI | 50406 | 4988\|5306\|3 | 1,29E-07 | GO:1902533 | 0,443297 |
| 2648 | 9 | GO Biological Proces | Regulation of developmental process | 3,83E-05 | STAT3\|ERE | 50406 | 4127\|4988\|5 | 1,36E-07 | GO:0050793 | 0,44168 |
| 1075 | 7 | GO Biological Proces | Cell activation | 4,38E-05 | STAT3\|EGI | 50406 | 4127\|5306\|4 | 1,60E-07 | GO:0001775 | 0,435853 |
| 1095 | 7 | GO Biological Proces | Regulation of anatomical structure morphogenesis | 4,82E-05 | STAT3\|ERE | 50406 | 4127\|4988\|3 | 1,82E-07 | GO:0022603 | 0,431695 |
| 2740 | 9 | GO Biological Proces | Regulation of localization | 4,82E-05 | STAT3\|ERE | 50406 | 4127\|4988\|5 | 1,84E-07 | GO:0032879 | 0,431695 |
| 21 | 3 | GO Biological Proces | Positive regulation of cytokine production involved i | 4,97E-05 | STAT3\|TLR | 50406 | 4127\|4079\|5 | 1,93E-07 | GO:1900017 | 0,430364 |
| 1807 | 8 | GO Biological Proces | Regulation of intracellular signal transduction | 5,18E-05 | ERBB2\|EGI | 50406 | 4988\|5306\|3 | 2,04E-07 | GO:1902531 | 0,428567 |
| 303 | 5 | GO Biological Proces | Regulation of angiogenesis | 5,52E-05 | STAT3\|ERE | 50406 | 4127\|4988\|3 | 2,20E-07 | GO:0045765 | 0,425806 |
| 23 | 3 | GO Biological Proces | Regulation of production of mirnas involved in gene | 6,03E-05 | STAT3\|EGI | 50406 | 4127\|5306\|5 | 2,48E-07 | GO:1903798 | 0,421968 |
| 1874 | 8 | GO Biological Proces | Regulation of cell differentiation | 6,49E-05 | STAT3\|ERE | 50406 | 4127\|4988\|5 | 2,71E-07 | GO:0045595 | 0,418776 |
| 316 | 5 | GO Biological Proces | Leukocyte migration | 6,49E-05 | FN1\|ITGB1 | 50406 | 5576\|5270\|5 | 2,71E-07 | GO:0050900 | 0,418776 |
| 24 | 3 | GO Biological Proces | Positive regulation of gene silencing by mirna | 6,51E-05 | STAT3\|EGI | 50406 | 4127\|5306\|5 | 2,79E-07 | GO:2000637 | 0,418642 |
| 1899 | 8 | GO Biological Proces | Interspecies interaction between organisms | 6,84E-05 | STAT3\|EGI | 50406 | 4127\|5306\|3 | 3,01E-07 | GO:0044419 | 0,416494 |
| 4426 | 10 | GO Biological Proces | System development | 7,75E-05 | STAT3\|ERE | 50406 | 4127\|4988\|5 | 3,54E-07 | GO:0048731 | 0,41107 |
| 127 | 4 | GO Biological Proces | Cell-matrix adhesion | 8,22E-05 | FN1\|ITGB1 | 50406 | 5576\|5270\|3 | 3,90E-07 | GO:0007160 | 0,408513 |
| 1231 | 7 | GO Biological Proces | Negative regulation of multicellular organismal proc | 8,40E-05 | STAT3\|ERE | 50406 | 4127\|4988\|5 | 4,03E-07 | GO:0051241 | 0,407572 |
| 726 | 6 | GO Biological Proces | Cell morphogenesis | 9,95E-05 | ERBB2\|EGI | 50406 | 4988\|5306\|3 | 4,95E-07 | GO:0000902 | 0,400218 |
| 1271 | 7 | GO Biological Proces | Negative regulation of signal transduction | 9,97E-05 | ERBB2\|EGI | 50406 | 4988\|5306\|3 | 5,01E-07 | GO:0009968 | 0,40013 |
| 144 | 4 | GO Biological Proces | Positive regulation of inflammatory response | 1,20E-04 | STAT3\|EGI | 50406 | 4127\|5306\|4 | 6,36E-07 | GO:0050729 | 0,392082 |
| 2096 | 8 | GO Biological Proces | Regulation of multicellular organismal development | 1,20E-04 | STAT3\|ERE | 50406 | 4127\|4988\|5 | 6,48E-07 | GO:2000026 | 0,392082 |
| 3197 | 9 | GO Biological Proces | Animal organ development | 1,30E-04 | STAT3\|ERE | 50406 | 4127\|4988\|5 | 7,16E-07 | GO:0048513 | 0,388606 |
| 34 | 3 | GO Biological Proces | Maintenance of blood-brain barrier | 1,40E-04 | ITGB1\|IL6\| | 50406 | 5270\|5153\|5 | 7,39E-07 | GO:0035633 | 0,385387 |
| 390 | 5 | GO Biological Proces | Lymphocyte activation | 1,40E-04 | STAT3\|TLR | 50406 | 4127\|4079\|5 | 7,56E-07 | GO:0046649 | 0,385387 |
| 2165 | 8 | GO Biological Proces | Anatomical structure morphogenesis | 1,50E-04 | STAT3\|ERE | 50406 | 4127\|4988\|5 | 8,34E-07 | GO:0009653 | 0,382391 |
| 44 | 3 | GO Biological Proces | Acute-phase response | 2,60E-04 | STAT3\|FN1 | 50406 | 4127\|5576\|5 | 1,54E-06 | GO:0006953 | 0,358503 |
| 893 | 6 | GO Biological Proces | Negative regulation of apoptotic process | 2,80E-04 | EGFR\|NOT | 50406 | 5306\|3485\|5 | 1,65E-06 | GO:0043066 | 0,355284 |
| 2386 | 8 | GO Biological Proces | Regulation of catalytic activity | 2,90E-04 | STAT3\|ERE | 50406 | 4127\|4988\|5 | 1,77E-06 | GO:0050790 | 0,35376 |
| 1551 | 7 | GO Biological Proces | Generation of neurons | 3,00E-04 | STAT3\|ERE | 50406 | 4127\|4988\|5 | 1,93E-06 | GO:0048699 | 0,352288 |
| 929 | 6 | GO Biological Proces | Leukocyte activation | 3,20E-04 | STAT3\|TLR | 50406 | 4127\|4079\|5 | 2,08E-06 | GO:0045321 | 0,349485 |
| 196 | 4 | GO Biological Proces | Positive regulation of peptidyl-tyrosine phosphoryla | 3,20E-04 | STAT3\|IL6 | 50406 | 4127\|5153\|3 | 2,12E-06 | GO:0050731 | 0,349485 |
| 3 | 2 | GO Biological Proces | Negative regulation of primary mirna processing | 3,50E-04 | STAT3\|IL6 | 50406 | 4127\|5153 | 2,35E-06 | GO:2000635 | 0,345593 |
| 2481 | 8 | GO Biological Proces | Immune system process | 3,60E-04 | STAT3\|NO | 50406 | 4127\|3485\|5 | 2,40E-06 | GO:0002376 | 0,34437 |
| 3702 | 9 | GO Biological Proces | Cell differentiation | 3,80E-04 | STAT3\|ERE | 50406 | 4127\|4988\|5 | 2,60E-06 | GO:0030154 | 0,342022 |
| 209 | 4 | GO Biological Proces | Positive regulation of erk1 and erk2 cascade | 3,90E-04 | EGFR\|NOT | 50406 | 5306\|3485\|4 | 2,72E-06 | GO:0070374 | 0,340894 |
| 988 | 6 | GO Biological Proces | Central nervous system development | 4,20E-04 | STAT3\|ERE | 50406 | 4127\|4988\|5 | 2,97E-06 | GO:0007417 | 0,337675 |
| 987 | 6 | GO Biological Proces | Response to organonitrogen compound | 4,20E-04 | STAT3\|EGI | 50406 | 4127\|5306\|3 | 2,96E-06 | GO:0010243 | 0,337675 |
| 56 | 3 | GO Biological Proces | Leukocyte cell-cell adhesion | 4,30E-04 | ITGB1\|CD4 | 50406 | 5270\|3263\|5 | 3,07E-06 | GO:0007159 | 0,336653 |
| 1676 | 7 | GO Biological Proces | Homeostatic process | 4,50E-04 | STAT3\|EGI | 50406 | 4127\|5306\|3 | 3,26E-06 | GO:0042592 | 0,334679 |
| 1013 | 6 | GO Biological Proces | Cellular response to cytokine stimulus | 4,70E-04 | STAT3\|FN1 | 50406 | 4127\|5576\|4 | 3,44E-06 | GO:0071345 | 0,33279 |
| 4 | 2 | GO Biological Proces | Calcium-independent cell-matrix adhesion | 4,80E-04 | FN1\|ITGB1 | 50406 | 5576\|5270 | 3,52E-06 | GO:0007161 | 0,331876 |
| 1019 | 6 | GO Biological Proces | Neuron differentiation | 4,80E-04 | STAT3\|ERE | 50406 | 4127\|4988\|5 | 3,56E-06 | GO:0030182 | 0,331876 |
| 5579 | 10 | GO Biological Proces | Positive regulation of cellular process | 4,80E-04 | STAT3\|ERE | 50406 | 4127\|4988\|5 | 3,58E-06 | GO:0048522 | 0,331876 |
| 566 | 5 | GO Biological Proces | Cell morphogenesis involved in differentiation | 6,10E-04 | ERBB2\|NO | 50406 | 4988\|3485\|5 | 4,63E-06 | GO:0000904 | 0,321467 |
| 1770 | 7 | GO Biological Proces | Positive regulation of multicellular organismal proce | 6,20E-04 | STAT3\|EGI | 50406 | 4127\|5306\|3 | 4,71E-06 | GO:0051240 | 0,320761 |
| 248 | 4 | GO Biological Proces | Cell fate commitment | 6,80E-04 | STAT3\|NO | 50406 | 4127\|3485\|5 | 5,30E-06 | GO:0045165 | 0,316749 |
| 68 | 3 | GO Biological Proces | Positive regulation of tyrosine phosphorylation of st | 6,90E-04 | STAT3\|IL6 | 50406 | 4127\|5153\|5 | 5,39E-06 | GO:0042531 | 0,316115 |
| 1842 | 7 | GO Biological Proces | Positive regulation of molecular function | 7,70E-04 | STAT3\|ERE | 50406 | 4127\|4988\|5 | 6,16E-06 | GO:0044093 | 0,311351 |
| 6 | 2 | GO Biological Proces | T-helper 17 cell lineage commitment | 8,20E-04 | STAT3\|IL6 | 50406 | 4127\|5153 | 6,57E-06 | GO:0072540 | 0,308619 |
| 267 | 4 | GO Biological Proces | Positive regulation of growth | 8,60E-04 | ERBB2\|EGI | 50406 | 4988\|5306\|3 | 7,07E-06 | GO:0045927 | 0,30655 |
| 626 | 5 | GO Biological Proces | Leukocyte activation involved in immune response | 9,20E-04 | STAT3\|TLR | 50406 | 4127\|4079\|5 | 7,55E-06 | GO:0002366 | 0,303621 |
| 1181 | 6 | GO Biological Proces | Cellular response to endogenous stimulus | 0,001 | STAT3\|ERE | 50406 | 4127\|4988\|5 | 8,34E-06 | GO:0071495 | 0,3 |
| 656 | 5 | GO Biological Proces | Tube morphogenesis | 0,0011 | EGFR\|NOT | 50406 | 5306\|3485\|5 | 9,46E-06 | GO:0035239 | 0,295861 |
| 678 | 5 | GO Biological Proces | Cytokine-mediated signaling pathway | 0,0013 | STAT3\|FN1 | 50406 | 4127\|5576\|5 | 1,11E-05 | GO:0019221 | 0,288606 |
| 680 | 5 | GO Biological Proces | Neuron projection development | 0,0013 | ERBB2\|EGI | 50406 | 4988\|5306\|3 | 1,13E-05 | GO:0031175 | 0,288606 |
| 676 | 5 | GO Biological Proces | Regulation of growth | 0,0013 | STAT3\|ERE | 50406 | 4127\|4988\|5 | 1,09E-05 | GO:0040008 | 0,288606 |
| 315 | 4 | GO Biological Proces | Angiogenesis | 0,0015 | NOTCH1\|F | 50406 | 3485\|5576\|5 | 1,34E-05 | GO:0001525 | 0,282391 |
| 712 | 5 | GO Biological Proces | Regulation of cell adhesion | 0,0015 | ERBB2\|NO | 50406 | 4988\|3485\|5 | 1,41E-05 | GO:0030155 | 0,282391 |
| 94 | 3 | GO Biological Proces | Positive regulation of interleukin-6 production | 0,0015 | STAT3\|TLR | 50406 | 4127\|4079\|5 | 1,38E-05 | GO:0032755 | 0,282391 |
| 1284 | 6 | GO Biological Proces | Regulation of hydrolase activity | 0,0015 | STAT3\|ERE | 50406 | 4127\|4988\|5 | 1,35E-05 | GO:0051336 | 0,282391 |
| 103 | 3 | GO Biological Proces | Regulation of notch signaling pathway | 0,0019 | STAT3\|EGI | 50406 | 4127\|5306\|3 | 1,80E-05 | GO:0008593 | 0,272125 |
| 338 | 4 | GO Biological Proces | Extracellular matrix organization | 0,0019 | FN1\|ITGB1 | 50406 | 5576\|5270\|3 | 1,77E-05 | GO:0030198 | 0,272125 |
| 11 | 2 | GO Biological Proces | Positive regulation of production of mirnas involved | 0,0019 | EGFR\|IL6 | 50406 | 5306\|5153 | 1,83E-05 | GO:1903800 | 0,272125 |
| 352 | 4 | GO Biological Proces | Positive regulation of protein transport | 0,0022 | ERBB2\|EGI | 50406 | 4988\|5306\|4 | 2,07E-05 | GO:0051222 | 0,265758 |
| 772 | 5 | GO Biological Proces | Positive regulation of hydrolase activity | 0,0022 | STAT3\|ERE | 50406 | 4127\|4988\|5 | 2,08E-05 | GO:0051345 | 0,265758 |
| 112 | 3 | GO Biological Proces | Lymphocyte activation involved in immune respons | 0,0024 | STAT3\|TLR | 50406 | 4127\|4079\|5 | 2,30E-05 | GO:0002285 | 0,261979 |
| 112 | 3 | GO Biological Proces | Positive regulation of peptidyl-serine phosphorylatic | 0,0024 | EGFR\|IL6\| | 50406 | 5306\|5153\|3 | 2,30E-05 | GO:0033138 | 0,261979 |
| 1437 | 6 | GO Biological Proces | Regulation of response to stress | 0,0026 | STAT3\|EGI | 50406 | 4127\|5306\|4 | 2,57E-05 | GO:0080134 | 0,258503 |
| 4874 | 9 | GO Biological Proces | Negative regulation of cellular process | 0,0028 | STAT3\|ERE | 50406 | 4127\|4988\|5 | 2,88E-05 | GO:0048523 | 0,255284 |
| 2337 | 7 | GO Biological Proces | Positive regulation of gene expression | 0,0029 | STAT3\|ERE | 50406 | 4127\|4988\|5 | 3,02E-05 | GO:0010628 | 0,25376 |
| 4913 | 9 | GO Biological Proces | Regulation of molecular function | 0,0029 | STAT3\|ERE | 50406 | 4127\|4988\|5 | 3,09E-05 | GO:0065009 | 0,25376 |
| 6948 | 10 | GO Biological Proces | Regulation of metabolic process | 0,003 | STAT3\|ERE | 50406 | 4127\|4988\|5 | 3,20E-05 | GO:0019222 | 0,252288 |
| 1489 | 6 | GO Biological Proces | Positive regulation of catalytic activity | 0,003 | STAT3\|ERE | 50406 | 4127\|4988\|5 | 3,15E-05 | GO:0043085 | 0,252288 |
| 1514 | 6 | GO Biological Proces | Regulation of immune system process | 0,0032 | STAT3\|ERE | 50406 | 4127\|4988\|3 | 3,47E-05 | GO:0002682 | 0,249485 |
| 858 | 5 | GO Biological Proces | Response to lipid | 0,0032 | STAT3\|EGI | 50406 | 4127\|5306\|3 | 3,45E-05 | GO:0033993 | 0,249485 |
| 16 | 2 | GO Biological Proces | interleukin-6-mediated signaling pathway | 0,0032 | STAT3\|IL6 | 50406 | 4127\|5153 | 3,58E-05 | GO:0070102 | 0,249485 |
| 129 | 3 | GO Biological Proces | Positive regulation of protein localization to membr | 0,0032 | ERBB2\|EGI | 50406 | 4988\|5306\|5 | 3,47E-05 | GO:1905477 | 0,249485 |
| 2402 | 7 | GO Biological Proces | Regulation of cellular component organization | 0,0033 | ERBB2\|EGI | 50406 | 4988\|5306\|3 | 3,62E-05 | GO:0051128 | 0,248149 |
| 872 | 5 | GO Biological Proces | Circulatory system development | 0,0033 | ERBB2\|NO | 50406 | 4988\|3485\|5 | 3,72E-05 | GO:0072359 | 0,248149 |
| 424 | 4 | GO Biological Proces | Regulation of cell-cell adhesion | 0,0037 | ERBB2\|NO | 50406 | 4988\|3485\|5 | 4,26E-05 | GO:0022407 | 0,24318 |
| 423 | 4 | GO Biological Proces | Positive regulation of cell adhesion | 0,0037 | ERBB2\|FN1 | 50406 | 4988\|5576\|5 | 4,22E-05 | GO:0045785 | 0,24318 |
| 18 | 2 | GO Biological Proces | Negative regulation of anoikis | 0,0038 | NOTCH1\|I | 50406 | 3485\|5270 | 4,44E-05 | GO:2000811 | 0,242022 |
| 1588 | 6 | GO Biological Proces | Immune response | 0,0039 | STAT3\|NO | 50406 | 4127\|3485\|4 | 4,55E-05 | GO:0006955 | 0,240894 |
| 1587 | 6 | GO Biological Proces | Positive regulation of transcription, dna-templated | 0,0039 | STAT3\|ERE | 50406 | 4127\|4988\|5 | 4,53E-05 | GO:0045893 | 0,240894 |
| 925 | 5 | GO Biological Proces | Cell adhesion | 0,0041 | EGFR\|FN1 | 50406 | 5306\|5576\|5 | 4,94E-05 | GO:0007155 | 0,238722 |
| 923 | 5 | GO Biological Proces | Positive regulation of transport | 0,0041 | ERBB2\|EGI | 50406 | 4988\|5306\|4 | 4,89E-05 | GO:0051050 | 0,238722 |
| 19 | 2 | GO Biological Proces | Glomerulus vasculature development | 0,0041 | NOTCH1\|P | 50406 | 3485\|5090 | 4,91E-05 | GO:0072012 | 0,238722 |
| 148 | 3 | GO Biological Proces | Adaptive immune response based on somatic recom | 0,0043 | STAT3\|TLR | 50406 | 4127\|4079\|5 | 5,19E-05 | GO:0002460 | 0,236653 |
| 934 | 5 | GO Biological Proces | Regulation of protein localization | 0,0043 | ERBB2\|EGI | 50406 | 4988\|5306\|4 | 5,17E-05 | GO:0032880 | 0,236653 |
| 156 | 3 | GO Biological Proces | Negative regulation of hemopoiesis | 0,0048 | ERBB2\|NO | 50406 | 4988\|3485\|4 | 6,05E-05 | GO:1903707 | 0,231876 |
| 21 | 2 | GO Biological Proces | Negative regulation of stem cell differentiation | 0,0048 | STAT3\|NO | 50406 | 4127\|3485 | 5,91E-05 | GO:2000737 | 0,231876 |
| 164 | 3 | GO Biological Proces | Response to corticosteroid | 0,0054 | EGFR\|NOT | 50406 | 5306\|3485\|5 | 7,00E-05 | GO:0031960 | 0,226761 |
| 167 | 3 | GO Biological Proces | Positive regulation of protein secretion | 0,0057 | EGFR\|TLR4 | 50406 | 5306\|4079\|5 | 7,38E-05 | GO:0050714 | 0,224413 |
| 5447 | 9 | GO Biological Proces | Cellular component organization | 0,0058 | STAT3\|ERE | 50406 | 4127\|4988\|5 | 7,57E-05 | GO:0016043 | 0,223657 |
| 171 | 3 | GO Biological Proces | Positive regulation of cell growth | 0,0059 | ERBB2\|EGI | 50406 | 4988\|5306\|5 | 7,91E-05 | GO:0030307 | 0,222915 |
| 1013 | 5 | GO Biological Proces | Regulation of response to external stimulus | 0,0059 | STAT3\|EGI | 50406 | 4127\|5306\|3 | 7,62E-05 | GO:0032101 | 0,222915 |
| 495 | 4 | GO Biological Proces | Neuron projection morphogenesis | 0,0059 | ERBB2\|EGI | 50406 | 4988\|5306\|3 | 7,75E-05 | GO:0048812 | 0,222915 |

| | | | | | | | | | | |
|---|---|---|---|---|---|---|---|---|---|---|
| 170 | 3 | GO Biological Proces | Positive regulation of protein kinase b signaling | 0,0059 | ERBB2\|EGI | 50406 | 4988\|5306\|5 | 7,78E-05 | GO:0051897 | 0,222915 |
| 494 | 4 | GO Biological Proces | Cellular response to growth factor stimulus | 0,0059 | ERBB2\|EGI | 50406 | 4988\|5306\|3 | 7,69E-05 | GO:0071363 | 0,222915 |
| 493 | 4 | GO Biological Proces | Regulation of hemopoiesis | 0,0059 | STAT3\|ERE | 50406 | 4127\|4988\|5 | 7,63E-05 | GO:1903706 | 0,222915 |
| 173 | 3 | GO Biological Proces | Ameboidal-type cell migration | 0,006 | FN1\|ITGB1 | 50406 | 5576\|5270\|5 | 8,18E-05 | GO:0001667 | 0,222185 |
| 1760 | 6 | GO Biological Proces | Tissue development | 0,006 | EGFR\|NOT | 50406 | 5306\|3485\|5 | 8,16E-05 | GO:0009888 | 0,222185 |
| 505 | 4 | GO Biological Proces | Cell-cell adhesion | 0,0061 | EGFR\|ITGE | 50406 | 5306\|5270\|3 | 8,37E-05 | GO:0098609 | 0,221467 |
| 1776 | 6 | GO Biological Proces | Regulation of transport | 0,0062 | ERBB2\|EGI | 50406 | 4988\|5306\|3 | 8,58E-05 | GO:0051049 | 0,220761 |
| 522 | 4 | GO Biological Proces | Heart development | 0,0068 | ERBB2\|NO | 50406 | 4988\|3485\|5 | 9,51E-05 | GO:0007507 | 0,216749 |
| 27 | 2 | GO Biological Proces | Regulation of astrocyte differentiation | 0,0068 | NOTCH1\|II | 50406 | 3485\|5153 | 9,47E-05 | GO:0048710 | 0,216749 |
| 5591 | 9 | GO Biological Proces | Localization | 0,0068 | STAT3\|EGI | 50406 | 4127\|5306\|3 | 9,49E-05 | GO:0051179 | 0,216749 |
| 28 | 2 | GO Biological Proces | Response to lipoprotein particle | 0,007 | TLR4\|ITGB | 50406 | 4079\|5270 | 1,00E-04 | GO:0055094 | 0,21549 |
| 4042 | 8 | GO Biological Proces | Regulation of biological quality | 0,007 | STAT3\|EGI | 50406 | 4127\|5306\|3 | 1,00E-04 | GO:0065008 | 0,21549 |
| 188 | 3 | GO Biological Proces | Regulation of reactive oxygen species metabolic pro | 0,0071 | STAT3\|EGI | 50406 | 4127\|5306\|4 | 1,00E-04 | GO:2000377 | 0,214874 |
| 189 | 3 | GO Biological Proces | Positive regulation of vasculature development | 0,0072 | STAT3\|NO | 50406 | 4127\|3485\|5 | 1,10E-04 | GO:1904018 | 0,214267 |
| 29 | 2 | GO Biological Proces | Regulation of interleukin-17 production | 0,0073 | TLR4\|IL6 | 50406 | 4079\|5153 | 1,10E-04 | GO:0032660 | 0,213668 |
| 30 | 2 | GO Biological Proces | Regulation of glial cell proliferation | 0,0077 | NOTCH1\|II | 50406 | 3485\|5153 | 1,20E-04 | GO:0060251 | 0,211351 |
| 30 | 2 | GO Biological Proces | Cellular response to lipoprotein particle stimulus | 0,0077 | TLR4\|ITGB | 50406 | 4079\|5270 | 1,20E-04 | GO:0071402 | 0,211351 |
| 2875 | 7 | GO Biological Proces | Negative regulation of macromolecule metabolic pr | 0,0079 | STAT3\|EGI | 50406 | 4127\|5306\|3 | 1,20E-04 | GO:0010605 | 0,210237 |
| 561 | 4 | GO Biological Proces | Tissue morphogenesis | 0,0082 | EGFR\|NOT | 50406 | 5306\|3485\|5 | 1,30E-04 | GO:0048729 | 0,208619 |
| 32 | 2 | GO Biological Proces | ERBB2 signaling pathway | 0,0085 | ERBB2\|EGI | 50406 | 4988\|5306 | 1,30E-04 | GO:0038128 | 0,207058 |
| 210 | 3 | GO Biological Proces | Phagocytosis | 0,0092 | TLR4\|ITGB | 50406 | 4079\|5270\|5 | 1,40E-04 | GO:0006909 | 0,203621 |
| 218 | 3 | GO Biological Proces | Regulation of lymphocyte proliferation | 0,0101 | ERBB2\|TLF | 50406 | 4988\|4079\|5 | 1,60E-04 | GO:0050670 | 0,199568 |
| 37 | 2 | GO Biological Proces | Astrocyte development | 0,0107 | EGFR\|TLR4 | 50406 | 5306\|4079 | 1,70E-04 | GO:0014002 | 0,197062 |
| 630 | 4 | GO Biological Proces | Regulation of secretion by cell | 0,0119 | EGFR\|NOT | 50406 | 5306\|3485\|4 | 2,00E-04 | GO:1903530 | 0,192445 |
| 6137 | 9 | GO Biological Proces | Macromolecule metabolic process | 0,0125 | STAT3\|ERE | 50406 | 4127\|4988\|5 | 2,10E-04 | GO:0043170 | 0,190309 |
| 1253 | 5 | GO Biological Proces | Positive regulation of transcription by rna polymera: | 0,0125 | STAT3\|EGI | 50406 | 4127\|5306\|3 | 2,10E-04 | GO:0045944 | 0,190309 |
| 41 | 2 | GO Biological Proces | Regulation of extracellular matrix organization | 0,0125 | NOTCH1\|II | 50406 | 3485\|5153 | 2,10E-04 | GO:1903053 | 0,190309 |
| 42 | 2 | GO Biological Proces | Regulation of collagen metabolic process | 0,0128 | ITGB1\|IL6 | 50406 | 5270\|5153 | 2,20E-04 | GO:0010712 | 0,189279 |
| 242 | 3 | GO Biological Proces | Lymphocyte differentiation | 0,0128 | STAT3\|ITG | 50406 | 4127\|5270\|5 | 2,20E-04 | GO:0030098 | 0,189279 |
| 243 | 3 | GO Biological Proces | T cell activation | 0,0128 | STAT3\|IL6 | 50406 | 4127\|5153\|3 | 2,20E-04 | GO:0042110 | 0,189279 |
| 42 | 2 | GO Biological Proces | Cellular extravasation | 0,0128 | ITGB1\|PEC | 50406 | 5270\|5090 | 2,20E-04 | GO:0045123 | 0,189279 |
| 4479 | 8 | GO Biological Proces | Establishment of localization | 0,0128 | STAT3\|EGI | 50406 | 4127\|5306\|5 | 2,20E-04 | GO:0051234 | 0,189279 |
| 43 | 2 | GO Biological Proces | Regulation of bone resorption | 0,0132 | EGFR\|IL6 | 50406 | 5306\|5153 | 2,30E-04 | GO:0045124 | 0,187943 |
| 43 | 2 | GO Biological Proces | Cellular response to epidermal growth factor stimul | 0,0132 | ERBB2\|EGI | 50406 | 4988\|5306 | 2,30E-04 | GO:0071364 | 0,187943 |
| 2172 | 6 | GO Biological Proces | Regulation of transcription by rna polymerase ii | 0,0151 | STAT3\|ERE | 50406 | 4127\|4988\|5 | 2,70E-04 | GO:0006357 | 0,182102 |
| 47 | 2 | GO Biological Proces | Positive regulation of epithelial to mesenchymal tra | 0,0154 | NOTCH1\|II | 50406 | 3485\|5153 | 2,70E-04 | GO:0010718 | 0,181248 |
| 47 | 2 | GO Biological Proces | Negative regulation of cold-induced thermogenesis | 0,0154 | NOTCH1\|T | 50406 | 3485\|4079 | 2,70E-04 | GO:0120163 | 0,181248 |
| 696 | 4 | GO Biological Proces | Negative regulation of cell population proliferation | 0,0159 | STAT3\|ERE | 50406 | 4127\|4988\|3 | 2,90E-04 | GO:0008285 | 0,17986 |
| 265 | 3 | GO Biological Proces | Positive regulation of map kinase activity | 0,0159 | ERBB2\|EGI | 50406 | 4988\|5306\|4 | 2,80E-04 | GO:0043406 | 0,17986 |
| 48 | 2 | GO Biological Proces | Digestive tract morphogenesis | 0,0159 | EGFR\|NOT | 50406 | 5306\|3485 | 2,80E-04 | GO:0048546 | 0,17986 |
| 49 | 2 | GO Biological Proces | Mononuclear cell migration | 0,0164 | IL6\|PECAN | 50406 | 5153\|5090 | 3,00E-04 | GO:0071674 | 0,178516 |
| 271 | 3 | GO Biological Proces | Positive regulation of dna-binding transcription fact | 0,0167 | STAT3\|TLR | 50406 | 4127\|4079\|5 | 3,00E-04 | GO:0051091 | 0,177728 |
| 50 | 2 | GO Biological Proces | Positive regulation of notch signaling pathway | 0,0169 | STAT3\|NO | 50406 | 4127\|3485 | 3,10E-04 | GO:0045747 | 0,177211 |
| 51 | 2 | GO Biological Proces | Morphogenesis of an epithelial sheet | 0,0174 | NOTCH1\|C | 50406 | 3485\|3263 | 3,20E-04 | GO:0002011 | 0,175945 |
| 719 | 4 | GO Biological Proces | Positive regulation of cell death | 0,0176 | NOTCH1\|T | 50406 | 3485\|4079\|5 | 3,20E-04 | GO:0010942 | 0,175449 |
| 280 | 3 | GO Biological Proces | Negative regulation of cytokine production | 0,018 | FN1\|TLR4\| | 50406 | 5576\|4079\|5 | 3,30E-04 | GO:0001818 | 0,174473 |
| 52 | 2 | GO Biological Proces | Positive regulation of fibroblast proliferation | 0,018 | EGFR\|FN1 | 50406 | 5306\|5576 | 3,30E-04 | GO:0048146 | 0,174473 |
| 1389 | 5 | GO Biological Proces | Positive regulation of developmental process | 0,0182 | STAT3\|NO | 50406 | 4127\|3485\|5 | 3,40E-04 | GO:0051094 | 0,173993 |
| 53 | 2 | GO Biological Proces | Negative regulation of erbb signaling pathway | 0,0185 | ERBB2\|EGI | 50406 | 4988\|5306 | 3,40E-04 | GO:1901185 | 0,173283 |
| 287 | 3 | GO Biological Proces | Negative regulation of neurogenesis | 0,019 | STAT3\|NO | 50406 | 4127\|3485\|5 | 3,60E-04 | GO:0050768 | 0,172125 |
| 742 | 4 | GO Biological Proces | Regulation of cytokine production | 0,0194 | STAT3\|FN1 | 50406 | 4127\|5576\|4 | 3,70E-04 | GO:0001817 | 0,17122 |
| 55 | 2 | GO Biological Proces | Regulation of macrophage activation | 0,0195 | TLR4\|IL6 | 50406 | 4079\|5153 | 3,70E-04 | GO:0043030 | 0,170997 |
| 2310 | 6 | GO Biological Proces | Response to external stimulus | 0,0197 | ERBB2\|EGI | 50406 | 4988\|5306\|3 | 3,70E-04 | GO:0009605 | 0,170553 |
| 59 | 2 | GO Biological Proces | Sprouting angiogenesis | 0,0222 | NOTCH1\|I | 50406 | 3485\|5270 | 4,20E-04 | GO:0002040 | 0,165365 |
| 59 | 2 | GO Biological Proces | Positive regulation of cytokine production involved i | 0,0222 | TLR4\|IL6 | 50406 | 4079\|5153 | 4,20E-04 | GO:0002720 | 0,165365 |
| 59 | 2 | GO Biological Proces | Positive regulation of wound healing | 0,0222 | TLR4\|ITGB | 50406 | 4079\|5270 | 4,20E-04 | GO:0090303 | 0,165365 |
| 61 | 2 | GO Biological Proces | Positive regulation of chemokine production | 0,0235 | TLR4\|IL6 | 50406 | 4079\|5153 | 4,50E-04 | GO:0032722 | 0,162893 |
| 313 | 3 | GO Biological Proces | Response to lipopolysaccharide | 0,0238 | NOTCH1\|T | 50406 | 3485\|4079\|5 | 4,60E-04 | GO:0032496 | 0,162342 |
| 315 | 3 | GO Biological Proces | Regulation of leukocyte cell-cell adhesion | 0,024 | ERBB2\|IL6 | 50406 | 4988\|5153\|3 | 4,70E-04 | GO:1903037 | 0,161979 |
| 62 | 2 | GO Biological Proces | Positive regulation of protein localization to plasma | 0,024 | EGFR\|ITGE | 50406 | 5306\|5270 | 4,70E-04 | GO:1903078 | 0,161979 |
| 4976 | 8 | GO Biological Proces | Cellular macromolecule metabolic process | 0,0243 | STAT3\|ERE | 50406 | 4127\|4988\|5 | 4,70E-04 | GO:0044260 | 0,161439 |
| 2429 | 6 | GO Biological Proces | Negative regulation of nitrogen compound metabol | 0,0252 | STAT3\|EGI | 50406 | 4127\|5306\|3 | 4,90E-04 | GO:0051172 | 0,15986 |
| 67 | 2 | GO Biological Proces | Endothelial cell migration | 0,0271 | ITGB1\|PEC | 50406 | 5270\|5090 | 5,40E-04 | GO:0043542 | 0,156703 |
| 333 | 3 | GO Biological Proces | Regulation of developmental growth | 0,0274 | STAT3\|NO | 50406 | 4127\|3485\|5 | 5,50E-04 | GO:0048638 | 0,156225 |
| 828 | 4 | GO Biological Proces | Regulation of neurogenesis | 0,0275 | STAT3\|NO | 50406 | 4127\|3485\|5 | 5,50E-04 | GO:0050767 | 0,156067 |
| 1567 | 5 | GO Biological Proces | Response to oxygen-containing compound | 0,0289 | STAT3\|EGI | 50406 | 4127\|5306\|3 | 5,90E-04 | GO:1901700 | 0,15391 |
| 849 | 4 | GO Biological Proces | Response to hormone | 0,0294 | STAT3\|EGI | 50406 | 4127\|5306\|3 | 6,10E-04 | GO:0009725 | 0,153165 |
| 72 | 2 | GO Biological Proces | Positive regulation of gliogenesis | 0,0299 | NOTCH1\|II | 50406 | 3485\|5153 | 6,20E-04 | GO:0014015 | 0,152433 |
| 865 | 4 | GO Biological Proces | Symbiotic process | 0,0312 | STAT3\|EGI | 50406 | 4127\|5306\|5 | 6,50E-04 | GO:0044403 | 0,150585 |
| 74 | 2 | GO Biological Proces | Hair follicle development | 0,0313 | EGFR\|NOT | 50406 | 5306\|3485 | 6,60E-04 | GO:0001942 | 0,150446 |
| 74 | 2 | GO Biological Proces | Oligodendrocyte differentiation | 0,0313 | ERBB2\|NO | 50406 | 4988\|3485 | 6,60E-04 | GO:0048709 | 0,150446 |
| 75 | 2 | GO Biological Proces | Animal organ regeneration | 0,0319 | EGFR\|NOT | 50406 | 5306\|3485 | 6,70E-04 | GO:0031100 | 0,149621 |
| 76 | 2 | GO Biological Proces | Endoderm development | 0,0327 | NOTCH1\|F | 50406 | 3485\|5576 | 6,90E-04 | GO:0007492 | 0,148545 |
| 76 | 2 | GO Biological Proces | Endothelial cell differentiation | 0,0327 | NOTCH1\|P | 50406 | 3485\|5090 | 6,90E-04 | GO:0045446 | 0,148545 |
| 79 | 2 | GO Biological Proces | Cellular response to mechanical stimulus | 0,0345 | EGFR\|TLR4 | 50406 | 5306\|4079 | 7,40E-04 | GO:0071260 | 0,146218 |
| 80 | 2 | GO Biological Proces | Mesenchymal cell development | 0,0353 | NOTCH1\|F | 50406 | 3485\|5576 | 7,60E-04 | GO:0014031 | 0,145223 |
| 911 | 4 | GO Biological Proces | Response to organic cyclic compound | 0,0366 | STAT3\|EGI | 50406 | 4127\|5306\|3 | 7,90E-04 | GO:0014070 | 0,143652 |
| 82 | 2 | GO Biological Proces | Positive regulation of nik/nf-kappab signaling | 0,0368 | EGFR\|TLR4 | 50406 | 5306\|4079 | 8,00E-04 | GO:1901224 | 0,143415 |
| 382 | 3 | GO Biological Proces | In utero embryonic development | 0,0375 | EGFR\|NOT | 50406 | 5306\|3485\|5 | 8,20E-04 | GO:0001701 | 0,142597 |
| 384 | 3 | GO Biological Proces | Axonogenesis | 0,0378 | ERBB2\|NO | 50406 | 4988\|3485\|5 | 8,30E-04 | GO:0007409 | 0,142251 |
| 85 | 2 | GO Biological Proces | Positive regulation of b cell activation | 0,0389 | TLR4\|IL6 | 50406 | 4079\|5153 | 8,60E-04 | GO:0050871 | 0,141005 |
| 87 | 2 | GO Biological Proces | Cell fate specification | 0,0405 | NOTCH1\|I | 50406 | 3485\|5270 | 9,00E-04 | GO:0001708 | 0,139254 |
| 398 | 3 | GO Biological Proces | Regulation of translation | 0,0412 | STAT3\|ERE | 50406 | 4127\|4988\|5 | 9,20E-04 | GO:0006417 | 0,13851 |
| 88 | 2 | GO Biological Proces | Positive regulation of smooth muscle cell proliferati | 0,0412 | EGFR\|IL6 | 50406 | 5306\|5153 | 9,20E-04 | GO:0048661 | 0,13851 |
| 979 | 4 | GO Biological Proces | Secretion by cell | 0,0456 | FN1\|IL6\|CI | 50406 | 5576\|5153\|3 | 0,001 | GO:0032940 | 0,134104 |
| 95 | 2 | GO Biological Proces | Viral entry into host cell | 0,0465 | EGFR\|ITGE | 50406 | 5306\|5270 | 0,0011 | GO:0046718 | 0,133255 |
| 95 | 2 | GO Biological Proces | Regulation of reactive oxygen species biosynthetic p | 0,0465 | STAT3\|TLR | 50406 | 4127\|4079 | 0,0011 | GO:1903426 | 0,133255 |
| 993 | 4 | GO Biological Proces | Positive regulation of cell differentiation | 0,0474 | STAT3\|NO | 50406 | 4127\|3485\|5 | 0,0011 | GO:0045597 | 0,132422 |
| 1002 | 4 | GO Biological Proces | Embryo development | 0,0487 | EGFR\|NOT | 50406 | 5306\|3485\|5 | 0,0011 | GO:0009790 | 0,131247 |
| 1805 | 5 | GO Biological Proces | Vesicle-mediated transport | 0,0487 | FN1\|TLR4\| | 50406 | 5576\|4079\|5 | 0,0011 | GO:0016192 | 0,131247 |
| 98 | 2 | GO Biological Proces | Defense response to gram-negative bacterium | 0,0487 | TLR4\|IL6 | 50406 | 4079\|5153 | 0,0011 | GO:0050829 | 0,131247 |
| 98 | 2 | GO Biological Proces | Nephron epithelium development | 0,0487 | NOTCH1\|P | 50406 | 3485\|5090 | 0,0011 | GO:0072009 | 0,131247 |
| 99 | 2 | GO Biological Proces | Gland morphogenesis | 0,0492 | EGFR\|NOT | 50406 | 5306\|3485 | 0,0012 | GO:0022612 | 0,130803 |
| 381 | 7 | GO Cellular Compon | Receptor complex | 3,19E-07 | ERBB2\|EGI | 50406 | 4988\|5306\|3 | 1,30E-10 | GO:0043235 | 0,649621 |
| 350 | 5 | GO Cellular Compon | Apical plasma membrane | 5,50E-04 | ERBB2\|EGI | 50406 | 4988\|5306\|3 | 4,46E-07 | GO:0016324 | 0,325964 |
| 5314 | 10 | GO Cellular Compon | Plasma membrane | 8,50E-04 | STAT3\|ERE | 50406 | 4127\|4988\|5 | 2,20E-06 | GO:0005886 | 0,307058 |
| 1623 | 7 | GO Cellular Compon | Integral component of plasma membrane | 8,50E-04 | ERBB2\|EGI | 50406 | 4988\|5306\|4 | 2,63E-06 | GO:0005887 | 0,307058 |
| 824 | 6 | GO Cellular Compon | Cell surface | 8,50E-04 | EGFR\|NOT | 50406 | 5306\|3485\|4 | 1,04E-06 | GO:0009986 | 0,307058 |

| | | | | | | | | | | |
|---|---|---|---|---|---|---|---|---|---|---|
| 2386 | 8 | GO Cellular Compon | Cytoplasmic vesicle | 8,50E-04 | ERBB2\|EGI | 50406 | 4988\|5306\|3 | 1,77E-06 | GO:0031410 | 0,307058 |
| 5073 | 10 | GO Cellular Compon | Protein-containing complex | 8,50E-04 | STAT3\|ERE | 50406 | 4127\|4988\|5 | 1,38E-06 | GO:0032991 | 0,307058 |
| 1219 | 6 | GO Cellular Compon | Plasma membrane region | 0,0021 | ERBB2\|EGI | 50406 | 4988\|5306\|3 | 1,00E-05 | GO:0098590 | 0,267778 |
| 4542 | 9 | GO Cellular Compon | Endomembrane system | 0,0029 | ERBB2\|EGI | 50406 | 4988\|5306\|3 | 1,56E-05 | GO:0012505 | 0,25376 |
| 820 | 5 | GO Cellular Compon | Anchoring junction | 0,0049 | EGFR\|NOT | 50406 | 5306\|3485\|5 | 2,77E-05 | GO:0070161 | 0,23098 |
| 169 | 3 | GO Cellular Compon | Plasma membrane signaling receptor complex | 0,0115 | ERBB2\|ITG | 50406 | 4988\|5270\|5 | 7,64E-05 | GO:0098802 | 0,19393 |
| 1715 | 6 | GO Cellular Compon | Whole membrane | 0,0115 | ERBB2\|EGI | 50406 | 4988\|5306\|4 | 7,04E-05 | GO:0098805 | 0,19393 |
| 547 | 4 | GO Cellular Compon | Plasma membrane protein complex | 0,0155 | ERBB2\|EGI | 50406 | 4988\|5306\|5 | 1,10E-04 | GO:0098797 | 0,180967 |
| 1141 | 5 | GO Cellular Compon | Membrane protein complex | 0,0173 | ERBB2\|EGI | 50406 | 4988\|5306\|4 | 1,30E-04 | GO:0098796 | 0,176195 |
| 237 | 3 | GO Cellular Compon | Basolateral plasma membrane | 0,0252 | ERBB2\|EGI | 50406 | 4988\|5306\|3 | 2,00E-04 | GO:0016323 | 0,15986 |
| 2075 | 6 | GO Cellular Compon | Cell junction | 0,0252 | STAT3\|EGI | 50406 | 4127\|5306\|3 | 2,10E-04 | GO:0030054 | 0,15986 |
| 2107 | 6 | GO Cellular Compon | Bounding membrane of organelle | 0,0252 | ERBB2\|EGI | 50406 | 4988\|5306\|3 | 2,20E-04 | GO:0098588 | 0,15986 |
| 44 | 2 | GO Cellular Compon | Basal plasma membrane | 0,0257 | ERBB2\|EGI | 50406 | 4988\|5306 | 2,40E-04 | GO:0009925 | 0,159007 |
| 46 | 2 | GO Cellular Compon | Myelin sheath | 0,0268 | ERBB2\|ITG | 50406 | 4988\|5270 | 2,60E-04 | GO:0043209 | 0,157187 |
| 727 | 4 | GO Cellular Compon | Perinuclear region of cytoplasm | 0,0332 | ERBB2\|EGI | 50406 | 4988\|5306\|4 | 3,40E-04 | GO:0048471 | 0,147886 |
| 331 | 3 | GO Cellular Compon | External side of plasma membrane | 0,0435 | TLR4\|ITGB | 50406 | 4079\|5270\|5 | 5,40E-04 | GO:0009897 | 0,136151 |
| 3548 | 7 | GO Cellular Compon | Organelle membrane | 0,0435 | STAT3\|ERE | 50406 | 4127\|4988\|5 | 4,60E-04 | GO:0031090 | 0,136151 |
| 324 | 3 | GO Cellular Compon | Membrane raft | 0,0435 | EGFR\|ITGE | 50406 | 5306\|5270\|5 | 5,10E-04 | GO:0045121 | 0,136151 |
| 845 | 4 | GO Cellular Compon | Secretory granule | 0,0445 | NOTCH1\|F | 50406 | 3485\|5576\|3 | 6,00E-04 | GO:0030141 | 0,135164 |
| 5181 | 8 | GO Cellular Compon | Integral component of membrane | 0,045 | ERBB2\|EGI | 50406 | 4988\|5306\|3 | 6,40E-04 | GO:0016021 | 0,134679 |
| 72 | 2 | GO Cellular Compon | Cell-cell contact zone | 0,045 | ITGB1\|PEC | 50406 | 5270\|5090 | 6,20E-04 | GO:0044291 | 0,134679 |
| 1581 | 8 | GO Molecular Functi | Signaling receptor binding | 3,80E-04 | STAT3\|ERE | 50406 | 4127\|4988\|5 | 7,19E-08 | GO:0005102 | 0,342022 |
| 1453 | 7 | GO Molecular Functi | Signaling receptor activity | 0,0032 | ERBB2\|EGI | 50406 | 4988\|5306\|3 | 1,24E-06 | GO:0038023 | 0,249485 |
| 68 | 3 | GO Molecular Functi | Collagen binding | 0,007 | FN1\|ITGB1 | 50406 | 5576\|5270\|3 | 5,39E-06 | GO:0005518 | 0,21549 |
| 1240 | 6 | GO Molecular Functi | Transmembrane signaling receptor activity | 0,0115 | ERBB2\|EGI | 50406 | 4988\|5306\|3 | 1,10E-05 | GO:0004888 | 0,19393 |
| 147 | 3 | GO Molecular Functi | Integrin binding | 0,0443 | EGFR\|FN1 | 50406 | 5306\|5576\|5 | 5,09E-05 | GO:0005178 | 0,13536 |
| 149 | 3 | GO Molecular Functi | Protein phosphatase binding | 0,0443 | STAT3\|ERE | 50406 | 4127\|4988\|5 | 5,29E-05 | GO:0019903 | 0,13536 |
| 196 | 7 | KEGG Pathways | Proteoglycans in cancer | 6,17E-10 | STAT3\|ERE | 50406 | 4127\|4988\|5 | 1,36E-12 | hsa05205 | 0,920971 |
| 106 | 5 | KEGG Pathways | HIF-1 signaling pathway | 2,40E-07 | STAT3\|ERE | 50406 | 4127\|4988\|5 | 1,32E-09 | hsa04066 | 0,661979 |
| 517 | 7 | KEGG Pathways | Pathways in cancer | 2,40E-07 | STAT3\|ERE | 50406 | 4127\|4988\|5 | 1,06E-09 | hsa05200 | 0,661979 |
| 350 | 6 | KEGG Pathways | PI3K-Akt signaling pathway | 7,77E-07 | ERBB2\|EGI | 50406 | 4988\|5306\|5 | 6,85E-09 | hsa04151 | 0,610958 |
| 160 | 5 | KEGG Pathways | MicroRNAs in cancer | 8,85E-07 | STAT3\|ERE | 50406 | 4127\|4988\|5 | 9,75E-09 | hsa05206 | 0,605306 |
| 78 | 4 | KEGG Pathways | EGFR tyrosine kinase inhibitor resistance | 4,46E-06 | STAT3\|ERE | 50406 | 4127\|4988\|5 | 5,89E-08 | hsa01521 | 0,535067 |
| 125 | 4 | KEGG Pathways | Yersinia infection | 2,38E-05 | FN1\|TLR4\| | 50406 | 5576\|4079\|5 | 3,67E-07 | hsa05135 | 0,462342 |
| 46 | 3 | KEGG Pathways | Malaria | 9,91E-05 | TLR4\|IL6\|F | 50406 | 4079\|5153\|5 | 1,75E-06 | hsa05144 | 0,400393 |
| 198 | 4 | KEGG Pathways | Focal adhesion | 1,10E-04 | ERBB2\|EGI | 50406 | 4988\|5306\|5 | 2,20E-06 | hsa04510 | 0,395861 |
| 218 | 4 | KEGG Pathways | Shigellosis | 1,50E-04 | EGFR\|TLR4 | 50406 | 5306\|4079\|5 | 3,20E-06 | hsa05131 | 0,382391 |
| 60 | 3 | KEGG Pathways | Inflammatory bowel disease | 1,50E-04 | STAT3\|TLR | 50406 | 4127\|4079\|5 | 3,75E-06 | hsa05321 | 0,382391 |
| 68 | 3 | KEGG Pathways | Non-small cell lung cancer | 2,00E-04 | STAT3\|ERE | 50406 | 4127\|4988\|5 | 5,39E-06 | hsa05223 | 0,369897 |
| 74 | 3 | KEGG Pathways | Pertussis | 2,30E-04 | TLR4\|ITGB | 50406 | 4079\|5270\|5 | 6,88E-06 | hsa05133 | 0,363827 |
| 73 | 3 | KEGG Pathways | Pancreatic cancer | 2,30E-04 | STAT3\|ERE | 50406 | 4127\|4988\|5 | 6,62E-06 | hsa05212 | 0,363827 |
| 88 | 3 | KEGG Pathways | ECM-receptor interaction | 3,40E-04 | FN1\|ITGB1 | 50406 | 5576\|5270\|3 | 1,14E-05 | hsa04512 | 0,346852 |
| 88 | 3 | KEGG Pathways | PD-L1 expression and PD-1 checkpoint pathway in ca | 3,40E-04 | STAT3\|EGI | 50406 | 4127\|5306\|4 | 1,14E-05 | hsa05235 | 0,346852 |
| 95 | 3 | KEGG Pathways | Endocrine resistance | 3,80E-04 | ERBB2\|EGI | 50406 | 4988\|5306\|3 | 1,42E-05 | hsa01522 | 0,342022 |
| 98 | 3 | KEGG Pathways | AGE-RAGE signaling pathway in diabetic complicatic | 3,80E-04 | STAT3\|FN1 | 50406 | 4127\|5576\|5 | 1,56E-05 | hsa04933 | 0,342022 |
| 100 | 3 | KEGG Pathways | Amoebiasis | 3,80E-04 | FN1\|TLR4\| | 50406 | 5576\|4079\|5 | 1,65E-05 | hsa05146 | 0,342022 |
| 325 | 4 | KEGG Pathways | Human papillomavirus infection | 3,80E-04 | EGFR\|NOT | 50406 | 5306\|3485\|5 | 1,52E-05 | hsa05165 | 0,342022 |
| 105 | 3 | KEGG Pathways | Toxoplasmosis | 4,10E-04 | STAT3\|TLR | 50406 | 4127\|4079\|5 | 1,91E-05 | hsa05145 | 0,338722 |
| 127 | 3 | KEGG Pathways | FoxO signaling pathway | 6,80E-04 | STAT3\|EGI | 50406 | 4127\|5306\|5 | 3,32E-05 | hsa04068 | 0,316749 |
| 138 | 3 | KEGG Pathways | Measles | 8,30E-04 | STAT3\|TLR | 50406 | 4127\|4079\|5 | 4,23E-05 | hsa05162 | 0,308092 |
| 145 | 3 | KEGG Pathways | Breast cancer | 9,20E-04 | ERBB2\|EGI | 50406 | 4988\|5306\|3 | 4,89E-05 | hsa05224 | 0,303621 |
| 160 | 3 | KEGG Pathways | JAK-STAT signaling pathway | 0,0012 | STAT3\|EGI | 50406 | 4127\|5306\|5 | 6,52E-05 | hsa04630 | 0,292082 |
| 159 | 3 | KEGG Pathways | Hepatitis B | 0,0012 | STAT3\|TLR | 50406 | 4127\|4079\|5 | 6,40E-05 | hsa05161 | 0,292082 |
| 187 | 3 | KEGG Pathways | Pathogenic Escherichia coli infection | 0,0017 | TLR4\|ITGB | 50406 | 4079\|5270\|5 | 1,00E-04 | hsa05130 | 0,276955 |
| 193 | 3 | KEGG Pathways | Epstein-Barr virus infection | 0,0018 | STAT3\|IL6 | 50406 | 4127\|5153\|3 | 1,10E-04 | hsa05169 | 0,274473 |
| 209 | 3 | KEGG Pathways | Regulation of actin cytoskeleton | 0,0022 | EGFR\|FN1 | 50406 | 5306\|5576\|5 | 1,40E-04 | hsa04810 | 0,265758 |
| 218 | 3 | KEGG Pathways | Human cytomegalovirus infection | 0,0024 | STAT3\|EGI | 50406 | 4127\|5306\|5 | 1,60E-04 | hsa05163 | 0,261979 |
| 41 | 2 | KEGG Pathways | Bladder cancer | 0,0031 | ERBB2\|EGI | 50406 | 4988\|5306 | 2,10E-04 | hsa05219 | 0,250864 |
| 55 | 2 | KEGG Pathways | Legionellosis | 0,0052 | TLR4\|IL6 | 50406 | 4079\|5153 | 3,70E-04 | hsa05134 | 0,2284 |
| 57 | 2 | KEGG Pathways | Endometrial cancer | 0,0054 | ERBB2\|EGI | 50406 | 4988\|5306 | 4,00E-04 | hsa05213 | 0,226761 |
| 67 | 2 | KEGG Pathways | Adherens junction | 0,0072 | ERBB2\|EGI | 50406 | 4988\|5306 | 5,40E-04 | hsa04520 | 0,214267 |
| 70 | 2 | KEGG Pathways | Bacterial invasion of epithelial cells | 0,0074 | FN1\|ITGB1 | 50406 | 5576\|5270 | 5,90E-04 | hsa05100 | 0,213077 |
| 70 | 2 | KEGG Pathways | Leishmaniasis | 0,0074 | TLR4\|ITGB | 50406 | 4079\|5270 | 5,90E-04 | hsa05140 | 0,213077 |
| 69 | 2 | KEGG Pathways | Central carbon metabolism in cancer | 0,0074 | ERBB2\|EGI | 50406 | 4988\|5306 | 5,70E-04 | hsa05230 | 0,213077 |
| 83 | 2 | KEGG Pathways | ErbB signaling pathway | 0,0098 | ERBB2\|EGI | 50406 | 4988\|5306 | 8,20E-04 | hsa04012 | 0,200877 |
| 85 | 2 | KEGG Pathways | Rheumatoid arthritis | 0,01 | TLR4\|IL6 | 50406 | 4079\|5153 | 8,60E-04 | hsa05323 | 0,2 |
| 89 | 2 | KEGG Pathways | Hypertrophic cardiomyopathy | 0,0107 | ITGB1\|IL6 | 50406 | 5270\|5153 | 9,40E-04 | hsa05410 | 0,197062 |
| 91 | 2 | KEGG Pathways | Hematopoietic cell lineage | 0,0109 | IL6\|CD44 | 50406 | 5153\|3263 | 9,80E-04 | hsa04640 | 0,196257 |
| 92 | 2 | KEGG Pathways | Small cell lung cancer | 0,0109 | FN1\|ITGB1 | 50406 | 5576\|5270 | 0,001 | hsa05222 | 0,196257 |
| 96 | 2 | KEGG Pathways | Prostate cancer | 0,0115 | ERBB2\|EGI | 50406 | 4988\|5306 | 0,0011 | hsa05215 | 0,19393 |
| 99 | 2 | KEGG Pathways | Chagas disease | 0,0119 | TLR4\|IL6 | 50406 | 4079\|5153 | 0,0012 | hsa05142 | 0,192445 |
| 101 | 2 | KEGG Pathways | Toll-like receptor signaling pathway | 0,0121 | TLR4\|IL6 | 50406 | 4079\|5153 | 0,0012 | hsa04620 | 0,191721 |
| 101 | 2 | KEGG Pathways | Th17 cell differentiation | 0,0121 | STAT3\|IL6 | 50406 | 4127\|5153 | 0,0012 | hsa04659 | 0,191721 |
| 107 | 2 | KEGG Pathways | Insulin resistance | 0,013 | STAT3\|IL6 | 50406 | 4127\|5153 | 0,0013 | hsa04931 | 0,188606 |
| 109 | 2 | KEGG Pathways | Leukocyte transendothelial migration | 0,0132 | ITGB1\|PEC | 50406 | 5270\|5090 | 0,0014 | hsa04670 | 0,187943 |
| 137 | 2 | KEGG Pathways | Cell adhesion molecules | 0,0201 | ITGB1\|PEC | 50406 | 5270\|5090 | 0,0022 | hsa04514 | 0,16968 |
| 142 | 2 | KEGG Pathways | Phagosome | 0,0211 | TLR4\|ITGB | 50406 | 4079\|5270 | 0,0023 | hsa04145 | 0,167572 |
| 144 | 2 | KEGG Pathways | Gastric cancer | 0,0213 | ERBB2\|EGI | 50406 | 4988\|5306 | 0,0024 | hsa05226 | 0,167162 |
| 149 | 2 | KEGG Pathways | Necroptosis | 0,0223 | STAT3\|TLR | 50406 | 4127\|4079 | 0,0026 | hsa04217 | 0,16517 |
| 156 | 2 | KEGG Pathways | Tight junction | 0,0239 | ERBB2\|ITG | 50406 | 4988\|5270 | 0,0028 | hsa04530 | 0,16216 |
| 156 | 2 | KEGG Pathways | Hepatitis C | 0,0239 | STAT3\|EGI | 50406 | 4127\|5306 | 0,0028 | hsa05160 | 0,16216 |
| 165 | 2 | KEGG Pathways | Influenza A | 0,0257 | TLR4\|IL6 | 50406 | 4079\|5153 | 0,0031 | hsa05164 | 0,159007 |
| 168 | 2 | KEGG Pathways | Tuberculosis | 0,0261 | TLR4\|IL6 | 50406 | 4079\|5153 | 0,0032 | hsa05152 | 0,158336 |
| 174 | 2 | KEGG Pathways | NOD-like receptor signaling pathway | 0,0275 | TLR4\|IL6 | 50406 | 4079\|5153 | 0,0034 | hsa04621 | 0,156067 |
| 187 | 2 | KEGG Pathways | Kaposi sarcoma-associated herpesvirus infection | 0,031 | STAT3\|IL6 | 50406 | 4127\|5153 | 0,004 | hsa05167 | 0,150864 |
| 193 | 2 | KEGG Pathways | Calcium signaling pathway | 0,0324 | ERBB2\|EGI | 50406 | 4988\|5306 | 0,0042 | hsa04020 | 0,148945 |
| 202 | 2 | KEGG Pathways | Rap1 signaling pathway | 0,0348 | EGFR\|ITGE | 50406 | 5306\|5270 | 0,0046 | hsa04015 | 0,145842 |
| 209 | 2 | KEGG Pathways | Salmonella infection | 0,0366 | TLR4\|IL6 | 50406 | 4079\|5153 | 0,0049 | hsa05132 | 0,143652 |
| 1548 | 8 | Reactome Pathways | Disease | 0,0011 | STAT3\|ERE | 50406 | 4127\|4988\|5 | 6,10E-08 | HSA-1643685 | 0,295861 |
| 85 | 4 | Reactome Pathways | Integrin cell surface interactions | 0,0011 | FN1\|ITGB1 | 50406 | 5576\|5270\|3 | 8,21E-08 | HSA-216083 | 0,295861 |
| 107 | 4 | Reactome Pathways | Interleukin-4 and Interleukin-13 signaling | 0,0012 | STAT3\|FN1 | 50406 | 4127\|5576\|5 | 2,00E-07 | HSA-6785807 | 0,292082 |
| 138 | 4 | Reactome Pathways | Cell surface interactions at the vascular wall | 0,0023 | FN1\|ITGB1 | 50406 | 5576\|5270\|3 | 5,39E-07 | HSA-202733 | 0,263827 |
| 392 | 5 | Reactome Pathways | Diseases of signal transduction by growth factor rec | 0,0027 | STAT3\|ERE | 50406 | 4127\|4988\|5 | 7,75E-07 | HSA-5663202 | 0,256864 |
| 4 | 2 | Reactome Pathways | PLCG1 events in ERBB2 signaling | 0,0075 | ERBB2\|EGI | 50406 | 4988\|5306 | 3,52E-06 | HSA-1251932 | 0,212494 |
| 53 | 3 | Reactome Pathways | Signaling by PTK6 | 0,0075 | STAT3\|ERE | 50406 | 4127\|4988\|5 | 2,62E-06 | HSA-8848021 | 0,212494 |
| 502 | 5 | Reactome Pathways | Signaling by Receptor Tyrosine Kinases | 0,0075 | STAT3\|ERE | 50406 | 4127\|4988\|5 | 2,59E-06 | HSA-9006934 | 0,212494 |
| 6 | 2 | Reactome Pathways | Fibronectin matrix formation | 0,0114 | FN1\|ITGB1 | 50406 | 5576\|5270 | 6,57E-06 | HSA-1566977 | 0,19431 |
| 1956 | 7 | Reactome Pathways | Immune System | 0,0131 | STAT3\|FN1 | 50406 | 4127\|5576\|4 | 9,21E-06 | HSA-168256 | 0,188273 |

| | | | | | | | | | | |
|---|---|---|---|---|---|---|---|---|---|---|
| 280 | 4 | Reactome Pathways | MAPK1/MAPK3 signaling | 0,0131 | ERBB2\|EGI | 50406 | 4988\|5306\|5 | 8,50E-06 | HSA-5684996 | 0,188273 |
| 79 | 3 | Reactome Pathways | Signaling by MET | 0,0131 | STAT3\|FN1 | 50406 | 4127\|5576\|5 | 8,32E-06 | HSA-6806834 | 0,188273 |
| 681 | 5 | Reactome Pathways | Cytokine Signaling in Immune system | 0,014 | STAT3\|FN1 | 50406 | 4127\|5576\|5 | 1,13E-05 | HSA-1280215 | 0,185387 |
| 11 | 2 | Reactome Pathways | Interleukin-6 signaling | 0,0187 | STAT3\|IL6 | 50406 | 4127\|5153 | 1,83E-05 | HSA-1059683 | 0,172816 |
| 12 | 2 | Reactome Pathways | TFAP2 (AP-2) family regulates transcription of growt | 0,0206 | ERBB2\|EGI | 50406 | 4988\|5306 | 2,13E-05 | HSA-8866910 | 0,168613 |
| 13 | 2 | Reactome Pathways | ERBB2 Activates PTK6 Signaling | 0,0225 | ERBB2\|EGI | 50406 | 4988\|5306 | 2,46E-05 | HSA-8847993 | 0,164782 |
| 15 | 2 | Reactome Pathways | ERBB2 Regulates Cell Motility | 0,0276 | ERBB2\|EGI | 50406 | 4988\|5306 | 3,18E-05 | HSA-6785631 | 0,155909 |
| 16 | 2 | Reactome Pathways | GRB2 events in ERBB2 signaling | 0,0296 | ERBB2\|EGI | 50406 | 4988\|5306 | 3,58E-05 | HSA-1963640 | 0,152871 |
| 16 | 2 | Reactome Pathways | PI3K events in ERBB2 signaling | 0,0296 | ERBB2\|EGI | 50406 | 4988\|5306 | 3,58E-05 | HSA-1963642 | 0,152871 |
| 16 | 2 | Reactome Pathways | Signaling by ERBB2 ECD mutants | 0,0296 | ERBB2\|EGI | 50406 | 4988\|5306 | 3,58E-05 | HSA-9665348 | 0,152871 |
| 21 | 2 | Reactome Pathways | Signal transduction by L1 | 0,0411 | EGFR\|ITGE | 50406 | 5306\|5270 | 5,91E-05 | HSA-445144 | 0,138616 |
| 22 | 2 | Reactome Pathways | SHC1 events in ERBB2 signaling | 0,0431 | ERBB2\|EGI | 50406 | 4988\|5306 | 6,44E-05 | HSA-1250196 | 0,136552 |
| 22 | 2 | Reactome Pathways | Signaling by ERBB2 TMD/JMD mutants | 0,0431 | ERBB2\|EGI | 50406 | 4988\|5306 | 6,44E-05 | HSA-9665686 | 0,136552 |
| 25 | 2 | Reactome Pathways | Signaling by ERBB2 KD Mutants | 0,0491 | ERBB2\|EGI | 50406 | 4988\|5306 | 8,19E-05 | HSA-9664565 | 0,130892 |
| 271 | 7 | TISSUES | Cervical carcinoma cell | 3,71E-08 | STAT3\|ERE | 50406 | 4127\|4988\|5 | 1,25E-11 | BTO:0000180 | 0,743063 |
| 1882 | 9 | TISSUES | Liver | 9,76E-06 | STAT3\|ERE | 50406 | 4127\|4988\|5 | 6,58E-09 | BTO:0000759 | 0,501055 |
| 398 | 5 | TISSUES | Vascular system | 6,20E-04 | NOTCH1\|F | 50406 | 3485\|5576\|5 | 8,35E-07 | BTO:0001085 | 0,320761 |
| 1436 | 7 | TISSUES | Respiratory system | 6,80E-04 | STAT3\|FN1 | 50406 | 4127\|5576\|4 | 1,15E-06 | BTO:0000203 | 0,316749 |
| 3 | 2 | TISSUES | Kupffer cell | 0,0012 | TLR4\|IL6 | 50406 | 4079\|5153 | 2,35E-06 | BTO:0000685 | 0,292082 |
| 949 | 6 | TISSUES | Leukemia cell | 0,0012 | STAT3\|ERE | 50406 | 4127\|4988\|3 | 2,35E-06 | BTO:0001271 | 0,292082 |
| 3 | 2 | TISSUES | BT-474 cell | 0,0012 | ERBB2\|EGI | 50406 | 4988\|5306 | 2,35E-06 | BTO:0001932 | 0,292082 |
| 3 | 2 | TISSUES | SK-BR-3 cell | 0,0012 | ERBB2\|EGI | 50406 | 4988\|5306 | 2,35E-06 | BTO:0002419 | 0,292082 |
| 624 | 5 | TISSUES | Epithelium | 0,0017 | ERBB2\|NO | 50406 | 4988\|3485\|5 | 7,43E-06 | BTO:0000416 | 0,276955 |
| 1229 | 6 | TISSUES | Fetus | 0,0022 | STAT3\|EGI | 50406 | 4127\|5306\|5 | 1,05E-05 | BTO:0000449 | 0,265758 |
| 9 | 2 | TISSUES | Macrophage cell line | 0,0026 | TLR4\|IL6 | 50406 | 4079\|5153 | 1,29E-05 | BTO:0002278 | 0,258503 |
| 9 | 2 | TISSUES | Monocytic leukemia cell line | 0,0026 | TLR4\|IL6 | 50406 | 4079\|5153 | 1,29E-05 | BTO:0002332 | 0,258503 |
| 361 | 4 | TISSUES | Lymphocytic leukemia cell | 0,0038 | NOTCH1\|I | 50406 | 3485\|5270\|3 | 2,28E-05 | BTO:0000744 | 0,242022 |
| 14 | 2 | TISSUES | Parenchyma | 0,0042 | FN1\|IL6 | 50406 | 5576\|5153 | 2,81E-05 | BTO:0001539 | 0,237675 |
| 129 | 3 | TISSUES | Epithelial cell line | 0,0045 | ERBB2\|EGI | 50406 | 4988\|5306\|5 | 3,47E-05 | BTO:0001120 | 0,234679 |
| 137 | 3 | TISSUES | Plasma cell | 0,0049 | FN1\|ITGB1 | 50406 | 5576\|5270\|3 | 4,14E-05 | BTO:0000392 | 0,23098 |
| 165 | 3 | TISSUES | Breast cancer cell | 0,008 | ERBB2\|FN1 | 50406 | 4988\|5576\|3 | 7,13E-05 | BTO:0000150 | 0,209691 |
| 170 | 3 | TISSUES | Endothelium | 0,008 | NOTCH1\|F | 50406 | 3485\|5576\|5 | 7,78E-05 | BTO:0000393 | 0,209691 |
| 482 | 4 | TISSUES | Mouth | 0,008 | ERBB2\|FN1 | 50406 | 4988\|5576\|5 | 7,00E-05 | BTO:0001090 | 0,209691 |
| 24 | 2 | TISSUES | Retinal pigment epithelium | 0,008 | FN1\|CD44 | 50406 | 5576\|3263 | 7,58E-05 | BTO:0001177 | 0,209691 |
| 193 | 3 | TISSUES | Chronic lymphocytic leukemia cell | 0,0101 | ITGB1\|CD4 | 50406 | 5270\|3263\|5 | 1,10E-04 | BTO:0001546 | 0,199568 |
| 32 | 2 | TISSUES | Aorta endothelium | 0,0114 | NOTCH1\|F | 50406 | 3485\|5576 | 1,30E-04 | BTO:0000394 | 0,19431 |
| 1162 | 5 | TISSUES | Lung | 0,0121 | FN1\|TLR4\| | 50406 | 5576\|4079\|5 | 1,50E-04 | BTO:0000763 | 0,191721 |
| 1970 | 6 | TISSUES | Integument | 0,0124 | ERBB2\|NO | 50406 | 4988\|3485\|5 | 1,50E-04 | BTO:0000634 | 0,190658 |
| 1176 | 5 | TISSUES | Placenta | 0,0124 | STAT3\|EGI | 50406 | 4127\|5306\|5 | 1,50E-04 | BTO:0001078 | 0,190658 |
| 1277 | 5 | TISSUES | Excretory gland | 0,0169 | STAT3\|ERE | 50406 | 4127\|4988\|5 | 2,30E-04 | BTO:0000431 | 0,177211 |
| 69 | 2 | TISSUES | Cartilage | 0,0384 | FN1\|CD44 | 50406 | 5576\|3263 | 5,70E-04 | BTO:0000206 | 0,141567 |
| 337 | 3 | TISSUES | Blood vessel | 0,0384 | NOTCH1\|F | 50406 | 3485\|5576\|5 | 5,70E-04 | BTO:0001102 | 0,141567 |
| 1553 | 5 | TISSUES | Immune system | 0,0384 | STAT3\|FN1 | 50406 | 4127\|5576\|4 | 5,70E-04 | BTO:0005810 | 0,141567 |
| 2543 | 6 | TISSUES | Hematopoietic system | 0,0394 | STAT3\|FN1 | 50406 | 4127\|5576\|4 | 6,40E-04 | BTO:0000570 | 0,14045 |
| 2593 | 6 | TISSUES | Internal female genital organ | 0,0429 | STAT3\|EGI | 50406 | 4127\|5306\|5 | 7,10E-04 | BTO:0003099 | 0,136754 |
| 1675 | 5 | TISSUES | Blood | 0,048 | FN1\|TLR4\| | 50406 | 5576\|4079\|5 | 8,10E-04 | BTO:0000089 | 0,131876 |
| 933 | 4 | TISSUES | Hematopoietic cell | 0,0484 | FN1\|TLR4\| | 50406 | 5576\|4079\|5 | 8,70E-04 | BTO:0000574 | 0,131515 |
| 941 | 4 | TISSUES | Kidney | 0,0484 | STAT3\|FN1 | 50406 | 4127\|5576\|4 | 9,00E-04 | BTO:0000671 | 0,131515 |
| 927 | 4 | TISSUES | Trunk | 0,0484 | ERBB2\|FN1 | 50406 | 4988\|5576\|5 | 8,50E-04 | BTO:0001493 | 0,131515 |
| 3233 | 9 | UniProt Keywords | Signal | 7,30E-04 | ERBB2\|EGI | 50406 | 4988\|5306\|3 | 7,90E-07 | KW-0732 | 0,313668 |
| 3304 | 9 | UniProt Keywords | Disulfide bond | 7,30E-04 | ERBB2\|EGI | 50406 | 4988\|5306\|3 | 9,57E-07 | KW-1015 | 0,313668 |
| 4349 | 9 | UniProt Keywords | Glycoprotein | 0,0033 | ERBB2\|EGI | 50406 | 4988\|5306\|3 | 1,07E-05 | KW-0325 | 0,248149 |
| 1422 | 6 | UniProt Keywords | Receptor | 0,0056 | ERBB2\|EGI | 50406 | 4988\|5306\|3 | 2,42E-05 | KW-0675 | 0,225181 |
| 19 | 2 | UniProt Keywords | Acute phase | 0,0091 | FN1\|IL6 | 50406 | 5576\|5153 | 4,91E-05 | KW-0011 | 0,204096 |
| 474 | 4 | UniProt Keywords | Cell adhesion | 0,0101 | FN1\|ITGB1 | 50406 | 5576\|5270\|3 | 6,56E-05 | KW-0130 | 0,199568 |
| 568 | 4 | UniProt Keywords | Endosome | 0,0174 | ERBB2\|EGI | 50406 | 4988\|5306\|4 | 1,30E-04 | KW-0967 | 0,175945 |
| 3246 | 7 | UniProt Keywords | Cell membrane | 0,0302 | ERBB2\|EGI | 50406 | 4988\|5306\|3 | 2,60E-04 | KW-1003 | 0,151999 |
| 67 | 5 | WikiPathways | RAC1/PAK1/p38/MMP2 pathway | 2,54E-07 | STAT3\|ERE | 50406 | 4127\|4988\|5 | 1,45E-10 | WP3303 | 0,659517 |
| 18 | 4 | WikiPathways | Extracellular vesicles in the crosstalk of cardiac cells | 2,54E-07 | STAT3\|EGI | 50406 | 4127\|5306\|4 | 2,50E-10 | WP4300 | 0,659517 |
| 83 | 4 | WikiPathways | EGFR tyrosine kinase inhibitor resistance | 4,36E-05 | STAT3\|ERE | 50406 | 4127\|4988\|5 | 7,49E-08 | WP4806 | 0,436051 |
| 114 | 4 | WikiPathways | Gastrin signaling pathway | 1,10E-04 | STAT3\|EGI | 50406 | 4127\|5306\|5 | 2,56E-07 | WP4659 | 0,395861 |
| 28 | 3 | WikiPathways | Nanoparticle-mediated activation of receptor signal | 1,30E-04 | EGFR\|FN1 | 50406 | 5306\|5576\|5 | 4,28E-07 | WP2643 | 0,388606 |
| 336 | 5 | WikiPathways | PI3K-Akt signaling pathway | 1,30E-04 | EGFR\|FN1 | 50406 | 5306\|5576\|4 | 3,65E-07 | WP4172 | 0,388606 |
| 196 | 4 | WikiPathways | Focal adhesion | 5,20E-04 | ERBB2\|EGI | 50406 | 4988\|5306\|5 | 2,12E-06 | WP306 | 0,3284 |
| 49 | 3 | WikiPathways | Hepatitis C and hepatocellular carcinoma | 5,20E-04 | STAT3\|IL6 | 50406 | 4127\|5153\|3 | 2,09E-06 | WP3646 | 0,3284 |
| 5 | 2 | WikiPathways | TCA cycle nutrient use and invasiveness of ovarian c | 9,60E-04 | STAT3\|EGI | 50406 | 4127\|5306 | 4,93E-06 | WP2868 | 0,301773 |
| 66 | 3 | WikiPathways | Physico-chemical features and toxicity-associated pa | 9,60E-04 | ERBB2\|EGI | 50406 | 4988\|5306\|5 | 4,94E-06 | WP3680 | 0,301773 |
| 76 | 3 | WikiPathways | Prolactin signaling pathway | 0,001 | STAT3\|ERE | 50406 | 4127\|4988\|5 | 7,44E-06 | WP2037 | 0,3 |
| 72 | 3 | WikiPathways | Primary focal segmental glomerulosclerosis (FSGS) | 0,001 | NOTCH1\|T | 50406 | 3485\|4079\|5 | 6,36E-06 | WP2572 | 0,3 |
| 72 | 3 | WikiPathways | Non-small cell lung cancer | 0,001 | STAT3\|ERE | 50406 | 4127\|4988\|5 | 6,36E-06 | WP4255 | 0,3 |
| 72 | 3 | WikiPathways | Head and neck squamous cell carcinoma | 0,001 | ERBB2\|EGI | 50406 | 4988\|5306\|3 | 6,36E-06 | WP4674 | 0,3 |
| 87 | 3 | WikiPathways | Pancreatic adenocarcinoma pathway | 0,0012 | STAT3\|ERE | 50406 | 4127\|4988\|5 | 1,10E-05 | WP4263 | 0,292082 |
| 86 | 3 | WikiPathways | Acute viral myocarditis | 0,0012 | STAT3\|TLR | 50406 | 4127\|4079\|5 | 1,07E-05 | WP4298 | 0,292082 |
| 105 | 3 | WikiPathways | Senescence and autophagy in cancer | 0,002 | FN1\|IL6\|C | 50406 | 5576\|5153\|3 | 1,91E-05 | WP615 | 0,269897 |
| 12 | 2 | WikiPathways | IL-10 anti-inflammatory signaling pathway | 0,0021 | STAT3\|IL6 | 50406 | 4127\|5153 | 2,13E-05 | WP4495 | 0,267778 |
| 116 | 3 | WikiPathways | Spinal cord injury | 0,0023 | EGFR\|TLR4 | 50406 | 5306\|4079\|5 | 2,55E-05 | WP2431 | 0,263827 |
| 13 | 2 | WikiPathways | Mammary gland development pathway - Puberty (S | 0,0023 | ERBB2\|FN1 | 50406 | 4988\|5576 | 2,46E-05 | WP2814 | 0,263827 |
| 13 | 2 | WikiPathways | ncRNAs involved in STAT3 signaling in hepatocellula | 0,0023 | STAT3\|IL6 | 50406 | 4127\|5153 | 2,46E-05 | WP4337 | 0,263827 |
| 119 | 3 | WikiPathways | Hippo-Merlin signaling dysregulation | 0,0023 | EGFR\|ITGE | 50406 | 5306\|5270\|3 | 2,74E-05 | WP4541 | 0,263827 |
| 15 | 2 | WikiPathways | FOXP3 in COVID-19 | 0,0024 | STAT3\|IL6 | 50406 | 4127\|5153 | 3,18E-05 | WP5063 | 0,261979 |
| 129 | 3 | WikiPathways | Ebola virus pathway in host | 0,0025 | EGFR\|TLR4 | 50406 | 5306\|4079\|5 | 3,47E-05 | WP4217 | 0,260206 |
| 17 | 2 | WikiPathways | Cells and molecules involved in local acute inflamma | 0,0028 | ITGB1\|IL6 | 50406 | 5270\|5153 | 4,00E-05 | WP4493 | 0,255284 |
| 19 | 2 | WikiPathways | Overview of nanoparticle effects | 0,0033 | FN1\|IL6 | 50406 | 5576\|5153 | 4,91E-05 | WP3287 | 0,248149 |
| 19 | 2 | WikiPathways | LTF danger signal response pathway | 0,0033 | TLR4\|IL6 | 50406 | 4079\|5153 | 4,91E-05 | WP4478 | 0,248149 |
| 20 | 2 | WikiPathways | BMP signaling in eyelid development | 0,0034 | EGFR\|NOT | 50406 | 5306\|3485 | 5,40E-05 | WP3927 | 0,246852 |
| 153 | 3 | WikiPathways | Breast cancer pathway | 0,0034 | ERBB2\|EGI | 50406 | 4988\|5306\|3 | 5,72E-05 | WP4262 | 0,246852 |
| 151 | 3 | WikiPathways | Hepatitis B infection | 0,0034 | STAT3\|TLR | 50406 | 4127\|4079\|5 | 5,50E-05 | WP4666 | 0,246852 |
| 162 | 3 | WikiPathways | EGF/EGFR signaling pathway | 0,0038 | STAT3\|ERE | 50406 | 4127\|4988\|5 | 6,76E-05 | WP437 | 0,242022 |
| 28 | 2 | WikiPathways | TLR4 signaling and tolerance | 0,0055 | TLR4\|IL6 | 50406 | 4079\|5153 | 1,00E-04 | WP3851 | 0,225964 |
| 28 | 2 | WikiPathways | Interactions between immune cells and microRNAs | 0,0055 | STAT3\|TLR | 50406 | 4127\|4079 | 1,00E-04 | WP4559 | 0,225964 |
| 30 | 2 | WikiPathways | Extracellular vesicle-mediated signaling in recipient | 0,0059 | ERBB2\|EGI | 50406 | 4988\|5306 | 1,20E-04 | WP2870 | 0,222915 |
| 31 | 2 | WikiPathways | Mammary gland development pathway - Pregnancy | 0,0061 | ERBB2\|EGI | 50406 | 4988\|5306 | 1,20E-04 | WP2817 | 0,221467 |
| 34 | 2 | WikiPathways | Hepatocyte growth factor receptor signaling | 0,0071 | STAT3\|ITG | 50406 | 4127\|5270 | 1,50E-04 | WP313 | 0,214874 |
| 37 | 2 | WikiPathways | miRNAs involvement in the immune response in sep | 0,0081 | TLR4\|IL6 | 50406 | 4079\|5153 | 1,70E-04 | WP4329 | 0,209151 |
| 40 | 2 | WikiPathways | Bladder cancer | 0,0092 | ERBB2\|EGI | 50406 | 4988\|5306 | 2,00E-04 | WP2828 | 0,203621 |
| 41 | 2 | WikiPathways | Fibrin complement receptor 3 signaling pathway | 0,0094 | TLR4\|IL6 | 50406 | 4079\|5153 | 2,10E-04 | WP4136 | 0,202687 |
| 43 | 2 | WikiPathways | IL-6 signaling pathway | 0,01 | STAT3\|IL6 | 50406 | 4127\|5153 | 2,30E-04 | WP364 | 0,2 |
| 47 | 2 | WikiPathways | Thymic stromal lymphopoietin (TSLP) signaling path | 0,0116 | STAT3\|IL6 | 50406 | 4127\|5153 | 2,70E-04 | WP2203 | 0,193554 |

| | | | | | | | | | | |
|---|---|---|---|---|---|---|---|---|---|---|
| 48 | 2 | WikiPathways | Pluripotent stem cell differentiation pathway | 0,0118 | NOTCH1\|II | 50406 | 3485\|5153 | 2,80E-04 | WP2848 | 0,192812 |
| 50 | 2 | WikiPathways | Photodynamic therapy-induced AP-1 survival signali | 0,0125 | EGFR\|IL6 | 50406 | 5306\|5153 | 3,10E-04 | WP3611 | 0,190309 |
| 54 | 2 | WikiPathways | Pathogenic Escherichia coli infection | 0,0142 | TLR4\|ITGB | 50406 | 4079\|5270 | 3,60E-04 | WP2272 | 0,184771 |
| 54 | 2 | WikiPathways | Hematopoietic stem cell differentiation | 0,0142 | NOTCH1\|II | 50406 | 3485\|5153 | 3,60E-04 | WP2849 | 0,184771 |
| 302 | 3 | WikiPathways | Focal adhesion: PI3K-Akt-mTOR-signaling pathway | 0,0157 | EGFR\|FN1 | 50406 | 5306\|5576\|5 | 4,10E-04 | WP3932 | 0,18041 |
| 61 | 2 | WikiPathways | Notch signaling pathway (Netpath) | 0,0168 | STAT3\|NO | 50406 | 4127\|3485 | 4,50E-04 | WP61 | 0,177469 |
| 63 | 2 | WikiPathways | Endometrial cancer | 0,0175 | ERBB2\|EG | 50406 | 4988\|5306 | 4,80E-04 | WP4155 | 0,175696 |
| 64 | 2 | WikiPathways | Pathways affected in adenoid cystic carcinoma | 0,0177 | ERBB2\|NO | 50406 | 4988\|3485 | 5,00E-04 | WP3651 | 0,175203 |
| 66 | 2 | WikiPathways | AGE/RAGE pathway | 0,0184 | STAT3\|EG | 50406 | 4127\|5306 | 5,30E-04 | WP2324 | 0,173518 |
| 71 | 2 | WikiPathways | Non-genomic actions of 1,25 dihydroxyvitamin D3 | 0,0207 | TLR4\|IL6 | 50406 | 4079\|5153 | 6,10E-04 | WP4341 | 0,168403 |
| 75 | 2 | WikiPathways | Leptin signaling pathway | 0,0226 | STAT3\|ER | 50406 | 4127\|4988 | 6,70E-04 | WP2034 | 0,164589 |
| 78 | 2 | WikiPathways | Regulatory circuits of the STAT3 signaling pathway | 0,024 | STAT3\|EG | 50406 | 4127\|5306 | 7,30E-04 | WP4538 | 0,161979 |
| 82 | 2 | WikiPathways | Glioblastoma signaling pathways | 0,0259 | ERBB2\|EG | 50406 | 4988\|5306 | 8,00E-04 | WP2261 | 0,15867 |
| 85 | 2 | WikiPathways | Clear cell renal cell carcinoma pathways | 0,0273 | STAT3\|EG | 50406 | 4127\|5306 | 8,60E-04 | WP4018 | 0,156384 |
| 88 | 2 | WikiPathways | Androgen receptor signaling pathway | 0,0287 | STAT3\|EG | 50406 | 4127\|5306 | 9,20E-04 | WP138 | 0,154212 |
| 88 | 2 | WikiPathways | Hair follicle development: cytodifferentiation - part | 0,0287 | EGFR\|NOT | 50406 | 5306\|3485 | 9,20E-04 | WP2840 | 0,154212 |
| 90 | 2 | WikiPathways | ErbB signaling pathway | 0,0289 | ERBB2\|EG | 50406 | 4988\|5306 | 9,60E-04 | WP673 | 0,15391 |
| 95 | 2 | WikiPathways | Small cell lung cancer | 0,0316 | FN1\|ITGB1 | 50406 | 5576\|5270 | 0,0011 | WP4658 | 0,150031 |
| 428 | 3 | WikiPathways | VEGFA-VEGFR2 signaling pathway | 0,033 | STAT3\|FN1 | 50406 | 4127\|5576\|5 | 0,0011 | WP3888 | 0,148149 |
| 100 | 2 | WikiPathways | Neural crest differentiation | 0,0337 | NOTCH1\|I | 50406 | 3485\|5270 | 0,0012 | WP2064 | 0,147237 |
| 103 | 2 | WikiPathways | Toll-like receptor signaling pathway | 0,0352 | TLR4\|IL6 | 50406 | 4079\|5153 | 0,0012 | WP75 | 0,145346 |
| 116 | 2 | WikiPathways | Embryonic stem cell pluripotency pathways | 0,0436 | STAT3\|EG | 50406 | 4127\|5306 | 0,0016 | WP3931 | 0,136051 |

Supplementary file 6: Protein-protein interaction network of upregulated genes; the top 10 most ranked hub genes are highlighted.

| node_name | MCC | DMNC | MNC | Degree | EPC | BottleNeck | EcCentricit | Closeness | Radiality | Betweenne | Stress | ClusteringCoefficient |
|---|---|---|---|---|---|---|---|---|---|---|---|---|
| CCK | 17 | 0,33284 | 6 | 9 | 14,304 | 26 | 0,03817 | 17,20952 | 2,26313 | 388,3333 | 608 | 0,19444 |
| GABRD | 12 | 0,28529 | 6 | 6 | 13,918 | 2 | 0,03817 | 15,12619 | 2,20813 | 149,3667 | 322 | 0,4 |
| SCN3B | 9 | 0,46346 | 3 | 6 | 14,091 | 35 | 0,05344 | 16,48333 | 2,33386 | 666,5333 | 1224 | 0,2 |
| GRIK1 | 20 | 0,38039 | 6 | 6 | 13,654 | 3 | 0,04453 | 14,46667 | 2,1767 | 91,63333 | 250 | 0,53333 |
| GRIK2 | 20 | 0,38039 | 6 | 6 | 13,527 | 4 | 0,04453 | 14,46667 | 2,1767 | 91,63333 | 250 | 0,53333 |
| CRHR1 | 6 | 0,30898 | 3 | 5 | 14,023 | 32 | 0,04453 | 15,98333 | 2,31814 | 507,6 | 914 | 0,2 |
| CALY | 8 | 0,37893 | 4 | 4 | 13,192 | 1 | 0,0334 | 12,69405 | 2,03525 | 2,46667 | 8 | 0,66667 |
| CACNG8 | 12 | 0,47366 | 4 | 4 | 12,7 | 1 | 0,03817 | 11,38571 | 1,94095 | 0,66667 | 2 | 0,83333 |
| CACNA1G | 12 | 0,47366 | 4 | 4 | 13,096 | 1 | 0,04453 | 13,13333 | 2,1374 | 17,06667 | 66 | 0,83333 |
| COL24A1 | 4 | 0,30779 | 2 | 4 | 10,495 | 9 | 0,04453 | 14,01667 | 2,20813 | 426 | 714 | 0,16667 |
| COL11A2 | 5 | 0,30898 | 3 | 4 | 8,711 | 6 | 0,03817 | 12,32619 | 2,03525 | 268 | 426 | 0,33333 |
| LY6H | 8 | 0,37893 | 4 | 4 | 13,02 | 1 | 0,0334 | 12,69405 | 2,03525 | 9,3 | 22 | 0,66667 |
| MYO15A | 3 | 0 | 1 | 3 | 5,992 | 4 | 0,0334 | 10,45357 | 1,82308 | 190 | 268 | 0 |
| NRGN | 3 | 0 | 1 | 3 | 10,048 | 3 | 0,0334 | 11,91071 | 2,02739 | 65,46667 | 128 | 0 |
| CARTPT | 4 | 0,30898 | 3 | 3 | 12,877 | 2 | 0,03817 | 13,20952 | 2,16098 | 12,66667 | 46 | 0,66667 |
| COL22A1 | 4 | 0,30898 | 3 | 3 | 8,358 | 1 | 0,03817 | 11,40952 | 2,00382 | 28 | 82 | 0,66667 |
| PKD2L1 | 3 | 0,30779 | 2 | 3 | 2,708 | 2 | 0,03053 | 3 | 0,0916 | 4 | 4 | 0,33333 |
| GRM2 | 3 | 0,30779 | 2 | 3 | 11,931 | 3 | 0,03817 | 12,05952 | 2,05097 | 73,13333 | 164 | 0,33333 |
| SOHLH1 | 3 | 0,30779 | 2 | 3 | 11,341 | 3 | 0,0334 | 12,61071 | 2,05097 | 128 | 244 | 0,33333 |
| SHISA8 | 6 | 0,46346 | 3 | 3 | 12,136 | 1 | 0,03817 | 10,88571 | 1,93309 | 0 | 0 | 1 |
| ACTL6B | 2 | 0,30779 | 2 | 2 | 10,769 | 1 | 0,0334 | 10,72738 | 1,95667 | 0 | 0 | 1 |
| TGM1 | 2 | 0,30779 | 2 | 2 | 2,23 | 1 | 0,0229 | 2 | 0,0458 | 0 | 0 | 1 |
| PDIA2 | 2 | 0 | 1 | 2 | 2,294 | 2 | 0,01527 | 2,5 | 0,12214 | 4 | 4 | 0 |
| PKD1 | 2 | 0,30779 | 2 | 2 | 2,523 | 1 | 0,01527 | 2,5 | 0,08142 | 0 | 0 | 1 |
| ATP4A | 2 | 0 | 1 | 2 | 8,399 | 2 | 0,0334 | 11,69405 | 2,01953 | 66 | 76 | 0 |
| CALB2 | 2 | 0 | 1 | 2 | 9,695 | 2 | 0,0334 | 12,37738 | 2,07454 | 44,33333 | 52 | 0 |
| KRT17 | 2 | 0,30779 | 2 | 2 | 2,234 | 1 | 0,0229 | 2 | 0,0458 | 0 | 0 | 1 |
| FGF8 | 2 | 0 | 1 | 2 | 8,25 | 1 | 0,02969 | 10,41349 | 1,88595 | 7,8 | 16 | 0 |
| SYCE1 | 2 | 0 | 1 | 2 | 6,634 | 2 | 0,02969 | 9,76349 | 1,80736 | 66 | 124 | 0 |
| NOXA1 | 2 | 0 | 1 | 2 | 2,298 | 2 | 0,01527 | 2,5 | 0,12214 | 4 | 4 | 0 |
| NEGR1 | 2 | 0 | 1 | 2 | 7,55 | 1 | 0,03817 | 10,69286 | 1,97238 | 16 | 18 | 0 |
| LPPR3 | 2 | 0 | 1 | 2 | 4,299 | 2 | 0,02969 | 8,49563 | 1,57948 | 66 | 92 | 0 |
| LOR | 2 | 0,30779 | 2 | 2 | 2,258 | 1 | 0,0229 | 2 | 0,0458 | 0 | 0 | 1 |
| SLC44A5 | 2 | 0 | 1 | 2 | 6,709 | 1 | 0,03817 | 10,05952 | 1,91738 | 2 | 2 | 0 |
| TCTE1 | 2 | 0 | 1 | 2 | 2,024 | 3 | 0,0229 | 2 | 0,08015 | 2 | 2 | 0 |
| SNCG | 2 | 0 | 1 | 2 | 8,749 | 2 | 0,04453 | 11,83333 | 2,09811 | 48 | 74 | 0 |
| TRPM2 | 2 | 0,30779 | 2 | 2 | 2,568 | 1 | 0,01527 | 2,5 | 0,08142 | 0 | 0 | 1 |
| COL28A1 | 2 | 0,30779 | 2 | 2 | 6,853 | 1 | 0,0334 | 9,37024 | 1,78379 | 0 | 0 | 1 |
| TAC3 | 2 | 0 | 1 | 2 | 9,268 | 2 | 0,0334 | 11,94405 | 2,03525 | 34 | 54 | 0 |
| CORO1A | 1 | 0 | 1 | 1 | 1,506 | 1 | 0,01527 | 1 | 0,0458 | 0 | 0 | 0 |
| PROC | 1 | 0 | 1 | 1 | 1,495 | 1 | 0,01527 | 1 | 0,0458 | 0 | 0 | 0 |
| TNNT2 | 1 | 0 | 1 | 1 | 1,495 | 1 | 0,01527 | 1 | 0,0458 | 0 | 0 | 0 |
| CHRNA10 | 1 | 0 | 1 | 1 | 3,75 | 1 | 0,02969 | 7,82897 | 1,56376 | 0 | 0 | 0 |
| MAST1 | 1 | 0 | 1 | 1 | 1,775 | 1 | 0,01145 | 1,5 | 0,0687 | 0 | 0 | 0 |
| CHTF18 | 1 | 0 | 1 | 1 | 1,481 | 1 | 0,01527 | 1 | 0,0458 | 0 | 0 | 0 |
| B4GALNT3 | 1 | 0 | 1 | 1 | 1,523 | 1 | 0,01527 | 1 | 0,0458 | 0 | 0 | 0 |
| LPCAT4 | 1 | 0 | 1 | 1 | 2,881 | 1 | 0,02672 | 6,75 | 1,32016 | 0 | 0 | 0 |
| KIAA1875 | 1 | 0 | 1 | 1 | 1,481 | 1 | 0,01527 | 1 | 0,0458 | 0 | 0 | 0 |
| P2RX2 | 1 | 0 | 1 | 1 | 2,097 | 1 | 0,01527 | 2 | 0,07125 | 0 | 0 | 0 |
| CORO6 | 1 | 0 | 1 | 1 | 1,506 | 1 | 0,01527 | 1 | 0,0458 | 0 | 0 | 0 |
| MICAL1 | 1 | 0 | 1 | 1 | 1,492 | 1 | 0,01527 | 1 | 0,0458 | 0 | 0 | 0 |
| RTEL1 | 1 | 0 | 1 | 1 | 1,481 | 1 | 0,01527 | 1 | 0,0458 | 0 | 0 | 0 |
| CYP2D6 | 1 | 0 | 1 | 1 | 5,155 | 1 | 0,02969 | 8,71349 | 1,76022 | 0 | 0 | 0 |
| NOXO1 | 1 | 0 | 1 | 1 | 1,899 | 1 | 0,01018 | 1,83333 | 0,10178 | 0 | 0 | 0 |
| PNMA5 | 1 | 0 | 1 | 1 | 1,506 | 1 | 0,01527 | 1 | 0,0458 | 0 | 0 | 0 |
| SEMA5B | 1 | 0 | 1 | 1 | 1,492 | 1 | 0,01527 | 1 | 0,0458 | 0 | 0 | 0 |
| CAMKV | 1 | 0 | 1 | 1 | 7,942 | 1 | 0,03817 | 10,82619 | 2,05882 | 0 | 0 | 0 |
| RTL1 | 1 | 0 | 1 | 1 | 1,506 | 1 | 0,01527 | 1 | 0,0458 | 0 | 0 | 0 |
| CYP2E1 | 1 | 0 | 1 | 1 | 3,909 | 1 | 0,02672 | 7,59603 | 1,54805 | 0 | 0 | 0 |
| AP3B2 | 1 | 0 | 1 | 1 | 1,783 | 1 | 0,01145 | 1,5 | 0,0687 | 0 | 0 | 0 |
| MSLNL | 1 | 0 | 1 | 1 | 1,523 | 1 | 0,01527 | 1 | 0,0458 | 0 | 0 | 0 |
| TMEM249 | 1 | 0 | 1 | 1 | 1,481 | 1 | 0,01527 | 1 | 0,0458 | 0 | 0 | 0 |
| AMY2A | 1 | 0 | 1 | 1 | 1,929 | 1 | 0,01018 | 1,83333 | 0,10178 | 0 | 0 | 0 |
| TARBP1 | 0 | 0 | 0 | 0 | 1 | 0 | 0 | 0 | 0 | 0 | 0 | 0 |
| HBQ1 | 0 | 0 | 0 | 0 | 1 | 0 | 0 | 0 | 0 | 0 | 0 | 0 |
| RASL10A | 0 | 0 | 0 | 0 | 1 | 0 | 0 | 0 | 0 | 0 | 0 | 0 |
| ORC6 | 0 | 0 | 0 | 0 | 1 | 0 | 0 | 0 | 0 | 0 | 0 | 0 |
| PRLHR | 0 | 0 | 0 | 0 | 1 | 0 | 0 | 0 | 0 | 0 | 0 | 0 |
| DUSP4 | 0 | 0 | 0 | 0 | 1 | 0 | 0 | 0 | 0 | 0 | 0 | 0 |
| SERPINF1 | 0 | 0 | 0 | 0 | 1 | 0 | 0 | 0 | 0 | 0 | 0 | 0 |
| ACY3 | 0 | 0 | 0 | 0 | 1 | 0 | 0 | 0 | 0 | 0 | 0 | 0 |
| ZFR2 | 0 | 0 | 0 | 0 | 1 | 0 | 0 | 0 | 0 | 0 | 0 | 0 |
| LTK | 0 | 0 | 0 | 0 | 1 | 0 | 0 | 0 | 0 | 0 | 0 | 0 |
| RAB17 | 0 | 0 | 0 | 0 | 1 | 0 | 0 | 0 | 0 | 0 | 0 | 0 |
| SLC22A9 | 0 | 0 | 0 | 0 | 1 | 0 | 0 | 0 | 0 | 0 | 0 | 0 |
| TRIM74 | 0 | 0 | 0 | 0 | 1 | 0 | 0 | 0 | 0 | 0 | 0 | 0 |

| | | | | | | | | | | | | |
|---|---|---|---|---|---|---|---|---|---|---|---|---|
| PSMG3 | 0 | 0 | 0 | 0 | 1 | 0 | 0 | 0 | 0 | 0 | 0 | 0 |
| TRIM54 | 0 | 0 | 0 | 0 | 1 | 0 | 0 | 0 | 0 | 0 | 0 | 0 |
| PGM2L1 | 0 | 0 | 0 | 0 | 1 | 0 | 0 | 0 | 0 | 0 | 0 | 0 |
| AKR1E2 | 0 | 0 | 0 | 0 | 1 | 0 | 0 | 0 | 0 | 0 | 0 | 0 |
| C1QTNF4 | 0 | 0 | 0 | 0 | 1 | 0 | 0 | 0 | 0 | 0 | 0 | 0 |
| NECAB2 | 0 | 0 | 0 | 0 | 1 | 0 | 0 | 0 | 0 | 0 | 0 | 0 |
| ABCC12 | 0 | 0 | 0 | 0 | 1 | 0 | 0 | 0 | 0 | 0 | 0 | 0 |
| LCN15 | 0 | 0 | 0 | 0 | 1 | 0 | 0 | 0 | 0 | 0 | 0 | 0 |
| RPRML | 0 | 0 | 0 | 0 | 1 | 0 | 0 | 0 | 0 | 0 | 0 | 0 |
| SLC16A8 | 0 | 0 | 0 | 0 | 1 | 0 | 0 | 0 | 0 | 0 | 0 | 0 |
| AHRR | 0 | 0 | 0 | 0 | 1 | 0 | 0 | 0 | 0 | 0 | 0 | 0 |
| NLRP3 | 0 | 0 | 0 | 0 | 1 | 0 | 0 | 0 | 0 | 0 | 0 | 0 |
| MAPK15 | 0 | 0 | 0 | 0 | 1 | 0 | 0 | 0 | 0 | 0 | 0 | 0 |
| FOXD4L3 | 0 | 0 | 0 | 0 | 1 | 0 | 0 | 0 | 0 | 0 | 0 | 0 |
| ASTL | 0 | 0 | 0 | 0 | 1 | 0 | 0 | 0 | 0 | 0 | 0 | 0 |
| RTP5 | 0 | 0 | 0 | 0 | 1 | 0 | 0 | 0 | 0 | 0 | 0 | 0 |
| SYT5 | 0 | 0 | 0 | 0 | 1 | 0 | 0 | 0 | 0 | 0 | 0 | 0 |
| ASGR2 | 0 | 0 | 0 | 0 | 1 | 0 | 0 | 0 | 0 | 0 | 0 | 0 |
| DLK2 | 0 | 0 | 0 | 0 | 1 | 0 | 0 | 0 | 0 | 0 | 0 | 0 |
| TRIM17 | 0 | 0 | 0 | 0 | 1 | 0 | 0 | 0 | 0 | 0 | 0 | 0 |
| AMY1A | 0 | 0 | 0 | 0 | 1 | 0 | 0 | 0 | 0 | 0 | 0 | 0 |
| LRRC26 | 0 | 0 | 0 | 0 | 1 | 0 | 0 | 0 | 0 | 0 | 0 | 0 |
| PTCH2 | 0 | 0 | 0 | 0 | 1 | 0 | 0 | 0 | 0 | 0 | 0 | 0 |
| LRRC73 | 0 | 0 | 0 | 0 | 1 | 0 | 0 | 0 | 0 | 0 | 0 | 0 |
| GNMT | 0 | 0 | 0 | 0 | 1 | 0 | 0 | 0 | 0 | 0 | 0 | 0 |
| KIF12 | 0 | 0 | 0 | 0 | 1 | 0 | 0 | 0 | 0 | 0 | 0 | 0 |
| CORT | 0 | 0 | 0 | 0 | 1 | 0 | 0 | 0 | 0 | 0 | 0 | 0 |
| TNFRSF25 | 0 | 0 | 0 | 0 | 1 | 0 | 0 | 0 | 0 | 0 | 0 | 0 |
| PLCH2 | 0 | 0 | 0 | 0 | 1 | 0 | 0 | 0 | 0 | 0 | 0 | 0 |
| SPEF1 | 0 | 0 | 0 | 0 | 1 | 0 | 0 | 0 | 0 | 0 | 0 | 0 |
| PRMT8 | 0 | 0 | 0 | 0 | 1 | 0 | 0 | 0 | 0 | 0 | 0 | 0 |
| CPNE9 | 0 | 0 | 0 | 0 | 1 | 0 | 0 | 0 | 0 | 0 | 0 | 0 |
| CCDC57 | 0 | 0 | 0 | 0 | 1 | 0 | 0 | 0 | 0 | 0 | 0 | 0 |
| DRP2 | 0 | 0 | 0 | 0 | 1 | 0 | 0 | 0 | 0 | 0 | 0 | 0 |
| CCDC64 | 0 | 0 | 0 | 0 | 1 | 0 | 0 | 0 | 0 | 0 | 0 | 0 |
| ZGLP1 | 0 | 0 | 0 | 0 | 1 | 0 | 0 | 0 | 0 | 0 | 0 | 0 |
| CHRNA2 | 0 | 0 | 0 | 0 | 1 | 0 | 0 | 0 | 0 | 0 | 0 | 0 |
| LYG1 | 0 | 0 | 0 | 0 | 1 | 0 | 0 | 0 | 0 | 0 | 0 | 0 |
| R3HDM1 | 0 | 0 | 0 | 0 | 1 | 0 | 0 | 0 | 0 | 0 | 0 | 0 |
| AGAP6 | 0 | 0 | 0 | 0 | 1 | 0 | 0 | 0 | 0 | 0 | 0 | 0 |
| CITED1 | 0 | 0 | 0 | 0 | 1 | 0 | 0 | 0 | 0 | 0 | 0 | 0 |
| PNCK | 0 | 0 | 0 | 0 | 1 | 0 | 0 | 0 | 0 | 0 | 0 | 0 |
| TBC1D26 | 0 | 0 | 0 | 0 | 1 | 0 | 0 | 0 | 0 | 0 | 0 | 0 |
| SLC10A5 | 0 | 0 | 0 | 0 | 1 | 0 | 0 | 0 | 0 | 0 | 0 | 0 |
| C6orf141 | 0 | 0 | 0 | 0 | 1 | 0 | 0 | 0 | 0 | 0 | 0 | 0 |
| CES4A | 0 | 0 | 0 | 0 | 1 | 0 | 0 | 0 | 0 | 0 | 0 | 0 |
| NLRP2 | 0 | 0 | 0 | 0 | 1 | 0 | 0 | 0 | 0 | 0 | 0 | 0 |
| BEX5 | 0 | 0 | 0 | 0 | 1 | 0 | 0 | 0 | 0 | 0 | 0 | 0 |
| DNASE1L2 | 0 | 0 | 0 | 0 | 1 | 0 | 0 | 0 | 0 | 0 | 0 | 0 |
| RNF112 | 0 | 0 | 0 | 0 | 1 | 0 | 0 | 0 | 0 | 0 | 0 | 0 |
| TBC1D3G | 0 | 0 | 0 | 0 | 1 | 0 | 0 | 0 | 0 | 0 | 0 | 0 |
| TGFBR3L | 0 | 0 | 0 | 0 | 1 | 0 | 0 | 0 | 0 | 0 | 0 | 0 |
| PNMA3 | 0 | 0 | 0 | 0 | 1 | 0 | 0 | 0 | 0 | 0 | 0 | 0 |
| MPV17L2 | 0 | 0 | 0 | 0 | 1 | 0 | 0 | 0 | 0 | 0 | 0 | 0 |
| ANKRD24 | 0 | 0 | 0 | 0 | 1 | 0 | 0 | 0 | 0 | 0 | 0 | 0 |

Supplementary file 6: Functional enrichment of the top 10 upregulated hub genes.

| # backgrou | # genes | category | description | FDR corrected p-value | genes | network.SI | nodes.SUID | p-value | term name | transferred FDR value |
|---|---|---|---|---|---|---|---|---|---|---|
| 782 | 7 | COMPARTMENTS | Integral component of plasma membrane | 5,08E-05 | CALY\|CACI | 1863 | 330\|384\|5 | 1,82E-08 | GOCC:0005887 | 0,761605 |
| 841 | 7 | COMPARTMENTS | Intrinsic component of plasma membrane | 5,08E-05 | CALY\|CACI | 1863 | 330\|384\|5 | 2,99E-08 | GOCC:0031226 | 0,761605 |
| 25 | 3 | COMPARTMENTS | Sodium channel complex | 2,90E-04 | CACNA1G\| | 1863 | 384\|486\|6 | 3,12E-07 | GOCC:0034706 | 0,627427 |
| 1456 | 7 | COMPARTMENTS | Integral component of membrane | 8,80E-04 | CALY\|CACI | 1863 | 330\|384\|5 | 1,26E-06 | GOCC:0016021 | 0,541924 |
| 1552 | 7 | COMPARTMENTS | Intrinsic component of membrane | 0,0011 | CALY\|CACI | 1863 | 330\|384\|5 | 1,94E-06 | GOCC:0031224 | 0,524737 |
| 499 | 5 | COMPARTMENTS | Synapse | 0,0012 | CALY\|GABI | 1863 | 330\|588\|3 | 2,51E-06 | GOCC:0045202 | 0,518034 |
| 235 | 4 | COMPARTMENTS | Postsynapse | 0,0017 | CALY\|GABI | 1863 | 330\|588\|5 | 4,30E-06 | GOCC:0098794 | 0,491206 |
| 248 | 4 | COMPARTMENTS | Ion channel complex | 0,0019 | CACNA1G\| | 1863 | 384\|588\|4 | 5,30E-06 | GOCC:0034702 | 0,482638 |
| 295 | 4 | COMPARTMENTS | Transmembrane transporter complex | 0,0032 | CACNA1G\| | 1863 | 384\|588\|4 | 1,04E-05 | GOCC:1902495 | 0,442485 |
| 97 | 3 | COMPARTMENTS | Postsynaptic membrane | 0,0042 | GABRD\|GF | 1863 | 588\|528\|6 | 1,51E-05 | GOCC:0045211 | 0,421539 |
| 335 | 4 | COMPARTMENTS | Transporter complex | 0,0043 | CACNA1G\| | 1863 | 384\|588\|4 | 1,71E-05 | GOCC:1990351 | 0,419726 |
| 17 | 2 | COMPARTMENTS | Voltage-gated sodium channel complex | 0,0093 | CACNA1G\| | 1863 | 384\|486 | 4,00E-05 | GOCC:0001518 | 0,360308 |
| 139 | 3 | COMPARTMENTS | Synaptic membrane | 0,0093 | GABRD\|GF | 1863 | 588\|528\|6 | 4,32E-05 | GOCC:0097060 | 0,360308 |
| 1045 | 5 | COMPARTMENTS | Cell junction | 0,0176 | CALY\|GABI | 1863 | 330\|588\|3 | 8,84E-05 | GOCC:0030054 | 0,311175 |
| 193 | 3 | COMPARTMENTS | Cation channel complex | 0,021 | CACNA1G\| | 1863 | 384\|486\|6 | 1,10E-04 | GOCC:0034703 | 0,29757 |
| 35 | 2 | COMPARTMENTS | Collagen trimer | 0,0271 | COL24A1\|( | 1863 | 660\|396 | 1,50E-04 | GOCC:0005581 | 0,277927 |
| 675 | 4 | COMPARTMENTS | Plasma membrane protein complex | 0,0419 | CACNA1G\| | 1863 | 384\|588\|4 | 2,50E-04 | GOCC:0098797 | 0,244363 |
| 440 | 6 | GO Biological Process | Regulation of membrane potential | 4,60E-04 | CACNA1G\| | 1863 | 384\|588\|4 | 2,63E-08 | GO:0042391 | 0,591891 |
| 1942 | 8 | GO Biological Process | System process | 0,0031 | CACNA1G\| | 1863 | 384\|396\|5 | 3,58E-07 | GO:0003008 | 0,44493 |
| 1352 | 7 | GO Biological Process | Nervous system process | 0,0044 | CACNA1G\| | 1863 | 384\|396\|5 | 7,62E-07 | GO:0050877 | 0,417956 |
| 45 | 3 | GO Biological Process | Regulation of membrane depolarization | 0,0055 | CACNA1G\| | 1863 | 384\|486\|3 | 1,64E-06 | GO:0003254 | 0,400768 |
| 1145 | 6 | GO Biological Process | Cell-cell signaling | 0,0175 | CACNA1G\| | 1863 | 384\|588\|4 | 6,98E-06 | GO:0007267 | 0,311614 |
| 7 | 2 | GO Biological Process | Regulation of atrial cardiac muscle cell membra | 0,0185 | CACNA1G\| | 1863 | 384\|486 | 8,44E-06 | GO:0060371 | 0,307333 |
| 8 | 2 | GO Biological Process | SA node cell action potential | 0,0205 | CACNA1G\| | 1863 | 384\|486 | 1,05E-05 | GO:0086015 | 0,299426 |
| 97 | 3 | GO Biological Process | Action potential | 0,0208 | CACNA1G\| | 1863 | 384\|486\|6 | 1,51E-05 | GO:0001508 | 0,298307 |
| 714 | 5 | GO Biological Process | Inorganic ion transmembrane transport | 0,0208 | CACNA1G\| | 1863 | 384\|588\|4 | 1,42E-05 | GO:0098660 | 0,298307 |
| 418 | 4 | GO Biological Process | Chemical synaptic transmission | 0,0437 | CACNA1G\| | 1863 | 384\|588\|5 | 4,03E-05 | GO:0007268 | 0,241123 |
| 137 | 3 | GO Biological Process | Multicellular organismal signaling | 0,0437 | CACNA1G\| | 1863 | 384\|486\|6 | 4,14E-05 | GO:0035637 | 0,241123 |
| 17 | 2 | GO Biological Process | Membrane depolarization during cardiac muscl | 0,0437 | CACNA1G\| | 1863 | 384\|486 | 4,00E-05 | GO:0086012 | 0,241123 |
| 26 | 4 | GO Cellular Component | Sodium channel complex | 2,30E-06 | CACNA1G\| | 1863 | 384\|486\|5 | 9,35E-10 | GO:0034706 | 1 |
| 289 | 5 | GO Cellular Component | Ion channel complex | 2,10E-04 | CACNA1G\| | 1863 | 384\|588\|4 | 1,75E-07 | GO:0034702 | 0,652289 |
| 1623 | 7 | GO Cellular Component | Integral component of plasma membrane | 0,0013 | CALY\|CACI | 1863 | 330\|384\|5 | 2,63E-06 | GO:0005887 | 0,511869 |
| 5 | 2 | GO Cellular Component | Kainate selective glutamate receptor complex | 0,0015 | GRIK1\|GRI | 1863 | 528\|642 | 4,93E-06 | GO:0032983 | 0,500846 |
| 1351 | 6 | GO Cellular Component | Synapse | 0,0049 | CALY\|CACI | 1863 | 330\|384\|5 | 1,81E-05 | GO:0045202 | 0,409665 |
| 17 | 2 | GO Cellular Component | Voltage-gated sodium channel complex | 0,0098 | CACNA1G\| | 1863 | 384\|486 | 4,00E-05 | GO:0001518 | 0,356275 |
| 547 | 4 | GO Cellular Component | Plasma membrane protein complex | 0,0254 | CACNA1G\| | 1863 | 384\|486\|5 | 1,10E-04 | GO:0098797 | 0,282918 |
| 643 | 4 | GO Cellular Component | Postsynapse | 0,0389 | CALY\|GABI | 1863 | 330\|588\|5 | 2,10E-04 | GO:0098794 | 0,250086 |
| 1366 | 5 | GO Cellular Component | Neuron projection | 0,0452 | CACNA1G\| | 1863 | 384\|588\|3 | 3,10E-04 | GO:0043005 | 0,238524 |
| 281 | 3 | GO Cellular Component | Postsynaptic membrane | 0,0459 | GABRD\|GF | 1863 | 588\|528\|6 | 3,40E-04 | GO:0045211 | 0,23734 |
| 55 | 2 | GO Cellular Component | Terminal bouton | 0,0478 | CCK\|GRIK2 | 1863 | 390\|642 | 3,70E-04 | GO:0043195 | 0,234216 |
| 44 | 4 | GO Molecular Function | Sodium channel activity | 3,45E-05 | CACNA1G\| | 1863 | 384\|486\|5 | 6,61E-09 | GO:0005272 | 0,791409 |
| 332 | 5 | GO Molecular Function | Gated channel activity | 9,00E-04 | CACNA1G\| | 1863 | 384\|588\|4 | 3,45E-07 | GO:0022836 | 0,540193 |
| 425 | 5 | GO Molecular Function | Ion channel activity | 0,0015 | CACNA1G\| | 1863 | 384\|588\|4 | 1,15E-06 | GO:0005216 | 0,500846 |
| 53 | 3 | GO Molecular Function | Transmitter-gated ion channel activity involved | 0,002 | GABRD\|GF | 1863 | 588\|528\|6 | 2,62E-06 | GO:1904315 | 0,478687 |
| 5 | 2 | GO Molecular Function | Kainate selective glutamate receptor activity | 0,0023 | GRIK1\|GRI | 1863 | 528\|642 | 4,93E-06 | GO:0015277 | 0,467922 |
| 23 | 2 | GO Molecular Function | Voltage-gated sodium channel activity | 0,0159 | CACNA1G\| | 1863 | 384\|486 | 7,00E-05 | GO:0005248 | 0,318999 |
| 330 | 5 | KEGG Pathways | Neuroactive ligand-receptor interaction | 1,50E-04 | GABRD\|CC | 1863 | 588\|390\|5 | 3,35E-07 | hsa04080 | 0,678206 |
| 2 | 2 | Reactome Pathways | Activation of Na-permeable kainate receptors | 0,0245 | GRIK1\|GRI | 1863 | 528\|642 | 1,41E-06 | HSA-451307 | 0,285696 |
| 10 | 2 | SMART Domains | Fibrillar collagens C-terminal domain | 0,0196 | COL24A1\|( | 1863 | 660\|396 | 1,55E-05 | SM00038 | 0,302884 |
| 23 | 2 | SMART Domains | Thrombospondin N-terminal -like domains. | 0,0443 | COL24A1\|( | 1863 | 660\|396 | 7,00E-05 | SM00210 | 0,240073 |
| 346 | 5 | UniProt Keywords | Ion channel | 3,90E-04 | CACNA1G\| | 1863 | 384\|588\|4 | 4,22E-07 | KW-0407 | 0,604606 |
| 4349 | 9 | UniProt Keywords | Glycoprotein | 0,0034 | CALY\|CACI | 1863 | 330\|384\|6 | 1,07E-05 | KW-0325 | 0,437815 |
| 3233 | 8 | UniProt Keywords | Signal | 0,0042 | COL24A1\|( | 1863 | 660\|396\|5 | 1,84E-05 | KW-0732 | 0,421539 |
| 144 | 3 | UniProt Keywords | Postsynaptic cell membrane | 0,0089 | GABRD\|GF | 1863 | 588\|528\|6 | 4,79E-05 | KW-0628 | 0,363695 |
| 21 | 2 | UniProt Keywords | RNA editing | 0,0091 | GRIK1\|GRI | 1863 | 528\|642 | 5,91E-05 | KW-0691 | 0,361983 |
