## Supplementary tables for "Transcriptome profiling of the dorsomedial prefrontal cortex in suicide victims": S7_Top_DEGs_DisGeNET.pdf

Supplementary file 7: Depression focused gene set enrichment of top DEGs .

| Gene | Disease | Disease ID | Disease Class |
|---|---|---|---|
| ND6 | Anxiety | C0003467 | Behavior and Behavior Mechanisms |
| IL6 | | C0003467 | |
| ND5 | | C0003467 | |
| CARTPT | | C0003467 | |
| IL6 | Bipolar Disorder | C0005586 | Mental Disorders |
| P2RX2 | | C0005586 | |
| CARTPT | | C0005586 | |
| IL6 | Depression, Postpartum | C0221074 | Mental Disorders |
| SERPINA3 | | C0221074 | |
| CSF3 | Depressive disorder | C0011581 | Mental Disorders |
| ND6 | | C0011581 | |
| IL6 | | C0011581 | |
| SERPINA3 | | C0011581 | |
| ND5 | | C0011581 | |
| P2RX2 | | C0011581 | |
| ATP4A | | C0011581 | |
| CARTPT | | C0011581 | |
| IL6 | Depressive Symptoms | C0086132 | Behavior and Behavior Mechanisms |
| SERPINA3 | | C0086132 | |
| P2RX2 | | C0086132 | |
| PROC | | C0086132 | |
| CARTPT | | C0086132 | |
| ND6 | Developmental regression | C1836830 | Mental Disorders |
| ND5 | | C1836830 | |
| IL6 | Eating Disorders | C0013473 | Mental Disorders |
| SERPINA3 | | C0013473 | |
| CARTPT | | C0013473 | |
| CSF3 | Impaired cognition | C0338656 | Mental Disorders |
| IL6 | | C0338656 | |
| SERPINA3 | | C0338656 | |
| ND5 | | C0338656 | |
| P2RX2 | | C0338656 | |
| CSF3 | Major Depressive Disorder | C1269683 | Mental Disorders |
| IL1R2 | | C1269683 | |
| IL6 | | C1269683 | |
| P2RX2 | | C1269683 | |
| CARTPT | | C1269683 | |
| IL1R2 | | C0036341 | |

| | | | |
|---|---|---|---|
| IL6 | Schizophrenia | C0036341 | Mental Disorders |
| SERPINA3 | | C0036341 | |
| ND5 | | C0036341 | |
| P2RX2 | | C0036341 | |
| CARTPT | | C0036341 | |

| Gene | Gene_id | Disease | Disease_id | Type | Disease_Class | Semantic_Type | N_genes_d | N_SNPs_d | Score_gda | EI_gda | N_PMIDs | N_SNPs_gda | First_Ref | Last_Ref |
|---|---|---|---|---|---|---|---|---|---|---|---|---|---|---|
| CSF3 | 1440 | Addictive E | C0085281 | phenotype | Behavior and Behavior Mechanisms | Mental or Behavioral Dysfunction | 332 | 56 | 0,01 | 1 | 1 | 0 | 2018 | 2018 |
| CSF3 | 1440 | Depressed | C0344315 | phenotype | Behavior and Behavior Mechanisms | Mental or Behavioral Dysfunction | 1461 | 269 | 0,01 | 1 | 1 | 0 | 2017 | 2017 |
| CSF3 | 1440 | Mental De | C0011570 | disease | Behavior and Behavior Mechanisms | Mental or Behavioral Dysfunction | 1478 | 271 | 0,01 | 1 | 1 | 0 | 2017 | 2017 |
| CSF3 | 1440 | Depressior | C0282126 | disease | Mental Disorders | Mental or Behavioral Dysfunction | 41 | 0 | 0,3 | 1 | 1 | 0 | 1996 | 1996 |
| CSF3 | 1440 | Depressive | C0011581 | disease | Mental Disorders | Mental or Behavioral Dysfunction | 1719 | 297 | 0,31 | 1 | 2 | 0 | 1996 | 2017 |
| CSF3 | 1440 | Depressive | C0086133 | disease | Mental Disorders | Mental or Behavioral Dysfunction | 45 | 3 | 0,3 | 1 | 1 | 0 | 1996 | 1996 |
| CSF3 | 1440 | Endogenou | C0011573 | disease | Mental Disorders | Mental or Behavioral Dysfunction | 53 | 0 | 0,3 | 1 | 1 | 0 | 1996 | 1996 |
| CSF3 | 1440 | Impaired c | C0338656 | disease | Mental Disorders | Mental or Behavioral Dysfunction | 1630 | 348 | 0,01 | 1 | 1 | 0 | 2017 | 2017 |
| CSF3 | 1440 | Major Dep | C1269683 | disease | Mental Disorders | Mental or Behavioral Dysfunction | 1236 | 1451 | 0,01 | 1 | 1 | 0 | 2019 | 2019 |
| CSF3 | 1440 | Melanchol | C0025193 | disease | Mental Disorders | Mental or Behavioral Dysfunction | 51 | 8 | 0,3 | 1 | 1 | 0 | 1996 | 1996 |
| CSF3 | 1440 | Post-Traun | C0038436 | disease | Mental Disorders | Mental or Behavioral Dysfunction | 418 | 117 | 0,01 | 1 | 1 | 0 | 2017 | 2017 |
| CSF3 | 1440 | Unipolar D | C0041696 | disease | Mental Disorders | Mental or Behavioral Dysfunction | 641 | 225 | 0,3 | 1 | 1 | 0 | 1996 | 1996 |
| IL1R2 | 7850 | Anxiety ne | C1279420 | disease | Mental Disorders | Mental or Behavioral Dysfunction | 75 | 5 | 0,01 | 1 | 1 | 0 | 2015 | 2015 |
| IL1R2 | 7850 | Major Dep | C1269683 | disease | Mental Disorders | Mental or Behavioral Dysfunction | 1236 | 1451 | 0,01 | 1 | 1 | 0 | 2016 | 2016 |
| IL1R2 | 7850 | Schizophre | C0036341 | disease | Mental Disorders | Mental or Behavioral Dysfunction | 2872 | 2897 | 0,01 | 1 | 1 | 0 | 2017 | 2017 |
| IL1R2 | 7850 | Anxiety sta | C0700613 | disease | Mental Disorders; Behavior and Behavior Mechanisms | Mental or Behavioral Dysfunction | 40 | 6 | 0,01 | 1 | 1 | 0 | 2015 | 2015 |
| ND6 | 4541 | Anxiety | C0003467 | disease | Behavior and Behavior Mechanisms | Mental or Behavioral Dysfunction | 1048 | 287 | 0,11 | 1 | 1 | 0 | 2015 | 2015 |
| ND6 | 4541 | Autistic Dis | C0004352 | disease | Mental Disorders | Mental or Behavioral Dysfunction | 1112 | 395 | 0,1 | | 0 | 0 | | |
| ND6 | 4541 | Depressive | C0011581 | disease | Mental Disorders | Mental or Behavioral Dysfunction | 1719 | 297 | 0,1 | | 0 | 0 | | |
| ND6 | 4541 | Psychotic e | C0338614 | disease | Mental Disorders | Mental or Behavioral Dysfunction | 31 | 6 | 0,1 | | 0 | 0 | | |
| ND6 | 4541 | Attention c | C1263846 | disease | Mental Disorders | Mental or Behavioral Dysfunction | 842 | 420 | 0,1 | | 0 | 0 | | |
| ND6 | 4541 | Developme | C1836830 | disease | Mental Disorders | Disease or Syndrome | 333 | 80 | 0,1 | | 0 | 0 | | |
| ND6 | 4541 | Anxiety Dis | C0003469 | group | Mental Disorders | Mental or Behavioral Dysfunction | 840 | 163 | 0,01 | 1 | 1 | 0 | 2015 | 2015 |
| ND6 | 4541 | Anxiety ne | C1279420 | disease | Mental Disorders | Mental or Behavioral Dysfunction | 75 | 5 | 0,01 | 1 | 1 | 0 | 2015 | 2015 |
| ND6 | 4541 | Anxiety sta | C0700613 | disease | Mental Disorders; Behavior and Behavior Mechanisms | Mental or Behavioral Dysfunction | 40 | 6 | 0,01 | 1 | 1 | 0 | 2015 | 2015 |
| IL6 | 3569 | Mental De | C0011570 | disease | Behavior and Behavior Mechanisms | Mental or Behavioral Dysfunction | 1478 | 271 | 0,6 | 0,8 | 108 | 1 | 1995 | 2020 |
| IL6 | 3569 | Anxiety | C0003467 | disease | Behavior and Behavior Mechanisms | Mental or Behavioral Dysfunction | 1048 | 287 | 0,1 | 1 | 21 | 0 | 2013 | 2020 |
| IL6 | 3569 | Depressive | C0086132 | phenotype | Behavior and Behavior Mechanisms | Sign or Symptom | 421 | 120 | 0,1 | 0,92 | 50 | 1 | 2009 | 2020 |
| IL6 | 3569 | Depressed | C0344315 | phenotype | Behavior and Behavior Mechanisms | Mental or Behavioral Dysfunction | 1461 | 269 | 0,1 | 0,935 | 108 | 1 | 1995 | 2020 |
| IL6 | 3569 | Stress, Psy | C0038443 | disease | Behavior and Behavior Mechanisms | Mental or Behavioral Dysfunction | 199 | 24 | 0,09 | 1 | 9 | 0 | 2006 | 2019 |
| IL6 | 3569 | Abnormal I | C0233514 | phenotype | Behavior and Behavior Mechanisms | Mental or Behavioral Dysfunction | 910 | 121 | 0,08 | 0,75 | 8 | 2 | 2015 | 2020 |
| IL6 | 3569 | social stres | C0871388 | disease | Behavior and Behavior Mechanisms | Mental or Behavioral Dysfunction | 45 | 22 | 0,03 | 1 | 3 | 0 | 2006 | 2019 |
| IL6 | 3569 | NEUROTIC | C1842981 | disease | Behavior and Behavior Mechanisms | Mental or Behavioral Dysfunction | 141 | 54 | 0,03 | 1 | 3 | 0 | 2017 | 2019 |
| IL6 | 3569 | Alexithymi | C0002020 | phenotype | Behavior and Behavior Mechanisms | Mental or Behavioral Dysfunction | 39 | 12 | 0,01 | 1 | 1 | 0 | 2019 | 2019 |
| IL6 | 3569 | Behavioral | C0004941 | group | Behavior and Behavior Mechanisms | Sign or Symptom | 45 | 9 | 0,01 | 1 | 1 | 0 | 2019 | 2019 |
| IL6 | 3569 | Hunger | C0020175 | phenotype | Behavior and Behavior Mechanisms | Sign or Symptom | 70 | 12 | 0,01 | 1 | 1 | 0 | 2019 | 2019 |
| IL6 | 3569 | Emotional | C0086209 | phenotype | Behavior and Behavior Mechanisms | Mental or Behavioral Dysfunction | 25 | 1 | 0,01 | 1 | 1 | 0 | 2018 | 2018 |
| IL6 | 3569 | Feeling des | C0233488 | phenotype | Behavior and Behavior Mechanisms | Sign or Symptom | 21 | 3 | 0,01 | 1 | 1 | 0 | 2019 | 2019 |
| IL6 | 3569 | Depressior | C0221074 | disease | Female Urogenital Diseases and Pregnancy Complications; Me | Mental or Behavioral Dysfunction | 54 | 6 | 0,01 | 1 | 1 | 0 | 2016 | 2016 |
| IL6 | 3569 | Depressive | C0011581 | disease | Mental Disorders | Mental or Behavioral Dysfunction | 1719 | 297 | 0,6 | 0,6 | 108 | 1 | 1995 | 2020 |
| IL6 | 3569 | Unipolar D | C0041696 | disease | Mental Disorders | Mental or Behavioral Dysfunction | 641 | 225 | 0,6 | 1 | 25 | 0 | 1994 | 2019 |
| IL6 | 3569 | Bipolar Dis | C0005586 | disease | Mental Disorders | Mental or Behavioral Dysfunction | 1183 | 839 | 0,4 | 1 | 18 | 0 | 2013 | 2019 |
| IL6 | 3569 | Schizophre | C0036341 | disease | Mental Disorders | Mental or Behavioral Dysfunction | 2872 | 2897 | 0,4 | 0,95 | 40 | 1 | 1994 | 2020 |
| IL6 | 3569 | Major Dep | C1269683 | disease | Mental Disorders | Mental or Behavioral Dysfunction | 1236 | 1451 | 0,4 | 1 | 57 | 1 | 1994 | 2020 |
| IL6 | 3569 | Autistic Dis | C0004352 | disease | Mental Disorders | Mental or Behavioral Dysfunction | 1112 | 395 | 0,39 | 0,9 | 10 | 0 | 1996 | 2020 |
| IL6 | 3569 | Endogenou | C0011573 | disease | Mental Disorders | Mental or Behavioral Dysfunction | 53 | 0 | 0,3 | 1 | 1 | 0 | 2009 | 2009 |
| IL6 | 3569 | Melanchol | C0025193 | disease | Mental Disorders | Mental or Behavioral Dysfunction | 51 | 8 | 0,3 | 1 | 1 | 0 | 2009 | 2009 |
| IL6 | 3569 | Depressive | C0086133 | disease | Mental Disorders | Mental or Behavioral Dysfunction | 45 | 3 | 0,3 | 1 | 1 | 0 | 2009 | 2009 |
| IL6 | 3569 | Depressior | C0282126 | disease | Mental Disorders | Mental or Behavioral Dysfunction | 41 | 0 | 0,3 | 1 | 1 | 0 | 2009 | 2009 |
| IL6 | 3569 | Anxiety Dis | C0003469 | group | Mental Disorders | Mental or Behavioral Dysfunction | 840 | 163 | 0,1 | 1 | 22 | 0 | 2013 | 2020 |
| IL6 | 3569 | Post-Traun | C0038436 | disease | Mental Disorders | Mental or Behavioral Dysfunction | 418 | 117 | 0,1 | 1 | 18 | 1 | 2008 | 2020 |
| IL6 | 3569 | Mental det | C0234985 | phenotype | Mental Disorders | Mental or Behavioral Dysfunction | 508 | 121 | 0,1 | 0,857 | 14 | 0 | 2003 | 2019 |
| IL6 | 3569 | Impaired c | C0338656 | disease | Mental Disorders | Mental or Behavioral Dysfunction | 1630 | 348 | 0,1 | 0,919 | 37 | 1 | 2003 | 2020 |
| IL6 | 3569 | Autism Spe | C1510586 | disease | Mental Disorders | Mental or Behavioral Dysfunction | 1071 | 331 | 0,1 | 1 | 15 | 0 | 2017 | 2020 |
| IL6 | 3569 | Psychotic I | C0033975 | group | Mental Disorders | Mental or Behavioral Dysfunction | 560 | 179 | 0,09 | 0,889 | 9 | 0 | 2014 | 2019 |
| IL6 | 3569 | Nonorgani | C0349204 | disease | Mental Disorders | Mental or Behavioral Dysfunction | 376 | 98 | 0,08 | 0,875 | 8 | 0 | 2014 | 2019 |
| IL6 | 3569 | Mental dis | C0004936 | group | Mental Disorders | Mental or Behavioral Dysfunction | 789 | 149 | 0,07 | 0,857 | 7 | 2 | 2015 | 2019 |
| IL6 | 3569 | Mild cognit | C1270972 | disease | Mental Disorders | Mental or Behavioral Dysfunction | 430 | 96 | 0,07 | 0,714 | 7 | 0 | 2017 | 2019 |
| IL6 | 3569 | Mood Diso | C0525045 | group | Mental Disorders | Mental or Behavioral Dysfunction | 580 | 308 | 0,06 | 0,833 | 6 | 0 | 2008 | 2019 |
| IL6 | 3569 | Cognition I | C0009241 | group | Mental Disorders | Mental or Behavioral Dysfunction | 607 | 47 | 0,04 | 1 | 4 | 1 | 2016 | 2019 |
| IL6 | 3569 | Disruptive, | C0021122 | group | Mental Disorders | Mental or Behavioral Dysfunction | 67 | 9 | 0,04 | 1 | 4 | 0 | 2018 | 2019 |
| IL6 | 3569 | Mixed anxi | C0338908 | disease | Mental Disorders | Mental or Behavioral Dysfunction | 146 | 13 | 0,04 | 1 | 4 | 0 | 2013 | 2019 |
| IL6 | 3569 | Paranoid S | C0036349 | disease | Mental Disorders | Mental or Behavioral Dysfunction | 53 | 23 | 0,03 | 1 | 3 | 0 | 2010 | 2013 |
| IL6 | 3569 | Difficulty sl | C0235162 | phenotype | Mental Disorders | Sign or Symptom | 40 | 4 | 0,03 | 1 | 3 | 0 | 2017 | 2019 |
| IL6 | 3569 | Manic | C0338831 | disease | Mental Disorders | Mental or Behavioral Dysfunction | 166 | 8 | 0,03 | 1 | 3 | 0 | 2017 | 2018 |
| IL6 | 3569 | Diagnosis, | C0376338 | disease | Mental Disorders | Mental or Behavioral Dysfunction | 46 | 1 | 0,03 | 1 | 3 | 0 | 2017 | 2019 |
| IL6 | 3569 | Depressive | C2063866 | disease | Mental Disorders | Mental or Behavioral Dysfunction | 66 | 19 | 0,03 | 1 | 3 | 0 | 2017 | 2019 |
| IL6 | 3569 | Anorexia N | C0003125 | disease | Mental Disorders | Mental or Behavioral Dysfunction | 202 | 72 | 0,02 | 1 | 2 | 0 | 2001 | 2020 |
| IL6 | 3569 | Obsessive- | C0028768 | disease | Mental Disorders | Mental or Behavioral Dysfunction | 175 | 112 | 0,02 | 1 | 2 | 0 | 2019 | 2019 |
| IL6 | 3569 | Panic Disor | C0030319 | disease | Mental Disorders | Mental or Behavioral Dysfunction | 175 | 101 | 0,02 | 1 | 2 | 0 | 2017 | 2019 |
| IL6 | 3569 | Chronic sch | C0221765 | disease | Mental Disorders | Mental or Behavioral Dysfunction | 48 | 7 | 0,02 | 1 | 2 | 0 | 2015 | 2019 |
| IL6 | 3569 | Developme | C0424605 | phenotype | Mental Disorders | Mental or Behavioral Dysfunction | 584 | 68 | 0,02 | 1 | 2 | 0 | 2017 | 2018 |
| IL6 | 3569 | Severe dep | C0588008 | disease | Mental Disorders | Mental or Behavioral Dysfunction | 46 | 2 | 0,02 | 1 | 2 | 0 | 2018 | 2019 |
| IL6 | 3569 | Attention c | C1263846 | disease | Mental Disorders | Mental or Behavioral Dysfunction | 842 | 420 | 0,02 | 1 | 2 | 0 | 2018 | 2019 |
| IL6 | 3569 | Disturbanc | C2939186 | disease | Mental Disorders | Mental or Behavioral Dysfunction | 26 | 2 | 0,02 | 1 | 2 | 0 | 2012 | 2018 |
| IL6 | 3569 | Developme | C0008073 | group | Mental Disorders | Mental or Behavioral Dysfunction | 355 | 19 | 0,01 | 1 | 1 | 0 | 2011 | 2011 |
| IL6 | 3569 | Eating Diso | C0013473 | group | Mental Disorders | Mental or Behavioral Dysfunction | 133 | 42 | 0,01 | 1 | 1 | 0 | 2018 | 2018 |
| IL6 | 3569 | Personality | C0031212 | group | Mental Disorders | Mental or Behavioral Dysfunction | 49 | 8 | 0,01 | 1 | 1 | 0 | 2019 | 2019 |
| IL6 | 3569 | Attention I | C0041671 | disease | Mental Disorders | Mental or Behavioral Dysfunction | 123 | 7 | 0,01 | 1 | 1 | 0 | 2018 | 2018 |
| IL6 | 3569 | Seasonal A | C0085159 | disease | Mental Disorders | Mental or Behavioral Dysfunction | 57 | 17 | 0,01 | 1 | 1 | 0 | 2017 | 2017 |
| IL6 | 3569 | Atypical de | C0154437 | disease | Mental Disorders | Mental or Behavioral Dysfunction | 7 | 2 | 0,01 | 1 | 1 | 0 | 2018 | 2018 |
| IL6 | 3569 | Ruminatio | C0154575 | group | Mental Disorders | Mental or Behavioral Dysfunction | 33 | 13 | 0,01 | 1 | 1 | 0 | 2018 | 2018 |
| IL6 | 3569 | Coprophili | C0270500 | disease | Mental Disorders | Mental or Behavioral Dysfunction | 12 | 0 | 0,01 | 1 | 1 | 0 | 2014 | 2014 |
| IL6 | 3569 | Acute psyc | C0281774 | disease | Mental Disorders | Mental or Behavioral Dysfunction | 3 | 0 | 0,01 | 1 | 1 | 0 | 2017 | 2017 |
| IL6 | 3569 | Recurrent | C0349218 | disease | Mental Disorders | Mental or Behavioral Dysfunction | 29 | 9 | 0,01 | 0 | 1 | 0 | 2017 | 2017 |
| IL6 | 3569 | Postpartun | C0520678 | disease | Mental Disorders | Mental or Behavioral Dysfunction | 19 | 0 | 0,01 | 1 | 1 | 0 | 2019 | 2019 |
| IL6 | 3569 | Mild depre | C0588006 | disease | Mental Disorders | Mental or Behavioral Dysfunction | 8 | 0 | 0,01 | 1 | 1 | 0 | 2018 | 2018 |
| IL6 | 3569 | Bipolar I di | C0853193 | disease | Mental Disorders | Mental or Behavioral Dysfunction | 83 | 46 | 0,01 | 1 | 1 | 0 | 2019 | 2019 |
| IL6 | 3569 | Anxiety ne | C1279420 | disease | Mental Disorders | Mental or Behavioral Dysfunction | 75 | 5 | 0,01 | 1 | 1 | 0 | 2018 | 2018 |
| IL6 | 3569 | Neurodeve | C1535926 | group | Mental Disorders | Mental or Behavioral Dysfunction | 535 | 14 | 0,01 | 1 | 1 | 0 | 2018 | 2018 |
| IL6 | 3569 | clinical dep | C2362914 | disease | Mental Disorders | Mental or Behavioral Dysfunction | 14 | 0 | 0,01 | 1 | 1 | 0 | 2016 | 2016 |
| IL6 | 3569 | Post stroke | C2938940 | disease | Mental Disorders | Disease or Syndrome | 39 | 23 | 0,01 | 1 | 1 | 0 | 2018 | 2018 |
| IL6 | 3569 | Memory d | C3887551 | disease | Mental Disorders | Mental or Behavioral Dysfunction | 70 | 3 | 0,01 | 1 | 1 | 0 | 2017 | 2017 |
| IL6 | 3569 | Neurocogr | C4041080 | group | Mental Disorders | Mental or Behavioral Dysfunction | 79 | 0 | 0,01 | 1 | 1 | 0 | 2018 | 2018 |
| IL6 | 3569 | Depressive | C0349217 | disease | Mental Disorders; Behavior and Behavior Mechanisms | Mental or Behavioral Dysfunction | 27 | 2 | 0,02 | 0,5 | 2 | 0 | 2016 | 2020 |
| IL6 | 3569 | Depressior | C0235876 | phenotype | Mental Disorders; Behavior and Behavior Mechanisms | Sign or Symptom | 1 | 0 | 0,01 | 1 | 1 | 0 | 2018 | 2018 |
| IL6 | 3569 | Anxiety sta | C0700613 | disease | Mental Disorders; Behavior and Behavior Mechanisms | Mental or Behavioral Dysfunction | 40 | 6 | 0,01 | 1 | 1 | 0 | 2018 | 2018 |
| SERPINA3 | 12 | Mental De | C0011570 | disease | Behavior and Behavior Mechanisms | Mental or Behavioral Dysfunction | 1478 | 271 | 0,01 | 1 | 1 | 0 | 2017 | 2017 |
| SERPINA3 | 12 | Depressive | C0086132 | phenotype | Behavior and Behavior Mechanisms | Sign or Symptom | 421 | 120 | 0,01 | 1 | 1 | 0 | 2018 | 2018 |
| SERPINA3 | 12 | Abnormal I | C0233514 | phenotype | Behavior and Behavior Mechanisms | Mental or Behavioral Dysfunction | 910 | 121 | 0,01 | 1 | 1 | 0 | 2018 | 2018 |
| SERPINA3 | 12 | Depressed | C0344315 | phenotype | Behavior and Behavior Mechanisms | Mental or Behavioral Dysfunction | 1461 | 269 | 0,01 | 1 | 1 | 0 | 2017 | 2017 |
| SERPINA3 | 12 | Depressior | C0221074 | disease | Female Urogenital Diseases and Pregnancy Complications; Me | Mental or Behavioral Dysfunction | 54 | 6 | 0,01 | 1 | 1 | 0 | 2018 | 2018 |
| SERPINA3 | 12 | Psychotic I | C0033975 | group | Mental Disorders | Mental or Behavioral Dysfunction | 560 | 179 | 0,33 | 0,667 | 3 | 0 | 2002 | 2018 |
| SERPINA3 | 12 | Nonorgani | C0349204 | disease | Mental Disorders | Mental or Behavioral Dysfunction | 376 | 98 | 0,33 | 0,667 | 3 | 0 | 2002 | 2018 |
| SERPINA3 | 12 | Schizophre | C0036341 | disease | Mental Disorders | Mental or Behavioral Dysfunction | 2872 | 2897 | 0,05 | 1 | 5 | 0 | 1999 | 2018 |
| SERPINA3 | 12 | Mental det | C0234985 | phenotype | Mental Disorders | Mental or Behavioral Dysfunction | 508 | 121 | 0,05 | 1 | 5 | 0 | 1996 | 2008 |
| SERPINA3 | 12 | Impaired c | C0338656 | disease | Mental Disorders | Mental or Behavioral Dysfunction | 1630 | 348 | 0,05 | 1 | 5 | 0 | 1996 | 2008 |
| SERPINA3 | 12 | Mild cognit | C1270972 | disease | Mental Disorders | Mental or Behavioral Dysfunction | 430 | 96 | 0,03 | 1 | 3 | 0 | 2009 | 2019 |
| SERPINA3 | 12 | Mental dis | C0004936 | group | Mental Disorders | Mental or Behavioral Dysfunction | 789 | 149 | 0,02 | 1 | 2 | 0 | 2018 | 2018 |
| SERPINA3 | 12 | Depressive | C0011581 | disease | Mental Disorders | Mental or Behavioral Dysfunction | 1719 | 297 | 0,02 | 0,5 | 2 | 0 | 1999 | 2017 |
| SERPINA3 | 12 | Eating Diso | C0013473 | group | Mental Disorders | Mental or Behavioral Dysfunction | 133 | 42 | 0,01 | 1 | 1 | 0 | 2017 | 2017 |
| ND5 | 4540 | Anxiety | C0003467 | disease | Behavior and Behavior Mechanisms | Mental or Behavioral Dysfunction | 1048 | 287 | 0,1 | | 0 | 0 | | |
| ND5 | 4540 | Autistic Dis | C0004352 | disease | Mental Disorders | Mental or Behavioral Dysfunction | 1112 | 395 | 0,1 | | 0 | 0 | | |
| ND5 | 4540 | Depressive | C0011581 | disease | Mental Disorders | Mental or Behavioral Dysfunction | 1719 | 297 | 0,1 | | 0 | 0 | | |
| ND5 | 4540 | Psychotic e | C0338614 | disease | Mental Disorders | Mental or Behavioral Dysfunction | 31 | 6 | 0,1 | | 0 | 0 | | |
| ND5 | 4540 | Impaired c | C0338656 | disease | Mental Disorders | Mental or Behavioral Dysfunction | 1630 | 348 | 0,1 | | 0 | 0 | | |
| ND5 | 4540 | Attention c | C1263846 | disease | Mental Disorders | Mental or Behavioral Dysfunction | 842 | 420 | 0,1 | | 0 | 0 | | |
| ND5 | 4540 | Developme | C1836830 | disease | Mental Disorders | Disease or Syndrome | 333 | 80 | 0,1 | | 0 | 0 | | |
| ND5 | 4540 | Psychotic I | C0033975 | group | Mental Disorders | Mental or Behavioral Dysfunction | 560 | 179 | 0,01 | 1 | 1 | 0 | 2017 | 2017 |
| ND5 | 4540 | Schizophre | C0036341 | disease | Mental Disorders | Mental or Behavioral Dysfunction | 2872 | 2897 | 0,01 | 1 | 1 | 0 | 2008 | 2008 |
| ND5 | 4540 | Post-Traun | C0038436 | disease | Mental Disorders | Mental or Behavioral Dysfunction | 418 | 117 | 0,01 | 1 | 1 | 0 | 2015 | 2015 |
| ND5 | 4540 | Nonorgani | C0349204 | disease | Mental Disorders | Mental or Behavioral Dysfunction | 376 | 98 | 0,01 | 1 | 1 | 0 | 2017 | 2017 |
| P2RX2 | 22953 | Mental De | C0011570 | disease | Behavior and Behavior Mechanisms | Mental or Behavioral Dysfunction | 1478 | 271 | 0,01 | 1 | 1 | 2 | 2013 | 2013 |
| P2RX2 | 22953 | Depressive | C0086132 | phenotype | Behavior and Behavior Mechanisms | Sign or Symptom | 421 | 120 | 0,01 | 1 | 1 | 0 | 2019 | 2019 |
| P2RX2 | 22953 | Depressed | C0344315 | phenotype | Behavior and Behavior Mechanisms | Mental or Behavioral Dysfunction | 1461 | 269 | 0,01 | 1 | 1 | 2 | 2013 | 2013 |
| P2RX2 | 22953 | Bipolar Dis | C0005586 | disease | Mental Disorders | Mental or Behavioral Dysfunction | 1183 | 839 | 0,02 | 1 | 2 | 2 | 2008 | 2009 |

| | | | | | | | | | | | | | | |
|---|---|---|---|---|---|---|---|---|---|---|---|---|---|---|
| P2RX2 | 22953 | Borderline | C0006012 | disease | Mental Disorders | Mental or Behavioral Dysfunction | 221 | 82 | 0,02 | 1 | 2 | 0 | 2009 | 2013 |
| P2RX2 | 22953 | Mood Diso | C0525045 | group | Mental Disorders | Mental or Behavioral Dysfunction | 580 | 308 | 0,02 | 1 | 2 | 0 | 2009 | 2018 |
| P2RX2 | 22953 | Major Dep | C1269683 | disease | Mental Disorders | Mental or Behavioral Dysfunction | 1236 | 1451 | 0,02 | 1 | 2 | 0 | 2009 | 2013 |
| P2RX2 | 22953 | Depressive | C0011581 | disease | Mental Disorders | Mental or Behavioral Dysfunction | 1719 | 297 | 0,01 | 1 | 1 | 2 | 2013 | 2013 |
| P2RX2 | 22953 | Schizophre | C0036341 | disease | Mental Disorders | Mental or Behavioral Dysfunction | 2872 | 2897 | 0,01 | 1 | 1 | 0 | 2008 | 2008 |
| P2RX2 | 22953 | Impaired c | C0338656 | disease | Mental Disorders | Mental or Behavioral Dysfunction | 1630 | 348 | 0,01 | 1 | 1 | 0 | 2017 | 2017 |
| ATP4A | 495 | Mental De | C0011570 | disease | Behavior and Behavior Mechanisms | Mental or Behavioral Dysfunction | 1478 | 271 | 0,02 | 1 | 2 | 0 | 2018 | 2019 |
| ATP4A | 495 | Depressed | C0344315 | phenotype | Behavior and Behavior Mechanisms | Mental or Behavioral Dysfunction | 1461 | 269 | 0,02 | 1 | 2 | 0 | 2018 | 2019 |
| ATP4A | 495 | Depressive | C0011581 | disease | Mental Disorders | Mental or Behavioral Dysfunction | 1719 | 297 | 0,02 | 1 | 2 | 0 | 2018 | 2019 |
| PROC | 5624 | Depressive | C0086132 | phenotype | Behavior and Behavior Mechanisms | Sign or Symptom | 421 | 120 | 0,01 | 1 | 1 | 0 | 2018 | 2018 |
| PROC | 5624 | Attention c | C1263846 | disease | Mental Disorders | Mental or Behavioral Dysfunction | 842 | 420 | 0,01 | 1 | 1 | 0 | 2020 | 2020 |
| CARTPT | 9607 | Mental De | C0011570 | disease | Behavior and Behavior Mechanisms | Mental or Behavioral Dysfunction | 1478 | 271 | 0,34 | 1 | 5 | 0 | 2006 | 2018 |
| CARTPT | 9607 | Anxiety | C0003467 | disease | Behavior and Behavior Mechanisms | Mental or Behavioral Dysfunction | 1048 | 287 | 0,04 | 0,75 | 4 | 0 | 2006 | 2013 |
| CARTPT | 9607 | Depressed | C0344315 | phenotype | Behavior and Behavior Mechanisms | Mental or Behavioral Dysfunction | 1461 | 269 | 0,04 | 1 | 4 | 0 | 2006 | 2018 |
| CARTPT | 9607 | Addictive E | C0085281 | phenotype | Behavior and Behavior Mechanisms | Mental or Behavioral Dysfunction | 332 | 56 | 0,03 | 1 | 3 | 0 | 2007 | 2018 |
| CARTPT | 9607 | Depressive | C0086132 | phenotype | Behavior and Behavior Mechanisms | Sign or Symptom | 421 | 120 | 0,01 | 1 | 1 | 0 | 2018 | 2018 |
| CARTPT | 9607 | Anxiety Dis | C0003469 | group | Mental Disorders | Mental or Behavioral Dysfunction | 840 | 163 | 0,34 | 0,8 | 5 | 0 | 2003 | 2013 |
| CARTPT | 9607 | Depressive | C0011581 | disease | Mental Disorders | Mental or Behavioral Dysfunction | 1719 | 297 | 0,34 | 1 | 5 | 0 | 2006 | 2018 |
| CARTPT | 9607 | Schizophre | C0036341 | disease | Mental Disorders | Mental or Behavioral Dysfunction | 2872 | 2897 | 0,33 | 0,667 | 3 | 0 | 2004 | 2018 |
| CARTPT | 9607 | Bipolar Dis | C0005586 | disease | Mental Disorders | Mental or Behavioral Dysfunction | 1183 | 839 | 0,31 | 0 | 1 | 0 | 2004 | 2004 |
| CARTPT | 9607 | Psychotic I | C0033975 | group | Mental Disorders | Mental or Behavioral Dysfunction | 560 | 179 | 0,31 | 0 | 1 | 0 | 2006 | 2006 |
| CARTPT | 9607 | Nonorgani | C0349204 | disease | Mental Disorders | Mental or Behavioral Dysfunction | 376 | 98 | 0,31 | 0 | 1 | 0 | 2006 | 2006 |
| CARTPT | 9607 | Anxiety Sta | C0376280 | disease | Mental Disorders | Mental or Behavioral Dysfunction | 44 | 0 | 0,3 | 1 | 1 | 0 | 2003 | 2003 |
| CARTPT | 9607 | Anxiety ne | C1279420 | disease | Mental Disorders | Mental or Behavioral Dysfunction | 75 | 5 | 0,3 | 1 | 1 | 0 | 2003 | 2003 |
| CARTPT | 9607 | Mental dis | C0004936 | group | Mental Disorders | Mental or Behavioral Dysfunction | 789 | 149 | 0,01 | 1 | 1 | 0 | 2012 | 2012 |
| CARTPT | 9607 | Eating Diso | C0013473 | group | Mental Disorders | Mental or Behavioral Dysfunction | 133 | 42 | 0,01 | 1 | 1 | 0 | 2007 | 2007 |
| CARTPT | 9607 | Attention I | C0041671 | disease | Mental Disorders | Mental or Behavioral Dysfunction | 123 | 7 | 0,01 | 1 | 1 | 0 | 2008 | 2008 |
| CARTPT | 9607 | Unipolar D | C0041696 | disease | Mental Disorders | Mental or Behavioral Dysfunction | 641 | 225 | 0,01 | 1 | 1 | 0 | 2018 | 2018 |
| CARTPT | 9607 | Attention-I | C0339002 | disease | Mental Disorders | Mental or Behavioral Dysfunction | 12 | 1 | 0,01 | 1 | 1 | 0 | 2008 | 2008 |
| CARTPT | 9607 | Attention c | C1263846 | disease | Mental Disorders | Mental or Behavioral Dysfunction | 842 | 420 | 0,01 | 1 | 1 | 0 | 2008 | 2008 |
| CARTPT | 9607 | Major Dep | C1269683 | disease | Mental Disorders | Mental or Behavioral Dysfunction | 1236 | 1451 | 0,01 | 1 | 1 | 0 | 2018 | 2018 |
