## Supplementary tables for "Transcriptome profiling of the dorsomedial prefrontal cortex in suicide victims": S8_Co-expression_networks_with_glay_clusters.pdf

Supplementary file 8: Co-expression network data of downregulated genes; the top 10 most ranked hub genes are highlighted.

| name | betweenness | degree | log2FoldChange | Node_Status | padj | Rank_stat | shared name |
|---|---|---|---|---|---|---|---|
| SORBS3 | 7004,400116 | 116 | -1,503254358 | Clustered | 5,77E-04 | 707 | SORBS3 |
| RHOC | 6409,495473 | 105 | -1,925958166 | Clustered | 6,56E-05 | 704,25 | RHOC |
| S100A16 | 4397,76169 | 114 | -1,645242746 | Clustered | 6,65E-04 | 700 | S100A16 |
| SZRD1 | 9088,385865 | 88 | -1,500872432 | Clustered | 4,05E-04 | 699,75 | SZRD1 |
| TRIP6 | 4704,286451 | 101 | -1,601503127 | Clustered | 5,77E-04 | 699 | TRIP6 |
| AHNAK | 5131,964826 | 85 | -2,312466754 | Clustered | 1,41E-05 | 694,25 | AHNAK |
| GRIN2C | 3580,24906 | 108 | -2,166181557 | Clustered | 2,20E-04 | 692,5 | GRIN2C |
| CHDH | 3625,789357 | 102 | -1,851193891 | Clustered | 9,39E-04 | 691,5 | CHDH |
| MAPKAPK2 | 4250,193284 | 89 | -1,610321727 | Clustered | 4,05E-04 | 691,5 | MAPKAPK2 |
| FTL | 7408,598658 | 75 | -2,146090078 | Clustered | 5,22E-05 | 691 | FTL |
| IRAG1 | 3455,790802 | 105 | -2,866111414 | Clustered | 8,96E-06 | 690,25 | IRAG1 |
| MMP28 | 3752,262903 | 96 | -1,671462463 | Clustered | 0,001923 | 690,25 | MMP28 |
| UBTD1 | 4152,925979 | 82 | -1,064582316 | Clustered | 0,002002 | 687,75 | UBTD1 |
| FGFRL1 | 3364,019788 | 97 | -2,382155201 | Clustered | 8,14E-05 | 687 | FGFRL1 |
| SLC44A2 | 3539,385145 | 94 | -1,047087943 | Clustered | 0,004957 | 686,75 | SLC44A2 |
| MCL1 | 6048,816892 | 71 | -1,436052385 | Clustered | 1,91E-04 | 686,5 | MCL1 |
| TCN2 | 8435,680133 | 66 | -2,092055168 | Clustered | 7,15E-05 | 685,5 | TCN2 |
| DDR2 | 4038,610066 | 80 | -1,611967921 | Clustered | 6,61E-04 | 684,75 | DDR2 |
| ATOH8 | 4595,459846 | 73 | -2,575134567 | Clustered | 9,71E-05 | 683,75 | ATOH8 |
| PSKH1 | 4865,755918 | 69 | -1,306231373 | Clustered | 6,34E-04 | 683 | PSKH1 |
| STAT3 | 3341,241971 | 88 | -1,295022634 | Clustered | 0,002404 | 681,75 | STAT3 |
| METRN | 5123,502418 | 63 | -2,076596027 | Clustered | 5,95E-04 | 678,5 | METRN |
| WASF2 | 15081,4143 | 56 | -1,844923416 | Clustered | 3,84E-04 | 677 | WASF2 |
| TTC38 | 2886,700001 | 90 | -1,671128466 | Clustered | 0,001762 | 676,5 | TTC38 |
| HLA-E | 6527,740127 | 58 | -1,679088265 | Clustered | 3,65E-04 | 676,25 | HLA-E |
| CDC42EP4 | 3091,497849 | 78 | -1,680137287 | Clustered | 5,90E-04 | 674 | CDC42EP4 |
| ARHGEF26 | 2751,167453 | 96 | -2,29578728 | Clustered | 2,39E-04 | 673,25 | ARHGEF26 |
| VAMP5 | 2918,870485 | 81 | -2,18349657 | Clustered | 1,98E-04 | 672,75 | VAMP5 |
| GNG5 | 3741,089492 | 66 | -1,309266502 | Clustered | 4,02E-04 | 672 | GNG5 |
| SNTA1 | 2553,317347 | 98 | -1,797359123 | Clustered | 9,29E-04 | 671 | SNTA1 |
| GNA12 | 3924,206615 | 62 | -2,082375488 | Clustered | 1,41E-05 | 669,75 | GNA12 |
| MSI2 | 2460,049574 | 99 | -1,54663506 | Clustered | 5,73E-04 | 668,5 | MSI2 |
| RELL1 | 11225,42679 | 50 | -2,026602544 | Clustered | 3,30E-05 | 668,5 | RELL1 |
| SNX33 | 2496,540401 | 94 | -1,885949238 | Clustered | 4,94E-04 | 667,25 | SNX33 |
| YAP1 | 2777,487937 | 78 | -2,100451257 | Clustered | 1,80E-04 | 665,5 | YAP1 |
| WBP1 | 2969,991053 | 68 | -1,078467595 | Clustered | 0,002764 | 665 | WBP1 |
| ADCYAP1R1 | 2704,108192 | 77 | -1,394679075 | Clustered | 6,24E-04 | 662,5 | ADCYAP1R1 |
| MLC1 | 2706,596537 | 73 | -1,972313085 | Clustered | 4,82E-04 | 660,25 | MLC1 |
| GPR4 | 5896,510292 | 48 | -3,413584789 | Clustered | 3,85E-06 | 659,75 | GPR4 |
| AMOT | 3749,500487 | 52 | -1,62568909 | Clustered | 3,36E-04 | 658 | AMOT |
| STON2 | 2124,365061 | 85 | -2,197802589 | Clustered | 1,59E-04 | 654,25 | STON2 |
| MAVS | 2782,029299 | 62 | -1,321887113 | Clustered | 0,00508 | 653,25 | MAVS |
| BCL2 | 1908,340841 | 91 | -1,262924751 | Clustered | 0,00222 | 653 | BCL2 |
| LRP10 | 3268,848006 | 52 | -2,055562099 | Clustered | 1,91E-04 | 653 | LRP10 |
| TNFRSF1A | 3524,886085 | 50 | -1,441670926 | Clustered | 0,00196 | 651,5 | TNFRSF1A |
| CABLES1 | 1996,899686 | 80 | -1,80114928 | Clustered | 7,41E-04 | 649,25 | CABLES1 |
| RRAS | 2601,869347 | 61 | -1,490579515 | Clustered | 0,001049 | 648,5 | RRAS |
| ANTXR1 | 3108,666443 | 50 | -1,3571581 | Clustered | 1,13E-04 | 647,5 | ANTXR1 |
| BCL6 | 2053,339223 | 76 | -1,51142377 | Clustered | 6,96E-04 | 647,25 | BCL6 |
| ECE1 | 3354,210593 | 47 | -1,278259864 | Clustered | 0,0016 | 644,75 | ECE1 |
| PALLD | 3177,386214 | 47 | -2,175044901 | Clustered | 3,11E-05 | 642,75 | PALLD |
| NECAP2 | 2281,426224 | 59 | -1,099164078 | Clustered | 0,002541 | 639 | NECAP2 |
| RAB13 | 2198,646058 | 60 | -2,234050673 | Clustered | 1,97E-04 | 638,75 | RAB13 |
| FAM107A | 2812,472368 | 48 | -2,798652524 | Clustered | 2,33E-06 | 637,75 | FAM107A |
| GDF1 | 1682,472027 | 76 | -1,517494328 | Clustered | 0,001327 | 637,75 | GDF1 |
| ACVRL1 | 6244,066379 | 34 | -1,805122896 | Clustered | 8,03E-05 | 636 | ACVRL1 |
| CTDSP1 | 2308,246671 | 55 | -1,76675975 | Clustered | 3,21E-04 | 635,25 | CTDSP1 |
| TBL1X | 1535,199196 | 78 | -1,65797519 | Clustered | 4,71E-04 | 635 | TBL1X |

| | | | | | | | |
|---|---|---|---|---|---|---|---|
| TMEM219 | 2479,669102 | 51 | -1,169229322 | Clustered | 0,007421 | 633,75 | TMEM219 |
| VEGFB | 1457,007308 | 82 | -1,714936371 | Seed | 5,48E-04 | 633,75 | VEGFB |
| NBPF14 | 1721,621156 | 66 | -2,172294861 | Clustered | 8,95E-04 | 632,5 | NBPF14 |
| NHSL1 | 2194,728441 | 52 | -1,286487277 | Clustered | 0,0035 | 630 | NHSL1 |
| SMO | 1534,173721 | 70 | -2,396446953 | Clustered | 1,91E-04 | 629,5 | SMO |
| CASKIN2 | 2860,827472 | 42 | -1,010456435 | Clustered | 0,004655 | 629 | CASKIN2 |
| EFEMP2 | 1674,06714 | 64 | -1,57352654 | Clustered | 0,001695 | 629 | EFEMP2 |
| ITPKB | 1659,997927 | 62 | -2,40306438 | Clustered | 1,12E-05 | 625,75 | ITPKB |
| PLEKHO2 | 1482,386446 | 68 | -2,124769449 | Clustered | 2,20E-04 | 625,5 | PLEKHO2 |
| PBXIP1 | 1620,291774 | 62 | -1,808086314 | Clustered | 9,27E-04 | 624,75 | PBXIP1 |
| VSIR | 1566,703965 | 63 | -2,305860353 | Clustered | 3,95E-04 | 624,5 | VSIR |
| SMOX | 1264,829668 | 81 | -1,21401308 | Seed | 0,004191 | 619,75 | SMOX |
| MSN | 2416,567901 | 43 | -1,43360871 | Clustered | 1,41E-05 | 618,75 | MSN |
| COPZ2 | 2176,72353 | 45 | -1,610047521 | Clustered | 0,0024 | 618 | COPZ2 |
| KANK2 | 1806,336045 | 50 | -1,250500978 | Clustered | 0,004019 | 618 | KANK2 |
| CERS1 | 1304,050356 | 74 | -1,330744813 | Clustered | 0,003762 | 617 | CERS1 |
| MAPKAPK3 | 1369,216886 | 68 | -1,703526834 | Clustered | 0,001481 | 616,5 | MAPKAPK3 |
| SIPA1 | 2306,91141 | 42 | -1,227637175 | Clustered | 0,003956 | 614,5 | SIPA1 |
| LRTOMT | 1790,184916 | 48 | -1,546764888 | Clustered | 4,05E-04 | 613,75 | LRTOMT |
| MOB3C | 3188,550223 | 31 | -1,663191248 | Clustered | 5,15E-04 | 613,75 | MOB3C |
| GPR137B | 2812,785003 | 34 | -1,379448133 | Unclustered | 8,63E-04 | 613,5 | GPR137B |
| SYDE1 | 2438,744218 | 40 | -1,633722456 | Unclustered | 0,002089 | 612,5 | SYDE1 |
| EZR | 1403,67433 | 57 | -1,612745371 | Clustered | 0,002692 | 610,5 | EZR |
| RAB31 | 1284,119015 | 65 | -1,791207411 | Clustered | 7,43E-04 | 609,5 | RAB31 |
| CFLAR | 2809,25821 | 32 | -1,355274887 | Unclustered | 0,003123 | 609,25 | CFLAR |
| SLC7A11 | 1307,177379 | 60 | -1,78412284 | Clustered | 0,001238 | 606,25 | SLC7A11 |
| TNS3 | 2488,932513 | 35 | -2,428588605 | Clustered | 1,41E-05 | 605,25 | TNS3 |
| TCF7L2 | 1165,183732 | 67 | -1,309761253 | Clustered | 0,002089 | 605 | TCF7L2 |
| HELZ2 | 2552,395285 | 33 | -2,466768427 | Clustered | 2,33E-05 | 603,5 | HELZ2 |
| NES | 2039,059428 | 40 | -1,384624347 | Clustered | 4,71E-04 | 603,5 | NES |
| FYCO1 | 4054,30061 | 24 | -1,38742622 | Unclustered | 4,42E-04 | 603,25 | FYCO1 |
| PLD2 | 1123,104275 | 63 | -1,364837333 | Clustered | 0,003462 | 599,5 | PLD2 |
| LRRC8A | 1729,357084 | 41 | -1,409545279 | Clustered | 0,0019 | 598,75 | LRRC8A |
| GGT5 | 1940,308756 | 38 | -2,34962623 | Clustered | 1,08E-04 | 598,5 | GGT5 |
| SCARA3 | 1258,43251 | 56 | -2,087154025 | Clustered | 1,79E-04 | 597,5 | SCARA3 |
| SERPINH1 | 1696,994162 | 41 | -2,622747343 | Clustered | 9,71E-05 | 596,75 | SERPINH1 |
| ARHGAP19 | 2215,155099 | 32 | -1,857186508 | Clustered | 1,65E-05 | 595,25 | ARHGAP19 |
| PTTG1IP | 2502,26623 | 30 | -1,25062545 | Unclustered | 7,28E-04 | 595,25 | PTTG1IP |
| RGS3 | 2120,077572 | 33 | -1,112065077 | Clustered | 0,001461 | 592,5 | RGS3 |
| RIN2 | 1135,678518 | 52 | -1,422849844 | Clustered | 9,43E-04 | 589 | RIN2 |
| CGNL1 | 1061,968462 | 56 | -2,274213883 | Clustered | 1,59E-05 | 588,5 | CGNL1 |
| PXN | 2587,174122 | 26 | -1,649859088 | Clustered | 6,66E-04 | 587,75 | PXN |
| ZCCHC24 | 9660,236534 | 19 | -1,959866173 | Unclustered | 8,14E-05 | 587,25 | ZCCHC24 |
| DHRS3 | 1569,57457 | 37 | -1,437737877 | Clustered | 0,003159 | 586,25 | DHRS3 |
| PAQR6 | 2944,289136 | 23 | -1,740577575 | Clustered | 3,01E-04 | 585,25 | PAQR6 |
| BCAS1 | 4572,80122 | 20 | -1,877126198 | Clustered | 3,74E-04 | 585 | BCAS1 |
| TMEM164 | 932,9568857 | 58 | -1,152614816 | Clustered | 0,001953 | 584,25 | TMEM164 |
| ANKRD40 | 4164,002231 | 20 | -1,569516982 | Clustered | 2,69E-04 | 583 | ANKRD40 |
| MID1 | 1007,506089 | 53 | -1,172266817 | Clustered | 0,004774 | 582,75 | MID1 |
| DOCK6 | 1762,500526 | 31 | -1,755078447 | Unclustered | 0,001454 | 581,25 | DOCK6 |
| VASP | 2319,844056 | 24 | -1,157985771 | Clustered | 0,007421 | 575,25 | VASP |
| CDKN1A | 2799,801586 | 22 | -3,149516104 | Unclustered | 1,41E-05 | 574,25 | CDKN1A |
| TGFB2 | 2796,69599 | 22 | -1,465901924 | Unclustered | 0,001441 | 573,75 | TGFB2 |
| IFI35 | 1209,187427 | 42 | -1,744744086 | Clustered | 0,001327 | 573,5 | IFI35 |
| MAP4 | 3043,316307 | 20 | -1,614046646 | Clustered | 1,41E-05 | 571 | MAP4 |
| PARD3 | 894,191376 | 50 | -2,300860234 | Clustered | 9,23E-04 | 570,5 | PARD3 |
| MT-CO2 | 1958,96782 | 25 | -3,491557623 | Clustered | 1,41E-05 | 570,25 | MT-CO2 |
| PARD3B | 1603,553526 | 28 | -2,365493936 | Unclustered | 0,002093 | 568,25 | PARD3B |
| MT-ND3 | 2133,000468 | 23 | -3,829002115 | Clustered | 3,85E-06 | 565,75 | MT-ND3 |
| CLIC1 | 3763,182409 | 17 | -1,745516837 | Unclustered | 1,18E-04 | 562,5 | CLIC1 |
| NWD1 | 774,5342857 | 52 | -2,196298753 | Clustered | 3,01E-04 | 562,5 | NWD1 |

| | | | | | | | |
|---|---|---|---|---|---|---|---|
| KANK3 | 1458,223559 | 28 | -1,425725286 | Clustered | 7,96E-04 | 562,25 | KANK3 |
| ACSS1 | 1062,291536 | 39 | -1,368687752 | Clustered | 0,007018 | 561,5 | ACSS1 |
| RHPN2 | 1955,584619 | 23 | -1,689159433 | Unclustered | 0,001571 | 561,25 | RHPN2 |
| ZC3HAV1 | 719,0427381 | 57 | -1,518203129 | Clustered | 0,001768 | 560,5 | ZC3HAV1 |
| GPAM | 809,5374221 | 48 | -1,813644981 | Clustered | 0,001159 | 559,25 | GPAM |
| APPL2 | 1049,182747 | 38 | -1,113932803 | Clustered | 0,002491 | 558,5 | APPL2 |
| ENHO | 706,7734746 | 57 | -2,205329168 | Clustered | 3,31E-04 | 557,5 | ENHO |
| CACFD1 | 877,8204049 | 42 | -1,089772553 | Clustered | 0,005261 | 556,5 | CACFD1 |
| HSPB1 | 834,0010747 | 43 | -1,664950693 | Clustered | 0,002858 | 555,25 | HSPB1 |
| ADAMTS9 | 1374,872717 | 27 | -2,790033365 | Clustered | 2,91E-05 | 554 | ADAMTS9 |
| OAF | 862,9681095 | 42 | -1,641995334 | Clustered | 0,002053 | 554 | OAF |
| FCGRT | 720,4410429 | 51 | -1,586972702 | Clustered | 0,002581 | 553,75 | FCGRT |
| OSMR | 3002,996991 | 17 | -2,405279518 | Unclustered | 1,41E-05 | 553 | OSMR |
| SIX5 | 1263,867674 | 29 | -1,711811794 | Unclustered | 0,004104 | 550,5 | SIX5 |
| ADGRG1 | 931,6740819 | 37 | -1,722325628 | Clustered | 9,99E-04 | 550,25 | ADGRG1 |
| LTBR | 1365,253559 | 26 | -1,457984618 | Clustered | 0,001473 | 548,25 | LTBR |
| GABRG1 | 1414,965368 | 24 | -1,394263072 | Unclustered | 0,003538 | 547,75 | GABRG1 |
| TNFSF13 | 787,8451555 | 42 | -2,003065952 | Clustered | 3,89E-04 | 547,5 | TNFSF13 |
| NPAS3 | 810,4879609 | 41 | -1,39385665 | Clustered | 0,004018 | 547,25 | NPAS3 |
| IL4R | 3087,737408 | 16 | -2,79415517 | Clustered | 1,28E-04 | 547 | IL4R |
| GALNT10 | 2982,559346 | 16 | -1,070825363 | Unclustered | 7,19E-04 | 545,5 | GALNT10 |
| DDAH2 | 1171,403916 | 28 | -1,054159272 | Clustered | 0,006085 | 544,25 | DDAH2 |
| RFX4 | 715,075795 | 44 | -1,868238083 | Clustered | 5,15E-04 | 540,5 | RFX4 |
| KANK1 | 839,3186422 | 35 | -1,203525245 | Clustered | 0,001507 | 539,75 | KANK1 |
| NACC2 | 1082,065943 | 27 | -1,509685056 | Unclustered | 1,73E-04 | 538 | NACC2 |
| EYA2 | 821,0650549 | 35 | -1,443586577 | Clustered | 0,003972 | 537,75 | EYA2 |
| CHST14 | 1233,731404 | 24 | -1,202806137 | Unclustered | 0,007122 | 535,25 | CHST14 |
| EIF4EBP2 | 697,0623208 | 43 | -1,232579129 | Clustered | 0,001603 | 535,25 | EIF4EBP2 |
| IFITM3 | 4026,059613 | 13 | -2,199506981 | Unclustered | 1,57E-04 | 533,75 | IFITM3 |
| SLC5A10 | 1701,171223 | 19 | -2,225646859 | Clustered | 1,54E-04 | 532,75 | SLC5A10 |
| CCDC69 | 590,2654949 | 53 | -1,397079522 | Clustered | 0,001416 | 530,75 | CCDC69 |
| PPP1R14B | 748,7689882 | 36 | -1,386864375 | Clustered | 0,006776 | 530,75 | PPP1R14B |
| AGTRAP | 742,1790291 | 36 | -1,76261202 | Clustered | 0,002862 | 529,75 | AGTRAP |
| RAB34 | 900,4710047 | 27 | -1,415624042 | Unclustered | 0,00821 | 526,5 | RAB34 |
| TEAD1 | 665,4730622 | 45 | -1,259174425 | Clustered | 0,002634 | 526 | TEAD1 |
| CDKN2C | 1582,829269 | 18 | -1,231610246 | Unclustered | 0,001172 | 523 | CDKN2C |
| DYNC1LI2 | 1304,57642 | 21 | -1,16276589 | Clustered | 2,20E-04 | 522,5 | DYNC1LI2 |
| NOTCH2NLC | 1431,906782 | 19 | -1,559898638 | Clustered | 0,002472 | 522,25 | NOTCH2NLC |
| BST2 | 1056,174209 | 23 | -2,334438589 | Unclustered | 4,01E-04 | 521,25 | BST2 |
| LRRC32 | 1416,924079 | 19 | -2,820078033 | Clustered | 1,41E-05 | 521,25 | LRRC32 |
| MAOA | 616,2547254 | 44 | -1,07287968 | Clustered | 0,004567 | 519 | MAOA |
| METRNL | 479,8661515 | 55 | -1,400729077 | Clustered | 0,003454 | 518,25 | METRNL |
| TNFRSF1B | 743,6087375 | 30 | -2,428576337 | Clustered | 4,58E-05 | 517,25 | TNFRSF1B |
| ITGA5 | 1777,730217 | 16 | -2,379084248 | Clustered | 6,68E-04 | 517 | ITGA5 |
| ABHD4 | 1226,480384 | 21 | -1,5180432 | Unclustered | 7,88E-04 | 516,5 | ABHD4 |
| SOD3 | 753,7859249 | 28 | -2,106208327 | Clustered | 0,001417 | 514,75 | SOD3 |
| TMBIM1 | 1027,489106 | 22 | -2,136064546 | Unclustered | 2,91E-05 | 512,75 | TMBIM1 |
| ACSL5 | 2569,281975 | 13 | -2,335391799 | Unclustered | 3,22E-04 | 512,25 | ACSL5 |
| EMILIN1 | 511,8876004 | 49 | -1,311488092 | Seed | 0,008341 | 512,25 | EMILIN1 |
| PXDC1 | 715,7359558 | 29 | -1,433432696 | Clustered | 0,001138 | 511 | PXDC1 |
| CENPB | 2189,433904 | 14 | -1,286100937 | Clustered | 0,002134 | 508,75 | CENPB |
| TPD52L1 | 512,0280685 | 45 | -1,605587855 | Clustered | 0,001181 | 508 | TPD52L1 |
| EMID1 | 616,4782757 | 38 | -1,59750008 | Clustered | 0,003155 | 507,5 | EMID1 |
| MYORG | 1226,596049 | 19 | -1,788528186 | Unclustered | 6,65E-04 | 507,25 | MYORG |
| APCDD1 | 1394,584818 | 17 | -1,173497526 | Unclustered | 0,007124 | 507 | APCDD1 |
| GRAMD2B | 450,8148102 | 49 | -1,202666264 | Clustered | 0,003645 | 506,75 | GRAMD2B |
| PPARA | 574,7743103 | 41 | -1,541728611 | Clustered | 0,004019 | 506,75 | PPARA |
| ENSG0000028! | 1725,822835 | 15 | -2,231869708 | Unclustered | 6,96E-04 | 506 | ENSG00000285526 |
| RARRES2 | 569,0701999 | 40 | -1,521503864 | Clustered | 0,005609 | 504 | RARRES2 |
| SLC9A3R1 | 476,8788777 | 45 | -1,677545586 | Clustered | 0,001324 | 504 | SLC9A3R1 |
| DIPK2B | 2804,255599 | 11 | -1,235404623 | Unclustered | 0,001168 | 503,5 | DIPK2B |

| | | | | | | | |
|---|---|---|---|---|---|---|---|
| CALD1 | 786,4550079 | 23 | -2,076030462 | Unclustered | 0,002735 | 501,25 | CALD1 |
| ENG | 2774,869619 | 11 | -1,485319208 | Seed | 7,79E-04 | 501 | ENG |
| PHF21B | 774,027326 | 23 | -1,734551282 | Unclustered | 0,003365 | 499,25 | PHF21B |
| SLC39A1 | 1745,343355 | 14 | -1,297596721 | Unclustered | 9,33E-04 | 498,75 | SLC39A1 |
| ZBTB47 | 4384,972413 | 9 | -1,123941295 | Clustered | 0,001171 | 498,75 | ZBTB47 |
| MRO | 665,1013599 | 31 | -2,025947943 | Clustered | 3,85E-06 | 497,75 | MRO |
| ITGB5 | 1134,955584 | 18 | -1,43075672 | Unclustered | 9,02E-04 | 497,5 | ITGB5 |
| ELP4 | 394,9495286 | 45 | -1,372014632 | Clustered | 0,004292 | 497 | ELP4 |
| ALDH4A1 | 461,6386368 | 42 | -1,898761889 | Clustered | 9,40E-04 | 496,5 | ALDH4A1 |
| MBP | 1345,106904 | 16 | -1,84035691 | Clustered | 0,001138 | 495,5 | MBP |
| TRIOBP | 584,3584399 | 34 | -1,416078227 | Clustered | 0,004572 | 495,5 | TRIOBP |
| IRF1 | 1340,054212 | 16 | -1,57096747 | Unclustered | 0,004468 | 495 | IRF1 |
| GIMAP5 | 736,1928515 | 23 | -2,133768299 | Clustered | 3,95E-04 | 494,75 | GIMAP5 |
| FGF11 | 326,1298486 | 50 | -2,104078336 | Clustered | 0,001138 | 494,5 | FGF11 |
| PAX6 | 391,7530512 | 44 | -1,55776464 | Clustered | 0,003096 | 493,5 | PAX6 |
| PRKX | 1273,993894 | 16 | -1,721502578 | Unclustered | 0,005089 | 491,5 | PRKX |
| SHE | 756,947506 | 22 | -1,848505529 | Clustered | 6,45E-04 | 491,25 | SHE |
| ZFP36L1 | 304,8291601 | 50 | -1,951339172 | Clustered | 0,001423 | 491 | ZFP36L1 |
| SLC12A7 | 826,3760294 | 20 | -1,860019738 | Clustered | 3,17E-04 | 490 | SLC12A7 |
| SLC6A11 | 820,4908564 | 20 | -1,42867791 | Unclustered | 0,002074 | 489 | SLC6A11 |
| NOTCH2NLB | 564,9762363 | 30 | -1,866288575 | Clustered | 0,002703 | 483,75 | NOTCH2NLB |
| FGF1 | 1279,794806 | 15 | -1,467360833 | Seed | 5,39E-04 | 483 | FGF1 |
| SFXN5 | 531,2031935 | 31 | -1,38940994 | Clustered | 0,005273 | 482,25 | SFXN5 |
| CLDN5 | 1071,120052 | 16 | -2,353107711 | Clustered | 1,84E-04 | 482 | CLDN5 |
| HEYL | 1858,185714 | 11 | -2,538127002 | Unclustered | 3,59E-05 | 481,5 | HEYL |
| LYPD1 | 862,9740172 | 18 | -1,200594942 | Clustered | 0,001571 | 481,5 | LYPD1 |
| CTNS | 752,7071627 | 20 | -1,182720818 | Clustered | 0,006719 | 481 | CTNS |
| HSDL2 | 480,0189443 | 32 | -1,115928511 | Clustered | 0,001251 | 480,25 | HSDL2 |
| TRIM41 | 903,0398078 | 17 | -1,016241271 | Seed | 9,44E-04 | 479 | TRIM41 |
| FAM167A | 657,469322 | 24 | -1,435617347 | Unclustered | 3,74E-04 | 478,25 | FAM167A |
| TNFSF12-TNFSF | 482,9663707 | 31 | -1,628718387 | Clustered | 5,09E-04 | 478,25 | TNFSF12-TNFSF13 |
| HIPK2 | 1019,719189 | 16 | -1,967162245 | Clustered | 2,64E-04 | 478 | HIPK2 |
| ADD3 | 514,6816103 | 30 | -1,394195951 | Clustered | 8,68E-05 | 477,75 | ADD3 |
| RAI14 | 1500,409391 | 12 | -1,347012214 | Clustered | 7,72E-04 | 477,5 | RAI14 |
| CSPG4 | 1498,15025 | 12 | -1,361765222 | Unclustered | 0,002509 | 477 | CSPG4 |
| MT-ND4 | 872,7194555 | 17 | -4,027050178 | Clustered | 2,33E-06 | 476,5 | MT-ND4 |
| SGMS2 | 1135,017701 | 15 | -1,502620368 | Clustered | 0,001171 | 476 | SGMS2 |
| TAX1BP3 | 526,5152811 | 28 | -1,251370638 | Clustered | 0,002858 | 474,75 | TAX1BP3 |
| DAAM2 | 1628,254826 | 11 | -1,237126429 | Unclustered | 0,00324 | 474 | DAAM2 |
| PECAM1 | 787,2034495 | 18 | -1,627381019 | Clustered | 3,30E-05 | 474 | PECAM1 |
| SLC25A18 | 333,2010216 | 37 | -1,733135343 | Clustered | 0,001988 | 473,75 | SLC25A18 |
| PTH1R | 330,2752825 | 37 | -2,024936975 | Clustered | 0,001138 | 473,25 | PTH1R |
| ASPH | 306,052853 | 39 | -1,207177235 | Clustered | 0,002862 | 472,5 | ASPH |
| GDF11 | 573,4473682 | 25 | -1,017926931 | Unclustered | 0,008962 | 472,25 | GDF11 |
| SULT1C4 | 252,1869732 | 43 | -1,573130154 | Clustered | 4,05E-04 | 471,75 | SULT1C4 |
| TNS1 | 2425,707692 | 9 | -1,237784263 | Unclustered | 8,86E-04 | 469,75 | TNS1 |
| ZBTB20 | 610,124068 | 23 | -1,604634585 | Unclustered | 0,006908 | 468,75 | ZBTB20 |
| SLC6A12 | 4920,220985 | 6 | -1,395309706 | Unclustered | 5,84E-04 | 467,75 | SLC6A12 |
| MTRNR2L12 | 780,6602503 | 17 | -3,672012641 | Clustered | 1,41E-05 | 466,5 | MTRNR2L12 |
| TMEM176A | 826,7983593 | 16 | -1,463671427 | Clustered | 0,006074 | 466 | TMEM176A |
| NEK6 | 298,4774613 | 36 | -1,491434503 | Clustered | 0,001635 | 464,25 | NEK6 |
| ADGRL4 | 624,3485383 | 22 | -2,03094986 | Unclustered | 9,33E-05 | 463,75 | ADGRL4 |
| SP100 | 913,6781748 | 15 | -1,962485973 | Unclustered | 7,28E-04 | 463,5 | SP100 |
| MT-ATP6 | 803,4209624 | 16 | -3,676488778 | Clustered | 5,47E-06 | 462,5 | MT-ATP6 |
| FOXO1 | 417,3559195 | 27 | -1,61670118 | Clustered | 9,99E-04 | 461,5 | FOXO1 |
| NTRK2 | 606,1425392 | 22 | -1,026677245 | Unclustered | 0,007432 | 461,25 | NTRK2 |
| TP53BP2 | 875,4874958 | 15 | -1,464917231 | Unclustered | 0,001873 | 461 | TP53BP2 |
| P2RY2 | 329,6559349 | 31 | -2,947100249 | Clustered | 8,14E-05 | 460,75 | P2RY2 |
| PIEZO1 | 446,6980409 | 26 | -1,295432474 | Unclustered | 0,007377 | 460,75 | PIEZO1 |
| SMARCC1 | 1002,830768 | 14 | -1,652264679 | Unclustered | 0,003231 | 459,75 | SMARCC1 |
| EPHA2 | 1482,196844 | 10 | -2,192518902 | Unclustered | 2,20E-04 | 457,5 | EPHA2 |

| | | | | | | | |
|---|---|---|---|---|---|---|---|
| HEPACAM | 526,3233052 | 23 | -1,278129502 | Clustered | 0,005522 | 457,25 | HEPACAM |
| IFITM1 | 1784,896715 | 9 | -1,912200671 | Clustered | 9,33E-05 | 456,75 | IFITM1 |
| PSMB8 | 338,4905615 | 28 | -1,23646869 | Clustered | 0,003075 | 455,25 | PSMB8 |
| TSPAN4 | 631,8457705 | 20 | -1,26399507 | Clustered | 0,005757 | 455 | TSPAN4 |
| LAMB2 | 488,9025361 | 23 | -1,260994773 | Unclustered | 0,008809 | 454,75 | LAMB2 |
| CDK2 | 1097,907628 | 12 | -2,093539009 | Unclustered | 7,72E-04 | 454,5 | CDK2 |
| HEPN1 | 559,7313588 | 22 | -1,330638349 | Seed | 0,006798 | 454,25 | HEPN1 |
| BGN | 1071,607209 | 12 | -1,412240232 | Unclustered | 6,17E-04 | 453,5 | BGN |
| FAM167B | 294,5332808 | 31 | -2,378540691 | Clustered | 0,001171 | 453,25 | FAM167B |
| FBXW4 | 943,0140169 | 13 | -1,235851852 | Clustered | 7,41E-04 | 452,25 | FBXW4 |
| TRIM38 | 534,0395826 | 22 | -1,881024329 | Clustered | 0,002134 | 451,75 | TRIM38 |
| KLHL5 | 1174,63267 | 11 | -1,000442389 | Clustered | 0,001793 | 449,5 | KLHL5 |
| AHCYL1 | 377,8679713 | 24 | -1,253678447 | Clustered | 0,005349 | 448,25 | AHCYL1 |
| BMPR1B | 252,4055409 | 31 | -1,773506384 | Clustered | 0,002151 | 448,25 | BMPR1B |
| FADS2 | 205,5568384 | 35 | -1,438260845 | Clustered | 5,03E-04 | 447,75 | FADS2 |
| GPR146 | 187,5487673 | 36 | -1,780060419 | Clustered | 6,66E-04 | 446,75 | GPR146 |
| RXRA | 325,5319294 | 26 | -1,197629865 | Clustered | 0,004191 | 446,75 | RXRA |
| SASH1 | 404,7911374 | 23 | -1,249271093 | Clustered | 7,07E-04 | 446,75 | SASH1 |
| PPP1R18 | 1503,746538 | 9 | -1,296193966 | Unclustered | 0,001768 | 446,25 | PPP1R18 |
| ALDH6A1 | 211,3437956 | 33 | -1,329269544 | Clustered | 0,00341 | 445,5 | ALDH6A1 |
| KTN1 | 595,3982559 | 19 | -1,311865822 | Clustered | 0,004104 | 444,75 | KTN1 |
| PLCD3 | 215,7109628 | 32 | -1,482202009 | Clustered | 7,39E-04 | 444,25 | PLCD3 |
| TGM2 | 632,0634306 | 18 | -3,10391131 | Clustered | 3,85E-06 | 444 | TGM2 |
| CASC3 | 2602,306606 | 6 | -1,231800035 | Unclustered | 1,12E-05 | 441,75 | CASC3 |
| ADIRF | 771,1309812 | 14 | -1,721681349 | Unclustered | 7,41E-04 | 440,75 | ADIRF |
| TNIP2 | 952,8540156 | 11 | -1,381970399 | Unclustered | 0,001553 | 438,5 | TNIP2 |
| RIN3 | 111,503154 | 39 | -1,808451219 | Clustered | 0,002664 | 437 | RIN3 |
| IL6R | 370,7201834 | 22 | -1,753860184 | Unclustered | 0,004889 | 435,25 | IL6R |
| HVCN1 | 250,6332674 | 26 | -1,106255747 | Clustered | 0,003916 | 434,75 | HVCN1 |
| ROCK1 | 599,1027851 | 17 | -1,540575981 | Clustered | 0,005294 | 433,5 | ROCK1 |
| SOCS3 | 1288,729352 | 9 | -5,057118924 | Unclustered | 3,85E-06 | 432,25 | SOCS3 |
| CSF3 | 1042,753032 | 10 | -7,665694665 | Clustered | 2,06E-05 | 432 | CSF3 |
| GSDMD | 547,6359681 | 18 | -1,702679736 | Clustered | 0,003222 | 432 | GSDMD |
| KIF1C | 429,5340481 | 20 | -1,877366373 | Clustered | 3,78E-04 | 431,5 | KIF1C |
| PLEKHB1 | 784,8738394 | 12 | -1,466662639 | Unclustered | 1,21E-04 | 431 | PLEKHB1 |
| ABHD15 | 354,6908602 | 21 | -1,217718283 | Unclustered | 0,007421 | 428,5 | ABHD15 |
| MOCS1 | 952,8052514 | 10 | -1,092260735 | Unclustered | 0,005916 | 428 | MOCS1 |
| DOK1 | 224,6242957 | 25 | -1,48130472 | Unclustered | 0,005349 | 427,75 | DOK1 |
| RMDN1 | 222,7906704 | 25 | -1,184390706 | Clustered | 7,83E-05 | 427,25 | RMDN1 |
| AL645922.1 | 933,1713979 | 10 | -1,841526071 | Clustered | 0,001632 | 427 | AL645922.1 |
| NDRG2 | 198,5988259 | 25 | -1,597002023 | Clustered | 1,88E-04 | 423,75 | NDRG2 |
| PIPOX | 391,8108697 | 19 | -1,310194259 | Unclustered | 0,005402 | 422,25 | PIPOX |
| ID4 | 276,9521559 | 22 | -1,193830252 | Clustered | 0,007501 | 420,75 | ID4 |
| MTRNR2L8 | 624,2569896 | 15 | -3,489508239 | Seed | 2,35E-05 | 420,5 | MTRNR2L8 |
| CSF1 | 1419,482572 | 7 | -1,774379524 | Unclustered | 4,03E-04 | 420 | CSF1 |
| AXL | 366,5796323 | 19 | -1,658079217 | Clustered | 0,002732 | 419,75 | AXL |
| NT5DC2 | 168,2668045 | 26 | -1,348990045 | Clustered | 0,00412 | 418,75 | NT5DC2 |
| WNK1 | 1397,602018 | 7 | -1,270258906 | Unclustered | 2,43E-04 | 418 | WNK1 |
| HSD3B7 | 252,5551469 | 22 | -1,625352373 | Clustered | 0,006803 | 417,75 | HSD3B7 |
| SRGAP1 | 127,5224642 | 27 | -1,371026249 | Clustered | 0,006967 | 417 | SRGAP1 |
| WWTR1 | 973,8616948 | 9 | -2,036078263 | Unclustered | 6,30E-06 | 415,75 | WWTR1 |
| SLCO2B1 | 2127,34654 | 5 | -1,798274093 | Unclustered | 2,91E-05 | 415,25 | SLCO2B1 |
| TLR4 | 268,7995974 | 21 | -1,296284539 | Unclustered | 0,005757 | 414,5 | TLR4 |
| IL17RD | 234,2778982 | 22 | -1,281779741 | Unclustered | 4,05E-04 | 413,75 | IL17RD |
| ANGPT2 | 358,1836958 | 18 | -2,317587665 | Unclustered | 3,77E-04 | 413,5 | ANGPT2 |
| C2orf72 | 231,707901 | 22 | -1,118852666 | Unclustered | 0,002949 | 413,25 | C2orf72 |
| FSTL1 | 715,2733339 | 11 | -1,179474578 | Unclustered | 4,65E-04 | 413 | FSTL1 |
| TRIM56 | 480,7456898 | 16 | -1,871992298 | Seed | 0,003519 | 413 | TRIM56 |
| IGDCC4 | 1523,874567 | 6 | -1,257773416 | Unclustered | 0,006058 | 412,75 | IGDCC4 |
| XAF1 | 321,5307453 | 19 | -2,65281085 | Unclustered | 5,45E-04 | 412,25 | XAF1 |
| MT-CO1 | 562,6436579 | 15 | -3,71831363 | Clustered | 5,69E-06 | 412 | MT-CO1 |

| | | | | | | | |
|---|---|---|---|---|---|---|---|
| BMP7 | 269,0801065 | 20 | -1,347673368 | Clustered | 0,001583 | 411 | BMP7 |
| PRICKLE3 | 391,9734094 | 17 | -1,376240557 | Clustered | 0,007214 | 411 | PRICKLE3 |
| RHOBTB3 | 553,4697693 | 15 | -1,345882883 | Unclustered | 5,84E-04 | 411 | RHOBTB3 |
| NOTCH2NLA | 127,2324351 | 25 | -1,538980578 | Clustered | 0,004498 | 410,75 | NOTCH2NLA |
| GNG12 | 84,18182677 | 29 | -1,389779247 | Clustered | 6,41E-04 | 410 | GNG12 |
| IL17D | 162,6442954 | 23 | -1,092785099 | Unclustered | 2,44E-04 | 406,25 | IL17D |
| CNN3 | 1226,614521 | 7 | -1,224689729 | Unclustered | 0,001635 | 406 | CNN3 |
| A4GALT | 1396,773503 | 6 | -2,444517226 | Unclustered | 9,71E-05 | 405,75 | A4GALT |
| EMP3 | 1371,110846 | 6 | -1,49446434 | Unclustered | 0,006982 | 403,75 | EMP3 |
| SQOR | 2127,869878 | 4 | -1,562265002 | Unclustered | 9,45E-04 | 403,5 | SQOR |
| SH3BP2 | 217,6499918 | 20 | -1,785236881 | Unclustered | 0,001971 | 401,5 | SH3BP2 |
| CYBRD1 | 453,735639 | 15 | -1,201400608 | Unclustered | 0,006798 | 401 | CYBRD1 |
| SPATA21 | 323,9798553 | 17 | -1,523160117 | Unclustered | 5,34E-04 | 401 | SPATA21 |
| TGFBR3 | 445,389639 | 15 | -1,285143463 | Seed | 0,002286 | 399 | TGFBR3 |
| ORAI3 | 92,86730893 | 24 | -1,134601436 | Unclustered | 0,008007 | 398,25 | ORAI3 |
| ASAP3 | 302,0039125 | 17 | -1,207589958 | Unclustered | 0,001327 | 397,5 | ASAP3 |
| ADCY4 | 328,7933804 | 16 | -2,630614353 | Clustered | 1,15E-04 | 395,5 | ADCY4 |
| PON2 | 577,793368 | 12 | -1,411935776 | Clustered | 0,003822 | 395 | PON2 |
| HIGD1B | 852,9444175 | 8 | -1,462751158 | Unclustered | 0,001759 | 393,75 | HIGD1B |
| GLIS2 | 731,8248514 | 9 | -1,440922021 | Unclustered | 7,26E-04 | 392,75 | GLIS2 |
| ITGA1 | 268,4150942 | 17 | -1,706161027 | Clustered | 0,002596 | 392,5 | ITGA1 |
| APOL3 | 191,7305642 | 19 | -1,539469272 | Seed | 2,22E-04 | 391,75 | APOL3 |
| CTNNA3 | 919,5358595 | 7 | -1,600902943 | Unclustered | 0,007447 | 390 | CTNNA3 |
| C1QTNF5 | 104,0203322 | 22 | -1,493078894 | Clustered | 0,005453 | 389 | C1QTNF5 |
| MFRP | 104,0203322 | 22 | -1,483316578 | Clustered | 0,005751 | 389 | MFRP |
| NFKB2 | 791,6354655 | 8 | -1,966132851 | Clustered | 5,84E-04 | 388,25 | NFKB2 |
| BCAM | 395,3812353 | 14 | -1,275198372 | Unclustered | 0,00456 | 387,75 | BCAM |
| ARSD | 115,0058698 | 21 | -1,30672668 | Clustered | 0,007944 | 387,5 | ARSD |
| YBX3 | 696,589638 | 9 | -2,341060932 | Unclustered | 9,03E-05 | 385,25 | YBX3 |
| GPR143 | 695,3216564 | 9 | -1,7761729 | Unclustered | 0,001387 | 384,75 | GPR143 |
| ZIC4 | 372,3002254 | 14 | -1,602833545 | Unclustered | 0,002273 | 384,75 | ZIC4 |
| LPP | 186,080049 | 18 | -1,164986154 | Clustered | 0,009576 | 384,5 | LPP |
| ENSG00000288 | 693,7371153 | 9 | -1,923555455 | Clustered | 1,36E-04 | 384,25 | ENSG00000288681 |
| FGFR3 | 193,0345532 | 17 | -1,648697102 | Seed | 0,004087 | 380,5 | FGFR3 |
| BTG2 | 1379,926348 | 4 | -1,475403235 | Unclustered | 3,74E-04 | 378,5 | BTG2 |
| CASP4 | 439,2052071 | 12 | -2,722900051 | Unclustered | 2,69E-04 | 378,5 | CASP4 |
| MSRB3 | 64,1678227 | 22 | -1,160739937 | Clustered | 0,006641 | 377,75 | MSRB3 |
| BAZ1A | 146,0823143 | 18 | -2,004547621 | Clustered | 0,004483 | 377,5 | BAZ1A |
| KIF5B | 510,7108499 | 11 | -1,654899222 | Clustered | 4,09E-04 | 377 | KIF5B |
| HIF3A | 112,2948376 | 19 | -2,532296748 | Unclustered | 7,31E-04 | 376,75 | HIF3A |
| FLT1 | 710,8870894 | 8 | -2,064834164 | Unclustered | 7,77E-06 | 375,25 | FLT1 |
| STK17B | 700,693815 | 8 | -1,757789751 | Unclustered | 0,006725 | 374,25 | STK17B |
| EMX2 | 37,05369112 | 24 | -1,656747333 | Clustered | 0,004646 | 373,75 | EMX2 |
| MT2A | 449,4518991 | 11 | -2,355274858 | Unclustered | 3,77E-04 | 371,5 | MT2A |
| FAM114A1 | 537,5770199 | 10 | -1,515963698 | Unclustered | 0,007579 | 371 | FAM114A1 |
| ROM1 | 693,2739039 | 8 | -1,621610977 | Unclustered | 0,001793 | 370,75 | ROM1 |
| GADD45G | 1024,454272 | 5 | -1,555314286 | Unclustered | 0,004456 | 369,75 | GADD45G |
| GPR37L1 | 107,543827 | 18 | -1,46602439 | Clustered | 0,004275 | 369,5 | GPR37L1 |
| MAFF | 720,7258486 | 7 | -2,251282752 | Unclustered | 2,91E-05 | 369,5 | MAFF |
| LPAR1 | 338,9291383 | 12 | -1,423647602 | Unclustered | 0,006003 | 369 | LPAR1 |
| PDGFRB | 221,6960242 | 15 | -1,382247482 | Clustered | 0,00154 | 368,5 | PDGFRB |
| PRELP | 172,1057729 | 16 | -1,363573172 | Unclustered | 0,001039 | 368,5 | PRELP |
| EPHX1 | 302,121365 | 13 | -1,568443478 | Unclustered | 0,003258 | 368,25 | EPHX1 |
| PARP12 | 300,3604474 | 13 | -1,906356531 | Clustered | 0,001181 | 367,25 | PARP12 |
| PPP1R3C | 65,09546358 | 19 | -2,062377727 | Unclustered | 6,96E-04 | 363,75 | PPP1R3C |
| NOTCH2 | 190,4128219 | 15 | -1,657838263 | Clustered | 0,002962 | 363,5 | NOTCH2 |
| NBPF10 | 50,06864564 | 20 | -2,313898165 | Unclustered | 3,74E-04 | 361,5 | NBPF10 |
| C3orf70 | 690,3990736 | 7 | -1,088806329 | Unclustered | 5,35E-04 | 361 | C3orf70 |
| PLIN4 | 564,4736736 | 9 | -2,024300177 | Unclustered | 0,002858 | 360,75 | PLIN4 |
| ERBB2 | 667,7399645 | 8 | -1,037186729 | Unclustered | 0,003444 | 360,25 | ERBB2 |
| TGFB1I1 | 175,3549302 | 15 | -1,52121859 | Clustered | 0,00414 | 360 | TGFB1I1 |

| | | | | | | | |
|---|---|---|---|---|---|---|---|
| S1PR1 | 736,2395487 | 6 | -1,50809941 | Unclustered | 0,002135 | 359,75 | S1PR1 |
| RBM38 | 169,4535297 | 15 | -1,017851137 | Unclustered | 0,007024 | 358,5 | RBM38 |
| RPGR | 112,4014675 | 16 | -2,101963651 | Unclustered | 0,001404 | 358,5 | RPGR |
| GIMAP1-GIMAI | 648,2465358 | 8 | -1,887303615 | Unclustered | 8,80E-04 | 358,25 | GIMAP1-GIMAP5 |
| BCL6B | 310,1083437 | 11 | -2,26565124 | Clustered | 4,54E-05 | 355,5 | BCL6B |
| DDR1 | 714,5635571 | 6 | -1,217029732 | Unclustered | 0,007698 | 354,75 | DDR1 |
| DEPP1 | 808,1120876 | 5 | -3,780524925 | Unclustered | 3,85E-06 | 354,25 | DEPP1 |
| REST | 241,1908026 | 12 | -1,434342328 | Unclustered | 0,008341 | 353 | REST |
| ZIC2 | 697,5676343 | 6 | -1,4931126 | Unclustered | 0,005091 | 352,25 | ZIC2 |
| TMEM176B | 863,251588 | 4 | -1,407875207 | Unclustered | 0,006719 | 348 | TMEM176B |
| ZHX2 | 686,9977742 | 6 | -1,458990584 | Unclustered | 3,20E-04 | 347,75 | ZHX2 |
| ACAA2 | 266,1487737 | 11 | -1,214383757 | Unclustered | 9,23E-04 | 347,5 | ACAA2 |
| CAPG | 686,2103144 | 6 | -1,191250852 | Unclustered | 0,00683 | 346,75 | CAPG |
| MT-ND6 | 102,1042644 | 15 | -4,456542453 | Clustered | 2,06E-05 | 345,5 | MT-ND6 |
| CFB | 399,6215049 | 9 | -2,101993573 | Unclustered | 9,84E-04 | 344,75 | CFB |
| MYO1C | 168,9851176 | 13 | -1,185501601 | Clustered | 0,008319 | 344,25 | MYO1C |
| GRAP | 240,8366927 | 11 | -2,368205574 | Unclustered | 1,15E-04 | 344 | GRAP |
| MT-CYB | 87,16308845 | 15 | -3,594818145 | Clustered | 1,07E-05 | 342 | MT-CYB |
| MT-ND1 | 87,16308845 | 15 | -3,620096698 | Clustered | 8,25E-06 | 342 | MT-ND1 |
| MT-ND2 | 87,16308845 | 15 | -3,677767932 | Clustered | 5,47E-06 | 342 | MT-ND2 |
| AKR1C3 | 36,40320344 | 18 | -1,640051357 | Unclustered | 0,003751 | 340,5 | AKR1C3 |
| PLA2G5 | 280,6139747 | 10 | -2,142213999 | Unclustered | 5,48E-04 | 340 | PLA2G5 |
| MT-CO3 | 73,59518739 | 15 | -3,295420852 | Clustered | 1,59E-05 | 338 | MT-CO3 |
| NOTCH4 | 205,8742711 | 11 | -1,258260133 | Clustered | 0,003096 | 338 | NOTCH4 |
| MMRN2 | 1370 | 2 | -1,146755225 | Unclustered | 5,92E-04 | 337,75 | MMRN2 |
| ANXA2 | 1370 | 2 | -2,569097094 | Unclustered | 3,31E-04 | 337,75 | ANXA2 |
| TGFBR2 | 1370 | 2 | -1,874318986 | Unclustered | 6,23E-04 | 337,75 | TGFBR2 |
| GPR39 | 115,0141078 | 13 | -1,184345831 | Clustered | 0,001374 | 336,75 | GPR39 |
| AL603832.3 | 549,780488 | 7 | -1,29605599 | Clustered | 3,50E-04 | 336,5 | AL603832.3 |
| GLI3 | 96,35313834 | 14 | -2,162783091 | Unclustered | 9,27E-04 | 335,75 | GLI3 |
| INHBB | 687,4145536 | 5 | -1,330488092 | Unclustered | 0,001668 | 334,25 | INHBB |
| TP53INP1 | 236,3582486 | 10 | -1,274833382 | Unclustered | 0,004908 | 333,5 | TP53INP1 |
| TNFRSF10D | 305,0890073 | 9 | -3,71601299 | Seed | 1,55E-04 | 331,25 | TNFRSF10D |
| PAIP2B | 73,35926896 | 14 | -1,844812379 | Clustered | 2,49E-04 | 329,25 | PAIP2B |
| CMTM6 | 26,93385773 | 17 | -1,439429832 | Unclustered | 0,004804 | 327,5 | CMTM6 |
| SEMA3F | 64,58671763 | 14 | -1,822020084 | Unclustered | 0,001377 | 327,25 | SEMA3F |
| HSPG2 | 196,6219285 | 10 | -1,490776419 | Unclustered | 0,006186 | 326,5 | HSPG2 |
| APOL6 | 32,18595333 | 16 | -1,879499042 | Unclustered | 0,001571 | 323,5 | APOL6 |
| MTURN | 88,16476925 | 12 | -1,392039484 | Clustered | 2,42E-04 | 323 | MTURN |
| NEK7 | 688,552381 | 4 | -1,98093825 | Unclustered | 3,30E-05 | 322,5 | NEK7 |
| TP53INP2 | 184,657017 | 10 | -1,785696679 | Clustered | 6,23E-04 | 322,5 | TP53INP2 |
| MOBP | 686,4166667 | 4 | -1,477051819 | Unclustered | 0,006463 | 321 | MOBP |
| NFATC2 | 76,81033776 | 12 | -1,978467444 | Clustered | 8,32E-04 | 318,5 | NFATC2 |
| TMCO3 | 722,7836207 | 3 | -1,041155316 | Unclustered | 1,34E-04 | 317,25 | TMCO3 |
| FOXO4 | 99,7333342 | 11 | -1,51608528 | Clustered | 3,82E-04 | 316,5 | FOXO4 |
| DTNA | 585,7583642 | 5 | -1,406438471 | Unclustered | 3,30E-05 | 316,25 | DTNA |
| DCHS1 | 461,9801813 | 6 | -1,052851108 | Unclustered | 0,006463 | 316,25 | DCHS1 |
| EMP2 | 341,3259354 | 7 | -1,630279381 | Unclustered | 0,001036 | 316 | EMP2 |
| SUCLG2 | 132,7241709 | 10 | -1,282802638 | Seed | 9,16E-04 | 315 | SUCLG2 |
| CMTM3 | 697,5739776 | 3 | -1,801812882 | Unclustered | 9,84E-04 | 311,75 | CMTM3 |
| TBX2 | 38,12104868 | 14 | -2,056241856 | Unclustered | 9,61E-04 | 311,25 | TBX2 |
| LGALS3 | 639,9539591 | 4 | -1,652500884 | Unclustered | 1,89E-04 | 310,5 | LGALS3 |
| GLUL | 72,52171778 | 11 | -1,510164512 | Clustered | 0,005294 | 308,5 | GLUL |
| CACHD1 | 234,8365961 | 8 | -1,300714966 | Unclustered | 0,001049 | 306,75 | CACHD1 |
| CTDSP2 | 298,0468111 | 7 | -1,02514443 | Unclustered | 0,002068 | 306 | CTDSP2 |
| FXYD3 | 98,52288769 | 10 | -2,006802357 | Unclustered | 0,003721 | 306 | FXYD3 |
| ASCL1 | 30,66675127 | 14 | -1,834153254 | Unclustered | 0,0095 | 305,25 | ASCL1 |
| B4GALT1 | 353,4783447 | 6 | -1,934418835 | Unclustered | 3,22E-04 | 304,75 | B4GALT1 |
| PLEKHG2 | 157,2743885 | 9 | -1,595442845 | Unclustered | 0,005273 | 304,25 | PLEKHG2 |
| MT-ND4L | 34,19243803 | 13 | -3,976062275 | Clustered | 1,36E-04 | 303,5 | MT-ND4L |
| MT-ND5 | 34,19243803 | 13 | -4,050647525 | Clustered | 1,03E-04 | 303,5 | MT-ND5 |

| | | | | | | | |
|---|---|---|---|---|---|---|---|
| NLRC5 | 143,2302823 | 9 | -1,608087108 | Clustered | 0,006555 | 303,25 | NLRC5 |
| ACACB | 136,7164889 | 9 | -1,237046031 | Unclustered | 0,008811 | 302,75 | ACACB |
| PARP14 | 133,5071171 | 9 | -1,437929422 | Unclustered | 0,001216 | 302,25 | PARP14 |
| FGR | 686 | 3 | -1,690078019 | Unclustered | 0,001138 | 302 | FGR |
| IRF7 | 686 | 3 | -2,301206327 | Unclustered | 8,68E-04 | 302 | IRF7 |
| MDFIC | 686 | 3 | -1,382639691 | Unclustered | 9,84E-04 | 302 | MDFIC |
| MINDY1 | 686 | 3 | -1,168484419 | Unclustered | 0,005627 | 302 | MINDY1 |
| HES6 | 34,12348035 | 12 | -1,503770258 | Unclustered | 0,008294 | 296,5 | HES6 |
| ASRGL1 | 12,57867202 | 15 | -1,406634716 | Unclustered | 0,004986 | 295,5 | ASRGL1 |
| NOL8 | 45,40129495 | 11 | -1,213074098 | Unclustered | 0,008979 | 295,5 | NOL8 |
| GAREM2 | 24,35358349 | 13 | -1,54920447 | Unclustered | 0,007018 | 294,25 | GAREM2 |
| PAQR8 | 60,15924722 | 10 | -1,502368997 | Unclustered | 0,001793 | 294 | PAQR8 |
| SEC14L2 | 16,59824901 | 14 | -1,363937408 | Unclustered | 0,004908 | 291,25 | SEC14L2 |
| ZBTB42 | 603,6222096 | 3 | -2,52904372 | Unclustered | 3,74E-04 | 291,25 | ZBTB42 |
| TIMP1 | 49,4660607 | 10 | -1,902564129 | Unclustered | 6,96E-04 | 289 | TIMP1 |
| ALDH7A1 | 6,478881781 | 15 | -1,082981247 | Unclustered | 0,008708 | 286,5 | ALDH7A1 |
| BBOX1 | 118,3109379 | 8 | -1,590643362 | Clustered | 5,43E-04 | 286,25 | BBOX1 |
| MYOM1 | 112,1780727 | 8 | -1,409166961 | Unclustered | 0,001646 | 283,75 | MYOM1 |
| SWAP70 | 63,99327625 | 9 | -1,073922606 | Unclustered | 0,001759 | 282,75 | SWAP70 |
| GPRC5C | 104,2637246 | 8 | -1,559481274 | Unclustered | 0,002537 | 282,25 | GPRC5C |
| MFNG | 60,90712522 | 9 | -2,035155605 | Clustered | 5,94E-04 | 281,75 | MFNG |
| MYL3 | 60,63119127 | 9 | -2,143994953 | Unclustered | 9,84E-04 | 281,25 | MYL3 |
| NCKAP5 | 17,87868759 | 12 | -2,346841224 | Clustered | 0,001321 | 281 | NCKAP5 |
| DAG1 | 59,51121103 | 9 | -1,141489043 | Unclustered | 0,004002 | 280,25 | DAG1 |
| PDLIM3 | 36,23031778 | 10 | -1,088211169 | Unclustered | 9,93E-04 | 279,5 | PDLIM3 |
| EMP1 | 686 | 2 | -2,825975072 | Unclustered | 8,03E-05 | 278 | EMP1 |
| APOLD1 | 686 | 2 | -1,449903822 | Unclustered | 3,95E-04 | 278 | APOLD1 |
| BTN2A2 | 686 | 2 | -1,45073207 | Unclustered | 0,001216 | 278 | BTN2A2 |
| IL1R1 | 686 | 2 | -1,931960733 | Unclustered | 0,00147 | 278 | IL1R1 |
| MT1F | 686 | 2 | -2,309297011 | Unclustered | 9,99E-04 | 278 | MT1F |
| NKD1 | 686 | 2 | -1,0644835 | Unclustered | 5,46E-04 | 278 | NKD1 |
| PODXL | 686 | 2 | -1,139823179 | Unclustered | 4,38E-04 | 278 | PODXL |
| S100A10 | 686 | 2 | -1,803158736 | Unclustered | 9,11E-04 | 278 | S100A10 |
| SOWAHC | 686 | 2 | -1,520179442 | Unclustered | 8,55E-04 | 278 | SOWAHC |
| TEAD3 | 686 | 2 | -1,754569434 | Unclustered | 0,004018 | 278 | TEAD3 |
| LRRCC1 | 56,10550257 | 9 | -1,481619664 | Clustered | 0,004599 | 277,75 | LRRCC1 |
| CDC45 | 55,487823 | 9 | -3,291957179 | Unclustered | 0,003369 | 277,25 | CDC45 |
| MGST1 | 32,97355365 | 10 | -1,570978797 | Unclustered | 0,002017 | 277 | MGST1 |
| GJA1 | 52,39421433 | 9 | -1,748579596 | Clustered | 0,004842 | 276,75 | GJA1 |
| ELK3 | 257,1524665 | 5 | -1,381735559 | Unclustered | 3,00E-04 | 275,75 | ELK3 |
| GJA4 | 175,5985257 | 6 | -2,636118441 | Seed | 1,91E-05 | 274,75 | GJA4 |
| C16orf74 | 11,66099617 | 12 | -2,288172019 | Unclustered | 7,26E-04 | 274 | C16orf74 |
| TRIP10 | 29,98762225 | 10 | -1,730634188 | Unclustered | 0,001579 | 274 | TRIP10 |
| ZNF366 | 46,14117497 | 9 | -2,06484845 | Unclustered | 0,00541 | 272,75 | ZNF366 |
| PODN | 26,71356938 | 10 | -1,956292454 | Clustered | 3,00E-04 | 272,5 | PODN |
| CYP11A1 | 9,862858753 | 12 | -1,72509171 | Unclustered | 0,007282 | 271 | CYP11A1 |
| ZFC3H1 | 14,14015512 | 11 | -1,822728269 | Clustered | 0,008522 | 268,5 | ZFC3H1 |
| RASSF8 | 22,68698945 | 10 | -1,183038145 | Unclustered | 2,91E-05 | 267,5 | RASSF8 |
| GFAP | 58,91869208 | 8 | -3,049486844 | Unclustered | 2,33E-06 | 266,75 | GFAP |
| PTPN21 | 126,8527266 | 6 | -1,024423151 | Unclustered | 4,85E-04 | 266,25 | PTPN21 |
| ALDH1L1 | 124,2174857 | 6 | -2,098528634 | Unclustered | 0,001802 | 265,75 | ALDH1L1 |
| VWF | 56,19484685 | 8 | -2,165853271 | Unclustered | 1,41E-05 | 265,25 | VWF |
| RGL3 | 10,41609617 | 11 | -1,904471676 | Unclustered | 0,00483 | 264,5 | RGL3 |
| OLIG1 | 340,9303006 | 3 | -1,042468016 | Unclustered | 5,42E-04 | 262,75 | OLIG1 |
| CSRP1 | 185,4450093 | 5 | -1,244654646 | Unclustered | 0,005273 | 262,25 | CSRP1 |
| CARHSP1 | 185,3259441 | 5 | -1,271435033 | Unclustered | 0,004889 | 261,75 | CARHSP1 |
| NAA16 | 16,36459636 | 10 | -2,162939401 | Clustered | 0,003787 | 260,5 | NAA16 |
| DENND2A | 170,2567452 | 5 | -1,059501974 | Unclustered | 7,28E-04 | 259,25 | DENND2A |
| PITPNC1 | 13,09614928 | 10 | -1,094189622 | Unclustered | 0,00341 | 258 | PITPNC1 |
| PHYHD1 | 166,820925 | 5 | -1,764836539 | Unclustered | 0,006625 | 257,25 | PHYHD1 |
| NECTIN2 | 41,16185511 | 8 | -1,042160996 | Unclustered | 0,007421 | 256,25 | NECTIN2 |

| | | | | | | | |
|---|---|---|---|---|---|---|---|
| OTUD7B | 80,11376757 | 6 | -1,184432345 | Unclustered | 0,008182 | 253,75 | OTUD7B |
| C4orf3 | 291,0586032 | 3 | -1,19896473 | Unclustered | 2,33E-06 | 252,25 | C4orf3 |
| SLC6A13 | 57,82283939 | 6 | -1,254881662 | Unclustered | 0,007549 | 244,75 | SLC6A13 |
| LPAR3 | 13,06708021 | 9 | -2,308006191 | Unclustered | 8,95E-04 | 244,25 | LPAR3 |
| MAML2 | 230,1191726 | 3 | -1,306280523 | Unclustered | 0,003522 | 243,25 | MAML2 |
| PLEKHA4 | 22,79703369 | 8 | -2,368811388 | Unclustered | 9,44E-04 | 241,75 | PLEKHA4 |
| MCM7 | 32,82380891 | 7 | -1,28544769 | Clustered | 0,002702 | 241 | MCM7 |
| CFAP53 | 3,651300011 | 10 | -2,067448387 | Clustered | 0,008347 | 240,5 | CFAP53 |
| STOM | 47,41884395 | 6 | -1,439131316 | Unclustered | 2,74E-04 | 240,25 | STOM |
| NUPR1 | 20,69092853 | 8 | -2,367210485 | Unclustered | 1,93E-04 | 239,75 | NUPR1 |
| RGS20 | 9,903352191 | 9 | -1,842573557 | Unclustered | 0,001049 | 239,75 | RGS20 |
| IFITM2 | 77,16114653 | 5 | -2,425609004 | Unclustered | 1,16E-04 | 239,25 | IFITM2 |
| FAM189A2 | 28,4527183 | 7 | -1,577624333 | Clustered | 0,006364 | 238 | FAM189A2 |
| EPS8 | 45,15169228 | 6 | -1,020449962 | Unclustered | 0,007421 | 237,75 | EPS8 |
| NDUFA4L2 | 0,313593352 | 11 | -1,17947081 | Clustered | 0,002979 | 237 | NDUFA4L2 |
| MYBPC1 | 6,473735905 | 9 | -1,386999789 | Unclustered | 0,004809 | 234,25 | MYBPC1 |
| TGIF1 | 6,198829156 | 9 | -1,878536163 | Unclustered | 1,91E-04 | 232,75 | TGIF1 |
| ZMYM5 | 58,20808294 | 5 | -1,437730852 | Unclustered | 0,006719 | 231,25 | ZMYM5 |
| TAGLN2 | 20,76726774 | 7 | -1,625228543 | Unclustered | 8,14E-05 | 231 | TAGLN2 |
| TBC1D16 | 5,397583133 | 9 | -1,408040852 | Unclustered | 9,99E-04 | 230,75 | TBC1D16 |
| C10orf105 | 19,80116838 | 7 | -3,477231703 | Unclustered | 3,85E-06 | 229,5 | C10orf105 |
| MAP4K4 | 82,82736524 | 4 | -1,607321216 | Unclustered | 0,005244 | 228,5 | MAP4K4 |
| NFATC4 | 3,411531394 | 9 | -1,738987951 | Unclustered | 0,005627 | 226,25 | NFATC4 |
| LDLRAD2 | 47,25388515 | 5 | -1,635764591 | Unclustered | 0,006739 | 225,75 | LDLRAD2 |
| NPL | 9,245901962 | 8 | -1,360717536 | Unclustered | 0,003321 | 225,25 | NPL |
| CRTAP | 68,47899219 | 4 | -1,016431922 | Unclustered | 0,003358 | 224,5 | CRTAP |
| EMCN | 14,87227132 | 7 | -1,475335048 | Unclustered | 0,004324 | 223,5 | EMCN |
| GAB2 | 24,9374365 | 6 | -1,316208468 | Seed | 0,003018 | 223,5 | GAB2 |
| SLCO1A2 | 24,9374365 | 6 | -1,413786169 | Clustered | 0,006308 | 223,5 | SLCO1A2 |
| NTSR2 | 7,082691248 | 8 | -1,513929972 | Unclustered | 0,00395 | 222,75 | NTSR2 |
| RREB1 | 44,78448947 | 5 | -1,821263449 | Unclustered | 0,004261 | 222,75 | RREB1 |
| BNIP2 | 41,33125111 | 5 | -1,117260991 | Unclustered | 4,02E-04 | 221,75 | BNIP2 |
| RASSF2 | 24,29026168 | 6 | -1,227209619 | Clustered | 8,40E-04 | 221,75 | RASSF2 |
| TAP1 | 38,53420194 | 5 | -1,379368262 | Unclustered | 0,003709 | 220,25 | TAP1 |
| MTSS2 | 19,94681327 | 6 | -1,441032266 | Clustered | 2,25E-04 | 218,25 | MTSS2 |
| MLKL | 3,707532643 | 8 | -2,345941544 | Clustered | 0,003373 | 214,75 | MLKL |
| PRR29 | 47,90637396 | 4 | -1,30971503 | Clustered | 0,001404 | 214,5 | PRR29 |
| N4BP2 | 15,45594859 | 6 | -1,530486343 | Unclustered | 0,007518 | 212,75 | N4BP2 |
| ATP1A2 | 26,24983118 | 5 | -1,369193516 | Unclustered | 0,00866 | 210,75 | ATP1A2 |
| WWC1 | 5,951023666 | 7 | -1,072287925 | Unclustered | 7,11E-04 | 210 | WWC1 |
| ITPRIPL2 | 5,827573468 | 7 | -1,529176082 | Unclustered | 0,00447 | 209,5 | ITPRIPL2 |
| NQO1 | 40,45558038 | 4 | -1,975316173 | Unclustered | 0,00115 | 208,5 | NQO1 |
| CHD7 | 1,58049397 | 8 | -1,993490846 | Clustered | 0,006268 | 207,25 | CHD7 |
| CEBPD | 62,63056369 | 3 | -1,91978246 | Unclustered | 0,002509 | 207,25 | CEBPD |
| PPP1R3G | 10,05765869 | 6 | -1,578894044 | Unclustered | 0,008862 | 206,25 | PPP1R3G |
| ANP32B | 19,768413 | 5 | -1,455503516 | Unclustered | 0,00489 | 203,25 | ANP32B |
| NIBAN2 | 7,697477579 | 6 | -1,839975179 | Unclustered | 1,34E-04 | 203,25 | NIBAN2 |
| EBF1 | 7,419334004 | 6 | -1,864375408 | Unclustered | 0,002176 | 202,75 | EBF1 |
| IFI27 | 19,49386335 | 5 | -1,323339484 | Unclustered | 0,001793 | 202,75 | IFI27 |
| SAMD4A | 49,16441082 | 3 | -1,791587171 | Unclustered | 1,82E-05 | 200,25 | SAMD4A |
| HERC5 | 15,03519929 | 5 | -1,341856143 | Unclustered | 8,46E-04 | 198,25 | HERC5 |
| ITGB4 | 24,53042968 | 4 | -2,267717271 | Unclustered | 1,36E-04 | 196,5 | ITGB4 |
| C11orf96 | 44,85412782 | 3 | -1,880298992 | Unclustered | 0,001517 | 196,25 | C11orf96 |
| LRP4 | 44,41565159 | 3 | -1,651912038 | Unclustered | 8,80E-04 | 195,25 | LRP4 |
| SLC7A5 | 23,28414923 | 4 | -1,510986045 | Unclustered | 3,02E-04 | 195 | SLC7A5 |
| KREMEN1 | 12,23797022 | 5 | -1,163781134 | Seed | 9,61E-04 | 194,75 | KREMEN1 |
| ZNF853 | 10,19419332 | 5 | -1,060779345 | Unclustered | 5,08E-04 | 192,75 | ZNF853 |
| BATF2 | 2,418747172 | 6 | -1,697439859 | Unclustered | 0,005864 | 190,25 | BATF2 |
| NOSTRIN | 19,06347015 | 4 | -1,767063986 | Unclustered | 0,003692 | 190 | NOSTRIN |
| CHST3 | 82,82604741 | 2 | -1,33981329 | Unclustered | 0,003313 | 189,25 | CHST3 |
| SLC2A4 | 2,063783627 | 6 | -2,139307209 | Clustered | 0,003966 | 188,75 | SLC2A4 |

| | | | | | | | |
|---|---|---|---|---|---|---|---|
| ENSG0000028[illegible] | 7,214577212 | 5 | -1,757835131 | Unclustered | 0,001191 | 188,25 | ENSG00000288656 |
| CD81 | 1,212137356 | 6 | -1,037626519 | Clustered | 0,009153 | 185,75 | CD81 |
| MT-ATP8 | 0 | 9 | -3,382311124 | Clustered | 3,28E-05 | 183,5 | MT-ATP8 |
| ROBO4 | 22,62454895 | 3 | -1,574395212 | Unclustered | 9,23E-04 | 178,75 | ROBO4 |
| SLC14A1 | 8,511664459 | 4 | -2,519456626 | Unclustered | 7,86E-04 | 177,5 | SLC14A1 |
| EFNB1 | 2,358539288 | 5 | -1,301085081 | Unclustered | 0,007609 | 175,75 | EFNB1 |
| PGGHG | 2,076925016 | 5 | -1,851659446 | Unclustered | 0,001953 | 175,25 | PGGHG |
| ADAMTSL4 | 46,74821227 | 2 | -1,755177351 | Unclustered | 0,004511 | 174,25 | ADAMTSL4 |
| CD109 | 17,50280831 | 3 | -1,653818343 | Unclustered | 0,002289 | 173,75 | CD109 |
| SLC4A4 | 1,838040919 | 5 | -1,41458825 | Unclustered | 0,00573 | 173,25 | SLC4A4 |
| SORCS2 | 16,74721794 | 3 | -1,103043887 | Unclustered | 0,004961 | 173,25 | SORCS2 |
| FAM181A | 6,245723457 | 4 | -1,48439922 | Unclustered | 0,004365 | 173 | FAM181A |
| EHD2 | 5,726916251 | 4 | -1,284837878 | Unclustered | 0,006233 | 171 | EHD2 |
| CCDC8 | 4,838358365 | 4 | -1,634474909 | Unclustered | 0,008345 | 170 | CCDC8 |
| GPER1 | 4,292181486 | 4 | -1,618821039 | Unclustered | 5,39E-04 | 169,5 | GPER1 |
| FLI1 | 12,38711726 | 3 | -1,235825394 | Unclustered | 0,006684 | 168,25 | FLI1 |
| AIF1L | 3,742255895 | 4 | -1,162635702 | Unclustered | 0,006803 | 168 | AIF1L |
| BAG3 | 33,32411028 | 2 | -2,062314859 | Unclustered | 0,001191 | 165,25 | BAG3 |
| SMIM3 | 31,72181725 | 2 | -1,751560359 | Unclustered | 4,01E-04 | 163,25 | SMIM3 |
| RAB20 | 30,13643448 | 2 | -2,171334543 | Unclustered | 1,95E-04 | 162,25 | RAB20 |
| CCDC9B | 1,849601083 | 4 | -1,338571635 | Unclustered | 0,004619 | 161,5 | CCDC9B |
| PKP4 | 0 | 7 | -1,300566039 | Clustered | 9,84E-04 | 161,25 | PKP4 |
| ITGA6 | 6,622893379 | 3 | -1,451713438 | Unclustered | 0,001192 | 160,25 | ITGA6 |
| LSR | 26,22163323 | 2 | -1,034921941 | Unclustered | 0,001873 | 159,25 | LSR |
| HSPB7 | 0,513311547 | 4 | -1,536566305 | Unclustered | 0,002355 | 156 | HSPB7 |
| AEBP1 | 0,421765529 | 4 | -1,804282111 | Unclustered | 0,001558 | 154,5 | AEBP1 |
| PRX | 3,931147963 | 3 | -1,462738303 | Unclustered | 0,007819 | 154,25 | PRX |
| MECOM | 0,386874237 | 4 | -2,237831711 | Unclustered | 0,001682 | 154 | MECOM |
| DLL4 | 3,777400959 | 3 | -1,802774835 | Unclustered | 0,002664 | 153,75 | DLL4 |
| SPX | 18,77431081 | 2 | -1,598224228 | Unclustered | 4,47E-05 | 150,75 | SPX |
| TRIM5 | 3,308432937 | 3 | -1,641404418 | Unclustered | 0,001793 | 150,75 | TRIM5 |
| SLC19A3 | 3,289008487 | 3 | -1,815870156 | Unclustered | 9,74E-04 | 150,25 | SLC19A3 |
| PGF | 1,991434738 | 3 | -1,712623827 | Unclustered | 0,004966 | 147,25 | PGF |
| SDC4 | 1,732940736 | 3 | -1,943169643 | Unclustered | 0,001635 | 145,75 | SDC4 |
| LRP5 | 1,122754595 | 3 | -1,089053089 | Unclustered | 0,008748 | 143,75 | LRP5 |
| RAB30 | 10,67547255 | 2 | -1,068104671 | Unclustered | 0,00578 | 142,75 | RAB30 |
| PMP2 | 0,505046328 | 3 | -1,427223601 | Unclustered | 4,42E-04 | 140,75 | PMP2 |
| SLC40A1 | 9,249831094 | 2 | -1,469448587 | Unclustered | 5,44E-04 | 139,75 | SLC40A1 |
| IL1R2 | 6,371566667 | 2 | -5,17688717 | Unclustered | 8,54E-04 | 134,75 | IL1R2 |
| SLC52A3 | 3,442609543 | 2 | -2,513996213 | Unclustered | 7,43E-04 | 127,75 | SLC52A3 |
| CDH23 | 2,805985376 | 2 | -1,540412346 | Unclustered | 0,004946 | 125,75 | CDH23 |
| AK4 | 1,206883857 | 2 | -1,042703104 | Unclustered | 0,00203 | 120,25 | AK4 |
| TMPRSS4 | 1 | 2 | -4,814013093 | Unclustered | 0,007916 | 119 | TMPRSS4 |
| JUNB | 1 | 2 | -1,251010841 | Unclustered | 0,00578 | 119 | JUNB |
| GCSH | 0,875 | 2 | -1,012071091 | Unclustered | 2,09E-05 | 118,25 | GCSH |
| MIDN | 0,797180088 | 2 | -1,112592113 | Unclustered | 0,007447 | 117,75 | MIDN |
| EPB41L5 | 0,5 | 2 | -1,25185149 | Unclustered | 0,004859 | 116,25 | EPB41L5 |
| LFNG | 0,1 | 2 | -1,206384744 | Unclustered | 0,003252 | 114,25 | LFNG |
| ICAM2 | 0 | 3 | -1,234651268 | Clustered | 0,001558 | 108,5 | ICAM2 |
| CFH | 0 | 3 | -1,599774366 | Unclustered | 0,001571 | 108,5 | CFH |
| MASP1 | 0 | 3 | -1,422347474 | Unclustered | 0,001765 | 108,5 | MASP1 |
| MT1X | 0 | 3 | -2,516431181 | Unclustered | 1,91E-04 | 108,5 | MT1X |
| CCDC190 | 0 | 3 | -2,227262291 | Unclustered | 0,007421 | 108,5 | CCDC190 |
| MMD2 | 0 | 3 | -1,94537676 | Unclustered | 7,86E-04 | 108,5 | MMD2 |
| SAMD9L | 0 | 3 | -1,936102872 | Clustered | 9,45E-04 | 108,5 | SAMD9L |
| SHTN1 | 0 | 3 | -1,285555961 | Unclustered | 0,003519 | 108,5 | SHTN1 |
| TGFB3 | 0 | 3 | -1,475855985 | Unclustered | 0,0025 | 108,5 | TGFB3 |
| CLEC2B | 0 | 2 | -2,212225907 | Unclustered | 7,46E-04 | 84,5 | CLEC2B |
| F11R | 0 | 2 | -1,493539224 | Unclustered | 0,004346 | 84,5 | F11R |
| GDPD2 | 0 | 2 | -1,891008328 | Unclustered | 0,001571 | 84,5 | GDPD2 |
| ICAM1 | 0 | 2 | -3,739200185 | Unclustered | 1,41E-05 | 84,5 | ICAM1 |

| | | | | | | | |
|---|---|---|---|---|---|---|---|
| ITGB1 | 0 | 2 | -1,054900554 | Unclustered | 5,42E-04 | 84,5 | ITGB1 |
| ITPKC | 0 | 2 | -1,803083924 | Unclustered | 9,64E-04 | 84,5 | ITPKC |
| ITPRIP | 0 | 2 | -1,635251983 | Unclustered | 0,002093 | 84,5 | ITPRIP |
| LEF1 | 0 | 2 | -1,41226395 | Unclustered | 1,73E-04 | 84,5 | LEF1 |
| LIMK2 | 0 | 2 | -1,732469478 | Unclustered | 2,20E-04 | 84,5 | LIMK2 |
| LYN | 0 | 2 | -1,187591011 | Unclustered | 0,006462 | 84,5 | LYN |
| MT1E | 0 | 2 | -1,850888415 | Unclustered | 0,002155 | 84,5 | MT1E |
| NDRG1 | 0 | 2 | -1,297166496 | Unclustered | 0,002235 | 84,5 | NDRG1 |
| ERBIN | 0 | 2 | -1,224465926 | Unclustered | 0,00483 | 84,5 | ERBIN |
| NUDT16 | 0 | 2 | -1,056980554 | Unclustered | 1,24E-04 | 84,5 | NUDT16 |
| PARP9 | 0 | 2 | -1,615259743 | Unclustered | 0,001873 | 84,5 | PARP9 |
| AL672142.1 | 0 | 2 | -1,800612227 | Unclustered | 0,009831 | 84,5 | AL672142.1 |
| PRSS35 | 0 | 2 | -1,486508658 | Unclustered | 0,006888 | 84,5 | PRSS35 |
| PSMB9 | 0 | 2 | -1,222951543 | Unclustered | 0,009335 | 84,5 | PSMB9 |
| APOL1 | 0 | 2 | -1,615113975 | Unclustered | 0,002134 | 84,5 | APOL1 |
| SH3TC1 | 0 | 2 | -1,669518564 | Unclustered | 0,002384 | 84,5 | SH3TC1 |
| SLC38A3 | 0 | 2 | -1,560707612 | Unclustered | 7,52E-04 | 84,5 | SLC38A3 |
| SLCO4A1 | 0 | 2 | -1,889303929 | Unclustered | 0,003252 | 84,5 | SLCO4A1 |
| GADD45B | 0 | 2 | -2,150634086 | Unclustered | 5,42E-04 | 84,5 | GADD45B |
| SRPX2 | 0 | 2 | -2,036232231 | Unclustered | 0,002093 | 84,5 | SRPX2 |
| SYTL4 | 0 | 2 | -2,373174122 | Unclustered | 8,68E-04 | 84,5 | SYTL4 |
| TCF7 | 0 | 2 | -1,429088318 | Unclustered | 0,003637 | 84,5 | TCF7 |
| VIM | 0 | 2 | -1,363957198 | Unclustered | 9,62E-04 | 84,5 | VIM |
| ZFP36 | 0 | 2 | -3,47263433 | Unclustered | 2,42E-05 | 84,5 | ZFP36 |
| SLC11A1 | 0 | 1 | -2,257598176 | Unclustered | 0,001426 | 49,75 | SLC11A1 |
| FCGBP | 0 | 1 | -2,19088102 | Unclustered | 0,005188 | 49,75 | FCGBP |
| GOLGA8F | 0 | 1 | -3,865731628 | Unclustered | 0,007641 | 49,75 | GOLGA8F |
| FOS | 0 | 1 | -1,274619211 | Unclustered | 0,007447 | 49,75 | FOS |
| ZAN | 0 | 1 | -4,569590401 | Unclustered | 0,002557 | 49,75 | ZAN |
| PDLIM1 | 0 | 1 | -1,51720027 | Unclustered | 0,00414 | 49,75 | PDLIM1 |
| BCL3 | 0 | 1 | -1,793057359 | Unclustered | 0,002317 | 49,75 | BCL3 |
| C4A | 0 | 1 | -1,772688637 | Unclustered | 0,004804 | 49,75 | C4A |
| C4B | 0 | 1 | -1,771903682 | Unclustered | 0,004908 | 49,75 | C4B |
| CD59 | 0 | 1 | -1,044269022 | Unclustered | 0,00448 | 49,75 | CD59 |
| BHMT2 | 0 | 1 | -1,692133349 | Unclustered | 0,00116 | 49,75 | BHMT2 |
| CTXND1 | 0 | 1 | -1,116800327 | Unclustered | 9,79E-04 | 49,75 | CTXND1 |
| CSRNP1 | 0 | 1 | -1,162244766 | Unclustered | 0,002683 | 49,75 | CSRNP1 |
| ARHGAP31 | 0 | 1 | -1,494001284 | Unclustered | 0,002053 | 49,75 | ARHGAP31 |
| CARD10 | 0 | 1 | -1,872859776 | Unclustered | 0,003365 | 49,75 | CARD10 |
| EPAS1 | 0 | 1 | -1,783423505 | Unclustered | 8,14E-05 | 49,75 | EPAS1 |
| ERLIN2 | 0 | 1 | -1,029933561 | Unclustered | 0,00679 | 49,75 | ERLIN2 |
| ARHGAP11A | 0 | 1 | -1,163178602 | Unclustered | 0,00187 | 49,75 | ARHGAP11A |
| CCDC103 | 0 | 1 | -1,425087864 | Unclustered | 0,006402 | 49,75 | CCDC103 |
| FAM187A | 0 | 1 | -1,432427029 | Unclustered | 0,006625 | 49,75 | FAM187A |
| CDH5 | 0 | 1 | -1,478212479 | Unclustered | 1,34E-04 | 49,75 | CDH5 |
| FLT4 | 0 | 1 | -1,596882824 | Unclustered | 0,004842 | 49,75 | FLT4 |
| FOXF1 | 0 | 1 | -2,424027544 | Unclustered | 3,87E-04 | 49,75 | FOXF1 |
| CCDC102A | 0 | 1 | -1,548654161 | Unclustered | 0,008319 | 49,75 | CCDC102A |
| FOXJ1 | 0 | 1 | -2,654641896 | Unclustered | 5,48E-04 | 49,75 | FOXJ1 |
| GNG11 | 0 | 1 | -1,192735778 | Unclustered | 1,24E-04 | 49,75 | GNG11 |
| ALDH3B1 | 0 | 1 | -1,691952461 | Unclustered | 0,002702 | 49,75 | ALDH3B1 |
| C2CD4B | 0 | 1 | -3,565663959 | Unclustered | 0,001975 | 49,75 | C2CD4B |
| IL32 | 0 | 1 | -1,786967483 | Unclustered | 0,002664 | 49,75 | IL32 |
| IRAK3 | 0 | 1 | -1,788485723 | Unclustered | 0,007122 | 49,75 | IRAK3 |
| CLEC1A | 0 | 1 | -2,246529107 | Unclustered | 0,001939 | 49,75 | CLEC1A |
| ITIH5 | 0 | 1 | -1,226085954 | Unclustered | 0,004456 | 49,75 | ITIH5 |
| KCNE4 | 0 | 1 | -2,791284354 | Unclustered | 8,22E-05 | 49,75 | KCNE4 |
| MDFI | 0 | 1 | -1,505522771 | Unclustered | 0,001668 | 49,75 | MDFI |
| ALDH1A1 | 0 | 1 | -1,008232492 | Unclustered | 0,003063 | 49,75 | ALDH1A1 |
| MFGE8 | 0 | 1 | -1,114736405 | Unclustered | 0,007377 | 49,75 | MFGE8 |
| APLN | 0 | 1 | -1,086458431 | Unclustered | 0,008515 | 49,75 | APLN |

| | | | | | | | |
|---|---|---|---|---|---|---|---|
| ARHGEF37 | 0 | 1 | -1,091545082 | Unclustered | 0,005694 | 49,75 | ARHGEF37 |
| MT1G | 0 | 1 | -2,302434721 | Unclustered | 0,00331 | 49,75 | MT1G |
| ID3 | 0 | 1 | -1,591307019 | Unclustered | 0,00963 | 49,75 | ID3 |
| NANOS1 | 0 | 1 | -1,287115341 | Unclustered | 0,007629 | 49,75 | NANOS1 |
| NEDD1 | 0 | 1 | -1,056498487 | Unclustered | 0,003155 | 49,75 | NEDD1 |
| DOCK1 | 0 | 1 | -1,429451544 | Unclustered | 9,59E-05 | 49,75 | DOCK1 |
| CAVIN1 | 0 | 1 | -1,20600666 | Unclustered | 0,003286 | 49,75 | CAVIN1 |
| PLA1A | 0 | 1 | -2,729647551 | Unclustered | 9,40E-04 | 49,75 | PLA1A |
| PLIN2 | 0 | 1 | -1,335445708 | Unclustered | 0,006549 | 49,75 | PLIN2 |
| ESAM | 0 | 1 | -1,270854487 | Unclustered | 0,001195 | 49,75 | ESAM |
| PPP1R13L | 0 | 1 | -1,388621439 | Unclustered | 0,007963 | 49,75 | PPP1R13L |
| PPP1R1B | 0 | 1 | -1,372705473 | Unclustered | 0,007967 | 49,75 | PPP1R1B |
| PREX2 | 0 | 1 | -1,09708132 | Unclustered | 9,16E-04 | 49,75 | PREX2 |
| RASL12 | 0 | 1 | -1,302829189 | Unclustered | 0,008534 | 49,75 | RASL12 |
| ADORA2B | 0 | 1 | -1,870108039 | Unclustered | 0,002953 | 49,75 | ADORA2B |
| NFATC1 | 0 | 1 | -1,599915997 | Unclustered | 0,003358 | 49,75 | NFATC1 |
| RPS27 | 0 | 1 | -1,045959996 | Unclustered | 0,007667 | 49,75 | RPS27 |
| S100A11 | 0 | 1 | -1,400947402 | Unclustered | 0,008979 | 49,75 | S100A11 |
| S100A8 | 0 | 1 | -3,389374203 | Unclustered | 5,84E-04 | 49,75 | S100A8 |
| S100A9 | 0 | 1 | -2,690645541 | Unclustered | 0,006967 | 49,75 | S100A9 |
| FAT1 | 0 | 1 | -1,268399754 | Unclustered | 0,008324 | 49,75 | FAT1 |
| SDS | 0 | 1 | -1,963745588 | Unclustered | 0,002962 | 49,75 | SDS |
| ISG20 | 0 | 1 | -1,375698557 | Unclustered | 0,001756 | 49,75 | ISG20 |
| SH2D2A | 0 | 1 | -2,569163918 | Unclustered | 0,001203 | 49,75 | SH2D2A |
| KCNJ16 | 0 | 1 | -1,698412379 | Unclustered | 0,002404 | 49,75 | KCNJ16 |
| SMAD6 | 0 | 1 | -1,326235665 | Unclustered | 0,008713 | 49,75 | SMAD6 |
| GIMAP2 | 0 | 1 | -1,770901905 | Unclustered | 0,002159 | 49,75 | GIMAP2 |
| SPATA1 | 0 | 1 | -1,16337369 | Unclustered | 5,46E-04 | 49,75 | SPATA1 |
| TEAD2 | 0 | 1 | -1,743559333 | Unclustered | 0,005406 | 49,75 | TEAD2 |
| TGFB1 | 0 | 1 | -1,139139912 | Unclustered | 0,006058 | 49,75 | TGFB1 |
| GOLIM4 | 0 | 1 | -1,086285922 | Unclustered | 0,003123 | 49,75 | GOLIM4 |
| GRAMD1C | 0 | 1 | -1,453108458 | Unclustered | 0,004275 | 49,75 | GRAMD1C |
| THSD4 | 0 | 1 | -1,975520956 | Unclustered | 0,002139 | 49,75 | THSD4 |
| TIPARP | 0 | 1 | -1,0739426 | Unclustered | 0,007916 | 49,75 | TIPARP |
| TM4SF1 | 0 | 1 | -1,765483369 | Unclustered | 4,02E-04 | 49,75 | TM4SF1 |
| PRKCH | 0 | 1 | -1,709716946 | Unclustered | 1,29E-04 | 49,75 | PRKCH |
| TTC23 | 0 | 1 | -1,027192634 | Unclustered | 0,004002 | 49,75 | TTC23 |
| TYMP | 0 | 1 | -2,359865705 | Unclustered | 0,002134 | 49,75 | TYMP |
| YES1 | 0 | 1 | -1,472300685 | Unclustered | 1,46E-05 | 49,75 | YES1 |
| ZEB2 | 0 | 1 | -1,182293567 | Unclustered | 0,003538 | 49,75 | ZEB2 |
| ZIC5 | 0 | 1 | -1,547463886 | Unclustered | 0,008452 | 49,75 | ZIC5 |
| ZNF680 | 0 | 1 | -1,0699721 | Unclustered | 0,009153 | 49,75 | ZNF680 |

Supplementary file 8: Co-expression network data of upregulated genes; the top 10 most ranked hub genes are highlighted.

| name | betweenness | degree | log2FoldChange | padj | Rank_stat | shared name |
|---|---|---|---|---|---|---|
| CALY | 308,5256301 | 39 | 1,485084433 | 0,002053 | 94 | CALY |
| SOHLH1 | 401,6324883 | 27 | 1,335836086 | 0,015748 | 87,75 | SOHLH1 |
| CPNE9 | 138,3551033 | 34 | 1,557307378 | 0,003438 | 87,5 | CPNE9 |
| NRGN | 116,6896206 | 33 | 1,338233206 | 0,00963 | 85,75 | NRGN |
| MAST1 | 148,6512848 | 28 | 1,027618848 | 0,016262 | 84,25 | MAST1 |
| GRIK1 | 143,5842487 | 28 | 1,161673399 | 0,001677 | 83,75 | GRIK1 |
| SYT5 | 99,10299657 | 33 | 1,259139581 | 0,017918 | 82,75 | SYT5 |
| TRPM2 | 223,710142 | 21 | 1,050754419 | 5,39E-04 | 81,25 | TRPM2 |
| BX276092.9 | 575,6169904 | 15 | 2,279984492 | 0,003222 | 79,5 | BX276092.9 |
| MYO15A | 219,0596195 | 20 | 1,199815203 | 0,002313 | 79,5 | MYO15A |
| PGM2L1 | 138,6646908 | 25 | 1,145629356 | 0,007447 | 79,5 | PGM2L1 |
| PNMA3 | 99,82262735 | 27 | 1,491351478 | 0,007421 | 77,75 | PNMA3 |
| AP3B2 | 85,2207987 | 31 | 1,053030273 | 0,036148 | 77,25 | AP3B2 |
| R3HDM1 | 86,29392858 | 30 | 1,036988626 | 0,036319 | 77 | R3HDM1 |
| PNCK | 80,29650161 | 32 | 1,059414629 | 0,003123 | 76,5 | PNCK |
| ZFR2 | 796,6497242 | 11 | 1,098006982 | 0,002862 | 76 | ZFR2 |
| CACNA1G | 132,0097555 | 21 | 1,098563359 | 0,039941 | 75,75 | CACNA1G |
| LPCAT4 | 81,85478506 | 29 | 1,061878554 | 0,029616 | 75 | LPCAT4 |
| RPRML | 72,05616844 | 31 | 1,44685866 | 0,017772 | 74,75 | RPRML |
| RASL10A | 183,4413388 | 14 | 1,010962143 | 0,005778 | 74,5 | RASL10A |
| CHTF18 | 738,822918 | 10 | 1,330956163 | 3,50E-04 | 74 | CHTF18 |
| MPV17L2 | 108,9928756 | 18 | 1,324605327 | 4,64E-04 | 72,25 | MPV17L2 |
| CAMKV | 63,44972888 | 29 | 1,024340961 | 0,023381 | 72 | CAMKV |
| CACNG8 | 138,9491715 | 14 | 1,065642072 | 0,004104 | 71,5 | CACNG8 |
| CCK | 57,10083047 | 29 | 1,301183717 | 0,045241 | 71 | CCK |
| TMEM249 | 405,2991378 | 8 | 1,638774655 | 0,001195 | 69,75 | TMEM249 |
| CORO6 | 432,4064197 | 7 | 1,362714826 | 0,01285 | 69 | CORO6 |
| SERPINF1 | 90,14882997 | 18 | 1,097374598 | 0,049892 | 68,25 | SERPINF1 |
| AHRR | 102,835614 | 12 | 1,307663688 | 0,023569 | 66,25 | AHRR |
| RNF112 | 35,95998436 | 26 | 1,163252396 | 0,002901 | 64,75 | RNF112 |
| PNMA5 | 35,01907769 | 26 | 1,41957014 | 0,017417 | 64,25 | PNMA5 |
| NOXA1 | 241,5728936 | 5 | 1,246952995 | 0,006364 | 64 | NOXA1 |
| BICDL1 | 80,63958738 | 17 | 1,184361037 | 0,022312 | 63,5 | BICDL1 |
| SYCE1 | 161,8129687 | 6 | 1,526958221 | 0,007301 | 63,5 | SYCE1 |
| CRHR1 | 89,05101286 | 12 | 1,021152651 | 0,030254 | 62,75 | CRHR1 |
| GABRD | 30,62461267 | 25 | 1,191681998 | 0,019507 | 62,5 | GABRD |
| SCN3B | 50,83633913 | 20 | 1,055496609 | 0,0191 | 62 | SCN3B |
| DRP2 | 26,09263102 | 25 | 1,067915409 | 0,018017 | 61,5 | DRP2 |
| PDIA2 | 154,9566368 | 5 | 1,577014259 | 0,005638 | 61 | PDIA2 |
| ABCC12 | 27,51008342 | 22 | 1,428346639 | 0,038467 | 60,25 | ABCC12 |
| CYP2E1 | 98,60957217 | 9 | 1,512524292 | 0,011809 | 59,75 | CYP2E1 |
| CCDC57 | 176 | 4 | 1,043935189 | 0,017168 | 59,5 | CCDC57 |
| PSMG3 | 25,48173032 | 22 | 1,004438853 | 0,003558 | 59,25 | PSMG3 |
| MICAL1 | 108,183074 | 5 | 1,299282584 | 0,003522 | 56,5 | MICAL1 |
| ACTL6B | 5,745156744 | 23 | 1,11873798 | 0,031653 | 55,5 | ACTL6B |
| PRMT8 | 16,21344752 | 20 | 1,017080216 | 0,016328 | 55,5 | PRMT8 |
| RTP5 | 22,4503443 | 17 | 1,087586804 | 0,006858 | 54 | RTP5 |
| CYP2D7 | 100,3638109 | 4 | 1,076935792 | 0,035293 | 53 | CYP2D7 |
| TCTE1 | 12,73940792 | 17 | 1,009968861 | 0,021259 | 52 | TCTE1 |
| GRIK2 | 48,98861613 | 8 | 1,374419396 | 6,82E-04 | 49,25 | GRIK2 |
| NECAB2 | 11,45930804 | 14 | 1,002687653 | 0,002134 | 49 | NECAB2 |
| TAC3 | 15,6929317 | 12 | 1,575138923 | 0,023231 | 48,75 | TAC3 |
| B4GALNT3 | 91,88484848 | 3 | 1,16564294 | 0,01214 | 48,5 | B4GALNT3 |
| TNNT2 | 59,80581362 | 6 | 1,335690278 | 0,00385 | 48,5 | TNNT2 |
| GRM2 | 20,95401413 | 10 | 1,188375678 | 0,017403 | 46,5 | GRM2 |

| | | | | | | |
|---|---|---|---|---|---|---|
| DLK2 | 6,050188894 | 11 | 1,118565201 | 0,006803 | 45 | DLK2 |
| ANKRD24 | 64,58403144 | 4 | 1,005194527 | 0,00131 | 45 | ANKRD24 |
| WDR97 | 37,61640733 | 5 | 1,407594933 | 0,00246 | 44 | WDR97 |
| P2RX2 | 89 | 2 | 3,010706157 | 4,85E-04 | 43,25 | P2RX2 |
| ZGLP1 | 89 | 2 | 1,119569712 | 0,025358 | 43,25 | ZGLP1 |
| TBC1D26 | 48,24647596 | 4 | 2,092324699 | 0,003222 | 42 | TBC1D26 |
| CALB2 | 2,135192582 | 11 | 1,124701501 | 0,004687 | 41,5 | CALB2 |
| CORT | 1,26565106 | 12 | 1,478951637 | 0,001321 | 41,25 | CORT |
| KRT17 | 3,390950962 | 10 | 1,308873708 | 0,019161 | 41 | KRT17 |
| PKD1 | 73,9 | 2 | 1,054550917 | 0,001195 | 40 | PKD1 |
| NEGR1 | 4,533359536 | 8 | 1,120314104 | 0,011588 | 39,25 | NEGR1 |
| BEX5 | 1,326261208 | 9 | 1,11339938 | 0,019043 | 37,25 | BEX5 |
| PLPPR3 | 1,408333333 | 8 | 1,144085695 | 0,010709 | 36,25 | PLPPR3 |
| LCN15 | 36,1273296 | 2 | 1,598961893 | 0,00634 | 35 | LCN15 |
| ASTL | 1,541111068 | 6 | 1,370631764 | 0,0102 | 34,5 | ASTL |
| SEMA5B | 3,878491017 | 3 | 1,219991267 | 0,006096 | 29,5 | SEMA5B |
| TGM1 | 12,5 | 2 | 1,5806132 | 0,008343 | 29 | TGM1 |
| TRIM54 | 2,884848485 | 3 | 1,280311696 | 0,008622 | 28,5 | TRIM54 |
| ATP4A | 0,2 | 4 | 2,393310718 | 0,001975 | 27 | ATP4A |
| RTEL1 | 5,333333333 | 2 | 1,115674959 | 0,001195 | 27 | RTEL1 |
| PKD2L1 | 0,698981808 | 3 | 1,322598614 | 0,009756 | 25,5 | PKD2L1 |
| CITED1 | 0 | 5 | 1,031057586 | 0,046929 | 24,25 | CITED1 |
| TRIM17 | 0,487179487 | 2 | 1,050234131 | 0,010564 | 21,5 | TRIM17 |
| PLCH2 | 0 | 3 | 1,08545462 | 0,021641 | 19,25 | PLCH2 |
| CES4A | 0 | 2 | 1,021312646 | 0,017312 | 15,75 | CES4A |
| NOXO1 | 0 | 2 | 1,38734254 | 5,34E-04 | 15,75 | NOXO1 |
| CYP2D6 | 0 | 1 | 1,156042333 | 0,045242 | 9,5 | CYP2D6 |
| CARTPT | 0 | 1 | 1,81980008 | 0,019466 | 9,5 | CARTPT |
| LORICRIN | 0 | 1 | 1,803926378 | 0,008381 | 9,5 | LORICRIN |
| KIF12 | 0 | 1 | 1,26860296 | 0,00115 | 9,5 | KIF12 |
| LTK | 0 | 1 | 1,492943739 | 7,41E-04 | 9,5 | LTK |
| LY6H | 0 | 1 | 1,07617123 | 0,001993 | 9,5 | LY6H |
| PTCH2 | 0 | 1 | 1,053060627 | 0,005349 | 9,5 | PTCH2 |
| PROC | 0 | 1 | 1,97239753 | 0,01961 | 9,5 | PROC |
| SLC44A5 | 0 | 1 | 1,064892142 | 0,030153 | 9,5 | SLC44A5 |
| RTL1 | 0 | 1 | 1,507033312 | 0,027174 | 9,5 | RTL1 |
| TARBP1 | 0 | 1 | 1,306970395 | 0,011976 | 9,5 | TARBP1 |
| TGFBR3L | 0 | 1 | 1,051353191 | 0,002557 | 9,5 | TGFBR3L |
| AGAP6 | 0 | 1 | 1,042080575 | 0,01563 | 9,5 | AGAP6 |
| TIMM23B-AGA | 0 | 1 | 1,056709194 | 0,013788 | 9,5 | TIMM23B-AGAP6 |
| TNFRSF25 | 0 | 1 | 1,581402716 | 0,006636 | 9,5 | TNFRSF25 |
| DNASE1L2 | 0 | 1 | 1,199218077 | 0,005496 | 9,5 | DNASE1L2 |

Supplementary file 8: Co-expression network clusters of downregulated genes.

| __glayCluster | betweenness | degree | log2FoldChange | name | padj | Rank_stat | shared name |
|---|---|---|---|---|---|---|---|
| 1 | 7004.40011615783 | 116 | -1.50325435758696 | SORBS3 | 5.7662016: | 707 | SORBS3 |
| | 9088.38586509955 | 88 | -1.50087243153436 | SZRD1 | 4.0548499: | 699,75 | SZRD1 |
| | 5131.96482561097 | 85 | -2.31246675377311 | AHNAK | 1.4105385( | 694,25 | AHNAK |
| | 3580.2490596035 | 108 | -2.16618155725699 | GRIN2C | 2.1951423 | 692,5 | GRIN2C |
| | 4250.19328383344 | 89 | -1.61032172729005 | MAPKAPK2 | 4.0548499: | 691,5 | MAPKAPK2 |
| | 3455.79080185914 | 105 | -2.86611141396491 | IRAG1 | 8.9608589! | 690,25 | IRAG1 |
| | 3364.01978764192 | 97 | -2.3821552012305 | FGFRL1 | 8.1389262! | 687 | FGFRL1 |
| | 3539.38514540332 | 94 | -1.0470879431495 | SLC44A2 | 0.0049569! | 686,75 | SLC44A2 |
| | 6048.81689166699 | 71 | -1.436052384764 | MCL1 | 1.9124371 | 686,5 | MCL1 |
| | 4038.6100657267 | 80 | -1.61196792079135 | DDR2 | 6.6123329! | 684,75 | DDR2 |
| | 4865.75591838579 | 69 | -1.30623137302227 | PSKH1 | 6.3423773! | 683 | PSKH1 |
| | 3091.49784940503 | 78 | -1.68013728710646 | CDC42EP4 | 5.9041835: | 674 | CDC42EP4 |
| | 3741.08949218619 | 66 | -1.30926650182874 | GNG5 | 4.0195288( | 672 | GNG5 |
| | 3924.20661457381 | 62 | -2.08237548823861 | GNA12 | 1.4053328: | 669,75 | GNA12 |
| | 2496.540400948 | 94 | -1.88594923771028 | SNX33 | 4.9384899( | 667,25 | SNX33 |
| | 2704.10819228082 | 77 | -1.3946790752735 | ADCYAP1R | 6.2351611 | 662,5 | ADCYAP1R1 |
| | 2706.59653673612 | 73 | -1.97231308503933 | MLC1 | 4.8245536: | 660,25 | MLC1 |
| | 5896.51029186093 | 48 | -3.41358478854833 | GPR4 | 3.8460093! | 659,75 | GPR4 |
| | 3749.5004866938 | 52 | -1.62568909009584 | AMOT | 3.3634485! | 658 | AMOT |
| | 2124.36506099125 | 85 | -2.19780258935763 | STON2 | 1.5878005! | 654,25 | STON2 |
| | 3268.84800629492 | 52 | -2.05556209920081 | LRP10 | 1.9124371 | 653 | LRP10 |
| | 1996.89968582971 | 80 | -1.80114928020846 | CABLES1 | 7.4060703( | 649,25 | CABLES1 |
| | 3108.66644315202 | 50 | -1.35715809956401 | ANTXR1 | 1.1322525! | 647,5 | ANTXR1 |
| | 3354.21059312952 | 47 | -1.27825986427286 | ECE1 | 0.0015999( | 644,75 | ECE1 |
| | 3177.38621420098 | 47 | -2.17504490068685 | PALLD | 3.1077606! | 642,75 | PALLD |
| | 2812.47236844943 | 48 | -2.79865252353504 | FAM107A | 2.3291787! | 637,75 | FAM107A |
| | 2194.72844109268 | 52 | -1.28648727667251 | NHSL1 | 0.0034998 | 630 | NHSL1 |
| | 1534.17372088929 | 70 | -2.39644695289864 | SMO | 1.9100529! | 629,5 | SMO |
| | 2860.82747179975 | 42 | -1.01045643545126 | CASKIN2 | 0.0046545( | 629 | CASKIN2 |
| | 1659.99792727696 | 62 | -2.40306438042635 | ITPKB | 1.1199223! | 625,75 | ITPKB |
| | 1482.38644630569 | 68 | -2.12476944930361 | PLEKHO2 | 2.1951423 | 625,5 | PLEKHO2 |
| | 1620.29177390687 | 62 | -1.80808631354268 | PBXIP1 | 9.2702773 | 624,75 | PBXIP1 |
| | 1264.82966790757 | 81 | -1.21401307970234 | SMOX | 0.0041914: | 619,75 | SMOX |
| | 2416.56790062556 | 43 | -1.43360870967626 | MSN | 1.4053328: | 618,75 | MSN |
| | 1369.21688616233 | 68 | -1.70352683439333 | MAPKAPK3 | 0.0014805! | 616,5 | MAPKAPK3 |
| | 3188.55022347471 | 31 | -1.66319124769708 | MOB3C | 5.1486767 | 613,75 | MOB3C |
| | 2812.78500275645 | 34 | -1.37944813330231 | GPR137B | 8.6263711! | 613,5 | GPR137B |
| | 1403.67433030724 | 57 | -1.61274537141908 | EZR | 0.0026919: | 610,5 | EZR |
| | 2488.93251268519 | 35 | -2.4285886051129 | TNS3 | 1.4053328: | 605,25 | TNS3 |
| | 2552.39528459303 | 33 | -2.46676842653017 | HELZ2 | 2.3343003 | 603,5 | HELZ2 |
| | 2039.05942833494 | 40 | -1.38462434742232 | NES | 4.7112953! | 603,5 | NES |
| | 4054.30060984098 | 24 | -1.38742622003794 | FYCO1 | 4.4201532: | 603,25 | FYCO1 |
| | 1729.3570844544 | 41 | -1.40954527859516 | LRRC8A | 0.0019003 | 598,75 | LRRC8A |
| | 1258.43250995327 | 56 | -2.08715402502349 | SCARA3 | 1.7870641! | 597,5 | SCARA3 |
| | 2215.15509913807 | 32 | -1.85718650787612 | ARHGAP19 | 1.6536729 | 595,25 | ARHGAP19 |
| | 2502.26622962378 | 30 | -1.25062545024913 | PTTG1IP | 7.2815330 | 595,25 | PTTG1IP |
| | 1135.67851814079 | 52 | -1.42284984417717 | RIN2 | 9.4330053( | 589 | RIN2 |
| | 1061.96846196322 | 56 | -2.27421388284922 | CGNL1 | 1.5873235 | 588,5 | CGNL1 |
| | 932.956885729062 | 58 | -1.15261481556733 | TMEM164 | 0.0019527 | 584,25 | TMEM164 |
| | 1007.50608909542 | 53 | -1.17226681672382 | MID1 | 0.0047736: | 582,75 | MID1 |
| | 2796.69599009667 | 22 | -1.46590192447041 | TGFB2 | 0.0014410 | 573,75 | TGFB2 |
| | 1062.29153561091 | 39 | -1.3686877522608 | ACSS1 | 0.0070184 | 561,5 | ACSS1 |
| | 1955.58461854116 | 23 | -1.68915943337733 | RHPN2 | 0.0015711( | 561,25 | RHPN2 |
| | 809.537422053625 | 48 | -1.81364498141472 | GPAM | 0.0011586( | 559,25 | GPAM |
| | 1049.18274655926 | 38 | -1.11393280324046 | APPL2 | 0.0024909( | 558,5 | APPL2 |
| | 862.96810953689 | 42 | -1.64199533358167 | OAF | 0.0020528: | 554 | OAF |
| | 1263.86767400192 | 29 | -1.71181179445338 | SIX5 | 0.0041041! | 550,5 | SIX5 |
| | 931.674081946478 | 37 | -1.72232562783873 | ADGRG1 | 9.9876776( | 550,25 | ADGRG1 |
| | 1414.96536778976 | 24 | -1.39426307202038 | GABRG1 | 0.0035375: | 547,75 | GABRG1 |
| | 787.845155527436 | 42 | -2.00306595165672 | TNFSF13 | 3.8881687! | 547,5 | TNFSF13 |
| | 715.075795031921 | 44 | -1.86823808291118 | RFX4 | 5.1486767 | 540,5 | RFX4 |
| | 839.318642222613 | 35 | -1.2035252450988 | KANK1 | 0.0015071 | 539,75 | KANK1 |
| | 1082.06594279229 | 27 | -1.50968505638063 | NACC2 | 1.7291800! | 538 | NACC2 |

| | | | | | | |
|---|---|---|---|---|---|---|
| 821.065054850119 | 35 | -1.4435865770089 | EYA2 | 0.0039724 | 537,75 | EYA2 |
| 1233.73140379676 | 24 | -1.20280613733767 | CHST14 | 0.0071216 | 535,25 | CHST14 |
| 697.062320767978 | 43 | -1.23257912930356 | EIF4EBP2 | 0.0016030 | 535,25 | EIF4EBP2 |
| 900.471004713386 | 27 | -1.41562404213888 | RAB34 | 0.0082096 | 526,5 | RAB34 |
| 665.473062186598 | 45 | -1.25917442453994 | TEAD1 | 0.0026340 | 526 | TEAD1 |
| 1582.82926867628 | 18 | -1.23161024643305 | CDKN2C | 0.0011717 | 523 | CDKN2C |
| 1431.90678169143 | 19 | -1.55989863848918 | NOTCH2NL | 0.0024723 | 522,25 | NOTCH2NLC |
| 616.25472537813 | 44 | -1.07287967994792 | MAOA | 0.0045670 | 519 | MAOA |
| 1226.48038416724 | 21 | -1.51804319996441 | ABHD4 | 7.8792796 | 516,5 | ABHD4 |
| 753.785924906943 | 28 | -2.10620832749647 | SOD3 | 0.0014166 | 514,75 | SOD3 |
| 1027.48910551192 | 22 | -2.13606454583144 | TMBIM1 | 2.9092641 | 512,75 | TMBIM1 |
| 511.887600365921 | 49 | -1.31148809207292 | EMILIN1 | 0.0083411 | 512,25 | EMILIN1 |
| 715.735955827579 | 29 | -1.43343269625626 | PXDC1 | 0.0011376 | 511 | PXDC1 |
| 512.028068508266 | 45 | -1.60558785491796 | TPD52L1 | 0.0011807 | 508 | TPD52L1 |
| 1226.59604894421 | 19 | -1.78852818636538 | MYORG | 6.6529865 | 507,25 | MYORG |
| 1394.58481797862 | 17 | -1.17349752575431 | APCDD1 | 0.0071243 | 507 | APCDD1 |
| 450.814810227649 | 49 | -1.20266626397812 | GRAMD2B | 0.0036453 | 506,75 | GRAMD2B |
| 476.878877701777 | 45 | -1.67754558620436 | SLC9A3R1 | 0.0013237 | 504 | SLC9A3R1 |
| 1745.34335509261 | 14 | -1.29759672096682 | SLC39A1 | 9.3338746 | 498,75 | SLC39A1 |
| 665.101359863067 | 31 | -2.02594794275979 | MRO | 3.8460093 | 497,75 | MRO |
| 1134.95558353021 | 18 | -1.43075671985785 | ITGB5 | 9.0170879 | 497,5 | ITGB5 |
| 461.638636760952 | 42 | -1.89876188920569 | ALDH4A1 | 9.4032749 | 496,5 | ALDH4A1 |
| 820.490856417145 | 20 | -1.42867790975904 | SLC6A11 | 0.0020742 | 489 | SLC6A11 |
| 480.018944299202 | 32 | -1.11592851118721 | HSDL2 | 0.0012506 | 480,25 | HSDL2 |
| 657.469321961273 | 24 | -1.43561734670711 | FAM167A | 3.7411225 | 478,25 | FAM167A |
| 482.96637070275 | 31 | -1.62871838711814 | TNFSF12-TI | 5.0940605 | 478,25 | TNFSF12-TNFSF13 |
| 514.681610256362 | 30 | -1.39419595070199 | ADD3 | 8.6836999 | 477,75 | ADD3 |
| 526.515281139702 | 28 | -1.25137063768245 | TAX1BP3 | 0.0028576 | 474,75 | TAX1BP3 |
| 333.201021579503 | 37 | -1.73313534324767 | SLC25A18 | 0.0019876 | 473,75 | SLC25A18 |
| 252.186973238649 | 43 | -1.57313015440511 | SULT1C4 | 4.0548499 | 471,75 | SULT1C4 |
| 298.477461300978 | 36 | -1.49143450269391 | NEK6 | 0.0016350 | 464,25 | NEK6 |
| 606.14253918105 | 22 | -1.02667724500593 | NTRK2 | 0.0074315 | 461,25 | NTRK2 |
| 875.487495813113 | 15 | -1.46491723079134 | TP53BP2 | 0.0018729 | 461 | TP53BP2 |
| 329.655934916115 | 31 | -2.94710024866011 | P2RY2 | 8.1389262 | 460,75 | P2RY2 |
| 526.323305181331 | 23 | -1.27812950224032 | HEPACAM | 0.0055219 | 457,25 | HEPACAM |
| 488.902536119115 | 23 | -1.26099477322261 | LAMB2 | 0.0088090 | 454,75 | LAMB2 |
| 1097.90762837803 | 12 | -2.09353900946569 | CDK2 | 7.7208845 | 454,5 | CDK2 |
| 559.73135884128 | 22 | -1.33063834882169 | HEPN1 | 0.0067975 | 454,25 | HEPN1 |
| 377.86797132988 | 24 | -1.25367844708301 | AHCYL1 | 0.0053493 | 448,25 | AHCYL1 |
| 252.405540941063 | 31 | -1.77350638429846 | BMPR1B | 0.0021509 | 448,25 | BMPR1B |
| 205.556838364865 | 35 | -1.43826084478538 | FADS2 | 5.0333876 | 447,75 | FADS2 |
| 187.548767296189 | 36 | -1.78006041941187 | GPR146 | 6.6613268 | 446,75 | GPR146 |
| 325.53192936579 | 26 | -1.19762986484132 | RXRA | 0.0041914 | 446,75 | RXRA |
| 404.791137422195 | 23 | -1.24927109339614 | SASH1 | 7.0741293 | 446,75 | SASH1 |
| 1503.74653795724 | 9 | -1.29619396597404 | PPP1R18 | 0.0017679 | 446,25 | PPP1R18 |
| 215.710962827392 | 32 | -1.48220200917166 | PLCD3 | 7.3854909 | 444,25 | PLCD3 |
| 952.854015571373 | 11 | -1.38197039862551 | TNIP2 | 0.0015530 | 438,5 | TNIP2 |
| 354.690860198963 | 21 | -1.21771828281517 | ABHD15 | 0.0074214 | 428,5 | ABHD15 |
| 224.624295652235 | 25 | -1.48130472019615 | DOK1 | 0.0053493 | 427,75 | DOK1 |
| 222.790670391897 | 25 | -1.18439070628434 | RMDN1 | 7.8263073 | 427,25 | RMDN1 |
| 198.59882585882 | 25 | -1.59700202293838 | NDRG2 | 1.8839658 | 423,75 | NDRG2 |
| 391.810869663479 | 19 | -1.31019425879179 | PIPOX | 0.0054019 | 422,25 | PIPOX |
| 276.952155862933 | 22 | -1.19383025167179 | ID4 | 0.0075013 | 420,75 | ID4 |
| 1419.4825715823 | 7 | -1.77437952443608 | CSF1 | 4.0328621 | 420 | CSF1 |
| 366.579632325713 | 19 | -1.65807921688154 | AXL | 0.0027317 | 419,75 | AXL |
| 268.799597441233 | 21 | -1.29628453872472 | TLR4 | 0.0057565 | 414,5 | TLR4 |
| 234.277898157644 | 22 | -1.28177974137987 | IL17RD | 4.0548499 | 413,75 | IL17RD |
| 231.707901004105 | 22 | -1.11885266595513 | C2orf72 | 0.0029488 | 413,25 | C2orf72 |
| 715.273333878012 | 11 | -1.17947457753827 | FSTL1 | 4.6477145 | 413 | FSTL1 |
| 269.080106458693 | 20 | -1.34767336824999 | BMP7 | 0.0015831 | 411 | BMP7 |
| 553.469769343557 | 15 | -1.34588288255823 | RHOBTB3 | 5.8425105 | 411 | RHOBTB3 |
| 127.232435104955 | 25 | -1.53898057773567 | NOTCH2NL | 0.0044981 | 410,75 | NOTCH2NLA |
| 84.1818267665275 | 29 | -1.38977924725095 | GNG12 | 6.4146788 | 410 | GNG12 |
| 162.644295430558 | 23 | -1.09278509921349 | IL17D | 2.4351527 | 406,25 | IL17D |
| 1226.61452103652 | 7 | -1.2246897291207 | CNN3 | 0.0016350 | 406 | CNN3 |
| 453.735639033539 | 15 | -1.20140060838761 | CYBRD1 | 0.0067975 | 401 | CYBRD1 |

| | | | | | | |
|---|---|---|---|---|---|---|
| 323.979855324177 | 17 | -1.52316011716425 | SPATA21 | 5.33895826 | 401 | SPATA21 |
| 577.793367971039 | 12 | -1.4119357763371 | PON2 | 0.0038224 | 395 | PON2 |
| 731.824851430521 | 9 | -1.44092202085778 | GLIS2 | 7.26344922 | 392,75 | GLIS2 |
| 695.32165639705 | 9 | -1.77617290010222 | GPR143 | 0.0013871 | 384,75 | GPR143 |
| 193.034553194903 | 17 | -1.64869710220548 | FGFR3 | 0.0040873 | 380,5 | FGFR3 |
| 700.693814975116 | 8 | -1.75778975105229 | STK17B | 0.0067245 | 374,25 | STK17B |
| 107.543826986855 | 18 | -1.46602439030714 | GPR37L1 | 0.0042749 | 369,5 | GPR37L1 |
| 221.696024150864 | 15 | -1.38224748201251 | PDGFRB | 0.0015395 | 368,5 | PDGFRB |
| 302.121364986382 | 13 | -1.56844347763082 | EPHX1 | 0.0032576 | 368,25 | EPHX1 |
| 65.0954635799151 | 19 | -2.0623777272964 | PPP1R3C | 6.9633907 | 363,75 | PPP1R3C |
| 190.412821877658 | 15 | -1.65783826265923 | NOTCH2 | 0.0029617 | 363,5 | NOTCH2 |
| 50.0686456375793 | 20 | -2.31389816459308 | NBPF10 | 3.7411225 | 361,5 | NBPF10 |
| 690.399073602552 | 7 | -1.088806329035 | C3orf70 | 5.3462928 | 361 | C3orf70 |
| 564.47367360213 | 9 | -2.02430017706791 | PLIN4 | 0.0028576 | 360,75 | PLIN4 |
| 667.739964451618 | 8 | -1.03718672927482 | ERBB2 | 0.0034441 | 360,25 | ERBB2 |
| 736.239548702692 | 6 | -1.50809941048292 | S1PR1 | 0.0021350 | 359,75 | S1PR1 |
| 169.453529690725 | 15 | -1.01785113697665 | RBM38 | 0.0070240 | 358,5 | RBM38 |
| 714.563557148094 | 6 | -1.21702973153375 | DDR1 | 0.0076980 | 354,75 | DDR1 |
| 697.567634334379 | 6 | -1.4931125995799 | ZIC2 | 0.0050910 | 352,25 | ZIC2 |
| 686.997774172639 | 6 | -1.45899058421892 | ZHX2 | 3.1975007 | 347,75 | ZHX2 |
| 266.148773704758 | 11 | -1.21438375729382 | ACAA2 | 9.2289517 | 347,5 | ACAA2 |
| 36.4032034377219 | 18 | -1.64005135693768 | AKR1C3 | 0.0037506 | 340,5 | AKR1C3 |
| 549.780487983796 | 7 | -1.29605598970903 | AL603832.3 | 3.5024771 | 336,5 | AL603832.3 |
| 96.3531383400149 | 14 | -2.16278309122397 | GLI3 | 9.2702773 | 335,75 | GLI3 |
| 687.41455361499 | 5 | -1.33048809178409 | INHBB | 0.0016678 | 334,25 | INHBB |
| 64.5867176313361 | 14 | -1.82202008388391 | SEMA3F | 0.0013770 | 327,25 | SEMA3F |
| 461.980181324065 | 6 | -1.05285110756084 | DCHS1 | 0.0064632 | 316,25 | DCHS1 |
| 132.724170896463 | 10 | -1.28280263815145 | SUCLG2 | 9.1605441 | 315 | SUCLG2 |
| 72.5217177800247 | 11 | -1.51016451226999 | GLUL | 0.0052940 | 308,5 | GLUL |
| 234.836596141776 | 8 | -1.30071496607375 | CACHD1 | 0.0010494 | 306,75 | CACHD1 |
| 298.046811079494 | 7 | -1.02514443030961 | CTDSP2 | 0.0020679 | 306 | CTDSP2 |
| 133.507117142521 | 9 | -1.43792942246562 | PARP14 | 0.0012155 | 302,25 | PARP14 |
| 686.0 | 3 | -1.16848441856197 | MINDY1 | 0.0056272 | 302 | MINDY1 |
| 34.123480350389 | 12 | -1.50377025781333 | HES6 | 0.0082939 | 296,5 | HES6 |
| 24.3535834895412 | 13 | -1.5492044703017 | GAREM2 | 0.0070184 | 294,25 | GAREM2 |
| 60.1592472225101 | 10 | -1.50236899734047 | PAQR8 | 0.0017928 | 294 | PAQR8 |
| 16.5982490075773 | 14 | -1.36393740812355 | SEC14L2 | 0.0049083 | 291,25 | SEC14L2 |
| 118.310937891831 | 8 | -1.59064336180612 | BBOX1 | 5.4259198 | 286,25 | BBOX1 |
| 63.9932762456167 | 9 | -1.07392260613988 | SWAP70 | 0.0017594 | 282,75 | SWAP70 |
| 104.263724644971 | 8 | -1.55948127431826 | GPRC5C | 0.0025365 | 282,25 | GPRC5C |
| 59.5112110317588 | 9 | -1.1414890434092 | DAG1 | 0.0040022 | 280,25 | DAG1 |
| 32.9735536514314 | 10 | -1.57097879686584 | MGST1 | 0.0020166 | 277 | MGST1 |
| 52.3942143302894 | 9 | -1.74857959609833 | GJA1 | 0.0048421 | 276,75 | GJA1 |
| 257.152466514805 | 5 | -1.38173555949399 | ELK3 | 2.9950698 | 275,75 | ELK3 |
| 22.6869894528809 | 10 | -1.18303814521483 | RASSF8 | 2.9092641 | 267,5 | RASSF8 |
| 124.217485687343 | 6 | -2.09852863368516 | ALDH1L1 | 0.0018019 | 265,75 | ALDH1L1 |
| 185.445009290433 | 5 | -1.24465464608421 | CSRP1 | 0.0052733 | 262,25 | CSRP1 |
| 13.0961492764123 | 10 | -1.09418962232307 | PITPNC1 | 0.0034104 | 258 | PITPNC1 |
| 166.820924984355 | 5 | -1.76483653923225 | PHYHD1 | 0.0066247 | 257,25 | PHYHD1 |
| 230.119172637804 | 3 | -1.30628052288618 | MAML2 | 0.0035223 | 243,25 | MAML2 |
| 47.4188439539503 | 6 | -1.43913131624472 | STOM | 2.7390148 | 240,25 | STOM |
| 9.90335219073917 | 9 | -1.84257355683505 | RGS20 | 0.0010494 | 239,75 | RGS20 |
| 45.151692277186 | 6 | -1.02044996234808 | EPS8 | 0.0074214 | 237,75 | EPS8 |
| 6.19882915573814 | 9 | -1.87853616299911 | TGIF1 | 1.9124371 | 232,75 | TGIF1 |
| 5.39758313267679 | 9 | -1.40804085211432 | TBC1D16 | 9.9876776 | 230,75 | TBC1D16 |
| 19.8011683816689 | 7 | -3.47723170252809 | C10orf105 | 3.8460093 | 229,5 | C10orf105 |
| 9.24590196194566 | 8 | -1.36071753553064 | NPL | 0.0033213 | 225,25 | NPL |
| 38.5342019351214 | 5 | -1.37936826200059 | TAP1 | 0.0037088 | 220,25 | TAP1 |
| 19.9468132671355 | 6 | -1.4410322655834 | MTSS2 | 2.2479664 | 218,25 | MTSS2 |
| 26.2498311826309 | 5 | -1.36919351589913 | ATP1A2 | 0.0086599 | 210,75 | ATP1A2 |
| 5.95102366631136 | 7 | -1.07228792537979 | WWC1 | 7.1070945 | 210 | WWC1 |
| 40.455580375792 | 4 | -1.97531617330847 | NQO1 | 0.0011502 | 208,5 | NQO1 |
| 10.0576586902026 | 6 | -1.5788940439182 | PPP1R3G | 0.0088615 | 206,25 | PPP1R3G |
| 44.4156515889801 | 3 | -1.65191203849137 | LRP4 | 8.7954351 | 195,25 | LRP4 |
| 23.2841492320583 | 4 | -1.5109860450731 | SLC7A5 | 3.0245620 | 195 | SLC7A5 |
| 12.2379702192029 | 5 | -1.16378113444695 | KREMEN1 | 9.6065794 | 194,75 | KREMEN1 |

| | | | | | | | |
|---|---|---|---|---|---|---|---|
| | 10.1941933205956 | 5 | -1.06077934504554 | ZNF853 | 5.0774850 | 192,75 | ZNF853 |
| | 1.21213735632557 | 6 | -1.03762651918512 | CD81 | 0.0091530 | 185,75 | CD81 |
| | 1.83804091881638 | 5 | -1.41458824979857 | SLC4A4 | 0.0057296 | 173,25 | SLC4A4 |
| | 16.7472179432704 | 3 | -1.10304388712142 | SORCS2 | 0.0049605 | 173,25 | SORCS2 |
| | 6.24572345657376 | 4 | -1.48439922048949 | FAM181A | 0.0043646 | 173 | FAM181A |
| | 4.83835836462034 | 4 | -1.63447490935747 | CCDC8 | 0.0083446 | 170 | CCDC8 |
| | 1.84960108266558 | 4 | -1.33857163518723 | CCDC9B | 0.0046186 | 161,5 | CCDC9B |
| | 6.62289337854365 | 3 | -1.45171343848521 | ITGA6 | 0.0011924 | 160,25 | ITGA6 |
| | 1.73294073609347 | 3 | -1.94316964301871 | SDC4 | 0.0016350 | 145,75 | SDC4 |
| | 1.12275459471538 | 3 | -1.08905308891937 | LRP5 | 0.0087475 | 143,75 | LRP5 |
| | 0.505046328123452 | 3 | -1.42722360068452 | PMP2 | 4.4201532 | 140,75 | PMP2 |
| | 9.24983109386472 | 2 | -1.46944858710577 | SLC40A1 | 5.4355191 | 139,75 | SLC40A1 |
| | 1.20688385688386 | 2 | -1.04270310350054 | AK4 | 0.0020304 | 120,25 | AK4 |
| | 0.5 | 2 | -1.25185149015969 | EPB41L5 | 0.0048593 | 116,25 | EPB41L5 |
| | 0.1 | 2 | -1.20638474379779 | LFNG | 0.0032524 | 114,25 | LFNG |
| | 0.0 | 3 | -1.42234747442956 | MASP1 | 0.0017649 | 108,5 | MASP1 |
| | 0.0 | 3 | -1.93610287196336 | SAMD9L | 9.4536784 | 108,5 | SAMD9L |
| | 0.0 | 2 | -1.89100832755208 | GDPD2 | 0.0015711 | 84,5 | GDPD2 |
| | 0.0 | 2 | -1.05490055437022 | ITGB1 | 5.4150102 | 84,5 | ITGB1 |
| | 0.0 | 2 | -1.63525198300406 | ITPRIP | 0.0020926 | 84,5 | ITPRIP |
| | 0.0 | 2 | -1.73246947831314 | LIMK2 | 2.1951423 | 84,5 | LIMK2 |
| | 0.0 | 2 | -1.18759101060646 | LYN | 0.0064620 | 84,5 | LYN |
| | 0.0 | 2 | -1.61525974337222 | PARP9 | 0.0018729 | 84,5 | PARP9 |
| | 0.0 | 2 | -1.80061222688355 | AL672142.1 | 0.0098308 | 84,5 | AL672142.1 |
| | 0.0 | 2 | -1.48650865823848 | PRSS35 | 0.0068882 | 84,5 | PRSS35 |
| | 0.0 | 1 | -1.04426902212779 | CD59 | 0.0044802 | 49,75 | CD59 |
| | 0.0 | 1 | -1.11680032656473 | CTXND1 | 9.7857658 | 49,75 | CTXND1 |
| | 0.0 | 1 | -1.78342350467469 | EPAS1 | 8.1389262 | 49,75 | EPAS1 |
| | 0.0 | 1 | -1.0299335608639 | ERLIN2 | 0.0067903 | 49,75 | ERLIN2 |
| | 0.0 | 1 | -1.16317860165668 | ARHGAP11 | 0.0018700 | 49,75 | ARHGAP11A |
| | 0.0 | 1 | -1.59688282441047 | FLT4 | 0.0048421 | 49,75 | FLT4 |
| | 0.0 | 1 | -1.19273577803333 | GNG11 | 1.2437414 | 49,75 | GNG11 |
| | 0.0 | 1 | -1.69195246061627 | ALDH3B1 | 0.0027021 | 49,75 | ALDH3B1 |
| | 0.0 | 1 | -1.11473640525705 | MFGE8 | 0.0073769 | 49,75 | MFGE8 |
| | 0.0 | 1 | -1.08645843071036 | APLN | 0.0085145 | 49,75 | APLN |
| | 0.0 | 1 | -1.05649848677197 | NEDD1 | 0.0031553 | 49,75 | NEDD1 |
| | 0.0 | 1 | -1.09708131996607 | PREX2 | 9.1605441 | 49,75 | PREX2 |
| | 0.0 | 1 | -1.87010803925481 | ADORA2B | 0.0029525 | 49,75 | ADORA2B |
| | 0.0 | 1 | -1.26839975385876 | FAT1 | 0.0083235 | 49,75 | FAT1 |
| | 0.0 | 1 | -1.96374558812779 | SDS | 0.0029617 | 49,75 | SDS |
| | 0.0 | 1 | -1.37569855666554 | ISG20 | 0.0017563 | 49,75 | ISG20 |
| | 0.0 | 1 | -1.69841237917578 | KCNJ16 | 0.0024044 | 49,75 | KCNJ16 |
| | 0.0 | 1 | -1.32623566458005 | SMAD6 | 0.0087134 | 49,75 | SMAD6 |
| | 0.0 | 1 | -1.13913991197711 | TGFB1 | 0.0060578 | 49,75 | TGFB1 |
| | 0.0 | 1 | -1.08628592233364 | GOLIM4 | 0.0031230 | 49,75 | GOLIM4 |
| | 0.0 | 1 | -1.45310845842759 | GRAMD1C | 0.0042749 | 49,75 | GRAMD1C |
| | 0.0 | 1 | -1.02719263419281 | TTC23 | 0.0040022 | 49,75 | TTC23 |
| | 0.0 | 1 | -1.54746388570014 | ZIC5 | 0.0084516 | 49,75 | ZIC5 |
| | 0.0 | 1 | -1.06997210026627 | ZNF680 | 0.0091530 | 49,75 | ZNF680 |
| 2 | 6409.49547319207 | 105 | -1.9259581657853 | RHOC | 6.5610360 | 704,25 | RHOC |
| | 4397.76169017423 | 114 | -1.64524274592556 | S100A16 | 6.6529865 | 700 | S100A16 |
| | 4704.28645092041 | 101 | -1.60150312721719 | TRIP6 | 5.7662016 | 699 | TRIP6 |
| | 3625.78935721865 | 102 | -1.85119389116736 | CHDH | 9.3859159 | 691,5 | CHDH |
| | 7408.59865845785 | 75 | -2.14609007755925 | FTL | 5.2176081 | 691 | FTL |
| | 3752.26290259575 | 96 | -1.67146246250672 | MMP28 | 0.0019234 | 690,25 | MMP28 |
| | 4152.9259786155 | 82 | -1.06458231649508 | UBTD1 | 0.0020024 | 687,75 | UBTD1 |
| | 4595.45984608255 | 73 | -2.57513456718036 | ATOH8 | 9.7074511 | 683,75 | ATOH8 |
| | 3341.2419714315 | 88 | -1.29502263363271 | STAT3 | 0.0024044 | 681,75 | STAT3 |
| | 5123.50241828684 | 63 | -2.07659602692966 | METRN | 5.9499969 | 678,5 | METRN |
| | 15081.4143033344 | 56 | -1.84492341586387 | WASF2 | 3.8364423 | 677 | WASF2 |
| | 2886.70000141679 | 90 | -1.67112846601115 | TTC38 | 0.0017624 | 676,5 | TTC38 |
| | 2751.16745282633 | 96 | -2.29578728003549 | ARHGEF26 | 2.3870210 | 673,25 | ARHGEF26 |
| | 2918.87048535771 | 81 | -2.18349657027976 | VAMP5 | 1.9832980 | 672,75 | VAMP5 |
| | 2553.31734739612 | 98 | -1.79735912347783 | SNTA1 | 9.2850312 | 671 | SNTA1 |
| | 2460.04957420359 | 99 | -1.54663505963214 | MSI2 | 5.7278660 | 668,5 | MSI2 |
| | 2777.48793662052 | 78 | -2.10045125731957 | YAP1 | 1.7955876 | 665,5 | YAP1 |

| | | | | | | |
|---|---|---|---|---|---|---|
| 2969.99105309959 | 68 | -1.07846759462088 | WBP1 | 0.0027637 | 665 | WBP1 |
| 2782.02929901286 | 62 | -1.32188711325386 | MAVS | 0.0050800 | 653,25 | MAVS |
| 1908.34084101253 | 91 | -1.26292475051986 | BCL2 | 0.0022197 | 653 | BCL2 |
| 3524.88608539564 | 50 | -1.44167092598017 | TNFRSF1A | 0.0019604 | 651,5 | TNFRSF1A |
| 2601.86934695861 | 61 | -1.49057951516286 | RRAS | 0.0010494 | 648,5 | RRAS |
| 2053.33922309439 | 76 | -1.51142376997642 | BCL6 | 6.9606381 | 647,25 | BCL6 |
| 2281.42622433148 | 59 | -1.09916407828511 | NECAP2 | 0.0025412 | 639 | NECAP2 |
| 2198.64605848726 | 60 | -2.23405067314945 | RAB13 | 1.9723266 | 638,75 | RAB13 |
| 1682.47202665029 | 76 | -1.51749432826524 | GDF1 | 0.0013271 | 637,75 | GDF1 |
| 2308.24667123905 | 55 | -1.7667597504713 | CTDSP1 | 3.2118333 | 635,25 | CTDSP1 |
| 1535.19919603857 | 78 | -1.65797519046315 | TBL1X | 4.7072903 | 635 | TBL1X |
| 2479.66910238534 | 51 | -1.16922932151196 | TMEM219 | 0.0074214 | 633,75 | TMEM219 |
| 1457.00730788802 | 82 | -1.71493637122238 | VEGFB | 5.4799574 | 633,75 | VEGFB |
| 1721.62115579012 | 66 | -2.17229486126769 | NBPF14 | 8.9523606 | 632,5 | NBPF14 |
| 1674.0671399711 | 64 | -1.57352654001462 | EFEMP2 | 0.0016948 | 629 | EFEMP2 |
| 1566.70396464181 | 63 | -2.3058603526514 | VSIR | 3.9456324 | 624,5 | VSIR |
| 2176.723529787 | 45 | -1.61004752058078 | COPZ2 | 0.0024004 | 618 | COPZ2 |
| 1806.33604458922 | 50 | -1.25050097818365 | KANK2 | 0.0040186 | 618 | KANK2 |
| 1304.05035649838 | 74 | -1.33074481341042 | CERS1 | 0.0037623 | 617 | CERS1 |
| 1790.18491596383 | 48 | -1.54676488840795 | LRTOMT | 4.0488943 | 613,75 | LRTOMT |
| 1284.11901511444 | 65 | -1.79120741144673 | RAB31 | 7.4343148 | 609,5 | RAB31 |
| 1307.1773791983 | 60 | -1.78412284007295 | SLC7A11 | 0.0012380 | 606,25 | SLC7A11 |
| 1165.18373240436 | 67 | -1.30976125261645 | TCF7L2 | 0.0020890 | 605 | TCF7L2 |
| 1123.10427451561 | 63 | -1.36483733338586 | PLD2 | 0.0034616 | 599,5 | PLD2 |
| 9660.23653367638 | 19 | -1.95986617287722 | ZCCHC24 | 8.1389262 | 587,25 | ZCCHC24 |
| 1569.57456955781 | 37 | -1.43773787666326 | DHRS3 | 0.0031594 | 586,25 | DHRS3 |
| 2944.28913589012 | 23 | -1.74057757474429 | PAQR6 | 3.0120904 | 585,25 | PAQR6 |
| 4572.80121989653 | 20 | -1.8771261984192 | BCAS1 | 3.7411225 | 585 | BCAS1 |
| 4164.00223125919 | 20 | -1.56951698235902 | ANKRD40 | 2.6903398 | 583 | ANKRD40 |
| 1209.187426915 | 42 | -1.74474408643972 | IFI35 | 0.0013271 | 573,5 | IFI35 |
| 3043.31630668952 | 20 | -1.61404664568187 | MAP4 | 1.4053328 | 571 | MAP4 |
| 894.191376045702 | 50 | -2.30086023424496 | PARD3 | 9.2289517 | 570,5 | PARD3 |
| 1603.55352580458 | 28 | -2.36549393572211 | PARD3B | 0.0020926 | 568,25 | PARD3B |
| 774.534285660528 | 52 | -2.19629875297377 | NWD1 | 3.0120904 | 562,5 | NWD1 |
| 719.042738088624 | 57 | -1.51820312912293 | ZC3HAV1 | 0.0017679 | 560,5 | ZC3HAV1 |
| 706.773474634858 | 57 | -2.20532916846353 | ENHO | 3.3107045 | 557,5 | ENHO |
| 877.820404913904 | 42 | -1.08977255339033 | CACFD1 | 0.0052606 | 556,5 | CACFD1 |
| 834.001074690484 | 43 | -1.66495069311379 | HSPB1 | 0.0028576 | 555,25 | HSPB1 |
| 1374.87271724638 | 27 | -2.79003336542742 | ADAMTS9 | 2.9092641 | 554 | ADAMTS9 |
| 720.441042866883 | 51 | -1.58697270239882 | FCGRT | 0.0025811 | 553,75 | FCGRT |
| 1365.2535589391 | 26 | -1.4579846182037 | LTBR | 0.0014727 | 548,25 | LTBR |
| 810.487960911063 | 41 | -1.39385665042152 | NPAS3 | 0.0040184 | 547,25 | NPAS3 |
| 2982.55934620718 | 16 | -1.07082536292997 | GALNT10 | 7.1883013 | 545,5 | GALNT10 |
| 590.265494886792 | 53 | -1.39707952181232 | CCDC69 | 0.0014160 | 530,75 | CCDC69 |
| 748.768988194375 | 36 | -1.38686437486299 | PPP1R14B | 0.0067759 | 530,75 | PPP1R14B |
| 742.179029126285 | 36 | -1.76261201960492 | AGTRAP | 0.0028620 | 529,75 | AGTRAP |
| 1304.57641974914 | 21 | -1.16276588977586 | DYNC1LI2 | 2.1951423 | 522,5 | DYNC1LI2 |
| 1056.17420850986 | 23 | -2.33443858897694 | BST2 | 4.0109922 | 521,25 | BST2 |
| 479.866151454173 | 55 | -1.40072907720971 | METRNL | 0.0034540 | 518,25 | METRNL |
| 2189.43390352404 | 14 | -1.28610093666449 | CENPB | 0.0021343 | 508,75 | CENPB |
| 616.478275707757 | 38 | -1.59750008044161 | EMID1 | 0.0031553 | 507,5 | EMID1 |
| 574.774310314906 | 41 | -1.54172861073276 | PPARA | 0.0040186 | 506,75 | PPARA |
| 569.070199888741 | 40 | -1.52150386361491 | RARRES2 | 0.0056087 | 504 | RARRES2 |
| 786.4550079049 | 23 | -2.07603046209163 | CALD1 | 0.0027348 | 501,25 | CALD1 |
| 774.027325969821 | 23 | -1.734551282242 | PHF21B | 0.0033652 | 499,25 | PHF21B |
| 4384.97241278317 | 9 | -1.12394129468834 | ZBTB47 | 0.0011708 | 498,75 | ZBTB47 |
| 394.949528645187 | 45 | -1.372014631637 | ELP4 | 0.0042924 | 497 | ELP4 |
| 1345.10690369161 | 16 | -1.84035691003327 | MBP | 0.0011376 | 495,5 | MBP |
| 584.358439944856 | 34 | -1.41607822710342 | TRIOBP | 0.0045722 | 495,5 | TRIOBP |
| 1340.05421157668 | 16 | -1.57096747008835 | IRF1 | 0.0044682 | 495 | IRF1 |
| 326.129848561821 | 50 | -2.10407833638831 | FGF11 | 0.0011376 | 494,5 | FGF11 |
| 391.753051158994 | 44 | -1.55776464047232 | PAX6 | 0.0030958 | 493,5 | PAX6 |
| 1273.99389384709 | 16 | -1.72150257817722 | PRKX | 0.0050886 | 491,5 | PRKX |
| 304.829160113602 | 50 | -1.95133917163769 | ZFP36L1 | 0.0014225 | 491 | ZFP36L1 |
| 564.97623629308 | 30 | -1.86628857461693 | NOTCH2NL | 0.0027029 | 483,75 | NOTCH2NLB |
| 1279.79480637619 | 15 | -1.46736083344518 | FGF1 | 5.3898564 | 483 | FGF1 |

| | | | | | | |
|---|---|---|---|---|---|---|
| 531.203193472302 | 31 | -1.38940993987489 | SFXN5 | 0.0052733 | 482,25 | SFXN5 |
| 752.707162703845 | 20 | -1.18272081784577 | CTNS | 0.0067187 | 481 | CTNS |
| 903.039807815327 | 17 | -1.01624127103948 | TRIM41 | 9.4425430 | 479 | TRIM41 |
| 1019.71918908418 | 16 | -1.96716224467448 | HIPK2 | 2.6397369 | 478 | HIPK2 |
| 1500.40939118477 | 12 | -1.3470122137622 | RAI14 | 7.7208845 | 477,5 | RAI14 |
| 1498.15025017808 | 12 | -1.36176522152276 | CSPG4 | 0.0025090 | 477 | CSPG4 |
| 1135.01770089121 | 15 | -1.5026203680723 | SGMS2 | 0.0011708 | 476 | SGMS2 |
| 1628.25482630149 | 11 | -1.23712642885881 | DAAM2 | 0.0032399 | 474 | DAAM2 |
| 330.275282474113 | 37 | -2.02493697484625 | PTH1R | 0.0011376 | 473,25 | PTH1R |
| 306.052852951331 | 39 | -1.20717723515961 | ASPH | 0.0028620 | 472,5 | ASPH |
| 573.447368246326 | 25 | -1.01792693122207 | GDF11 | 0.0089619 | 472,25 | GDF11 |
| 2425.70769170862 | 9 | -1.2377842626596 | TNS1 | 8.8590426 | 469,75 | TNS1 |
| 610.12406795995 | 23 | -1.60463458459506 | ZBTB20 | 0.0069084 | 468,75 | ZBTB20 |
| 4920.22098538825 | 6 | -1.39530970634437 | SLC6A12 | 5.8425105 | 467,75 | SLC6A12 |
| 417.35591945666 | 27 | -1.61670117987215 | FOXO1 | 9.9907083 | 461,5 | FOXO1 |
| 446.698040948614 | 26 | -1.29543247418315 | PIEZO1 | 0.0073769 | 460,75 | PIEZO1 |
| 1002.83076805522 | 14 | -1.65226467914237 | SMARCC1 | 0.0032309 | 459,75 | SMARCC1 |
| 338.490561518396 | 28 | -1.23646869023637 | PSMB8 | 0.0030750 | 455,25 | PSMB8 |
| 294.53328080202 | 31 | -2.37854069140498 | FAM167B | 0.0011708 | 453,25 | FAM167B |
| 943.014016891677 | 13 | -1.23585185218222 | FBXW4 | 7.4060703 | 452,25 | FBXW4 |
| 534.039582619118 | 22 | -1.88102432892611 | TRIM38 | 0.0021343 | 451,75 | TRIM38 |
| 1174.6326700615 | 11 | -1.00044238899913 | KLHL5 | 0.0017928 | 449,5 | KLHL5 |
| 211.343795596236 | 33 | -1.32926954413336 | ALDH6A1 | 0.0034104 | 445,5 | ALDH6A1 |
| 595.398255941715 | 19 | -1.31186582211396 | KTN1 | 0.0041041 | 444,75 | KTN1 |
| 771.130981180817 | 14 | -1.72168134925702 | ADIRF | 7.4060703 | 440,75 | ADIRF |
| 111.503153965741 | 39 | -1.80845121945401 | RIN3 | 0.0026638 | 437 | RIN3 |
| 370.720183419192 | 22 | -1.75386018411093 | IL6R | 0.0048888 | 435,25 | IL6R |
| 250.633267372099 | 26 | -1.10625574737323 | HVCN1 | 0.0039160 | 434,75 | HVCN1 |
| 599.102785059051 | 17 | -1.54057598131795 | ROCK1 | 0.0052940 | 433,5 | ROCK1 |
| 429.534048141499 | 20 | -1.87736637306933 | KIF1C | 3.7753794 | 431,5 | KIF1C |
| 784.87383941342 | 12 | -1.46666263913369 | PLEKHB1 | 1.2052065 | 431 | PLEKHB1 |
| 952.805251391111 | 10 | -1.09226073474914 | MOCS1 | 0.0059162 | 428 | MOCS1 |
| 168.266804469879 | 26 | -1.34899004467958 | NT5DC2 | 0.0041204 | 418,75 | NT5DC2 |
| 1397.60201839151 | 7 | -1.27025890583465 | WNK1 | 2.4263087 | 418 | WNK1 |
| 252.555146878848 | 22 | -1.6253523734586 | HSD3B7 | 0.0068032 | 417,75 | HSD3B7 |
| 127.522464239269 | 27 | -1.37102624880074 | SRGAP1 | 0.0069665 | 417 | SRGAP1 |
| 358.183695825312 | 18 | -2.31758766524186 | ANGPT2 | 3.7684378 | 413,5 | ANGPT2 |
| 480.745689800428 | 16 | -1.87199229828239 | TRIM56 | 0.0035193 | 413 | TRIM56 |
| 321.53074531711 | 19 | -2.65281084974373 | XAF1 | 5.4514639 | 412,25 | XAF1 |
| 391.973409362523 | 17 | -1.37624055693146 | PRICKLE3 | 0.0072137 | 411 | PRICKLE3 |
| 217.649991833226 | 20 | -1.78523688113485 | SH3BP2 | 0.0019707 | 401,5 | SH3BP2 |
| 92.8673089276522 | 24 | -1.13460143569743 | ORAI3 | 0.0080073 | 398,25 | ORAI3 |
| 919.535859486852 | 7 | -1.60090294261068 | CTNNA3 | 0.0074473 | 390 | CTNNA3 |
| 104.02033217787 | 22 | -1.49307889410142 | C1QTNF5 | 0.0054531 | 389 | C1QTNF5 |
| 104.02033217787 | 22 | -1.48331657805591 | MFRP | 0.0057505 | 389 | MFRP |
| 115.005869803215 | 21 | -1.30672668010038 | ARSD | 0.0079443 | 387,5 | ARSD |
| 372.30022538361 | 14 | -1.60283354454546 | ZIC4 | 0.0022734 | 384,75 | ZIC4 |
| 186.08004903727 | 18 | -1.16498615405275 | LPP | 0.0095760 | 384,5 | LPP |
| 64.1678226986348 | 22 | -1.16073993711222 | MSRB3 | 0.0066414 | 377,75 | MSRB3 |
| 146.082314312161 | 18 | -2.00454762086486 | BAZ1A | 0.0044833 | 377,5 | BAZ1A |
| 510.710849850568 | 11 | -1.65489922160113 | KIF5B | 4.0879921 | 377 | KIF5B |
| 112.294837638194 | 19 | -2.5322967483855 | HIF3A | 7.3104425 | 376,75 | HIF3A |
| 710.887089435948 | 8 | -2.06483416353654 | FLT1 | 7.7728115 | 375,25 | FLT1 |
| 37.053691118586 | 24 | -1.65674733328184 | EMX2 | 0.0046462 | 373,75 | EMX2 |
| 537.577019945143 | 10 | -1.51596369782731 | FAM114A1 | 0.0075790 | 371 | FAM114A1 |
| 338.929138283824 | 12 | -1.42364760201825 | LPAR1 | 0.0060034 | 369 | LPAR1 |
| 172.105772931878 | 16 | -1.36357317223974 | PRELP | 0.0010387 | 368,5 | PRELP |
| 112.401467493783 | 16 | -2.1019636506586 | RPGR | 0.0014038 | 358,5 | RPGR |
| 241.190802559887 | 12 | -1.43434232802962 | REST | 0.0083411 | 353 | REST |
| 686.210314395459 | 6 | -1.19125085193138 | CAPG | 0.0068297 | 346,75 | CAPG |
| 73.3592689604174 | 14 | -1.84481237857072 | PAIP2B | 2.4949648 | 329,25 | PAIP2B |
| 26.9338577334677 | 17 | -1.43942983176881 | CMTM6 | 0.0048038 | 327,5 | CMTM6 |
| 32.1859533288228 | 16 | -1.87949904239039 | APOL6 | 0.0015711 | 323,5 | APOL6 |
| 88.1647692542281 | 12 | -1.39203948448754 | MTURN | 2.4203180 | 323 | MTURN |
| 184.657016959599 | 10 | -1.78569667872469 | TP53INP2 | 6.2283965 | 322,5 | TP53INP2 |
| 686.416666666667 | 4 | -1.47705181915748 | MOBP | 0.0064627 | 321 | MOBP |

| | | | | | | |
|---|---|---|---|---|---|---|
| 76.8103377593929 | 12 | -1.97846744424301 | NFATC2 | 8.3180261 | 318,5 | NFATC2 |
| 99.7333342027999 | 11 | -1.5160852797275 | FOXO4 | 3.8229985 | 316,5 | FOXO4 |
| 341.325935419369 | 7 | -1.63027938099549 | EMP2 | 0.0010363 | 316 | EMP2 |
| 38.1210486789734 | 14 | -2.05624185578331 | TBX2 | 9.6098136 | 311,25 | TBX2 |
| 98.5228876883964 | 10 | -2.00680235715677 | FXYD3 | 0.0037214 | 306 | FXYD3 |
| 30.6667512700756 | 14 | -1.83415325358248 | ASCL1 | 0.0095002 | 305,25 | ASCL1 |
| 136.716488869797 | 9 | -1.23704603139524 | ACACB | 0.0088110 | 302,75 | ACACB |
| 686.0 | 3 | -2.30120632746891 | IRF7 | 8.6821682 | 302 | IRF7 |
| 12.5786720208364 | 15 | -1.40663471600074 | ASRGL1 | 0.0049864 | 295,5 | ASRGL1 |
| 45.4012949512483 | 11 | -1.21307409761418 | NOL8 | 0.0089791 | 295,5 | NOL8 |
| 49.4660606960052 | 10 | -1.90256412903563 | TIMP1 | 6.9606381 | 289 | TIMP1 |
| 6.47888178063624 | 15 | -1.0829812466221 | ALDH7A1 | 0.0087083 | 286,5 | ALDH7A1 |
| 112.178072681005 | 8 | -1.40916696147333 | MYOM1 | 0.0016462 | 283,75 | MYOM1 |
| 60.6311912677944 | 9 | -2.14399495297601 | MYL3 | 9.8350899 | 281,25 | MYL3 |
| 17.8786875860193 | 12 | -2.34684122406246 | NCKAP5 | 0.0013210 | 281 | NCKAP5 |
| 36.2303177773589 | 10 | -1.0882111689258 | PDLIM3 | 9.9320251 | 279,5 | PDLIM3 |
| 56.1055025727432 | 9 | -1.48161966359868 | LRRCC1 | 0.0045992 | 277,75 | LRRCC1 |
| 55.4878230045393 | 9 | -3.29195717856976 | CDC45 | 0.0033686 | 277,25 | CDC45 |
| 11.6609961650363 | 12 | -2.28817201886308 | C16orf74 | 7.2634492 | 274 | C16orf74 |
| 29.9876222488279 | 10 | -1.73063418760247 | TRIP10 | 0.0015785 | 274 | TRIP10 |
| 46.1411749683075 | 9 | -2.06484844958194 | ZNF366 | 0.0054096 | 272,75 | ZNF366 |
| 9.86285875345084 | 12 | -1.7250917102392 | CYP11A1 | 0.0072820 | 271 | CYP11A1 |
| 14.1401551211803 | 11 | -1.8227282694143 | ZFC3H1 | 0.0085219 | 268,5 | ZFC3H1 |
| 10.4160961651991 | 11 | -1.90447167584143 | RGL3 | 0.0048297 | 264,5 | RGL3 |
| 185.325944149409 | 5 | -1.2714350329415 | CARHSP1 | 0.0048888 | 261,75 | CARHSP1 |
| 16.364596363363 | 10 | -2.16293940076962 | NAA16 | 0.0037873 | 260,5 | NAA16 |
| 170.256745171068 | 5 | -1.05950197401333 | DENND2A | 7.2815330 | 259,25 | DENND2A |
| 80.1137675703882 | 6 | -1.18443234527593 | OTUD7B | 0.0081819 | 253,75 | OTUD7B |
| 13.0670802102118 | 9 | -2.30800619054932 | LPAR3 | 8.9523606 | 244,25 | LPAR3 |
| 22.797033691928 | 8 | -2.36881138820375 | PLEKHA4 | 9.4425430 | 241,75 | PLEKHA4 |
| 32.8238089091772 | 7 | -1.28544769000699 | MCM7 | 0.0027021 | 241 | MCM7 |
| 3.65130001130032 | 10 | -2.06744838707335 | CFAP53 | 0.0083466 | 240,5 | CFAP53 |
| 6.47373590542912 | 9 | -1.38699978907263 | MYBPC1 | 0.0048090 | 234,25 | MYBPC1 |
| 58.2080829427788 | 5 | -1.43773085161921 | ZMYM5 | 0.0067187 | 231,25 | ZMYM5 |
| 82.8273652392635 | 4 | -1.60732121581016 | MAP4K4 | 0.0052435 | 228,5 | MAP4K4 |
| 3.41153139400833 | 9 | -1.73898795076374 | NFATC4 | 0.0056272 | 226,25 | NFATC4 |
| 68.4789921939023 | 4 | -1.01643192181752 | CRTAP | 0.0033582 | 224,5 | CRTAP |
| 14.8722713181417 | 7 | -1.47533504769718 | EMCN | 0.0043239 | 223,5 | EMCN |
| 24.9374365023861 | 6 | -1.31620846755252 | GAB2 | 0.0030178 | 223,5 | GAB2 |
| 24.9374365023861 | 6 | -1.41378616850165 | SLCO1A2 | 0.0063079 | 223,5 | SLCO1A2 |
| 7.08269124773333 | 8 | -1.51392997199306 | NTSR2 | 0.0039504 | 222,75 | NTSR2 |
| 44.7844894724576 | 5 | -1.82126344890581 | RREB1 | 0.0042609 | 222,75 | RREB1 |
| 24.2902616837941 | 6 | -1.2272096187708 | RASSF2 | 8.4037975 | 221,75 | RASSF2 |
| 15.455948586482 | 6 | -1.53048634310238 | N4BP2 | 0.0075180 | 212,75 | N4BP2 |
| 5.82757346781259 | 7 | -1.52917608239298 | ITPRIPL2 | 0.0044697 | 209,5 | ITPRIPL2 |
| 1.58049396956441 | 8 | -1.99349084648135 | CHD7 | 0.0062681 | 207,25 | CHD7 |
| 19.7684129994353 | 5 | -1.45550351565983 | ANP32B | 0.0048897 | 203,25 | ANP32B |
| 15.0351992865971 | 5 | -1.34185614339726 | HERC5 | 8.4617028 | 198,25 | HERC5 |
| 2.41874717199424 | 6 | -1.69743985893769 | BATF2 | 0.0058642 | 190,25 | BATF2 |
| 2.35853928765729 | 5 | -1.30108508140125 | EFNB1 | 0.0076091 | 175,75 | EFNB1 |
| 2.07692501577082 | 5 | -1.8516594464784 | PGGHG | 0.0019527 | 175,25 | PGGHG |
| 17.5028083141302 | 3 | -1.65381834346194 | CD109 | 0.0022894 | 173,75 | CD109 |
| 3.74225589547836 | 4 | -1.16263570169251 | AIF1L | 0.0068032 | 168 | AIF1L |
| 0.0 | 7 | -1.30056603937012 | PKP4 | 9.8350899 | 161,25 | PKP4 |
| 0.421765528895567 | 4 | -1.80428211101832 | AEBP1 | 0.0015583 | 154,5 | AEBP1 |
| 0.386874236874237 | 4 | -2.23783171063382 | MECOM | 0.0016821 | 154 | MECOM |
| 1.99143473797341 | 3 | -1.71262382670212 | PGF | 0.0049656 | 147,25 | PGF |
| 0.797180088424931 | 2 | -1.11259211347206 | MIDN | 0.0074473 | 117,75 | MIDN |
| 0.0 | 3 | -2.22726229052159 | CCDC190 | 0.0074214 | 108,5 | CCDC190 |
| 0.0 | 3 | -1.28555596118857 | SHTN1 | 0.0035193 | 108,5 | SHTN1 |
| 0.0 | 3 | -1.475855984882 | TGFB3 | 0.0025000 | 108,5 | TGFB3 |
| 0.0 | 2 | -1.29716649599625 | NDRG1 | 0.0022347 | 84,5 | NDRG1 |
| 0.0 | 2 | -1.05698055443371 | NUDT16 | 1.2437414 | 84,5 | NUDT16 |
| 0.0 | 2 | -1.2229515431702 | PSMB9 | 0.0093347 | 84,5 | PSMB9 |
| 0.0 | 2 | -1.56070761244132 | SLC38A3 | 7.5239403 | 84,5 | SLC38A3 |
| 0.0 | 2 | -1.42908831840161 | TCF7 | 0.0036373 | 84,5 | TCF7 |

| | | | | | | | |
|---|---|---|---|---|---|---|---|
| | 0.0 | 1 | -1.50552277102552 | MDFI | 0.0016678 | 49,75 | MDFI |
| | 0.0 | 1 | -1.09154508215119 | ARHGEF37 | 0.0056938 | 49,75 | ARHGEF37 |
| | 0.0 | 1 | -1.04595999565824 | RPS27 | 0.0076673 | 49,75 | RPS27 |
| | 0.0 | 1 | -1.97552095609579 | THSD4 | 0.0021391 | 49,75 | THSD4 |
| | 0.0 | 1 | -2.35986570486936 | TYMP | 0.0021343 | 49,75 | TYMP |
| | 0.0 | 1 | -1.47230068509477 | YES1 | 1.4575726 | 49,75 | YES1 |
| | 0.0 | 1 | -1.18229356652983 | ZEB2 | 0.0035375 | 49,75 | ZEB2 |
| 3 | 8435.68013317779 | 66 | -2.09205516761737 | TCN2 | 7.1521628 | 685,5 | TCN2 |
| | 6527.74012675882 | 58 | -1.67908826459219 | HLA-E | 3.6504628 | 676,25 | HLA-E |
| | 11225.4267941402 | 50 | -2.02660254360037 | RELL1 | 3.3026633 | 668,5 | RELL1 |
| | 6244.06637884893 | 34 | -1.80512289565505 | ACVRL1 | 8.0328097 | 636 | ACVRL1 |
| | 2306.91141048684 | 42 | -1.22763717526537 | SIPA1 | 0.0039556 | 614,5 | SIPA1 |
| | 2438.74421760483 | 40 | -1.63372245620827 | SYDE1 | 0.0020890 | 612,5 | SYDE1 |
| | 2809.25821005007 | 32 | -1.35527488656327 | CFLAR | 0.0031230 | 609,25 | CFLAR |
| | 1940.30875628408 | 38 | -2.34962622968297 | GGT5 | 1.0843279 | 598,5 | GGT5 |
| | 1696.9941622929 | 41 | -2.62274734325206 | SERPINH1 | 9.7074511 | 596,75 | SERPINH1 |
| | 2120.07757156666 | 33 | -1.11206507687372 | RGS3 | 0.0014611 | 592,5 | RGS3 |
| | 2587.1741219399 | 26 | -1.64985908805594 | PXN | 6.6646126 | 587,75 | PXN |
| | 1762.50052625904 | 31 | -1.75507844747474 | DOCK6 | 0.0014543 | 581,25 | DOCK6 |
| | 2319.84405627734 | 24 | -1.15798577114641 | VASP | 0.0074214 | 575,25 | VASP |
| | 2799.8015859298 | 22 | -3.14951610423915 | CDKN1A | 1.4053328 | 574,25 | CDKN1A |
| | 3763.18240919726 | 17 | -1.74551683712102 | CLIC1 | 1.1772056 | 562,5 | CLIC1 |
| | 1458.22355877365 | 28 | -1.42572528587952 | KANK3 | 7.9613938 | 562,25 | KANK3 |
| | 3002.99699141679 | 17 | -2.40527951773192 | OSMR | 1.4053328 | 553 | OSMR |
| | 3087.73740772789 | 16 | -2.79415517022974 | IL4R | 1.2821704 | 547 | IL4R |
| | 4026.05961339389 | 13 | -2.19950698128038 | IFITM3 | 1.5677255 | 533,75 | IFITM3 |
| | 1701.17122335467 | 19 | -2.2256468594815 | SLC5A10 | 1.5389879 | 532,75 | SLC5A10 |
| | 1416.92407916451 | 19 | -2.82007803302423 | LRRC32 | 1.4105385 | 521,25 | LRRC32 |
| | 743.608737538677 | 30 | -2.42857633659642 | TNFRSF1B | 4.5817353 | 517,25 | TNFRSF1B |
| | 1777.73021678137 | 16 | -2.37908424765224 | ITGA5 | 6.6795604 | 517 | ITGA5 |
| | 2569.28197468803 | 13 | -2.33539179895772 | ACSL5 | 3.2152877 | 512,25 | ACSL5 |
| | 2804.25559855198 | 11 | -1.23540462302653 | DIPK2B | 0.0011684 | 503,5 | DIPK2B |
| | 2774.86961857902 | 11 | -1.48531920776416 | ENG | 7.7858826 | 501 | ENG |
| | 736.192851527493 | 23 | -2.13376829930063 | GIMAP5 | 3.9456324 | 494,75 | GIMAP5 |
| | 756.947505990529 | 22 | -1.84850552883225 | SHE | 6.4485871 | 491,25 | SHE |
| | 826.376029417443 | 20 | -1.86001973805955 | SLC12A7 | 3.1741357 | 490 | SLC12A7 |
| | 1071.1200515062 | 16 | -2.35310771084964 | CLDN5 | 1.8359146 | 482 | CLDN5 |
| | 1858.18571369084 | 11 | -2.53812700242786 | HEYL | 3.5884739 | 481,5 | HEYL |
| | 862.974017186413 | 18 | -1.20059494186644 | LYPD1 | 0.0015711 | 481,5 | LYPD1 |
| | 787.203449530921 | 18 | -1.62738101903769 | PECAM1 | 3.3026633 | 474 | PECAM1 |
| | 624.348538277959 | 22 | -2.03094986009588 | ADGRL4 | 9.3312729 | 463,75 | ADGRL4 |
| | 913.678174797771 | 15 | -1.962485973169 | SP100 | 7.2815330 | 463,5 | SP100 |
| | 1482.19684356625 | 10 | -2.1925189021406 | EPHA2 | 2.1951423 | 457,5 | EPHA2 |
| | 1784.89671458911 | 9 | -1.91220067120532 | IFITM1 | 9.3312729 | 456,75 | IFITM1 |
| | 631.845770533647 | 20 | -1.26399507020203 | TSPAN4 | 0.0057565 | 455 | TSPAN4 |
| | 1071.60720862588 | 12 | -1.41224023166731 | BGN | 6.1681994 | 453,5 | BGN |
| | 632.063430635049 | 18 | -3.10391130986004 | TGM2 | 3.8460093 | 444 | TGM2 |
| | 2602.30660649475 | 6 | -1.23180003537352 | CASC3 | 1.1199223 | 441,75 | CASC3 |
| | 1288.7293523985 | 9 | -5.05711892423029 | SOCS3 | 3.8460093 | 432,25 | SOCS3 |
| | 1042.75303158723 | 10 | -7.66569466514523 | CSF3 | 2.0614406 | 432 | CSF3 |
| | 547.635968119566 | 18 | -1.70267973569072 | GSDMD | 0.0032215 | 432 | GSDMD |
| | 933.171397937935 | 10 | -1.84152607065168 | AL645922.1 | 0.0016318 | 427 | AL645922.1 |
| | 973.861694813282 | 9 | -2.03607826308843 | WWTR1 | 6.3040216 | 415,75 | WWTR1 |
| | 2127.34653996755 | 5 | -1.79827409324316 | SLCO2B1 | 2.9092641 | 415,25 | SLCO2B1 |
| | 1396.77350277573 | 6 | -2.44451722635744 | A4GALT | 9.7074511 | 405,75 | A4GALT |
| | 2127.86987848707 | 4 | -1.56226500227436 | SQOR | 9.4536784 | 403,5 | SQOR |
| | 328.793380388104 | 16 | -2.63061435316473 | ADCY4 | 1.1451180 | 395,5 | ADCY4 |
| | 852.944417464949 | 8 | -1.46275115815857 | HIGD1B | 0.0017587 | 393,75 | HIGD1B |
| | 268.415094223322 | 17 | -1.70616102659325 | ITGA1 | 0.0025961 | 392,5 | ITGA1 |
| | 191.730564190429 | 19 | -1.53946927201716 | APOL3 | 2.2219832 | 391,75 | APOL3 |
| | 791.635465506794 | 8 | -1.9661328505425 | NFKB2 | 5.8425105 | 388,25 | NFKB2 |
| | 395.381235327303 | 14 | -1.27519837217534 | BCAM | 0.0045595 | 387,75 | BCAM |
| | 696.58963799712 | 9 | -2.34106093215089 | YBX3 | 9.0344713 | 385,25 | YBX3 |
| | 693.737115345412 | 9 | -1.92355545485273 | ENSG0000 | 1.3634941 | 384,25 | ENSG00000288681 |
| | 1379.92634753715 | 4 | -1.47540323492893 | BTG2 | 3.7411225 | 378,5 | BTG2 |
| | 439.205207083799 | 12 | -2.72290005090829 | CASP4 | 2.6903398 | 378,5 | CASP4 |

| | | | | | | |
|---|---|---|---|---|---|---|
| 1024.45427203383 | 5 | -1.55531428630503 | GADD45G | 0.0044563{ | 369,75 | GADD45G |
| 720.725848647934 | 7 | -2.25128275168003 | MAFF | 2.9077415( | 369,5 | MAFF |
| 300.360447429256 | 13 | -1.90635653127579 | PARP12 | 0.0011807. | 367,25 | PARP12 |
| 175.354930182543 | 15 | -1.52121858995063 | TGFB1I1 | 0.0041400. | 360 | TGFB1I1 |
| 648.246535804207 | 8 | -1.88730361519795 | GIMAP1-GI | 8.7954351. | 358,25 | GIMAP1-GIMAP5 |
| 310.108343683558 | 11 | -2.26565124041641 | BCL6B | 4.5428113( | 355,5 | BCL6B |
| 808.112087560934 | 5 | -3.78052492473572 | DEPP1 | 3.8460093{ | 354,25 | DEPP1 |
| 399.62150491635 | 9 | -2.10199357294576 | CFB | 9.8350899! | 344,75 | CFB |
| 168.985117590525 | 13 | -1.18550160144134 | MYO1C | 0.0083193: | 344,25 | MYO1C |
| 240.836692721564 | 11 | -2.36820557390759 | GRAP | 1.1451180. | 344 | GRAP |
| 205.874271057141 | 11 | -1.25826013309674 | NOTCH4 | 0.0030958{ | 338 | NOTCH4 |
| 1370.0 | 2 | -1.14675522469493 | MMRN2 | 5.9213956. | 337,75 | MMRN2 |
| 1370.0 | 2 | -2.56909709380804 | ANXA2 | 3.3107045! | 337,75 | ANXA2 |
| 1370.0 | 2 | -1.87431898611158 | TGFBR2 | 6.2320028. | 337,75 | TGFBR2 |
| 115.014107829691 | 13 | -1.18434583148845 | GPR39 | 0.0013739< | 336,75 | GPR39 |
| 236.358248647215 | 10 | -1.27483338167479 | TP53INP1 | 0.0049083! | 333,5 | TP53INP1 |
| 305.089007273409 | 9 | -3.71601298981144 | TNFRSF10D | 1.5481559( | 331,25 | TNFRSF10D |
| 196.621928546143 | 10 | -1.49077641862608 | HSPG2 | 0.0061858: | 326,5 | HSPG2 |
| 722.783620663551 | 3 | -1.04115531631322 | TMCO3 | 1.3405532. | 317,25 | TMCO3 |
| 697.573977555976 | 3 | -1.80181288234891 | CMTM3 | 9.8350899! | 311,75 | CMTM3 |
| 353.478344699502 | 6 | -1.93441883512116 | B4GALT1 | 3.2152877! | 304,75 | B4GALT1 |
| 157.274388526768 | 9 | -1.59544284481383 | PLEKHG2 | 0.0052733: | 304,25 | PLEKHG2 |
| 143.230282298904 | 9 | -1.60808710787493 | NLRC5 | 0.0065553: | 303,25 | NLRC5 |
| 686.0 | 3 | -1.69007801856872 | FGR | 0.0011376( | 302 | FGR |
| 603.622209574478 | 3 | -2.52904371987814 | ZBTB42 | 3.7411225: | 291,25 | ZBTB42 |
| 60.9071252233344 | 9 | -2.03515560476669 | MFNG | 5.9417943. | 281,75 | MFNG |
| 686.0 | 2 | -1.44990382198491 | APOLD1 | 3.9456324{ | 278 | APOLD1 |
| 686.0 | 2 | -1.93196073320727 | IL1R1 | 0.0014696: | 278 | IL1R1 |
| 686.0 | 2 | -1.06448350006985 | NKD1 | 5.4633186< | 278 | NKD1 |
| 686.0 | 2 | -1.1398231785388 | PODXL | 4.3760612. | 278 | PODXL |
| 686.0 | 2 | -1.80315873569536 | S100A10 | 9.1126397< | 278 | S100A10 |
| 686.0 | 2 | -1.52017944242075 | SOWAHC | 8.5499633! | 278 | SOWAHC |
| 175.598525693697 | 6 | -2.63611844060247 | GJA4 | 1.9101967: | 274,75 | GJA4 |
| 26.7135693809391 | 10 | -1.95629245365858 | PODN | 2.9950698: | 272,5 | PODN |
| 126.852726568029 | 6 | -1.02442315056246 | PTPN21 | 4.8466630< | 266,25 | PTPN21 |
| 56.1948468501793 | 8 | -2.16585327087127 | VWF | 1.4053328: | 265,25 | VWF |
| 340.930300607276 | 3 | -1.04246801563214 | OLIG1 | 5.4224796! | 262,75 | OLIG1 |
| 41.1618551073624 | 8 | -1.04216099648206 | NECTIN2 | 0.0074214< | 256,25 | NECTIN2 |
| 291.058603157595 | 3 | -1.19896473047846 | C4orf3 | 2.3291787! | 252,25 | C4orf3 |
| 77.1611465306075 | 5 | -2.42560900387559 | IFITM2 | 1.1605952< | 239,25 | IFITM2 |
| 20.7672677357671 | 7 | -1.62522854257014 | TAGLN2 | 8.1389262! | 231 | TAGLN2 |
| 47.2538851451615 | 5 | -1.6357645913845 | LDLRAD2 | 0.0067385! | 225,75 | LDLRAD2 |
| 41.3312511055447 | 5 | -1.11726099101365 | BNIP2 | 4.0195288( | 221,75 | BNIP2 |
| 3.70753264262126 | 8 | -2.34594154368817 | MLKL | 0.0033728{ | 214,75 | MLKL |
| 47.9063739560331 | 4 | -1.30971502960339 | PRR29 | 0.0014038. | 214,5 | PRR29 |
| 62.6305636874712 | 3 | -1.91978246041933 | CEBPD | 0.0025090! | 207,25 | CEBPD |
| 7.69747757909759 | 6 | -1.83997517932624 | NIBAN2 | 1.3405532. | 203,25 | NIBAN2 |
| 19.4938633478683 | 5 | -1.3233394843732 | IFI27 | 0.0017928. | 202,75 | IFI27 |
| 49.1644108183312 | 3 | -1.79158717092164 | SAMD4A | 1.8191691< | 200,25 | SAMD4A |
| 44.8541278180876 | 3 | -1.88029899162079 | C11orf96 | 0.0015168{ | 196,25 | C11orf96 |
| 19.0634701545636 | 4 | -1.76706398642632 | NOSTRIN | 0.0036918: | 190 | NOSTRIN |
| 82.826047414921 | 2 | -1.33981328969455 | CHST3 | 0.0033132! | 189,25 | CHST3 |
| 22.6245489484285 | 3 | -1.5743952123132 | ROBO4 | 9.2289517( | 178,75 | ROBO4 |
| 46.7482122678183 | 2 | -1.75517735067124 | ADAMTSL4 | 0.0045111! | 174,25 | ADAMTSL4 |
| 5.72691625147611 | 4 | -1.28483787847202 | EHD2 | 0.0062331. | 171 | EHD2 |
| 4.29218148645765 | 4 | -1.61882103850134 | GPER1 | 5.3898564! | 169,5 | GPER1 |
| 12.3871172606129 | 3 | -1.23582539428532 | FLI1 | 0.0066836( | 168,25 | FLI1 |
| 31.7218172473848 | 2 | -1.75156035919616 | SMIM3 | 4.0109922( | 163,25 | SMIM3 |
| 30.1364344759789 | 2 | -2.17133454292137 | RAB20 | 1.9506902{ | 162,25 | RAB20 |
| 26.2216332285387 | 2 | -1.03492194140998 | LSR | 0.0018729( | 159,25 | LSR |
| 0.513311546840959 | 4 | -1.5365663045387 | HSPB7 | 0.0023554. | 156 | HSPB7 |
| 3.93114796251456 | 3 | -1.46273830250557 | PRX | 0.0078186: | 154,25 | PRX |
| 3.77740095930969 | 3 | -1.80277483517717 | DLL4 | 0.0026638{ | 153,75 | DLL4 |
| 3.3084329365547 | 3 | -1.64140441752229 | TRIM5 | 0.0017925< | 150,75 | TRIM5 |
| 3.28900848669312 | 3 | -1.8158701560232 | SLC19A3 | 9.7427715( | 150,25 | SLC19A3 |
| 10.6754725482543 | 2 | -1.06810467121904 | RAB30 | 0.0057802( | 142,75 | RAB30 |

| | | | | | | | |
|---|---|---|---|---|---|---|---|
| | 6.37156666721821 | 2 | -5.17688716981005 | IL1R2 | 8.5385529( | 134,75 | IL1R2 |
| | 3.44260954315022 | 2 | -2.513996213292 | SLC52A3 | 7.4343148: | 127,75 | SLC52A3 |
| | 0.875 | 2 | -1.01207109136606 | GCSH | 2.0923739( | 118,25 | GCSH |
| | 0.0 | 3 | -1.23465126785256 | ICAM2 | 0.0015584( | 108,5 | ICAM2 |
| | 0.0 | 3 | -1.59977436581419 | CFH | 0.0015711( | 108,5 | CFH |
| | 0.0 | 2 | -2.21222590741957 | CLEC2B | 7.4613662: | 84,5 | CLEC2B |
| | 0.0 | 2 | -1.49353922427667 | F11R | 0.0043464: | 84,5 | F11R |
| | 0.0 | 2 | -3.73920018496575 | ICAM1 | 1.4105385( | 84,5 | ICAM1 |
| | 0.0 | 2 | -1.8030839241682 | ITPKC | 9.6448987! | 84,5 | ITPKC |
| | 0.0 | 2 | -1.41226394978417 | LEF1 | 1.7291800! | 84,5 | LEF1 |
| | 0.0 | 2 | -1.61511397484986 | APOL1 | 0.0021343! | 84,5 | APOL1 |
| | 0.0 | 2 | -1.66951856373615 | SH3TC1 | 0.0023839( | 84,5 | SH3TC1 |
| | 0.0 | 2 | -1.88930392872807 | SLCO4A1 | 0.0032524 | 84,5 | SLCO4A1 |
| | 0.0 | 2 | -2.15063408571488 | GADD45B | 5.4150102! | 84,5 | GADD45B |
| | 0.0 | 2 | -3.47263433048376 | ZFP36 | 2.4220452! | 84,5 | ZFP36 |
| | 0.0 | 1 | -1.79305735877701 | BCL3 | 0.0023172 | 49,75 | BCL3 |
| | 0.0 | 1 | -1.69213334877263 | BHMT2 | 0.0011599! | 49,75 | BHMT2 |
| | 0.0 | 1 | -1.16224476595095 | CSRNP1 | 0.0026830! | 49,75 | CSRNP1 |
| | 0.0 | 1 | -1.87285977599936 | CARD10 | 0.0033648: | 49,75 | CARD10 |
| | 0.0 | 1 | -1.47821247934219 | CDH5 | 1.3405532 | 49,75 | CDH5 |
| | 0.0 | 1 | -2.42402754437134 | FOXF1 | 3.8665538( | 49,75 | FOXF1 |
| | 0.0 | 1 | -1.78848572266238 | IRAK3 | 0.0071216( | 49,75 | IRAK3 |
| | 0.0 | 1 | -2.79128435430761 | KCNE4 | 8.2244756 | 49,75 | KCNE4 |
| | 0.0 | 1 | -1.20600666001913 | CAVIN1 | 0.0032864( | 49,75 | CAVIN1 |
| | 0.0 | 1 | -2.72964755124708 | PLA1A | 9.4032749: | 49,75 | PLA1A |
| | 0.0 | 1 | -1.33544570770949 | PLIN2 | 0.0065487: | 49,75 | PLIN2 |
| | 0.0 | 1 | -1.27085448660136 | ESAM | 0.0011946: | 49,75 | ESAM |
| | 0.0 | 1 | -1.38862143949395 | PPP1R13L | 0.0079628! | 49,75 | PPP1R13L |
| | 0.0 | 1 | -1.30282918931787 | RASL12 | 0.0085339! | 49,75 | RASL12 |
| | 0.0 | 1 | -1.40094740212194 | S100A11 | 0.0089791( | 49,75 | S100A11 |
| | 0.0 | 1 | -2.56916391761183 | SH2D2A | 0.0012025: | 49,75 | SH2D2A |
| | 0.0 | 1 | -1.77090190486544 | GIMAP2 | 0.0021585! | 49,75 | GIMAP2 |
| | 0.0 | 1 | -1.07394259991702 | TIPARP | 0.0079161! | 49,75 | TIPARP |
| | 0.0 | 1 | -1.76548336921289 | TM4SF1 | 4.0195288( | 49,75 | TM4SF1 |
| | 0.0 | 1 | -1.70971694567118 | PRKCH | 1.2944890! | 49,75 | PRKCH |
| 4 | 1171.40391617754 | 28 | -1.0541592720369 | DDAH2 | 0.0060853: | 544,25 | DDAH2 |
| | 1725.822834735 | 15 | -2.23186970808633 | ENSG0000( | 6.9633907( | 506 | ENSG00000285526 |
| | 826.798359293441 | 16 | -1.46367142660967 | TMEM176/ | 0.0060738( | 466 | TMEM176A |
| | 1523.87456719122 | 6 | -1.25777341564043 | IGDCC4 | 0.0060578: | 412,75 | IGDCC4 |
| | 445.389639036451 | 15 | -1.28514346302836 | TGFBR3 | 0.0022857: | 399 | TGFBR3 |
| | 302.003912542023 | 17 | -1.20758995797419 | ASAP3 | 0.0013271: | 397,5 | ASAP3 |
| | 449.45189908276 | 11 | -2.35527485764981 | MT2A | 3.7684378( | 371,5 | MT2A |
| | 693.273903891543 | 8 | -1.6216109771929 | ROM1 | 0.0017928: | 370,75 | ROM1 |
| | 863.251587983312 | 4 | -1.40787520718553 | TMEM176E | 0.0067187 | 348 | TMEM176B |
| | 585.758364191544 | 5 | -1.40643847127611 | DTNA | 3.3026633! | 316,25 | DTNA |
| | 639.953959147925 | 4 | -1.65250088398547 | LGALS3 | 1.8946355: | 310,5 | LGALS3 |
| | 686.0 | 2 | -2.82597507217439 | EMP1 | 8.0328097! | 278 | EMP1 |
| | 58.9186920796665 | 8 | -3.04948684447332 | GFAP | 2.3291787! | 266,75 | GFAP |
| | 20.6909285348314 | 8 | -2.36721048450951 | NUPR1 | 1.9254263( | 239,75 | NUPR1 |
| | 28.4527182984847 | 7 | -1.57762433273662 | FAM189A2 | 0.0063644! | 238 | FAM189A2 |
| | 24.5304296843379 | 4 | -2.2677172713947 | ITGB4 | 1.3634941: | 196,5 | ITGB4 |
| | 2.06378362698564 | 6 | -2.13930720936761 | SLC2A4 | 0.0039660( | 188,75 | SLC2A4 |
| | 7.21457721168496 | 5 | -1.75783513050492 | ENSG0000( | 0.0011911: | 188,25 | ENSG00000288656 |
| | 8.51166445938175 | 4 | -2.51945662614134 | SLC14A1 | 7.8583325( | 177,5 | SLC14A1 |
| | 33.3241102846811 | 2 | -2.06231485940594 | BAG3 | 0.0011914! | 165,25 | BAG3 |
| | 18.7743108138253 | 2 | -1.5982242284845 | SPX | 4.4717466: | 150,75 | SPX |
| | 2.80598537567588 | 2 | -1.54041234601811 | CDH23 | 0.0049463: | 125,75 | CDH23 |
| | 0.0 | 3 | -2.51643118106797 | MT1X | 1.9124371: | 108,5 | MT1X |
| | 0.0 | 2 | -1.85088841501704 | MT1E | 0.0021550! | 84,5 | MT1E |
| | 0.0 | 2 | -1.36395719763325 | VIM | 9.6214204! | 84,5 | VIM |
| | 0.0 | 1 | -1.49400128382016 | ARHGAP31 | 0.0020528: | 49,75 | ARHGAP31 |
| | 0.0 | 1 | -1.3727054728055 | PPP1R1B | 0.0079671! | 49,75 | PPP1R1B |
| | 0.0 | 1 | -1.59991599726418 | NFATC1 | 0.0033582! | 49,75 | NFATC1 |
| 5 | 1958.967819506 | 25 | -3.49155762255721 | MT-CO2 | 1.4053328: | 570,25 | MT-CO2 |
| | 2133.0004682603 | 23 | -3.82900211501828 | MT-ND3 | 3.8460093! | 565,75 | MT-ND3 |
| | 872.719455458243 | 17 | -4.02705017841421 | MT-ND4 | 2.3291787! | 476,5 | MT-ND4 |

| | | | | | | | |
|---|---|---|---|---|---|---|---|
| | 780.66025025138 | 17 | -3.67201264127201 | MTRNR2L1 | 1.4053328 | 466,5 | MTRNR2L12 |
| | 803.420962351457 | 16 | -3.67648877830083 | MT-ATP6 | 5.4667654 | 462,5 | MT-ATP6 |
| | 624.256989625028 | 15 | -3.48950823862666 | MTRNR2L8 | 2.3480679 | 420,5 | MTRNR2L8 |
| | 562.643657897886 | 15 | -3.71831362974904 | MT-CO1 | 5.6914995 | 412 | MT-CO1 |
| | 102.10426438393 | 15 | -4.45654245307143 | MT-ND6 | 2.0576293 | 345,5 | MT-ND6 |
| | 87.1630884462679 | 15 | -3.59481814482255 | MT-CYB | 1.0694563 | 342 | MT-CYB |
| | 87.1630884462679 | 15 | -3.62009669824603 | MT-ND1 | 8.2505500 | 342 | MT-ND1 |
| | 87.1630884462679 | 15 | -3.6777679318453 | MT-ND2 | 5.4667654 | 342 | MT-ND2 |
| | 73.5951873878428 | 15 | -3.2954208523675 | MT-CO3 | 1.5922116 | 338 | MT-CO3 |
| | 34.1924380283781 | 13 | -3.97606227526756 | MT-ND4L | 1.3634941 | 303,5 | MT-ND4L |
| | 34.1924380283781 | 13 | -4.05064752476071 | MT-ND5 | 1.0289311 | 303,5 | MT-ND5 |
| | 57.82283938919 | 6 | -1.2548816622772 | SLC6A13 | 0.0075489 | 244,75 | SLC6A13 |
| | 0.313593352172977 | 11 | -1.17947081008856 | NDUFA4L2 | 0.0029788 | 237 | NDUFA4L2 |
| | 7.41933400355687 | 6 | -1.86437540815578 | EBF1 | 0.0021759 | 202,75 | EBF1 |
| | 0.0 | 9 | -3.38231112430034 | MT-ATP8 | 3.2847614 | 183,5 | MT-ATP8 |
| 6 | 0.0 | 1 | -2.25759817556733 | SLC11A1 | 0.0014261 | 49,75 | SLC11A1 |
| | 0.0 | 1 | -2.19088102004539 | FCGBP | 0.0051876 | 49,75 | FCGBP |
| 7 | 0.0 | 1 | -3.86573162848575 | GOLGA8F | 0.0076406 | 49,75 | GOLGA8F |
| | 1.0 | 2 | -4.81401309347573 | TMPRSS4 | 0.0079161 | 119 | TMPRSS4 |
| | 0.0 | 1 | -4.5695904011861 | ZAN | 0.0025573 | 49,75 | ZAN |
| 8 | 0.0 | 1 | -1.27461921103092 | FOS | 0.0074473 | 49,75 | FOS |
| | 1.0 | 2 | -1.2510108411128 | JUNB | 0.0057802 | 119 | JUNB |
| | 0.0 | 1 | -1.51720026963311 | PDLIM1 | 0.0041400 | 49,75 | PDLIM1 |
| 9 | 0.0 | 1 | -1.77268863715784 | C4A | 0.0048037 | 49,75 | C4A |
| | 0.0 | 1 | -1.77190368246383 | C4B | 0.0049083 | 49,75 | C4B |
| 10 | 1371.11084648195 | 6 | -1.49446434004589 | EMP3 | 0.0069821 | 403,75 | EMP3 |
| | 0.0 | 2 | -2.0362322309025 | SRPX2 | 0.0020926 | 84,5 | SRPX2 |
| | 0.0 | 2 | -2.37317412238245 | SYTL4 | 8.6821682 | 84,5 | SYTL4 |
| 11 | 0.0 | 1 | -1.42508786371628 | CCDC103 | 0.0064023 | 49,75 | CCDC103 |
| | 0.0 | 1 | -1.43242702903811 | FAM187A | 0.0066247 | 49,75 | FAM187A |
| 12 | 0.0 | 1 | -1.54865416094747 | CCDC102A | 0.0083193 | 49,75 | CCDC102A |
| | 0.0 | 1 | -2.65464189604975 | FOXJ1 | 5.4809478 | 49,75 | FOXJ1 |
| 13 | 686.0 | 2 | -1.45073206986754 | BTN2A2 | 0.0012155 | 278 | BTN2A2 |
| | 0.0 | 1 | -1.16337369015397 | SPATA1 | 5.4633186 | 49,75 | SPATA1 |
| 14 | 0.0 | 1 | -3.56566395865448 | C2CD4B | 0.0019752 | 49,75 | C2CD4B |
| | 0.0 | 1 | -1.78696748251542 | IL32 | 0.0026638 | 49,75 | IL32 |
| 15 | 0.0 | 1 | -2.24652910706168 | CLEC1A | 0.0019385 | 49,75 | CLEC1A |
| | 0.0 | 1 | -1.22608595351249 | ITIH5 | 0.0044563 | 49,75 | ITIH5 |
| 16 | 0.0 | 1 | -1.00823249184325 | ALDH1A1 | 0.0030629 | 49,75 | ALDH1A1 |
| | 686.0 | 3 | -1.38263969140886 | MDFIC | 9.8350899 | 302 | MDFIC |
| 16 | 280.613974687857 | 10 | -2.14221399914819 | PLA2G5 | 5.4838582 | 340 | PLA2G5 |
| | 0.0 | 3 | -1.94537676017578 | MMD2 | 7.8561315 | 108,5 | MMD2 |
| 17 | 686.0 | 2 | -2.30929701100462 | MT1F | 9.9876776 | 278 | MT1F |
| | 0.0 | 1 | -2.30243472092841 | MT1G | 0.0033098 | 49,75 | MT1G |
| 18 | 0.0 | 1 | -1.59130701923226 | ID3 | 0.0096300 | 49,75 | ID3 |
| | 0.0 | 1 | -1.28711534104511 | NANOS1 | 0.0076290 | 49,75 | NANOS1 |
| 19 | 688.552380952381 | 4 | -1.98093824997082 | NEK7 | 3.3026633 | 322,5 | NEK7 |
| | 0.0 | 1 | -1.42945154387013 | DOCK1 | 9.5944070 | 49,75 | DOCK1 |
| | 0.0 | 2 | -1.22446592630776 | ERBIN | 0.0048297 | 84,5 | ERBIN |
| 20 | 0.0 | 1 | -3.38937420253889 | S100A8 | 5.8425105 | 49,75 | S100A8 |
| | 0.0 | 1 | -2.69064554098678 | S100A9 | 0.0069665 | 49,75 | S100A9 |
| 21 | 686.0 | 2 | -1.75456943373289 | TEAD3 | 0.0040184 | 278 | TEAD3 |
| | 0.0 | 1 | -1.74355933341358 | TEAD2 | 0.0054055 | 49,75 | TEAD2 |

Supplementary file 8: ClueGo functional annotation of the three main co-expression network clusters of downregulated genes.

| #Cluster | GOID | GOTerm | Ontology Source | Term PValue | Term PValu | Group PVa | Group PVa | GOLevels | GOGroups | % Associat | Nr. Genes | Associated Genes Found |
|---|---|---|---|---|---|---|---|---|---|---|---|---|
| | GO:0042063 | gliogenesis | GO_BiologicalProces | 2,69906E-08 | 0,022468 | 0,00014 | 0,000281 | [3, 4, 5, 6, | Group00 | 9,090909 | 6 | [LFNG, NOTCH2, SWAP70, TGFB1, TLR4, TNFSF13] |
| | GO:0042063 | gliogenesis | GO_BiologicalProces | 2,69906E-08 | 0,012694 | 7,47E-05 | 0,000224 | [4, 5, 6, 7] | Group01 | 10,16949 | 6 | [BMP7, BMPR1B, DDR2, GDPD2, GLI3, ID4] |
| | GO:2000179 | positive regulation | GO_BiologicalProces | 8,05515E-08 | 0,002311 | 1,24E-05 | 8,65E-05 | [7, 8] | Group02 | 5,13834 | 13 | [ANTXR1, DDR1, DDR2, ERBB2, FGFR3, FLT4, LYN, MAPKAPK2, MAPKAPK3, NEK6, NTRK2, PDGFRB, STK17B] |
| | GO:0022612 | gland morphogenes | GO_BiologicalProces | 8,46087E-08 | 0,044645 | 0,00036 | 0,00036 | [4, 5, 6, 7, | Group03 | 10,20408 | 5 | [CD81, FGFR3, LYN, PDGFRB, TGFB1] |
| | GO:0022612 | gland morphogenes | GO_BiologicalProces | 8,46087E-08 | 0,012924 | 0,00036 | 0,00036 | [4, 5, 6, 7, | Group03 | 13,51351 | 5 | [CD81, FGFR3, LYN, PDGFRB, TGFB1] |
| | GO:0010001 | glial cell differentiat | GO_BiologicalProces | 7,86248E-07 | 0,011699 | 6,84E-05 | 0,000274 | [6] | Group04 | 21,05263 | 4 | [AXL, MAPKAPK2, MAPKAPK3, SNX33] |
| | GO:0010001 | glial cell differentiat | GO_BiologicalProces | 7,86248E-07 | 0,015252 | 6,84E-05 | 0,000274 | [7] | Group04 | 37,5 | 3 | [MAPKAPK2, MAPKAPK3, SNX33] |
| | GO:0021910 | smoothened signali | GO_BiologicalProces | 2,44448E-06 | 0,003266 | 1,77E-05 | 0,000106 | [4, 5, 6, 7] | Group05 | 5,357143 | 12 | [DCHS1, DDR1, ECE1, GLI3, LRP10, LRP4, MAPKAPK2, PDGFRB, SDC4, SLC9A3R1, TGFB1, TGFB2] |
| | GO:0003179 | heart valve morpho | GO_BiologicalProces | 3,4195E-06 | 0,022497 | 1,77E-05 | 0,000106 | [3, 5, 6, 7, | Group05 | 5,076142 | 10 | [DCHS1, GLI3, LRP10, LRP4, MAPKAPK2, PDGFRB, SDC4, SLC9A3R1, TGFB1, TGFB2] |
| | GO:0070723 | response to cholest | GO_BiologicalProces | 5,34992E-06 | 0,047785 | 2,03E-05 | 0,000101 | [5, 6] | Group06 | 5,027933 | 9 | [AMOT, CNN3, EPB41L5, ITGB1, ITGB5, PDGFRB, S1PR1, SDC4, SORBS3] |
| | GO:0070723 | response to cholest | GO_BiologicalProces | 5,34992E-06 | 0,047659 | 2,03E-05 | 0,000101 | [4, 6, 7, 8] | Group06 | 5,633803 | 8 | [AMOT, EPS8, EZR, ITGB5, S1PR1, SDC4, SORBS3, SWAP70] |
| | GO:0070723 | response to cholest | GO_BiologicalProces | 5,34992E-06 | 0,001748 | 9,25E-06 | 7,4E-05 | [5, 6, 7, 8] | Group07 | 11,29032 | 7 | [ADCYAP1R1, ADGRG1, CSF1, GPR4, ITPKB, NOTCH2, PDGFRB] |
| | GO:0003170 | heart valve develop | GO_BiologicalProces | 6,60342E-06 | 0,007964 | 9,25E-06 | 7,4E-05 | [6, 7, 8, 9] | Group07 | 11,11111 | 6 | [ADGRG1, CSF1, GPR4, ITPKB, NOTCH2, PDGFRB] |
| | GO:0051057 | positive regulation | GO_BiologicalProces | 9,24817E-06 | 0,009487 | 1,43E-08 | 1,57E-07 | [4, 5, 6] | Group08 | 22,22222 | 4 | [DAG1, DDR2, TGFB1, TGFB2] |
| | GO:0048709 | oligodendrocyte dif | GO_BiologicalProces | 1,17444E-05 | 0,009487 | 1,43E-08 | 1,57E-07 | [5, 6] | Group08 | 22,22222 | 4 | [DDR1, DDR2, TGFB1, TGFB2] |
| | GO:0046777 | protein autophosph | GO_BiologicalProces | 1,23564E-05 | 0,001022 | 1,43E-08 | 1,57E-07 | [5, 6] | Group08 | 22,72727 | 5 | [DAG1, MLC1, SMO, SMOX, TGFB1] |
| | GO:0060411 | cardiac septum mor | GO_BiologicalProces | 1,26988E-05 | 0,037302 | 1,43E-08 | 1,57E-07 | [5, 6, 7, 8, | Group08 | 15,38461 | 4 | [CSF1, DDR1, LRP5, TGFB1] |
| | GO:0021782 | glial cell developme | GO_BiologicalProces | 1,27585E-05 | 0,004904 | 8,04E-07 | 7,24E-06 | [4, 5, 6] | Group09 | 5,583756 | 11 | [BMP7, CSF1, DAG1, DCHS1, DDR1, GLI3, LRP5, RXRA, SMO, SMOX, TGFB1] |
| | GO:0021782 | glial cell developme | GO_BiologicalProces | 1,27585E-05 | 0,009487 | 8,04E-07 | 7,24E-06 | [5, 6] | Group09 | 22,22222 | 4 | [DDR1, DDR2, TGFB1, TGFB2] |
| | GO:0043583 | ear development | GO_BiologicalProces | 1,77477E-05 | 1,65E-05 | 8,04E-07 | 7,24E-06 | [4, 5, 6, 7] | Group09 | 8,888889 | 12 | [BMP7, CSF1, DAG1, DDR1, GLI3, ID4, LRP5, MSN, RXRA, SLC9A3R1, TGFB1, TGFB2] |
| | GO:0071397 | cellular response to | GO_BiologicalProces | 1,85227E-05 | 0,004888 | 8,04E-07 | 7,24E-06 | [4, 5, 6, 7] | Group09 | 6,17284 | 10 | [BMP7, CSF1, DAG1, DCHS1, DDR1, GLI3, LRP5, SMO, SMOX, TGFB1] |
| | GO:0071397 | cellular response to | GO_BiologicalProces | 1,85227E-05 | 0,045575 | 8,04E-07 | 7,24E-06 | [5, 6, 7, 8] | Group09 | 7,894737 | 6 | [CSF1, DDR1, LRP5, SMO, SMOX, TGFB1] |
| | GO:0003206 | cardiac chamber mo | GO_BiologicalProces | 1,90663E-05 | 0,037302 | 8,04E-07 | 7,24E-06 | [5, 6, 7, 8, | Group09 | 15,38461 | 4 | [CSF1, DDR1, LRP5, TGFB1] |
| | GO:0042476 | odontogenesis | GO_BiologicalProces | 1,90663E-05 | 0,00762 | 1,01E-07 | 1,01E-06 | [5, 6] | Group10 | 23,52941 | 4 | [DAG1, MLC1, SMO, SMOX] |
| | GO:0048754 | branching morphog | GO_BiologicalProces | 2,70033E-05 | 0,001022 | 1,01E-07 | 1,01E-06 | [5, 6] | Group10 | 22,72727 | 5 | [DAG1, MLC1, SMO, SMOX, TGFB1] |
| | GO:0048754 | branching morphog | GO_BiologicalProces | 2,70033E-05 | 0,037001 | 1,01E-07 | 1,01E-06 | [3, 4, 5, 6, | Group10 | 5,882353 | 8 | [GLI3, GLIS2, KANK1, SMO, SMOX, TGFB1, TGFB2, TLR4] |
| | GO:0061138 | morphogenesis of a | GO_BiologicalProces | 2,72464E-05 | 0,00236 | 1,01E-07 | 1,01E-06 | [4, 5, 6, 8, | Group10 | 8,888889 | 8 | [DAG1, ID4, LAMB2, LYN, SMO, SMOX, TGFB1, TLR4] |
| | GO:0061138 | morphogenesis of a | GO_BiologicalProces | 2,72464E-05 | 5,32E-06 | 1,01E-07 | 1,01E-06 | [6, 7] | Group10 | 6,343284 | 17 | [ADGRG1, APCDD1, CDKN2C, CSF1, DAG1, ERBB2, GLI3, GPR37L1, ID4, LAMB2, LYN, NTRK2, SMO, SMOX, TGFB1, TGFB2, TLR4] |
| | GO:0009953 | dorsal/ventral patte | GO_BiologicalProces | 3,19135E-05 | 0,00339 | 1,01E-07 | 1,01E-06 | [6, 7] | Group10 | 28,57143 | 4 | [DAG1, MLC1, SMO, SMOX] |
| | GO:0036315 | cellular response to | GO_BiologicalProces | 4,28091E-05 | 0,000153 | 1,01E-07 | 1,01E-06 | [5, 7, 8] | Group10 | 6,598985 | 13 | [CDKN2C, DAG1, ERBB2, GLI3, GPR37L1, ID4, LAMB2, LYN, NTRK2, SMO, SMOX, TGFB1, TLR4] |
| | GO:0036315 | cellular response to | GO_BiologicalProces | 4,28091E-05 | 0,037299 | 1,01E-07 | 1,01E-06 | [5, 6, 7, 8, | Group10 | 8,219178 | 6 | [GPR37L1, ID4, LAMB2, SMO, SMOX, TLR4] |
| | GO:0060413 | atrial septum morpl | GO_BiologicalProces | 4,28091E-05 | 0,002208 | 1,01E-07 | 1,01E-06 | [5, 6, 7, 8, | Group10 | 8,988764 | 8 | [CDKN2C, DAG1, ERBB2, GLI3, ID4, LYN, NTRK2, TGFB1] |
| | GO:0046579 | positive regulation | GO_BiologicalProces | 4,49925E-05 | 0,048475 | 1,01E-07 | 1,01E-06 | [5, 6, 7, 8, | Group10 | 14,28571 | 4 | [LAMB2, SMO, SMOX, TLR4] |
| | GO:0001657 | ureteric bud develo | GO_BiologicalProces | 4,50649E-05 | 0,012924 | 5,23E-12 | 6,28E-11 | [3, 4, 5, 6] | Group11 | 13,51351 | 5 | [CSF1, EZR, GPAM, SMO, SMOX] |
| | GO:0072163 | mesonephric epithe | GO_BiologicalProces | 4,8174E-05 | 0,037001 | 5,23E-12 | 6,28E-11 | [3, 4, 5, 6] | Group11 | 5,882353 | 8 | [DCHS1, EPB41L5, GLI3, ITGA6, SMO, SMOX, TGFB1, TGFB2] |
| | GO:0072164 | mesonephric tubule | GO_BiologicalProces | 4,8174E-05 | 0,001255 | 5,23E-12 | 6,28E-11 | [3, 4, 5, 6, | Group11 | 15,38461 | 6 | [DCHS1, FGFRL1, NOTCH2, SMAD6, TGFB1, TGFB2] |
| | GO:1903055 | positive regulation | GO_BiologicalProces | 5,45231E-05 | 0,024068 | 5,23E-12 | 6,28E-11 | [3, 4, 5, 6, | Group11 | 5,555555 | 9 | [BMP7, FGFRL1, NOTCH2, RXRA, SMAD6, SMO, SMOX, TGFB1, TGFB2] |
| | GO:0010715 | regulation of extrac | GO_BiologicalProces | 5,45231E-05 | 0,022895 | 5,23E-12 | 6,28E-11 | [4, 6] | Group11 | 100 | 2 | [SMO, SMOX] |
| | GO:0010715 | regulation of extrac | GO_BiologicalProces | 5,45231E-05 | 0,004904 | 5,23E-12 | 6,28E-11 | [4, 5, 6] | Group11 | 5,583756 | 11 | [BMP7, CSF1, DAG1, DCHS1, DDR1, GLI3, LRP5, RXRA, SMO, SMOX, TGFB1] |
| | GO:1903055 | positive regulation | GO_BiologicalProces | 5,45231E-05 | 0,009487 | 5,23E-12 | 6,28E-11 | [4, 5, 6] | Group11 | 22,22222 | 4 | [DAG1, DDR2, TGFB1, TGFB2] |
| Cluster1 | GO:0001823 | mesonephros devel | GO_BiologicalProces | 6,65314E-05 | 1,58E-05 | 5,23E-12 | 6,28E-11 | [4, 5, 6] | Group11 | 17,02128 | 8 | [ADGRG1, GLI3, GNG5, GPR37L1, LYN, NES, SMO, SMOX] |
| | GO:0072028 | nephron morphoge | GO_BiologicalProces | 6,81357E-05 | 0,000657 | 5,23E-12 | 6,28E-11 | [3, 4, 5, 6, | Group11 | 17,14286 | 6 | [DCHS1, FGFRL1, NOTCH2, SMAD6, TGFB1, TGFB2] |
| | GO:0006907 | pinocytosis | GO_BiologicalProces | 6,84138E-05 | 0,00347 | 5,23E-12 | 6,28E-11 | [3, 4, 5, 6, | Group11 | 7,258065 | 9 | [BMP7, FGFRL1, NOTCH2, RXRA, SMAD6, SMO, SMOX, TGFB1, TGFB2] |
| | GO:0045669 | positive regulation | GO_BiologicalProces | 7,46719E-05 | 0,030016 | 5,23E-12 | 6,28E-11 | [3, 4, 5, 6, | Group11 | 7,070707 | 7 | [BMP7, FGFRL1, NOTCH2, SMAD6, SMO, SMOX, TGFB2] |
| | GO:0090218 | positive regulation | GO_BiologicalProces | 7,6475E-05 | 0,032643 | 5,23E-12 | 6,28E-11 | [4, 5, 6, 7] | Group11 | 8,450705 | 6 | [GLI3, GLIS2, GPR37L1, RFX4, SMO, SMOX] |
| | GO:0040018 | positive regulation | GO_BiologicalProces | 7,6475E-05 | 0,005713 | 5,23E-12 | 6,28E-11 | [5, 6] | Group11 | 7,843137 | 8 | [BMPR1B, GLI3, LRP4, LRP5, RFX4, SMAD6, SMO, SMOX] |
| | GO:0072006 | nephron developme | GO_BiologicalProces | 8,09235E-05 | 1,65E-05 | 5,23E-12 | 6,28E-11 | [4, 5, 6, 7] | Group11 | 8,888889 | 12 | [BMP7, CSF1, DAG1, DDR1, GLI3, ID4, LRP5, MSN, RXRA, SLC9A3R1, TGFB1, TGFB2] |
| | GO:0042475 | odontogenesis of d | GO_BiologicalProces | 9,17075E-05 | 0,00762 | 5,23E-12 | 6,28E-11 | [5, 6] | Group11 | 23,52941 | 4 | [DAG1, MLC1, SMO, SMOX] |
| | GO:0044351 | macropinocytosis | GO_BiologicalProces | 9,18793E-05 | 0,00347 | 5,23E-12 | 6,28E-11 | [4, 5, 6, 7] | Group11 | 7,258065 | 9 | [BMP7, CSF1, GLI3, ITGA6, LRP4, SMO, SMOX, TGFB1, TGFB2] |
| | GO:0061005 | cell differentiation i | GO_BiologicalProces | 9,88761E-05 | 0,004888 | 5,23E-12 | 6,28E-11 | [4, 5, 6, 7] | Group11 | 6,17284 | 10 | [BMP7, CSF1, DAG1, DCHS1, DDR1, GLI3, LRP5, SMO, SMOX, TGFB1] |
| | GO:0003283 | atrial septum devel | GO_BiologicalProces | 0,000103665 | 0,001022 | 5,23E-12 | 6,28E-11 | [5, 6] | Group11 | 22,72727 | 5 | [DAG1, MLC1, SMO, SMOX, TGFB1] |
| | GO:0072073 | kidney epithelium d | GO_BiologicalProces | 0,000104285 | 0,013595 | 5,23E-12 | 6,28E-11 | [3, 4, 5, 6, | Group11 | 6,040268 | 9 | [BMP7, DCHS1, GLI3, GPR4, LAMB2, PDGFRB, SMO, SMOX, TGFB1] |
| | GO:0021940 | positive regulation | GO_BiologicalProces | 0,000136597 | 0,037001 | 5,23E-12 | 6,28E-11 | [3, 4, 5, 6, | Group11 | 5,882353 | 8 | [GLI3, GLIS2, KANK1, SMO, SMOX, TGFB1, TGFB2, TLR4] |
| | GO:0048839 | inner ear developm | GO_BiologicalProces | 0,000139734 | 0,01151 | 5,23E-12 | 6,28E-11 | [4, 5, 6, 7, | Group11 | 7,079646 | 8 | [BMP7, DCHS1, GLI3, SDC4, SMAD6, SMO, SMOX, TGFB1] |
| | GO:0002312 | B cell activation inv | GO_BiologicalProces | 0,000140426 | 0,022895 | 5,23E-12 | 6,28E-11 | [4, 5, 6, 7, | Group11 | 100 | 2 | [SMO, SMOX] |
| | GO:0060248 | detection of cell de | GO_BiologicalProces | 0,000143991 | 0,037302 | 5,23E-12 | 6,28E-11 | [4, 5, 6, 7, | Group11 | 15,38461 | 4 | [GLI3, GLIS2, GPR37L1, RFX4] |
| | GO:0007371 | ventral midline dete | GO_BiologicalProces | 0,000143991 | 0,002362 | 5,23E-12 | 6,28E-11 | [3, 4, 5, 6, | Group11 | 10,76923 | 7 | [BMP7, FGFRL1, NOTCH2, SMAD6, SMO, SMOX, TGFB2] |
| | GO:0072036 | mesenchymal to ep | GO_BiologicalProces | 0,000143991 | 0,030016 | 5,23E-12 | 6,28E-11 | [4, 5, 6, 7, | Group11 | 7,070707 | 7 | [BMP7, DCHS1, GLI3, PDGFRB, SMO, SMOX, TGFB1] |
| | GO:0072285 | mesenchymal to ep | GO_BiologicalProces | 0,000143991 | 0,016315 | 5,23E-12 | 6,28E-11 | [4, 5, 6, 7, | Group11 | 9,67742 | 6 | [GLI3, GLIS2, GPR4, LAMB2, SMO, SMOX] |
| | GO:0021513 | spinal cord dorsal/v | GO_BiologicalProces | 0,000150511 | 0,011719 | 5,23E-12 | 6,28E-11 | [3, 4, 5, 6, | Group11 | 8,333333 | 7 | [BMP7, DCHS1, GLI3, PDGFRB, SMO, SMOX, TGFB1] |
| | GO:0003205 | cardiac chamber de | GO_BiologicalProces | 0,000153301 | 0,016998 | 5,23E-12 | 6,28E-11 | [4, 5, 6, 7, | Group11 | 5,844156 | 9 | [BMP7, DCHS1, GLI3, LAMB2, SDC4, SMAD6, SMO, SMOX, TGFB1] |
| | GO:0003279 | cardiac septum dev | GO_BiologicalProces | 0,000192409 | 0,00236 | 5,23E-12 | 6,28E-11 | [4, 5, 6, 8, | Group11 | 8,888889 | 8 | [DAG1, ID4, LAMB2, LYN, SMO, SMOX, TGFB1, TLR4] |
| | GO:0060993 | kidney morphogene | GO_BiologicalProces | 0,000192409 | 0,017001 | 5,23E-12 | 6,28E-11 | [4, 5, 6, 7, | Group11 | 19,04762 | 4 | [NOTCH2, SMO, SMOX, TGFB2] |
| | GO:0008589 | regulation of smoot | GO_BiologicalProces | 0,0002106 | 0,032494 | 5,23E-12 | 6,28E-11 | [5, 6, 7, 8] | Group11 | 16 | 4 | [GLI3, RFX4, SMO, SMOX] |
| | GO:0021511 | spinal cord patterni | GO_BiologicalProces | 0,000211003 | 5,32E-06 | 5,23E-12 | 6,28E-11 | [6, 7] | Group11 | 6,343284 | 17 | [ADGRG1, APCDD1, CDKN2C, CSF1, DAG1, ERBB2, GLI3, GPR37L1, ID4, LAMB2, LYN, NTRK2, SMO, SMOX, TGFB1, TGFB2, TLR4] |
| | GO:1900182 | positive regulation | GO_BiologicalProces | 0,000241836 | 0,015315 | 5,23E-12 | 6,28E-11 | [5, 6, 7, 8] | Group11 | 7,954545 | 7 | [BMP7, CSF1, GLI3, ITGA6, LRP4, SMO, SMOX] |
| | GO:0048565 | digestive tract deve | GO_BiologicalProces | 0,000241836 | 0,045575 | 5,23E-12 | 6,28E-11 | [5, 6, 7, 8] | Group11 | 7,894737 | 6 | [CSF1, DDR1, LRP5, SMO, SMOX, TGFB1] |
| | GO:1900182 | positive regulation | GO_BiologicalProces | 0,000241836 | 0,00339 | 5,23E-12 | 6,28E-11 | [6, 7] | Group11 | 28,57143 | 4 | [DAG1, MLC1, SMO, SMOX] |
| | GO:0048708 | astrocyte differenti | GO_BiologicalProces | 0,000245387 | 0,000153 | 5,23E-12 | 6,28E-11 | [5, 7, 8] | Group11 | 6,598985 | 13 | [CDKN2C, DAG1, ERBB2, GLI3, GPR37L1, ID4, LAMB2, LYN, NTRK2, SMO, SMOX, TGFB1, TLR4] |
| | GO:0048708 | astrocyte differenti | GO_BiologicalProces | 0,000245387 | 0,037299 | 5,23E-12 | 6,28E-11 | [5, 6, 7, 8, | Group11 | 8,219178 | 6 | [GPR37L1, ID4, LAMB2, SMO, SMOX, TLR4] |
| | GO:0060444 | branching involved | GO_BiologicalProces | 0,000247031 | 0,00762 | 5,23E-12 | 6,28E-11 | [4, 5, 6, 7, | Group11 | 23,52941 | 4 | [NOTCH2, SMO, SMOX, TGFB2] |
| | GO:0060444 | branching involved | GO_BiologicalProces | 0,000247031 | 0,00843 | 5,23E-12 | 6,28E-11 | [5, 6, 7, 8, | Group11 | 7,407407 | 8 | [BMP7, DCHS1, GLI3, SDC4, SMAD6, SMO, SMOX, TGFB1] |
| | GO:0045879 | negative regulation | GO_BiologicalProces | 0,000247031 | 0,00843 | 5,23E-12 | 6,28E-11 | [4, 5, 6, 7, | Group11 | 7,407407 | 8 | [BMP7, DCHS1, GLI3, SDC4, SMAD6, SMO, SMOX, TGFB1] |
| | GO:0060444 | branching involved | GO_BiologicalProces | 0,000247031 | 0,048475 | 5,23E-12 | 6,28E-11 | [5, 6, 7, 8, | Group11 | 14,28571 | 4 | [LAMB2, SMO, SMOX, TLR4] |
| | GO:1903727 | positive regulation | GO_BiologicalProces | 0,000297632 | 0,023781 | 5,23E-12 | 6,28E-11 | [6, 7, 8, 9] | Group11 | 17,3913 | 4 | [GLI3, RFX4, SMO, SMOX] |
| | GO:0061180 | mammary gland epi | GO_BiologicalProces | 0,000305874 | 0,000472 | 5,23E-12 | 6,28E-11 | [5, 6, 7, 8, | Group11 | 44,44444 | 4 | [GLI3, RFX4, SMO, SMOX] |
| | GO:0061180 | mammary gland epi | GO_BiologicalProces | 0,000305874 | 0,037302 | 5,23E-12 | 6,28E-11 | [5, 6, 7, 8, | Group11 | 15,38461 | 4 | [CSF1, DDR1, LRP5, TGFB1] |
| | GO:0031032 | actomyosin structu | GO_BiologicalProces | 0,00032287 | 0,007931 | 5,23E-12 | 6,28E-11 | [5, 6, 7, 8, | Group11 | 7,476635 | 8 | [BMP7, DCHS1, GLI3, SDC4, SMAD6, SMO, SMOX, TGFB1] |
| | GO:0051017 | actin filament bund | GO_BiologicalProces | 0,000324212 | 0,022129 | 5,23E-12 | 6,28E-11 | [4, 5, 6, 7, | Group11 | 33,33333 | 3 | [GPR37L1, SMO, SMOX] |
| | GO:0014002 | astrocyte developm | GO_BiologicalProces | 0,000332018 | 0,022895 | 5,23E-12 | 6,28E-11 | [4, 5, 6, 7, | Group11 | 100 | 2 | [SMO, SMOX] |
| | GO:0014002 | astrocyte developm | GO_BiologicalProces | 0,000332018 | 0,022895 | 5,23E-12 | 6,28E-11 | [5, 6, 7, 8, | Group11 | 100 | 2 | [SMO, SMOX] |
| | GO:0030856 | regulation of epithe | GO_BiologicalProces | 2,91774E-06 | 0,00028 | 2,92E-06 | 1,75E-05 | [4, 5, 6] | Group0 | 7,751938 | 10 | [ASCL1, ATOH8, CD109, PAX6, PRKX, ROCK1, TNFRSF1A, YAP1, ZEB2, ZFP36L1] |
| | GO:0045636 | positive regulation | GO_BiologicalProces | 1,56724E-05 | 0,004209 | 4,63E-05 | 0,000139 | [5, 6] | Group1 | 8,641975 | 7 | [BST2, IFI35, IRF1, IRF7, MAVS, PSMB8, XAF1] |
| | GO:0032648 | regulation of interfe | GO_BiologicalProces | 1,74934E-05 | 0,004209 | 4,63E-05 | 0,000139 | [5, 6, 7] | Group1 | 8,641975 | 7 | [BST2, IFI35, IRF1, IRF7, MAVS, PSMB8, XAF1] |
| | GO:0045765 | regulation of angiog | GO_BiologicalProces | 2,24477E-05 | 0,012071 | 0,000137 | 0,000137 | [5, 6] | Group2 | 100 | 2 | [ALDH7A1, CHDH] |
| | GO:0032728 | positive regulation | GO_BiologicalProces | 2,81466E-05 | 0,012071 | 0,000137 | 0,000137 | [5, 6, 7] | Group2 | 100 | 2 | [ALDH7A1, CHDH] |
| | GO:0071357 | cellular response to | GO_BiologicalProces | 4,62544E-05 | 0,001644 | 1,75E-05 | 8,75E-05 | [5, 6] | Group3 | 12,76596 | 6 | [IRF1, IRF7, MAVS, TRIM38, ZBTB20, ZC3HAV1] |
| Cluster2 | GO:0060337 | type I interferon sig | GO_BiologicalProces | 4,62544E-05 | 0,002589 | 1,75E-05 | 8,75E-05 | [5, 6, 7] | Group3 | 16,12903 | 5 | [IRF1, IRF7, MAVS, ZBTB20, ZC3HAV1] |
| | GO:0050942 | positive regulation | GO_BiologicalProces | 5,39068E-05 | 0,039138 | 4,28E-05 | 0,000171 | [3, 4, 5, 6, | Group4 | 5,228758 | 8 | [ANGPT2, FGF1, FLT1, HIPK2, HSPB1, PGF, RRAS, VEGFB] |
| | GO:0045634 | regulation of melan | GO_BiologicalProces | 5,39068E-05 | 0,002088 | 4,28E-05 | 0,000171 | [4, 5, 6, 7, | Group4 | 5,106383 | 12 | [ANGPT2, EMP2, FGF1, FLT1, FOXO4, HIPK2, HSPB1, PGF, ROCK1, RRAS, TCF7L2, VEGFB] |
| | GO:0050932 | regulation of pigme | GO_BiologicalProces | 0,000127146 | 0,016967 | 4,28E-05 | 0,000171 | [4, 5, 6, 7, | Group4 | 5,925926 | 8 | [ANGPT2, FGF1, FLT1, HIPK2, HSPB1, PGF, RRAS, VEGFB] |
| | GO:0031456 | glycine betaine bios | GO_BiologicalProces | 0,000137168 | 0,011316 | 0,000127 | 0,000254 | [3, 4, 5, 6] | Group5 | 33,33333 | 3 | [ADAMTS9, BCL2, ZEB2] |
| | GO:0019285 | glycine betaine bios | GO_BiologicalProces | 0,000137168 | 0,004852 | 0,000127 | 0,000254 | [3, 4, 5, 6, | Group5 | 42,85714 | 3 | [ADAMTS9, BCL2, ZEB2] |
| | GO:0045766 | positive regulation | GO_BiologicalProces | 0,000195021 | 0,004852 | 0,000127 | 0,000254 | [4, 5, 6, 7] | Group5 | 42,85714 | 3 | [ADAMTS9, BCL2, ZEB2] |
| | GO:1904018 | positive regulation | GO_BiologicalProces | 0,000455095 | 0,001489 | 0,000127 | 0,000254 | [4, 5, 6, 7, | Group5 | 60 | 3 | [ADAMTS9, BCL2, ZEB2] |
| | GO:0045446 | endothelial cell diffe | GO_BiologicalProces | 1,16896E-07 | 1,27E-05 | 1,52E-10 | 1,37E-09 | [6] | Group09 | 9,090909 | 9 | [ACVRL1, APOLD1, CDH5, CLDN5, ENG, F11R, ICAM1, NOTCH4, PECAM1] |
| | GO:0002064 | epithelial cell devel | GO_BiologicalProces | 1,91191E-07 | 2,06E-05 | 1,52E-10 | 1,37E-09 | [4, 5, 6] | Group09 | 5,607477 | 12 | [ADAMTSL4, B4GALT1, CDH5, CDKN1A, CLDN5, ENG, EPHA2, F11R, ICAM1, NOTCH4, PECAM1, PODXL] |
| | GO:0001885 | endothelial cell dev | GO_BiologicalProces | 4,45493E-07 | 4,77E-05 | 1,52E-10 | 1,37E-09 | [5, 6, 7] | Group09 | 12,06897 | 7 | [CDH5, CLDN5, ENG, F11R, ICAM1, NOTCH4, PECAM1] |
| | GO:0031348 | negative regulation | GO_BiologicalProces | 1,02695E-06 | 0,000109 | 9,09E-08 | 6,36E-07 | [3, 4, 5, 6] | Group07 | 6,060606 | 10 | [FOXF1, GPER1, HLA-E, IL1R2, IRAK3, MYO1C, NLRC5, SOCS3, TNFRSF1B, ZFP36] |
| | GO:0030856 | regulation of epithe | GO_BiologicalProces | 1,1251E-06 | 0,000118 | 1,52E-10 | 1,37E-09 | [4, 5, 6] | Group09 | 6,976744 | 9 | [ACVRL1, APOLD1, CDH5, CLDN5, MAFF, NOTCH4, PRKCH, WWTR1, ZFP36] |
| | GO:0072132 | mesenchyme morpl | GO_BiologicalProces | 2,80575E-06 | 0,000292 | 1,07E-07 | 6,39E-07 | [4, 5, 6, 7] | Group11 | 12,2449 | 6 | [ACVRL1, ENG, FOXF1, HEYL, LEF1, TGFBR2] |
| | GO:0045601 | regulation of endotl | GO_BiologicalProces | 4,10575E-06 | 0,000423 | 1,52E-10 | 1,37E-09 | [5, 6, 7] | Group09 | 16,66667 | 5 | [ACVRL1, APOLD1, CDH5, CLDN5, NOTCH4] |
| | GO:0071357 | cellular response to | GO_BiologicalProces | 4,40633E-06 | 0,000449 | 5,54E-07 | 2,77E-06 | [5, 6] | Group10 | 8,641975 | 7 | [HLA-E, IFI27, IFITM1, IFITM2, IFITM3, NLRC5, SP100] |
| | GO:0060337 | type I interferon sig | GO_BiologicalProces | 4,40633E-06 | 0,000449 | 5,54E-07 | 2,77E-06 | [5, 6, 7] | Group10 | 8,641975 | 7 | [HLA-E, IFI27, IFITM1, IFITM2, IFITM3, NLRC5, SP100] |
| | GO:0001569 | branching involved | GO_BiologicalProces | 4,86326E-06 | 0,000491 | 6,6E-09 | 5,28E-08 | [4, 5, 6, 7, | Group08 | 16,12903 | 5 | [DLL4, ENG, LEF1, NOTCH4, TGFBR2] |
| | GO:0001569 | branching involved | GO_BiologicalProces | 4,86326E-06 | 0,000491 | 1,07E-07 | 6,39E-07 | [4, 5, 6, 7, | Group11 | 16,12903 | 5 | [DLL4, ENG, LEF1, NOTCH4, TGFBR2] |
| | GO:0061138 | morphogenesis of a | GO_BiologicalProces | 5,07155E-06 | 0,000507 | 1,07E-07 | 6,39E-07 | [4, 5, 6] | Group11 | 5,076142 | 10 | [B4GALT1, DLL4, ENG, EPHA2, FOXF1, LEF1, NOTCH4, SOCS3, TGFBR2, TGM2] |
| | GO:0046718 | viral entry into host | GO_BiologicalProces | 5,60265E-06 | 0,000555 | 5,54E-07 | 2,77E-06 | [5, 6, 7, 8] | Group10 | 6,779661 | 8 | [F11R, ICAM1, IFITM1, IFITM2, IFITM3, ITGA5, NECTIN2, TRIM5] |
| | GO:0046596 | regulation of viral e | GO_BiologicalProces | 7,80896E-06 | 0,000765 | 5,54E-07 | 2,77E-06 | [5, 6, 7, 8, | Group10 | 14,70588 | 5 | [IFITM1, IFITM2, IFITM3, NECTIN2, TRIM5] |
| | GO:0060333 | interferon-gamma- | GO_BiologicalProces | 1,03025E-05 | 0,000999 | 9,09E-08 | 6,36E-07 | [5, 6, 7] | Group07 | 7,608696 | 7 | [HLA-E, ICAM1, MYO1C, NLRC5, SOCS3, SP100, TRIM5] |
| | GO:0010718 | positive regulation | GO_BiologicalProces | 1,19944E-05 | 0,001151 | 1,07E-07 | 6,39E-07 | [3, 4, 5, 6, | Group11 | 13,51351 | 5 | [ENG, LEF1, TGFB1I1, TGFBR2, WWTR1] |
| | GO:0003197 | endocardial cushion | GO_BiologicalProces | 1,56363E-05 | 0,001485 | 6,6E-09 | 5,28E-08 | [4, 5, 6, 7] | Group08 | 12,82051 | 5 | [ACVRL1, ENG, FOXF1, HEYL, TGFBR2] |
| | GO:0003197 | endocardial cushion | GO_BiologicalProces | 1,56363E-05 | 0,001485 | 1,07E-07 | 6,39E-07 | [4, 5, 6, 7] | Group11 | 12,82051 | 5 | [ACVRL1, ENG, FOXF1, HEYL, TGFBR2] |
| | GO:0044409 | entry into host | GO_BiologicalProces | 1,59339E-05 | 0,001498 | 5,54E-07 | 2,77E-06 | [5, 6] | Group10 | 5,882353 | 8 | [F11R, ICAM1, IFITM1, IFITM2, IFITM3, ITGA5, NECTIN2, TRIM5] |
| | GO:0051806 | entry into cell of ot | GO_BiologicalProces | 1,59339E-05 | 0,001498 | 5,54E-07 | 2,77E-06 | [5, 6] | Group10 | 5,882353 | 8 | [F11R, ICAM1, IFITM1, IFITM2, IFITM3, ITGA5, NECTIN2, TRIM5] |
| | GO:0030260 | entry into host cell | GO_BiologicalProces | 1,59339E-05 | 0,001498 | 5,54E-07 | 2,77E-06 | [6, 7] | Group10 | 5,882353 | 8 | [F11R, ICAM1, IFITM1, IFITM2, IFITM3, ITGA5, NECTIN2, TRIM5] |
| | GO:0061028 | establishment of en | GO_BiologicalProces | 1,77532E-05 | 0,001651 | 1,52E-10 | 1,37E-09 | [6, 7, 8] | Group09 | 12,5 | 5 | [CDH5, CLDN5, ENG, F11R, ICAM1] |
| | GO:0046597 | negative regulation | GO_BiologicalProces | 1,87098E-05 | 0,001721 | 5,54E-07 | 2,77E-06 | [5, 6, 7, 8, | Group10 | 20 | 4 | [IFITM1, IFITM2, IFITM3, TRIM5] |
| | GO:0035912 | dorsal aorta morph | GO_BiologicalProces | 2,90395E-05 | 0,002643 | 6,6E-09 | 5,28E-08 | [6, 7, 9, 10, | Group08 | 37,5 | 3 | [ACVRL1, DLL4, ENG] |
| | GO:0035912 | dorsal aorta morph | GO_BiologicalProces | 2,90395E-05 | 0,002643 | 1,07E-07 | 6,39E-07 | [6, 7, 9, 10, | Group11 | 37,5 | 3 | [ACVRL1, DLL4, ENG] |
| | GO:0002040 | sprouting angiogen | GO_BiologicalProces | 2,90834E-05 | 0,002618 | 6,6E-09 | 5,28E-08 | [4, 5, 6, 8, | Group08 | 8,219178 | 6 | [ACVRL1, DLL4, EPHA2, ITGA5, LEF1, MMRN2] |
| | GO:0003208 | cardiac ventricle mo | GO_BiologicalProces | 2,90834E-05 | 0,002618 | 1,07E-07 | 6,39E-07 | [4, 5, 6, 7, | Group11 | 8,219178 | 6 | [DLL4, ENG, FOXF1, HEYL, PPP1R13L, TGFBR2] |
| | GO:0035907 | dorsal aorta develo | GO_BiologicalProces | 4,32983E-05 | 0,003854 | 6,6E-09 | 5,28E-08 | [6, 8, 9, 10, | Group08 | 33,33333 | 3 | [ACVRL1, DLL4, ENG] |
| | GO:0035907 | dorsal aorta develo | GO_BiologicalProces | 4,32983E-05 | 0,003854 | 1,07E-07 | 6,39E-07 | [6, 8, 9, 10, | Group11 | 33,33333 | 3 | [ACVRL1, DLL4, ENG] |
| Cluster3 | GO:1903901 | negative regulation | GO_BiologicalProces | 8,37541E-05 | 0,00737 | 5,54E-07 | 2,77E-06 | [4, 5, 6, 7] | Group10 | 6,818182 | 6 | [IFITM1, IFITM2, IFITM3, SP100, TRIM5, ZFP36] |
| | GO:0003203 | endocardial cushion | GO_BiologicalProces | 8,66261E-05 | 0,007536 | 6,6E-09 | 5,28E-08 | [5, 6, 7, 8] | Group08 | 13,7931 | 4 | [ACVRL1, ENG, HEYL, TGFBR2] |
| | GO:0003203 | endocardial cushion | GO_BiologicalProces | 8,66261E-05 | 0,007536 | 1,07E-07 | 6,39E-07 | [5, 6, 7, 8] | Group11 | 13,7931 | 4 | [ACVRL1, ENG, HEYL, TGFBR2] |
| | GO:0002043 | blood vessel endoth | GO_BiologicalProces | 0,000111376 | 0,009578 | 6,6E-09 | 5,28E-08 | [5, 6, 7, 9, | Group08 | 25 | 3 | [ACVRL1, DLL4, EPHA2] |
| | GO:0032835 | glomerulus develop | GO_BiologicalProces | 0,000218345 | 0,018559 | 0,000218 | 0,000873 | [3, 4, 5, 6, | Group00 | 7,462687 | 5 | [CFLAR, HEYL, PECAM1, PODXL, WWTR1] |
| | GO:0033005 | positive regulation | GO_BiologicalProces | 0,000276784 | 0,02325 | 0,000342 | 0,001027 | [4, 5, 6, 7, | Group05 | 18,75 | 3 | [FGR, IL4R, NECTIN2] |
| | GO:0060841 | venous blood vesse | GO_BiologicalProces | 0,000276784 | 0,02325 | 6,6E-09 | 5,28E-08 | [4, 6, 7, 8, | Group08 | 18,75 | 3 | [ACVRL1, ENG, FOXF1] |
| | GO:0045603 | positive regulation | GO_BiologicalProces | 0,000276784 | 0,02325 | 1,52E-10 | 1,37E-09 | [5, 6, 7, 8] | Group09 | 18,75 | 3 | [ACVRL1, CDH5, CLDN5] |
| | GO:0060841 | venous blood vesse | GO_BiologicalProces | 0,000276784 | 0,02325 | 1,07E-07 | 6,39E-07 | [4, 6, 7, 8, | Group11 | 18,75 | 3 | [ACVRL1, ENG, FOXF1] |
| | GO:0048525 | negative regulation | GO_BiologicalProces | 0,000300161 | 0,024913 | 5,54E-07 | 2,77E-06 | [3, 4, 5, 6] | Group10 | 5,405406 | 6 | [IFITM1, IFITM2, IFITM3, SP100, TRIM5, ZFP36] |
| | GO:0010717 | regulation of epithe | GO_BiologicalProces | 0,000306165 | 0,025106 | 1,07E-07 | 6,39E-07 | [4, 5, 6, 7, | Group11 | 6,944445 | 5 | [ENG, LEF1, TGFB1I1, TGFBR2, WWTR1] |
| | GO:0003198 | epithelial to mesen | GO_BiologicalProces | 0,000334088 | 0,027061 | 1,07E-07 | 6,39E-07 | [4, 5, 6, 7, | Group11 | 17,64706 | 3 | [ENG, HEYL, TGFBR2] |
| | GO:0033003 | regulation of mast c | GO_BiologicalProces | 0,000342335 | 0,027387 | 0,000342 | 0,001027 | [4, 5, 6, 7] | Group05 | 9,756098 | 4 | [FGR, FOXF1, IL4R, NECTIN2] |
| | GO:0045824 | negative regulation | GO_BiologicalProces | 0,000342335 | 0,027387 | 9,09E-08 | 6,36E-07 | [4, 5, 6, 7] | Group07 | 9,756098 | 4 | [HLA-E, IRAK3, MYO1C, NLRC5] |
| | GO:0050728 | negative regulation | GO_BiologicalProces | 0,000346549 | 0,027377 | 9,09E-08 | 6,36E-07 | [4, 5, 6, 7] | Group07 | 5,263158 | 6 | [FOXF1, GPER1, IL1R2, SOCS3, TNFRSF1B, ZFP36] |
| | GO:0010596 | negative regulation | GO_BiologicalProces | 0,000375985 | 0,029327 | 6,6E-09 | 5,28E-08 | [4, 5, 6, 7, | Group08 | 9,523809 | 4 | [ACVRL1, DLL4, MMRN2, SP100] |
| | GO:0001837 | epithelial to mesen | GO_BiologicalProces | 0,000416929 | 0,032103 | 1,07E-07 | 6,39E-07 | [6, 7, 8] | Group11 | 5,084746 | 6 | [ENG, HEYL, LEF1, TGFB1I1, TGFBR2, WWTR1] |

| | | | | | | | | | | | |
|---|---|---|---|---|---|---|---|---|---|---|---|
| GO:0001570 | vasculogenesis | GO_BiologicalProces | 0,000444221 | 0,033761 | 1,07E-07 | 6,39E-07 | [4, 5, 7, 8, 9 | Group11 | 6,410256 | 5 | [ENG, EPHA2, FOXF1, TGFBR2, TIPARP] |
| GO:0060438 | trachea developme | GO_BiologicalProces | 0,000470409 | 0,035281 | 1,07E-07 | 6,39E-07 | [3, 4, 5, 6] | Group11 | 15,78947 | 3 | [FOXF1, LEF1, TGFBR2] |
| GO:0070498 | interleukin-1-media | GO_BiologicalProces | 0,00055012 | 0,040709 | 0,00055 | 0,0011 | [5, 6, 7] | Group03 | 15 | 3 | [IL1R1, IL1R2, IRAK3] |
| GO:1901739 | regulation of myobl | GO_BiologicalProces | 0,00055012 | 0,040709 | 0,00055 | 0,0011 | [5, 6, 7, 8, 9 | Group04 | 15 | 3 | [CFLAR, EHD2, IL4R] |
| GO:1901552 | positive regulation o | GO_BiologicalProces | 0,000652558 | 0,047637 | 1,52E-10 | 1,37E-09 | [5, 6, 7, 8, 9 | Group09 | 40 | 2 | [CDH5, CLDN5] |
| GO:1903142 | positive regulation o | GO_BiologicalProces | 0,000652558 | 0,047637 | 1,52E-10 | 1,37E-09 | [6, 7, 8, 9, 1 | Group09 | 40 | 2 | [CDH5, CLDN5] |
| GO:1905007 | positive regulation o | GO_BiologicalProces | 0,000652558 | 0,047637 | 1,07E-07 | 6,39E-07 | [4, 5, 6, 7, 8 | Group11 | 40 | 2 | [ENG, TGFBR2] |
| GO:0050701 | interleukin-1 secret | GO_BiologicalProces | 0,042256366 | 0,042256 | 0,042256 | 0,042256 | [5, 6, 7, 8, 9 | Group01 | 5 | 2 | [GSDMD, IL1R2] |
| GO:0051898 | negative regulation | GO_BiologicalProces | 0,042256366 | 0,042256 | 0,042256 | 0,042256 | [5, 6, 7, 8] | Group02 | 5 | 2 | [EPHA2, GPER1] |
| GO:0035890 | exit from host | GO_BiologicalProces | 0,042256366 | 0,042256 | 0,042256 | 0,042256 | [4, 5, 6] | Group06 | 5 | 2 | [PODXL, TRIM5] |
| GO:0019076 | viral release from ho | GO_BiologicalProces | 0,042256366 | 0,042256 | 0,042256 | 0,042256 | [4, 5, 6, 7, 8 | Group06 | 5 | 2 | [PODXL, TRIM5] |
| GO:0035891 | exit from host cell | GO_BiologicalProces | 0,042256366 | 0,042256 | 0,042256 | 0,042256 | [5, 6, 7] | Group06 | 5 | 2 | [PODXL, TRIM5] |
| GO:0046638 | positive regulation o | GO_BiologicalProces | 0,042256366 | 0,042256 | 1,07E-07 | 6,39E-07 | [7, 8, 9, 10, | Group11 | 5 | 2 | [IL4R, TGFBR2] |

Supplementary file 8: Co-expression network clusters of upregulated genes.

| __glayCluster | betweenness | degree | log2FoldChange | name | padj | Rank_stat | shared name |
|---|---|---|---|---|---|---|---|
| 1 | 308,5256301 | 39 | 1,485084433 | CALY | 0,002053 | 94 | CALY |
| | 116,6896206 | 33 | 1,338233206 | NRGN | 0,00963 | 85,75 | NRGN |
| | 138,6646908 | 25 | 1,145629356 | PGM2L1 | 0,007447 | 79,5 | PGM2L1 |
| | 183,4413388 | 14 | 1,010962143 | RASL10A | 0,005778 | 74,5 | RASL10A |
| | 108,9928756 | 18 | 1,324605327 | MPV17L2 | 4,64E-04 | 72,25 | MPV17L2 |
| | 63,44972888 | 29 | 1,024340961 | CAMKV | 0,023381 | 72 | CAMKV |
| | 138,9491715 | 14 | 1,065642072 | CACNG8 | 0,004104 | 71,5 | CACNG8 |
| | 90,14882997 | 18 | 1,097374598 | SERPINF1 | 0,049892 | 68,25 | SERPINF1 |
| | 30,62461267 | 25 | 1,191681998 | GABRD | 0,019507 | 62,5 | GABRD |
| | 50,83633913 | 20 | 1,055496609 | SCN3B | 0,0191 | 62 | SCN3B |
| | 27,51008342 | 22 | 1,428346639 | ABCC12 | 0,038467 | 60,25 | ABCC12 |
| | 25,48173032 | 22 | 1,004438853 | PSMG3 | 0,003558 | 59,25 | PSMG3 |
| | 12,73940792 | 17 | 1,009968861 | TCTE1 | 0,021259 | 52 | TCTE1 |
| | 11,45930804 | 14 | 1,002687653 | NECAB2 | 0,002134 | 49 | NECAB2 |
| | 89 | 2 | 3,010706157 | P2RX2 | 4,85E-04 | 43,25 | P2RX2 |
| | 2,135192582 | 11 | 1,124701501 | CALB2 | 0,004687 | 41,5 | CALB2 |
| | 1,26565106 | 12 | 1,478951637 | CORT | 0,001321 | 41,25 | CORT |
| | 3,390950962 | 10 | 1,308873708 | KRT17 | 0,019161 | 41 | KRT17 |
| | 4,533359536 | 8 | 1,120314104 | NEGR1 | 0,011588 | 39,25 | NEGR1 |
| | 1,326261208 | 9 | 1,11339938 | BEX5 | 0,019043 | 37,25 | BEX5 |
| | 1,408333333 | 8 | 1,144085695 | PLPPR3 | 0,010709 | 36,25 | PLPPR3 |
| | 0 | 5 | 1,031057586 | CITED1 | 0,046929 | 24,25 | CITED1 |
| | 0 | 1 | 1,07617123 | LY6H | 0,001993 | 9,5 | LY6H |
| | 0 | 1 | 1,97239753 | PROC | 0,01961 | 9,5 | PROC |
| | 0 | 1 | 1,051353191 | TGFBR3L | 0,002557 | 9,5 | TGFBR3L |
| 2 | 401,6324883 | 27 | 1,335836086 | SOHLH1 | 0,015748 | 87,75 | SOHLH1 |
| | 138,3551033 | 34 | 1,557307378 | CPNE9 | 0,003438 | 87,5 | CPNE9 |
| | 148,6512848 | 28 | 1,027618848 | MAST1 | 0,016262 | 84,25 | MAST1 |
| | 143,5842487 | 28 | 1,161673399 | GRIK1 | 0,001677 | 83,75 | GRIK1 |
| | 99,10299657 | 33 | 1,259139581 | SYT5 | 0,017918 | 82,75 | SYT5 |
| | 223,710142 | 21 | 1,050754419 | TRPM2 | 5,39E-04 | 81,25 | TRPM2 |
| | 219,0596195 | 20 | 1,199815203 | MYO15A | 0,002313 | 79,5 | MYO15A |
| | 99,82262735 | 27 | 1,491351478 | PNMA3 | 0,007421 | 77,75 | PNMA3 |
| | 85,2207987 | 31 | 1,053030273 | AP3B2 | 0,036148 | 77,25 | AP3B2 |
| | 86,29392858 | 30 | 1,036988626 | R3HDM1 | 0,036319 | 77 | R3HDM1 |
| | 80,29650161 | 32 | 1,059414629 | PNCK | 0,003123 | 76,5 | PNCK |
| | 132,0097555 | 21 | 1,098563359 | CACNA1G | 0,039941 | 75,75 | CACNA1G |
| | 81,85478506 | 29 | 1,061878554 | LPCAT4 | 0,029616 | 75 | LPCAT4 |
| | 72,05616844 | 31 | 1,44685866 | RPRML | 0,017772 | 74,75 | RPRML |
| | 57,10083047 | 29 | 1,301183717 | CCK | 0,045241 | 71 | CCK |
| | 102,835614 | 12 | 1,307663688 | AHRR | 0,023569 | 66,25 | AHRR |
| | 35,95998436 | 26 | 1,163252396 | RNF112 | 0,002901 | 64,75 | RNF112 |
| | 35,01907769 | 26 | 1,41957014 | PNMA5 | 0,017417 | 64,25 | PNMA5 |
| | 80,63958738 | 17 | 1,184361037 | BICDL1 | 0,022312 | 63,5 | BICDL1 |
| | 89,05101286 | 12 | 1,021152651 | CRHR1 | 0,030254 | 62,75 | CRHR1 |
| | 26,09263102 | 25 | 1,067915409 | DRP2 | 0,018017 | 61,5 | DRP2 |
| | 98,60957217 | 9 | 1,512524292 | CYP2E1 | 0,011809 | 59,75 | CYP2E1 |
| | 5,745156744 | 23 | 1,11873798 | ACTL6B | 0,031653 | 55,5 | ACTL6B |
| | 16,21344752 | 20 | 1,017080216 | PRMT8 | 0,016328 | 55,5 | PRMT8 |
| | 22,4503443 | 17 | 1,087586804 | RTP5 | 0,006858 | 54 | RTP5 |
| | 15,6929317 | 12 | 1,575138923 | TAC3 | 0,023231 | 48,75 | TAC3 |
| | 20,95401413 | 10 | 1,188375678 | GRM2 | 0,017403 | 46,5 | GRM2 |
| | 6,050188894 | 11 | 1,118565201 | DLK2 | 0,006803 | 45 | DLK2 |
| | 1,541111068 | 6 | 1,370631764 | ASTL | 0,0102 | 34,5 | ASTL |
| | 3,878491017 | 3 | 1,219991267 | SEMA5B | 0,006096 | 29,5 | SEMA5B |
| | 0,2 | 4 | 2,393310718 | ATP4A | 0,001975 | 27 | ATP4A |

| | | | | | | | |
|---|---|---|---|---|---|---|---|
| | 0,698981808 | 3 | 1,322598614 | PKD2L1 | 0,009756 | 25,5 | PKD2L1 |
| | 0,487179487 | 2 | 1,050234131 | TRIM17 | 0,010564 | 21,5 | TRIM17 |
| | 0 | 3 | 1,08545462 | PLCH2 | 0,021641 | 19,25 | PLCH2 |
| | 0 | 1 | 1,81980008 | CARTPT | 0,019466 | 9,5 | CARTPT |
| | 0 | 1 | 1,064892142 | SLC44A5 | 0,030153 | 9,5 | SLC44A5 |
| 3 | 575,6169904 | 15 | 2,279984492 | BX276092.! | 0,003222 | 79,5 | BX276092.9 |
| | 796,6497242 | 11 | 1,098006982 | ZFR2 | 0,002862 | 76 | ZFR2 |
| | 738,822918 | 10 | 1,330956163 | CHTF18 | 3,50E-04 | 74 | CHTF18 |
| | 405,2991378 | 8 | 1,638774655 | TMEM249 | 0,001195 | 69,75 | TMEM249 |
| | 432,4064197 | 7 | 1,362714826 | CORO6 | 0,01285 | 69 | CORO6 |
| | 241,5728936 | 5 | 1,246952995 | NOXA1 | 0,006364 | 64 | NOXA1 |
| | 161,8129687 | 6 | 1,526958221 | SYCE1 | 0,007301 | 63,5 | SYCE1 |
| | 154,9566368 | 5 | 1,577014259 | PDIA2 | 0,005638 | 61 | PDIA2 |
| | 176 | 4 | 1,043935189 | CCDC57 | 0,017168 | 59,5 | CCDC57 |
| | 108,183074 | 5 | 1,299282584 | MICAL1 | 0,003522 | 56,5 | MICAL1 |
| | 100,3638109 | 4 | 1,076935792 | CYP2D7 | 0,035293 | 53 | CYP2D7 |
| | 48,98861613 | 8 | 1,374419396 | GRIK2 | 6,82E-04 | 49,25 | GRIK2 |
| | 59,80581362 | 6 | 1,335690278 | TNNT2 | 0,00385 | 48,5 | TNNT2 |
| | 91,88484848 | 3 | 1,16564294 | B4GALNT3 | 0,01214 | 48,5 | B4GALNT3 |
| | 64,58403144 | 4 | 1,005194527 | ANKRD24 | 0,00131 | 45 | ANKRD24 |
| | 37,61640733 | 5 | 1,407594933 | WDR97 | 0,00246 | 44 | WDR97 |
| | 89 | 2 | 1,119569712 | ZGLP1 | 0,025358 | 43,25 | ZGLP1 |
| | 48,24647596 | 4 | 2,092324699 | TBC1D26 | 0,003222 | 42 | TBC1D26 |
| | 73,9 | 2 | 1,054550917 | PKD1 | 0,001195 | 40 | PKD1 |
| | 36,1273296 | 2 | 1,598961893 | LCN15 | 0,00634 | 35 | LCN15 |
| | 12,5 | 2 | 1,5806132 | TGM1 | 0,008343 | 29 | TGM1 |
| | 2,884848485 | 3 | 1,280311696 | TRIM54 | 0,008622 | 28,5 | TRIM54 |
| | 5,333333333 | 2 | 1,115674959 | RTEL1 | 0,001195 | 27 | RTEL1 |
| | 0 | 2 | 1,021312646 | CES4A | 0,017312 | 15,75 | CES4A |
| | 0 | 2 | 1,38734254 | NOXO1 | 5,34E-04 | 15,75 | NOXO1 |
| | 0 | 1 | 1,156042333 | CYP2D6 | 0,045242 | 9,5 | CYP2D6 |
| | 0 | 1 | 1,803926378 | LORICRIN | 0,008381 | 9,5 | LORICRIN |
| | 0 | 1 | 1,053060627 | PTCH2 | 0,005349 | 9,5 | PTCH2 |
| | 0 | 1 | 1,581402716 | TNFRSF25 | 0,006636 | 9,5 | TNFRSF25 |
| | 0 | 1 | 1,199218077 | DNASE1L2 | 0,005496 | 9,5 | DNASE1L2 |
| 4 | 0 | 1 | 1,26860296 | KIF12 | 0,00115 | 9,5 | KIF12 |
| | 0 | 1 | 1,492943739 | LTK | 7,41E-04 | 9,5 | LTK |
| 5 | 0 | 1 | 1,507033312 | RTL1 | 0,027174 | 9,5 | RTL1 |
| | 0 | 1 | 1,306970395 | TARBP1 | 0,011976 | 9,5 | TARBP1 |
| 6 | 0 | 1 | 1,042080575 | AGAP6 | 0,01563 | 9,5 | AGAP6 |
| | 0 | 1 | 1,056709194 | TIMM23B-/ | 0,013788 | 9,5 | TIMM23B-AGAP6 |

Supplementary file 8: ClueGo functional annotation of the three main co-expression network clusters of upregulated genes.

| #Cluster | ID | Term | Ontology Source | Term PValu | Term PValu | Group PVa | Group PVa | GOLevels | GOGroups | % Associat | Nr. Genes | Associated Genes Found |
|---|---|---|---|---|---|---|---|---|---|---|---|---|
| Cluster1 | GO:0099590 | neurotransmitter | GO_BiologicalProcess-EBI- | 0,00 | 0,00 | 0,00 | 0,00 | [3, 6, 7, 8] | Group0 | 6,90 | 2,00 | [CACNG8, CALY] |
| | GO:0019076 | viral release from | GO_BiologicalProcess-EBI- | 0,00 | 0,00 | 0,00 | 0,00 | [2, 3, 7] | Group1 | 5,13 | 2,00 | [KRT17, PROC] |
| | GO:0044794 | positive regulatior | GO_BiologicalProcess-EBI- | 0,00 | 0,00 | 0,00 | 0,00 | [2, 6] | Group1 | 11,76 | 2,00 | [KRT17, PROC] |
| | GO:1902186 | regulation of viral | GO_BiologicalProcess-EBI- | 0,00 | 0,00 | 0,00 | 0,00 | [3, 4, 5, 8] | Group1 | 5,88 | 2,00 | [KRT17, PROC] |
| | GO:0006768 | biotin metabolic p | GO_BiologicalProcess-EBI- | 0,00 | 0,00 | 0,00 | 0,00 | [4, 5, 7] | Group1 | 15,38 | 2,00 | [KRT17, PROC] |
| | GO:1902188 | positive regulatior | GO_BiologicalProcess-EBI- | 0,00 | 0,00 | 0,00 | 0,00 | [3, 4, 5, 6, | Group1 | 10,53 | 2,00 | [KRT17, PROC] |
| | GO:0035891 | exit from host cell | GO_BiologicalProcess-EBI- | 0,00 | 0,00 | 0,00 | 0,00 | [6] | Group1 | 5,13 | 2,00 | [KRT17, PROC] |
| | GO:0044789 | modulation by hos | GO_BiologicalProcess-EBI- | 0,00 | 0,00 | 0,00 | 0,00 | [4, 5, 6, 9] | Group1 | 28,57 | 2,00 | [KRT17, PROC] |
| | GO:0044791 | positive regulatior | GO_BiologicalProcess-EBI- | 0,00 | 0,00 | 0,00 | 0,00 | [3, 4, 5, 6, | Group1 | 28,57 | 2,00 | [KRT17, PROC] |
| Cluster2 | GO:0005272 | sodium channel ac | GO_MolecularFunction-GC | 0,00 | 0,01 | 0,00 | 0,01 | [3, 4, 5, 6, | Group0 | 4,44 | 2,00 | [CACNA1G, CCK] |
| | GO:0032096 | negative regulatio | GO_BiologicalProcess-GOA | 0,00 | 0,01 | 0,00 | 0,01 | [5, 6, 7] | Group1 | 4,26 | 2,00 | [PKD2L1, RTP5] |
| | GO:0032099 | negative regulatio | GO_BiologicalProcess-GOA | 0,00 | 0,00 | 0,00 | 0,00 | [7, 8, 9] | Group2 | 8,33 | 4,00 | [CACNA1G, GRIK1, PKD2L1, TRPM2] |
| | GO:0032105 | negative regulatio | GO_BiologicalProcess-GOA | 0,00 | 0,01 | 0,00 | 0,01 | [4, 5] | Group3 | 7,14 | 2,00 | [GRIK1, GRM2] |
| | GO:0032108 | negative regulatio | GO_BiologicalProcess-GOA | 0,00 | 0,00 | 0,00 | 0,01 | [4, 6, 7, 8, | Group4 | 4,08 | 2,00 | [CCK, GRM2] |
| | GO:0032095 | regulation of resp | GO_BiologicalProcess-GOA | 0,00 | 0,01 | 0,02 | 0,02 | [3, 5] | Group5 | 4,76 | 2,00 | [CARTPT, CCK] |
| | GO:0032098 | regulation of appe | GO_BiologicalProcess-GOA | 0,00 | 0,01 | 0,02 | 0,02 | [3, 5] | Group5 | 8,00 | 2,00 | [CARTPT, CCK] |
| | GO:0032104 | regulation of resp | GO_BiologicalProcess-GOA | 0,00 | 0,01 | 0,02 | 0,02 | [4, 5] | Group5 | 7,69 | 2,00 | [CARTPT, CCK] |
| | GO:0032107 | regulation of resp | GO_BiologicalProcess-GOA | 0,00 | 0,01 | 0,02 | 0,02 | [5] | Group5 | 4,55 | 2,00 | [CARTPT, CCK] |
| | GO:0008066 | glutamate recepto | GO_MolecularFunction-GC | 0,00 | 0,00 | 0,02 | 0,02 | [4, 5, 6] | Group5 | 14,29 | 2,00 | [CARTPT, CCK] |
| | GO:0051954 | positive regulatior | GO_BiologicalProcess-GOA | 0,00 | 0,01 | 0,02 | 0,02 | [3, 4, 5, 6, | Group5 | 5,13 | 2,00 | [CARTPT, CCK] |
| | GO:0032094 | response to food | GO_BiologicalProcess-GOA | 0,00 | 0,01 | 0,02 | 0,02 | [5, 6] | Group5 | 7,69 | 2,00 | [CARTPT, CCK] |
| | GO:0005184 | neuropeptide horr | GO_MolecularFunction-GC | 0,00 | 0,01 | 0,02 | 0,02 | [4, 6, 7] | Group5 | 10,53 | 2,00 | [CARTPT, CCK] |
| | GO:0003254 | regulation of mem | GO_BiologicalProcess-GOA | 0,00 | 0,00 | 0,02 | 0,02 | [4, 5, 6, 7, | Group5 | 16,67 | 2,00 | [CARTPT, CCK] |
| | GO:0050912 | detection of chem | GO_BiologicalProcess-GOA | 0,00 | 0,00 | 0,02 | 0,02 | [5, 6, 7] | Group5 | 14,29 | 2,00 | [CARTPT, CCK] |
| | GO:0014047 | glutamate secretic | GO_BiologicalProcess-GOA | 0,00 | 0,00 | 0,02 | 0,02 | [4, 5, 6, 7, | Group5 | 16,67 | 2,00 | [CARTPT, CCK] |
| Cluster3 | GO:0070330 | aromatase activity | GO_MolecularFunction-GC | 0,00 | 0,00 | 0,00 | 0,00 | [5] | Group0 | 7,41 | 2,00 | [CYP2D6, CYP2D7] |
| | GO:0010310 | regulation of hydr | GO_BiologicalProcess-GOA | 0,00 | 0,00 | 0,00 | 0,00 | [5, 6] | Group1 | 12,50 | 2,00 | [NOXA1, NOXO1] |
| | GO:0043020 | NADPH oxidase co | GO_CellularComponent-G | 0,00 | 0,00 | 0,00 | 0,00 | [3, 4, 5, 6, | Group1 | 20,00 | 2,00 | [NOXA1, NOXO1] |
