## Supplementary tables for "Transcriptome profiling of the dorsomedial prefrontal cortex in suicide victims": S9_List_of_PCR_primers.pdf

Supplementary file 10: Primer sequences of validated genes

| Target name | Primer | Sequence (5' - 3') | Amplicon length (bp) |
|---|---|---|---|
| **GLUL** | GLUL-forward | tgatctccgtggctttggtt | 302 |
| | GLUL-reverse | tgttttgactccagccctgt | |
| **GJA1** | GJA1-forward | tcaagggcgttaaggatcgg | 309 |
| | GJA1-reverse | caaaaggctgtgcatgggag | |
| **GRM2** | GRM2-forward | aagctgagtgacaagtcccg | 307 |
| | GRM2-reverse | cagaacgggtgaacaggaca | |
| **GRIK1** | GRIK1-forward | ttccacacatacagacccgc | 374 |
| | GRIK1-reverse | gtactcggtcatcatgccca | |
| **GRIK2** | GRIK2-forward | tacaaacccaggcgtcttctc | 283 |
| | GRIK2-reverse | ccaccaaatgcctcccactat | |
| **SYT5** | SYT5-forward | acaatcagccatcctcgtctg | 374 |
| | SYT5-reverse | tggacaagcaagggagttctc | |
| **NRGN** | NRGN-forward | cggagactaggccagaactg | 263 |
| | NRGN-reverse | gtttagggtggggtggagtg | |
| **EPHA2** | EPHA2-forward | ggcaaggaagtggtactgct | 327 |
| | EPHA2-reverse | gaagttggtgccgtagtcca | |
| **ITPKB** | ITPKB-forward | acaagccttactgcctggag | 298 |
| | ITPKB-reverse | ctcccagctcctgtacctct | |
| **AQP1** | AQP1-forward | gctatgcgtgctggctacta | 323 |
| | AQP1-reverse | cggcatccaggtcatactcc | |
| **ITGB4** | ITGB4-forward | tccccatctttgctgtcacc | 244 |
| | ITGB4-reverse | cggatgtgaaaggacccagt | |
| **PRKCH** | PRKCH-forward | tcacccccttcctcactcagt | 279 |
| | PRKCH-reverse | atcccctccttgcacattcc | |
| **SLCO2B1** | SLCO2B1-forward | cctggggcagtgtggataat | 294 |
| | SLCO2B1-reverse | gcgaggtggggtggtaatag | |
| **S100B** | S100B-forward | tctggaagggagggagacaa | 206 |
| | S100B-reverse | tcgtggcaggcagtagtaac | |
| **NECAB2** | NECAB2-forward | caggatcttggtgccagct | 360 |
| | NECAB2-reverse | tgtggtcagtgtgggtcatg | |
| **Actin-beta** | Actin-beta-forward | gtgctatccctgtacgcctc | 271 |
| | Actin-beta-reverse | tggccatctcttgctcgaag | |
| **GAPDH** | GAPDH-forward | ccagaacatcatccctgc | 275 |
| | GAPDH-reverse | gtgggtgtcgctgttgaa | |
| **LDHA** | LDHA-forward | ggaggttgtgcatgttgtcc | 290 |
| | LDHA-reverse | cagtgaaggagccaggaagt | |

| GeneBank accession number |
|---|
| NM_002065.7 |
| NM_000165.5 |
| NM_000839.5 |
| NM_000830.5 |
| NM_021956.5 |
| NM_003180.3 |
| NM_006176.3 |
| NM_004431.5 |
| NM_002221.3 |
| NM_198098.3 |
| NM_000213.5 |
| NM_006255.4 |
| NM_007256.4 |
| NM_006272.3 |
| NM_019065.2 |
| NM_001101.5 |
| NM_002046.7 |
| NM_005566.4 |
